# Cell cycle-associated expression patterns predict gene function in mycobacteria

**DOI:** 10.1101/2020.02.10.942268

**Authors:** Aditya C. Bandekar, Sishir Subedi, Thomas R. Ioerger, Christopher M. Sassetti

**Affiliations:** Department of Microbiology and Physiological Systems, University of Massachusetts Medical School, Worcester, MA, 01605, USA; Department of Computer Science and Engineering, Texas A&M University, College Station, TX, 77843, USA

**Keywords:** Mycobacterium tuberculosis, prokaryotic cell cycle, cell cycle associated transcription, just-in-time transcription, divisome assembly and cytokinesis, nucleotide metabolism, cytokinesis

## Abstract

While the major events in prokaryotic cell cycle progression are likely to be coordinated with transcriptional and metabolic changes, these processes remain poorly characterized. Unlike many rapidly-growing bacteria, DNA replication and cell division are temporally-resolved in mycobacteria, making these slow-growing organisms a potentially useful system to investigate the prokaryotic cell cycle. To determine if cell-cycle dependent gene regulation occurs in mycobacteria, we characterized the temporal changes in the transcriptome of synchronously replicating populations of *Mycobacterium tuberculosis* (*Mtb*). By enriching for genes that display a sinusoidal expression pattern, we discover 485 genes that oscillate with a period consistent with the cell cycle. During cytokinesis, the timing of gene induction could be used to predict the timing of gene function, as mRNA abundance was found to correlate with the order in which proteins were recruited to the developing septum. Similarly, the expression pattern of primary metabolic genes could be used to predict the relative importance of these pathways for different cell cycle processes. Pyrimidine synthetic genes peaked during DNA replication and their depletion caused a filamentation phenotype that phenocopied defects in this process. In contrast, the IMP dehydrogenase *guaB2* dedicated to guanosine synthesis displayed the opposite expression pattern and its depletion perturbed septation. Together, these data imply obligate coordination between primary metabolism and cell division, and identify periodically regulated genes that can be related to specific cell biological functions.

## Introduction

Much of prokaryotic cell biology has been elucidated under rapid-growth conditions in which chromosomal replication takes longer than the doubling time of the cell [1], [2]. Under these conditions, the production of complete chromosomes for daughter cells is ensured via the simultaneous initiation of multiple rounds of DNA replication, and it is not possible for cells to temporally segregate DNA replication from cytokinesis as is seen in the eukaryotic cell cycle. However, this paradigm may not apply to many, if not most, of the bacteria in the environment. For example, *Caulobacter crescentus* exploits a specialized developmental program that produces distinct sessile and motile cells, which is associated with a strict cell cycle that segregates DNA replication from cytokinesis. More generally, most of the bacterial biomass in nutrient poor natural environments is likely to persist in slow-growing states [3]. When these conditions are modeled in nutrient-restricted *Escherichia coli*, major cellular events become restricted into distinct cell cycle periods designated, “B”, “C”, and “D”; in which DNA replication is restricted to the C period. The B and D periods occupy the time before or after C, respectively, and are divided by cytokinesis[4] [5]. Similarly, mycobacteria are slow-growing organisms, with doubling times that range from 3 hours to several days, which appear to constitutively employ this type of segregated cell cycle. While an ordered cell cycle may be more common in prokaryotes than generally appreciated, it has only been investigated in a limited number of systems.

The prokaryotic cell cycle has been most thoroughly studied in the aquatic bacterium, *C. crescentus*, largely because it is possible to produce bacterial cultures in which cells are replicating synchronously with respect to the cell cycle. As a result, the only genome-wide transcriptional profiles of synchronously-replicating bacteria have been produced in this species. These studies identified periodic fluctuations in mRNA abundance that correlated with the cell cycle [6], [7] and led to the elucidation of a regulatory cascade that controls cell cycle progression and cellular differentiation that appears to be conserved in other alphaproteobacteria [8]. In addition to these regulators of cell cycle progression, 19% of the genome was found to be periodically expressed [6]. These genes included those involved in primary metabolic processes that are not directly associated with cell cycle progression, suggesting that major cellular events, such as DNA replication and cytokinesis, may be coordinated with other aspects of cellular physiology. These apparent links between metabolism and cell cycle are consistent with a number of studies in model organisms, where metabolites such as NAD(P)H [9] and ATP [10] oscillate according to the cell cycle in *E. coli*; and UDP-glucose levels coordinate cell division timing with nutrient availability in both B. *subtilis* [11] and *E. coli* [12]. While these data indicate that cell cycle progression is likely coupled with some aspects of primary metabolism, it remains unclear how these processes interact and whether insights from transcriptional profiling in *C. crescentus* are generalizable to more diverse bacteria.

We sought to extend these paradigms to mycobacteria, a genus that includes relatively rapid growing environmental organisms, such as *M. smegmatis*, and slower growing pathogenic species such as *M. tuberculosis* (Mtb) and *M. bovis*. The cell cycle has been extensively characterized in both fast- and slow-growing mycobacteria using single-cell analyses of strains engineered to express fluorescent markers of DNA replication and cytokinesis. These studies show that DNA replication occurs only once per cycle in the majority of cells, and re-initiation of replication before division occurs only rarely [13] [14][15][16][17]. While the relative durations of the cell cycle phases can be influenced by the environment and stochastic factors such as birth length [16], the average duration of the B, C, and D periods in Mtb are generally in the range of 6-8 hours, 9-12 hours, and 6-9 hours, respectively [18].

These observations from time-lapse microscopy are consistent with metabolic labeling studies in synchronously-replicating cultures of *Mtb*, which can be generated using a mutant strain that harbors a cold-sensitive (cos) allele of the DNA replication initiator DnaA [19]. This *Mtbcos* strain is unable to initiate a new round of DNA replication at 30°C. Upon release into the permissive temperature (37°C), cultures synchronously incorporate 5,6-^3^H-uracil into alkali stable DNA for 11 hours, consistent with the C period length observed in single cells. The ability to produce synchronously replicating cultures that recapitulate the behavior of single cells makes mycobacteria an attractive system to investigate cell cycle-associated transcriptional changes.

Using the *Mtbcos* strain, we determined the transcriptional profile of synchronously replicating cultures across the cell cycle, and report that 485 genes meet strict criteria for periodic gene expression. Only a small fraction of the cell cycle-regulated gene sets of *Mtb* and *C. crescentus* overlap, suggesting that the links between cell cycle and metabolism are species-specific. We demonstrate that mRNA expression patterns in *Mtb* reflect the time at which the encoded proteins are incorporated into the developing septum, suggesting that functional information can be inferred from the kinetics of gene expression. Using this framework, we discover an unanticipated functional specialization of distinct nucleotide anabolic pathways. These observations show that DNA replication and cytokinesis are coordinated with different primary metabolic pathways, expanding the processes that are associated with these essential cellular events.

## Results

### DNA replication and cytokinesis are segregated in synchronously growing populations of *Mtb*

We generated synchronously replicating cultures of *Mtb* using the temperature-sensitive *Mtbcos* strain [19]. Chromosomal replication was inhibited by incubating this strain at 30°C for 36 hours. Upon shift to the permissive temperature (37°C), the optical density (Absorbance_600_) of both the *Mtbcos* mutant and a wild type control culture increased throughout a 54 hour time course, demonstrating that nutrients did not become limiting. While the control culture grew at a constant rate over this period, the *dnaAcos* strain showed a reproducible multiphasic growth pattern, an initial indication that cellular metabolism may be linked to cell cycle events (**Fig1A**).

**Figure 1:**
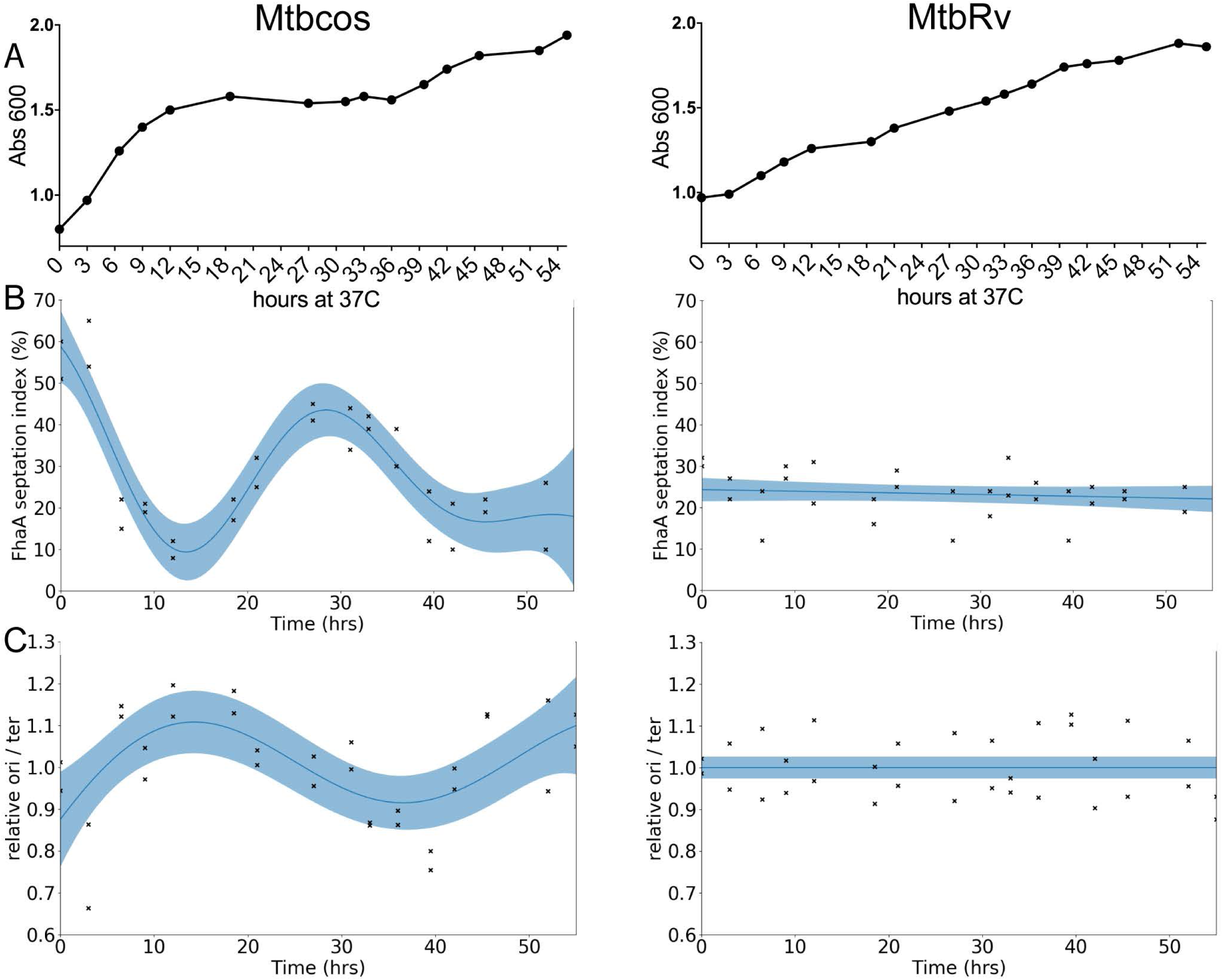
DNA replication and cytokinesis are segregated in synchronously growing populations of Mtb. (A) Growth of *Mtbcos* (left) and *MtbRv* (right) after release into permissive temperature 37°C. X axis: hours at 37°C. Y axis: Absorbance_600_ (B) FhaA septation index assay to determine cytokinesis phase. Percentage of *Mtbcos* (left) and *MtbRv* (right) populations containing an FhaA-venus focus localized at midcell after release into permissive temperature. Data points are representative of two biological replicates. Blue line is smoothed via the Gaussian process. The blue band indicates 95% confidence interval. Significant difference between *Mtbcos* and *MtbRv* curves was determined using a likelihood ratio test which determines if the data is fit best by a combined model (null hypothesis) or separate strain specific models (alternate hypothesis). Log_likelihood difference between combined and separate models = −38.489, p-value (chi-squared distribution; df=3) = 1.1e-16. (C) Origin/terminus assay to determine the DNA replication phase. Relative ori/ter ratio of *Mtbcos* (left) and *MtbRv* (right) populations after release in permissive temperature. Data points are representative of two biological replicates. Blue line is smoothed via the Gaussian Process. The blue band indicates 95% confidence interval. Significant difference between *Mtbcos* and *MtbRv* curves was determined using a likelihood ratio test Log_likelihood difference between combined and separate models = −12.412, p-value (chi-squared distribution; df=3) = 1.679e-05.

In order to estimate the efficiency of the synchronization and to delineate cell cycle periods, we collected cells every three hours and monitored chromosomal replication and cytokinesis. The phosphothreonine-binding protein, FhaA, marks sites of division [20], and we used a fluorescent allele of this protein to calculate a “septation index” that corresponded to the fraction of cells with FhaA localization at midcell. While the septation index of a control culture of asynchronous cells (*MtbRv*) was constant throughout the time course, this metric varied in a periodic manner in the *Mtbcos* strain. The majority of cells arrested at the non-permissive temperature had an FhaA focus at midcell, which is likely an artifact of the DnaA inactivation. The septation index of *Mtbcos* cells quickly decreased upon shift to the permissive temperature, falling below the index of unsynchronized cultures by 12 hours. Septation reached a peak in the synchronized cultures between 27 and 33 hours after release, marking the cytokinesis phase of the cell cycle (**Fig1B**).

To monitor chromosomal replication, we devised a PCR assay to quantify the relative abundance of DNA at the origin (ori) and terminus (ter) of replication. Upon initiation of replication, the cell will have an ori:ter ratio of 2:1, and this ratio should be maintained until the terminus is duplicated. As we observed for septation index, the ori:ter ratio remained constant in unsynchronized cultures. In contrast, the ori:ter ratio peaked twice in the synchronized cultures (**Fig 1C**). The first peak lasted for a duration of approximately 12 hours (between 15h-27h post release) and second one lasted between 48-hrs post release and the end of the study. Based on these data, we estimate that our time course captured approximately 1.5 cell cycles. Both the septation index and ori/ter ratio varied by approximately 50% of the range expected of fully synchronized cells, indicating that the synchrony of our cultures was incomplete. Regardless, these observations indicated that DNA replication is temporally segregated from cytokinesis, and the cultures were sufficiently synchronized to perform transcriptional profiling.

### Periodic gene expression correlates with cell cycle progression

In order to investigate whether gene expression changes are associated with major cellular events like DNA replication and cytokinesis, we profiled mRNA abundance in synchronized cultures every 3 hours across a 54 hour time course. After normalization and scaling, we first assessed correlation patterns in the dataset. The initial time point after temperature shift to 37°C (0hr) was uncorrelated with the rest of the datasets, presumably due to an adjustment to the temperature shift and was omitted from subsequent analyses. For the remaining data, we found the highest degree of correlation between adjacent time points, as expected for a time-resolved dataset **(Fig 2A)**. While this was generally true for both the synchronized *Mtbcos* and unsynchronized *MtbRv* strains, the correlation matrix from the *Mtbcos* cultures displayed a distinct three-block structure suggesting the presence of transcriptionally distinct phases. This structure is even more apparent upon hierarchical clustering, which revealed a pattern of gene expression consistent with an ordered progression of events throughout the time course (**FigS1A**).

**Figure 2:**
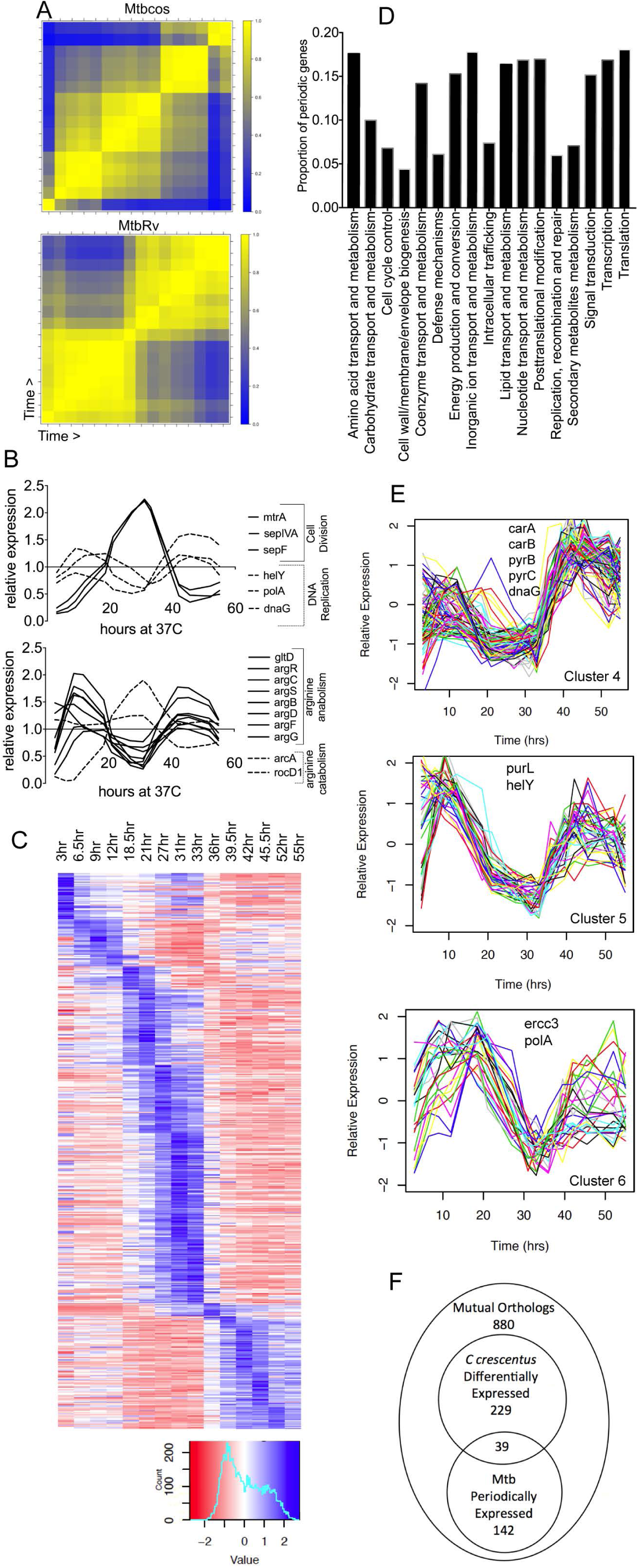
Periodic gene expression correlates with cell cycle progression. (A) Correlation matrix of DESeq normalized counts for single replicates of *Mtbcos* (top) and *MtbRv* (bottom) for all 16 timepoints. (Blue: Pearson’s correlation coefficient=0; Yellow: Pearson’s correlation coefficient=1). (B) Relative expression (GP smoothed, DESeq normalized read count for each time point divided by the mean value for that gene across time) of genes involved in DNA replication & cell division (top panel); arginine catabolism & anabolism (bottom panel). (C) Relative expression (standard normalized DESeq counts - each value is subtracted by the mean for that gene across time and then divided by the standard deviation) of 485 periodically expressed genes in *Mtbcos* (rows) sorted by peak expression time (columns). (D)Fraction of periodically expressed genes present in different Gene Ontology categories. (E) Clusters containing periodic genes with expression patterns consistent with a role in DNA replication. Known DNA replication/nucleotide biogenesis genes found in these clusters are listed. X axis: hours post release into permissive temperature. Y axis Relative expression (standard normalized DESeq counts - each value is subtracted by the mean for that gene across time and then divided by the standard deviation) (F) Overlap between periodically expressed (*Mtbcos*) and differentially expressed (*C. crescentus*) mutual orthologs.

To take advantage of both replicate measurements and the relatedness of adjacent time points, we used a Gaussian Process (GP) smoothing approach to estimate relative expression levels of each gene across the time course **(Fig S1B)**. The expression of many genes with known cell-cycle related functions were found to peak during the appropriate period. For example, genes that are important for cell division, such as the regulator, *mtrA* [21] or the septal components, *sepF* [22] and *sepIVA* [23], peak in expression once during cytokinesis. Similarly, genes important for DNA replication, such as those encoding DNA primase (*dnaG)* and the replicative polymerase (*polA)* display two expression peaks during this time course, corresponding to DNA replication (**Fig2B**). In addition, we found that the expression pattern of several primary metabolic pathways mirrored these cell-cycle related genes, and suggested alterations in metabolic flux. For example, genes necessary for arginine biosynthesis were co-regulated and had opposing expression patterns to genes involved in arginine catabolism (**Fig2B**).

We sought to more formally define genes with an expression pattern consistent with cell cycle progression. First, we removed genes with correlated expression patterns in synchronized and unsynchronized cultures to minimize the effect of changes in culture conditions during the time course. Next, we fit the raw data from both replicates for each gene to a sinusoidal function with the expected period of the *Mtb* cell cycle, optimizing the parameters for trend, amplitude, period and phase. Genes with a period outside the range of reasonable expectations based on the *Mtb* cell cycle were filtered out, along with genes with low overall expression or high variance between replicates. A goodness-of-fit criterion based on curve fitting residuals was applied, which maximized the difference in genes discovered in synchronized versus unsynchronized cultures. These criteria were determined to produce a false discovery rate of 0.35% using a permuted dataset, and a 2.6- fold enrichment in genes discovered in the synchronized data set, compared to the unsynchronized set. 485 genes were categorized as periodically expressed (**Fig 2C**; **Supplementary Table 2),** and all major functional categories were represented in this gene set (**Fig2D**).

Clustering this set of periodically-regulated genes further highlighted the association between gene function and cell cycle stage. Genes were distributed into 8 clusters using hierarchical clustering of the *Mtbcos* expression profiles **(FigS2),** producing groups of coordinately regulated genes with peak expression values ranging across the time course. Three of these clusters reflected expression profiles that peak during DNA replication (**Fig2E**). In these clusters, we find 9 genes with defined roles in DNA replication (*dnaG, polA, helY, ercc3*) and nucleotide biogenesis (*carA, carB, pyrB, pyrC, purL*), further indicating that expression patterns were consistent with gene function.

The periodically regulated gene set (485 genes) in *Mtb* represents 12% of the genome, whereas 19% of the *C crescentus* genome was previously found to be cell cycle associated [6][7]. In order to investigate whether similar cellular functions are periodically regulated in these phylogenetically diverse organisms, we compared the expression patterns of orthologous genes. Out of the 880 mutual orthologs identified as being reciprocal best BLAST matches, 229 genes were defined as cell cycle regulated in *C. crescentus* by virtue of being differentially expressed over time in synchronized cultures [7] and we found 142 to be periodically expressed in *Mtb* with an overlap of 39 genes (**Fig2F, Supplementary Table 3).** This overlap contains a number of genes with known cell cycle associated functions, such as the DNA replication initiator, *dnaA*, the DNA primase *dnaG* and nucleotide biogenesis (*nrdZ, purl, pyrB*). However, the modest degree of overlap also suggests that independent sets of genes are periodically expressed in these two phylogenetically distinct organisms.

### mRNA abundance predicts the order of assembly of mycobacterial divisome components and regulators

The order in which large multicomponent structures, such as the flagellum, are assembled in bacteria can be predicted based on the transcriptional regulation of the corresponding genes[24]. Based on this “just in time” transcription model [25], we hypothesized that the assembly of the large complex of proteins necessary for cell division, aka the “divisome”, may follow the same principles and provide a model to test whether mRNA abundance could be used to predict the timing of gene function in our system. To test this model, we assessed the temporal coincidence between mRNA abundance and protein localization at the developing and maturing septum.

To broadly identify genes that are induced during cytokinesis (27-33h), we identified genes that only peak once in the time course **(Supplementary Table 1**) and clustered genes based on similar expression patterns **(Fig3A).** Within the clusters that peak in expression at the appropriate times, we found a number of genes known to be involved in cytokinesis (**Fig3A**). The first to be induced was *ftsZ*, the tubulin-like nucleator of the septum, and the septally-localized Ser/Thr kinase, *pknD*. This was followed by the expression of genes encoding several divisome-associated proteins, FtsW, SepIVA, and LamA. After these, we found the gene encoding the new pole landmark protein, DivIVA, was induced. To determine if the timing of expression predicted the order of assembly, we chose three genes, *pknD*, *ftsW* and *divIVA* which peaked in expression early, in the middle, and late during the cytokinesis window, respectively. Pairs of these proteins were fused with fluorescent tags and their cellular location was determined by time lapse microscopy in *M. smegmatis*, a related mycobacterial species that expresses orthologs of these proteins and is an experimentally tractable model of mycobacterial division [26]. Dual-color fluorescence imaging revealed that PknD, FtsW, and DivIVA appear at the developing septum in the order predicted by mRNA abundance (**Fig 3B-D**). This order was not dictated by the maturation times of the two different fluorophores. Although not directly addressed in our experiments, it is likely that order of localization at the septum is independent of transcriptional regulation because PknD, and DivIVA fusion proteins were expressed from constitutive promoters. Instead, the transcriptional order correlates with assembly, which appears to be dictated at the posttranscriptional level.

**Figure 3:**
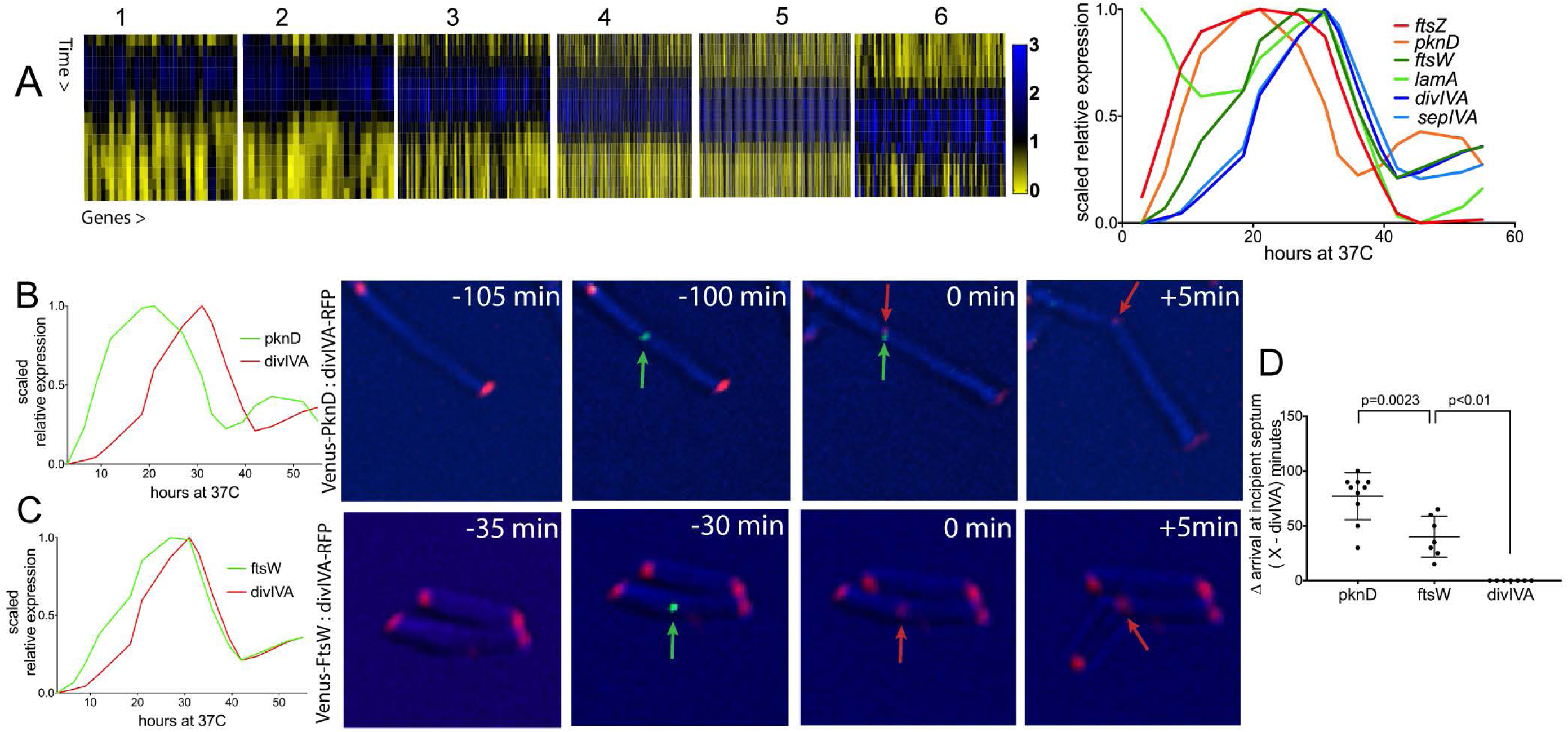
mRNA abundance predicts the order of assembly of mycobacterial divisome components and regulators. (A) (Left) Clusters of *Mtbcos* genes with expression patterns that peak during the cytokinesis period. (Right) Scaled relative expression (GP smoothed, DESeq normalized read count for each time point divided by the mean value for that gene across time, then scaled) of known cytokinesis genes from these clusters. (B) (Left) Scaled relative expression of PknD and DivIVA. (Right) Time-lapse imaging of *M smegmatis* expressing PknD-Venus (green) and DivIVA-RFP (red). Time (minutes) before the arrival of DivIVA at midcell is indicated. (C) (Left) Scaled relative expression of FtsW and DivIVA. (Right) Time-lapse imaging of *M smegmatis* expressing FtsW-Venus (green) and DivIVA-RFP (red). Time (minutes) before the arrival of DivIVA at midcell is indicated. (D) Time (minutes) between initial arrival of PknD (n=10), FtsW (n=7) and DivIVA at midcell. Error bars indicate mean ± SD. Statistically significant difference between pknD and ftsW determined using an unpaired T-test (α =0.05; p=0.0023). Statistically significant difference between ftsW and divIVA determined using a chi-squared test (α =0.05; χ^2^=7; df=1; p<0.01).

### Guanosine synthesis influences cytokinesis in mycobacteria

Having demonstrated that gene expression can predict the timing of gene function, at least in the case of the developing septum, we investigated whether obligate coordination exists between cellular events, such as DNA replication and cytokinesis, and upstream pathways that produce the precursors for these processes. In particular, we focused on nucleotide metabolism by analyzing the expression patterns of enzymes catalyzing anabolic reactions beginning from the tricarboxylic acid cycle precursor, glutamate to the final nucleos(t)ide products. Pyrimidine biogenesis from the very early stages of the *carAB-*encoded reactions down to the *pyrBCDEF-*encoded reactions were most highly expressed during the C phase at ∼12 hours and ∼48 hours (**Fig4A**), consistent with previous reports of an increased production of thymine nucleotides during DNA replication in *E coli* [27]. This also corroborates reports of nucleotide pool sizes in *E coli* being insufficient for chromosome replication and requiring *de novo* synthesis before the onset of the DNA replication phase [28]. Unexpectedly, the converse expression pattern was observed for the first reaction unique to guanosine synthesis *(guaB2)*. While both adenosine and guanosine purine rings are synthesized from the common precursor inosine monophosphate (IMP), the IMP dehydrogenase, GuaB2, catalyses the *first* dedicated reaction unique to GMP synthesis. *guaB2* peaked in expression during cytokinesis at ∼21hours (**Fig4A**). Genes dedicated to synthesizing adenosine from IMP, *purB* and *amk,* did not appear to be cell cycle regulated.

**Figure 4:**
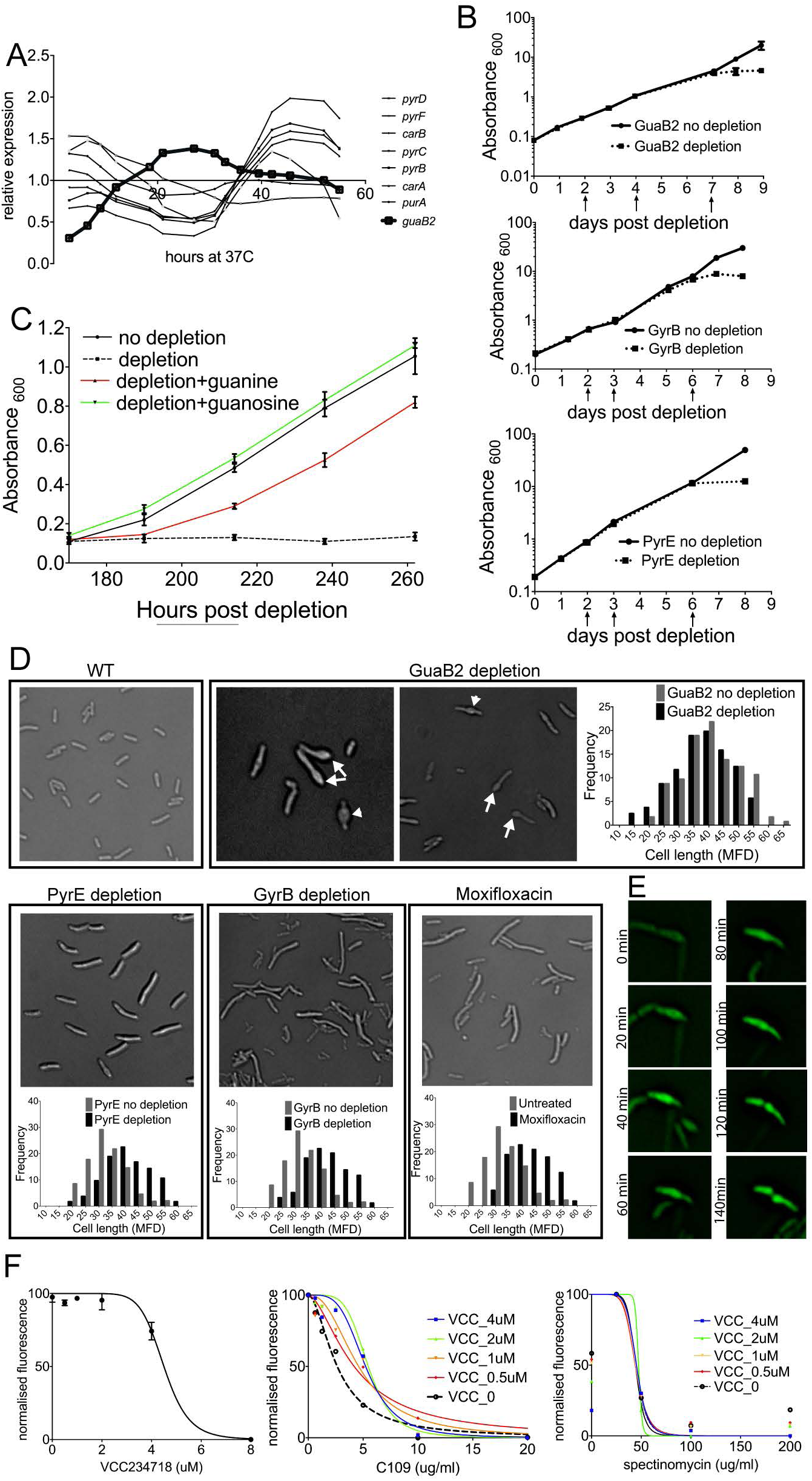
Guanosine synthesis influences cytokinesis in mycobacteria. (A) Relative expression (GP smoothed, DESeq normalized read count for each time point divided by the mean value for that gene across time) of IMP dehydrogenase *guaB2* compared to pyrimidine biosynthesis genes *carA*, *carB*, *pyrB*, *pyrC, pyrD, pyrF* (B) Cumulative growth (Absorbance_600_) of *M tuberculosis guaB-2*DAS (top), *gyrB-*DAS (center), *pyrE-*DAS (bottom) without depletion (solid line) and with depletion (dotted line). Arrows indicate the time during the pre-depletion period when cultures were back diluted into fresh growth medium. Data are represented as mean ± SD of two biological replicates (C) Growth (Absorbance_600_) of *M tuberculosis guaB2*-DAS ±depletion in the presence of either 200μM guanine or guanosine. Data are represented as mean ± SD of two biological replicates. (D) *M tuberculosis* cellular phenotypes upon genetic depletion of GuaB2, PyrE, GyrB. Images were obtained after the cessation of growth in depleted cells. In the case of GuaB2, septal bulges (arrowheads) and polar bulges (arrows) are indicated. Histograms indicate the cell length distribution of cells in which the target was either not depleted (gray) or depleted (black). *Mtb* was treated with 0.2μM moxifloxacin for 24 hours and imaged. Histograms indicate the cell length distribution of cells in untreated (gray) or treated cells (black). MFD = Maximum Feret Diameter (1μM ∼ 0.11MFD). Statistically significant/non-significant difference between the cell length distributions was determined using the Mann-Whitney test. (p_guaB2_=0.214; p_pyrE_<0.001; p_gyrB_<0.001; p_moxifloxacin_<0.001) (E) Time-lapse imaging at 20 minute intervals of GFP-expressing *M smegmatis* treated with 2μM VCC234718. (F) Left: Susceptibility of *M smegmatis* to VCC234718. Data are represented as mean ± SD Center: Cross titration assay on GFP-expressing *M smegmatis* with the indicated concentrations of VCC234718 and C109. Statistically significant difference between the VCC alone curve and other curves was determined using an extra-sum-of-squares F-test (α=0.05). p_(VCC 0.5μm)_= 0.0215; p_(VCC 1μM)_= 0.0284; p_(VCC2μM)_= 0.0001; p_(VCC4μM)_= 0.0019 Right: Cross titration assay on GFP-expressing *M smegmatis* with the indicated concentrations of VCC234718 and spectinomycin. The differences between the VCC alone curve and other curves were not significant, as determined using an extra-sum-of-squares F-test (α =0.05). p_(VCC0.5μM)_= 0.9989; p_(VCC1μM)_= 0.9999; p_(VCC2μM)_=ambiguous; p_(VCC4μM)_= 0.9978

The reciprocal expression patterns of pyrimidine and guanosine synthetic genes suggested that the requirement for these metabolites varied across the cell cycle and these requirements were associated with distinct cellular events. To investigate this hypothesis, we generated mutant *Mtb* strains in which synthesis of pyrimidines or guanosine was inhibited via the inducible genetic depletion of the PyrE or GuaB2 proteins, respectively. Each gene was fused to a C-terminal DAS+4 tag (DAS) that facilitated Clp protease-mediated degradation upon removal of anhydrotetracycline (aTc). In both cases, protein depletion inhibited bacterial growth (**Fig4B**), consistent with the essentiality of these pathways [29]. As *Mtb* expresses three GuaB paralogs, we verified that GuaB2 is essential for guanosine synthesis by supplementing the *guaB2-DAS* strain with 200uM guanine or guanosine. Consistent with previous studies [30], guanine partially rescued the growth defect of the depleted strain, whereas guanosine supplementation led to complete rescue (**Fig4C**).

The inverse expression patterns of *pyr* genes and *guaB2* implied increased *de novo* synthesis of pyrimidine nucleotides and guanosine was preferentially required for DNA synthesis or cytokinesis, respectively. To test this hypothesis, we used morphological criteria to infer which cellular processes were primarily impacted by the inhibition of these nucleotide synthetic pathways. PyrE depletion resulted in cell elongation before growth arrest. The mean cell length increased from 3.3uM to 4.2uM upon PyrE depletion (**Fig4D**). A similar phenotype was observed upon DNA gyrase GyrB depletion and in cells treated with the gyrase inhibiting fluoroquinolone, moxifloxacin (**Fig4D**), which disrupts DNA replication and causes cell filamentation in *E coli* [33]. Cell elongation upon inhibition of DNA replication is consistent with a number of previous observations in *B. subtilis* [31] and *M. smegmatis* [32].

In contrast to the elongation observed upon PyrE depletion, GuaB2-depleted cells were the same length as wild type, but many of these growth arrested cells had bulges at midcell or one pole, suggesting that GuaB2 depletion may influence cell division (**Fig4D**). To determine if the polar bulges were derived from misshapen septa, we performed time-lapse microscopy in *M. smegmatis* cells treated with a specific chemical inhibitor of GuaB2 (VCC234718). Similar to genetic depletion, chemical inhibition of the GuaB2 enzyme also inhibited growth and produced bulges at midcell or one pole (**Fig4E**). Time-lapse microscopy revealed that cells began to bulge at midcell by the completion of 1-2 cell cycles after the initiation of VCC234718 treatment, and that misshapen poles were derived from these bulges. Taken together, these observations imply that the cellular requirement for guanosine increases during cytokinesis, and that this requirement is reflected in the aberrant septation of guanosine-depleted cells.

FtsZ is among the most abundant proteins in the cell and this tubulin-like protein binds and hydrolyzes GTP [34] as it undergoes the cycles of polymerization and depolymerization necessary for septation [35]. This GTP requirement suggested a possible mechanistic connection between guanosine nucleotide levels and septation. To investigate whether the effect of guanosine depletion on septation could be attributed to altered FtsZ dynamics, we determined the effect of inhibiting both processes simultaneously using the GuaB2 inhibitor VCC234718 and C109, an inhibitor of FtsZ GTPase activity and polymerization [36]. Consistent with the hypothesized mechanistic link, we observed significant antagonism between these compounds. Even at concentrations of VCC234718 that alone had no effect on growth (0.5 - 4uM), this compound consistently increased the IC50 of C109 (**Fig4F**). In contrast, we found no interaction between VCC234718 and spectinomycin, an inhibitor of another major GTP consuming pathway, translation. C109, has previously been found to act additively with PCI90723, a compound that stabilizes the FtsZ filament. The opposite antagonistic interaction we observed between C109 and a GuaB2 inhibitor implies that guanosine inhibition inhibits polymerization, an effect that is consistent with the known GTP requirement for FtsZ polymerization.[34], [37]. Taken together, these data are consistent with a model in which transcriptional induction of GuaB2 during cytokinesis coincides with the increased consumption of GTP by FtsZ, and the septal defects observed on guanosine depletion are related to defects in FtsZ dynamics.

## Discussion

Previously, global transcriptomic studies of the prokaryotic cell cycle have been limited to the *C. crescentus* model. In this study, we leveraged a genetic strategy to synchronize *Mtb* cultures, and characterized cell cycle associated transcriptional changes in this organism. Comparisons between our *Mtb* studies and previously generated *C. cresentus* data are limited by a number of technical differences, including the method and degree of synchronization, that likely limited our ability to discern periodic patterns of relatively small amplitude in the *Mtb* dataset. Regardless, our findings are broadly comparable to observations in *C. crescentus*, as we found that a similar fraction of the genome is differentially expressed across the cell cycle in both systems, and a small fraction of orthologous genes are periodically-regulated in both. Despite these similarities, the majority of cell-cycle associated transcriptional changes (82% of *C. crescentus* genes and 72% of *Mtb* genes) were unique to each organism. Thus, despite some similarities, our analysis indicates that cell cycle progression is associated with distinct transcriptional networks in these structurally and phylogenetically divergent organisms.

In a number of cases, we found that increases in mRNA abundance could be used to associate genes with temporally-resolved cell cycle events, such as septation. The sequential expression of divisome components as cell division progresses has been observed previously in *C. crescentus* [6]. We provide functional evidence that this hierarchical expression pattern is associated with the order of divisome assembly in mycobacteria by demonstrating that the timing of gene induction correlates with the recruitment of the encoded proteins to the developing septum. Based on transcriptional data, we inferred the following order of assembly at the mycobacterial septum: FtsZ>PknD>FtsW>LamA>SepIVA>DivIVA. The recruitment of these proteins spans sequential processes of divisome assembly and new pole biogenesis. FtsZ initially marks the future division site [38], facilitating the recruitment of divisome components such as FtsW [39] and SepIVA [23]. The arrival of LamA at the later stages of assembly is consistent with its role in delaying septation and thereby promoting asymmetric cell division[40]. DivIVA is thought to be recruited to the negative curvature of the new pole after septation[41][42][43] and the segregation of daughter cell cytoplasm [13]. While these observations highlight that gene expression patterns can be used to predict the order of complex assembly, the outcome of this coordination remains unclear. As only a subset of currently known septal components were found to be periodically expressed, transcriptional regulation is unlikely to be the primary determinant of assembly order. Instead, this type of hierarchical gene expression has also been proposed as a mechanism to maximize efficiency by restricting protein expression to the period when it is needed [24][25]. Regulation of divisome assembly and function likely involves additional posttranslational mechanisms, as we found that the Ser/Thr kinase, PknD, is recruited relatively early in septal development and previous work described an important role for the Ser/Thr phosphatase, PstP, in cell division[44], [45]. While it remains possible that transcriptional regulation controls some aspects of septation, our data only show that expression pattern can predict the timing of gene function.

The observed periodic expression of primary metabolic functions suggested the importance of coordinating cell cycle events with the upstream pathways that provide their precursors. This model is supported by our finding that pyrimidine synthetic genes peak during a distinct cell cycle period than the dedicated guanosine synthetic gene, GuaB2; and that the depletion of these genes primarily disrupts different cellular processes. It is not surprising that the requirement for pyrimidines would increase during DNA synthesis, as *de novo* nucleotide synthesis is necessary to replicate the chromosome[28]. However, the distinct cytokinesis defect observed upon GuaB2 depletion was unanticipated. Guanine nucleotides are important for a myriad of cellular processes, including DNA replication, transcription, and macromolecular synthesis. We speculate that the preferential septation defect we observe upon GuaB2 depletion is related to the relatively low affinity of FtsZ for GTP. An accurate determination of nucleotide pool sizes in mycobacteria is still an open question [46], but in *E coli,* FtsZ has ∼500-fold lower affinity for GTP than the DnaE1 replicative DNA polymerase (K_m_ FtsZ_GTP_ =1mM[47]; K_m_DnaE1_GTP_=2uM[48]). Thus, the reported intracellular concentration of GTP[49] would support only one-half of the V_max_ of FtsZ [47]. These data indicate that GTP levels may limit FtsZ dynamics. It is likely that GuaB2 depletion leads to aberrant FtsZ activity and not a complete loss of function, since genetic depletion of FtsZ leads to filamentation [50], a phenotype distinct from the one we observe upon GuaB2 depletion. Instead, alterations in FtsZ filament length or rate of turnover may underly these defects.

Both DNA replication and cell division are essential processes that have been the focus of antibacterial drug discovery efforts[46][51]. In most cases, these efforts focus on inhibiting a limited number of physical components of the bacterial replisome or divisome. Our transcriptional data identified a wide variety of genes that are coordinately expressed with the genes encoding these complexes, and therefore may be required for their activity. While we have only investigated these functional dependencies in the context of nucleotide synthesis, our data suggest that many similar dependencies exist and can be predicted from transcriptional profiles. If so, these data could be used to identify a wealth of new strategies for inhibiting these specific essential cellular processes.

## Acknowledgements

We would like to thank Aashish Srivastava under the guidance of James Sacchettini and the Texas A&M Genomics and Bioinformatics Service for sequencing the RNASeq libraries. VCC234718 was a generous gift from Vinayak Singh & Valerie Mizrahi. C109 was a generous gift from Vadim Makarov. Genetic depletion strains were generated as part of a larger collaborative effort including the labs of Deborah Hung, Eric Rubin, Sabine Ehrt and Dirk Schnappinger. Kenan Murphy and Charlotte Reames contributed to strain generation from the Sassetti lab. We would like to thank Megan Proulx for critical reading of the manuscript. This work was supported by grant AI095208 to CMS.

## Author contributions

Conceptualization- ACB, TRI, CMS

Methodology- ACB, TRI

Investigation- ACB

Validation- ACB

Formal Analysis- ACB, SS, TRI

Writing Original Draft- ACB

Writing Review & Editing- ACB, TRI, CMS

Funding acquisition- CMS

Supervision- CMS

## Materials and Methods

### Strains

The *Mtbcos* strain was obtained from [19]. *MtbRv* is the H37Rv strain used as an unsynchronized control. Mtbcos and MtbRv expressing FhaA m-venus were transformed with pKP887 (mycobacterial replicating plasmid MEH expressing MSMEG FhaA-Venus expressed from the MSMEG *fhaA* native promoter (from K.P. Sundaram). *M smegmatis* expressing ftsW-mVenus and divIVA-RFP was transformed with ptb21-ftsW-mVenus-MEK and tb21-divIVA-RFP-MCtH. *M smegmatis* expressing pknD-mVenus and DivIVA-RFP was transformed with p16-pknD-mVenus-MEK [52] and tb21-DivIVA-RFP-MCtH. *Mtb* hypomorphs used in this study were generated as part of an earlier study [53] using a controlled protein degradation system described previously[54]. Three strains were used in this study: *Mtb guaB2*-DAS-Hyg^R^+Giles-TetON1-sspB-str^R^; *Mtb* gyrB-DAS-Hyg^R^+Giles-TetON6-sspB-str^R^ *; Mtb* pyrE-DAS-Hyg^R^+Giles-TetON1-sspB-str^R^. *M smegmatis* expressing green fluorescence contains the plasmid CT161 (m-Venus pMV261 Hyg^R^) obtained from the Eric Rubin Lab.

### Mtbcos synchronization

Cultures of Mtb*dnaAcos115* generated in a previous study [19], MtbH37Rv, Mtb*dnaAcos115*-FhaA-Venus and MtbH37Rv-fhaA-Venus were grown in standard culture media at 37°C under shaking conditions till OD_600_ 0.4. The cells were shifted to 30°C for 36 hours. The cultures were then shifted to 37°C and the cultures were processed for either DNA isolation, RNA isolation or fluorescent microscopy at the following times: 0h, 3h, 6.5h, 9h, 12h, 18.5h, 21h, 27h, 31h, 33h, 36h, 39.5h, 42h, 45.5h, 52h and 55h.

### Chromosomal DNA isolation

Chromosomal DNA was isolated from the cell pellet of 5ml culture from each timepoint. Briefly, 0.5 ml of chloroform:methanol (2:1) was added and the mixture was vortexed 5X 1min. 0.5ml of phenol:chloroform was added and the mixture was vortexed for 30 seconds. Finally, 0.5ml of TE buffer was added. This was centrifuged at 12,000g at4C for 5 minutes. The upper phase was mixed with 1 volume of chloroform and vortexed. After centrifugation, the upper phase was added to a new tube and 1/10 volume of 3M sodium acetate and 1 volume of isopropanol was added. Precipitated DNA was spun out of solution and resuspended in 20ul of TE buffer.

### Origin:terminus assay

Multiple primer sets (designed using the Primer3 design tool) amplifying 150bp at each location (Origin-0MB region surrounding Rv0001; Terminus –2.2MB region surrounding Rv1949c) of the *Mtb*H37Rv genome were tested for amplification efficiency. Efficiency was calculated from the negative slope of the standard curve of C_T_ v/s template concentration. The primer sets with the highest and most similar efficiencies for both loci were selected (95% for the origin and 93% for the terminus). Quantitative PCR was done using SYBR green (Biorad iQ SYBR Green Supermix) with 2ng of gDNA template per reaction. Delta Ct values were calculated as dCt= Ct_ori_-Ct_ter_. 2^-dCt values were then calculated for each timepoint. These values were then divided by the mean 2^-dCt across all timepoints to generate a relative ori/ter ratio for each timepoint.

### Microscopy

#### Static imaging

At each time point post release into 37°C (Fig1B) or timepoint post genetic depletion of GyrB, GuaB2 and PyrE (Fig4D), 1ml of *Mtb* culture was centrifuged and cells were re-suspended in a phosphate buffered saline solution containing 0.05% Tween80 and 4% paraformaldehyde. These fixed cells were then placed onto an agarose pad and DIC (Fig4D) or wide field fluorescence imaging (Fig1B) was performed with a DeltaVision Personal DV microscope (GE Healthcare) using a 60X oil immersion objective (AP). Cell lengths in Fig4D were determined using CellProfiler™ [55] which calculates a MFD (Maximum Feret Diameter) which is a measurement of the largest number of pixels between the two ends of the cell obtained while rotating a caliper along all possible angles. The approximate conversion factor of MFD to microns is 0.11. Calculating an MFD is especially useful for measuring mycobacteria since all cells are not strict rods (cells undergo V-snapping prior to resolution of cytokinesis and daughter cell separation). The cell debris observed during GyrB depletion in Fig4D was excluded from cell length quantification by training CellProfiler using CP Analyst™.

#### Live cell imaging

10ul of cells in logarithmic phase (OD_600_ 0.2-0.5) were spotted on a glass bottom 24-well plate (MatTek Corporation). 500ul of molten Luria Bertani medium (40-50C) was spread over the cells and allowed to solidify. For experiments with VCC234718, molten LB containing 2uM final concentration of VCC234718 was prepared before layering over the cells. Time-resolved imaging was performed with a DeltaVision Personal DV wide field fluorescence microscope equipped with Ultimate Focus™ capabilities and an environmental chamber warmed to 37 °C (Applied Precision). Images were taken at 5 or 10 minute intervals.

### RNA isolation, library preparation and sequencing

At each timepoint, 45ml culture was pelleted and resuspended in 1ml of TRIzol (Invitrogen) and transferred to lysing matrix tubes (MP Biomedicals: Lysing Matrix B). Cells were lysed in a MP Biomedicals Fast Prep-24 homogenizer (maximum power-6.5, 4 × 30s cycles, rest on ice for 5 minutes in between cycles). RNA was purified according to the manufacturer’s directions. RNA cleanup was performed with Qiagen RNeasy Mini kit (74104) omitting the DNase step. Instead, after elution, in-tube DNase treatment was performed using Ambion DNase Turbo. RNeasy cleanup was repeated again with double volumes of RLT and ethanol. RNA was subjected to rRNA removal with Ribozero Bacteria kit (Illumina-MRZB12424). Deep sequencing library was prepared using KAPA Stranded RNASeq kit (KK8401). The RNAseq libraries were sequenced on an Illumina 2500 instrument in paired-end mode, using a read-length of 150+150bp. The mean number of reads per sample was 8.9M (range 4.2-16.5M). The reads were mapped to the H37Rv genome using Burroughs Wheeler Alignment [56] with default parameter settings. Reads mapping to each ORF were totaled (sense strand only). Because certain loci were over-represented (e.g. rrs, rnpB, ssr, Rv3661, which had counts ∼0.5-1M), counts were truncated to a maximum coverage of 10,000 (reads/nt).

### Data normalization, filtering and centering

The global expression profiles of *Mtbcos* samples showed a gradual increase in expression of a few genes that dominate expression at latter time-points. Consequently, a compensatory decrease was observed in expression of other genes, making normalization by traditional reads per kilobase per million (RPKM) mis-representative. To correct for the bias induced by these outliers, the normalization method implemented in DESeq2 [57] was used, which first normalizes counts by the geometric mean for each gene across samples, and then scales each dataset to have a common median (which is less sensitive to outliers). This was applied to all 64 datasets (2 strains X 2 replicates X 16 time-points) in parallel. As a result, the expression patterns were well-calibrated between time-points, with the medians matched.

To identify a subset of genes with meaningful expression, the average expression over all time-points was calculated for each gene and divided by gene length (in nucleotides). 1070 genes out of 4018 with coverage<0.25 were dropped because expression patterns for genes with low expression are inherently noisy, leaving 2948 genes with coverage>0.25. **(Supplementary Table 1)**. Additionally, we removed 127 genes out of 2948 genes whose expression was >90% correlated between *Mtbcos* and *MtbRv* from subsequent analysis, as their expression patterns were assumed to be determined more by time than by difference in the strains. To center the expression values, the counts were divided by the mean for that gene across all the time points. This was done independently for *Mtbcos* and *MtbRv*.

### Gaussian Process Smoothing

In order to meaningfully integrate the data from the two replicates and to smooth out profiles over time, we used a Gaussian Process (GP) to fit the raw data (septation index and ori/ter – Fig1, gene expression-Fig2, Fig3, Fig4).

A GP model is a Bayesian model that estimates the probability distribution over functions using Gaussian distributions for likelihood functions. The advantage of a GP is that it is unbiased and therefore does not require assumptions of form of function. Instead, it only assumes that adjacent time points are better coupled than distant time points and that this correlation is based on Gaussian distributions.

A Gaussian Process is specified by a mean function and a covariance function

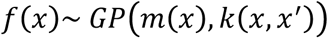

A prior mean m(x)=0 and a covariance function, squared exponential is given as:

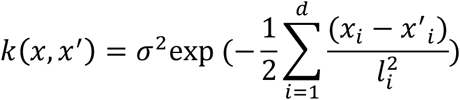

where l^2^ = lengthscale, *σ*^2^ = variance, d = input dimension

We normalized the expression value e(g,t) (with addition of pseudocounts of 10) of each gene g at each time point t by dividing the mean across all time points, and then taking log base *e* transformation so that the normalized value *e’(g,t)* fluctuates with a mean of 0. The formula is given as:

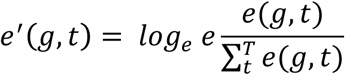

Gaussian estimation of the expression levels for a gene at different time points, subject to noise is given as:

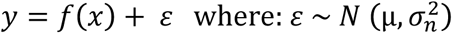

The predictive distribution for 15 test time points (∼ 3 hour intervals, 3-55 hours), {*x*_1_, *x*_2_, … ., *x*_*_} is specified as:

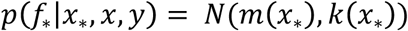

where:

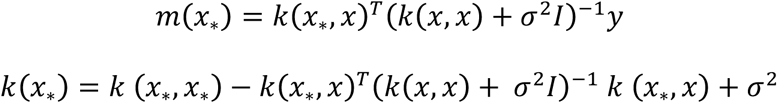

We utilized the GPy Python package to fit the relative expression data (value for cos1 and cos2 simultaneously normalized by the mean expression level across all 60 time points for each gene using the following hyperparameters: variance = 1.0, noise variance = 0.1 and lengthscale (range 1 ∼ 50) optimized to Maximum Likelihood Estimate (MLE) using a grid search method. After fitting the model, the predicted value (i.e. posterior mean) for each time point can be extracted. **Fig S1B** shows the GP regression obtained for *polA* (Rv 1629: DNA polymerase). Not only do the fitted values from the GP model generally interpolate between the observed data at each time point, they also present a smoother profile by averaging between adjacent time points to reduce noise. The error bands show the uncertainty in the model (95% confidence interval which can be denoted as ±1.96*s, where o is the estimated standard deviation at each X-coordinate (time point) from the Gaussian Process model based on variance of the training data and surrounding points).

### Periodicity Analysis

Traditional signal analysis methods like Fourier analysis, Fisher’s g-test, etc. as suggested by Wichert et. al.[58] performed poorly on our dataset because our experiment captured only about one-and-a-half cell-cycles. Thus, to identify periodic genes, we took an approach of sinusoidal curve-fitting, reminiscent of the non-linear curve fitting method described by Straume et.al. (COSIN2NL in COSOPT) [59]. We fit the expression profiles for each gene to a sin curve with free parameters (including frequency, phase, and trend), and selected genes with frequencies and amplitudes in a reasonable range. Goodness-of-fit was measured using residual sum-of-squares (RSS). Importantly, it is difficult to draw an absolute cutoff for significance based on RSS, since any data can be fit to a sin in some way, and RSS incorporates intrinsic noise in the data (E.g. between replicate observations). Hence, we took a comparative approach by also fitting the data to a quadratic curve, which captures the general trend of the expression profiles. We then compared the RSS of the sin fit to the RSS of the quadratic fit (which must also pay a similar price for noise in the same data). Periodic genes are defined as those that exhibit oscillatory behavior above and beyond the trend that can be represented by a quadratic. The curve fitting for each gene was applied to the DESeq-normalized read counts (15 time-points, 2 replicates each). The sinusoidal function implemented is written as:

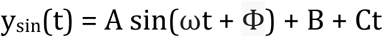

where: A = Amplitude; ω= Frequency; B= Mean offset; Φ= phase shift; Ct= a linear term to capture a net increasing or decreasing trend in the expression. The parameters in this function were optimized using the *curve_fit*() function in SciPy using non-linear least-squares. We then selected genes based on period length (27.5 hours< period < 55 hours) and amplitude (≥ 0.7). We also removed genes with a correlation coefficient of > 0.9 between expression profiles in Cos versus Rv. The residual sum-of-squares (goodness-of-fit) was calculated as follows:

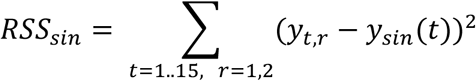

where y_sin_ are the sin function estimates for each time point.

A similar curve-fitting approach was used to fit the data to a quadratic curve:

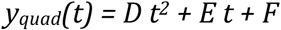

using *curve_fit*() to optimize the parameters D, E, and F for each gene, and the residual was calculated as:

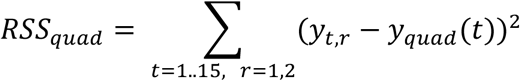

Finally, a score was calculated for each gene based on the ratio of residuals. To meaningfully enrich periodic genes in *Mtbcos,* we used a Receiver Operating Characteristic (ROC) curve to determine the RSS*_sin_*/RSS*_quad_* range where we optimally enrich for periodic genes in *Mtbcos.* The RSS*_sin_*/RSS*_quad_* range was determined to be 0.35-0.45. A threshold of 0.45 was chosen based on examining plots (**Fig S3, S4**) that visually exhibit clear oscillatory behavior (beyond the general trend). Thus, genes with a ratio of less than 0.45 were identified as periodic:

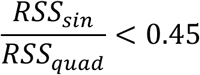

which means that the sinusoidal fit reduces the residual error by more than two-fold over a quadratic curve and hence fits the data better.

Using this comparative curve-fitting approach, 485 genes were identified as periodic in *Mtbcos* (**Supplementary Table 2)**, and only 183 genes in *MtbRv*, a ∼2.6 fold enrichment. To estimate the number of false positives in the set of 485 genes, we randomized the data (by shuffling the genes and timepoints) and subjected the randomized dataset to the same analysis as described above. This permutation analysis yielded only 14 periodic genes under the null hypothesis. Thus, we estimate the false discovery rate (FDR) at approximately 14/4019 = 0.35%.

### Clustering

Genes were clustered based on their expression profiles using hierarchical clustering (*hclust()* in R), using the ward.D2 method [60] based on the Euclidean distance between the vectors of expression values averaged between replicates over the 15 time points, which were standard-normalized for each gene (subtract mean and divide by standard deviation) to make the mean expression level equal to zero for each. The optimal number of clusters was determined based on the Bayesian Information Criterion (BIC) using *mclustBIC*() in the *mclust* R package [61], which showed that the optimal number of clusters among the 485 *Mtbcos* periodic genes was 8 (using the ‘VEE’ model). The dendogram was then divided into 8 disjoint clusters using *cutree()*.

### Peak Assignment

Using the GP fit data, we applied the following criteria to assign a peak to a gene’s expression profile. The time series *T* with *n* observations for each gene with smoothed expression values {*x*_1_, *x*_2_, … ., *x_n_*} at different time points {*t*_1_, *t*_2_, … . , *t_n_*} was defined as:

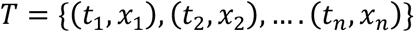

First, to screen out the increasing or decreasing trend at the beginning and end of the time series, and to focus on the cytokinesis phase in the middle of the time course, we excluded the first and last two time points from the peak assignment. Second, to identify well-spaced major peaks across time points, we defined a point *x_i_* as a peak if it has a greater magnitude than its two nearest neighbors on both sides. This is defined as:

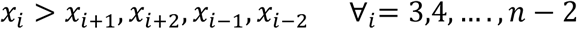

Furthermore, to filter out the genes with lower fluctuations, the difference between the magnitude of the highest peak *x_h_* and the global minimum *g_min_* was restricted to be greater than 0.5. Additionally, in the case of more than one peak in the time series, all the peaks were constrained to have at least a half magnitude of the highest peak in the expression profile. Finally, a set of peaks *P* for a time series was identified as:

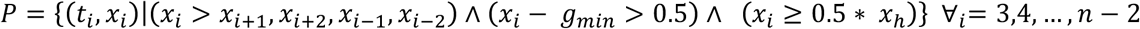

Among the significantly expressed genes, the peak assignment identified 1620 genes with a single peak and 71 genes with two peaks in the *Mtbcos* strain compared to 903 genes with a single peak and 8 genes with two peaks in *MtbRv*. Similarly, 1222 genes in the cos strain and 2344 genes in the wild type did not have any major peak. This once again confirmed that the gene expression levels in the *Mtbcos* strain show significantly higher fluctuations than *MtbRv*.

## Supplemental figure legends

**Fig S1A:**
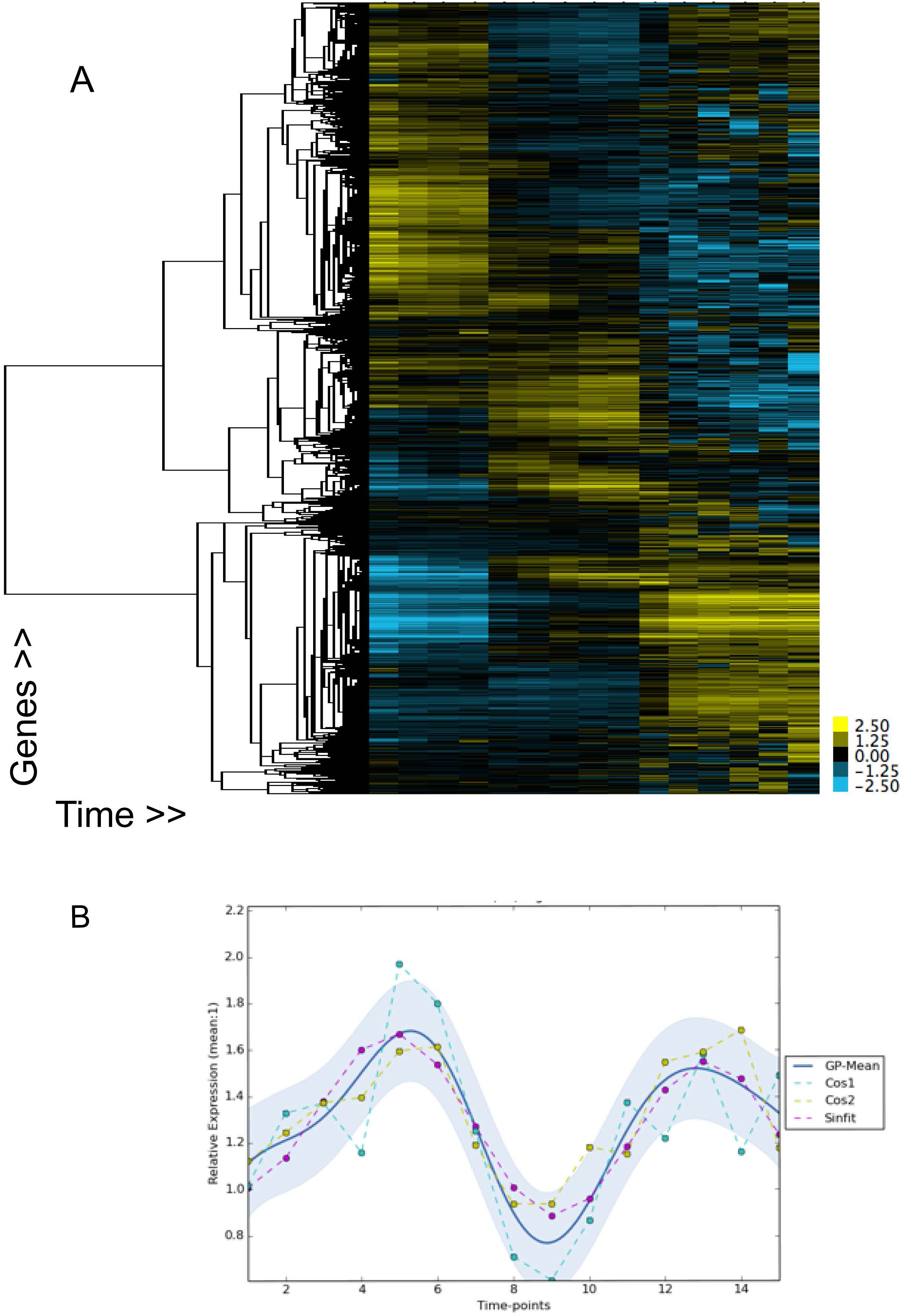
Hierarchical clustering of relative expression of genes (rows) in *Mtbcos* (log transformed DESeq normalized counts, centered around the mean, similarity metric: centered correlation, clustering method: centroid linkage). Columns - hours at 37°C. FigS1B: Expression profile of the DNA polymerase *polA*. Data from two replicates (yellow and blue dashed lines), GP fit (solid blue line) and sinusoidal fit (pink dotted line) are shown. Y axis: Relative expression (DESeq normalized value for each time point divided by the mean polA expression value across all time points.

**Fig S2:**
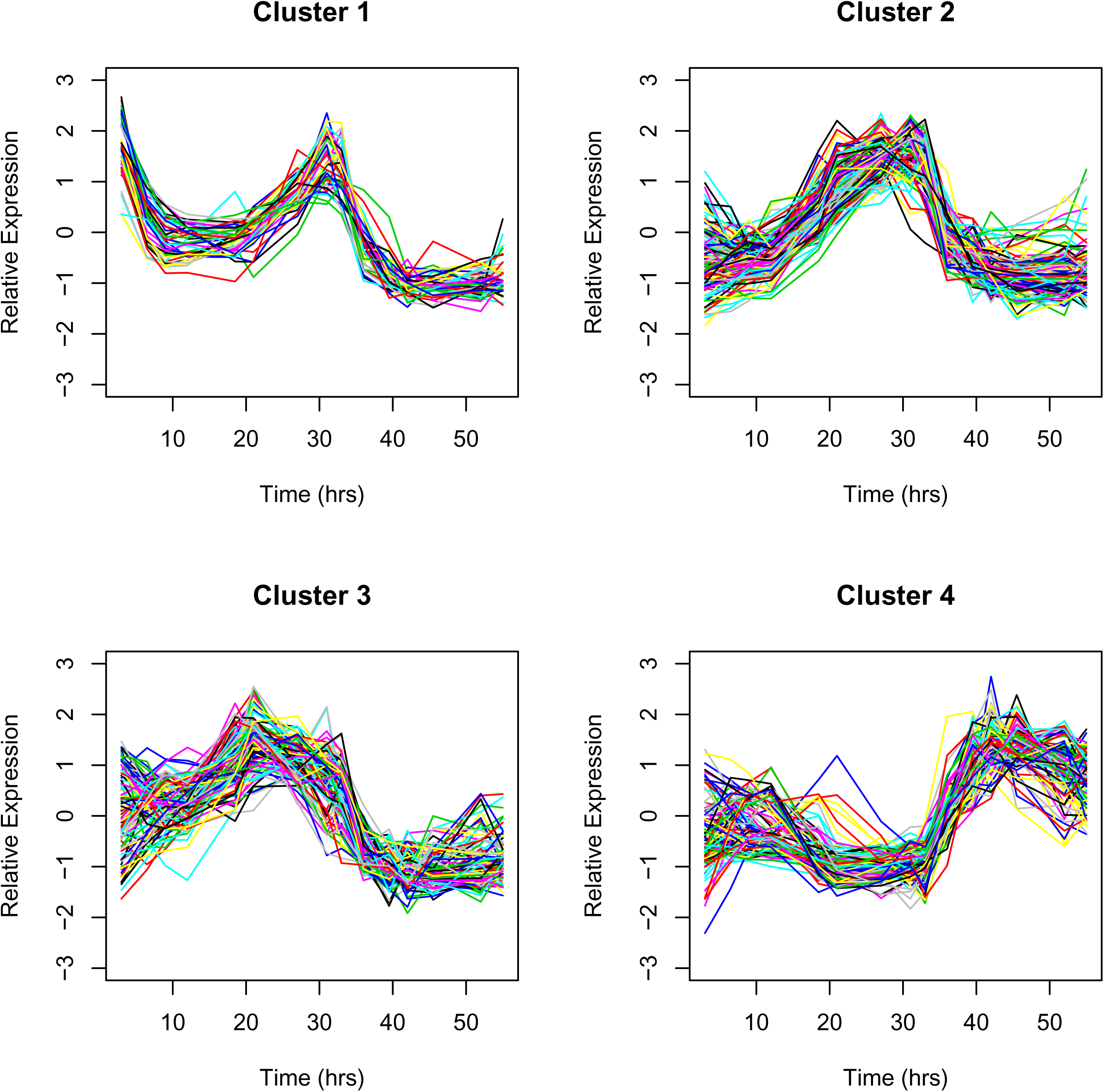
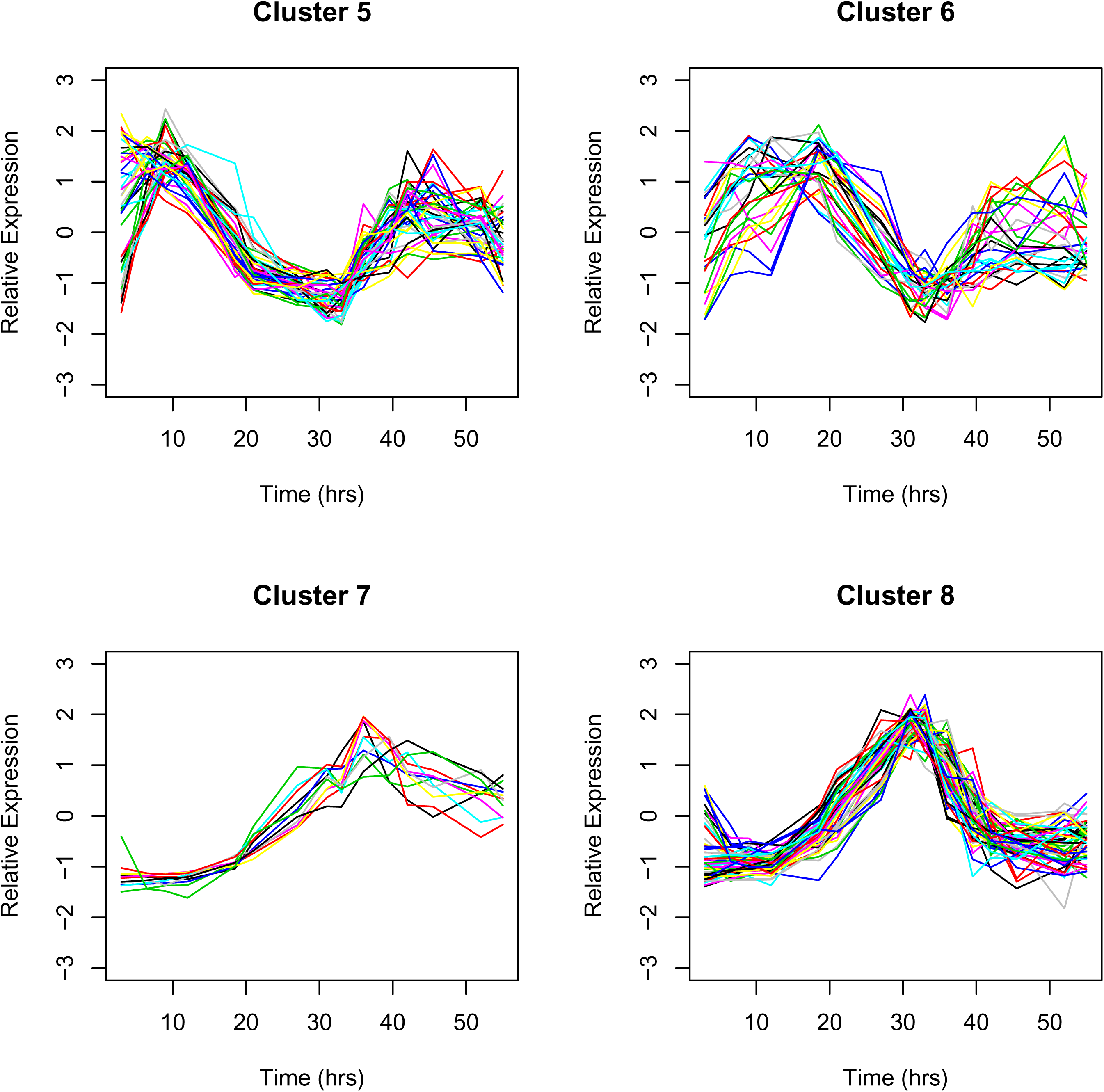
Eight clusters containing the 485 periodic genes in *Mtbcos* (Distance matrix: Euclidean, Clustering method: Hierarchical).

**Fig S3:**
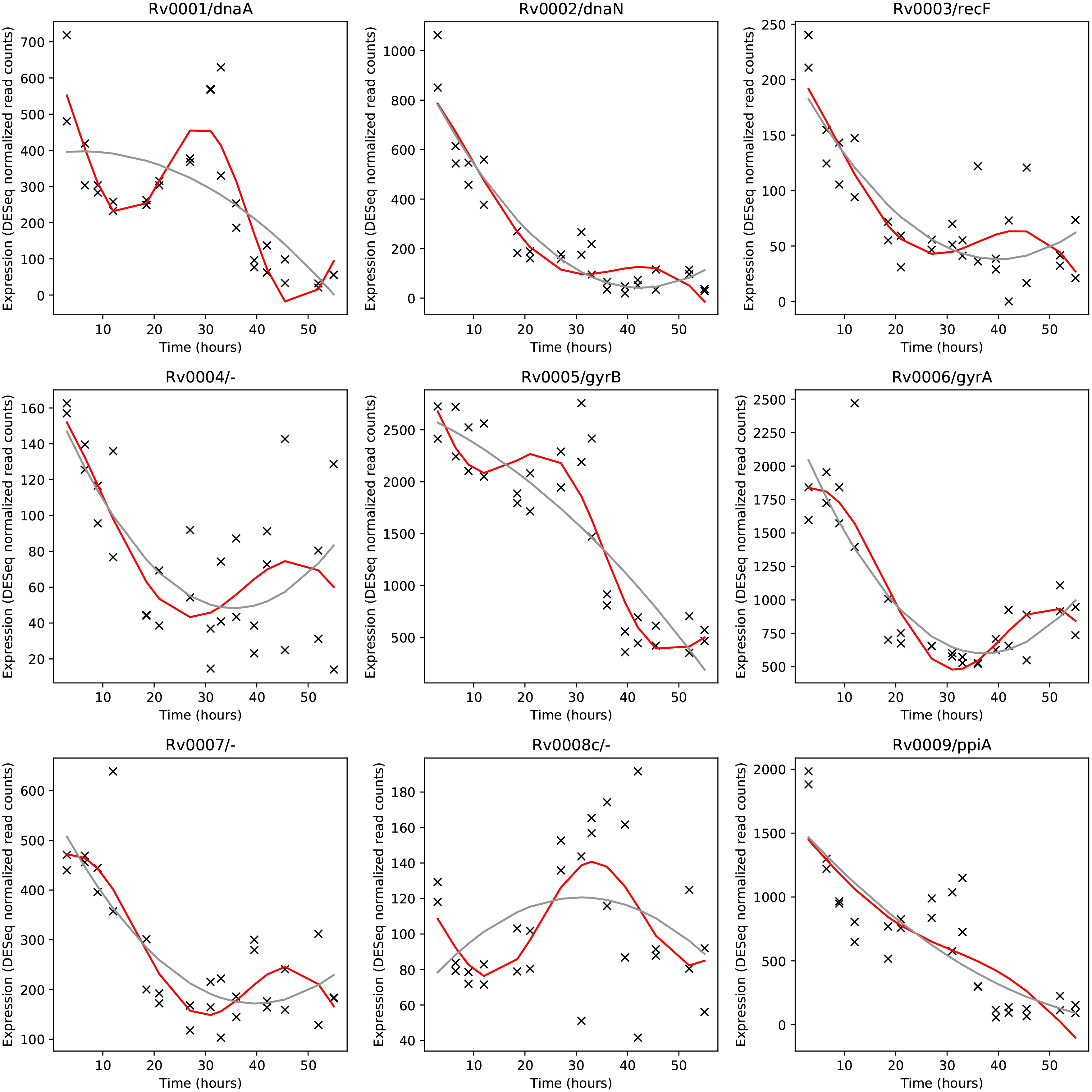
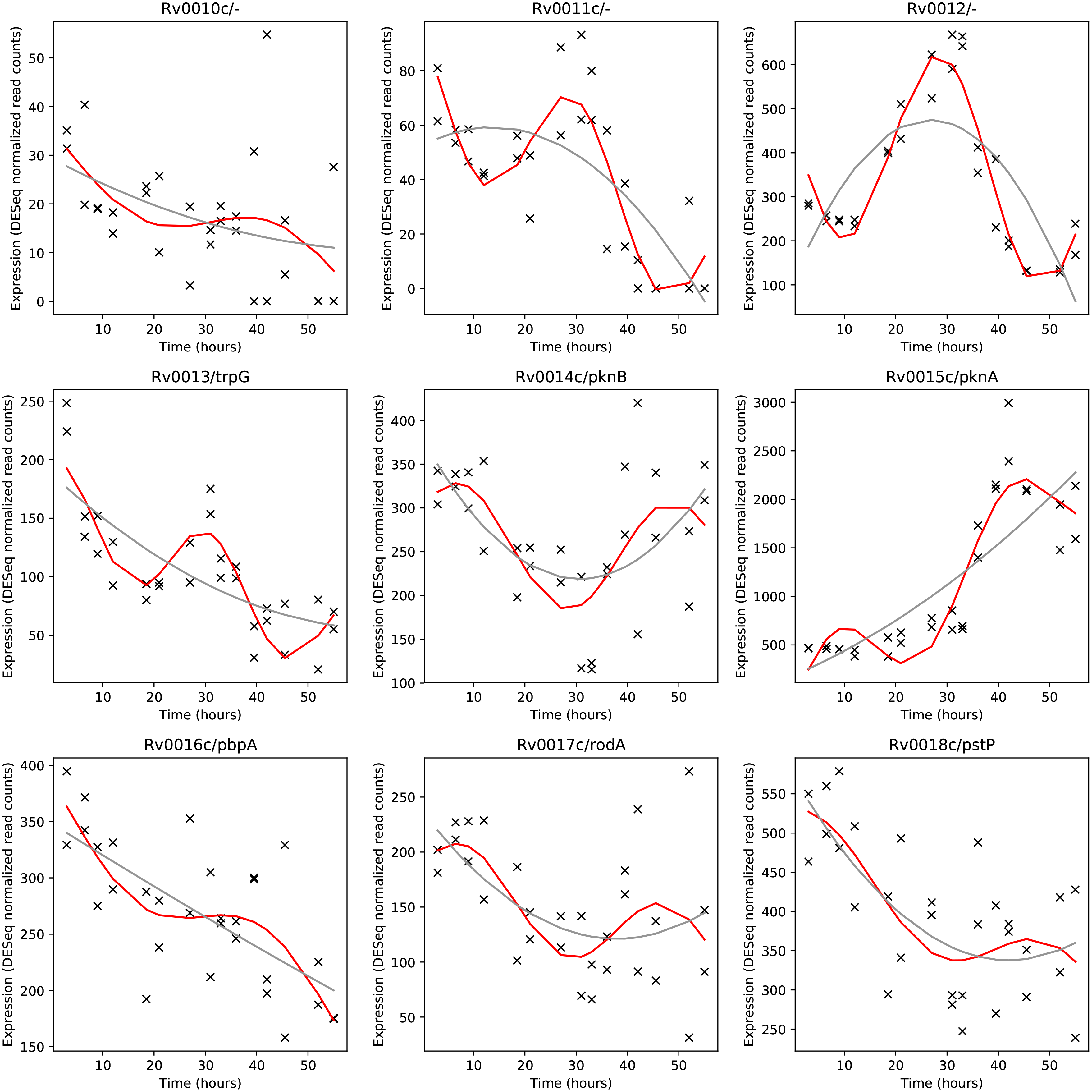
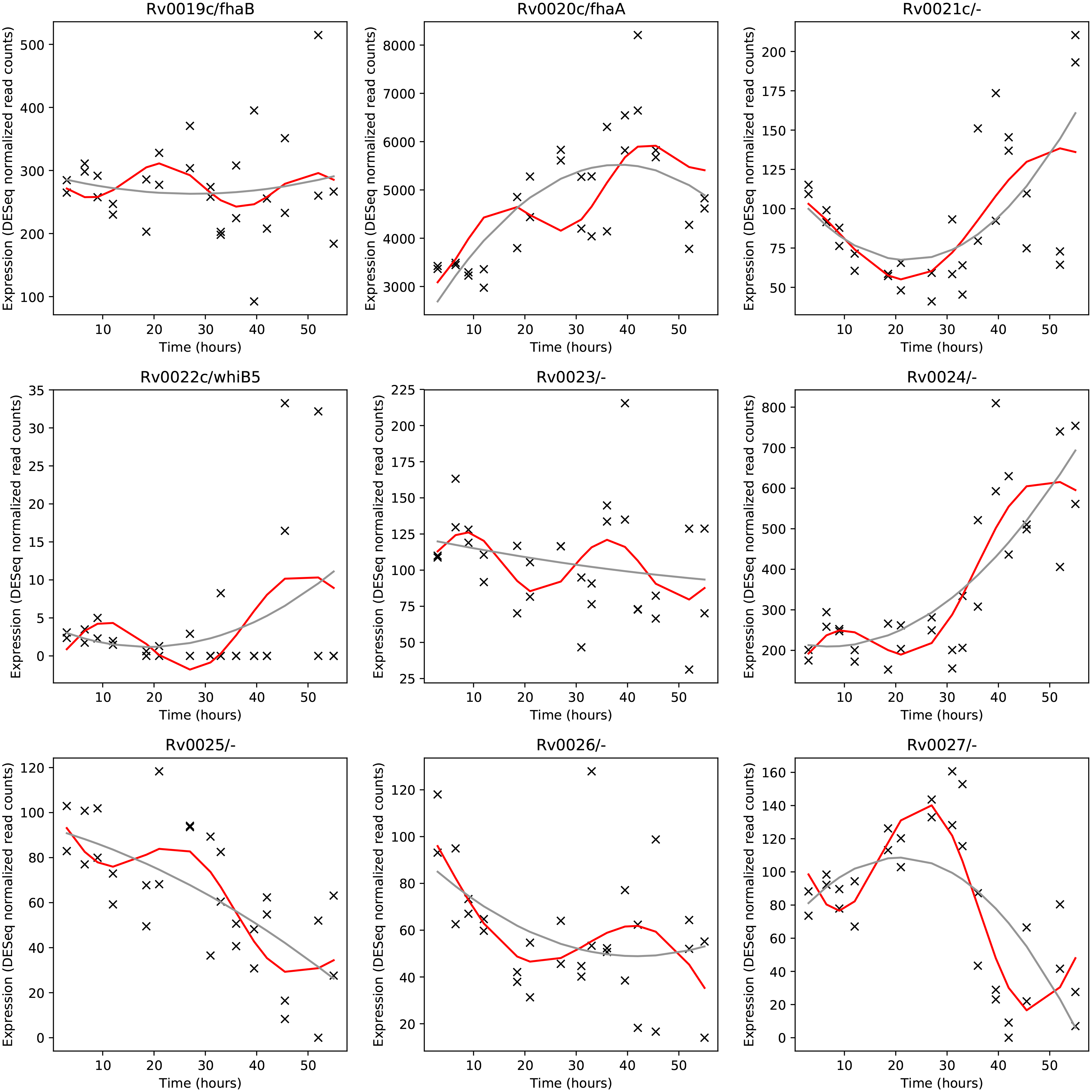
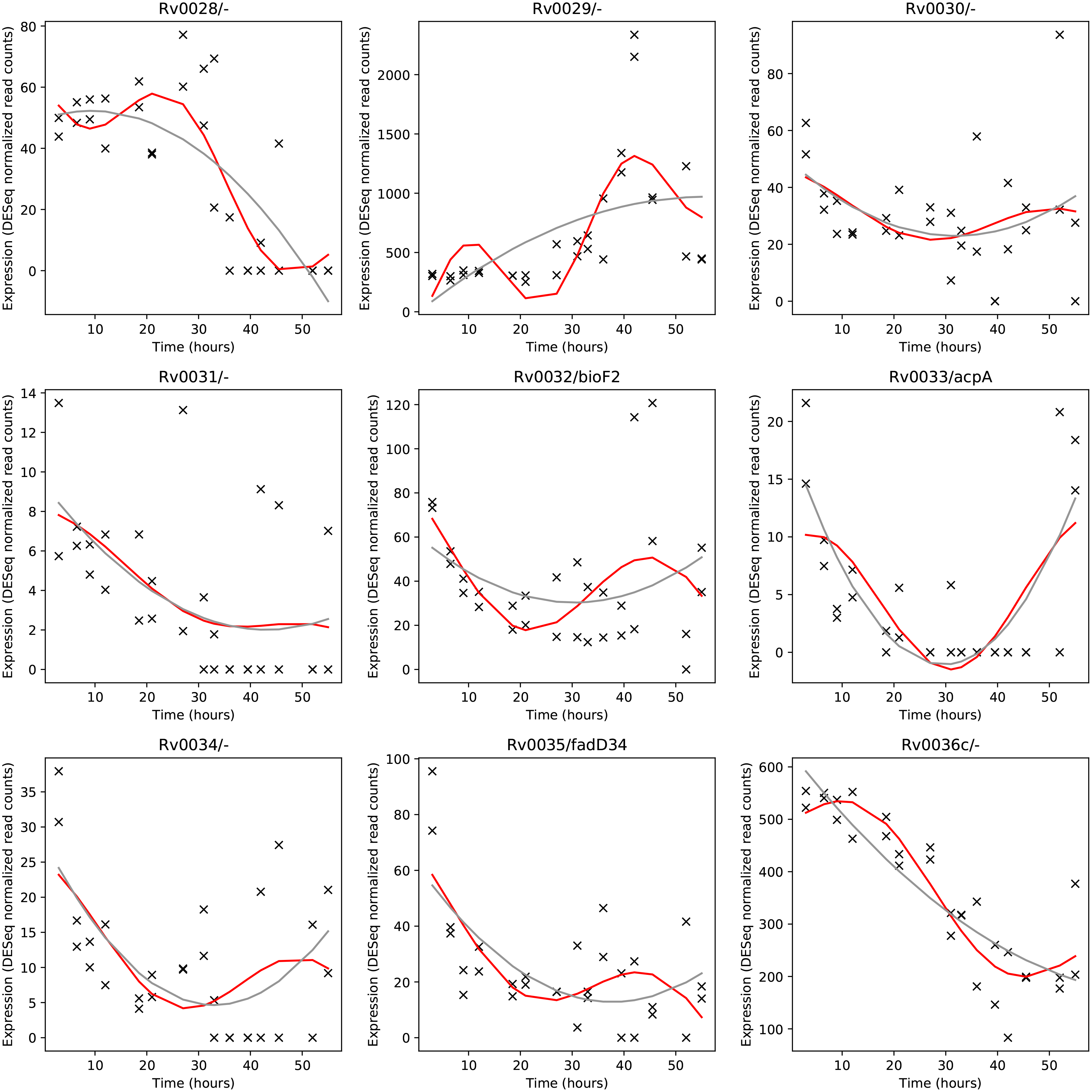
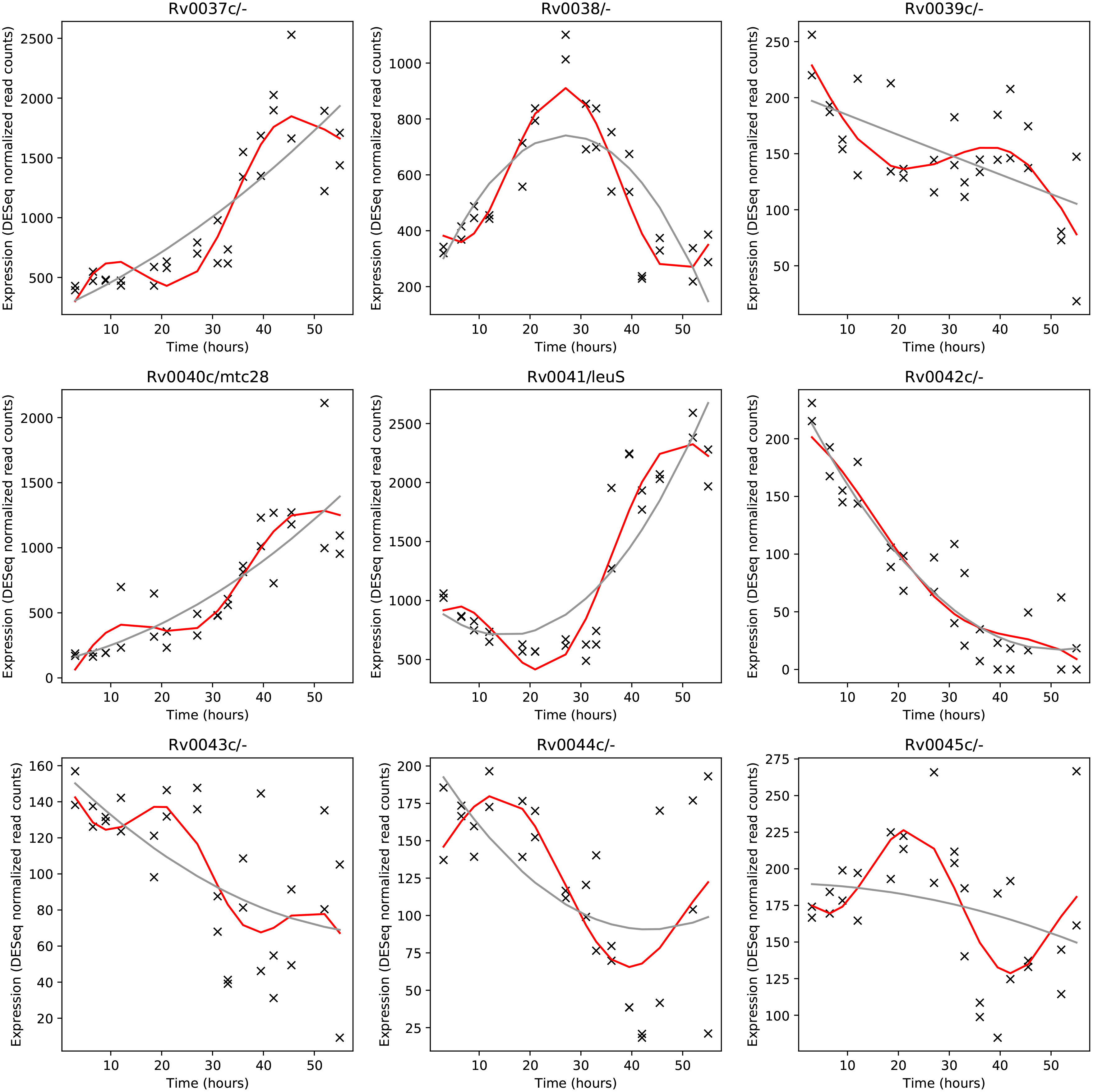
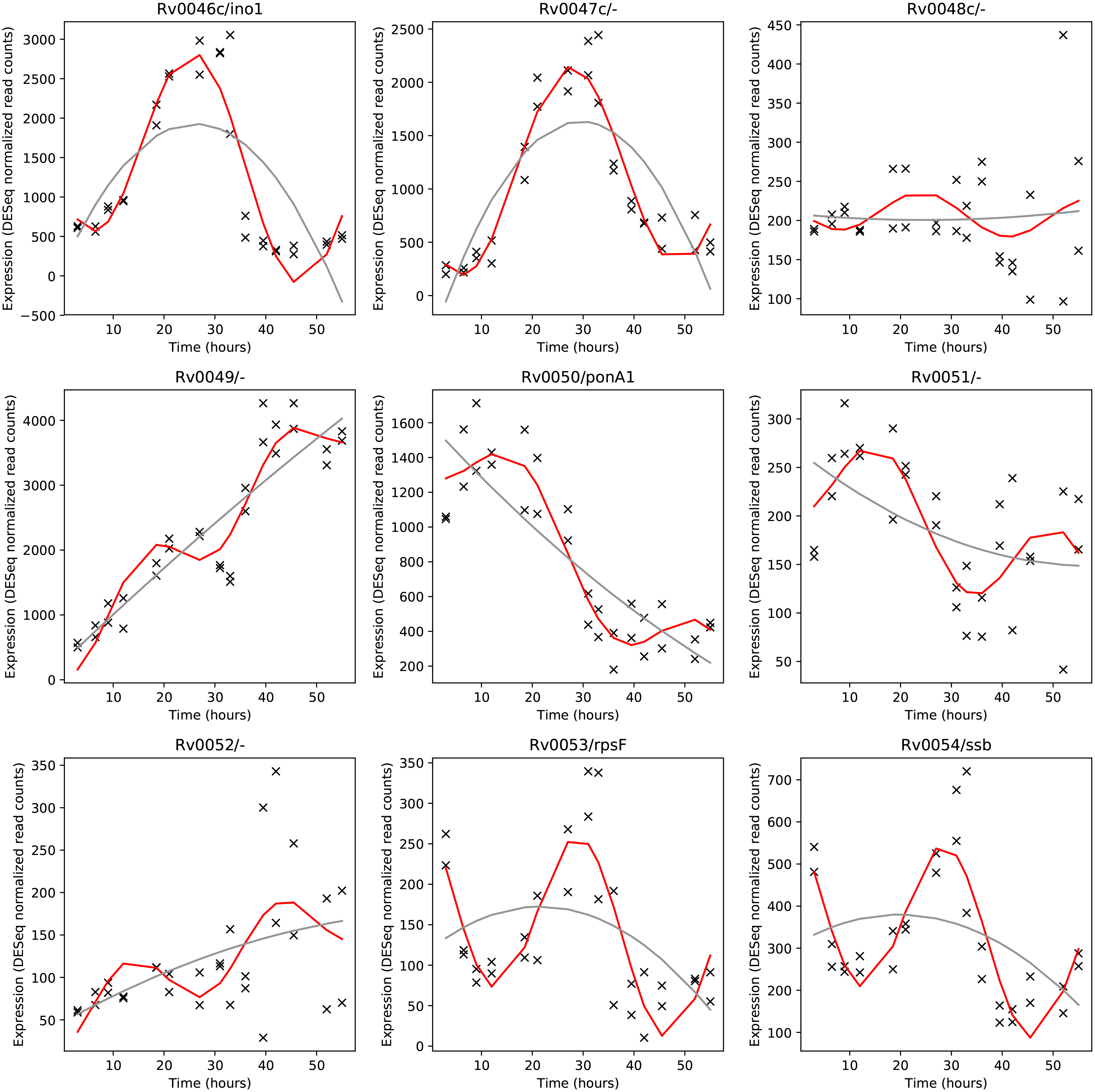
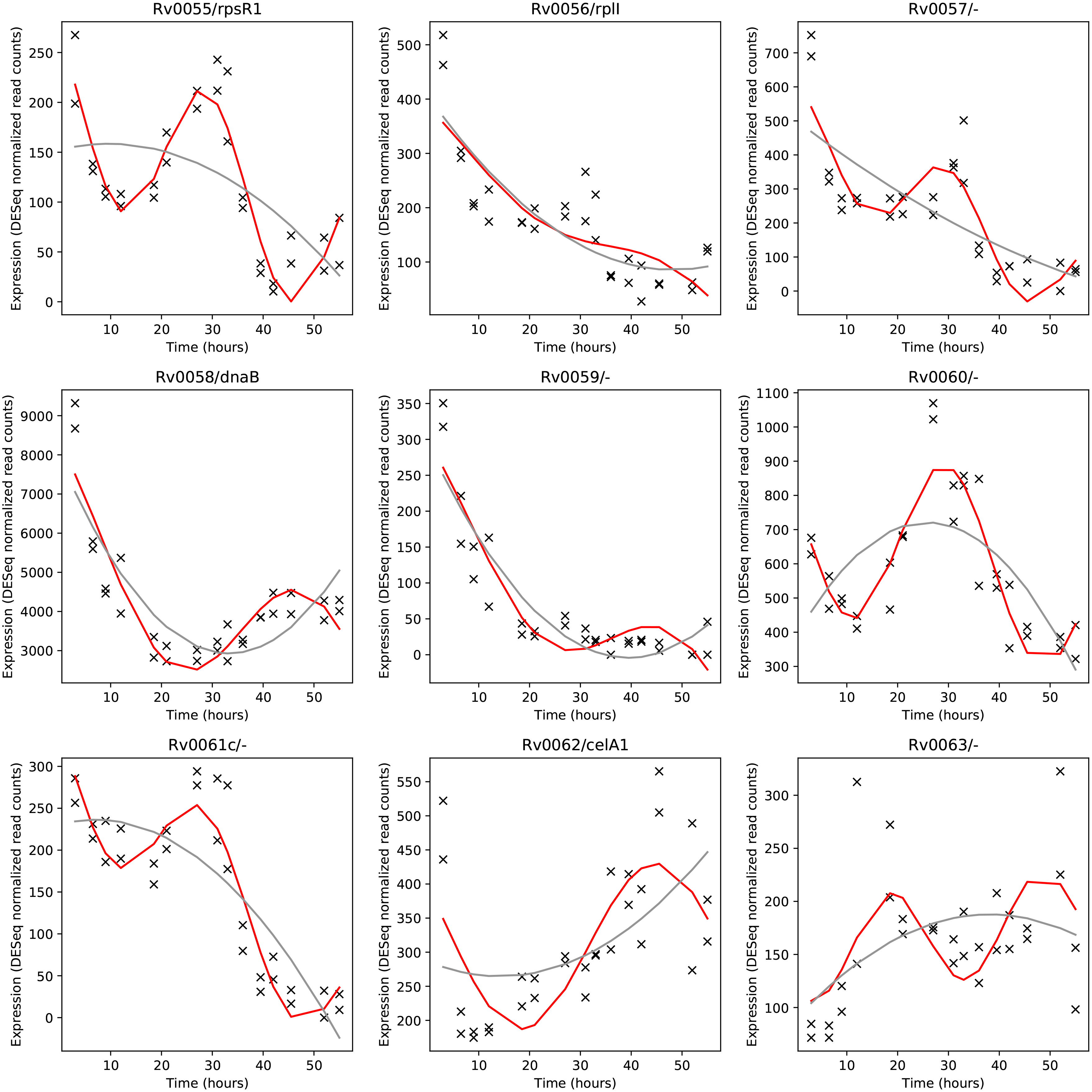
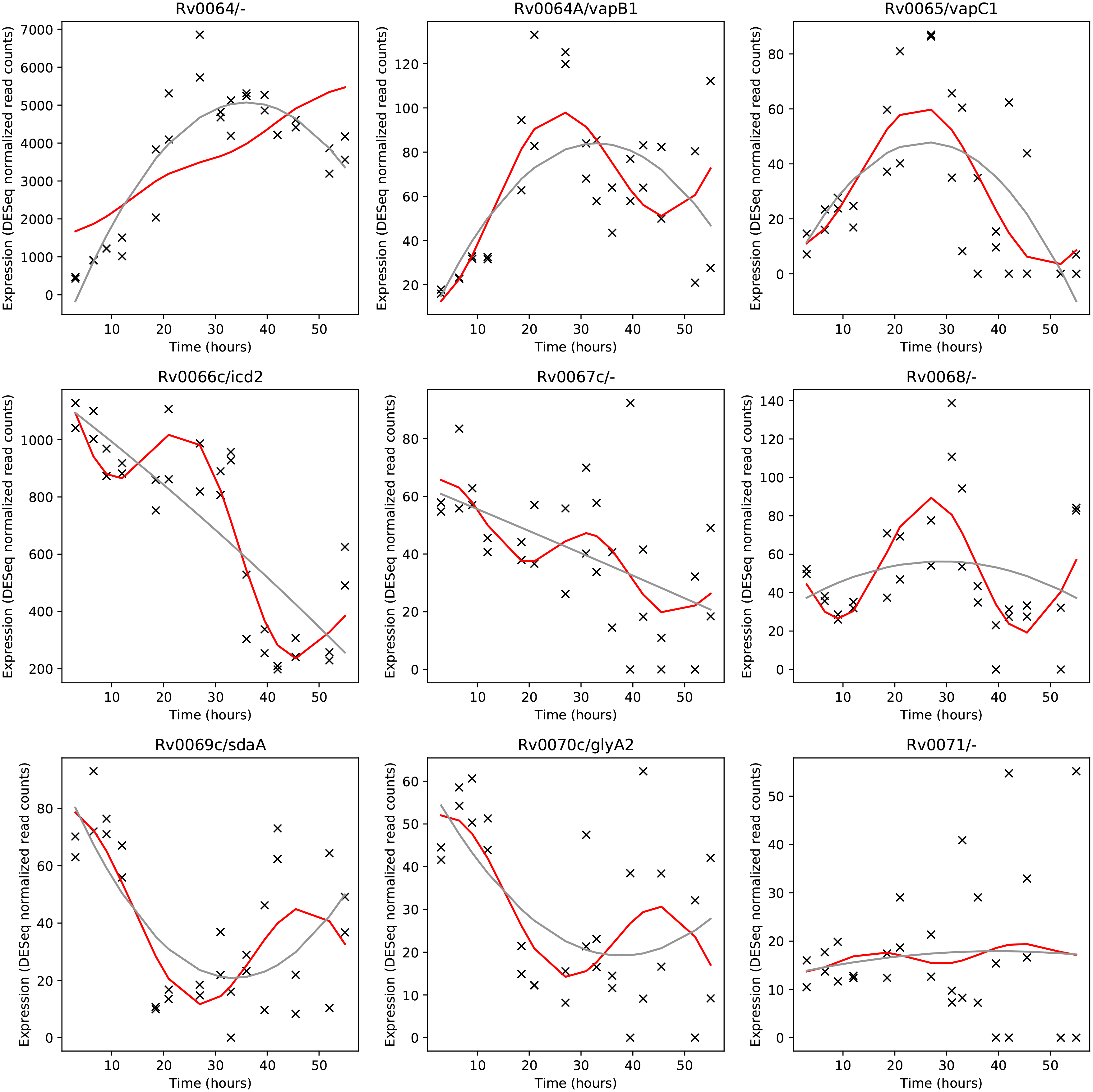
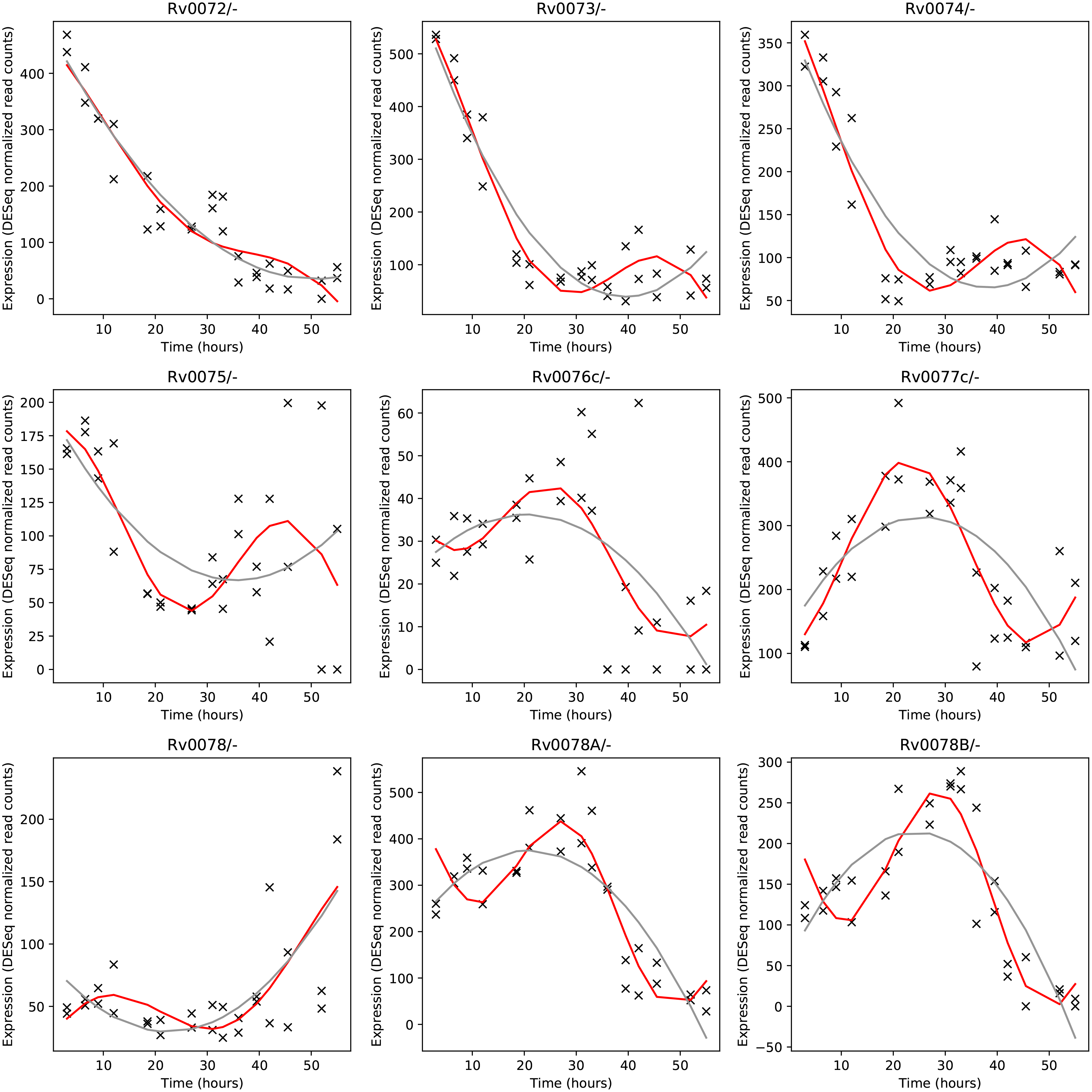
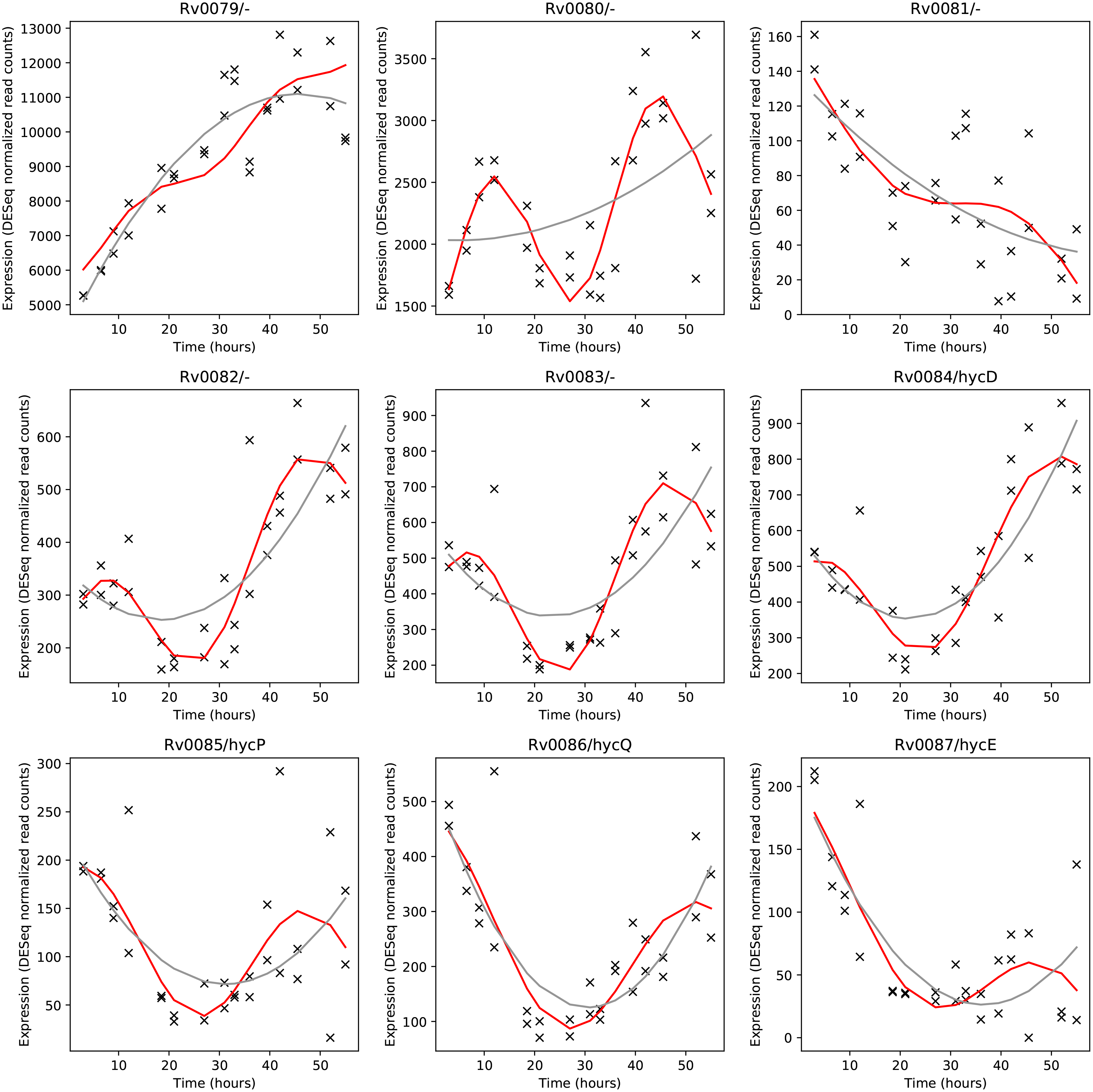
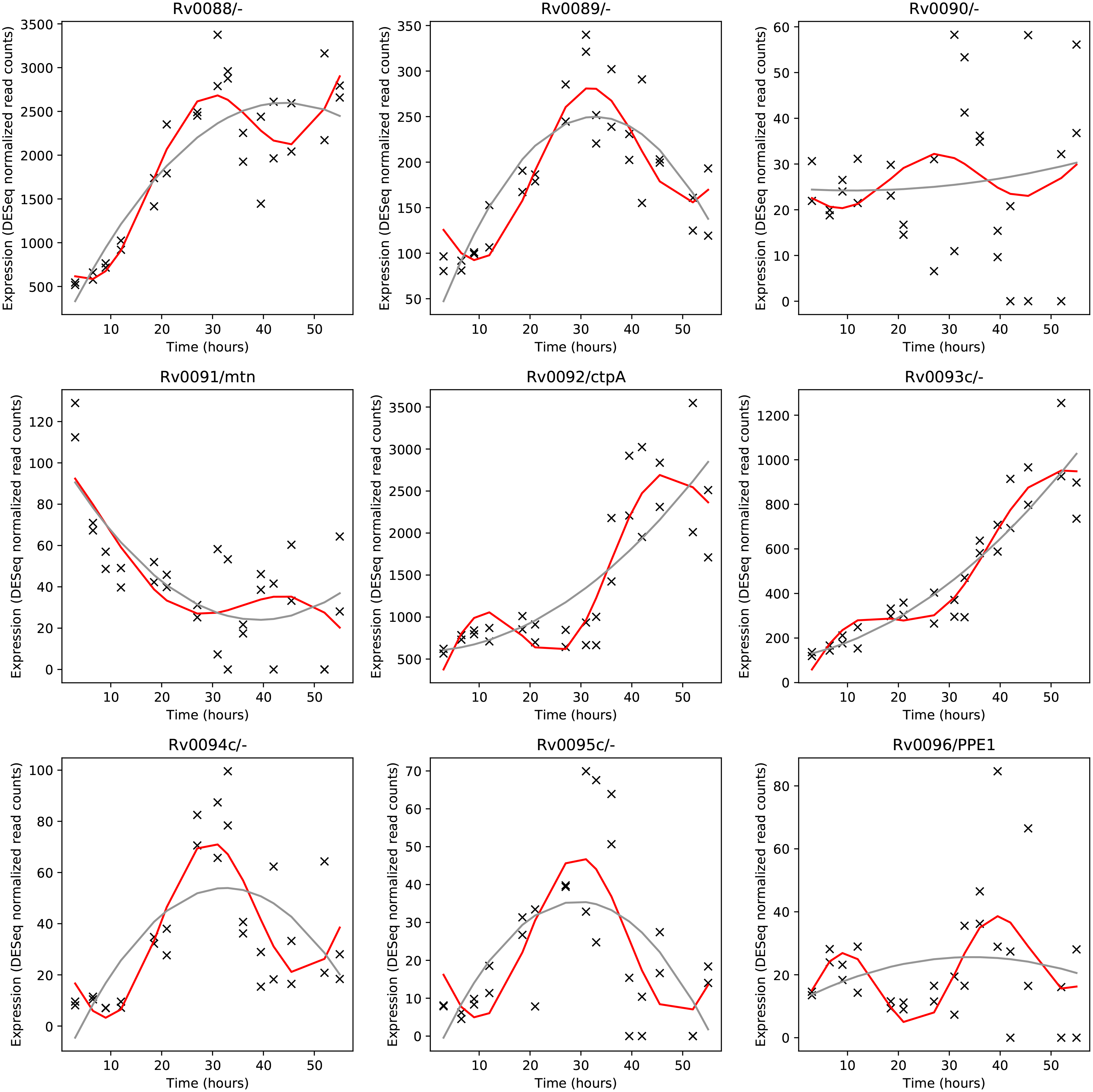
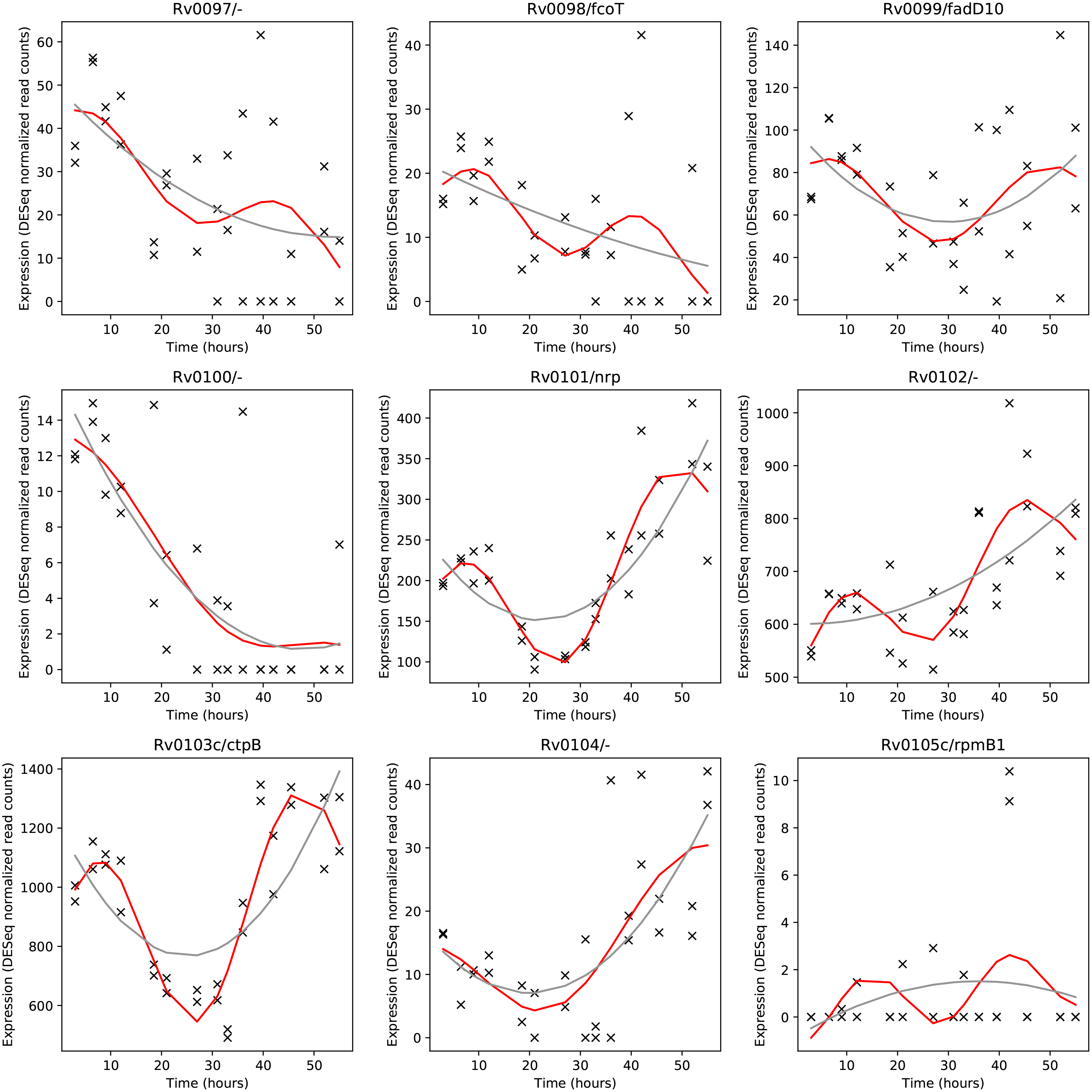
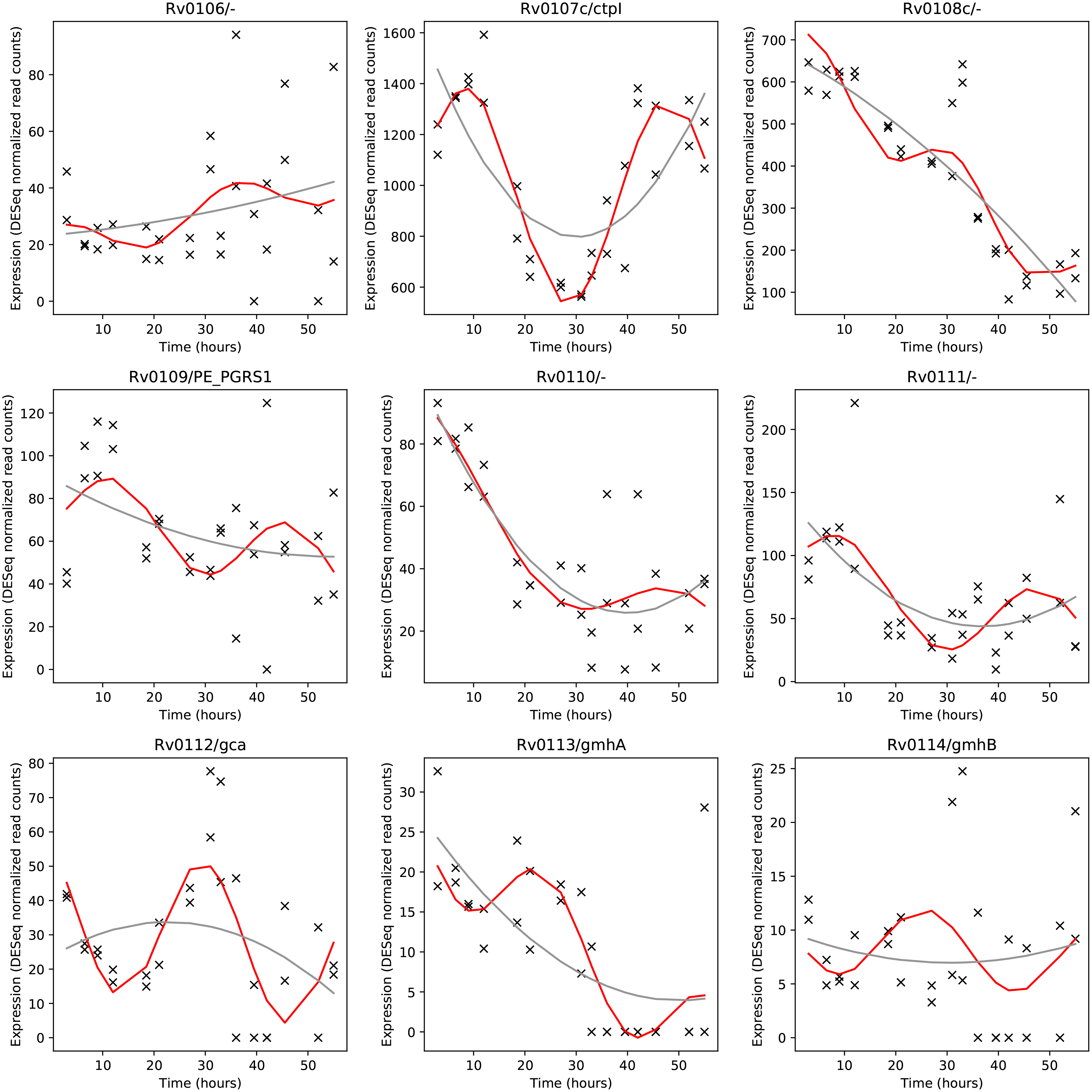
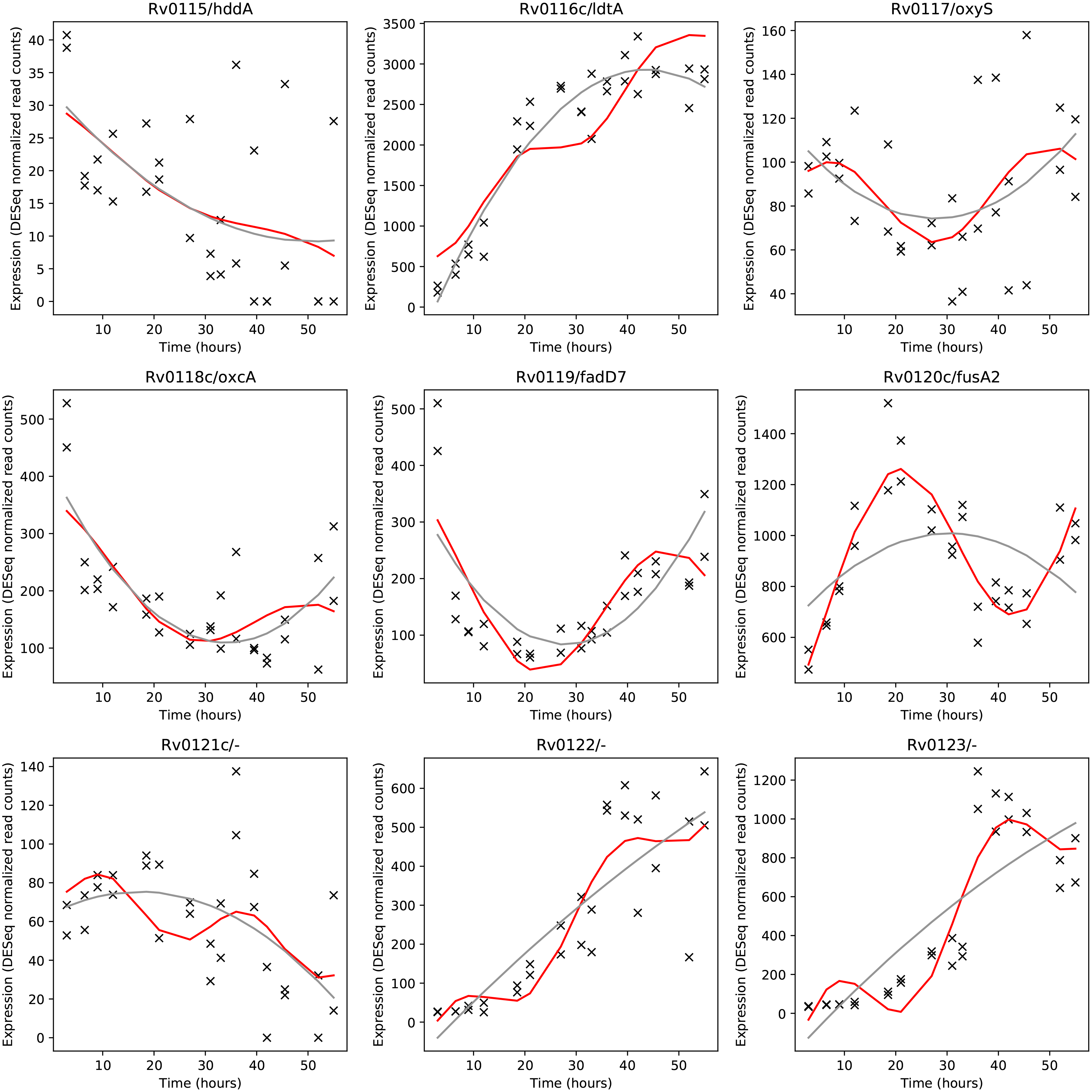
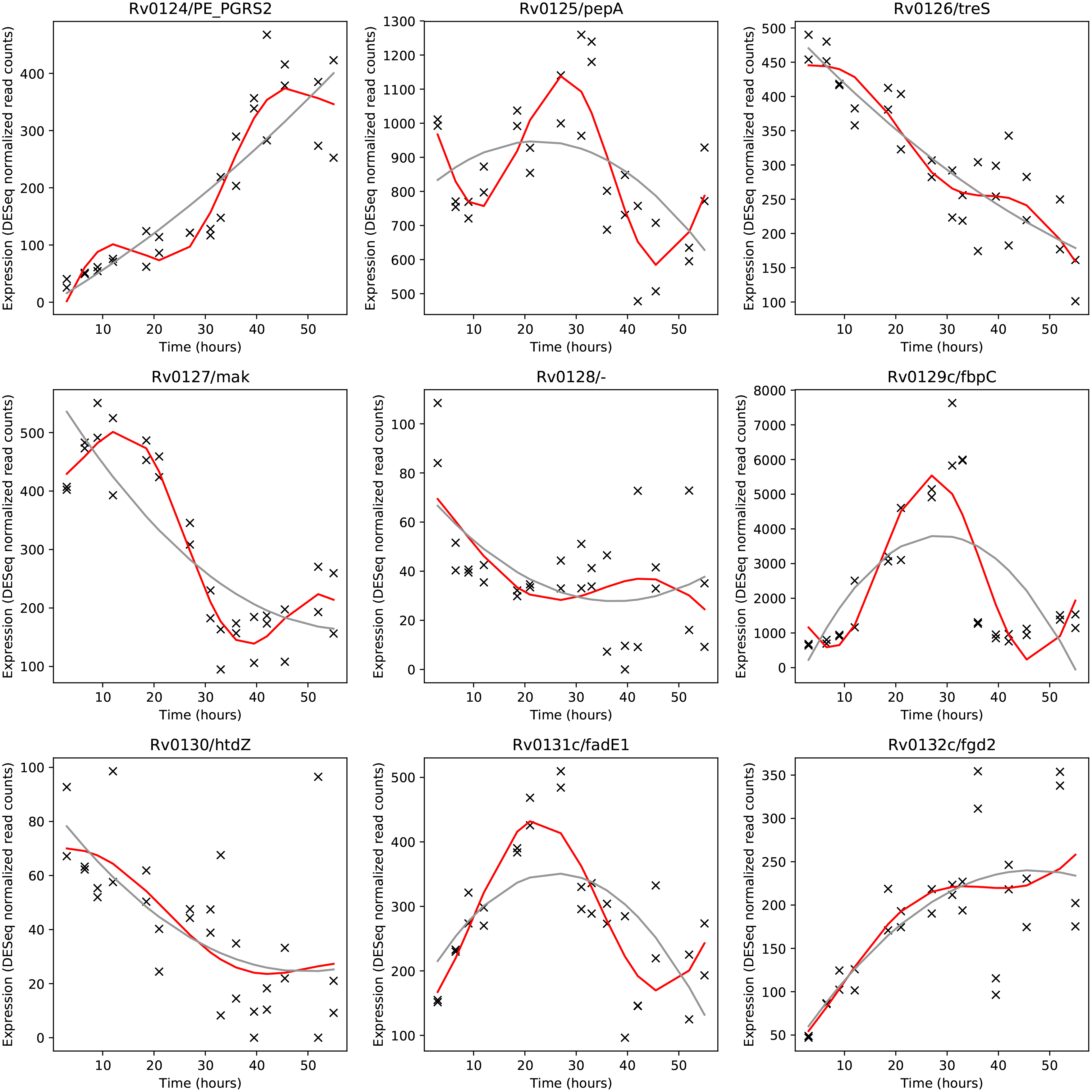
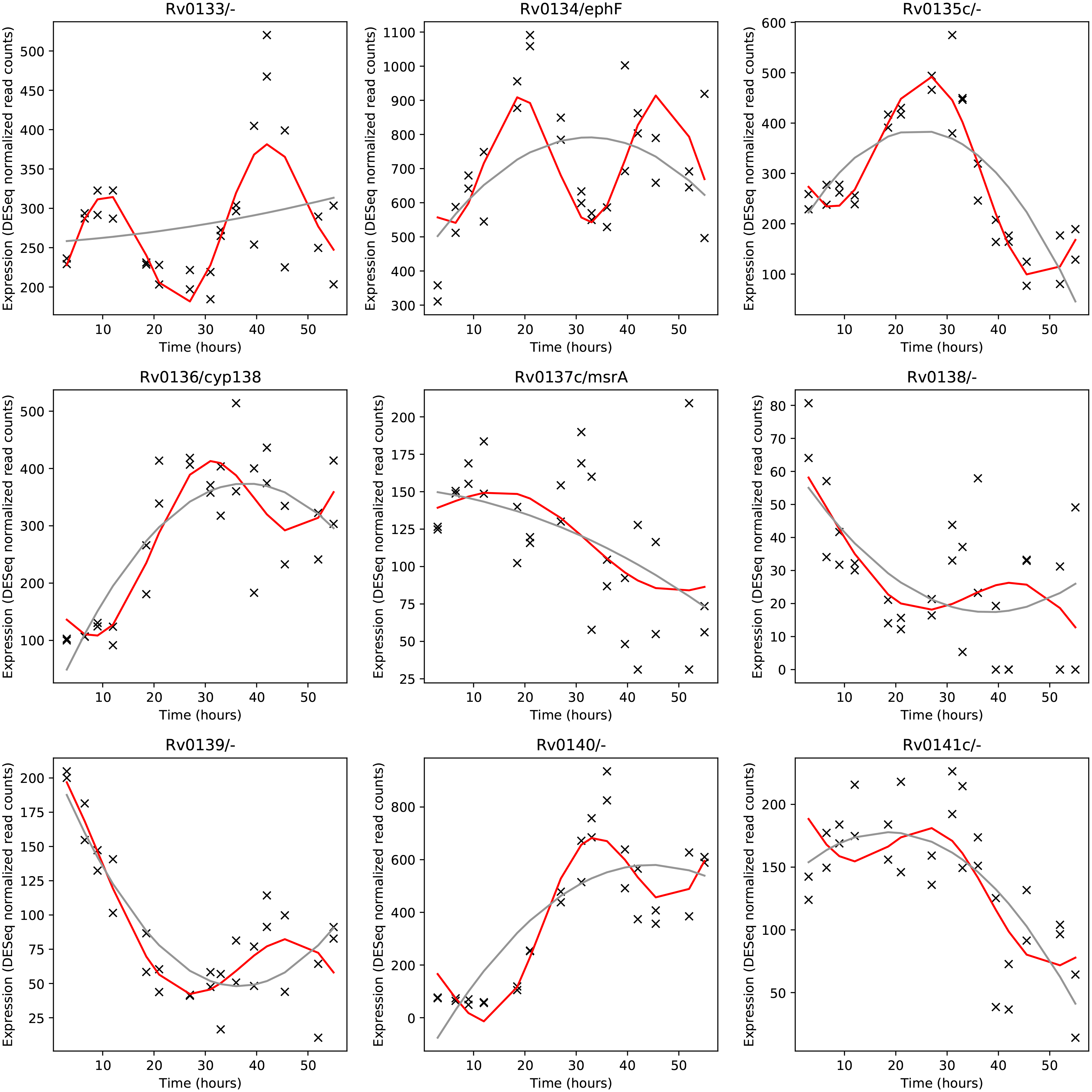
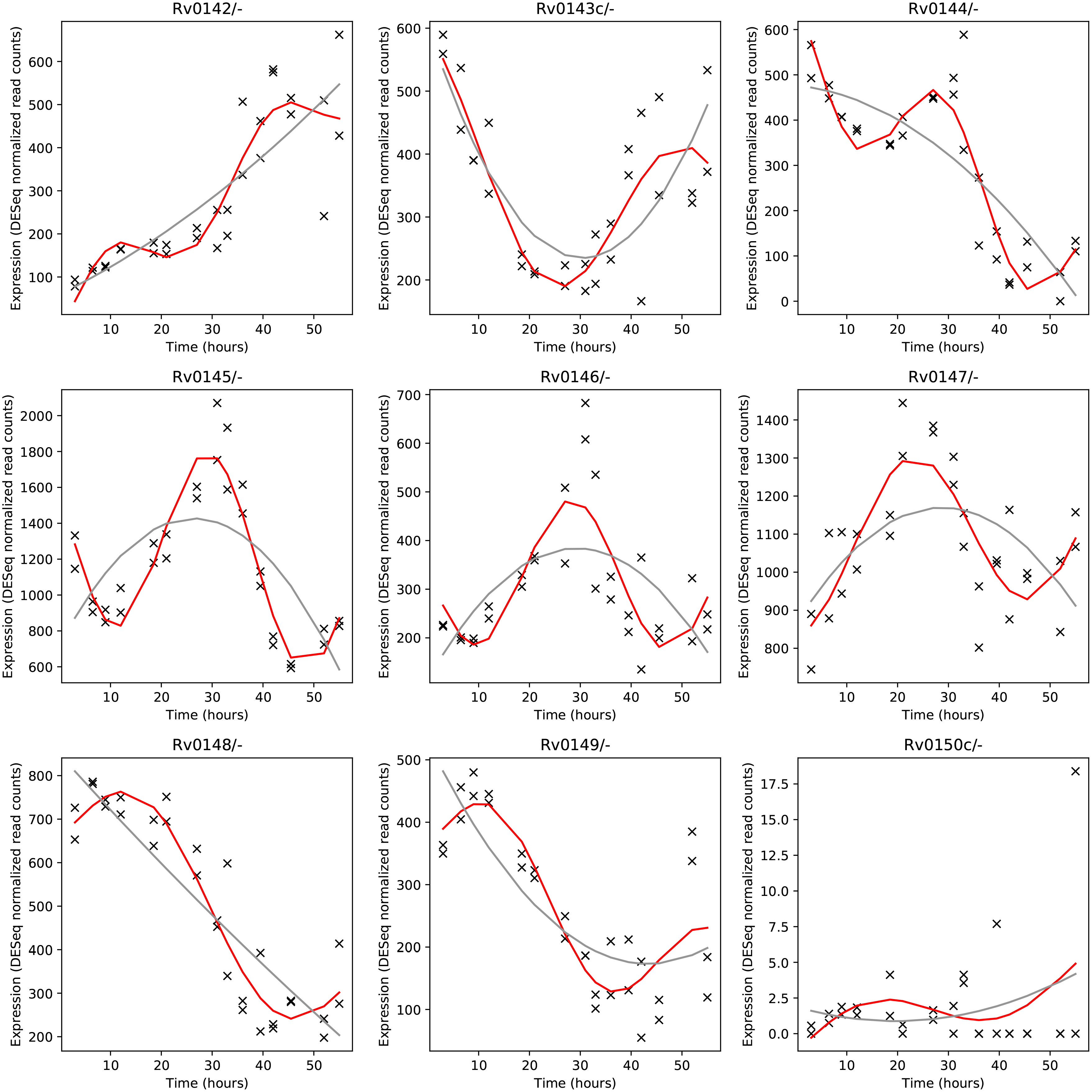
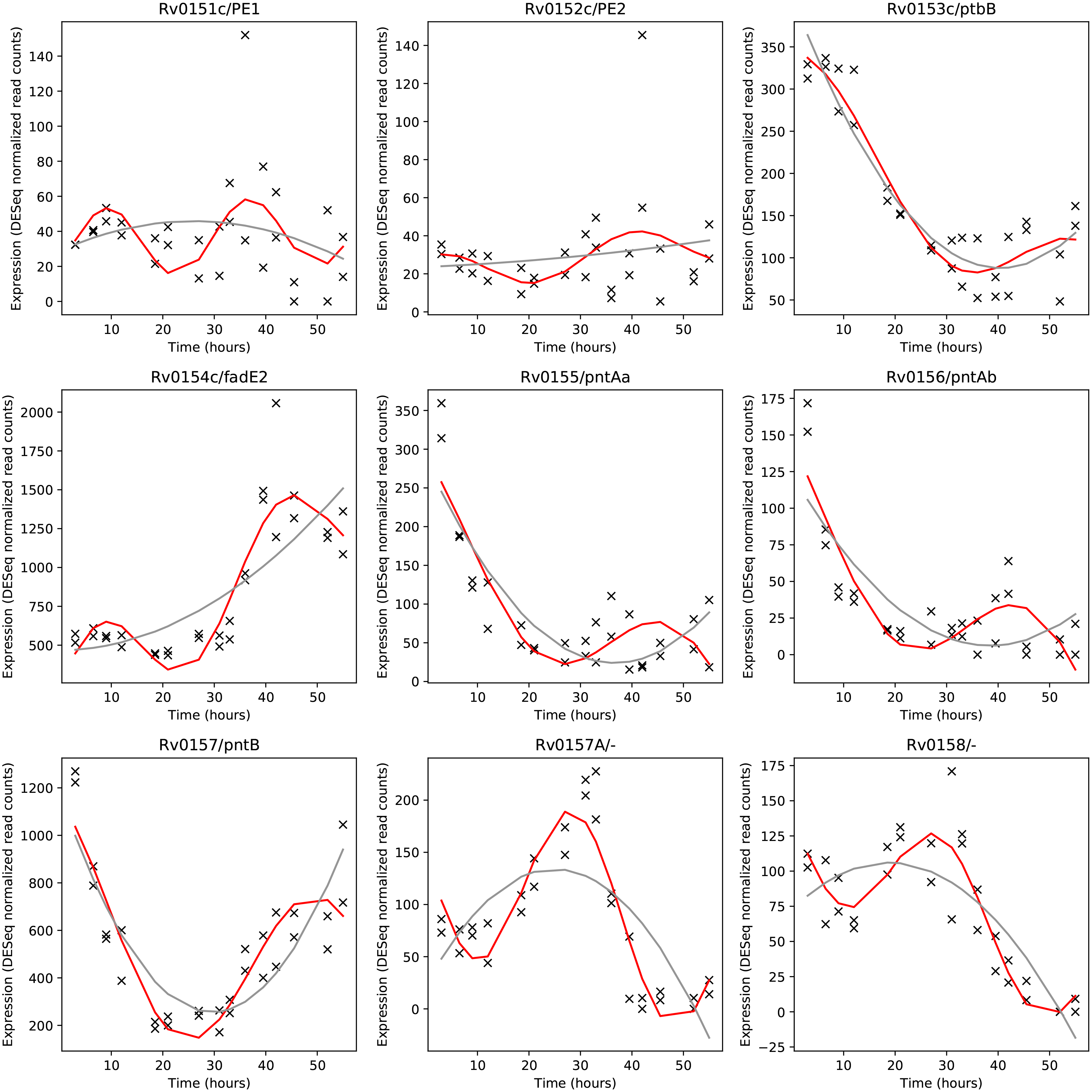
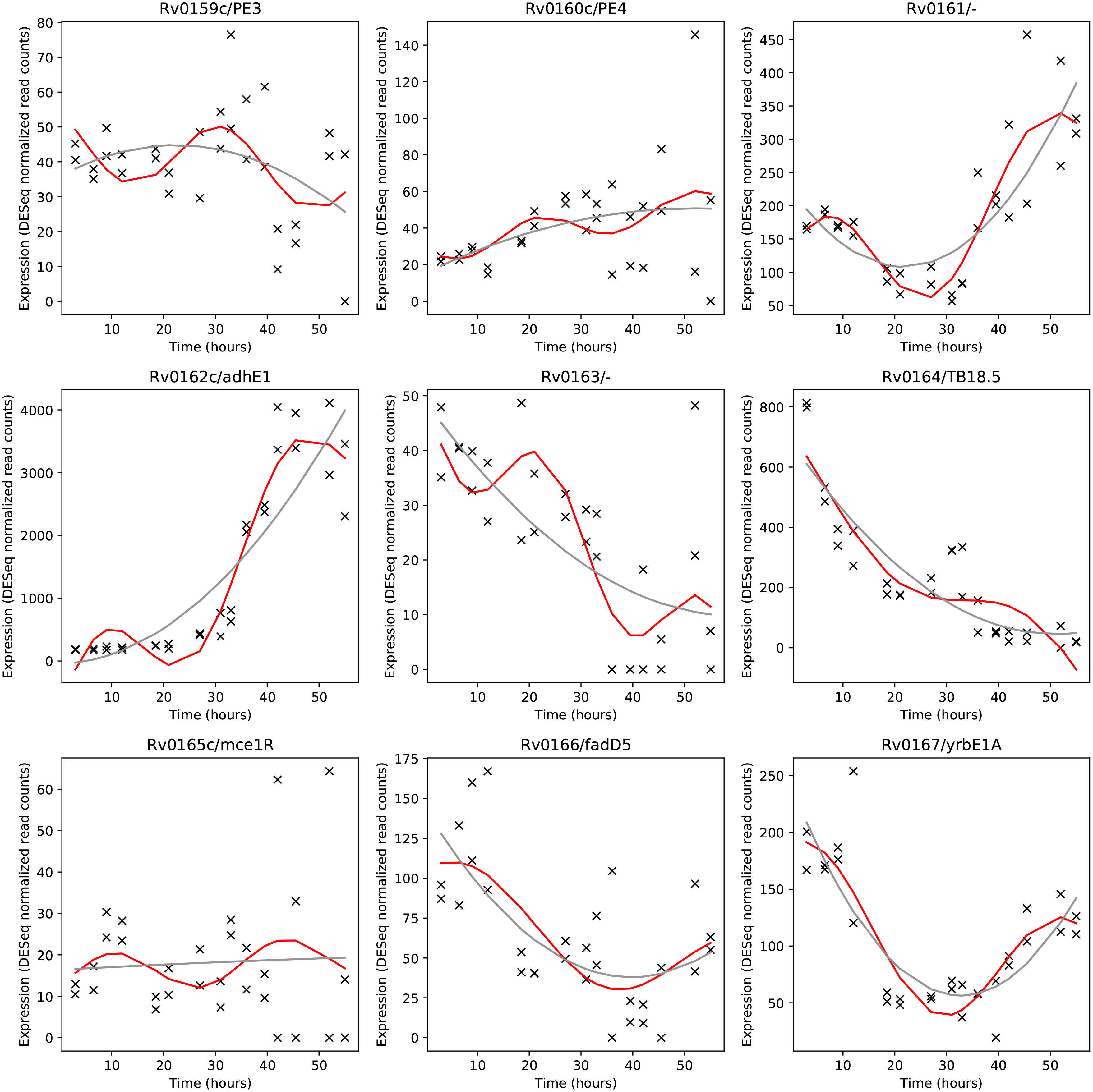
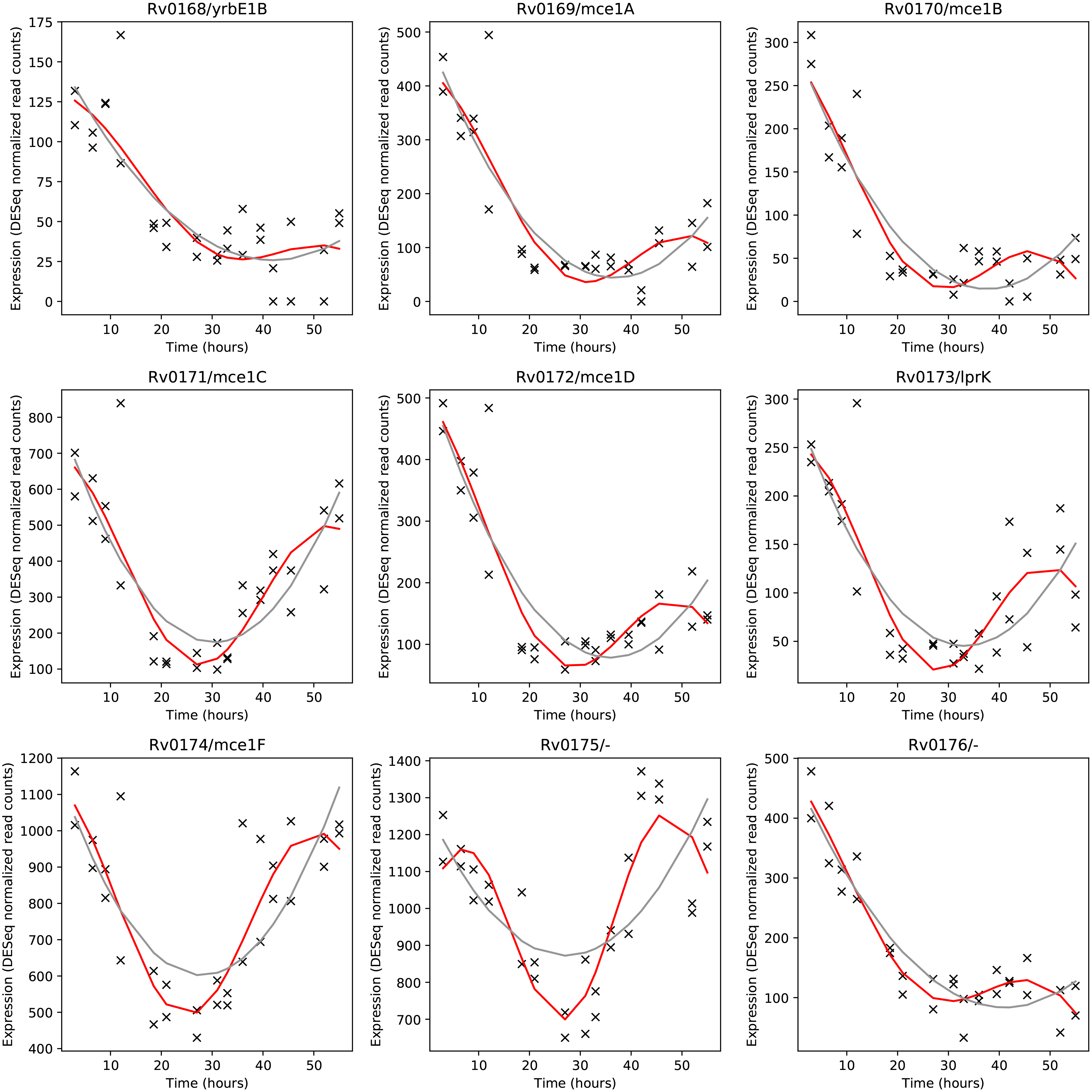
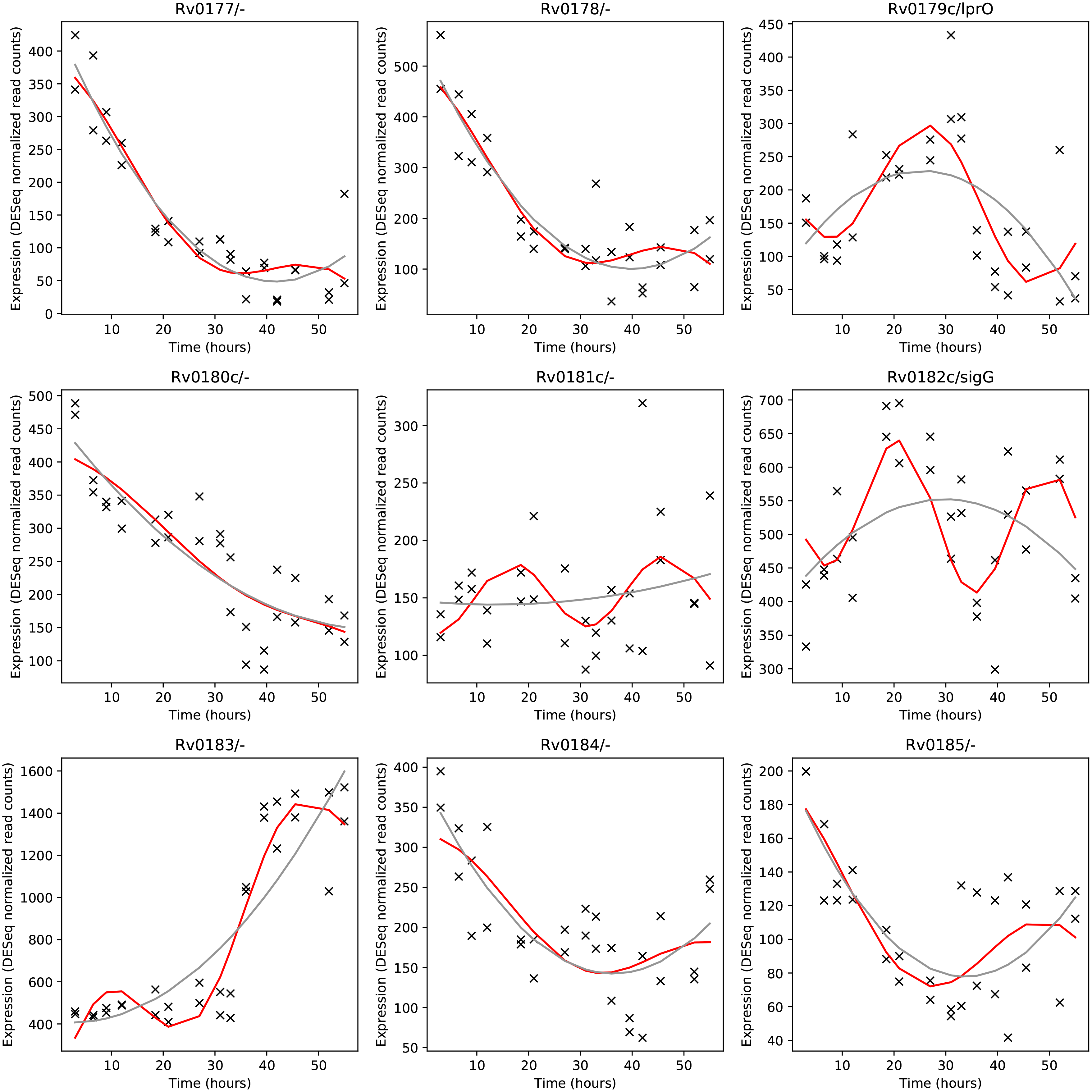
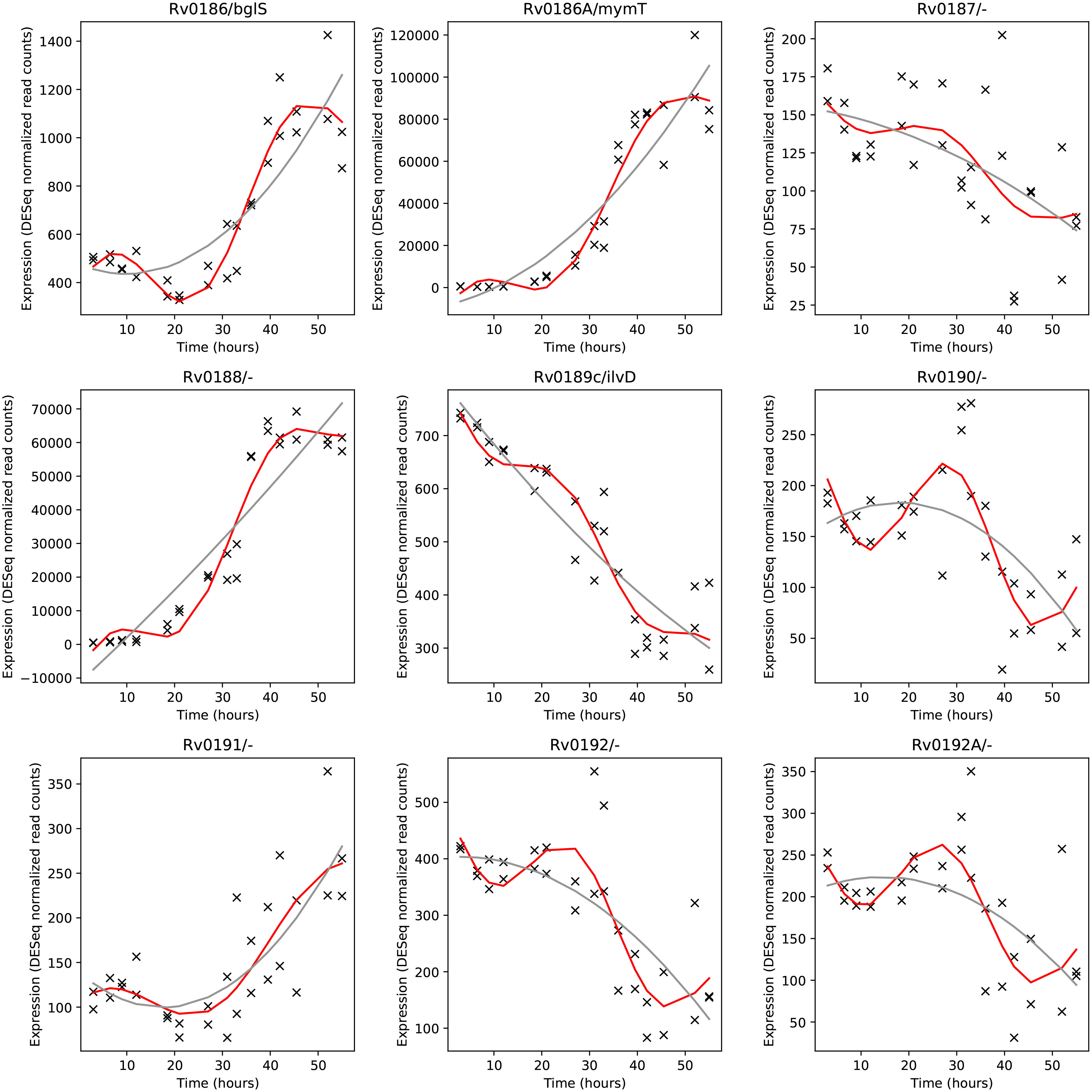
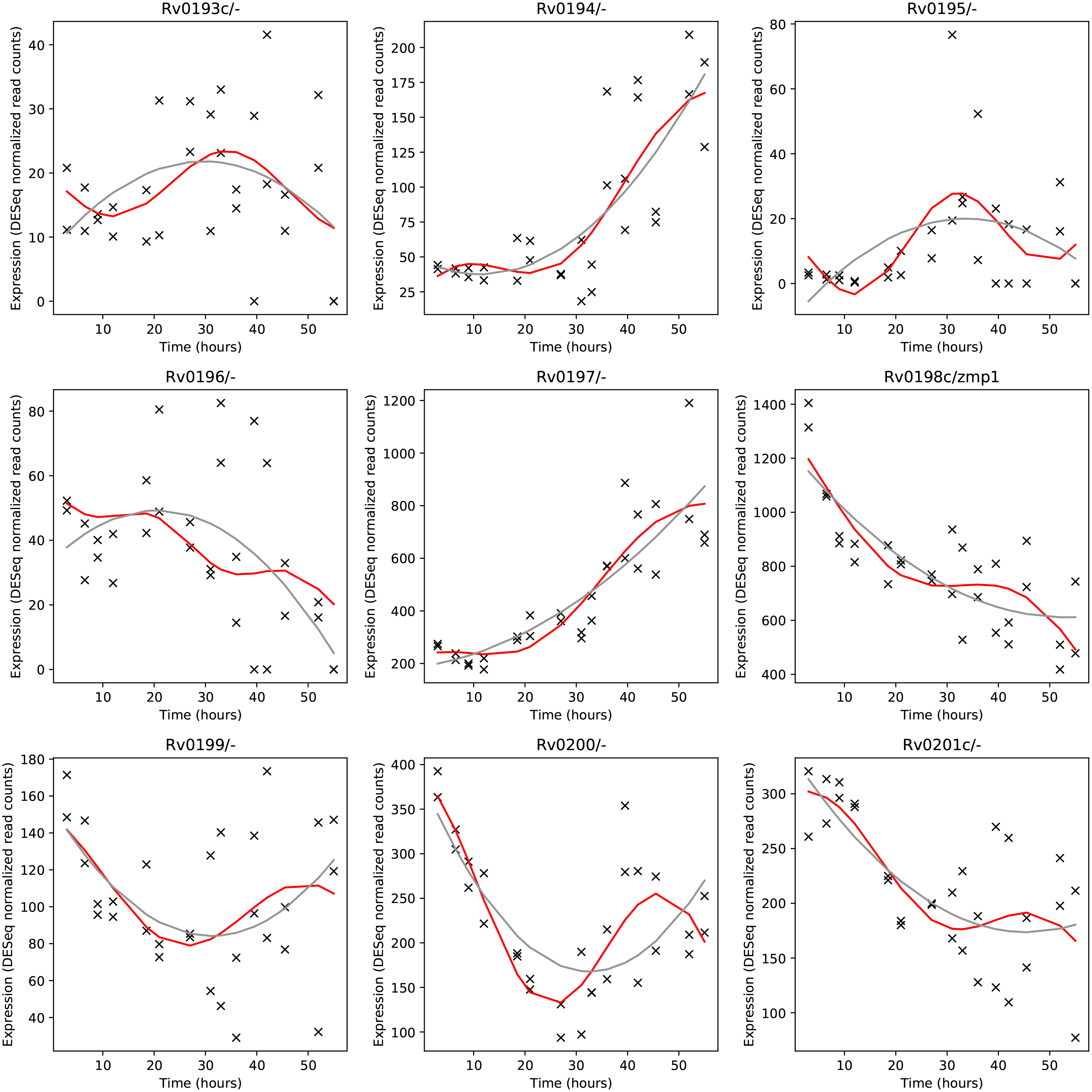
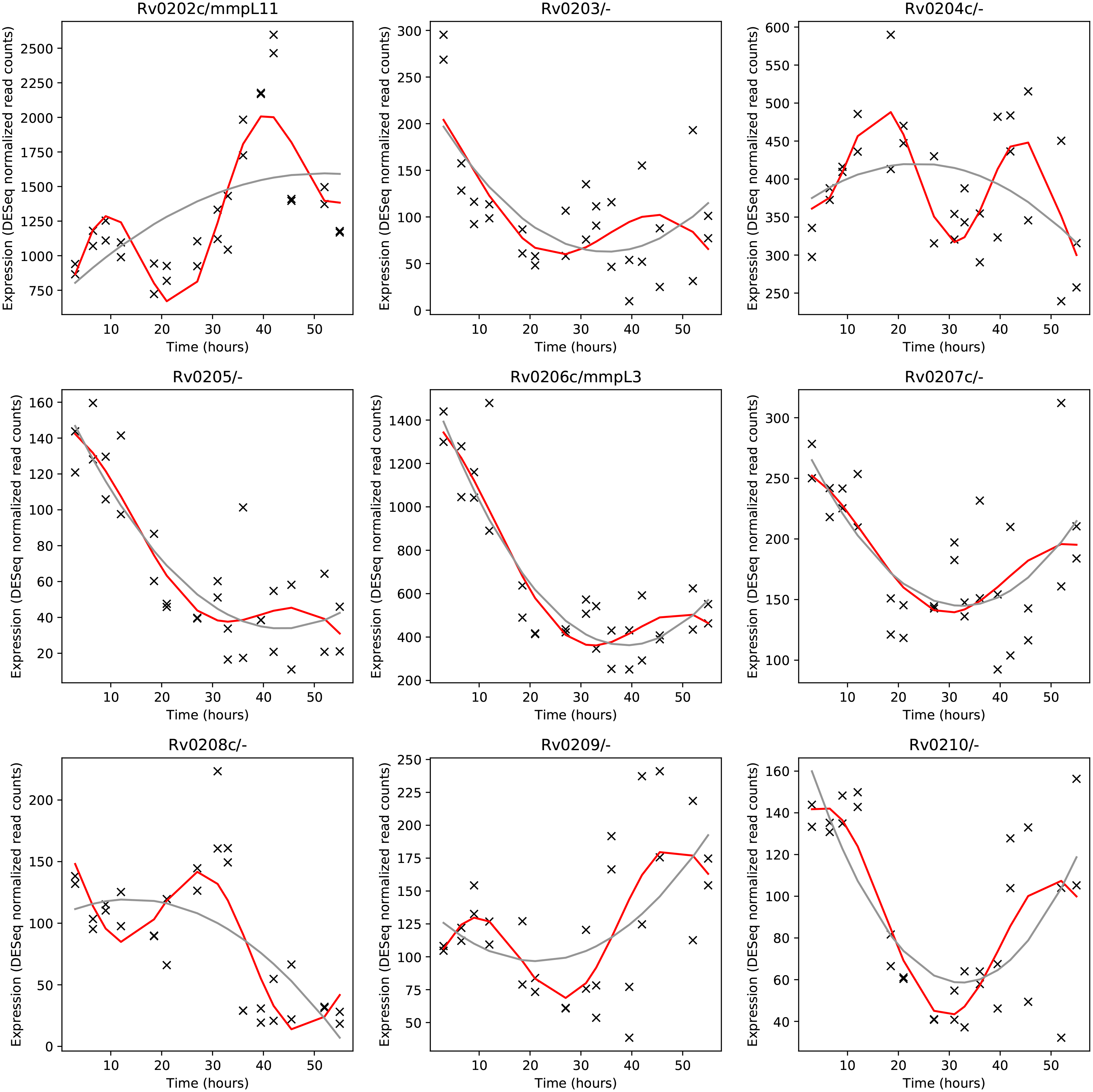
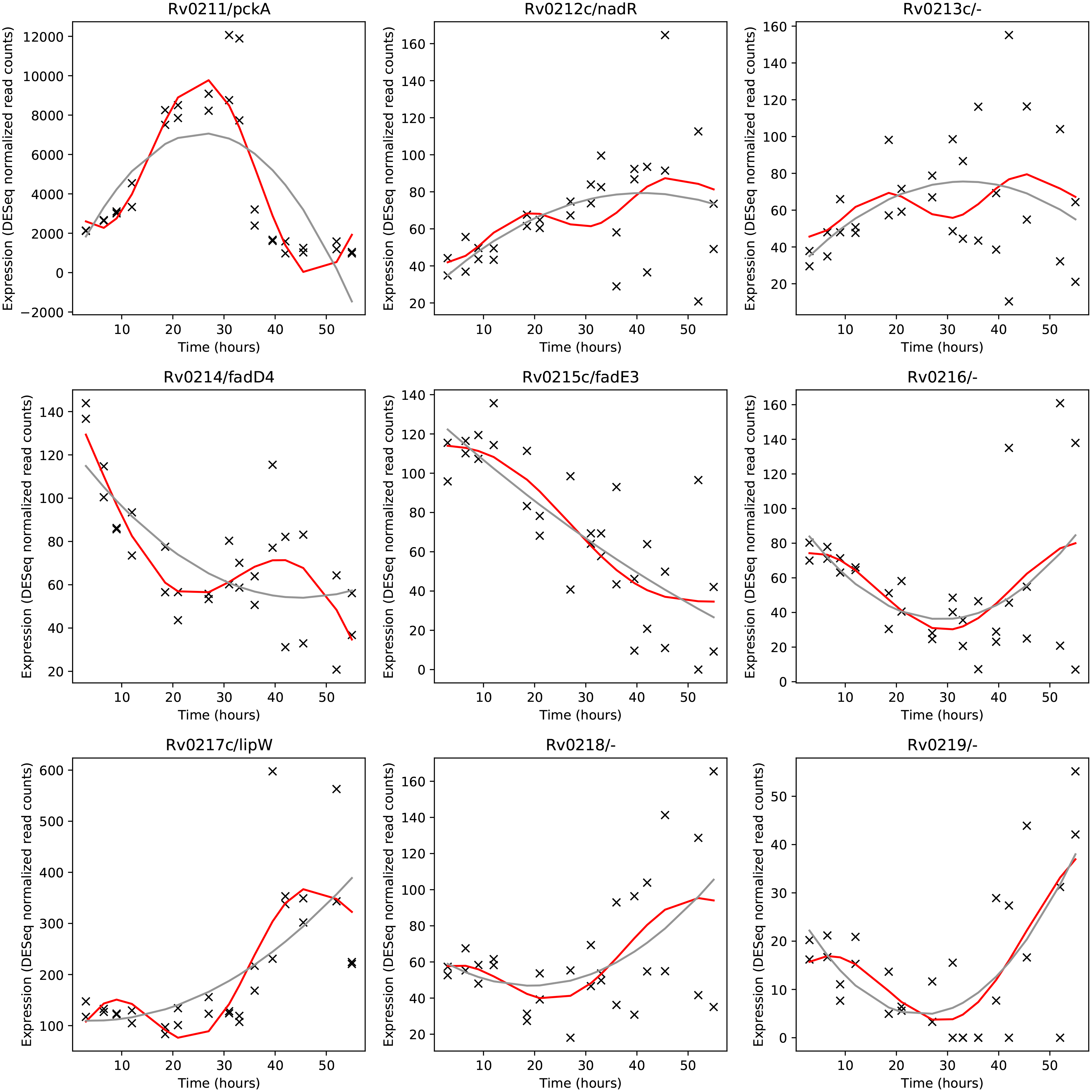
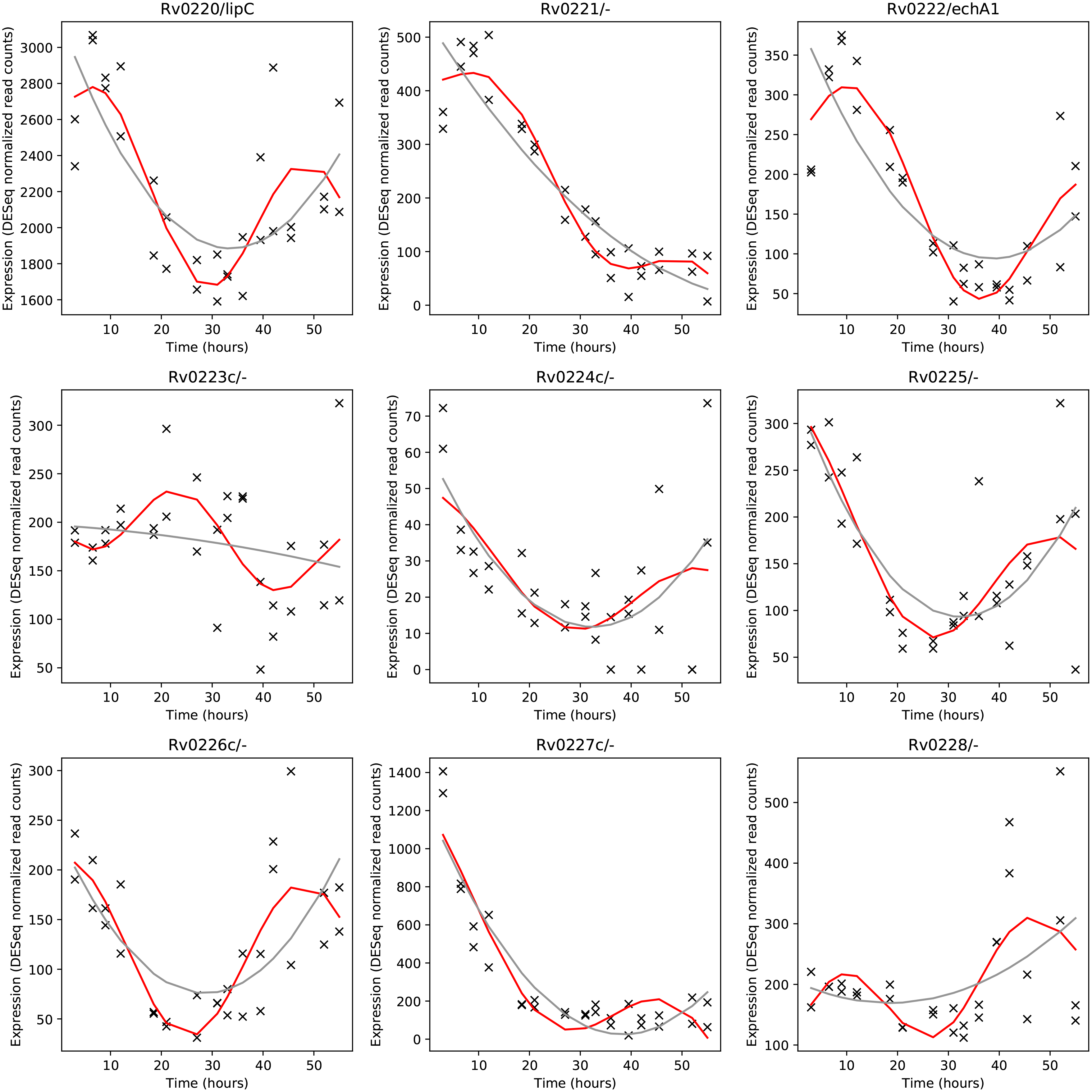
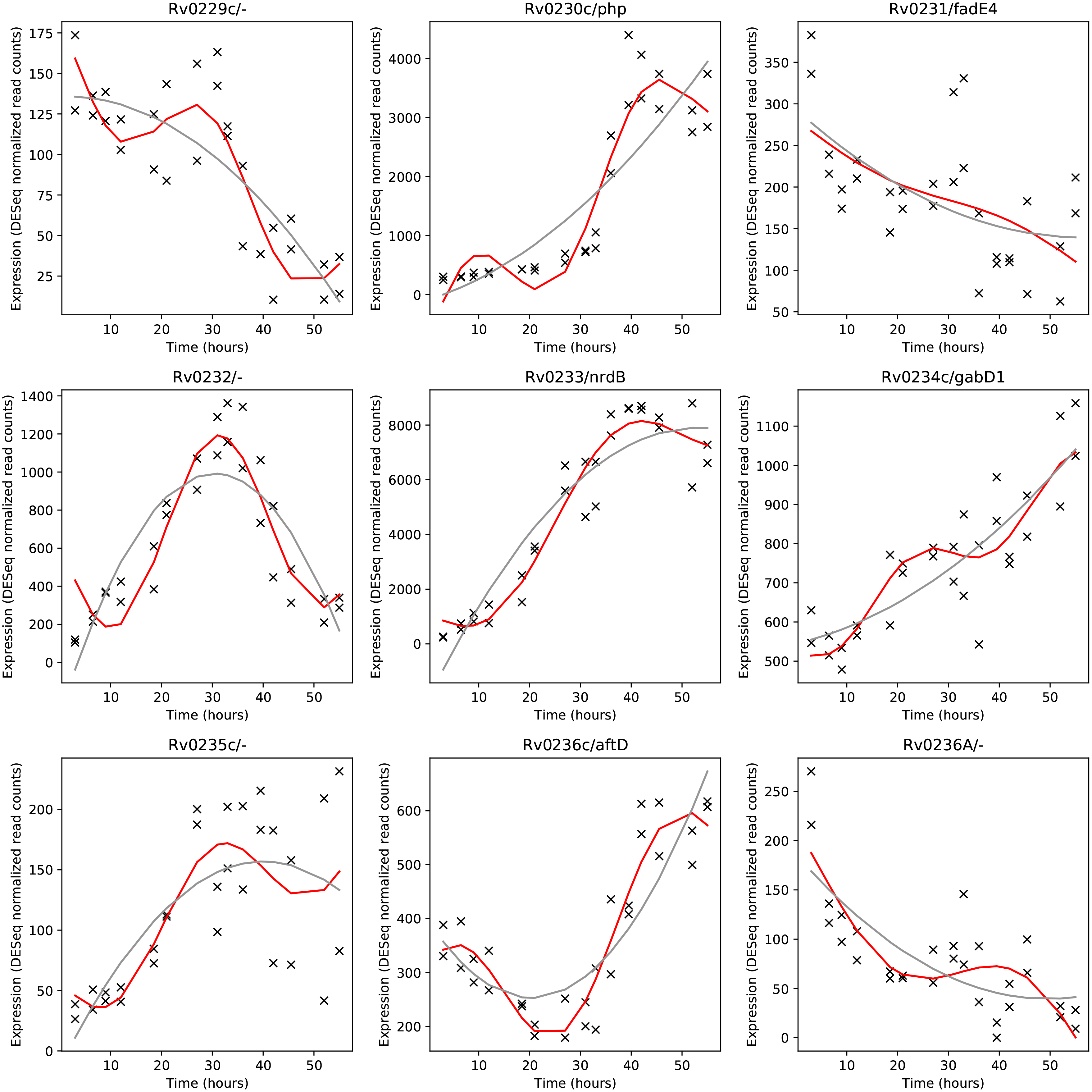
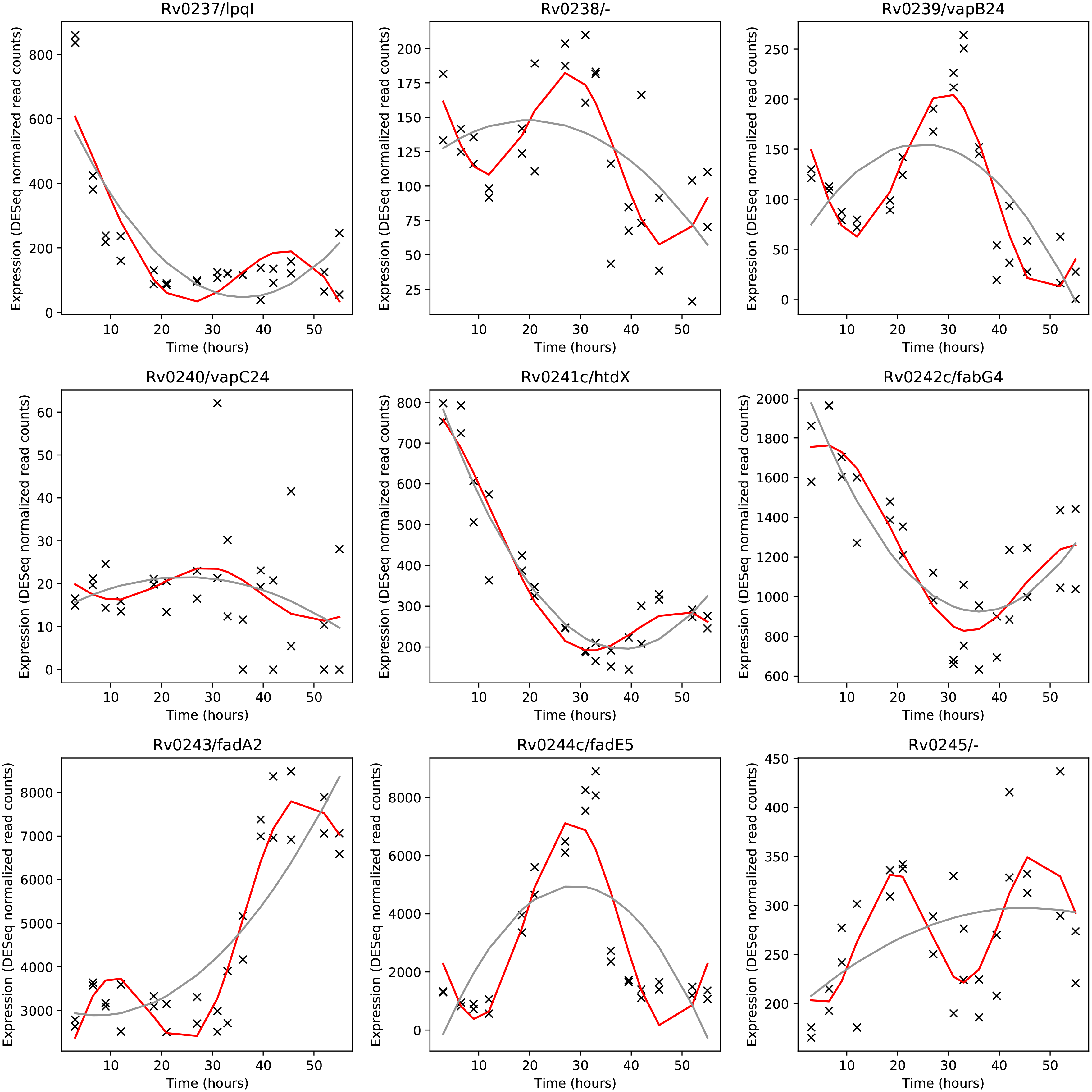
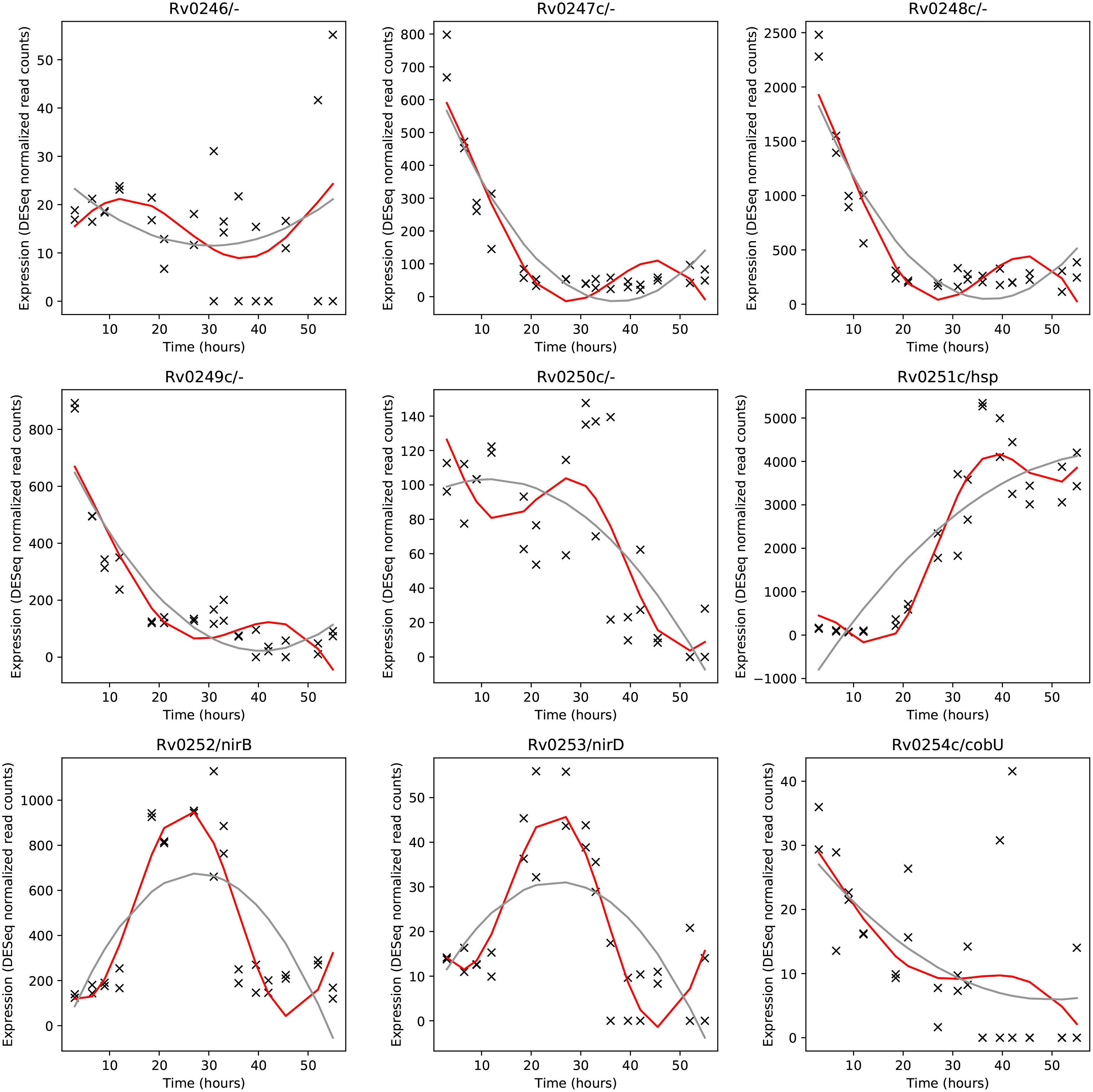
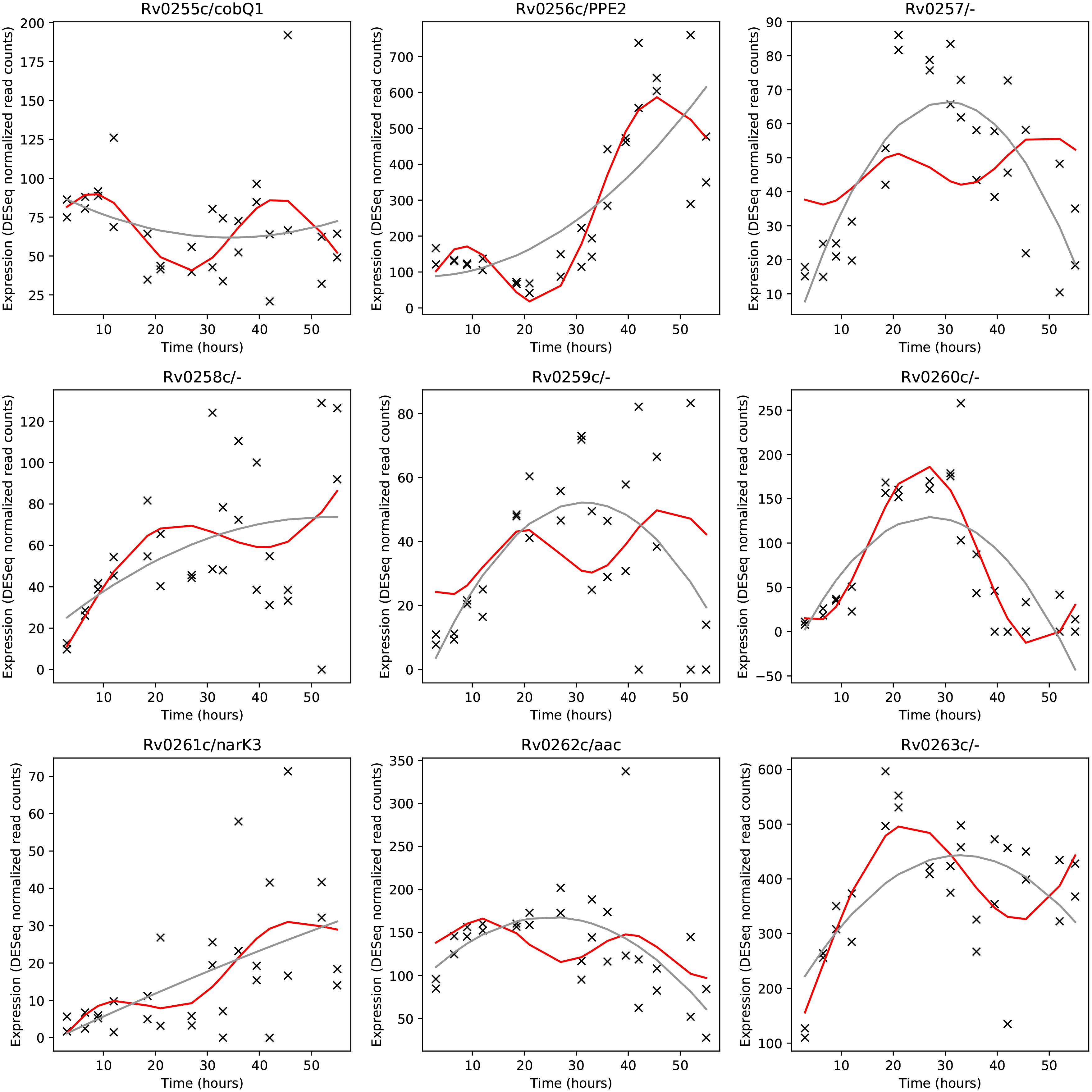
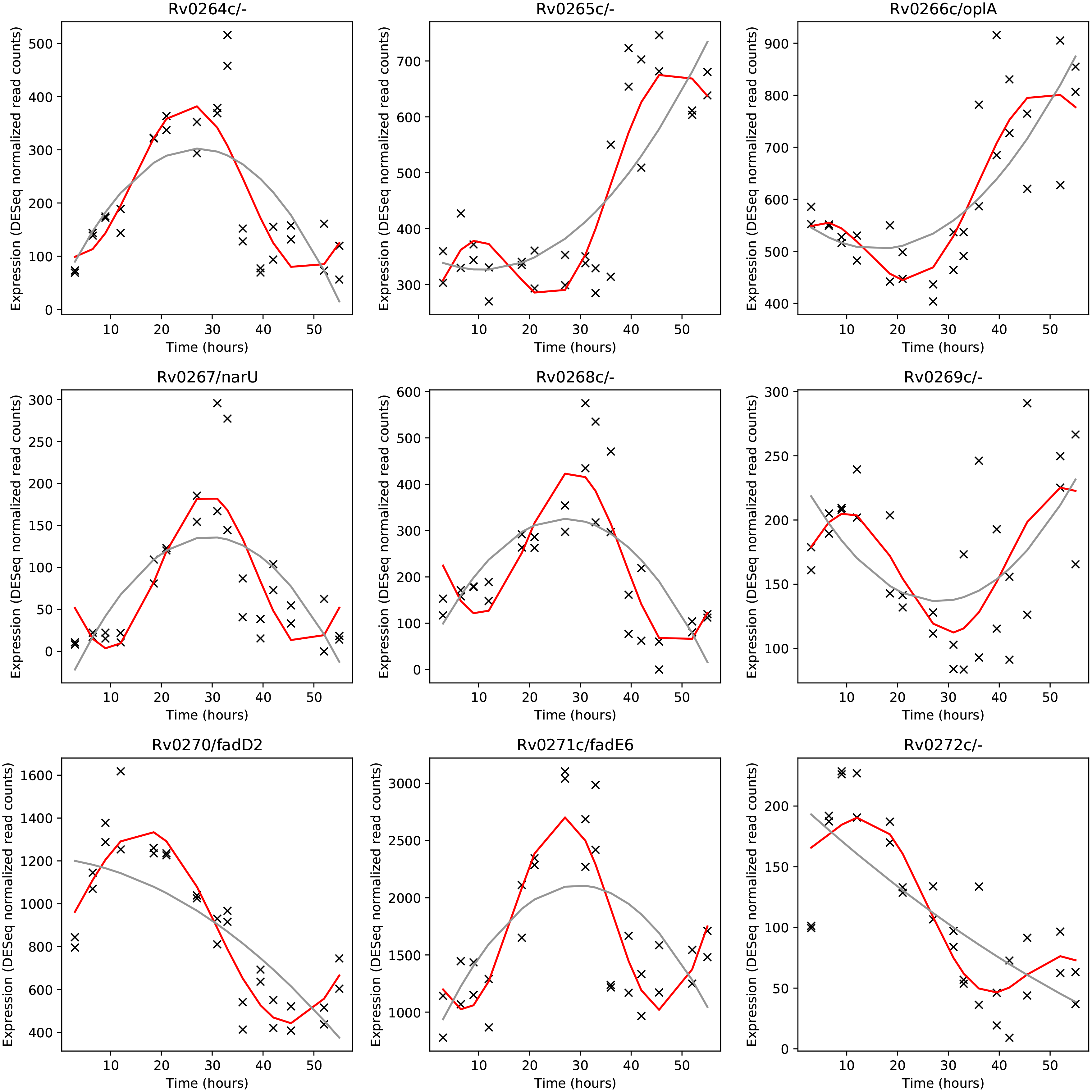
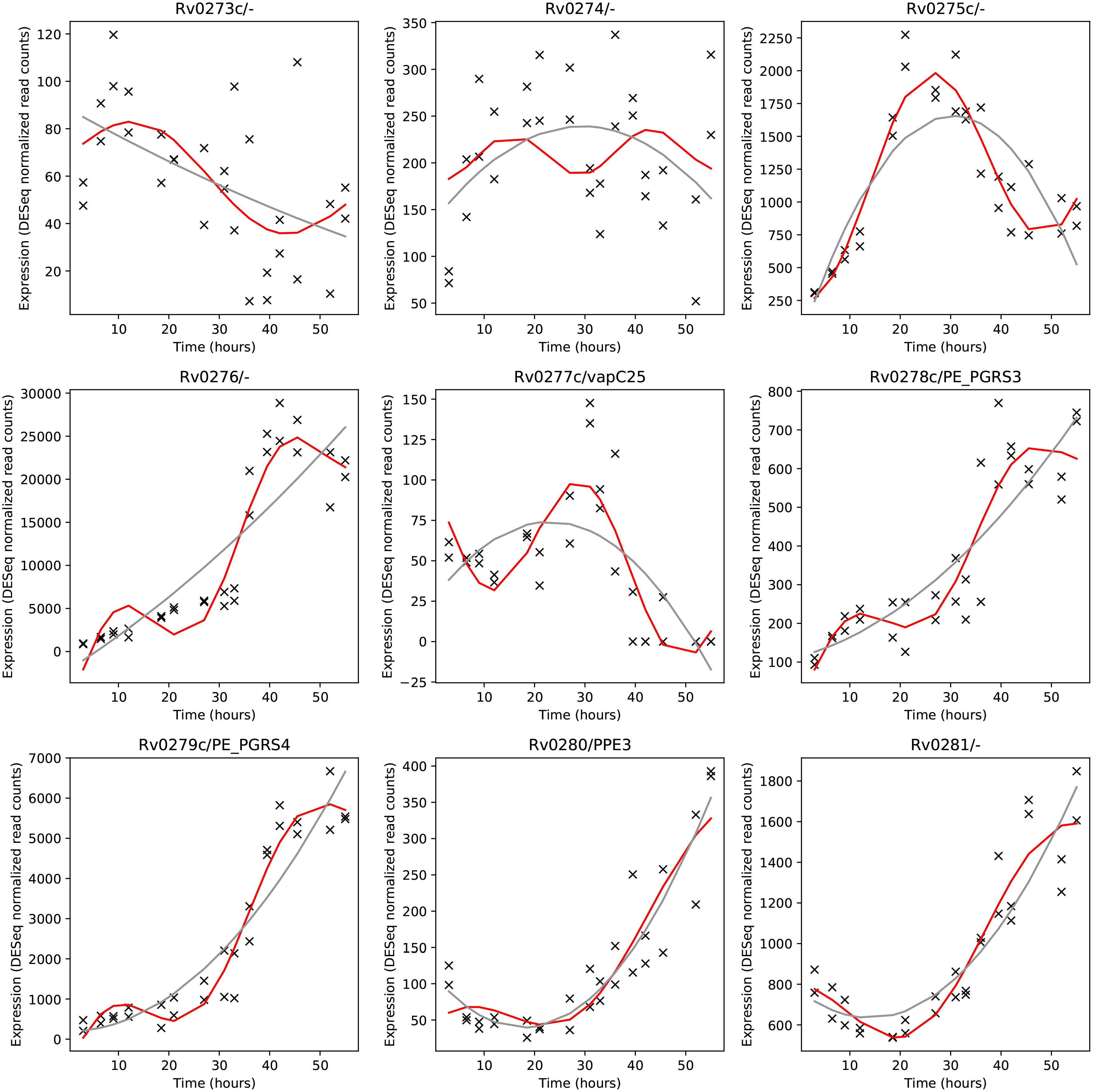
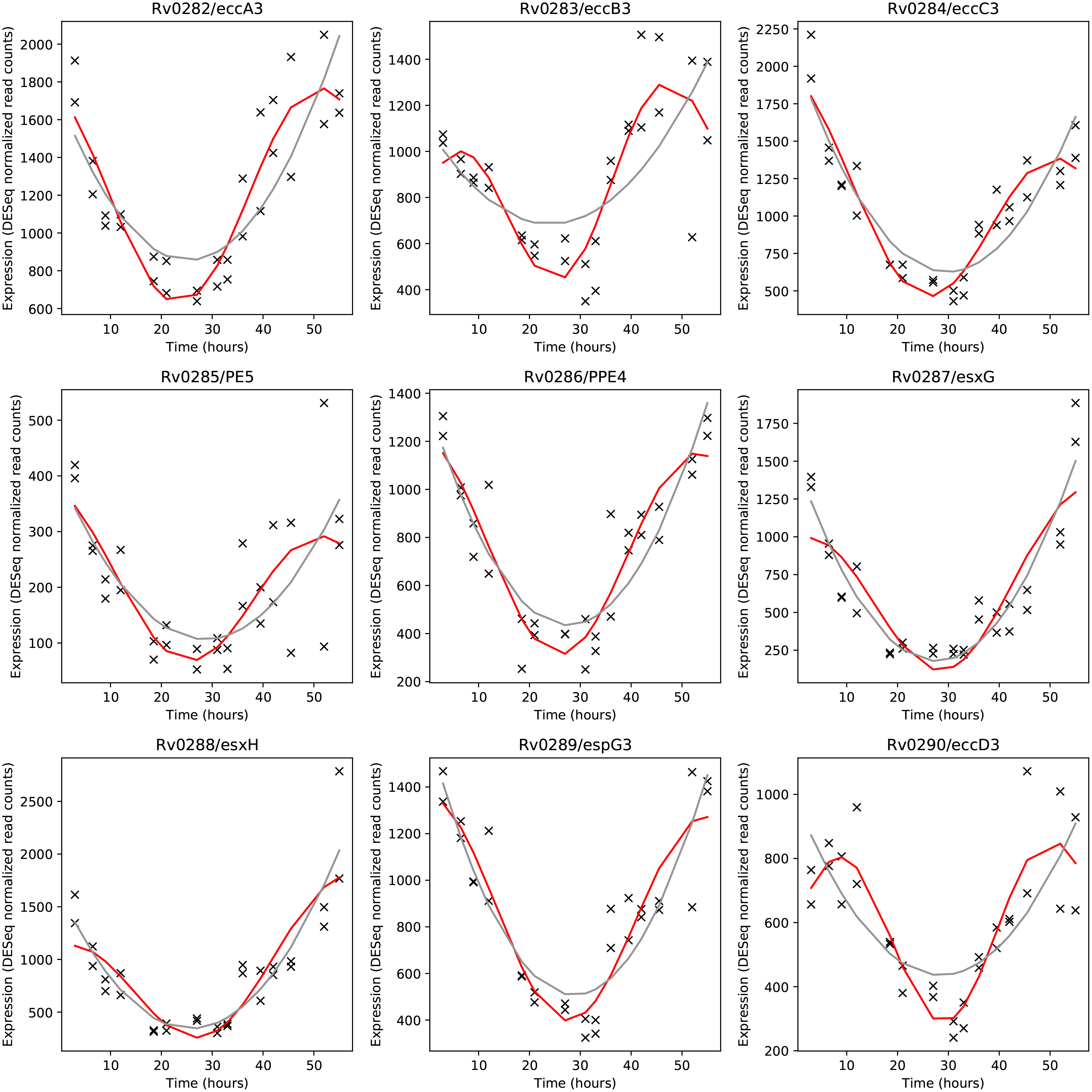
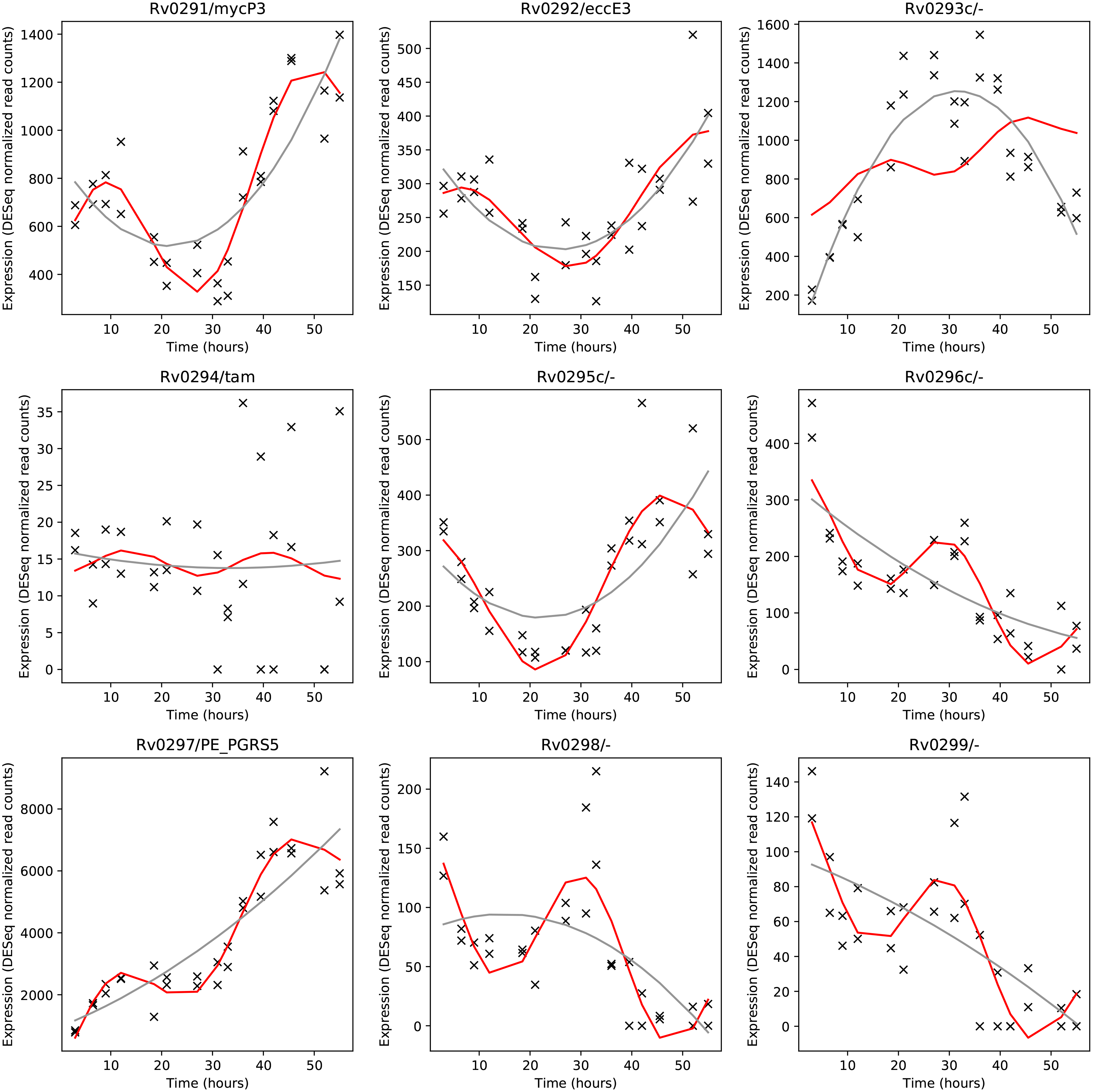
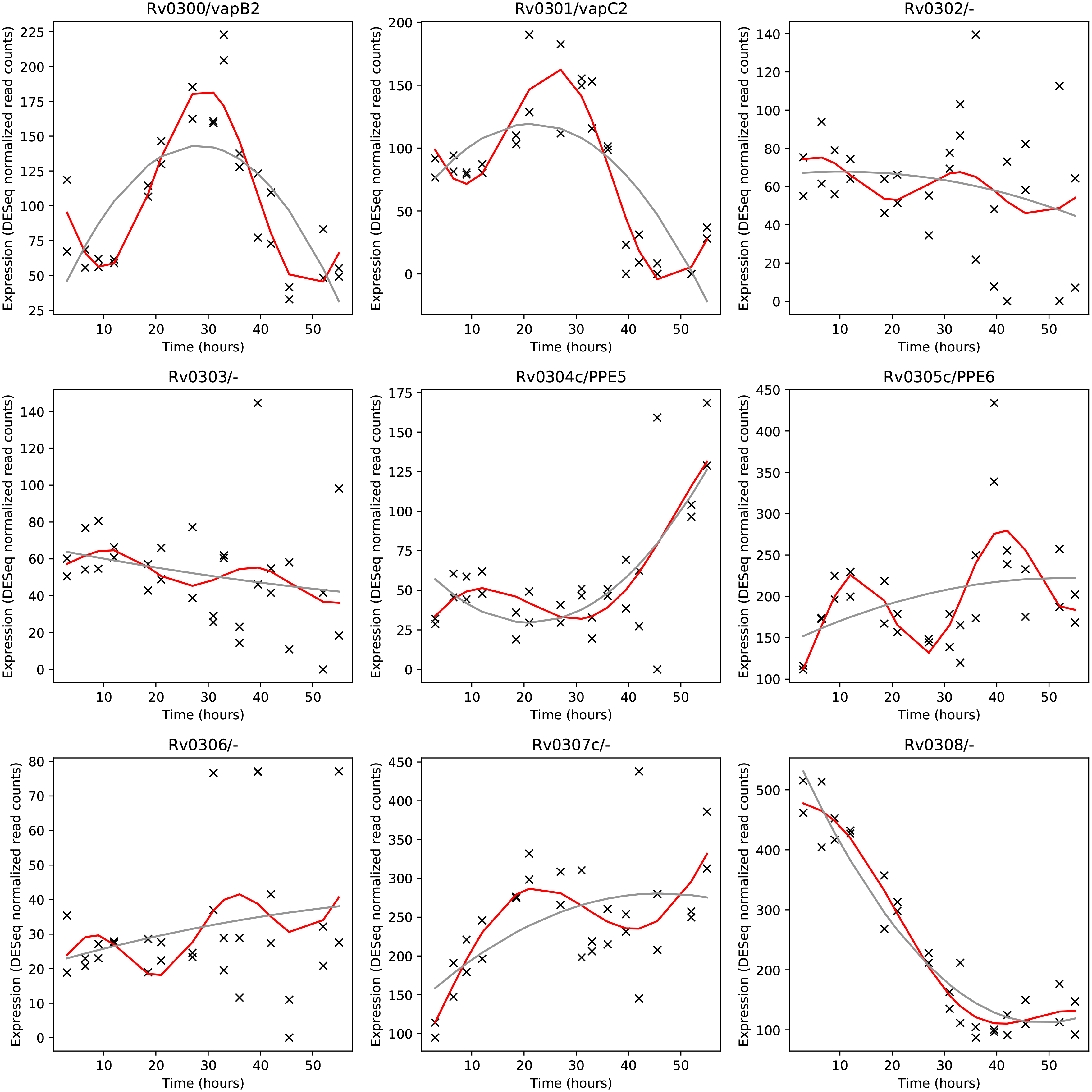
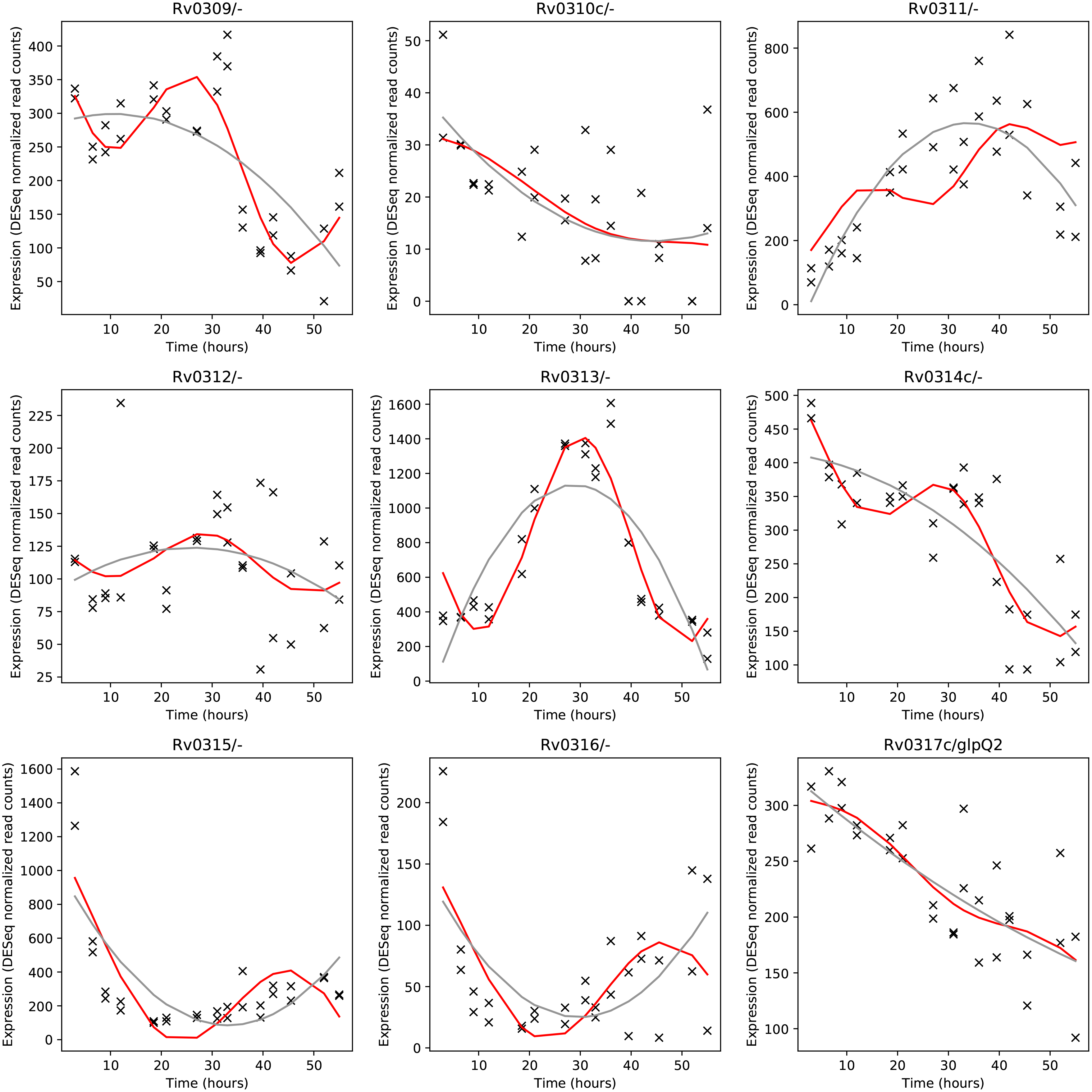
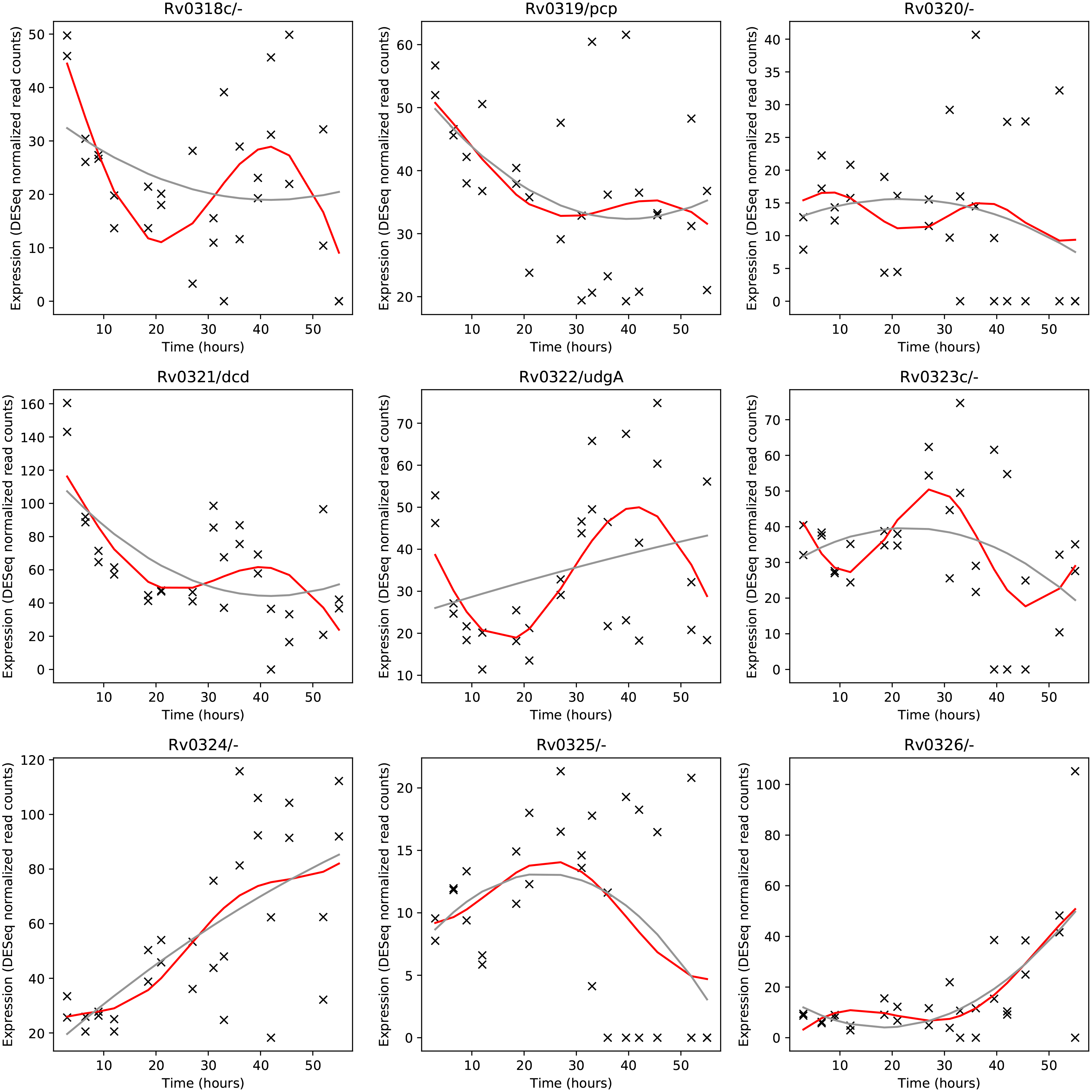
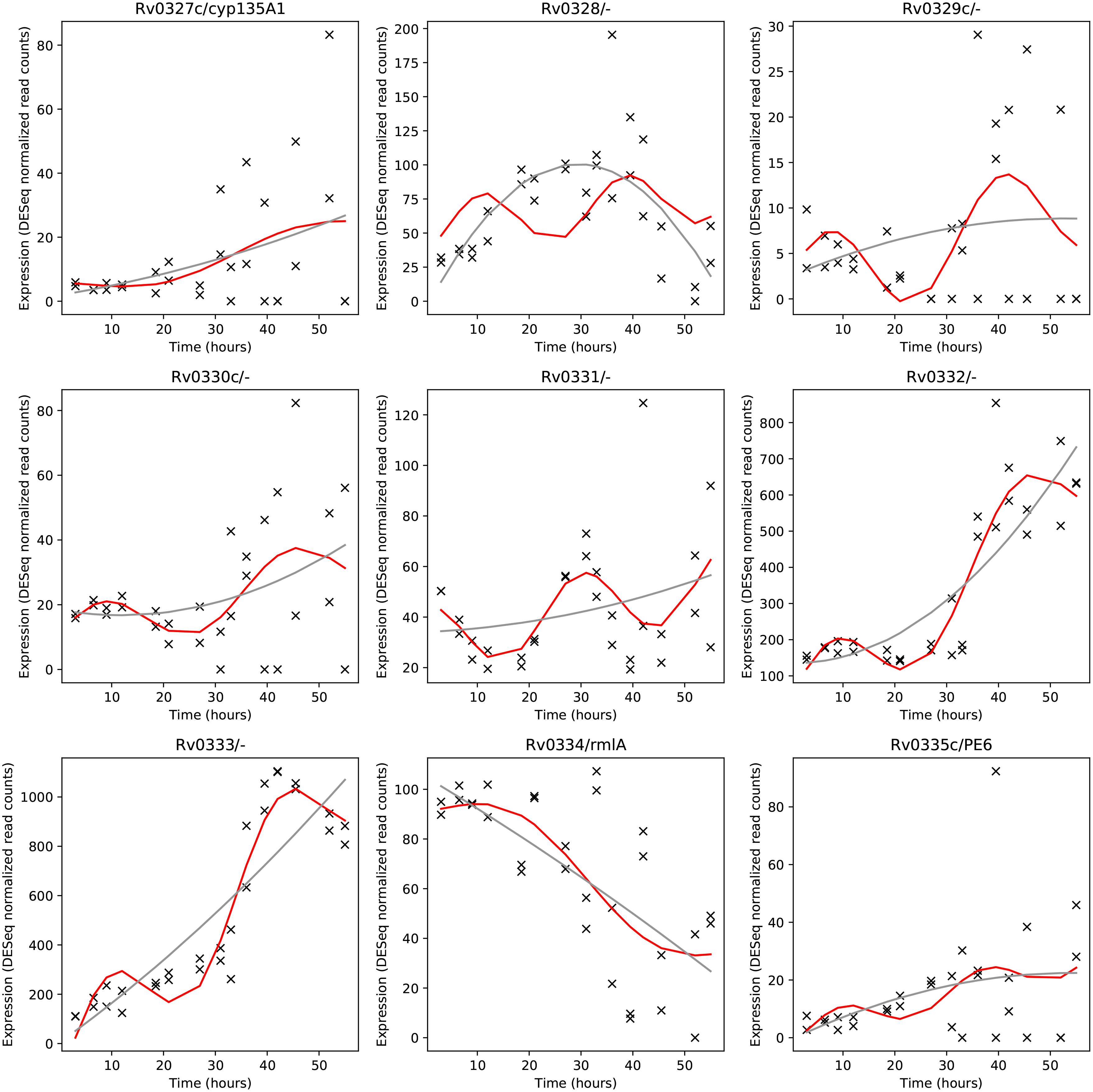
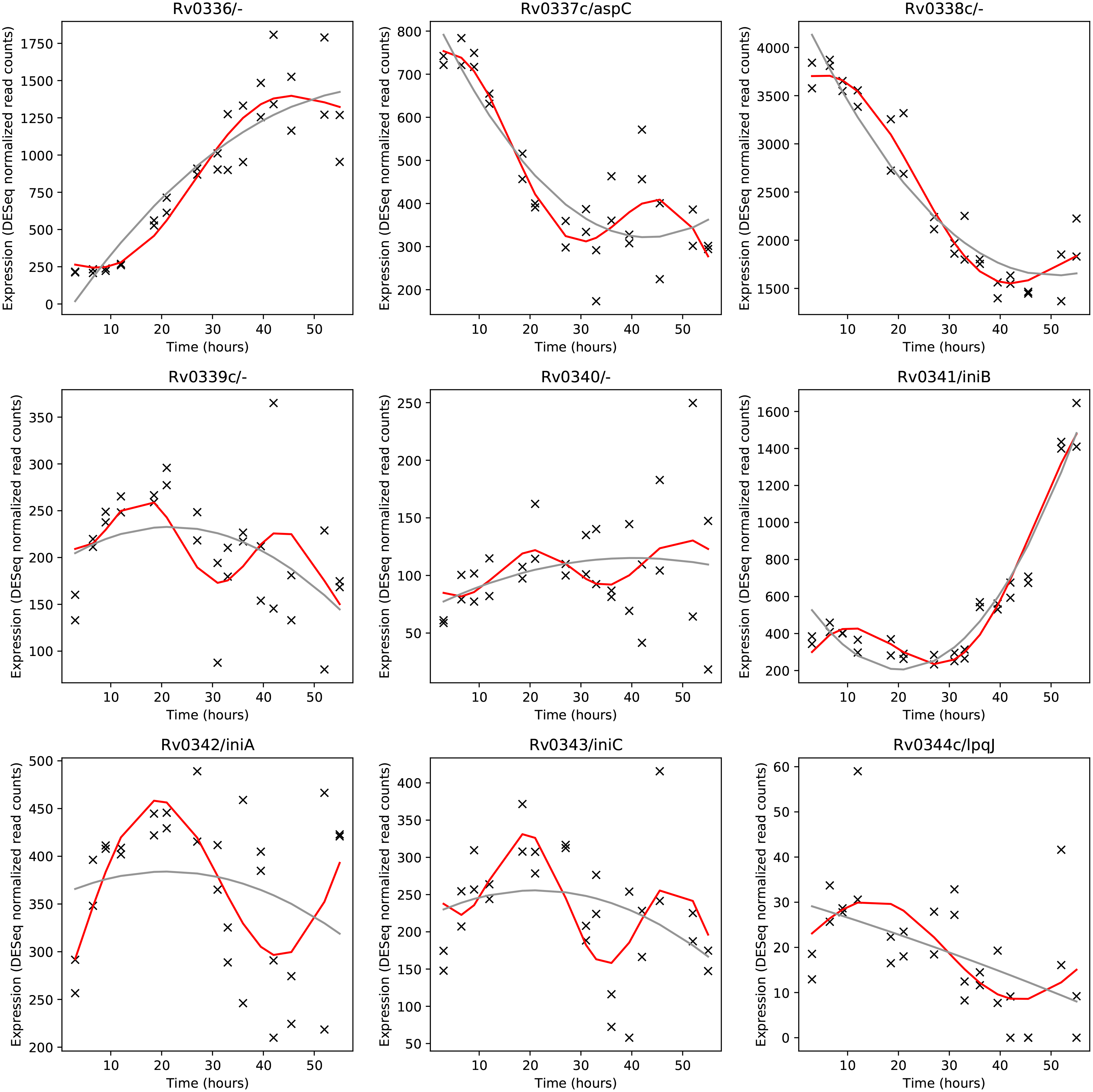
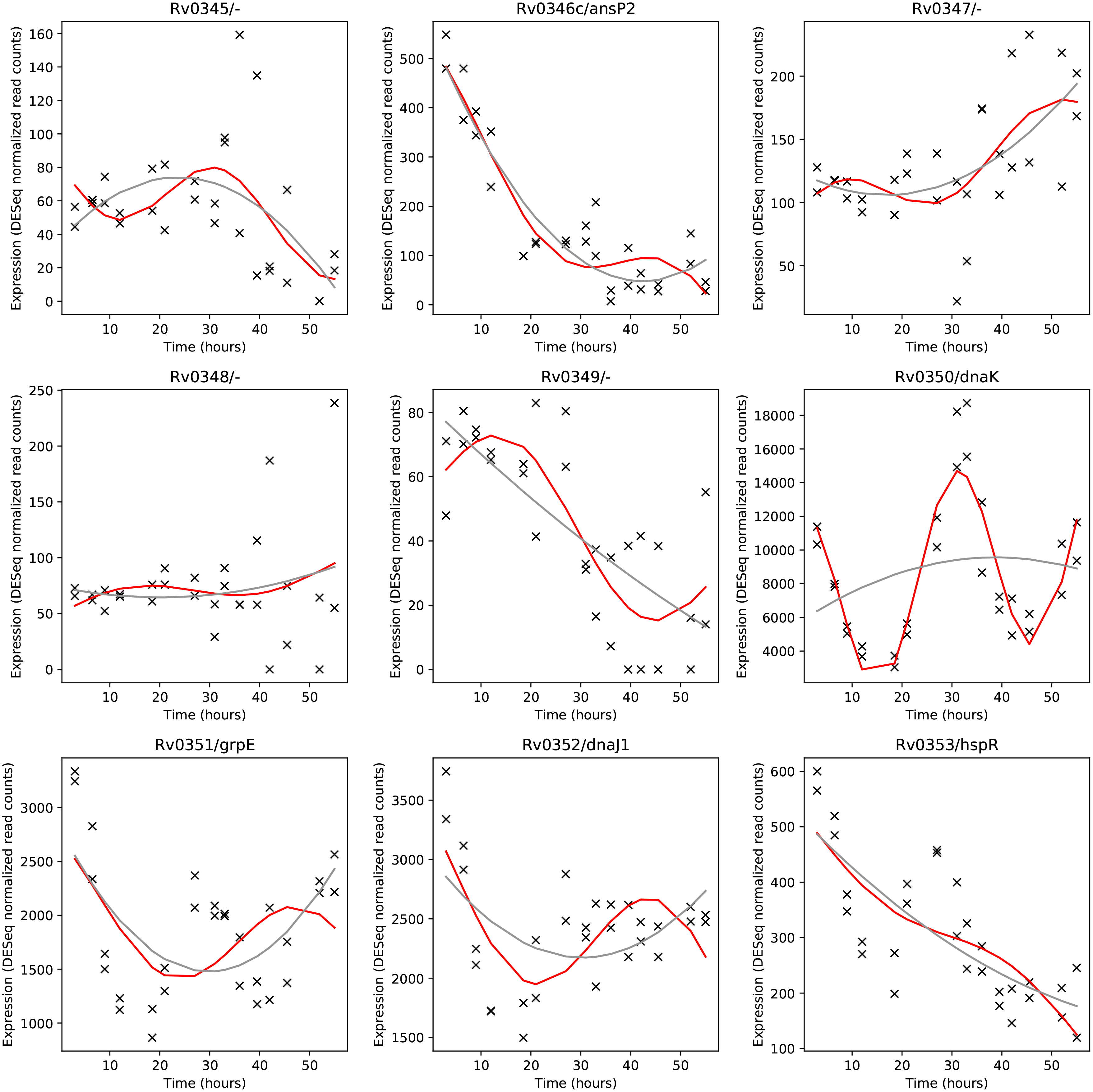
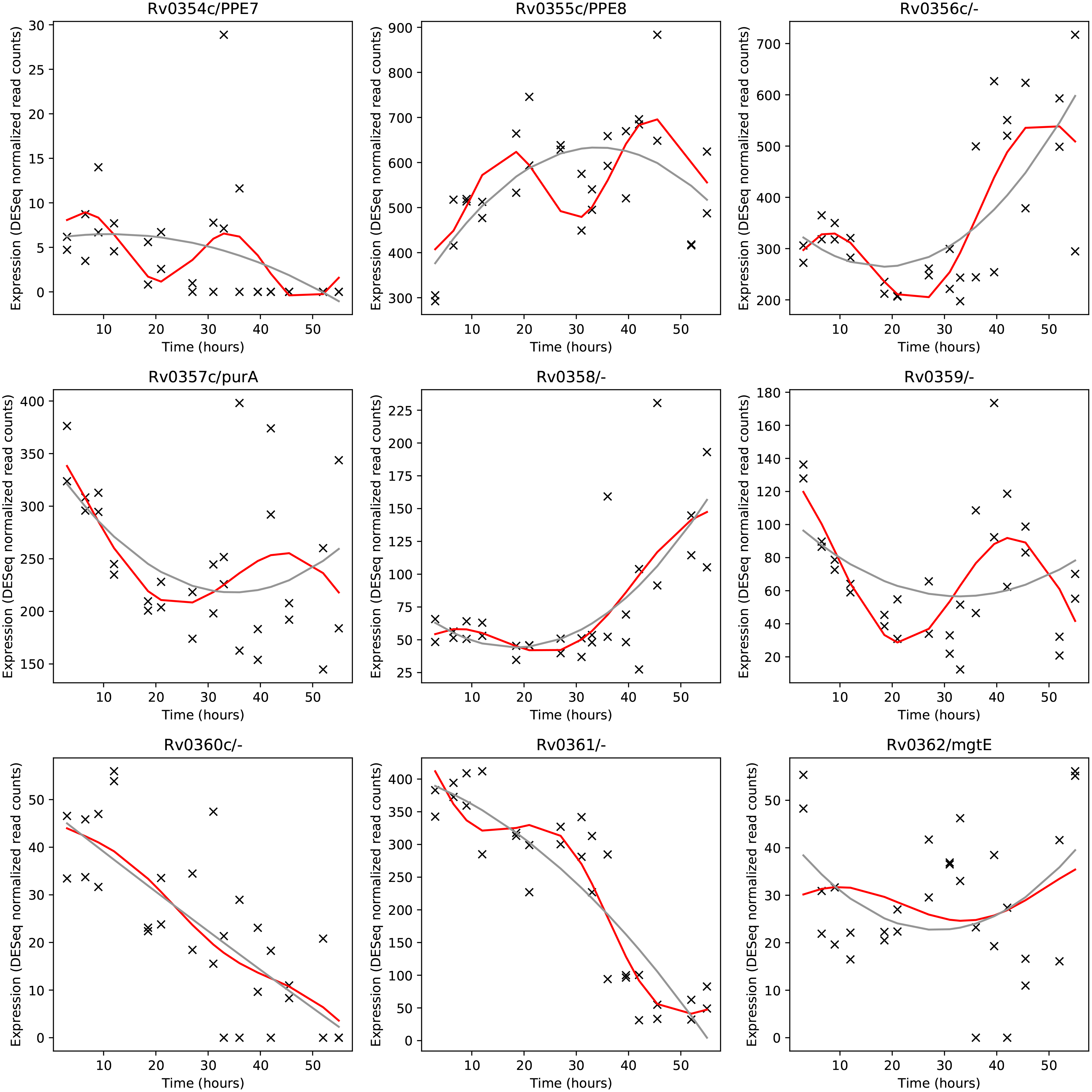
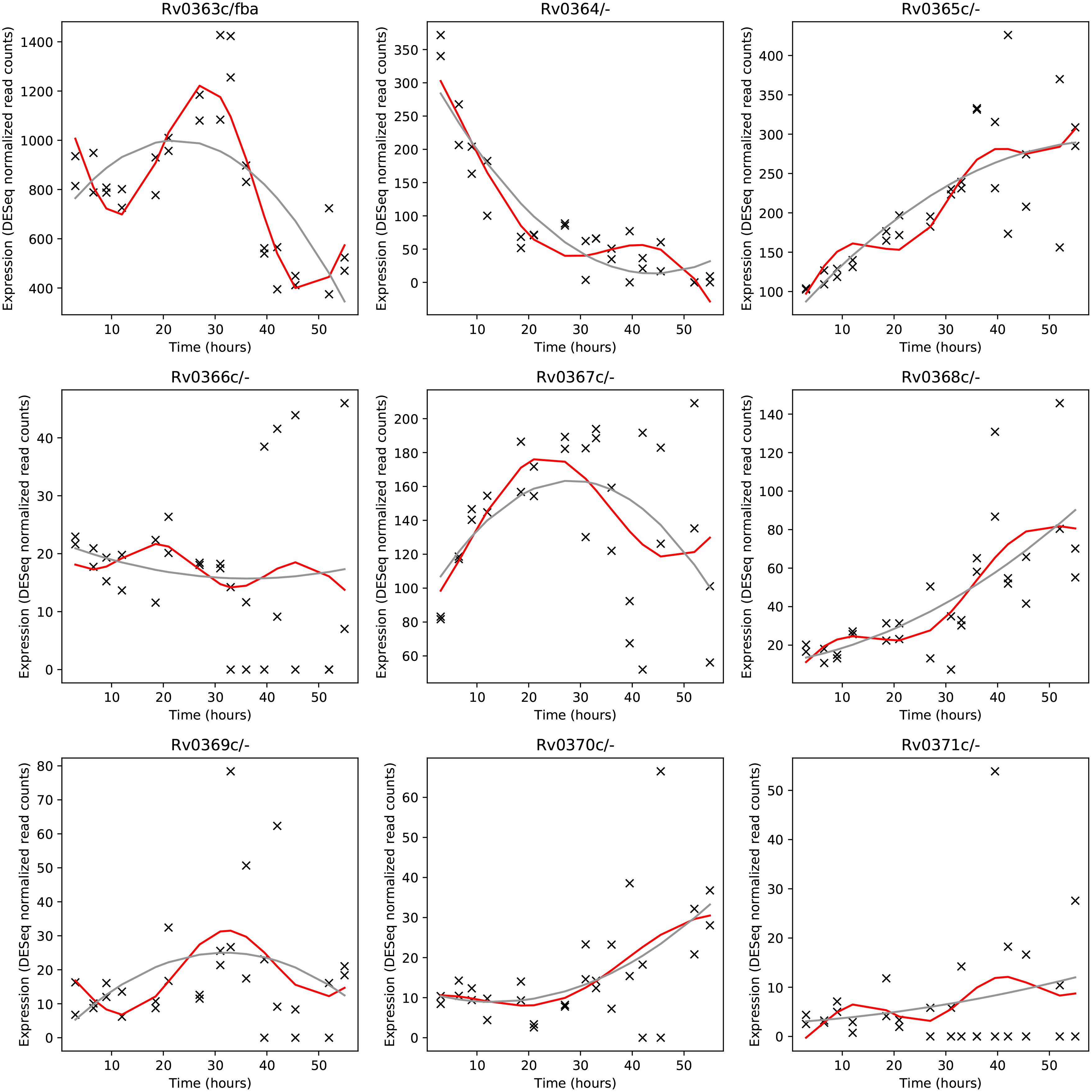
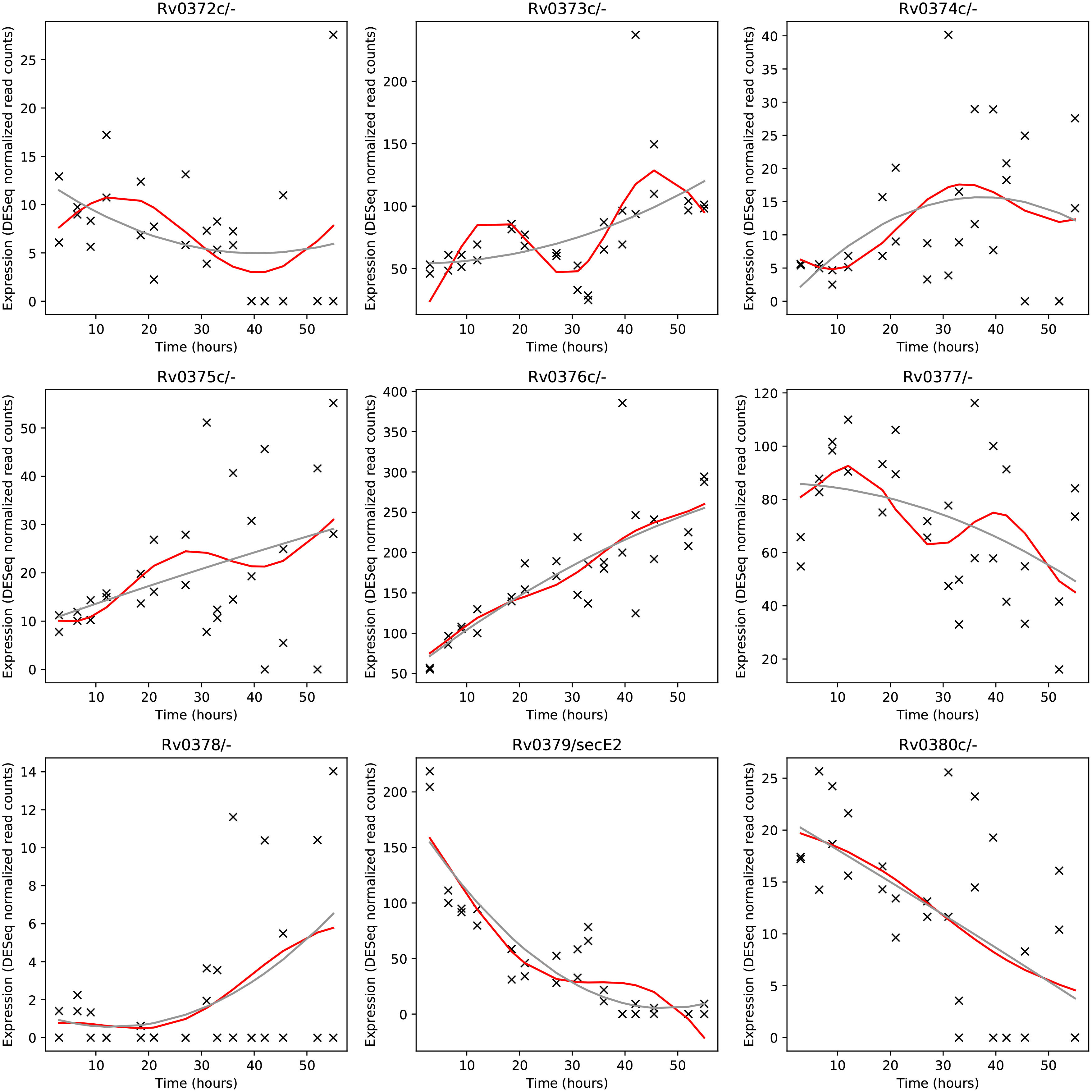
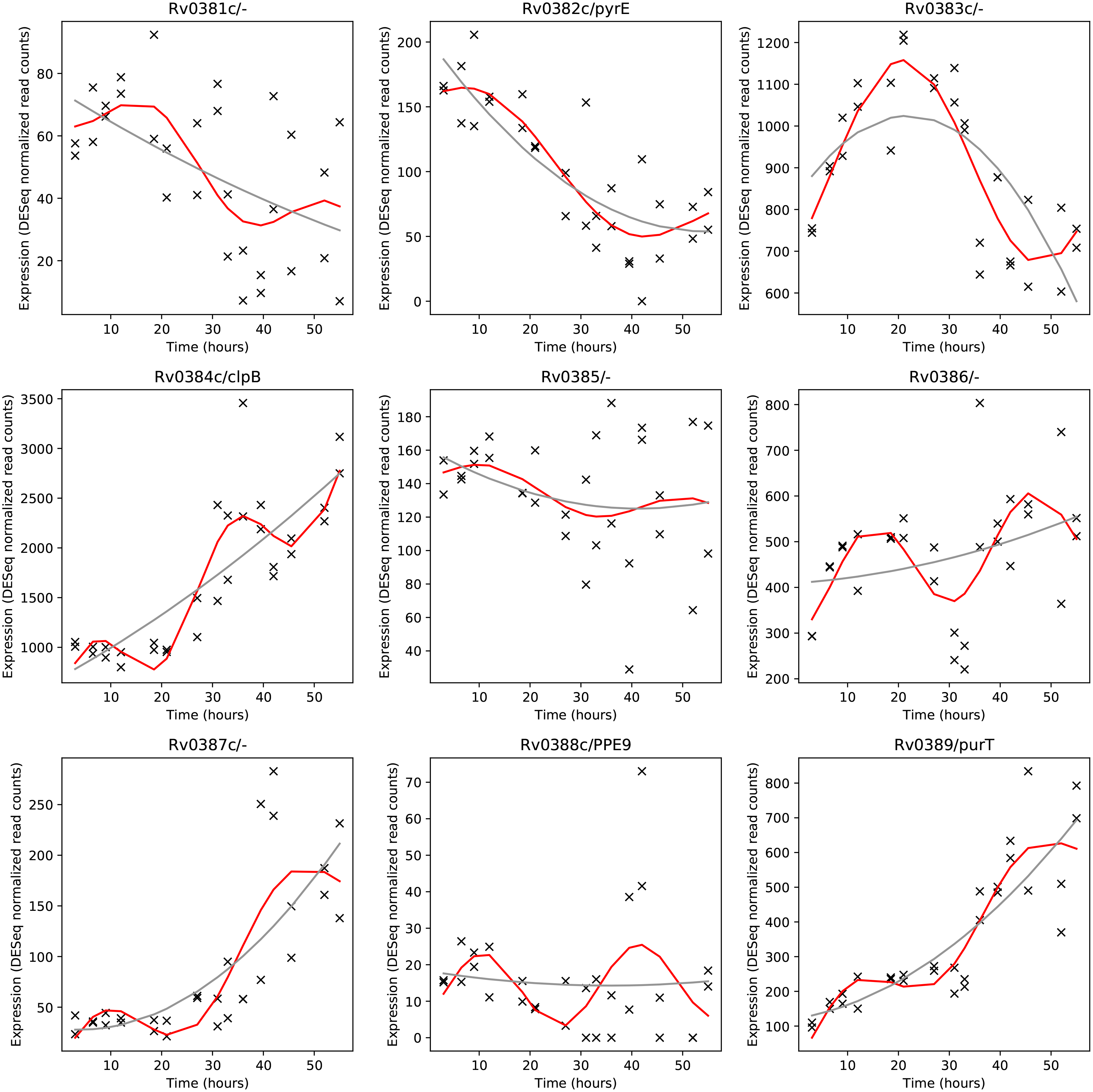
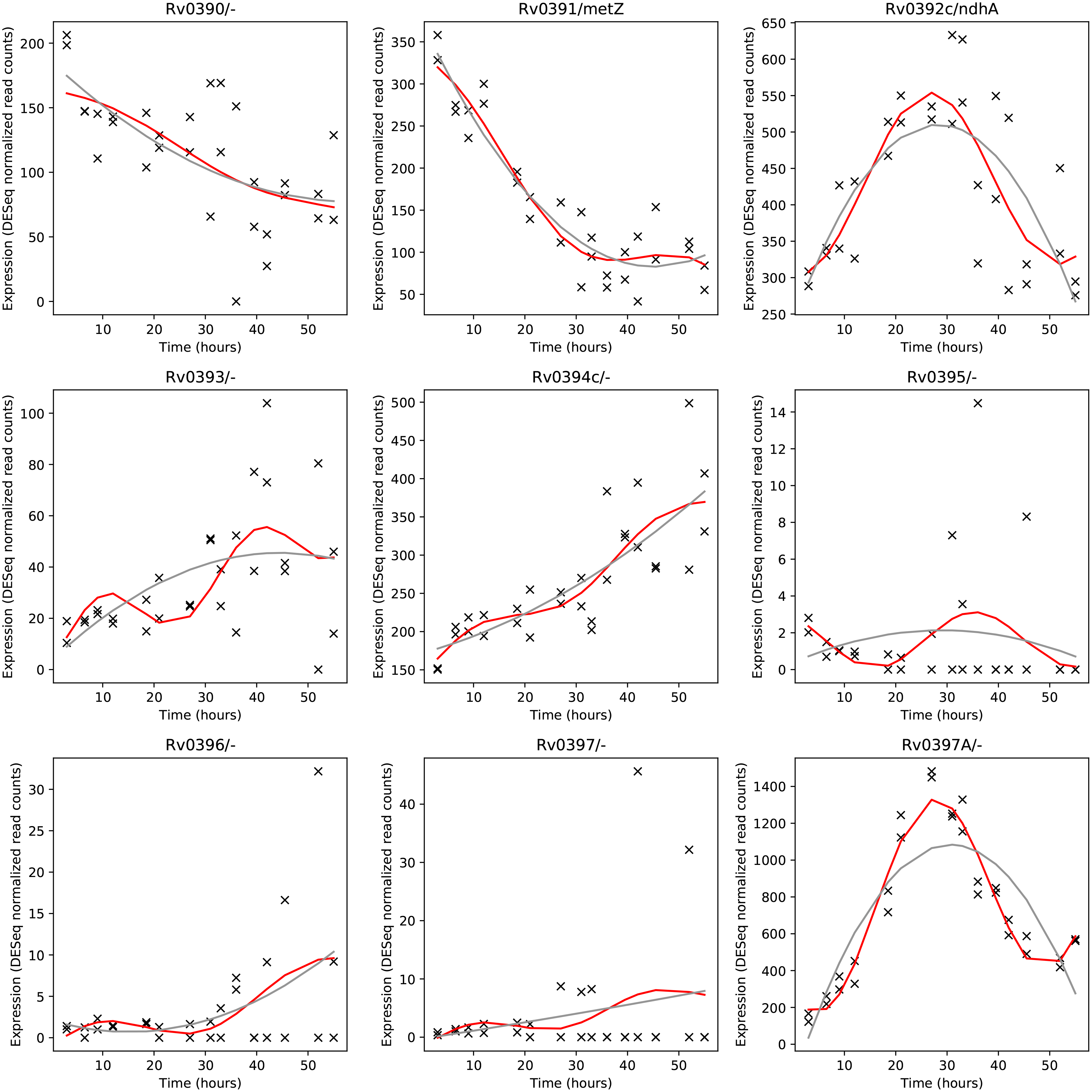
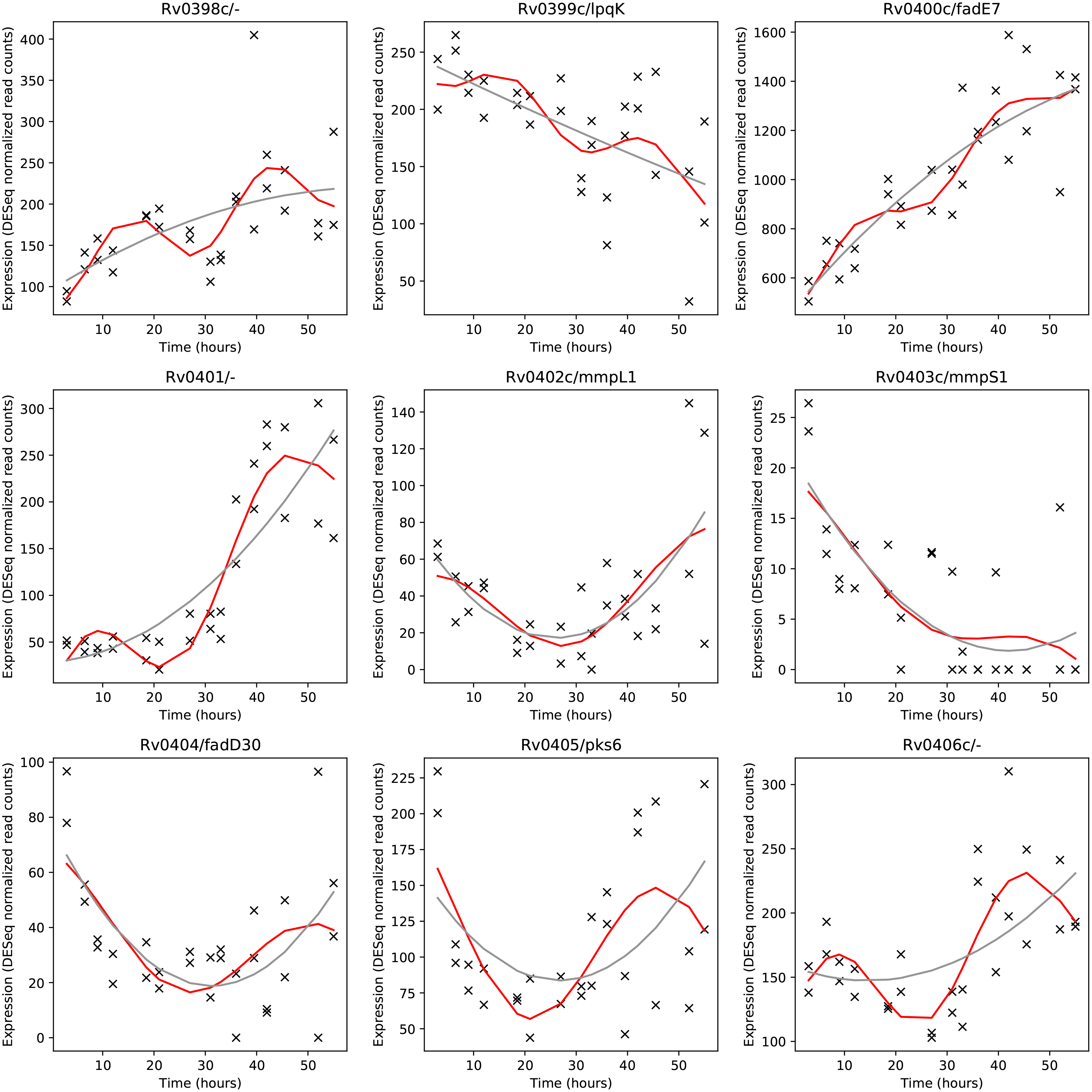
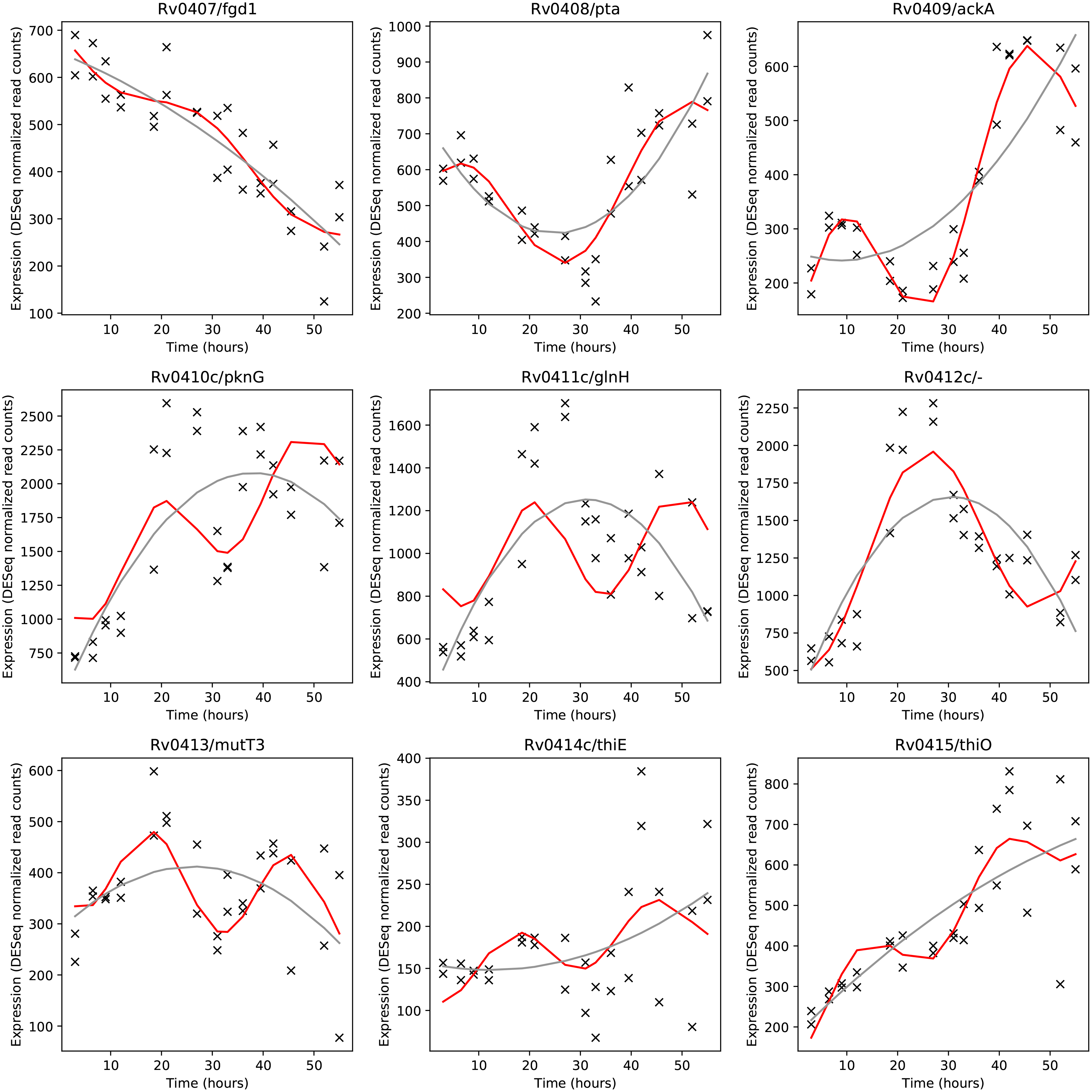
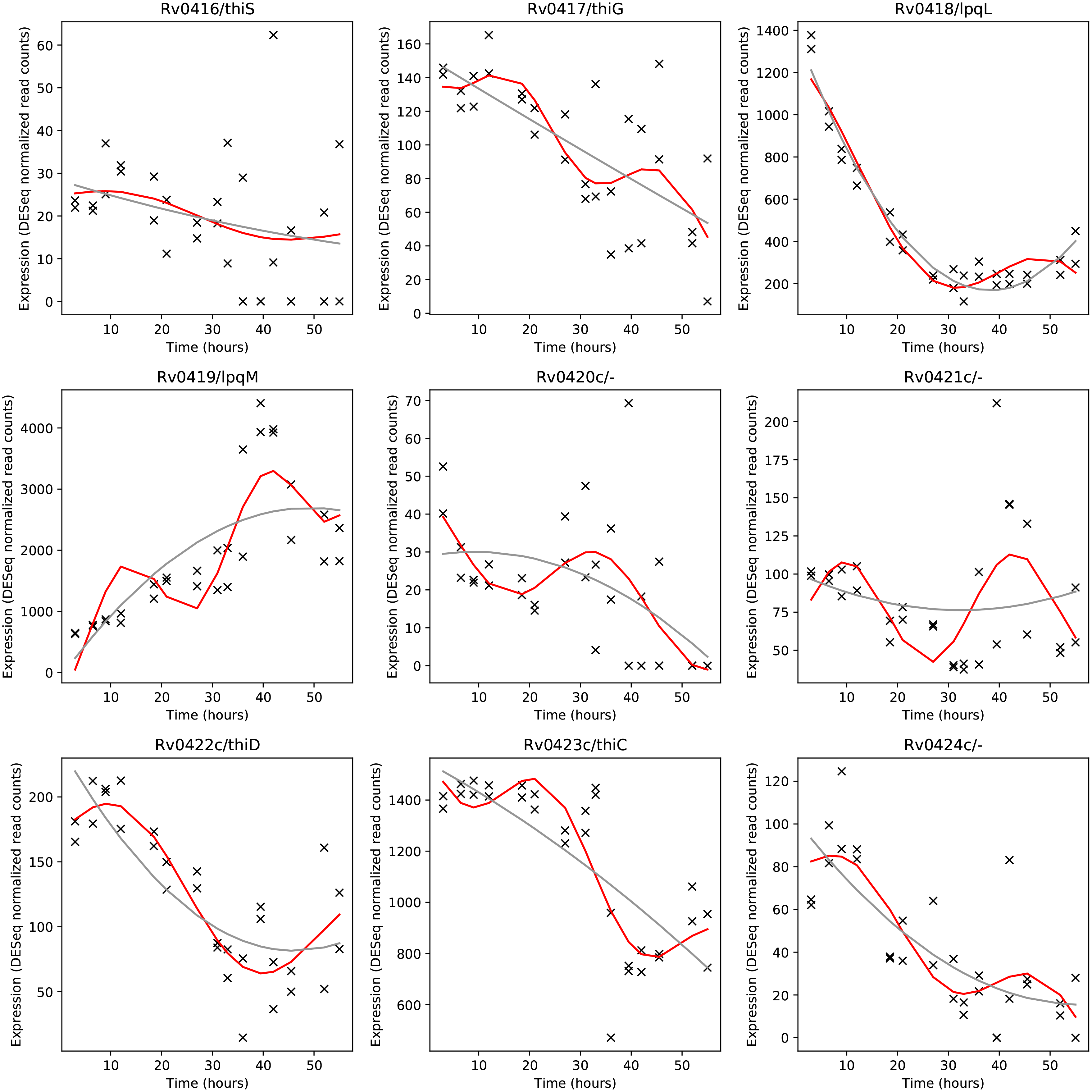
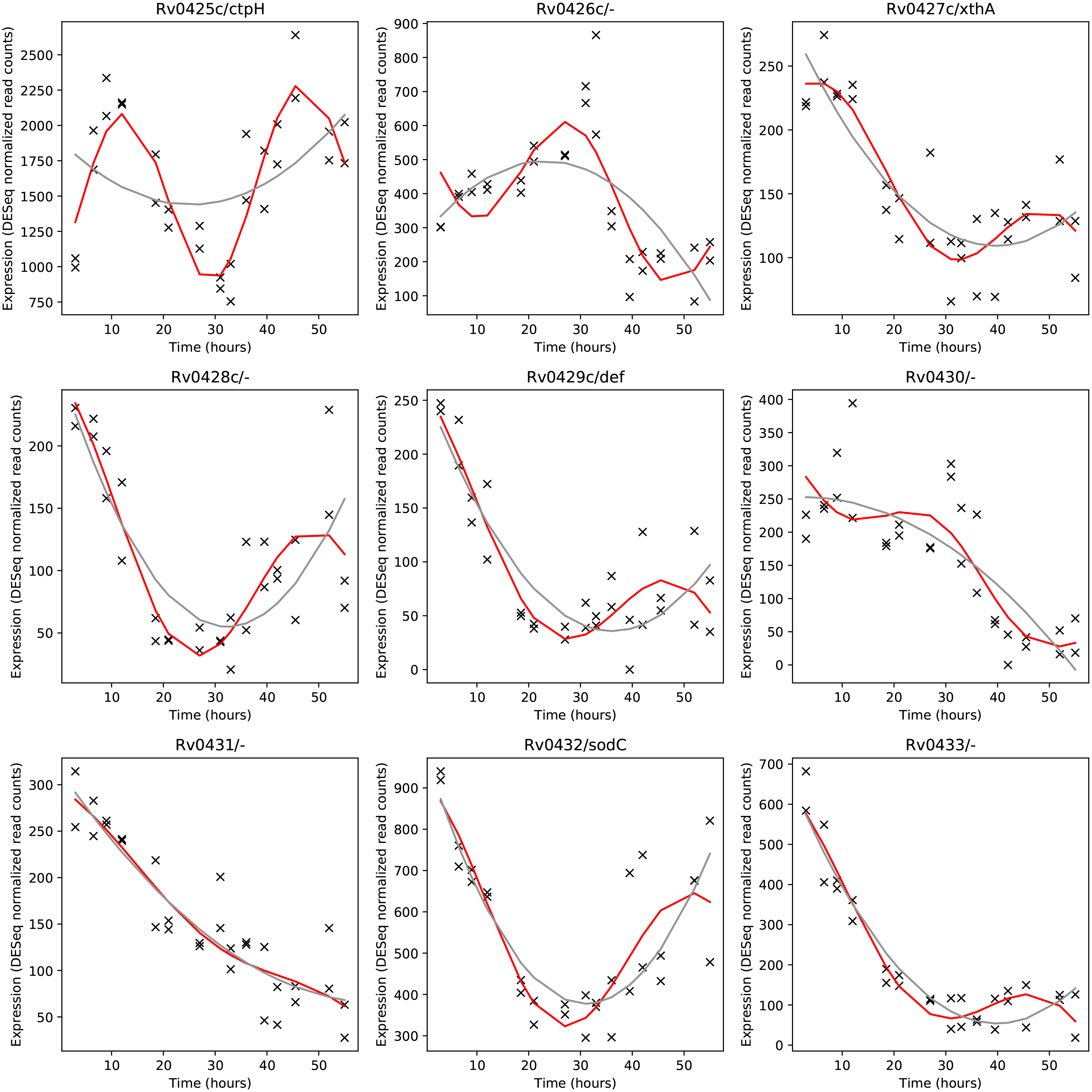
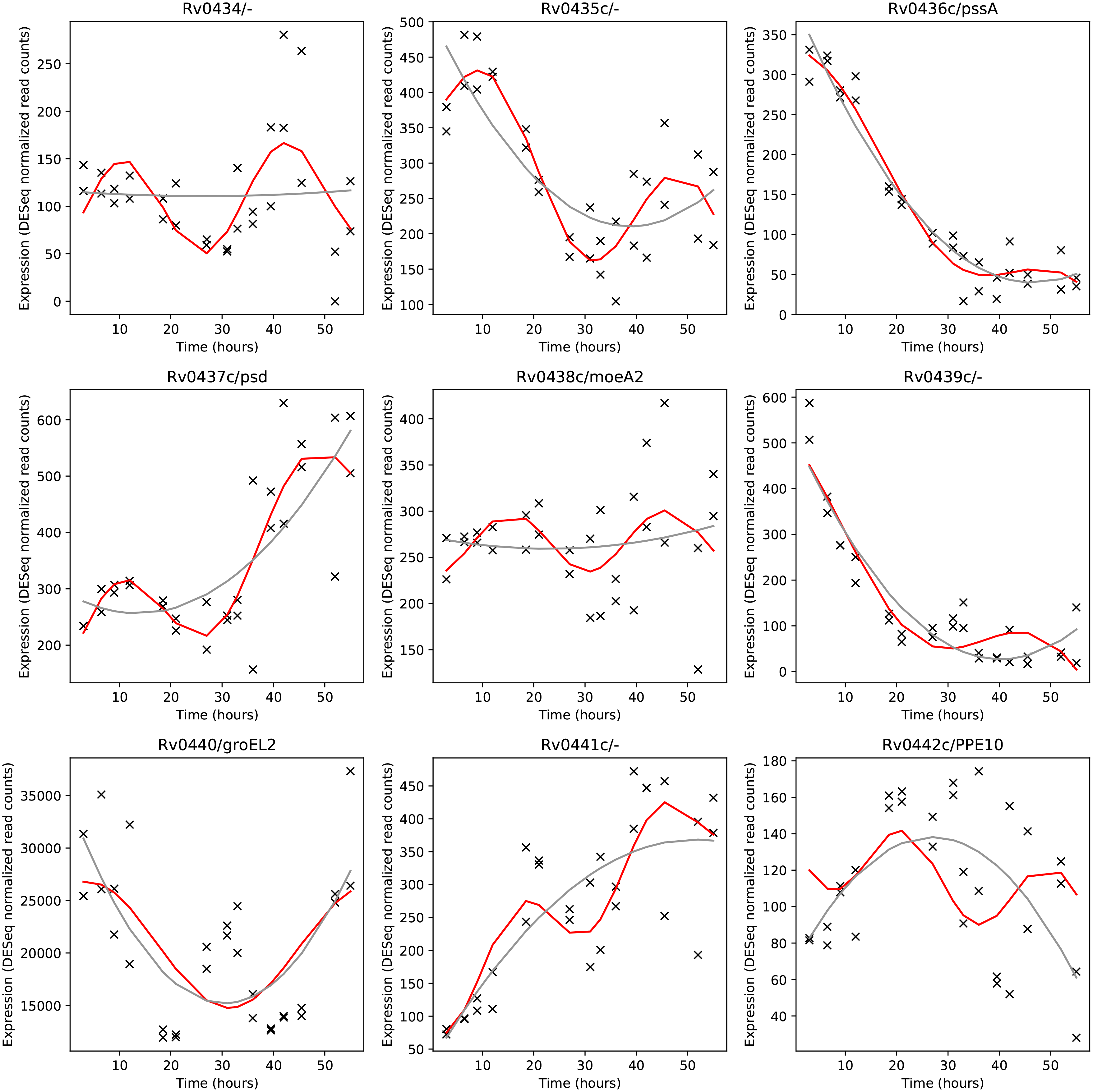
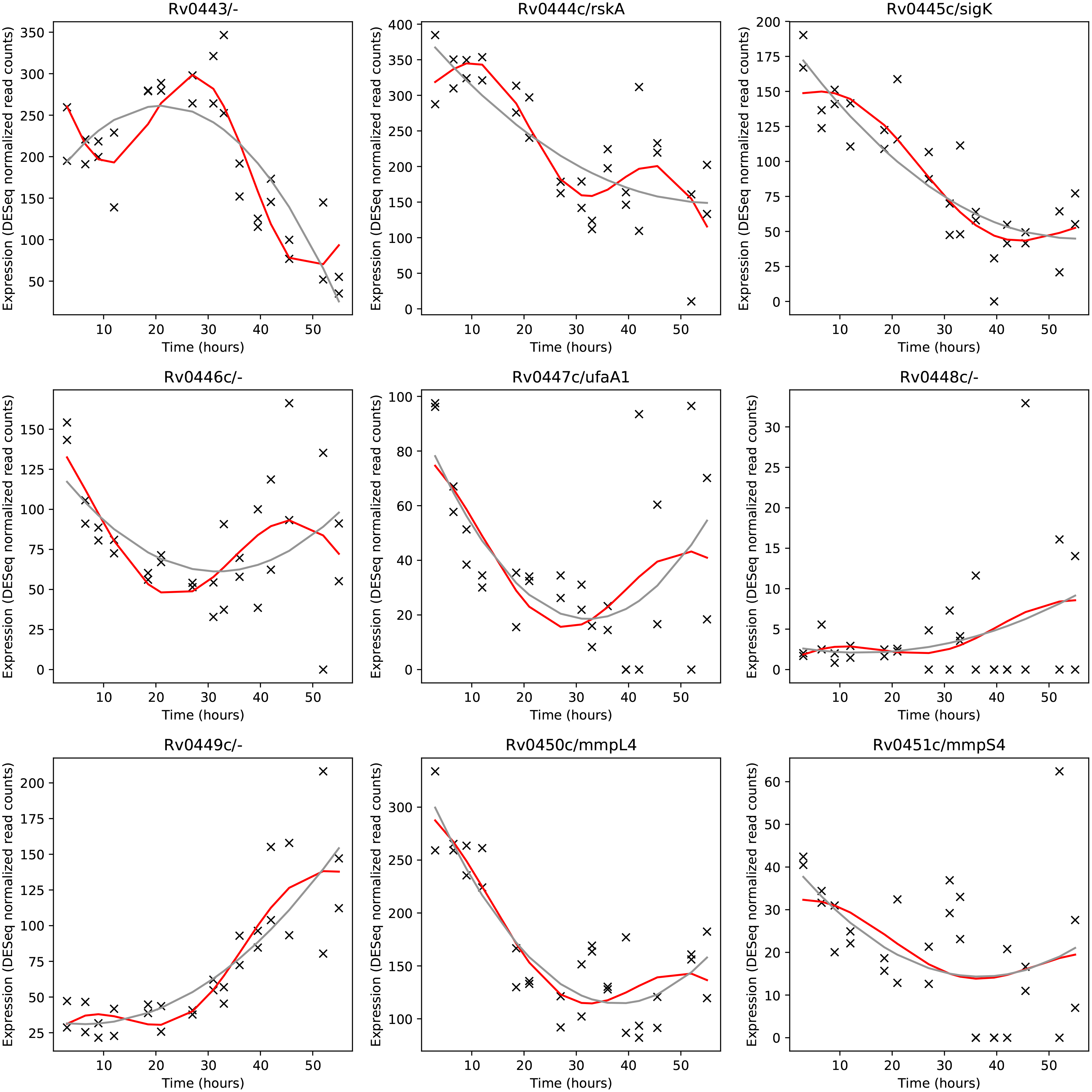
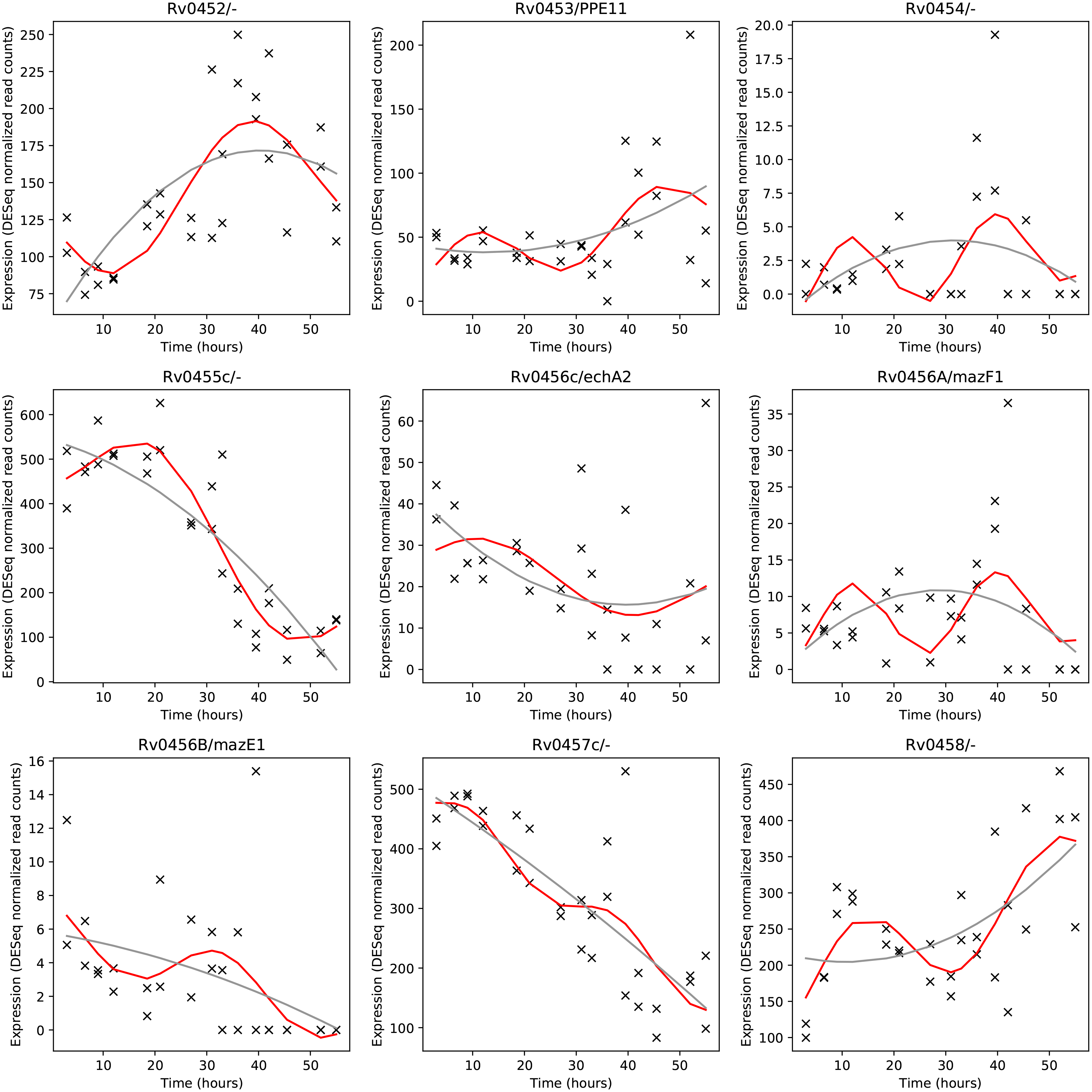
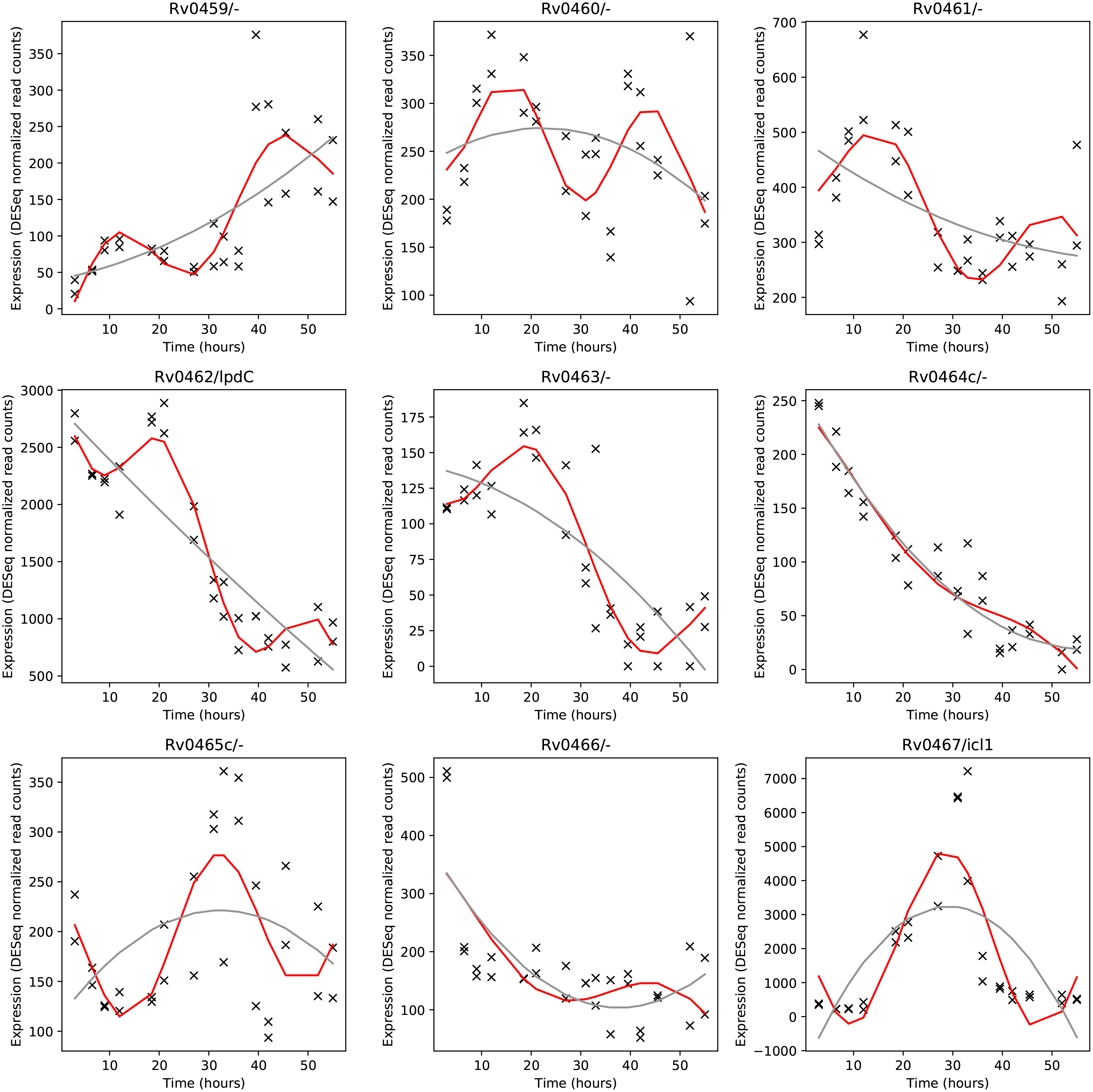
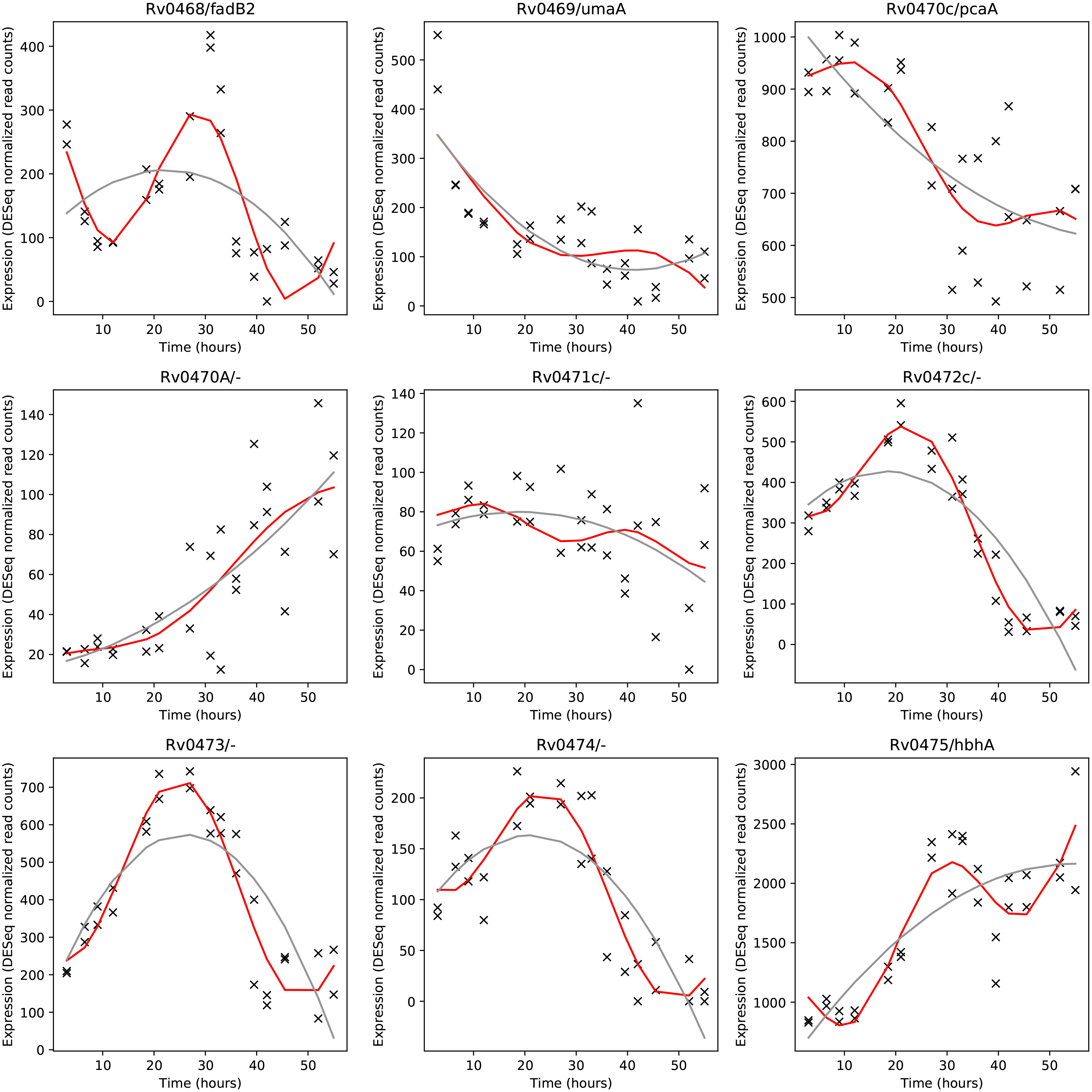
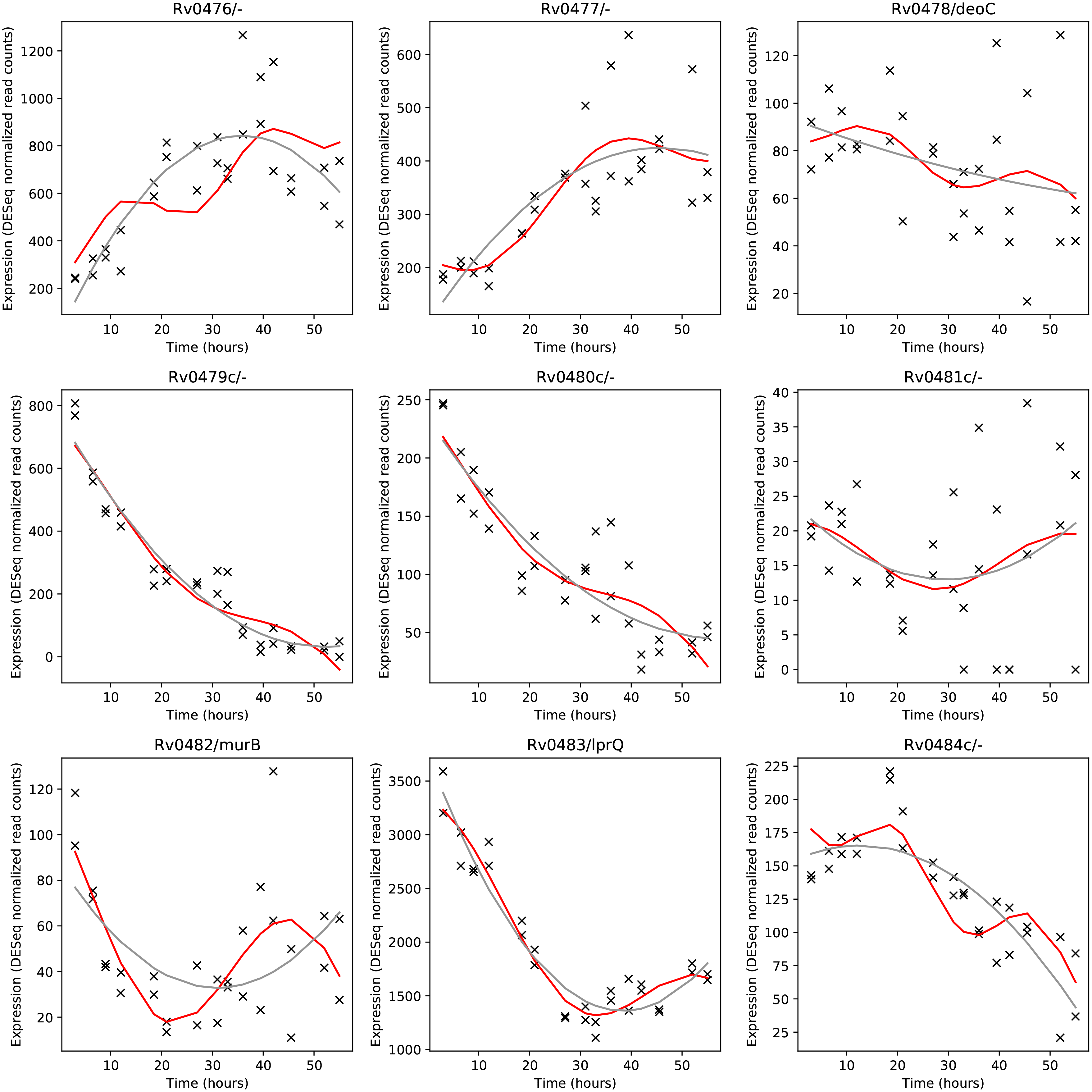
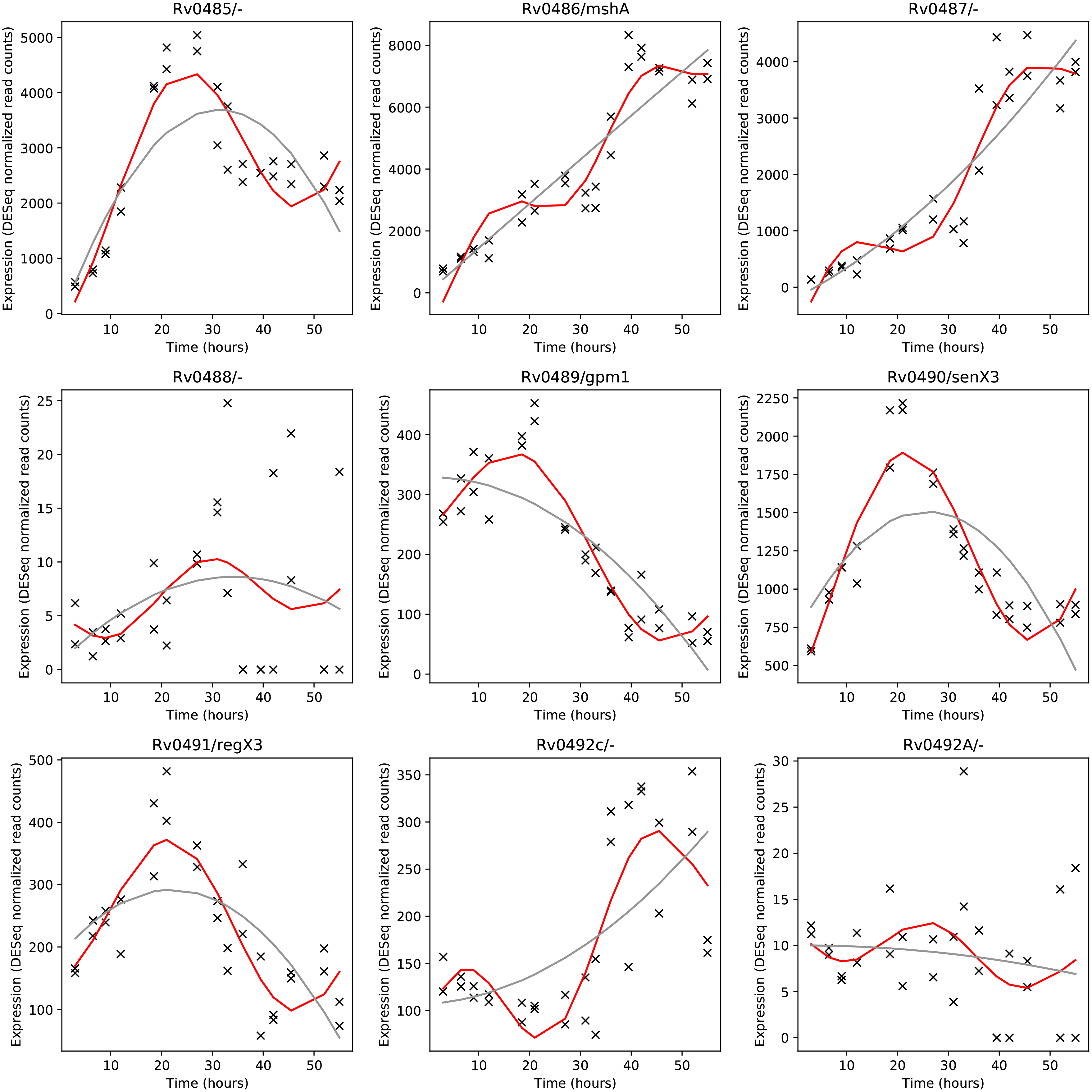
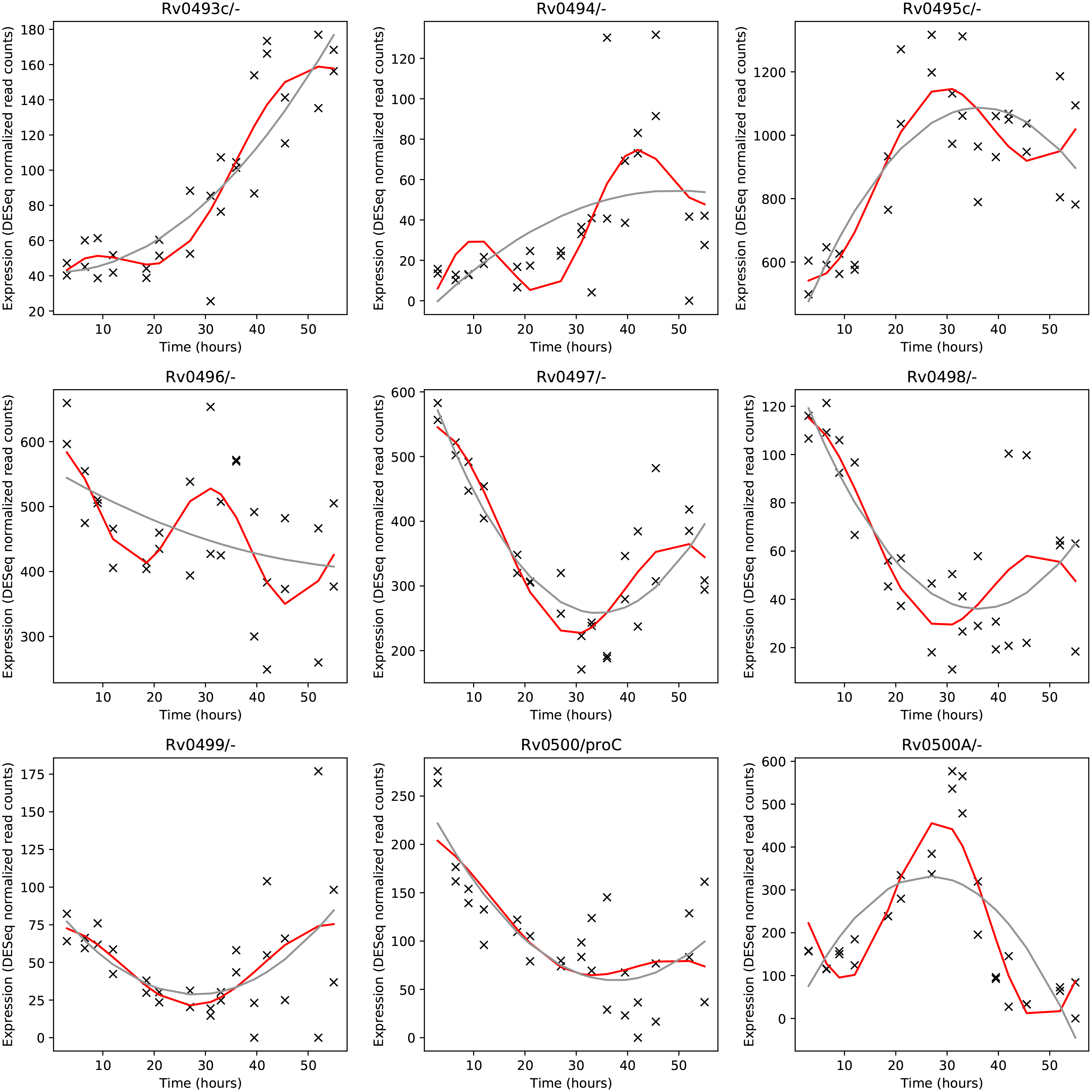
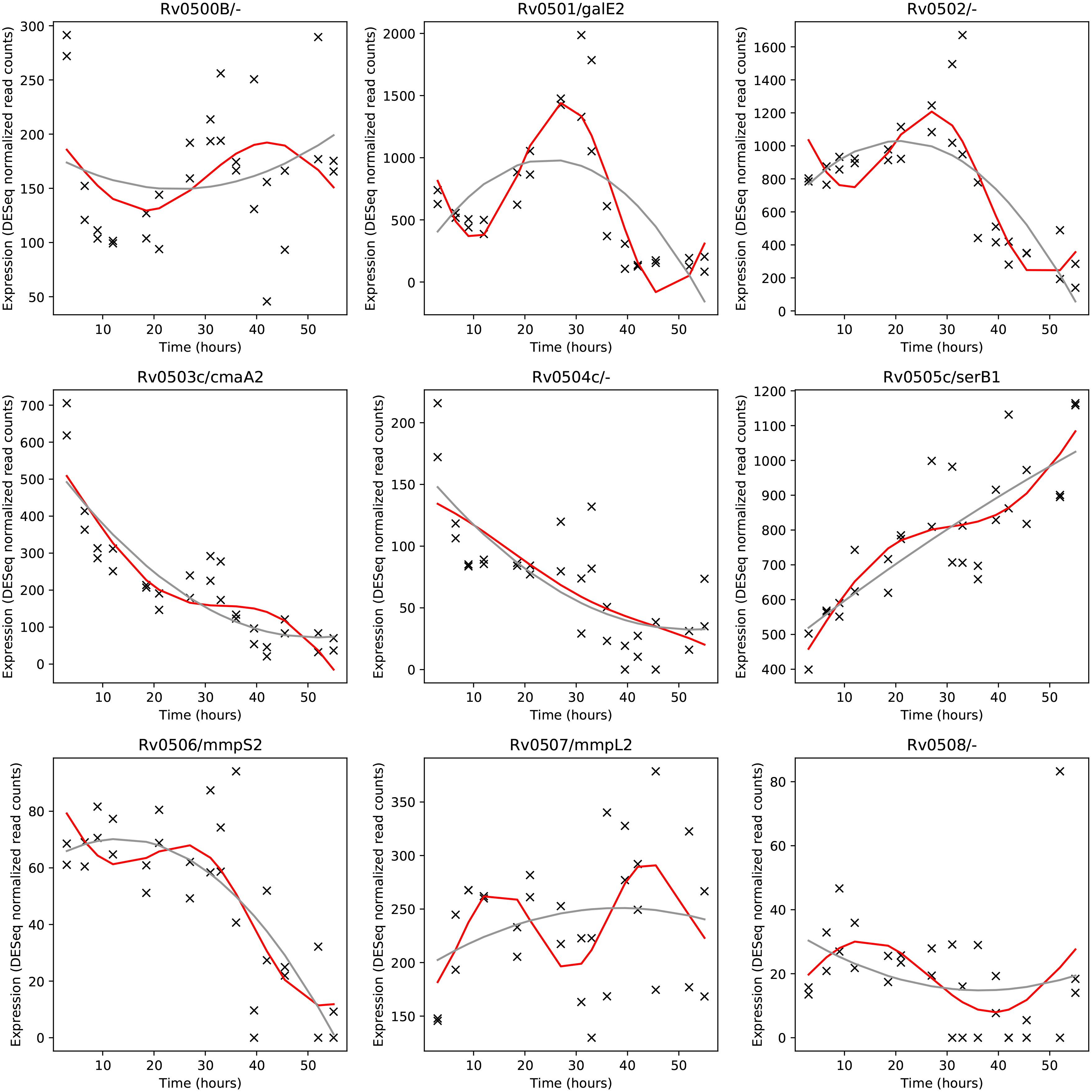
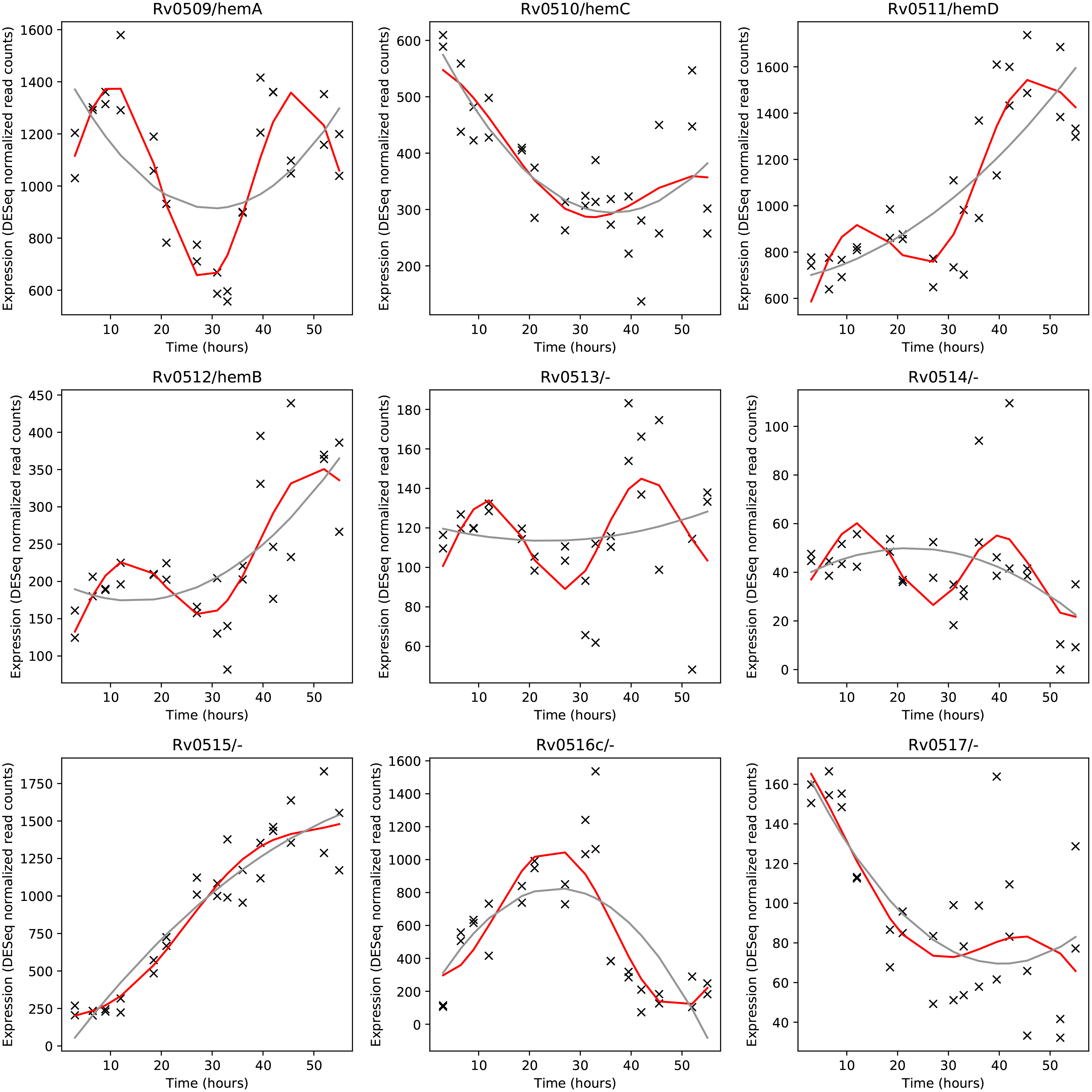
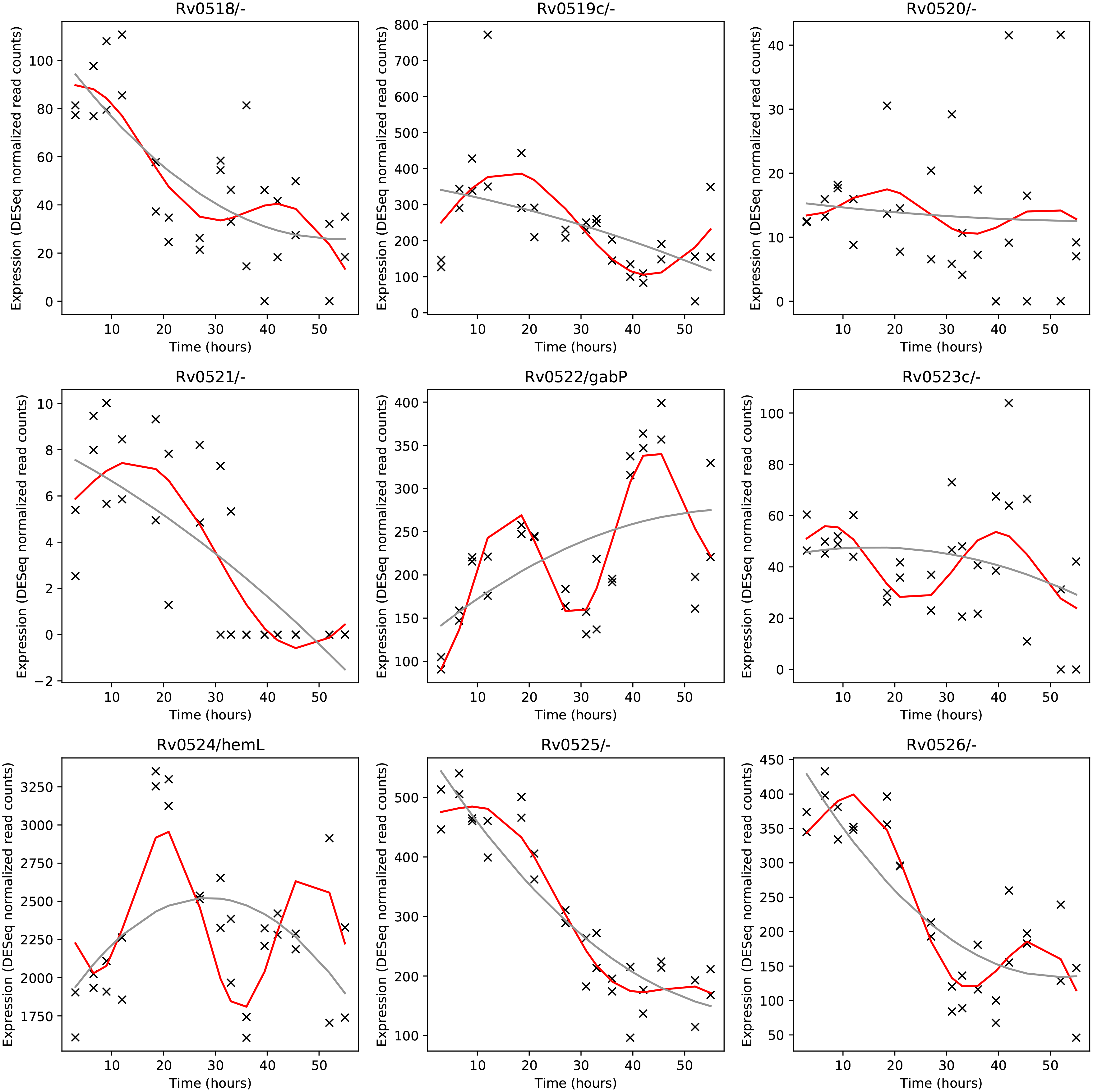
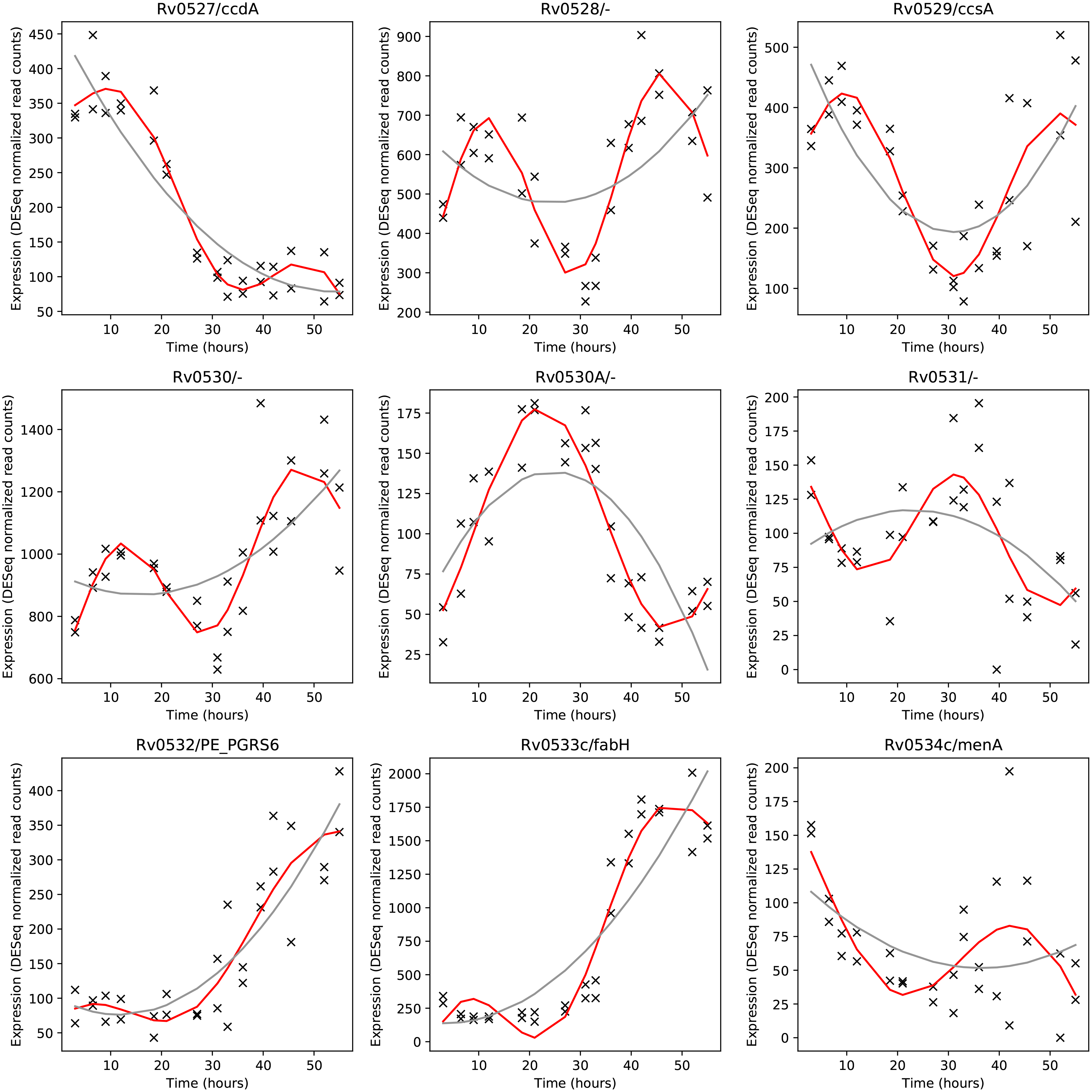
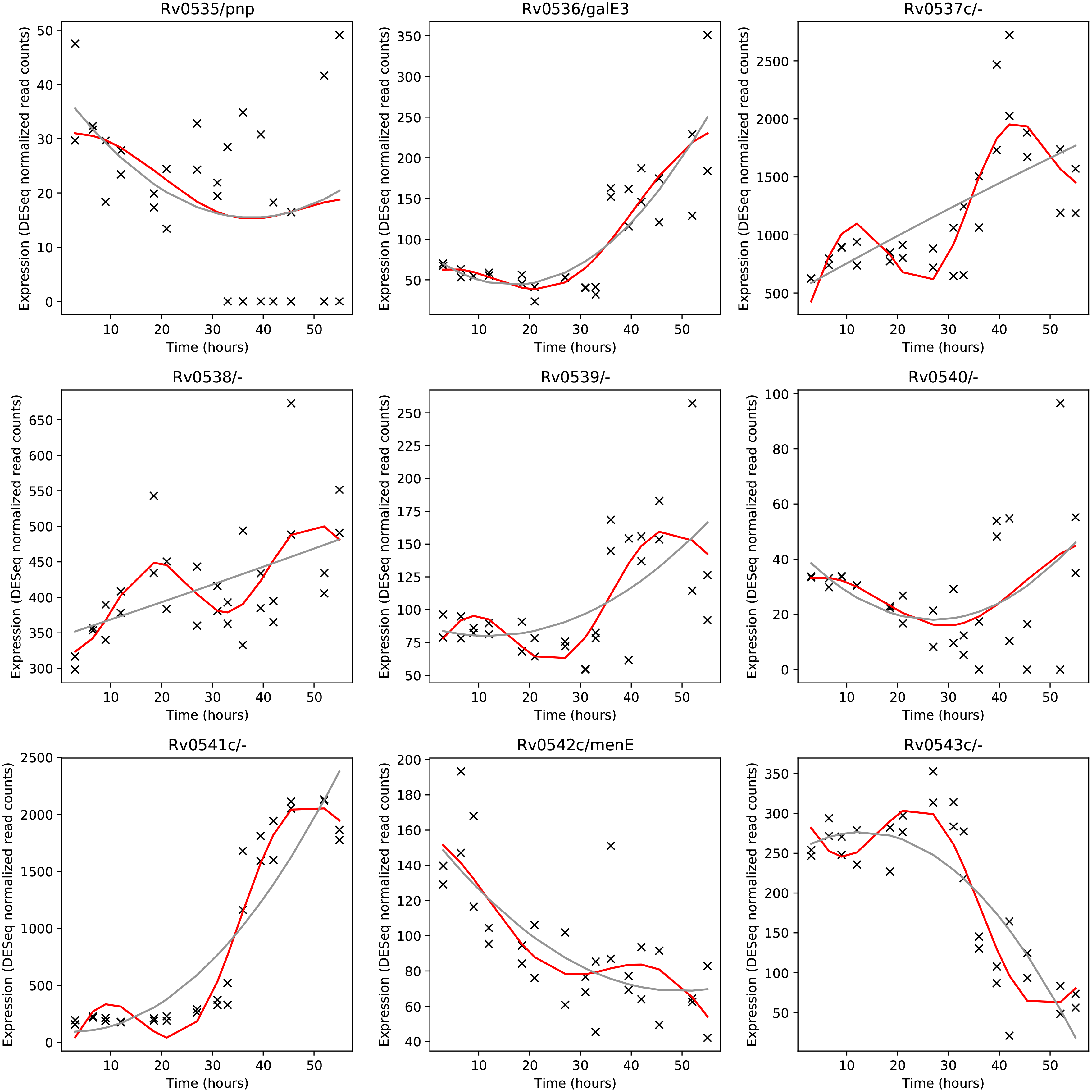
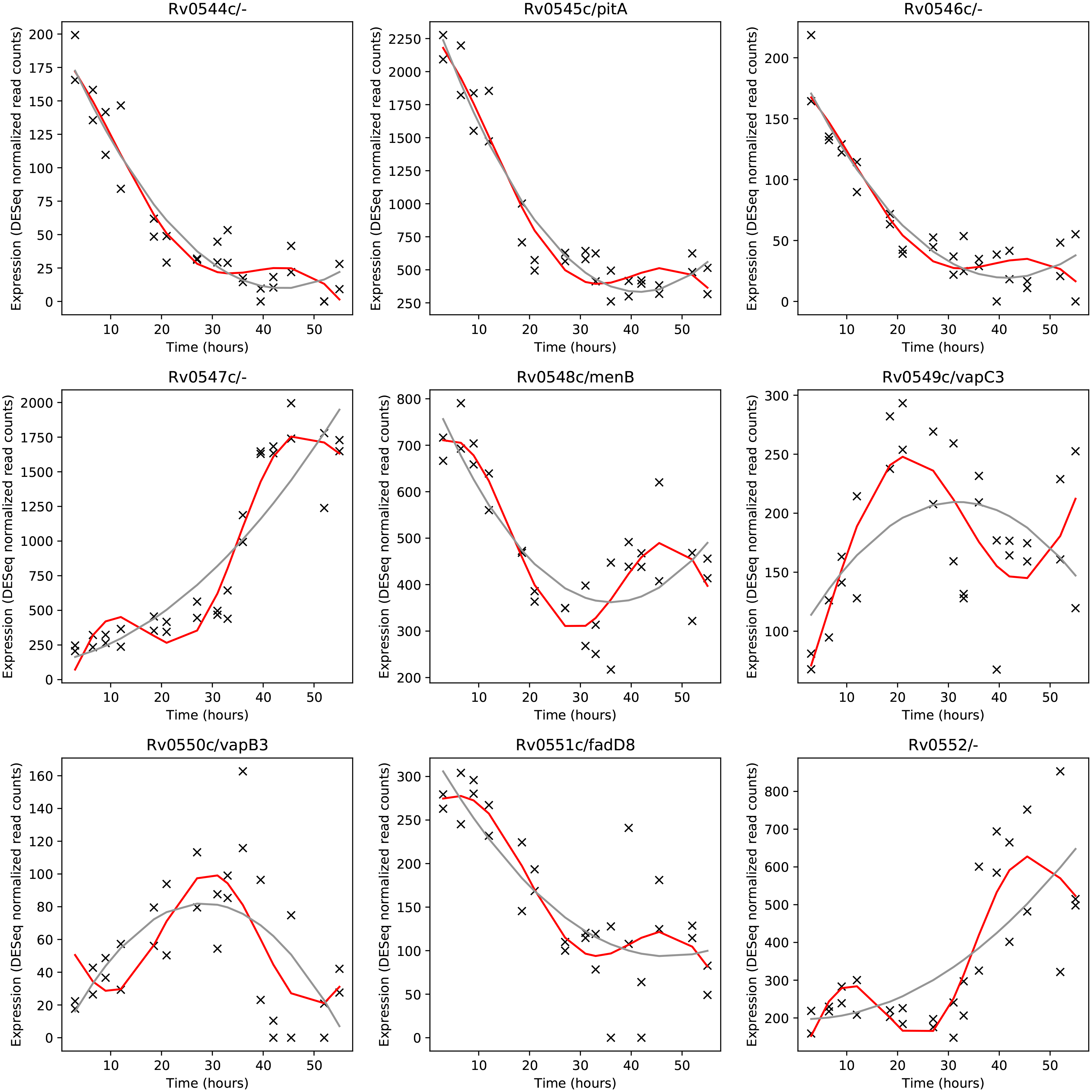
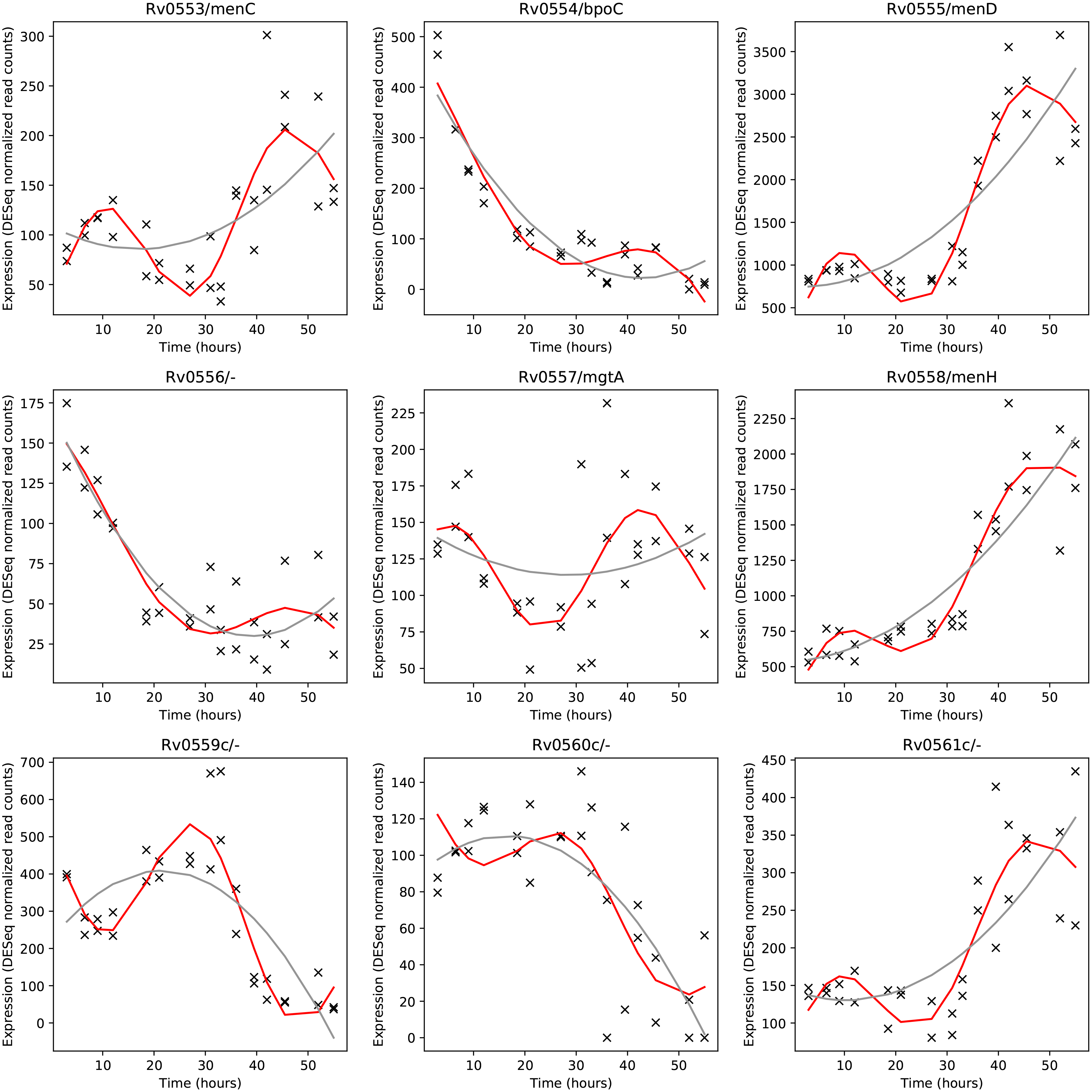
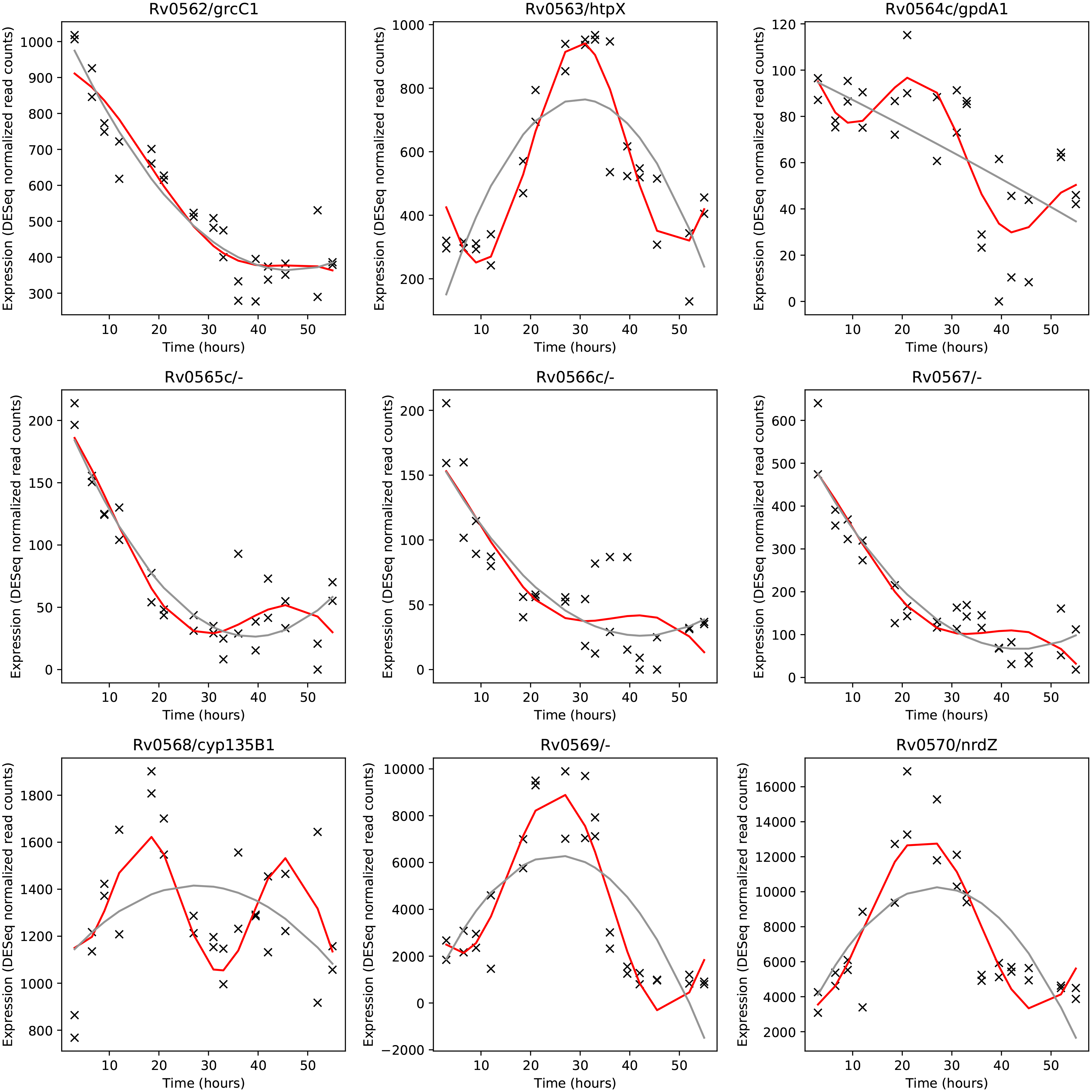
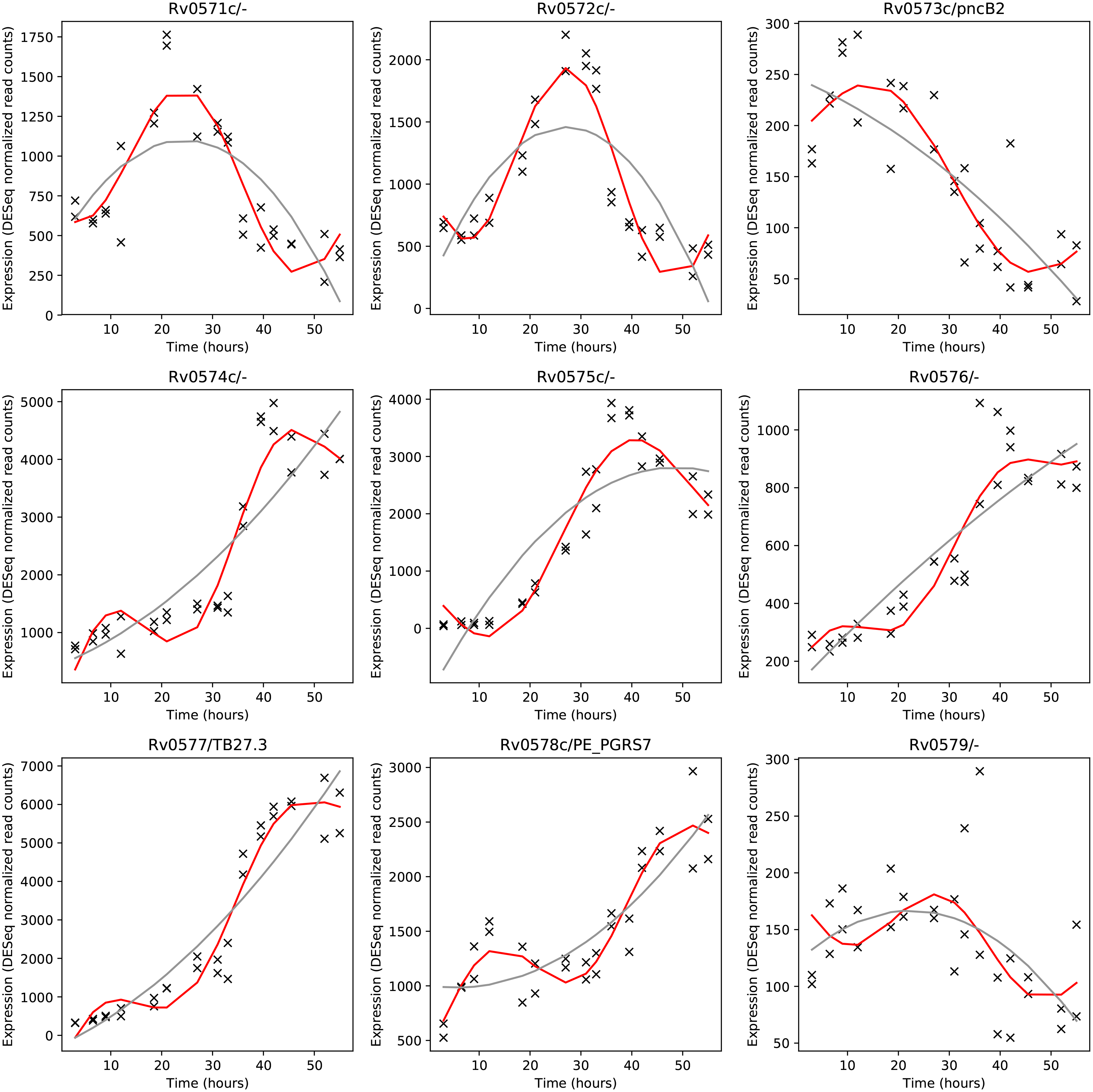
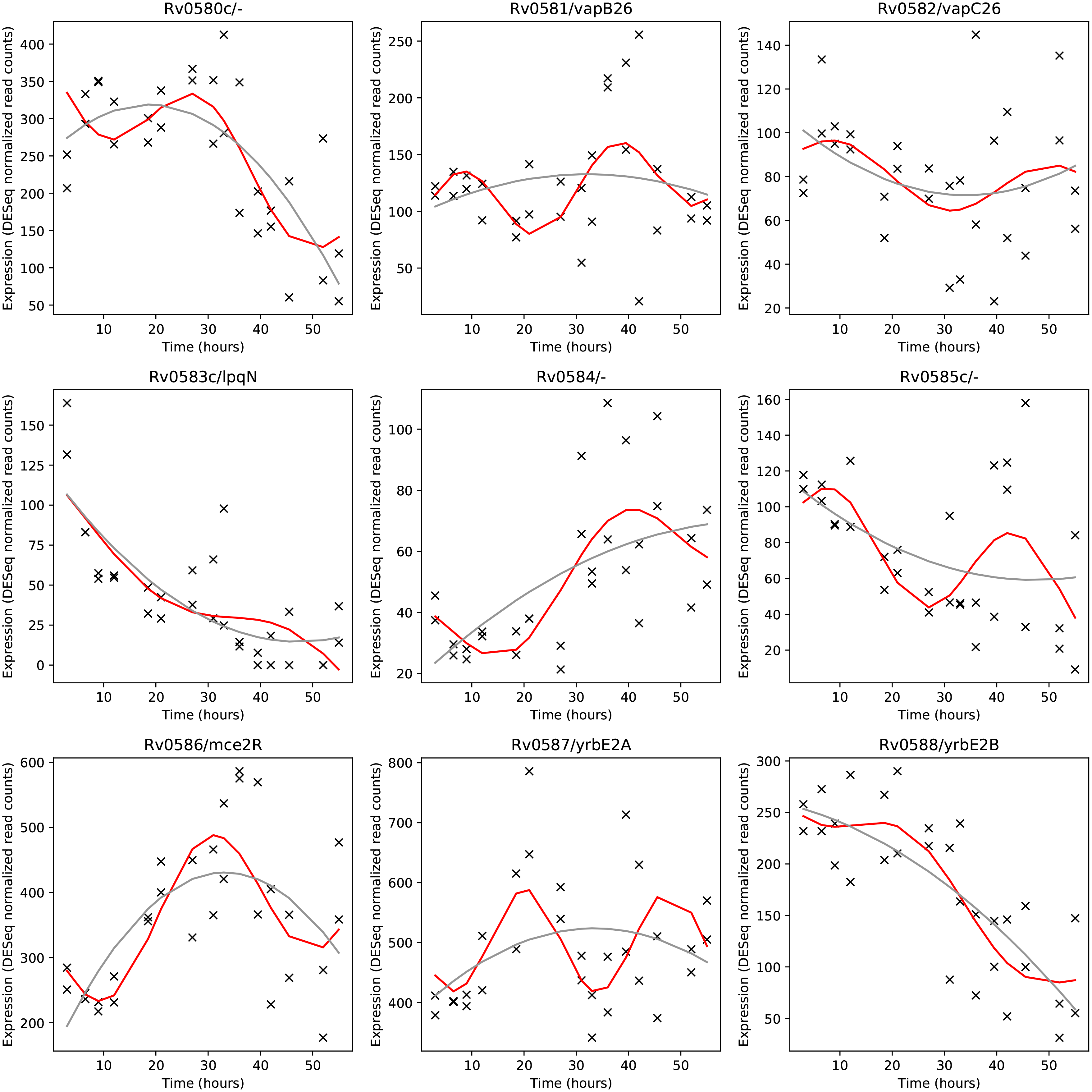
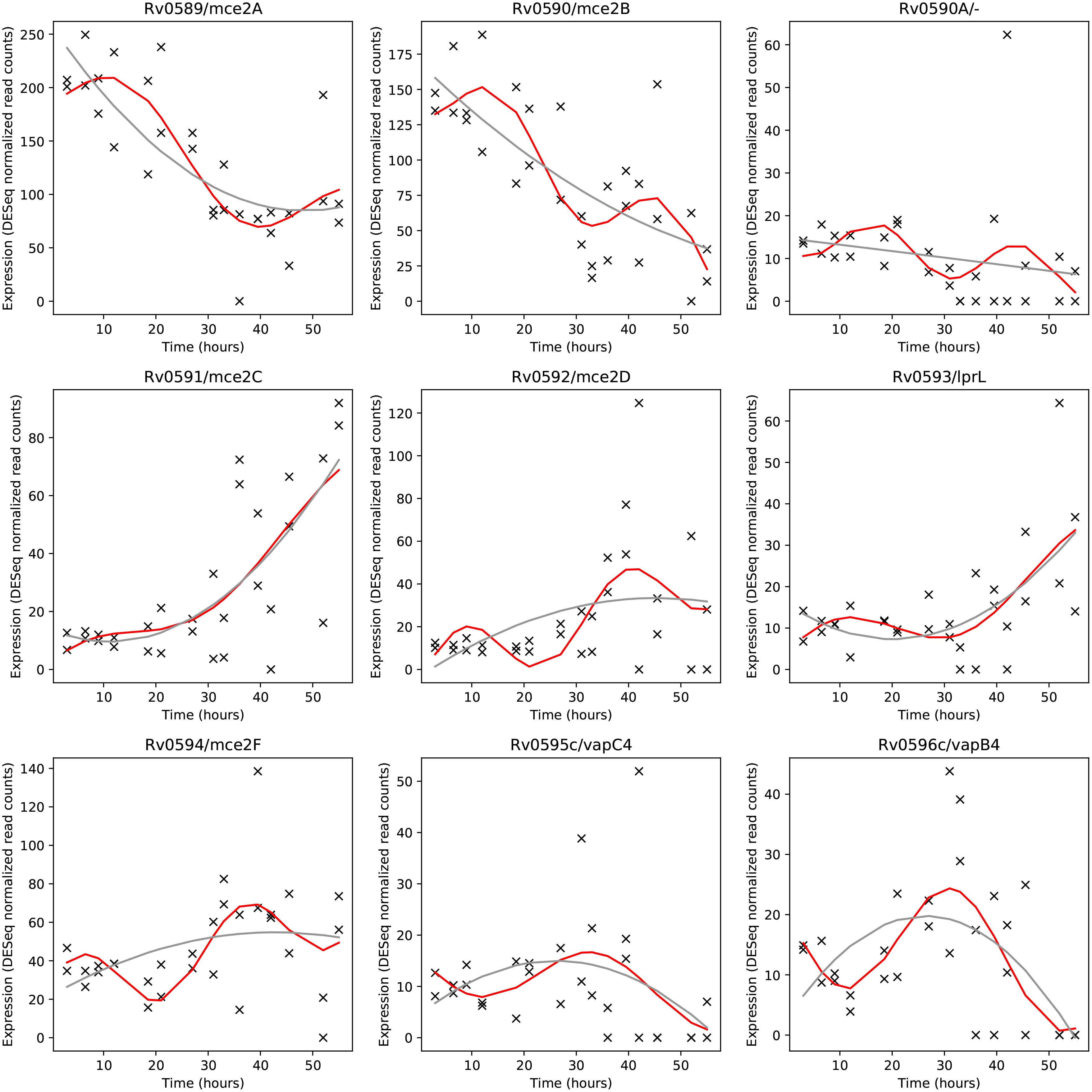
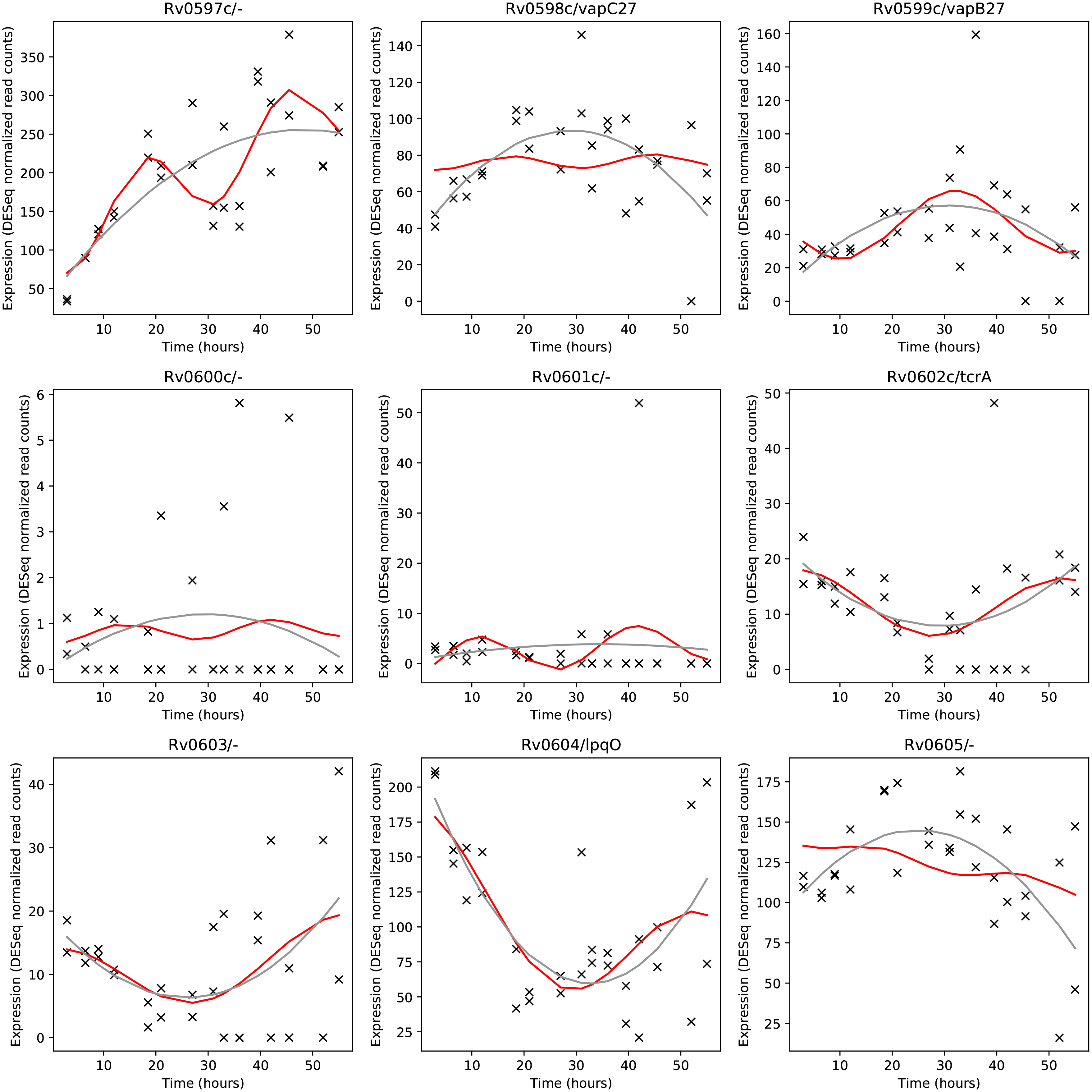
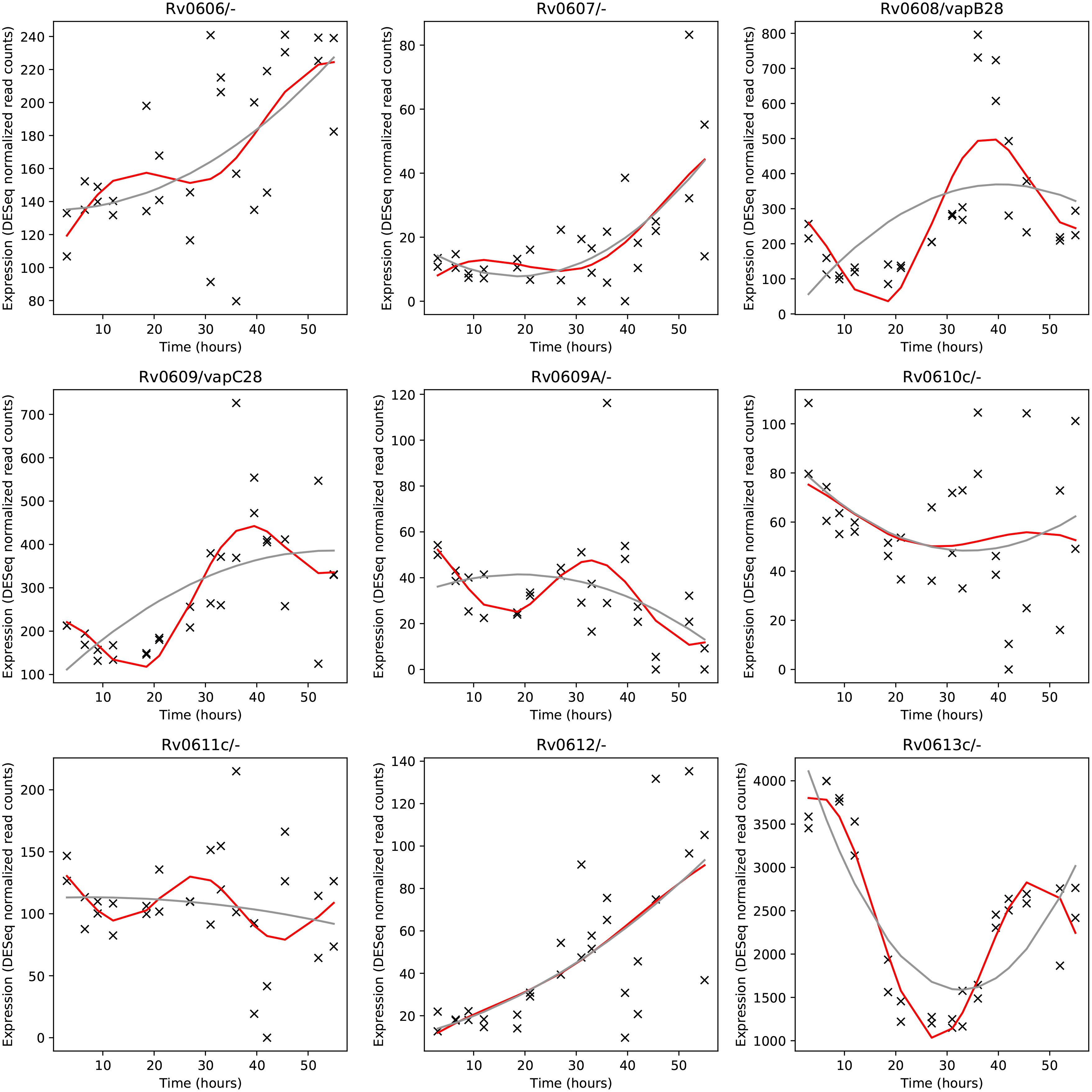
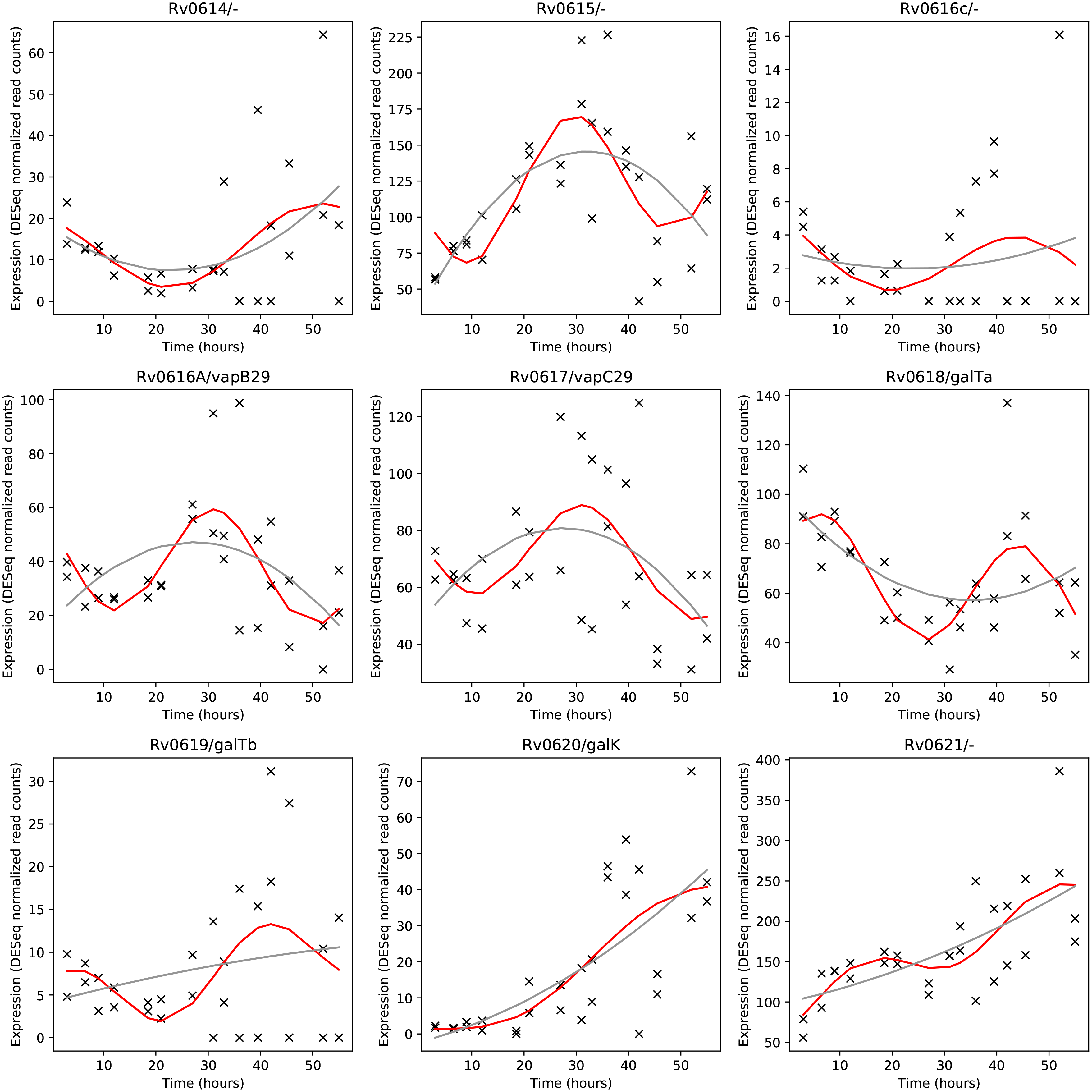
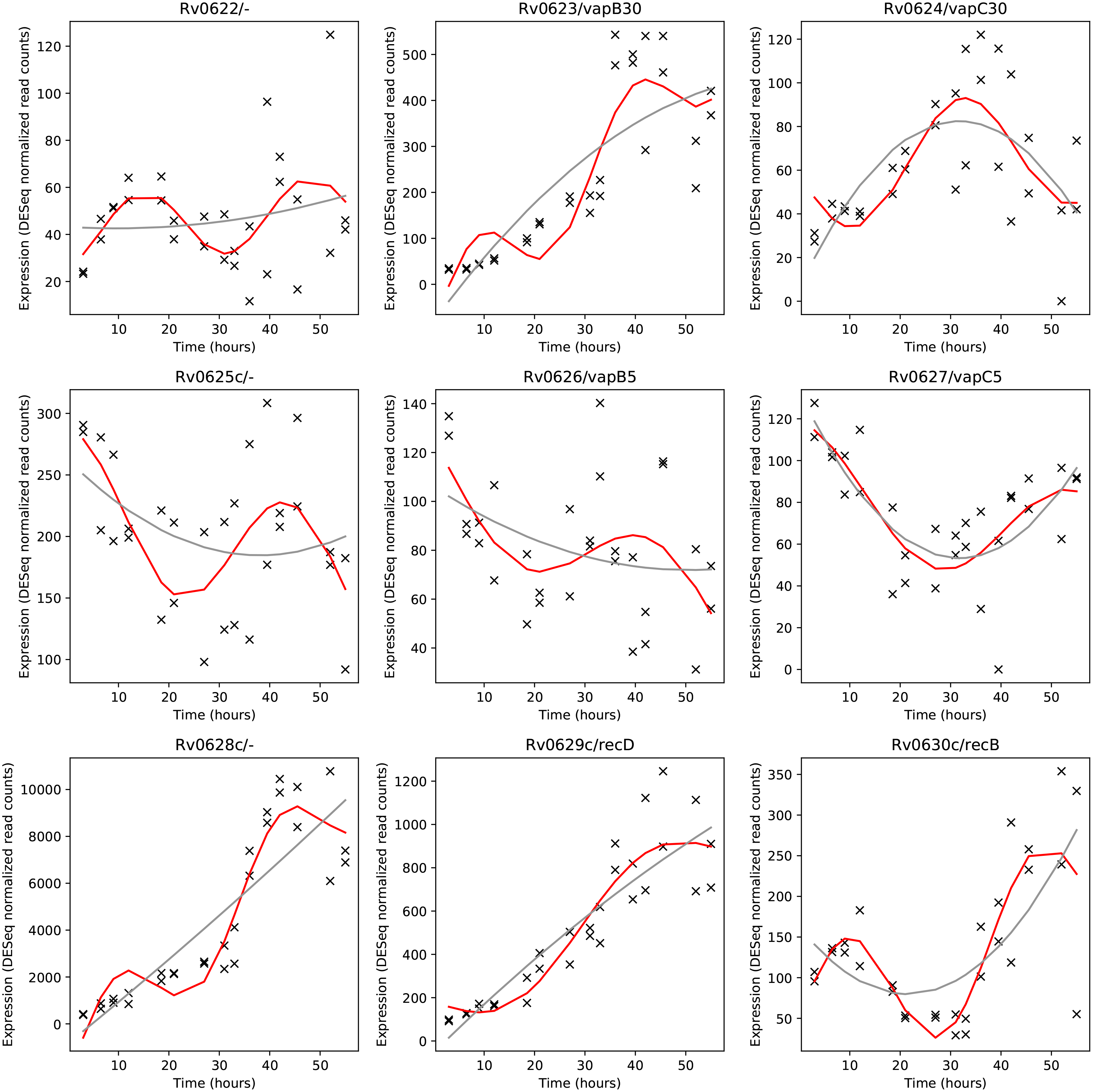
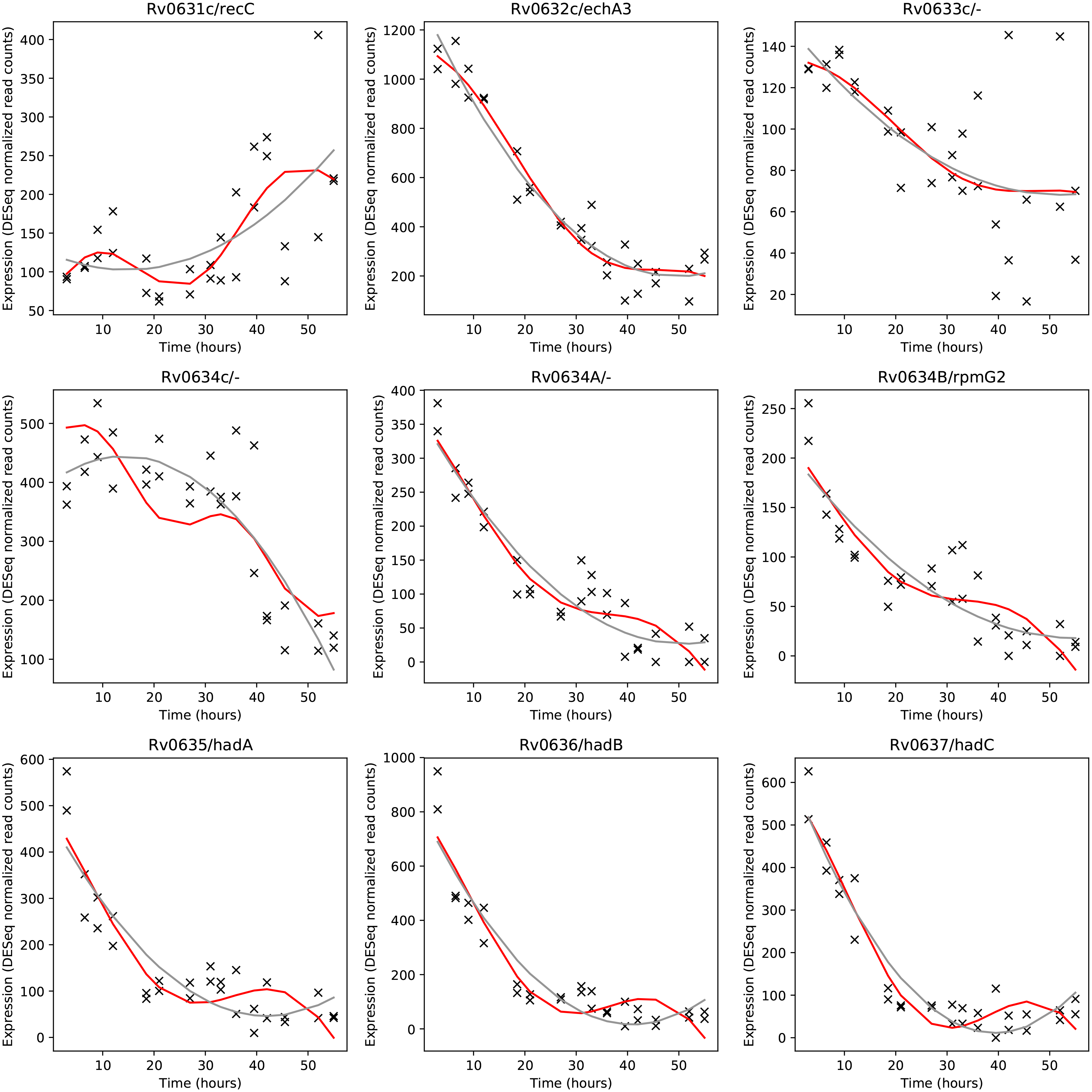
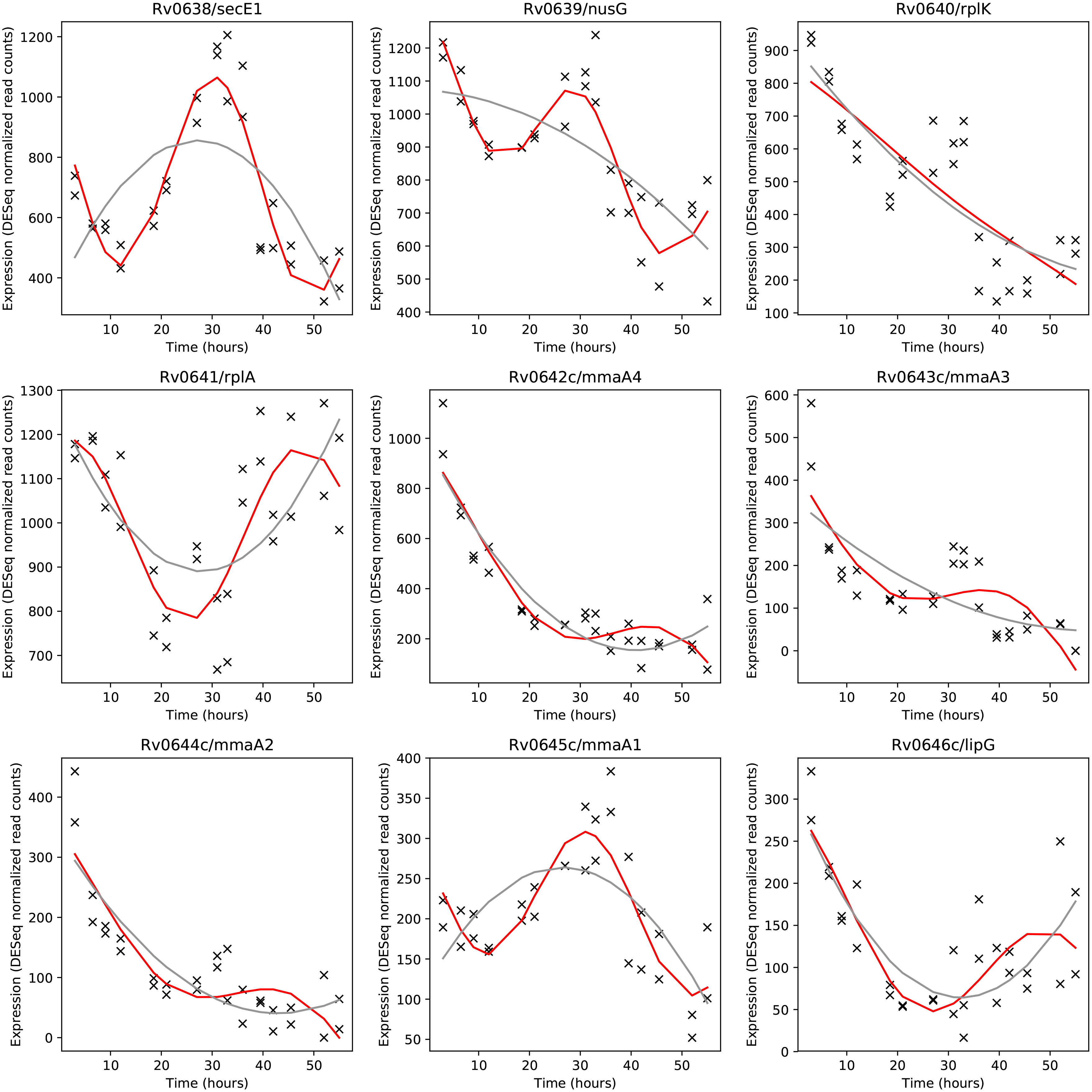
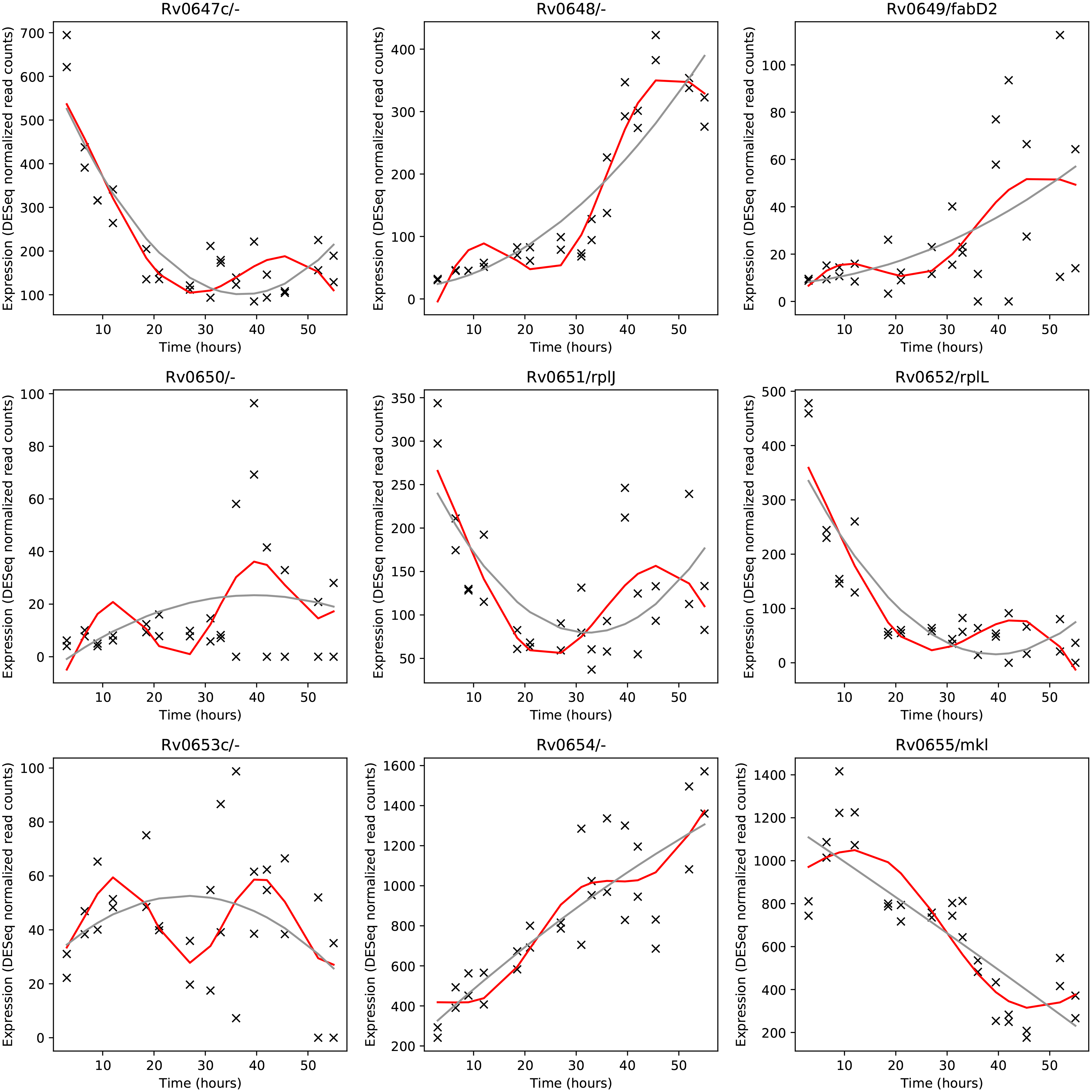
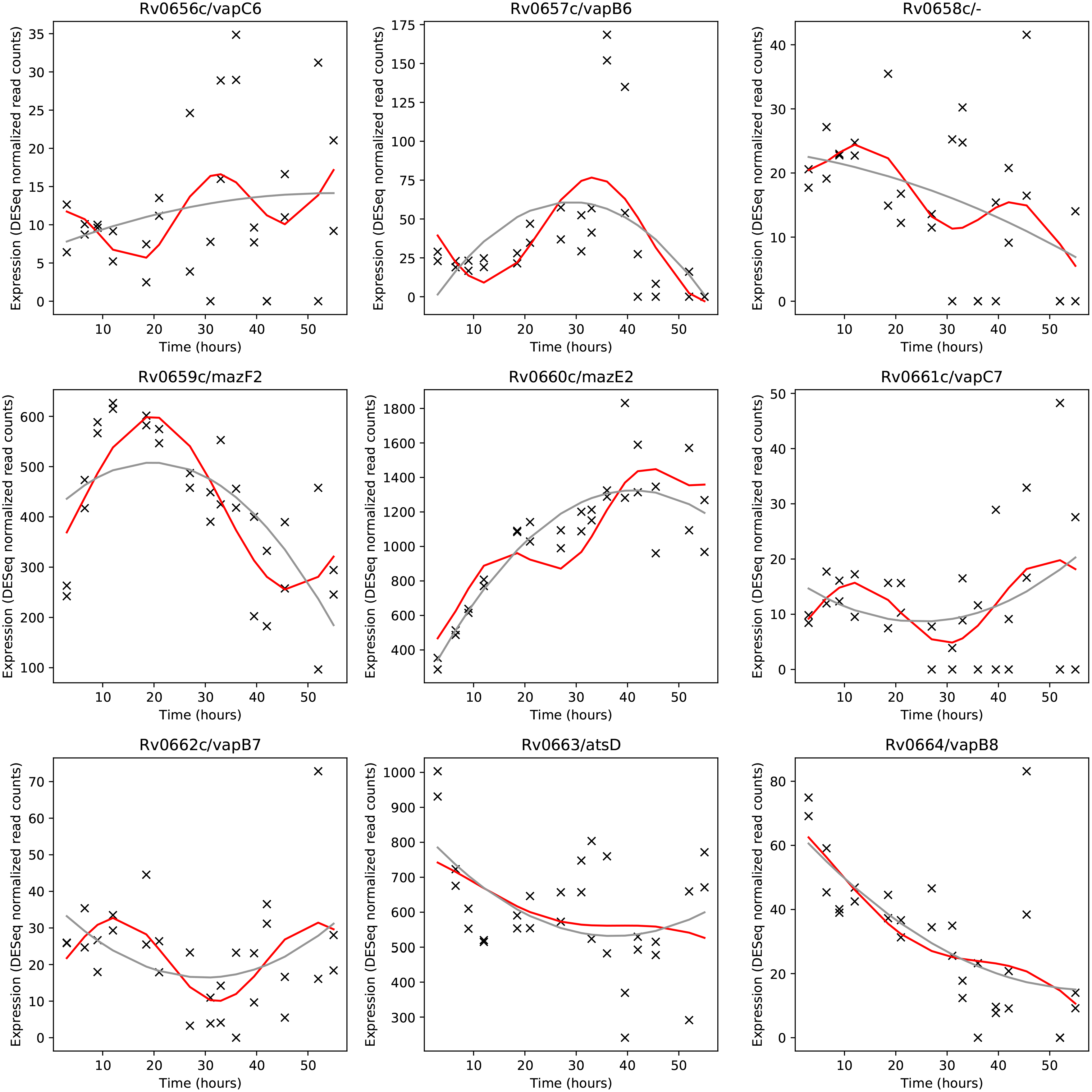
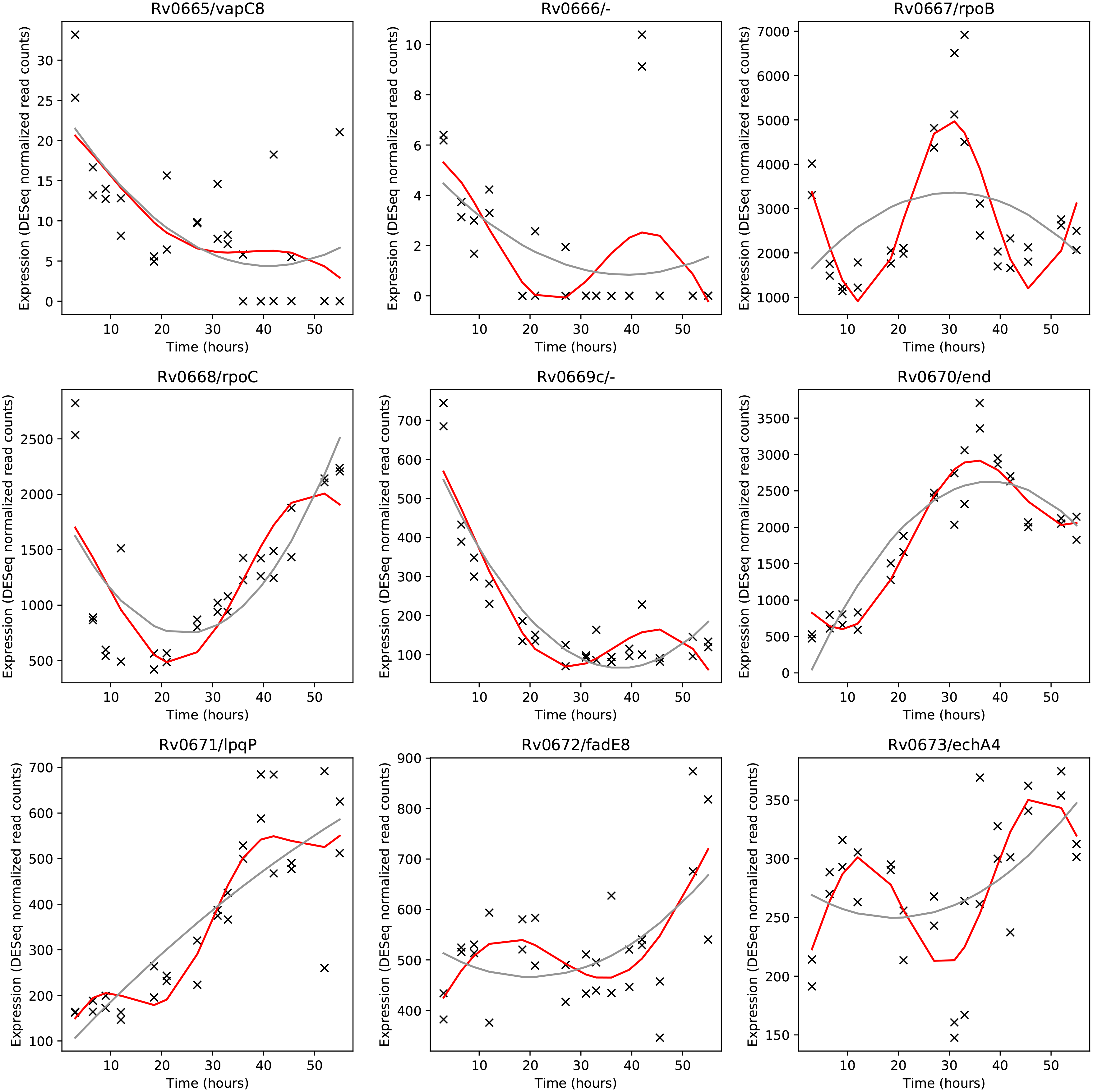
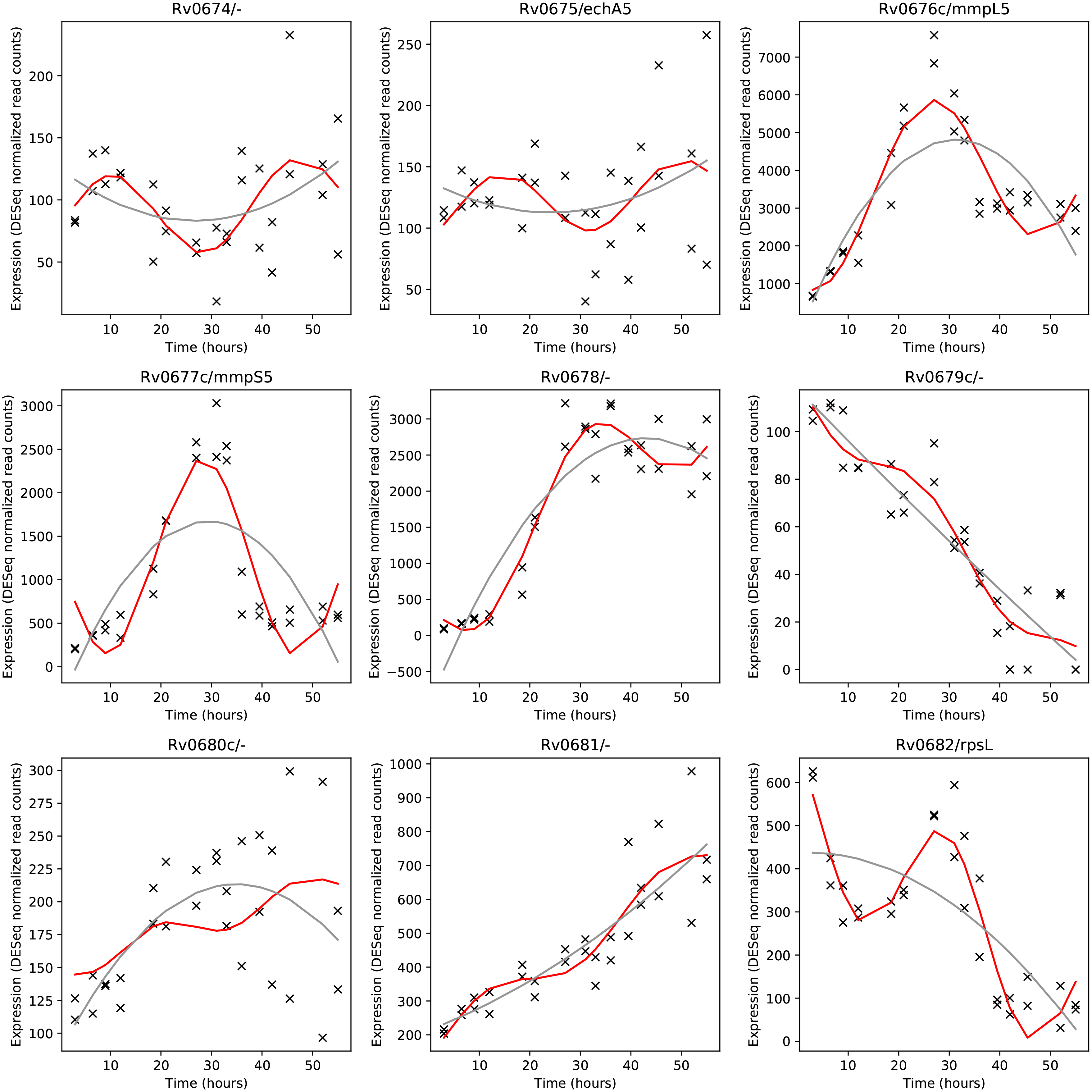
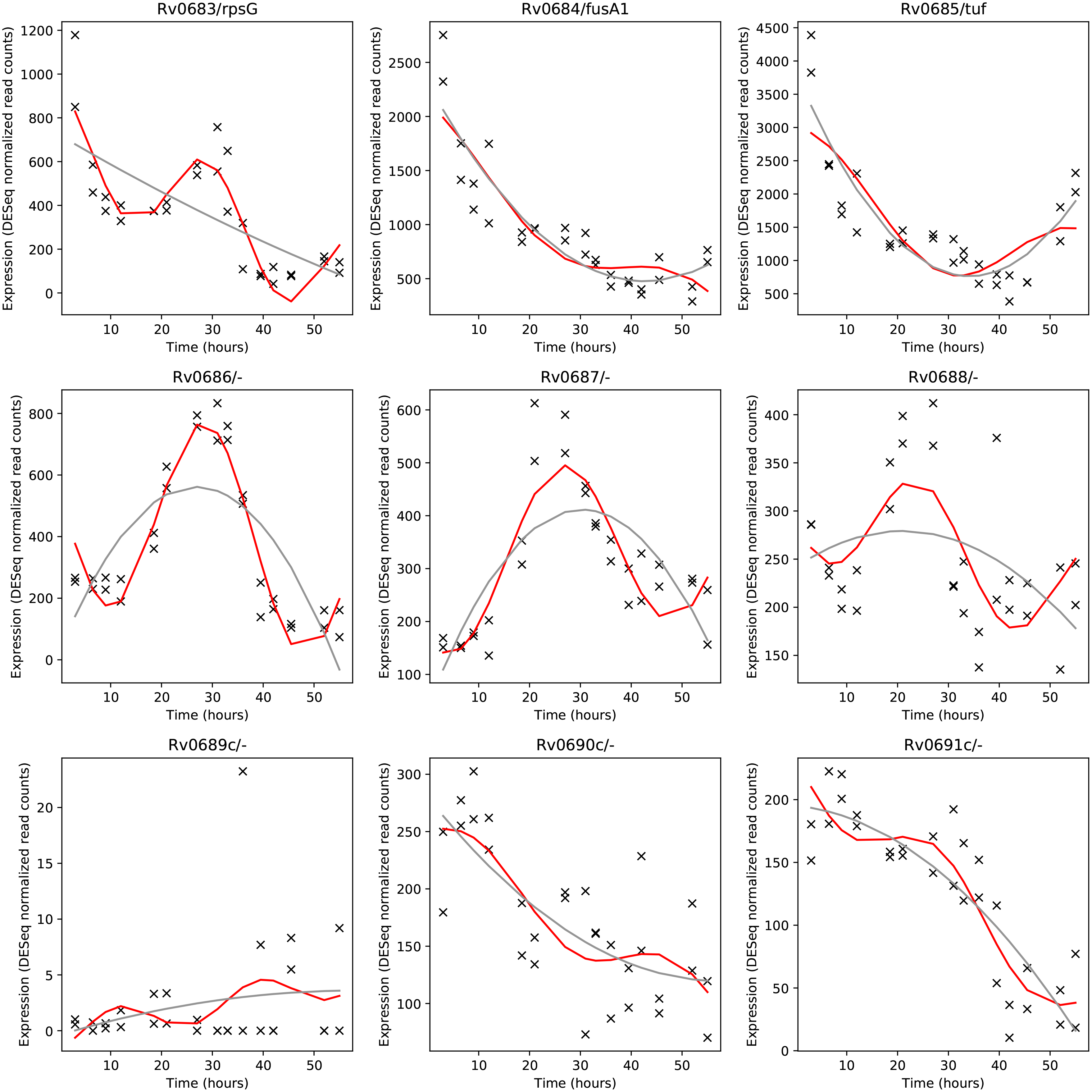
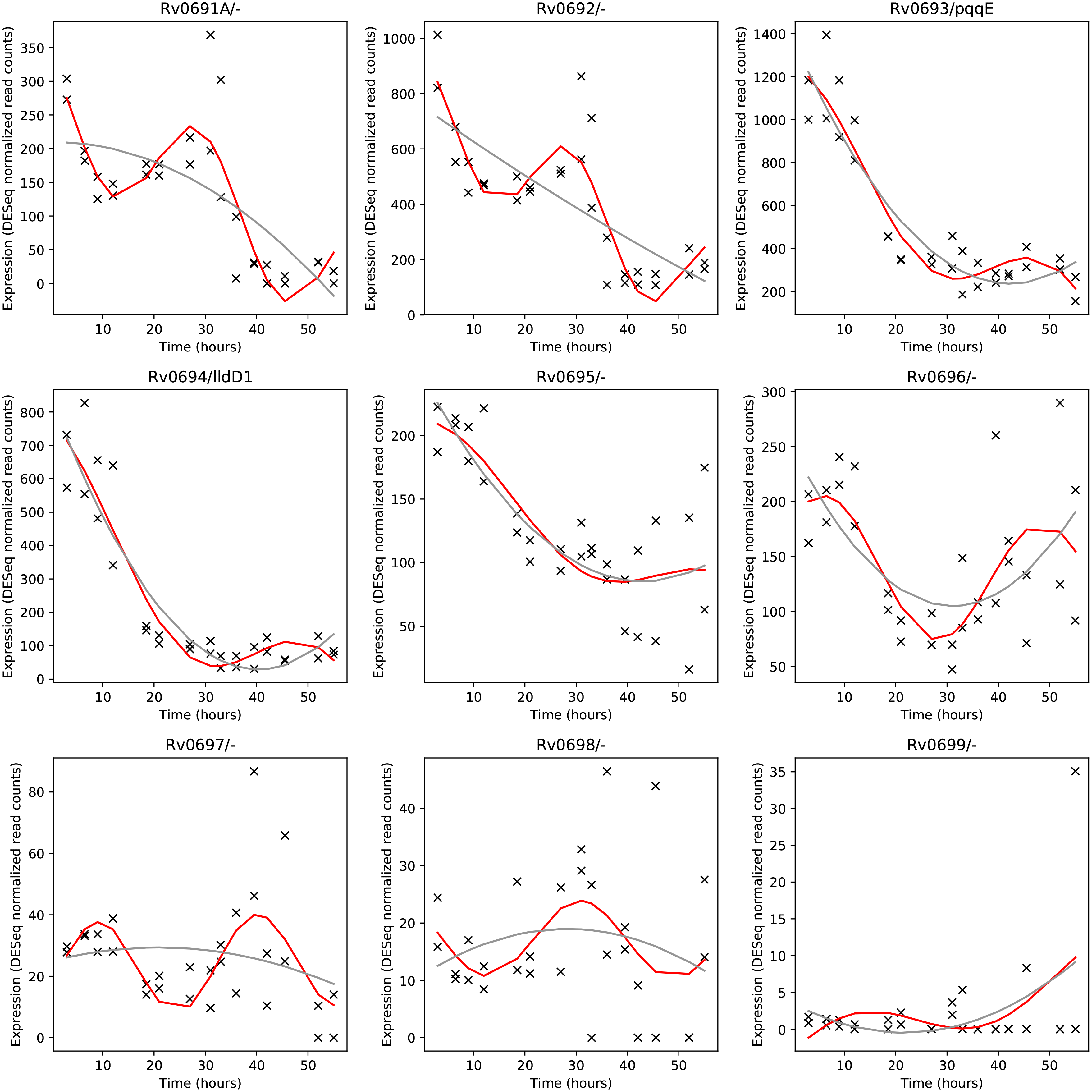
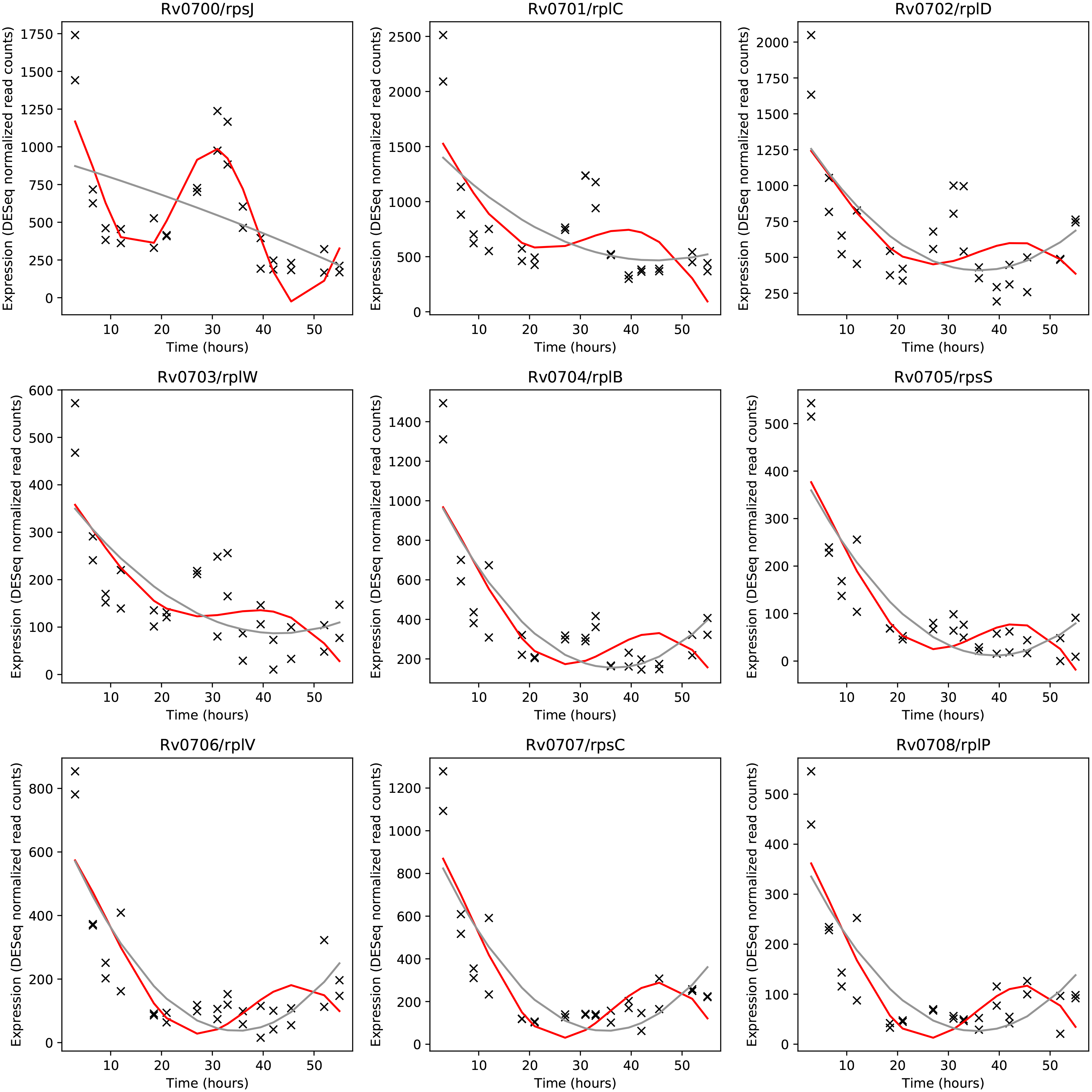
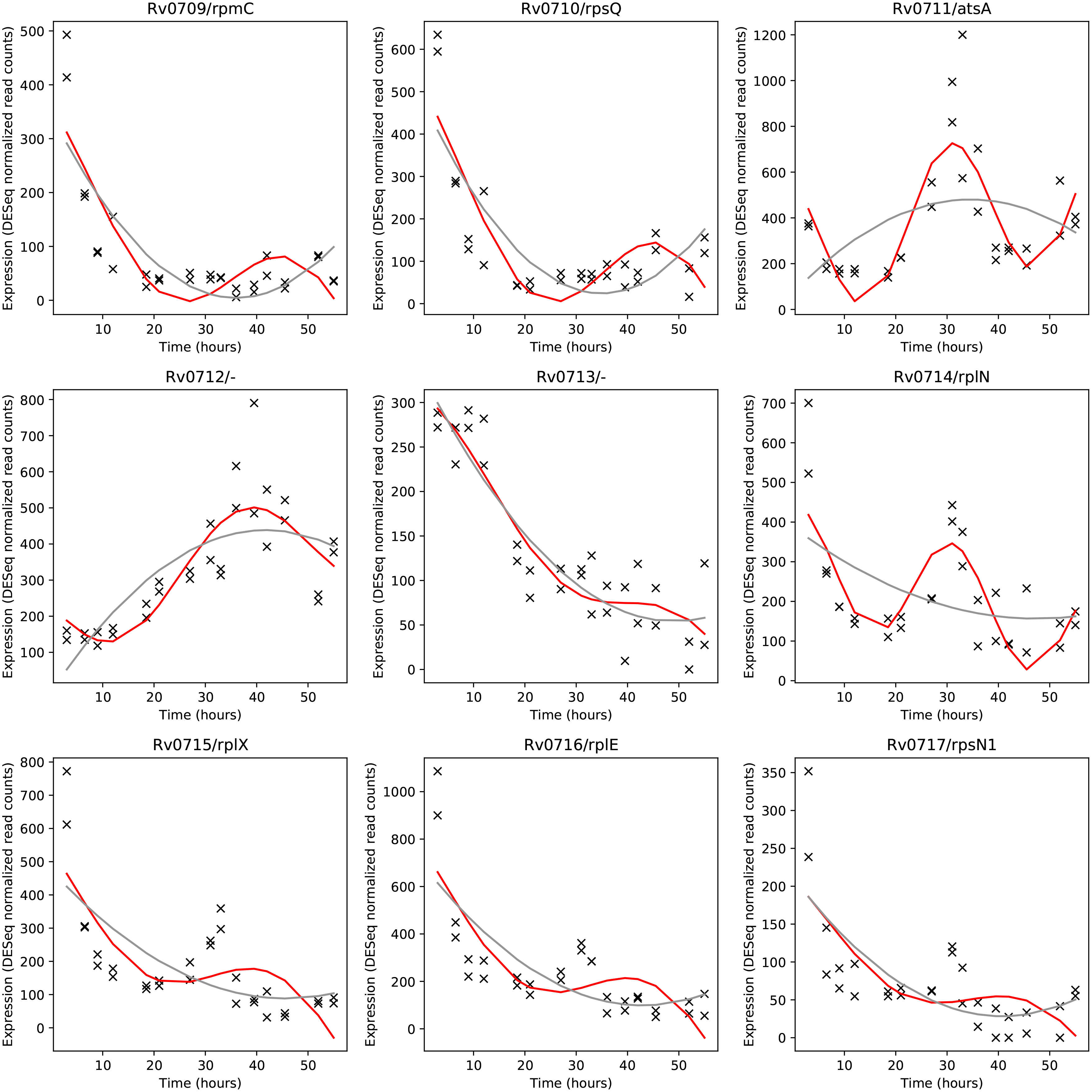
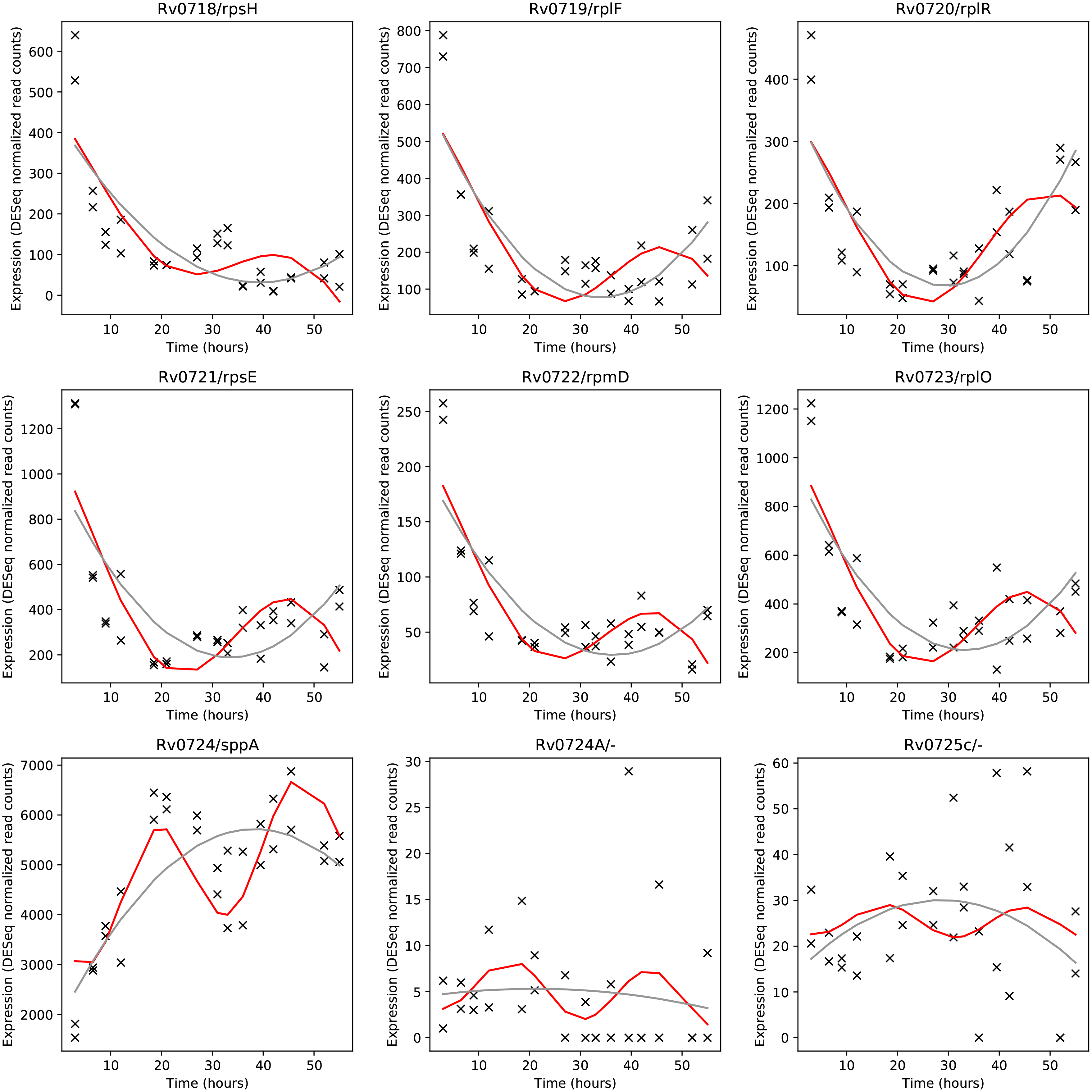
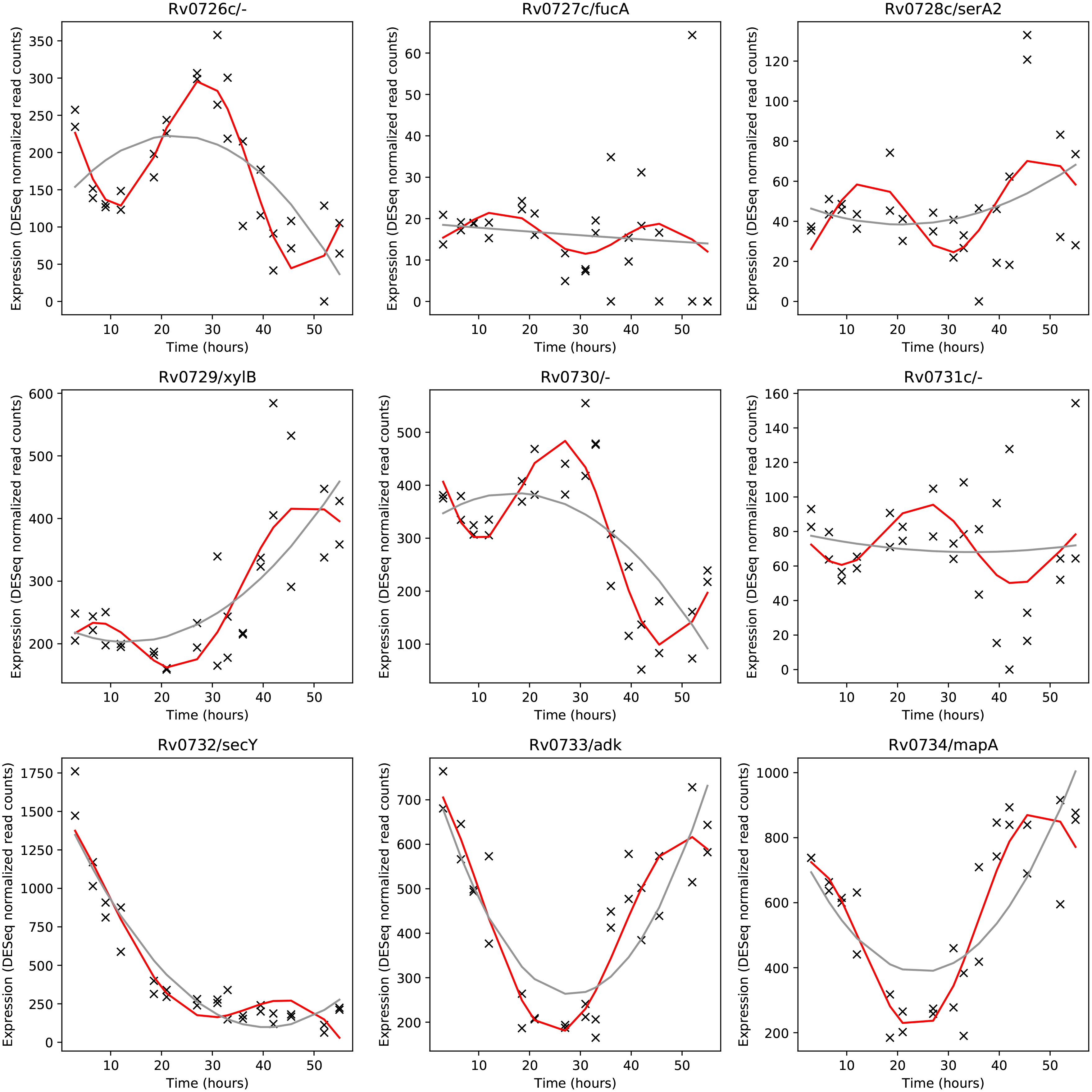
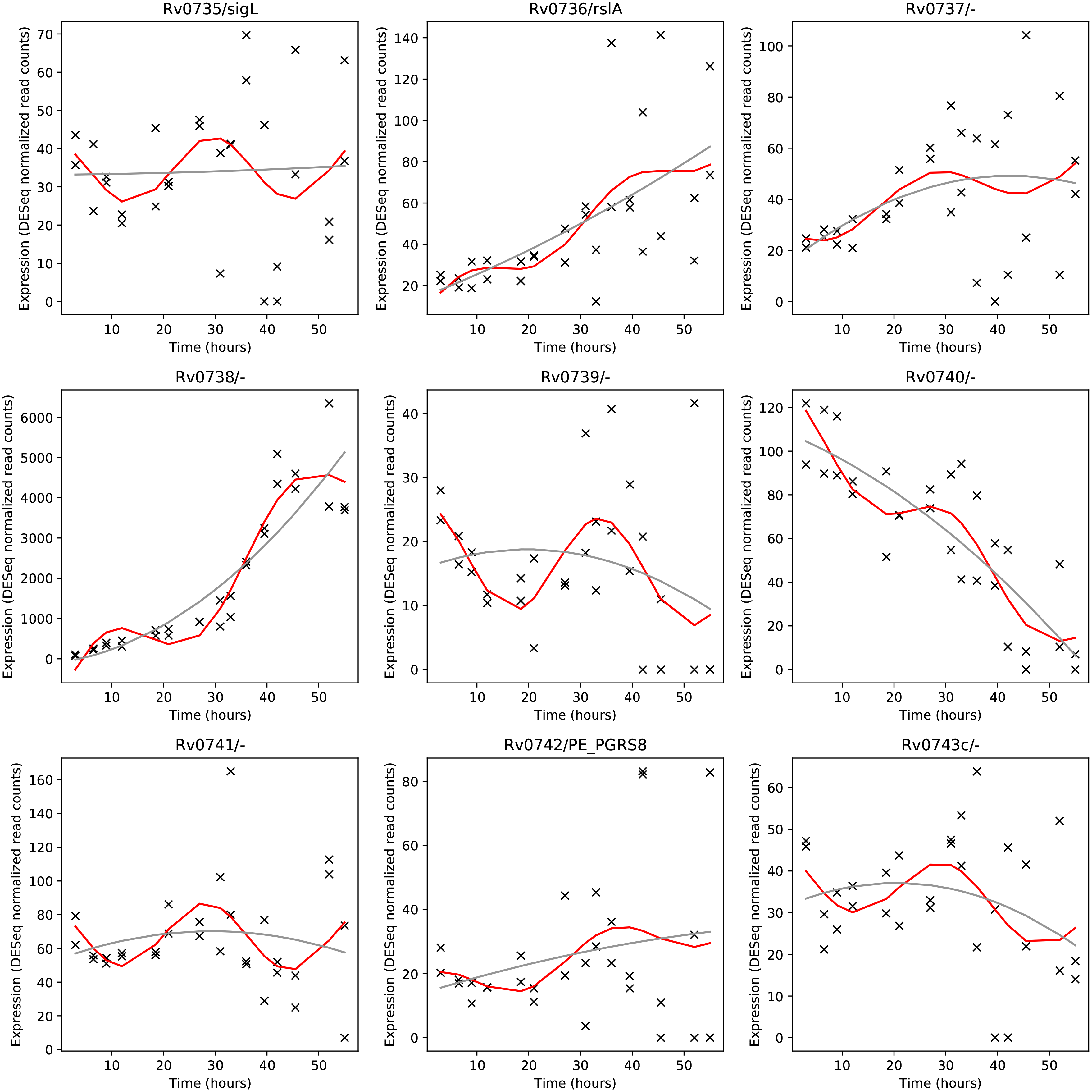
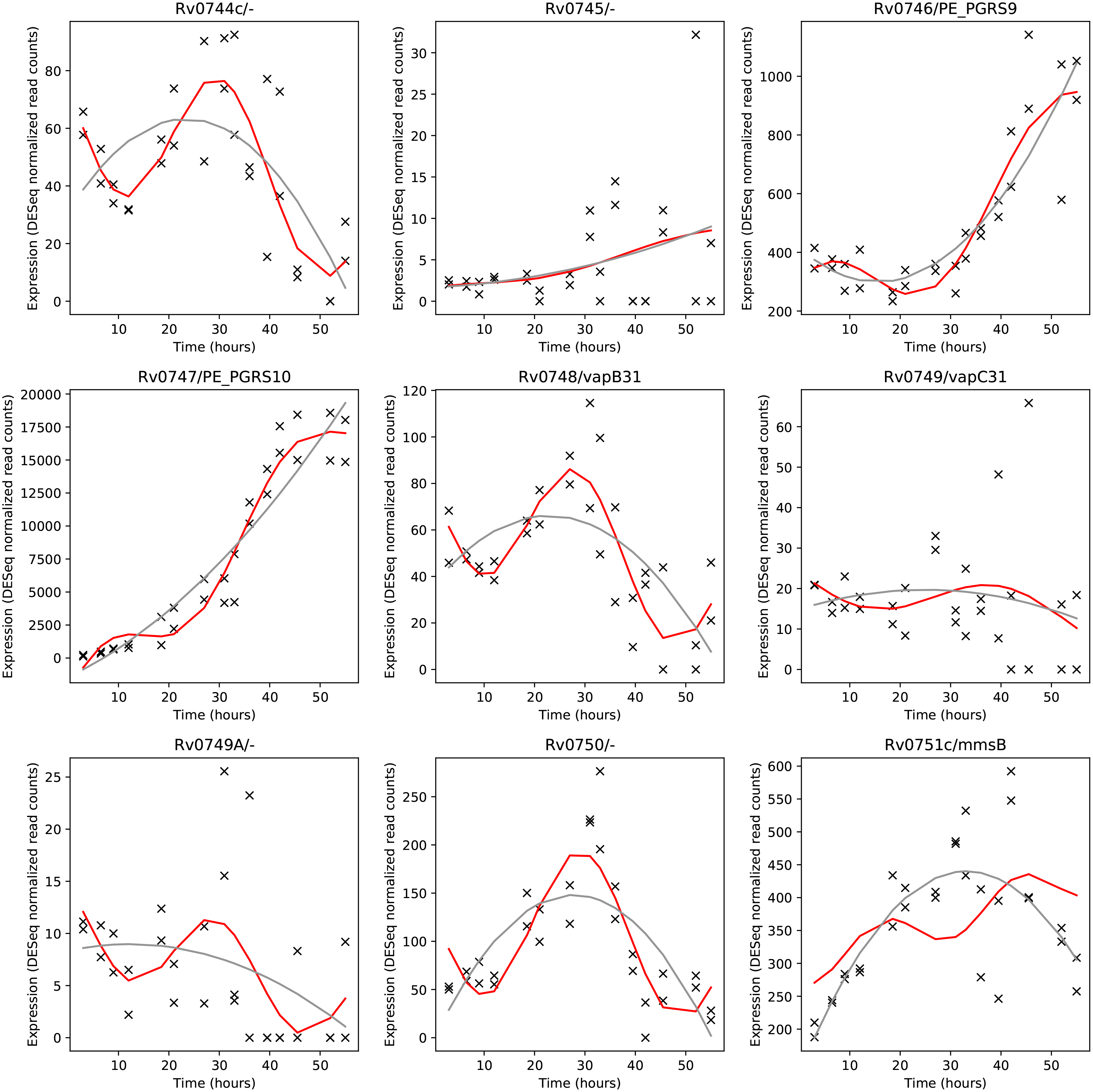
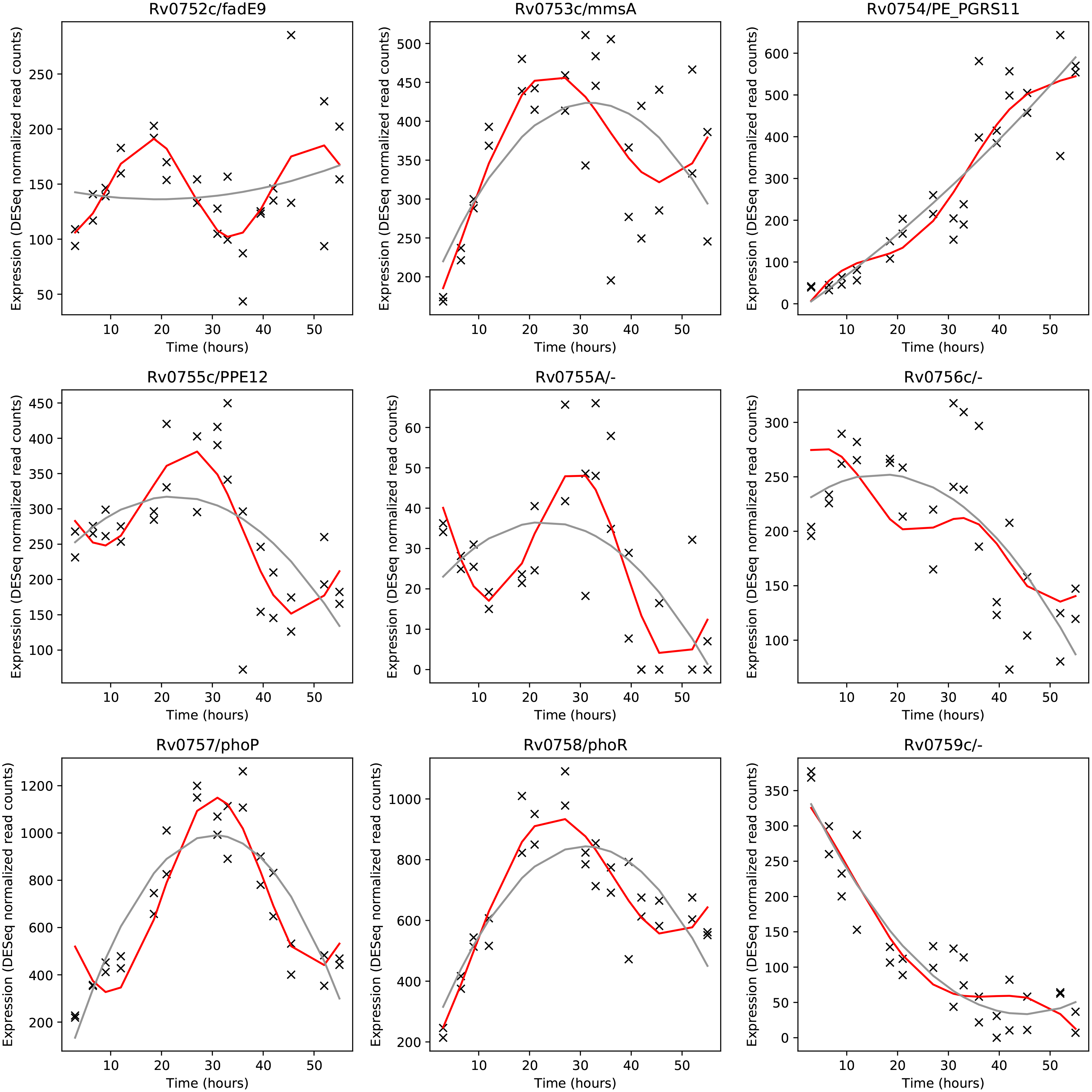
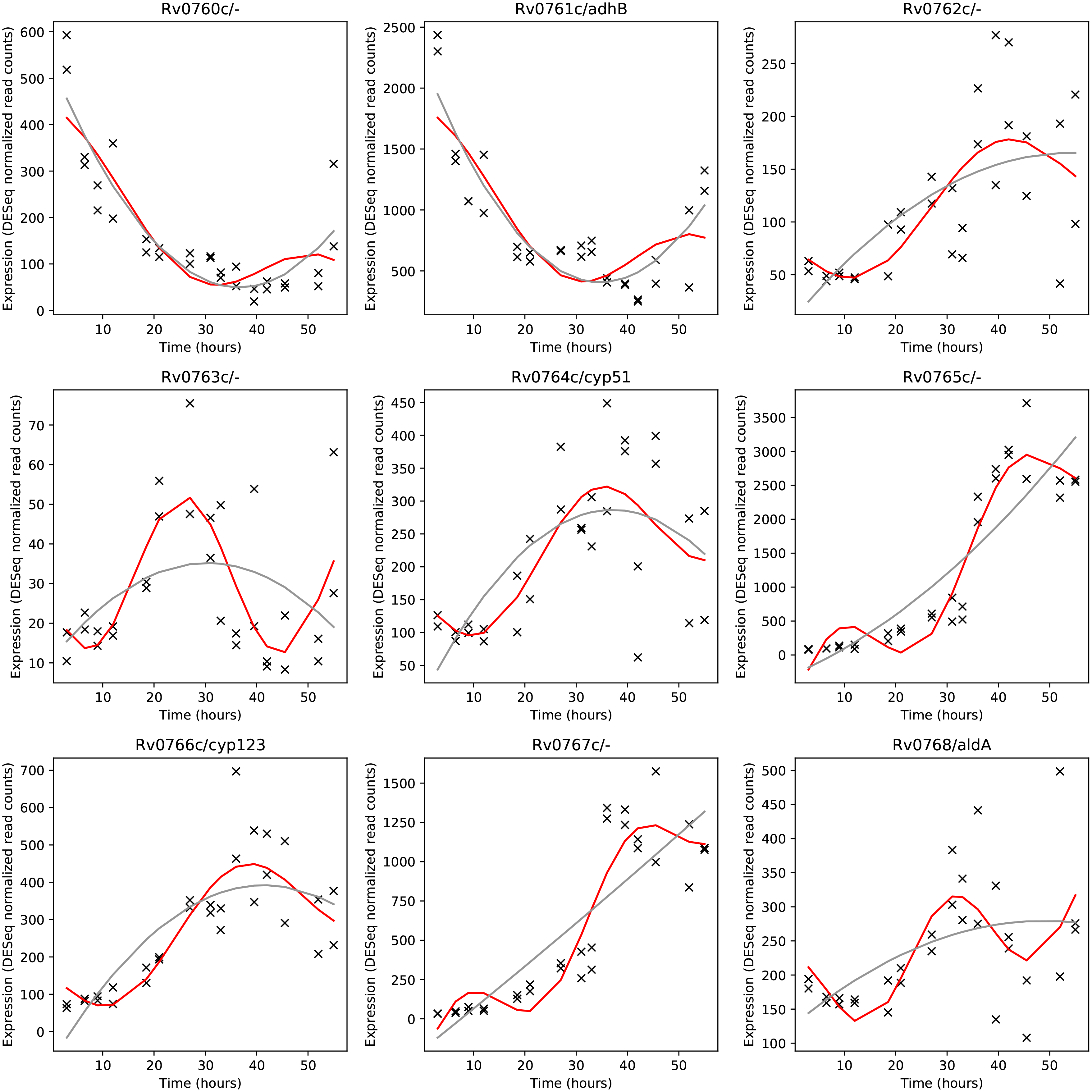
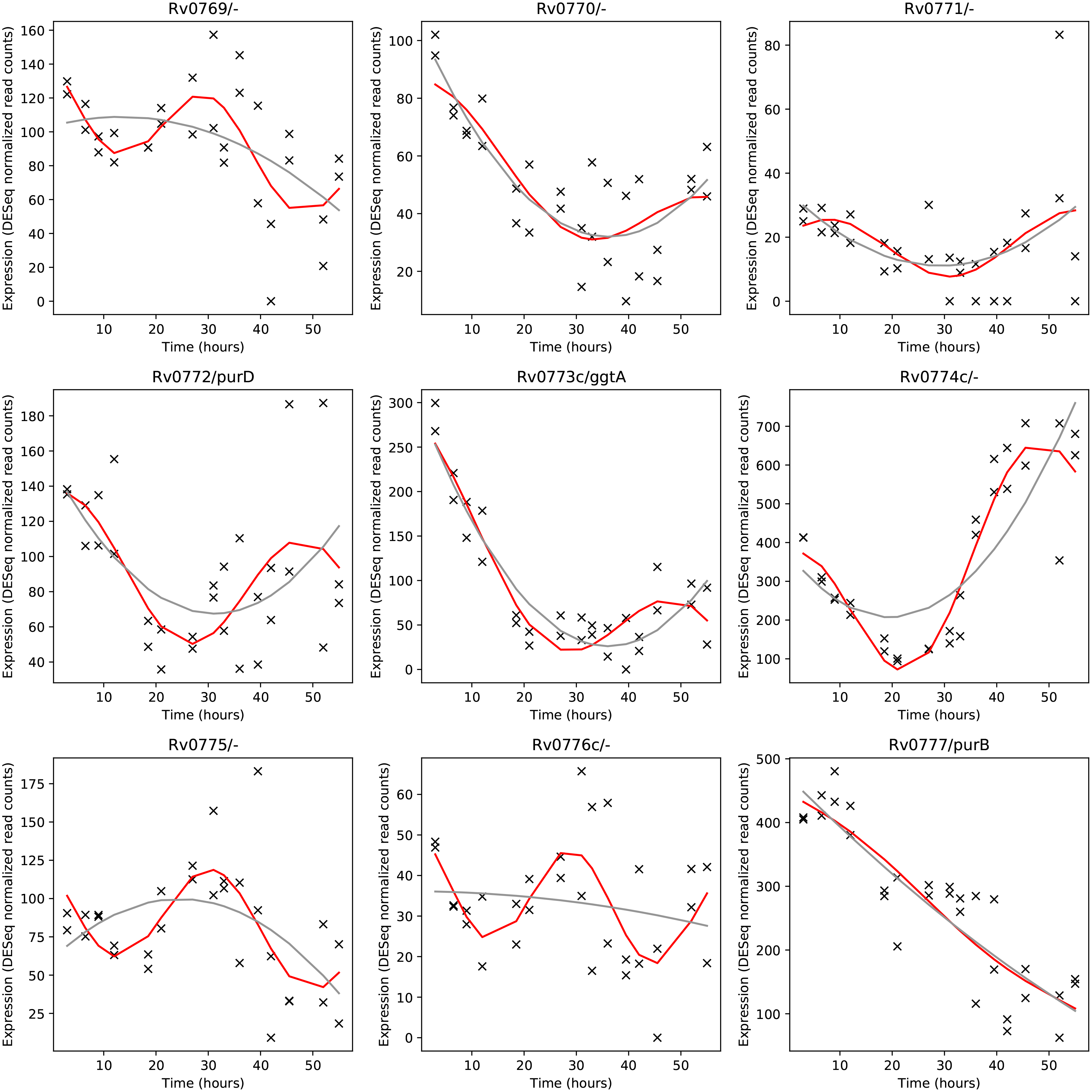
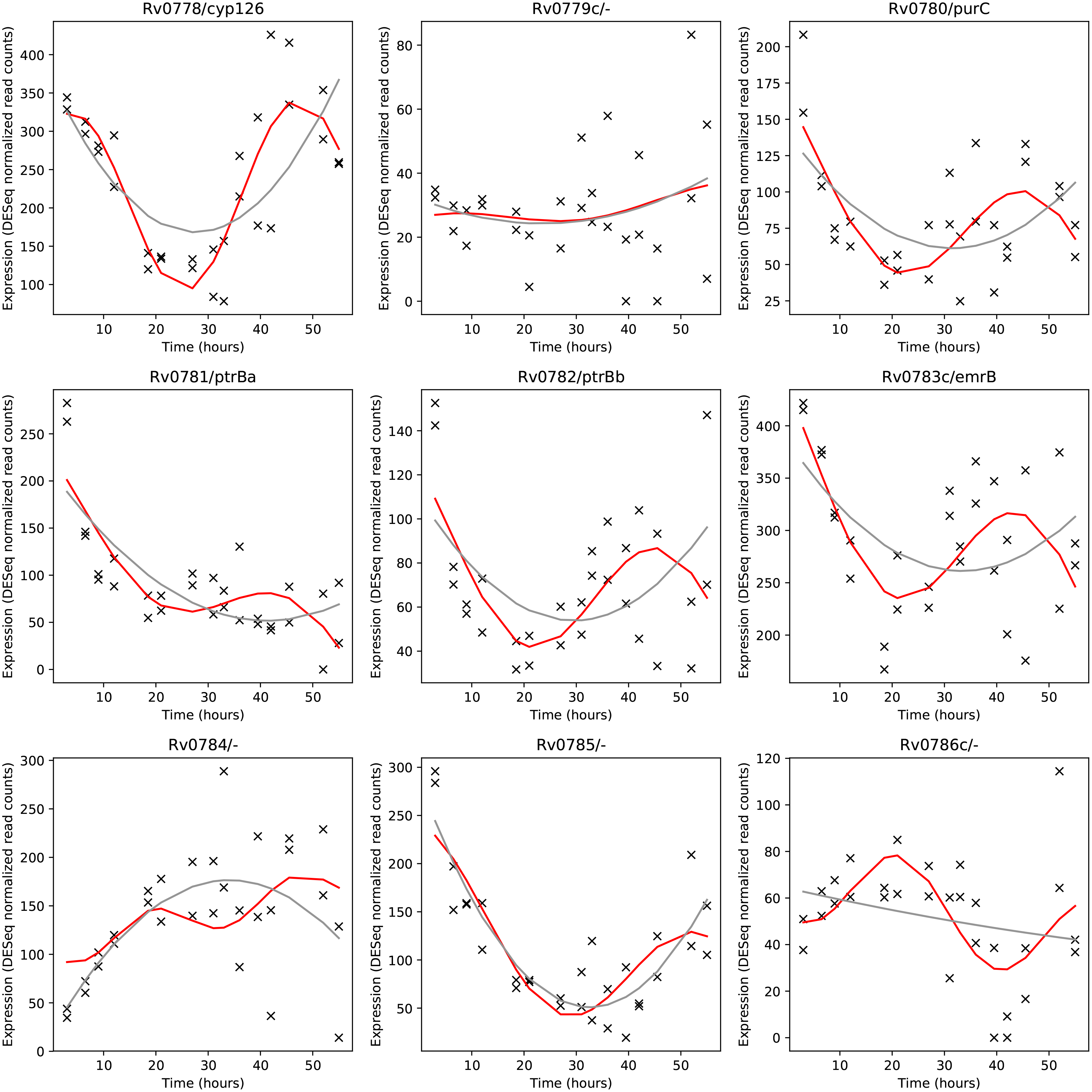
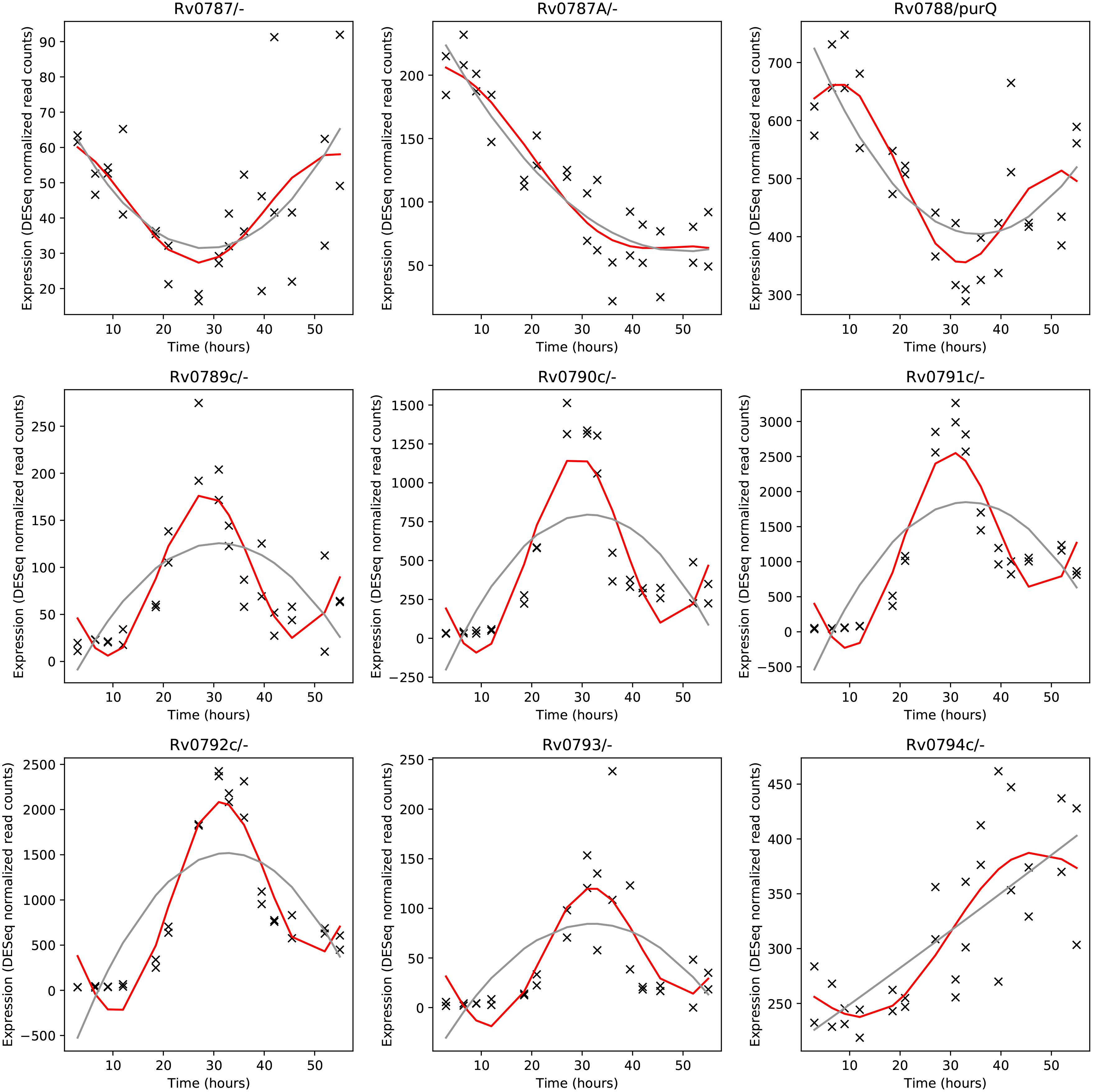
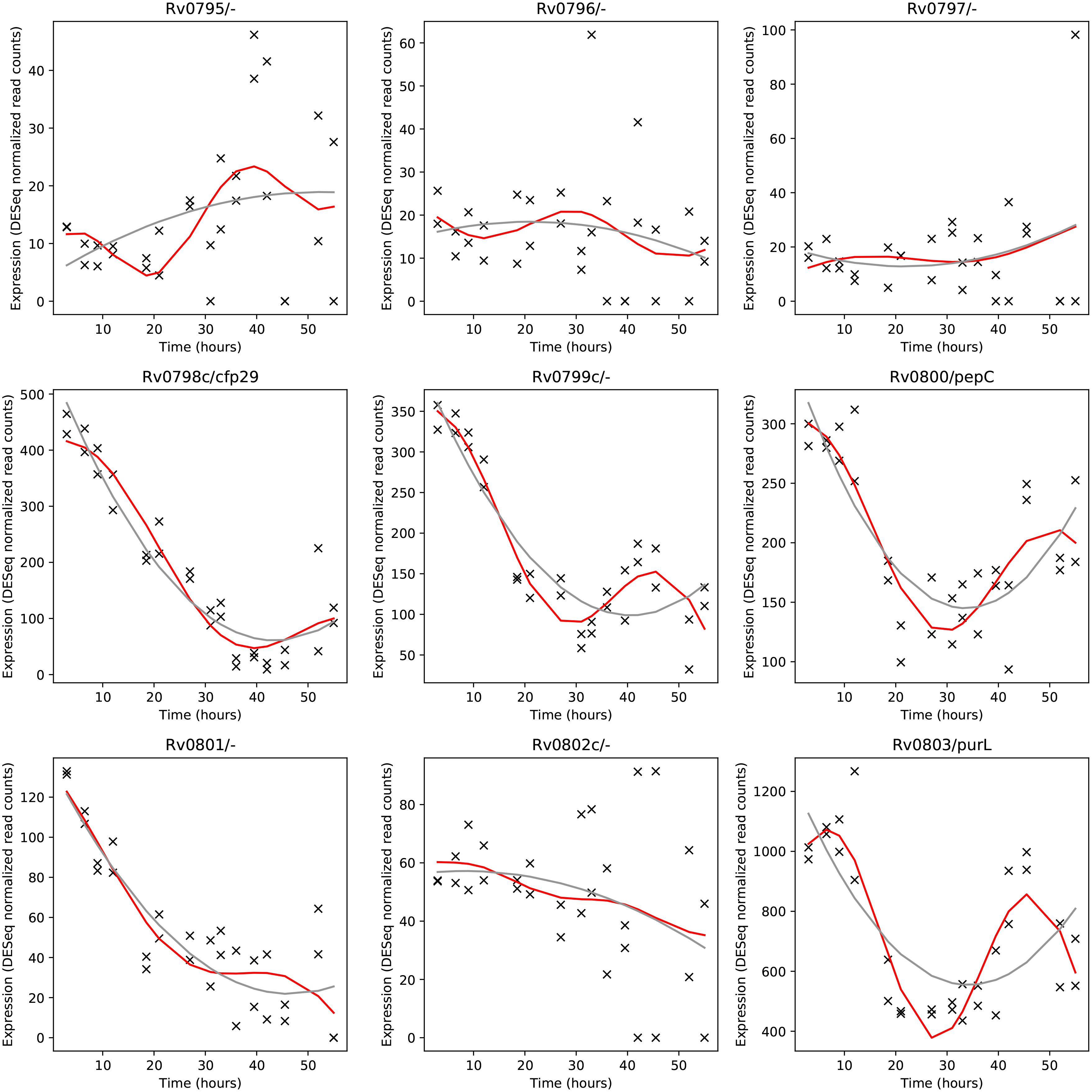
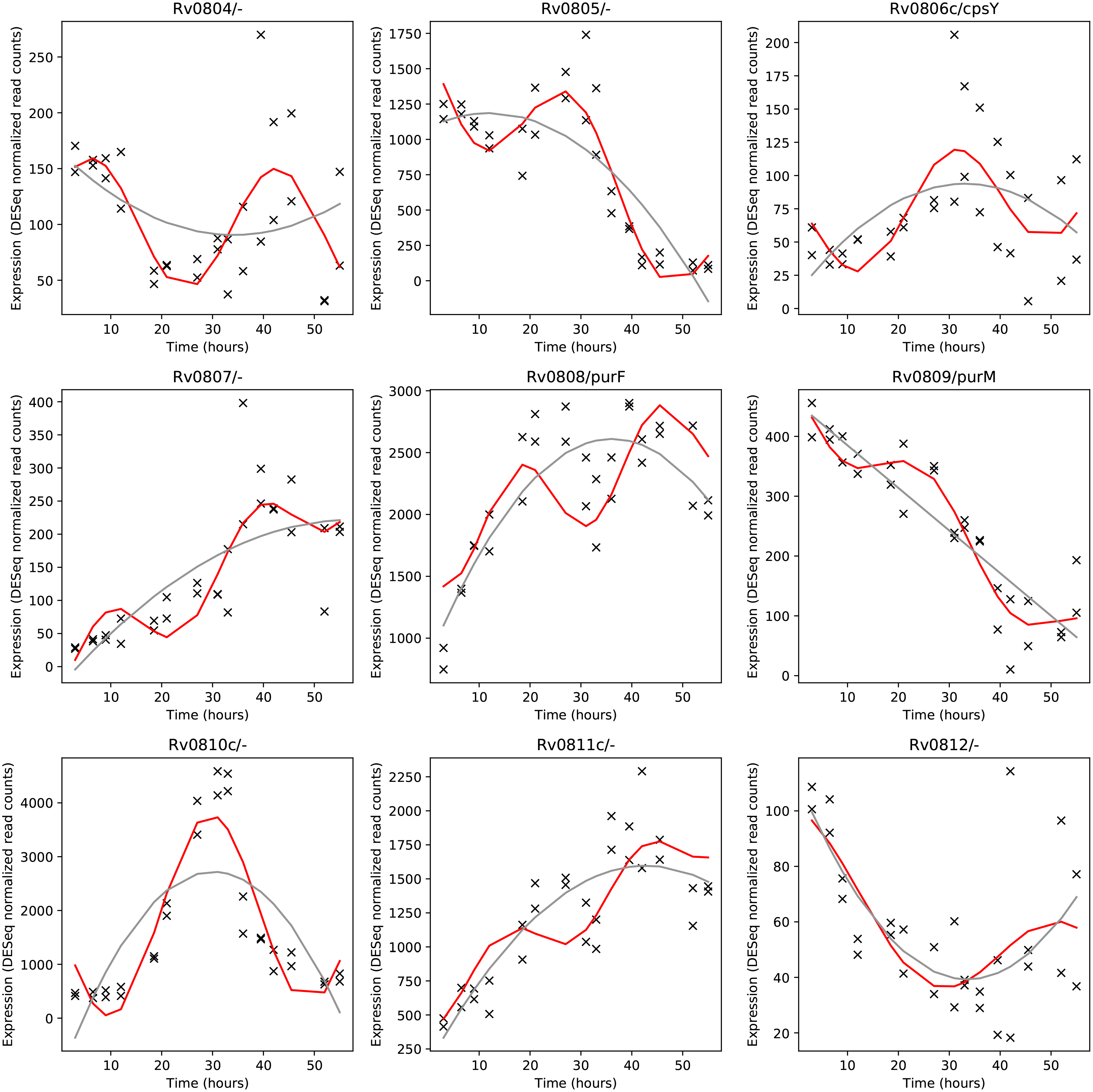
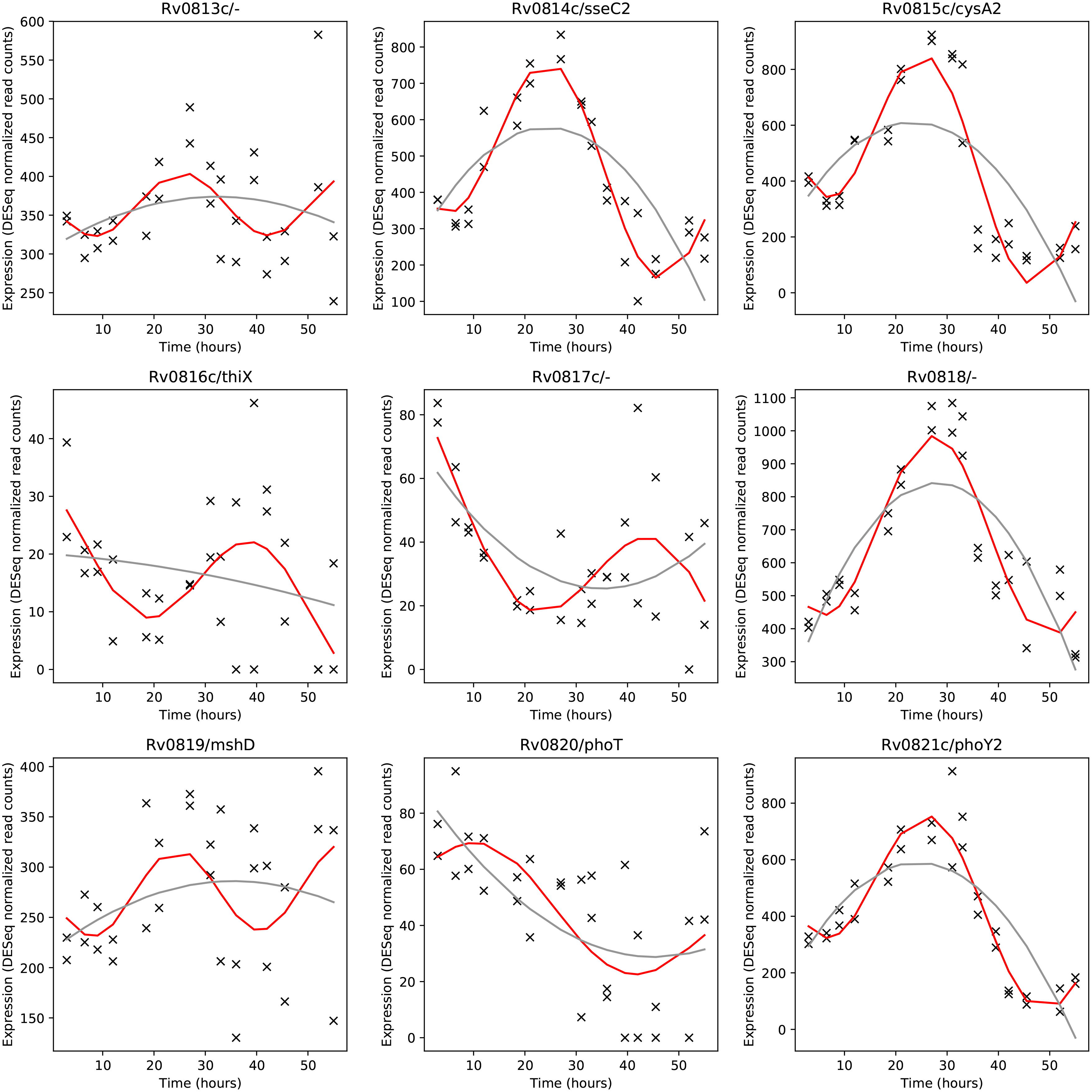
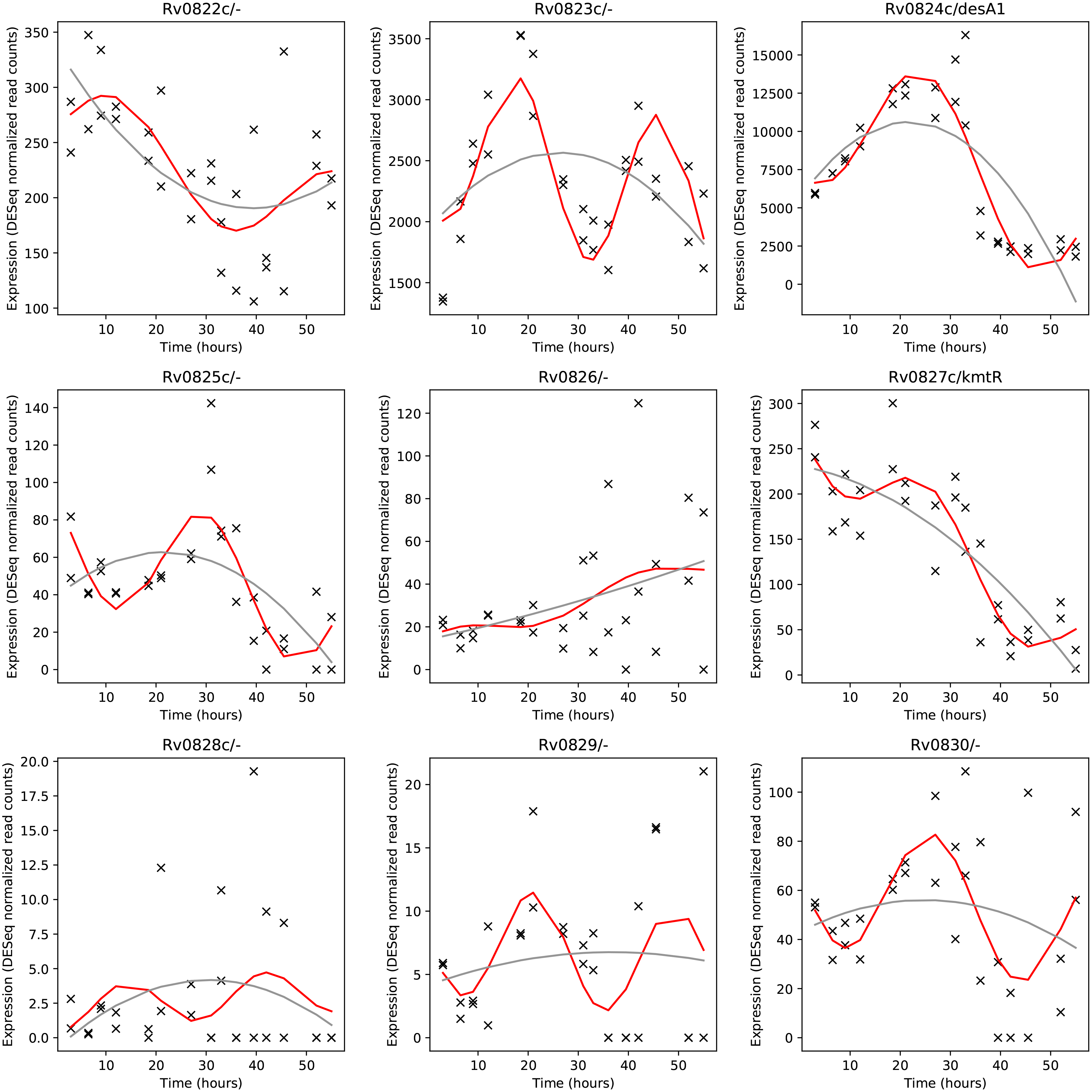
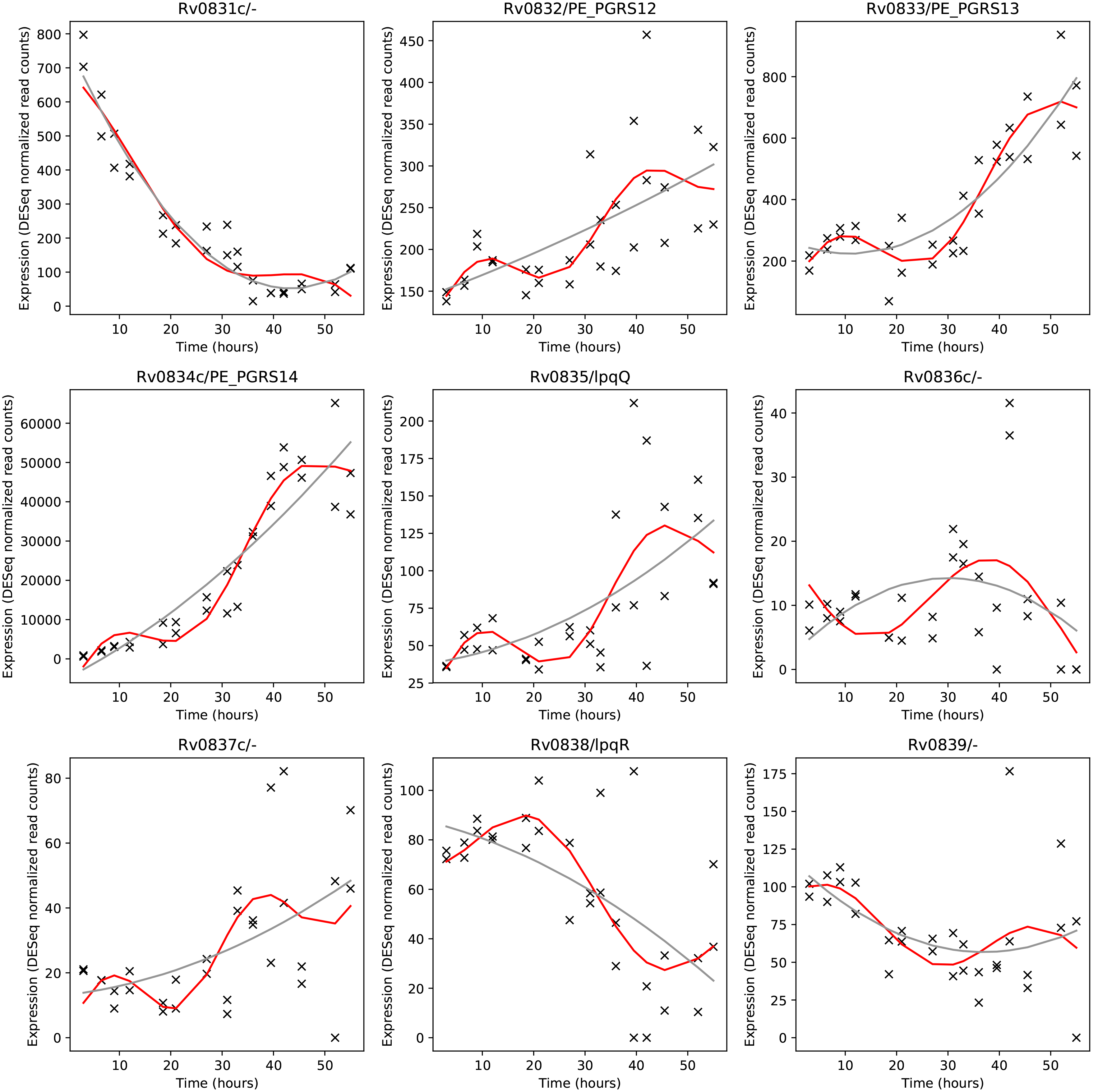
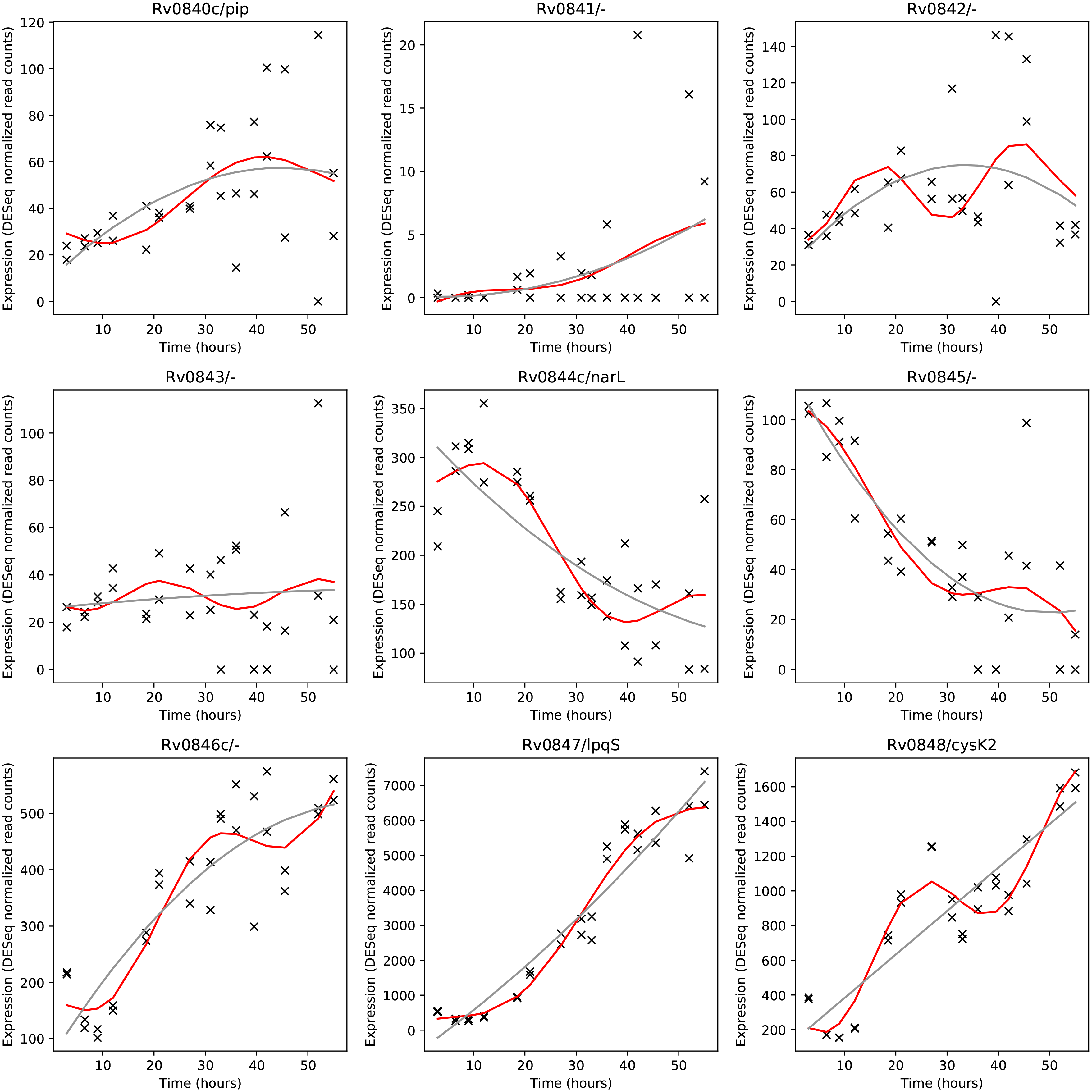
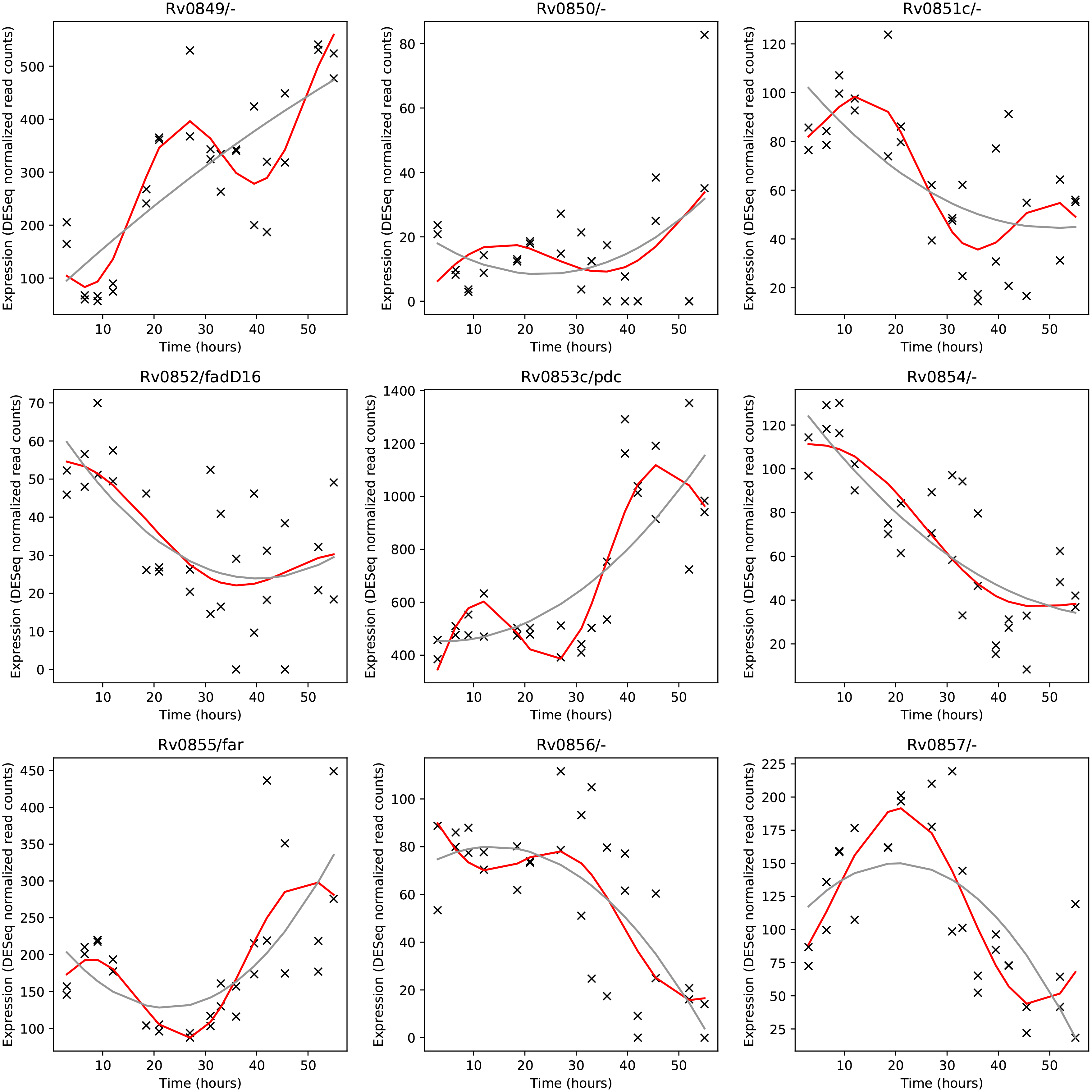
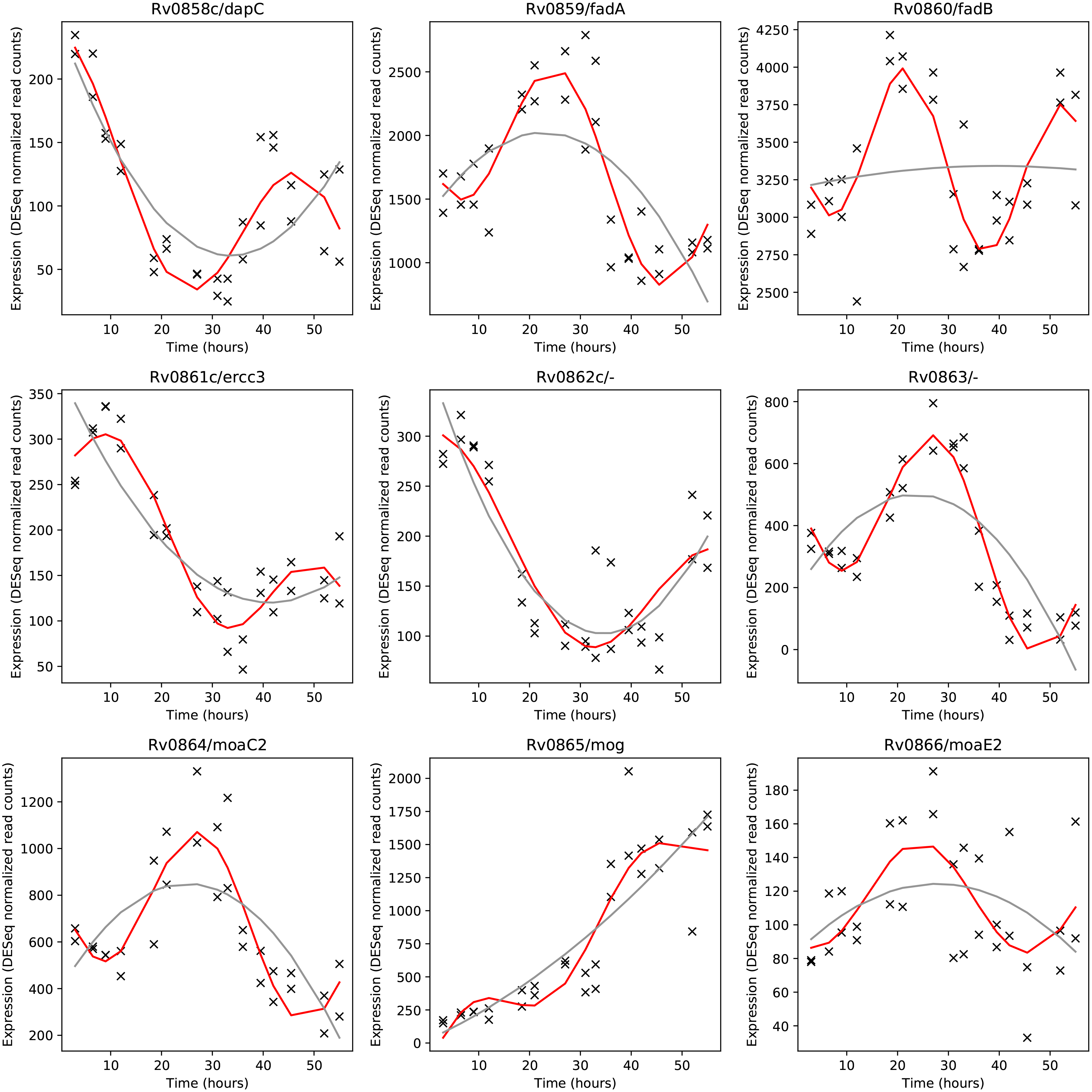
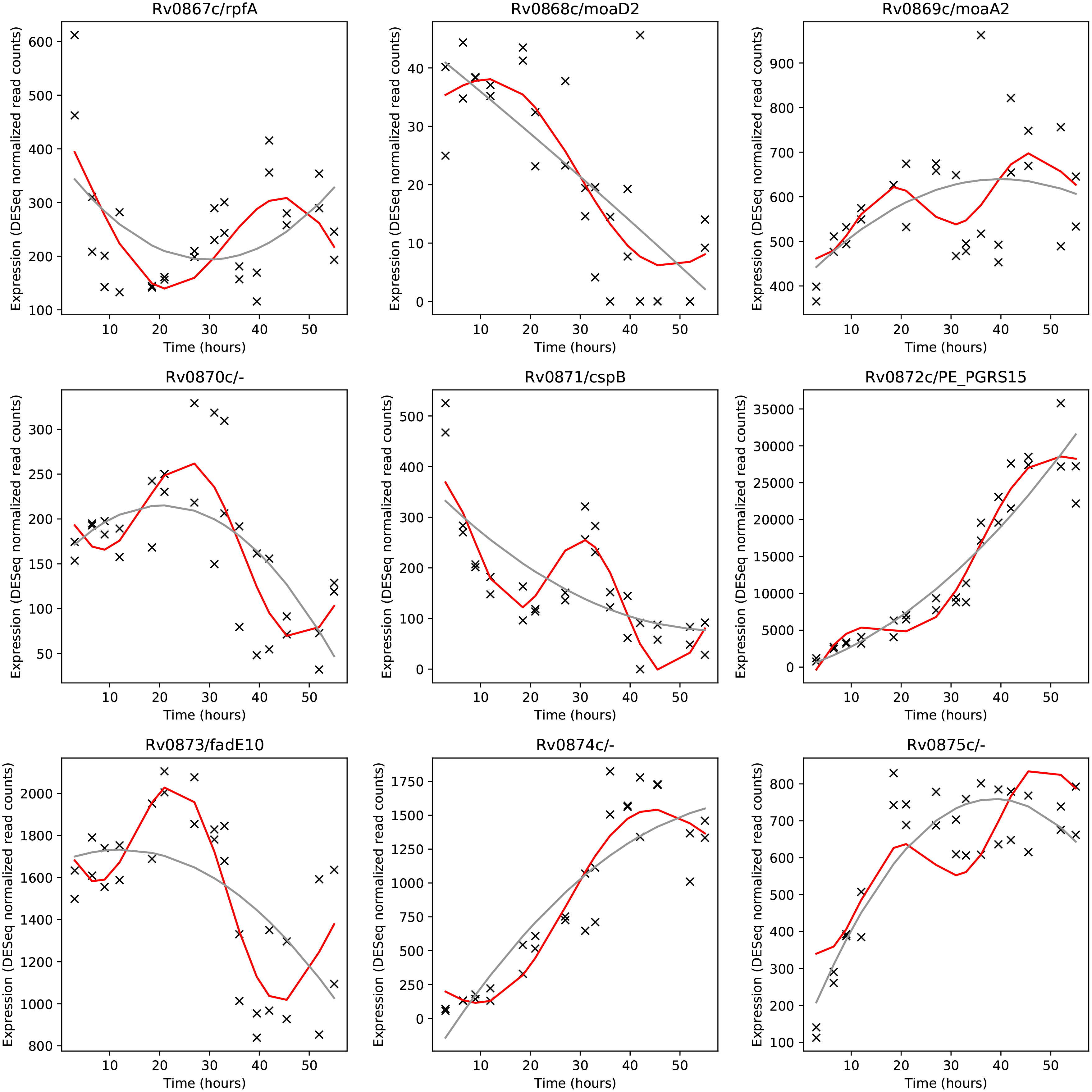
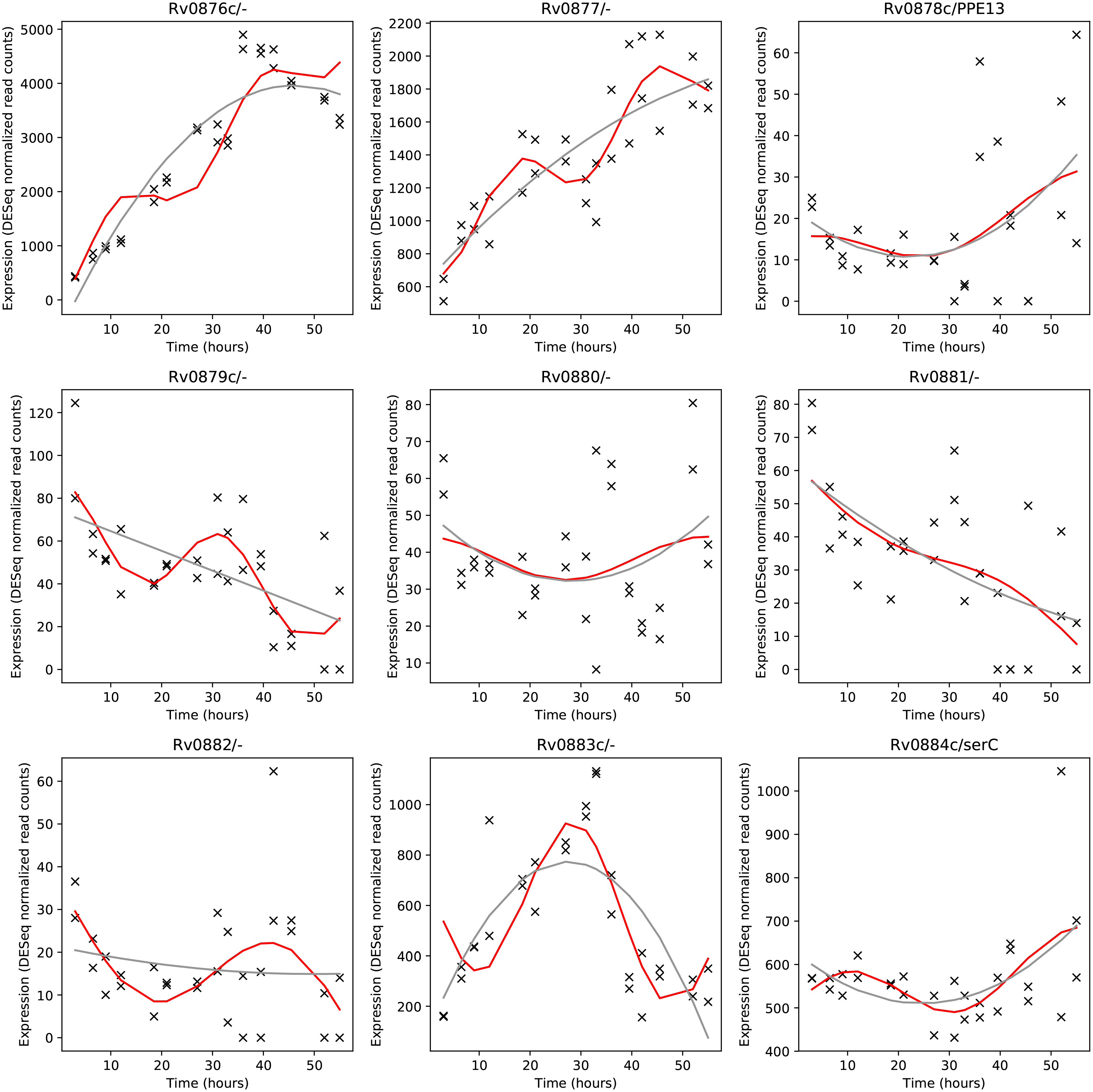
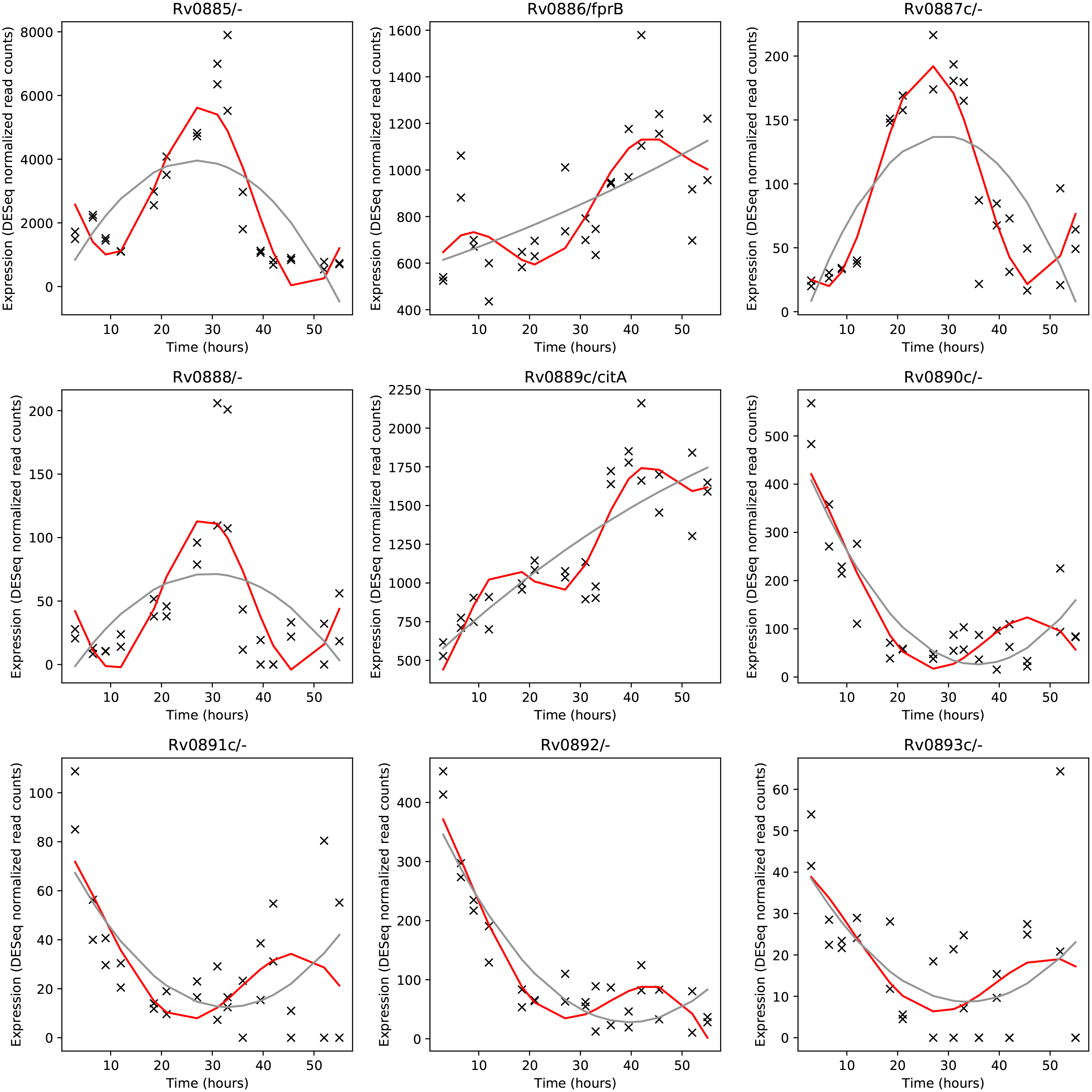
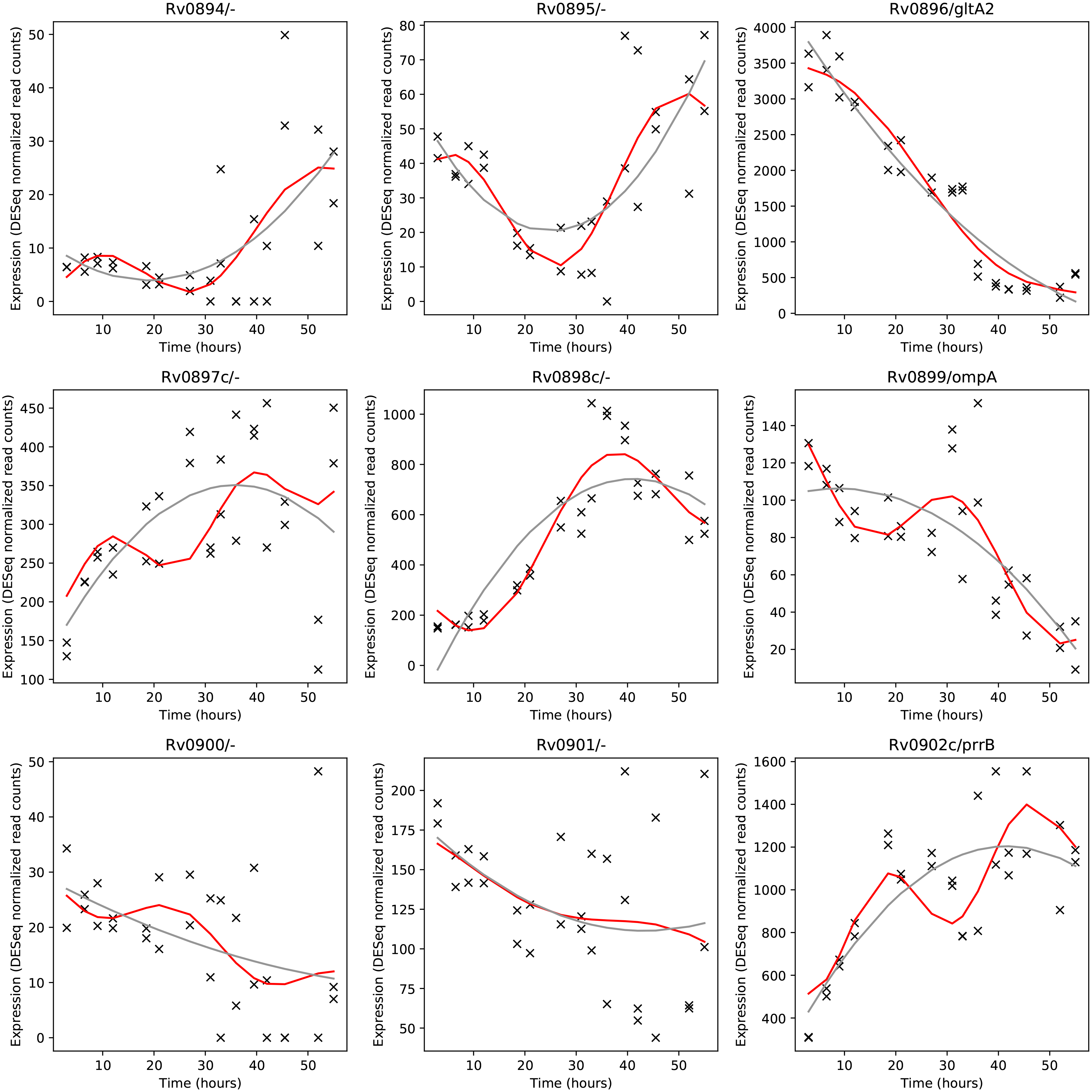
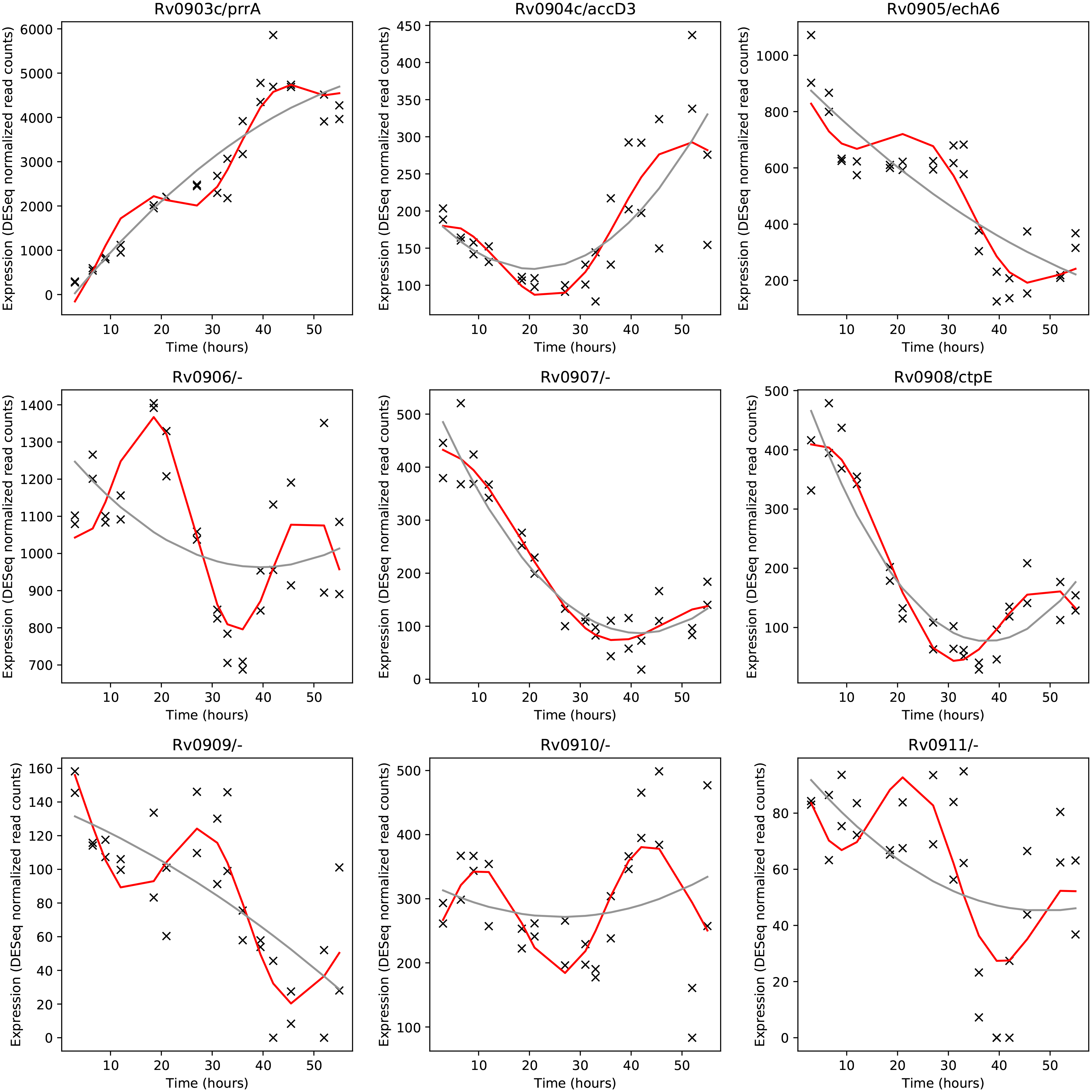
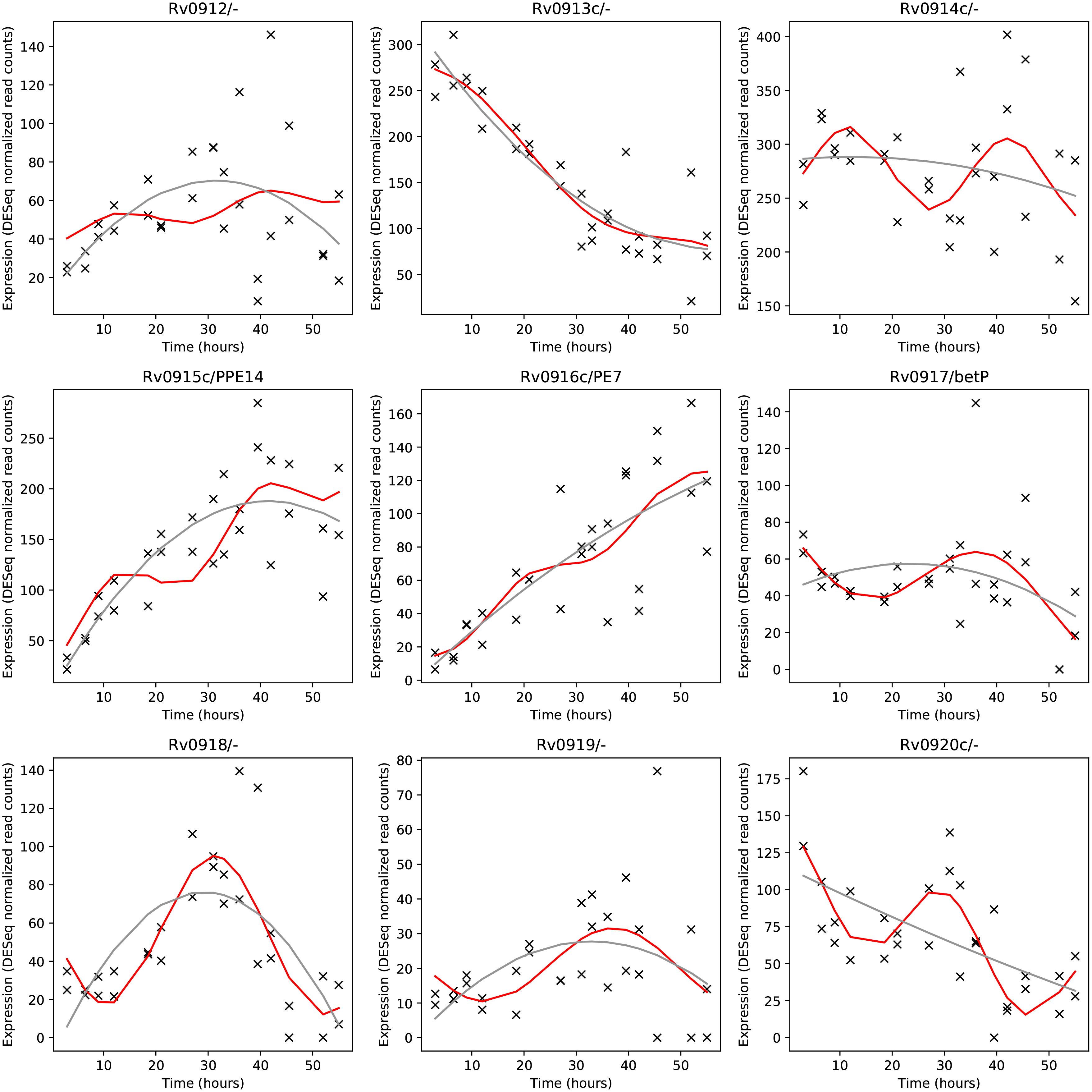
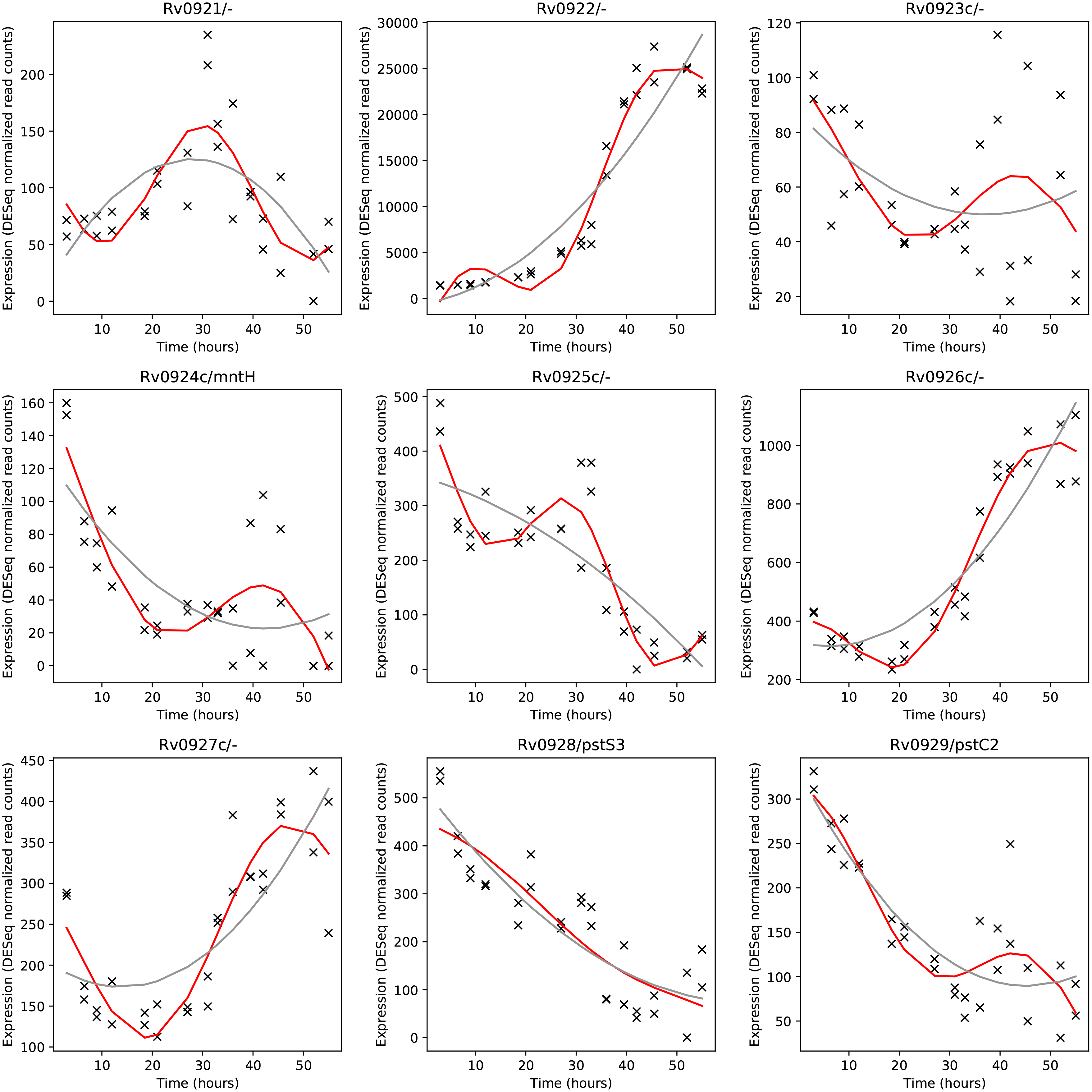
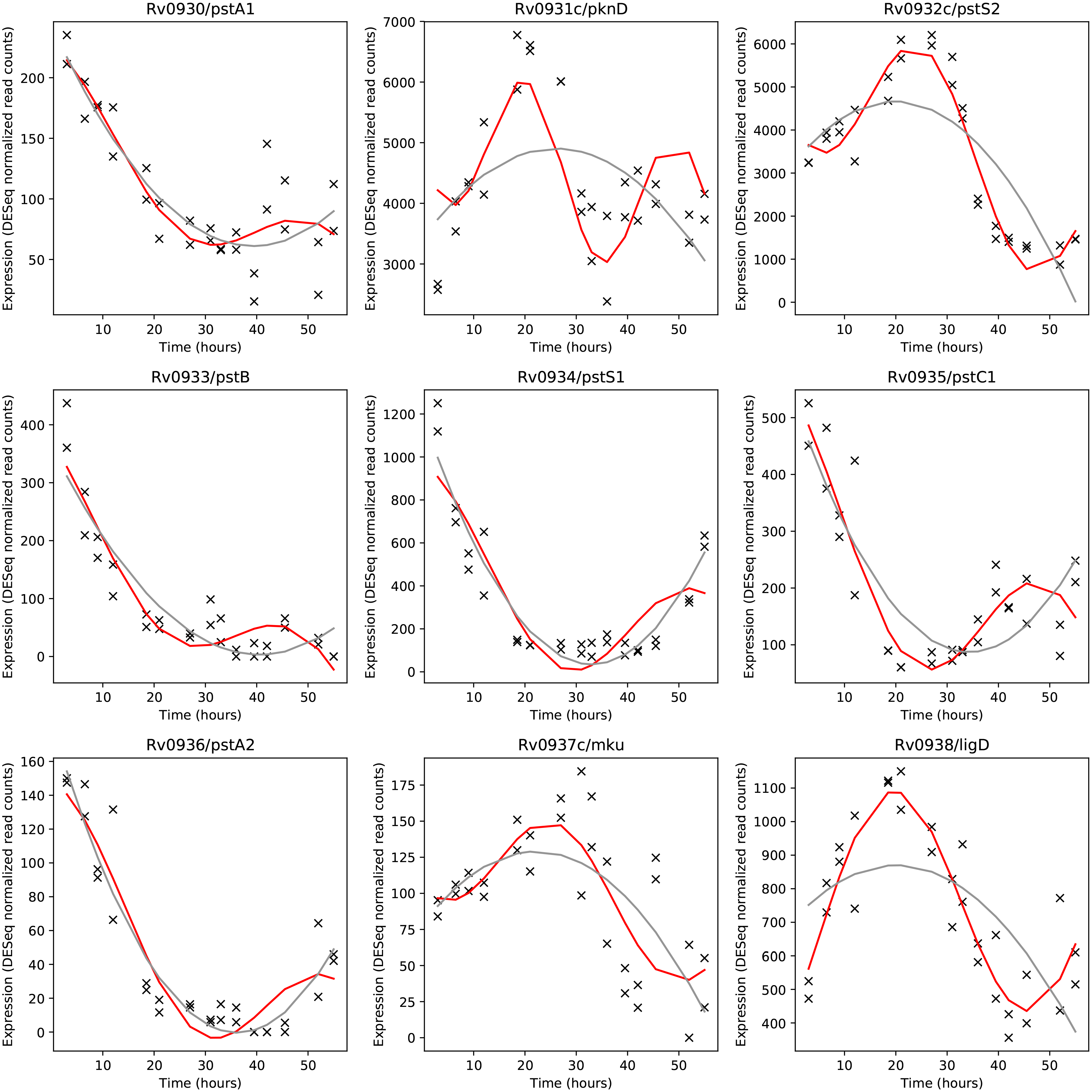
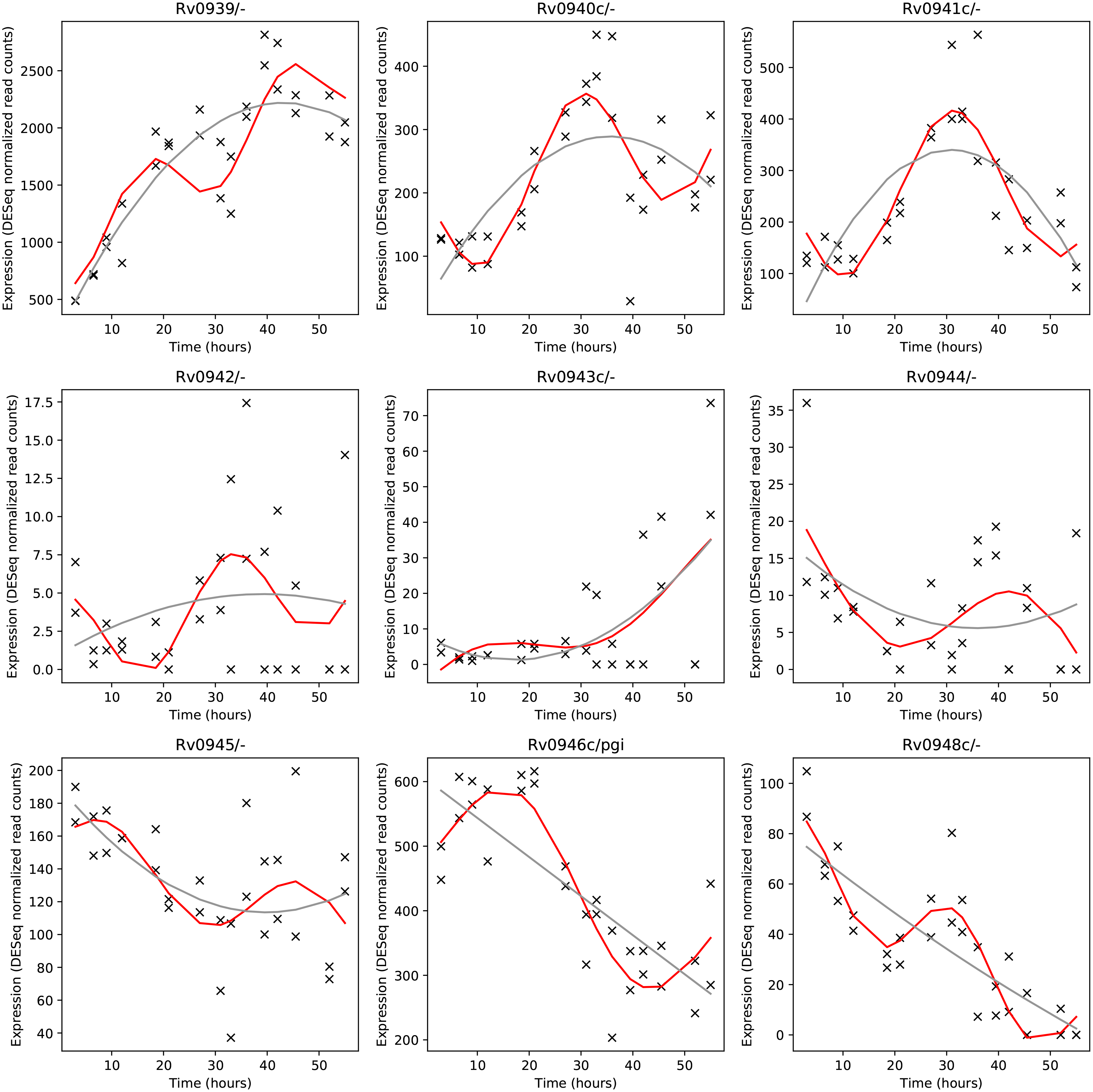
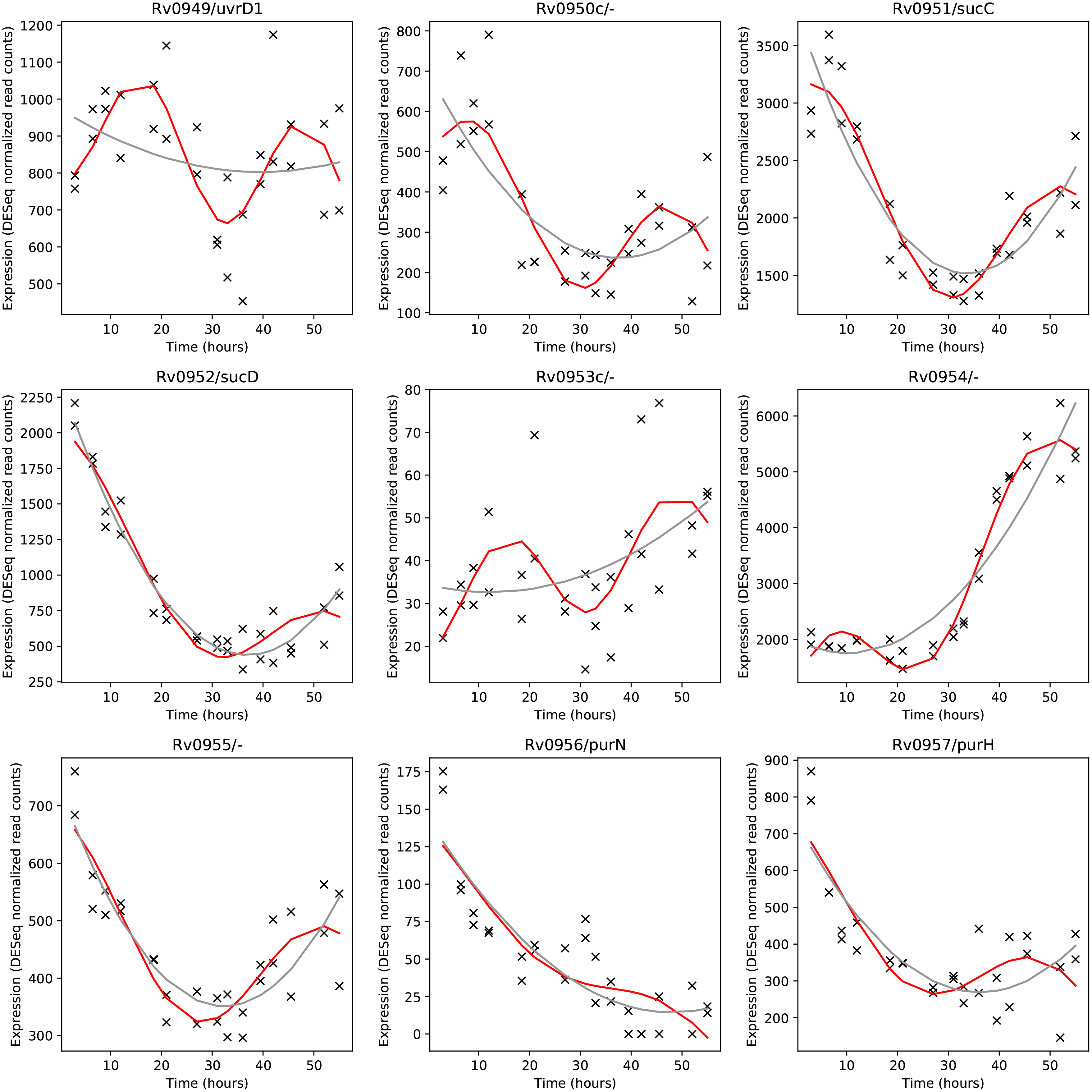
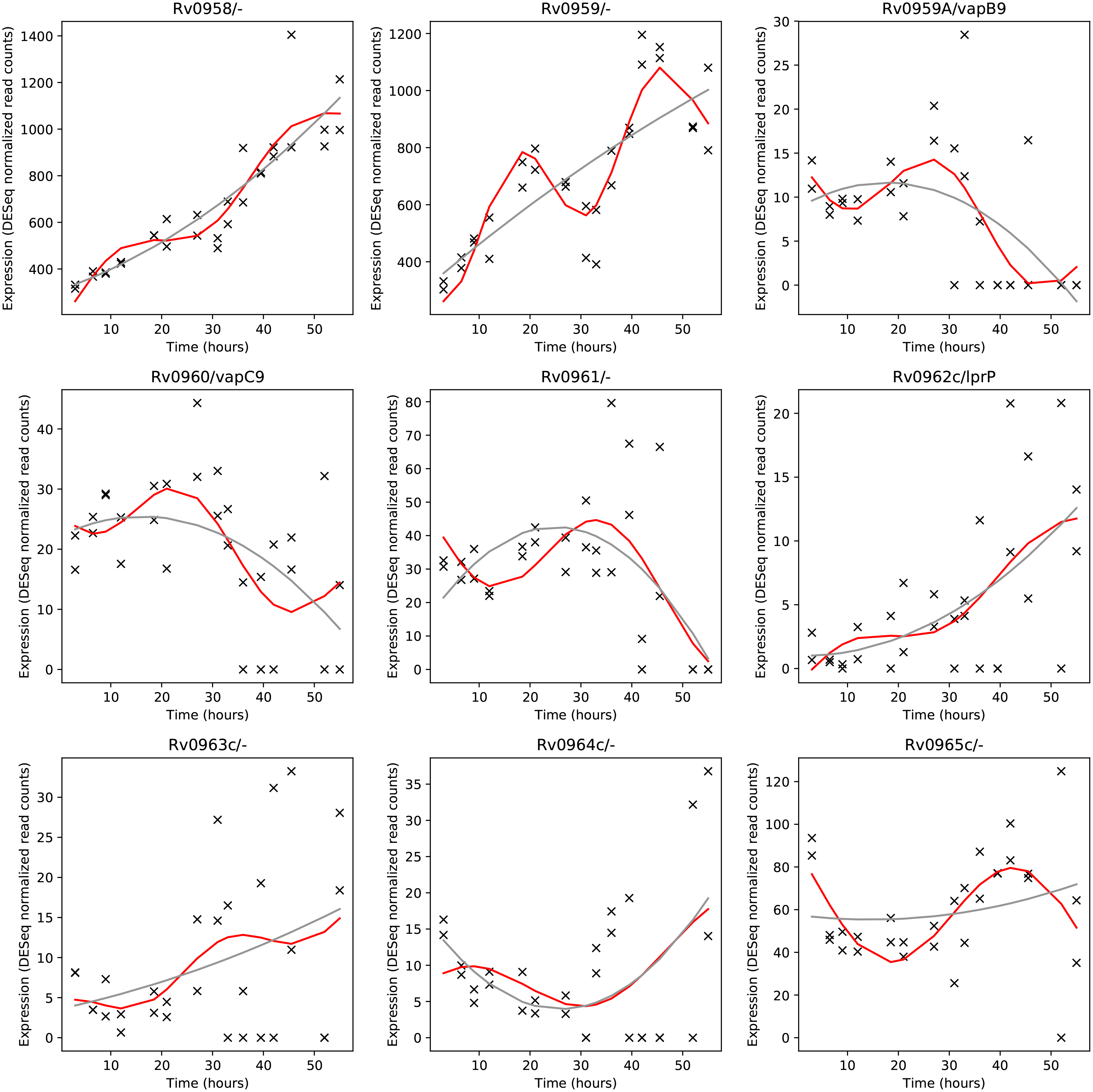
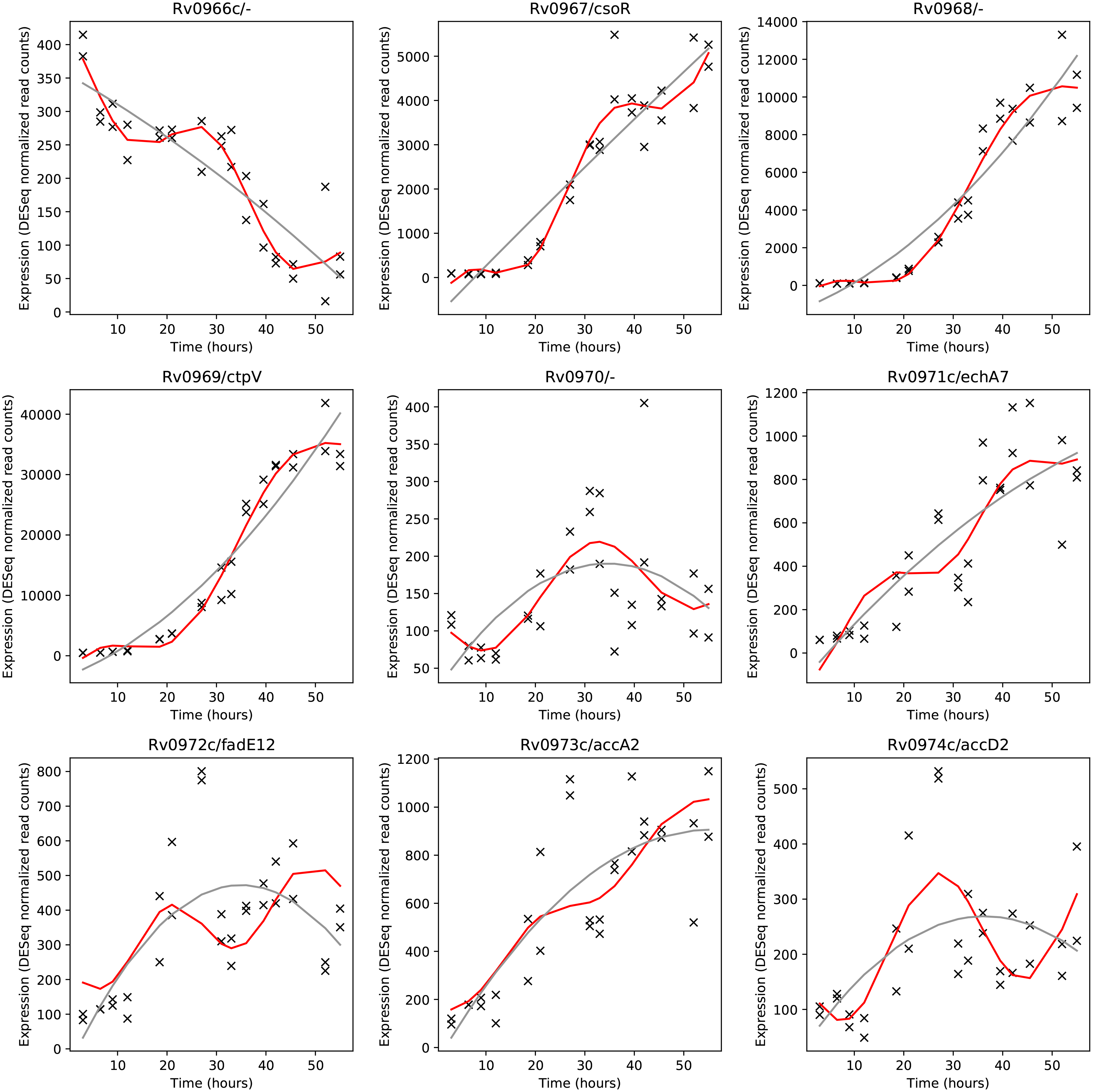
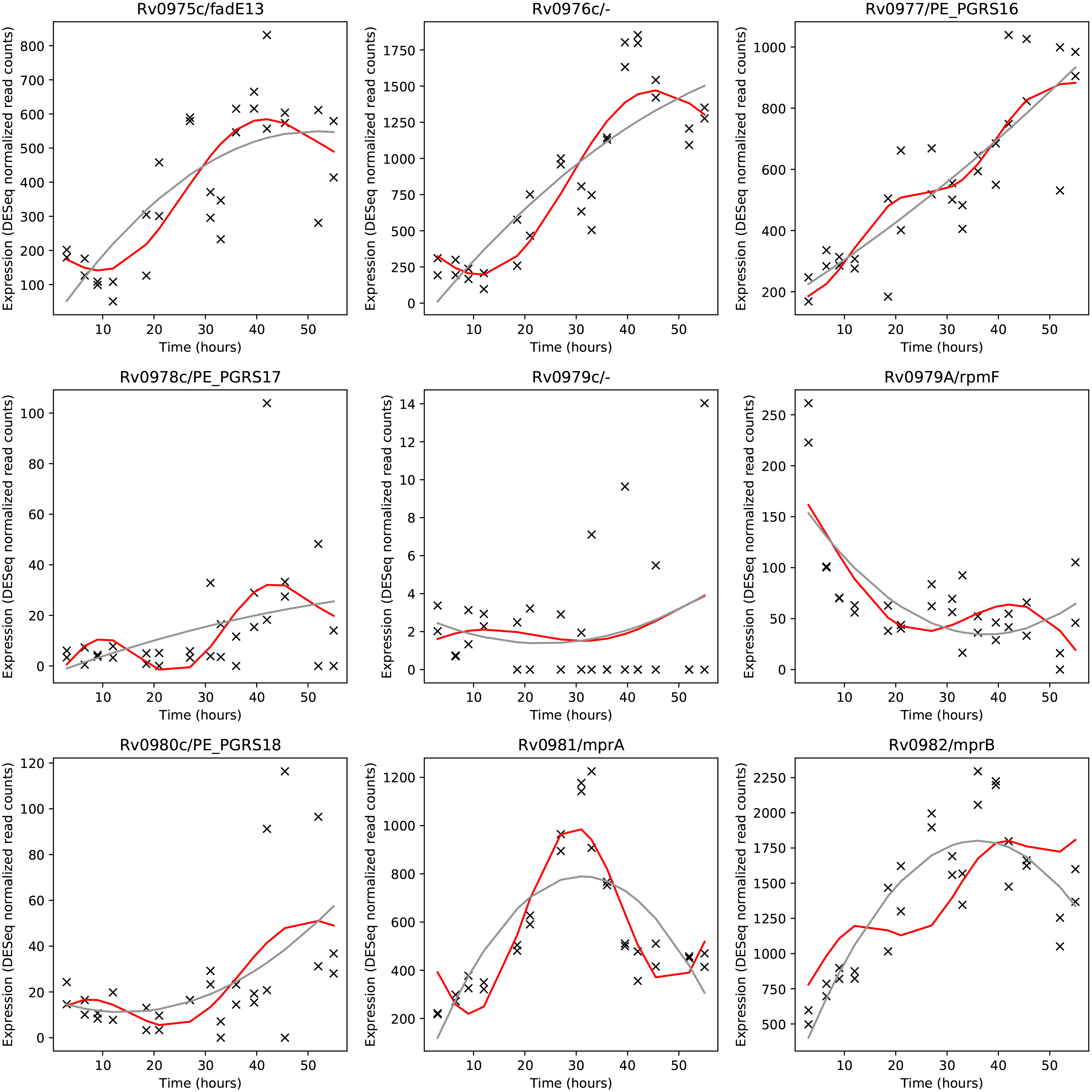
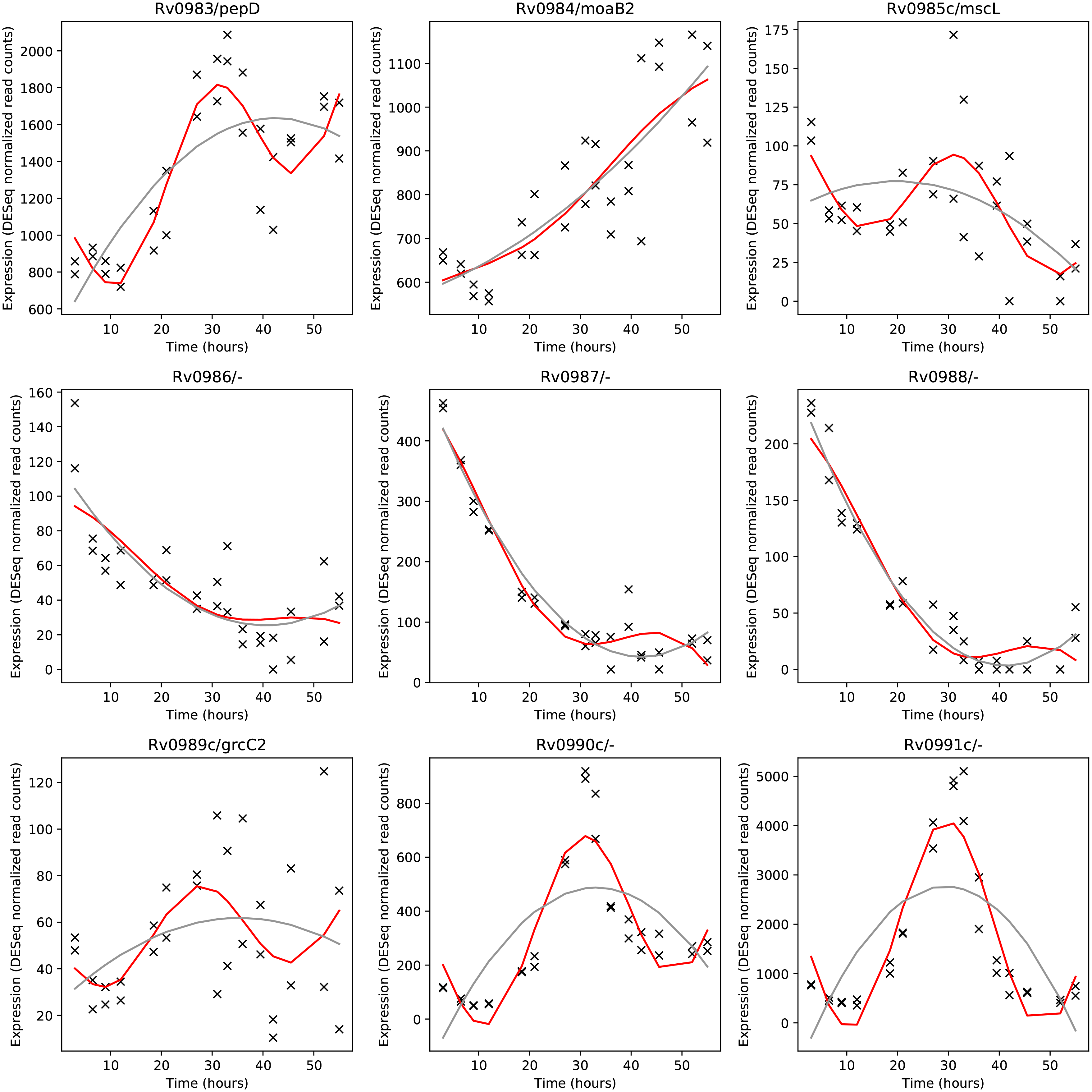
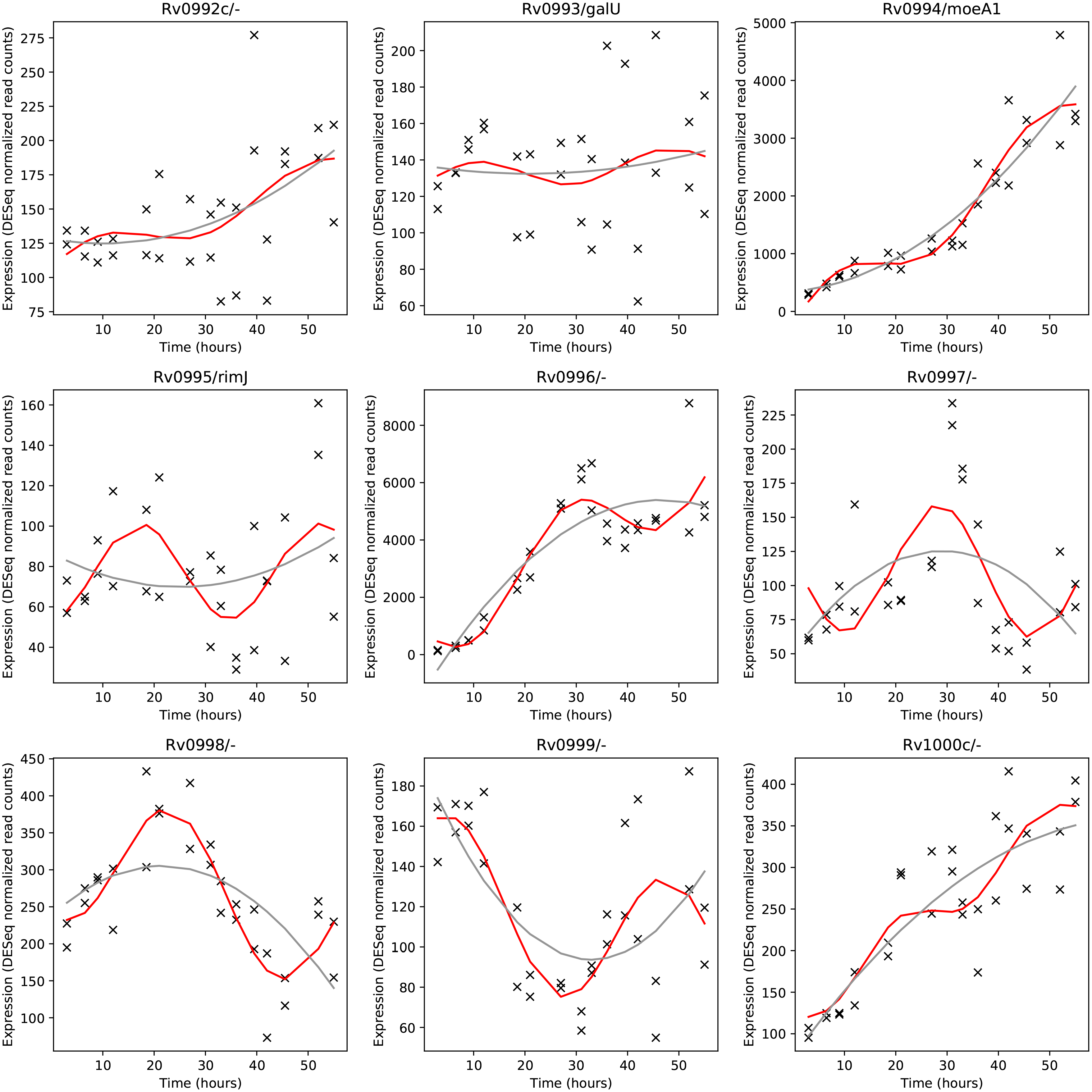
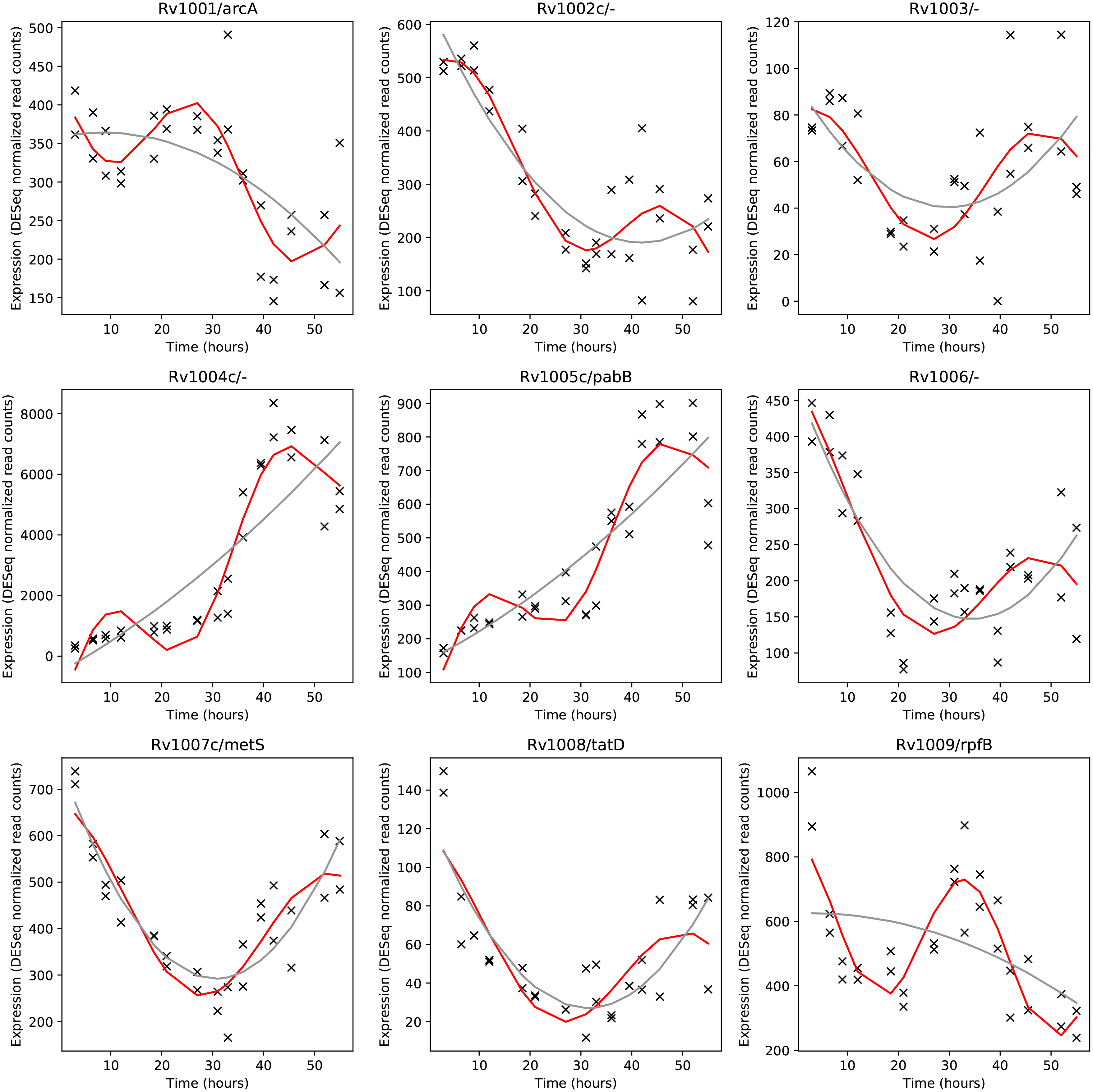
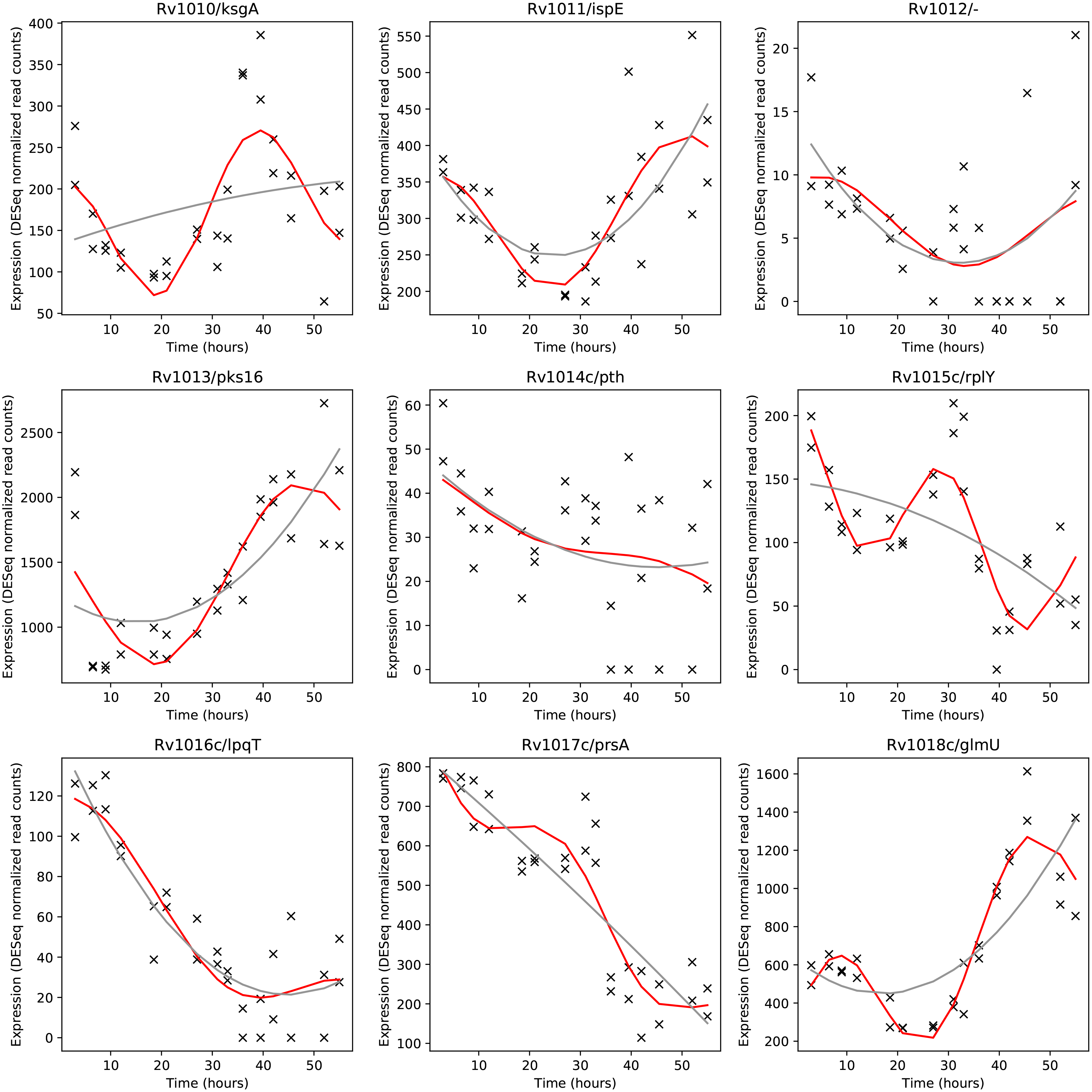
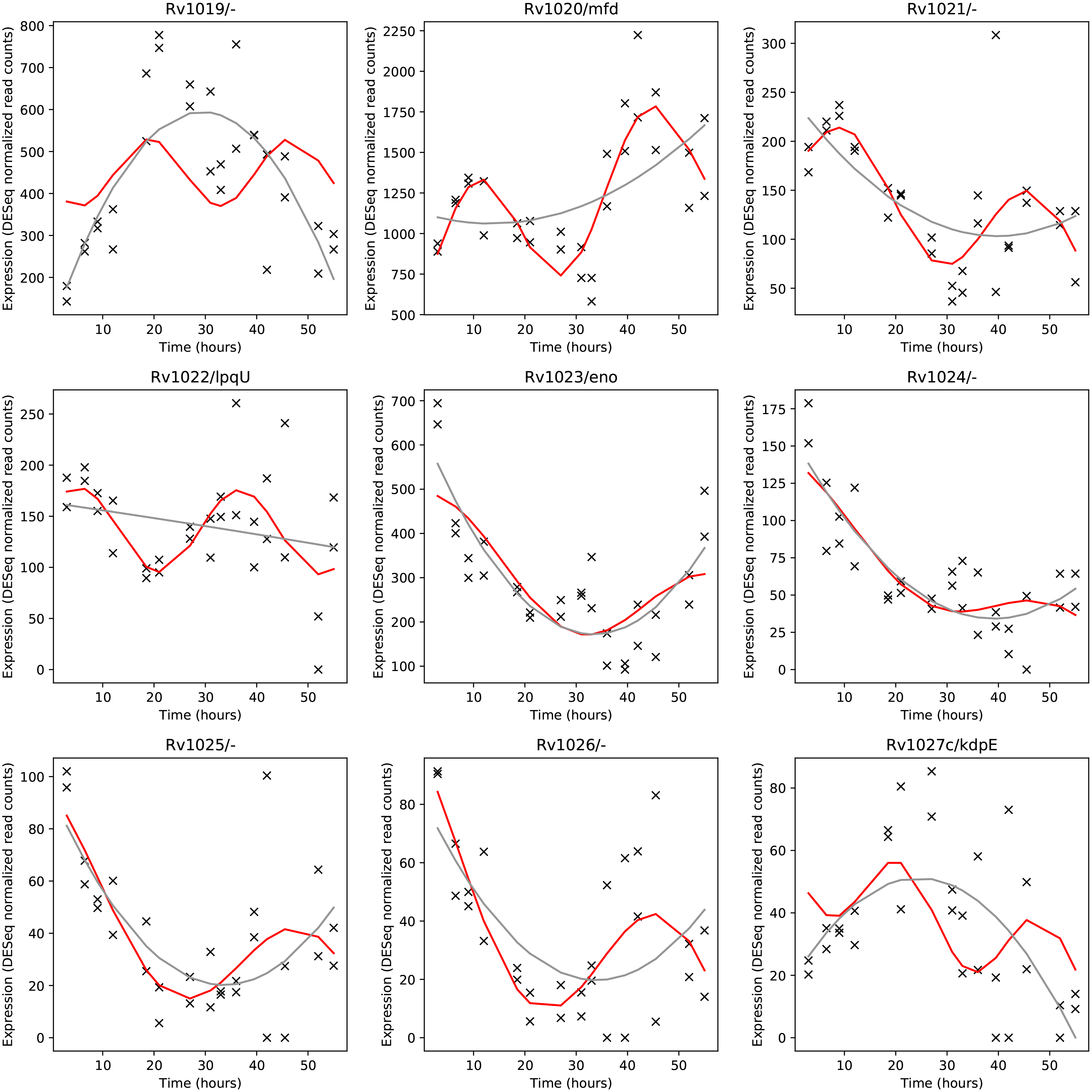
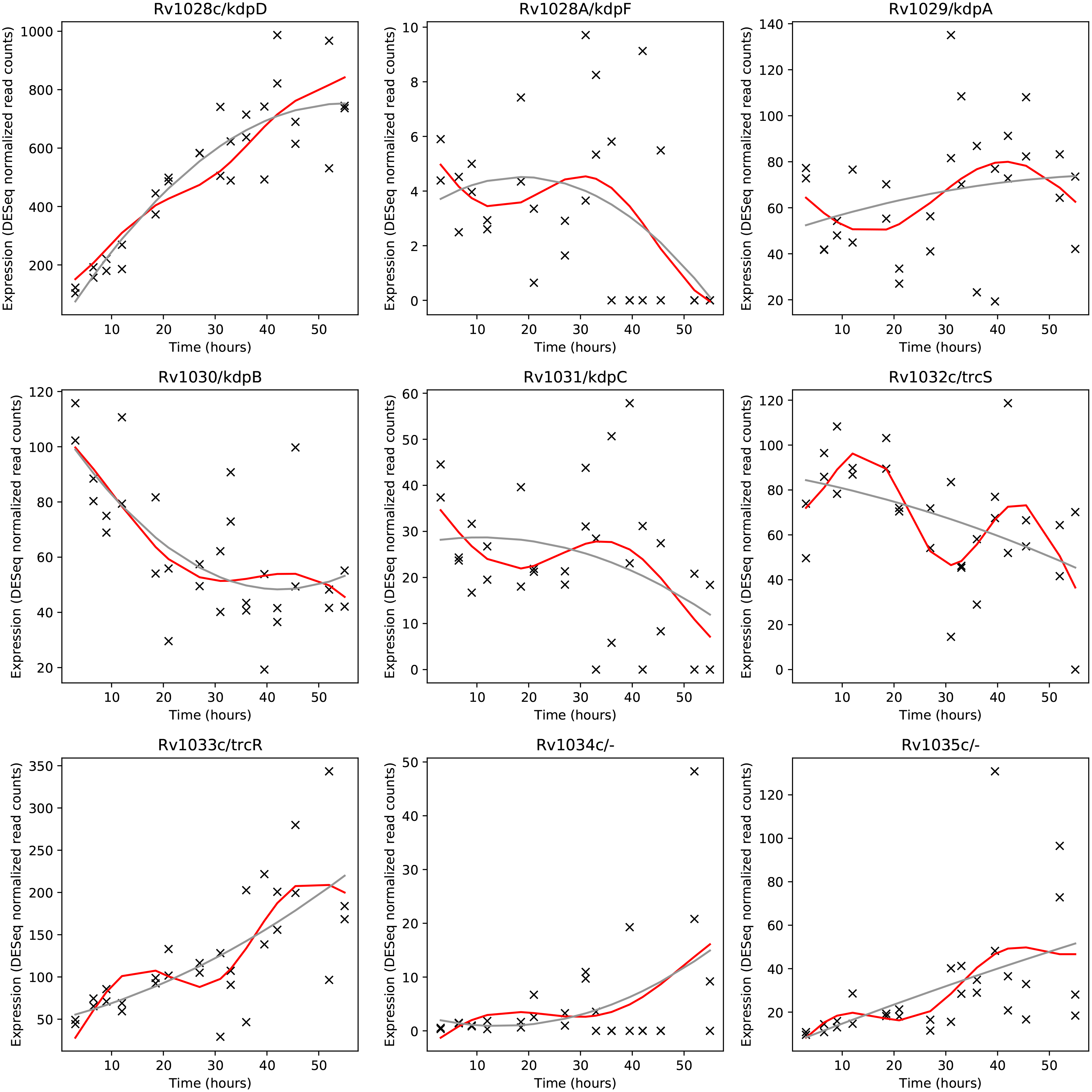
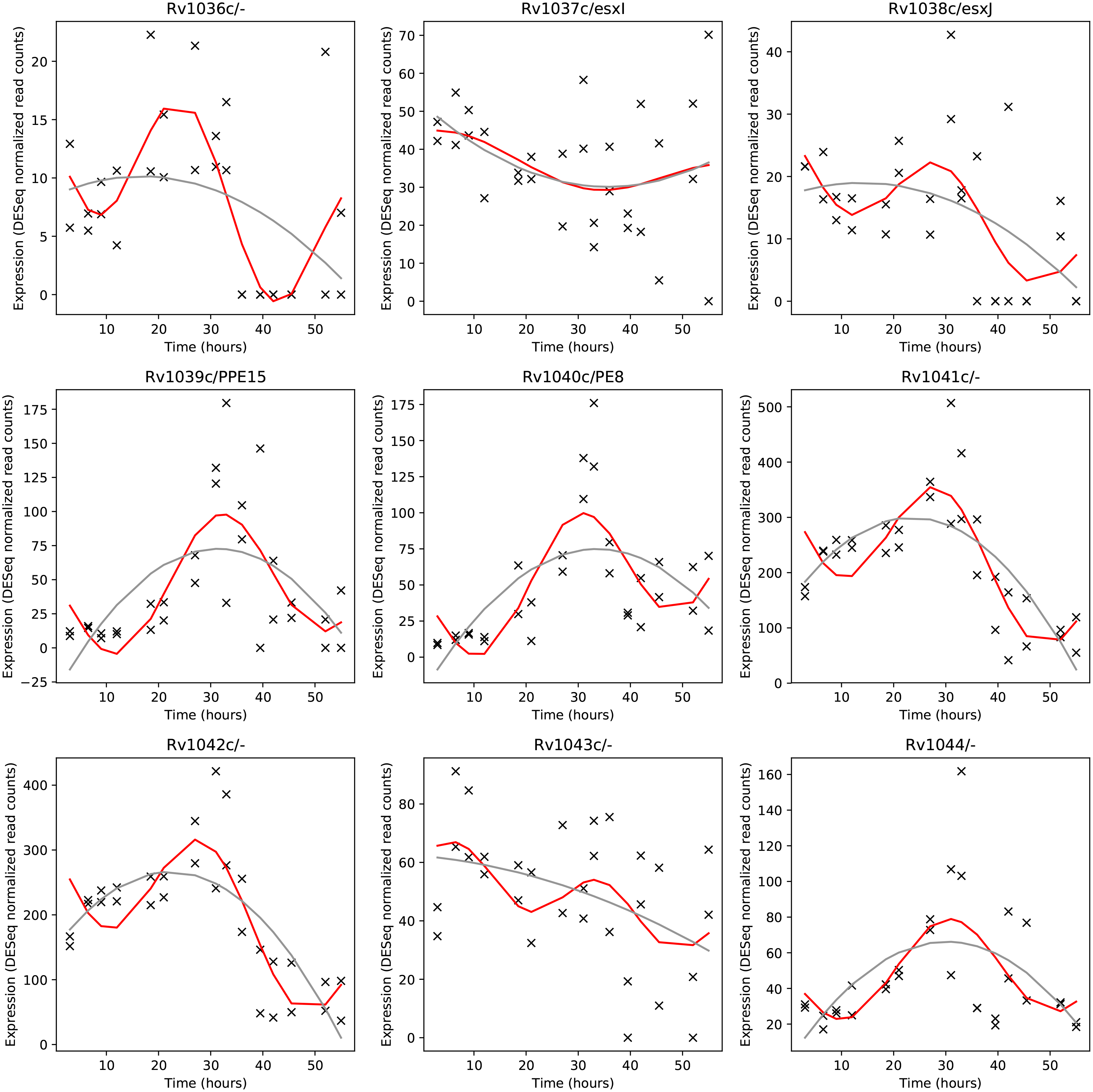
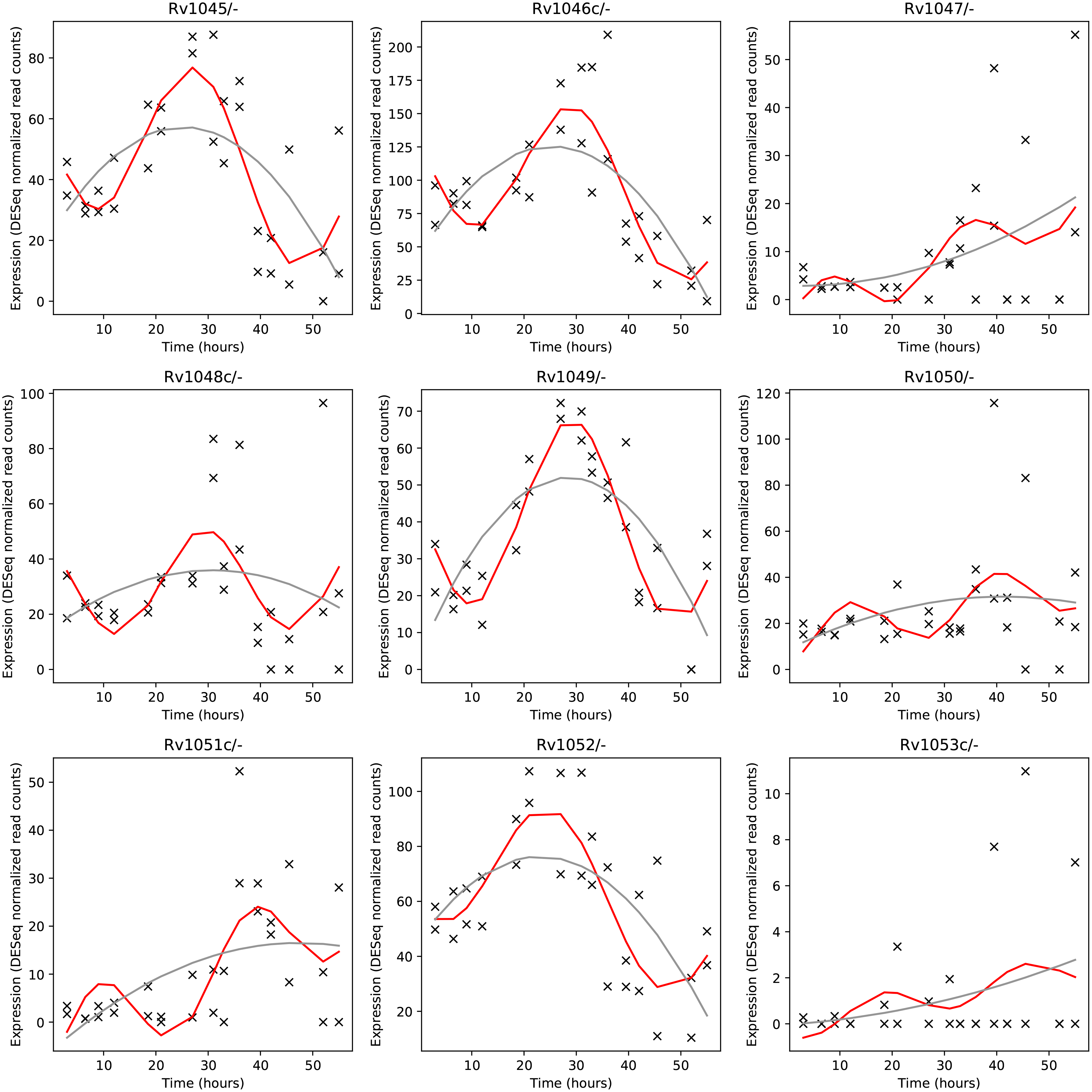
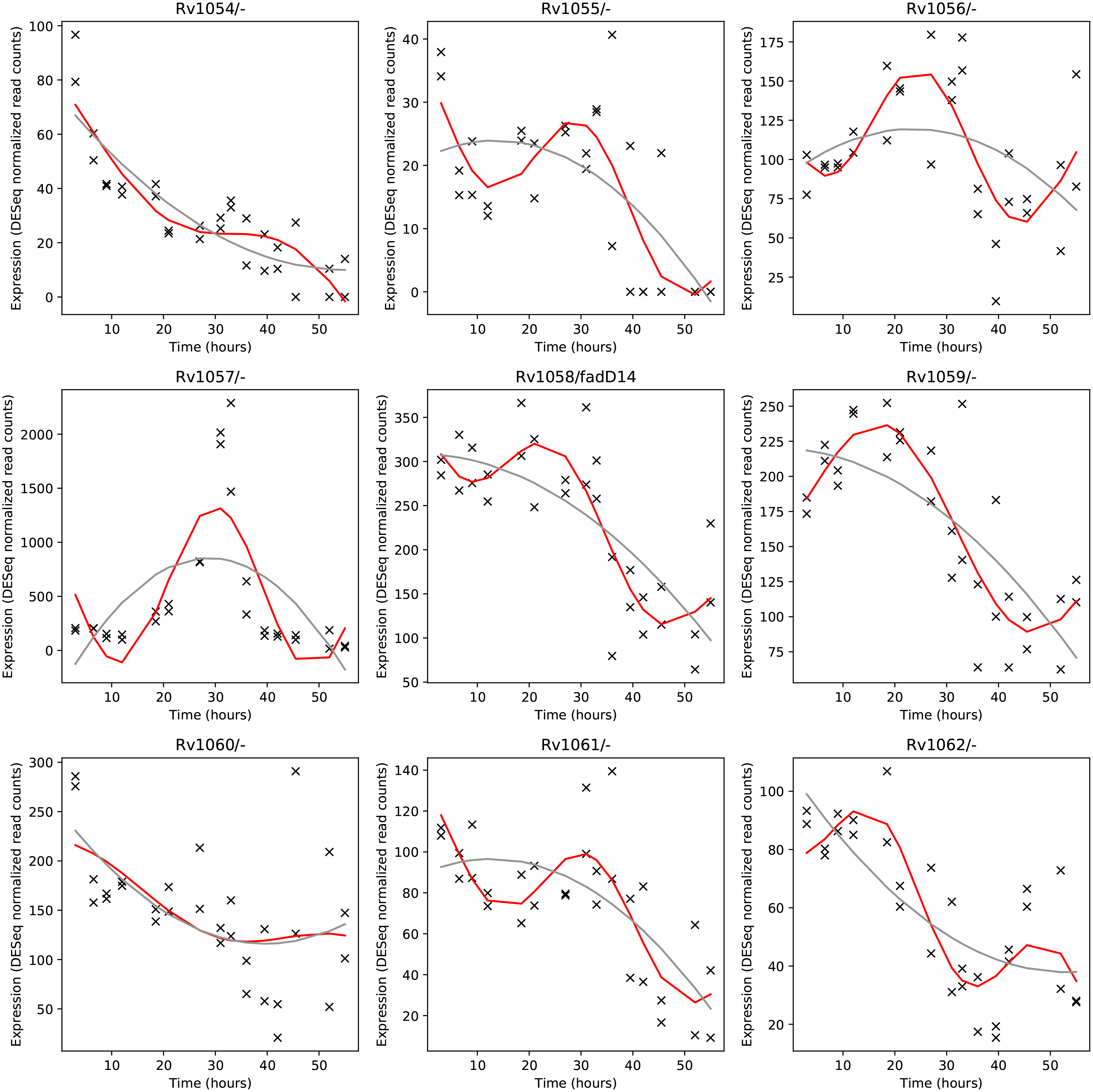
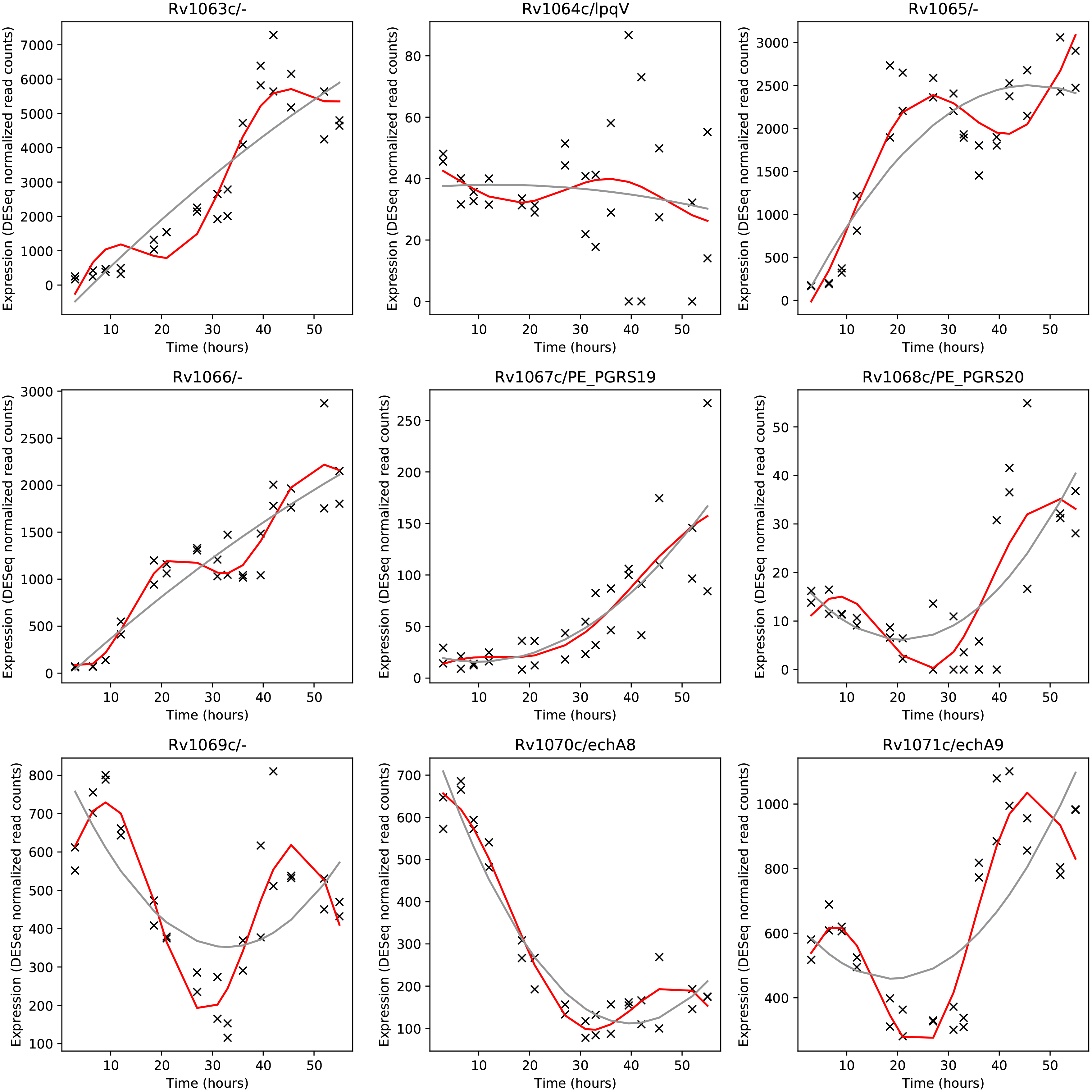
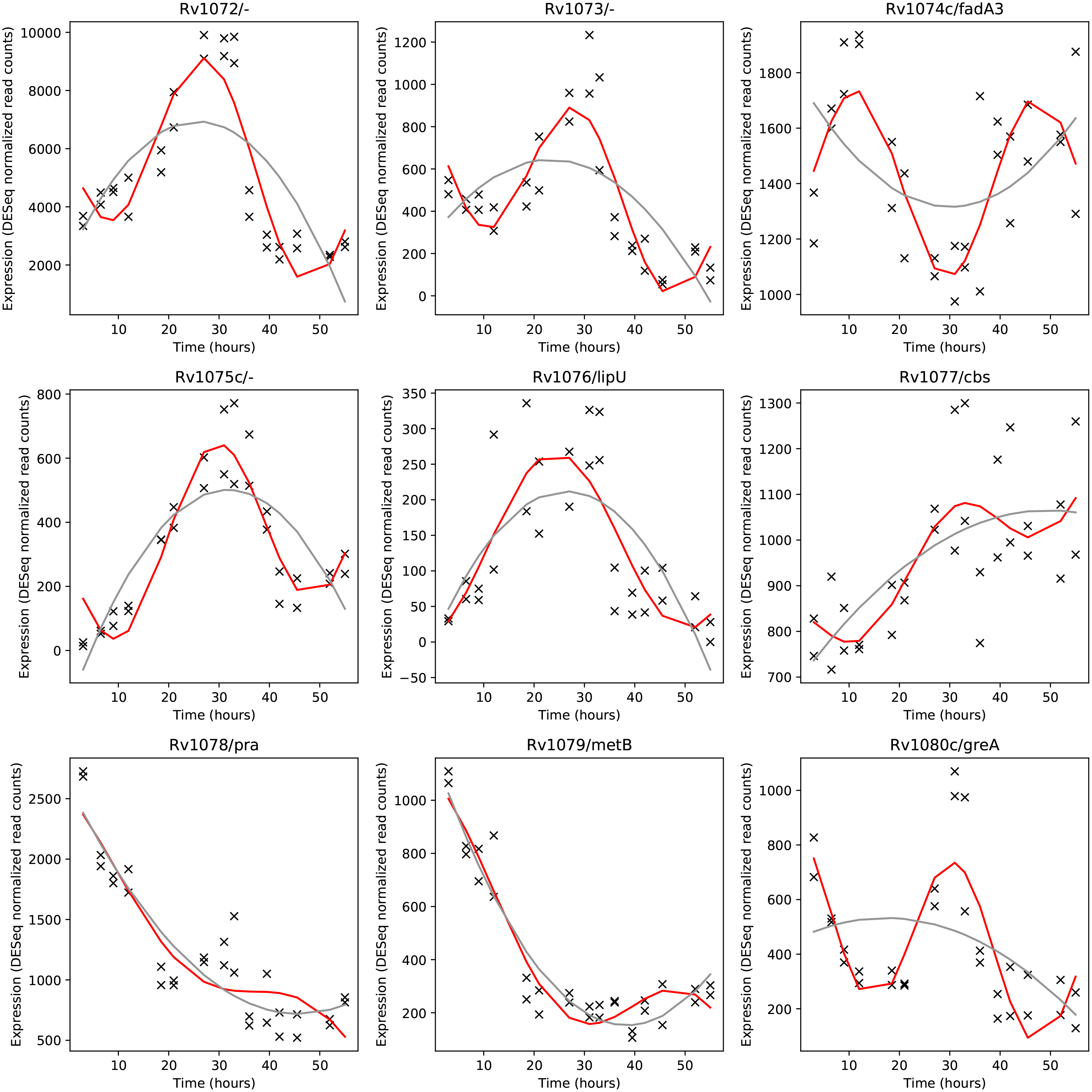
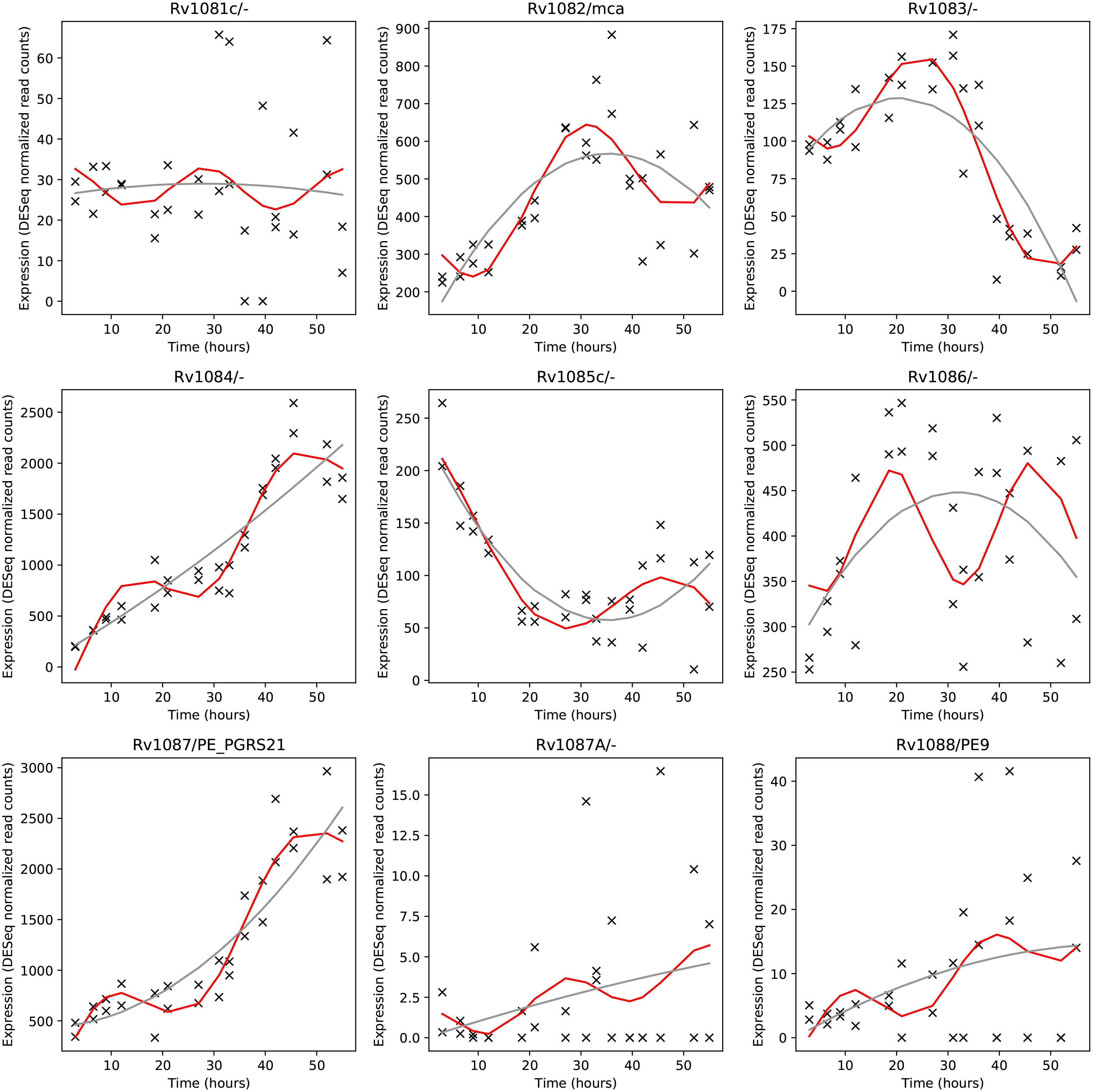
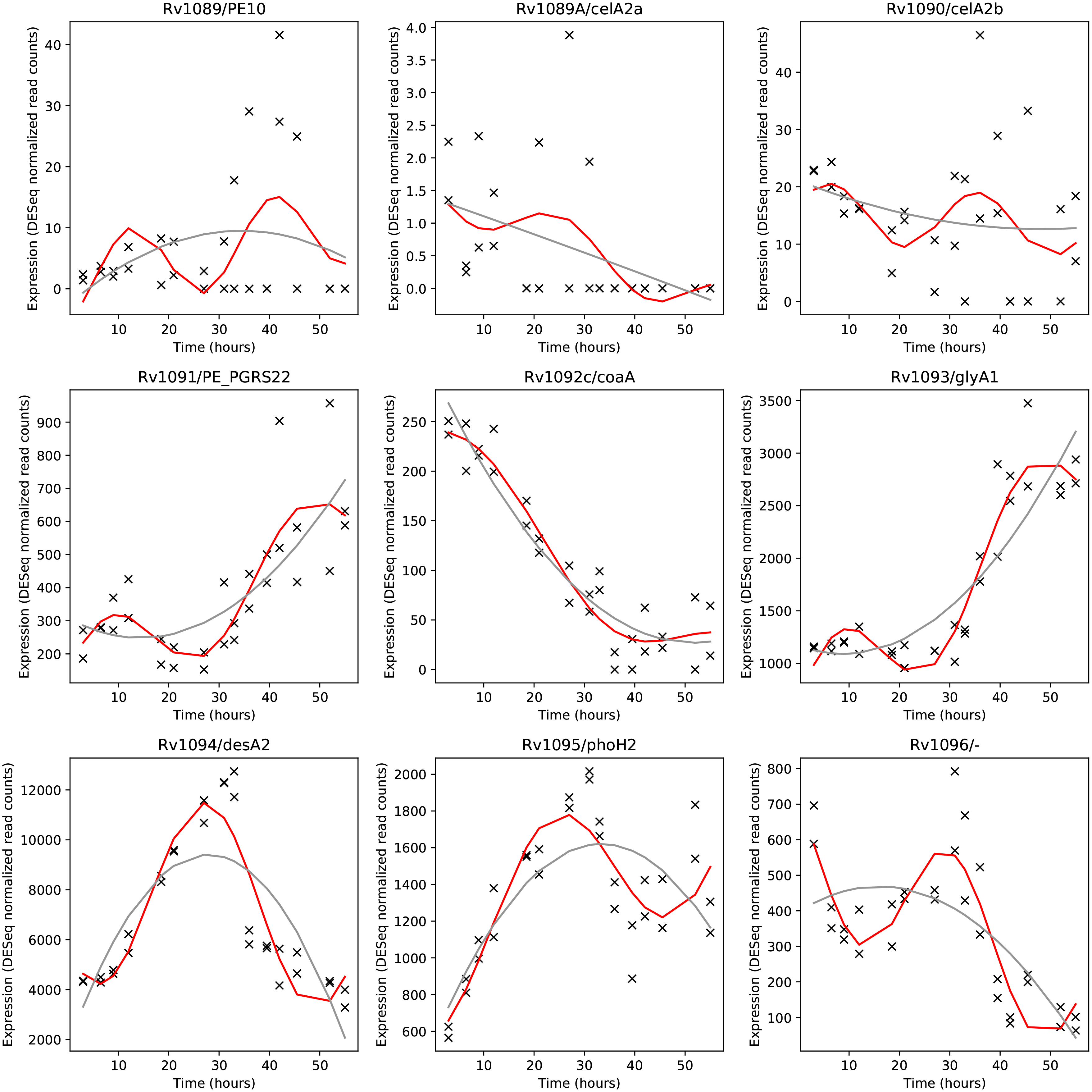
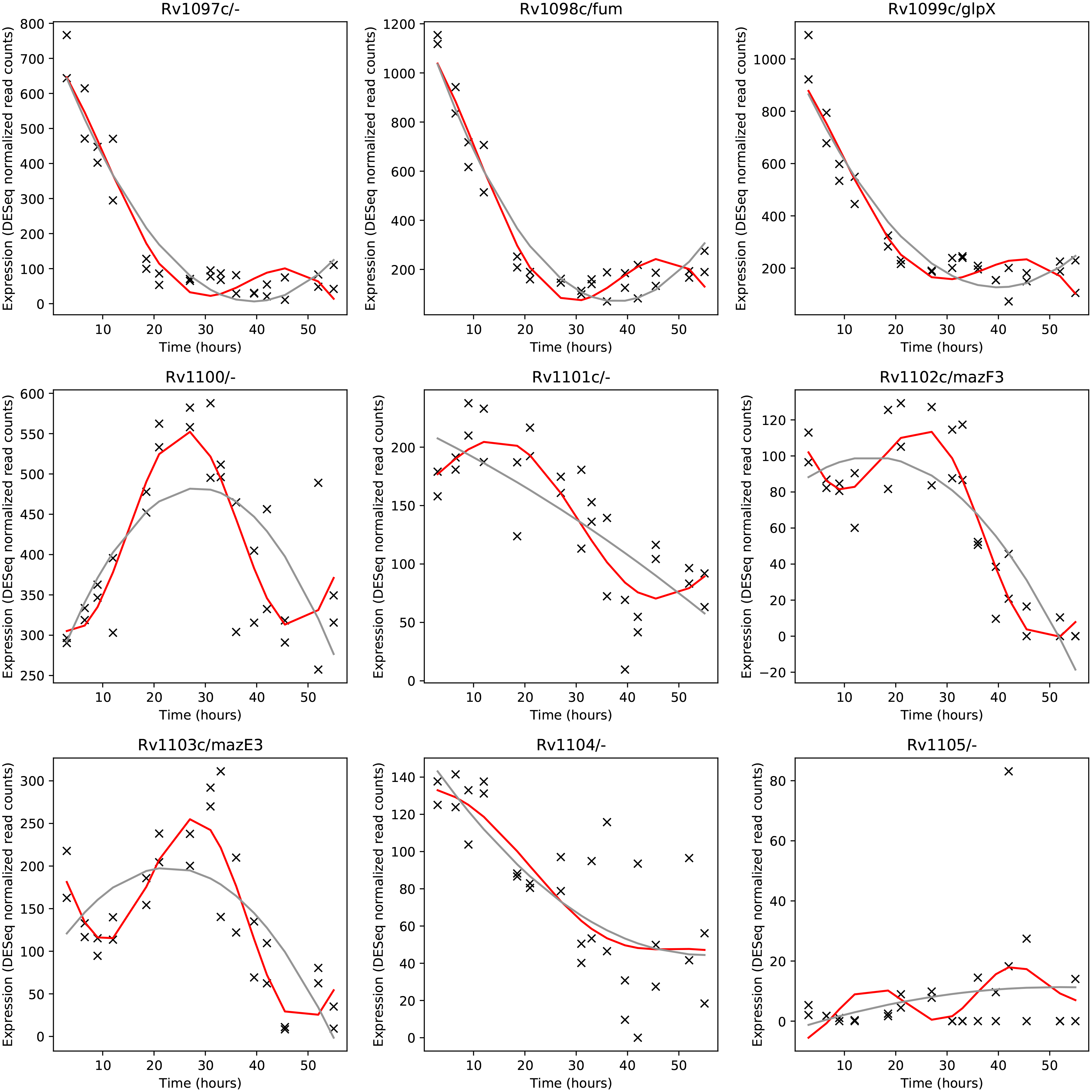
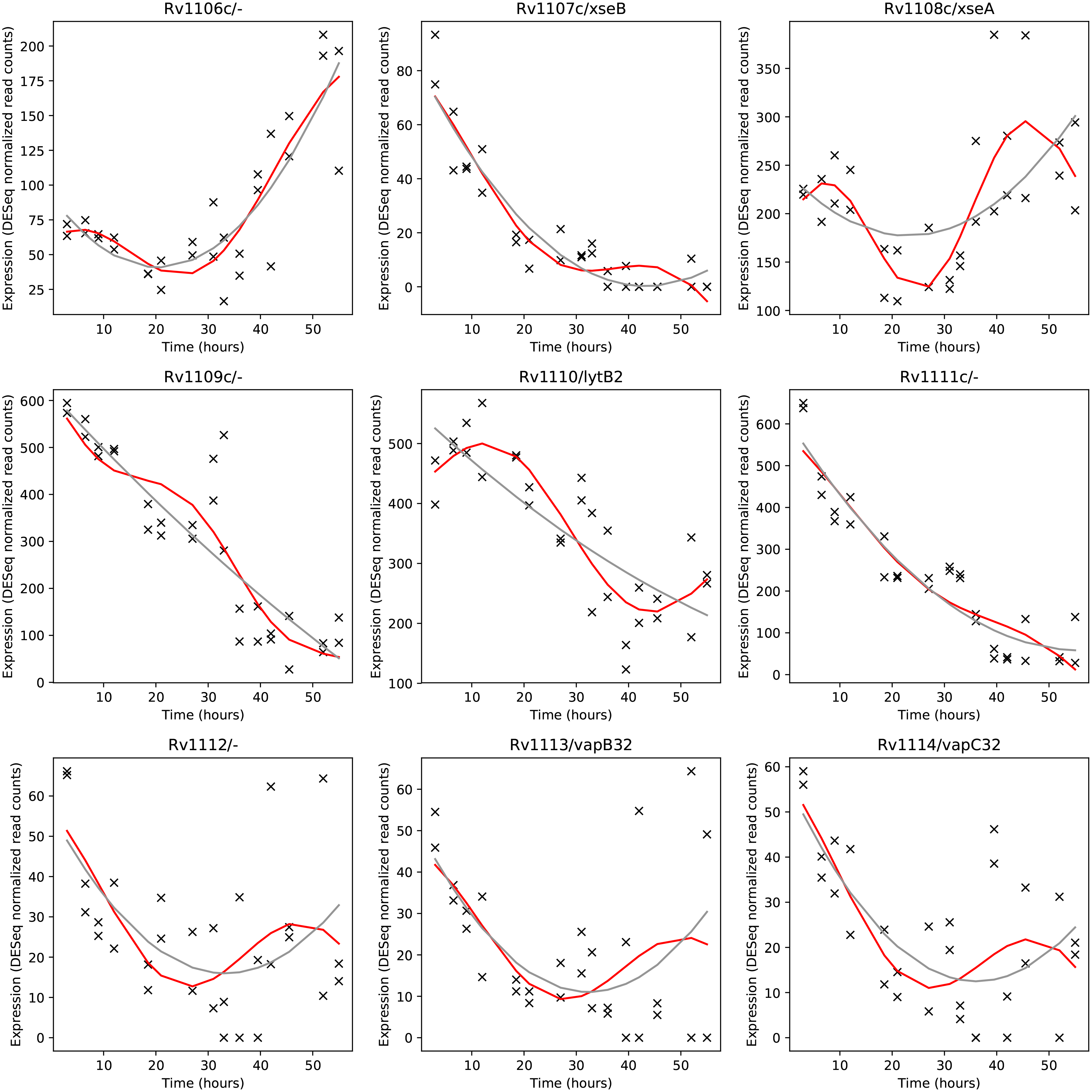
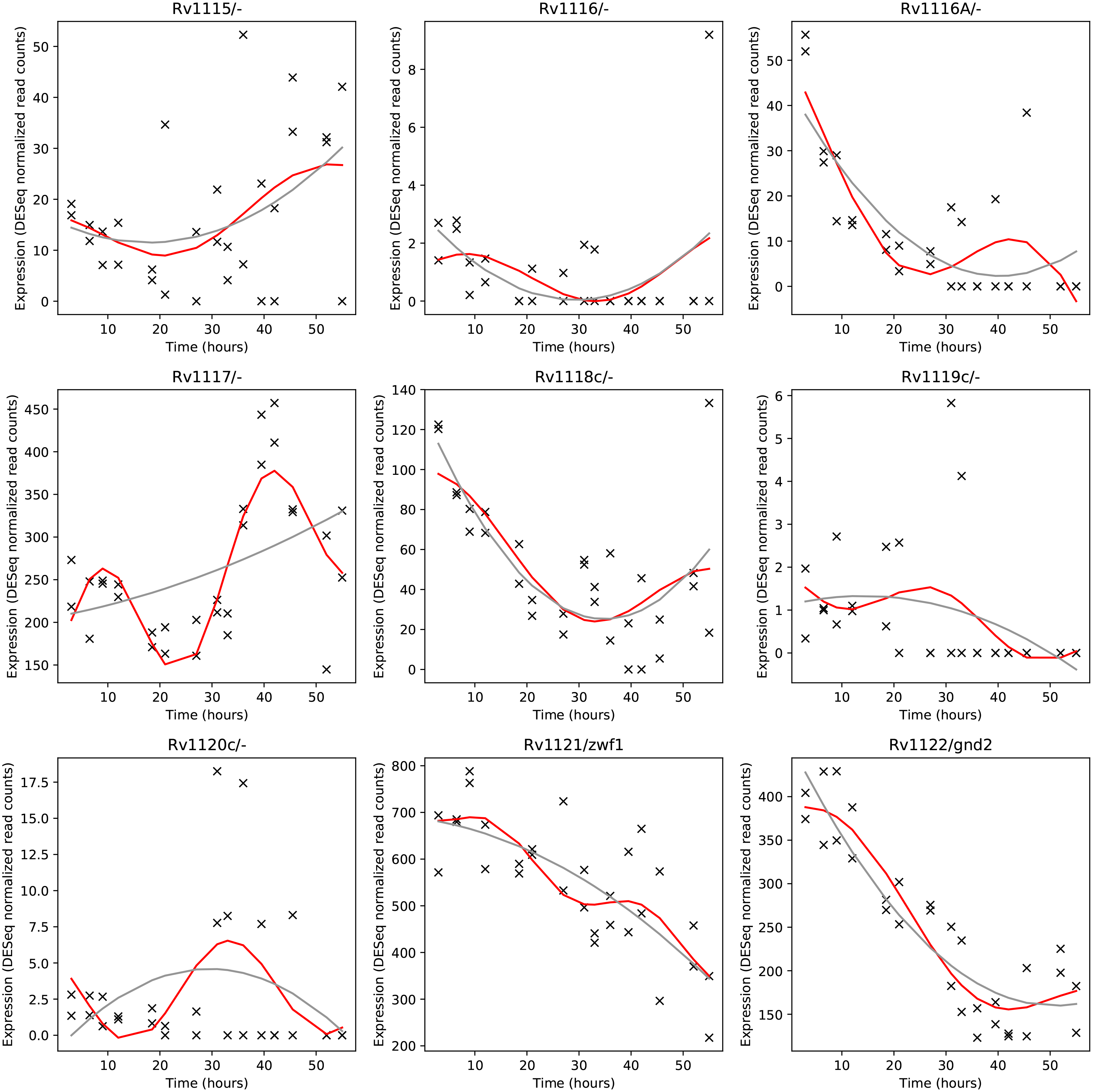
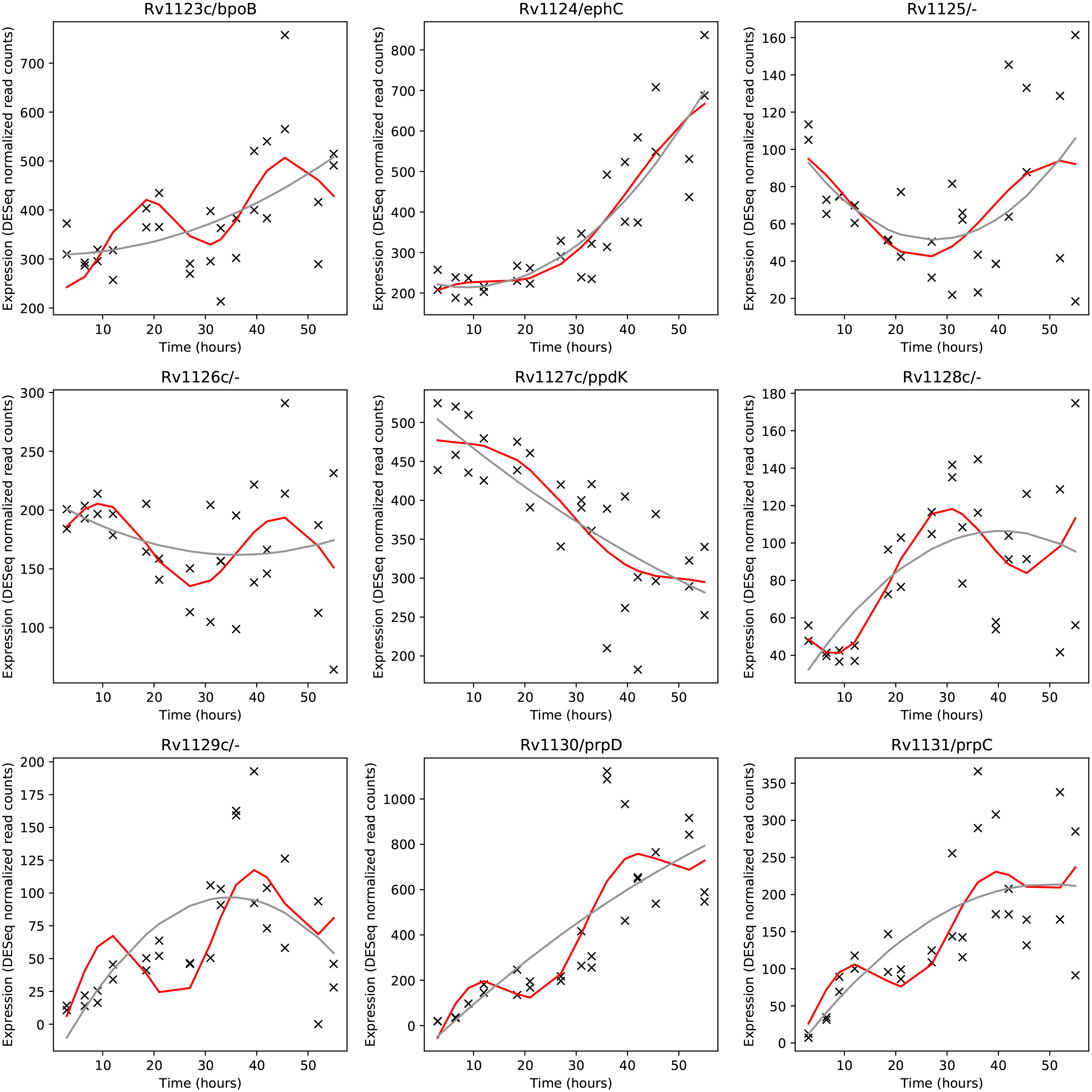
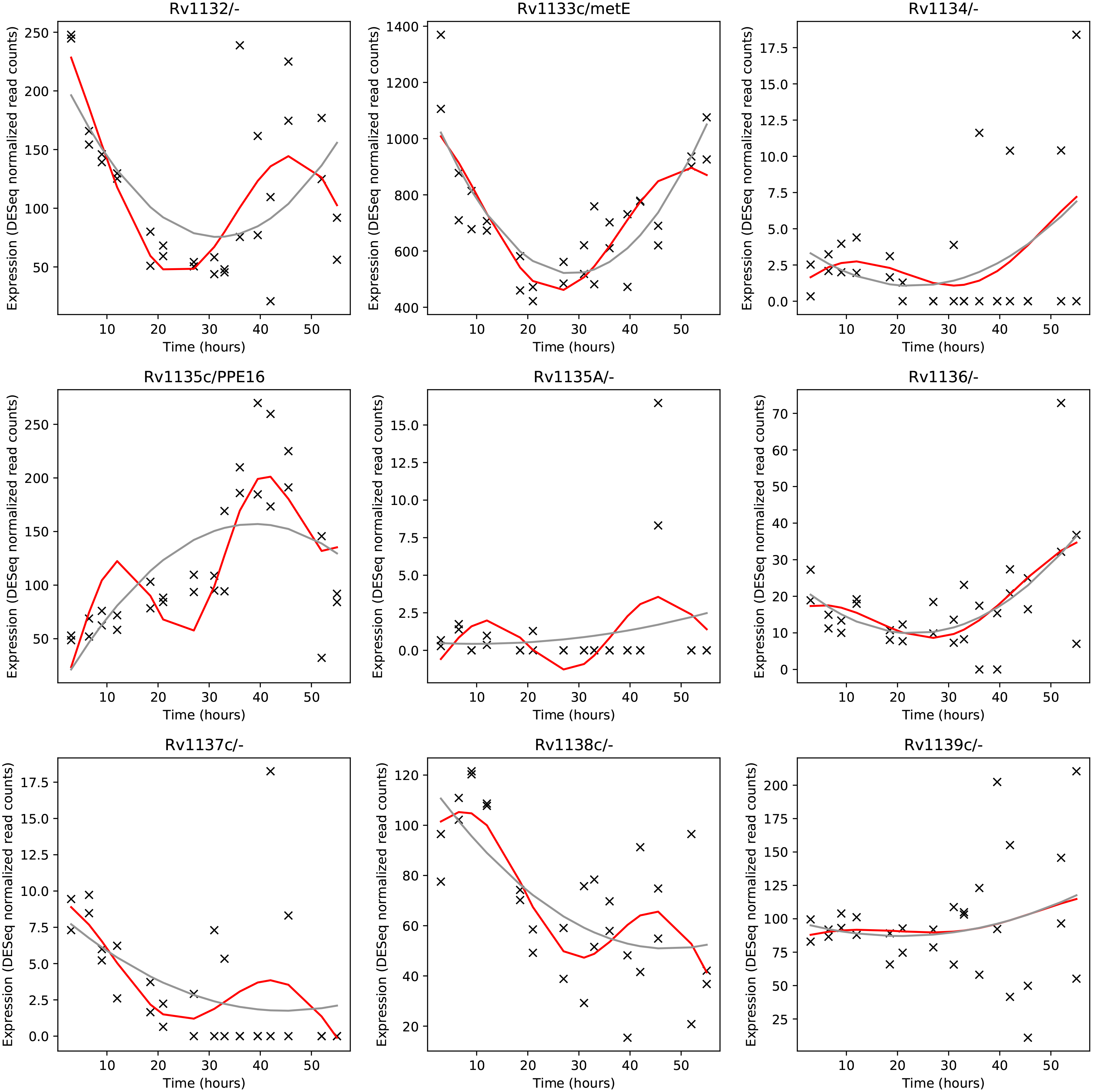
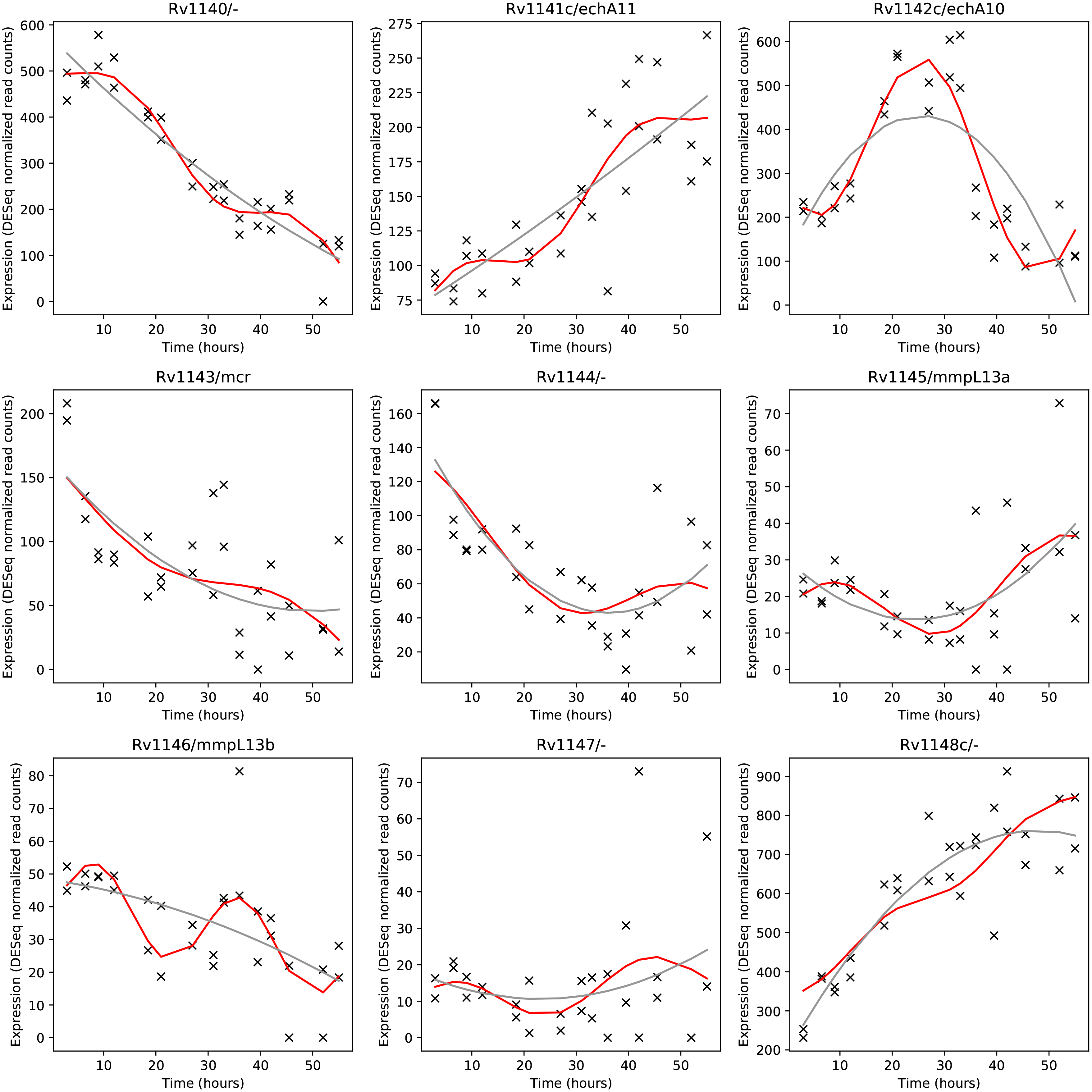
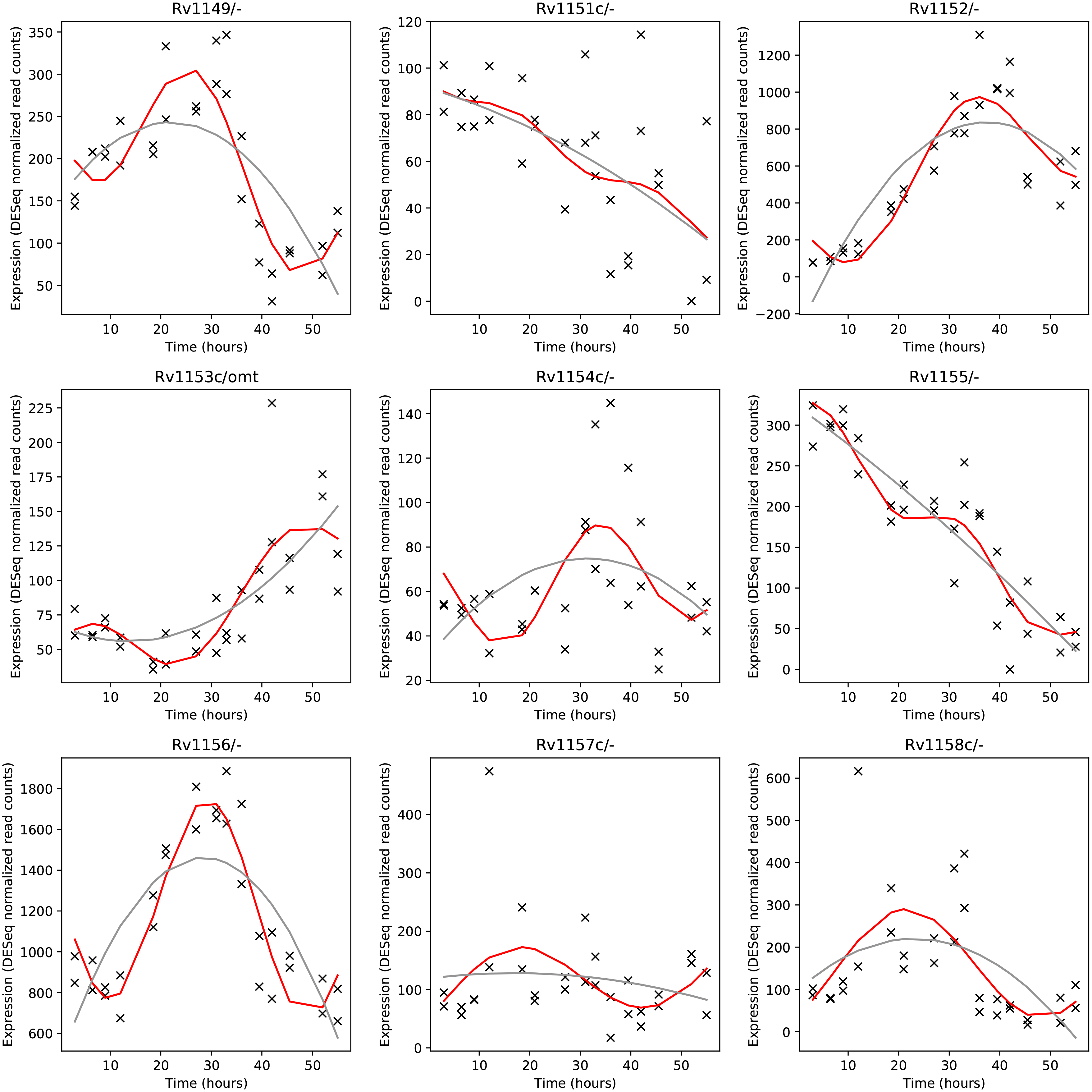
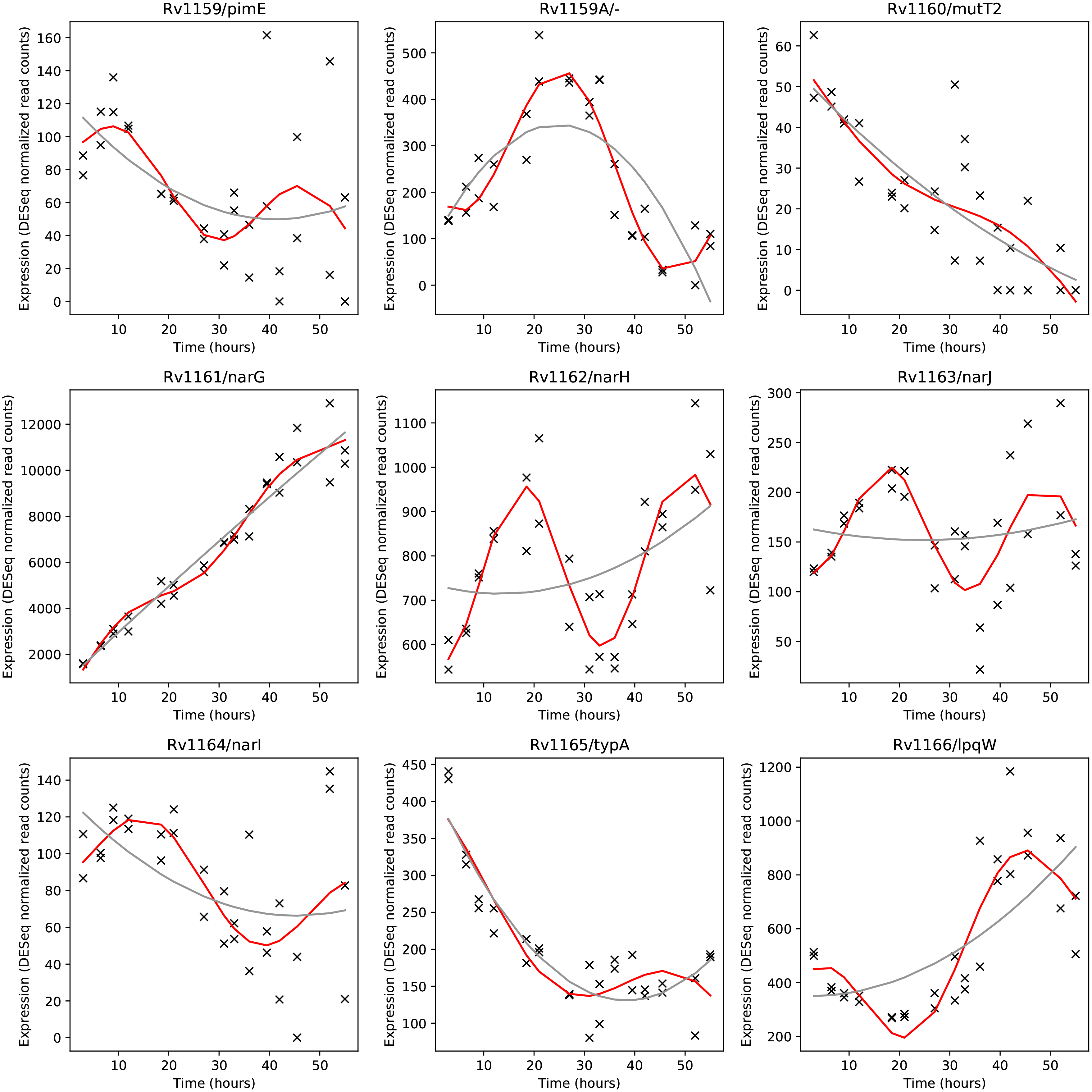
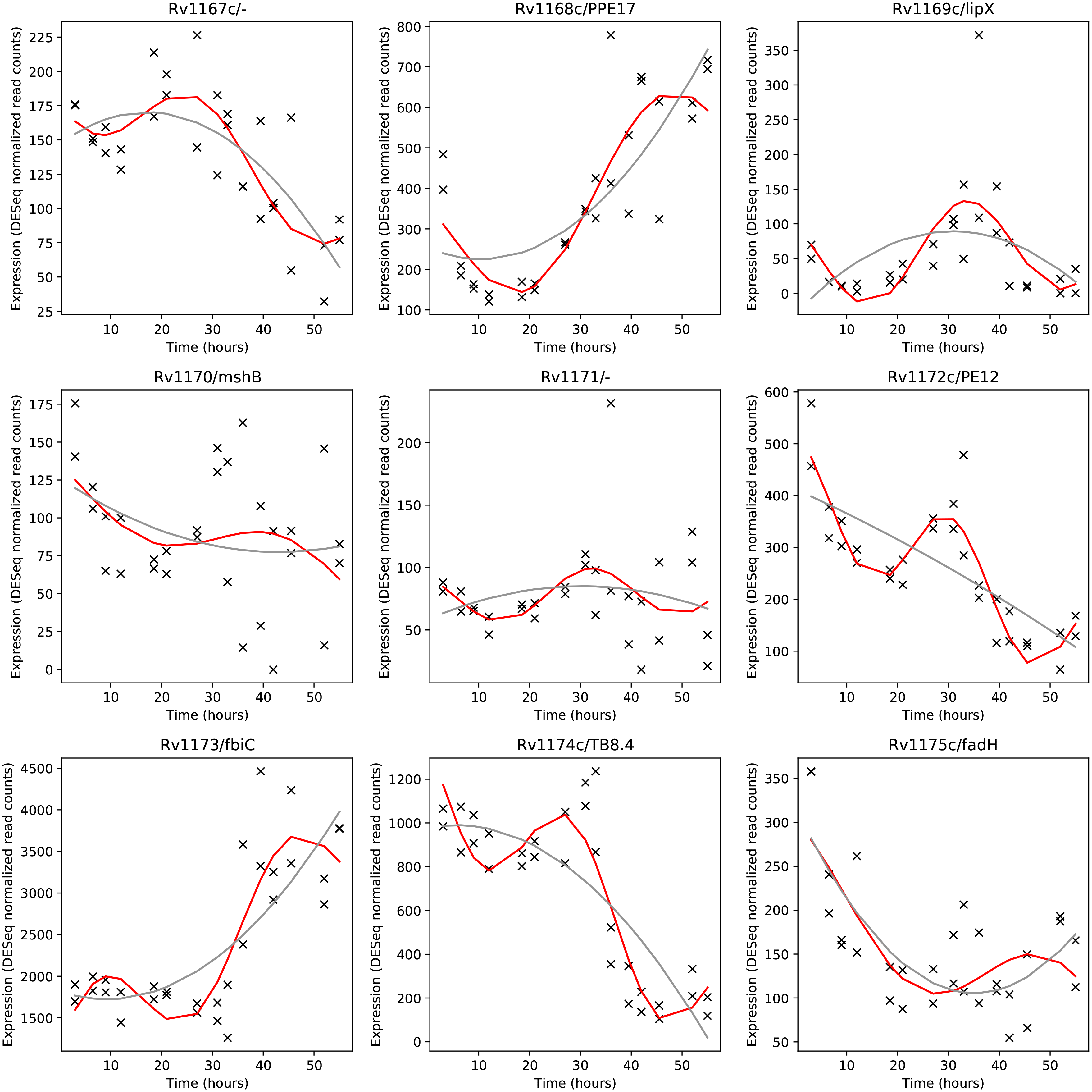
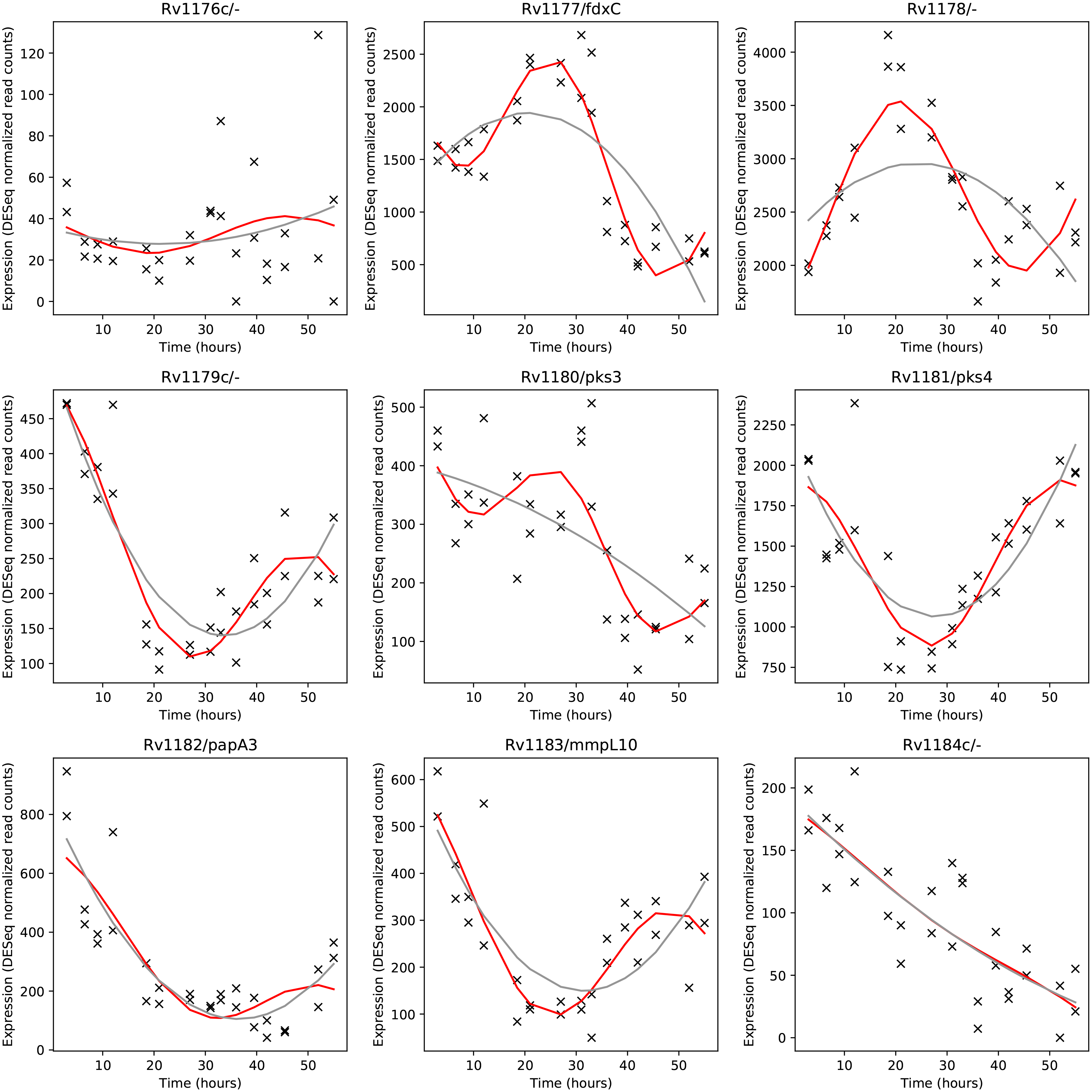
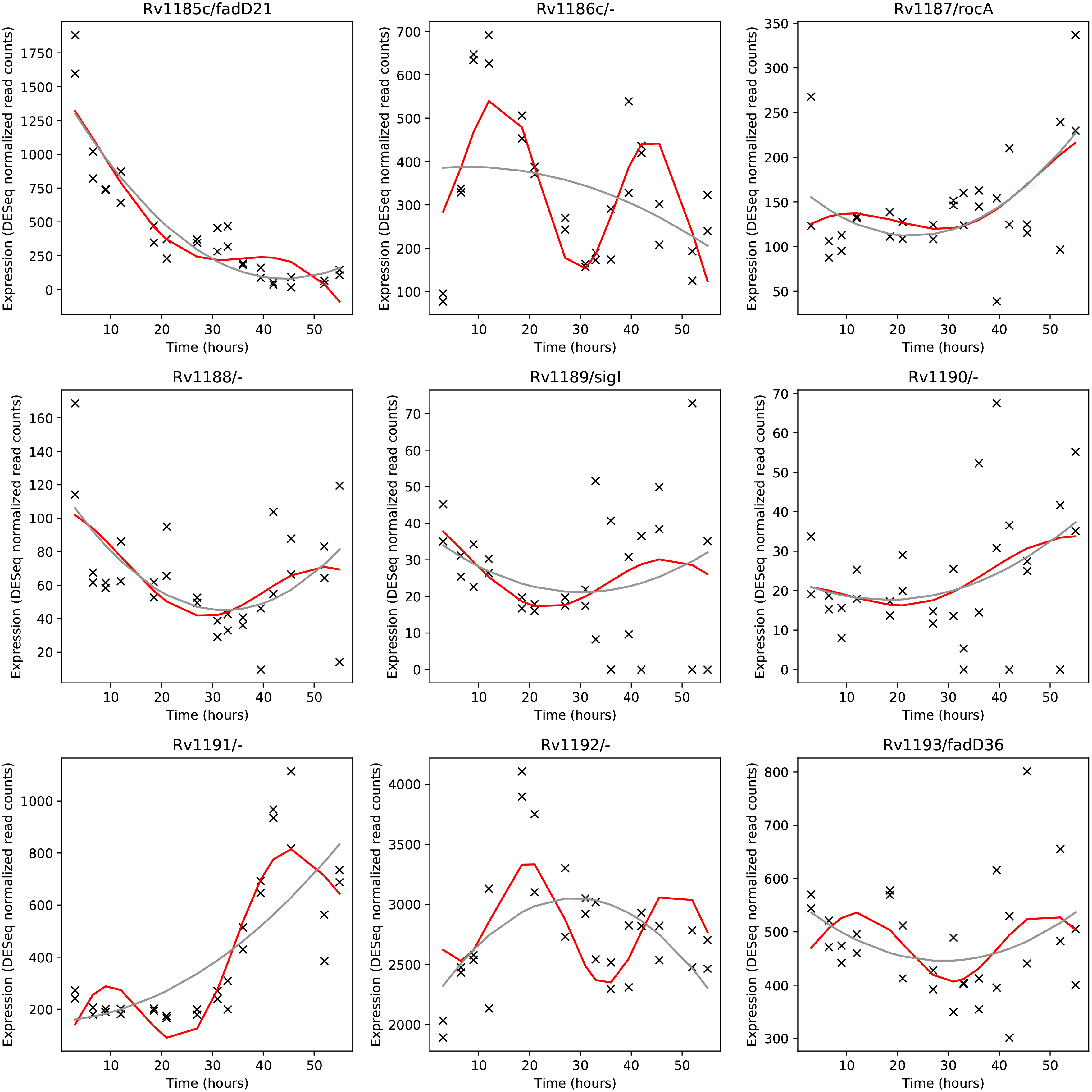
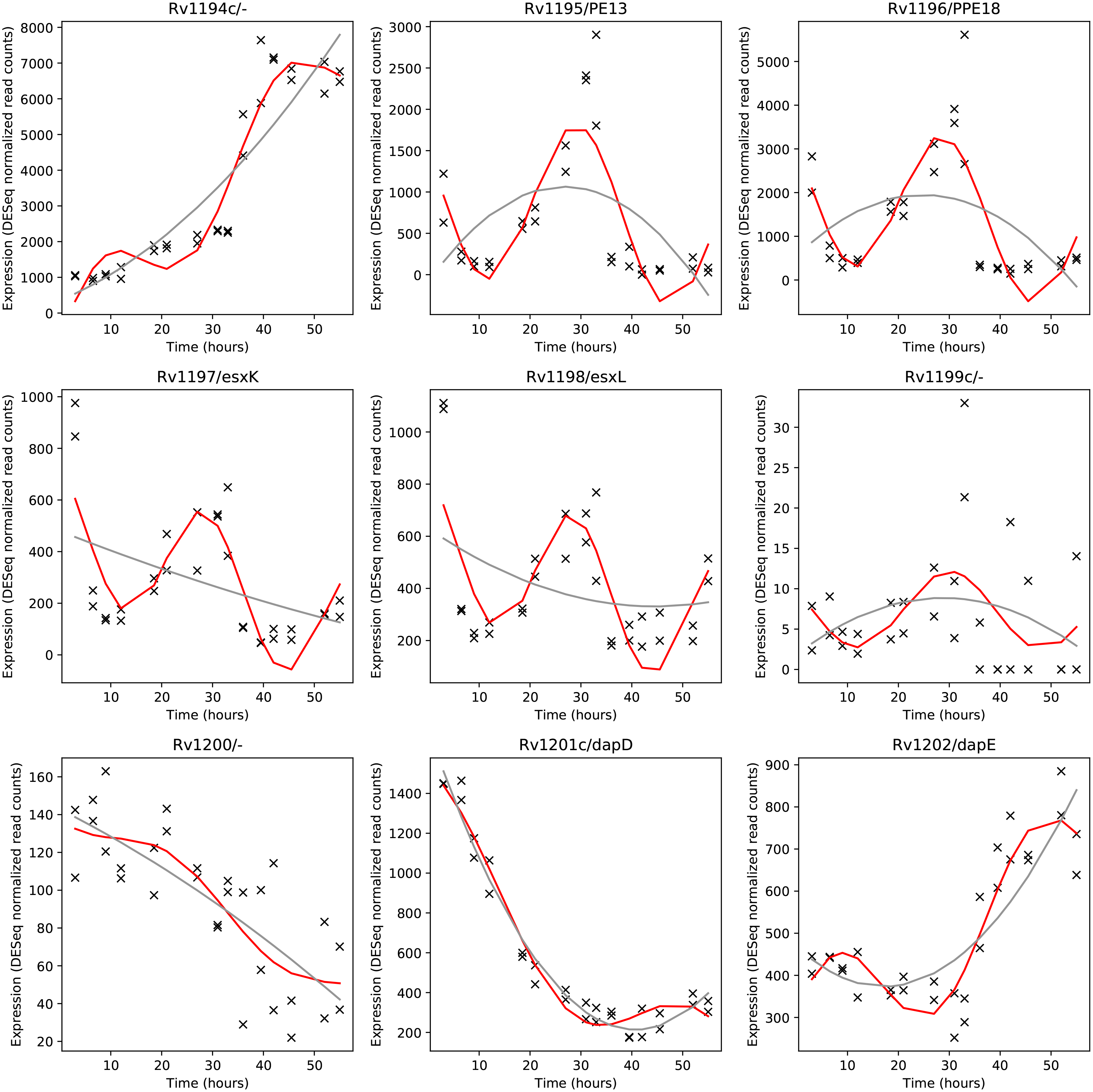
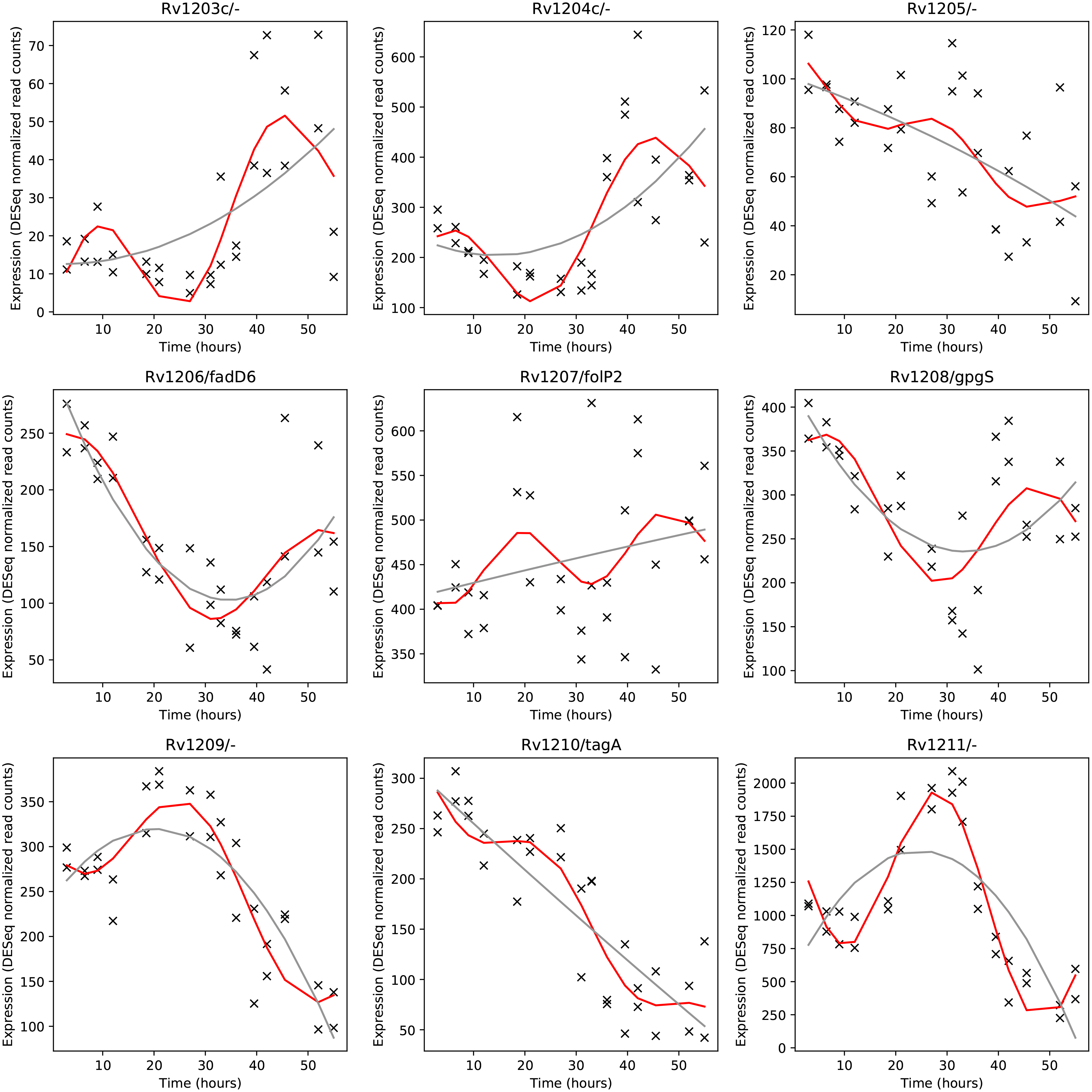
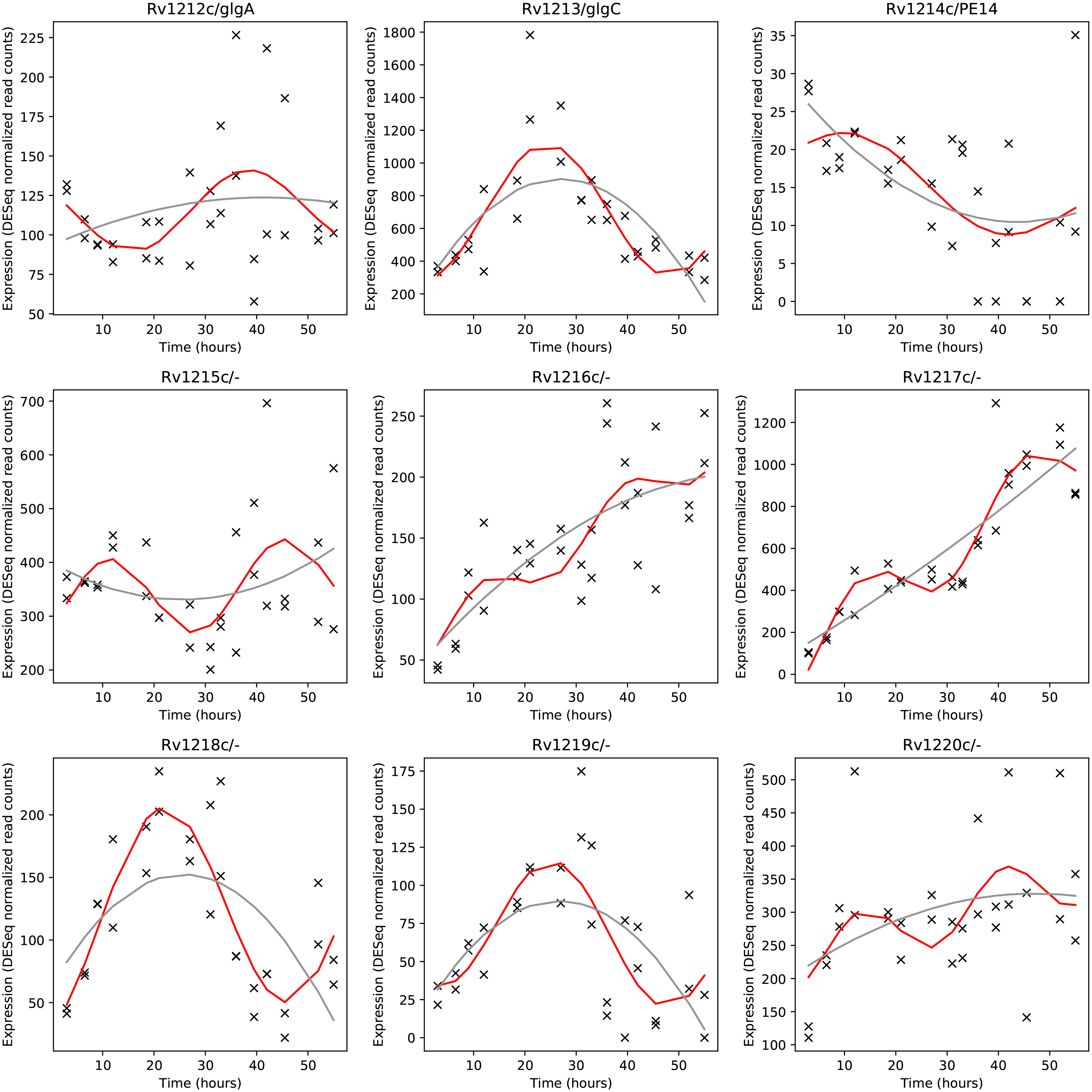
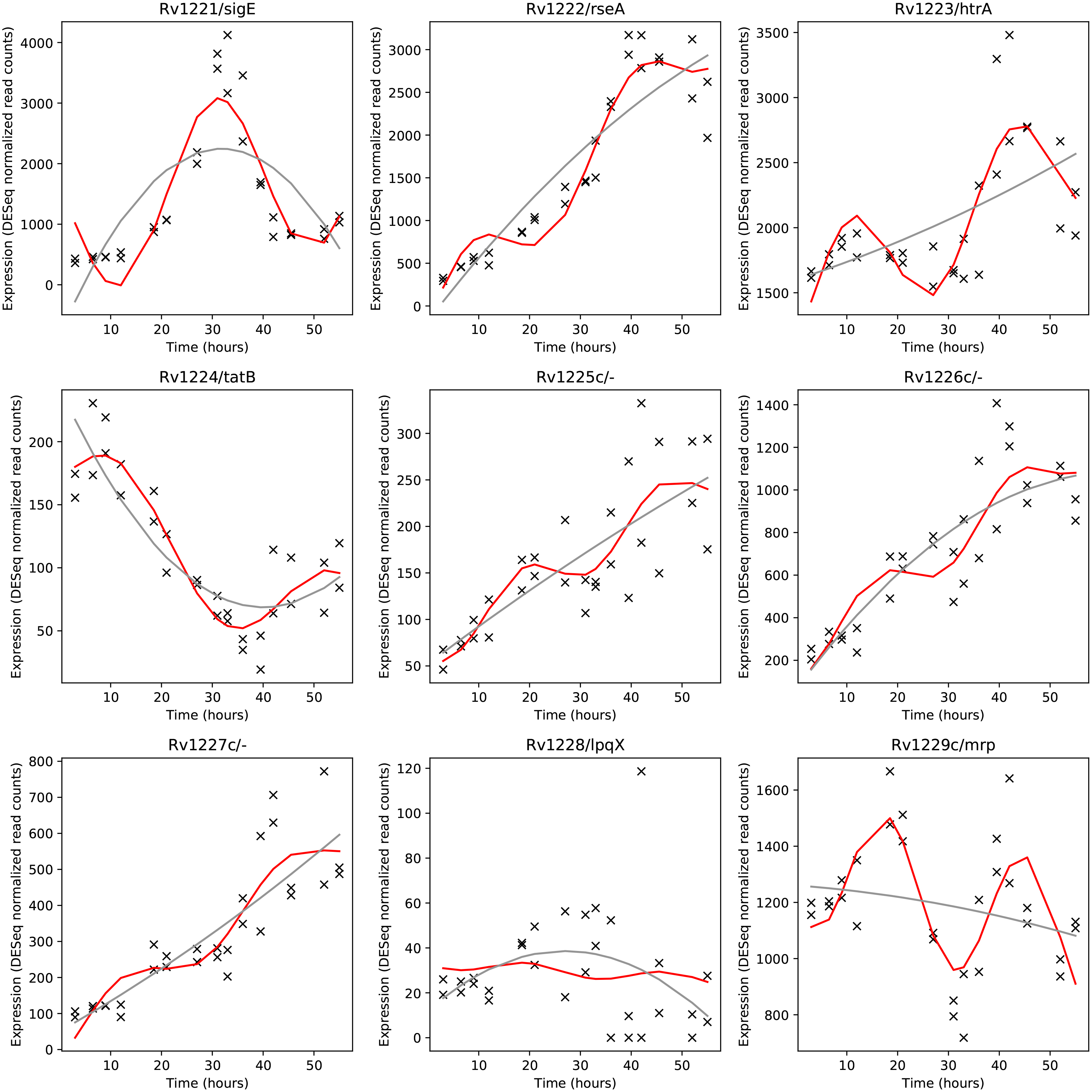
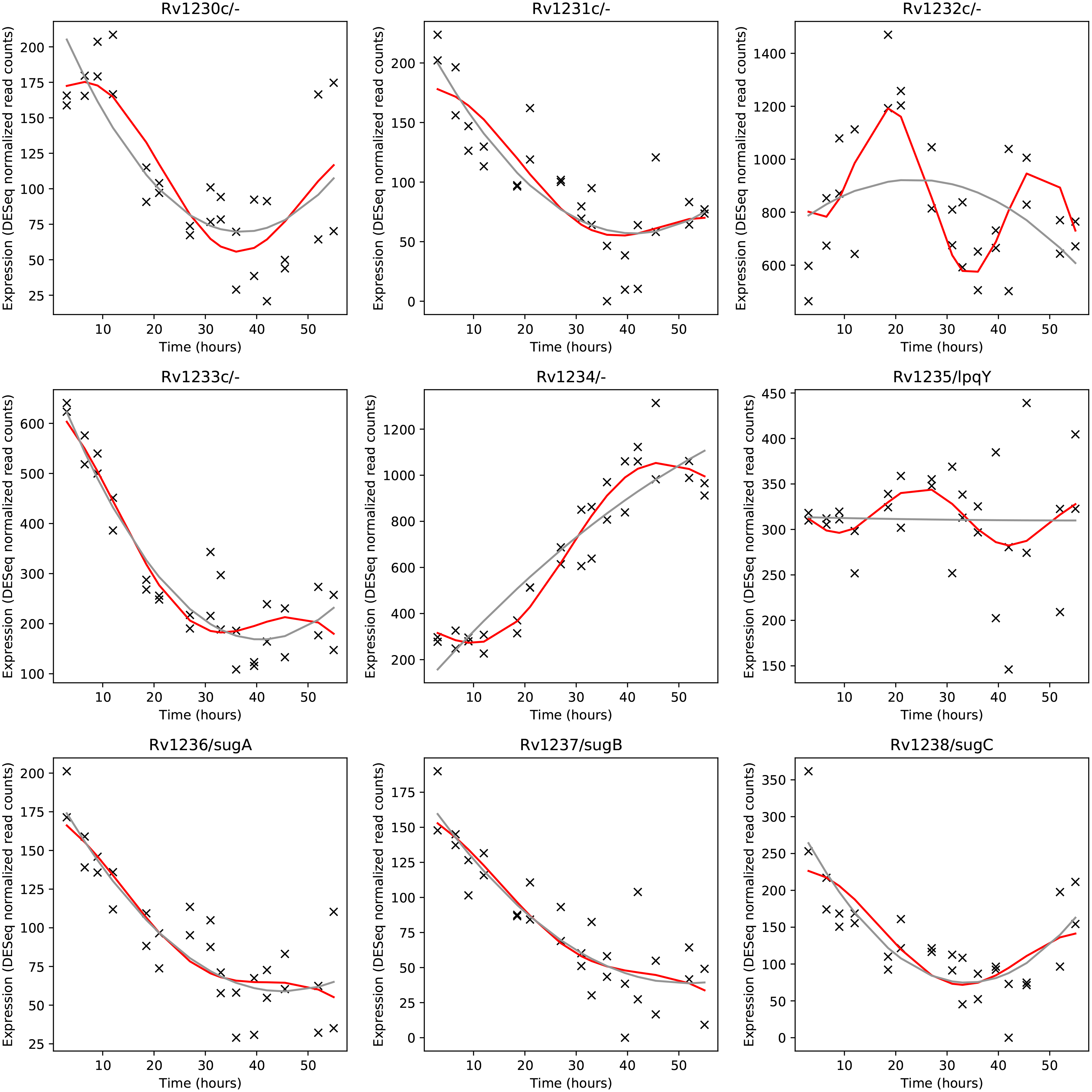
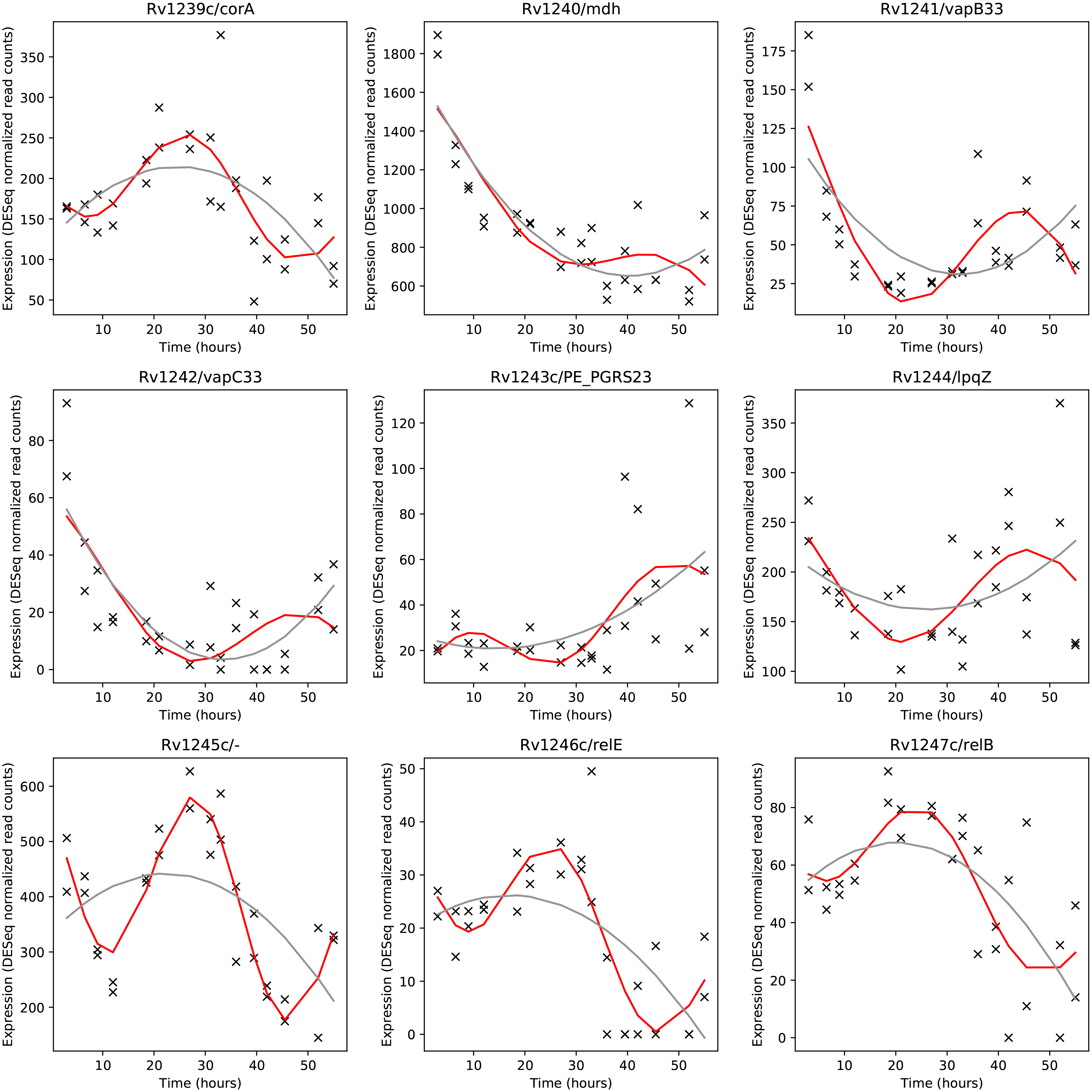
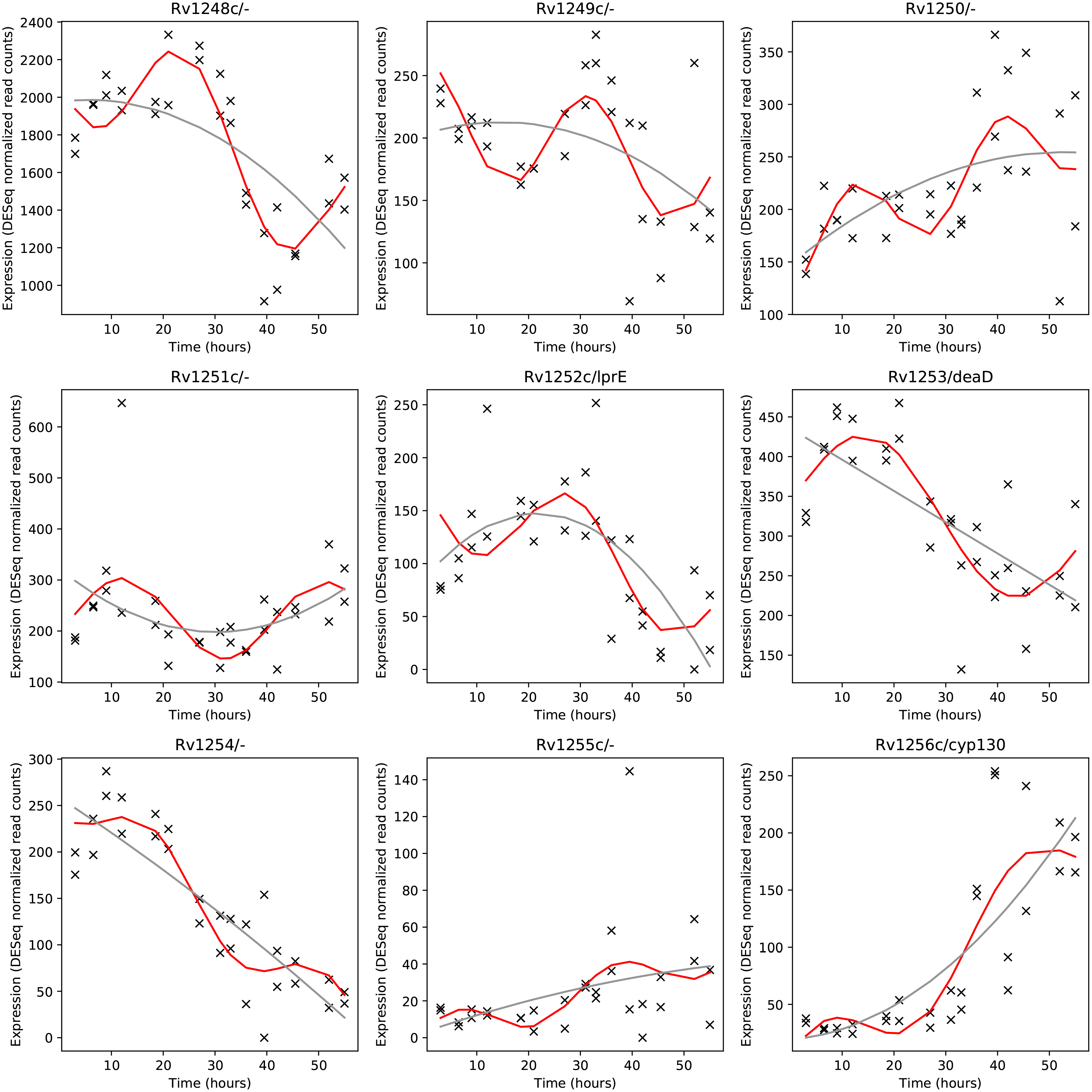
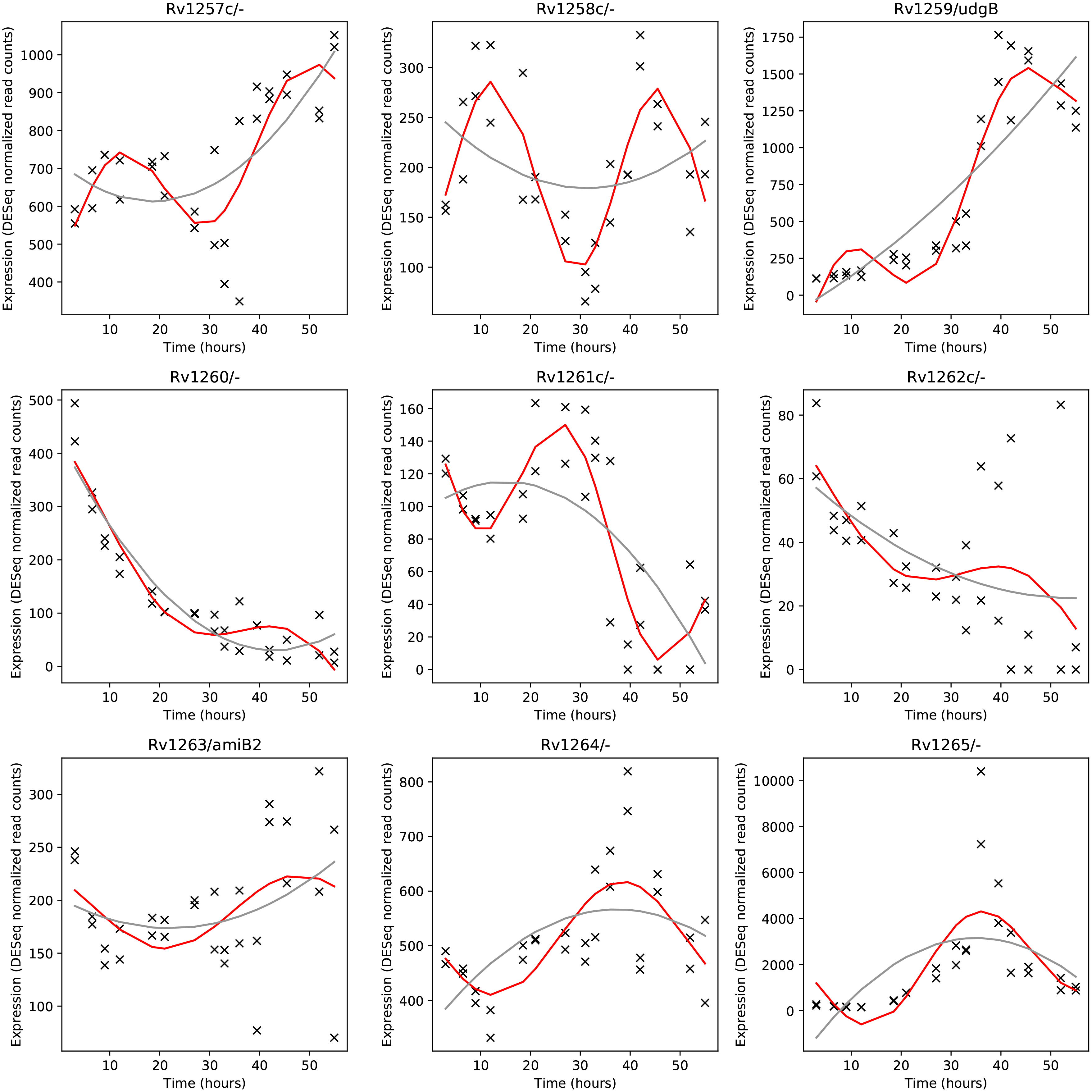
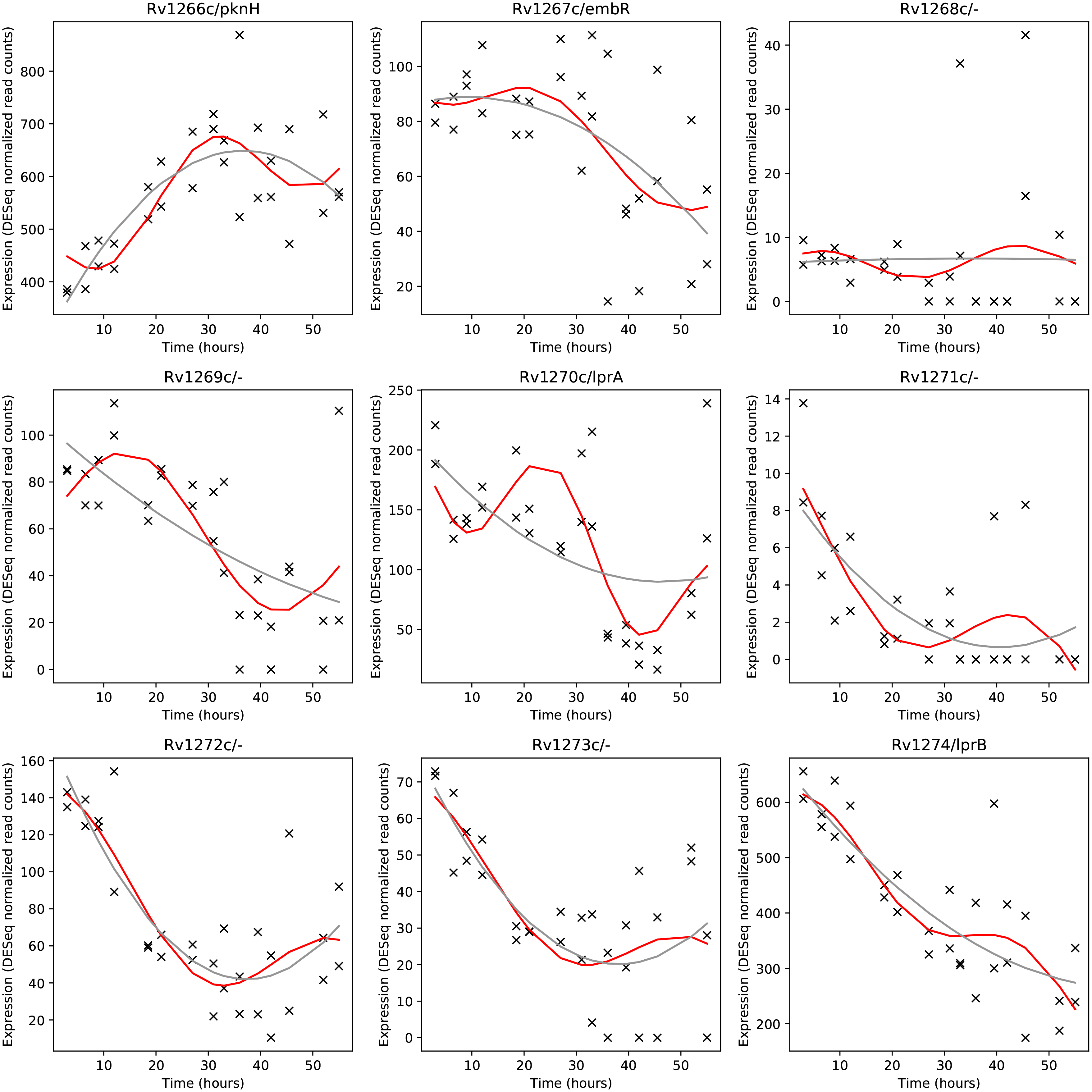
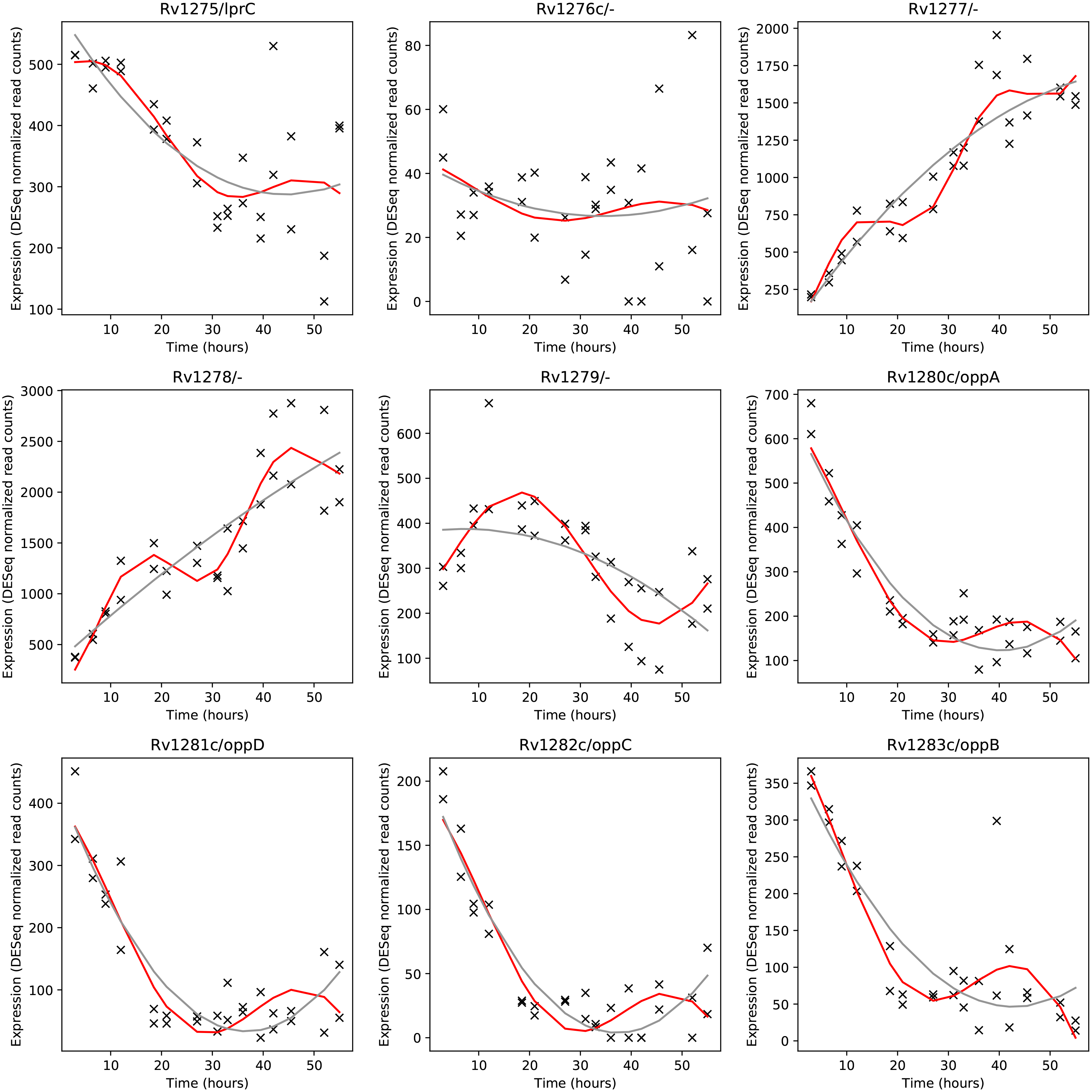
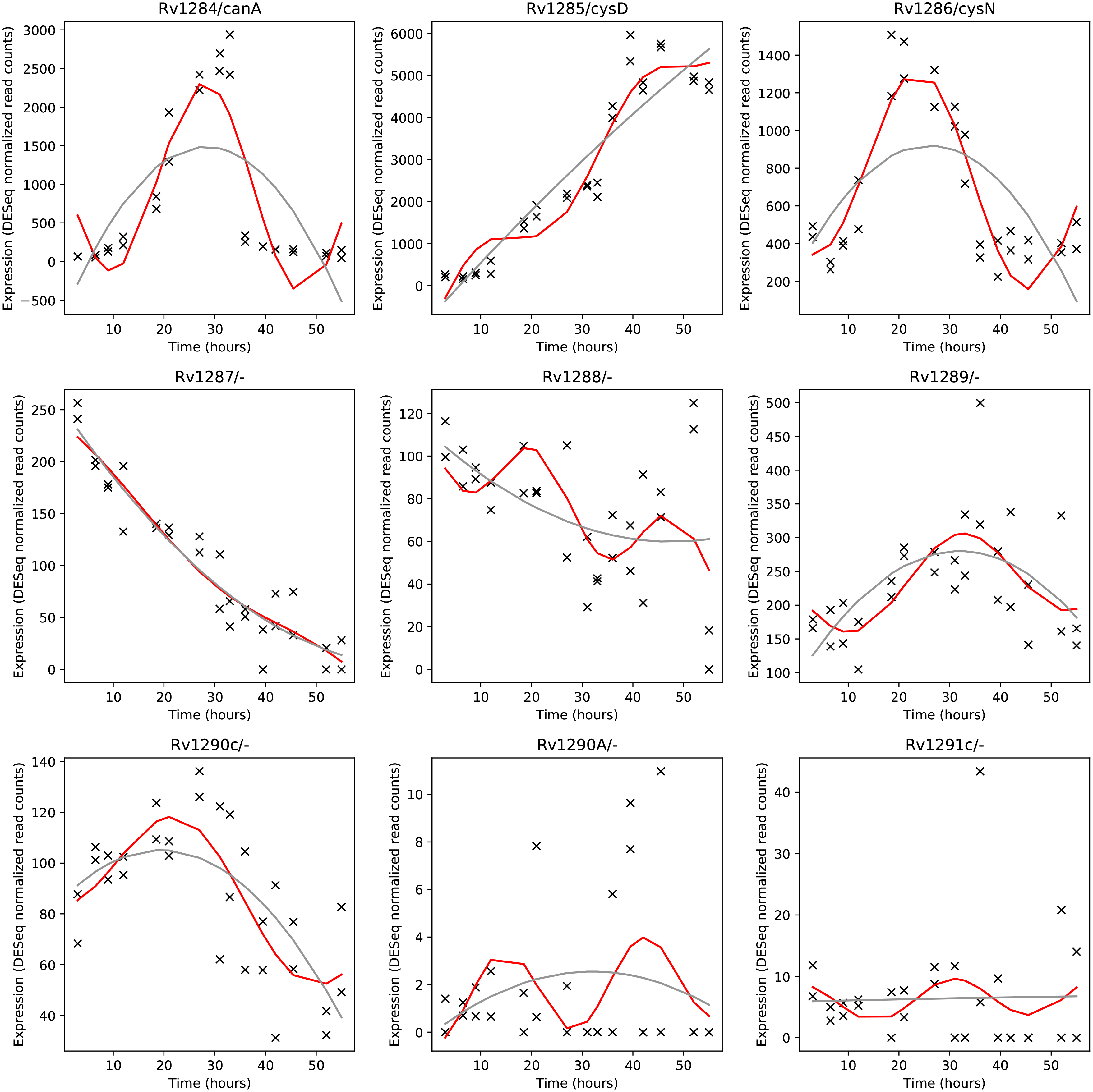
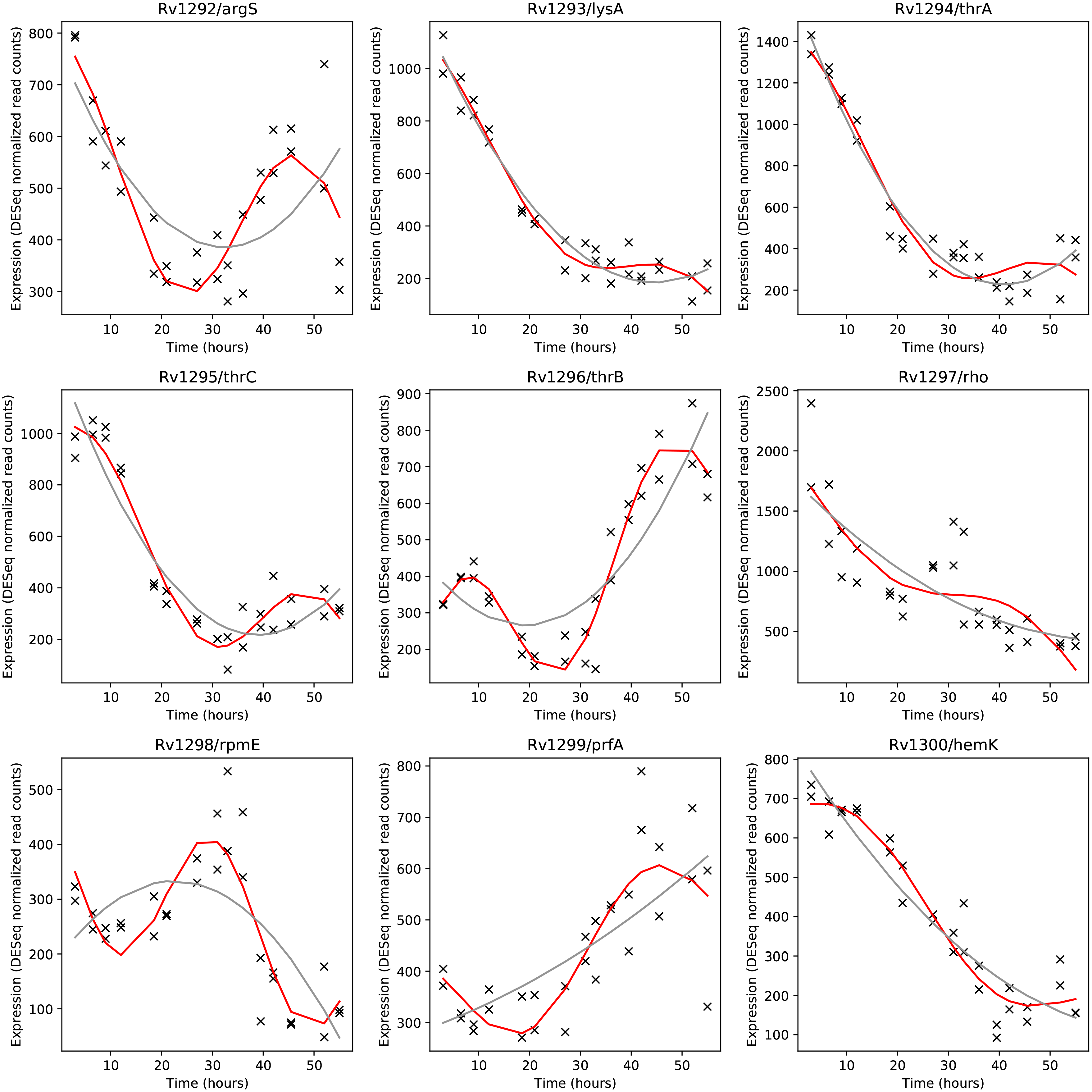
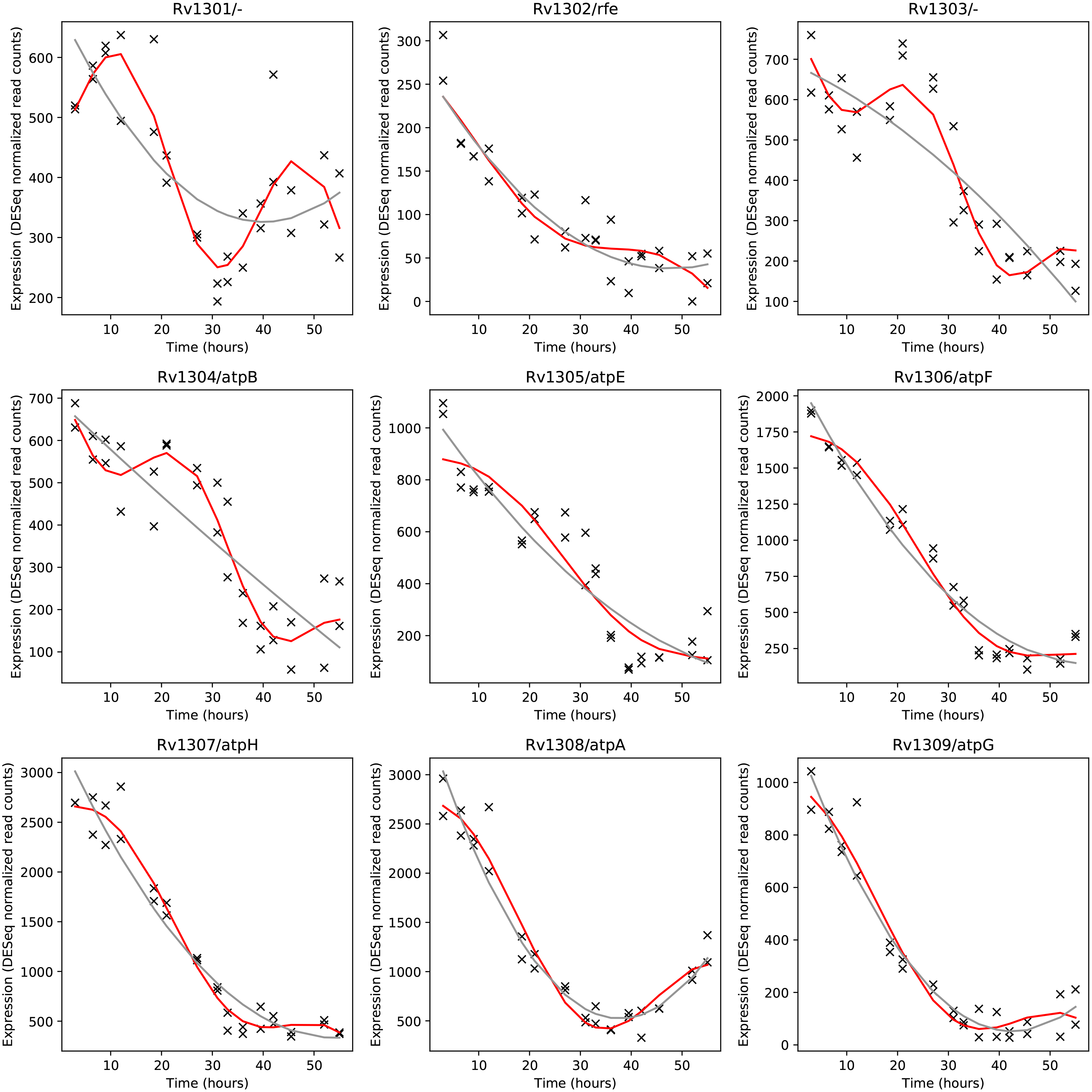
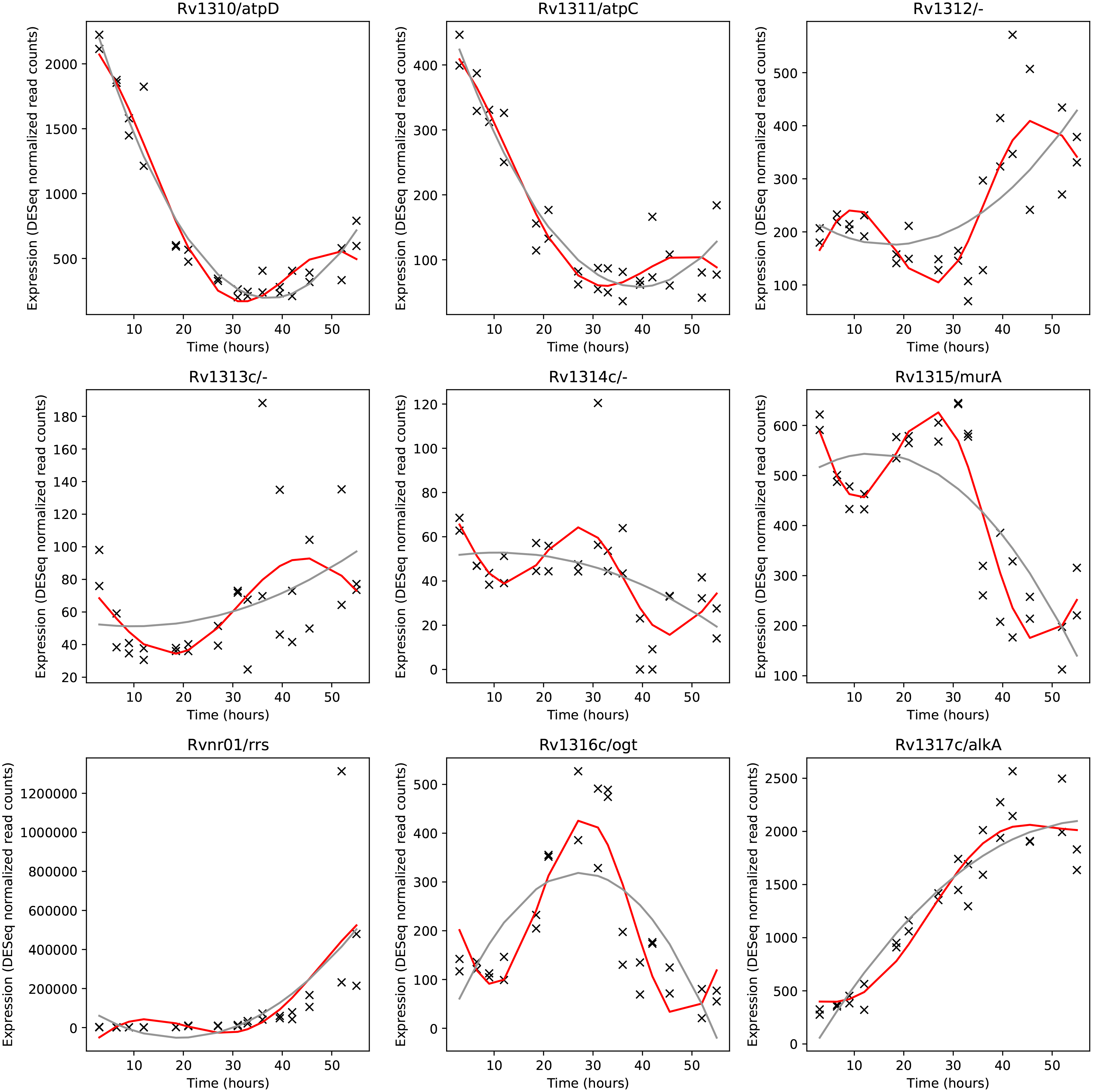
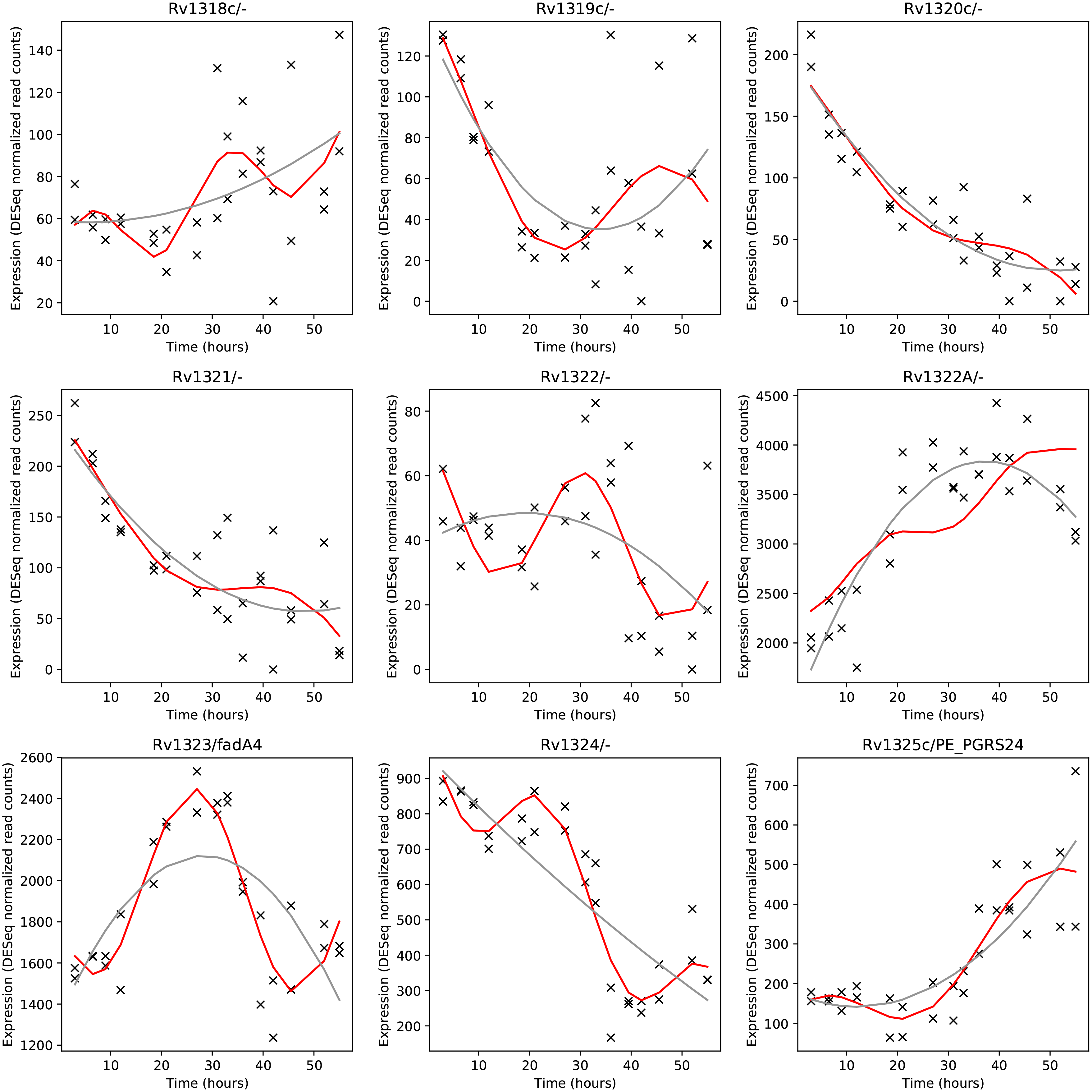
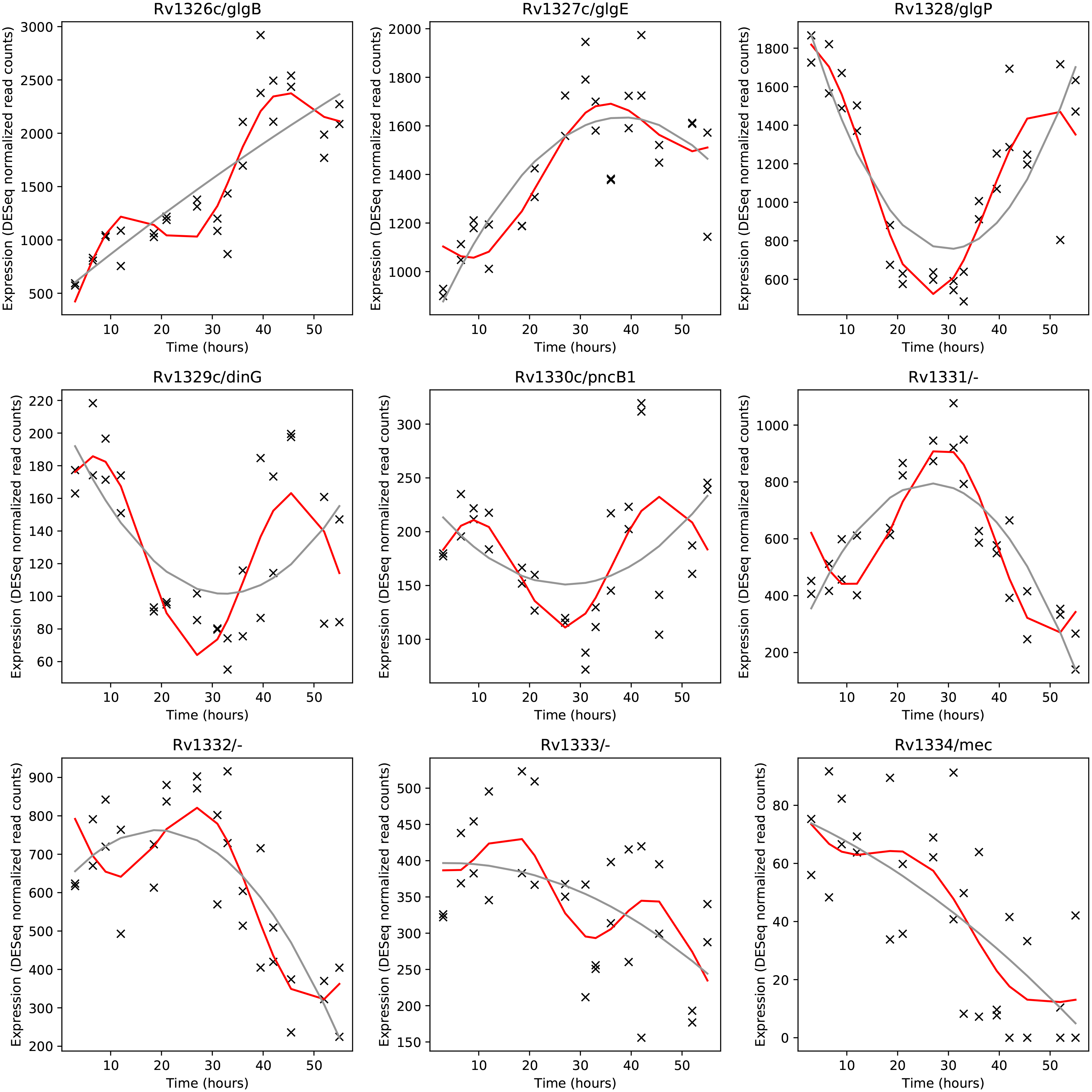
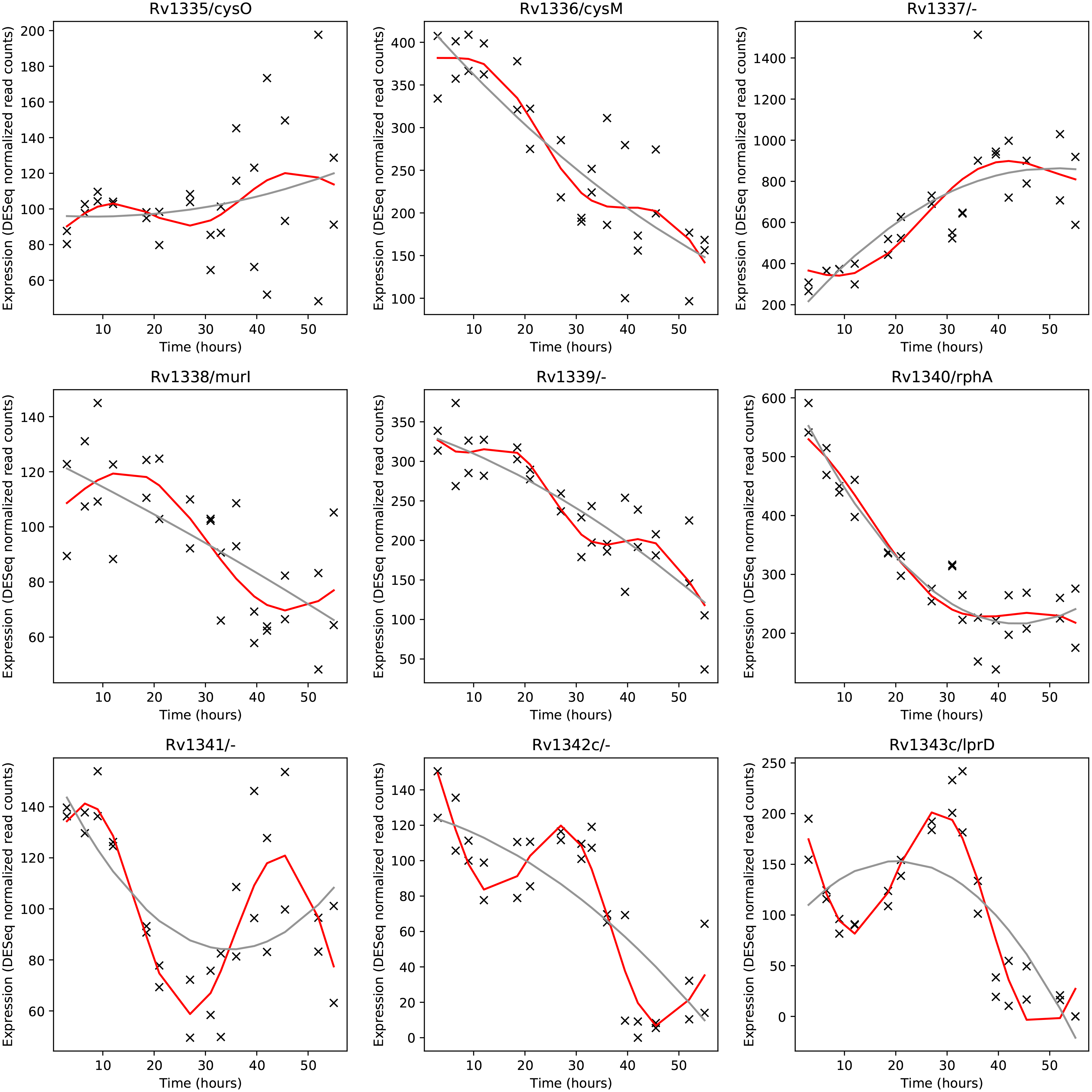
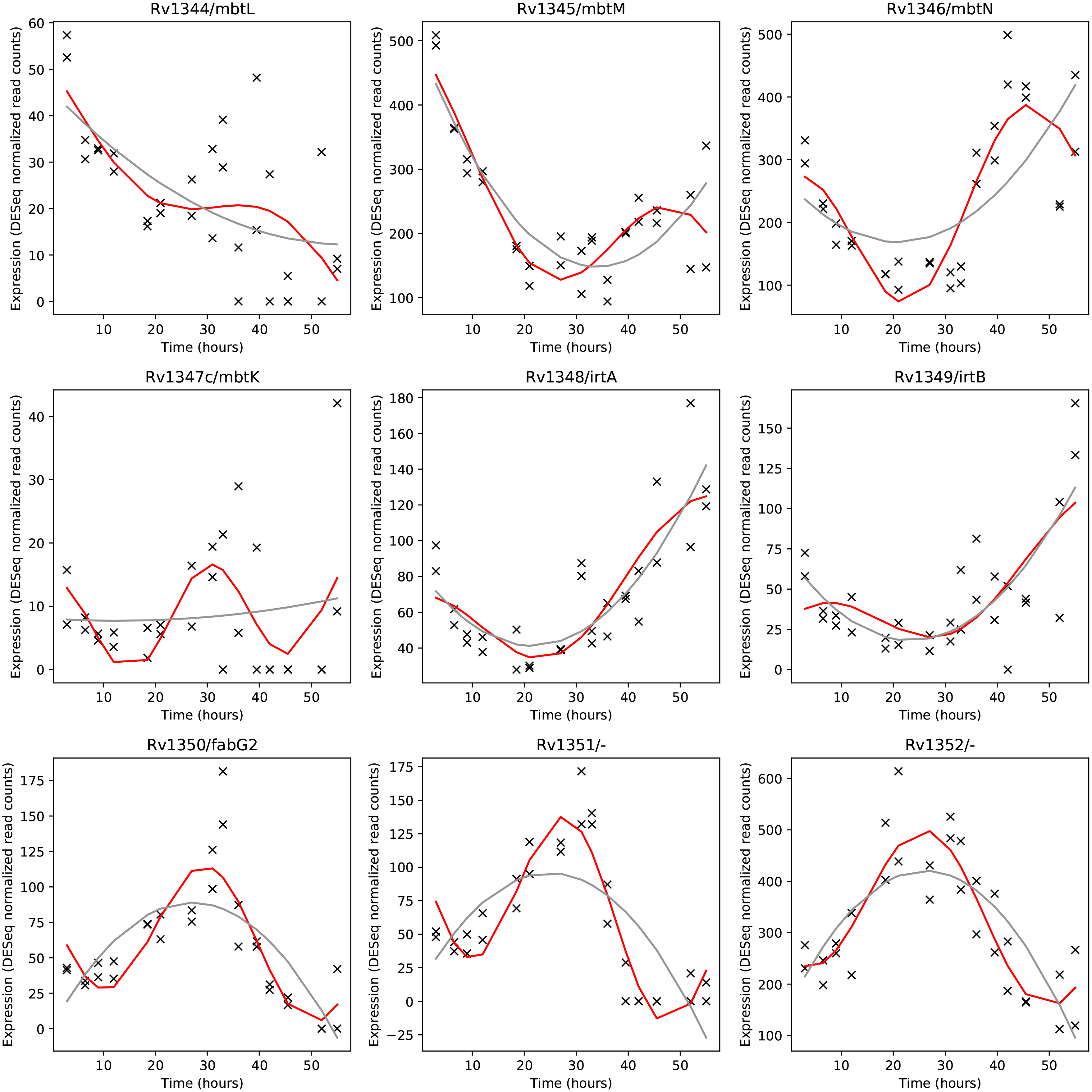
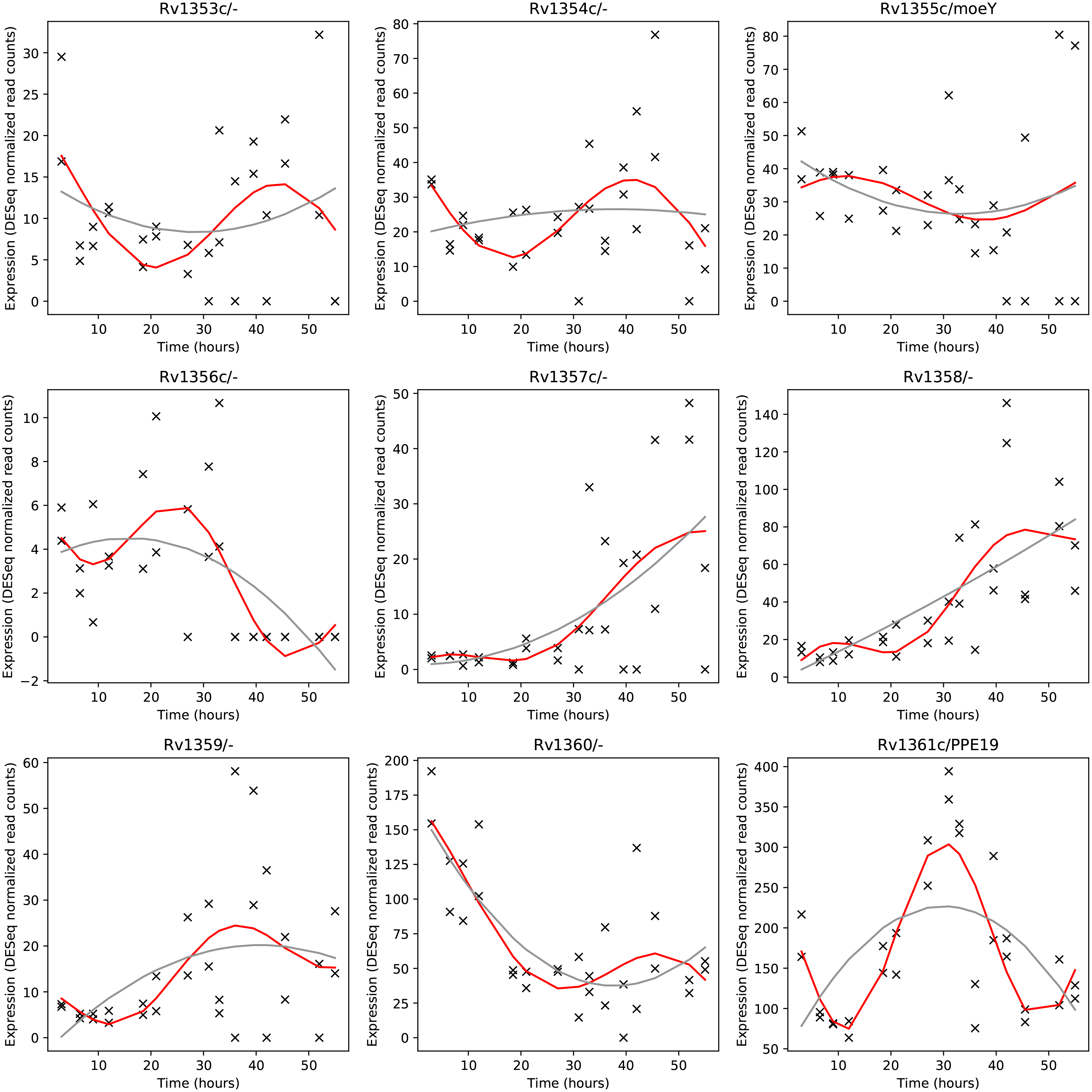
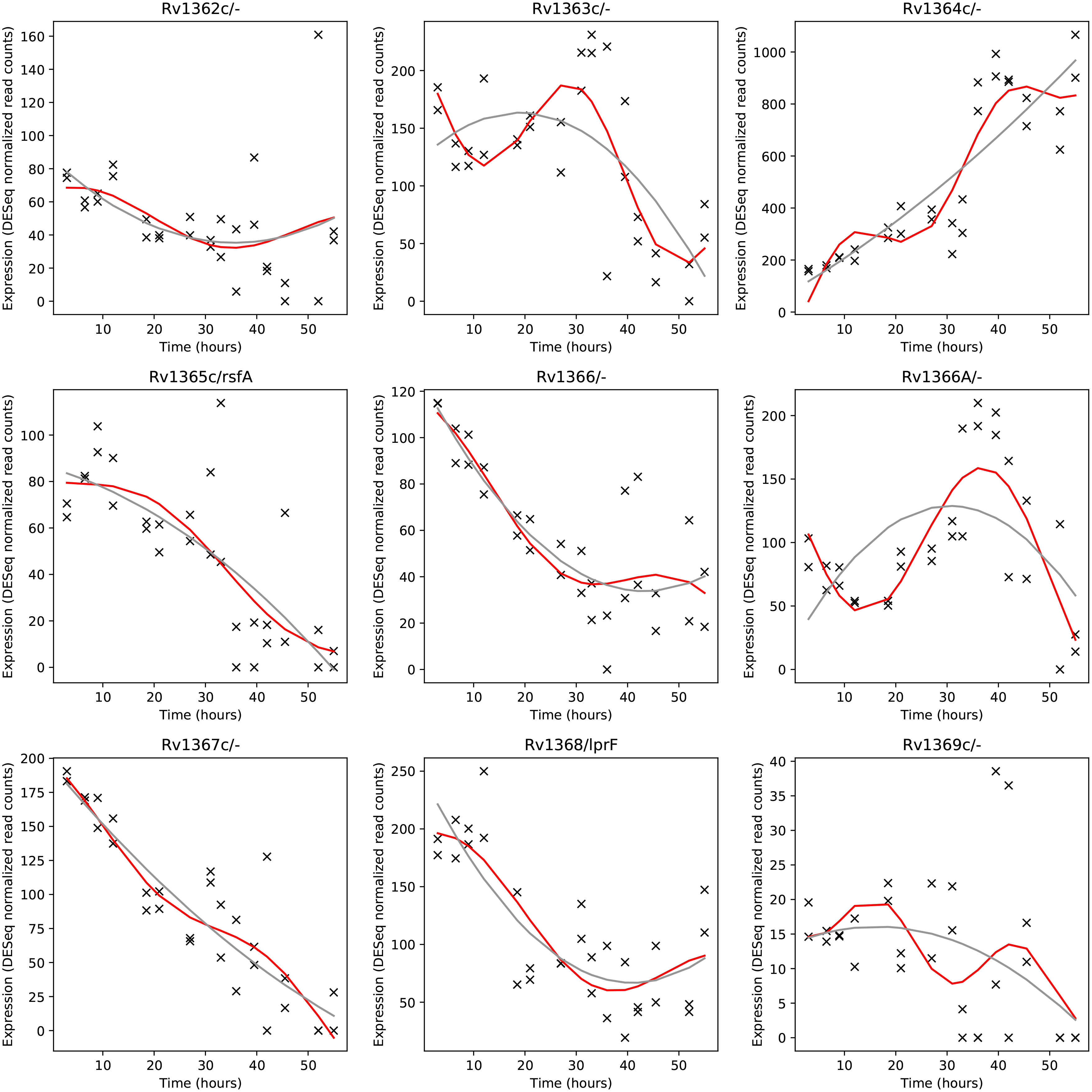
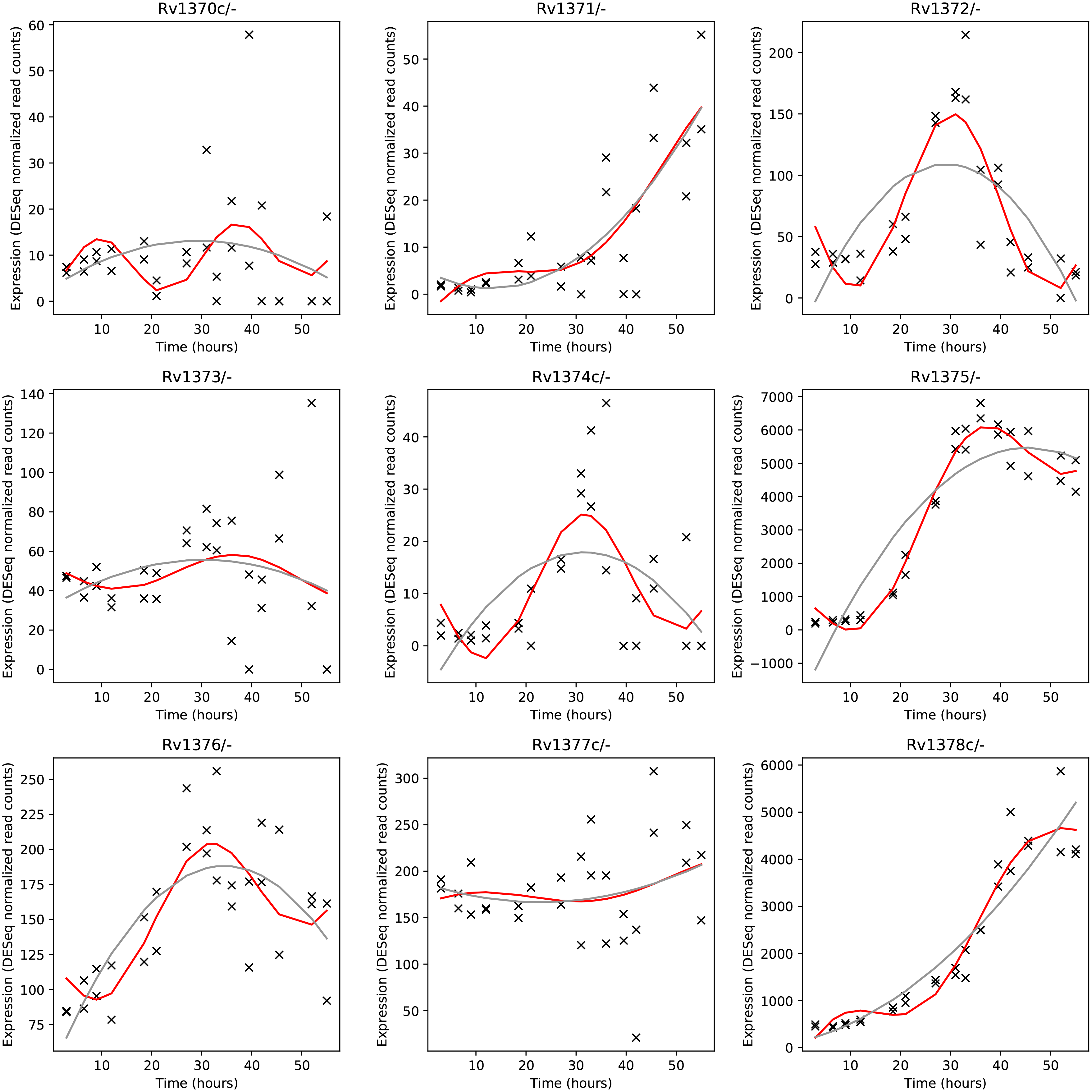
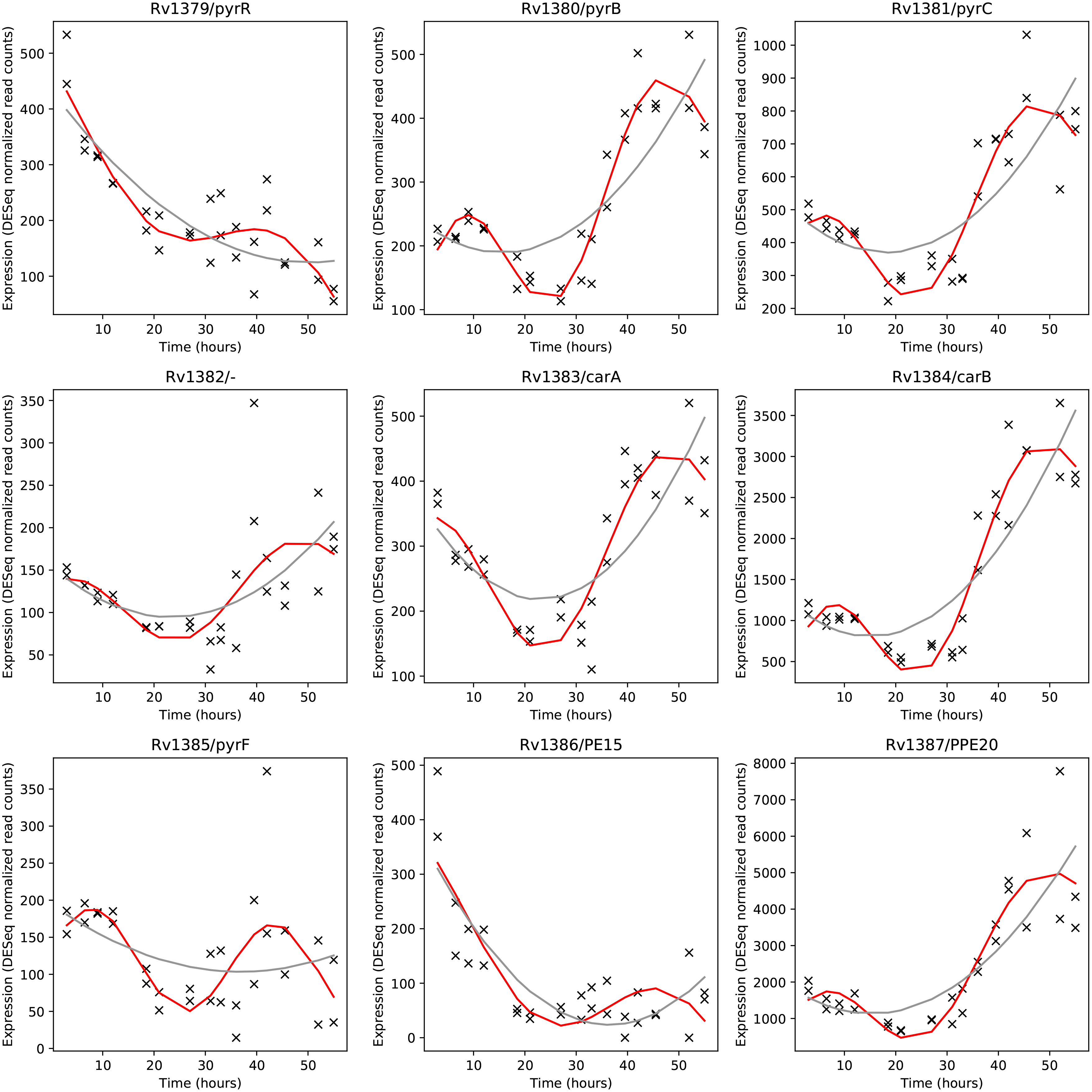
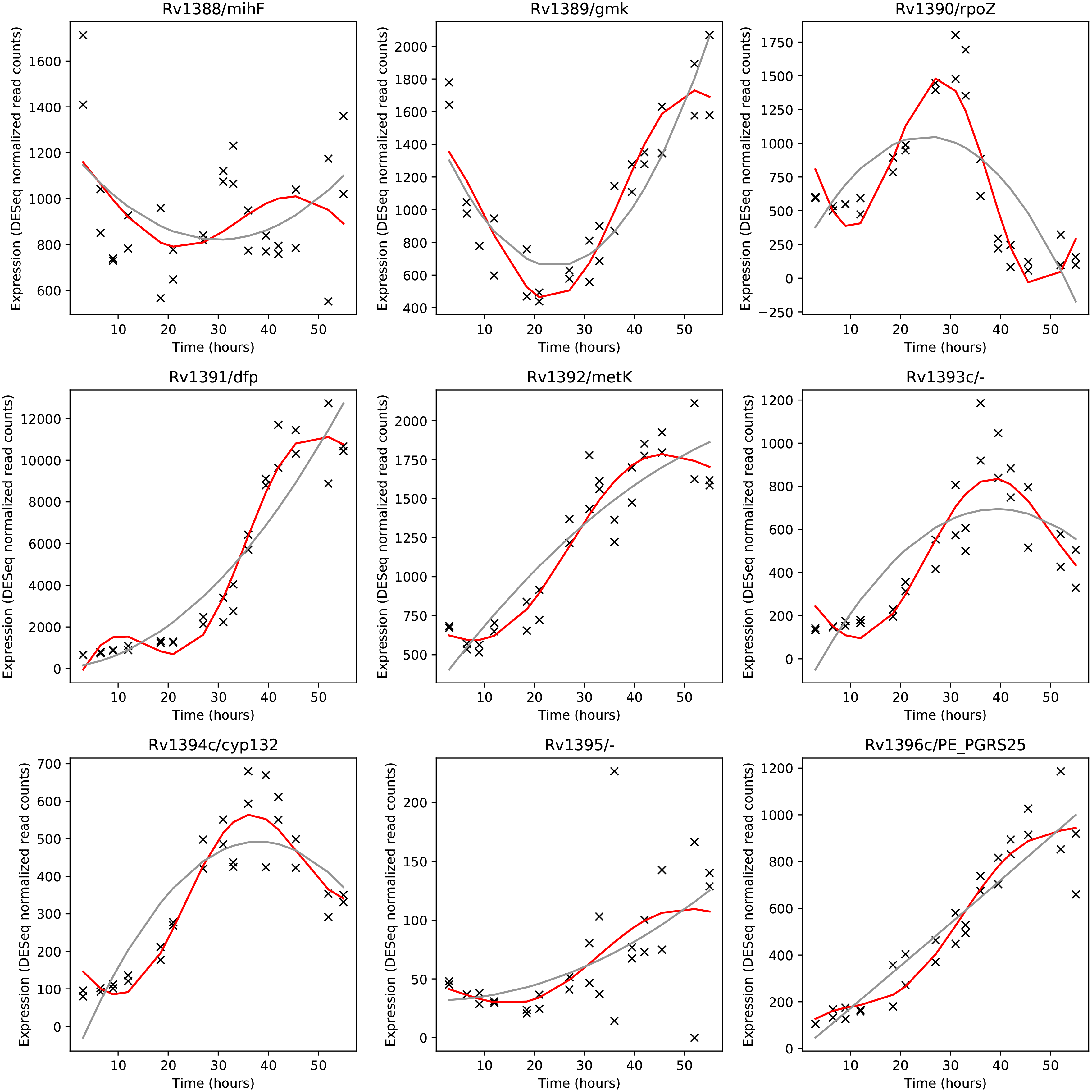
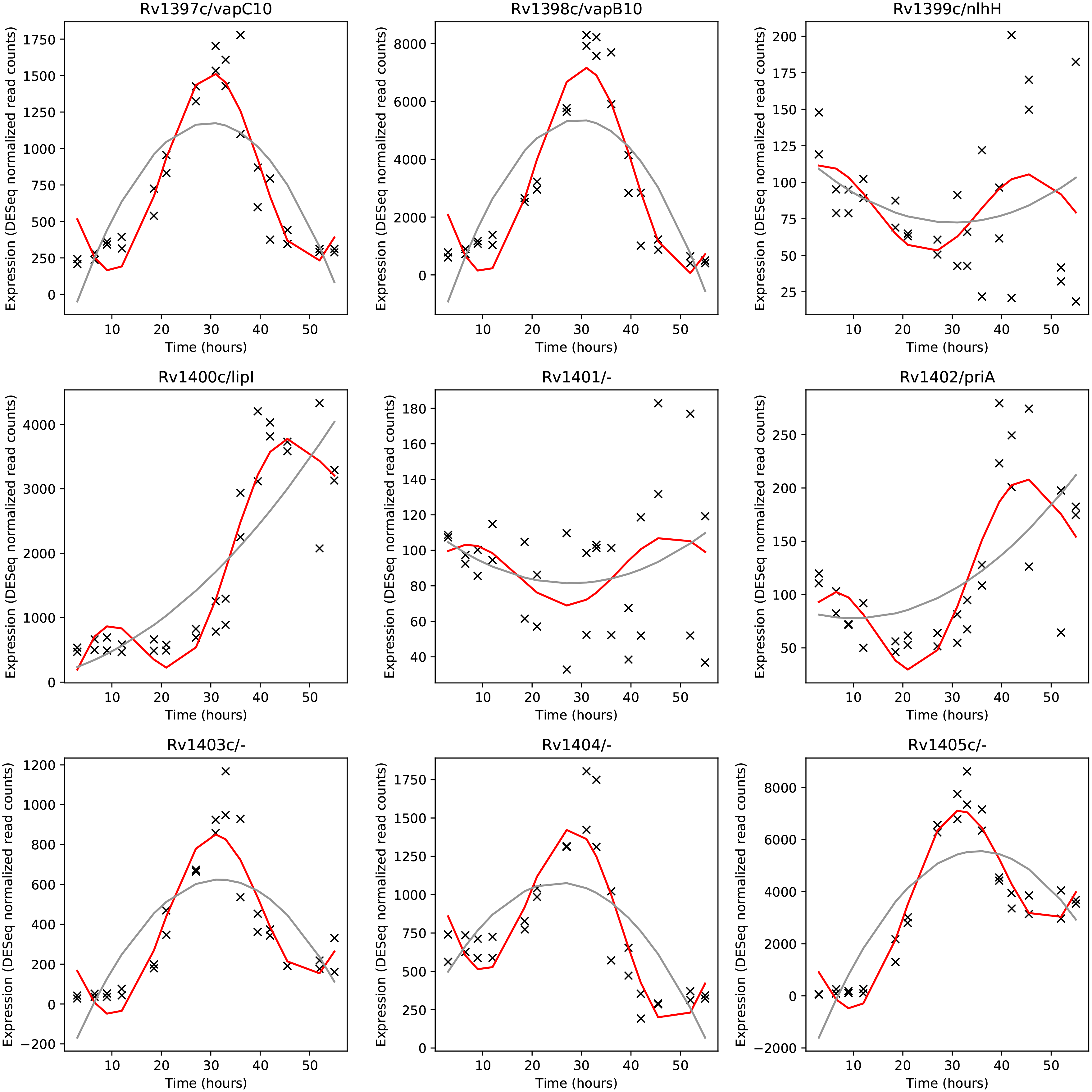
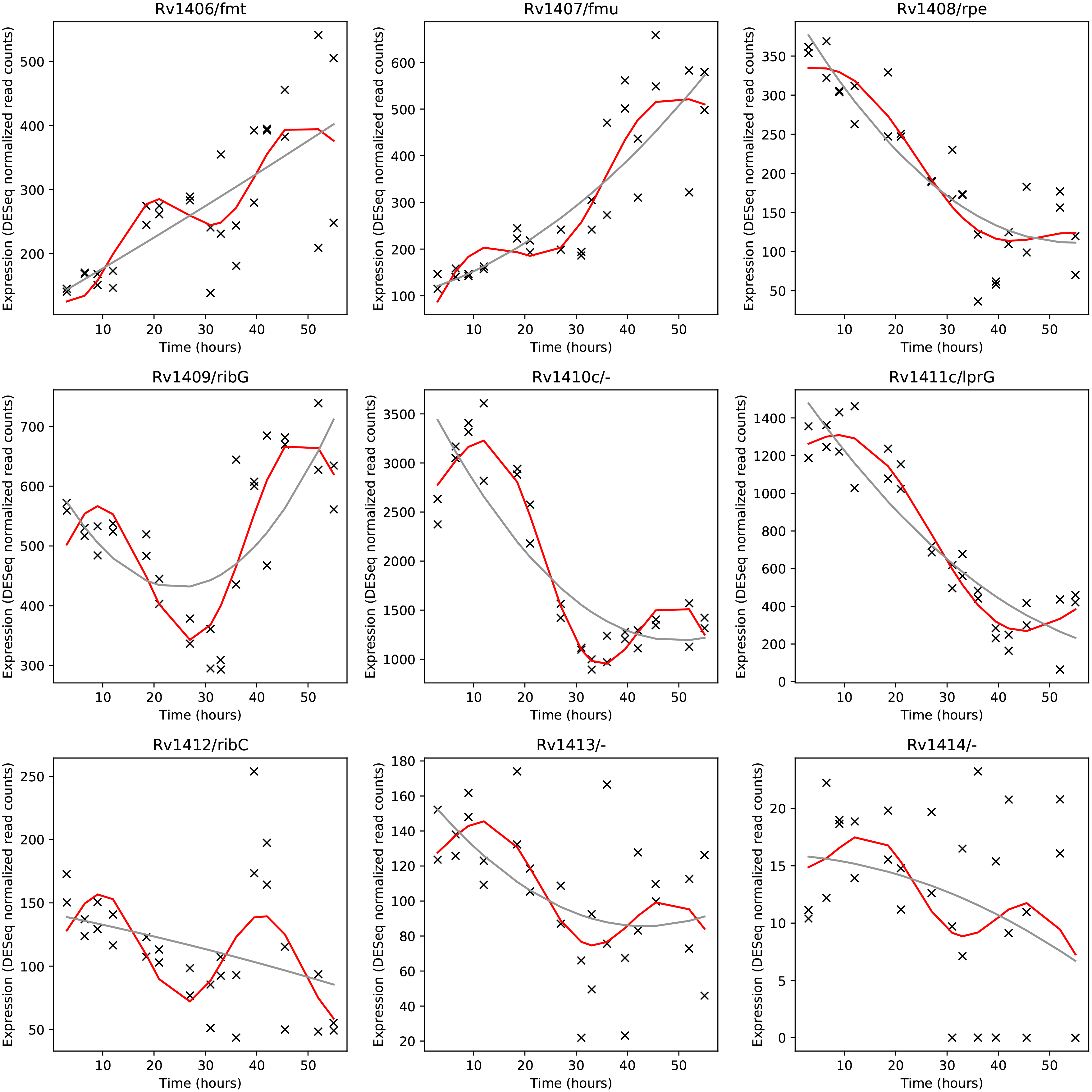
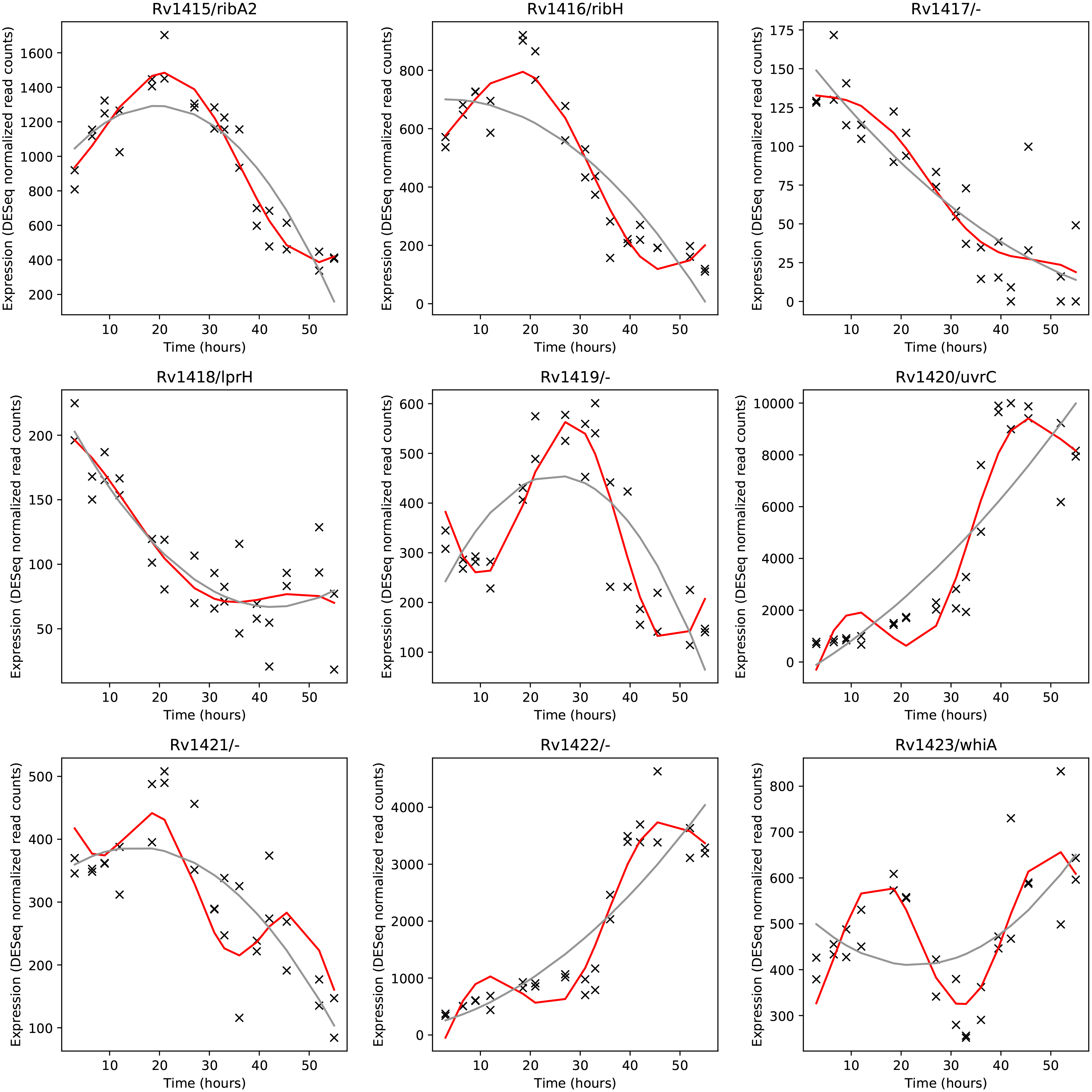
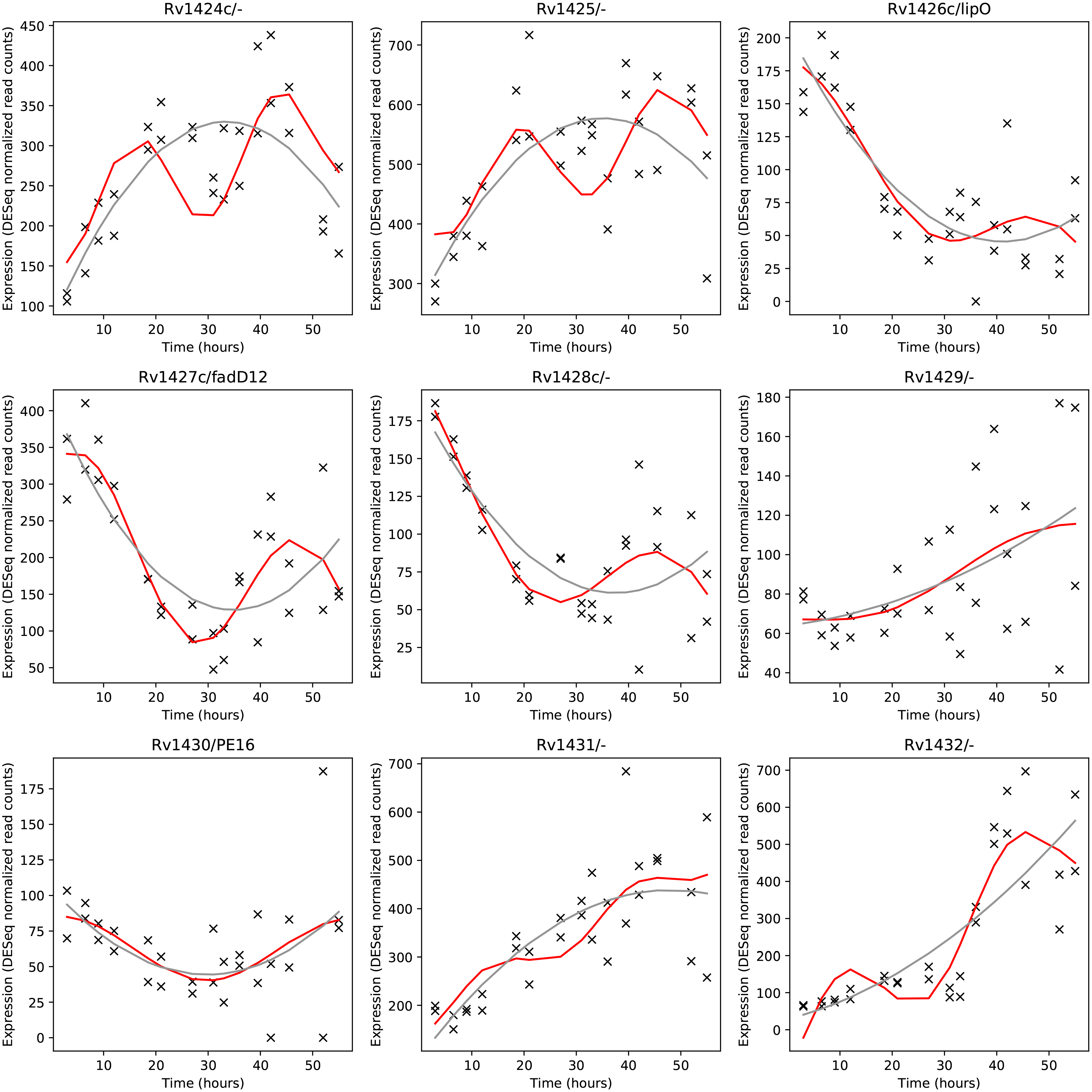
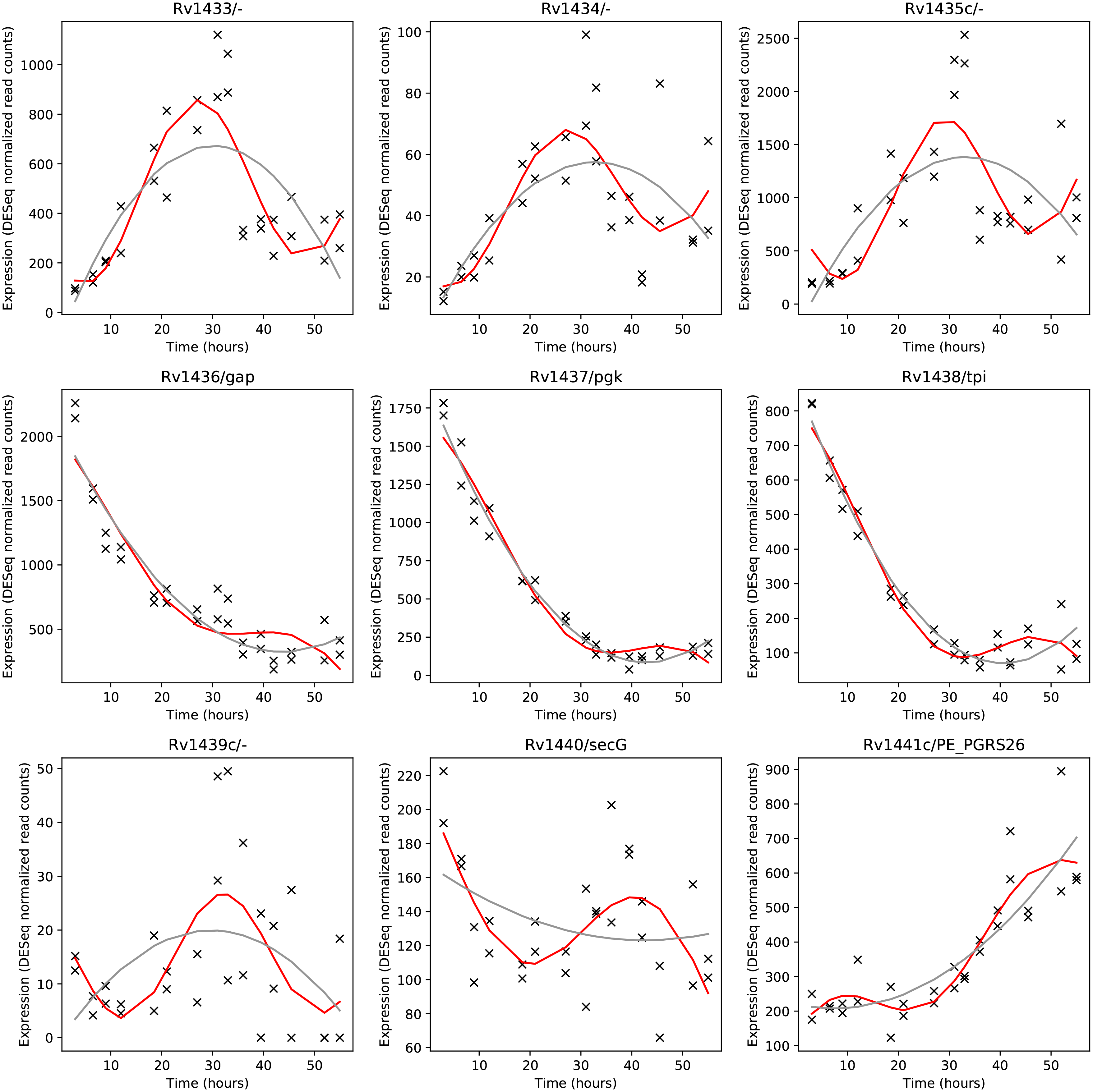
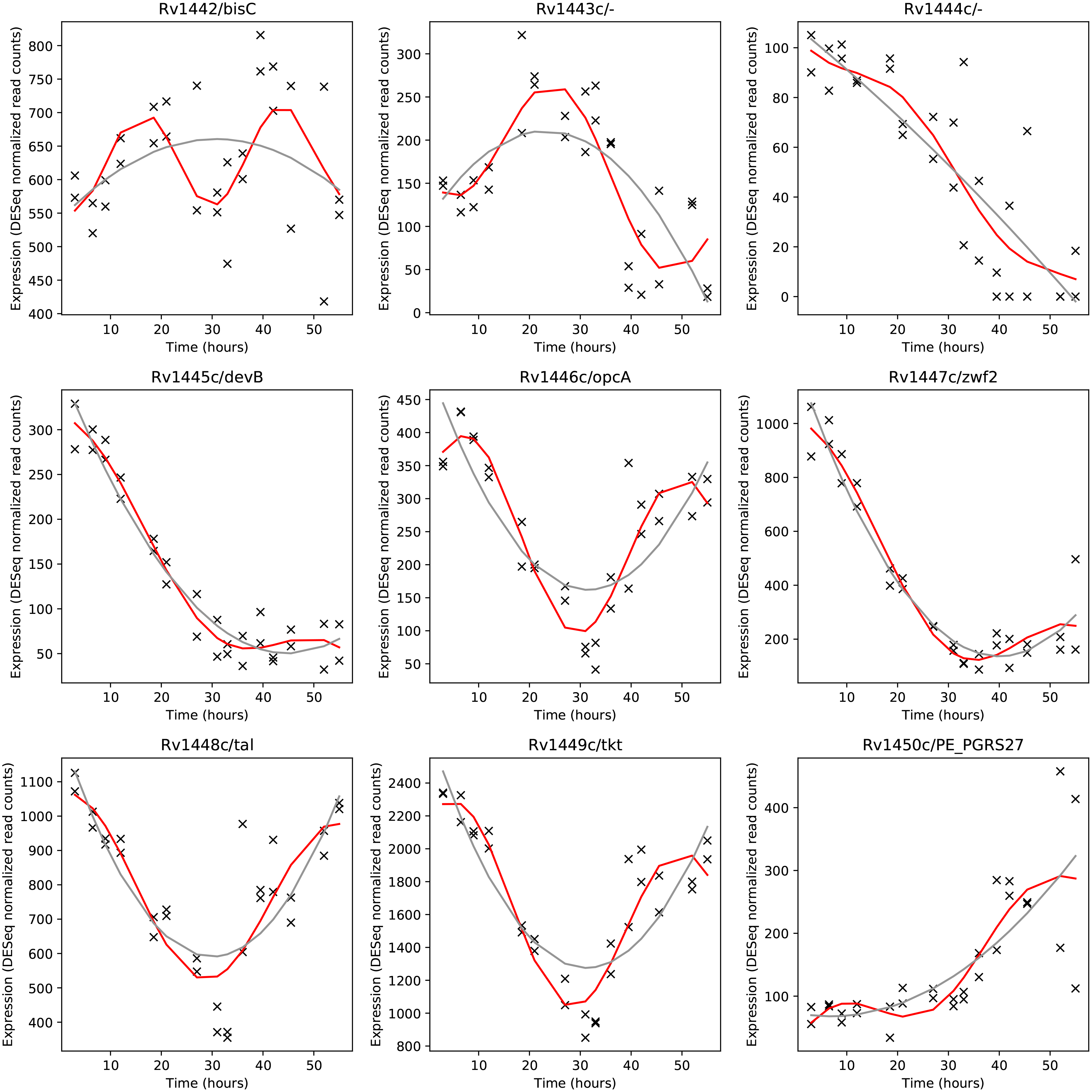
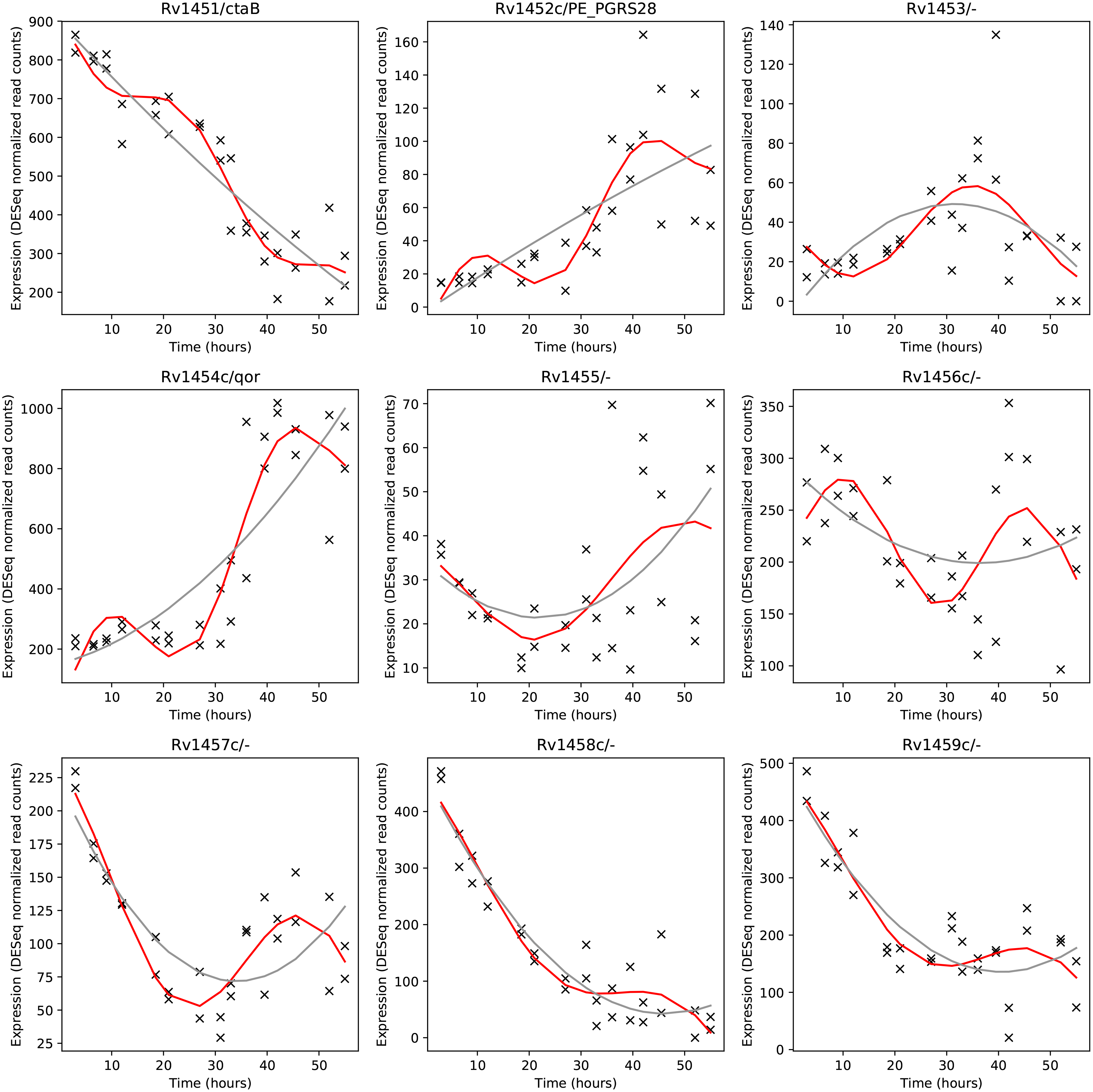
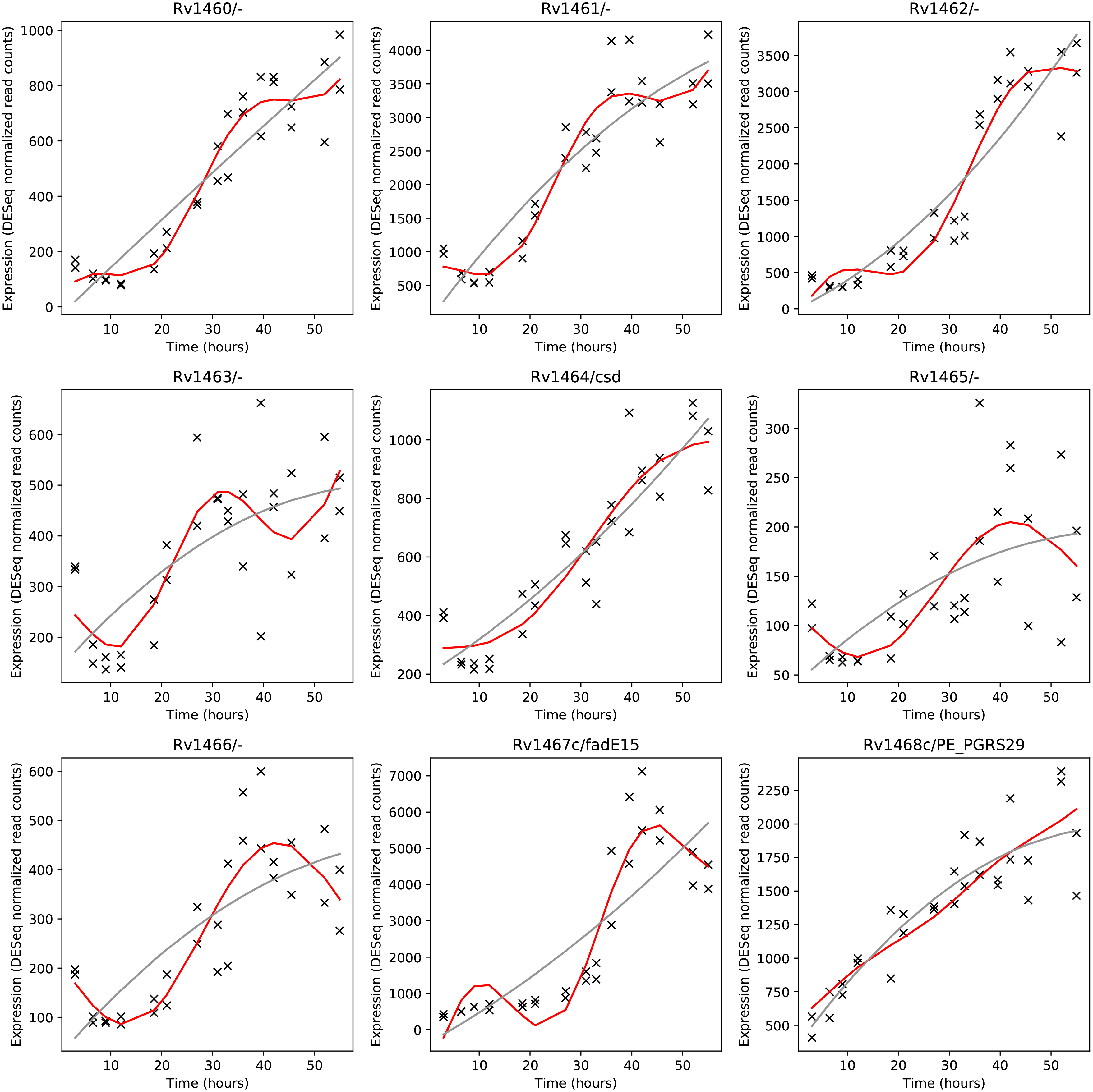
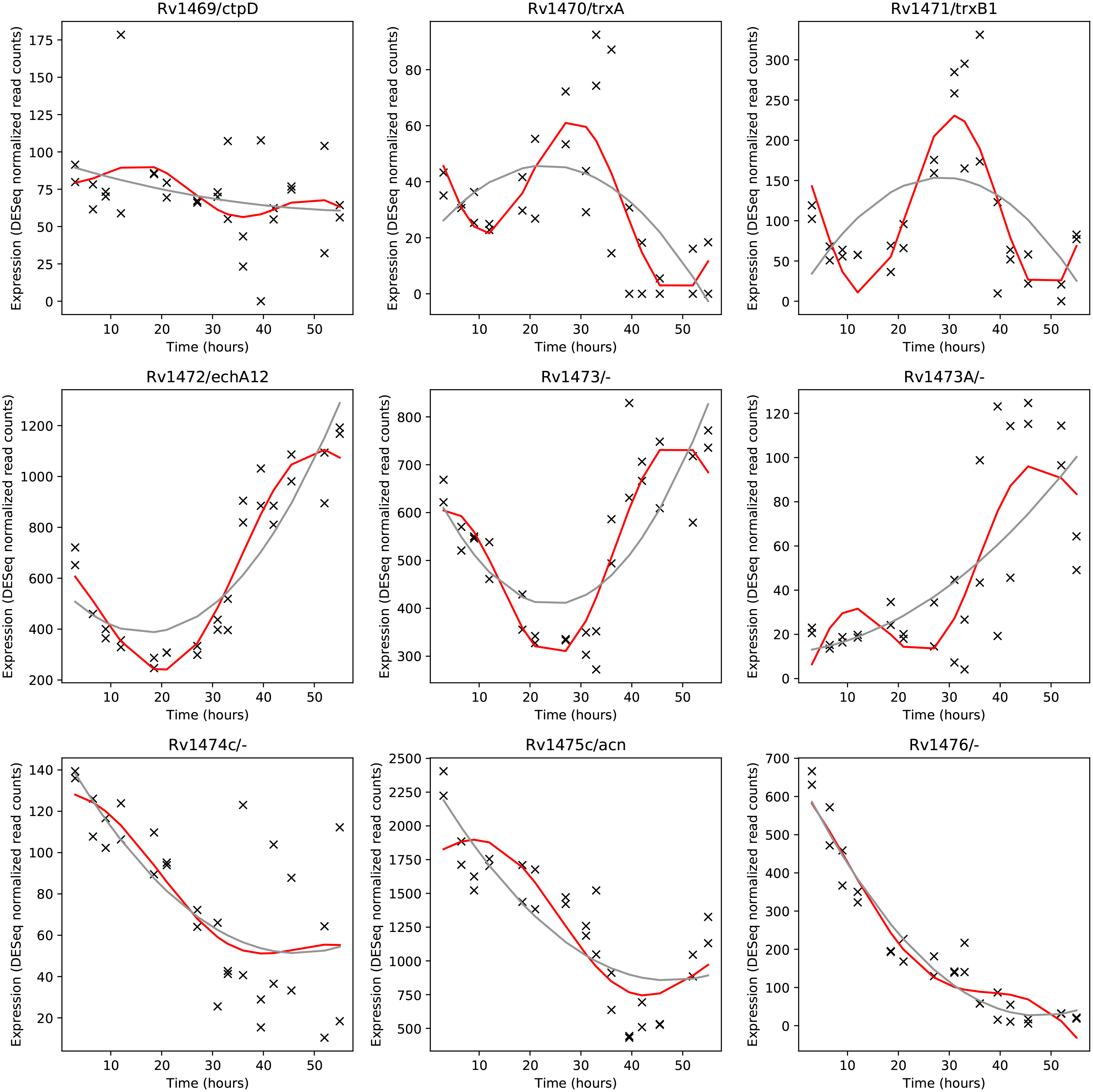
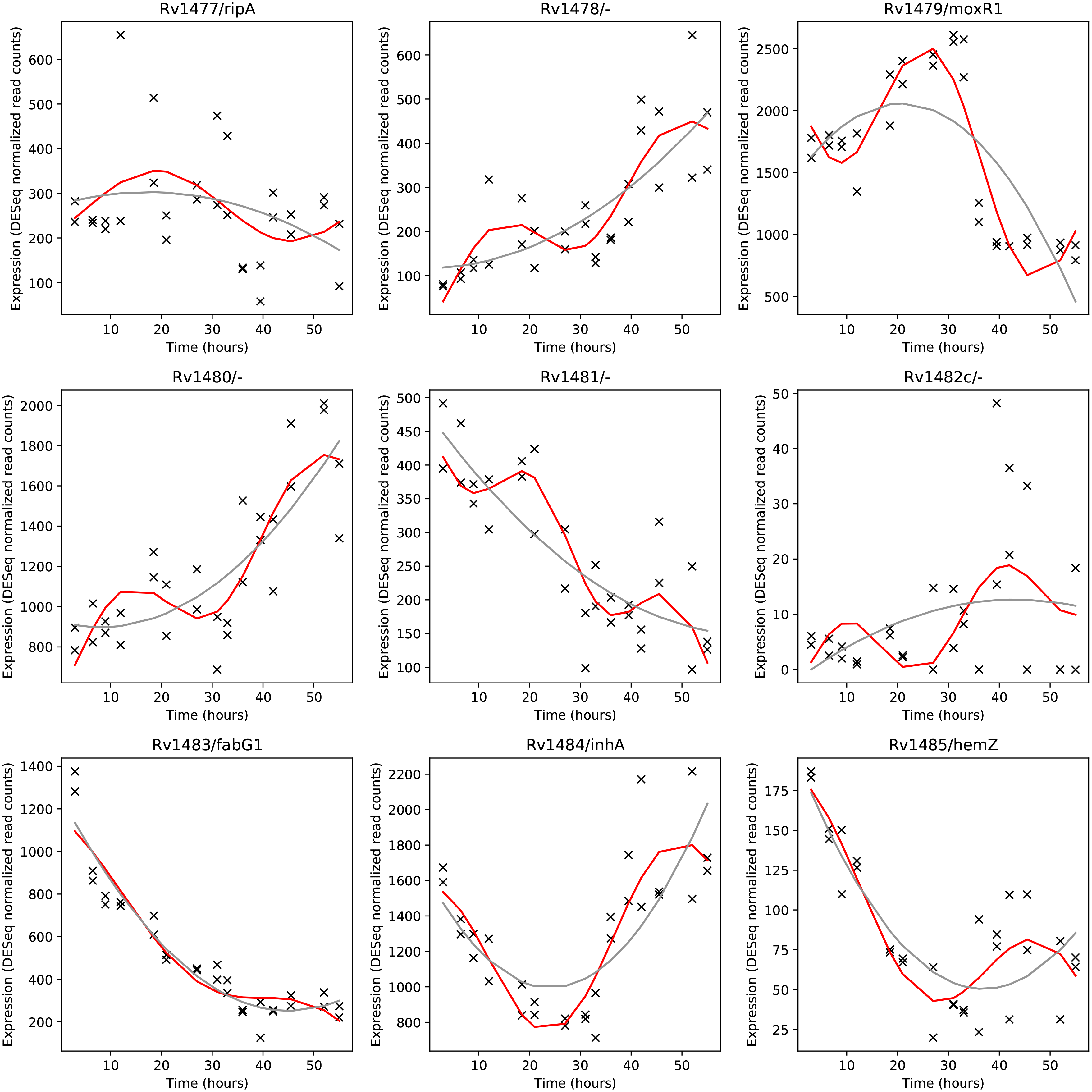
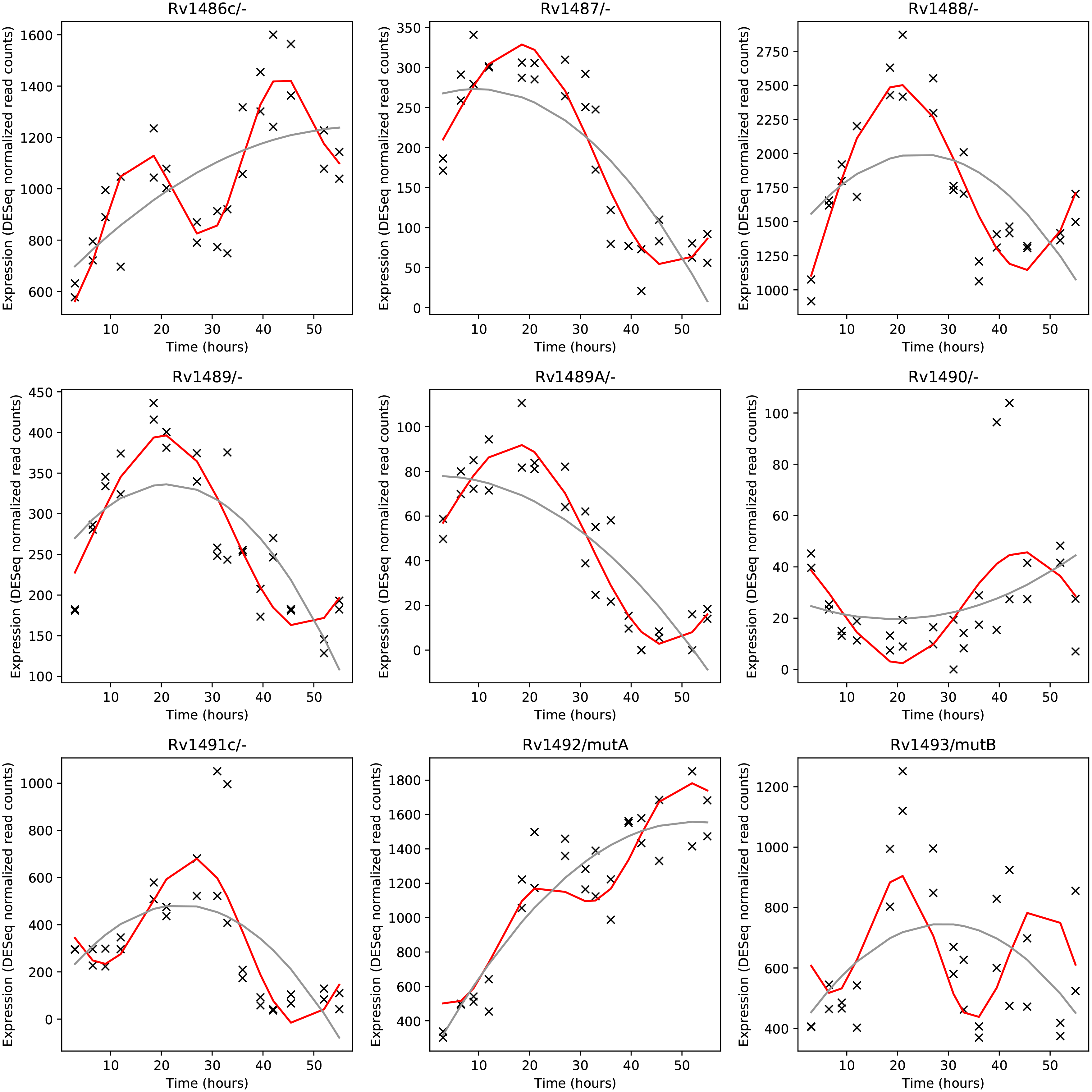
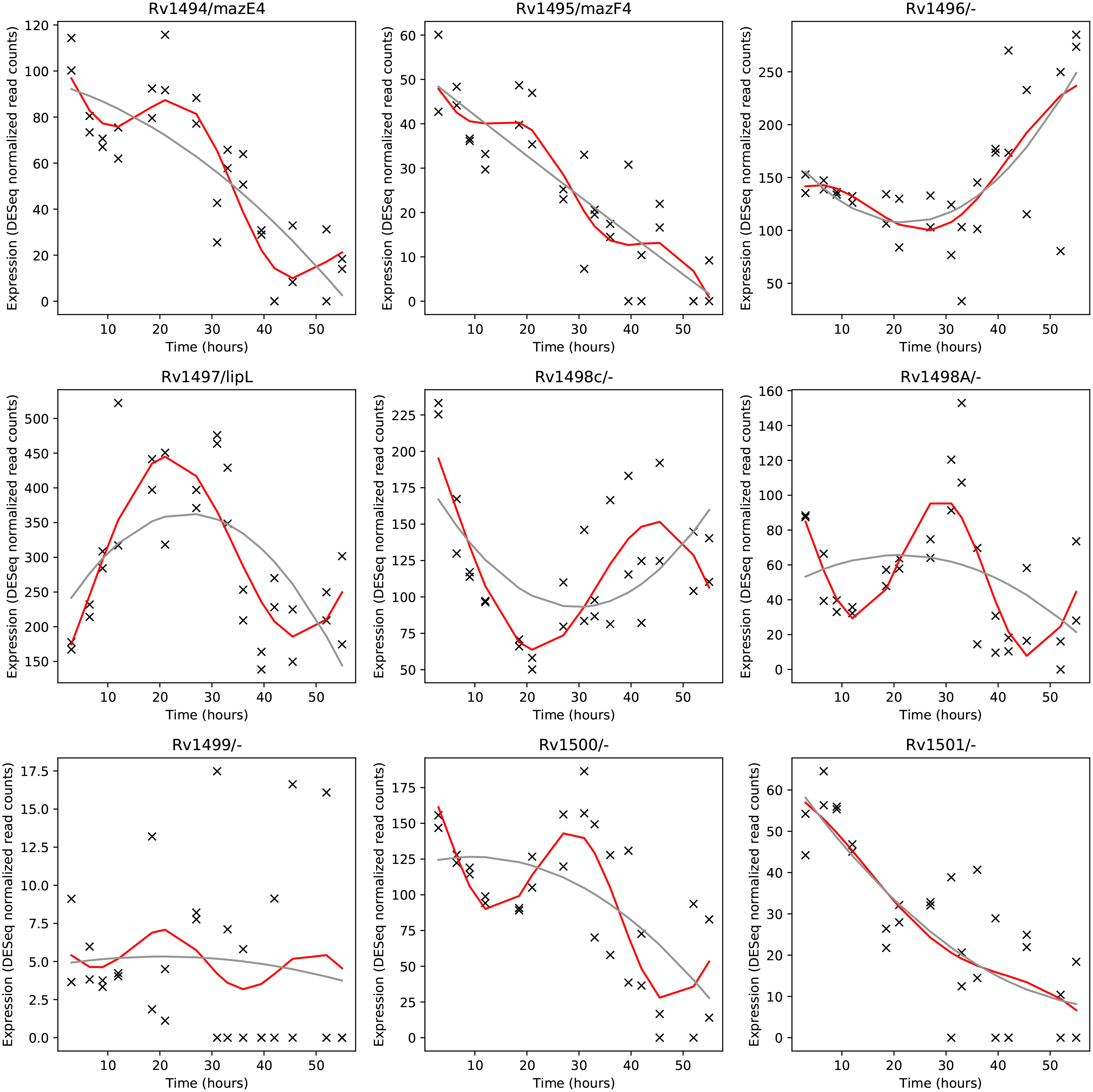
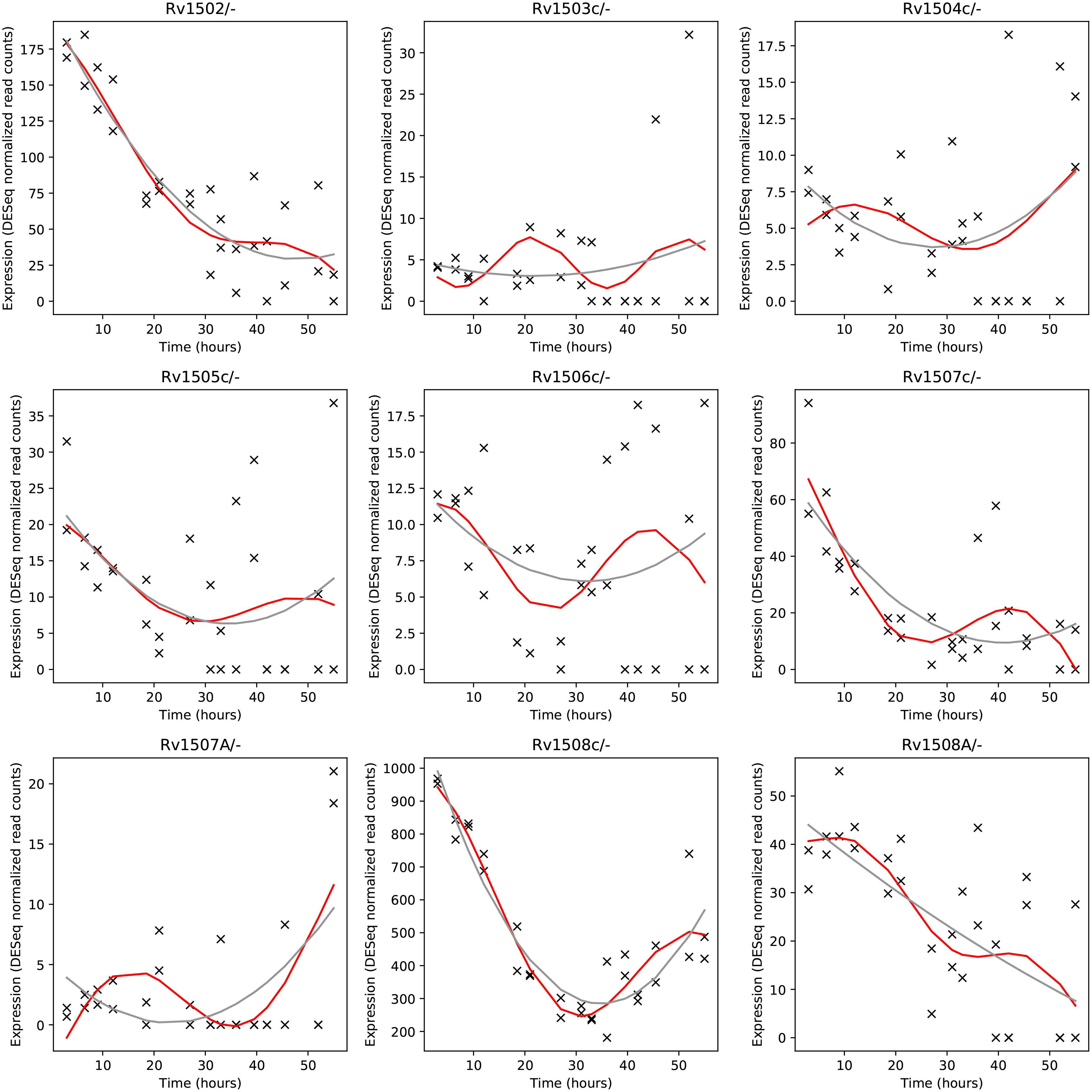
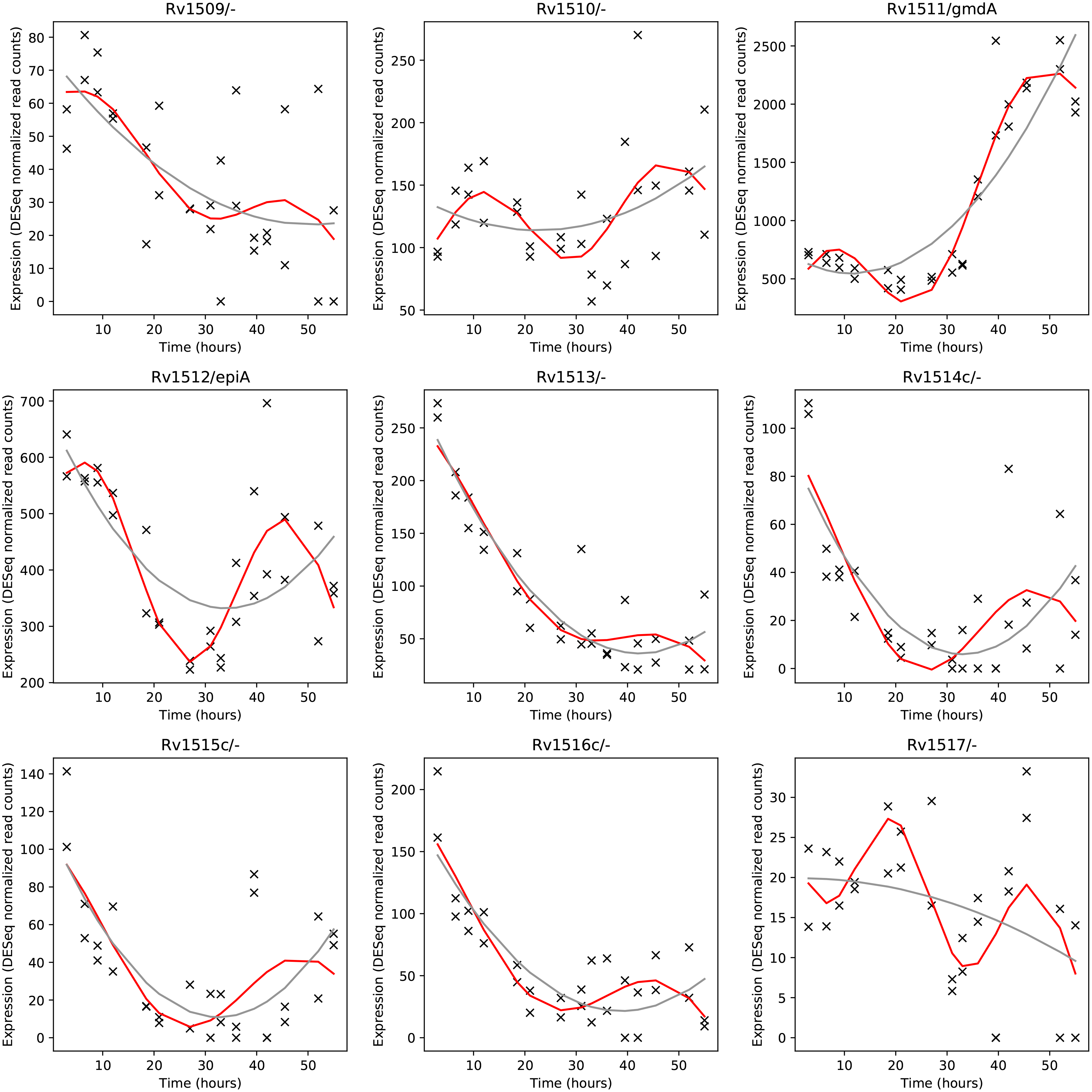
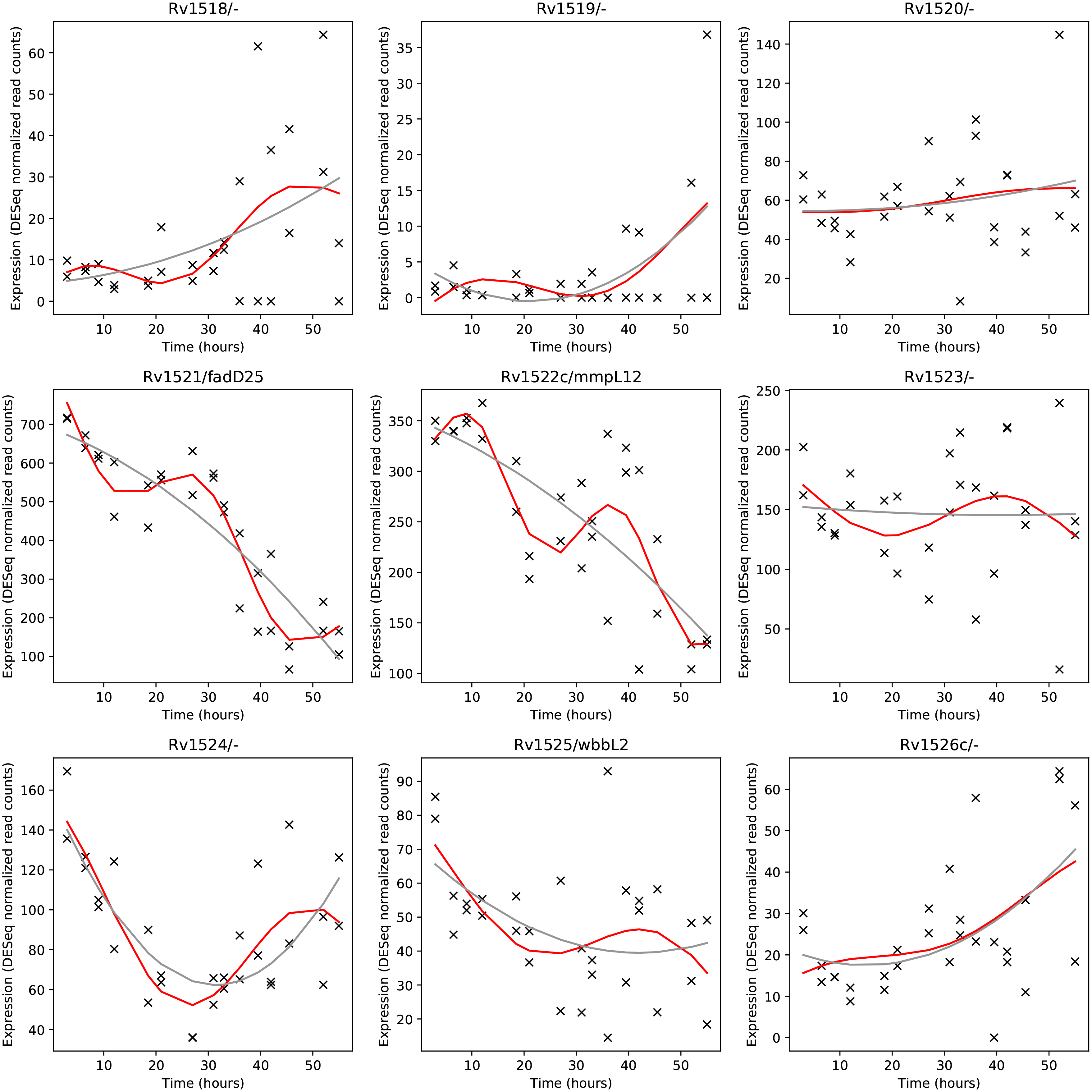
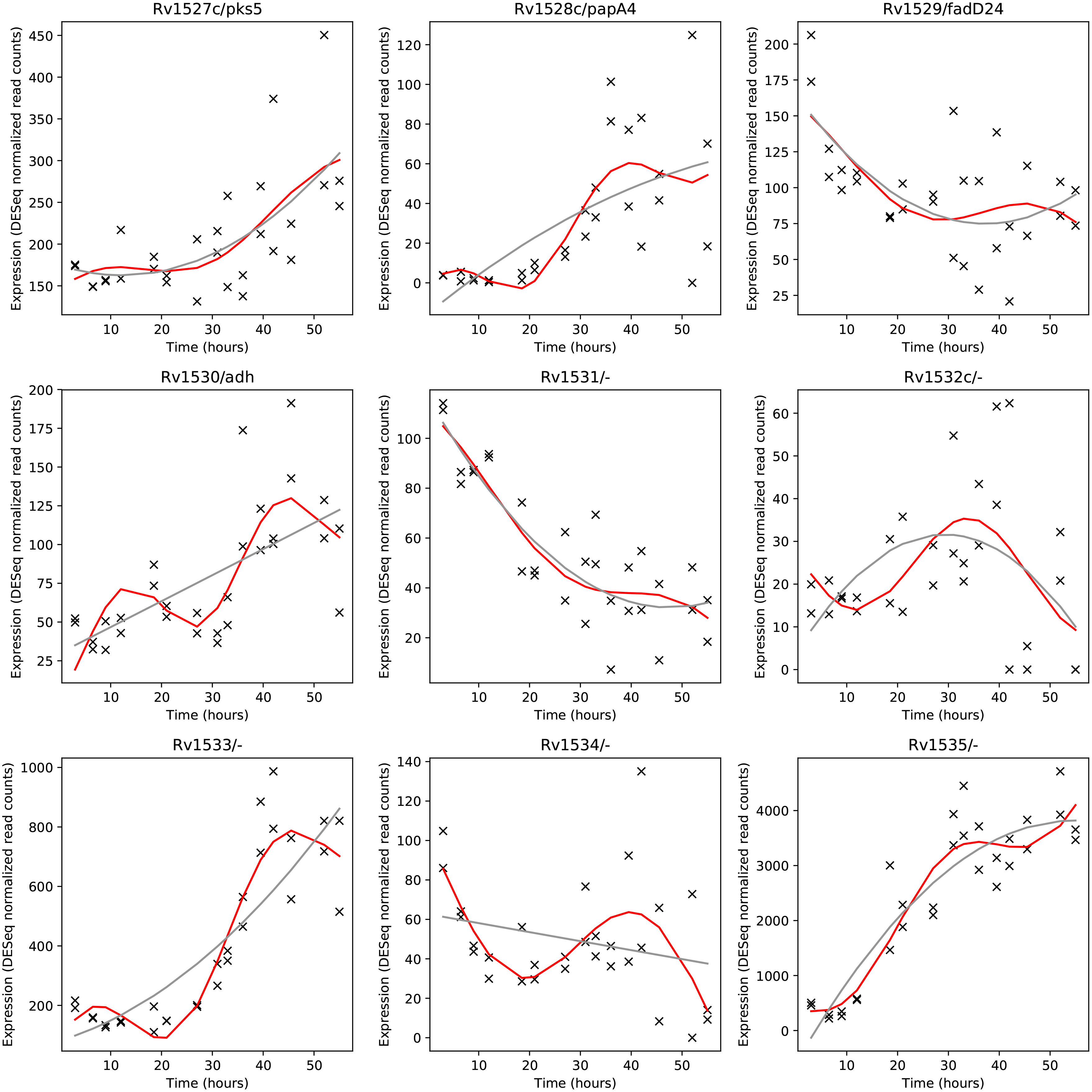
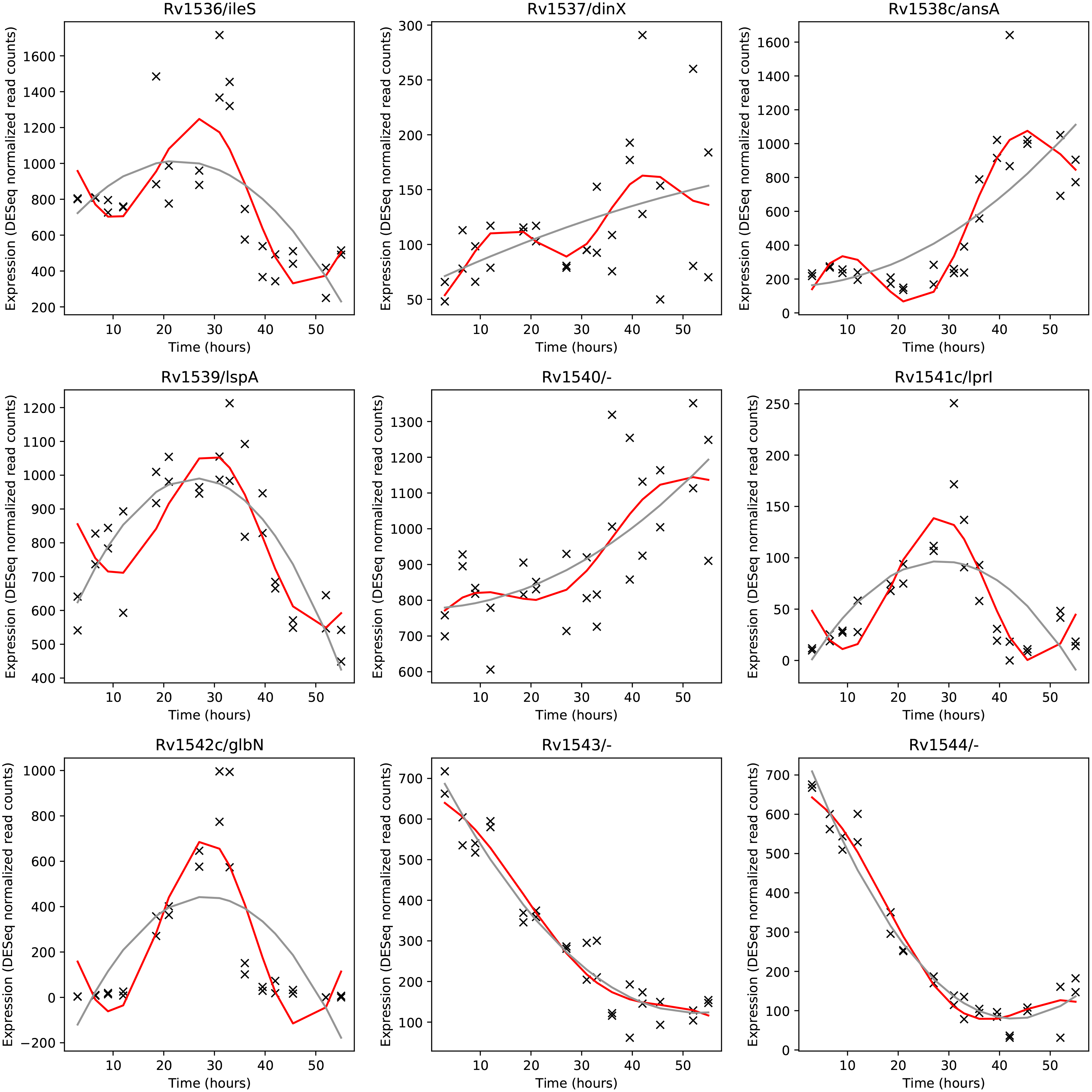
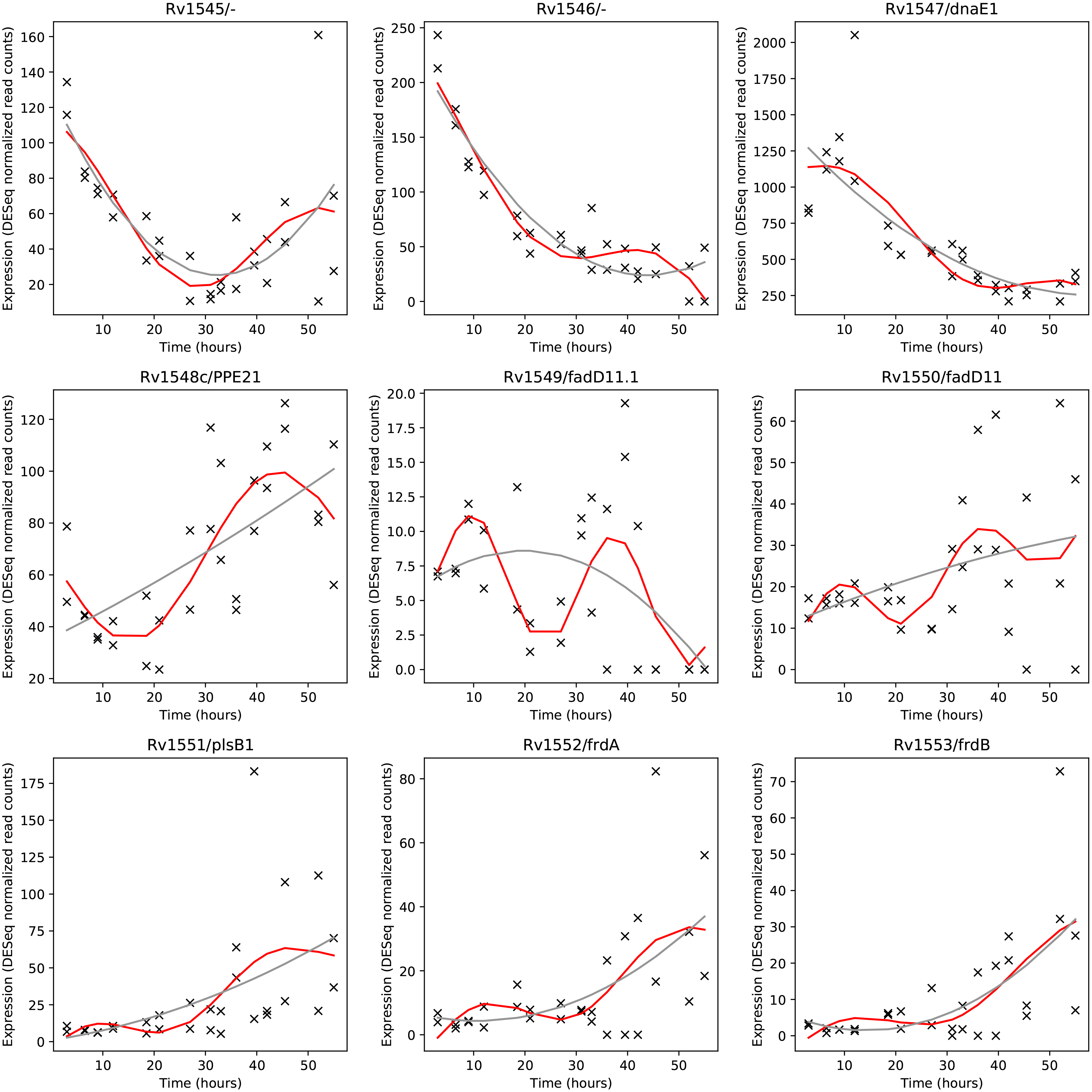
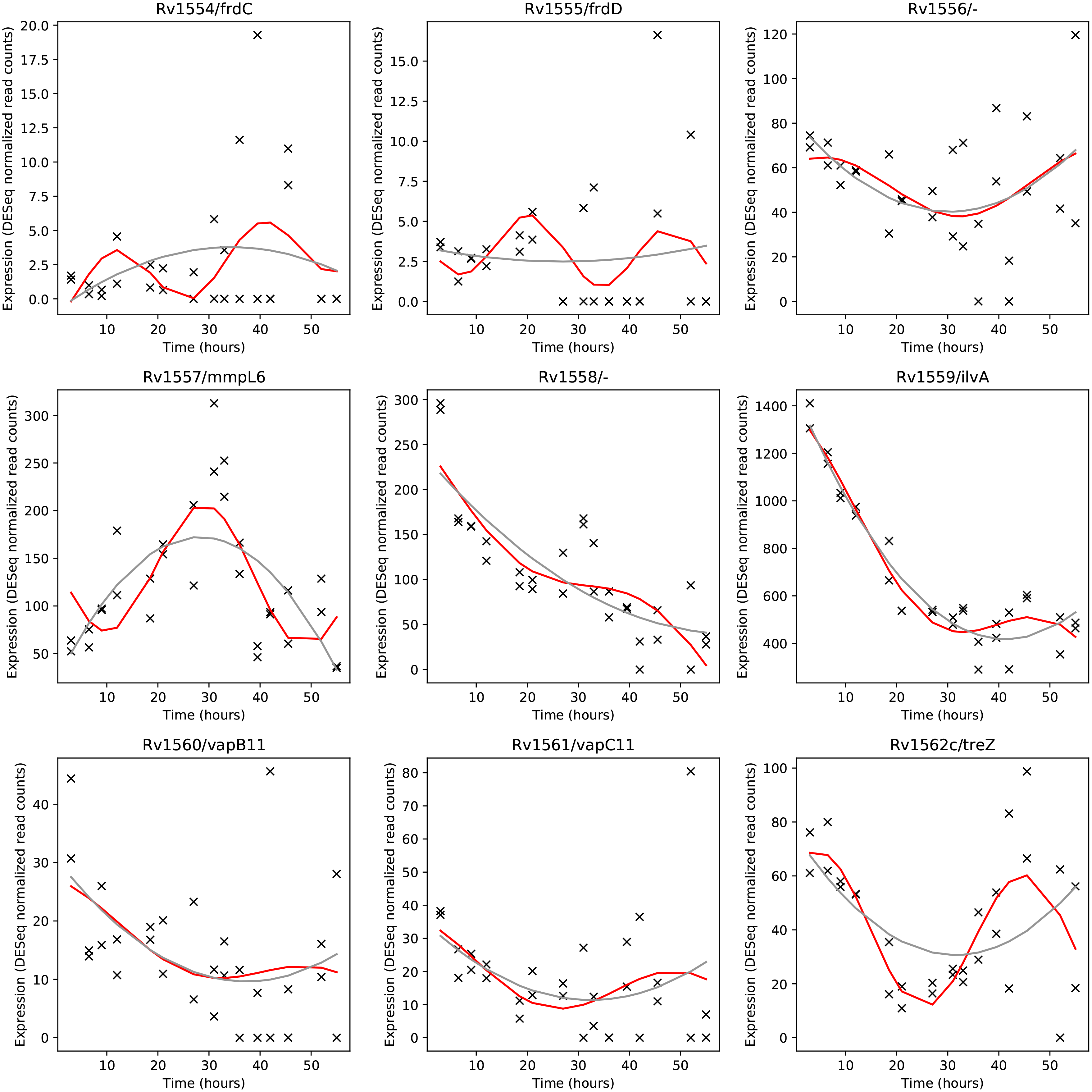
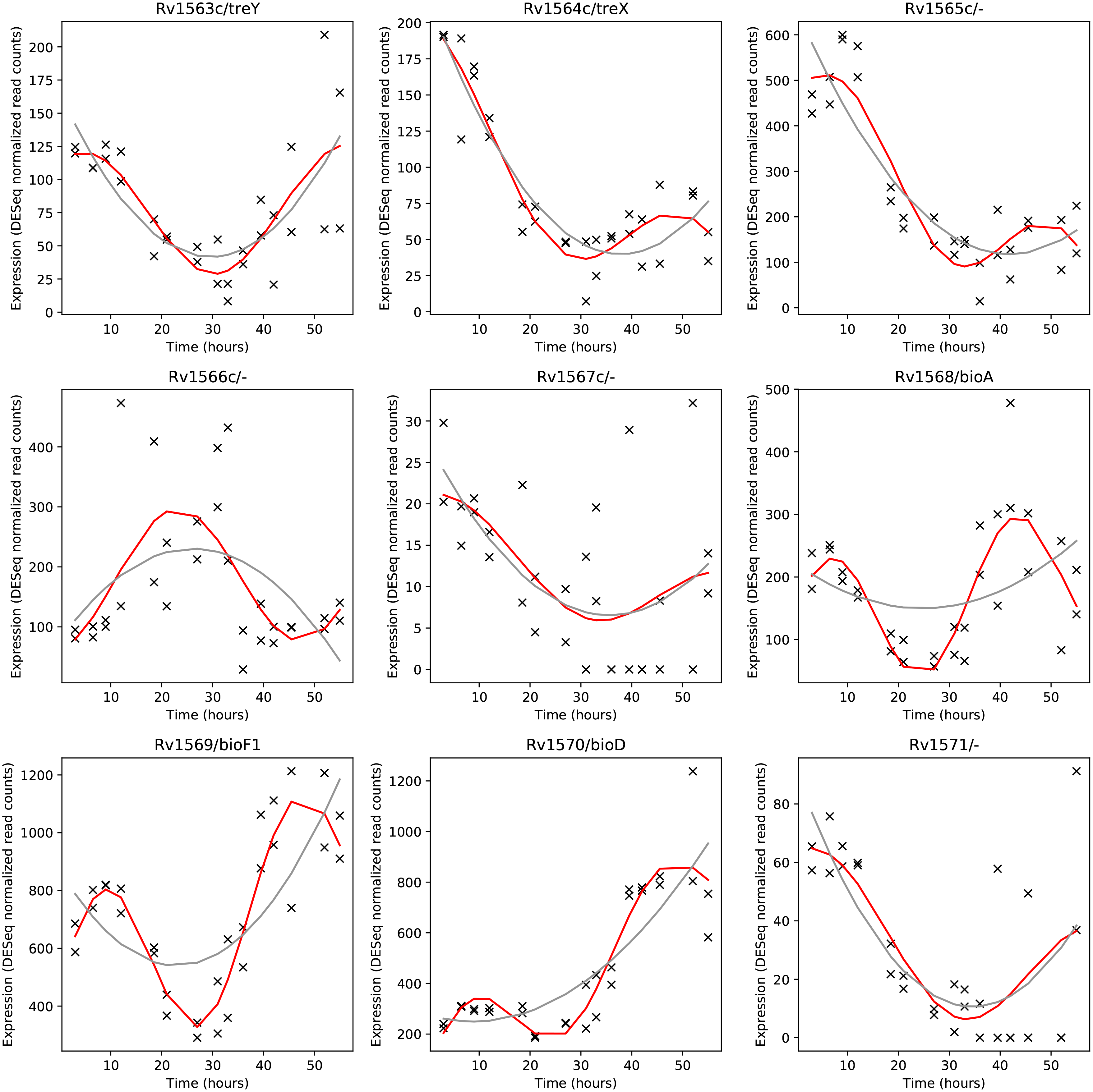
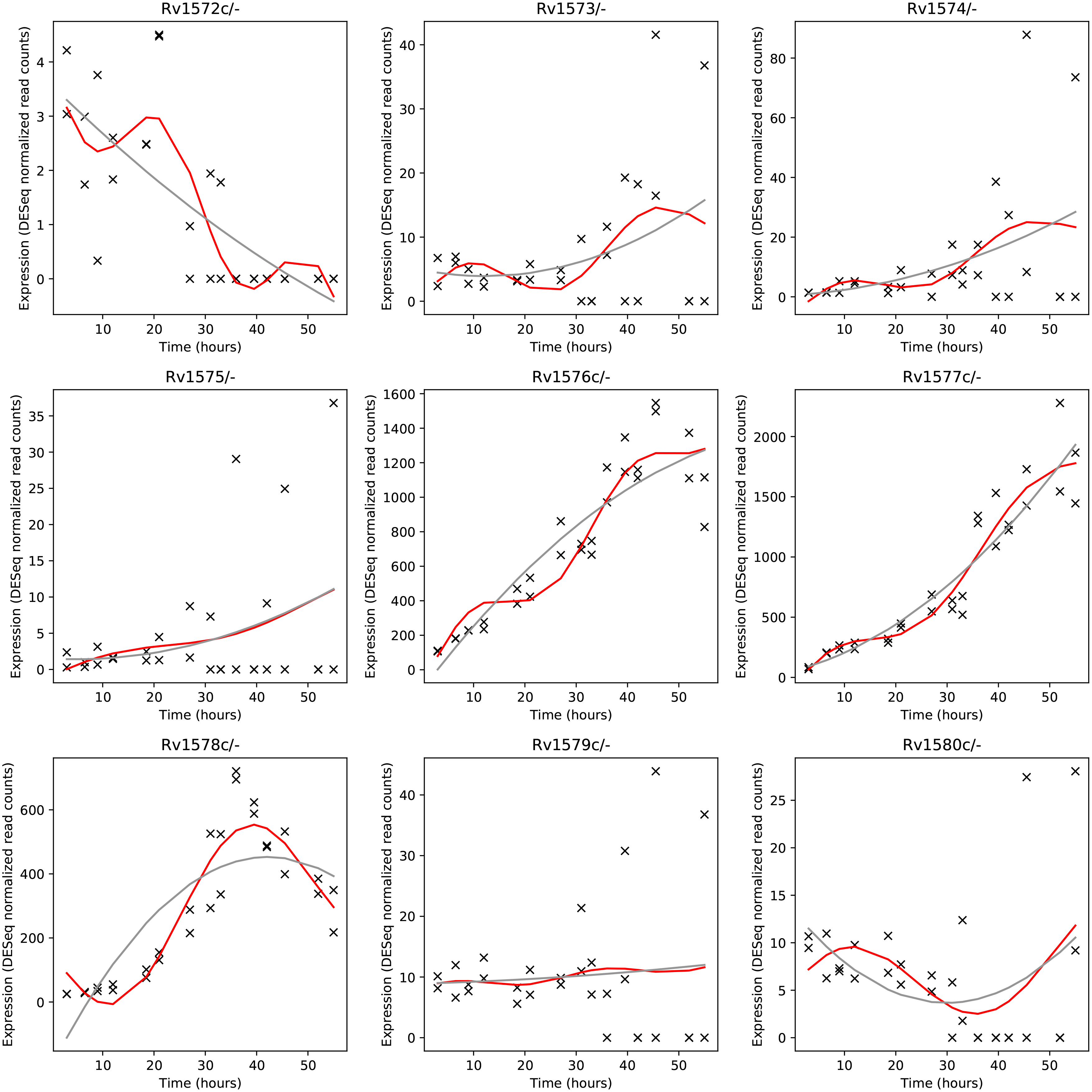
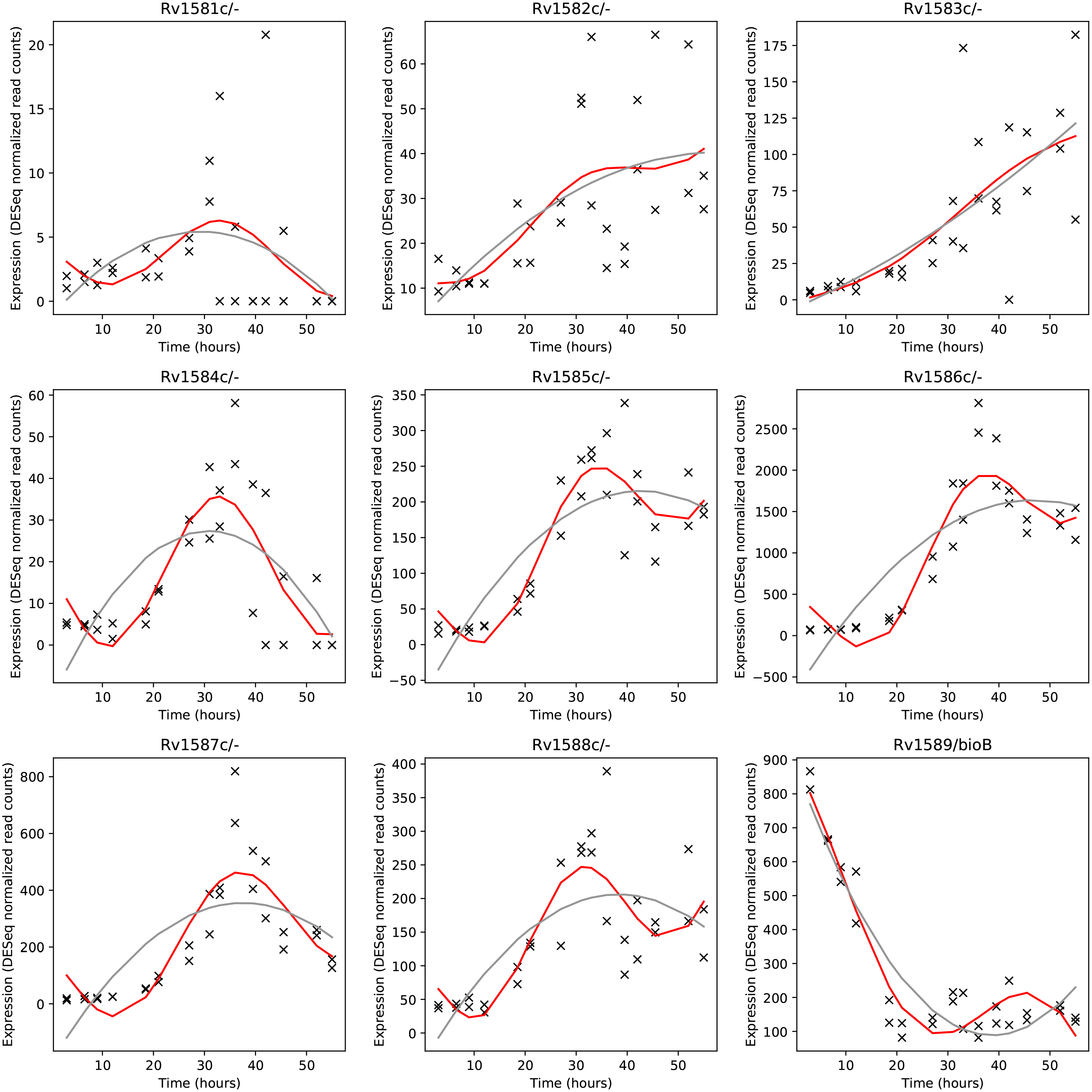
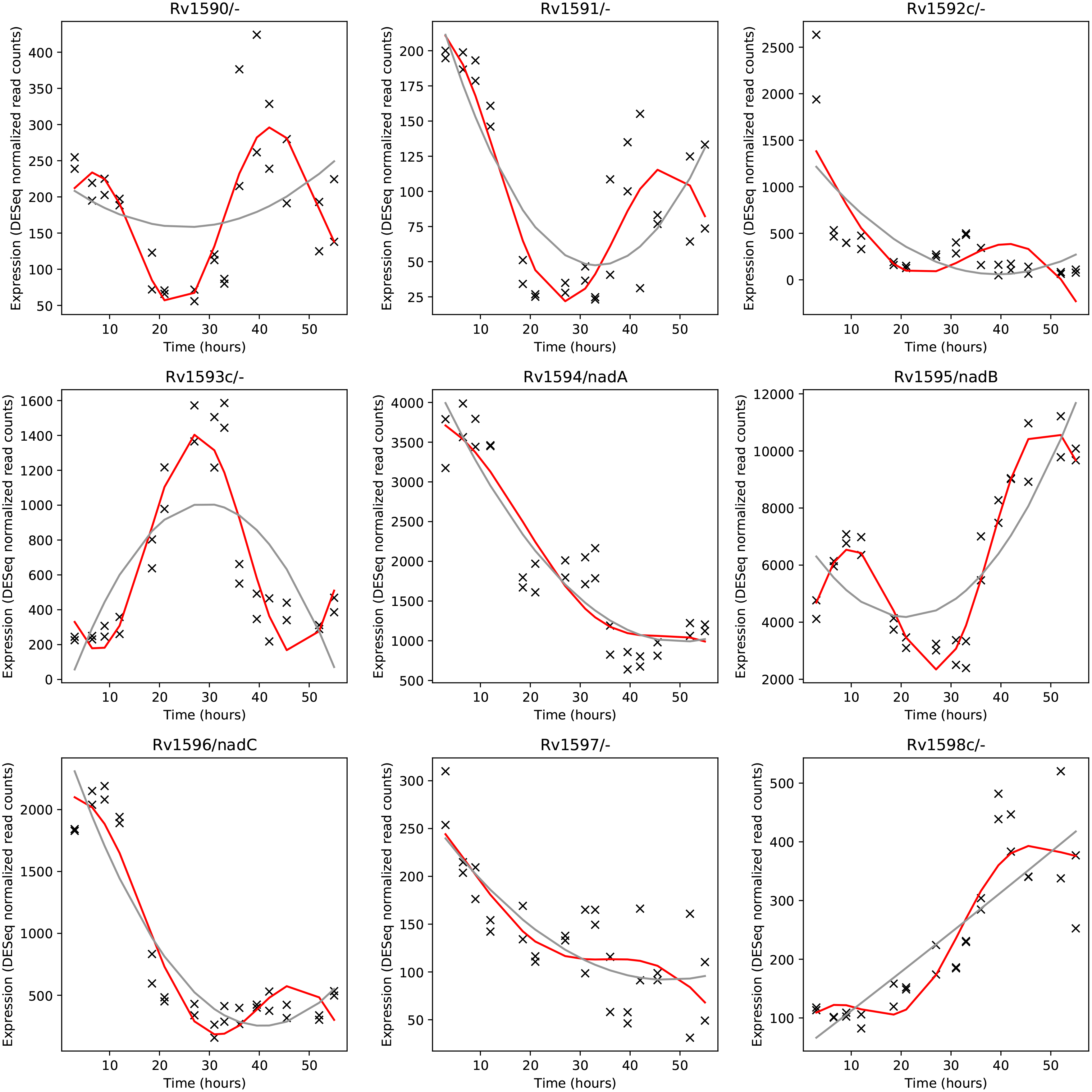
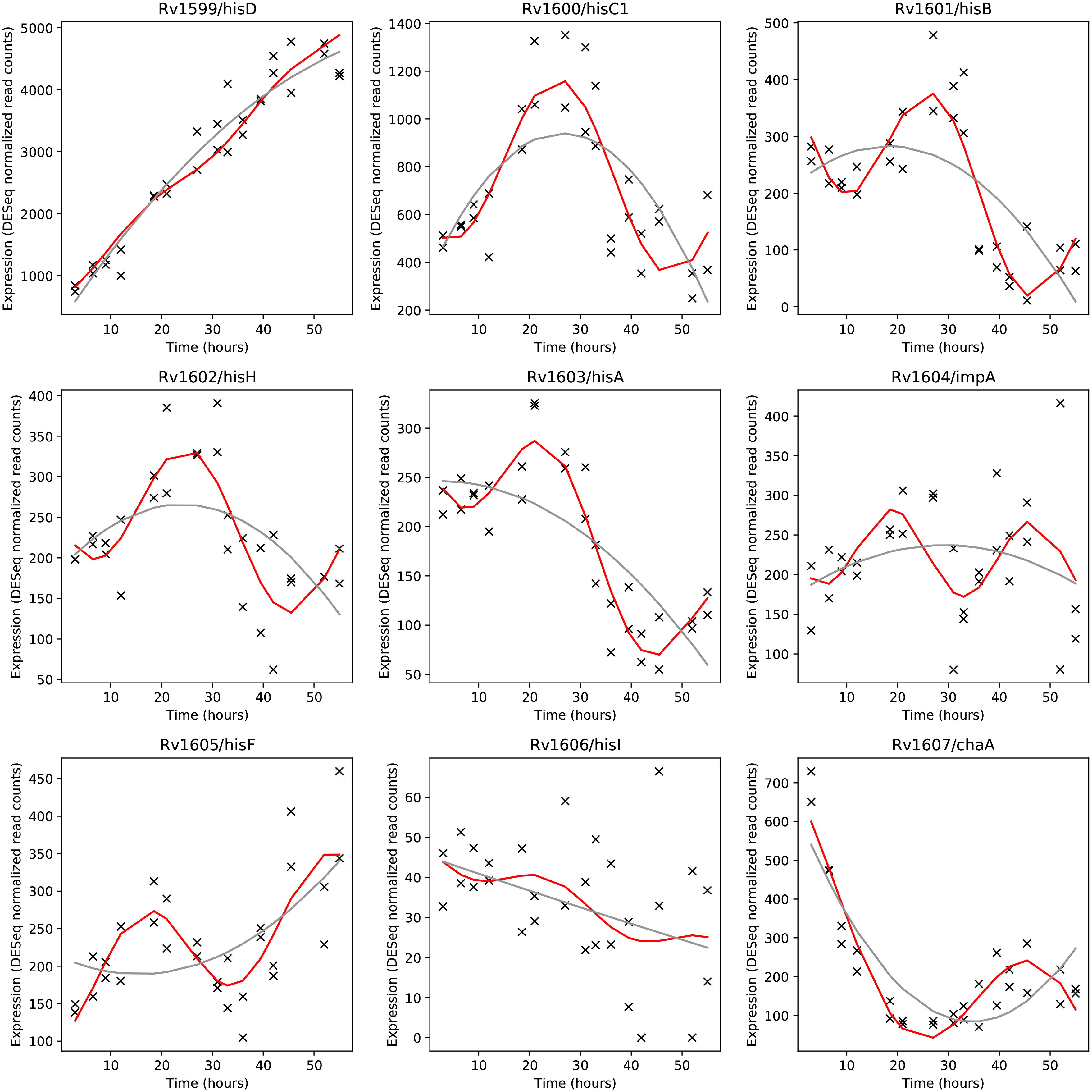
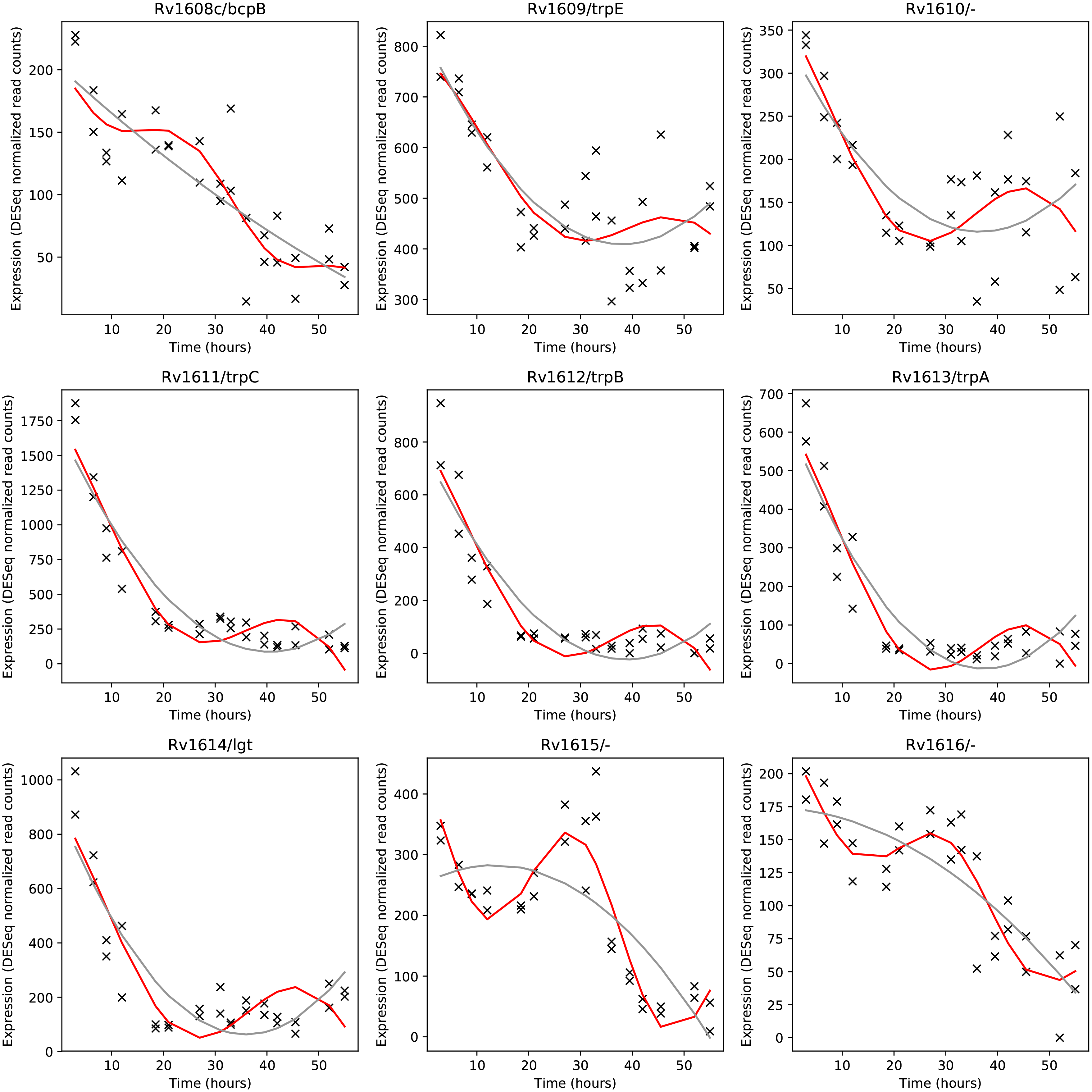
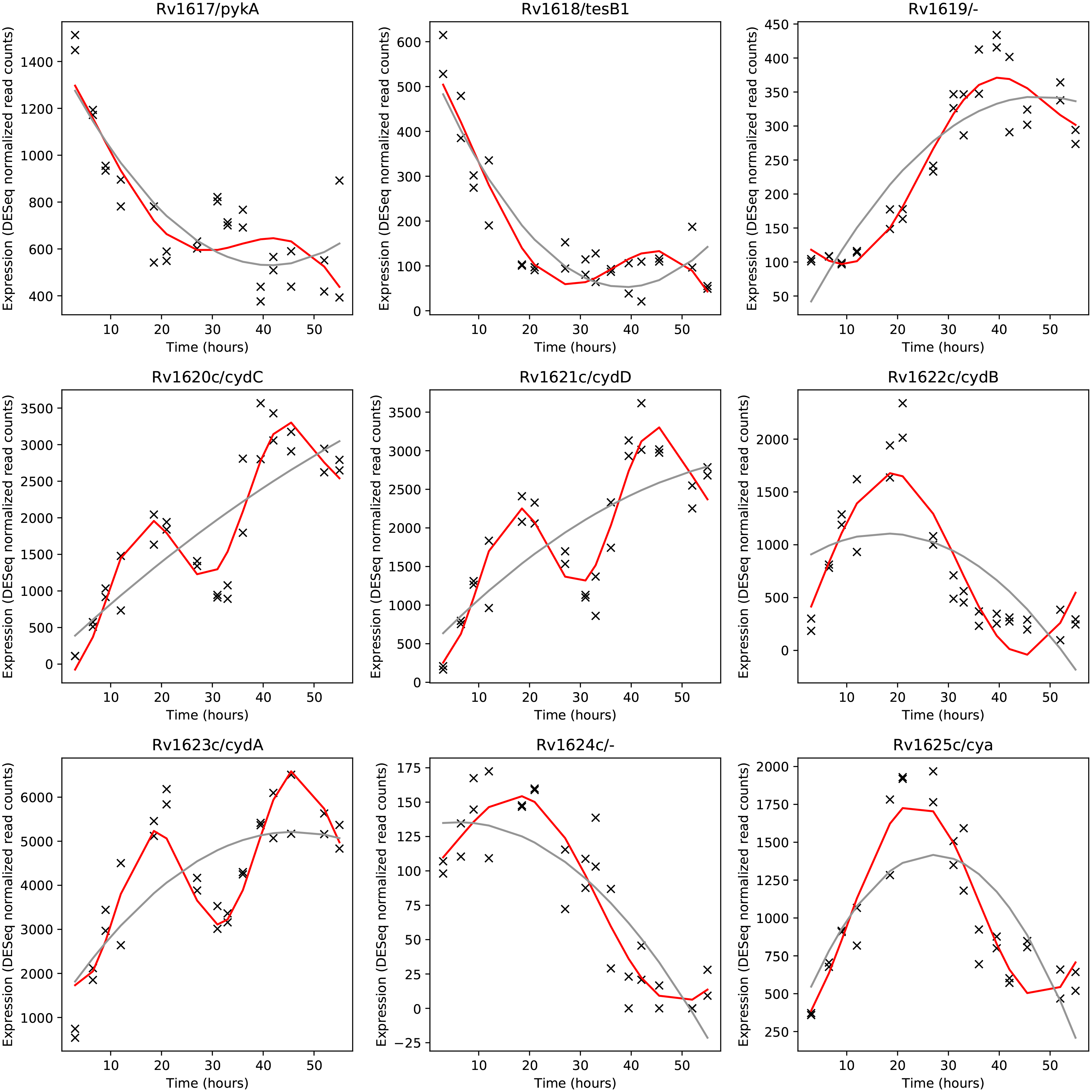
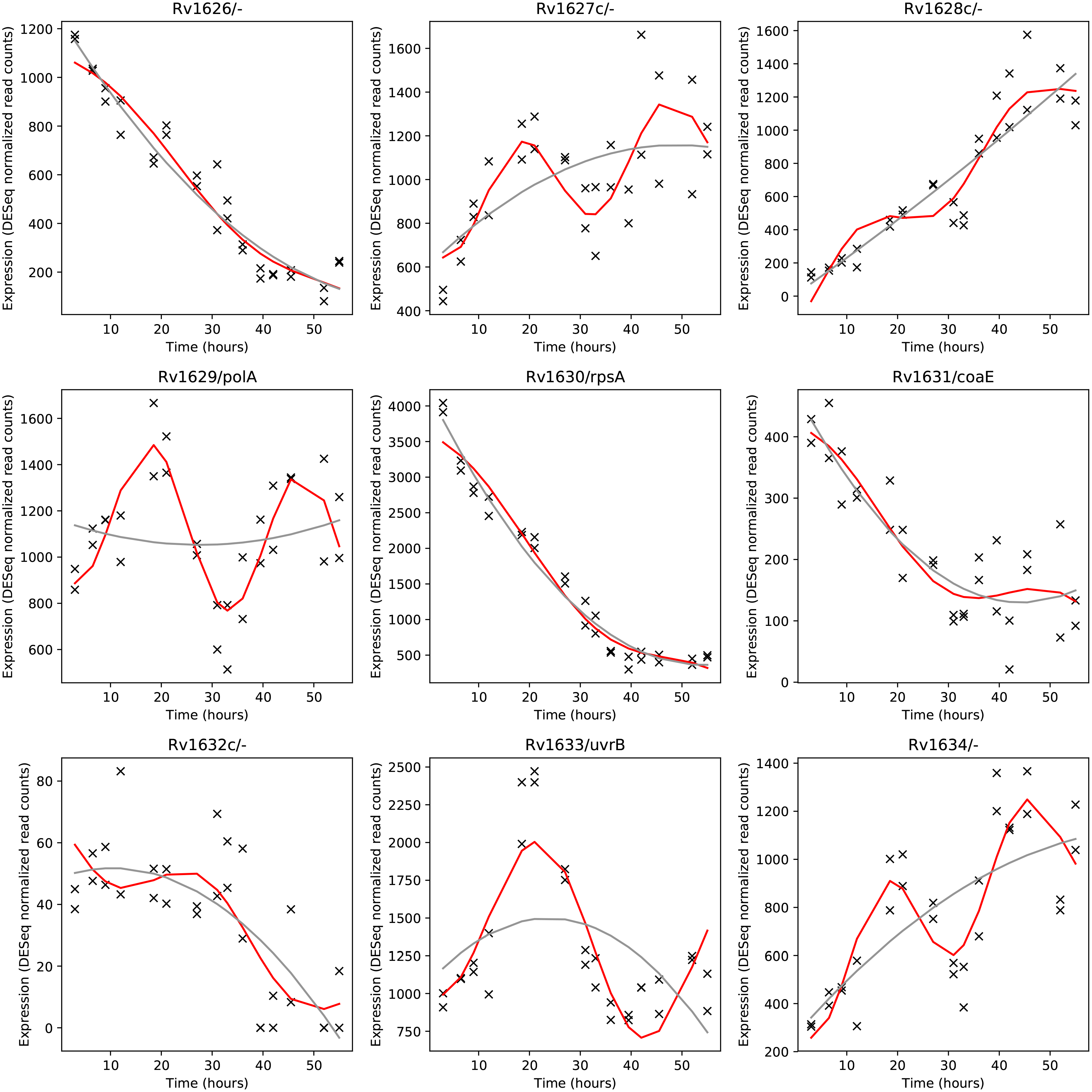
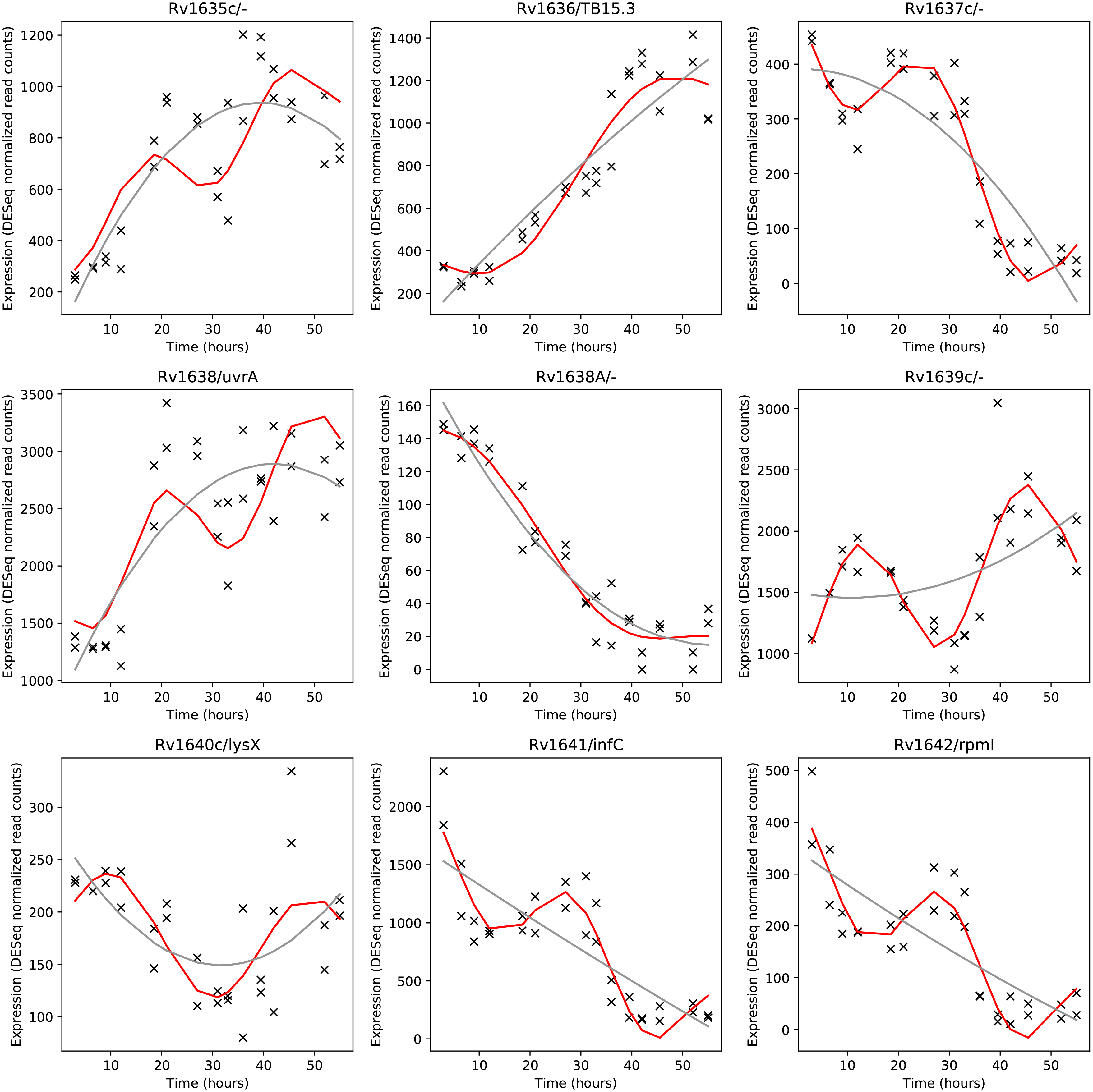
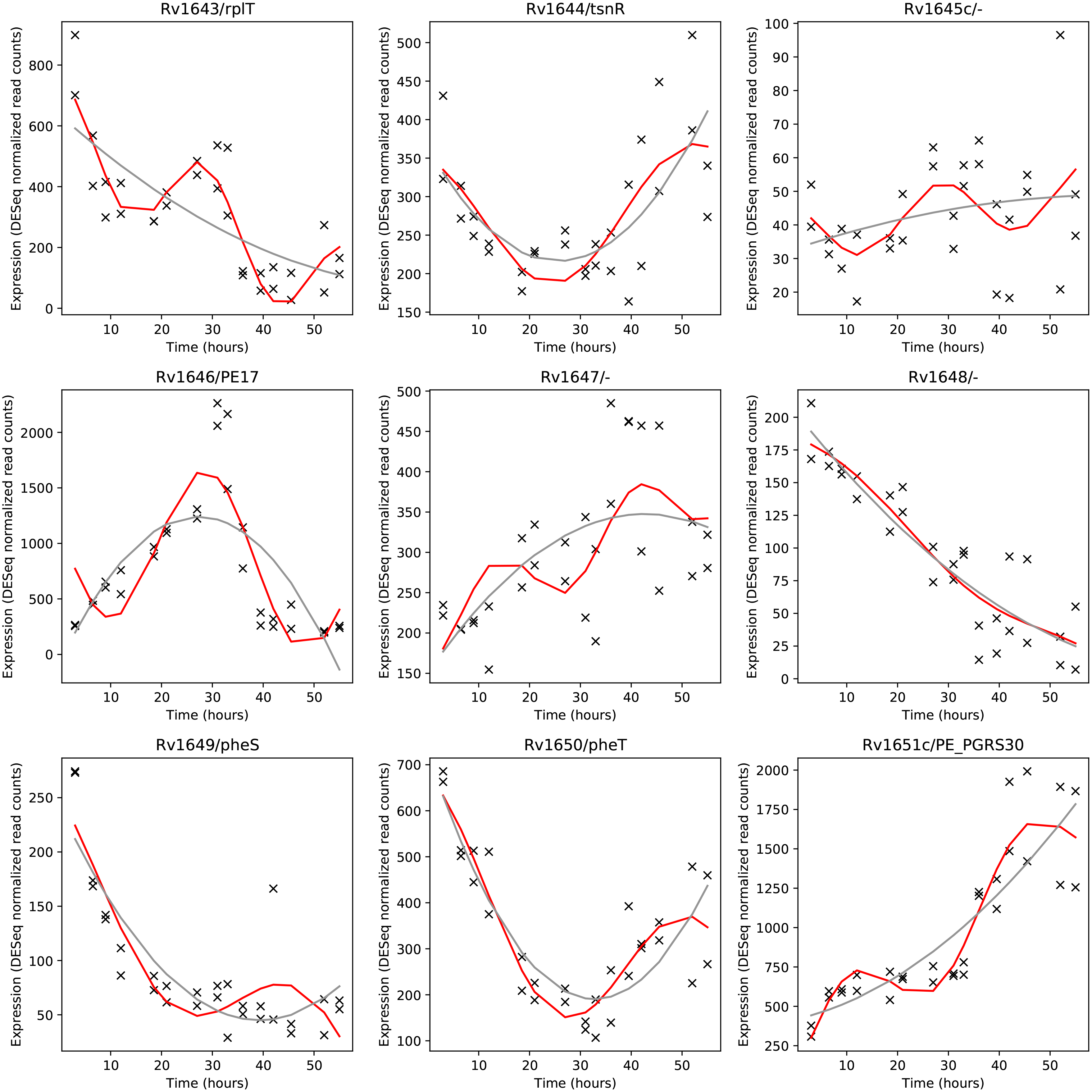
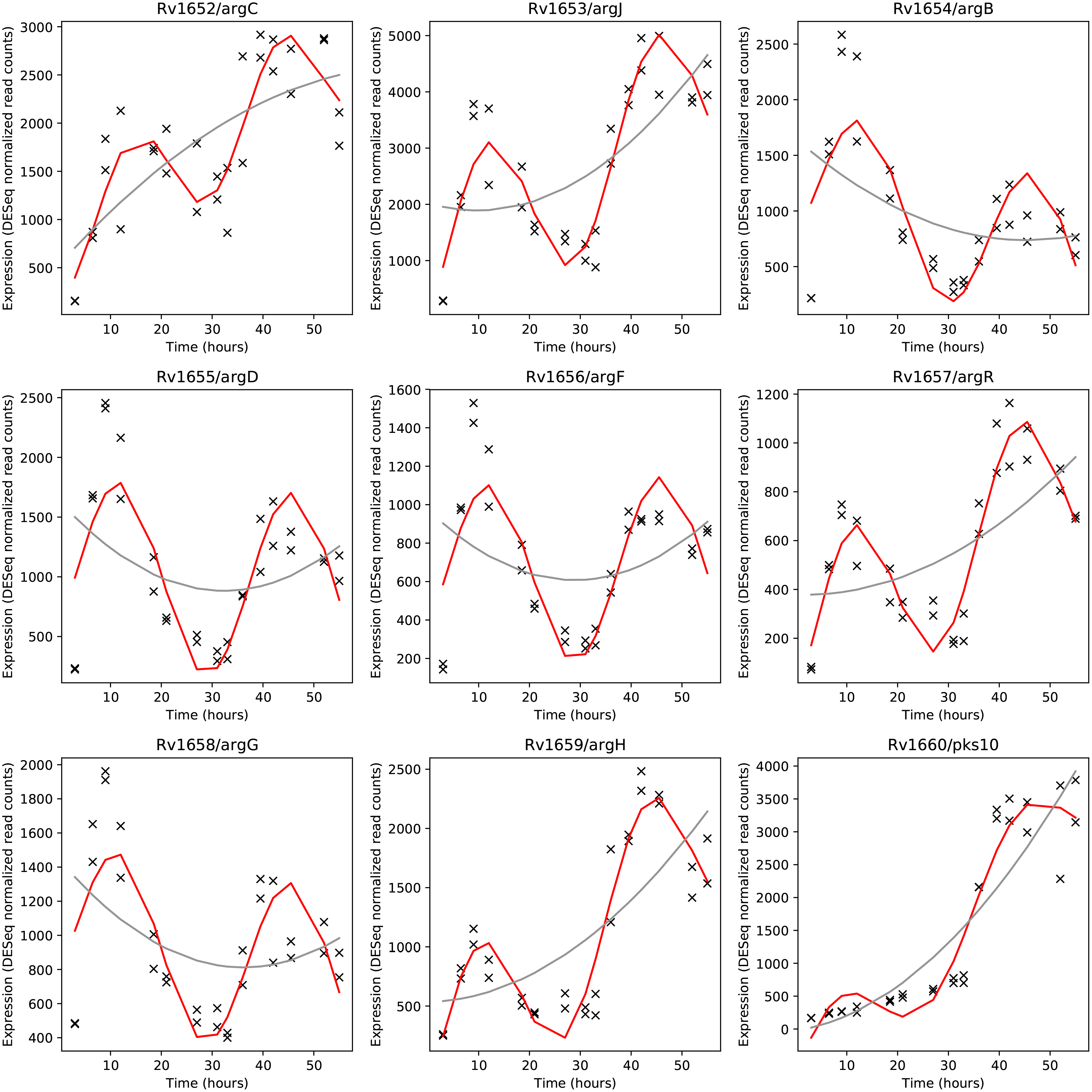
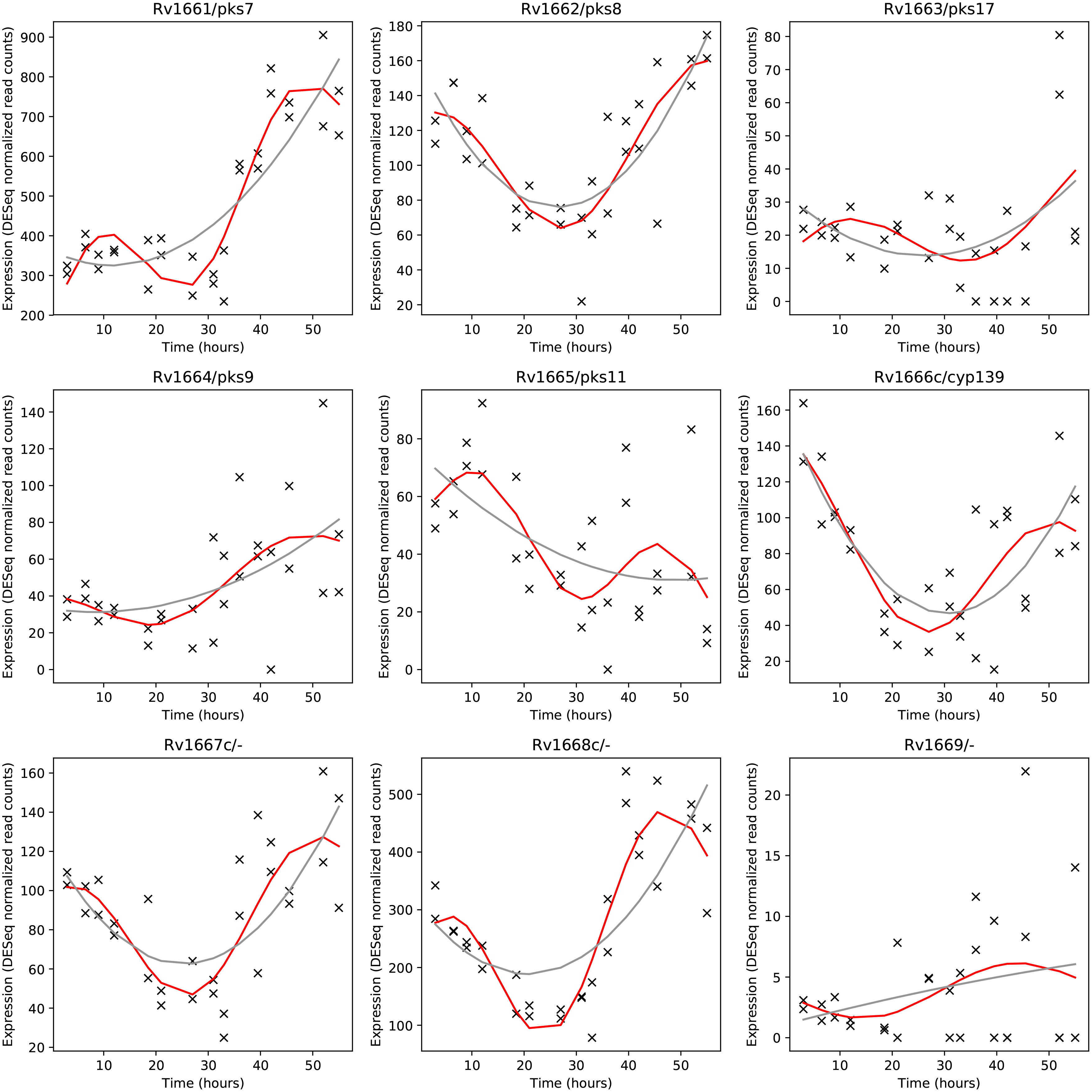
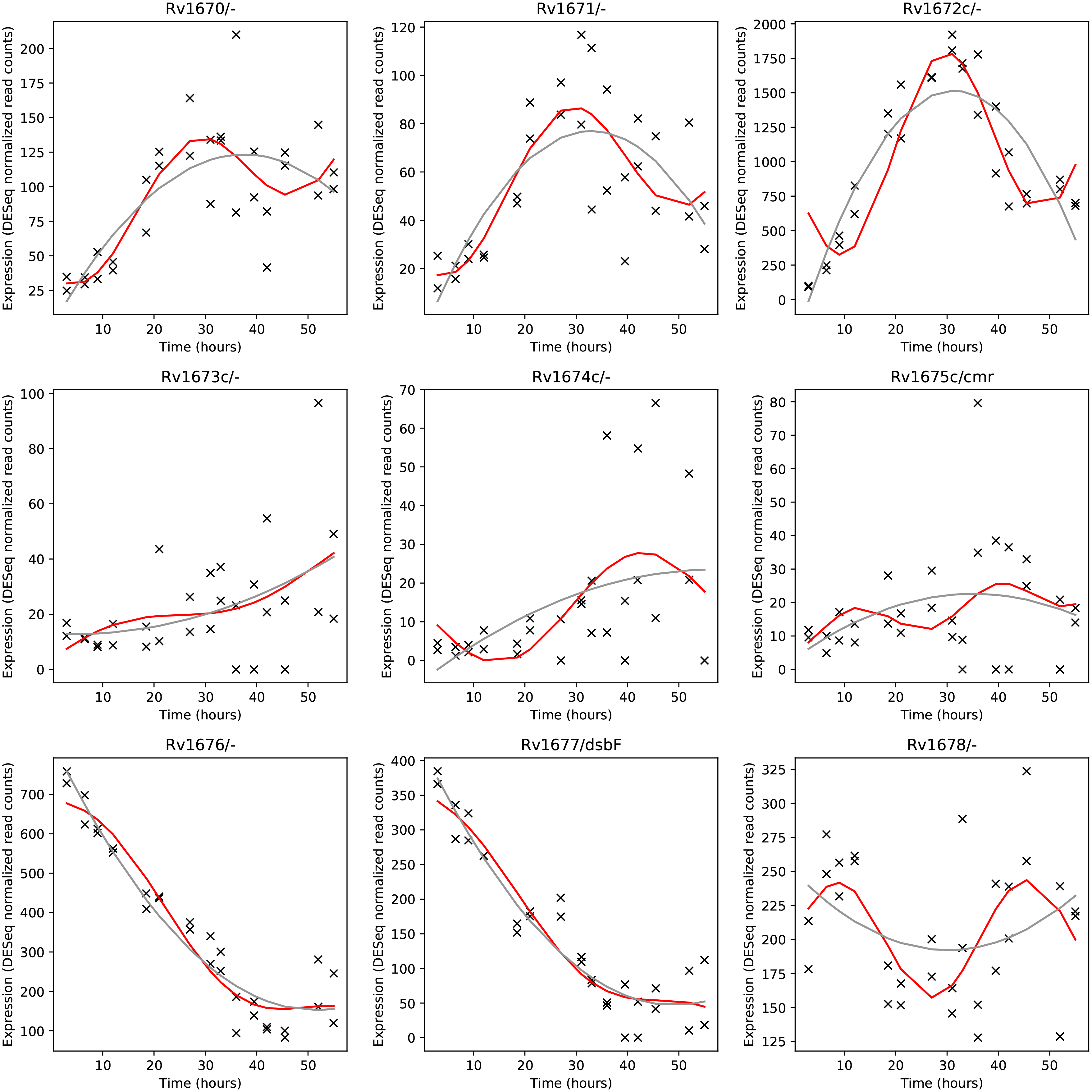
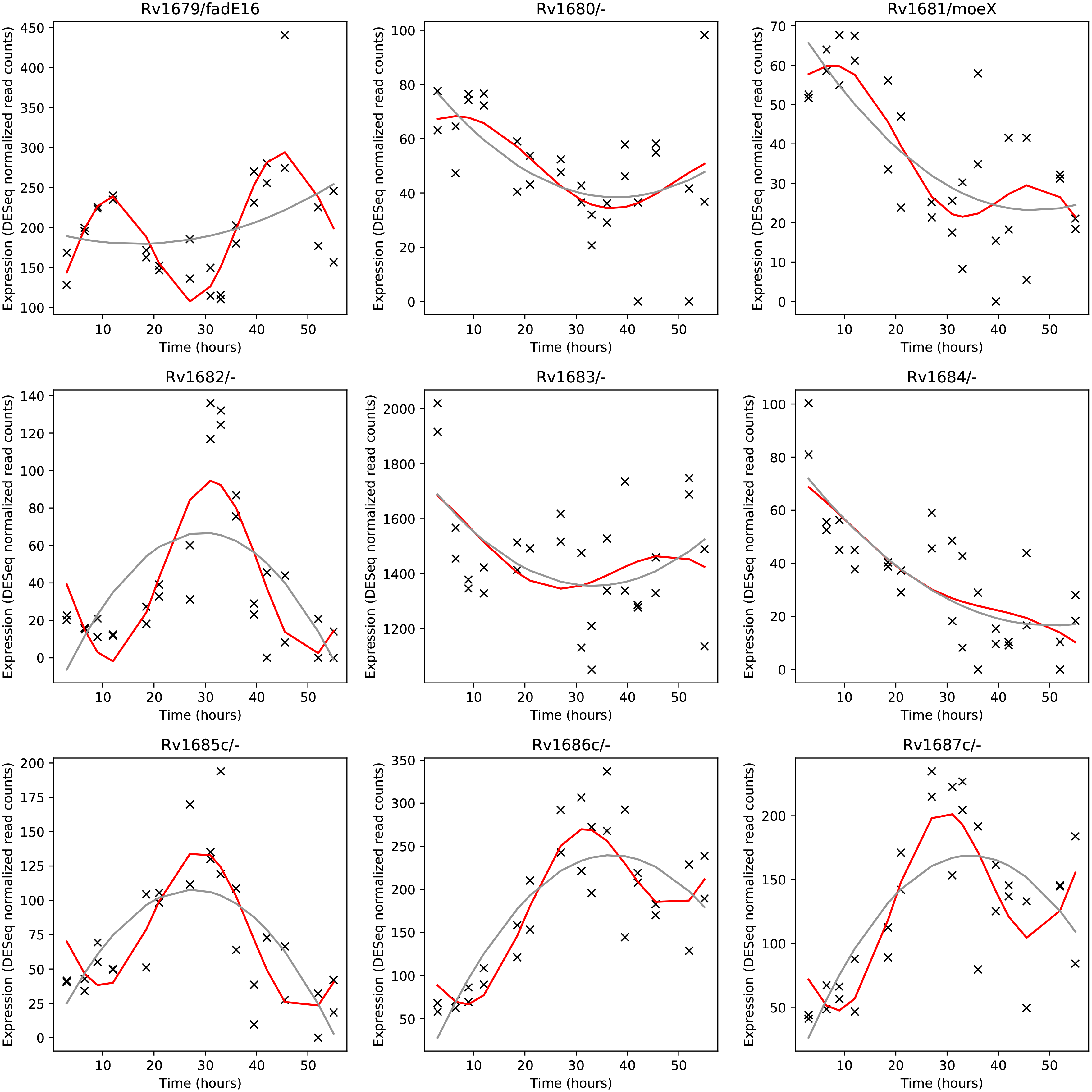
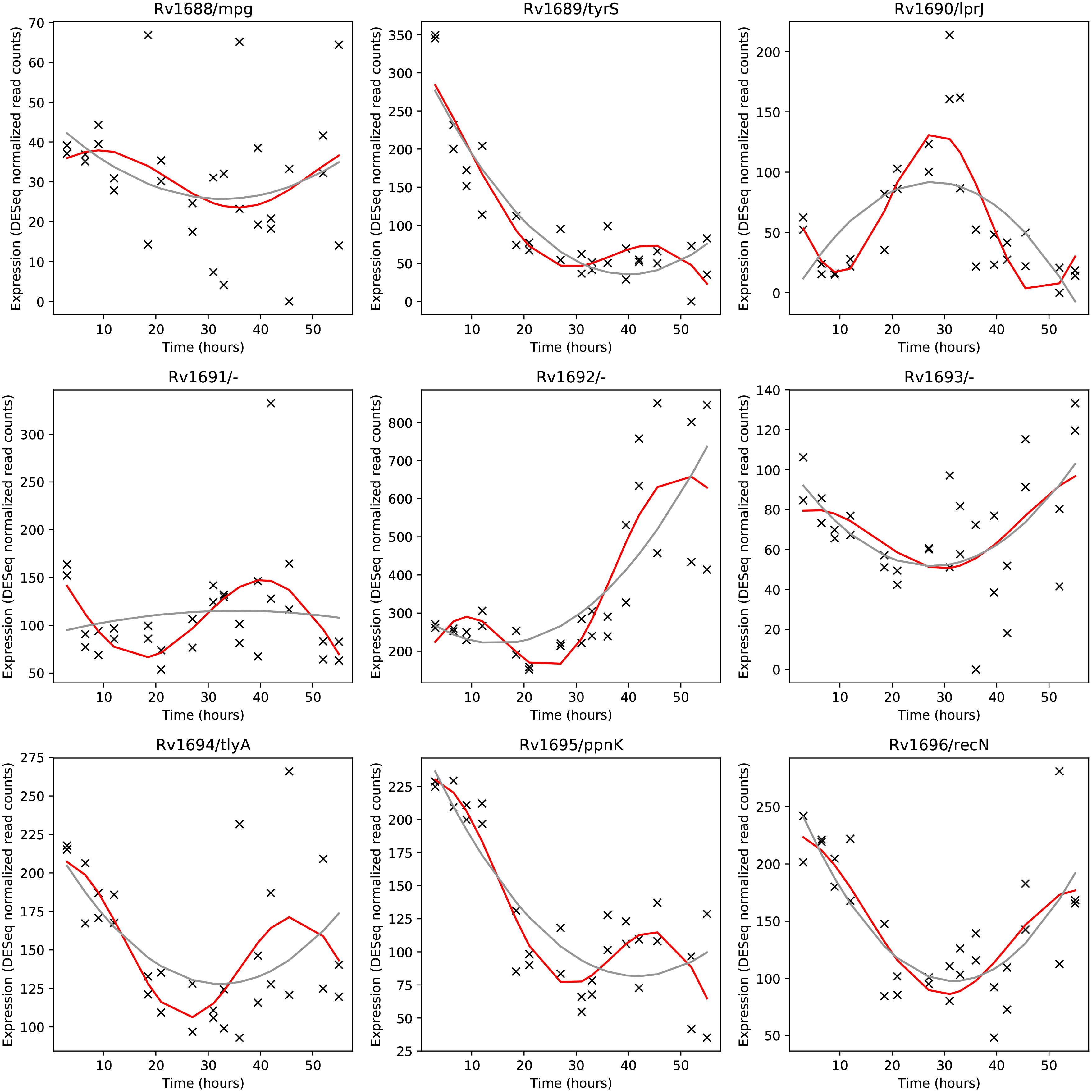
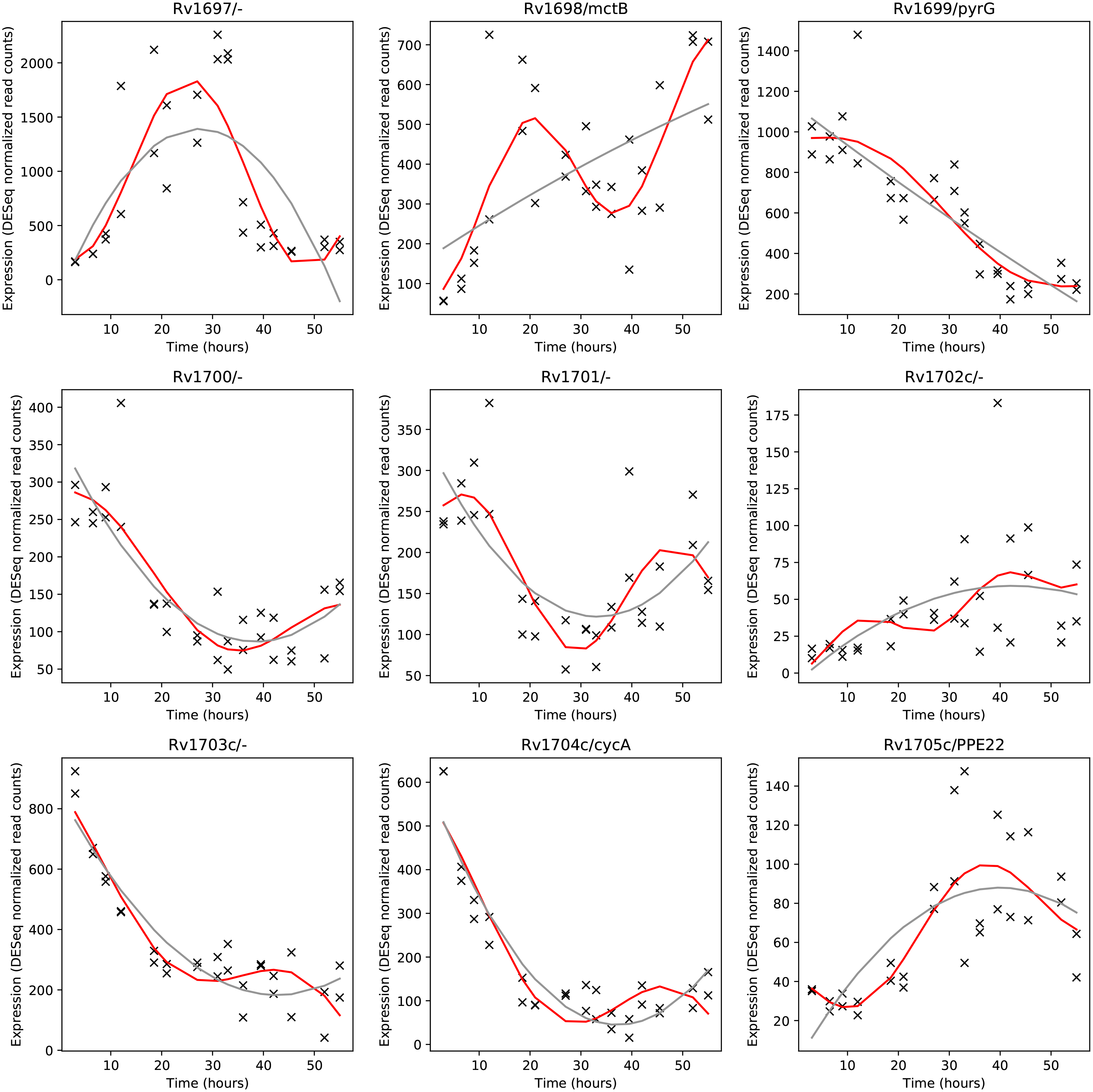
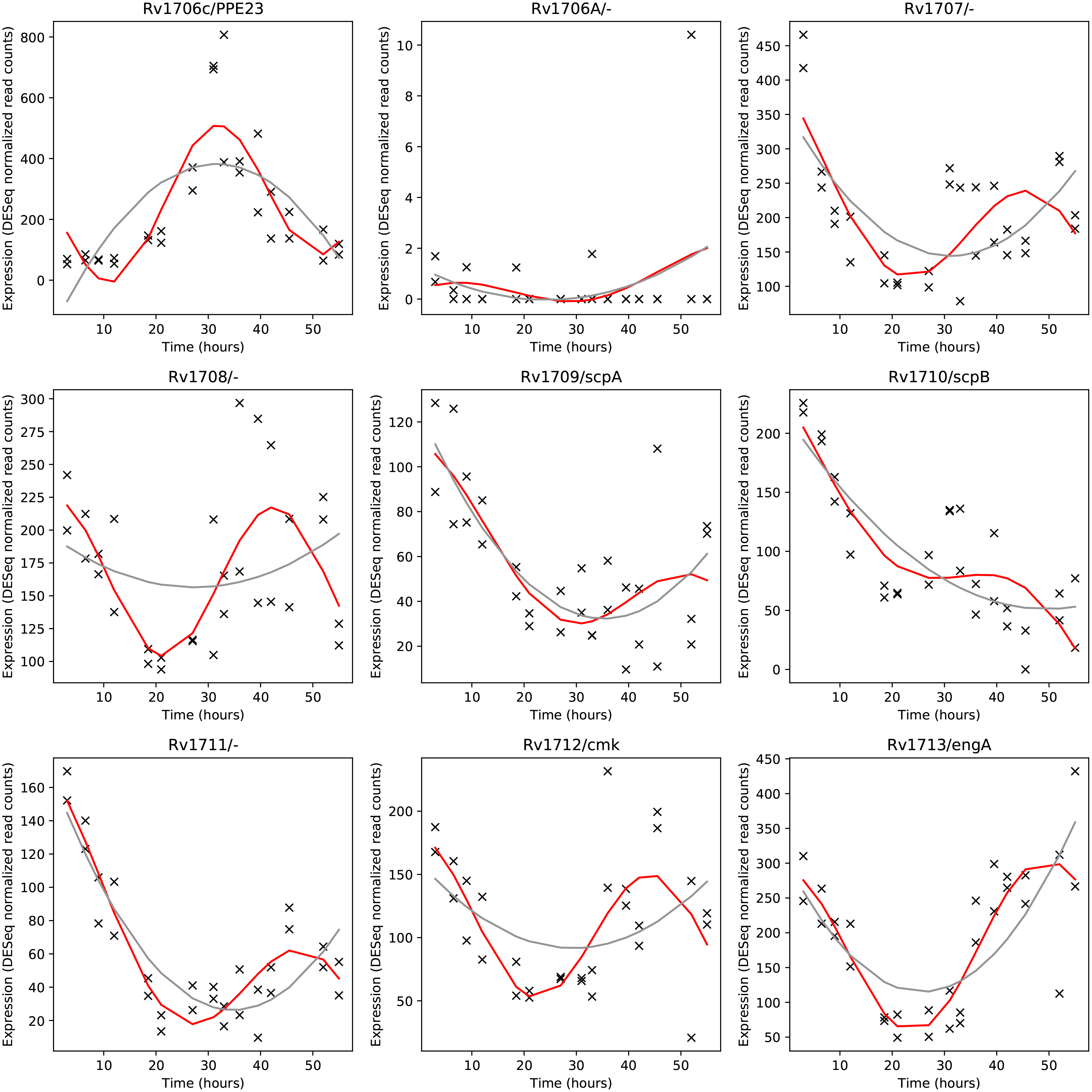
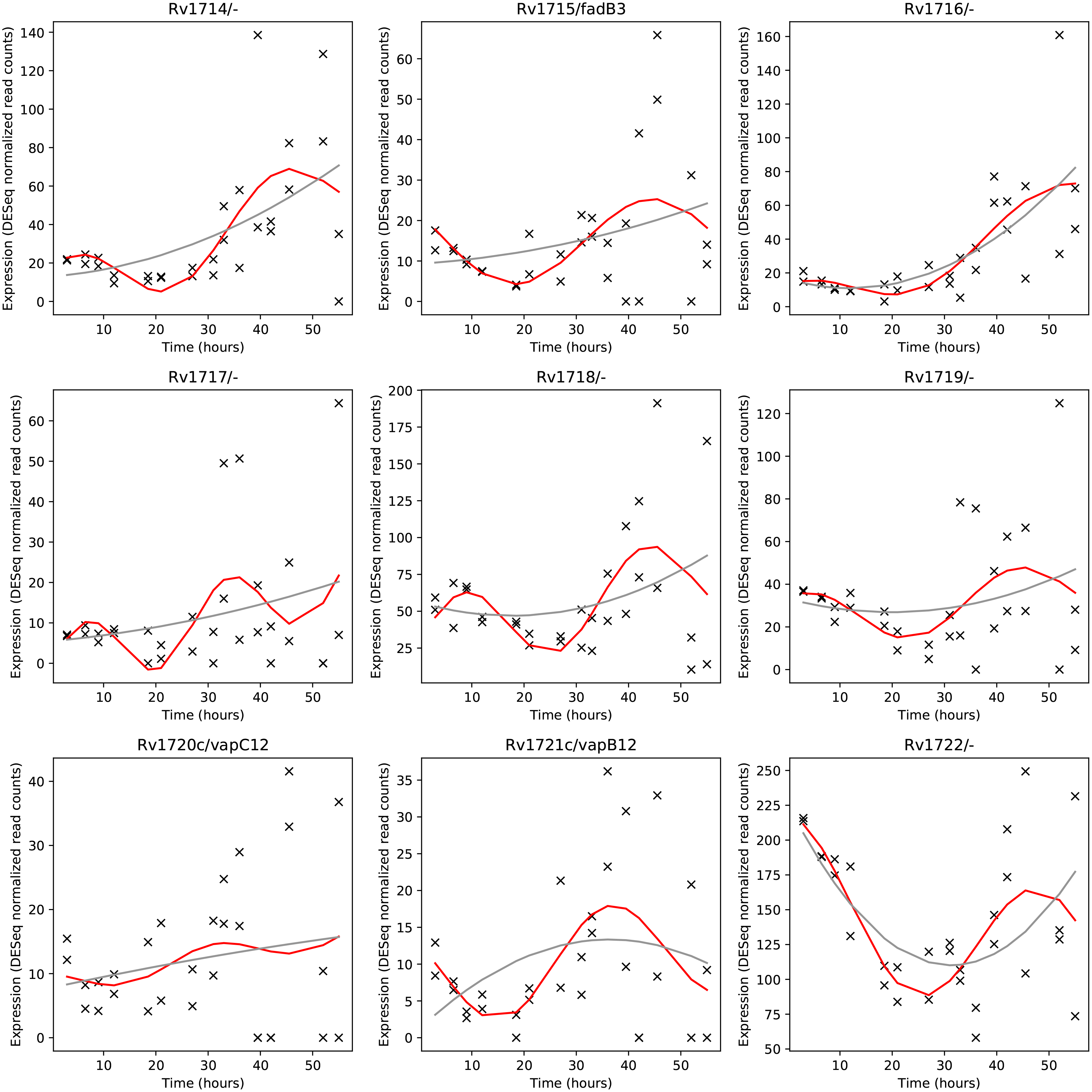
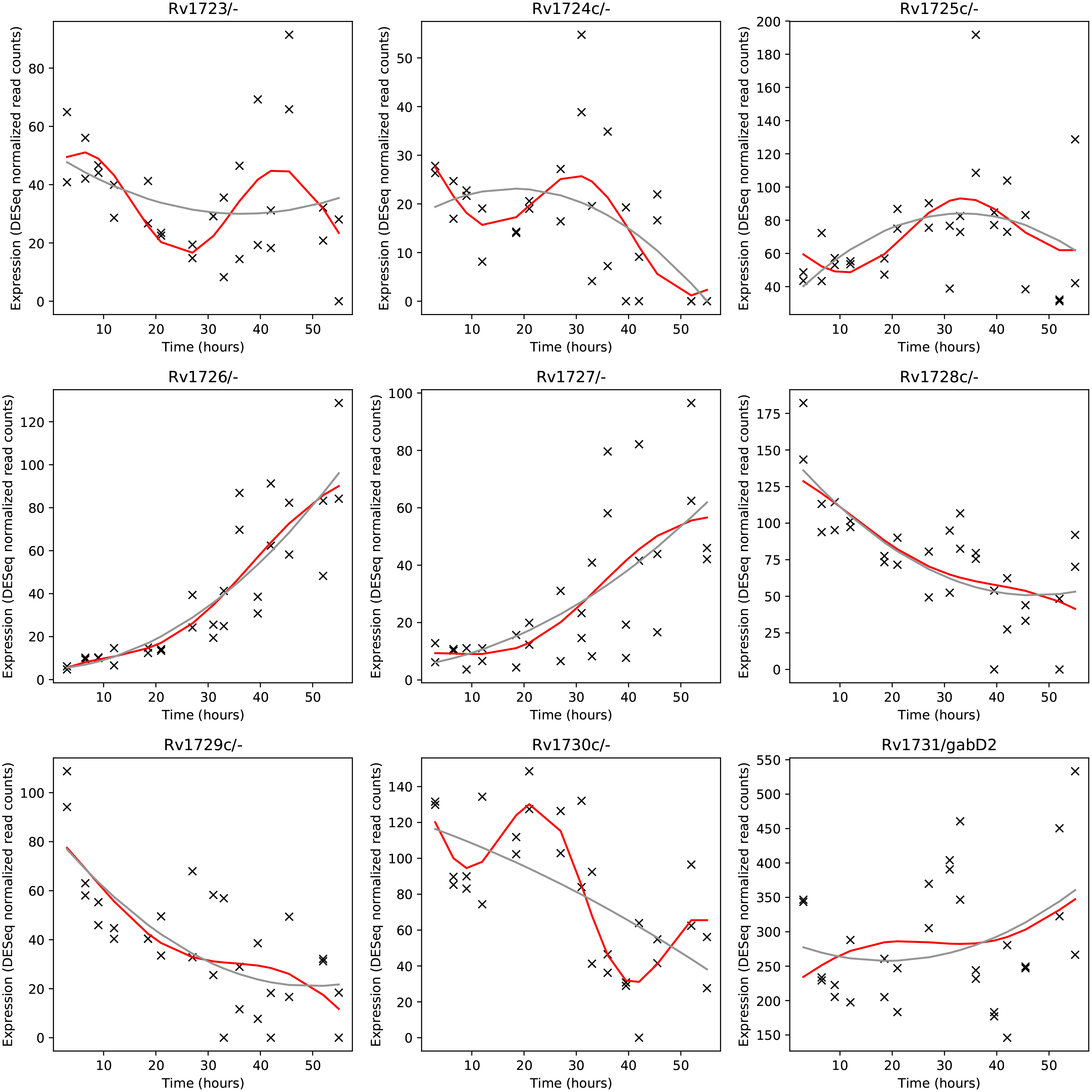
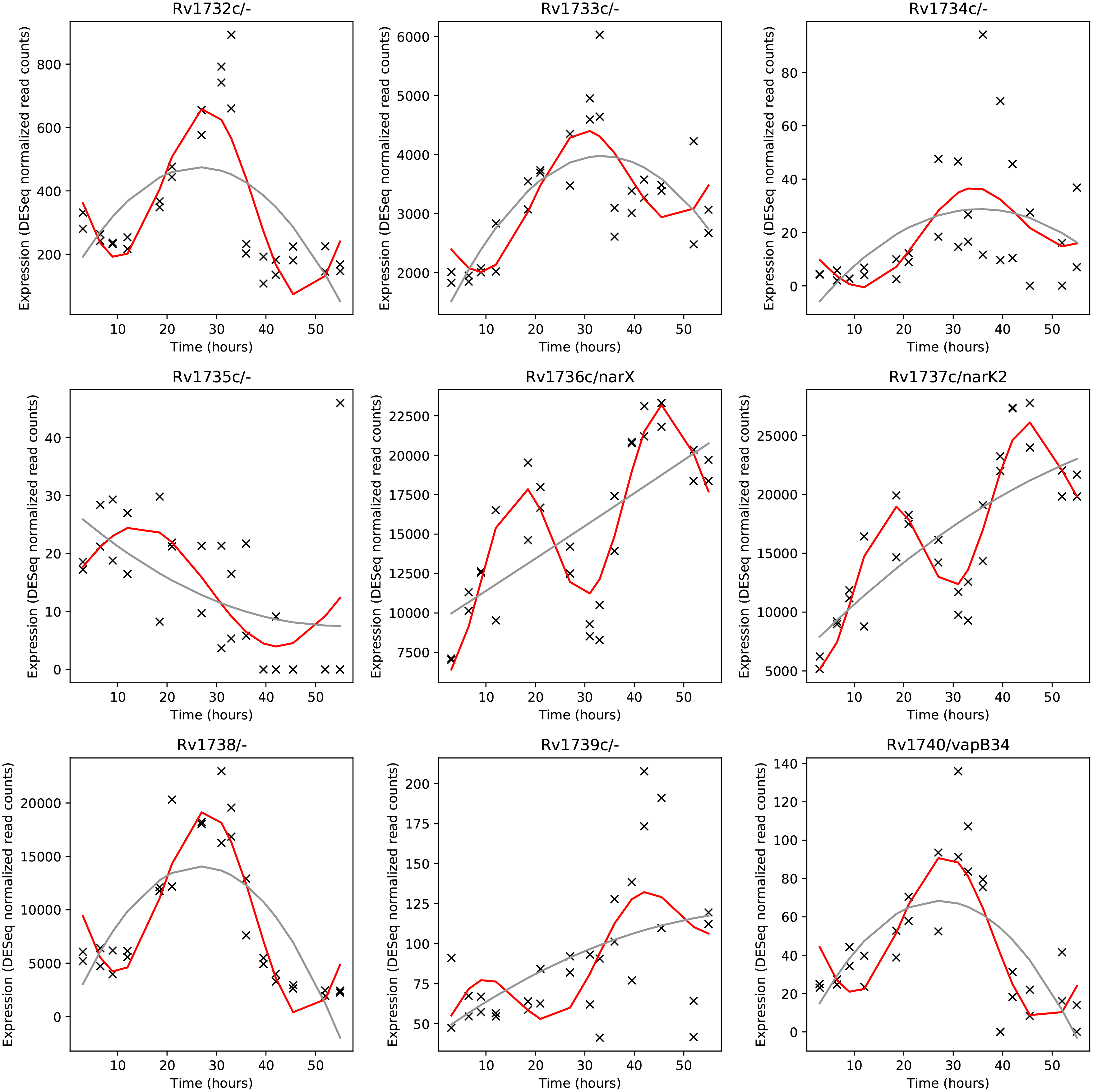
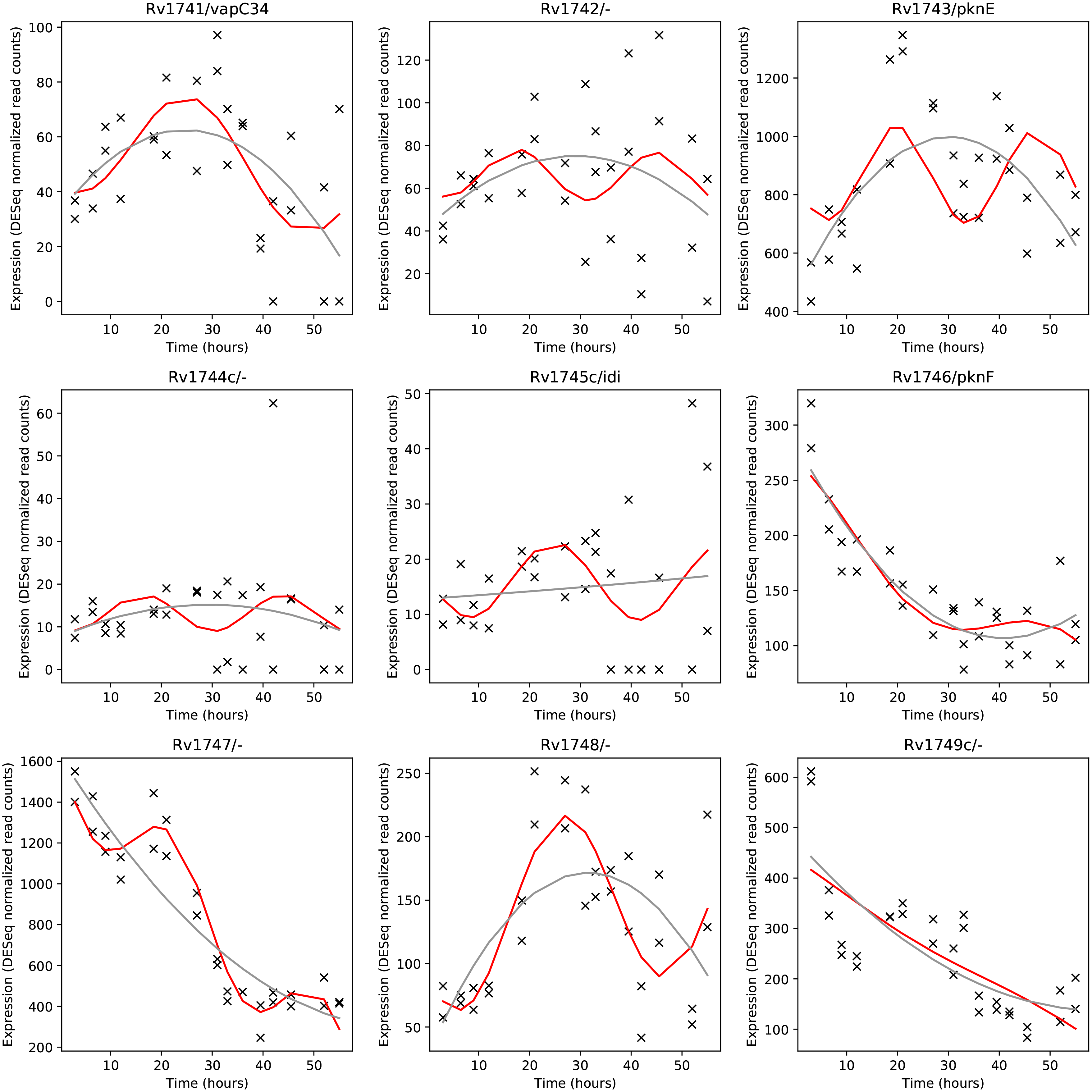
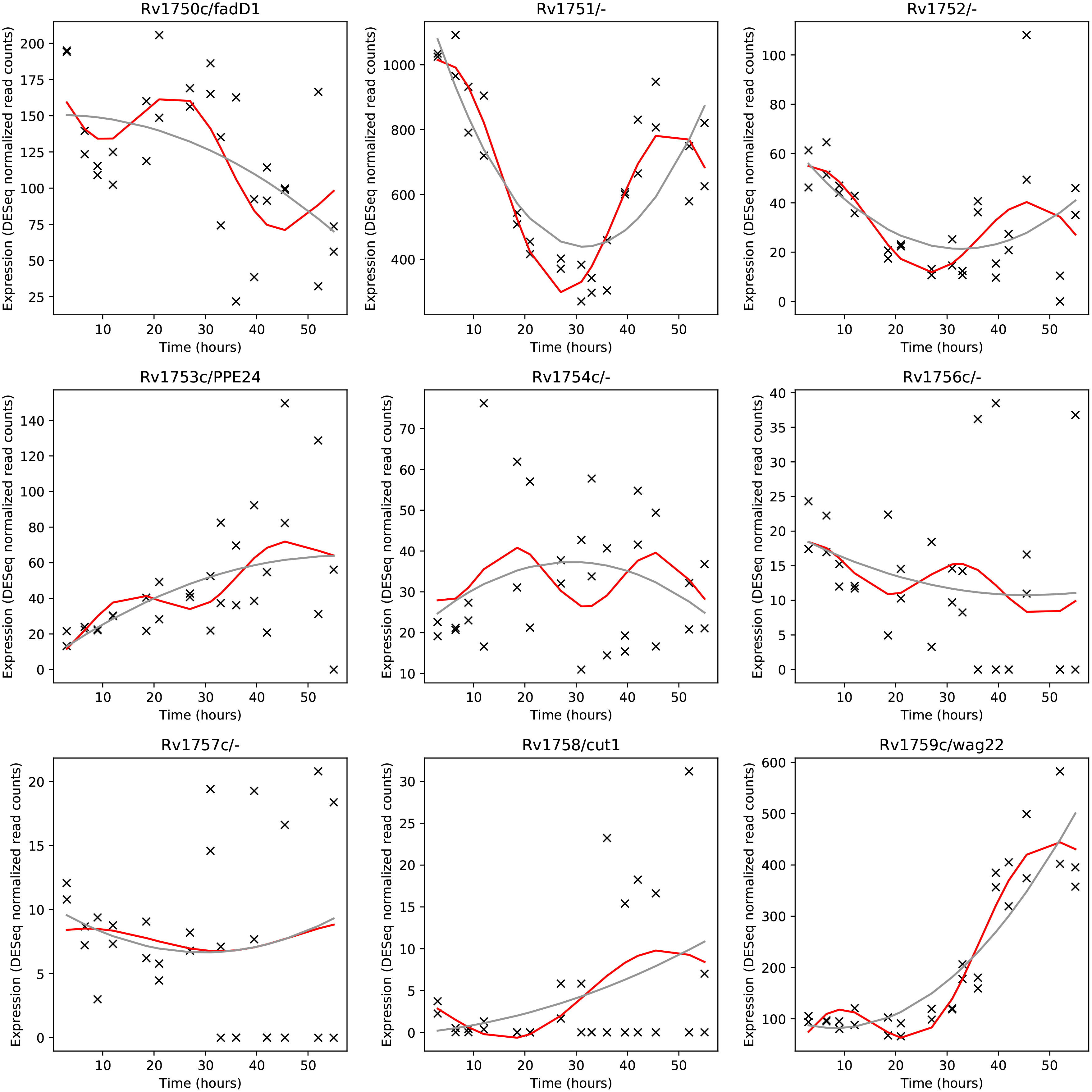
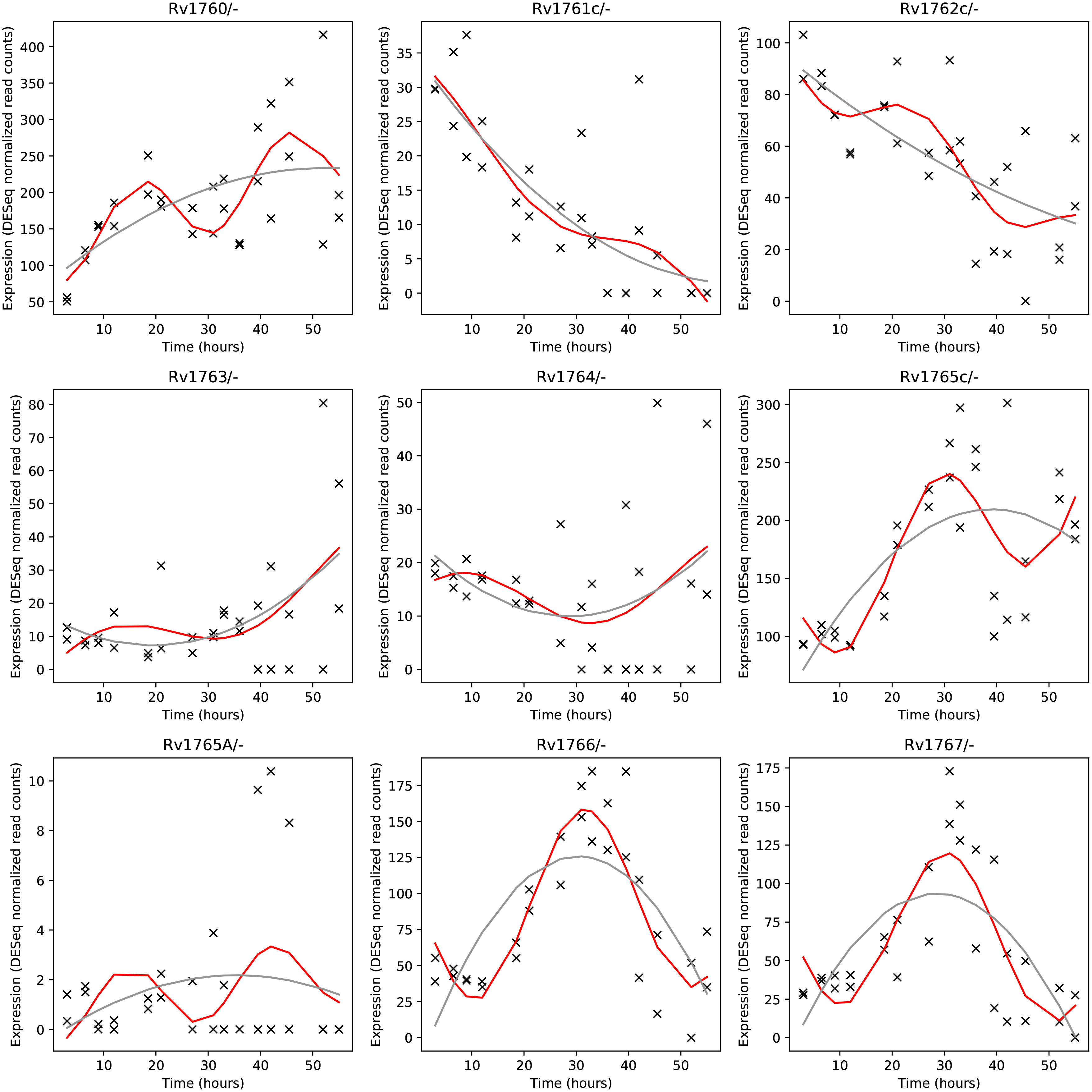
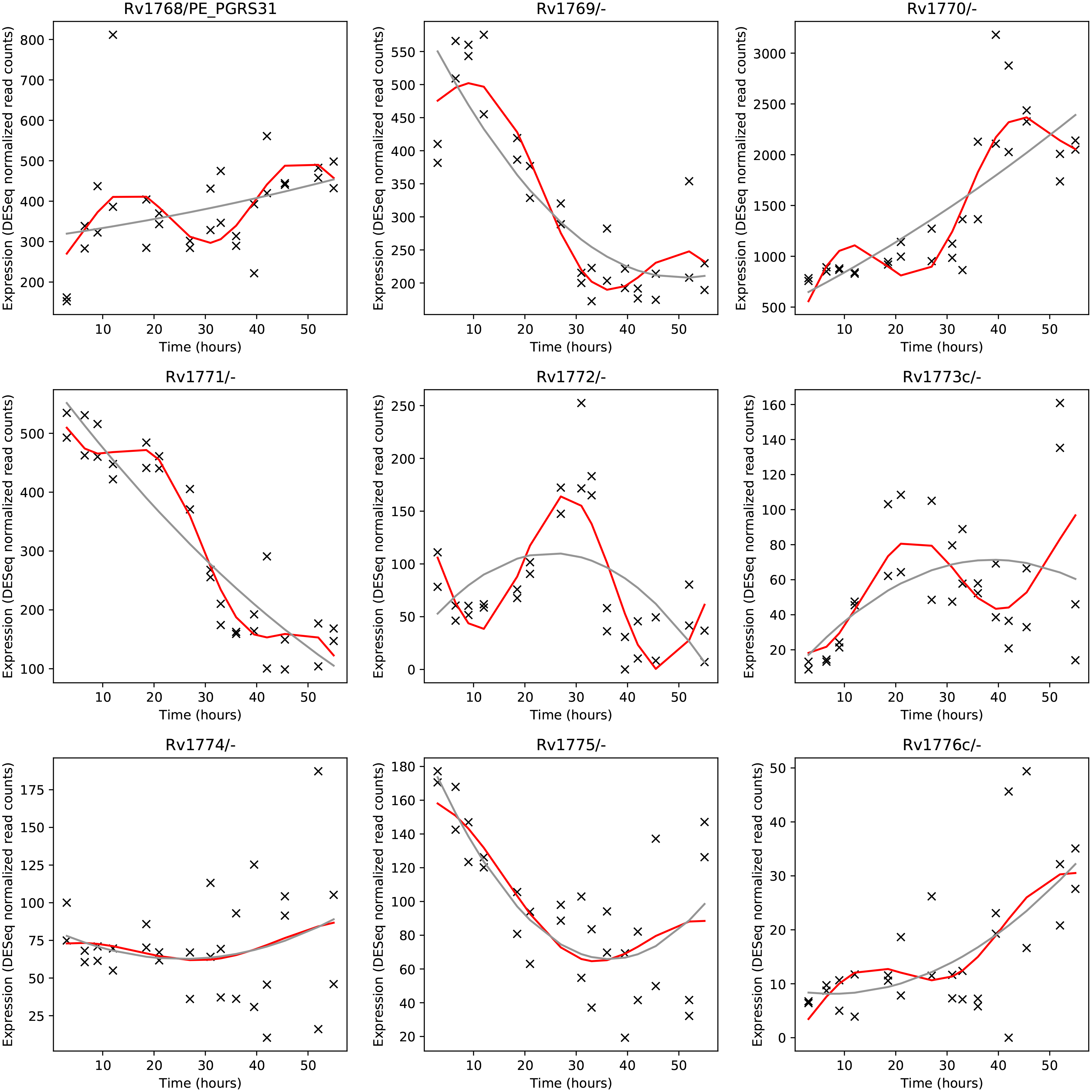
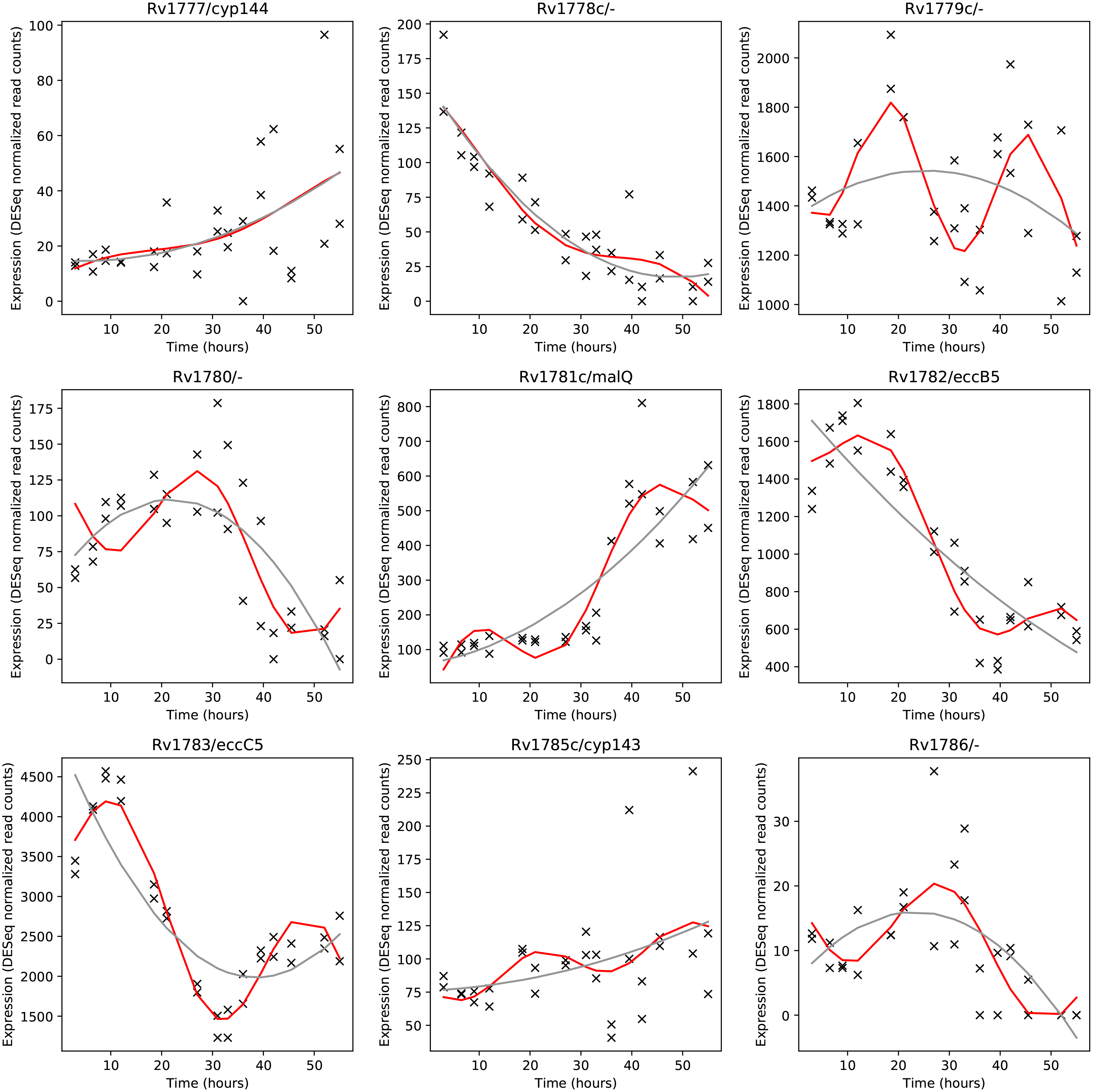
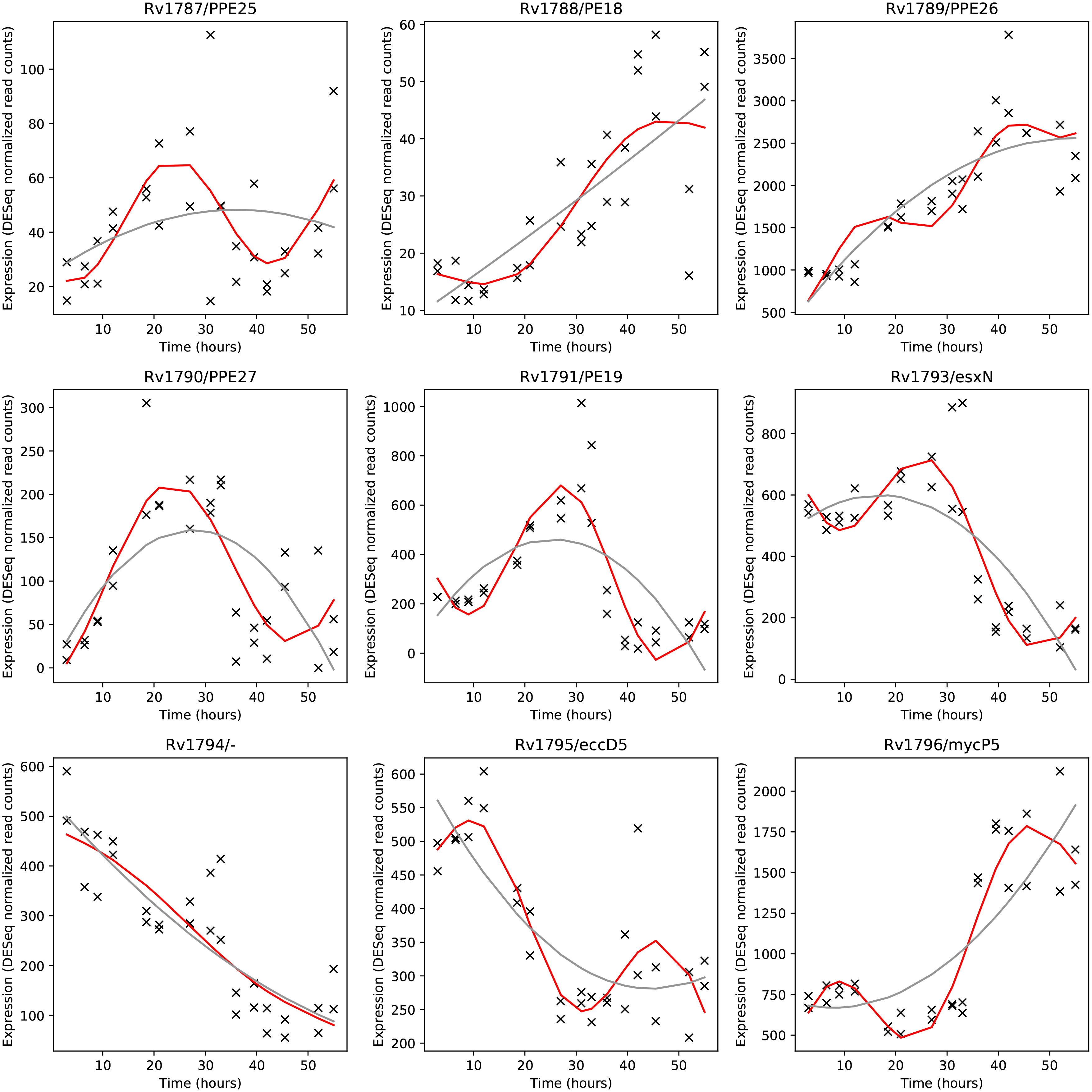
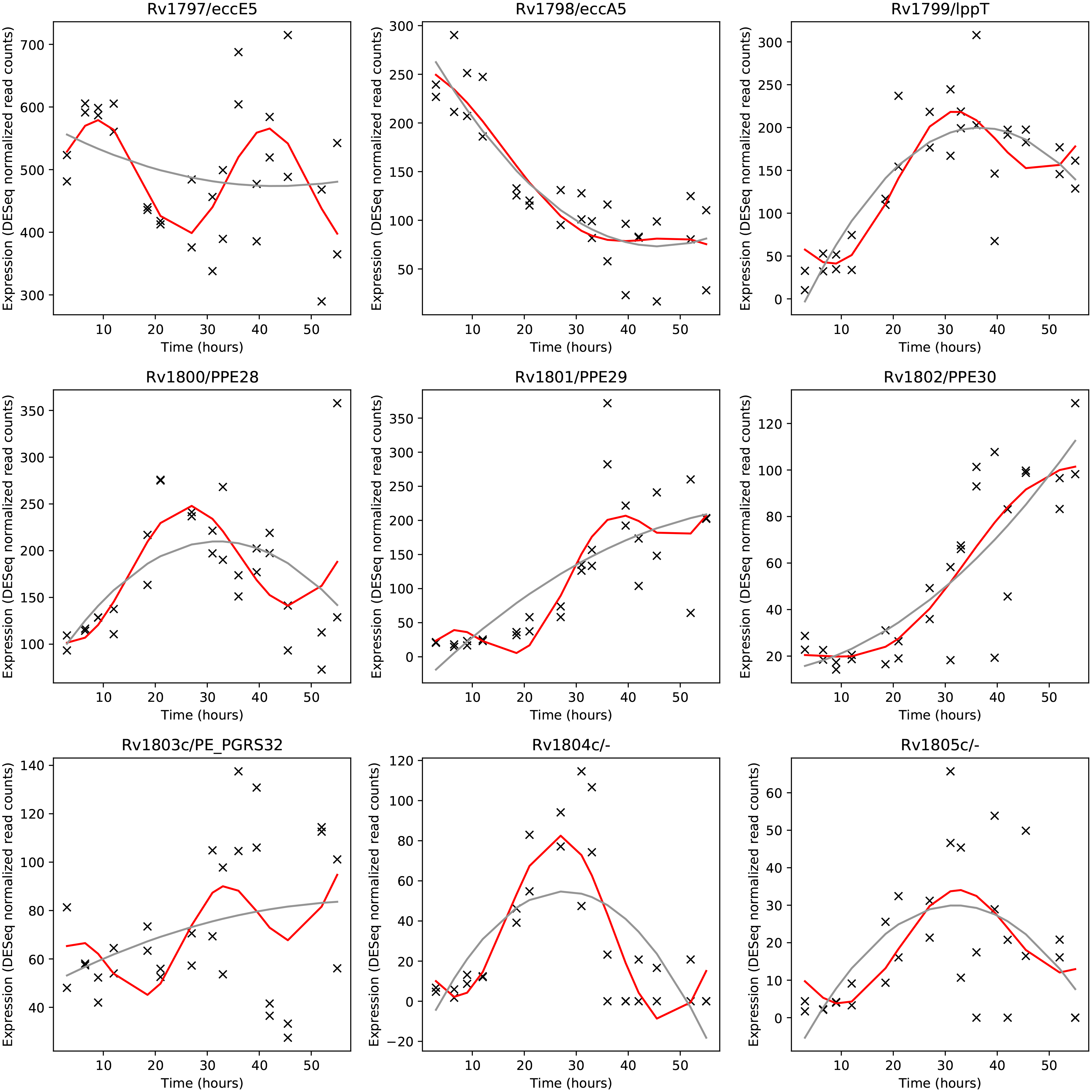
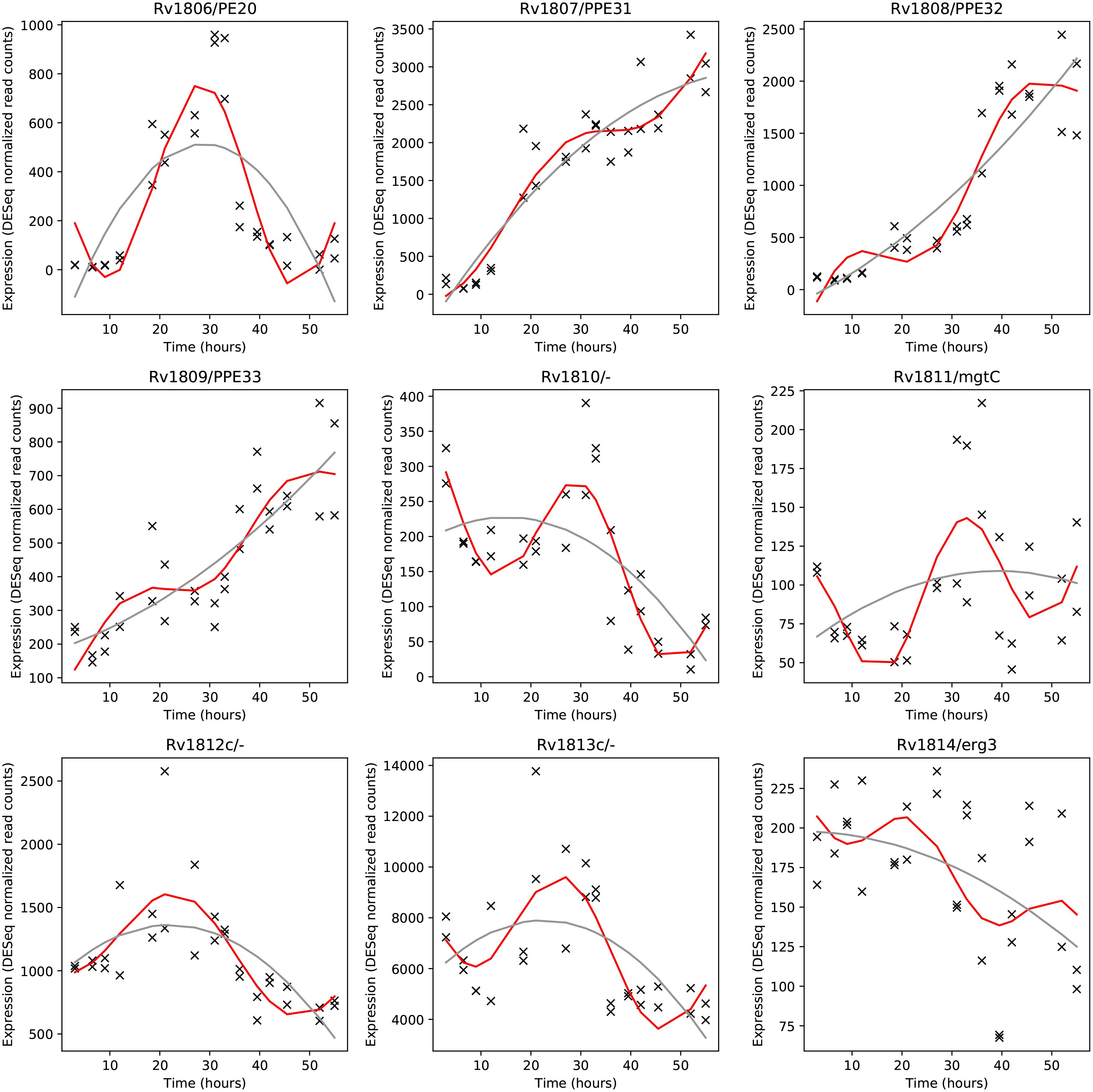
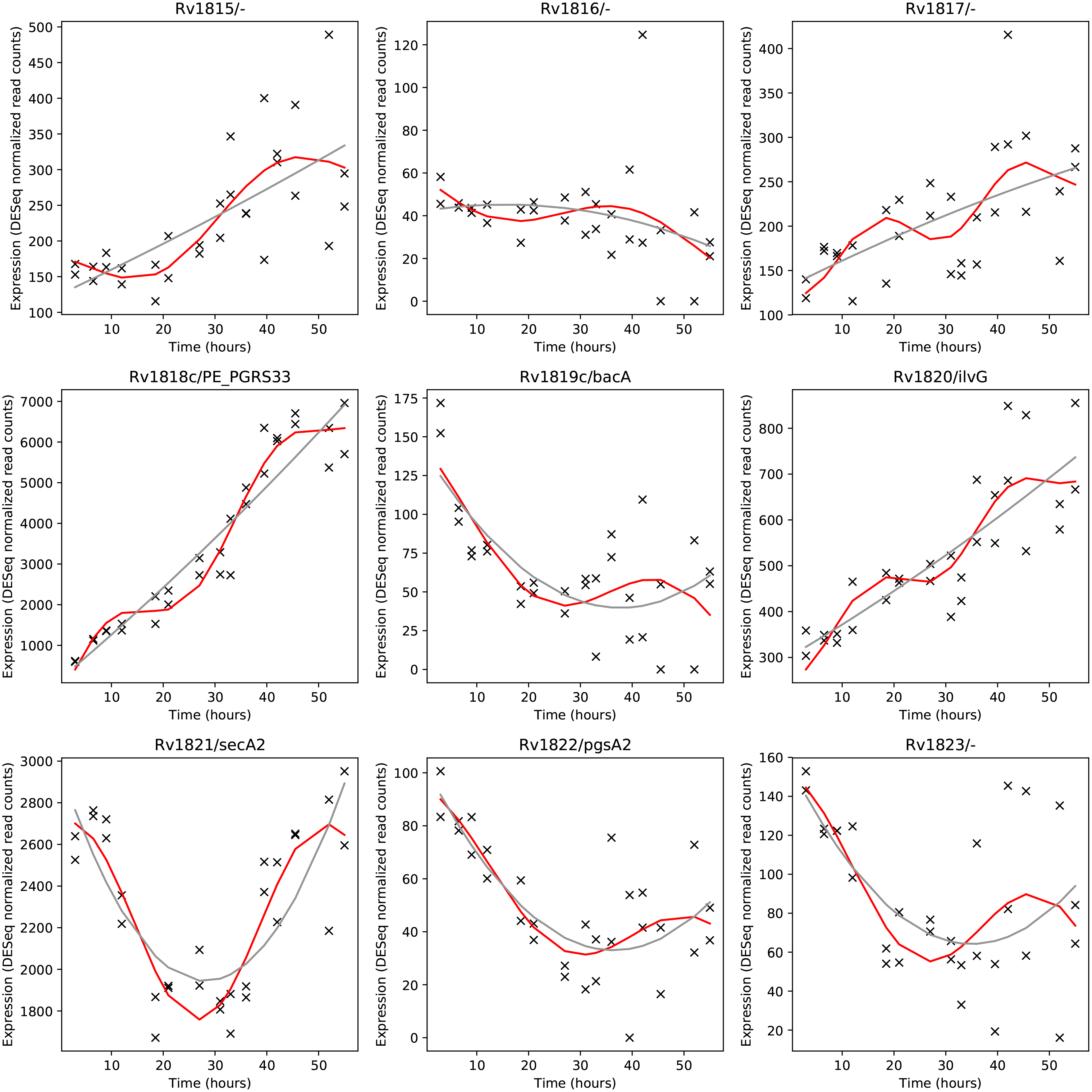
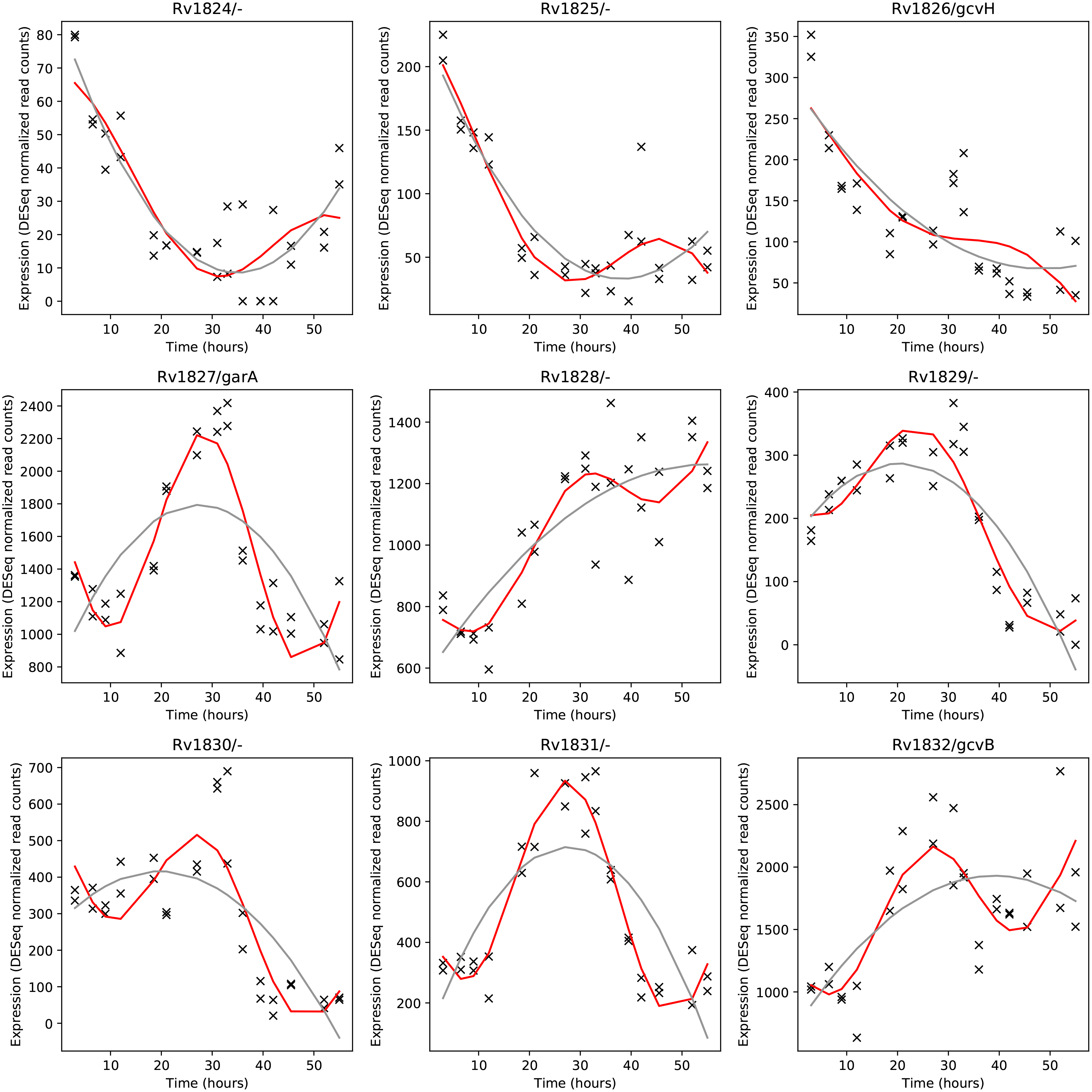
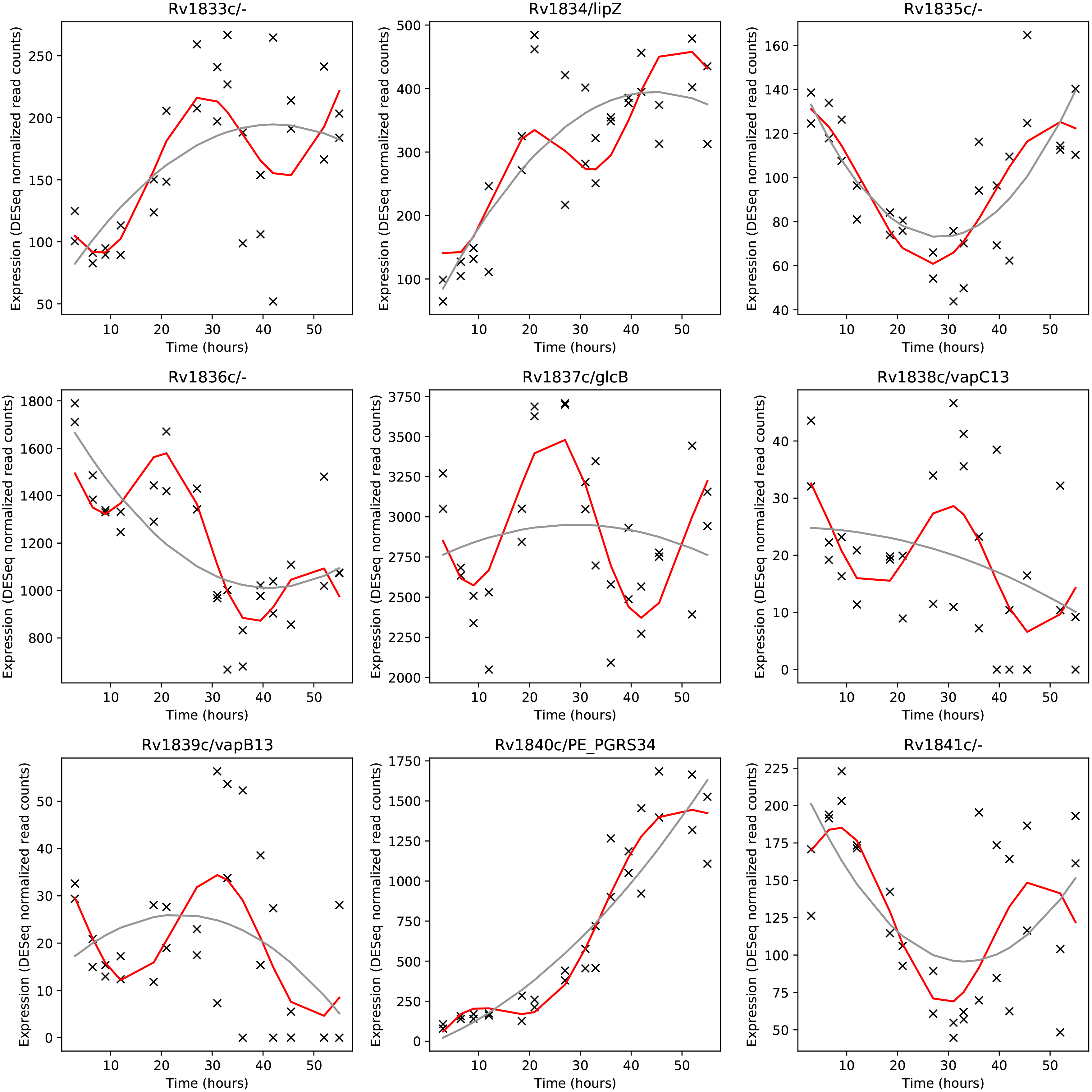
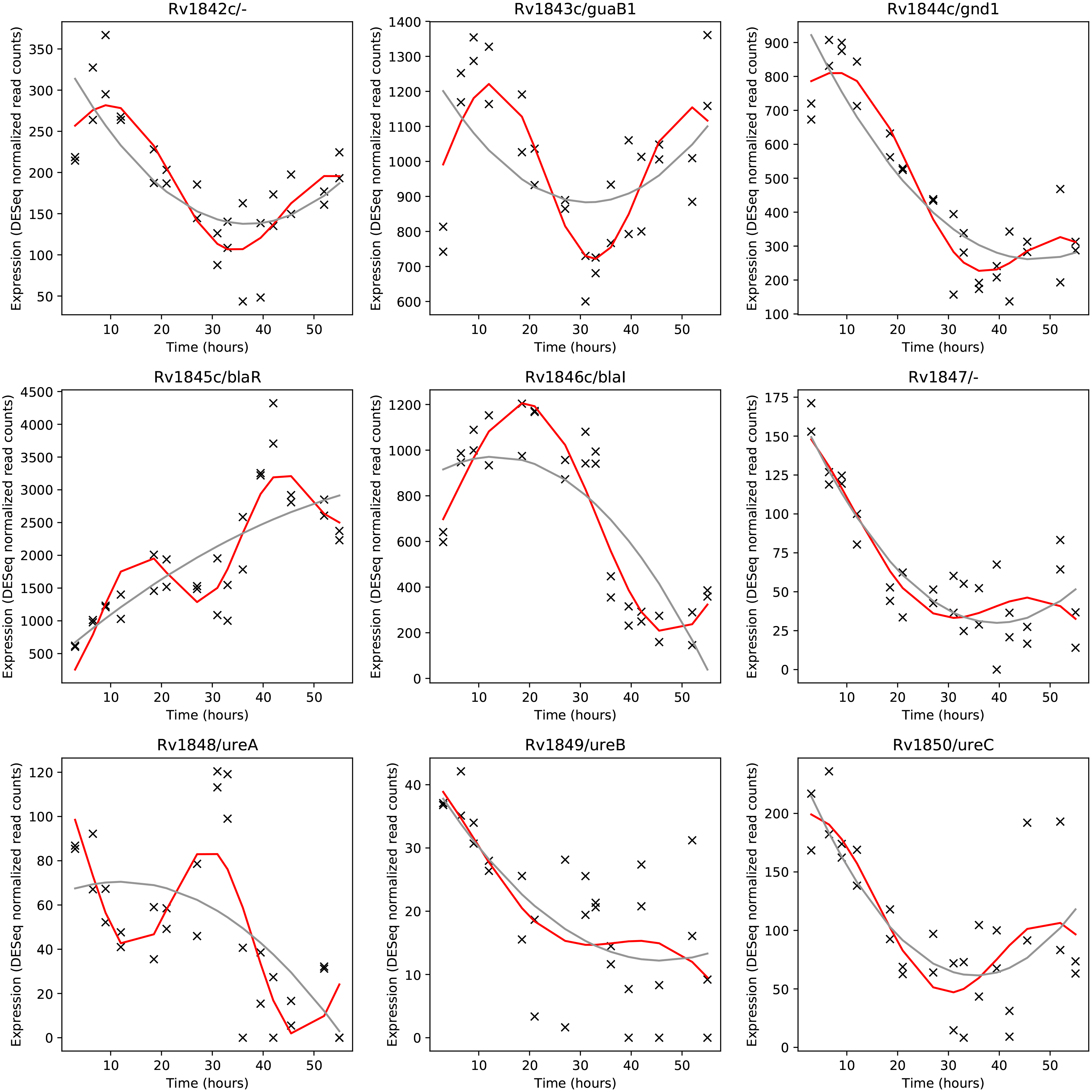
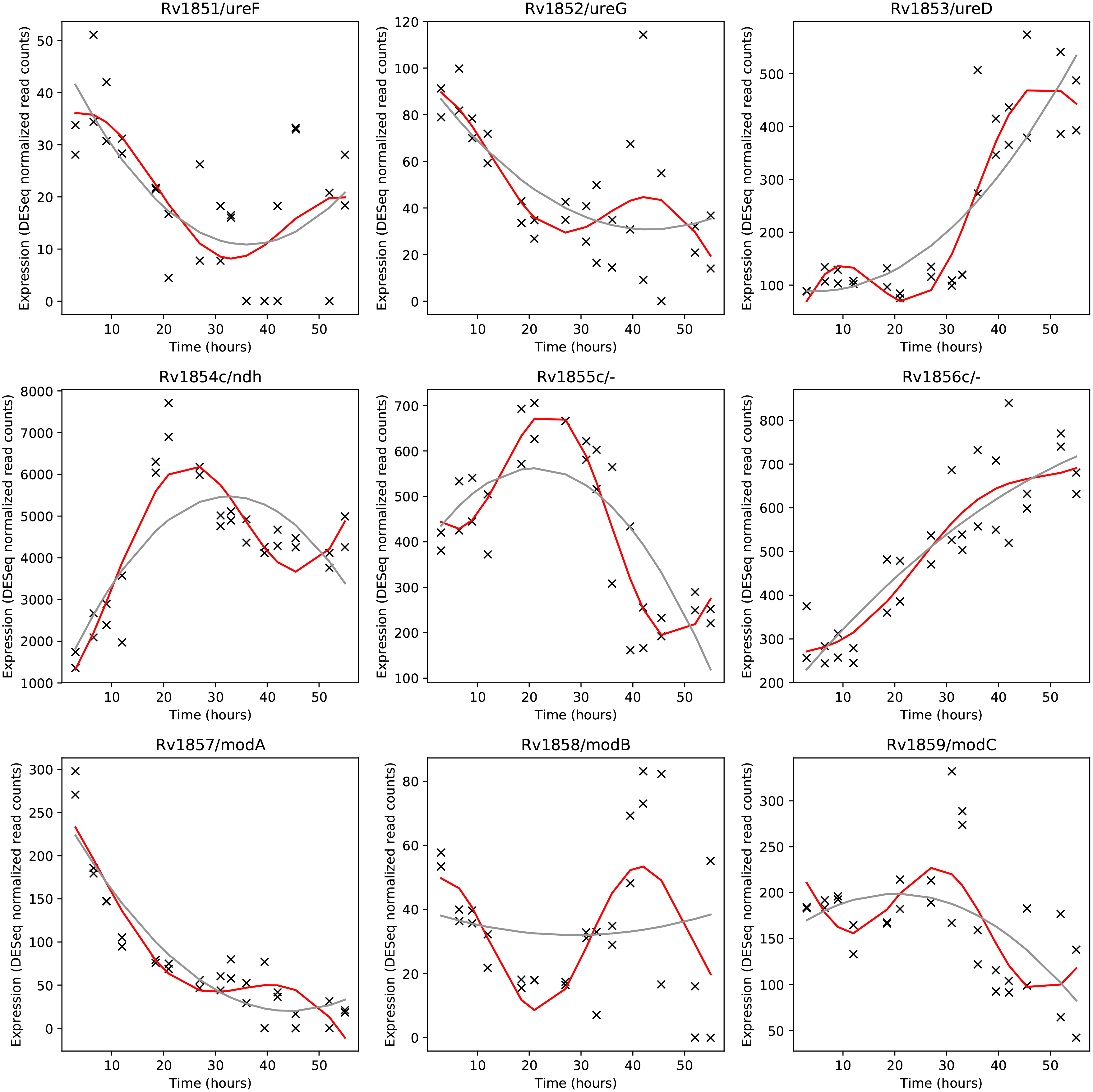
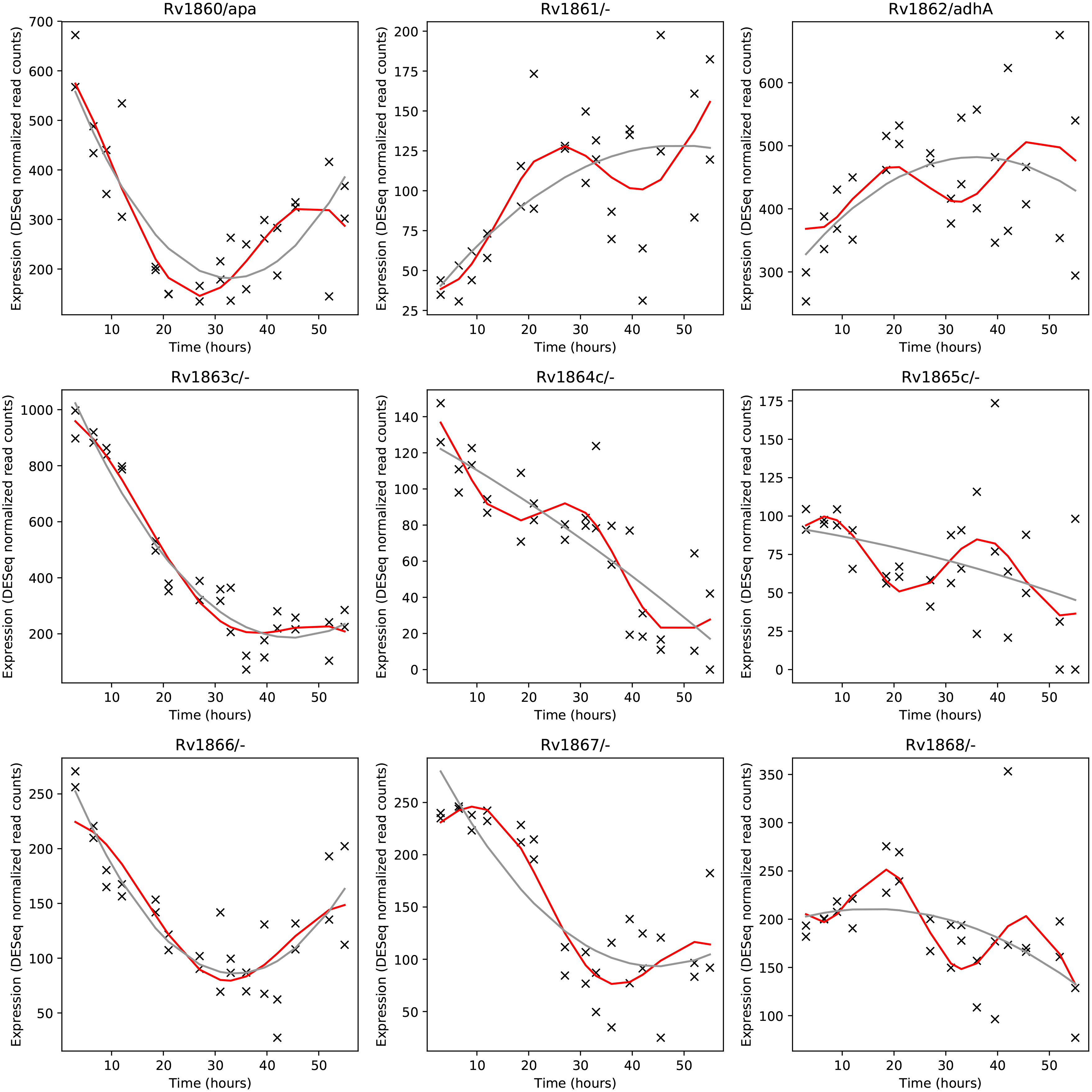
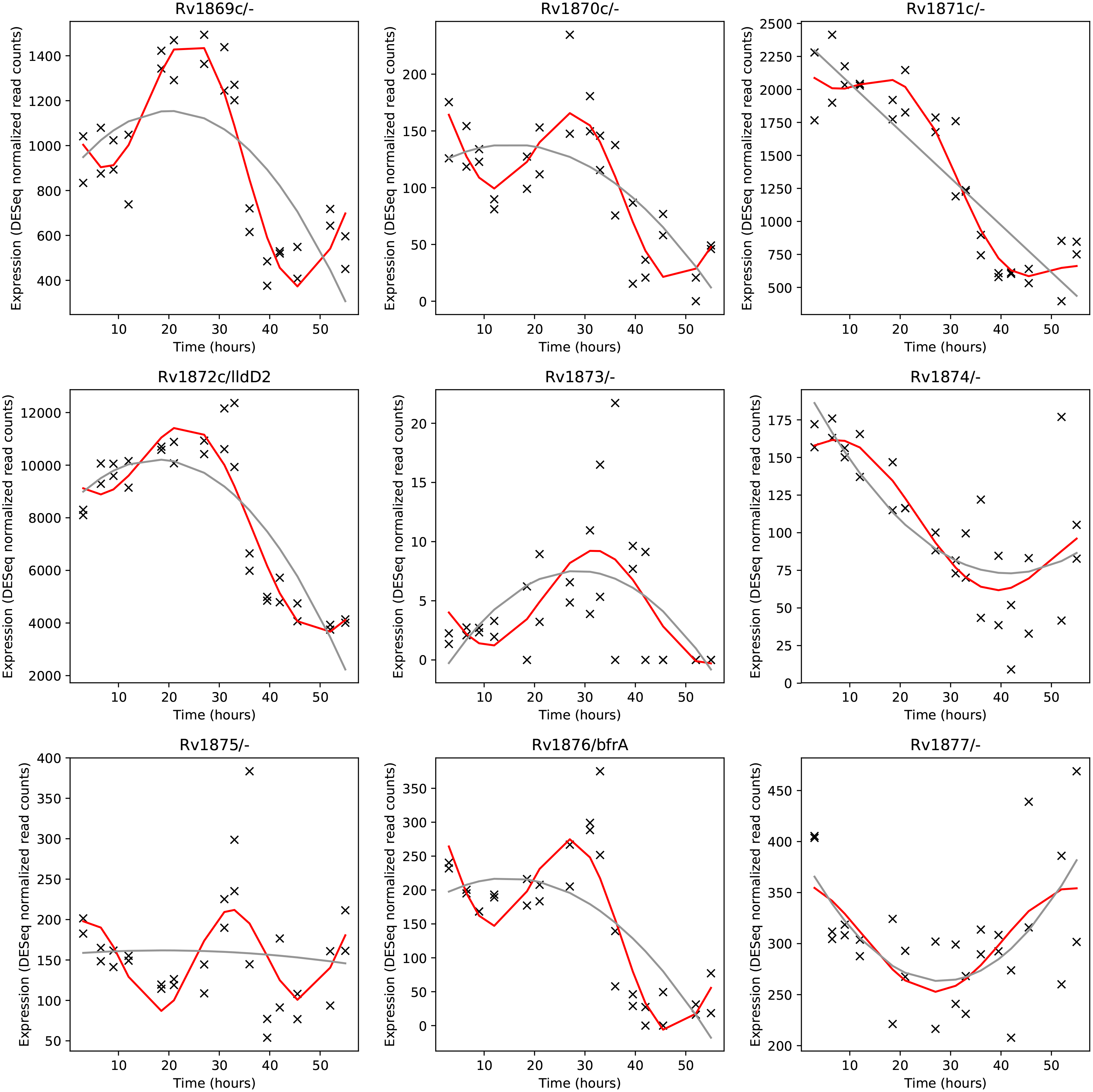
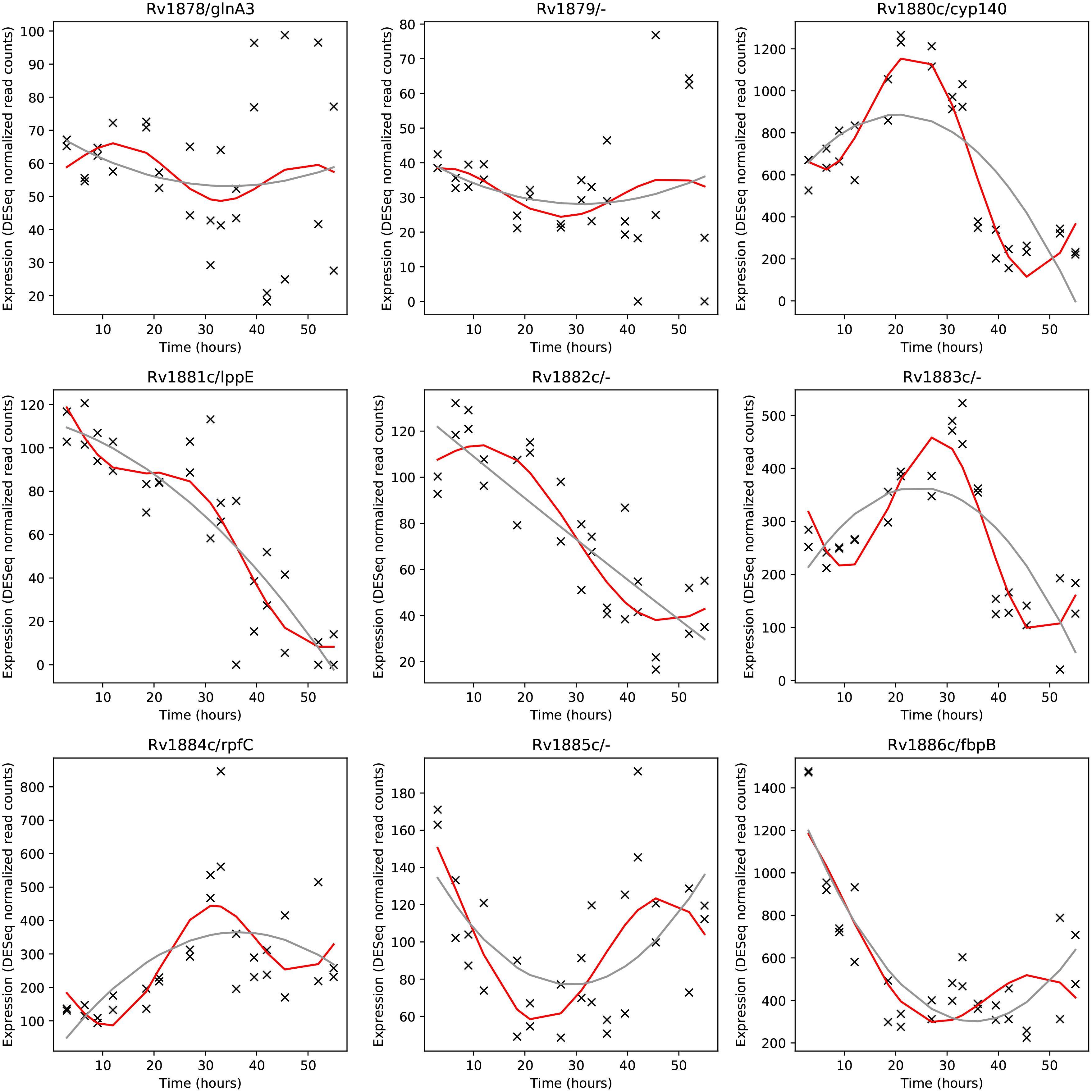
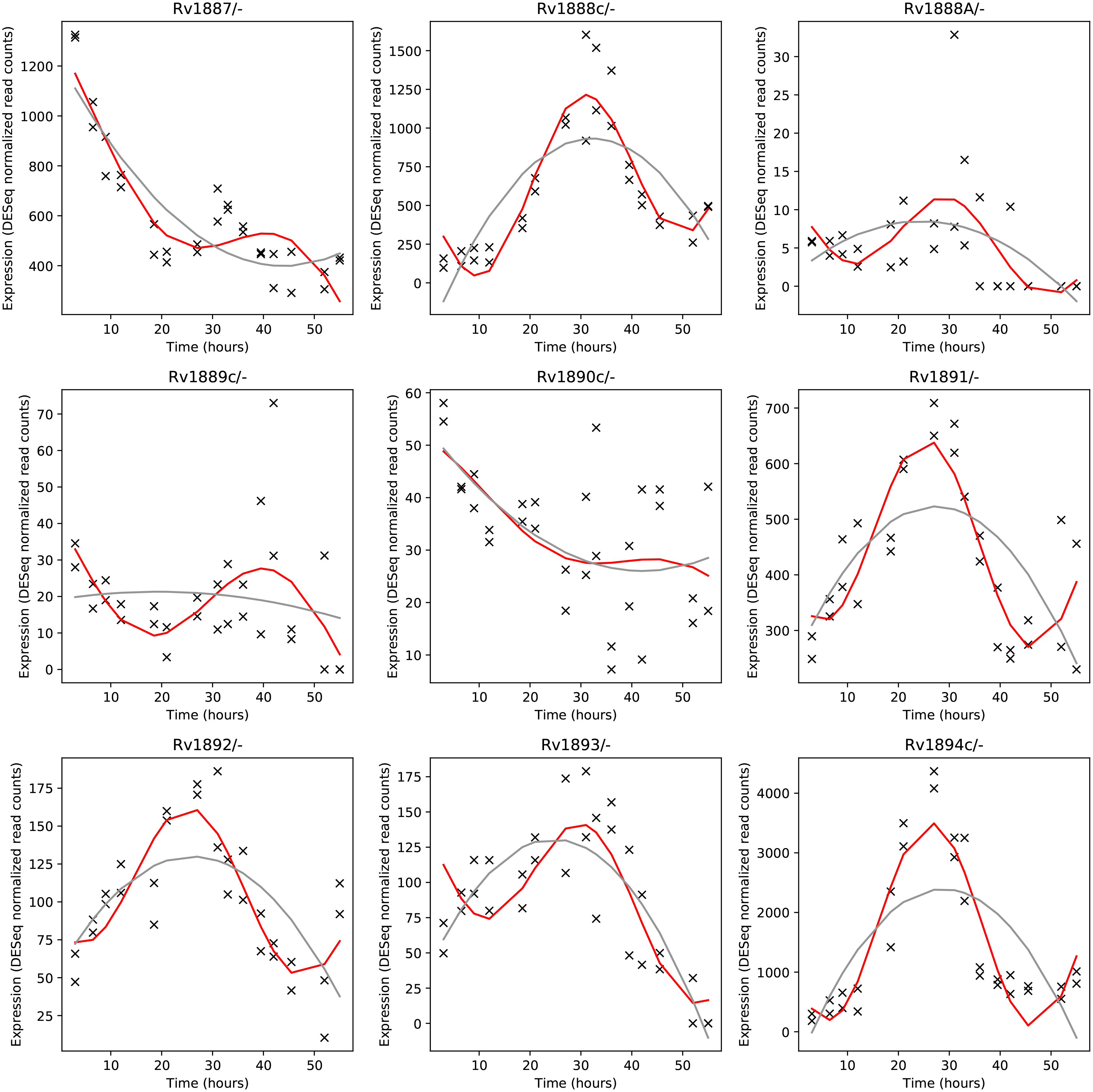
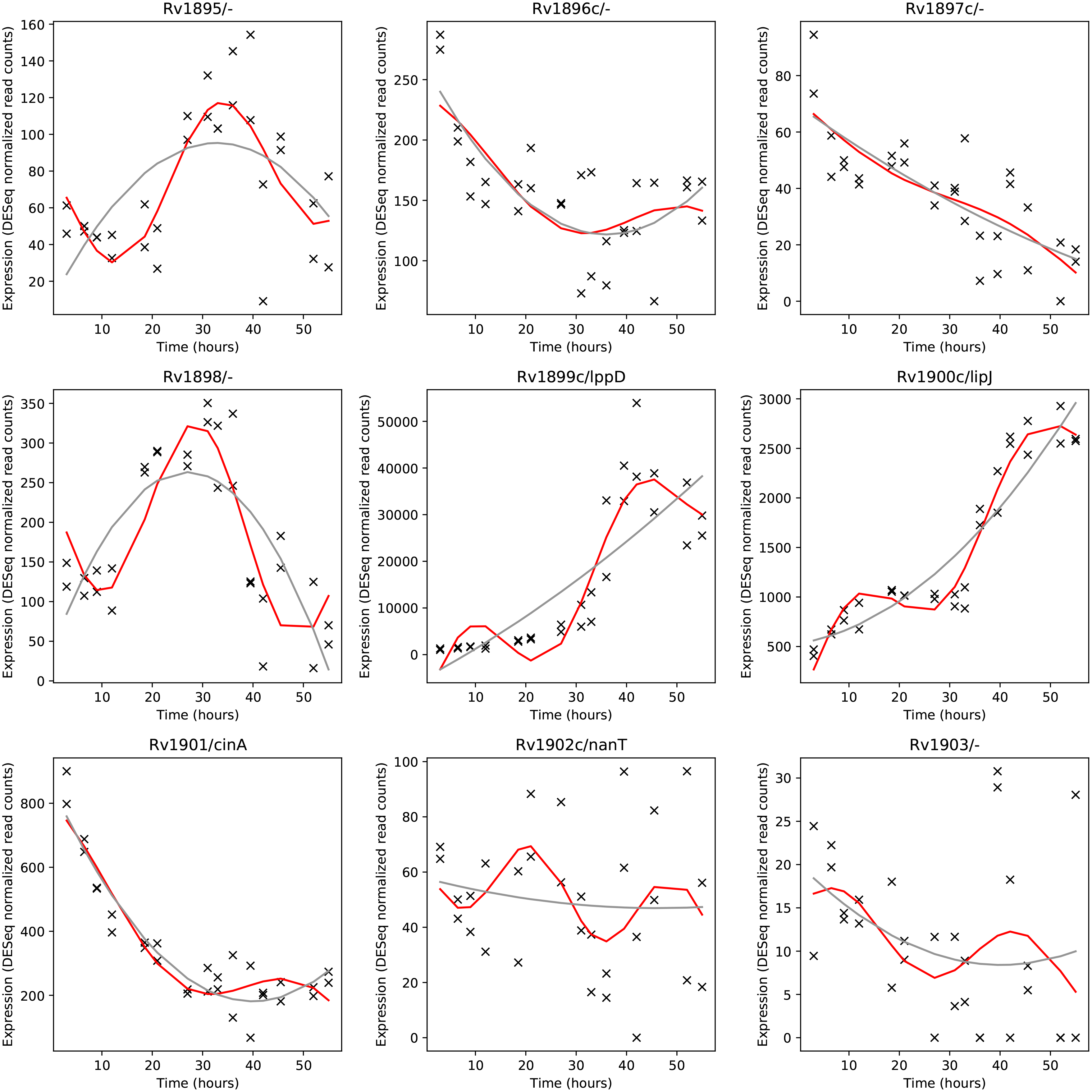
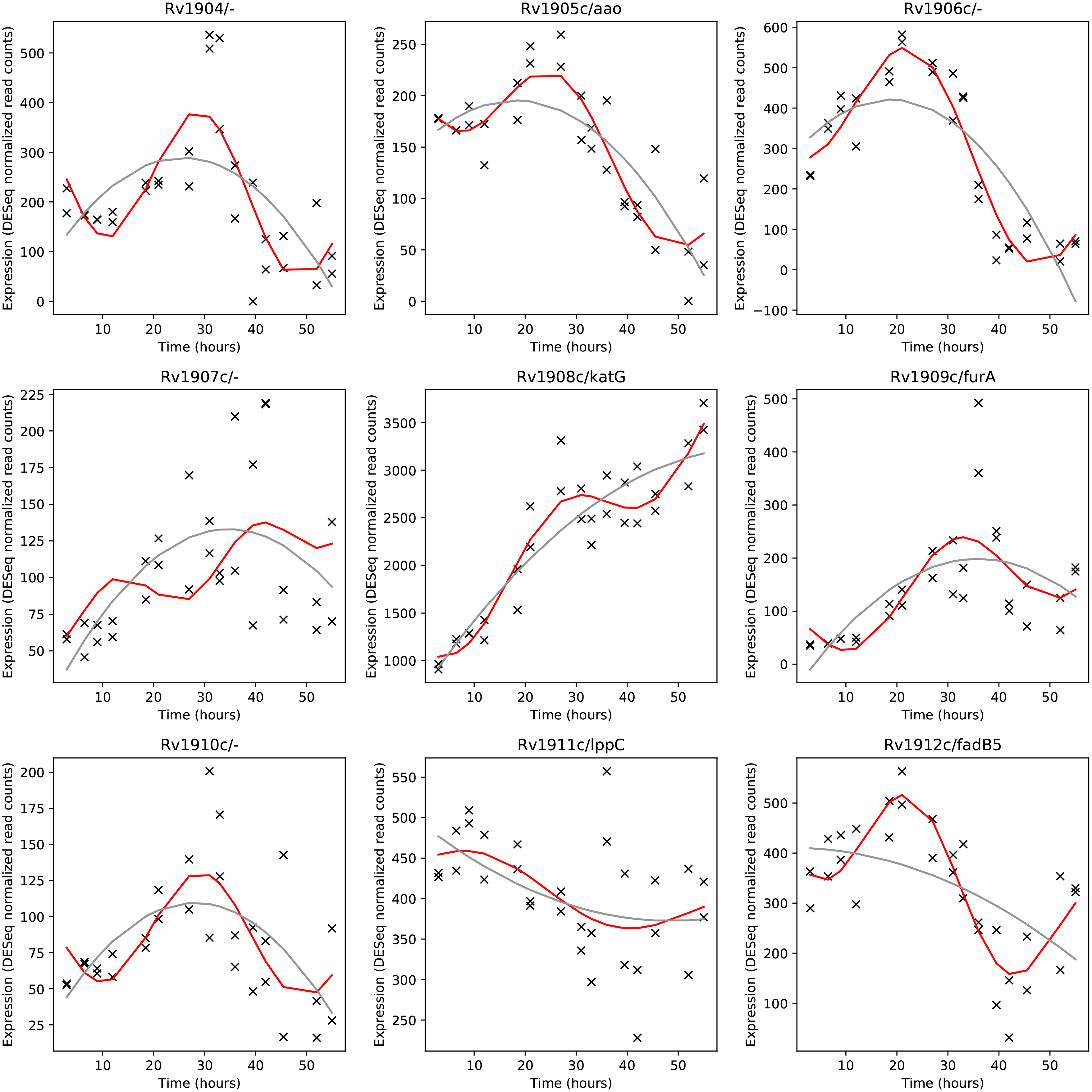
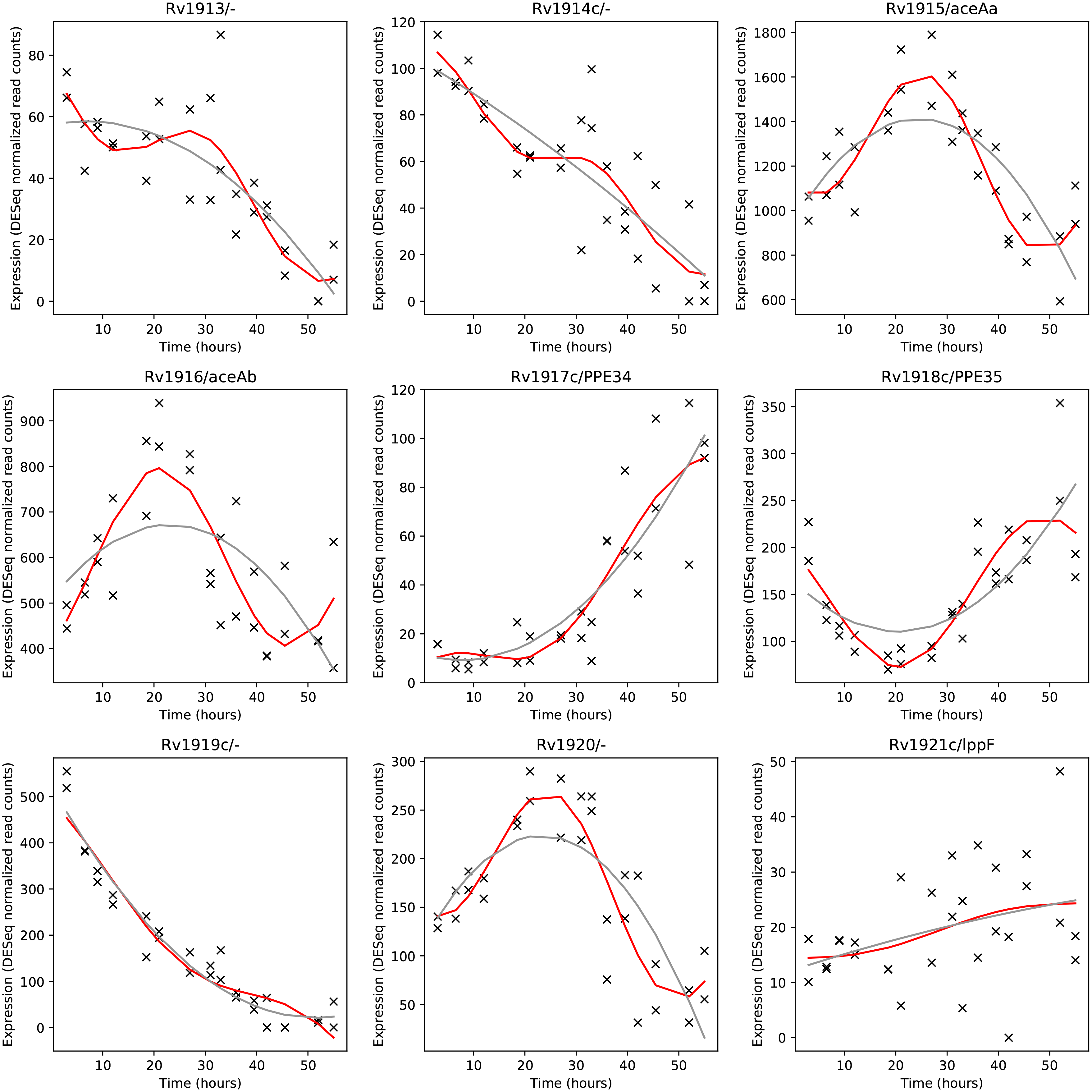
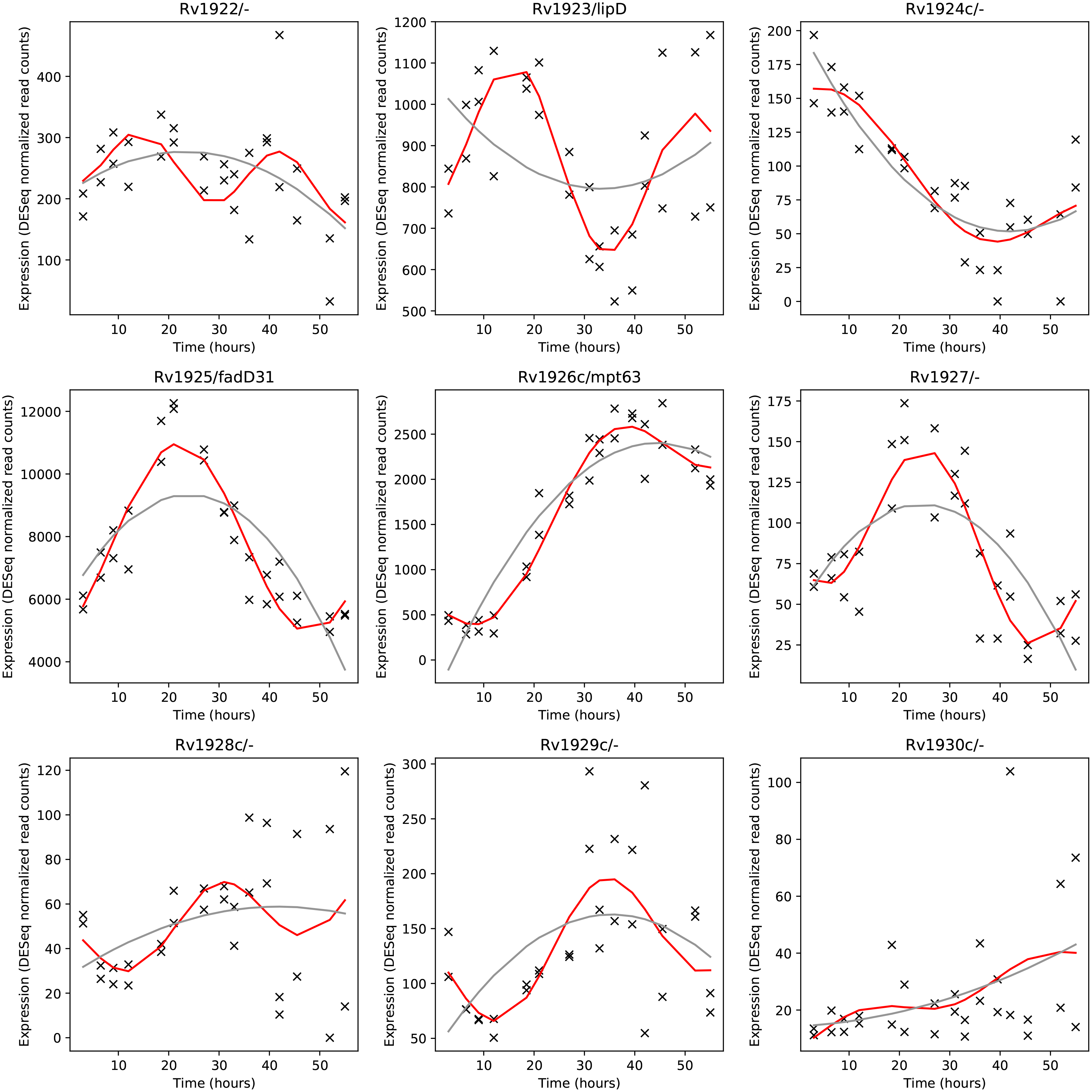
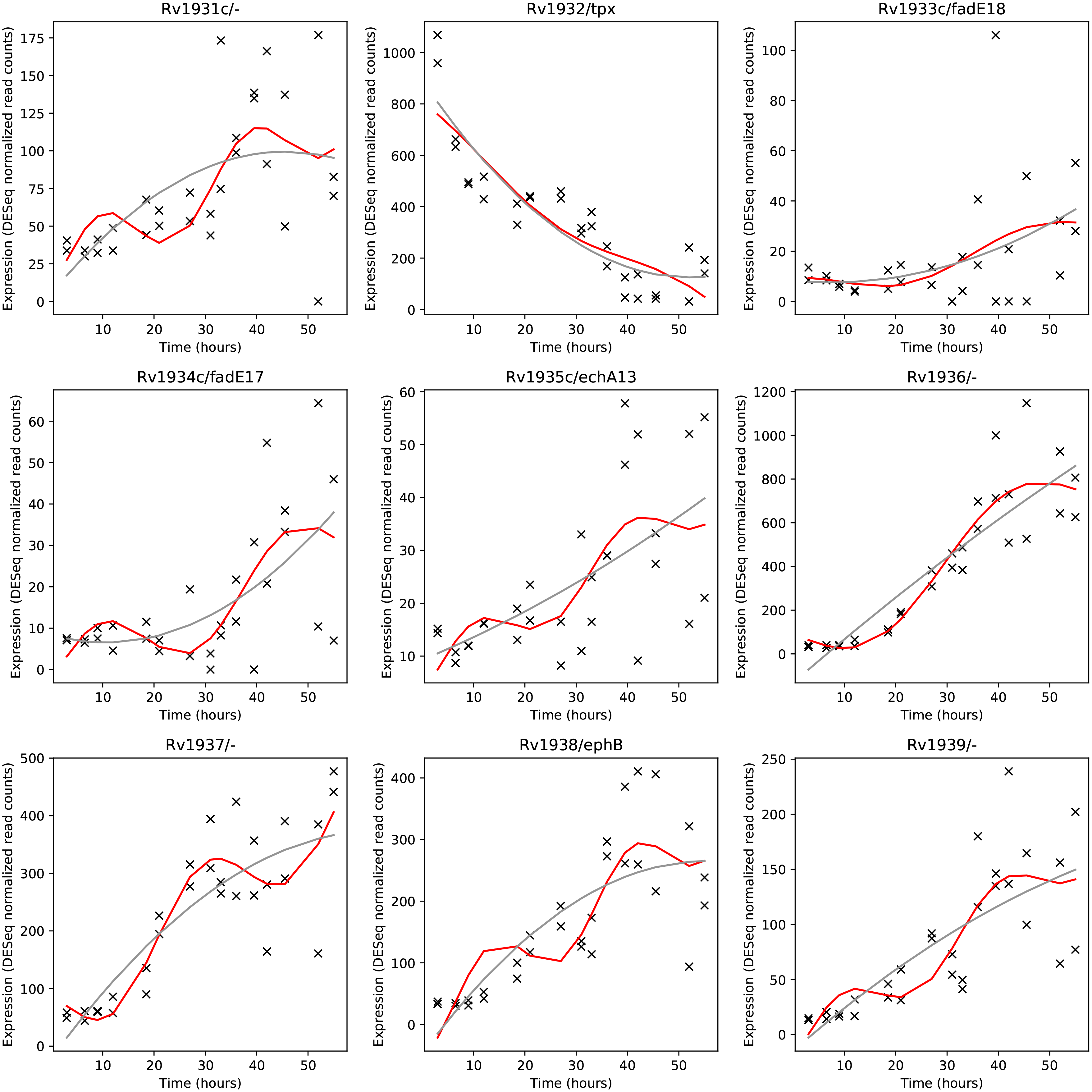
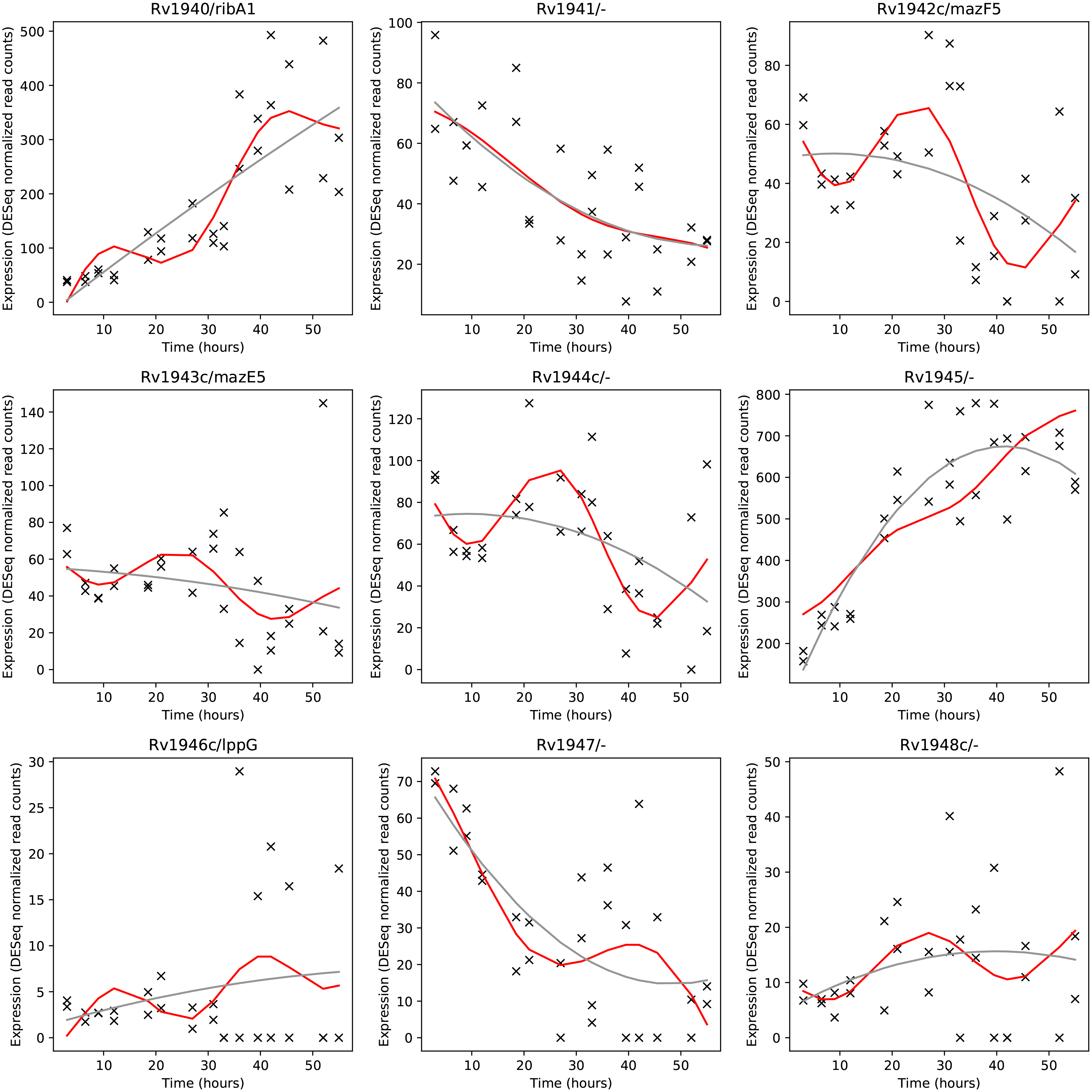
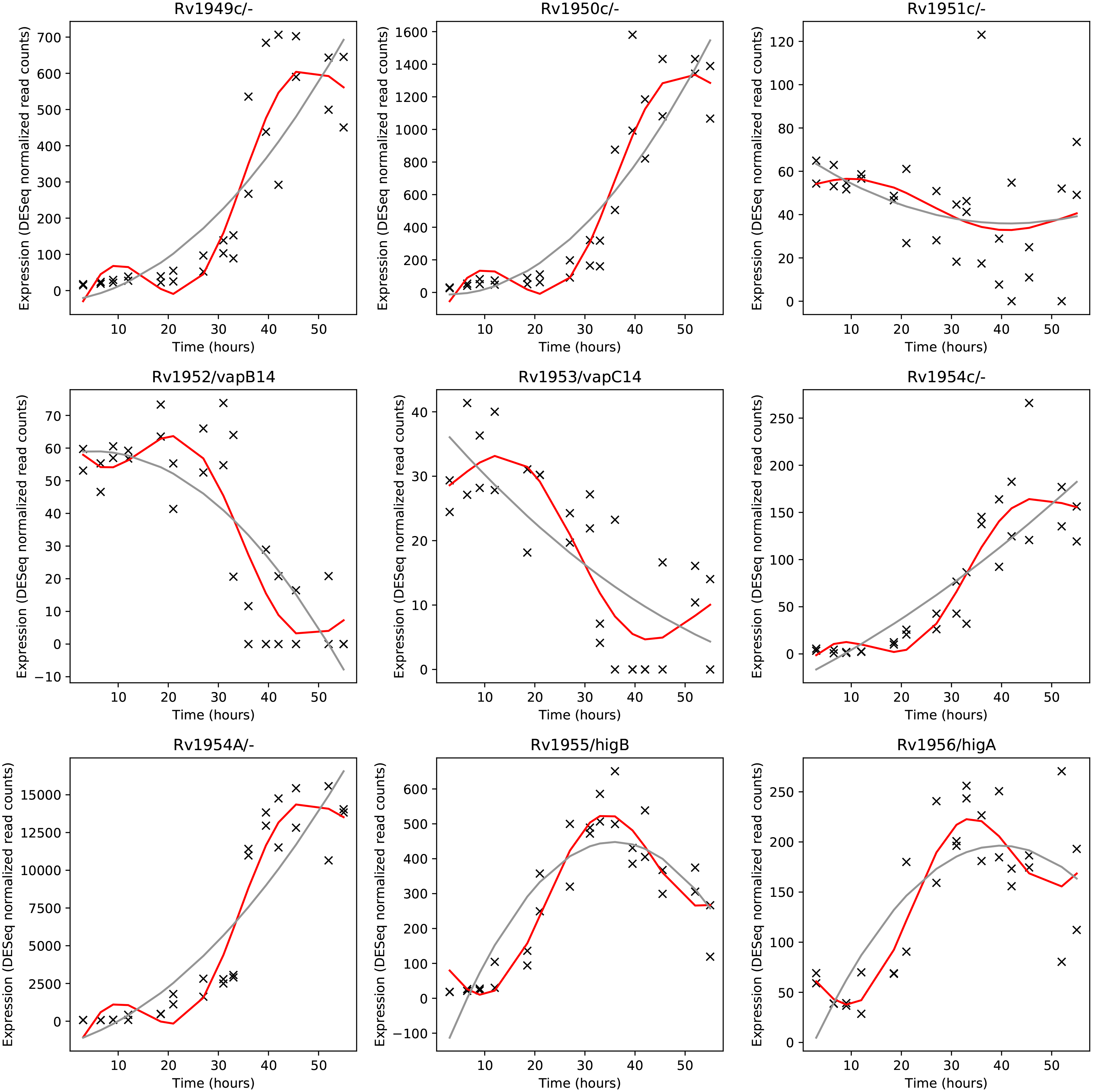
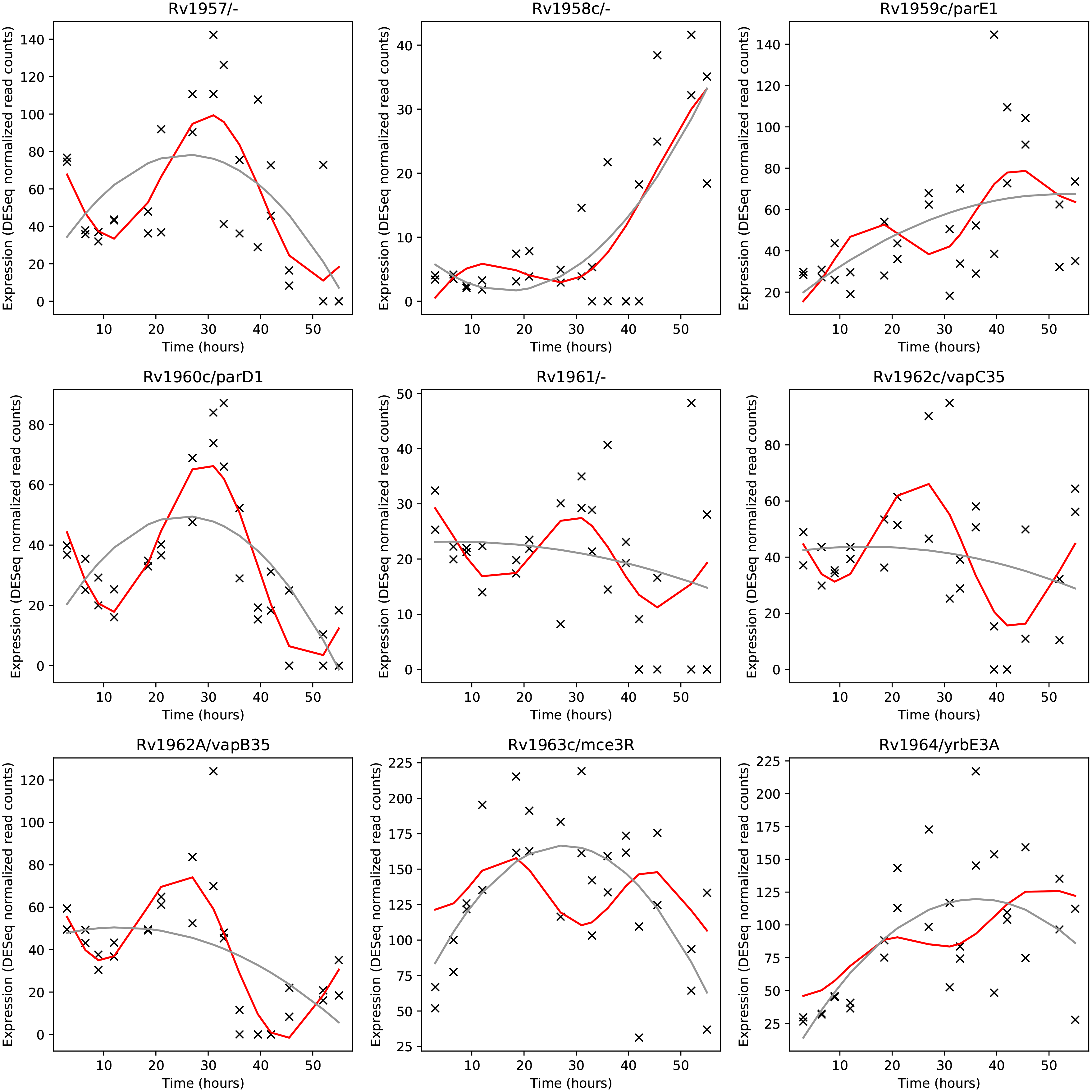
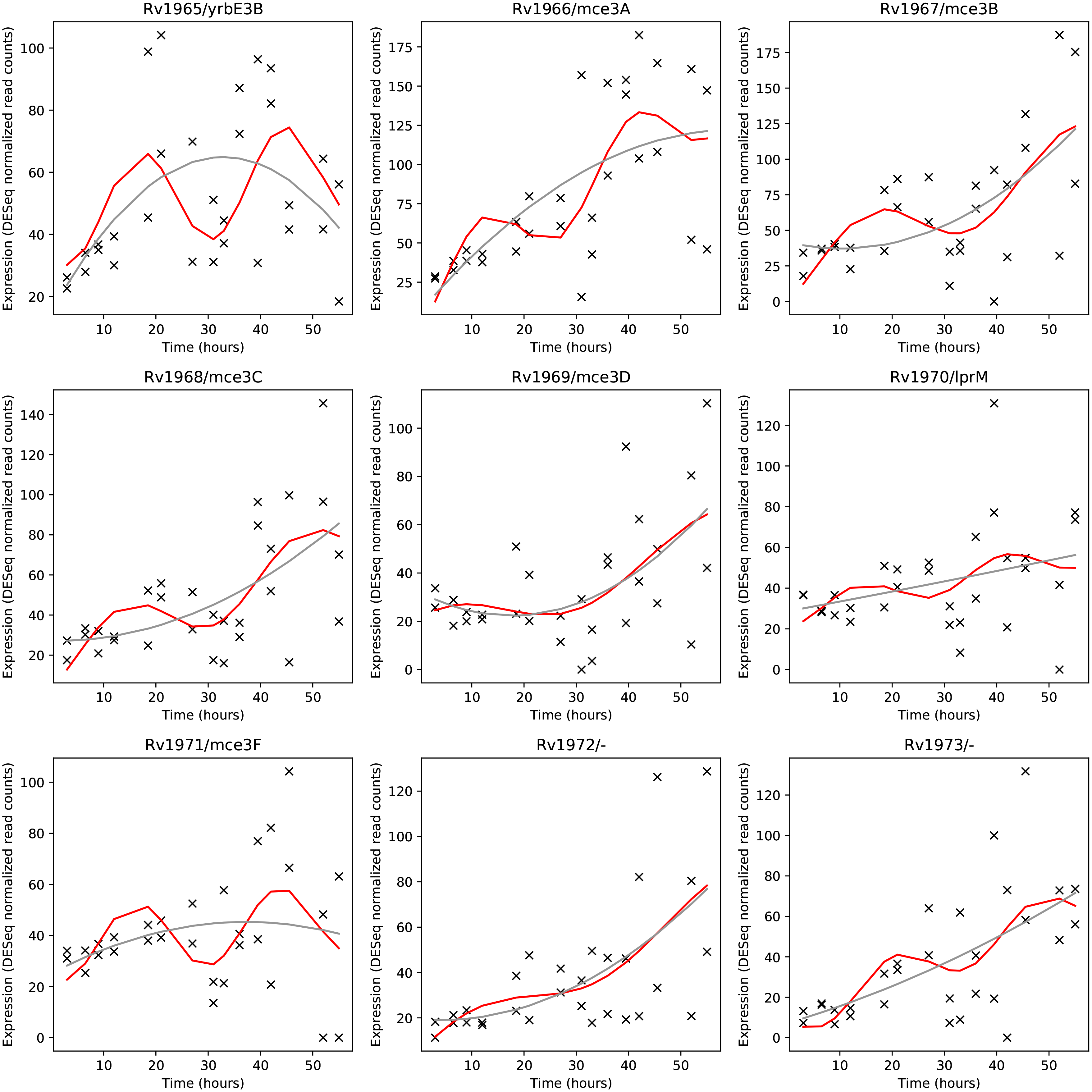
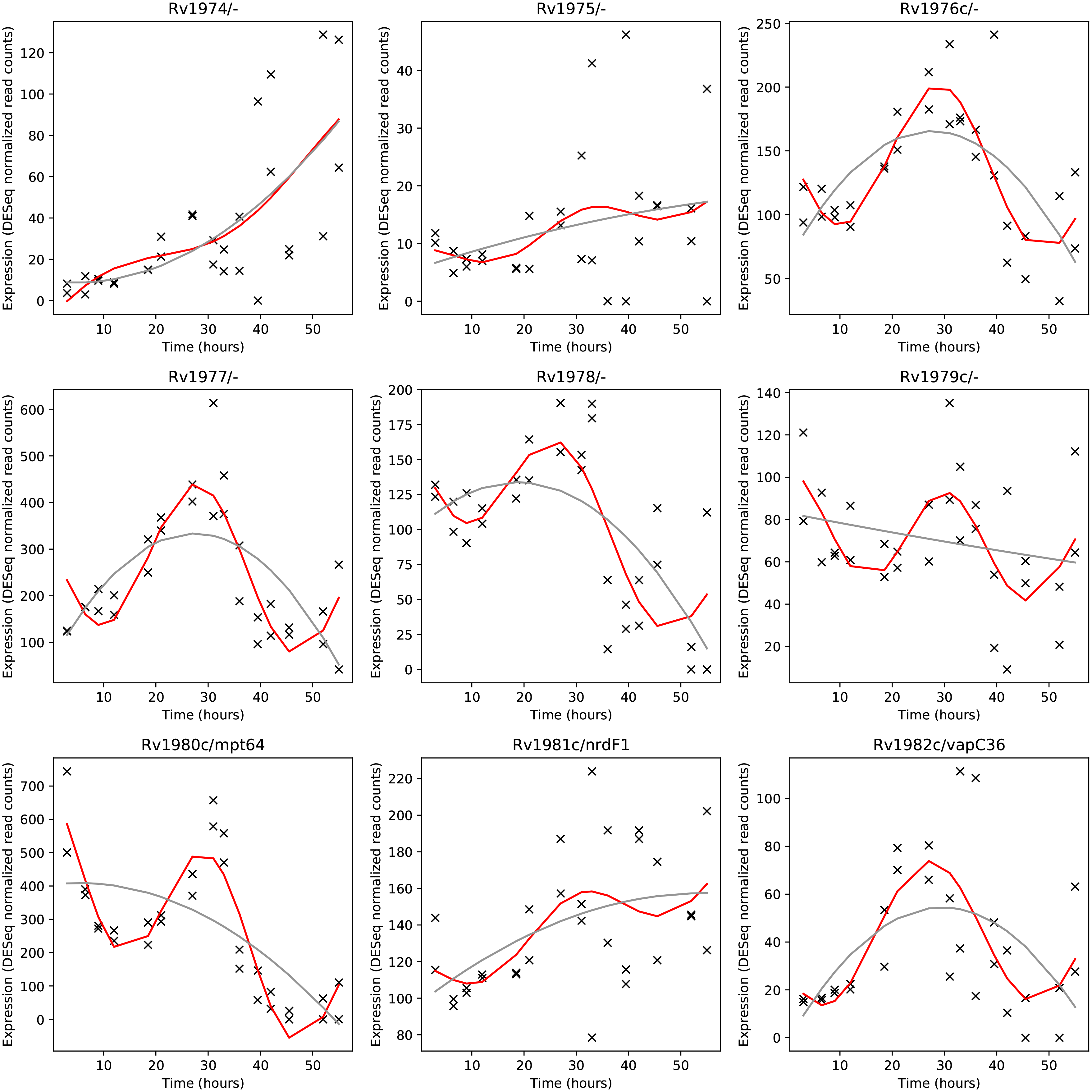
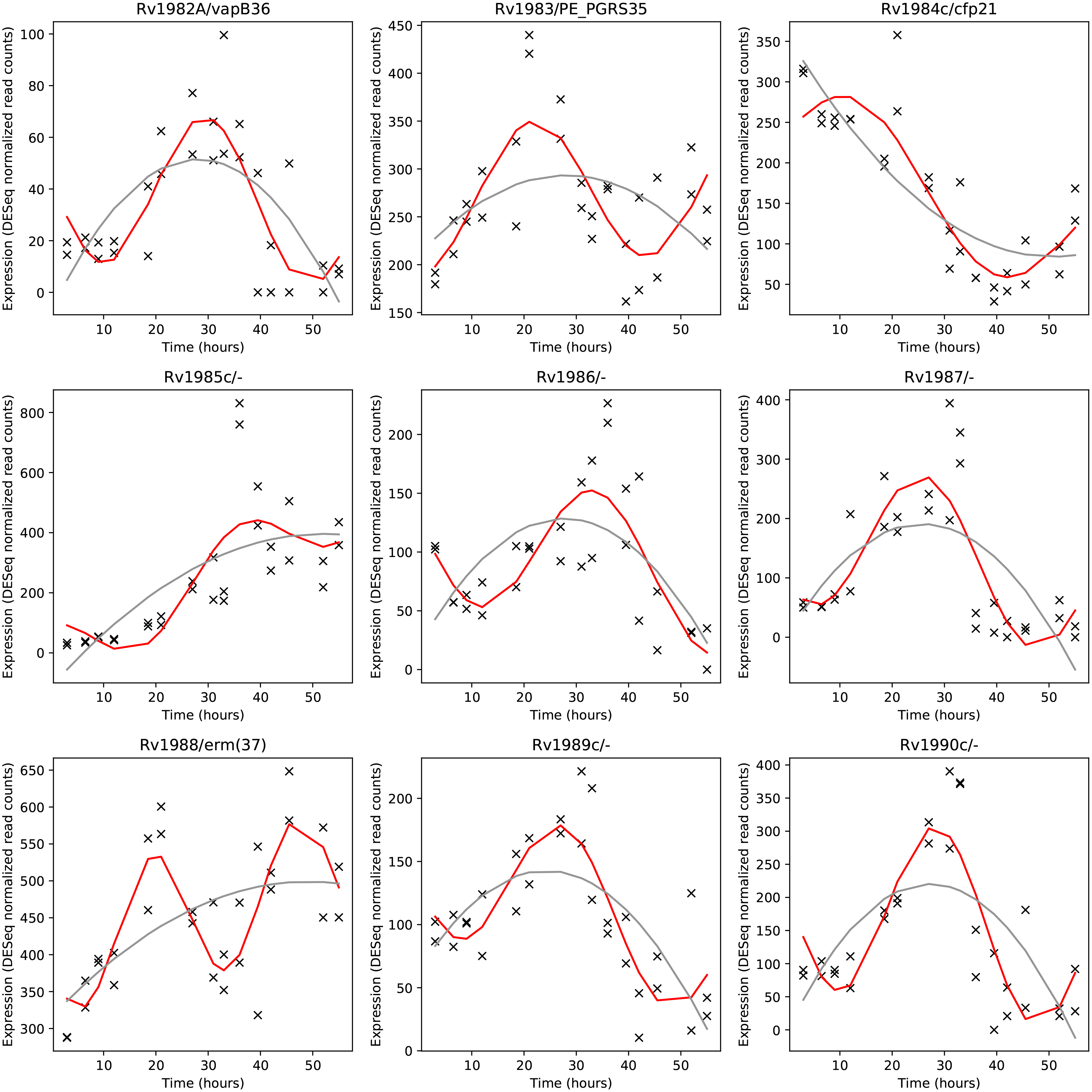
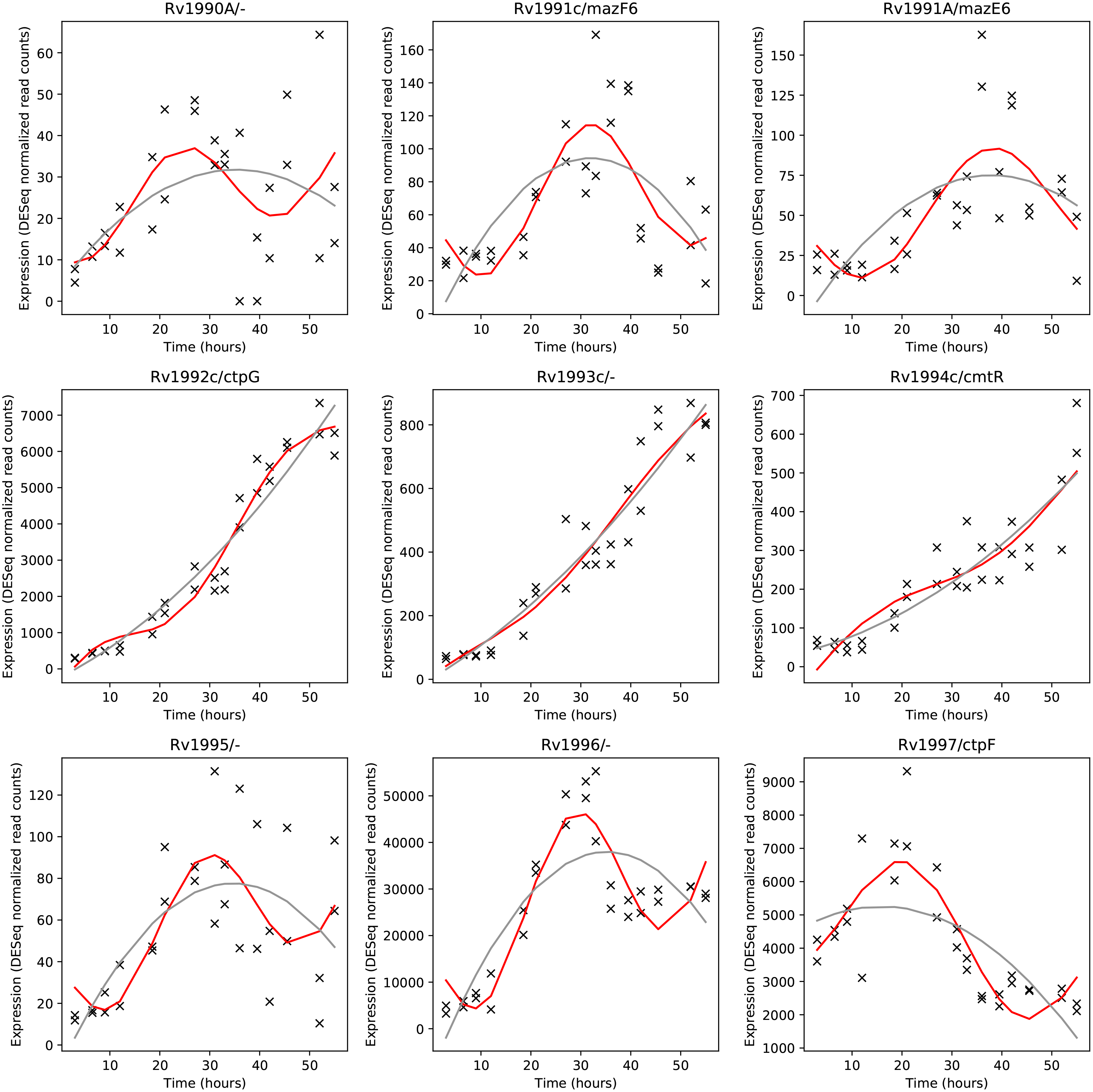
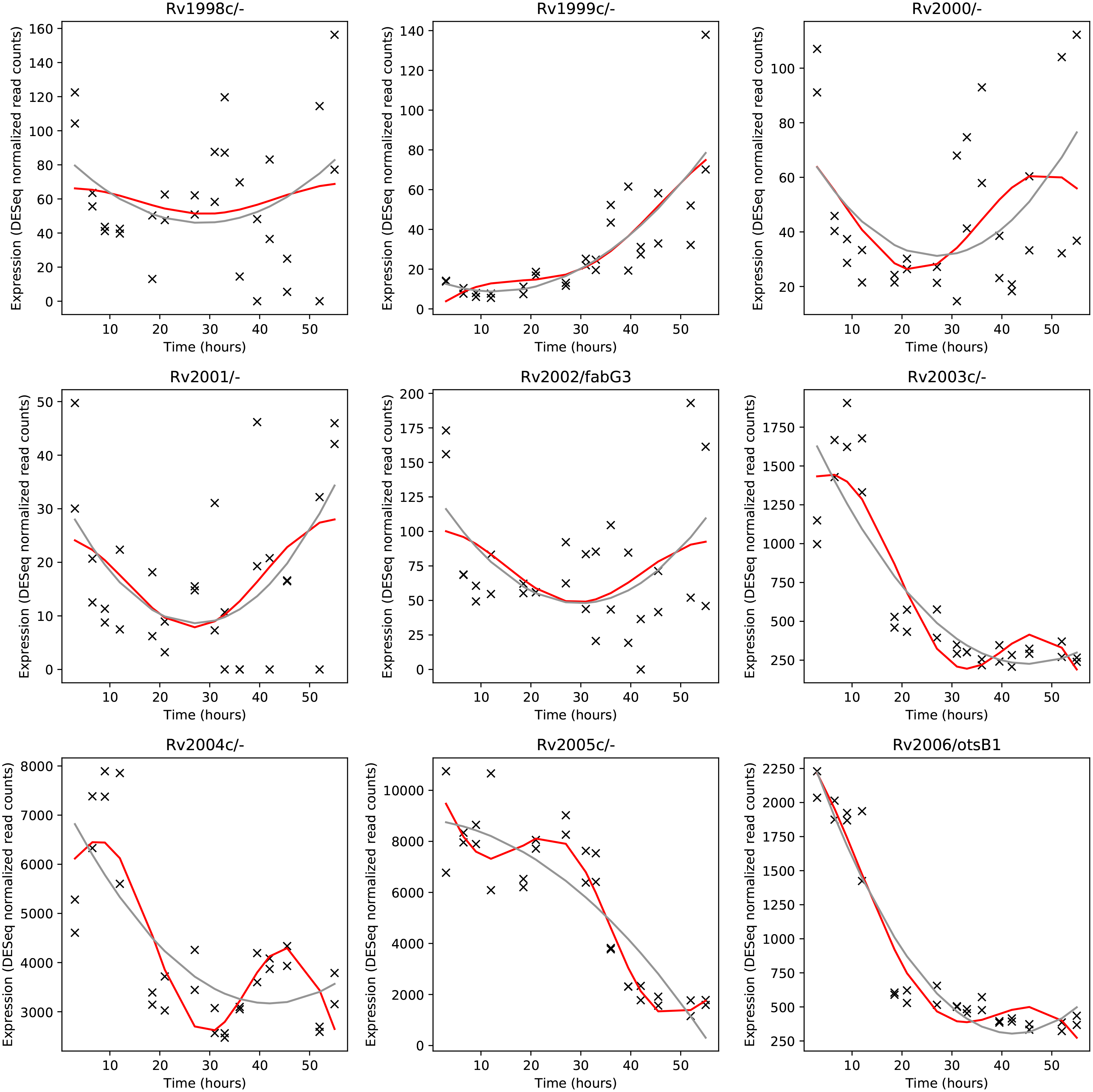
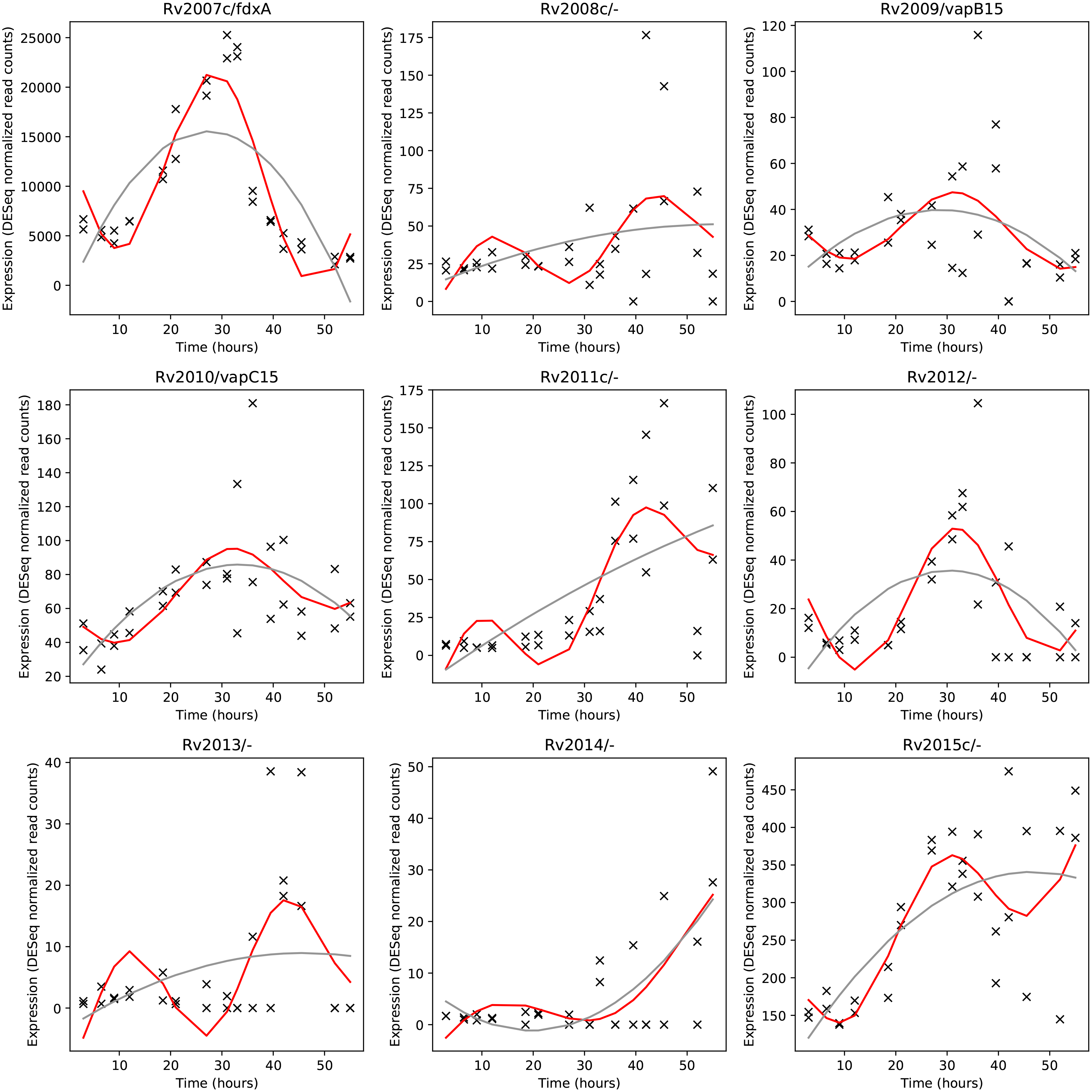
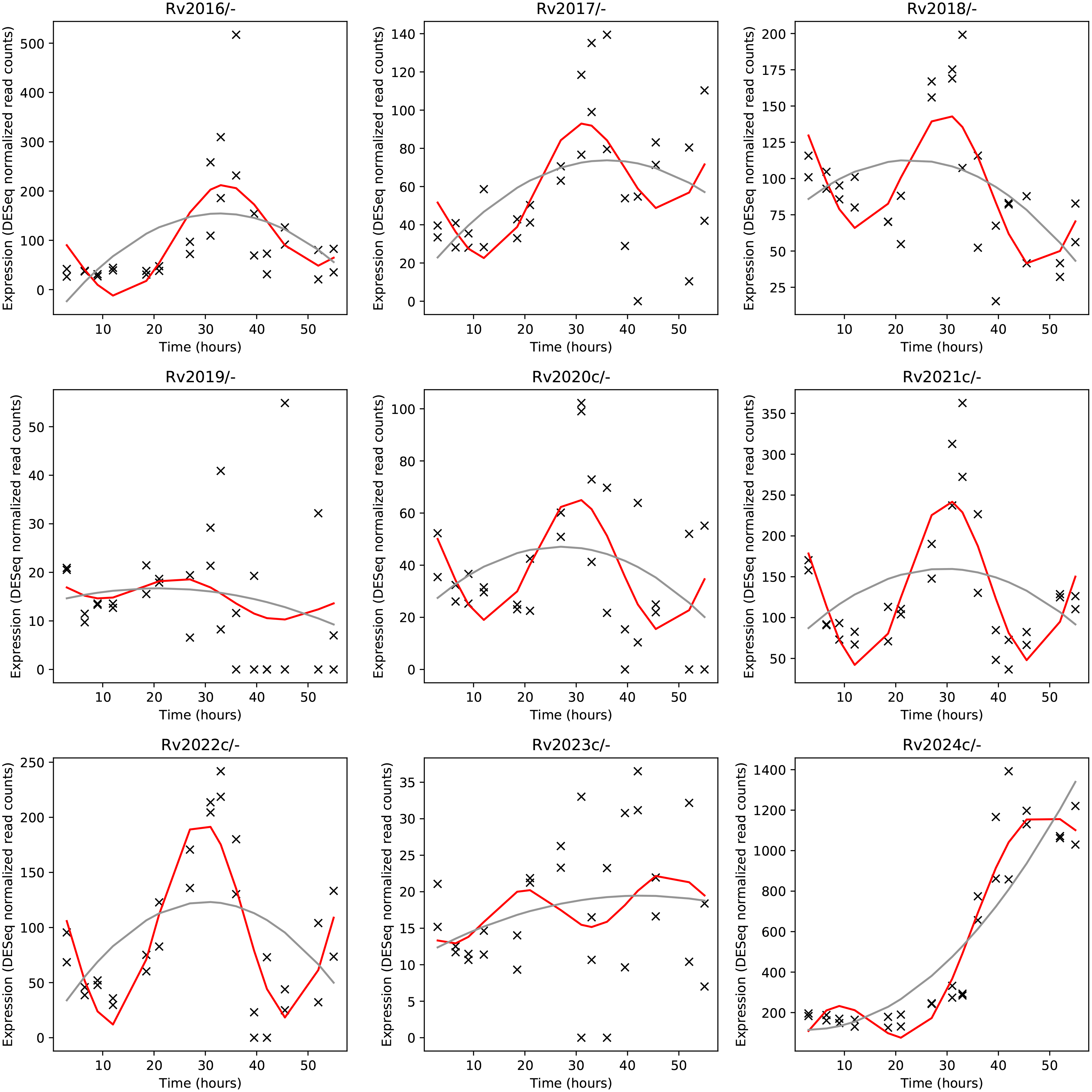
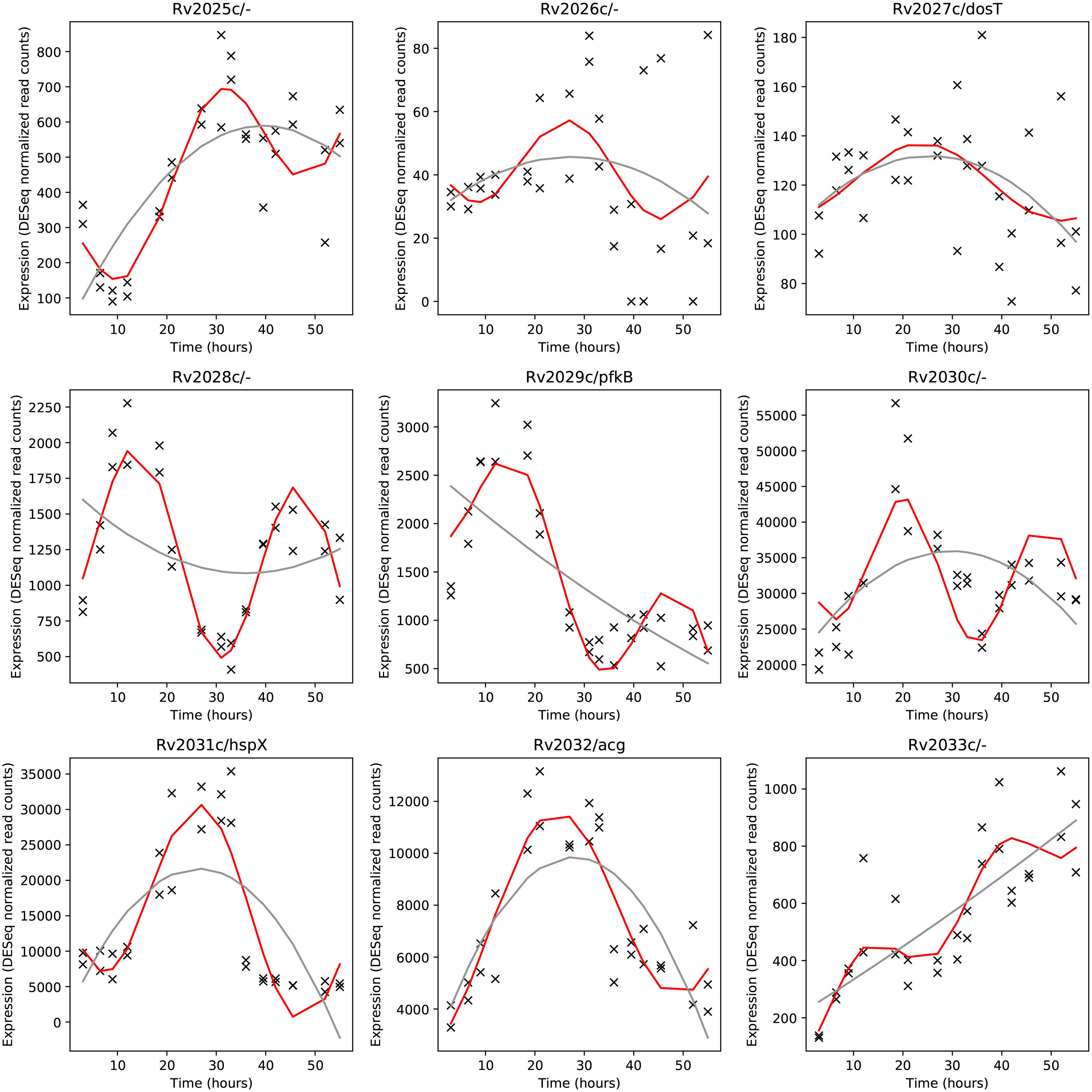
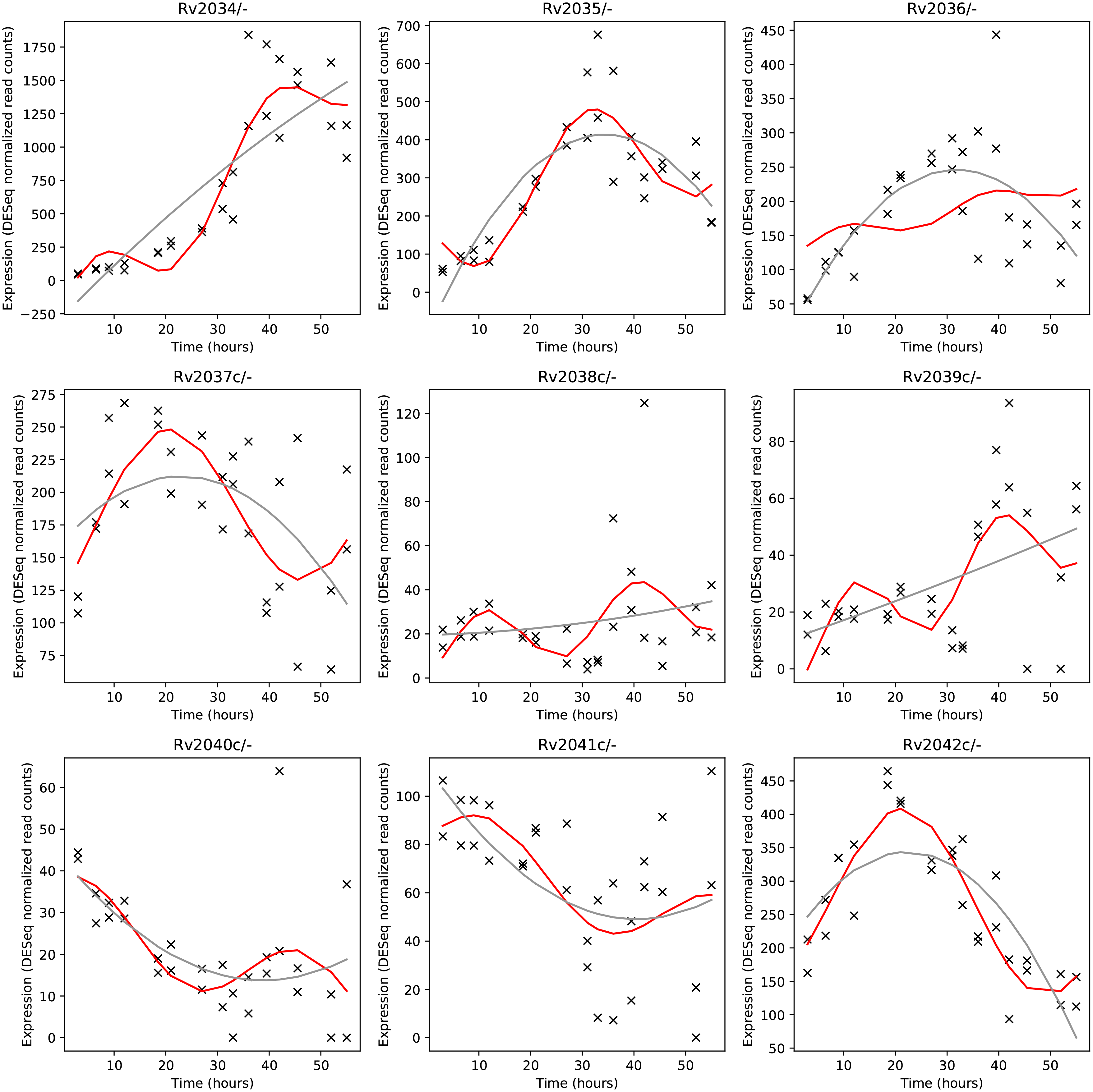
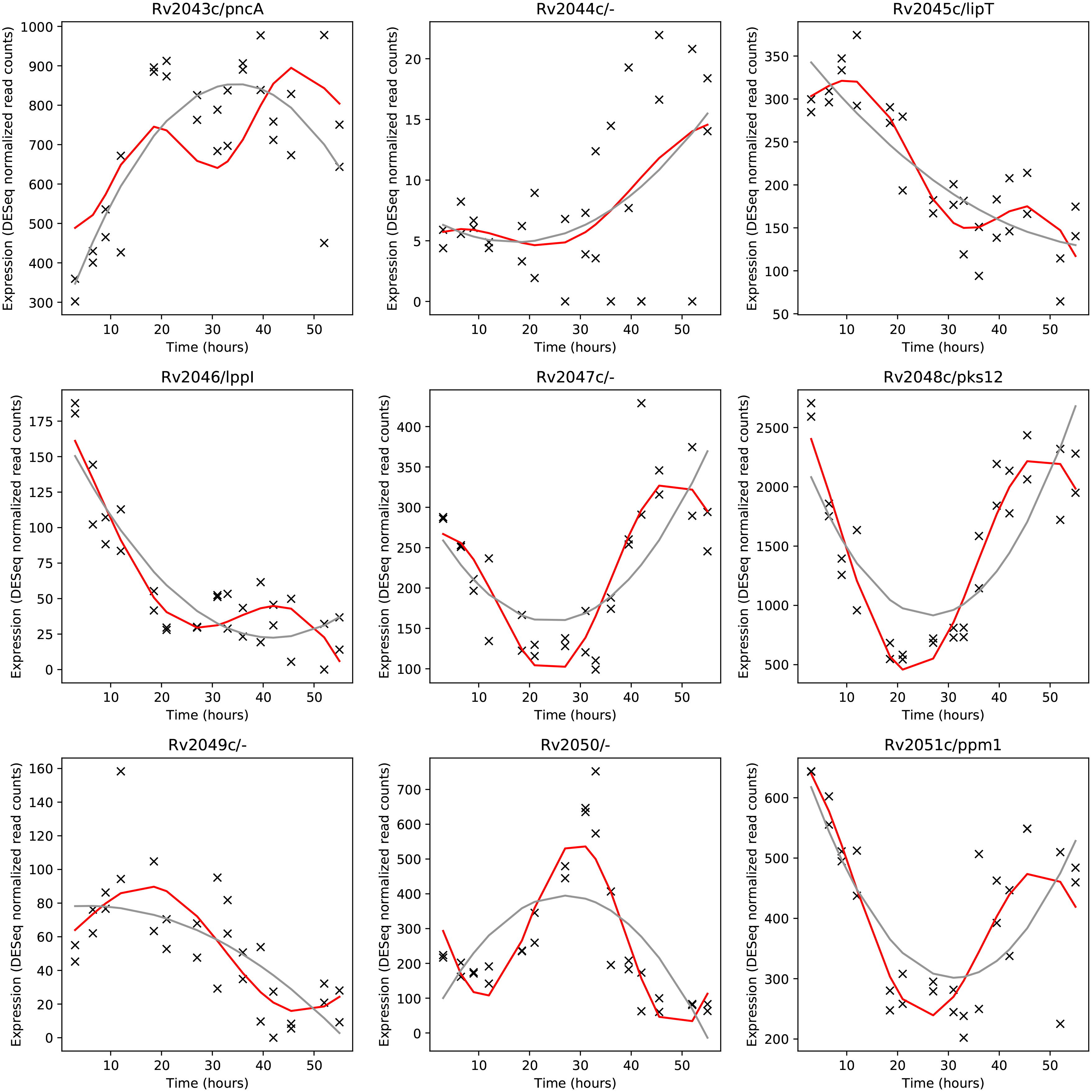
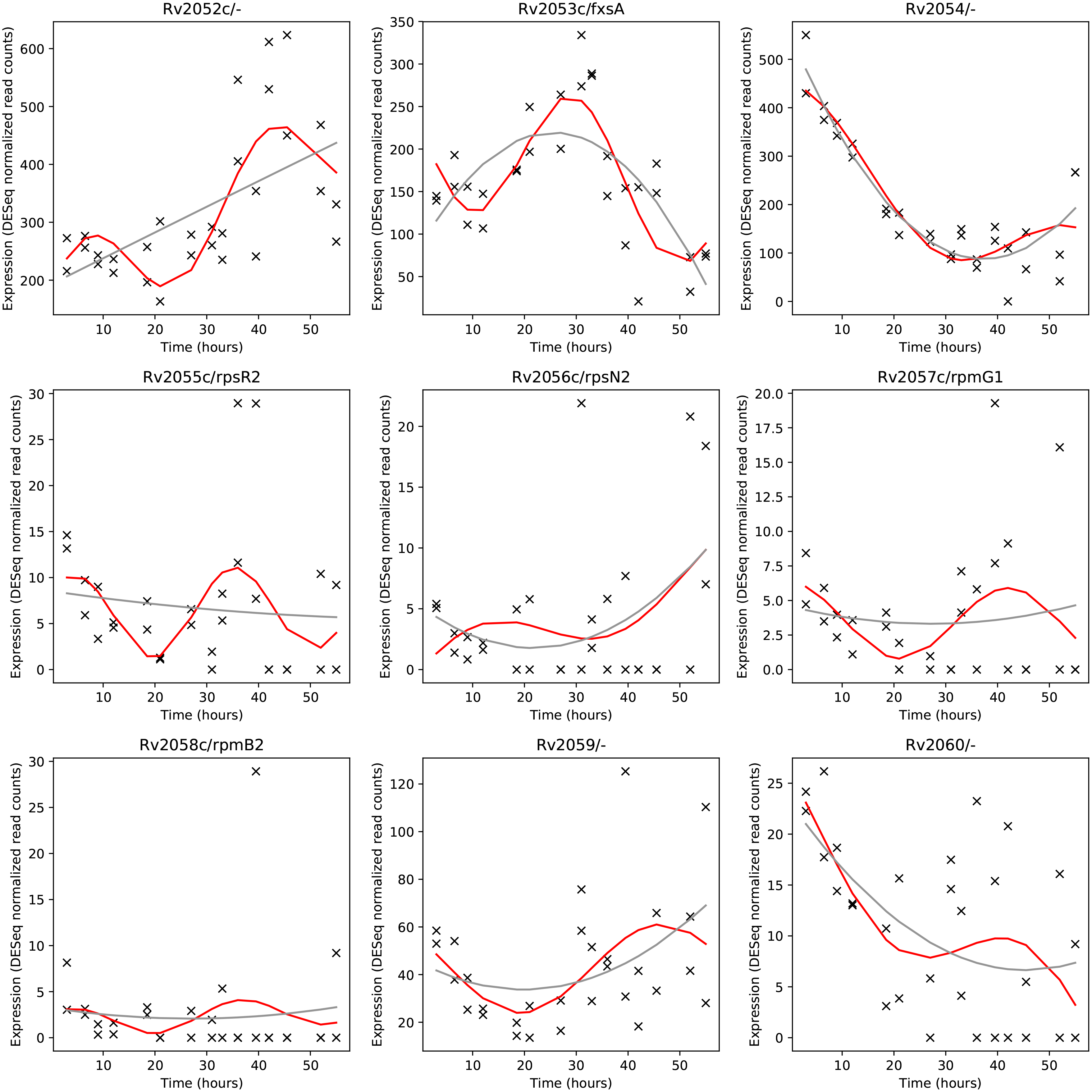
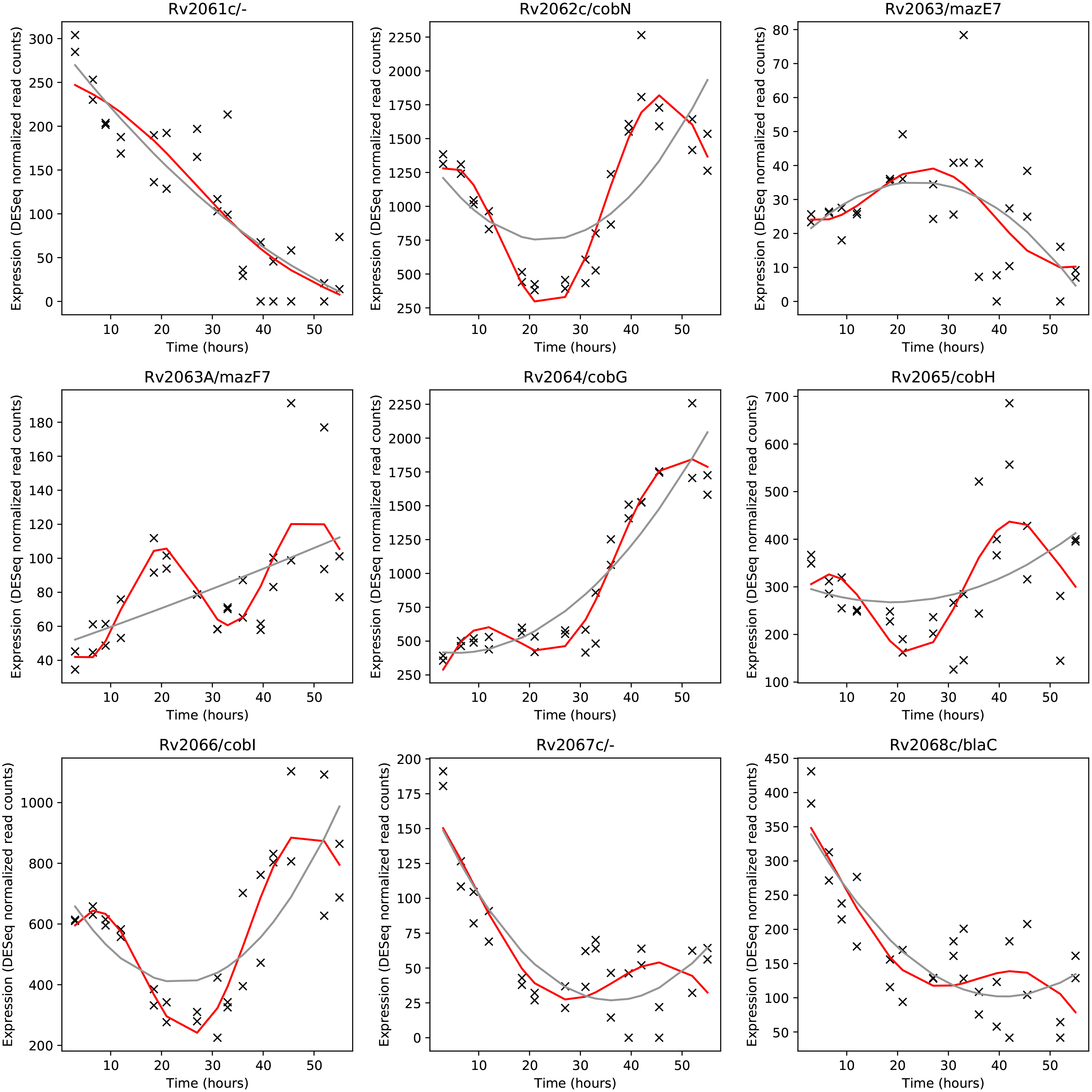
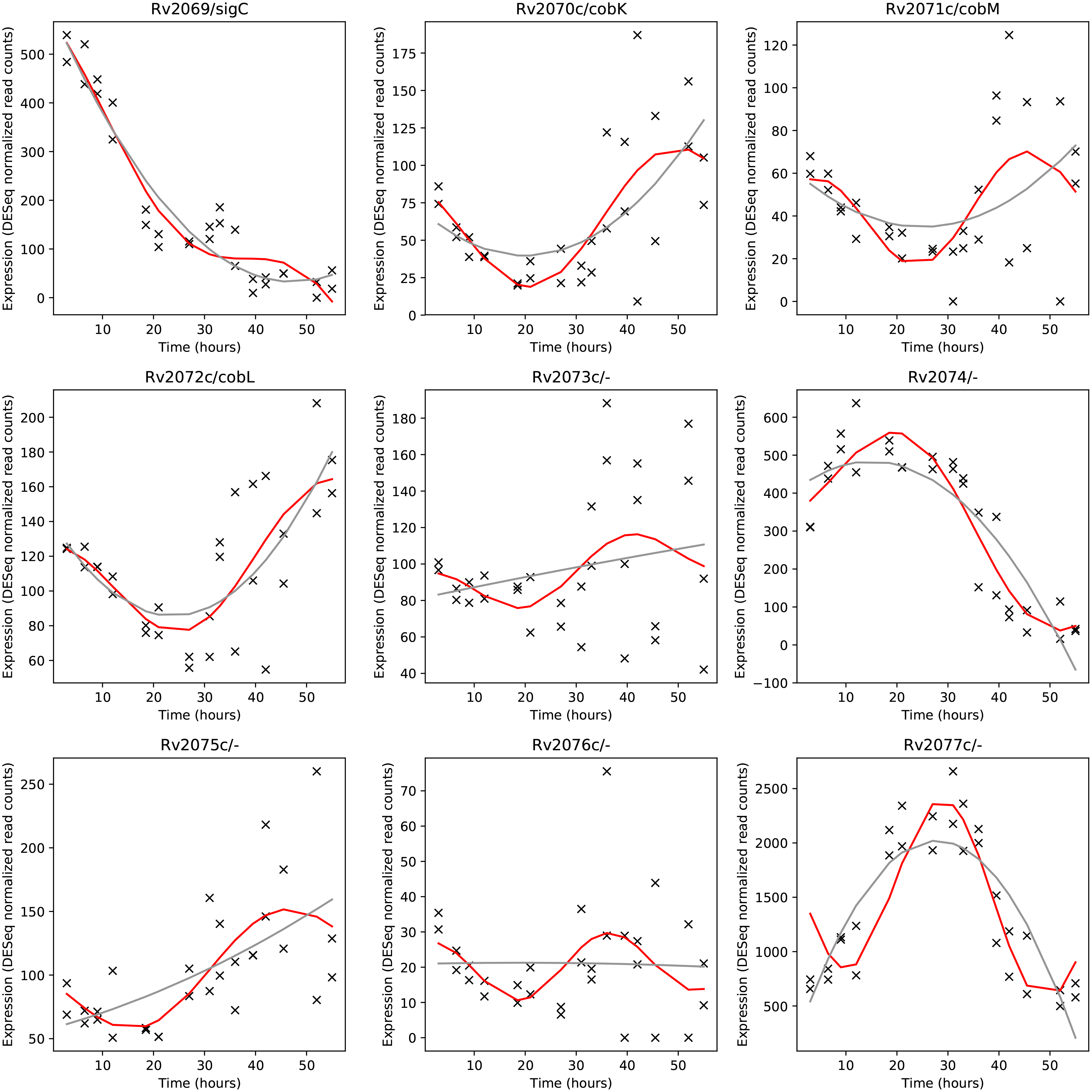
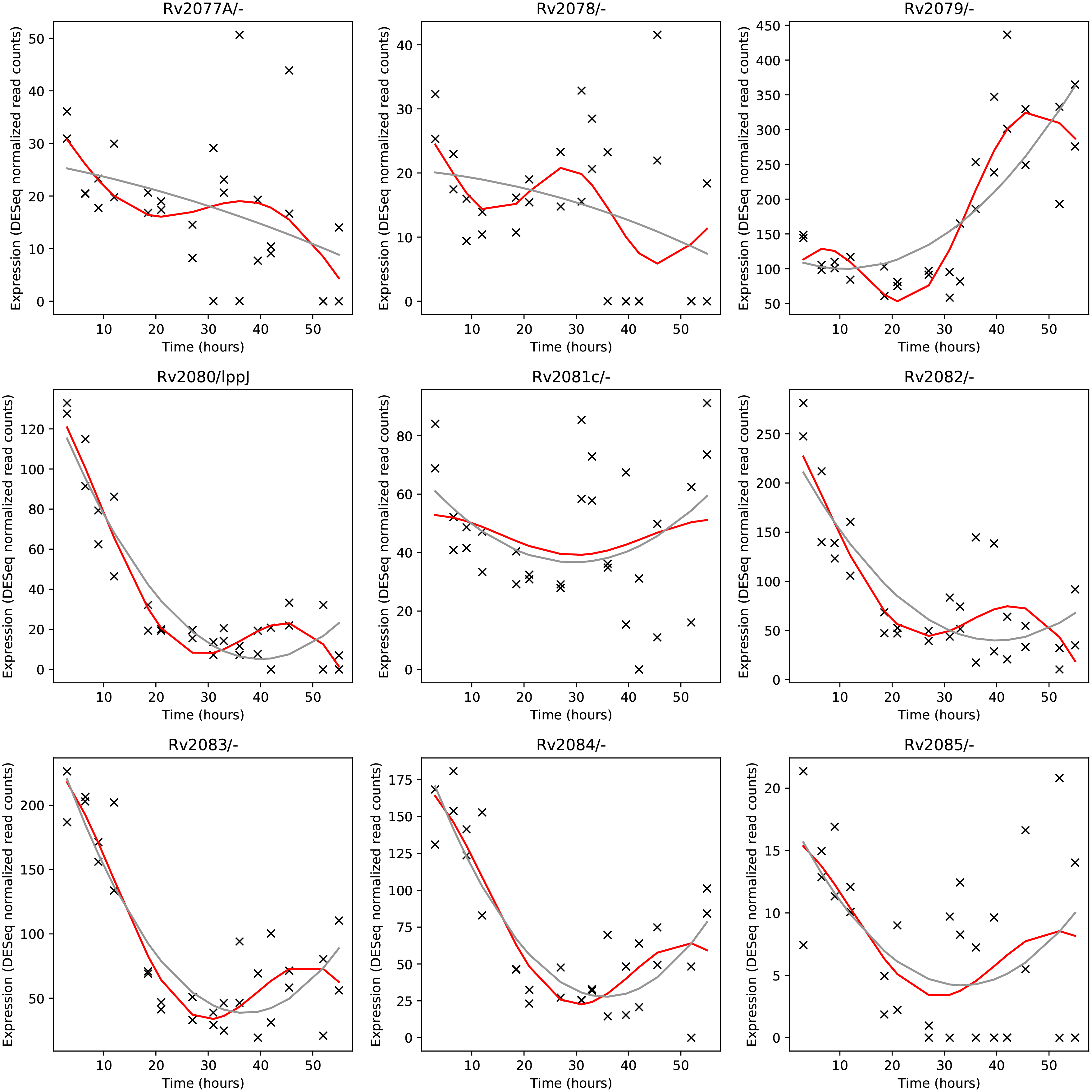
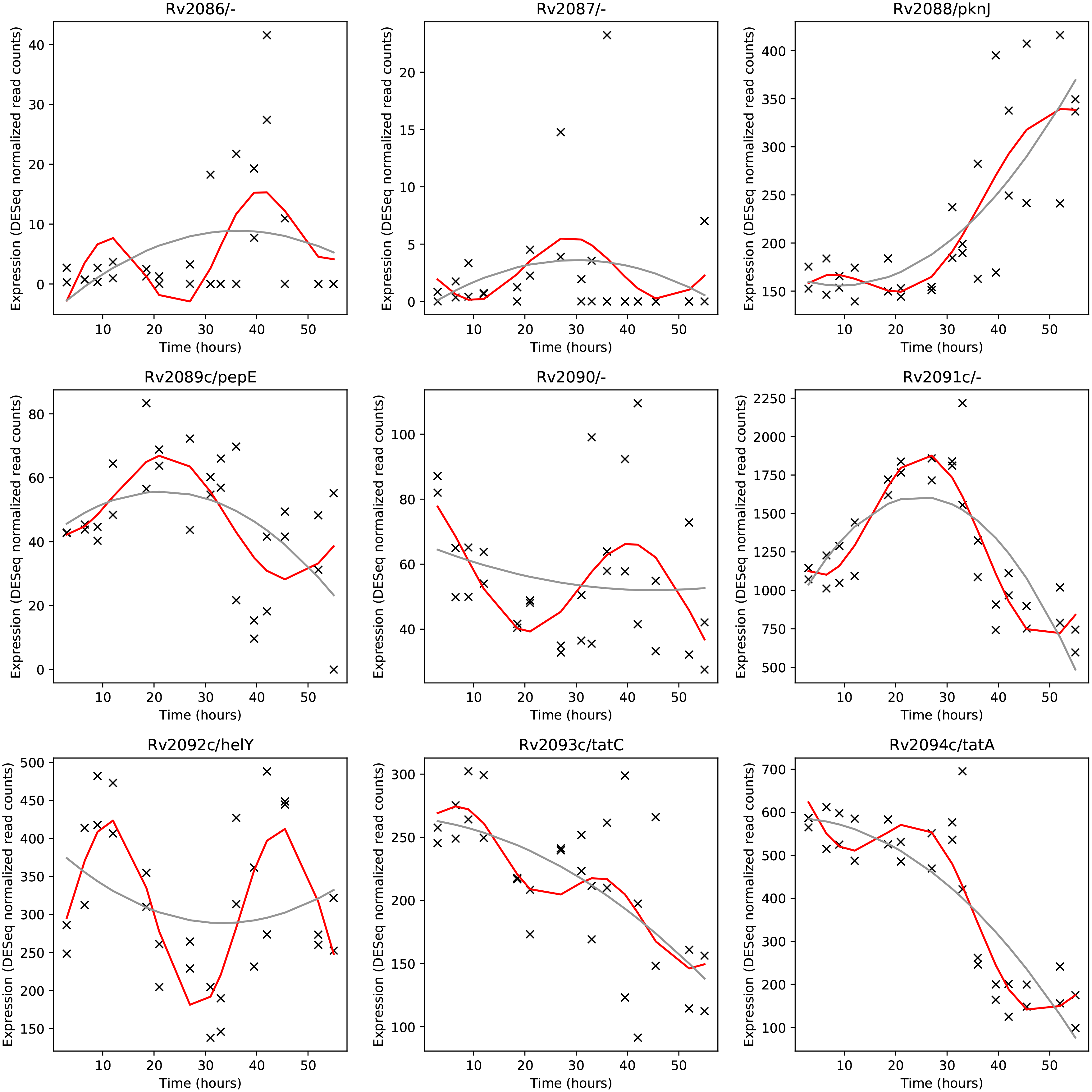
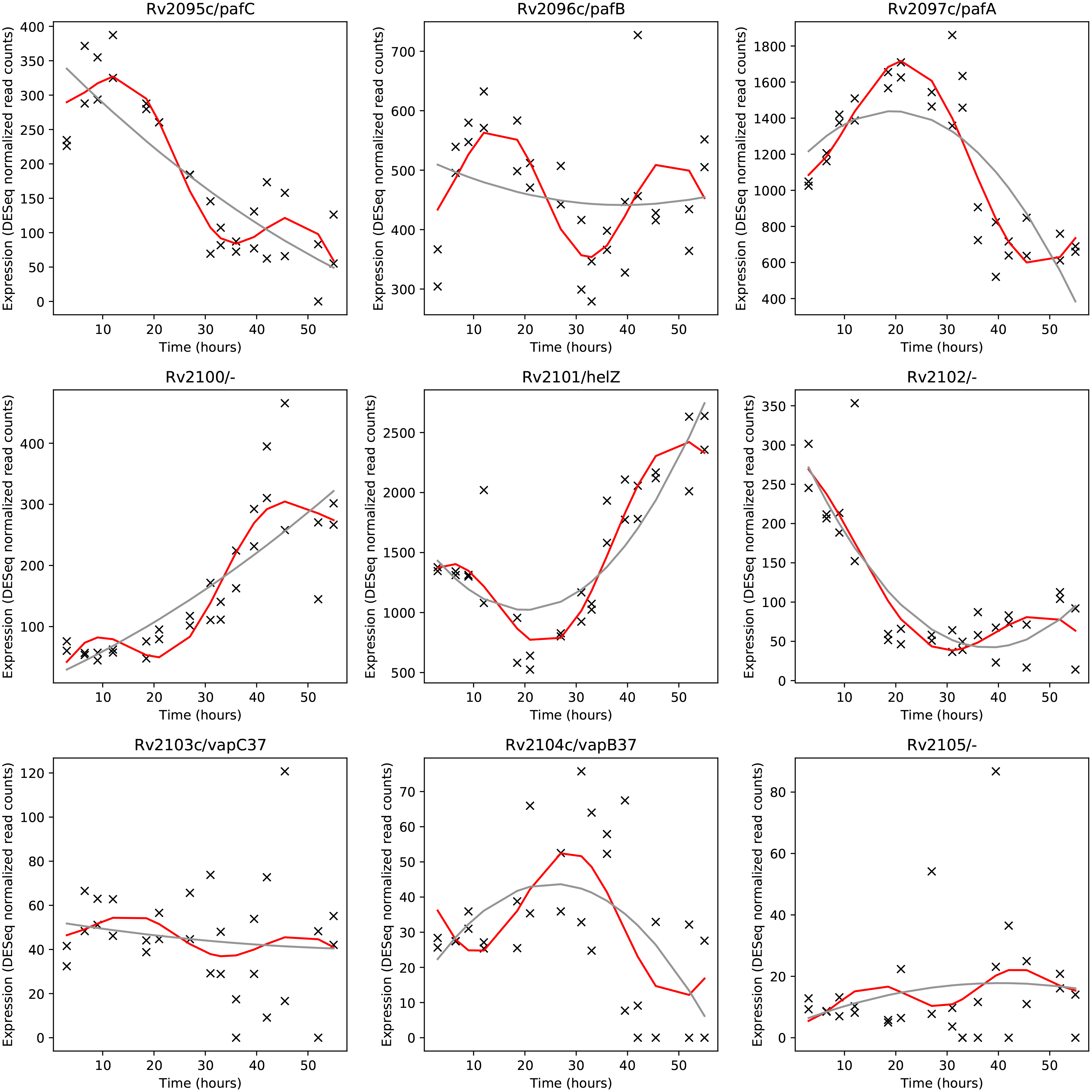
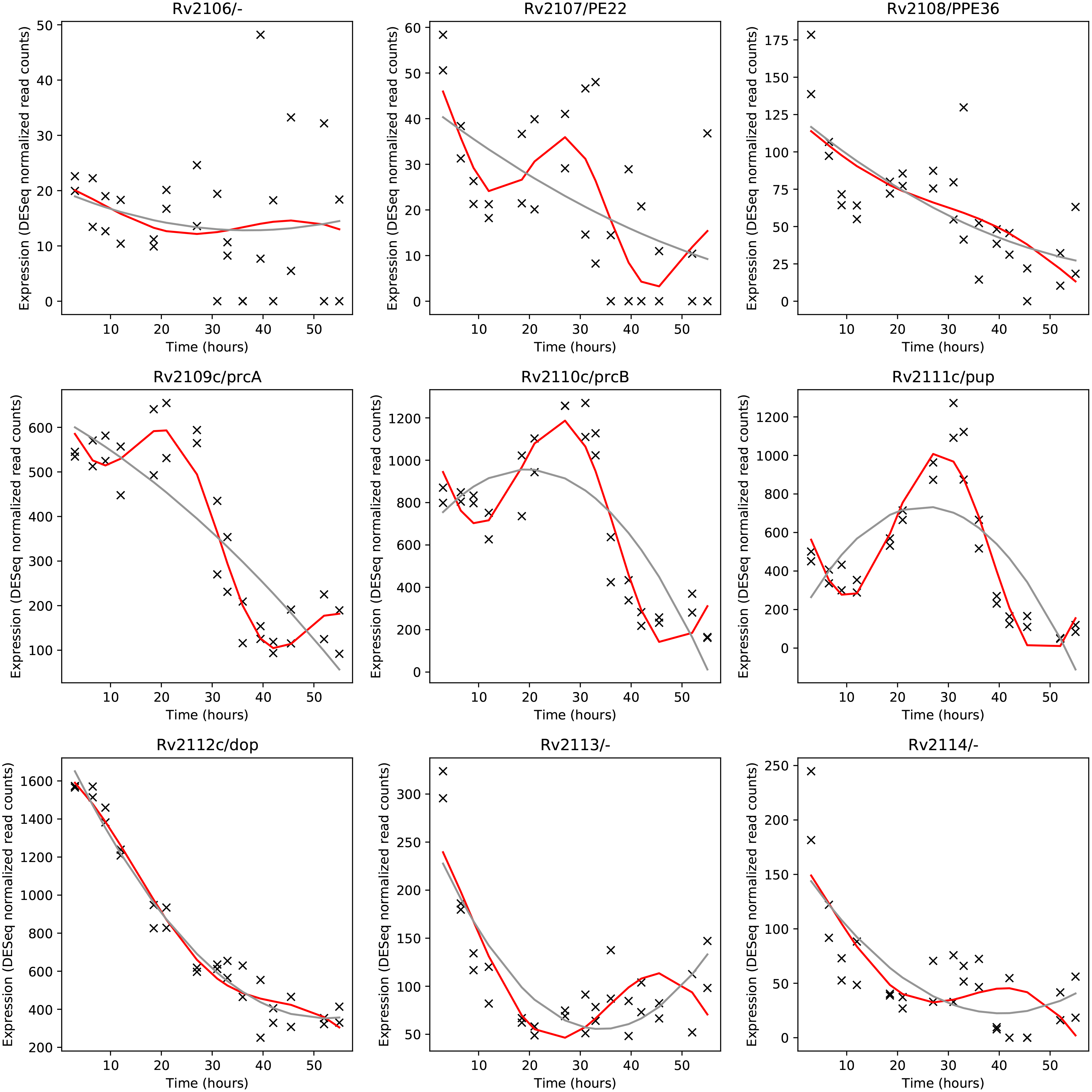
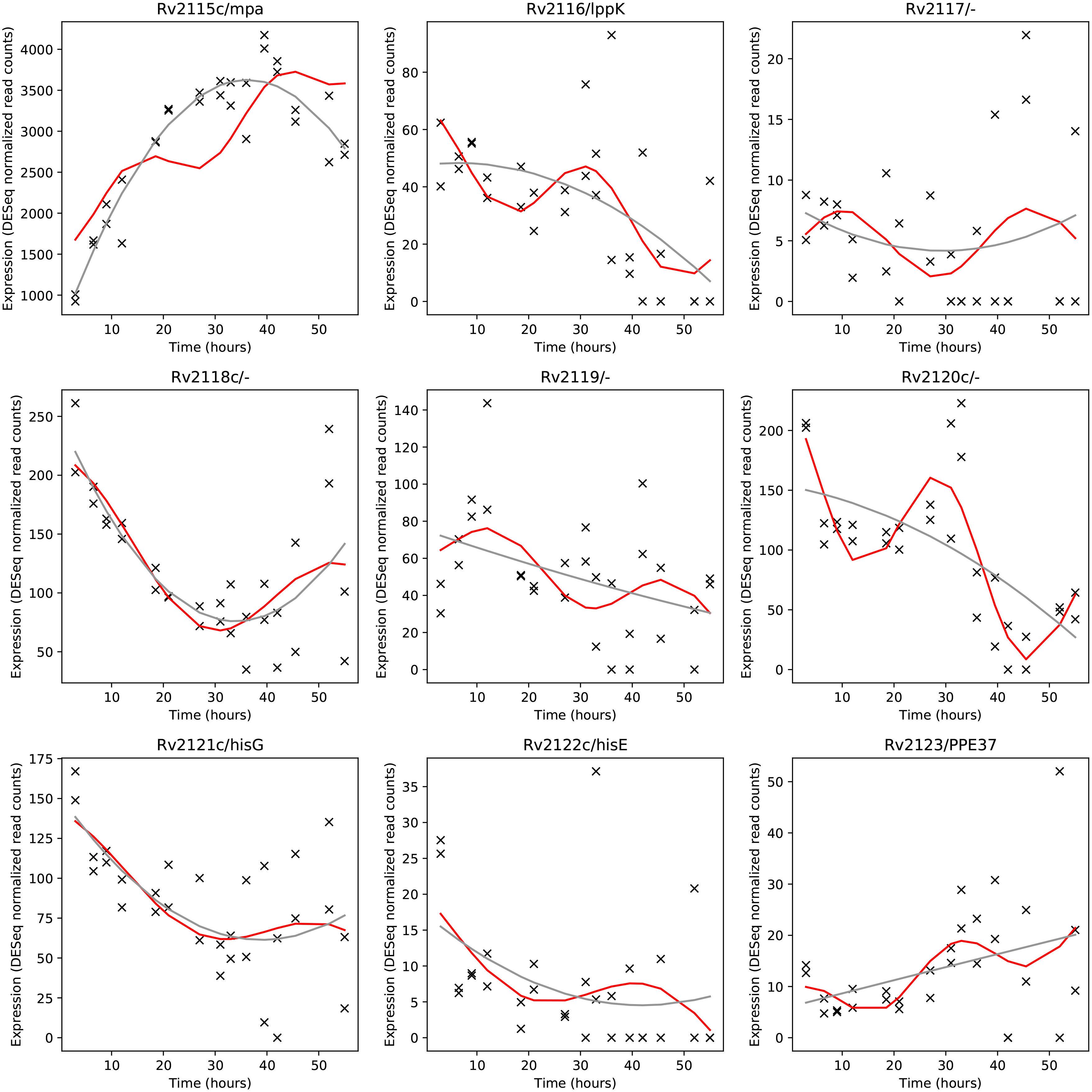
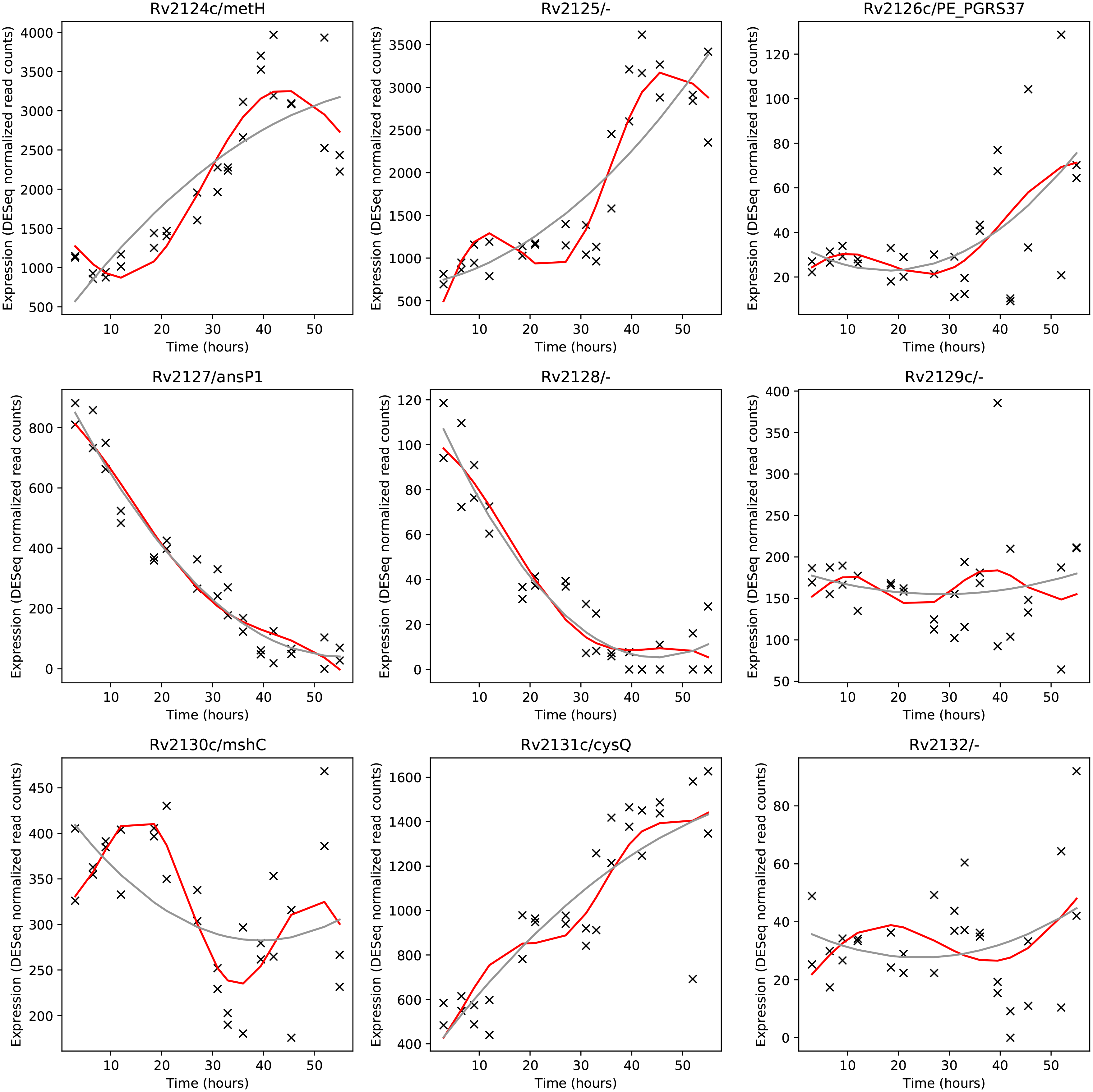
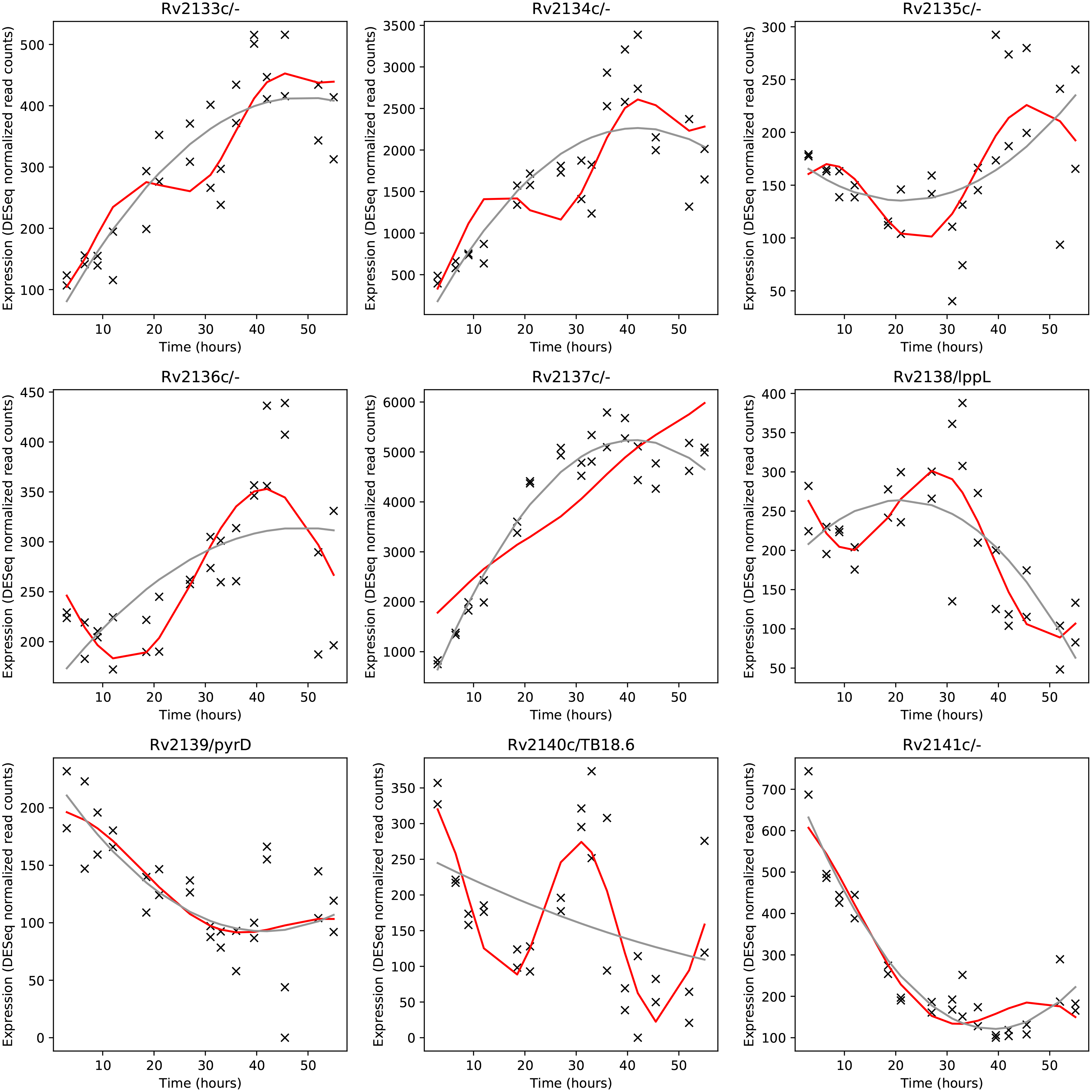
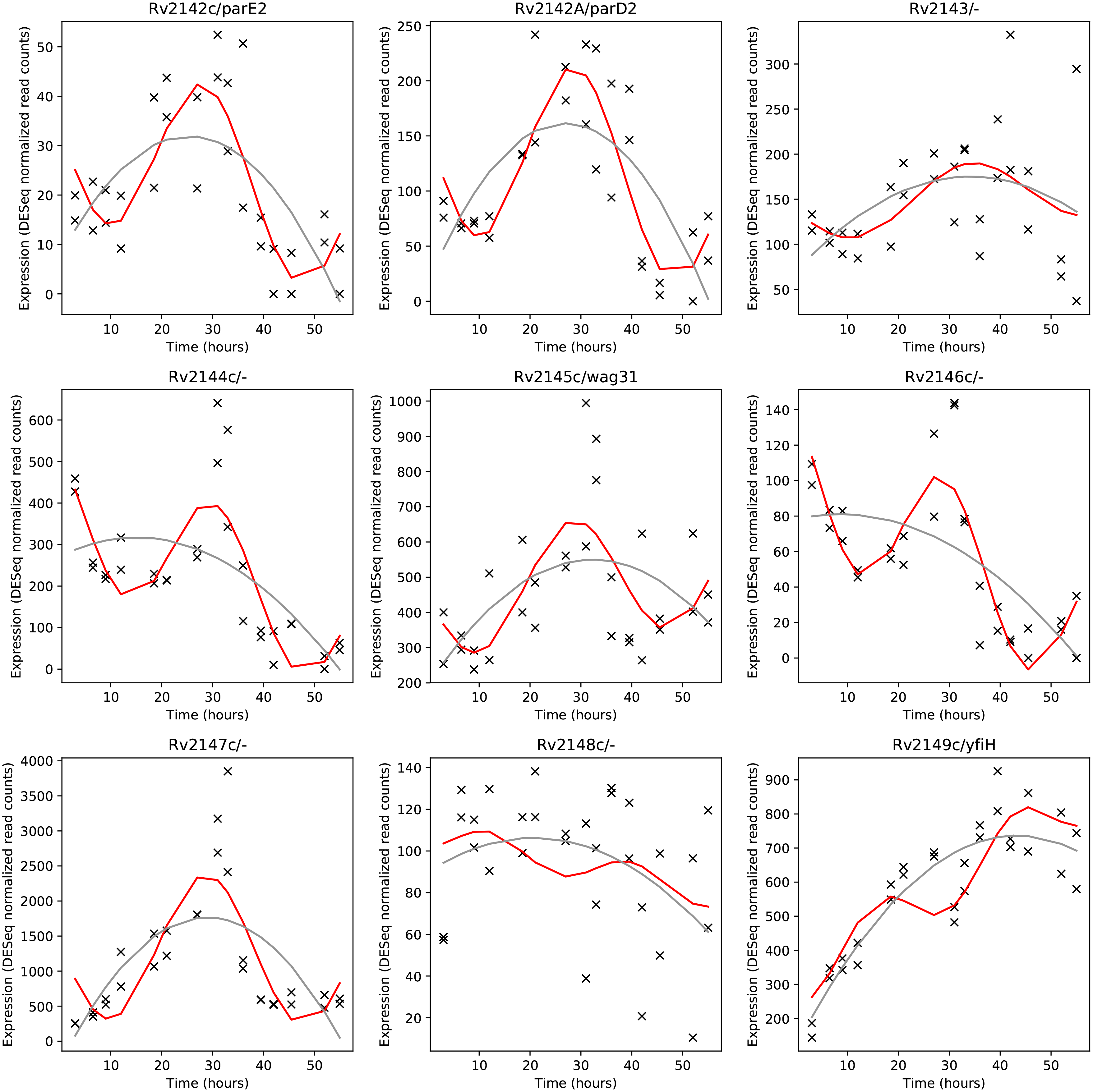
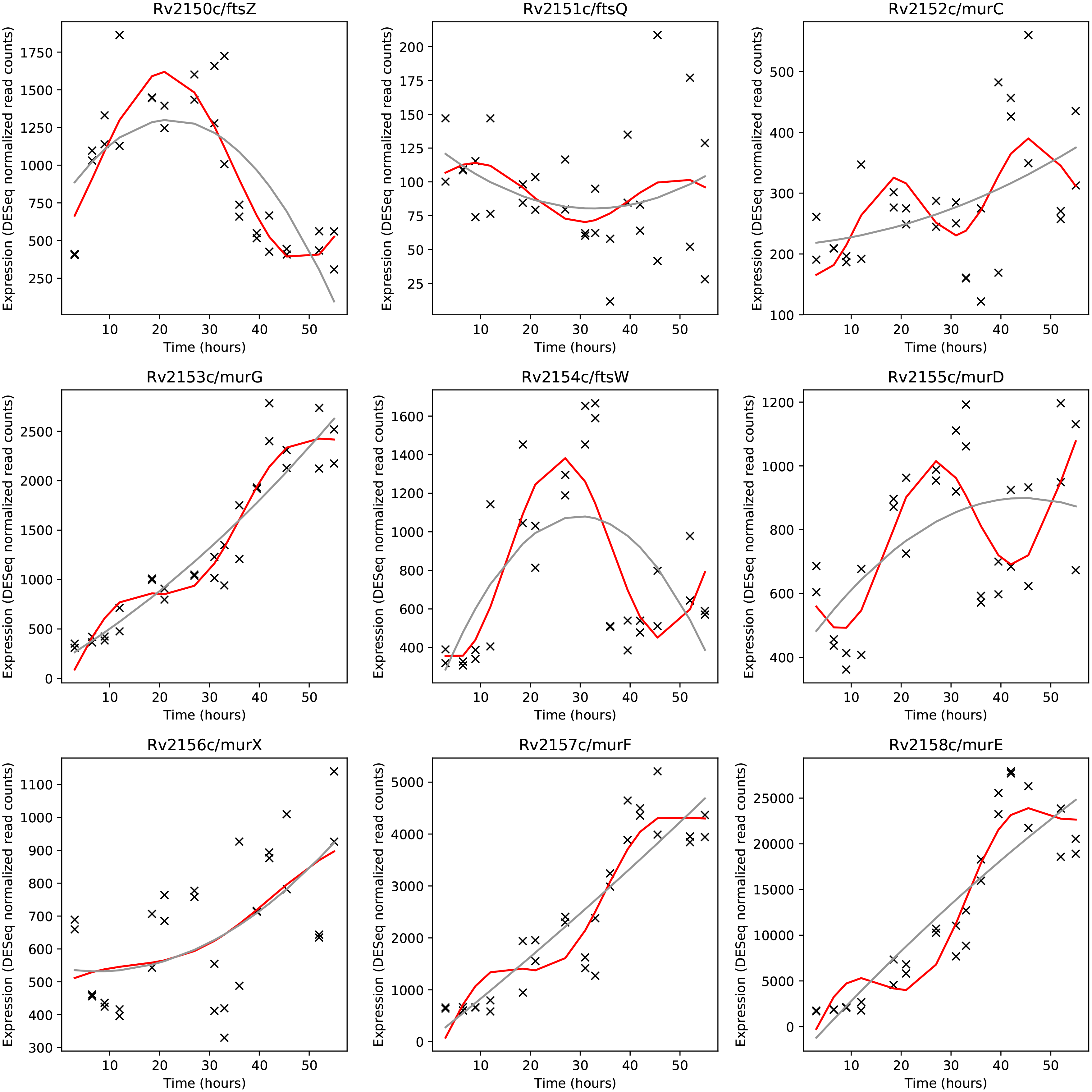
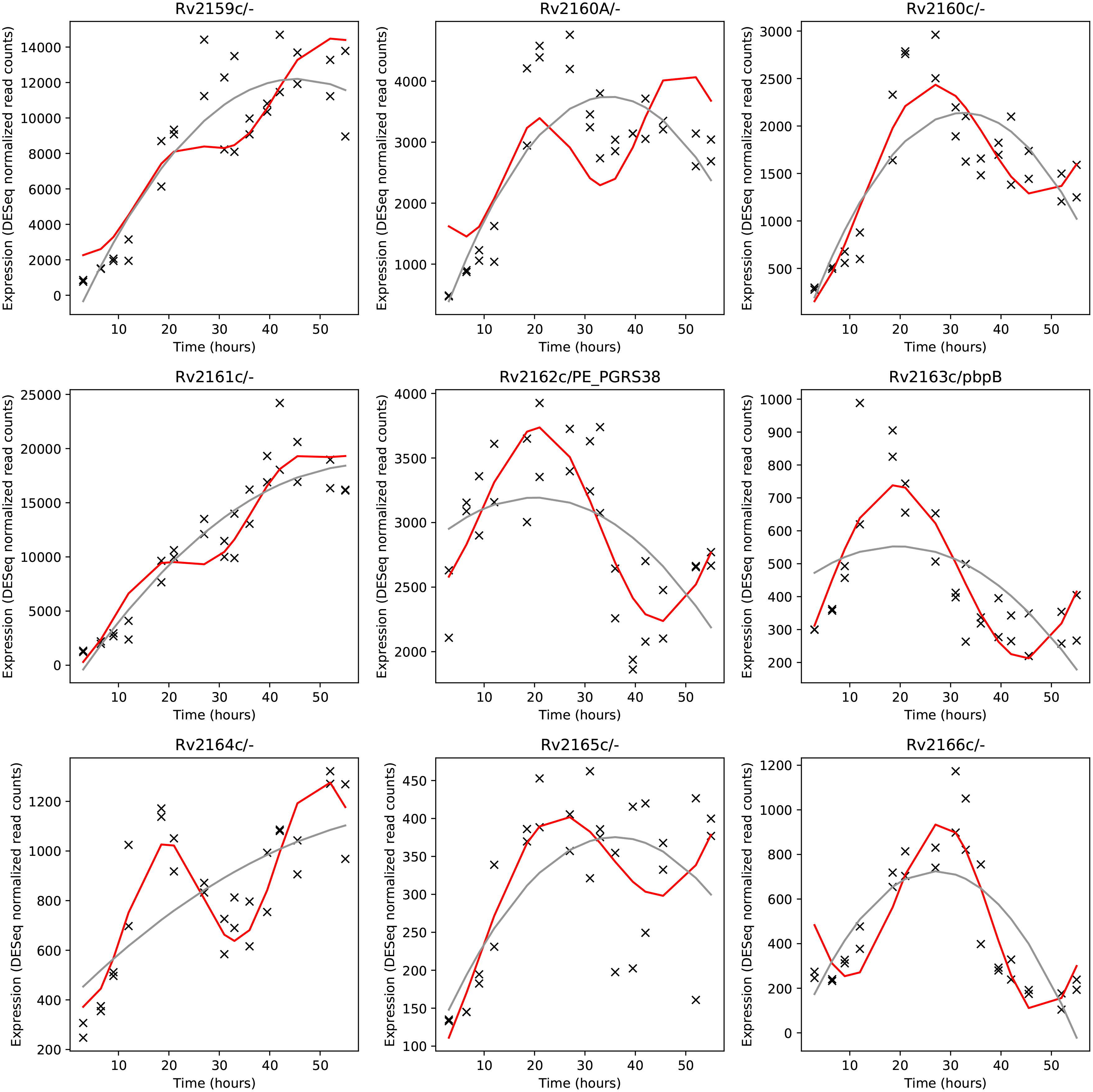
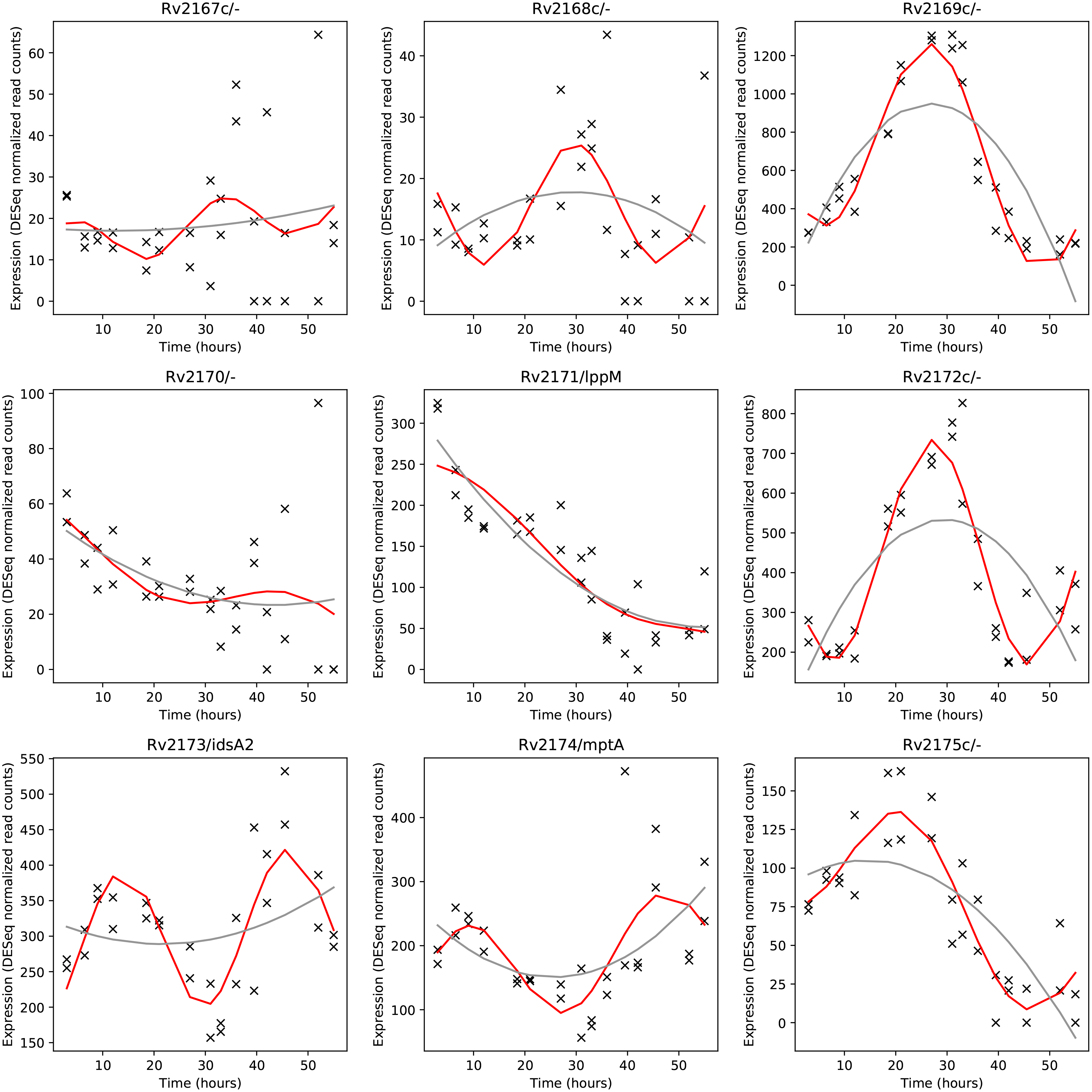
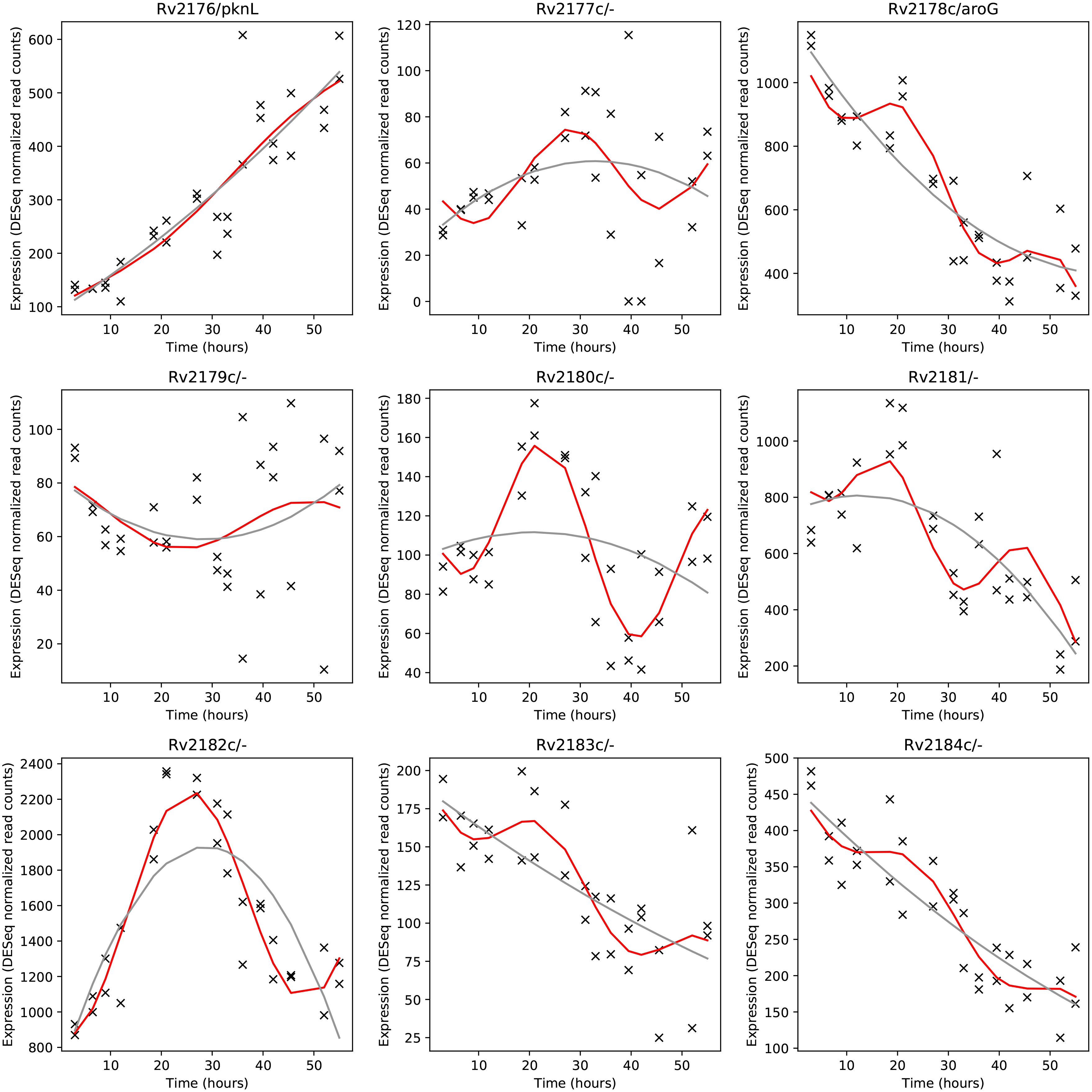
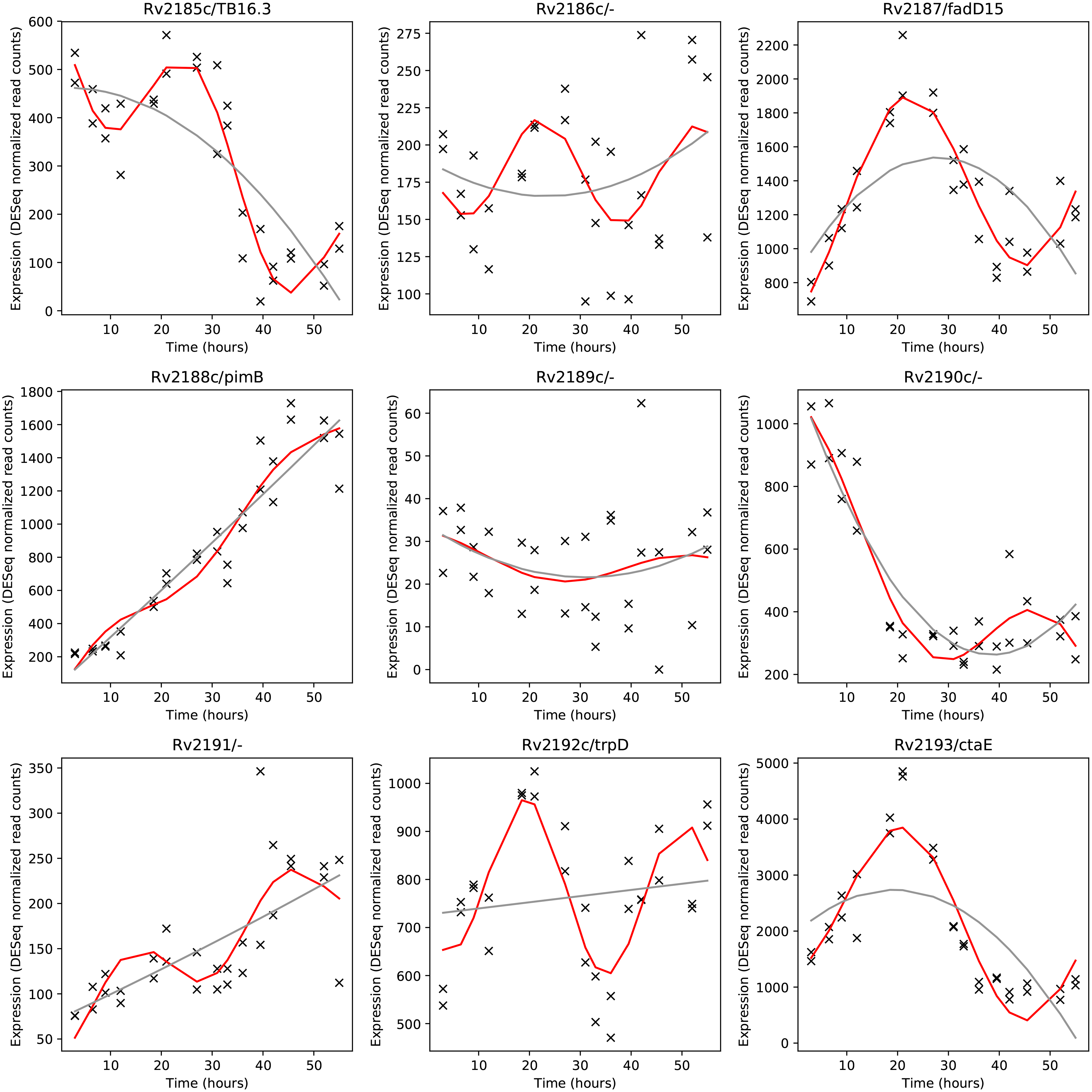
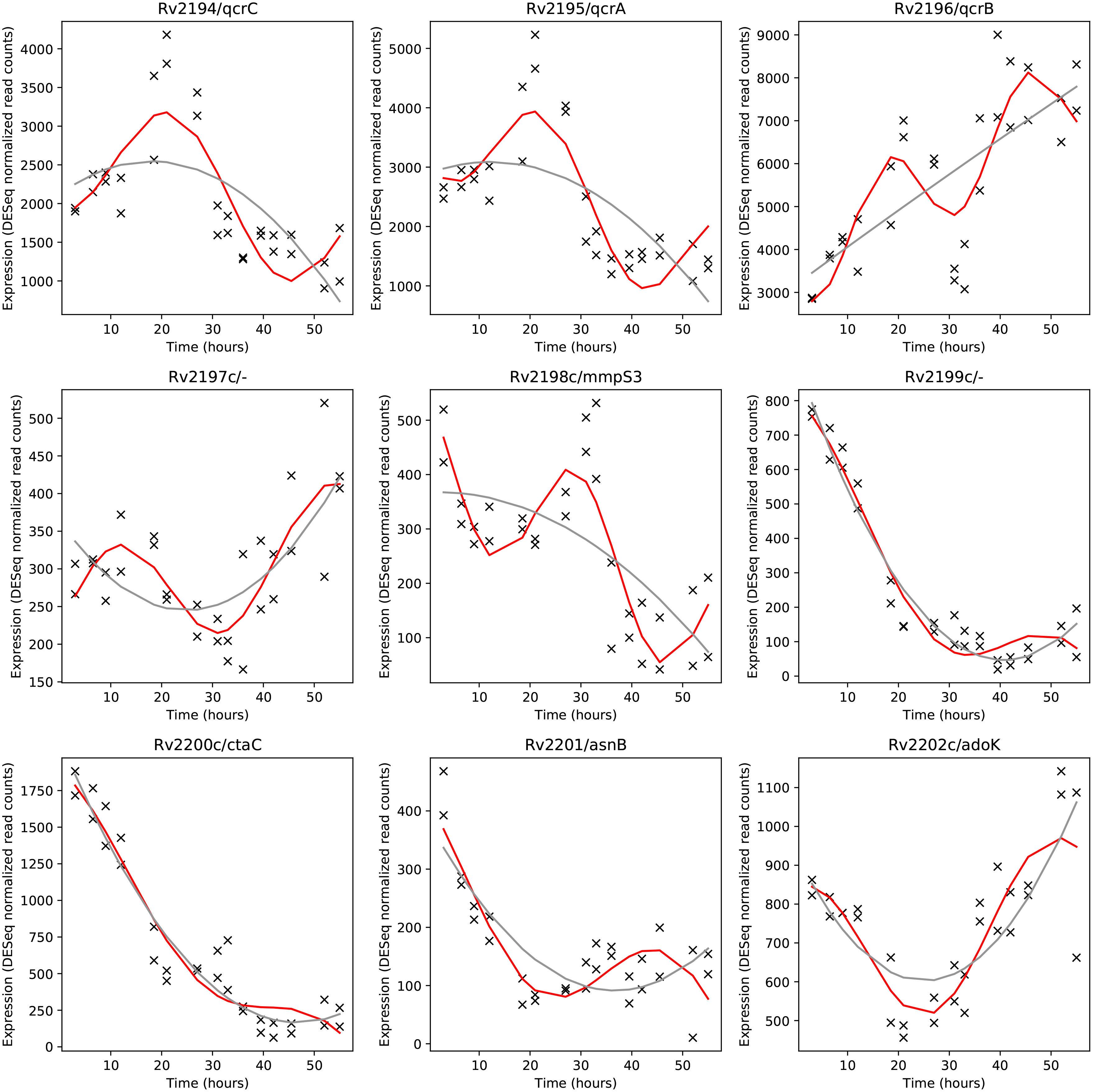
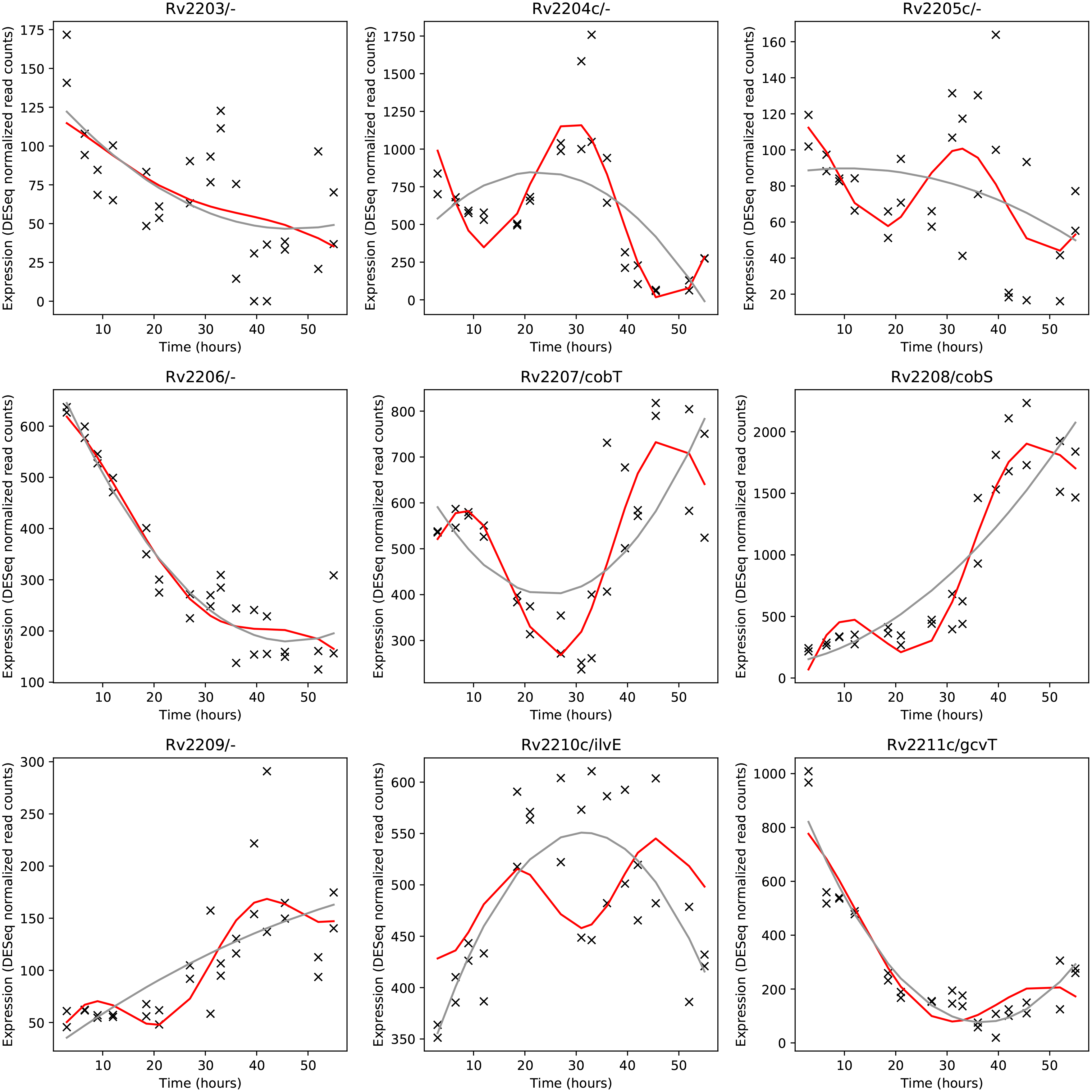
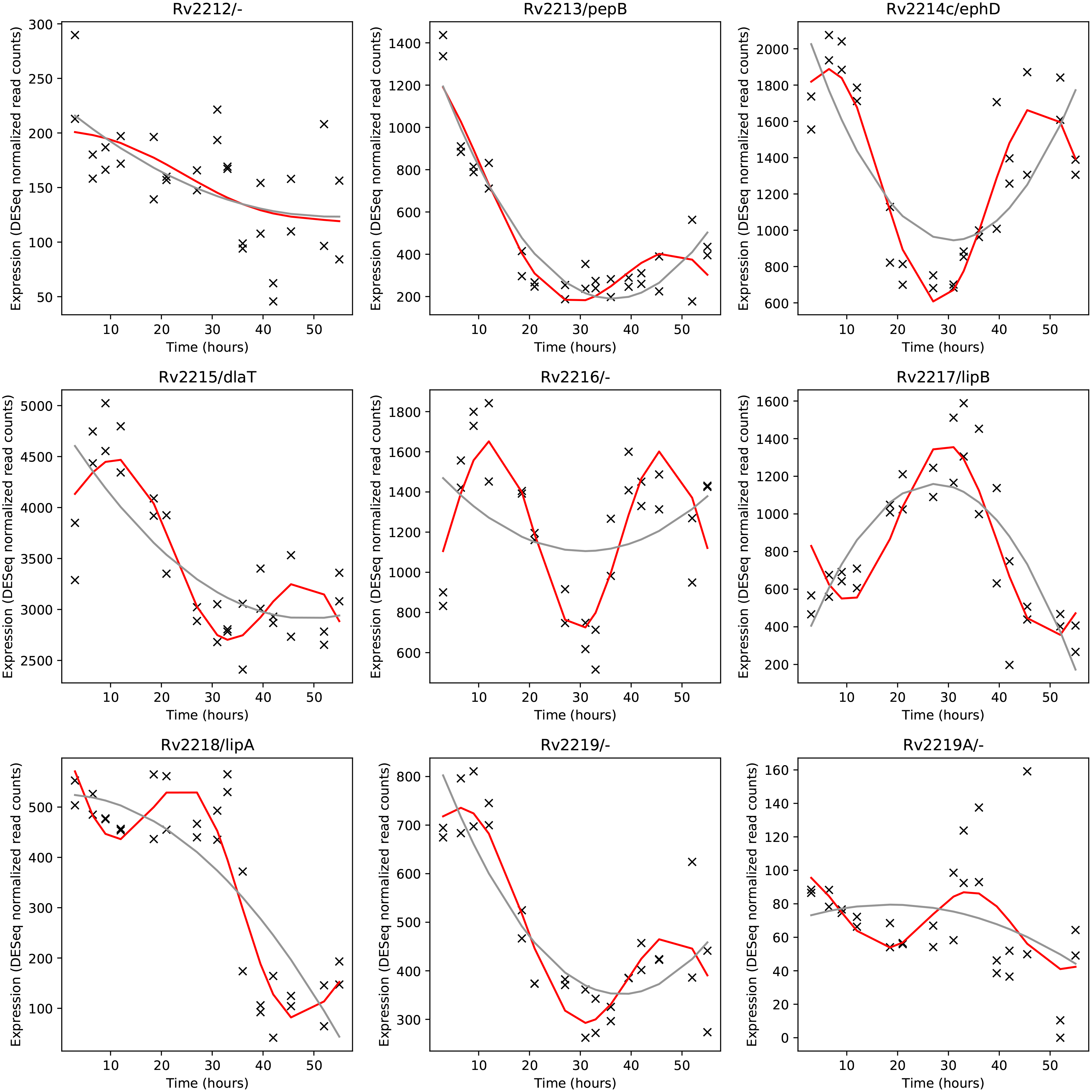
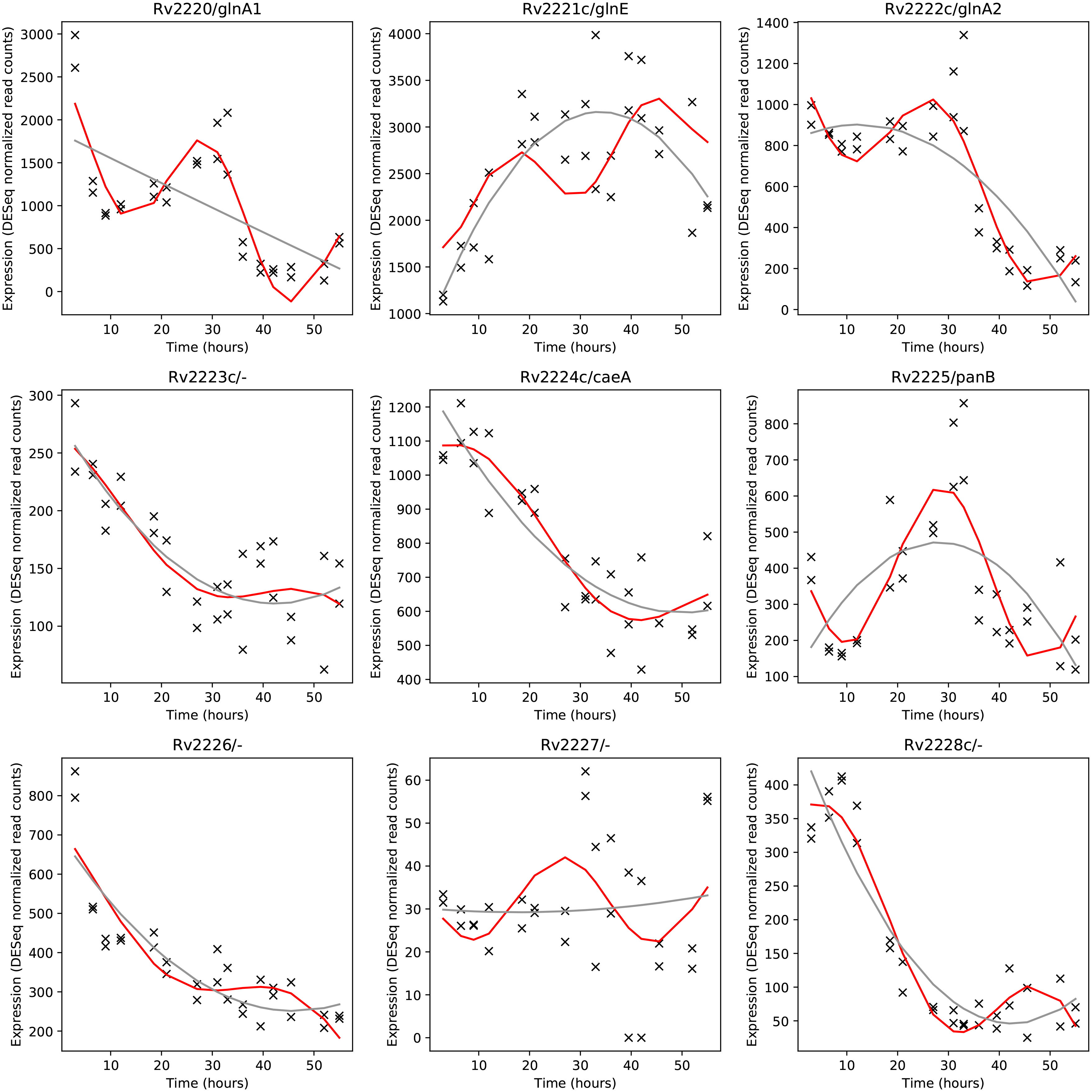
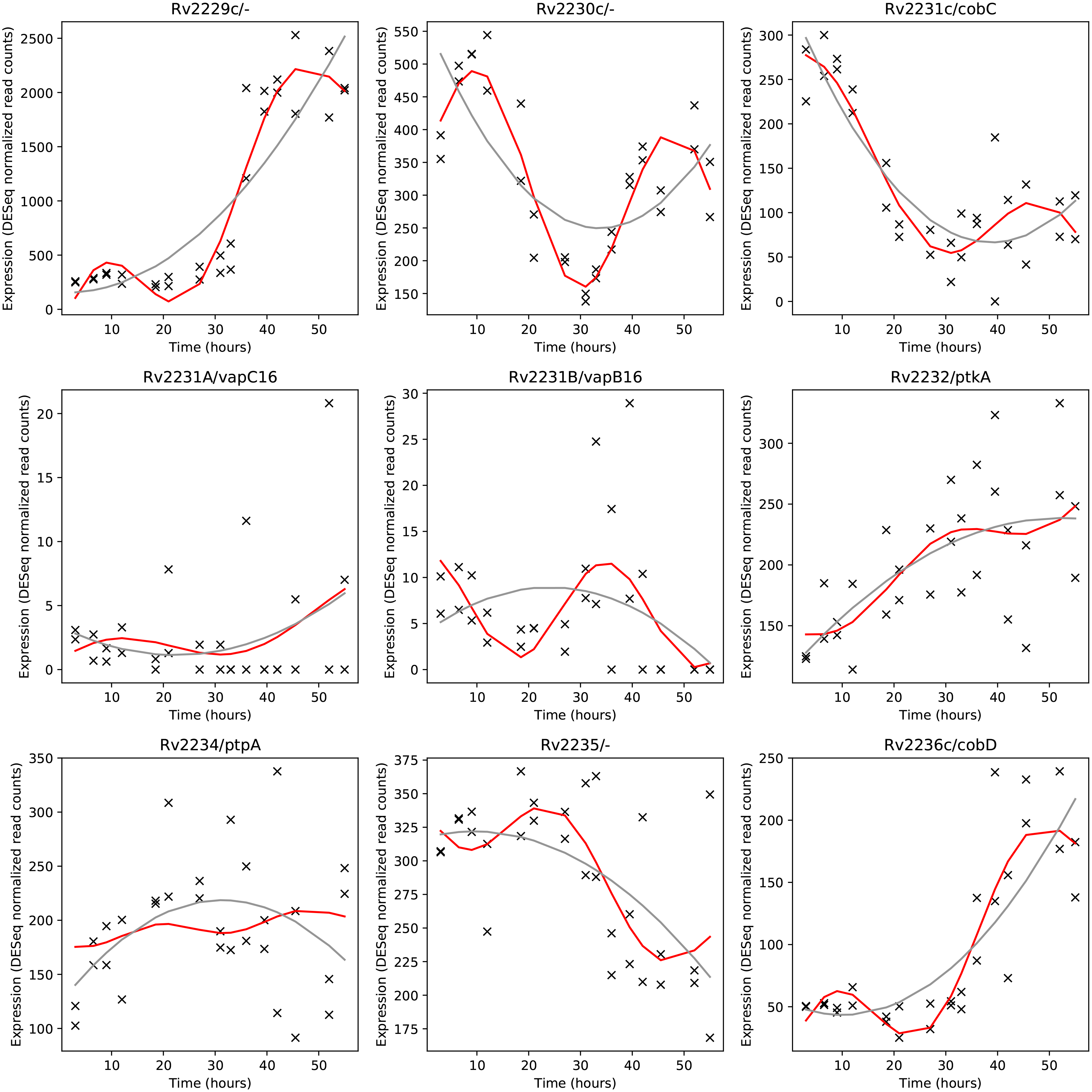
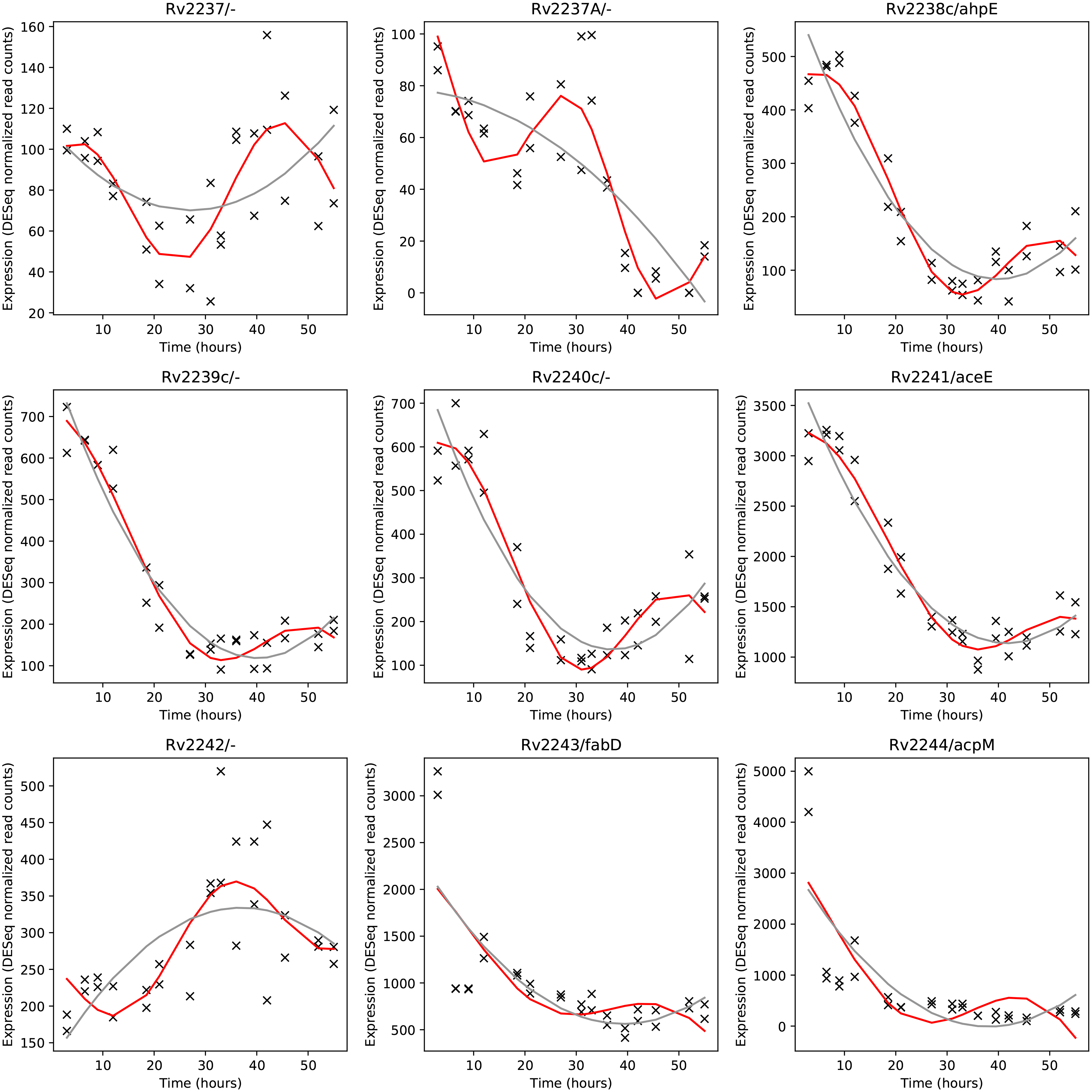
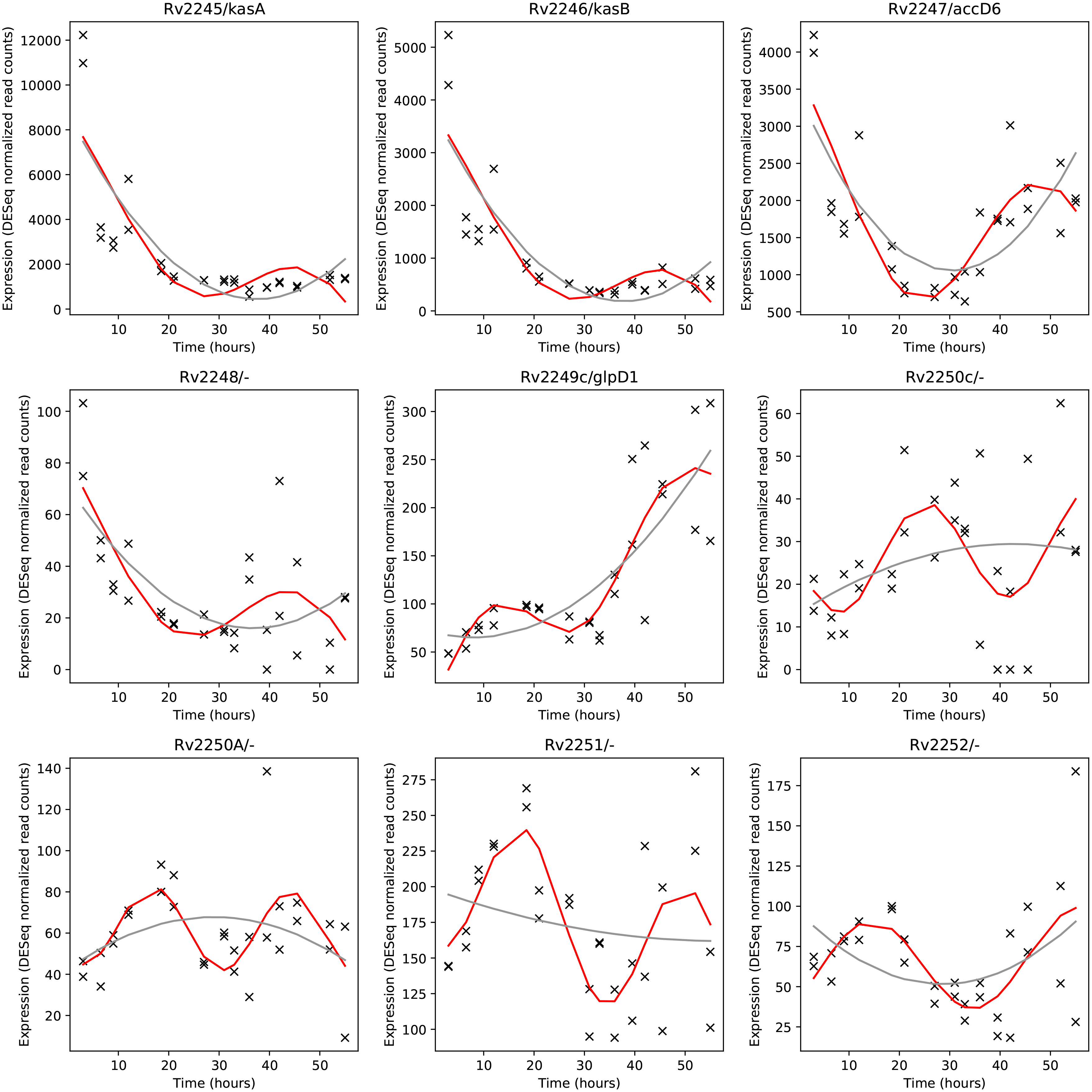
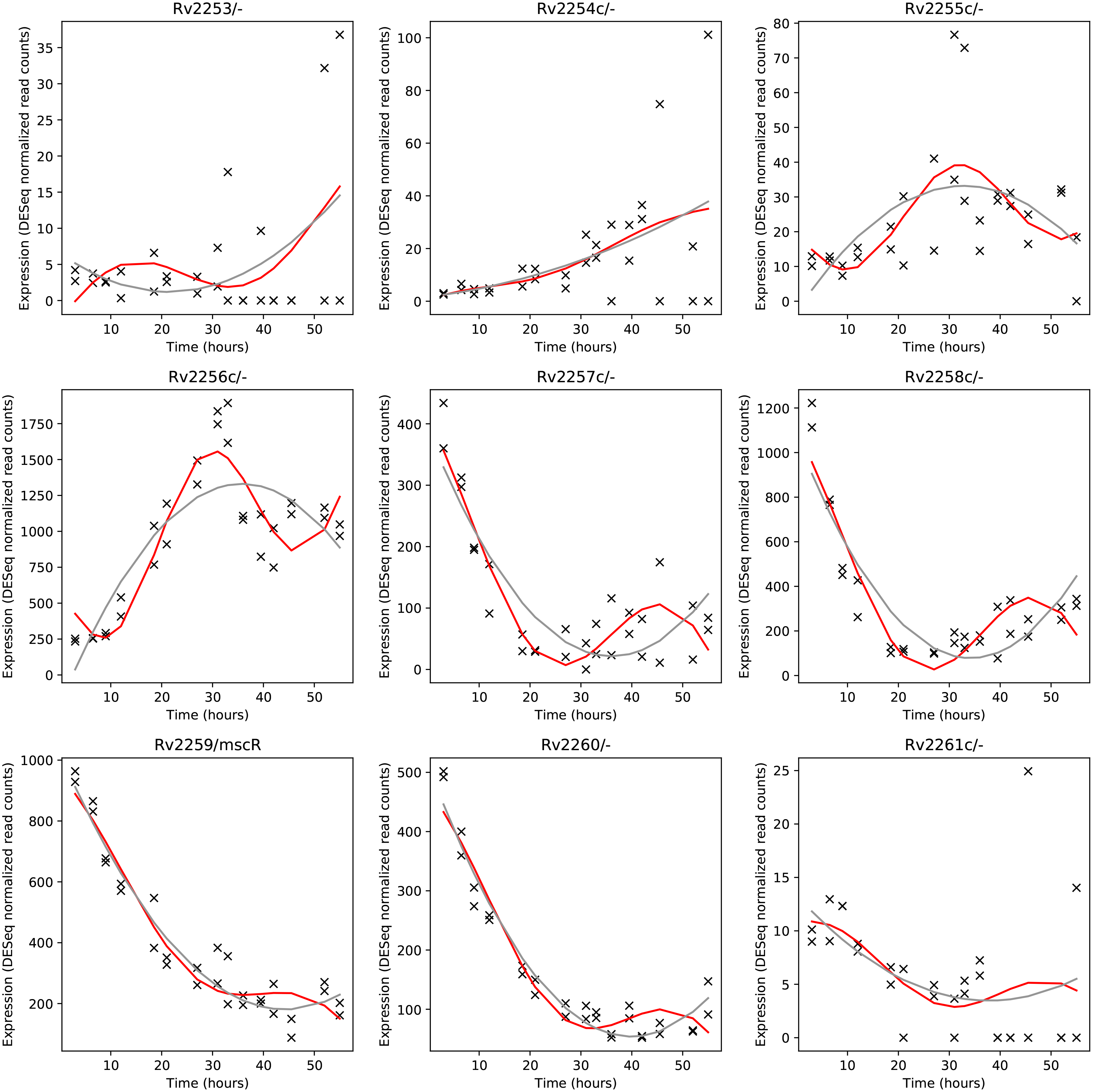
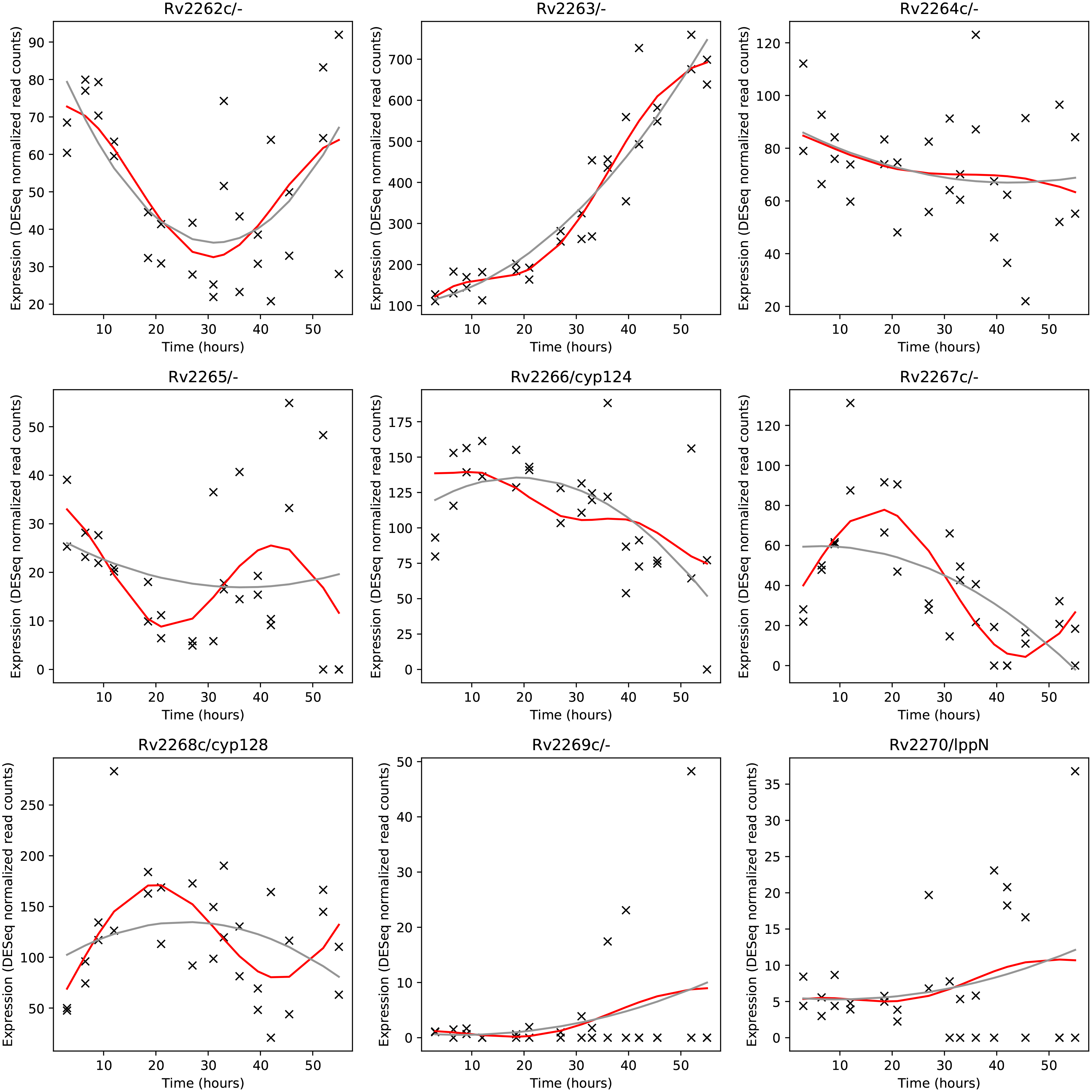
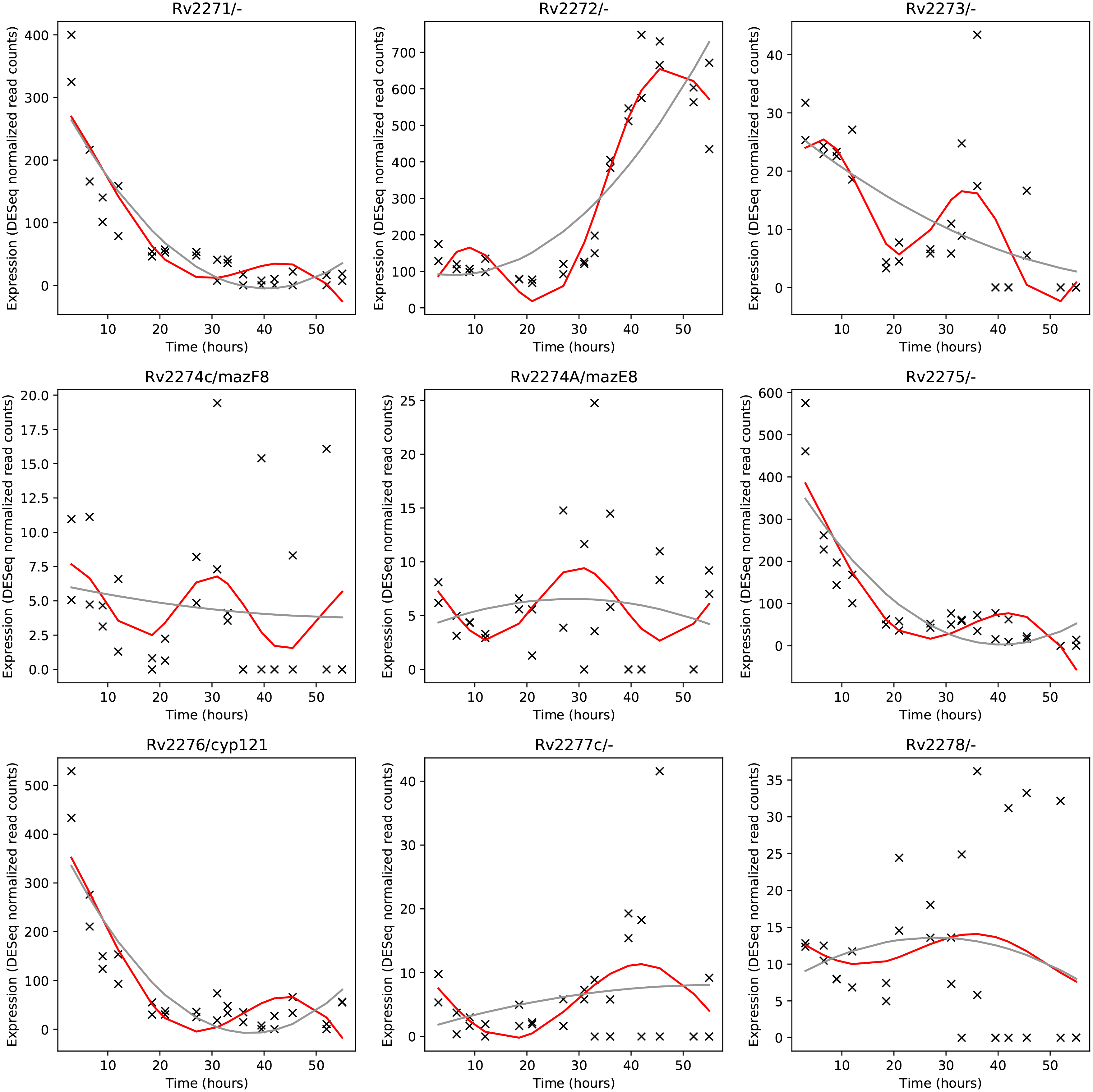
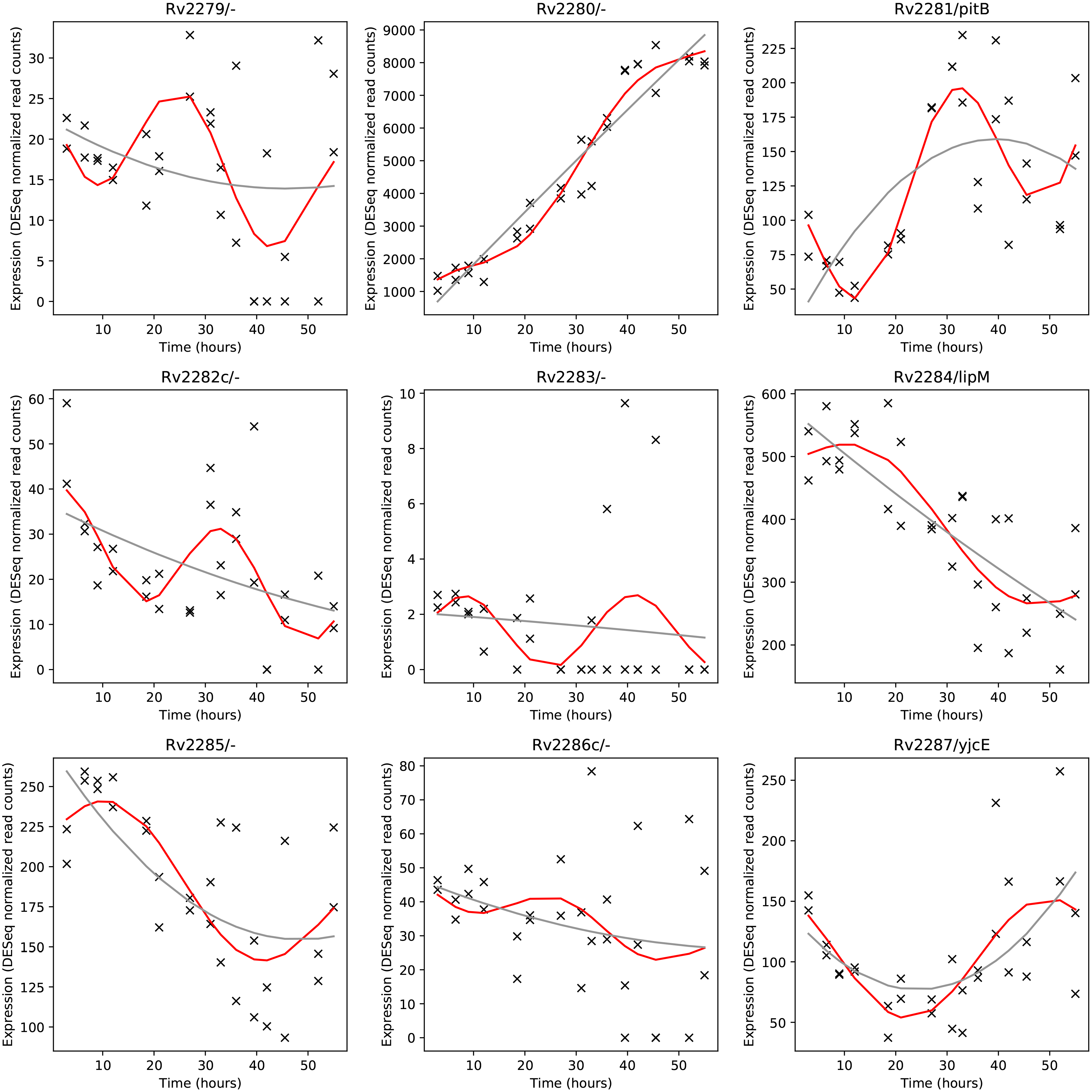
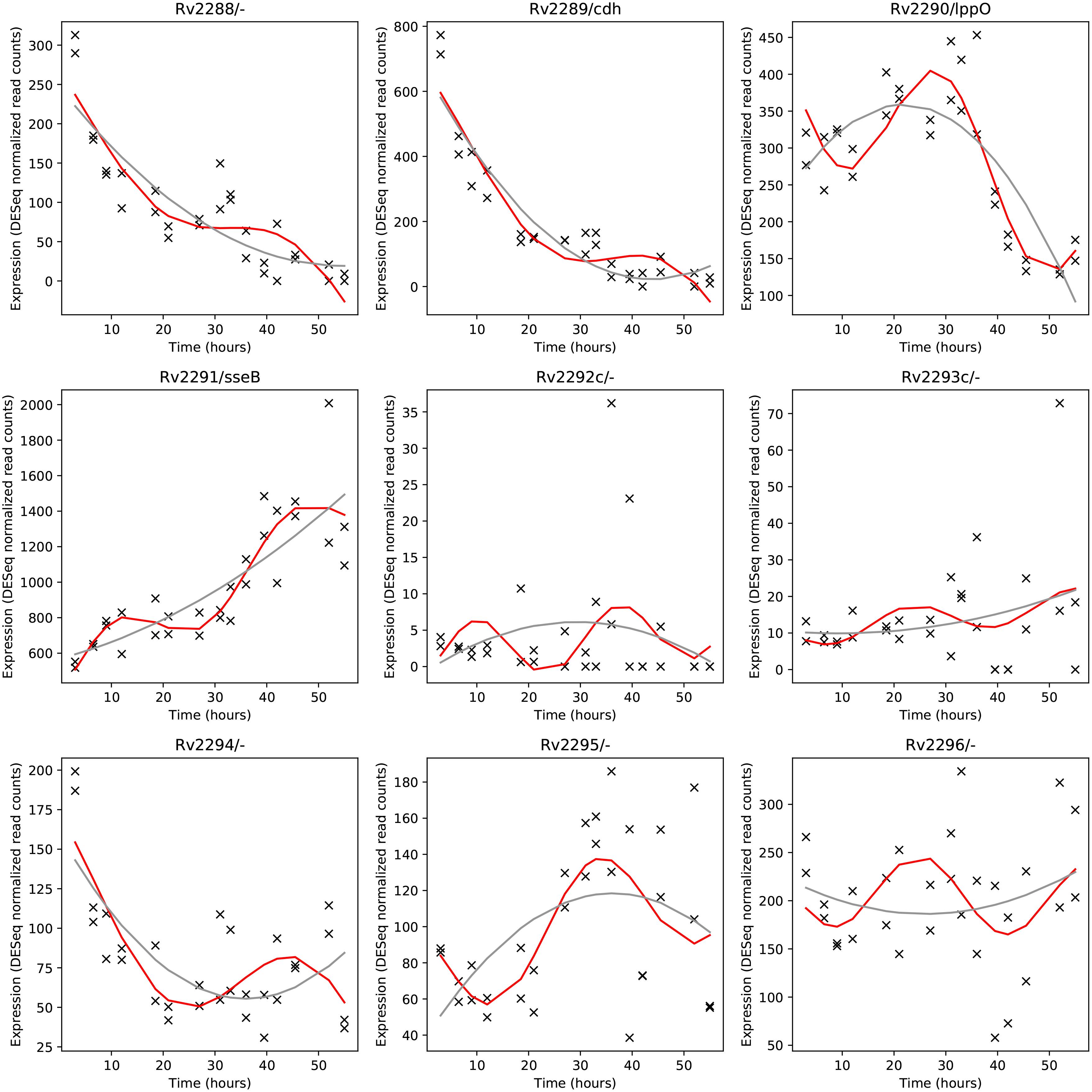
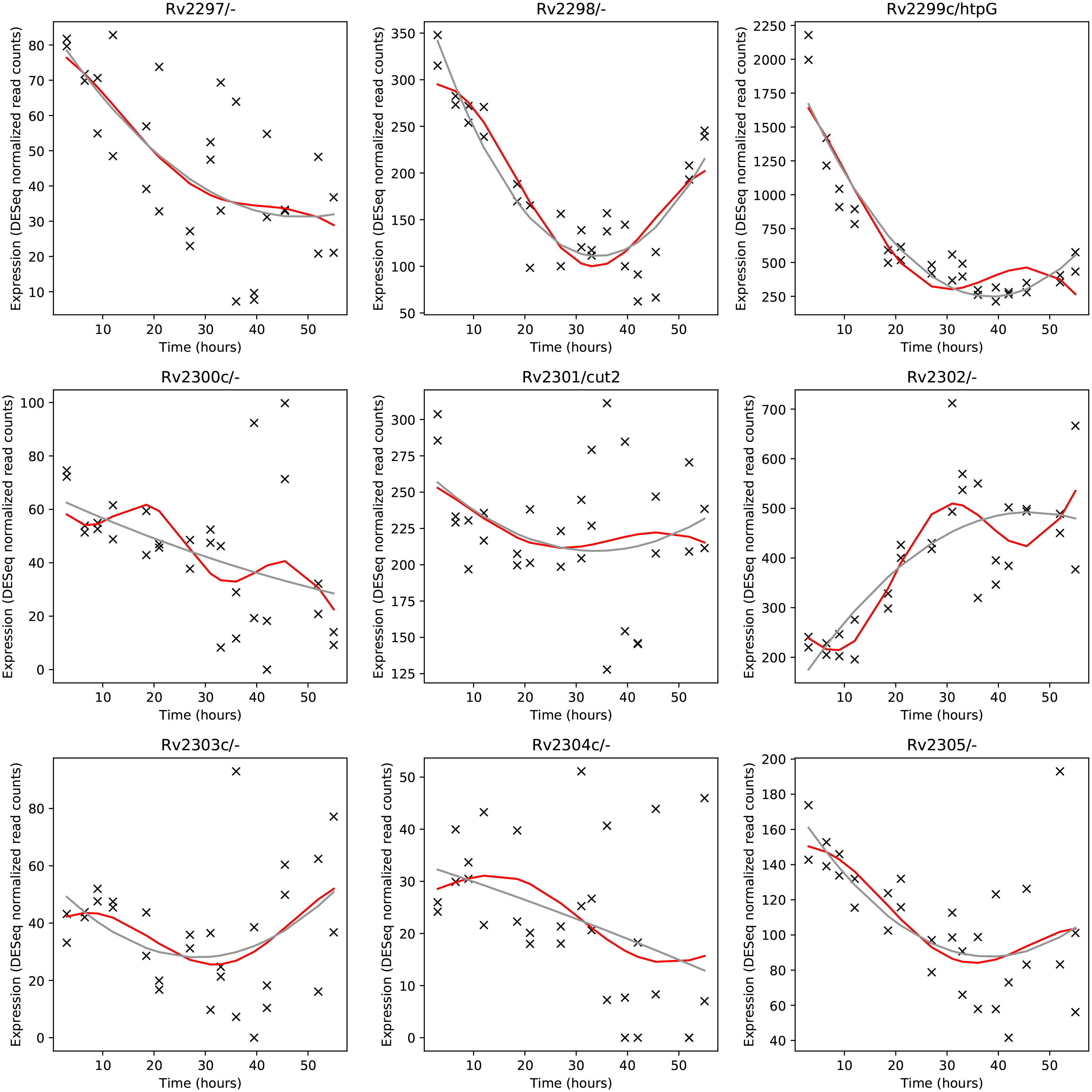
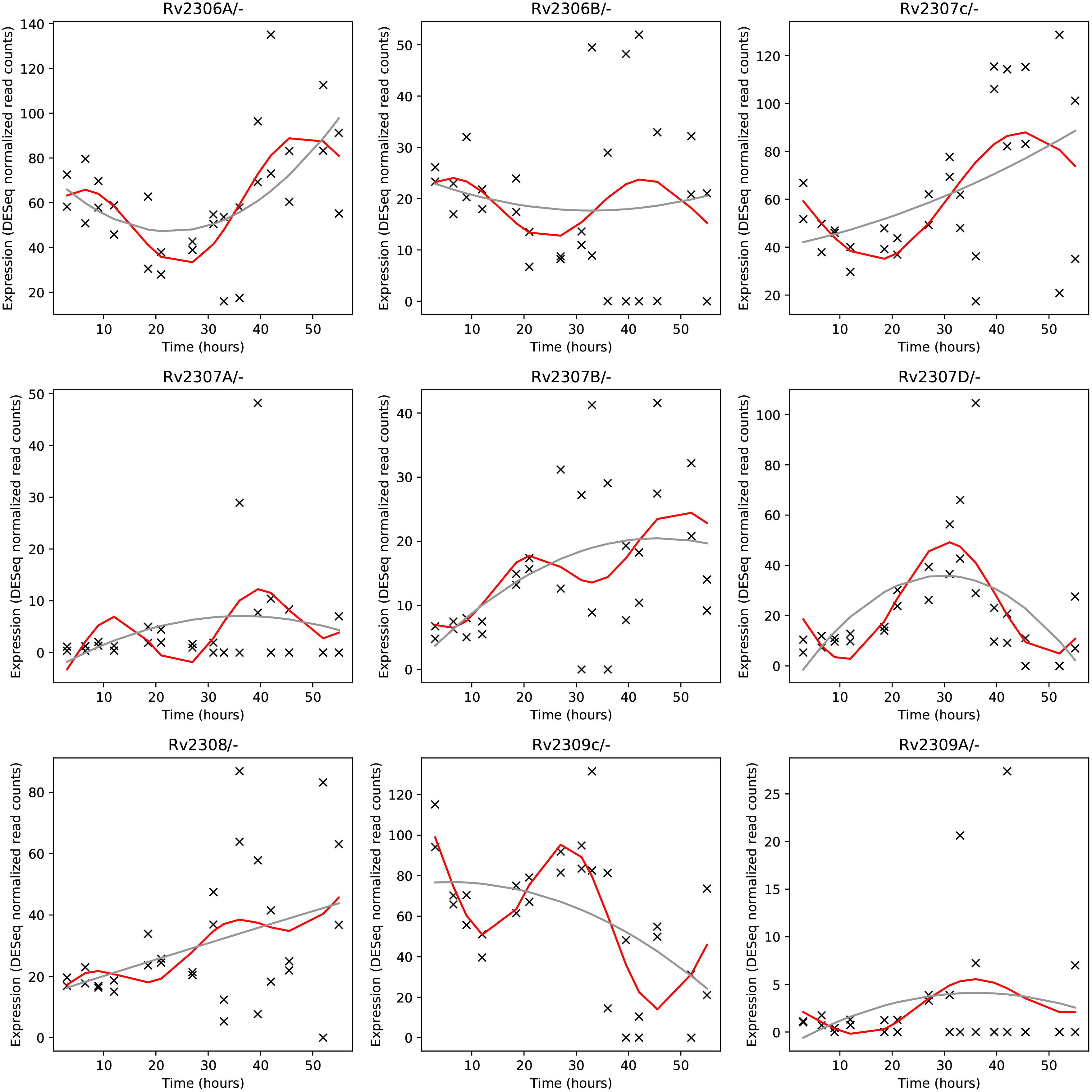
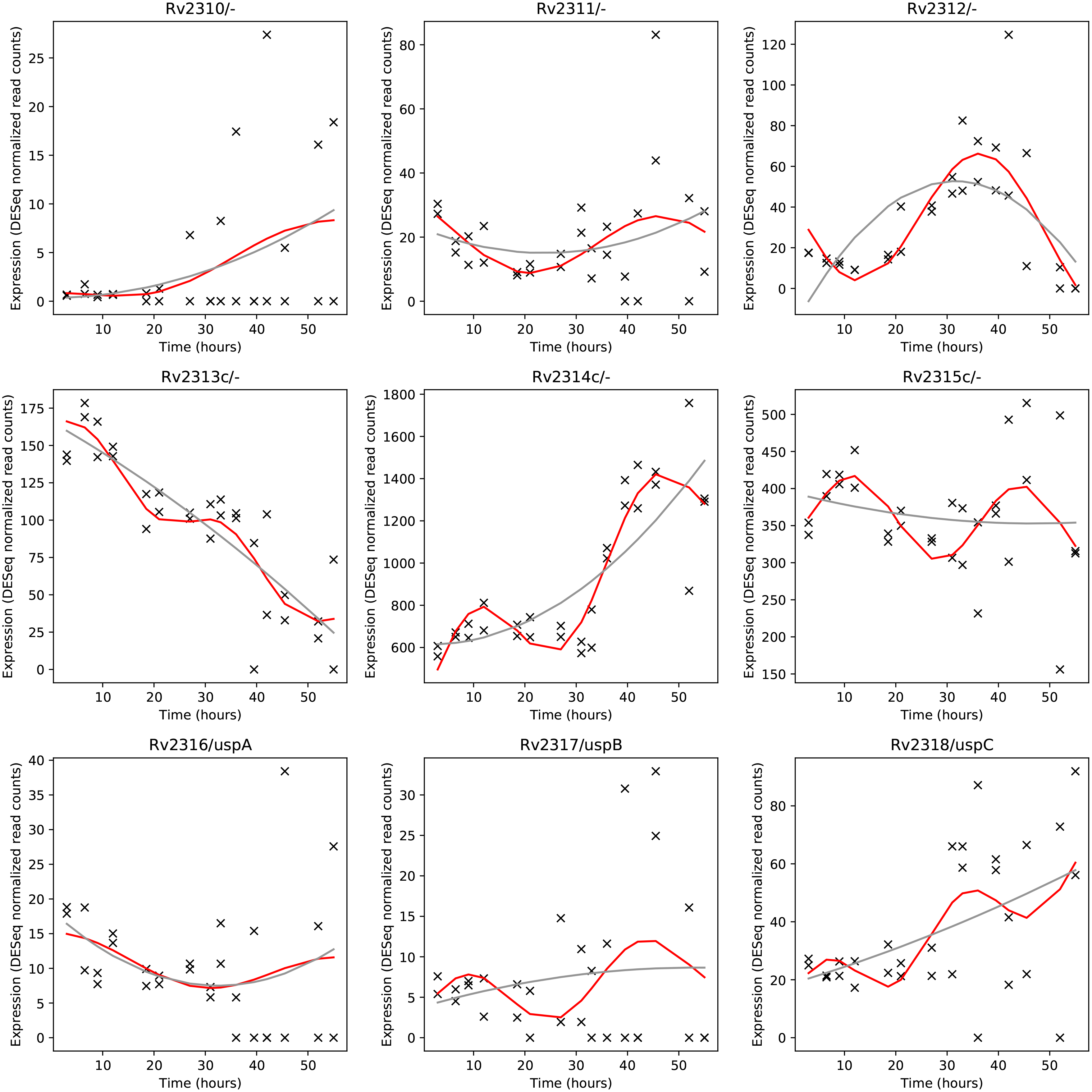
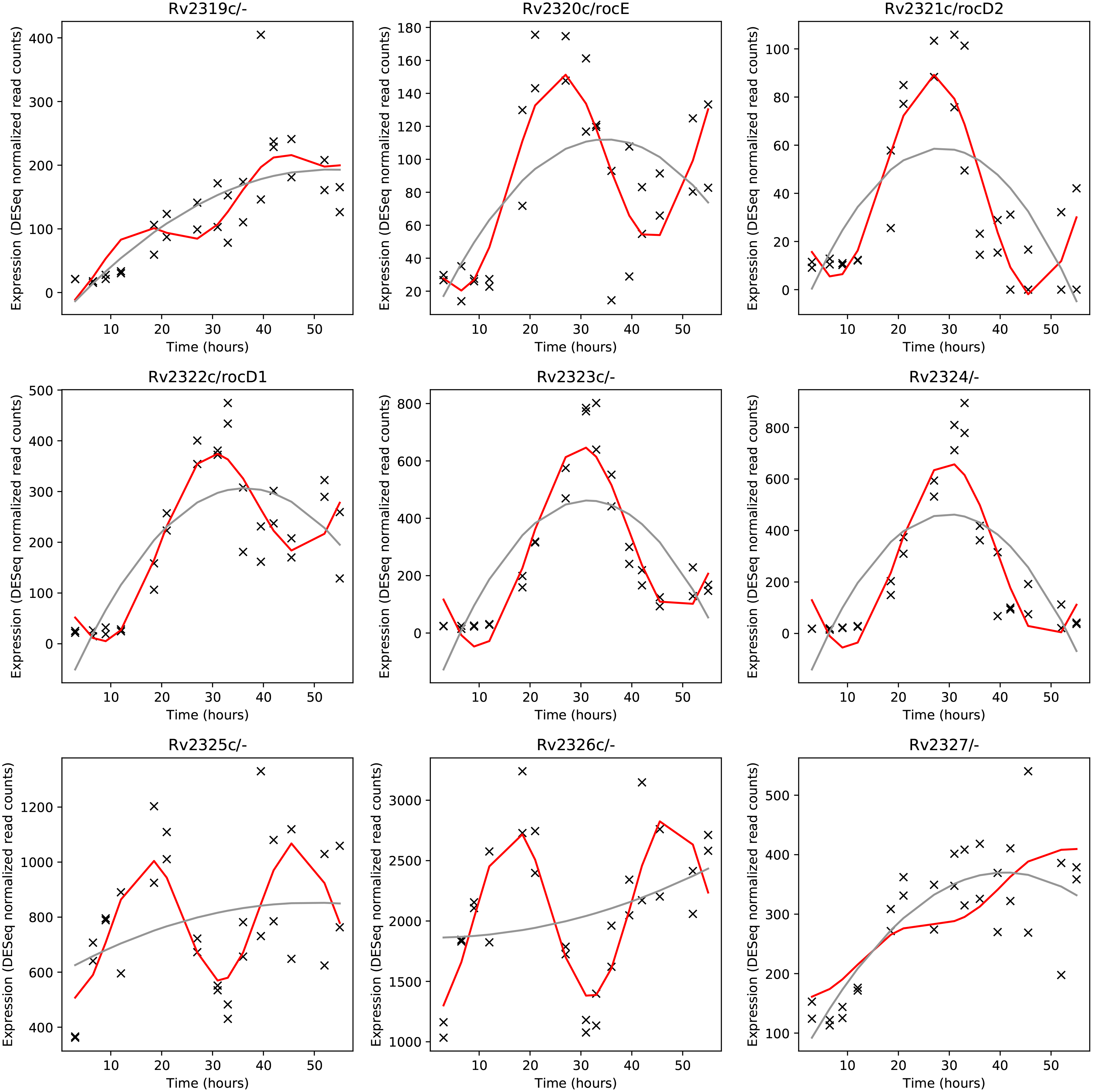
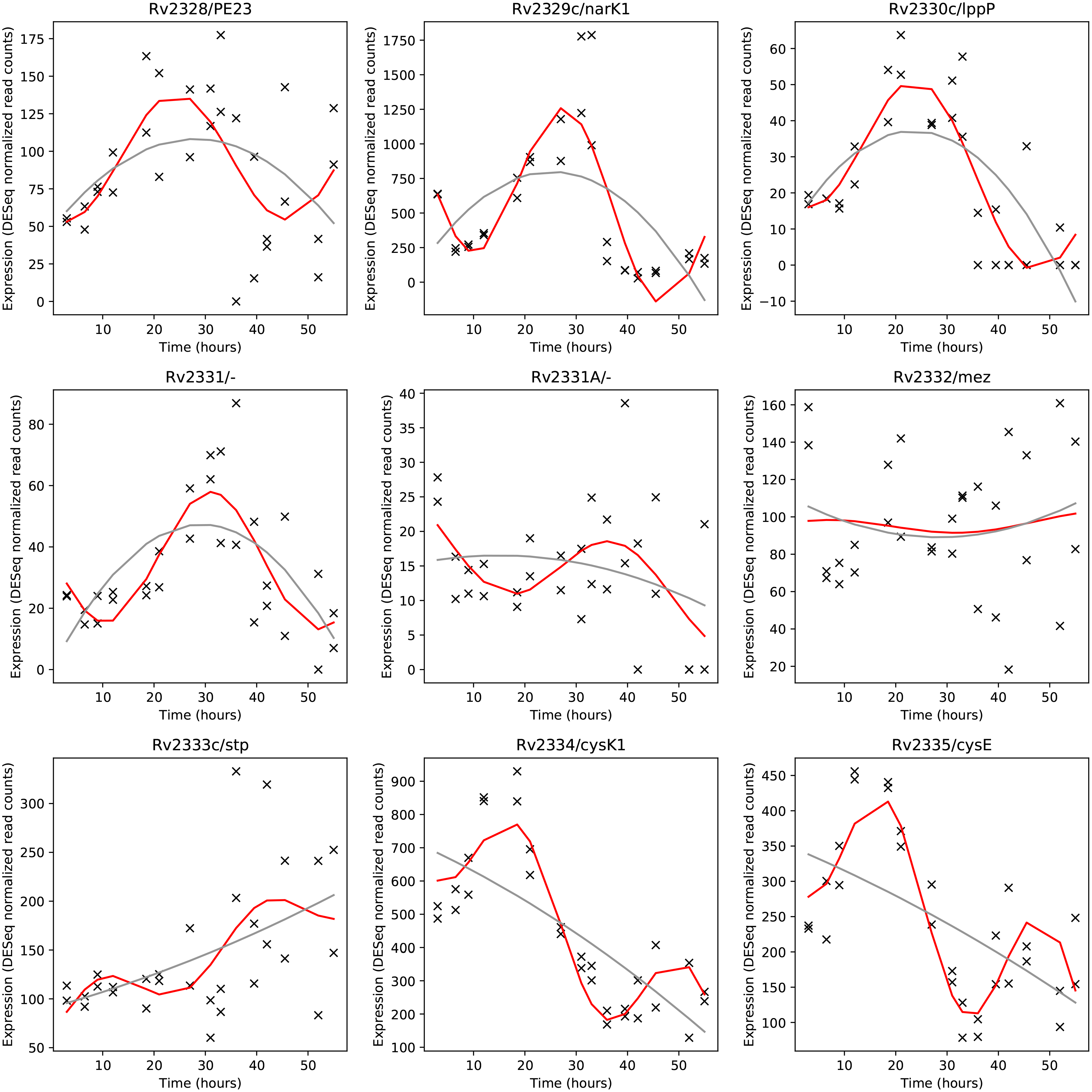
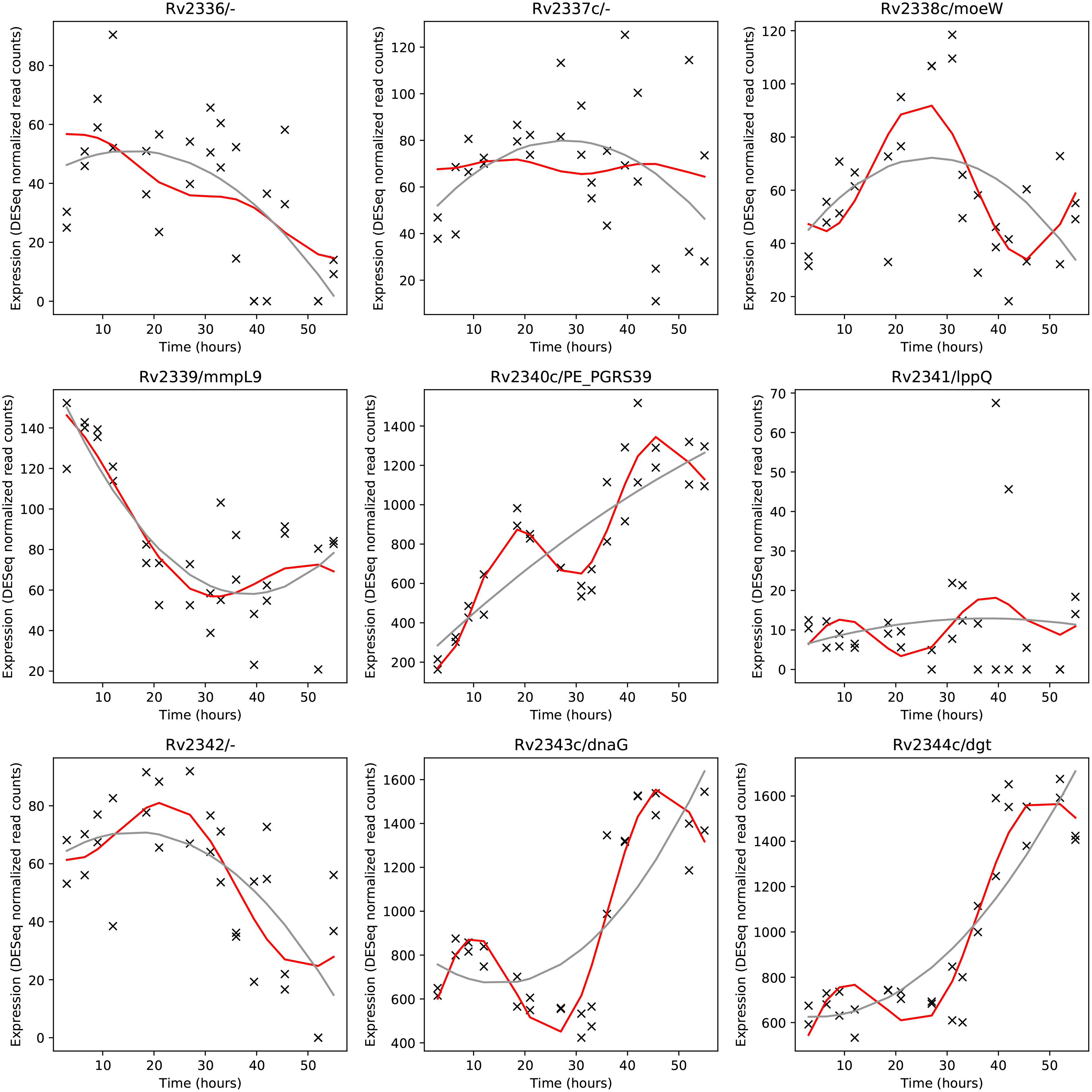
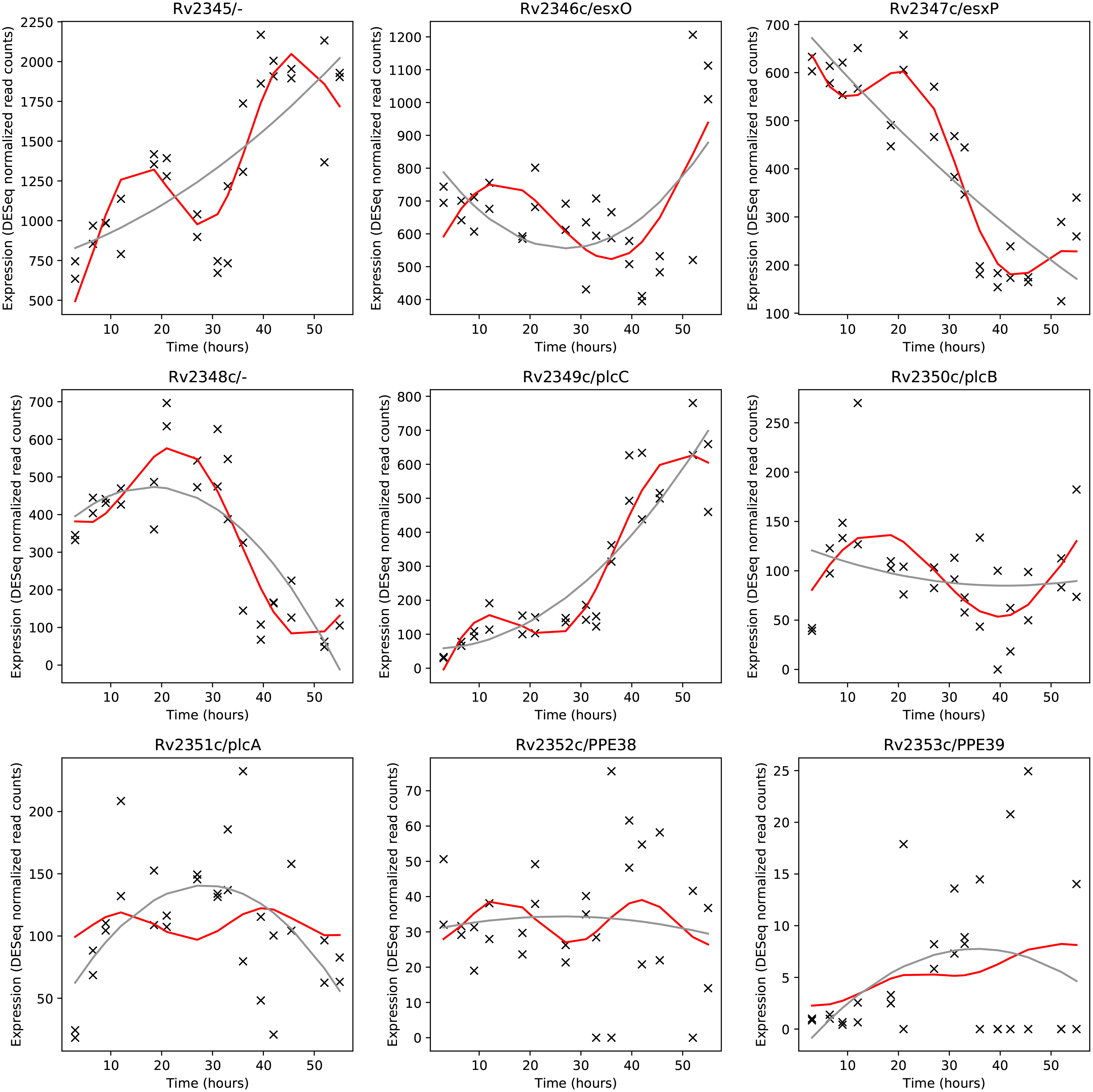
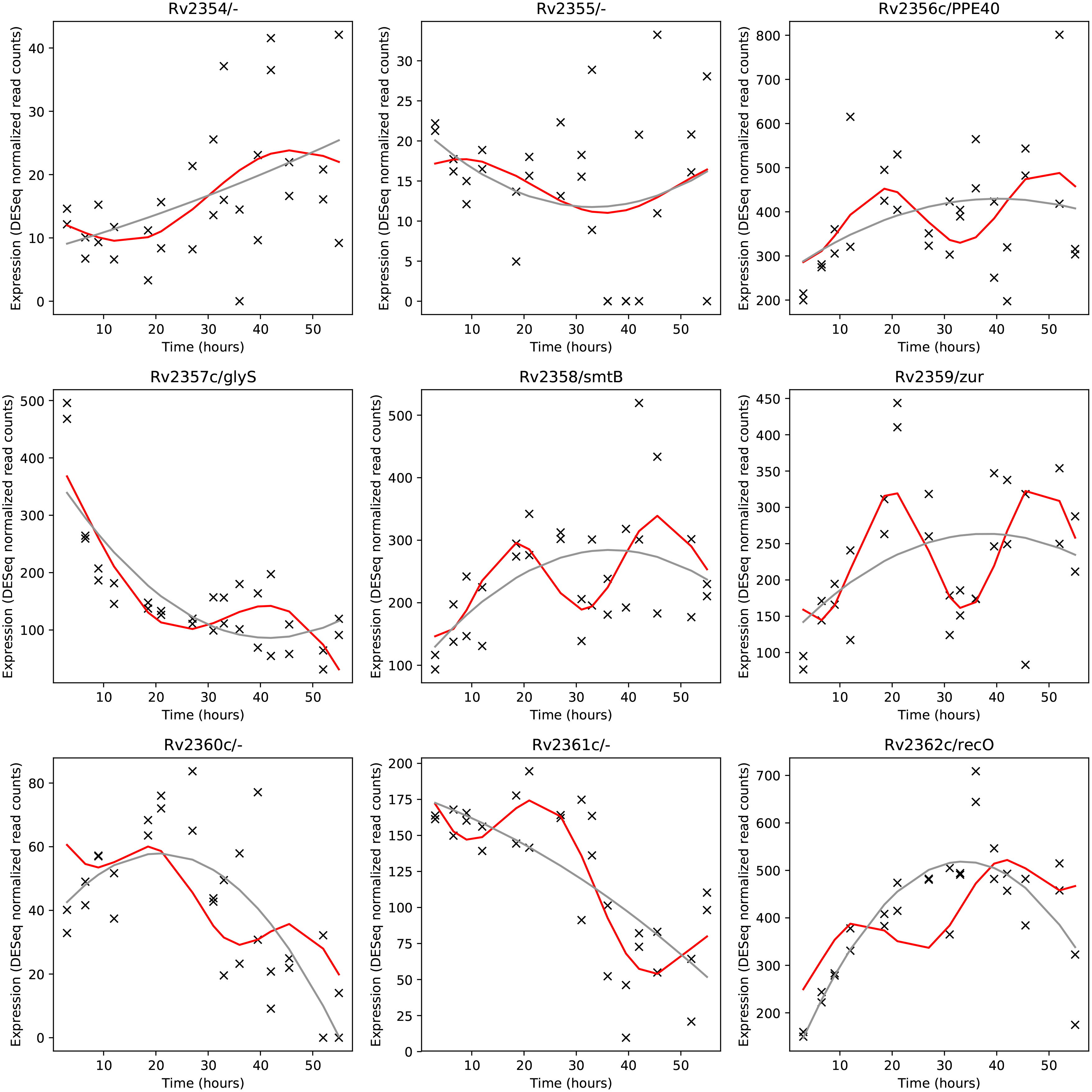
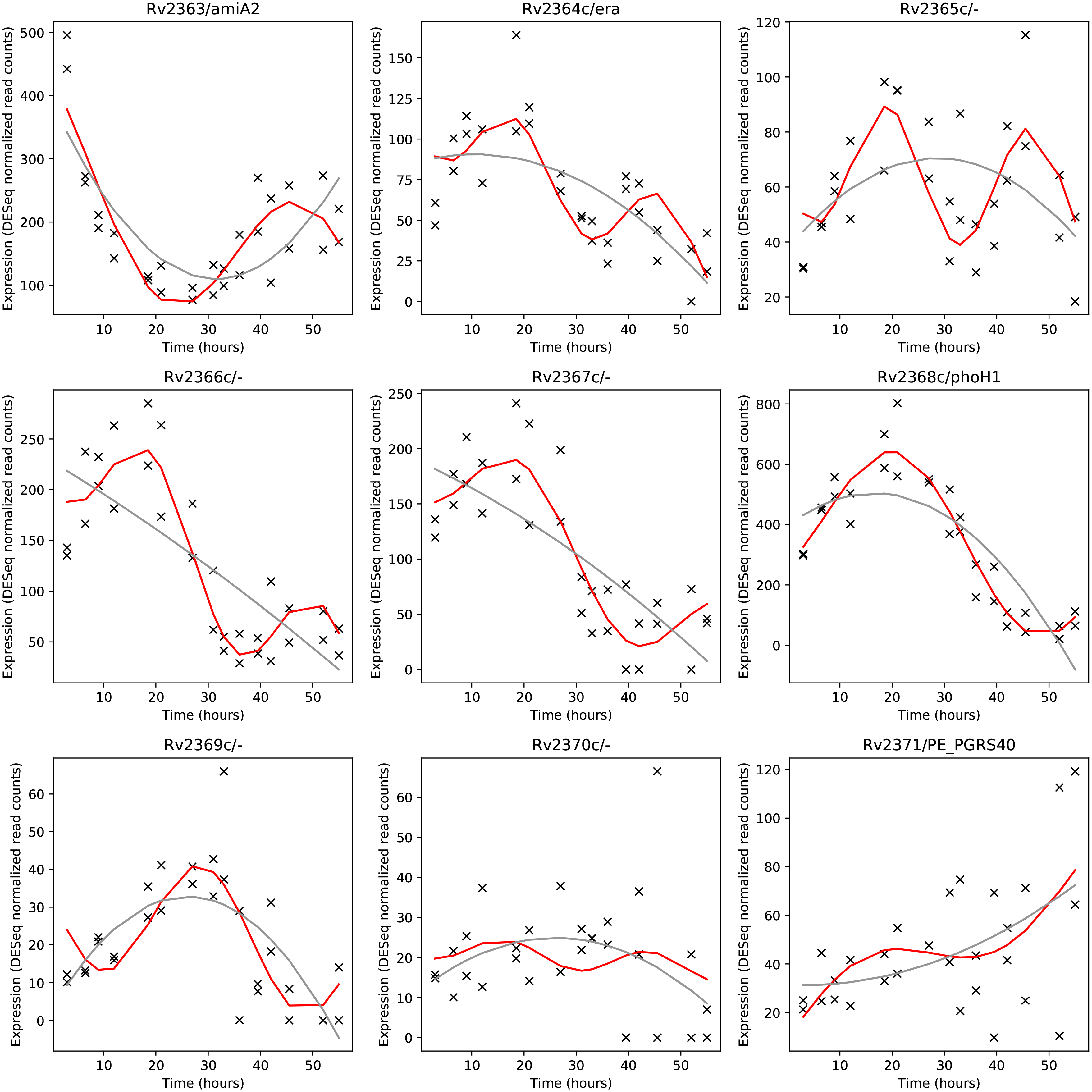
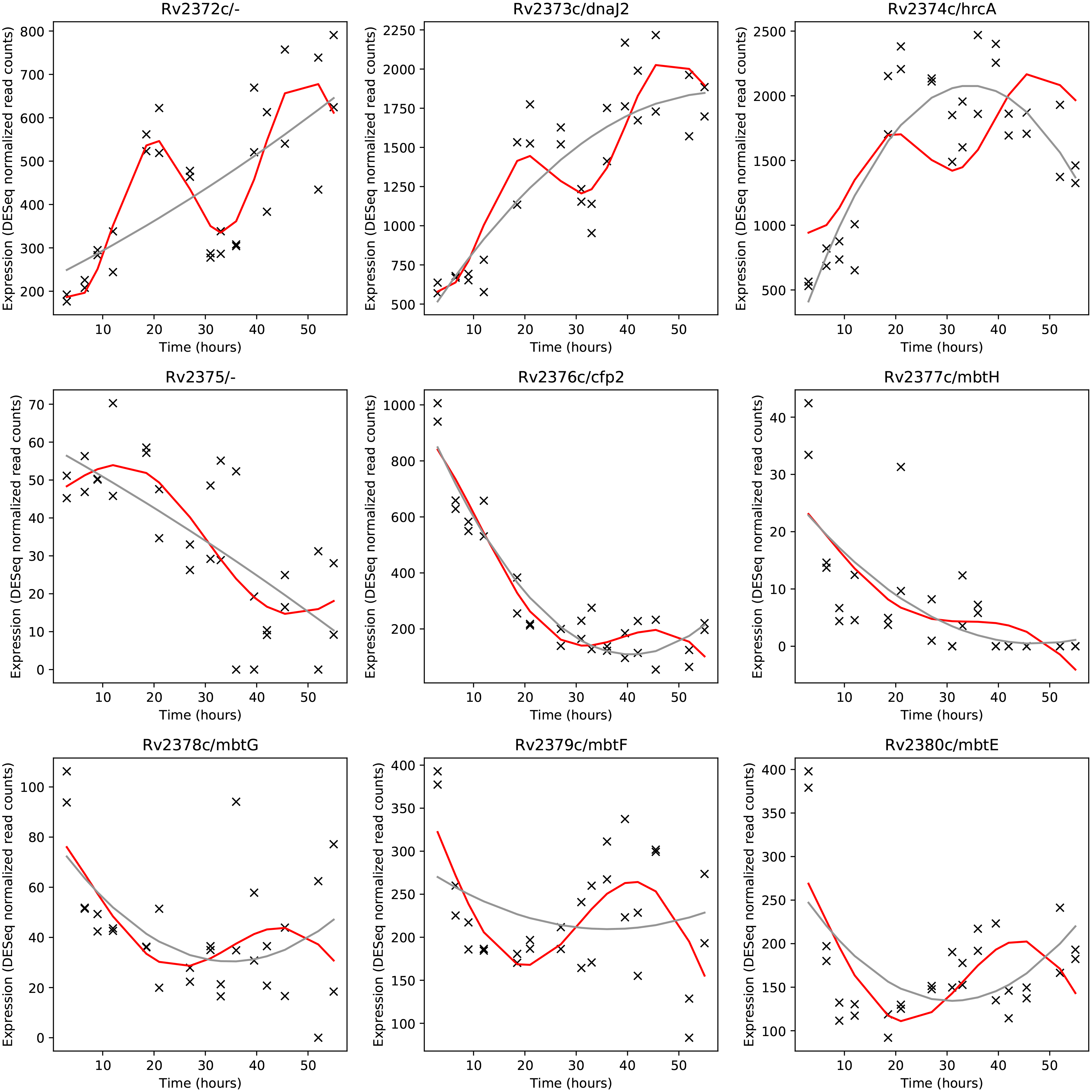
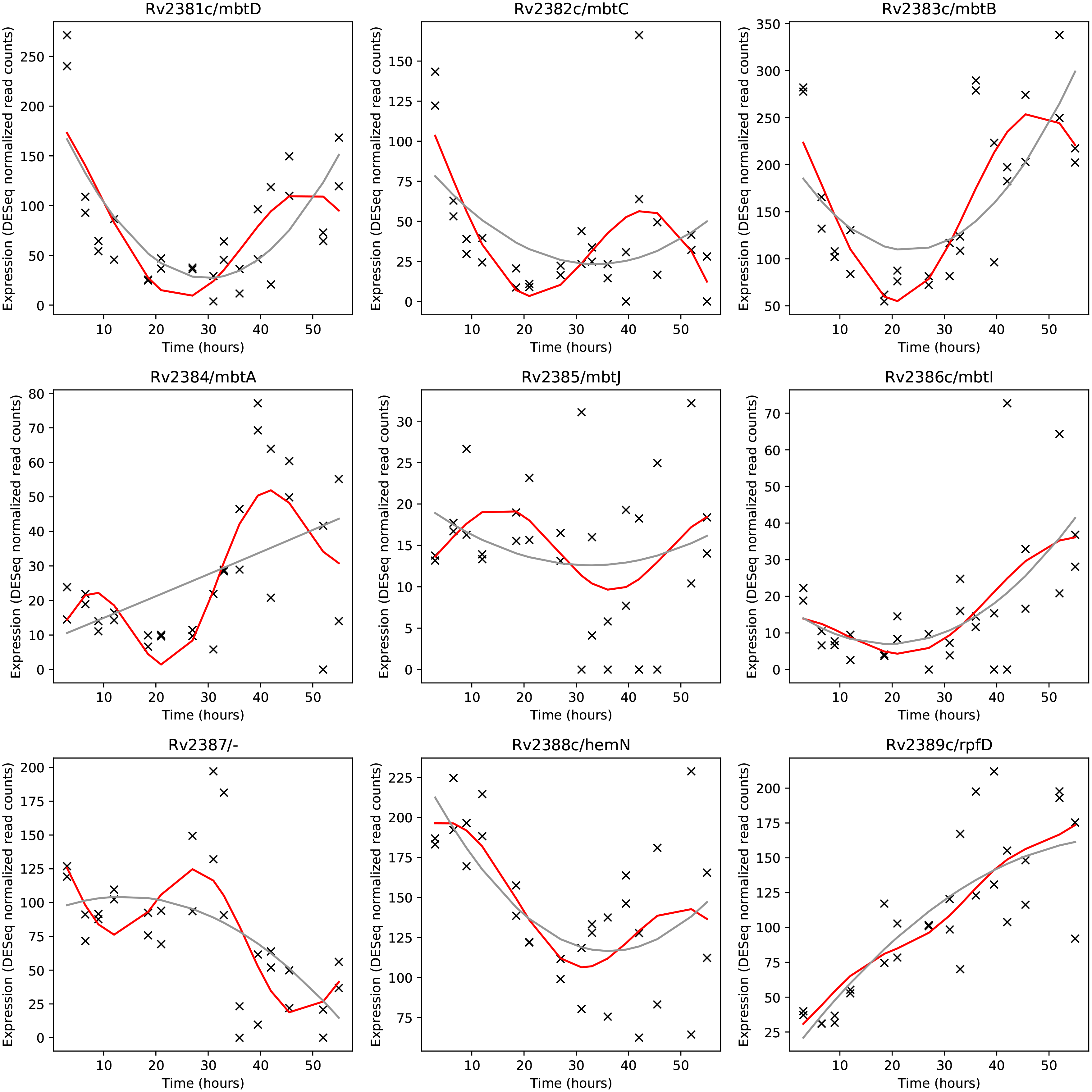
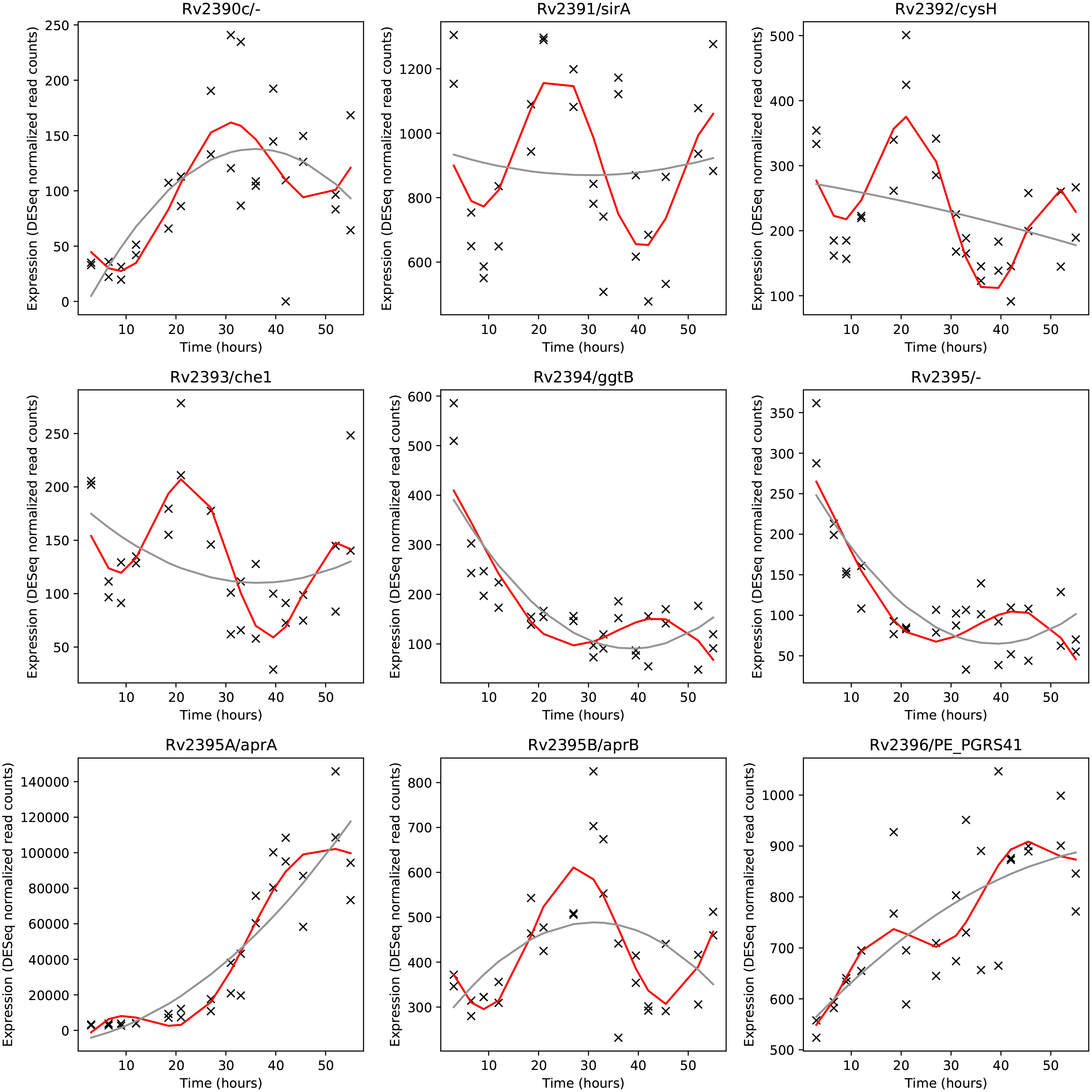
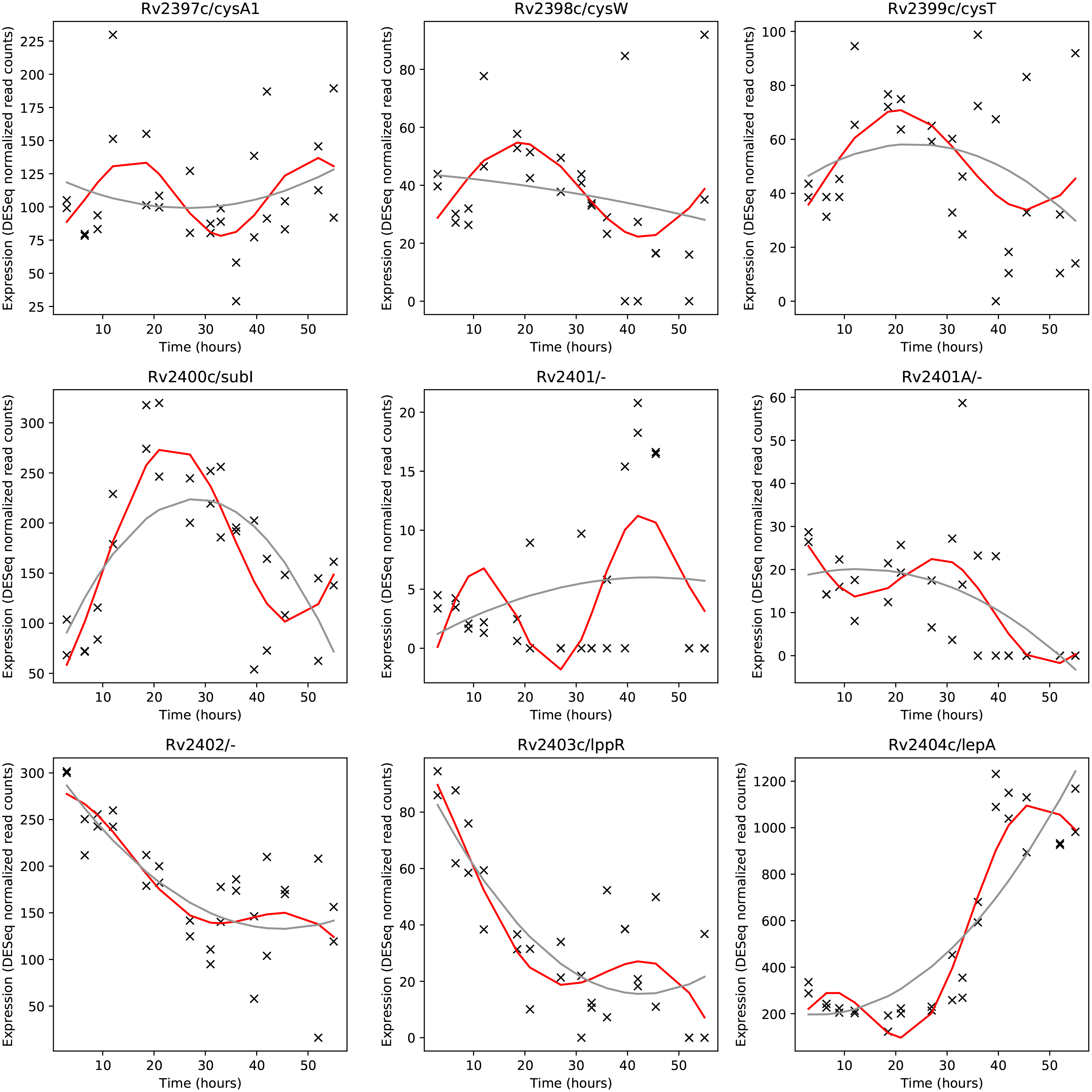
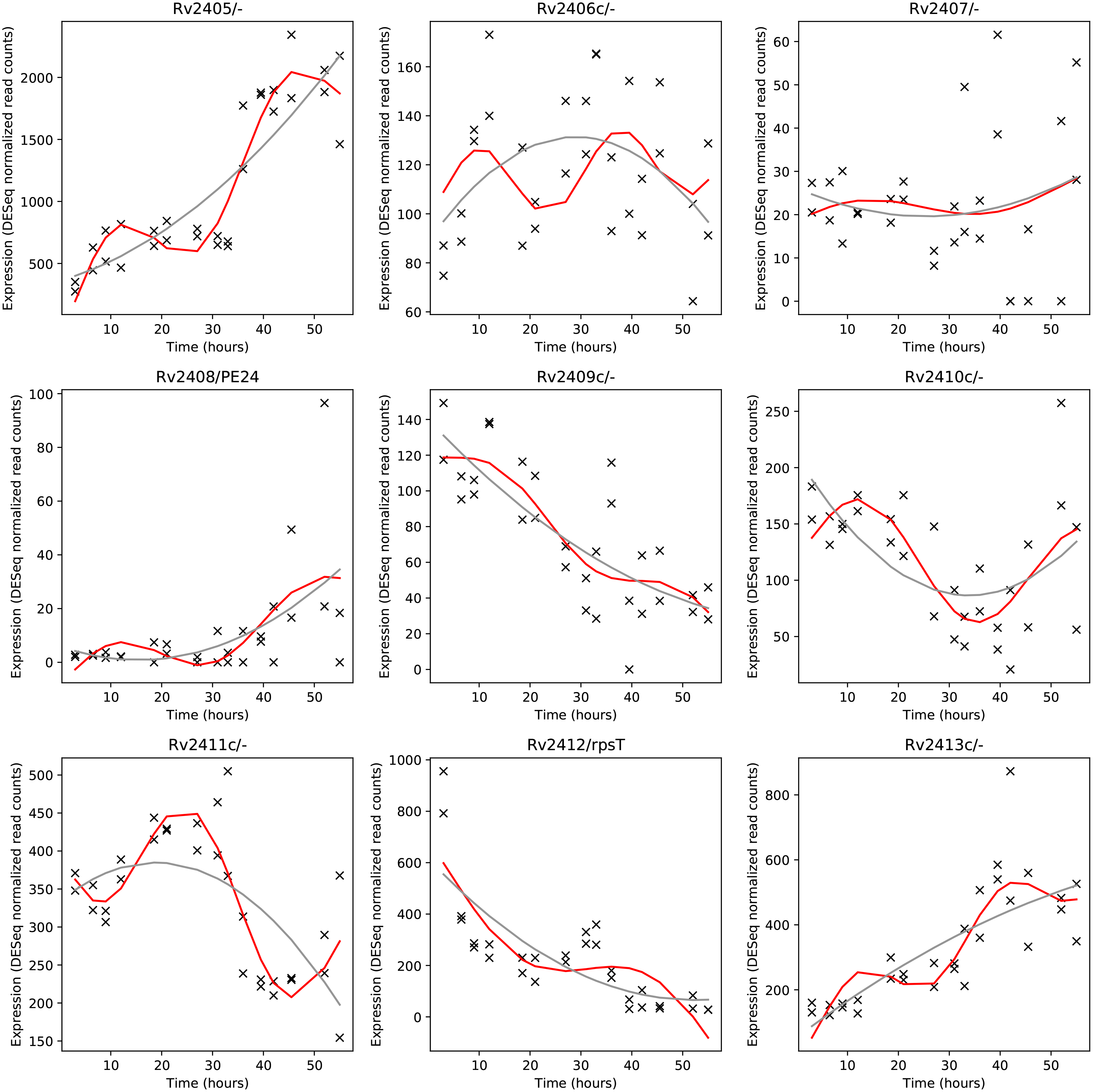
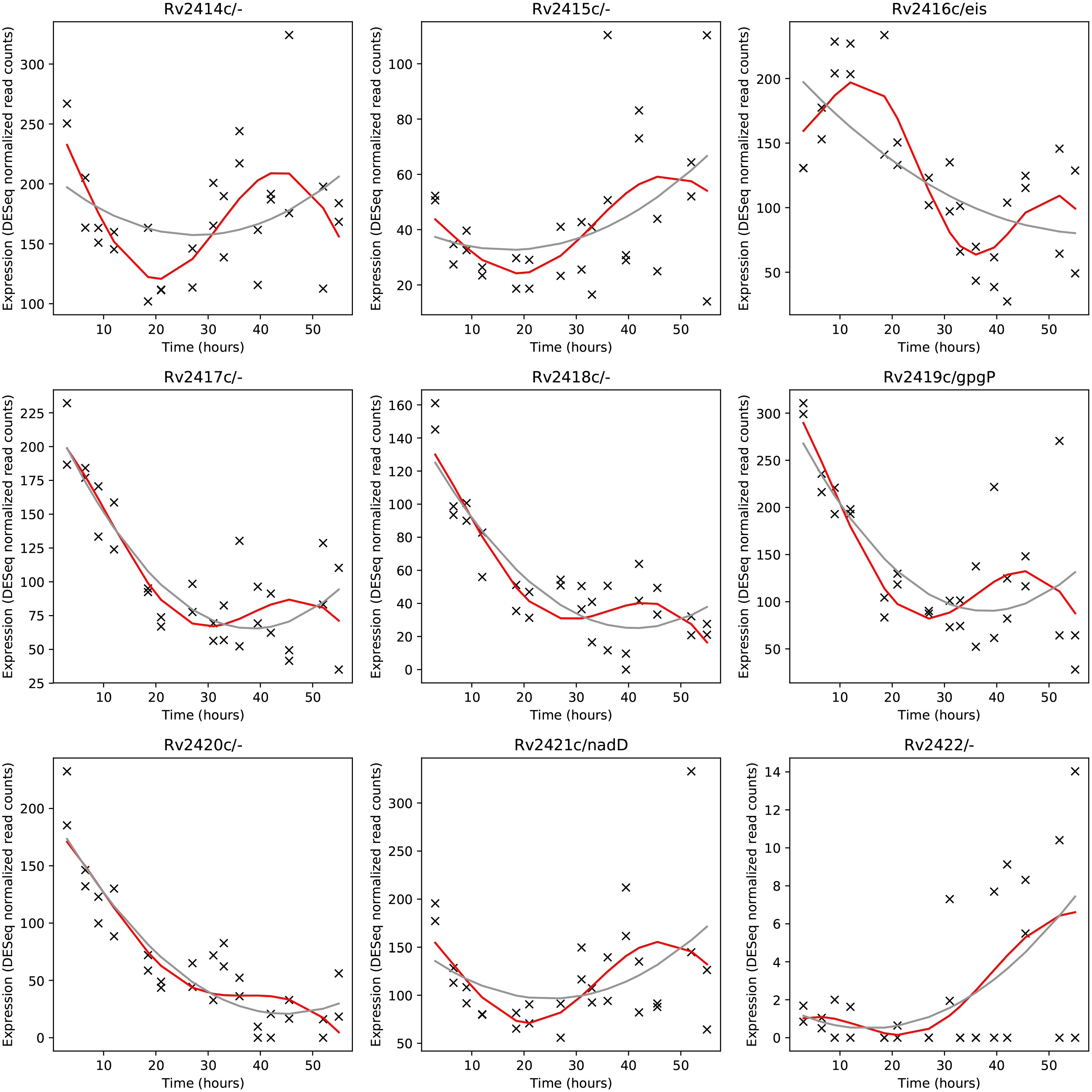
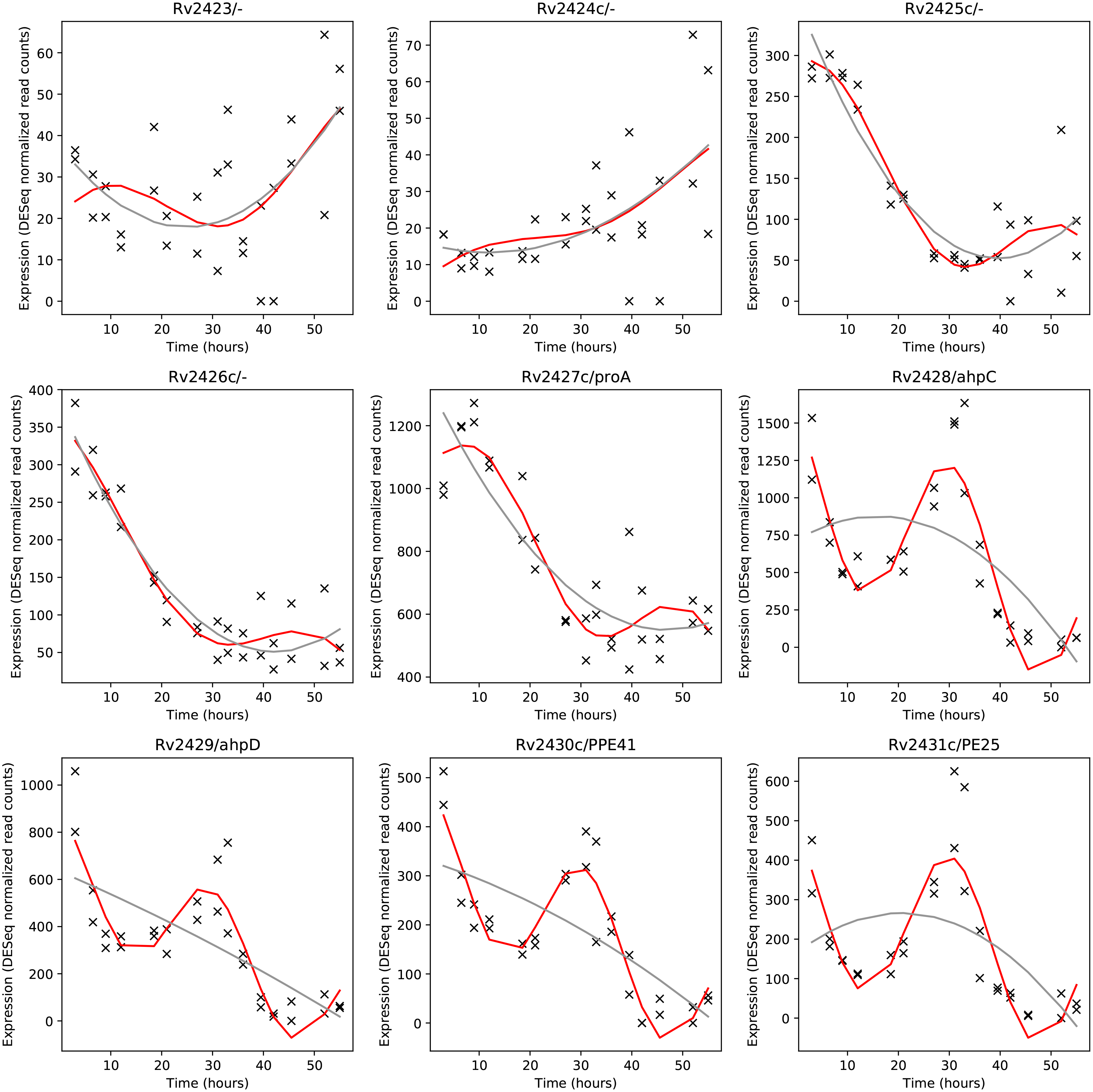
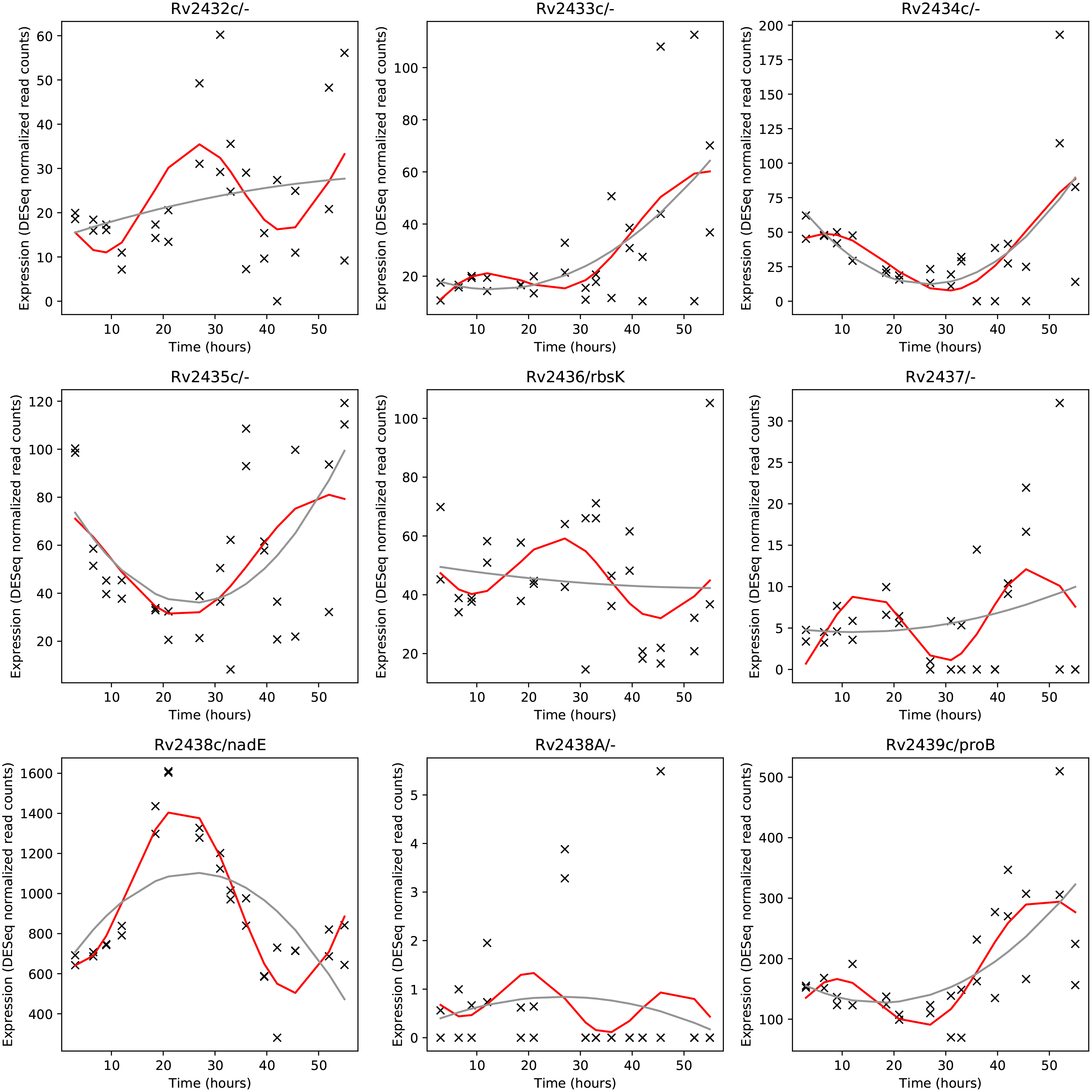
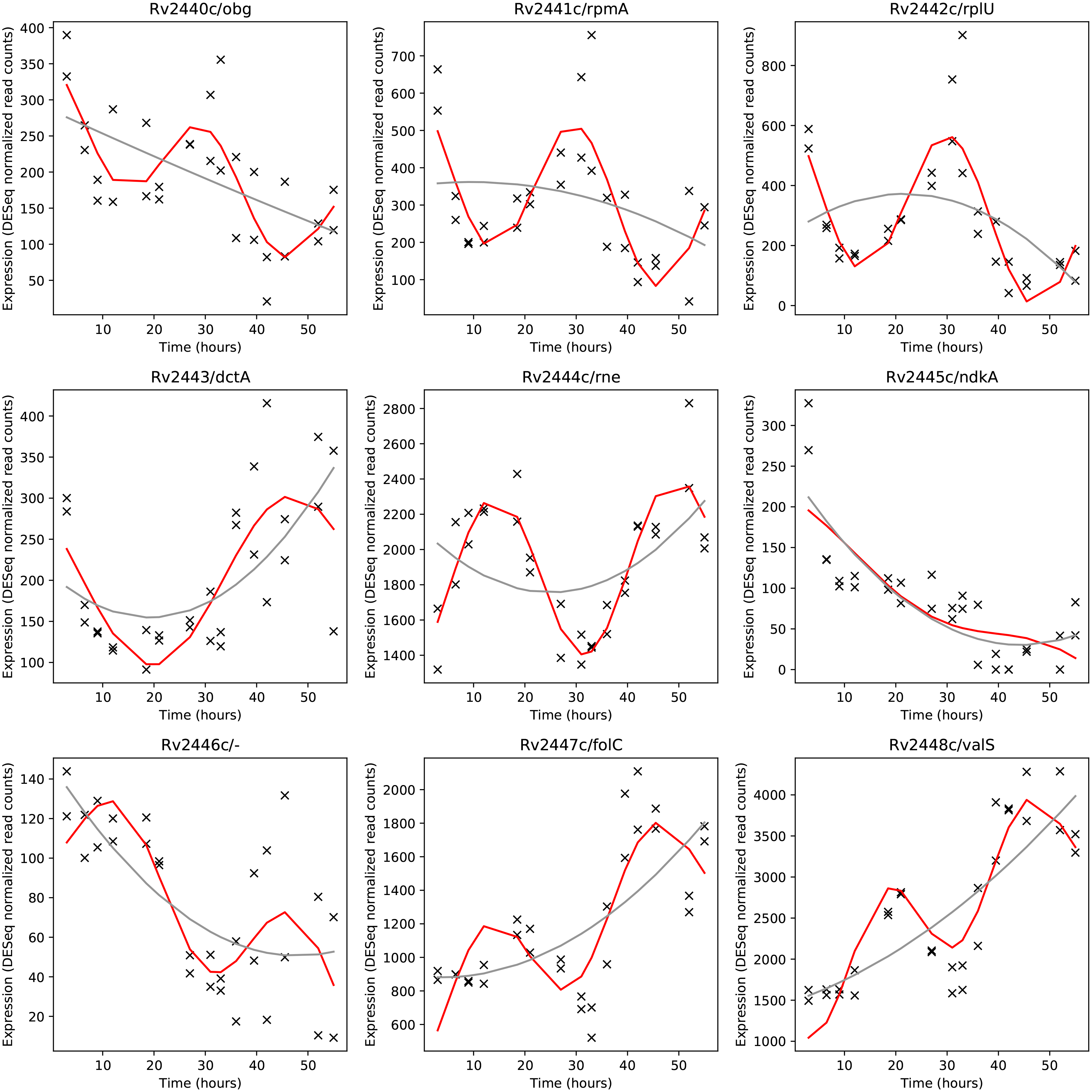
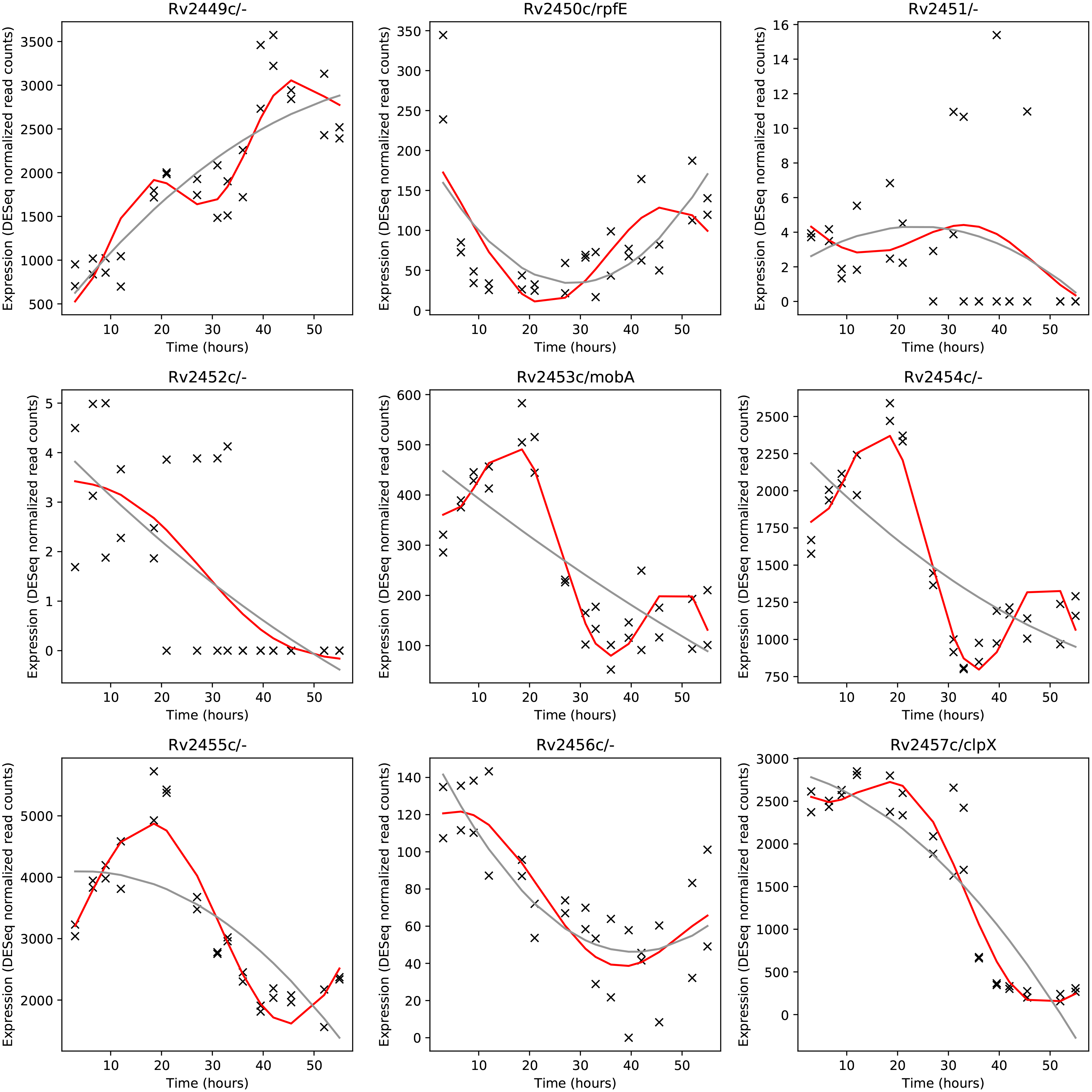
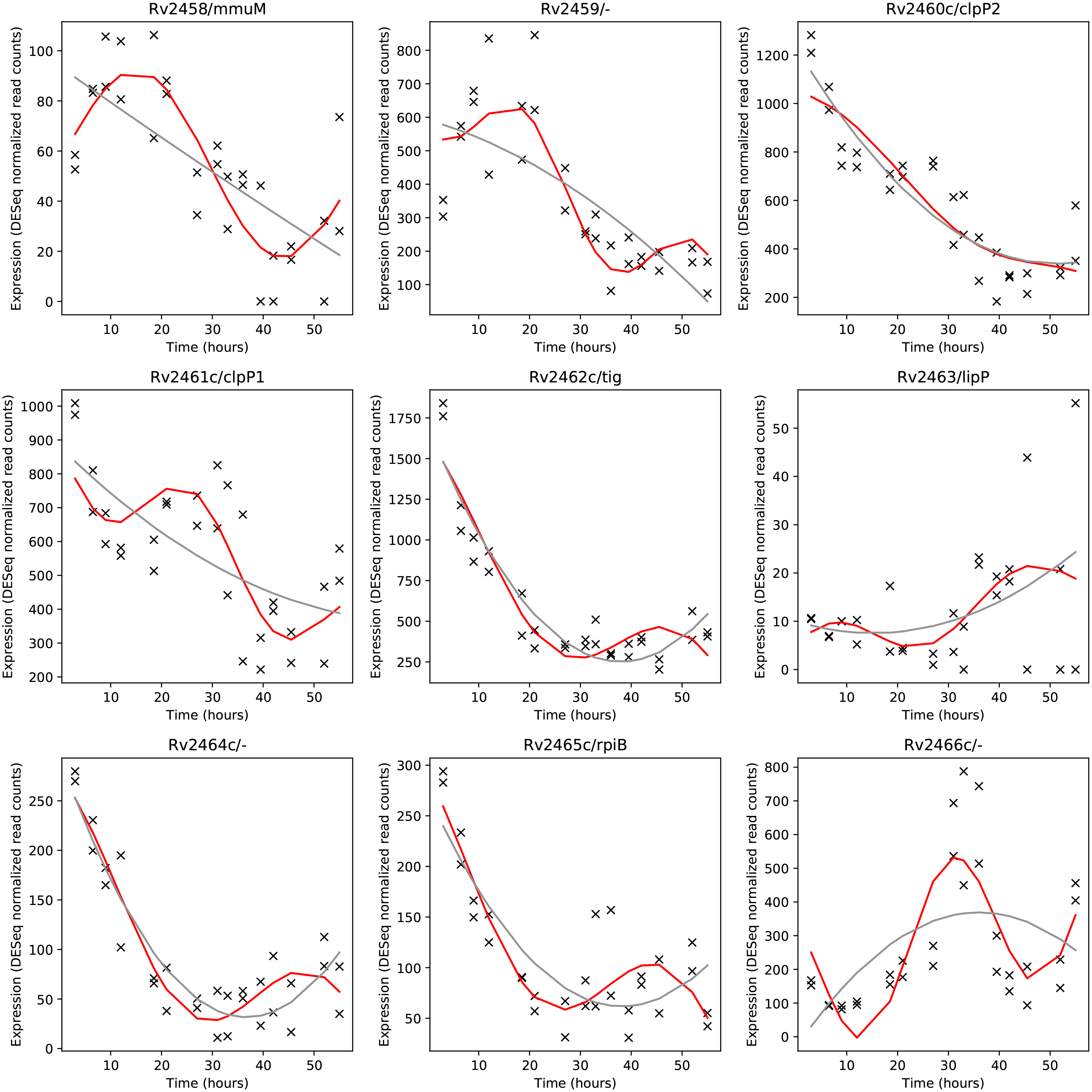
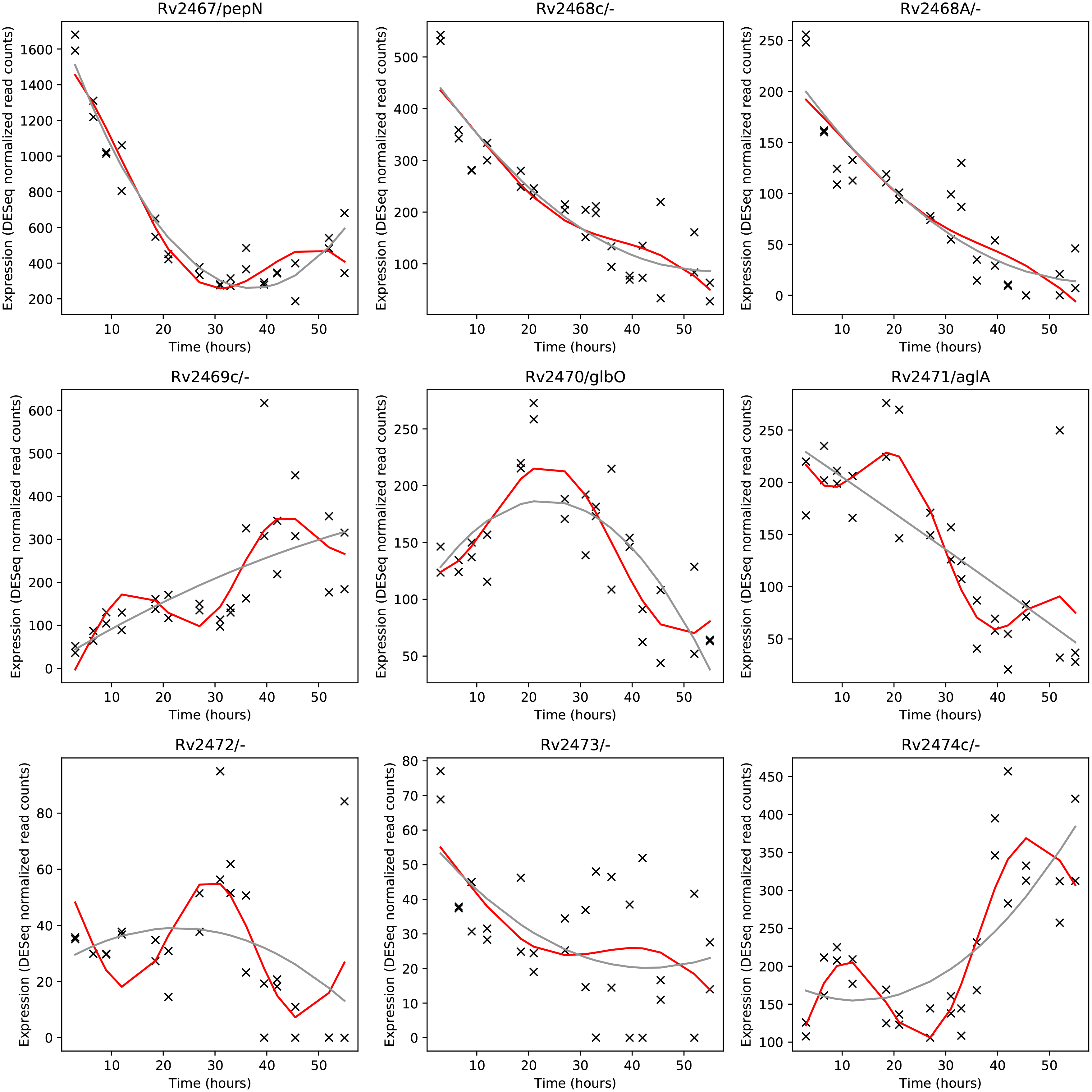
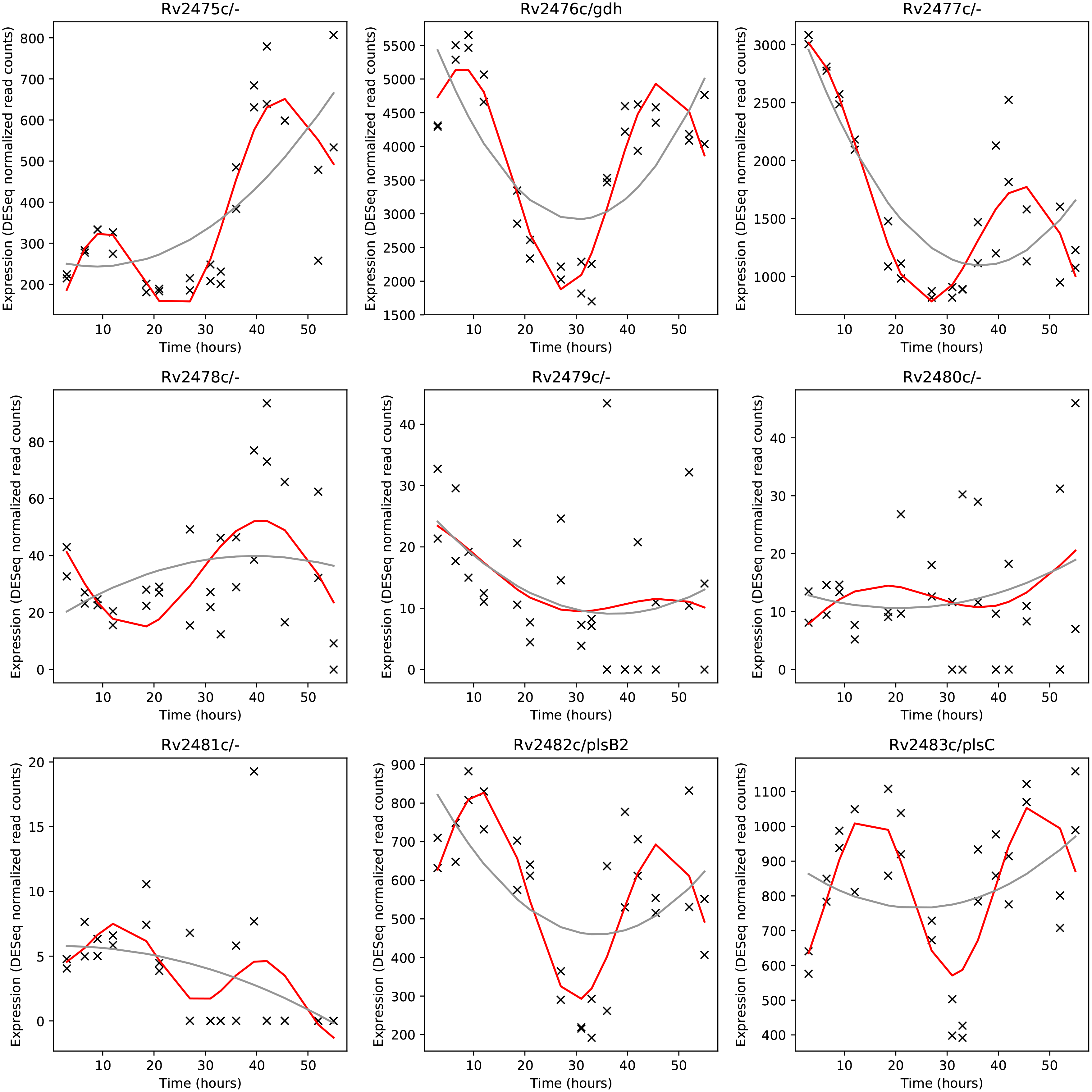
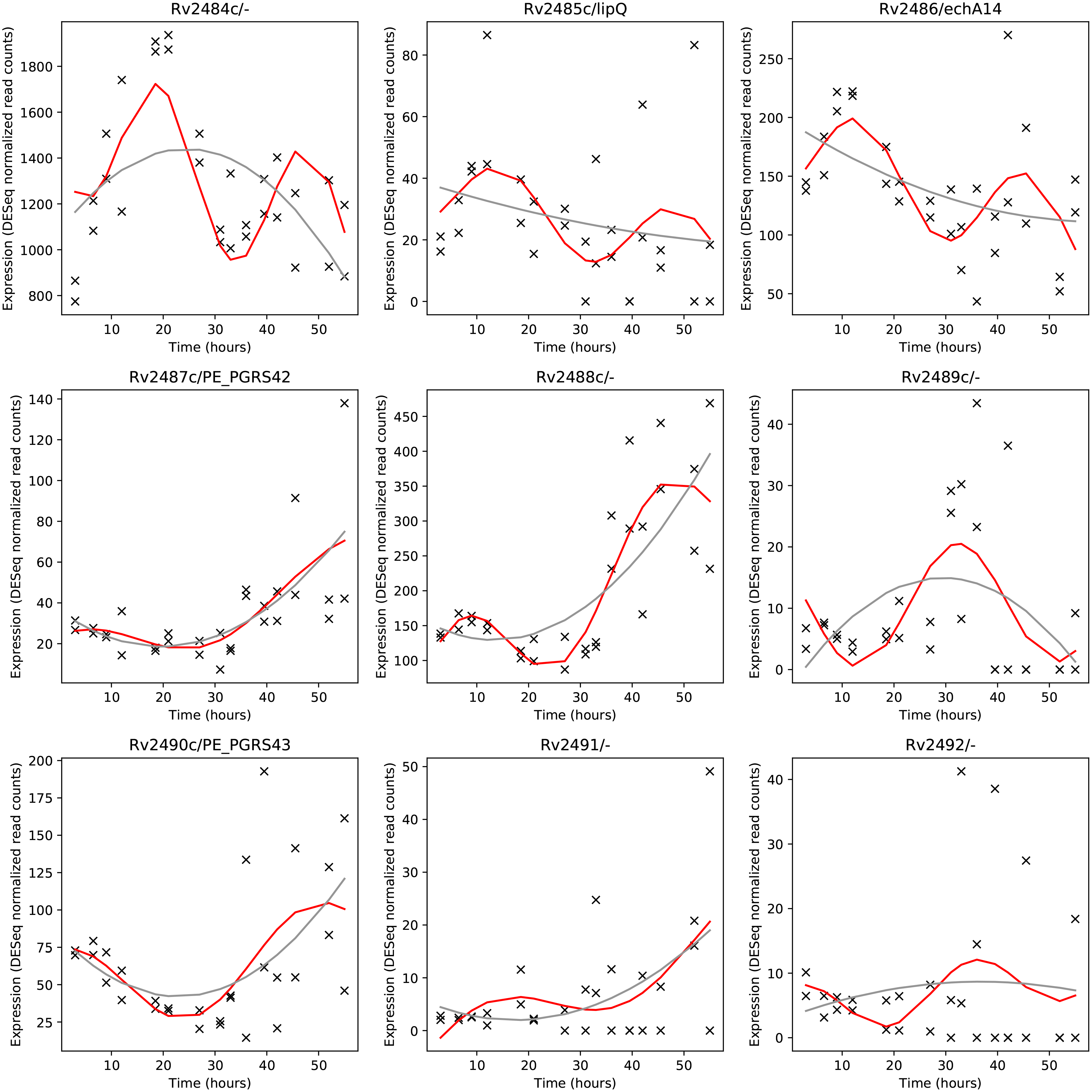
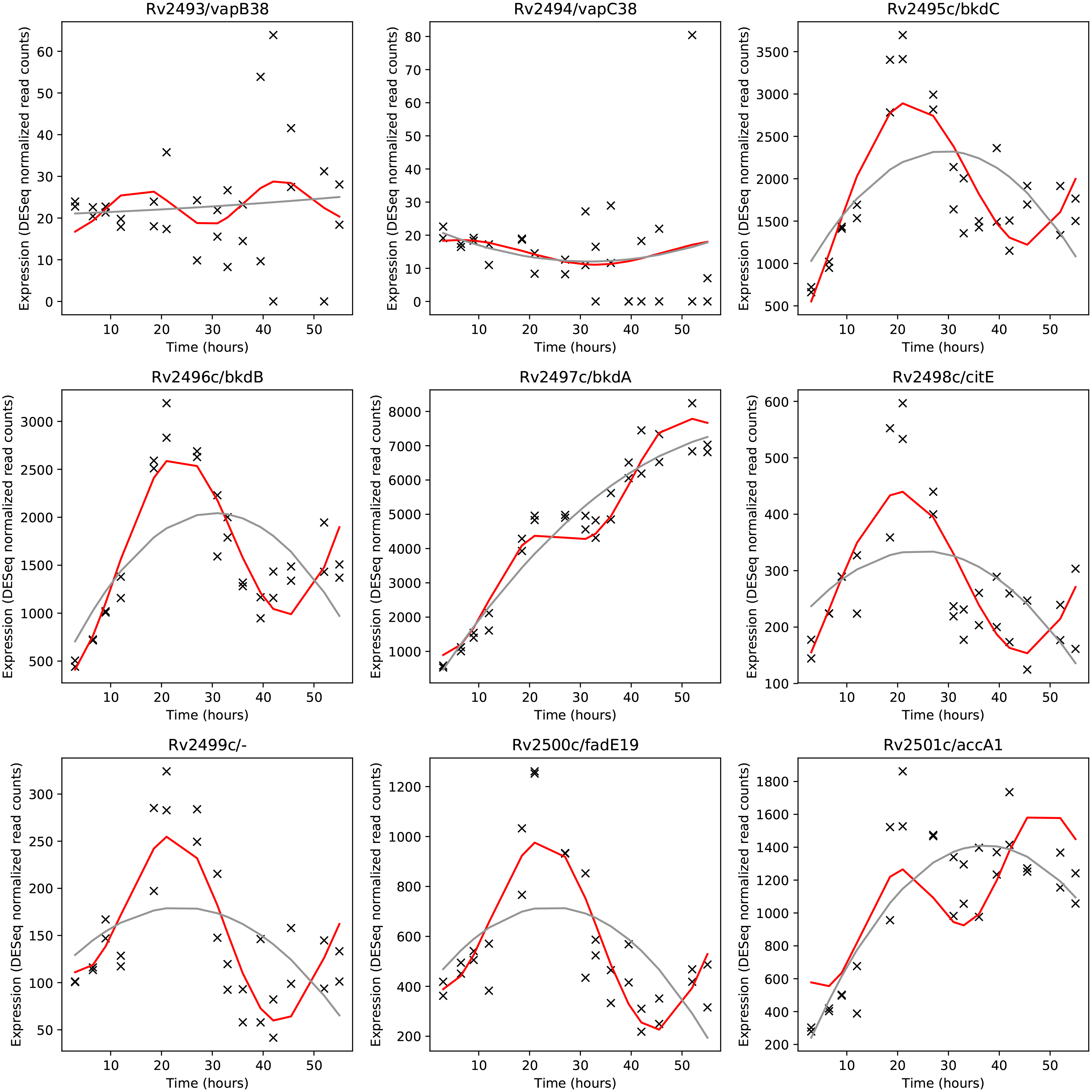
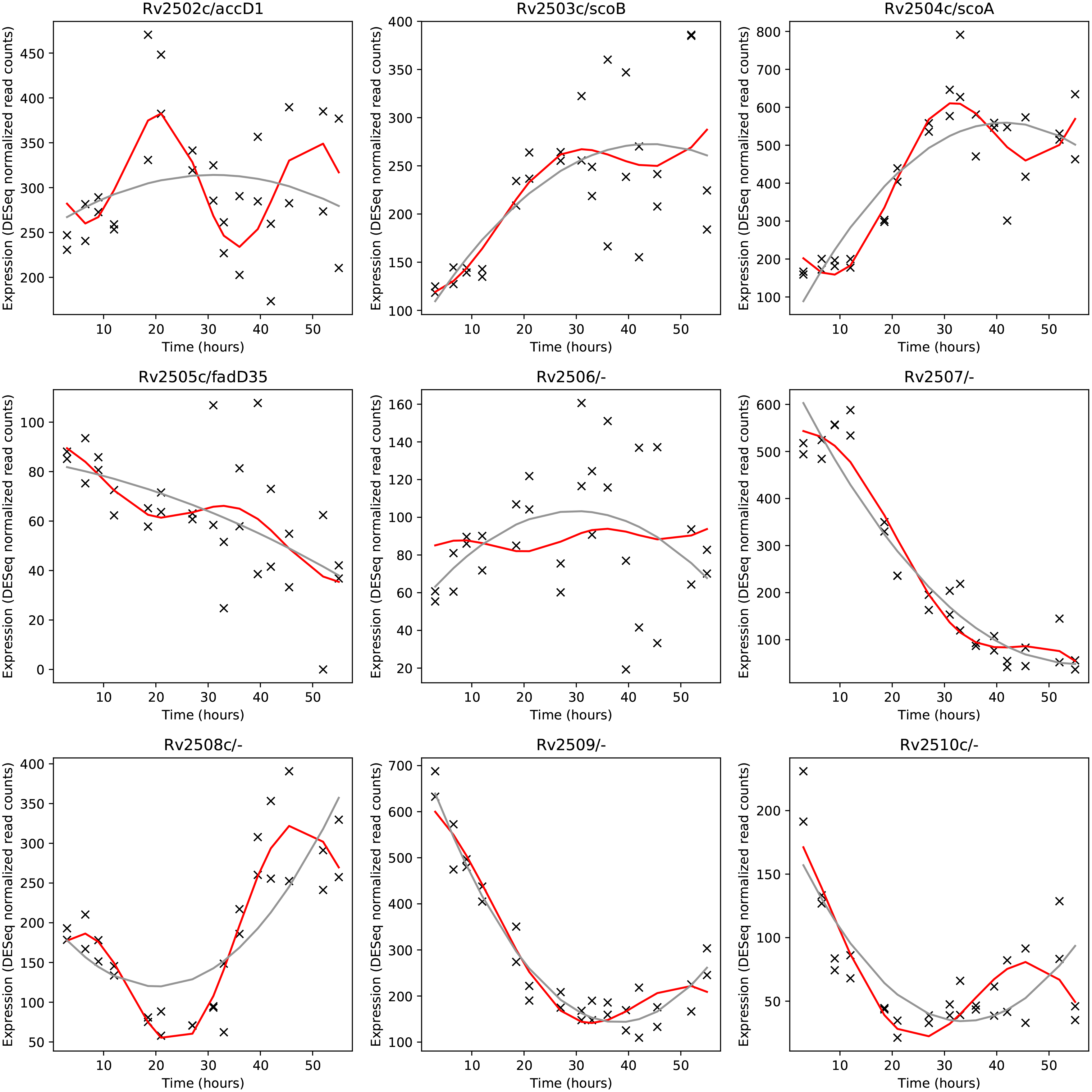
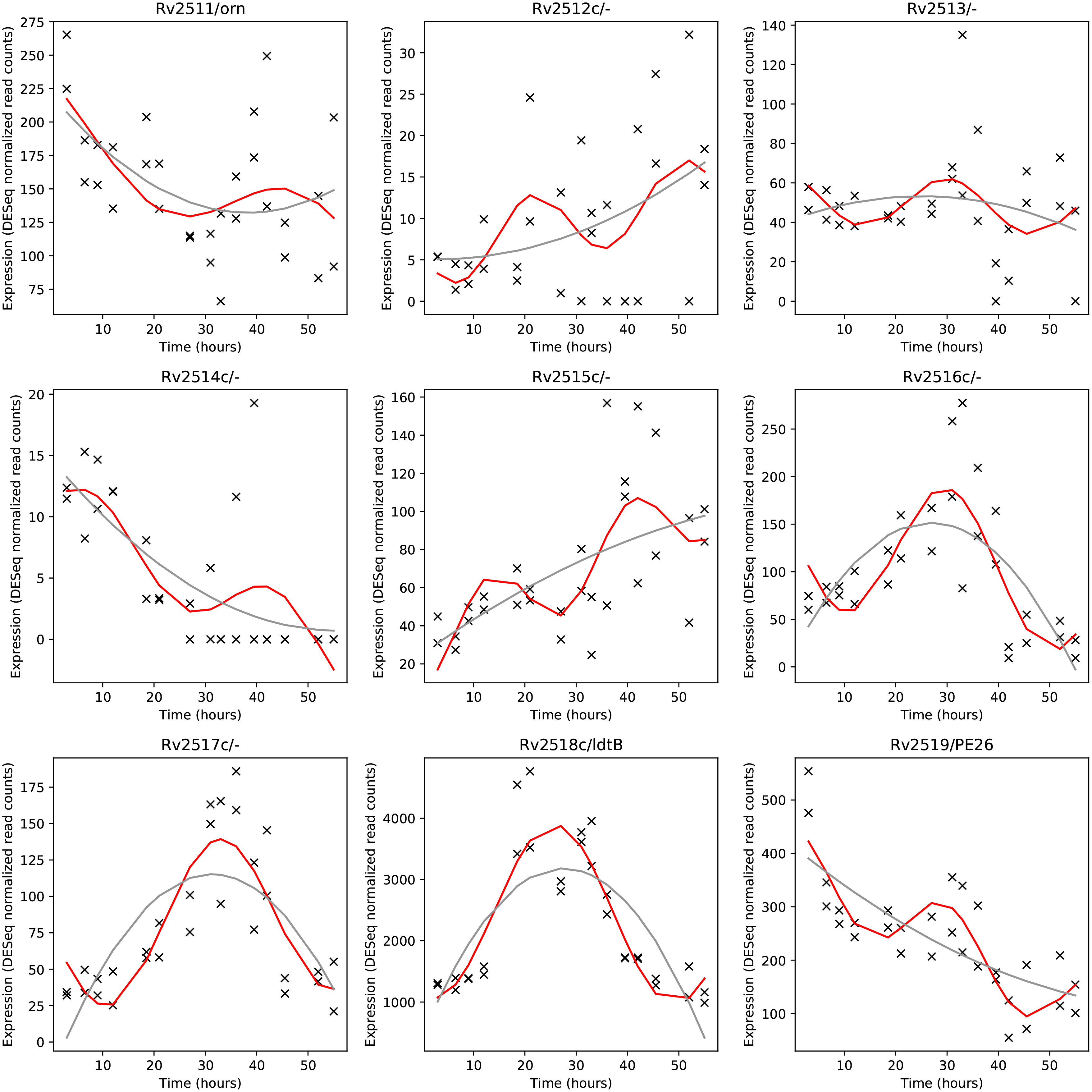
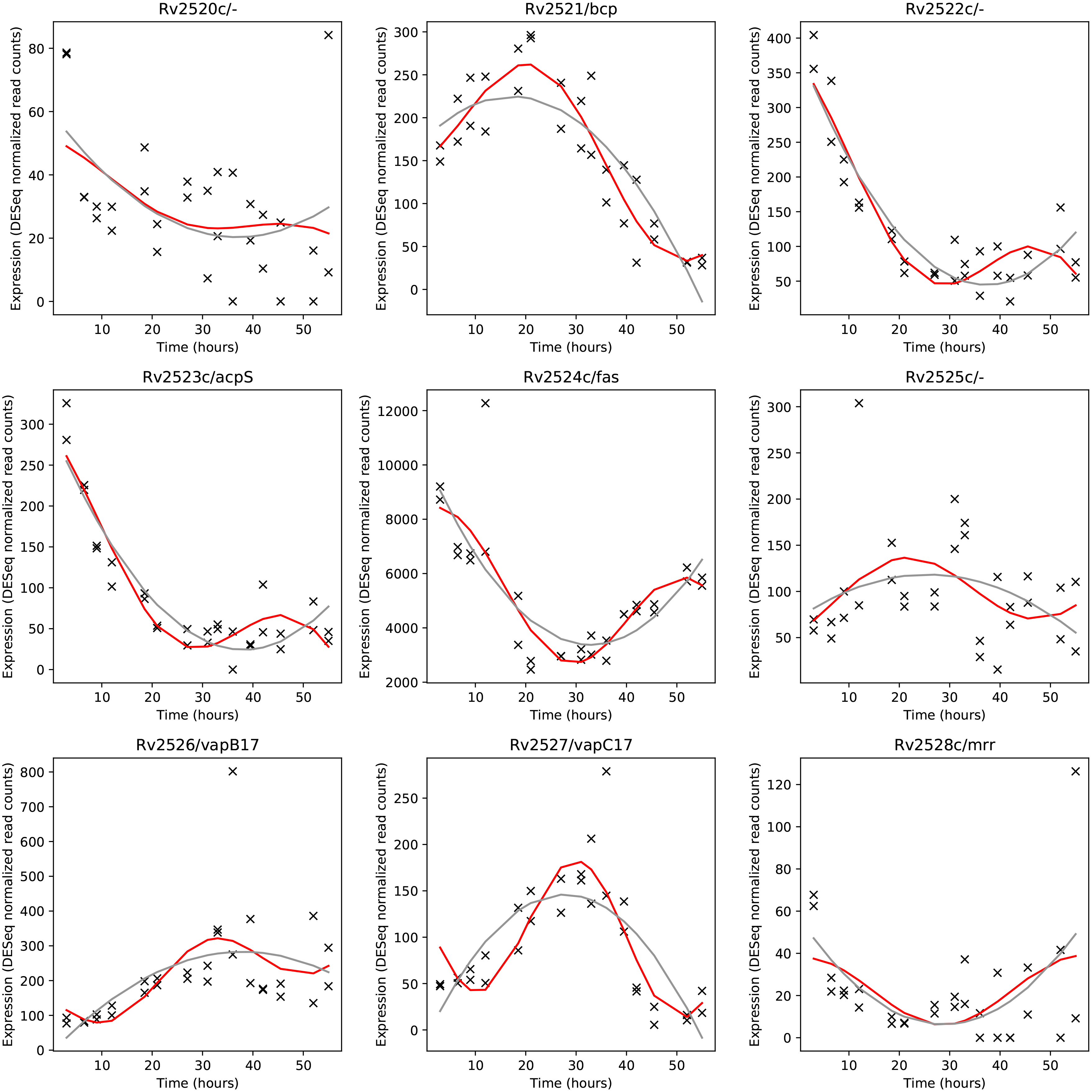
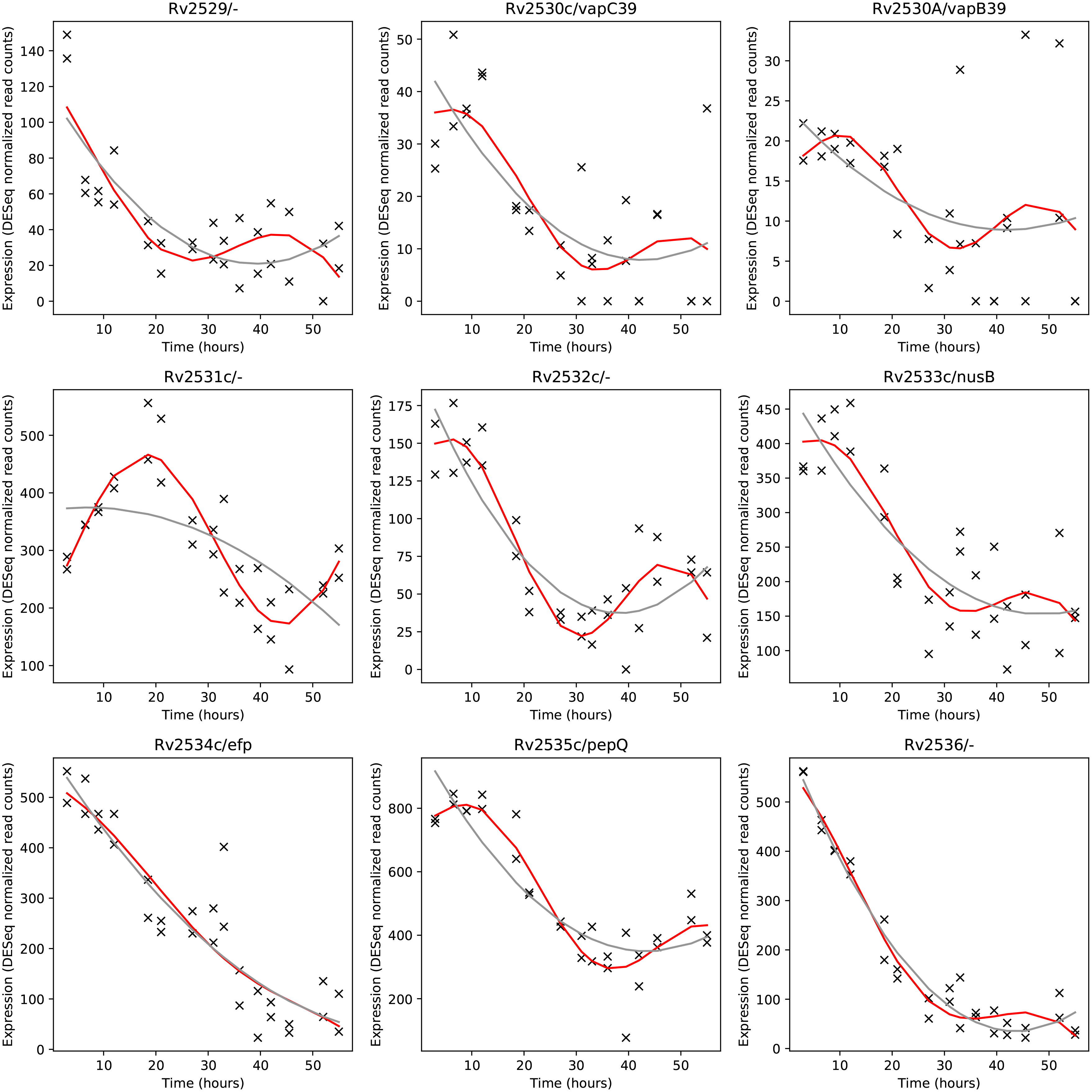
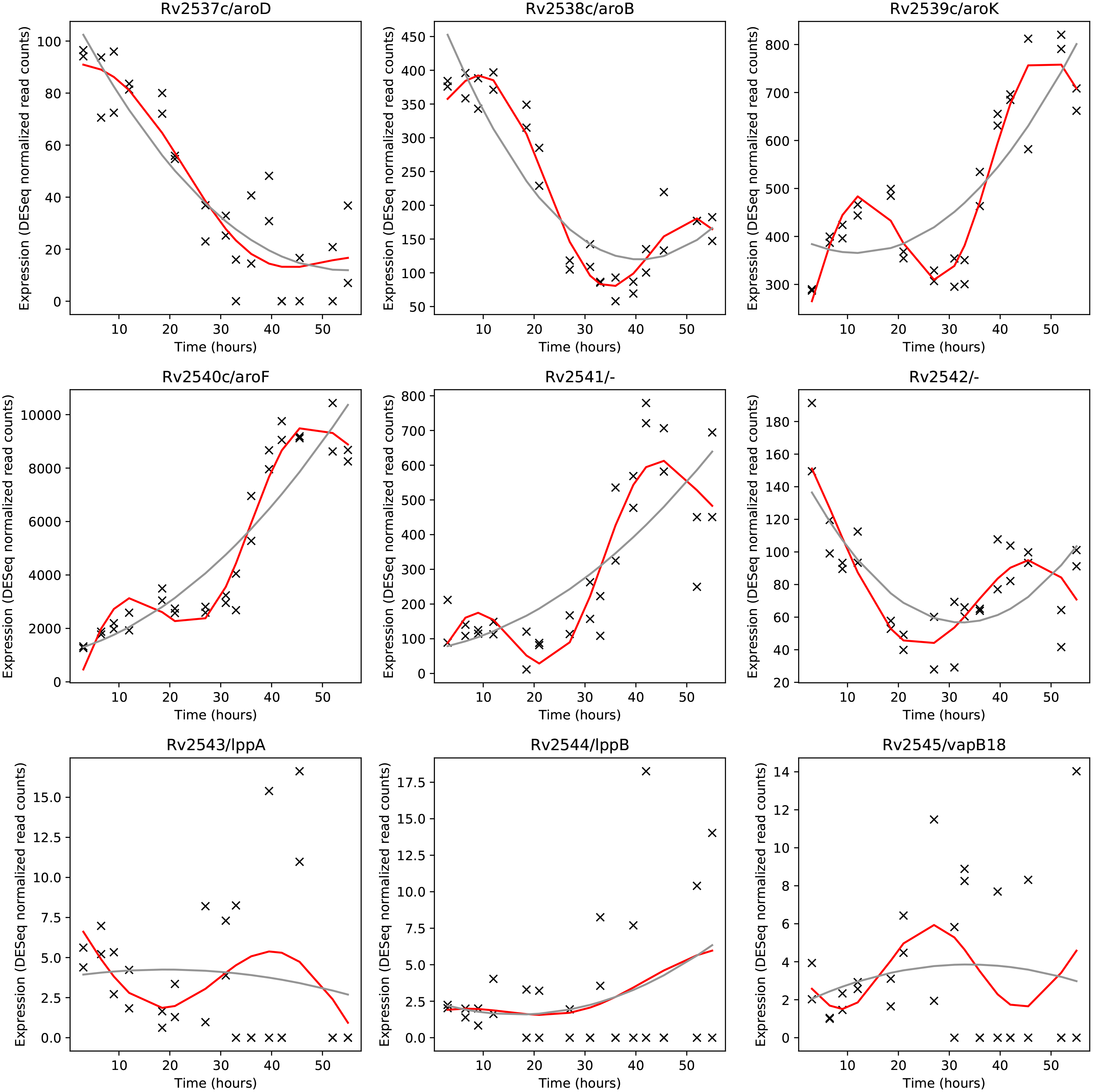
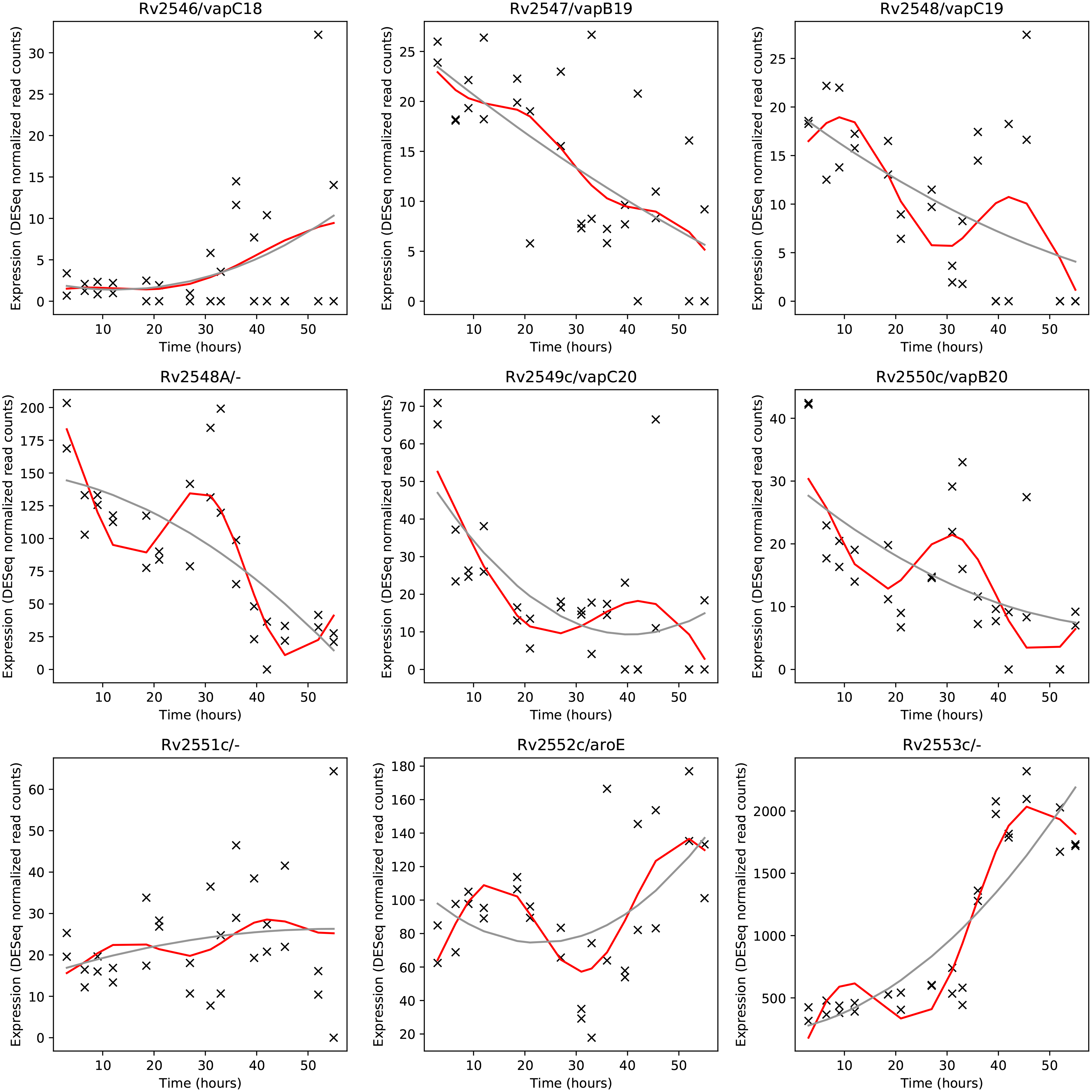
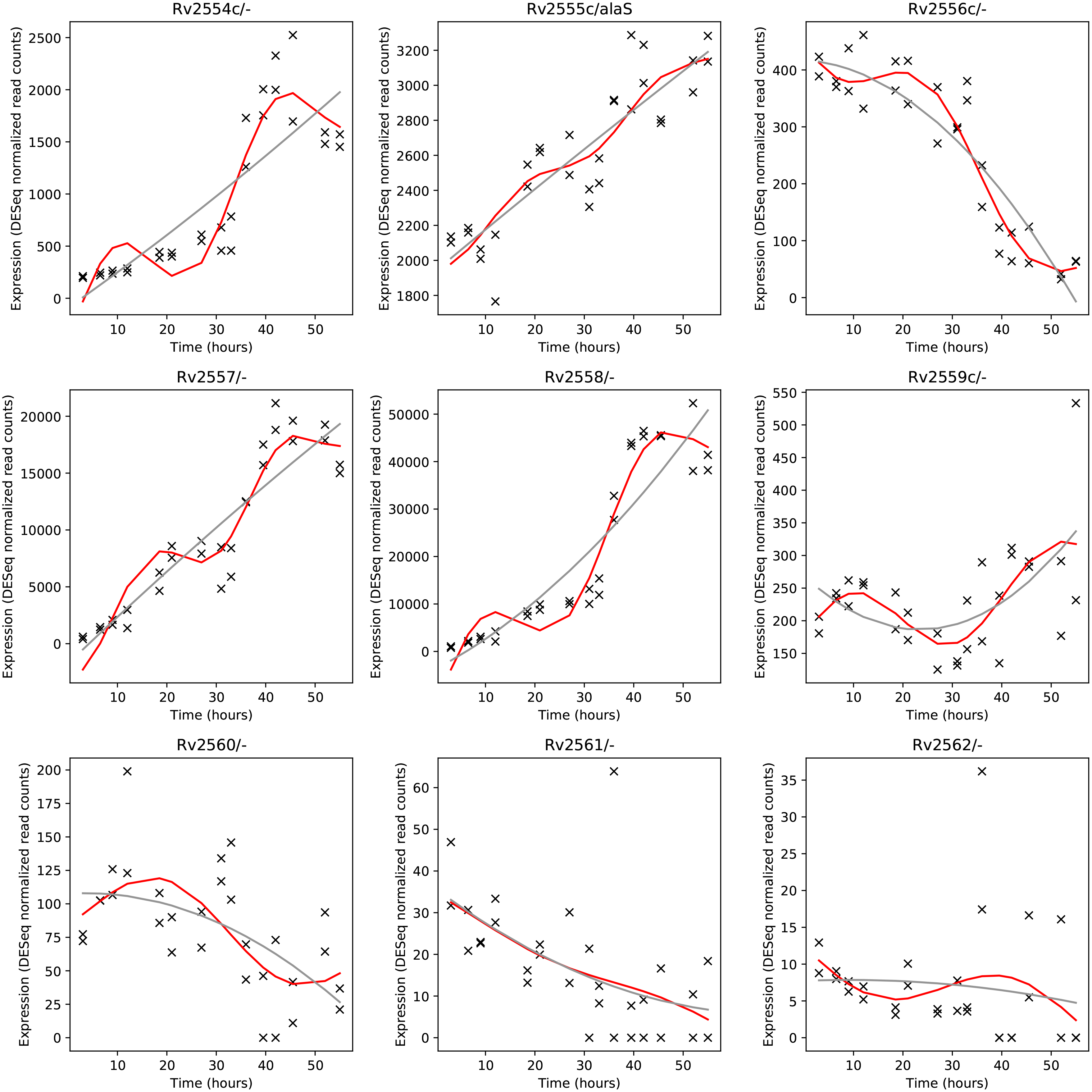
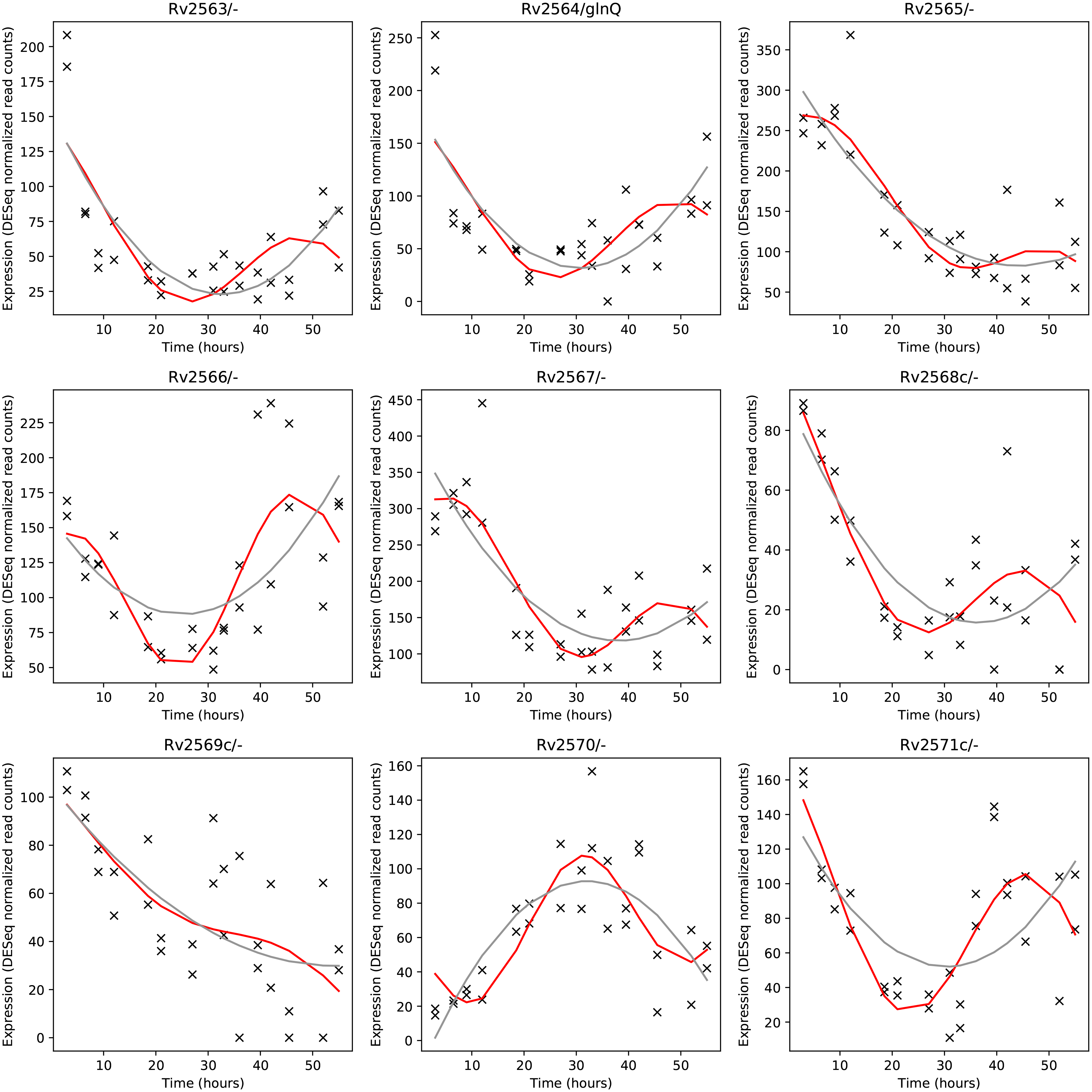
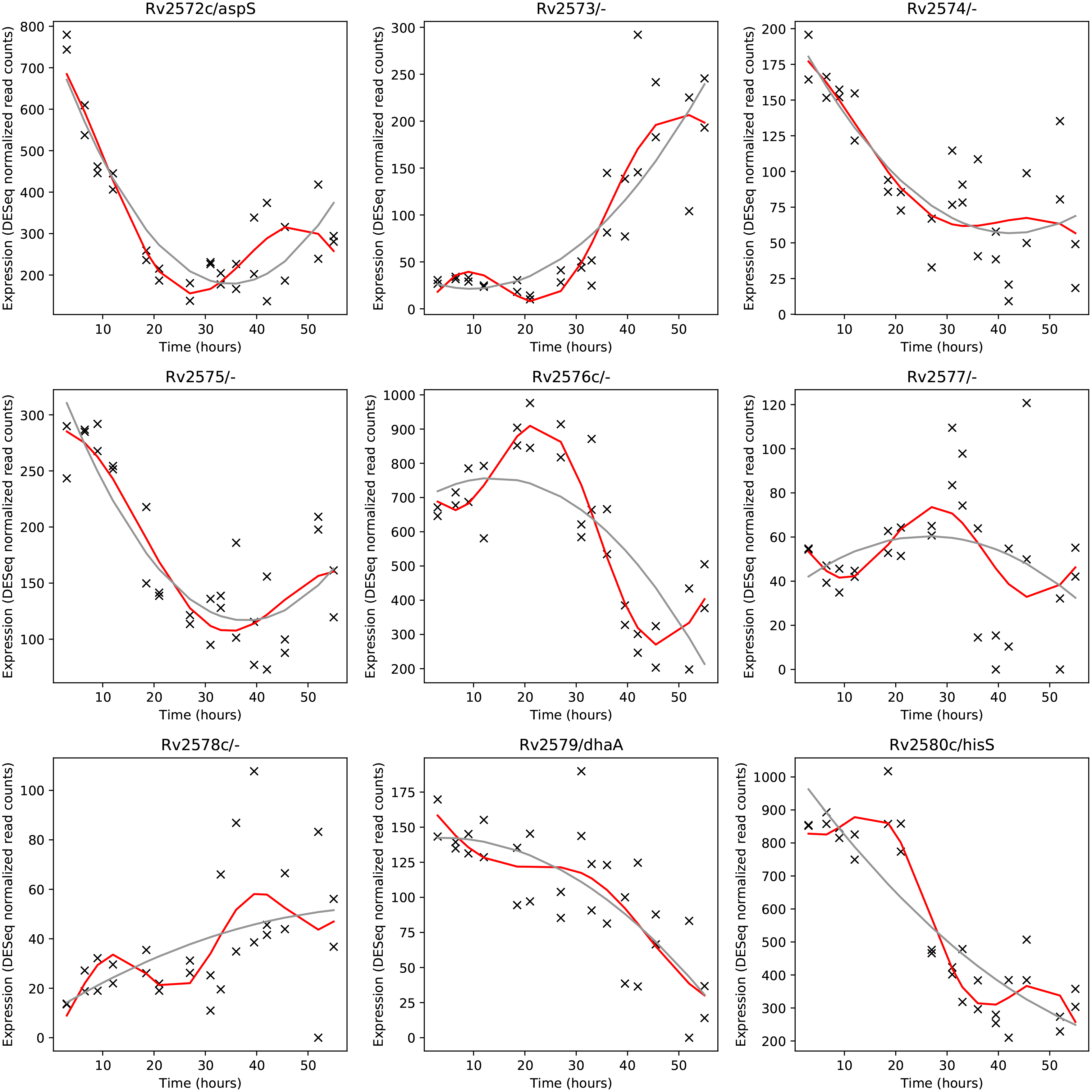
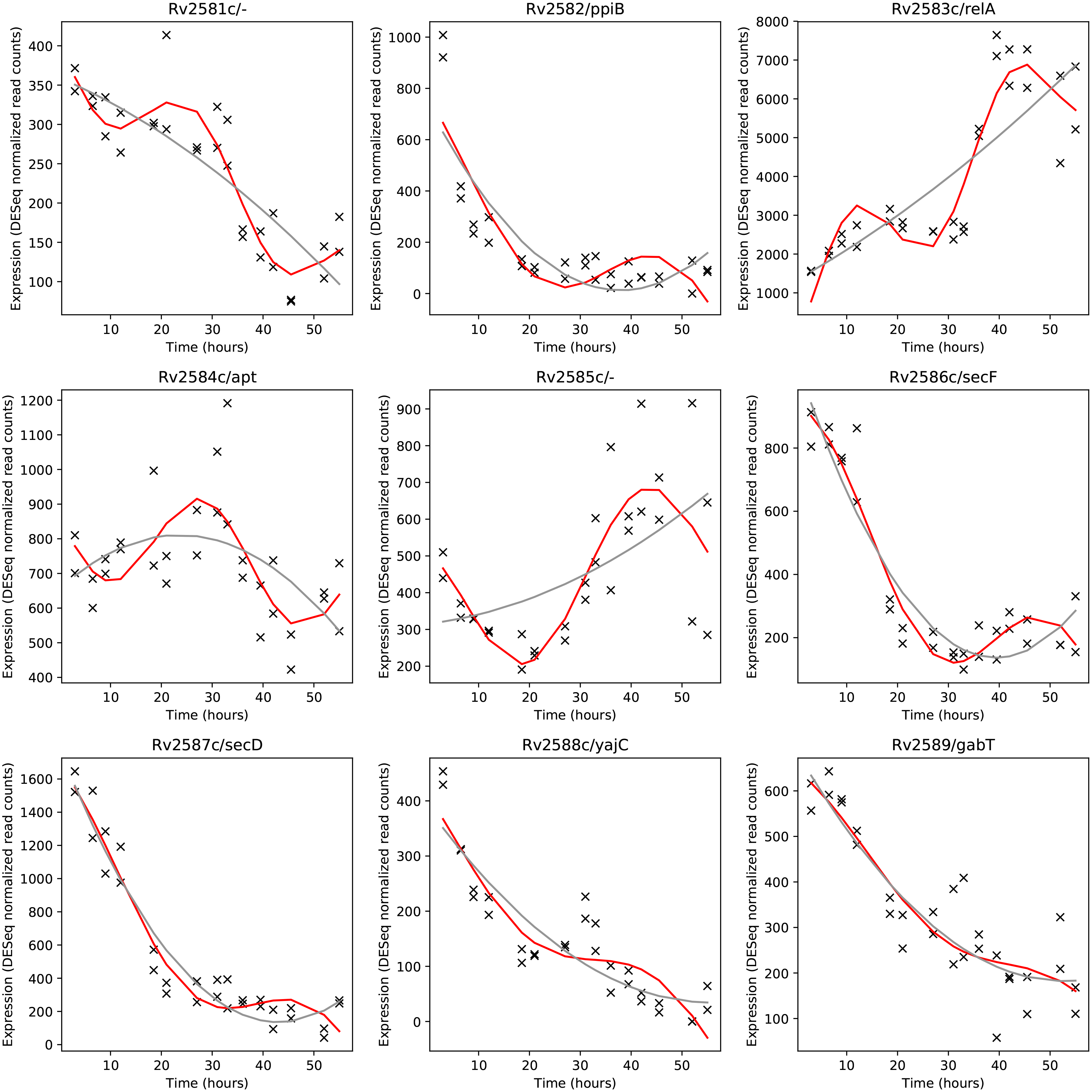
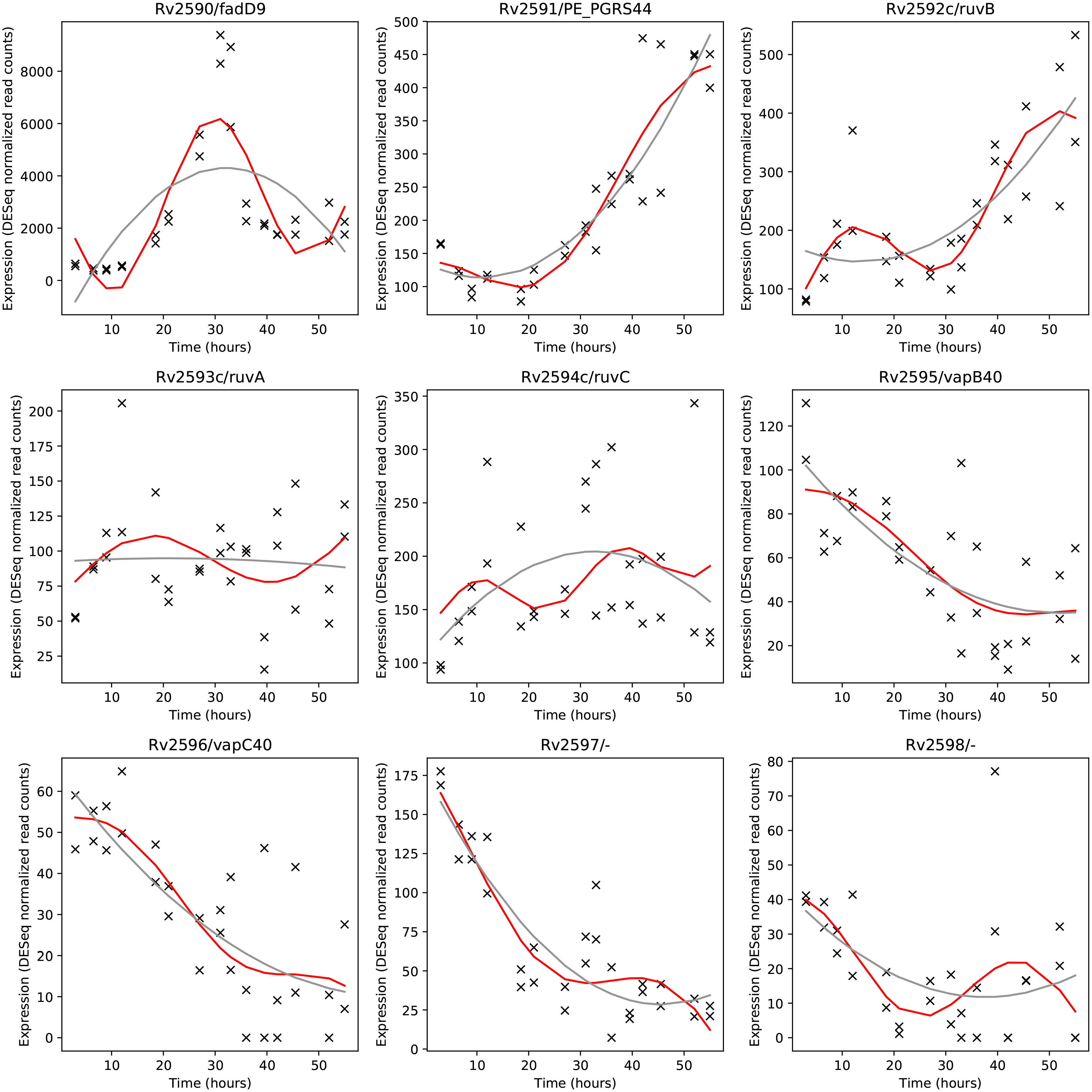
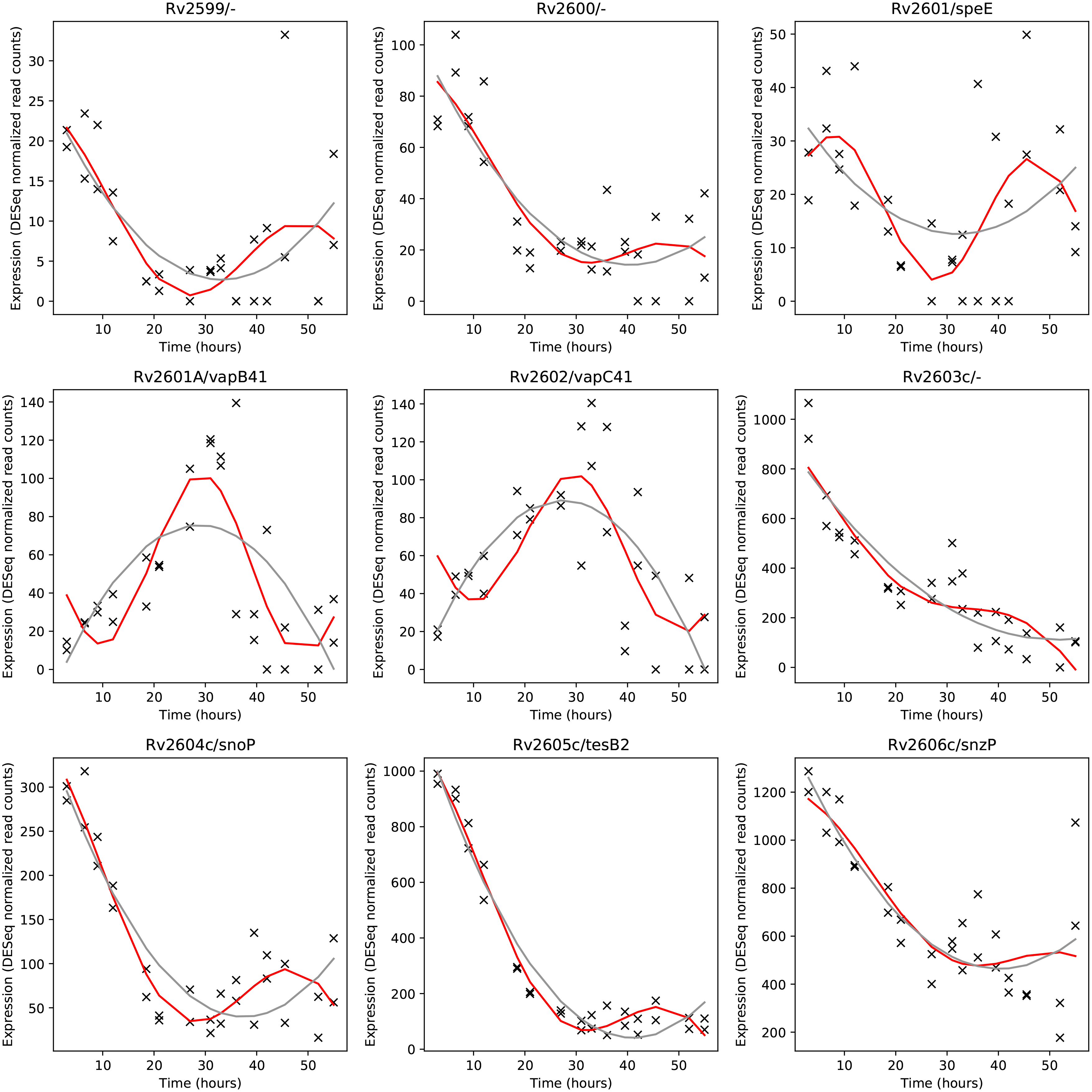
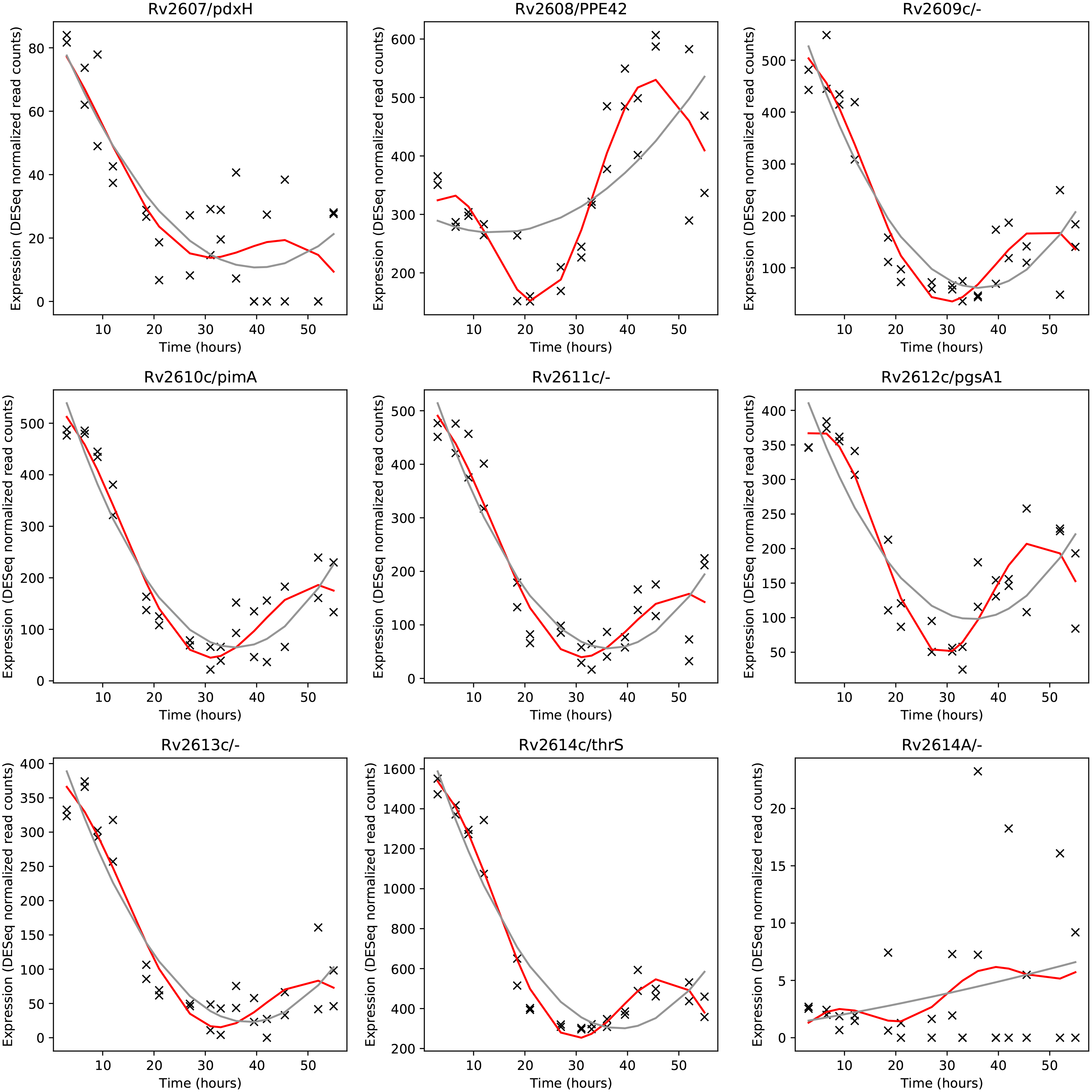
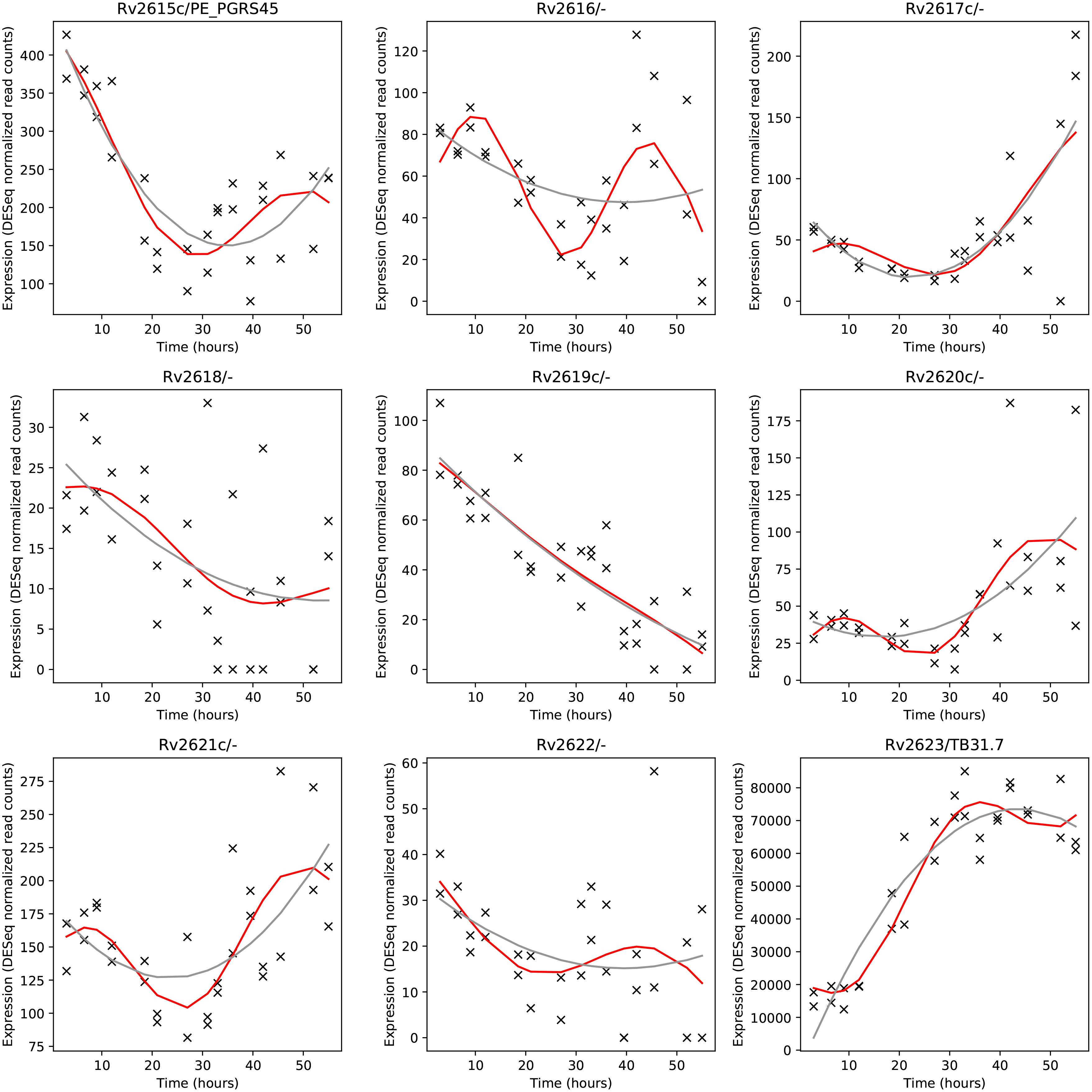
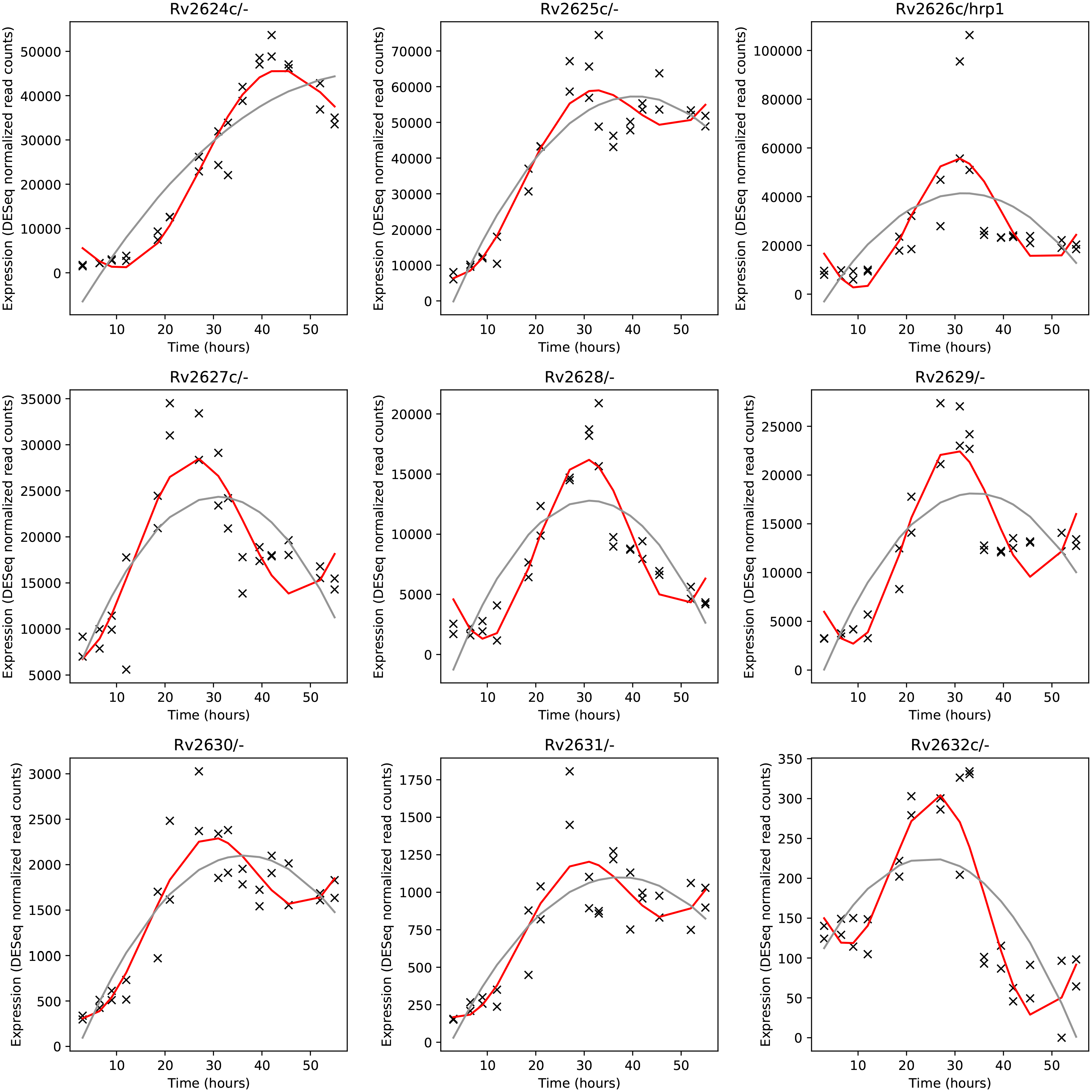
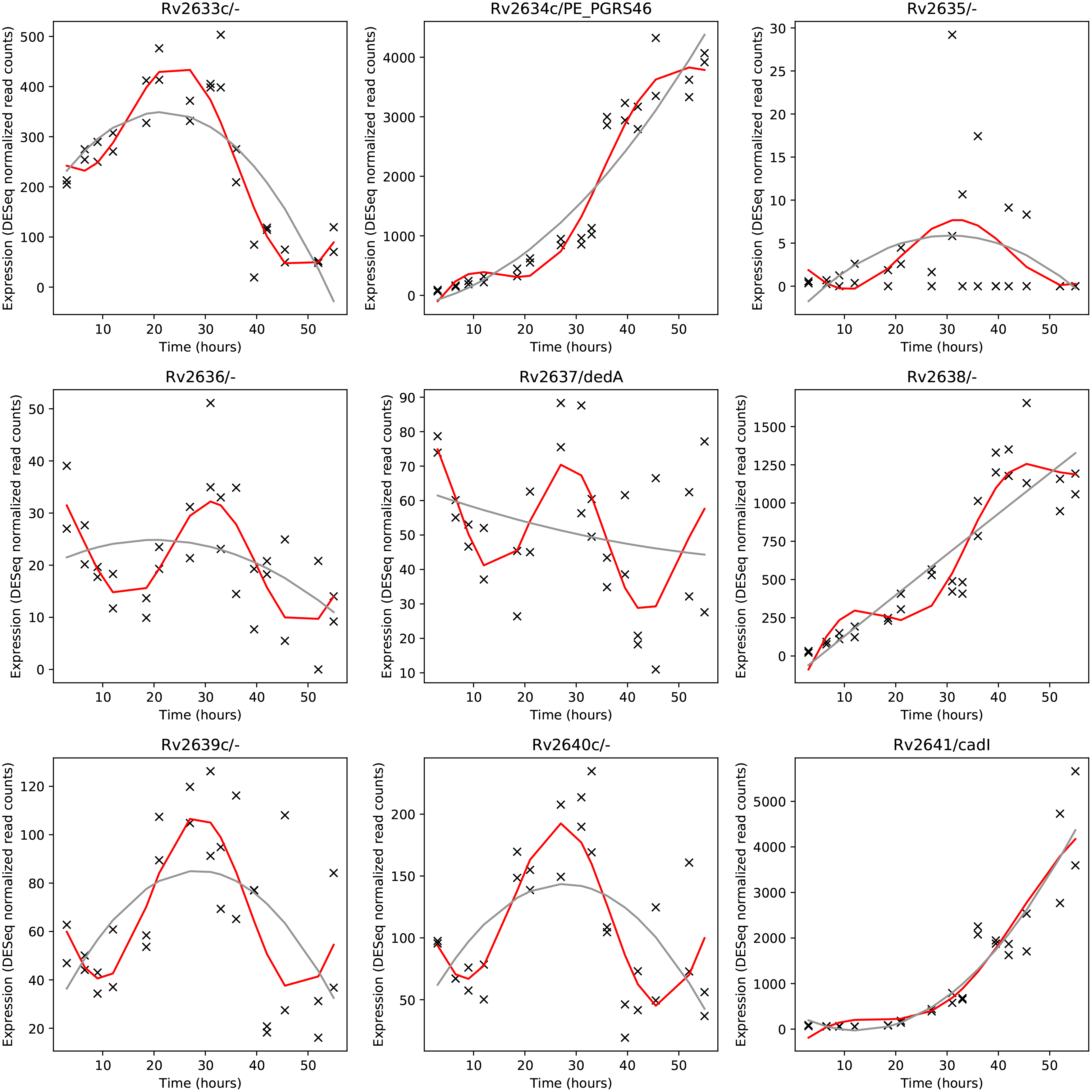
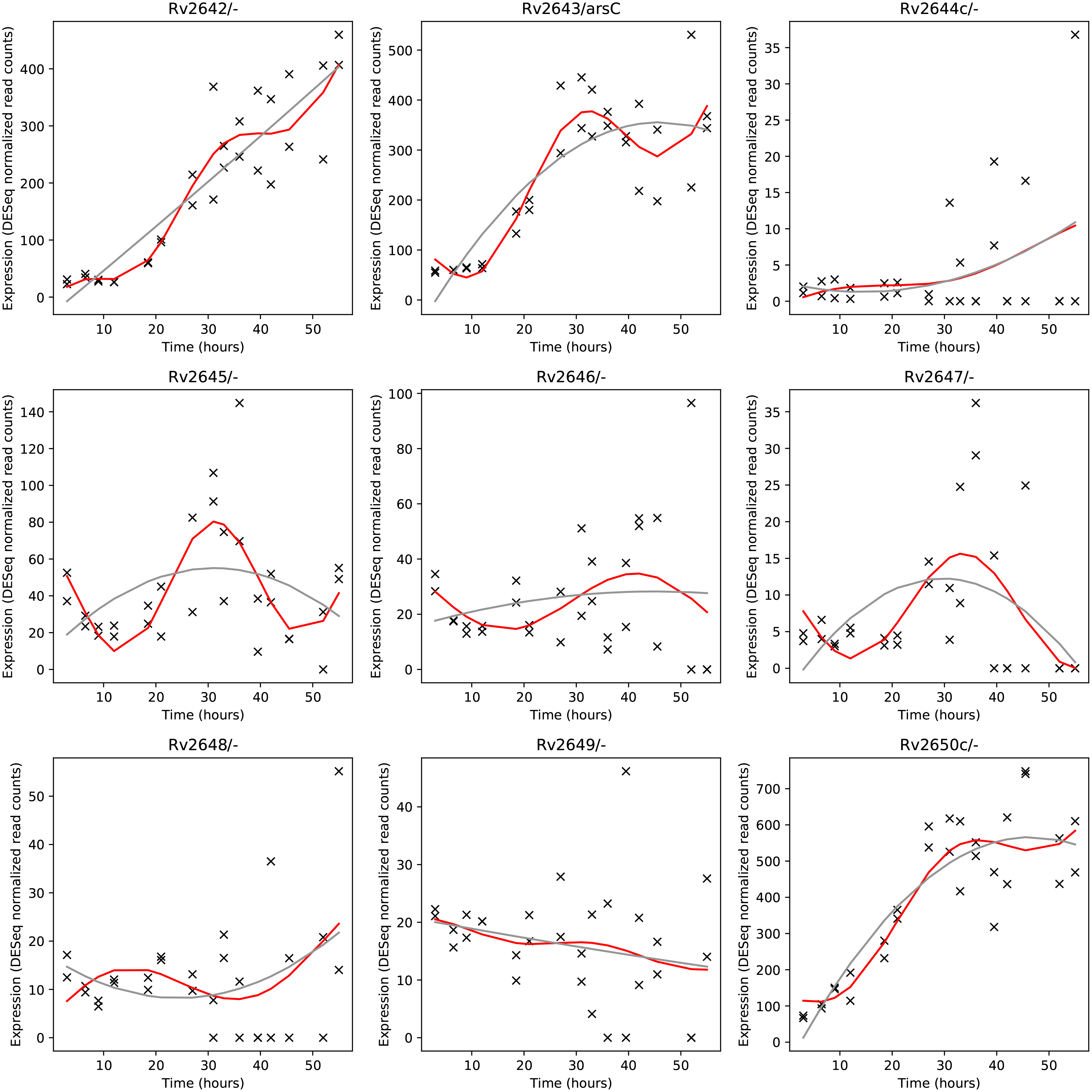
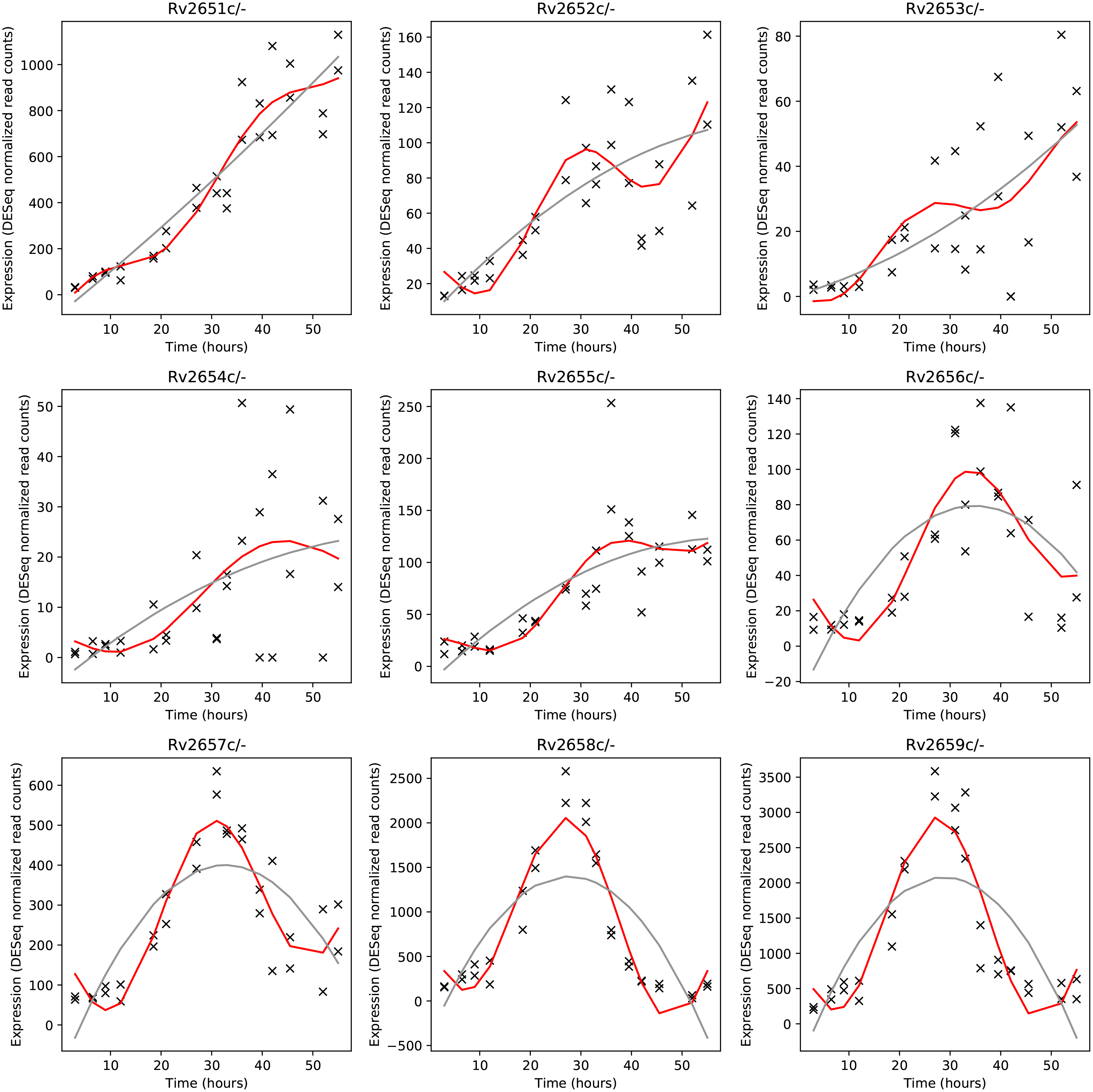
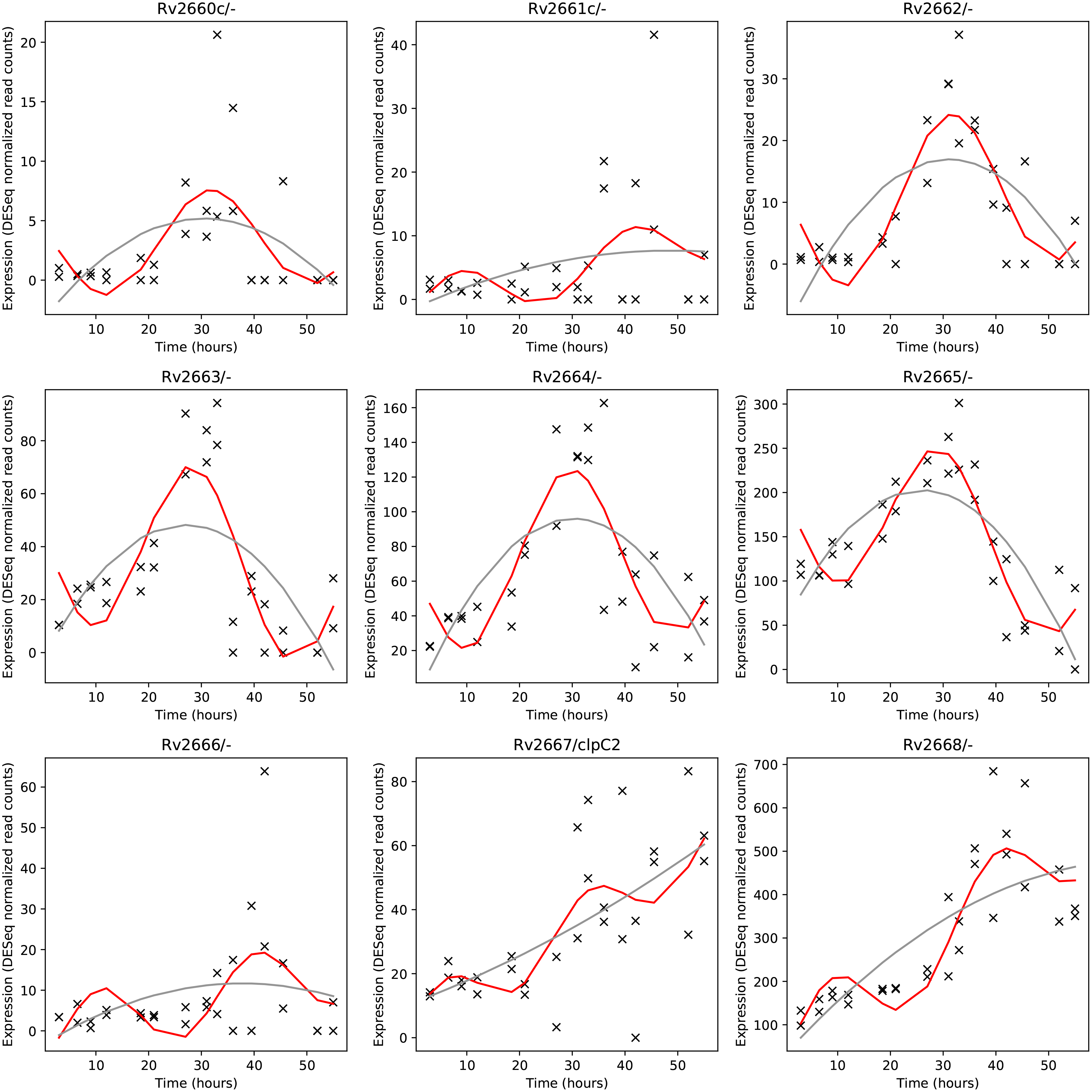
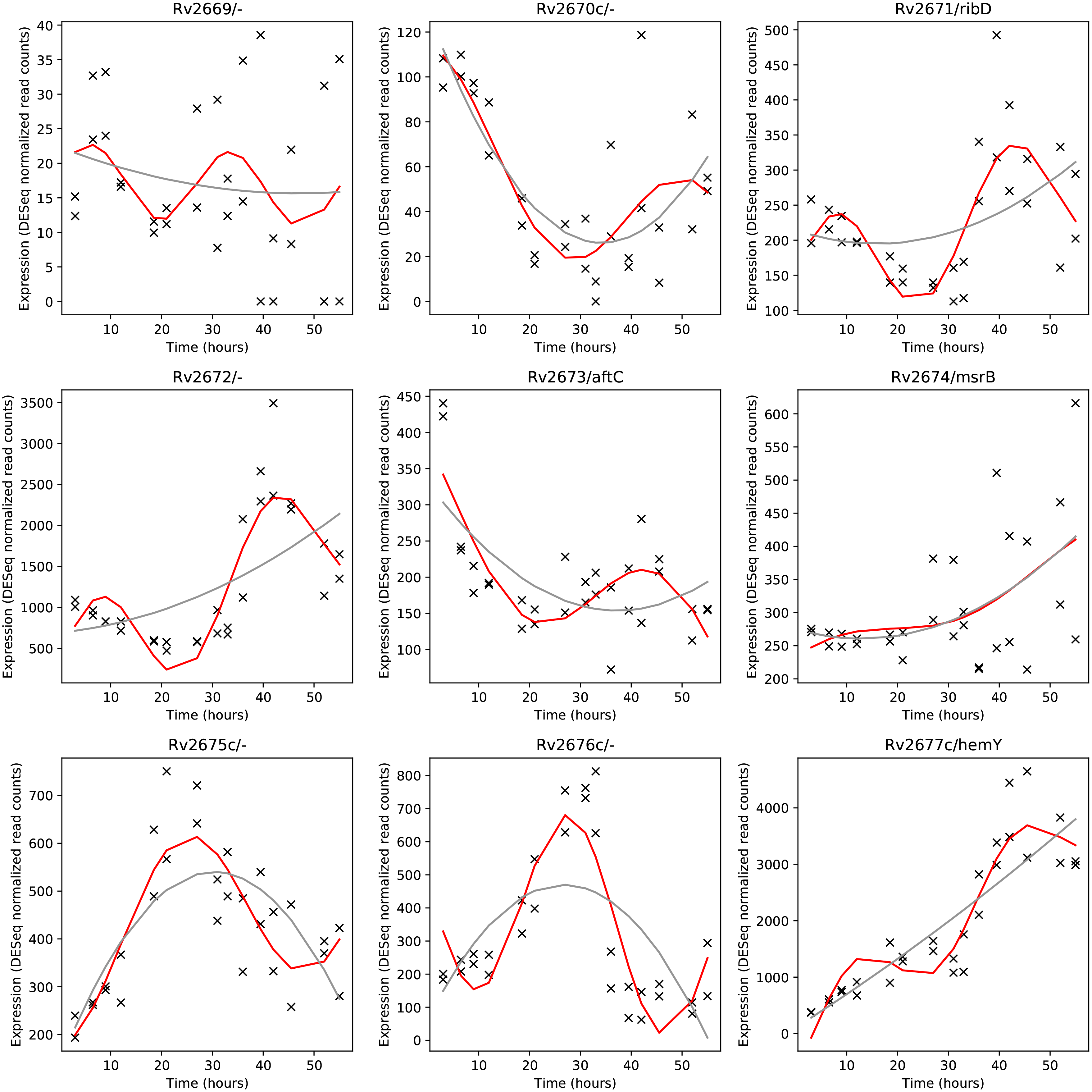
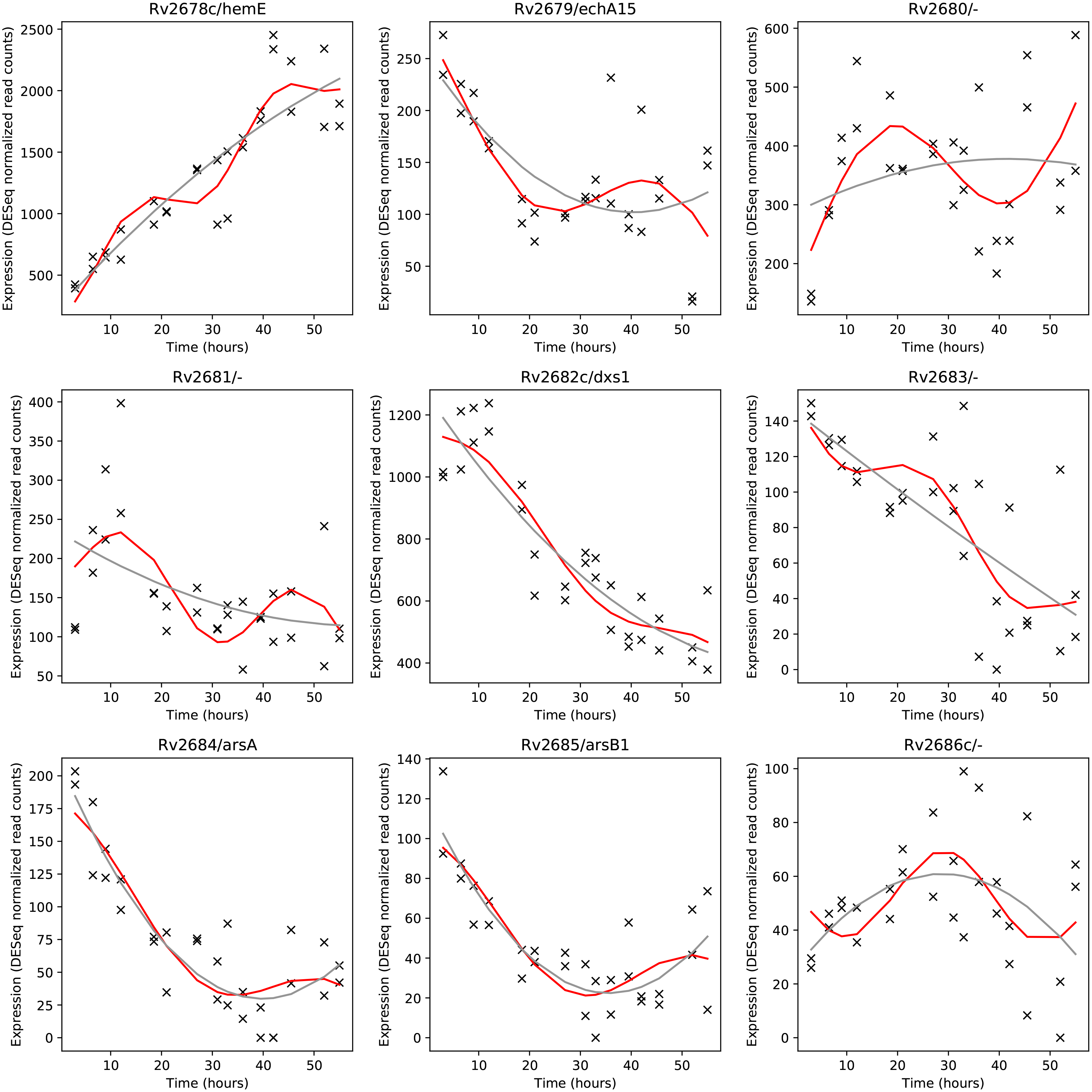
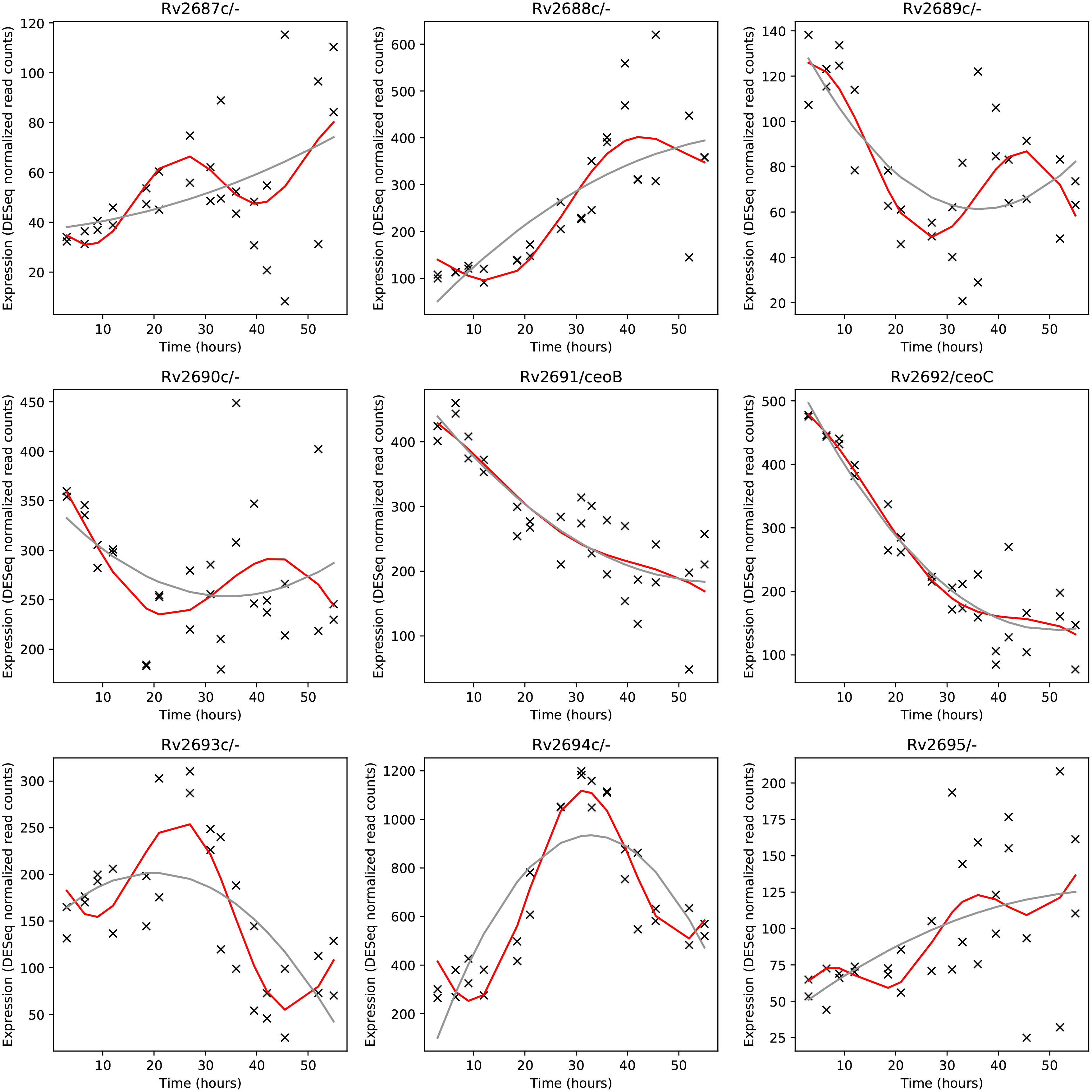
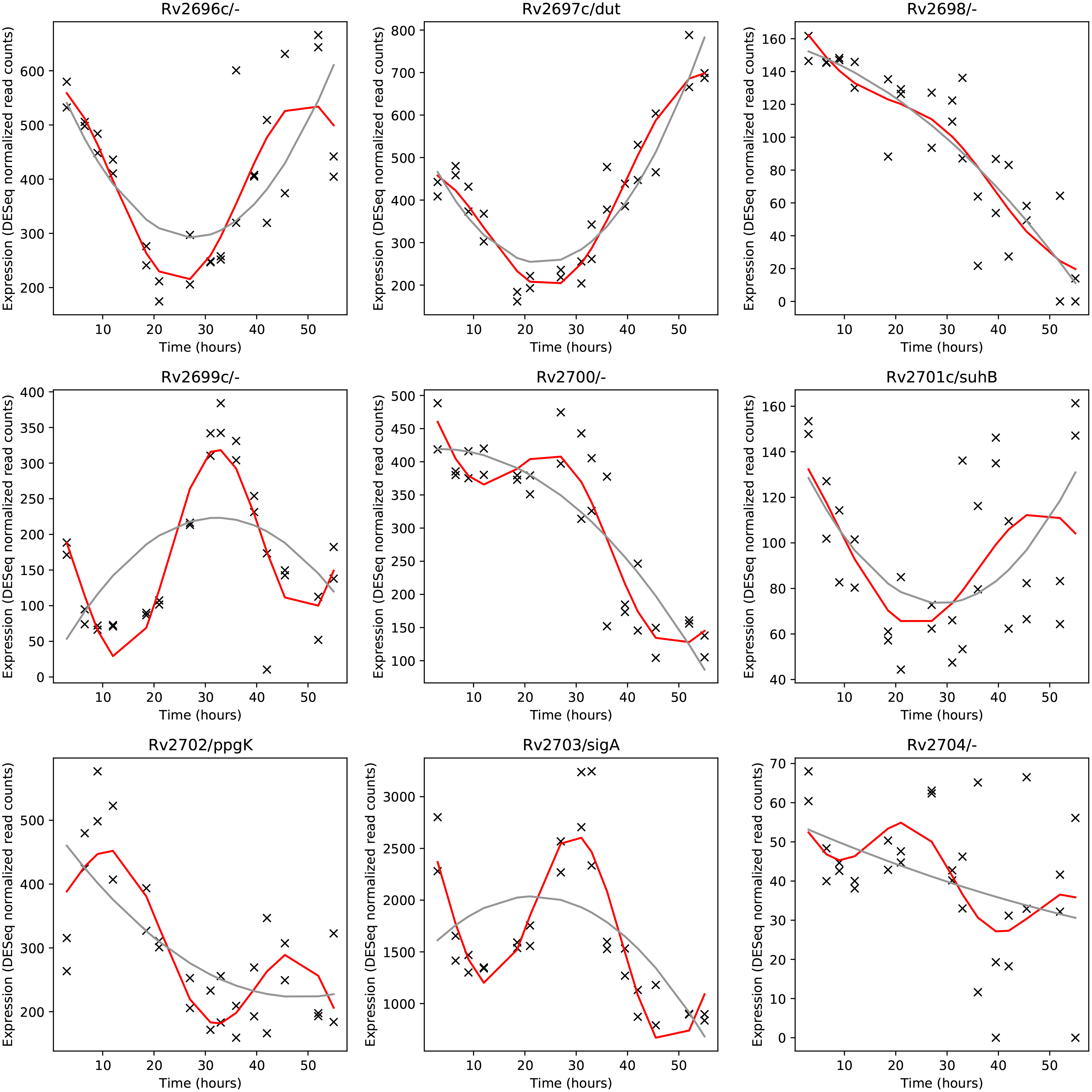
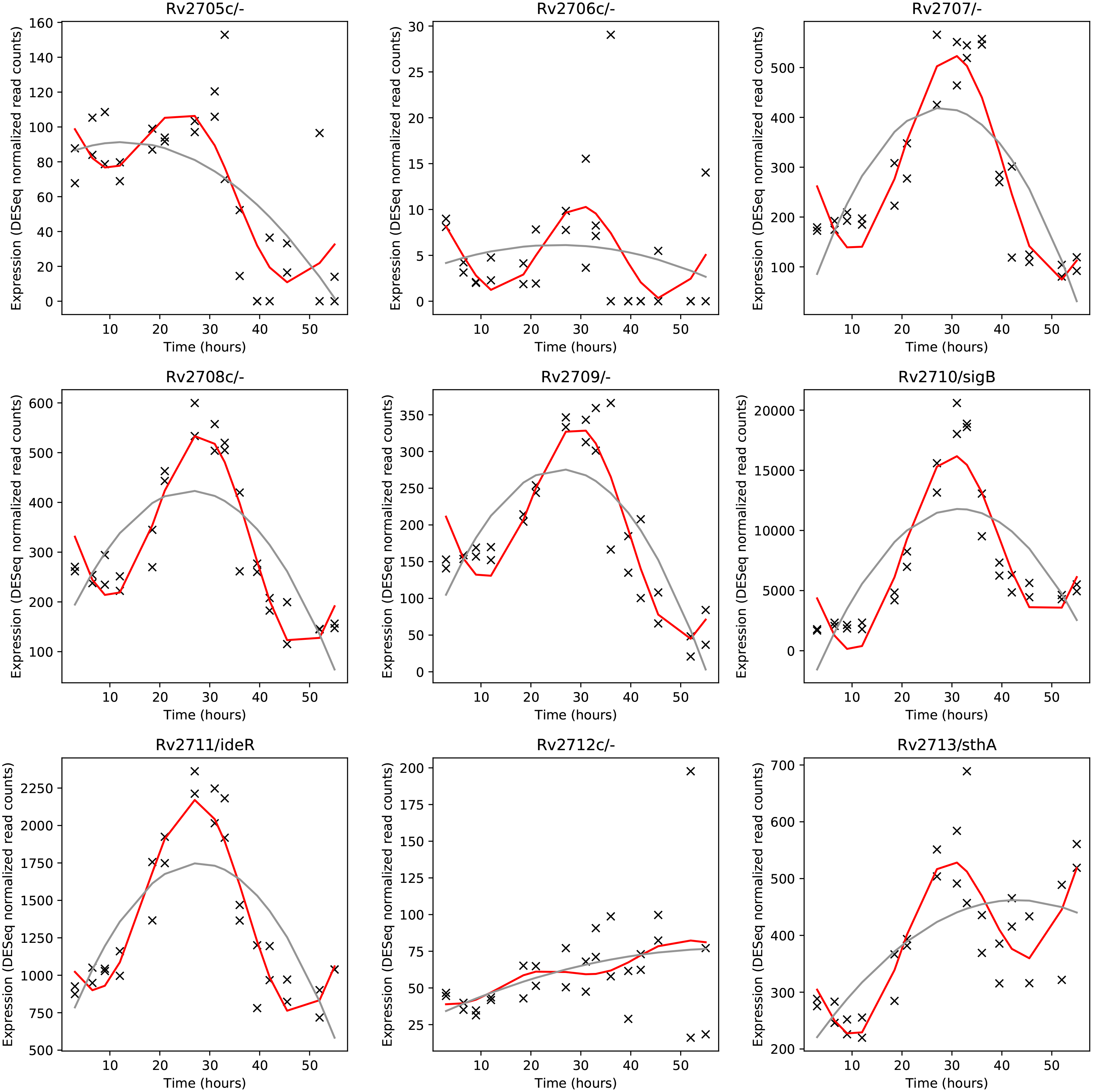
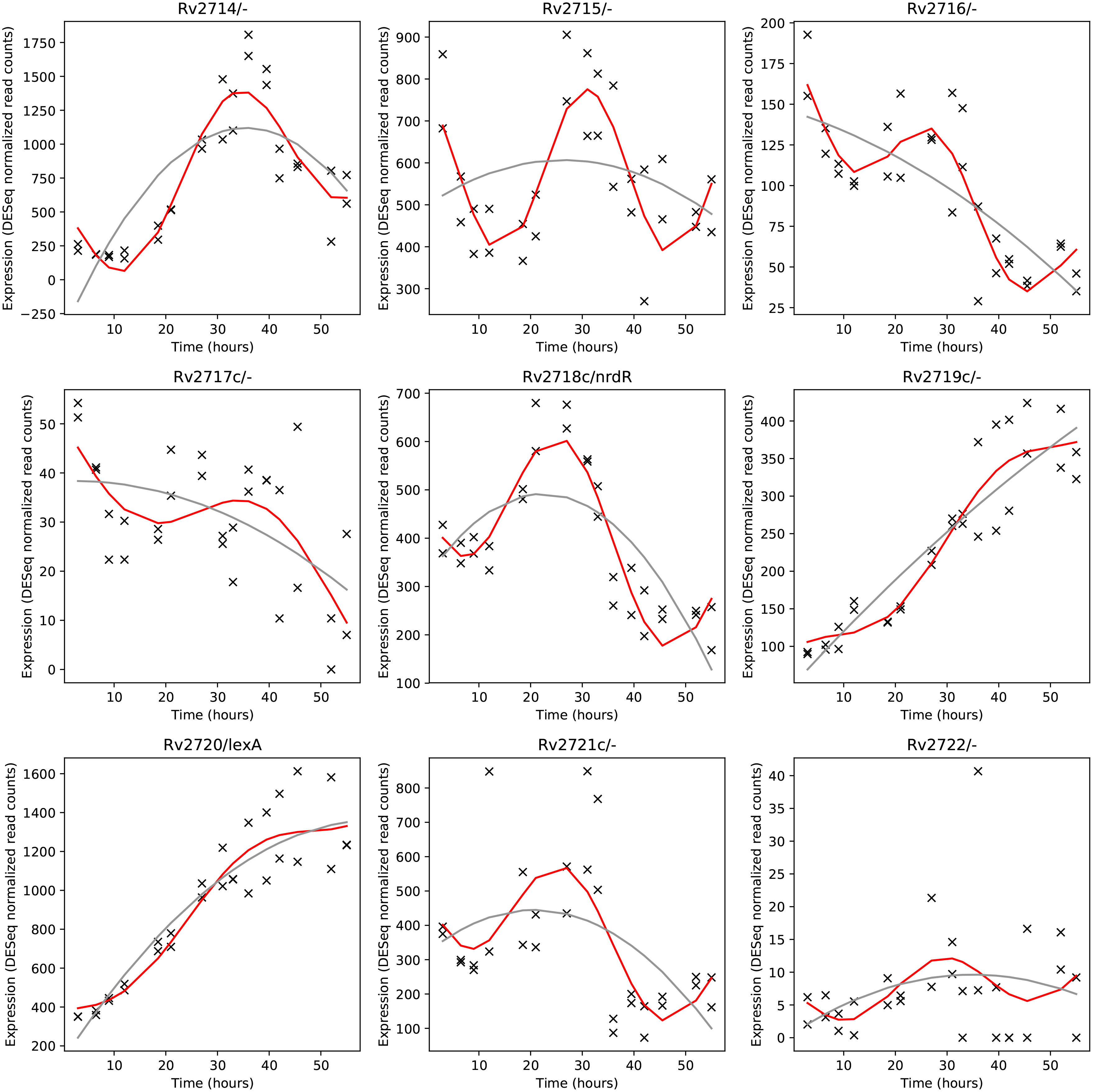
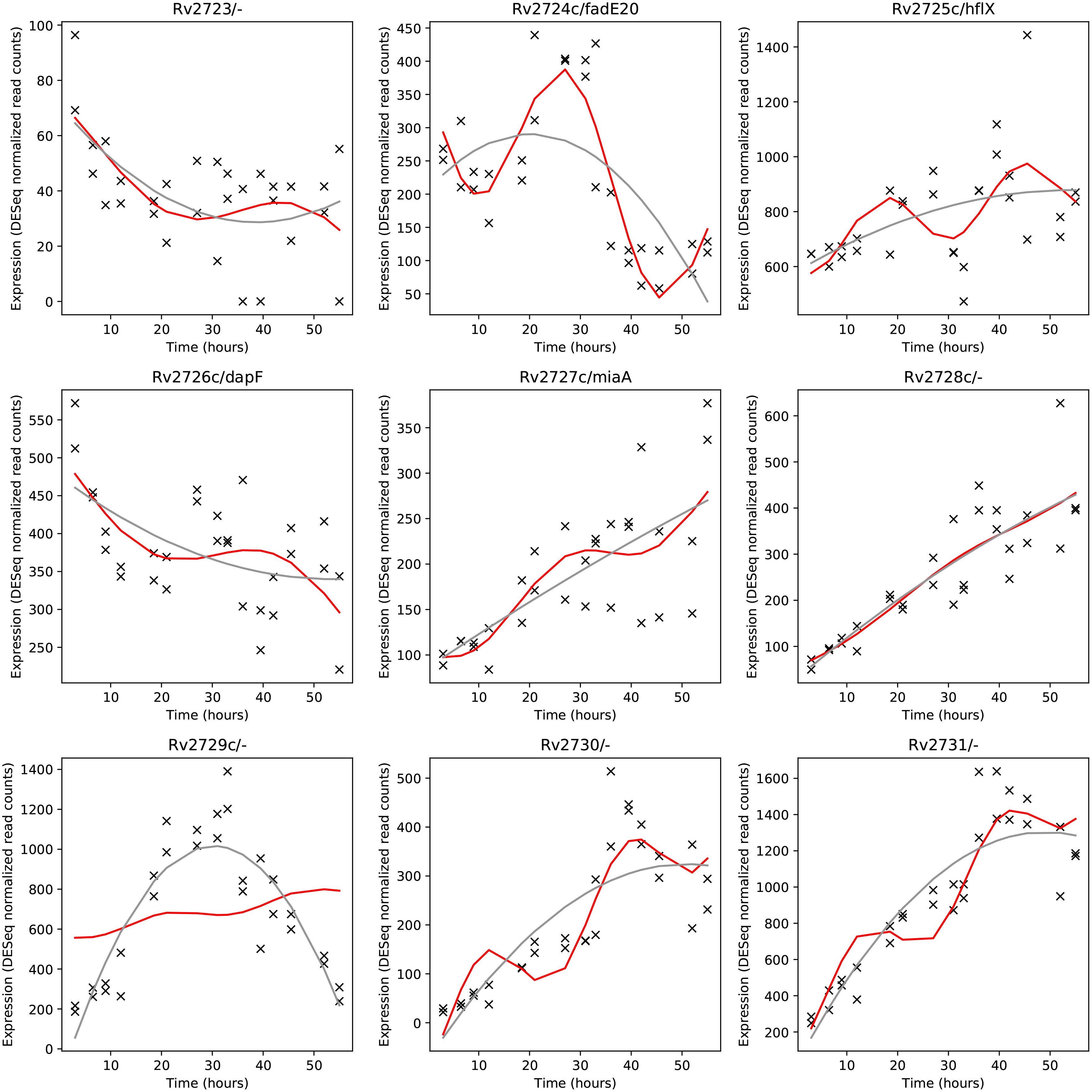
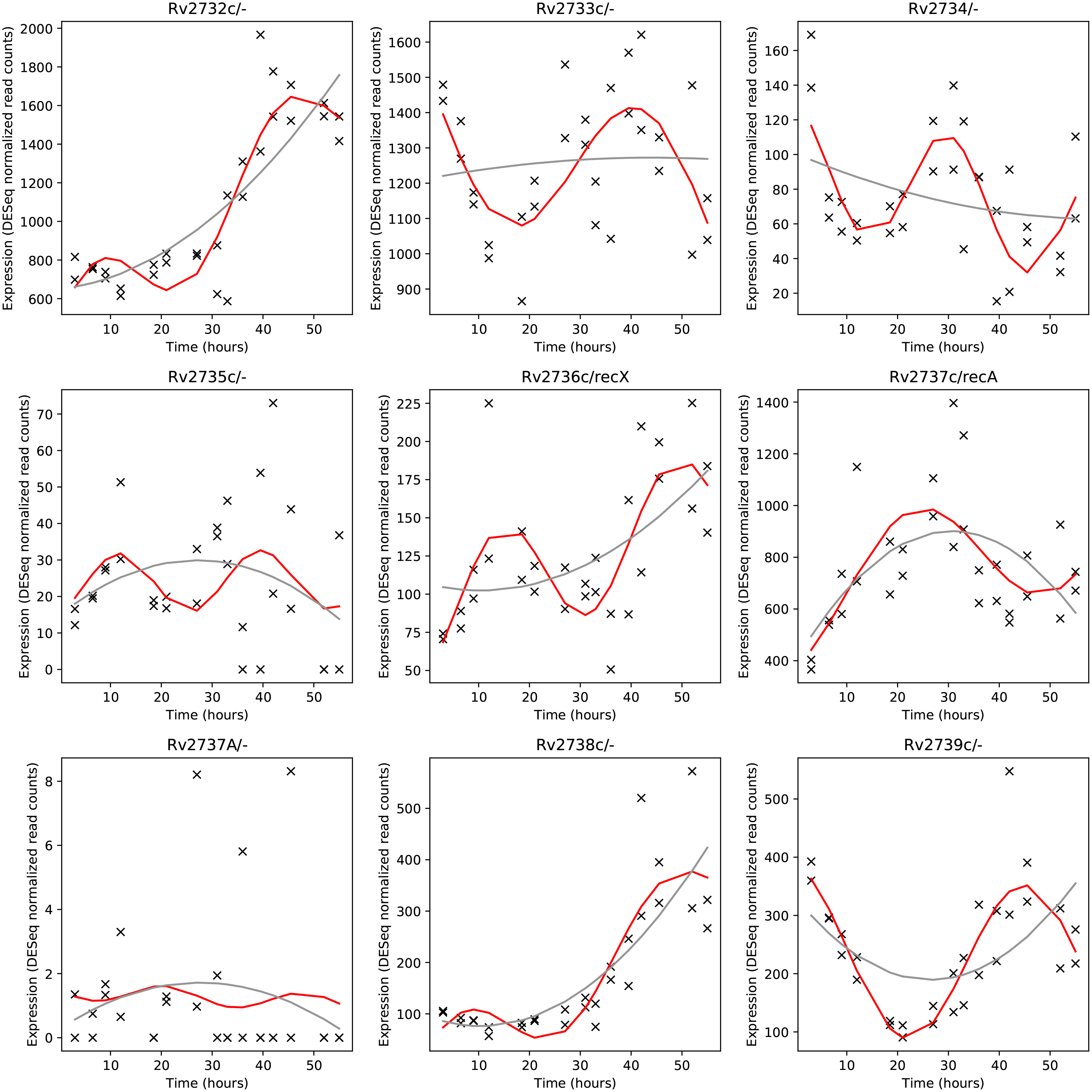
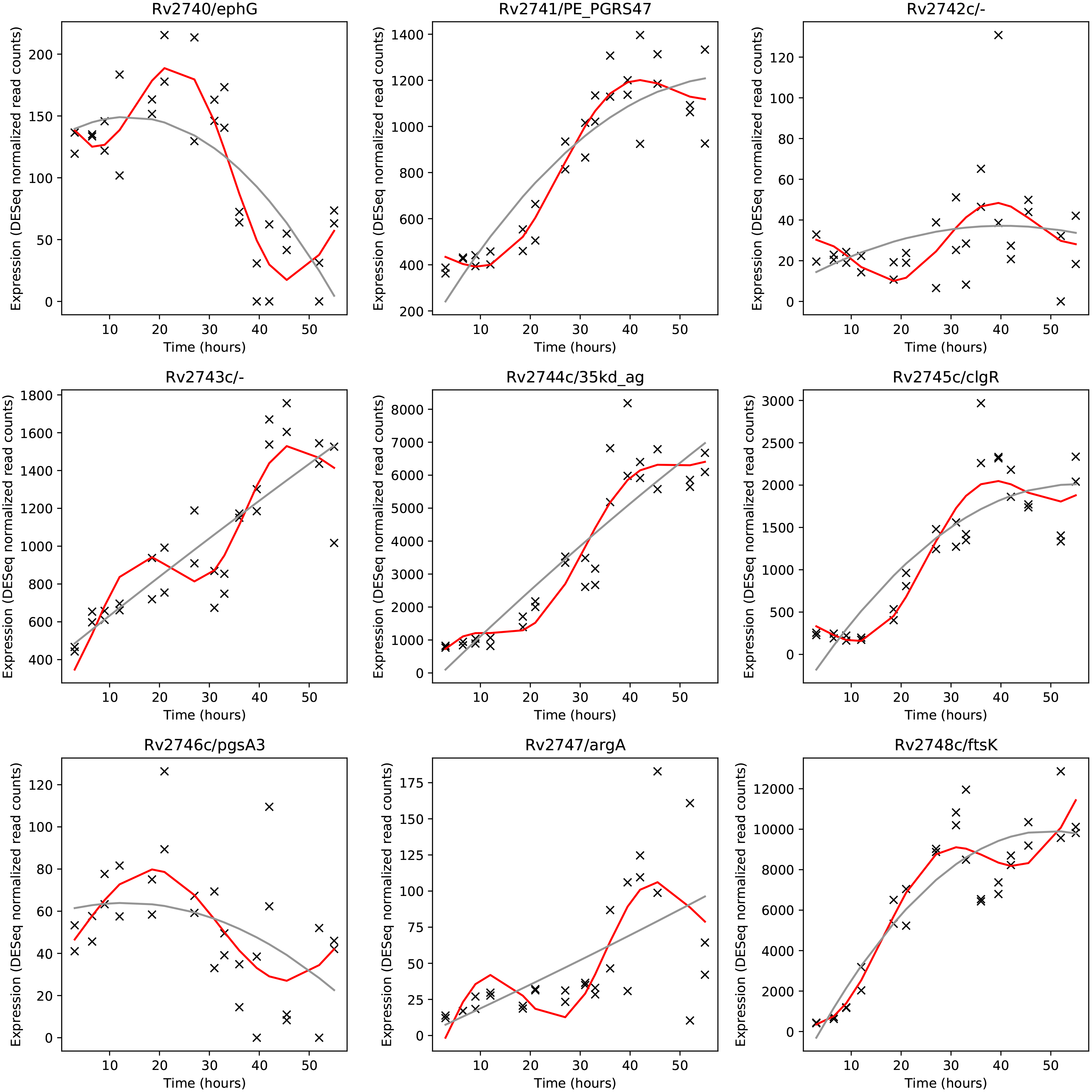
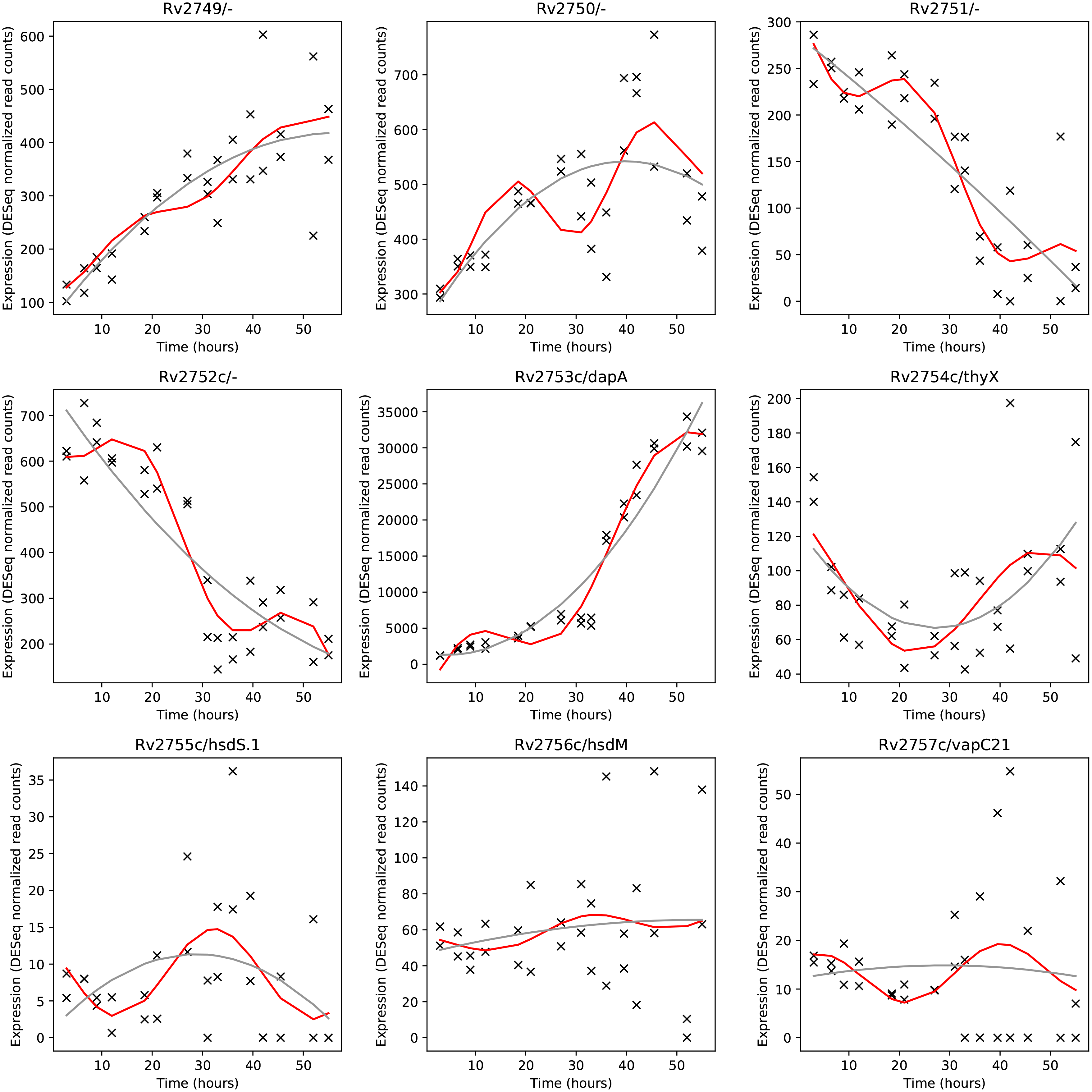
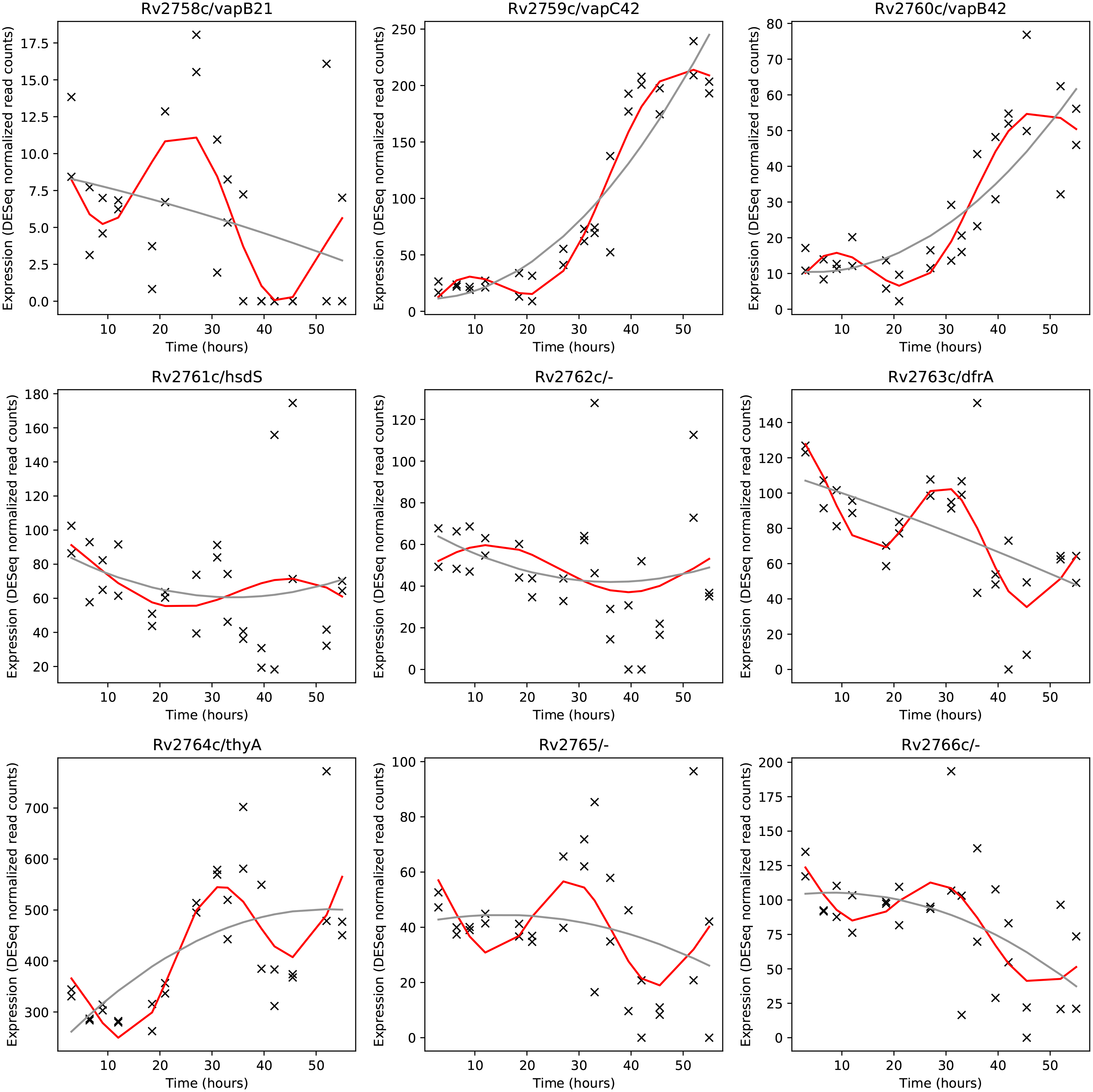
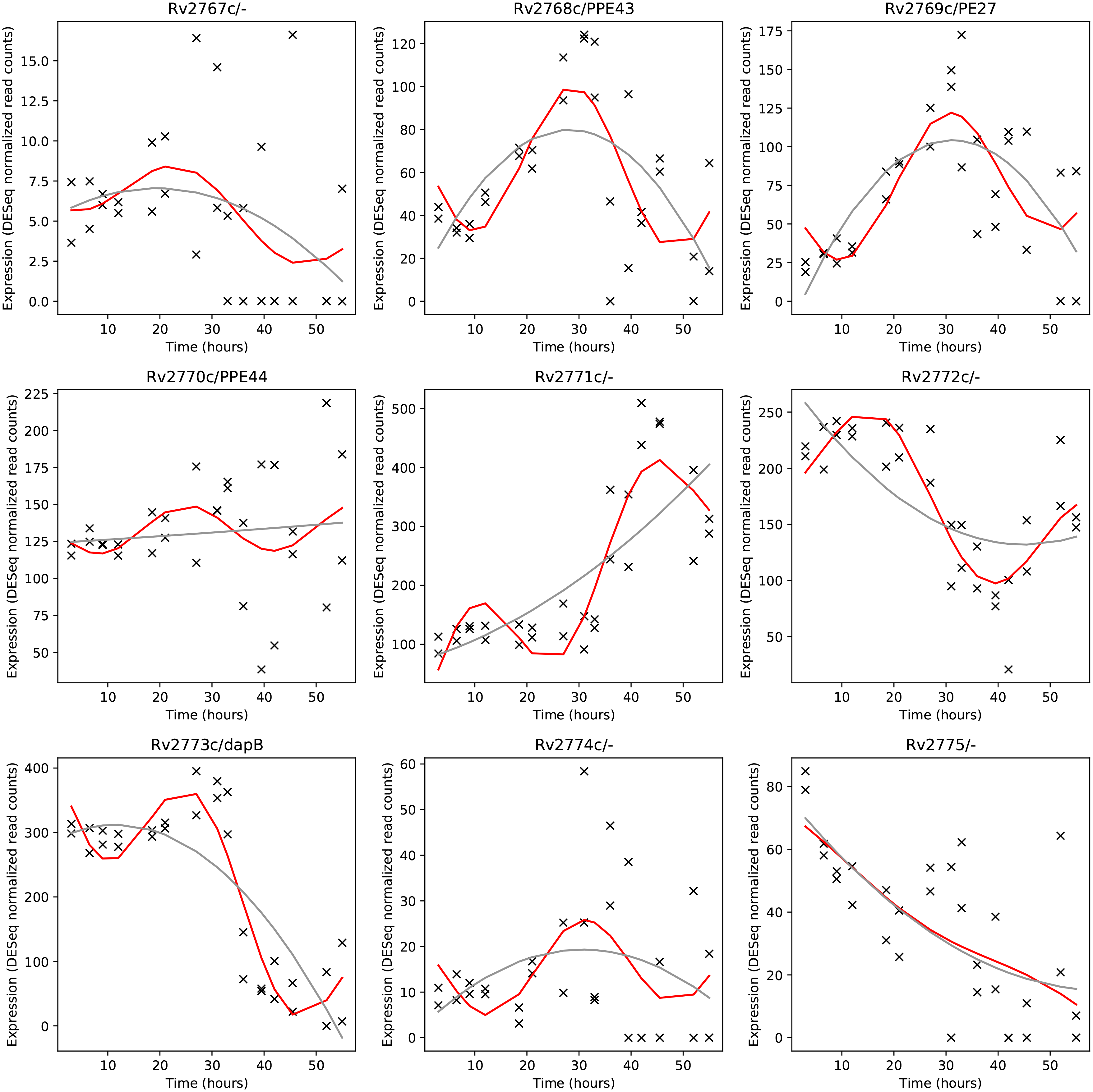
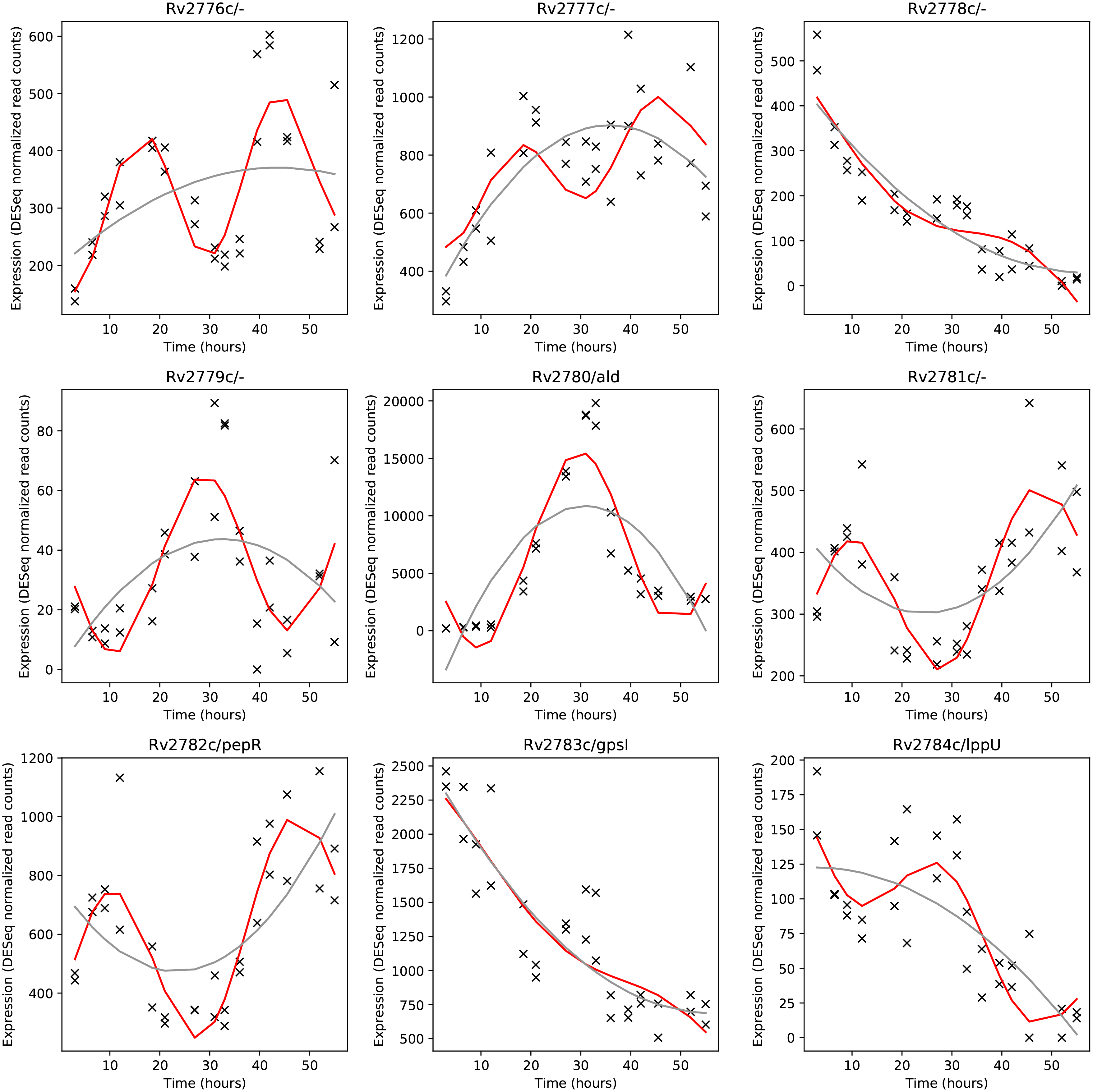
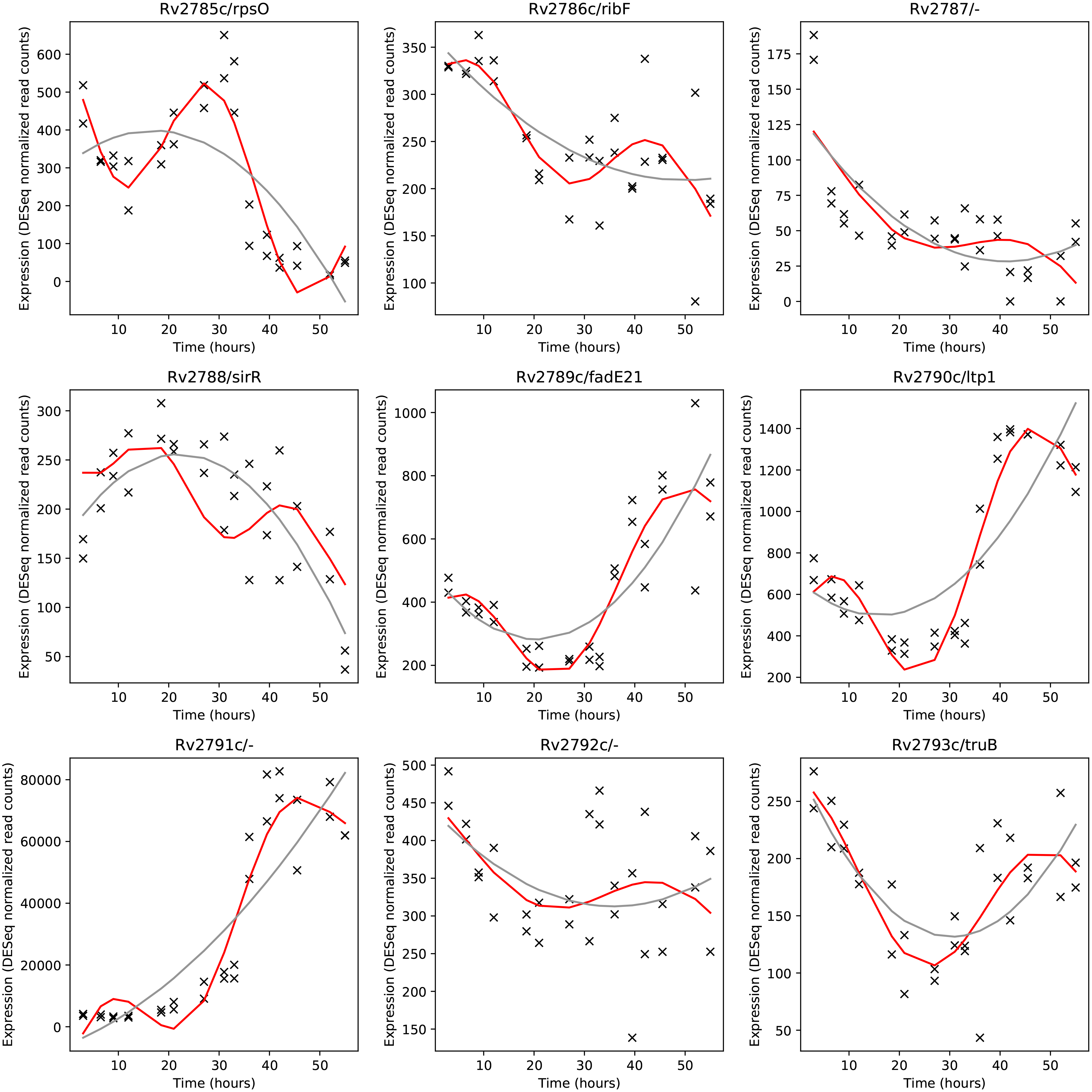
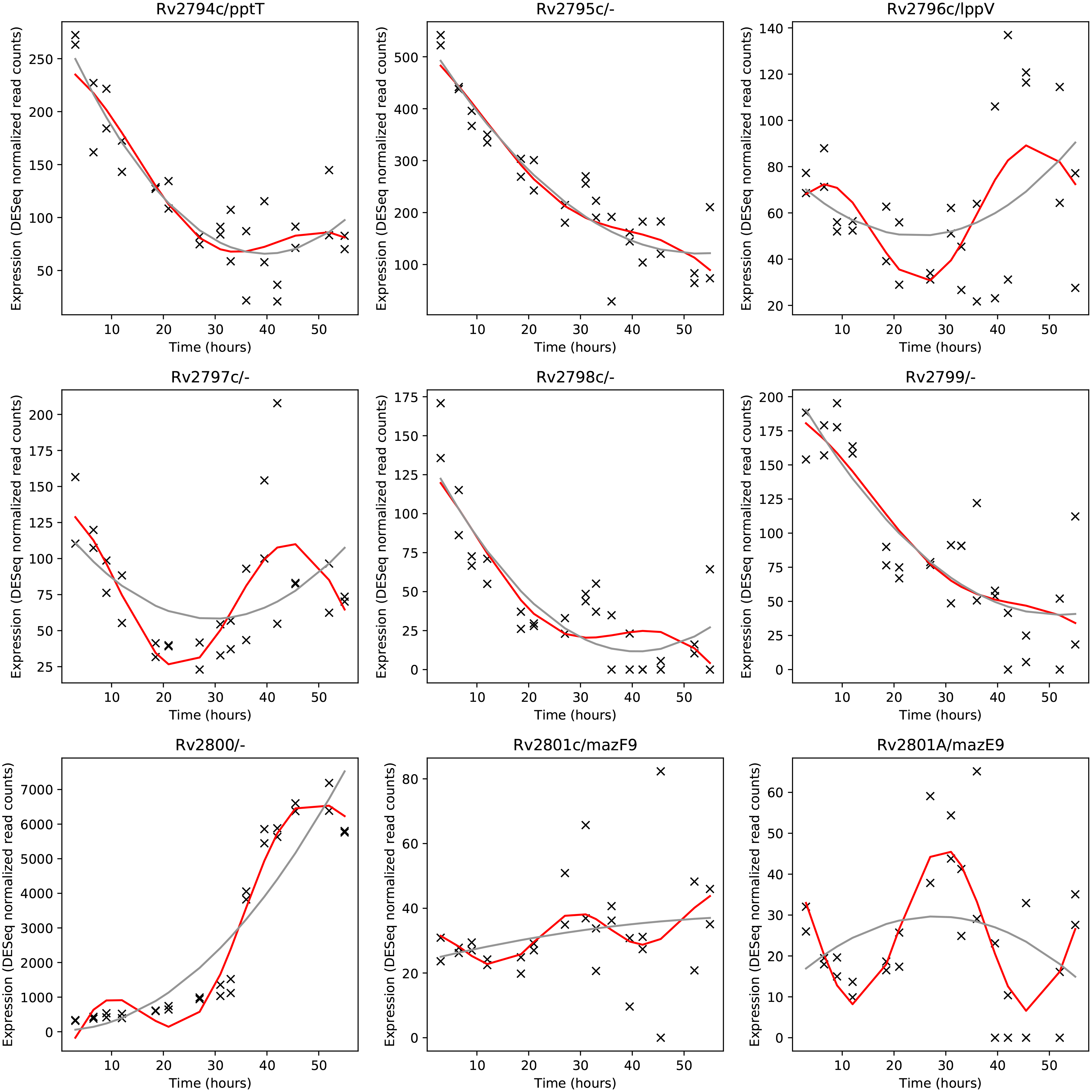
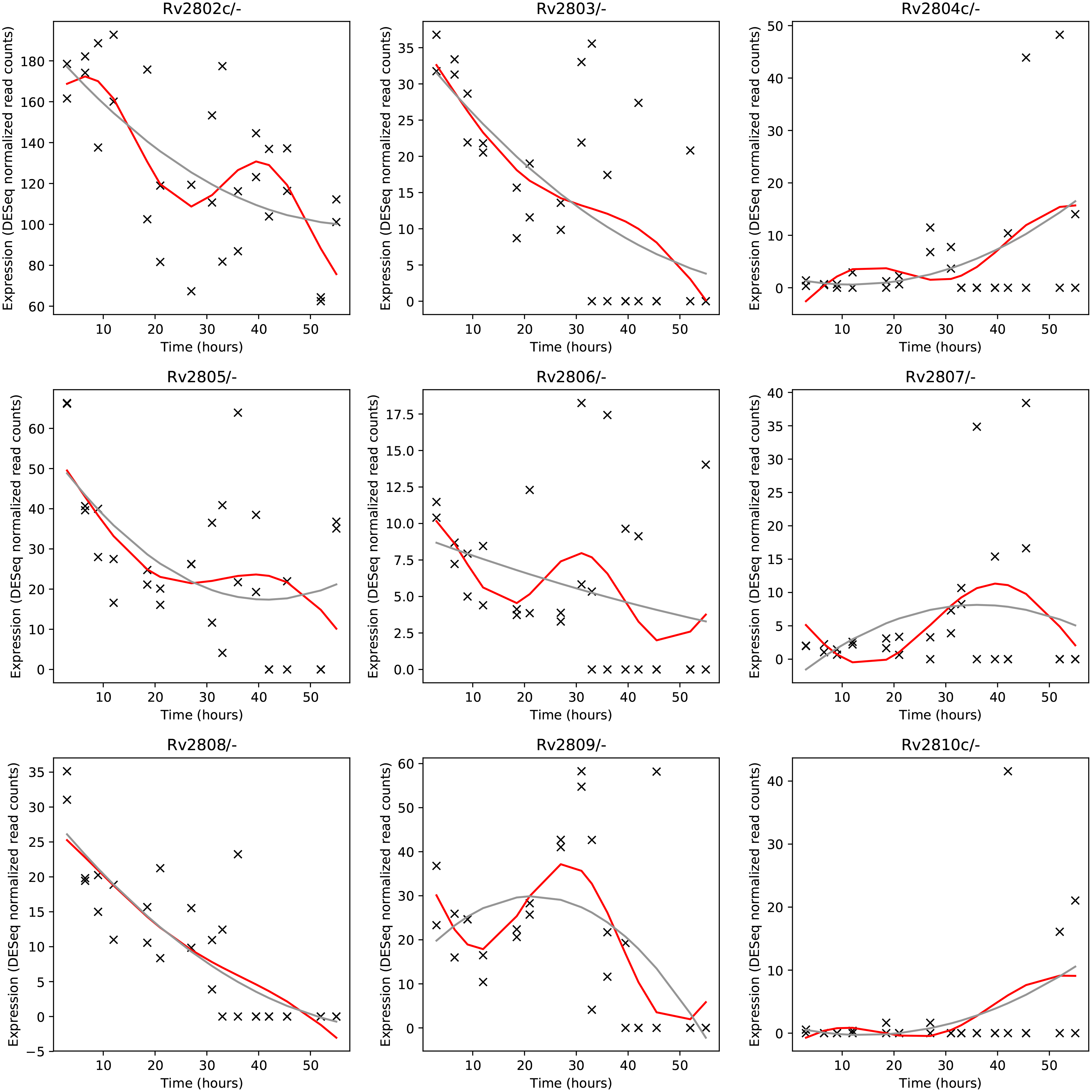
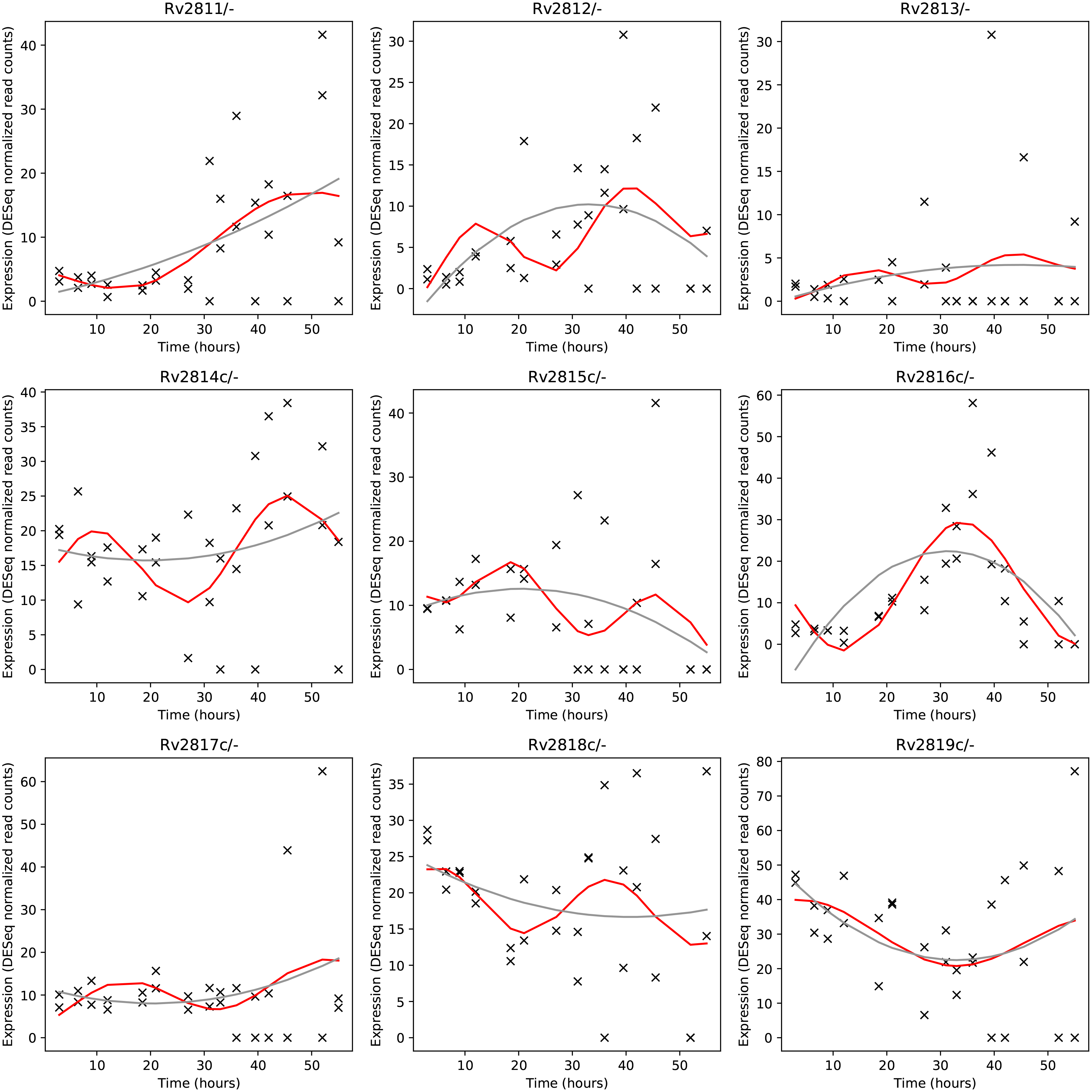
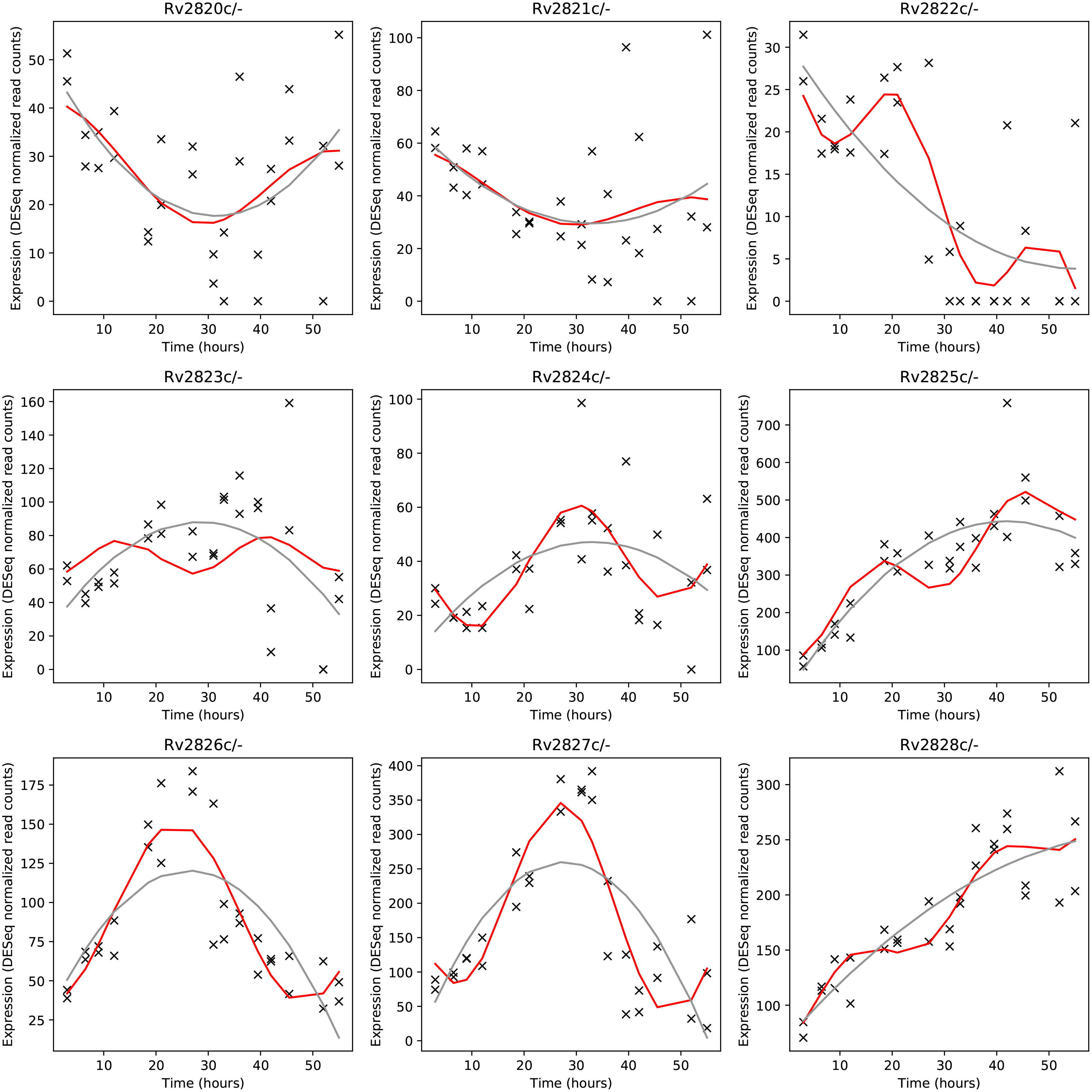
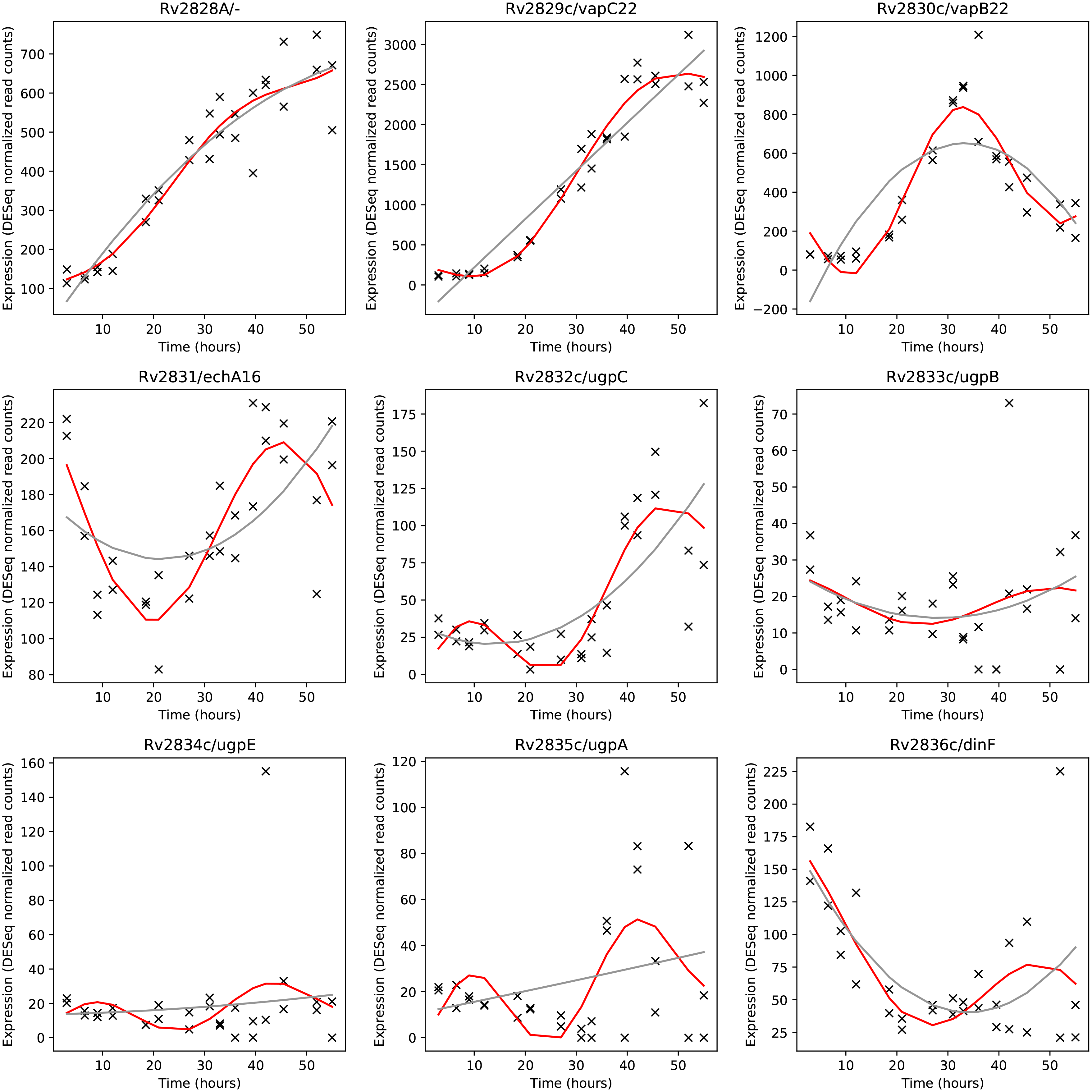
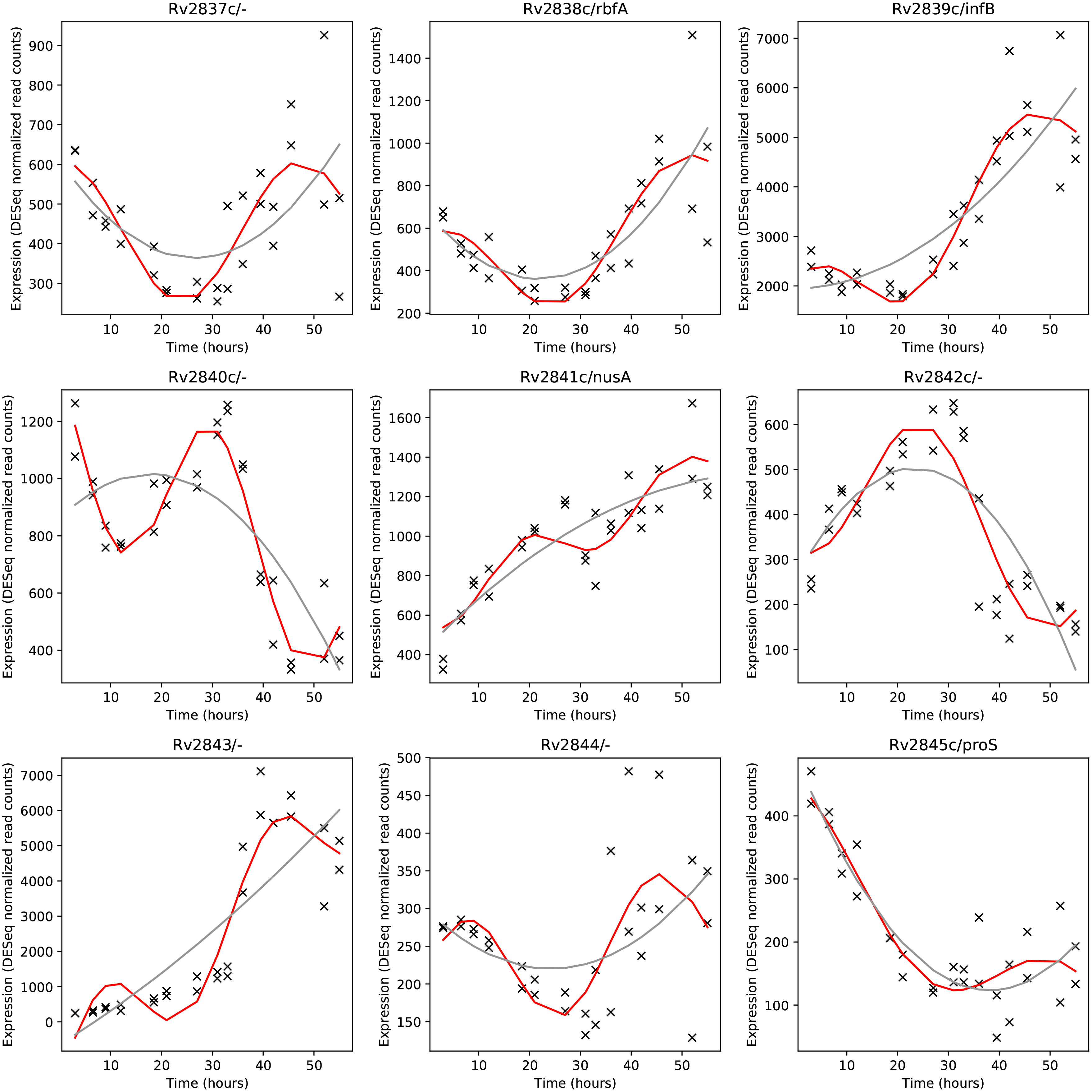
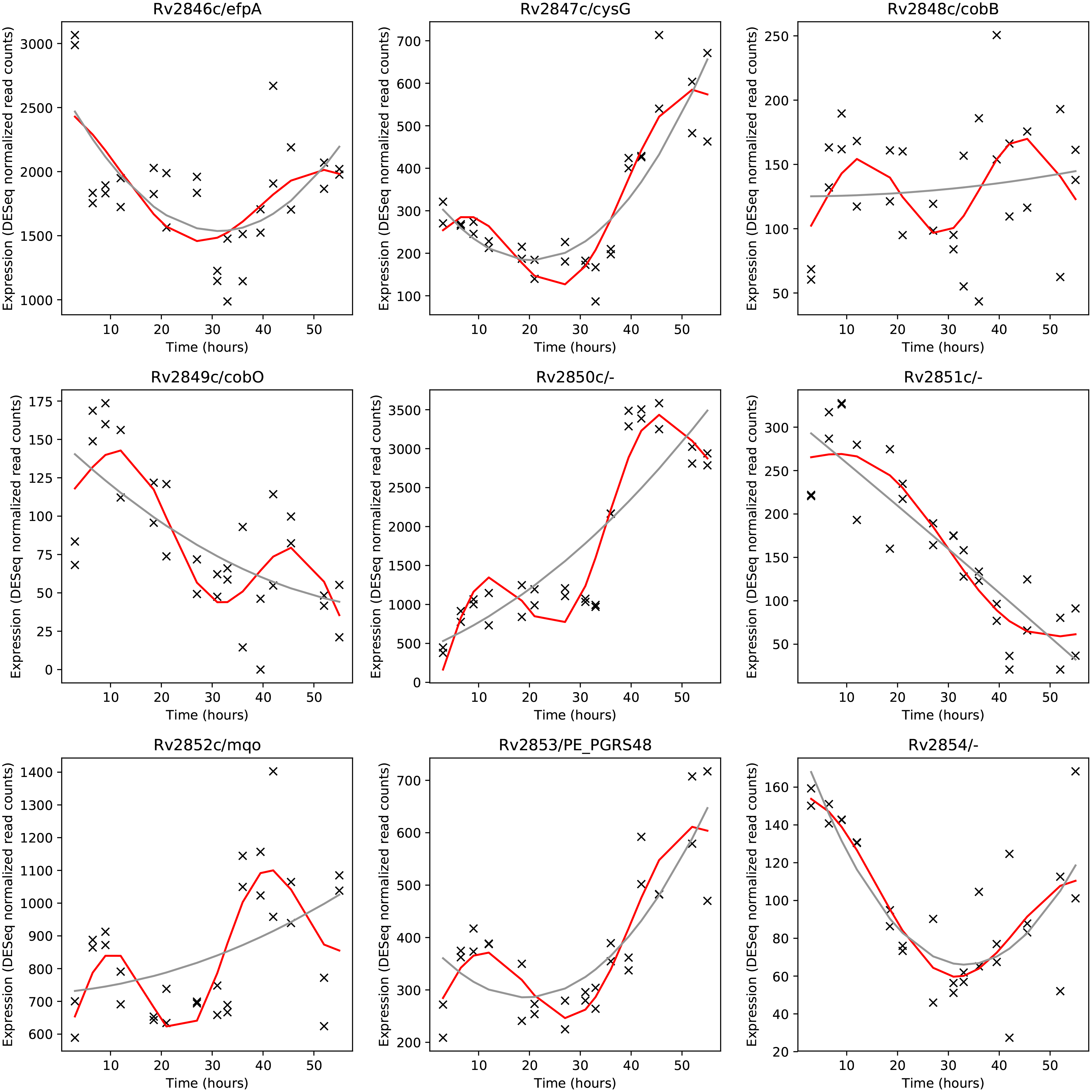
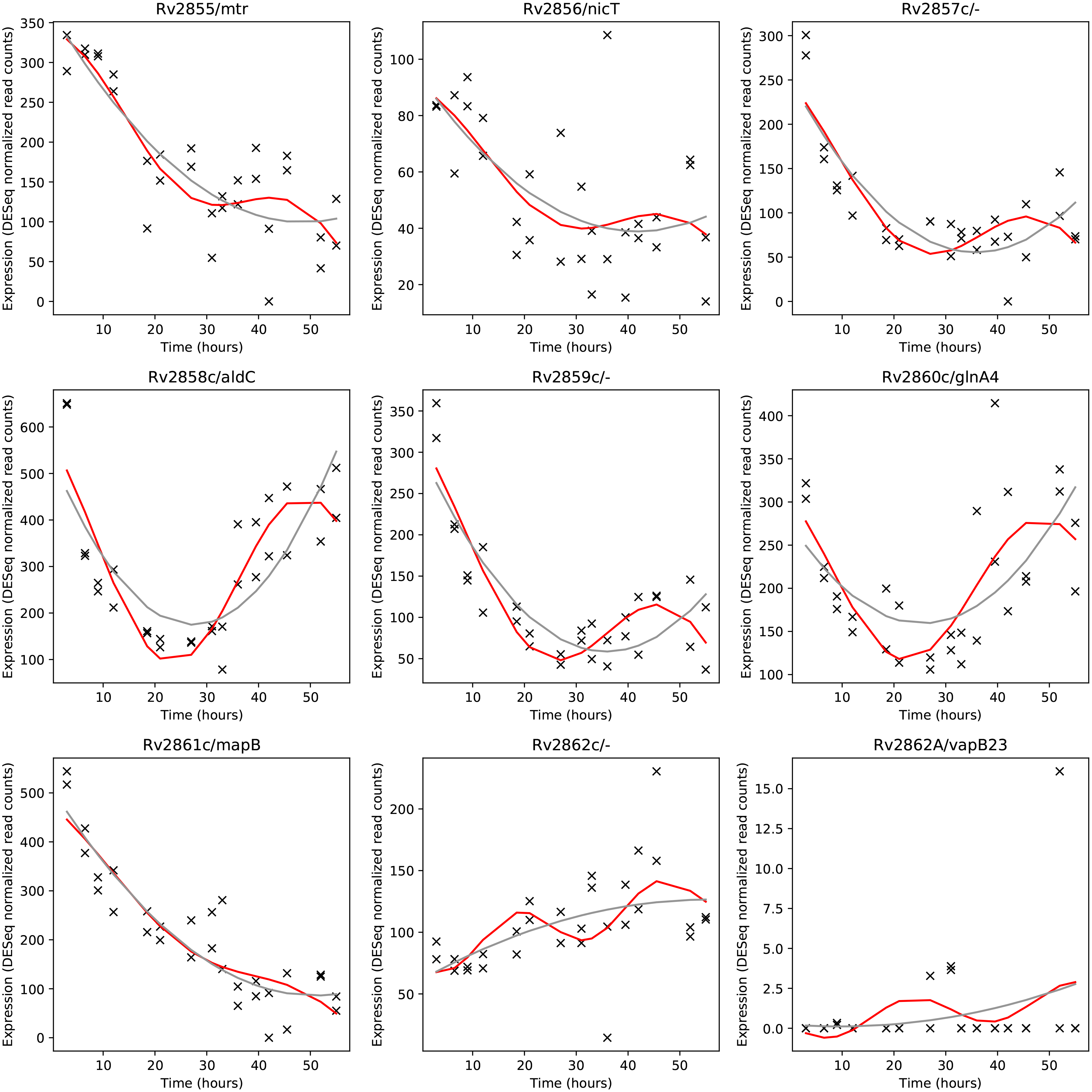
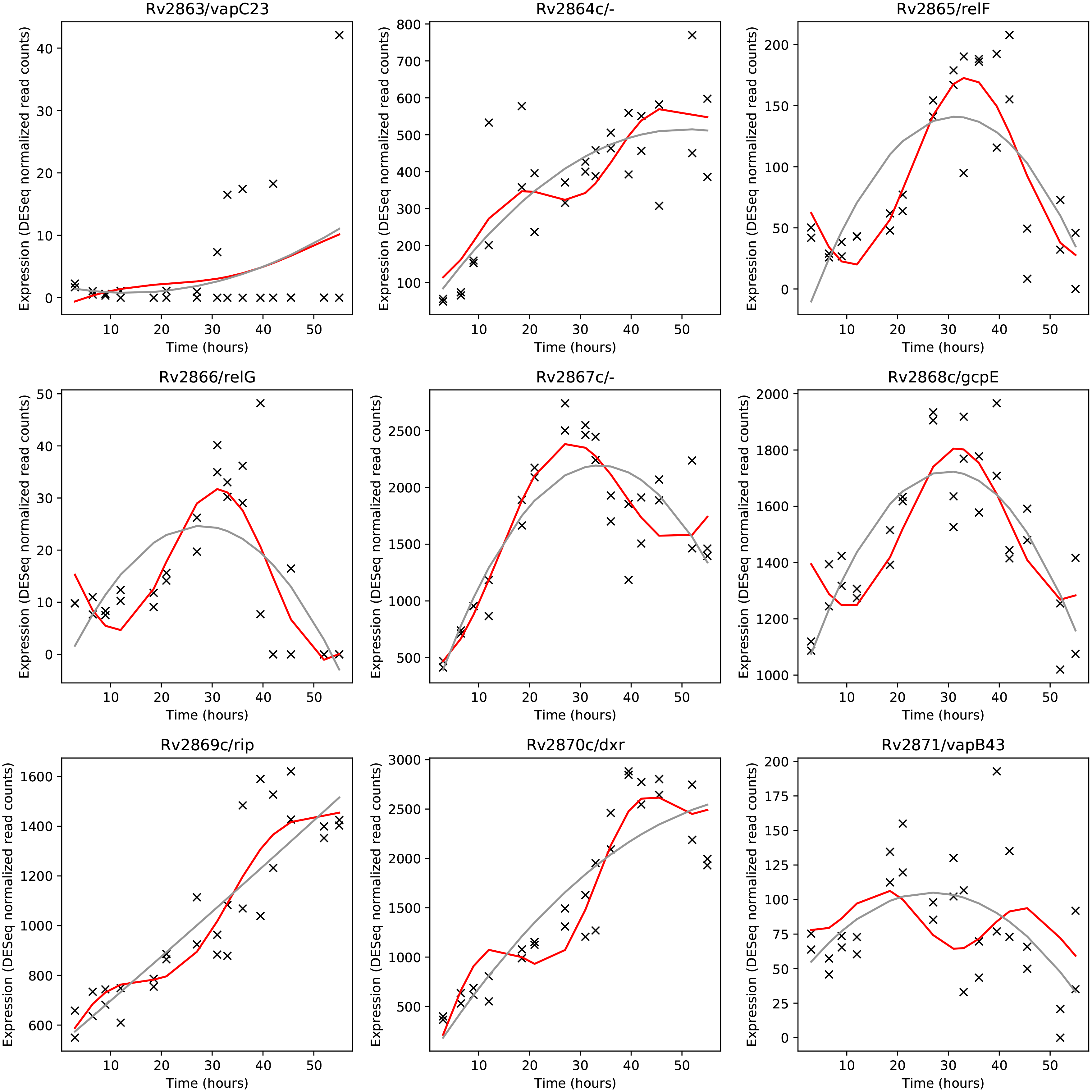
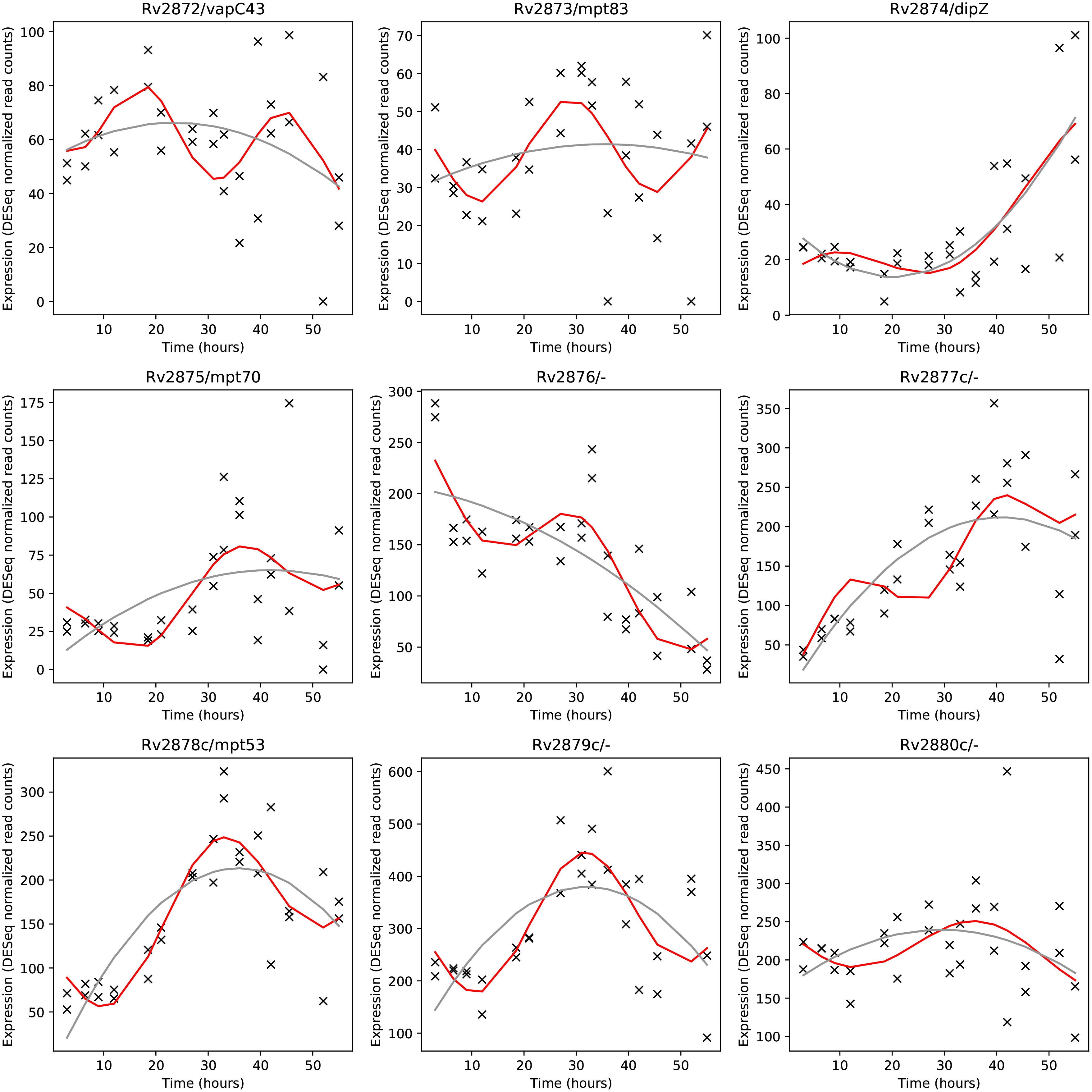
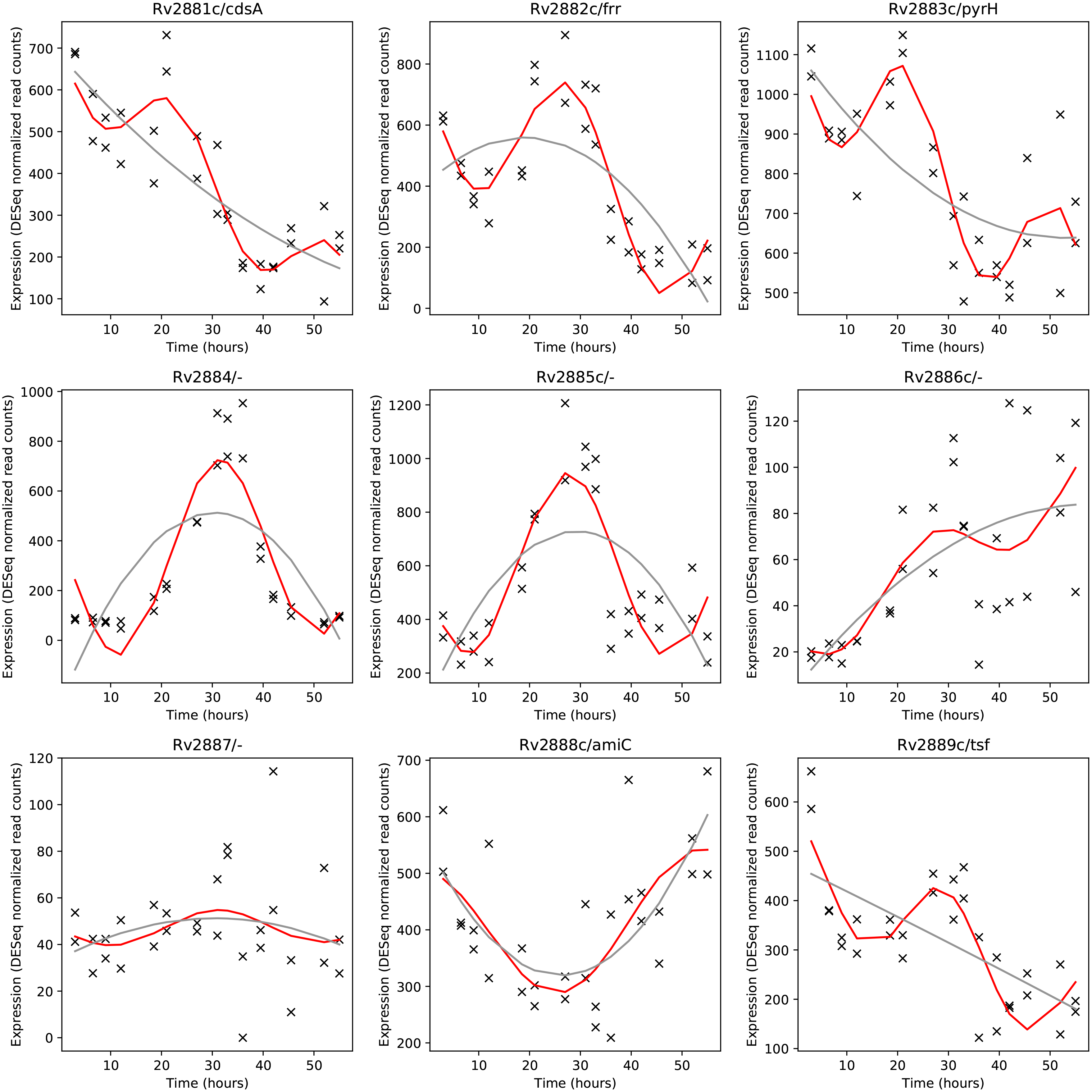
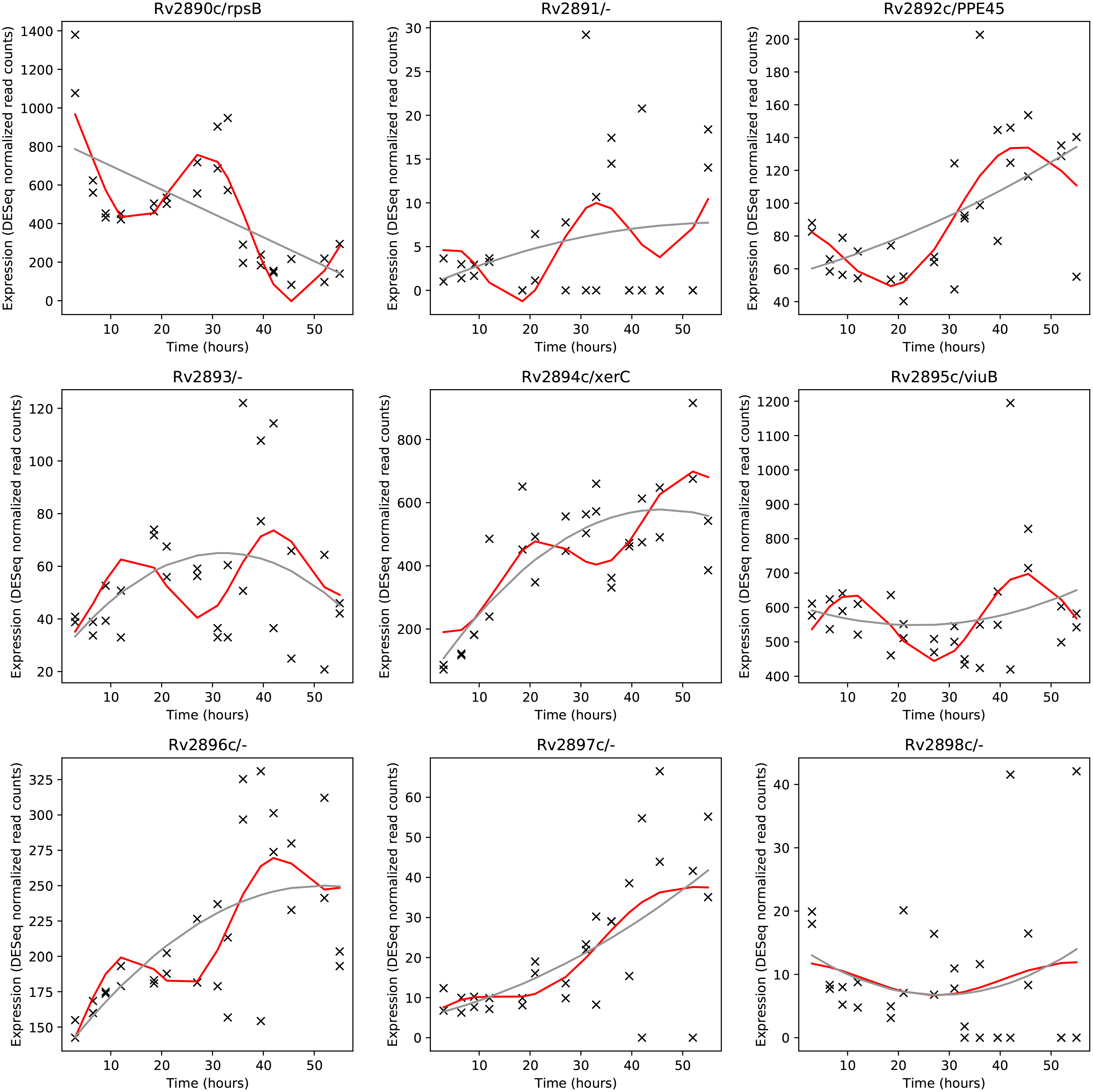
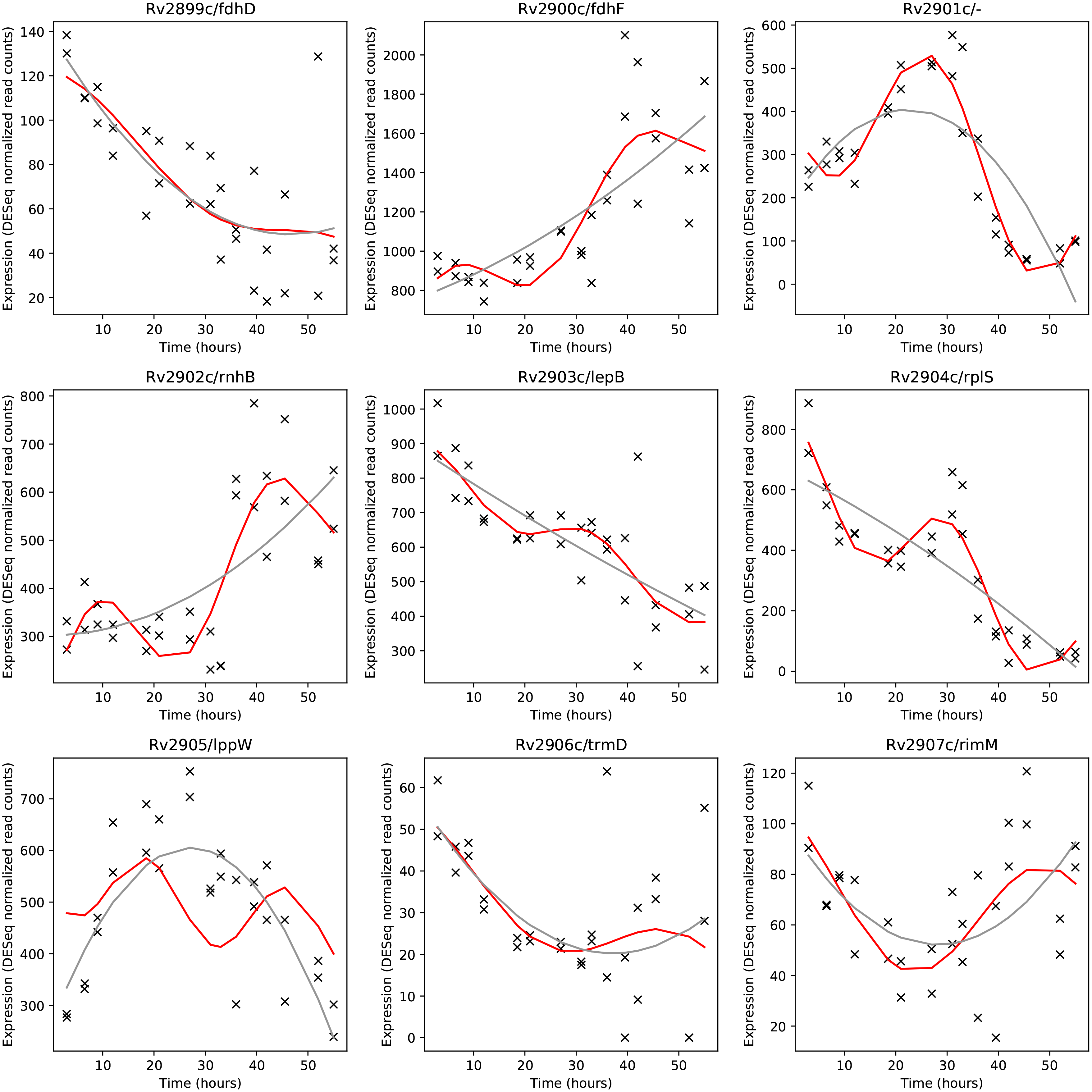
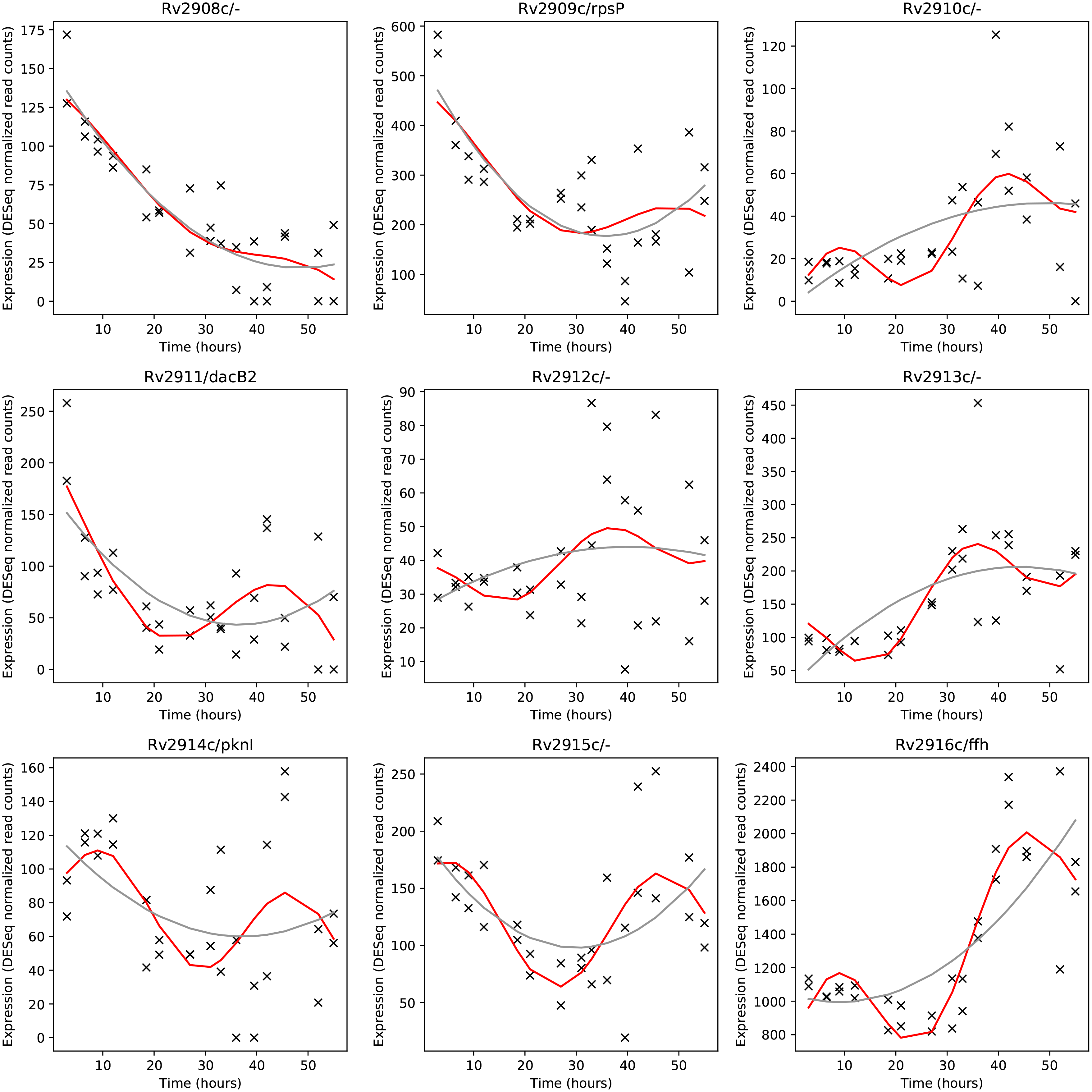
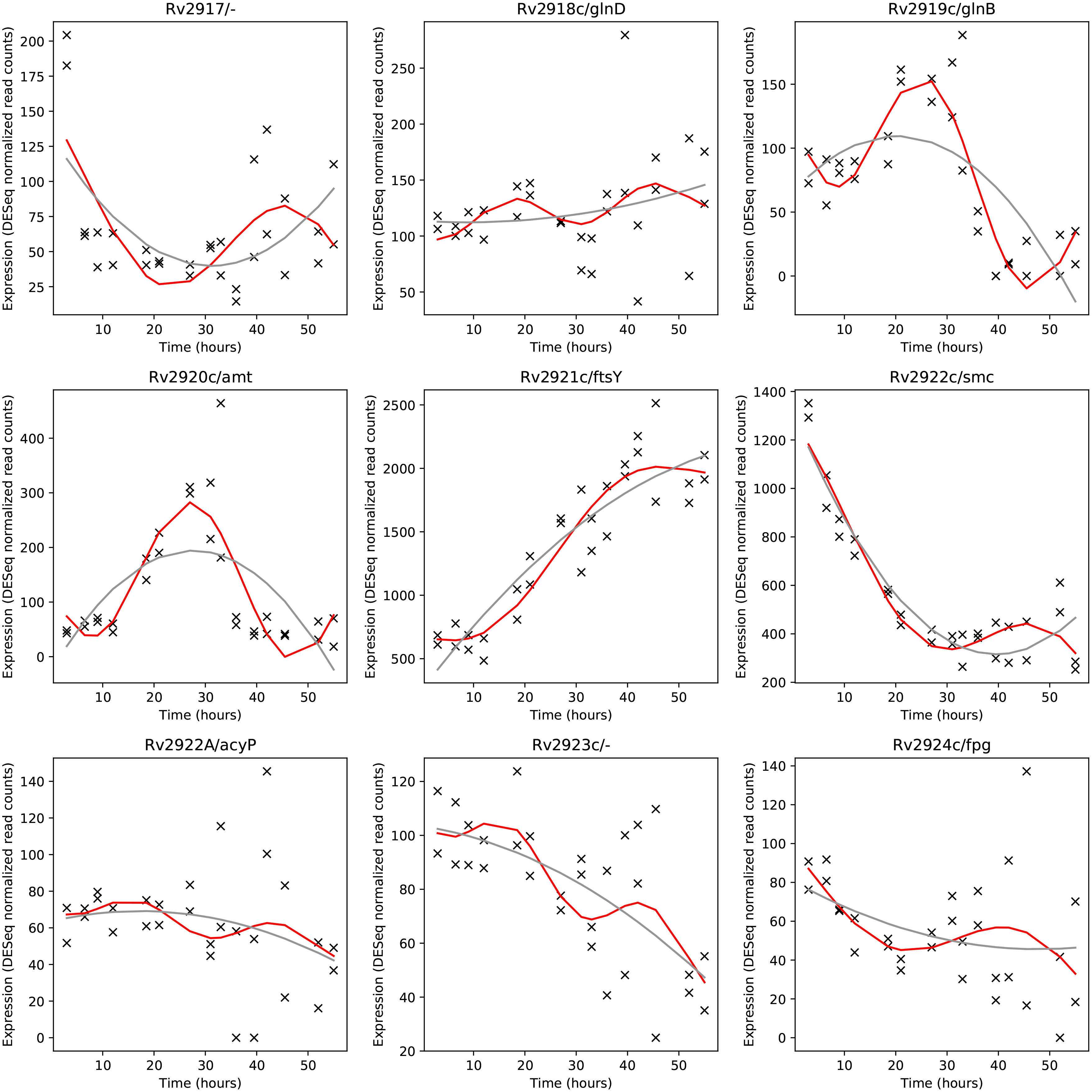
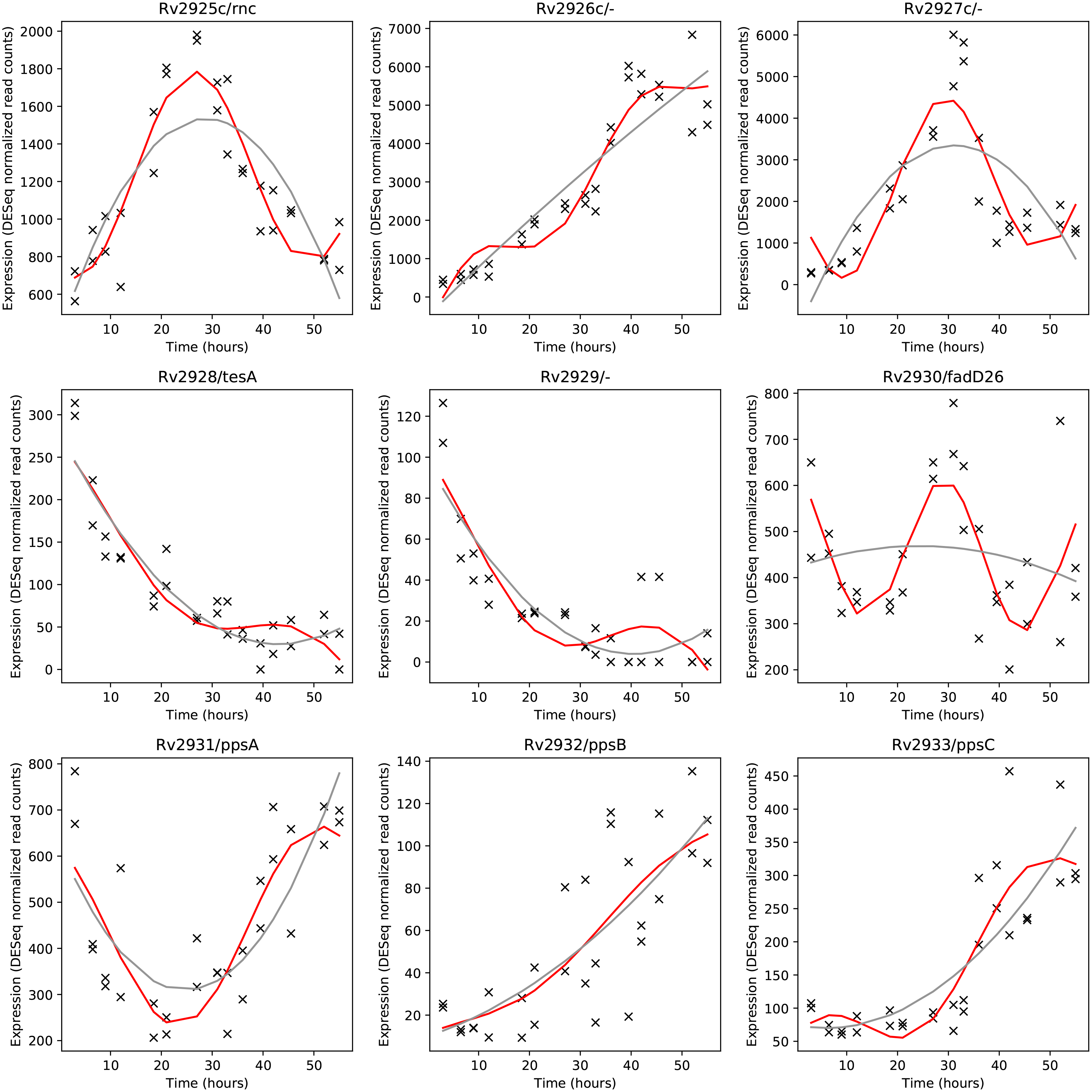
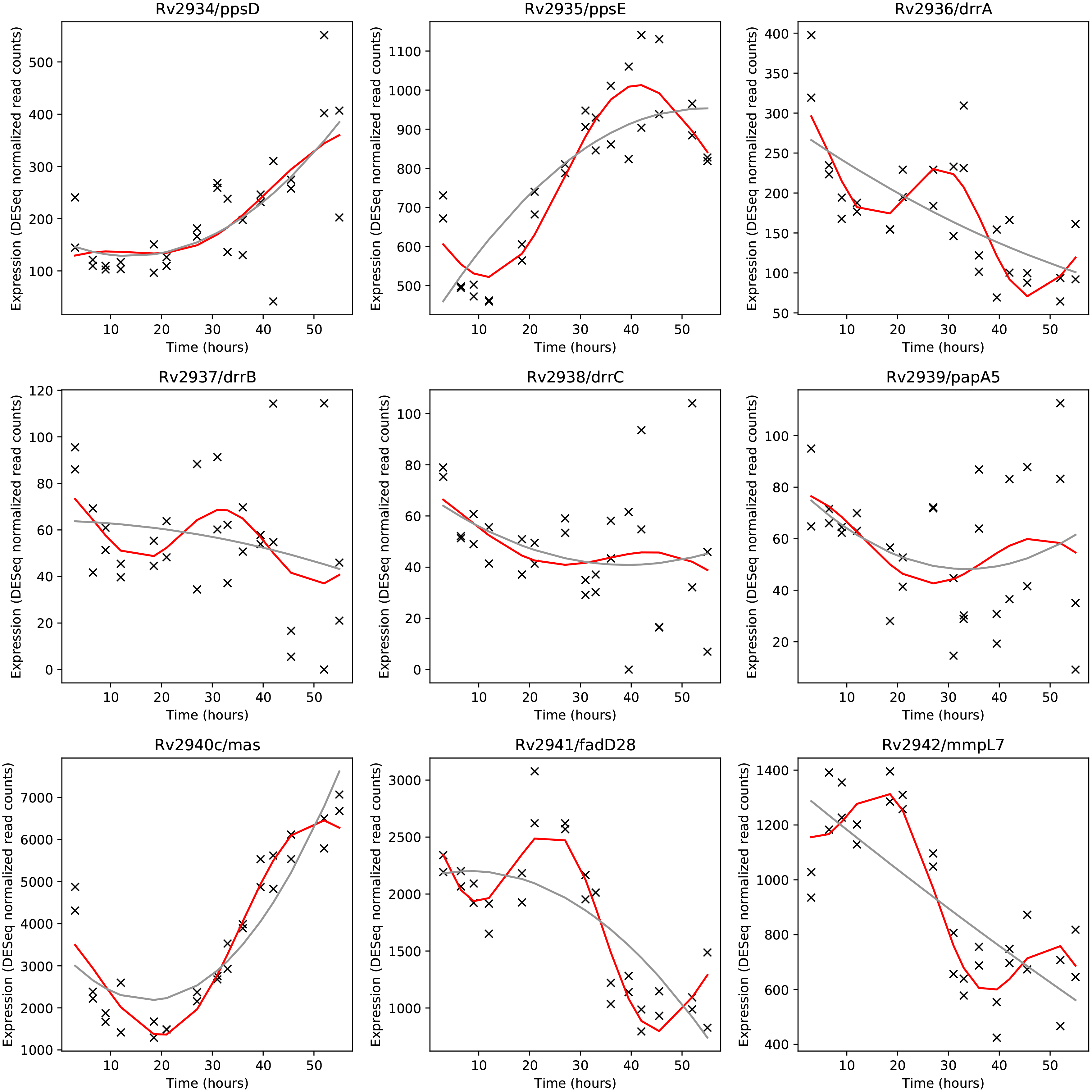
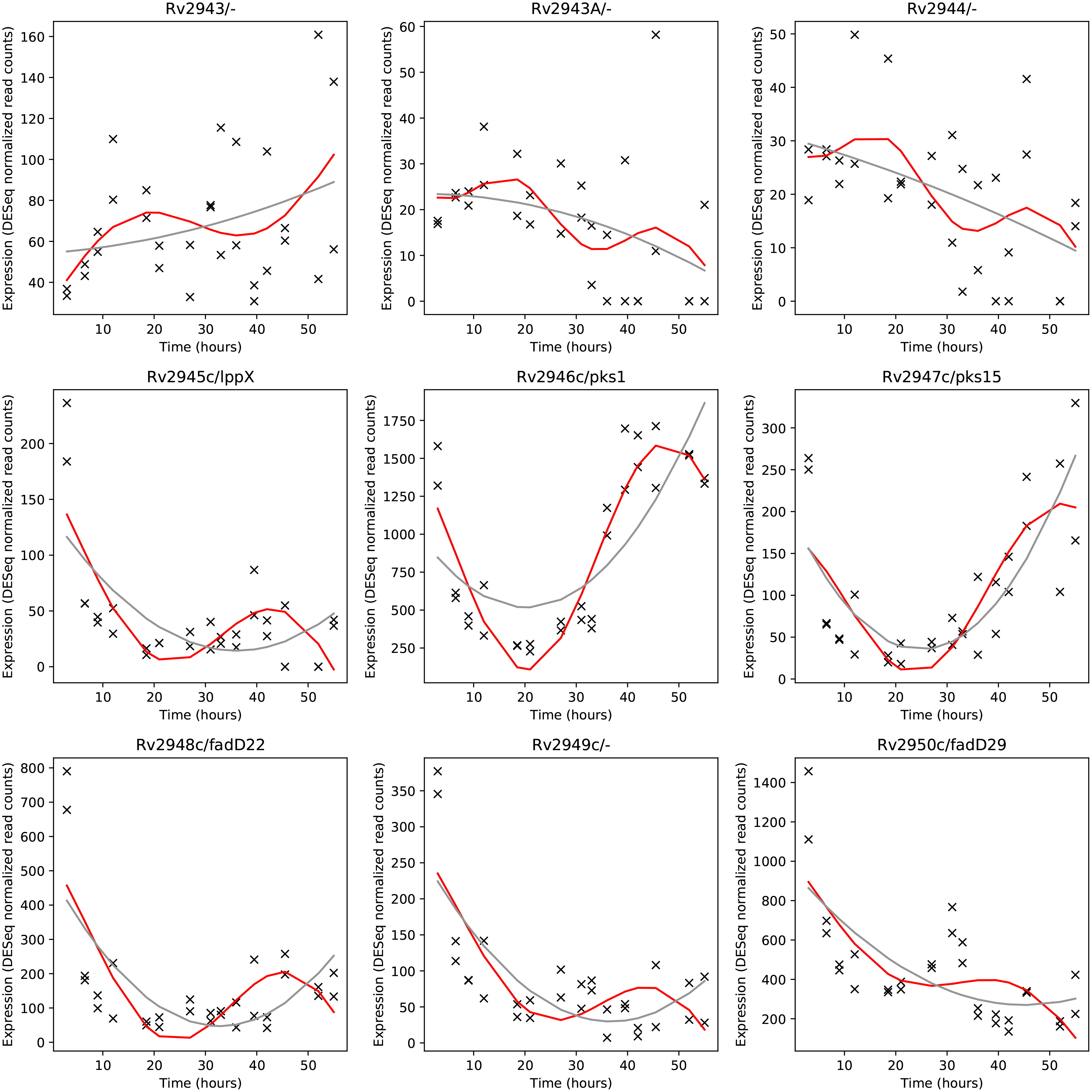
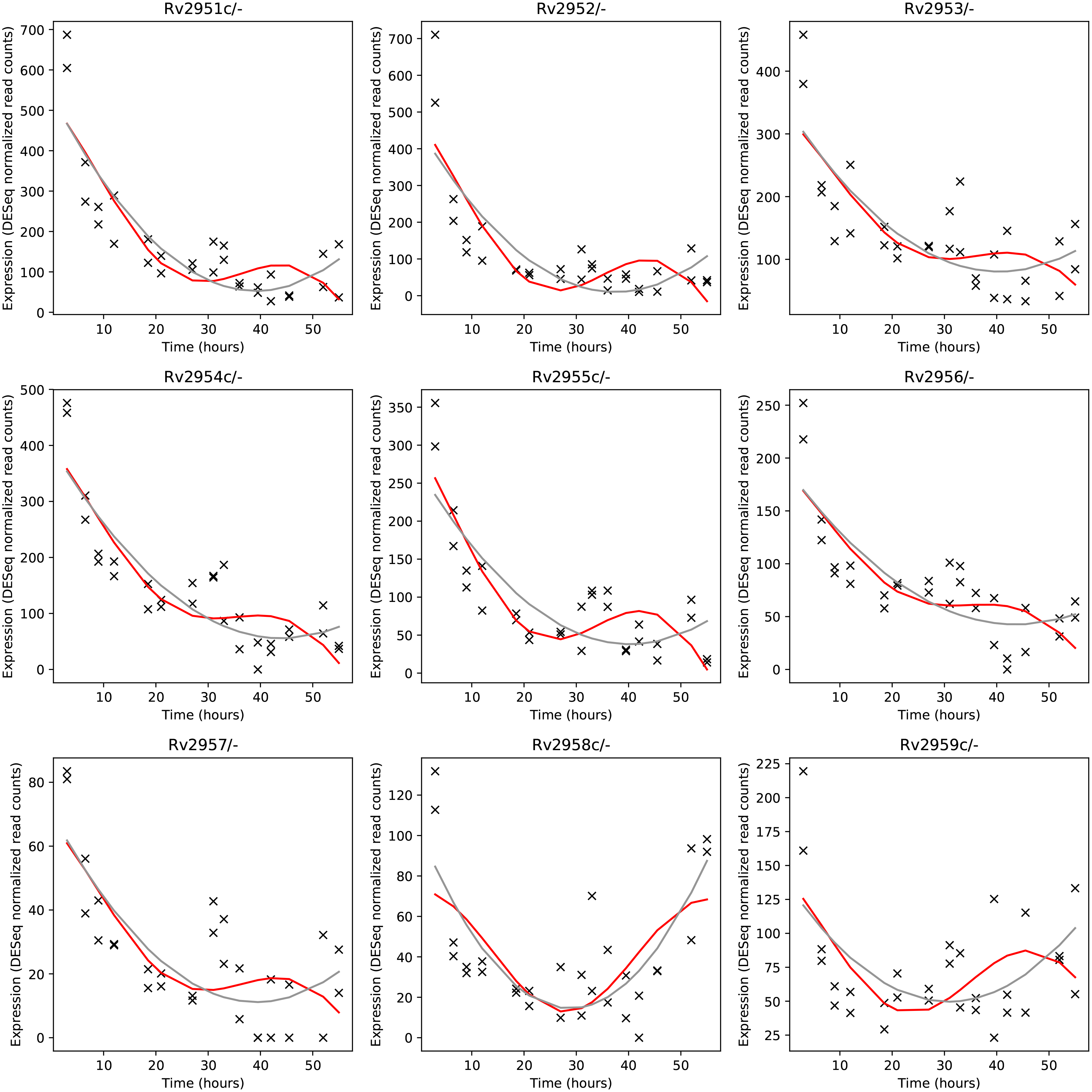
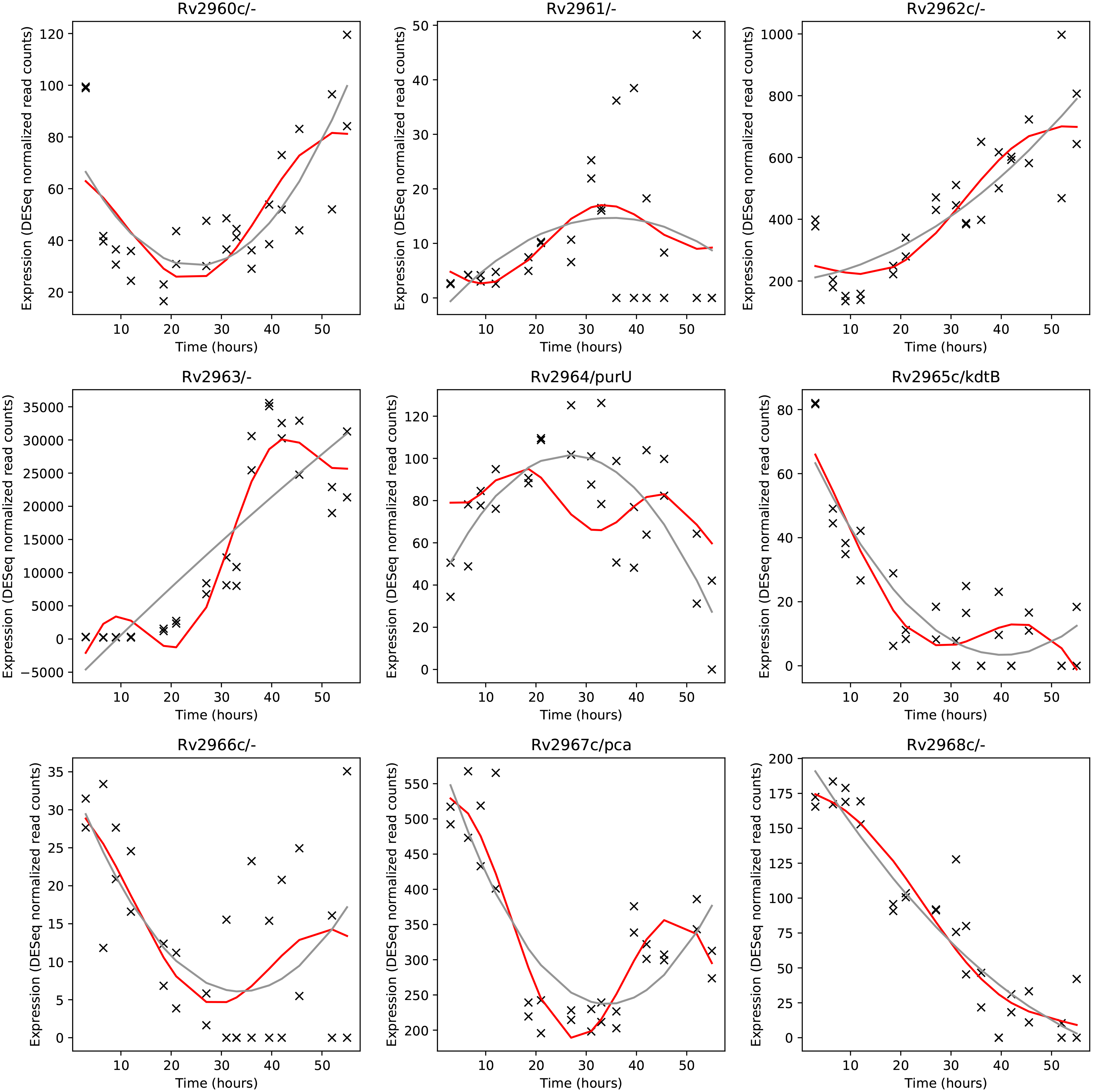
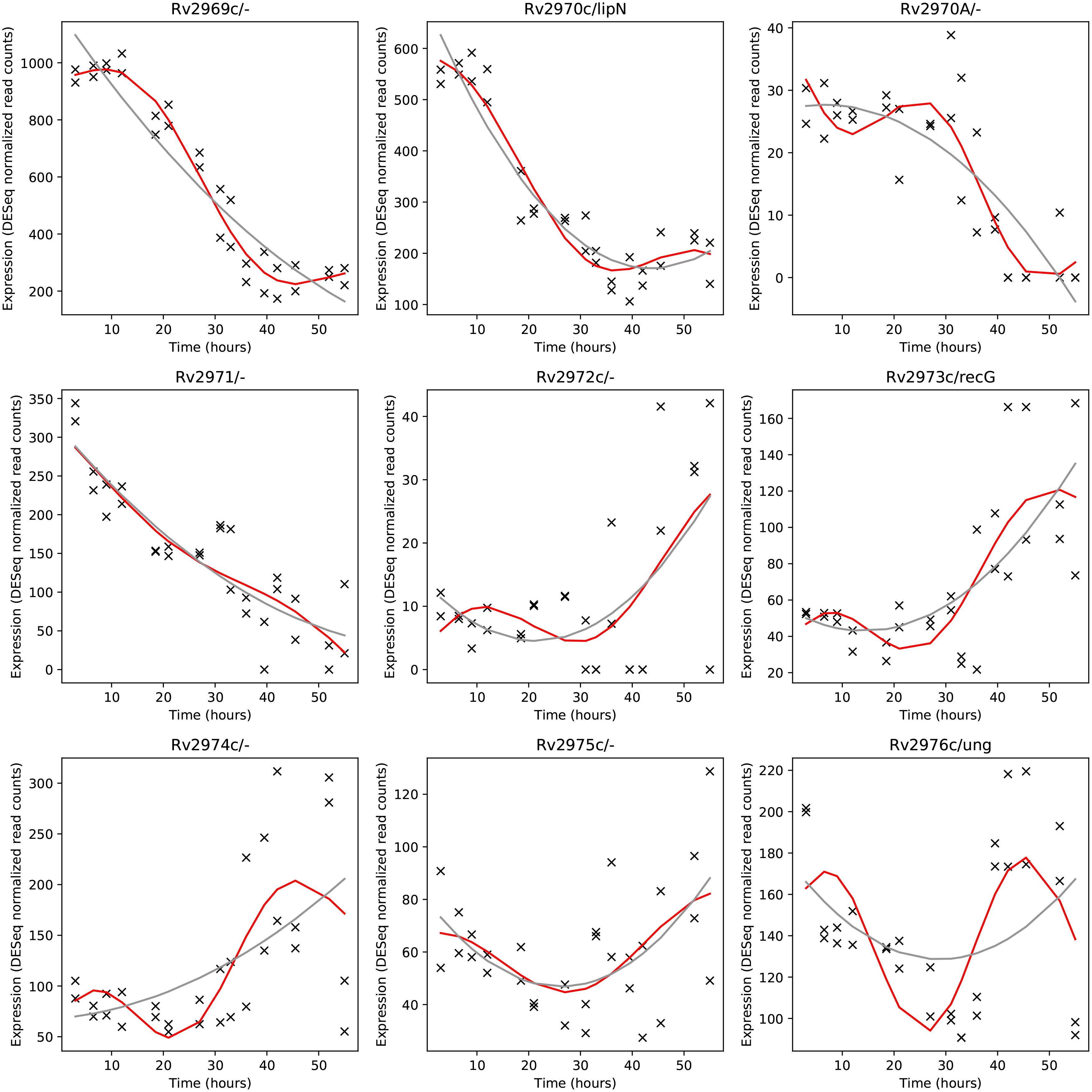
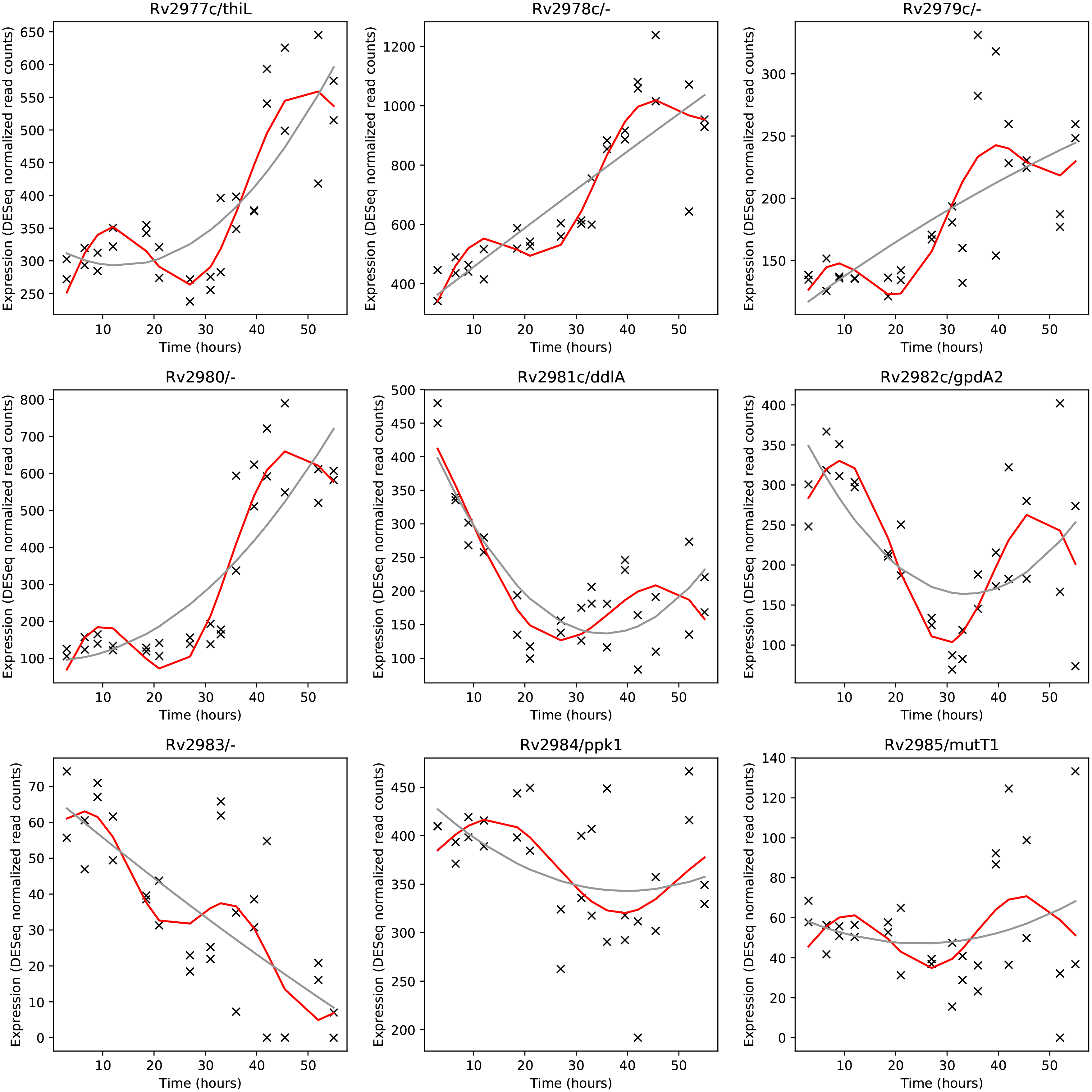
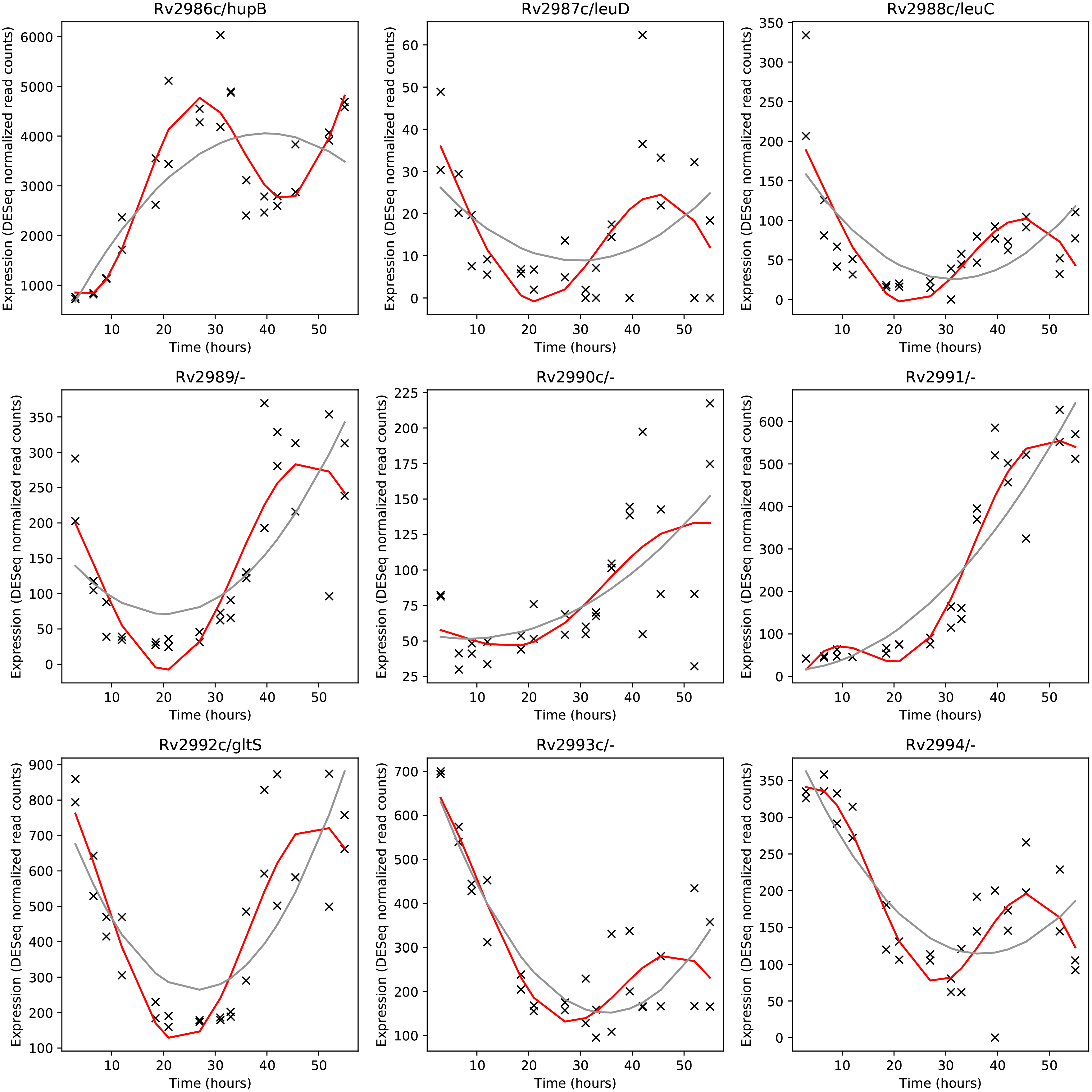
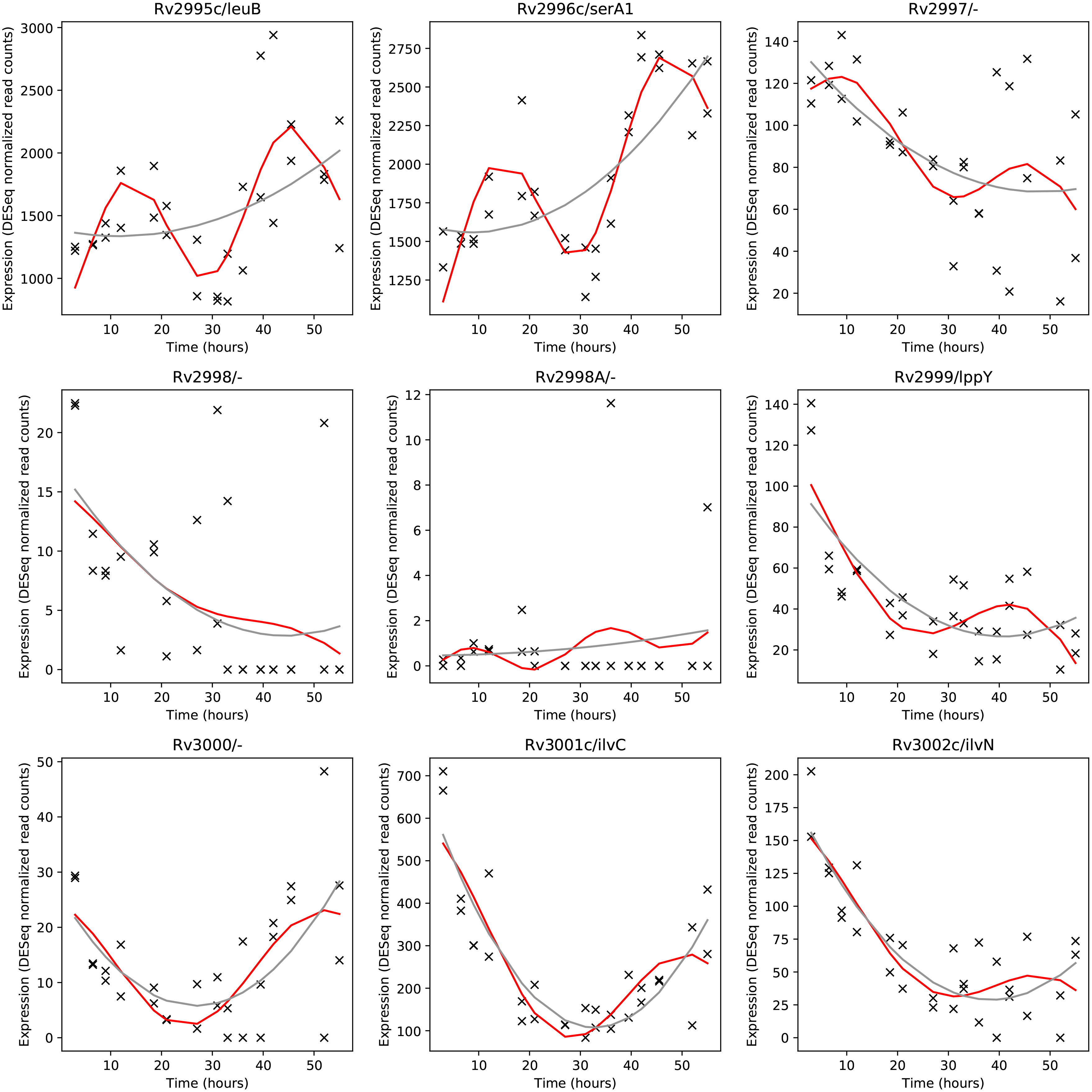
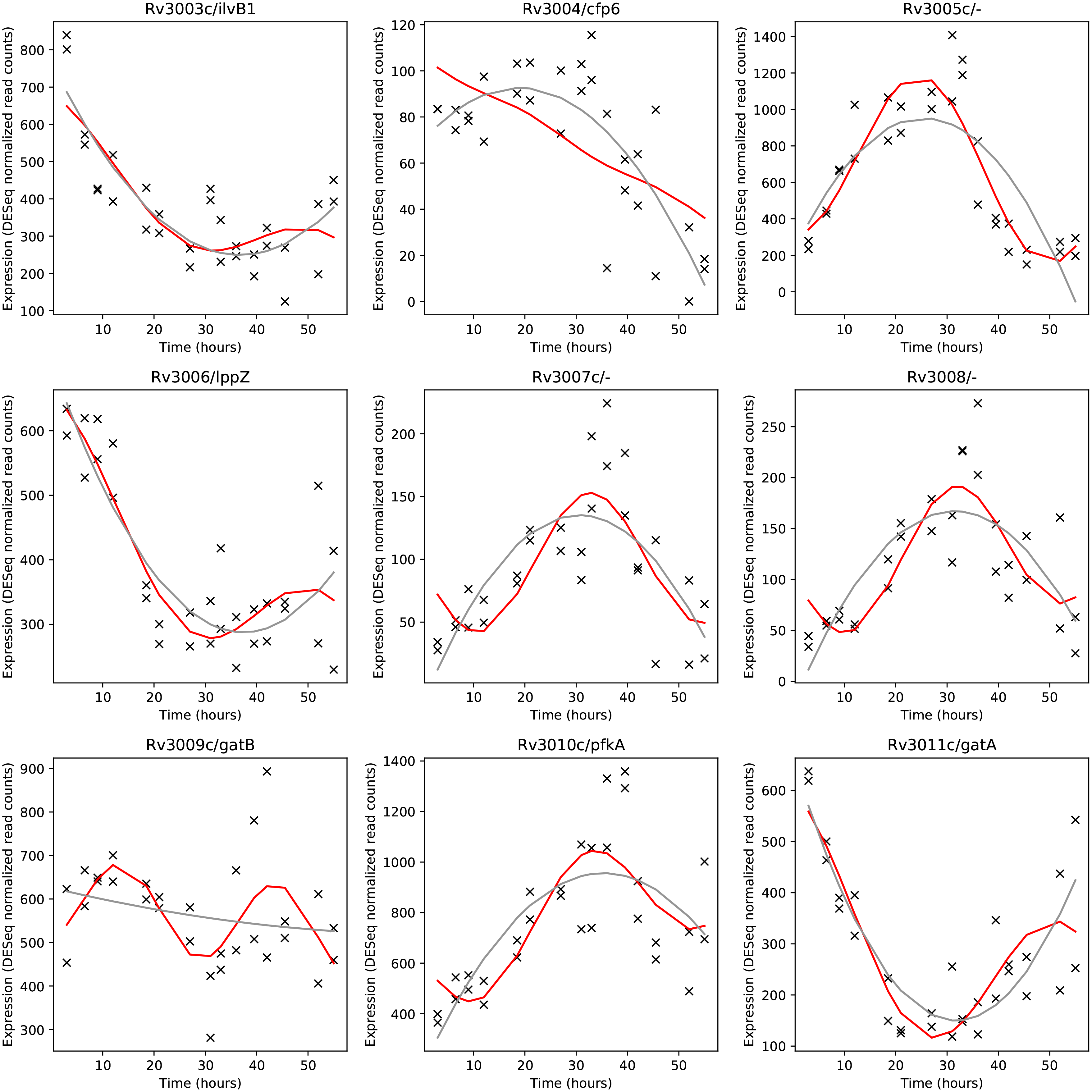
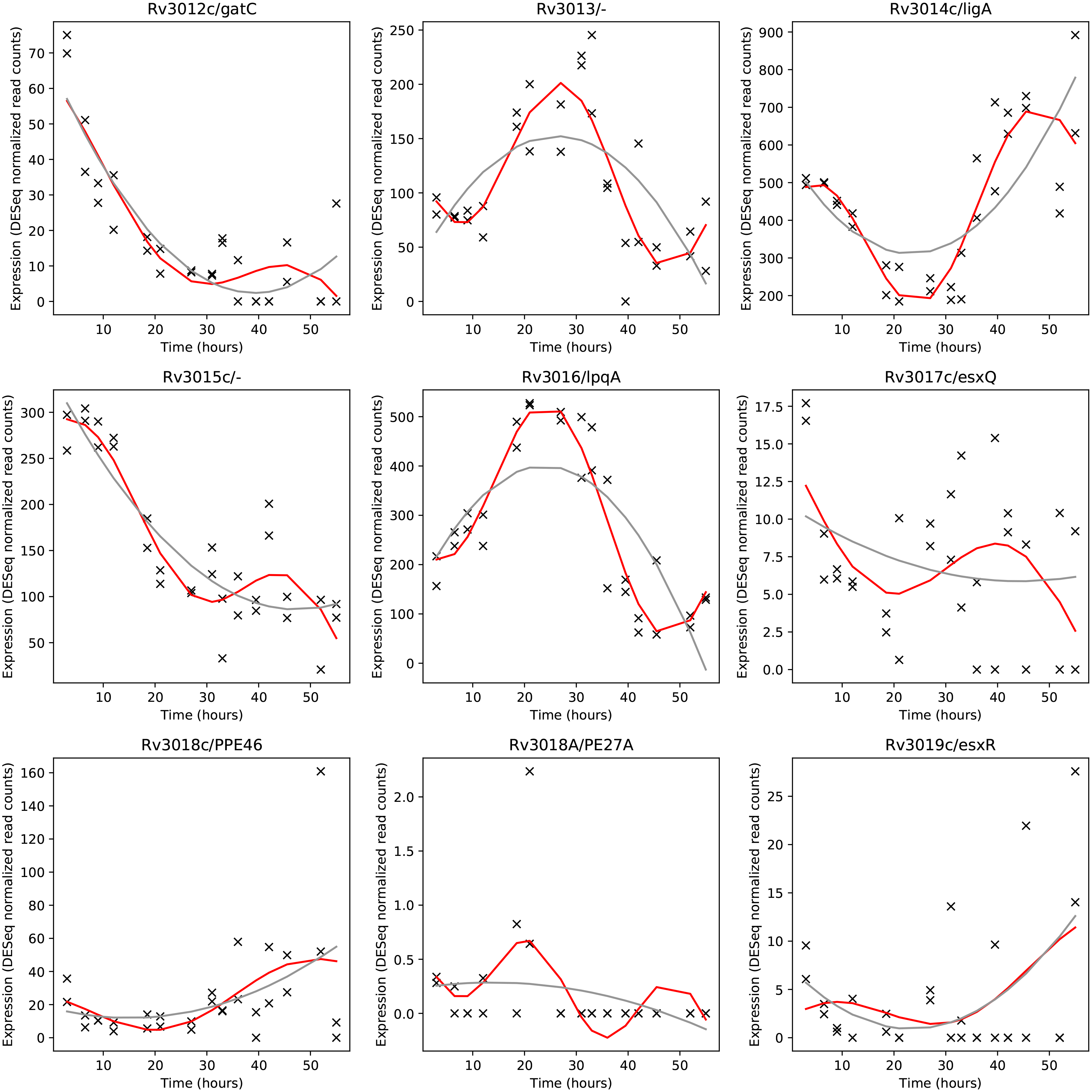
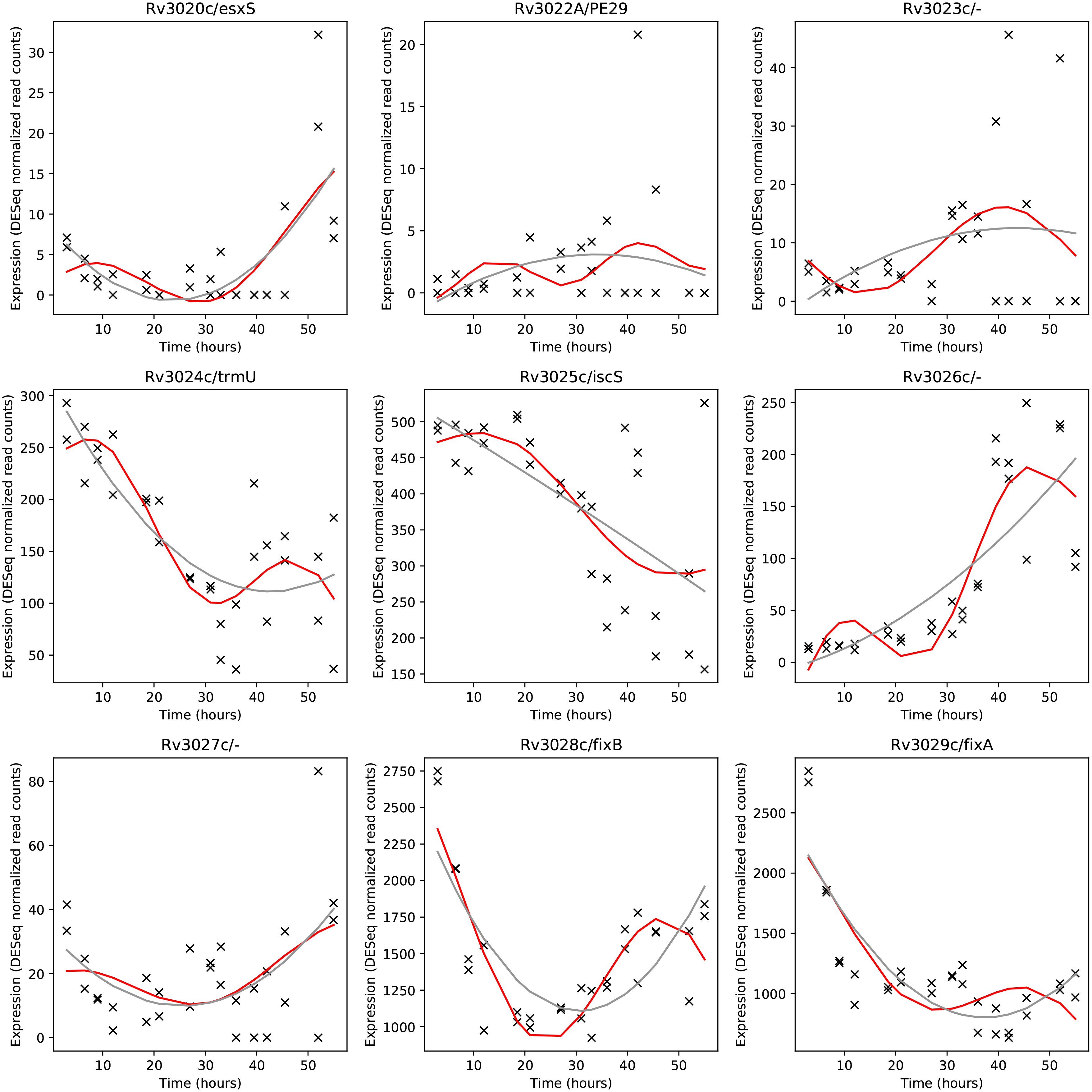
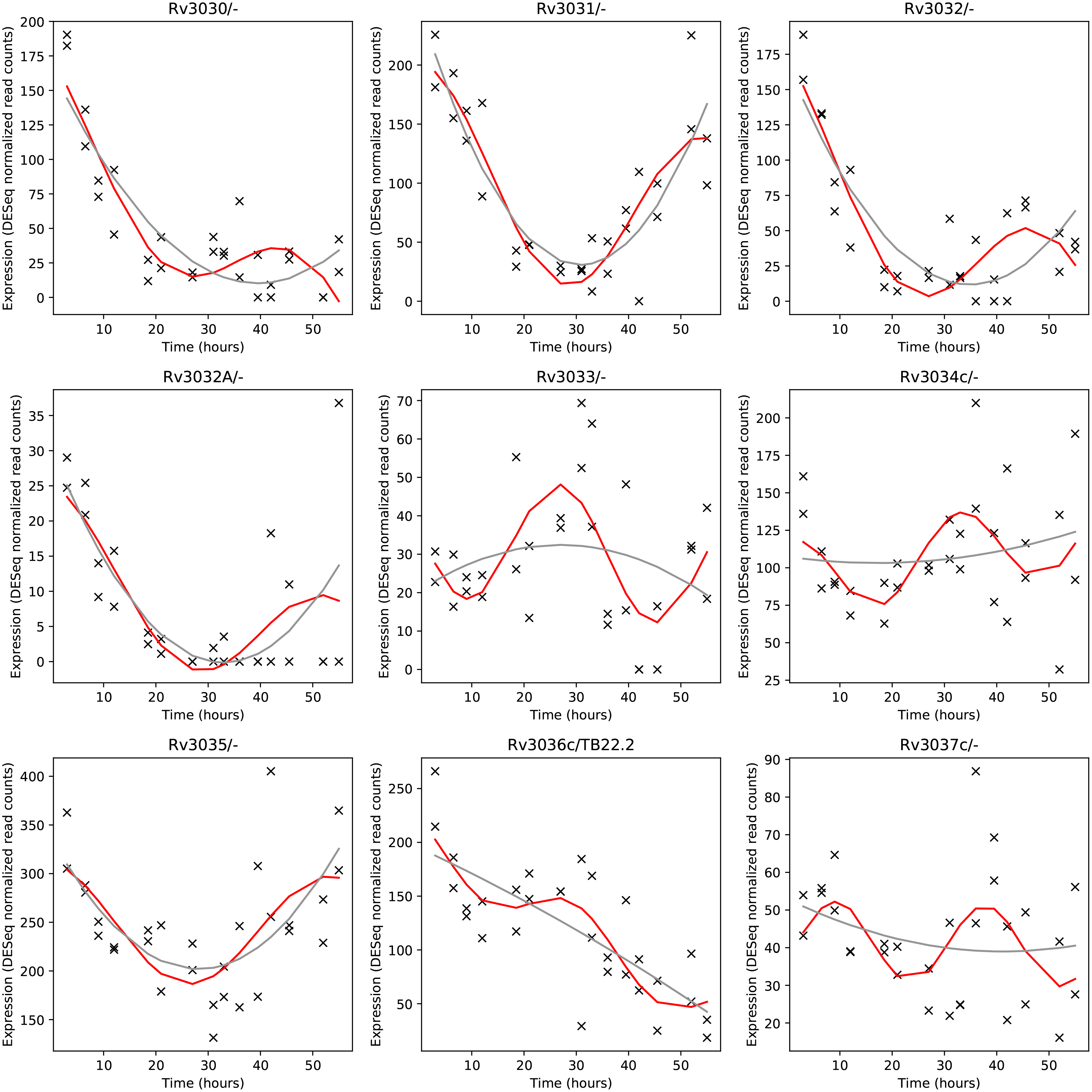
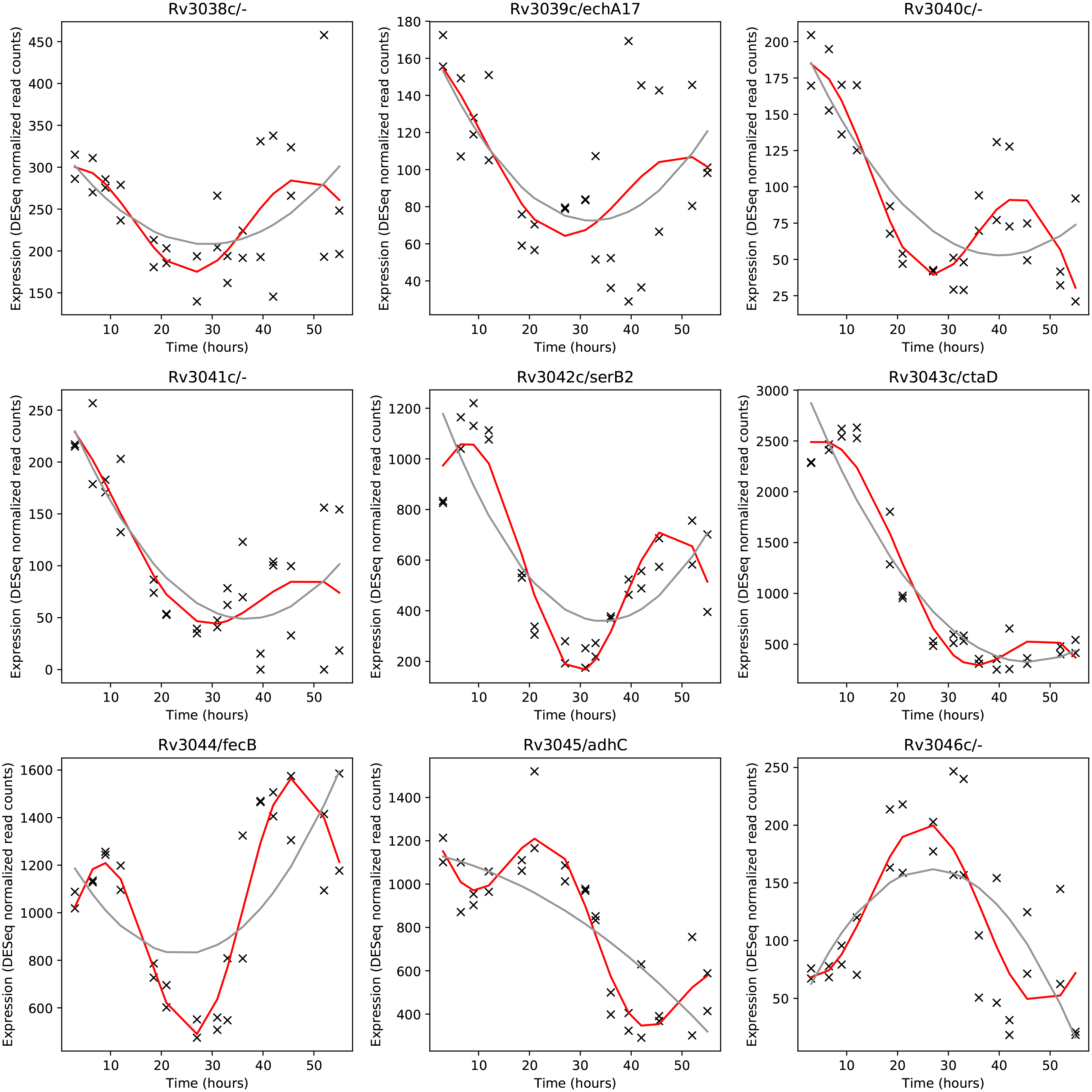
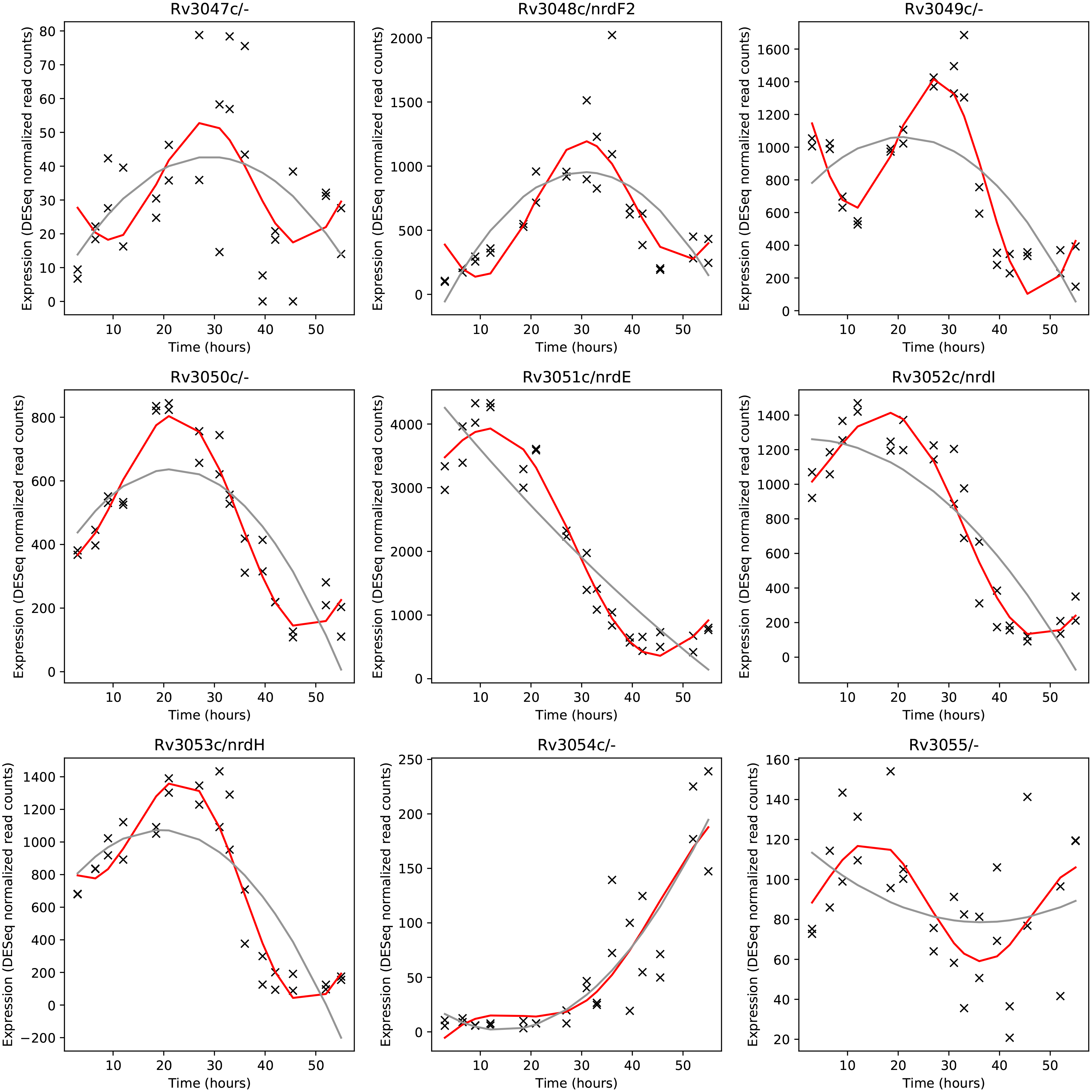
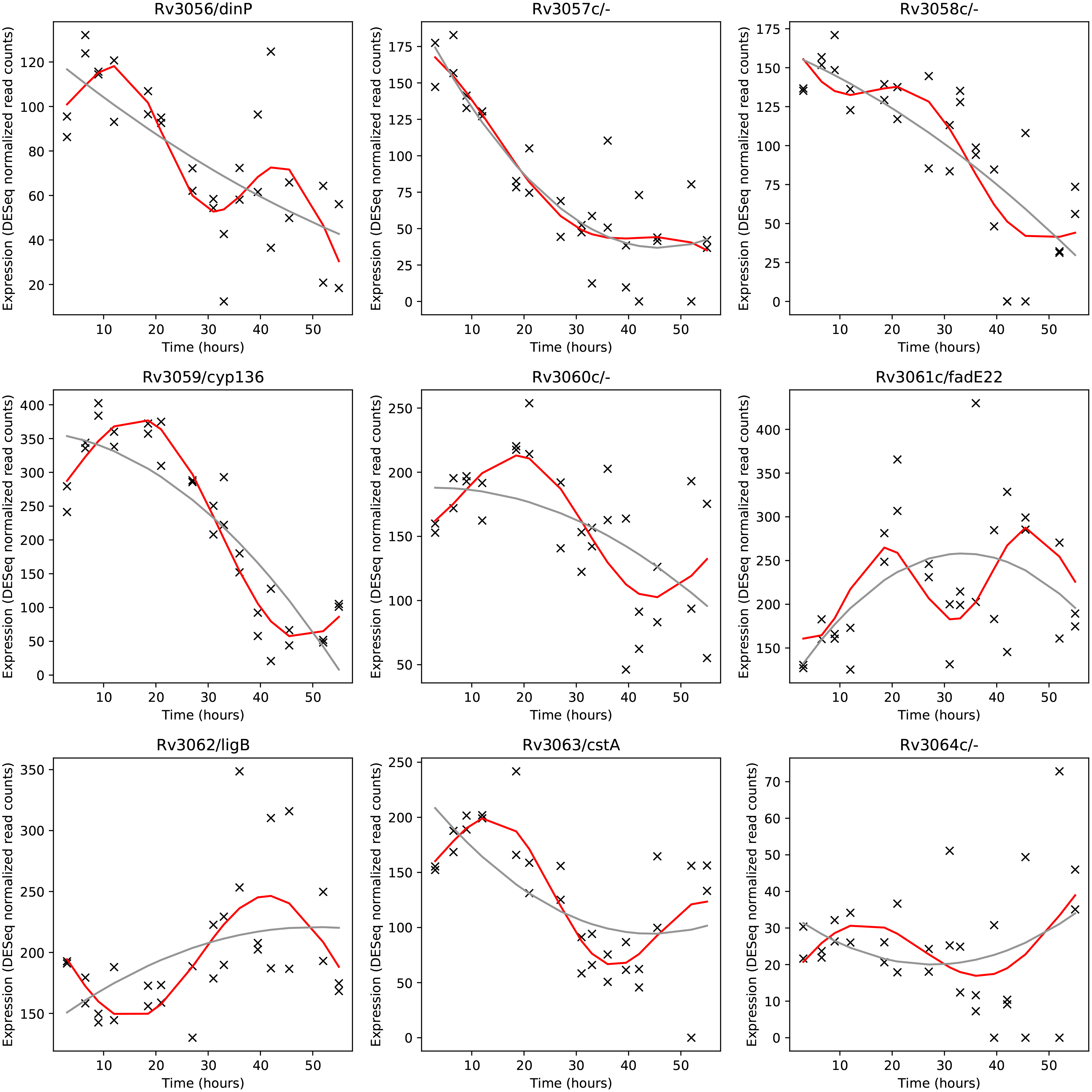
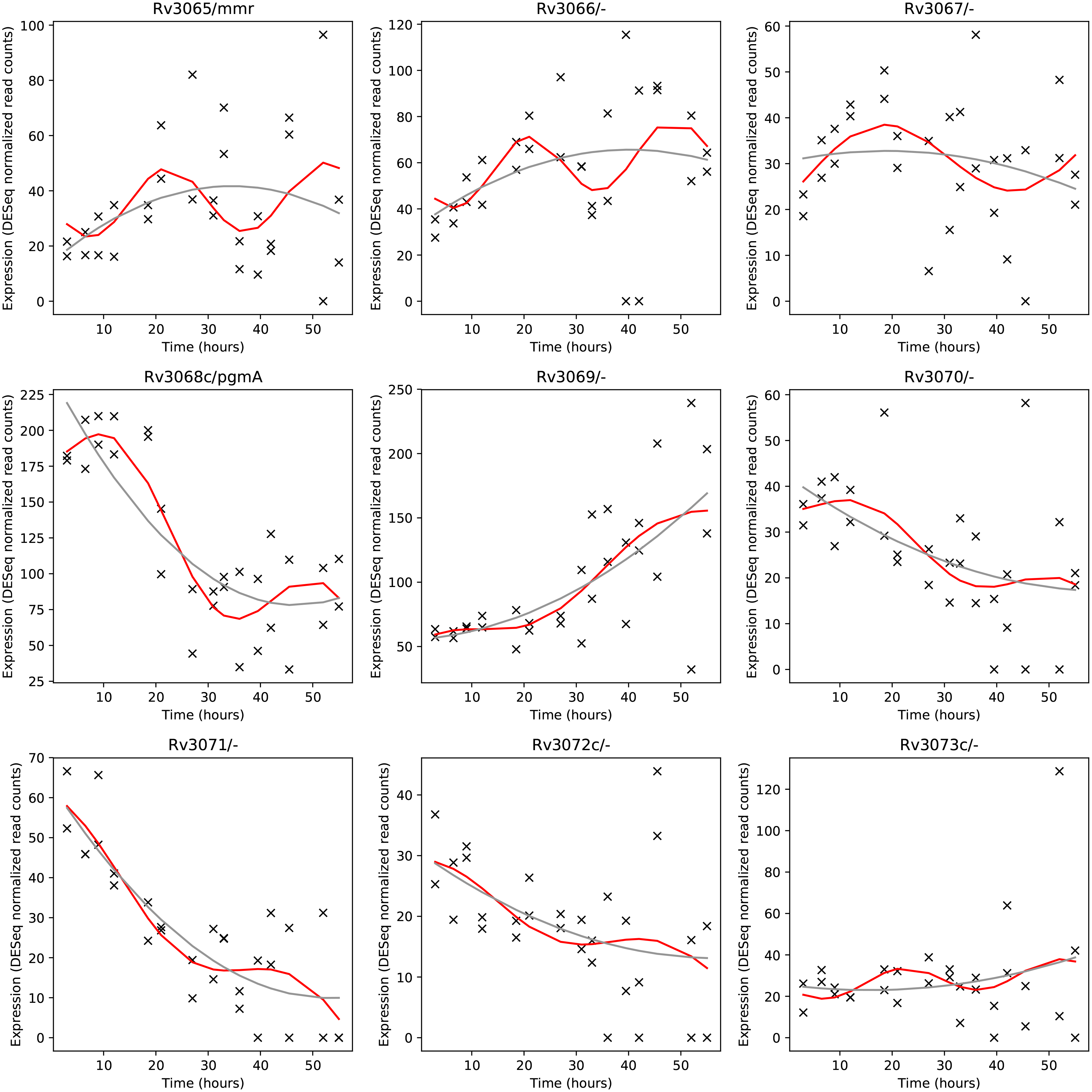
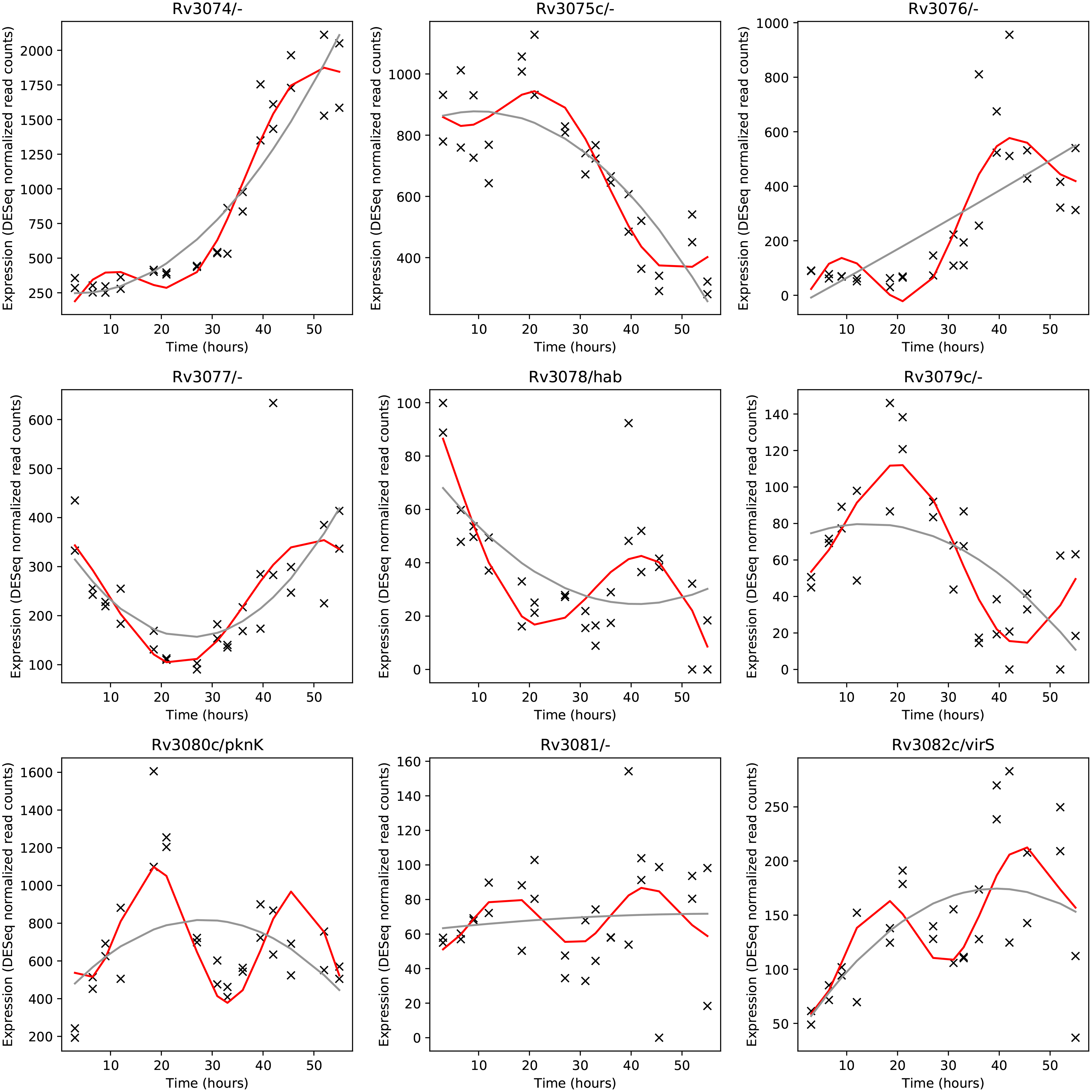
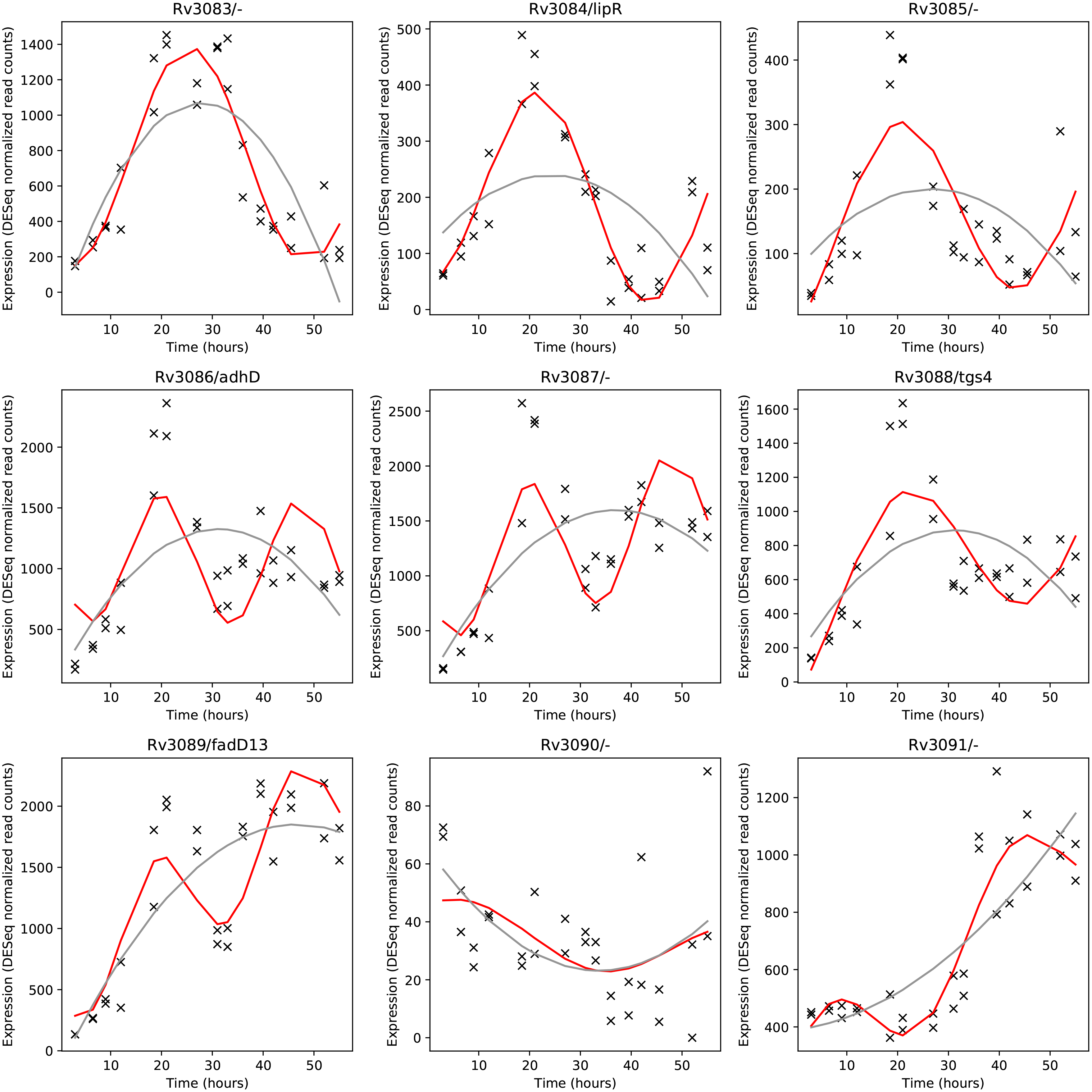
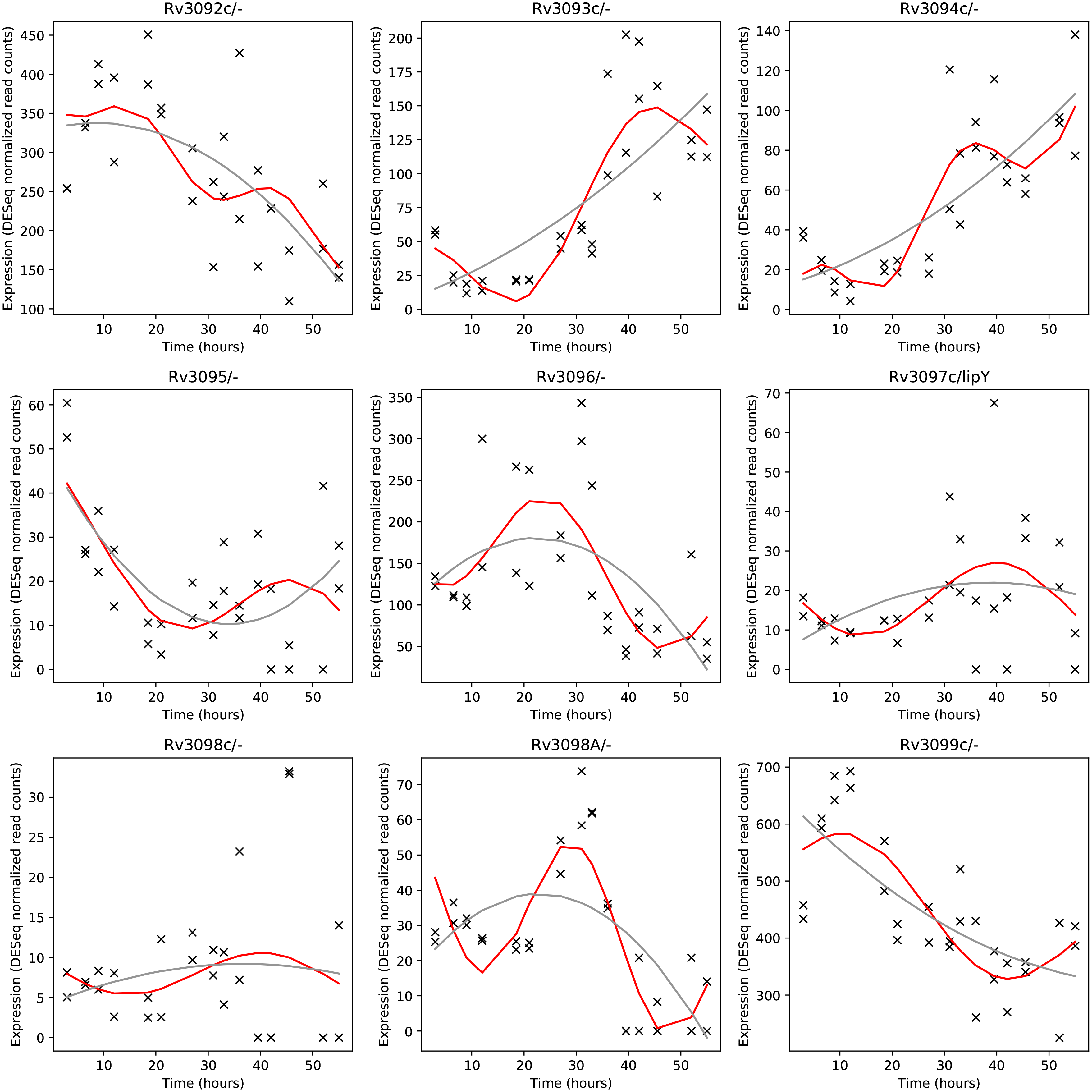
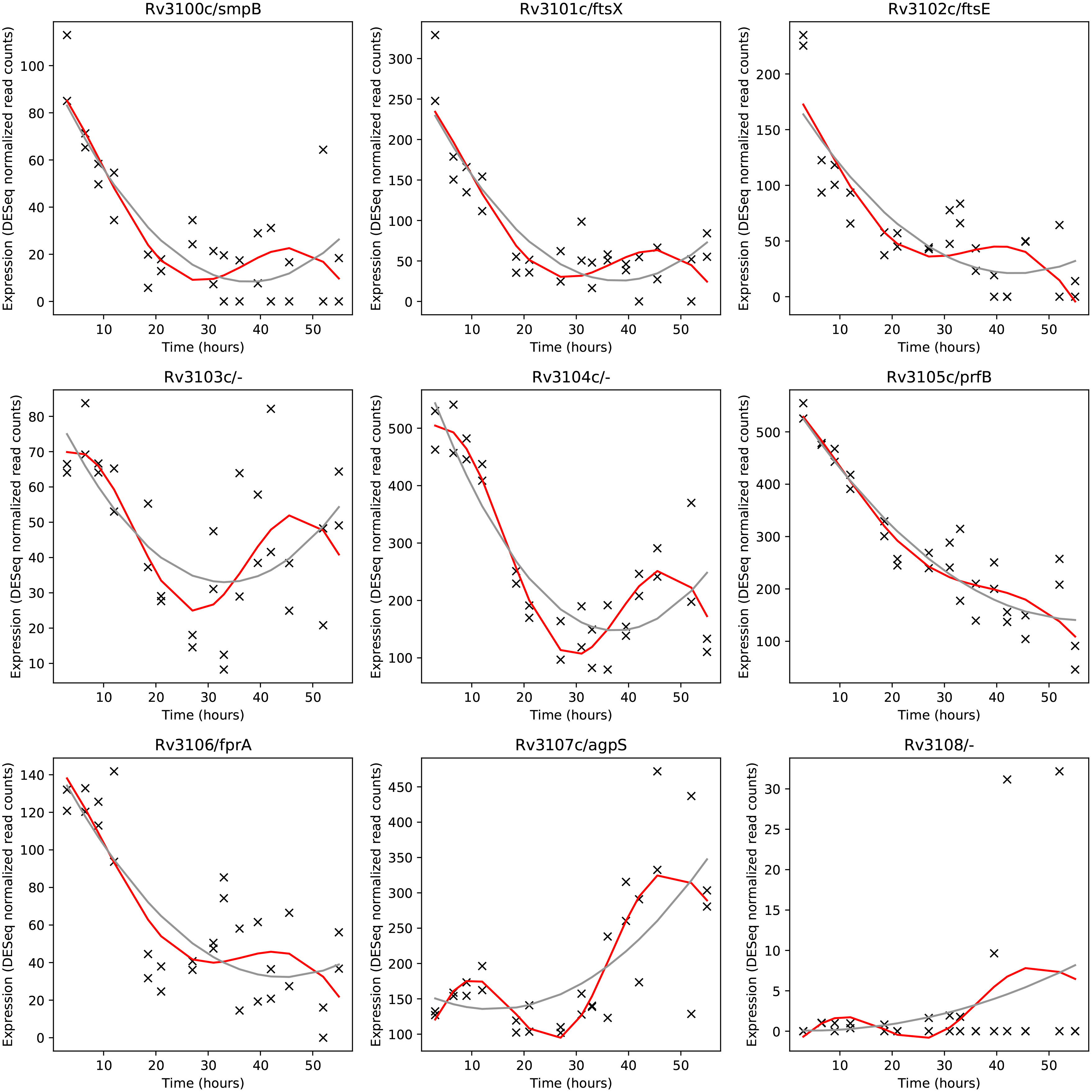
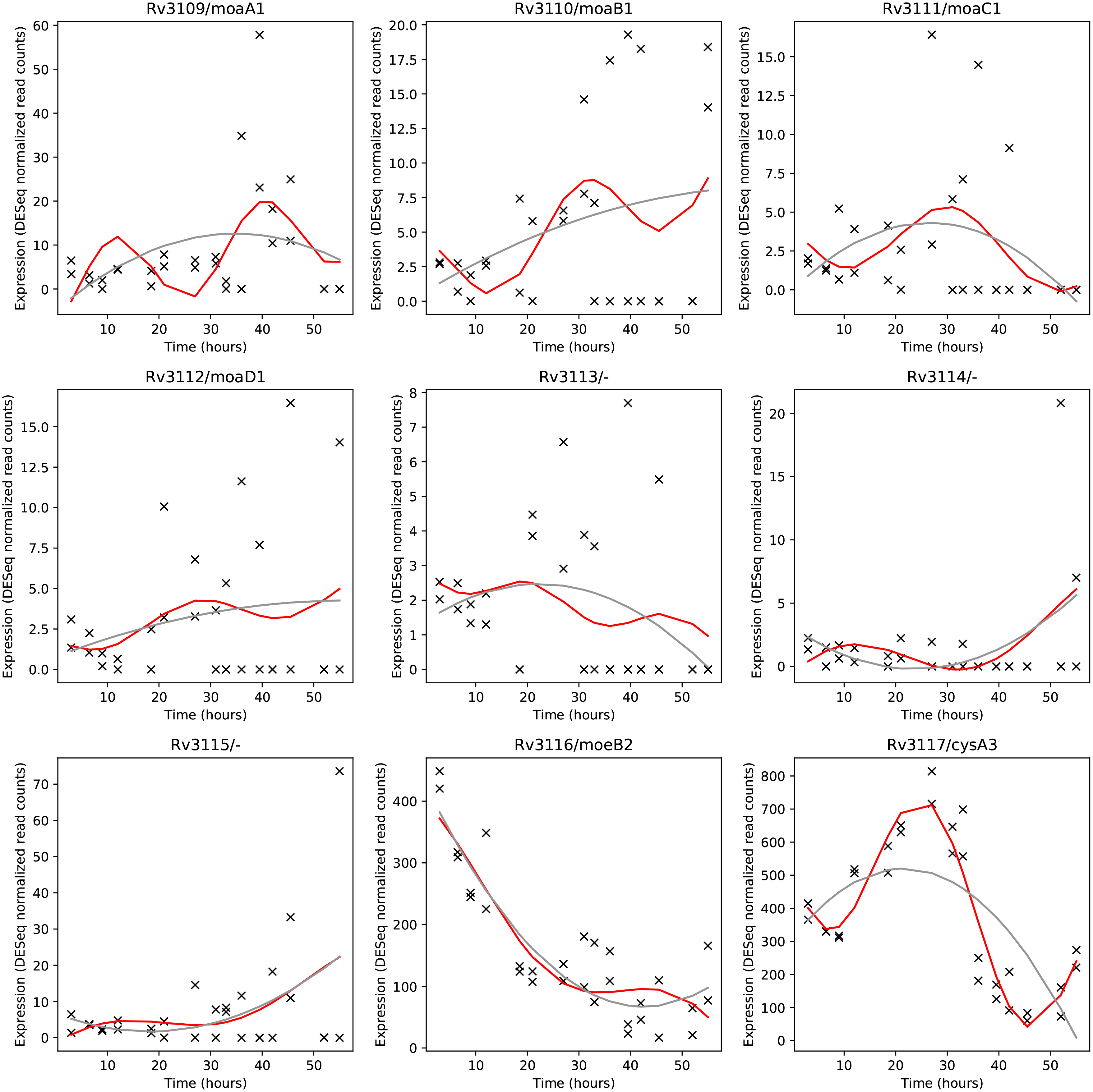
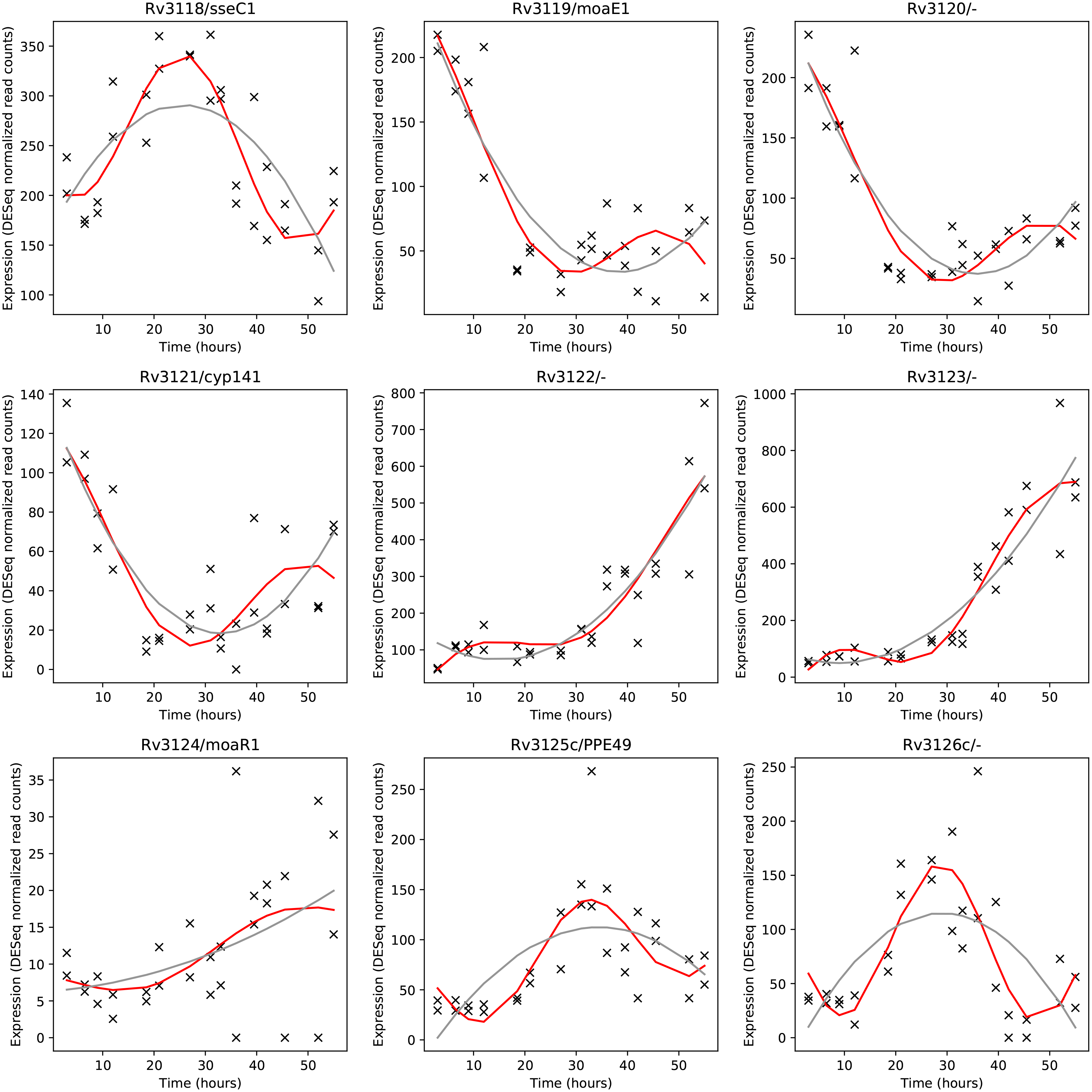
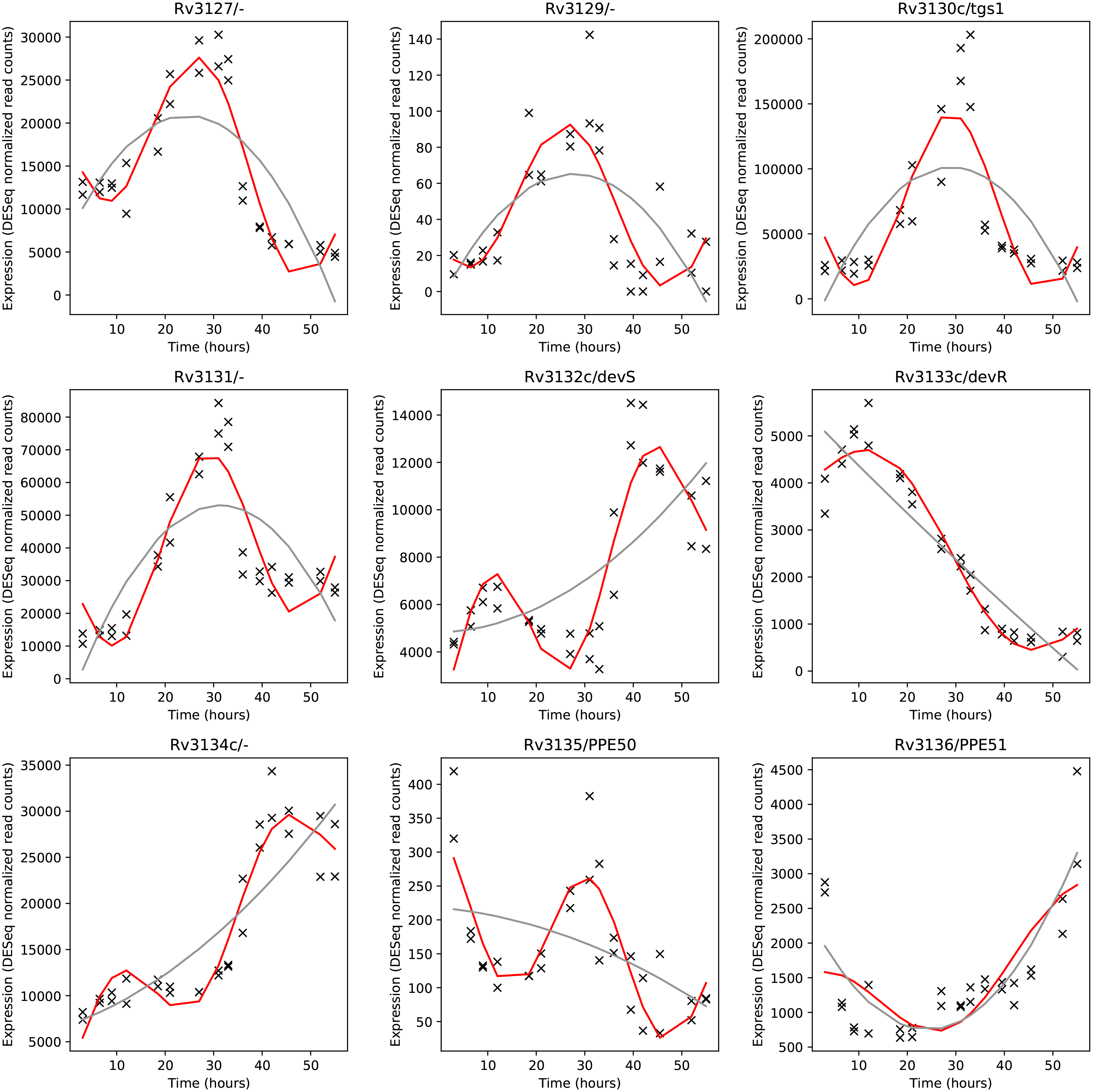
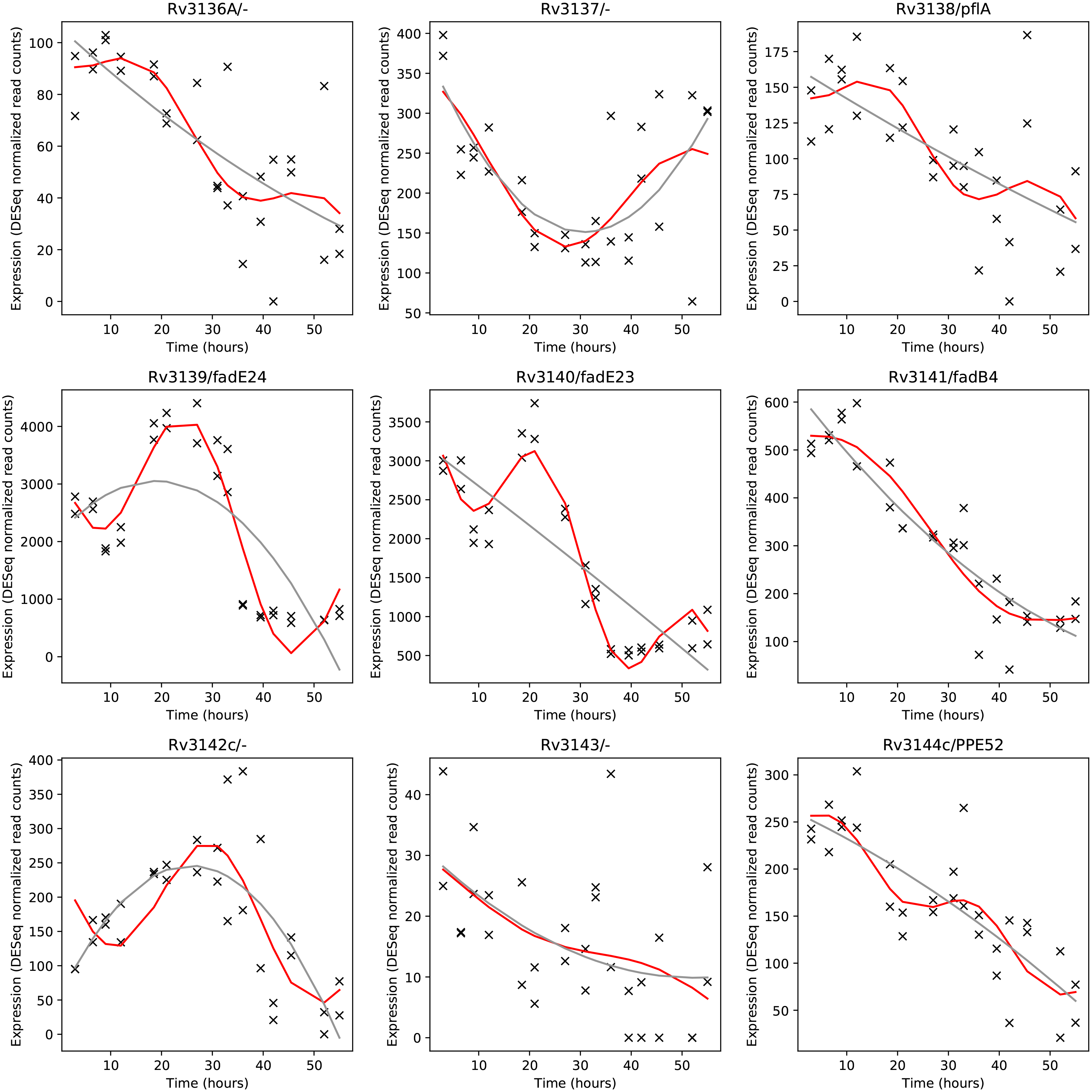
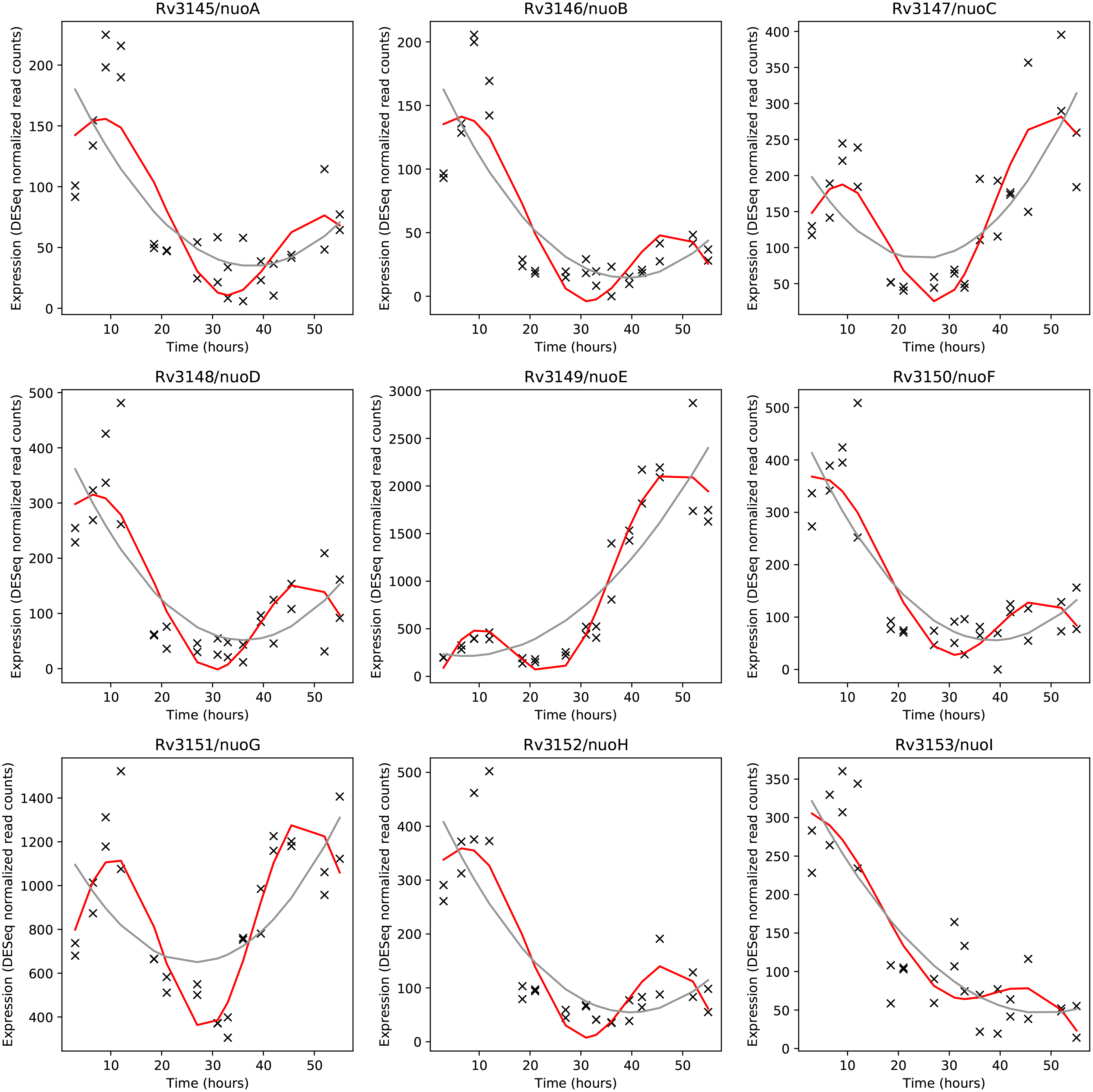
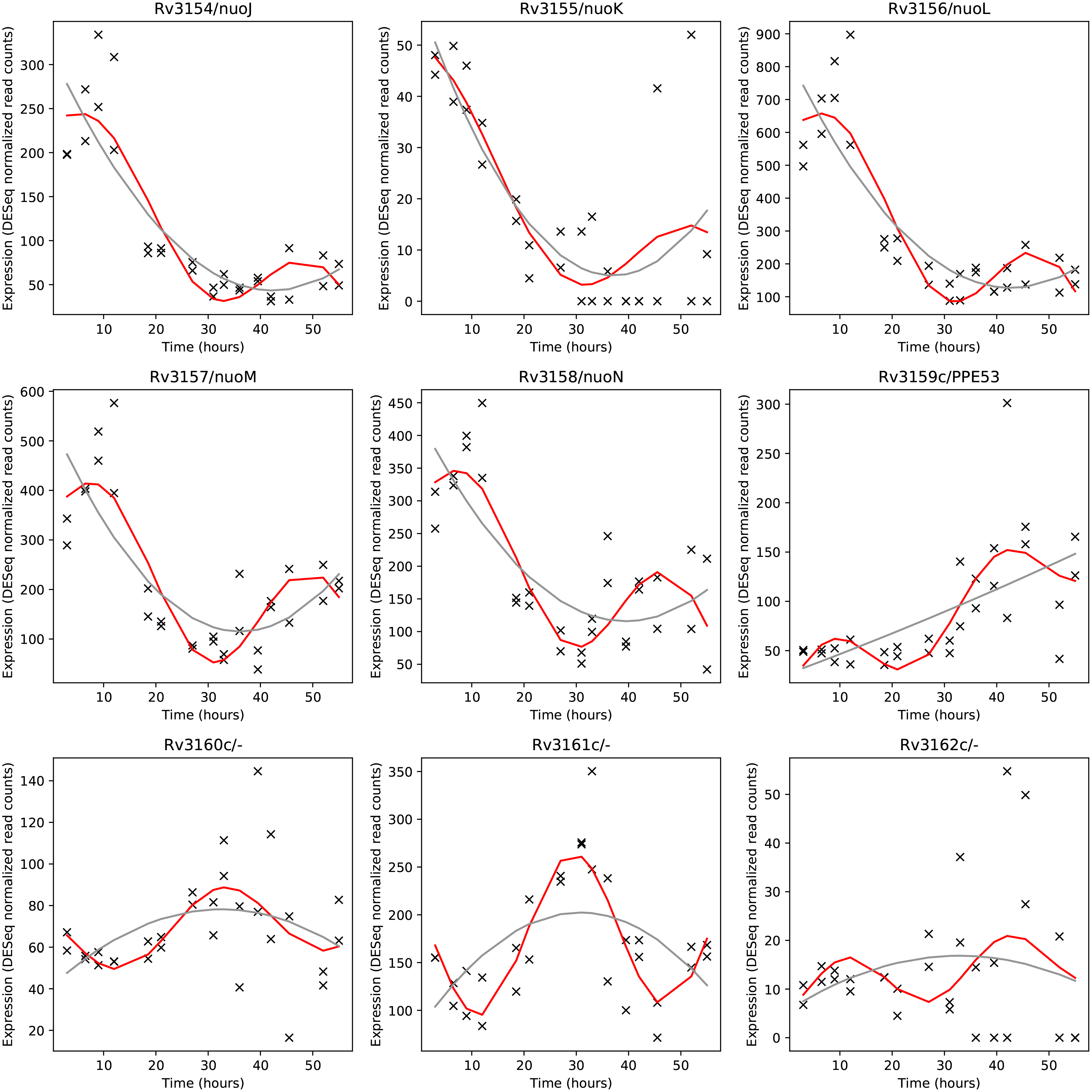
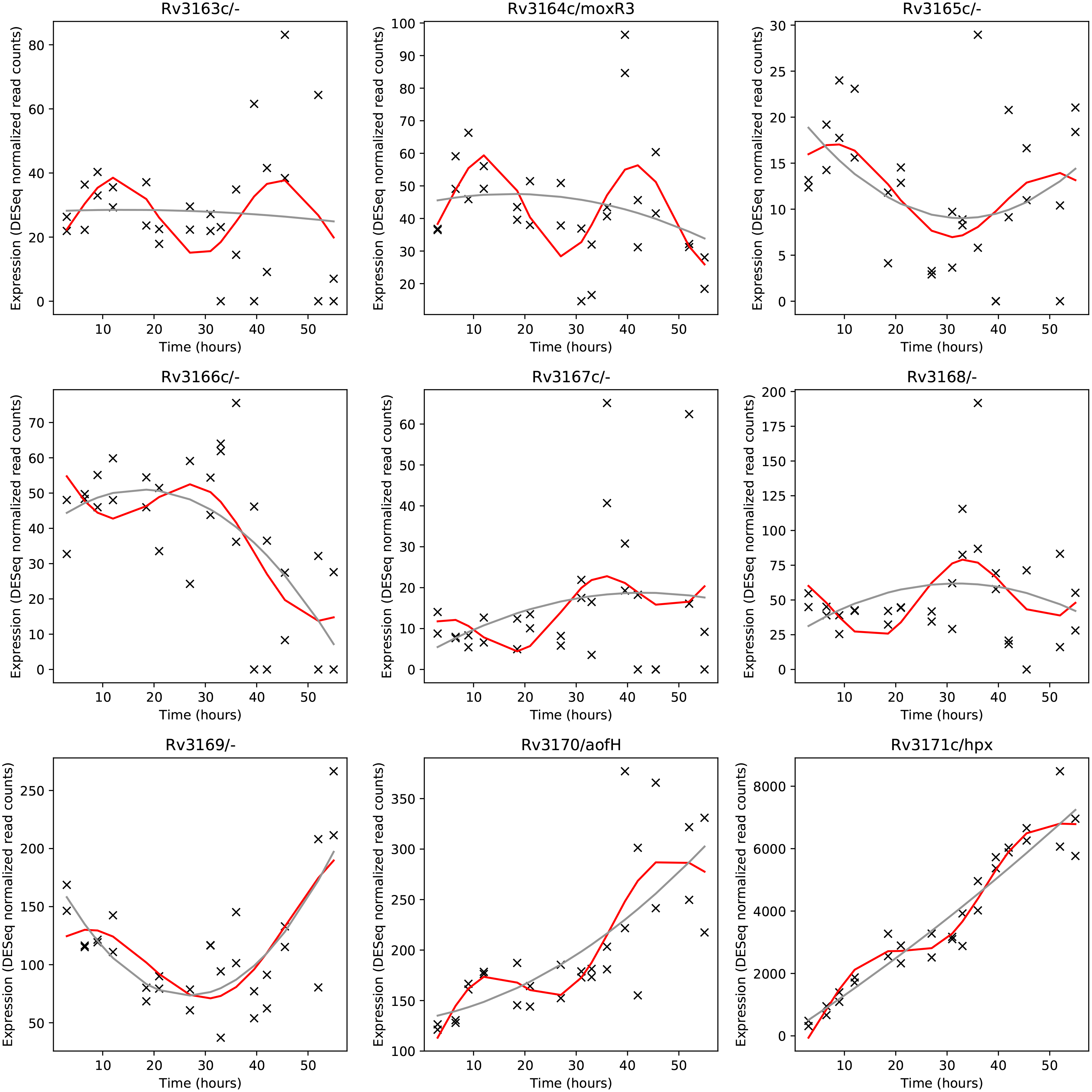
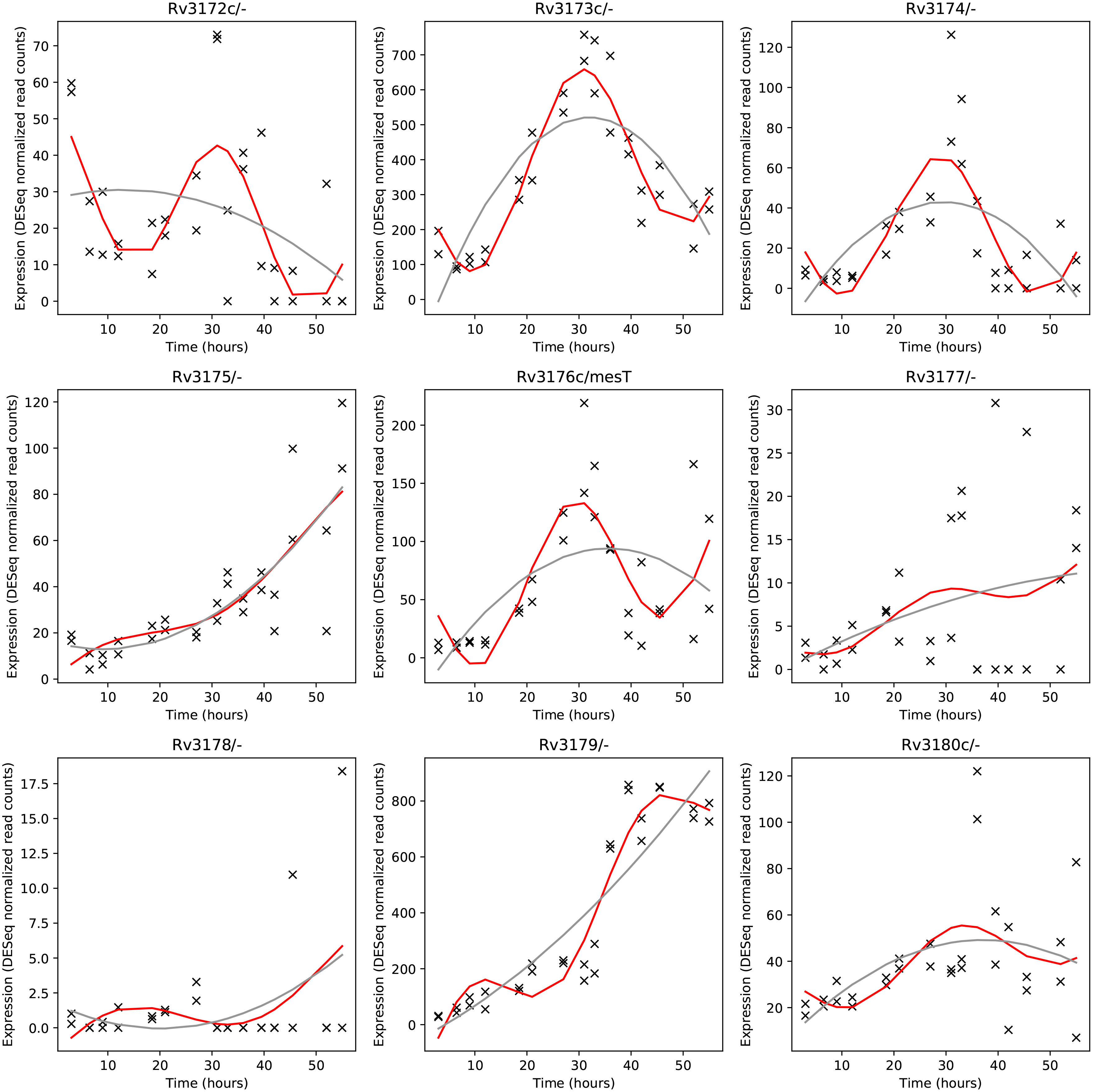
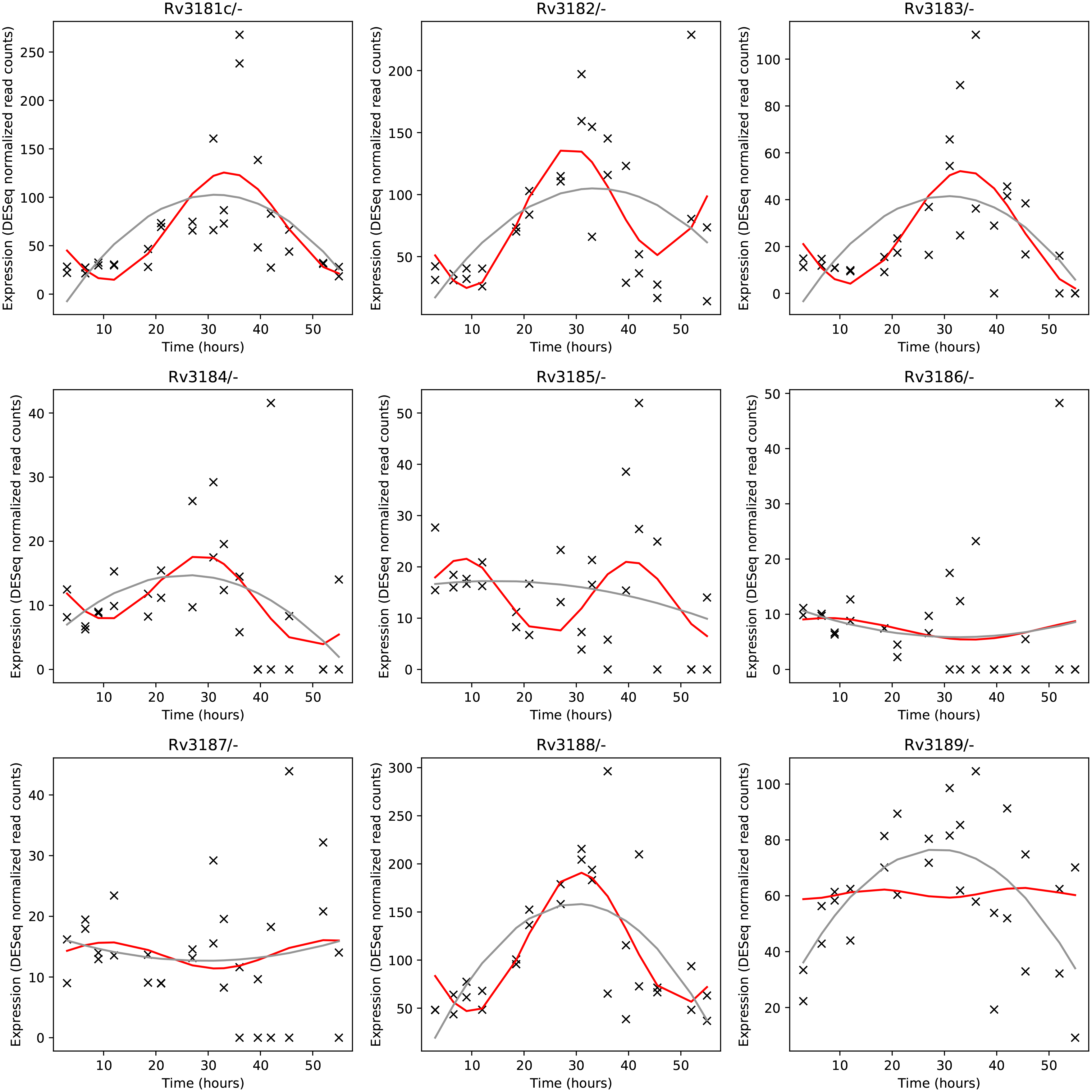
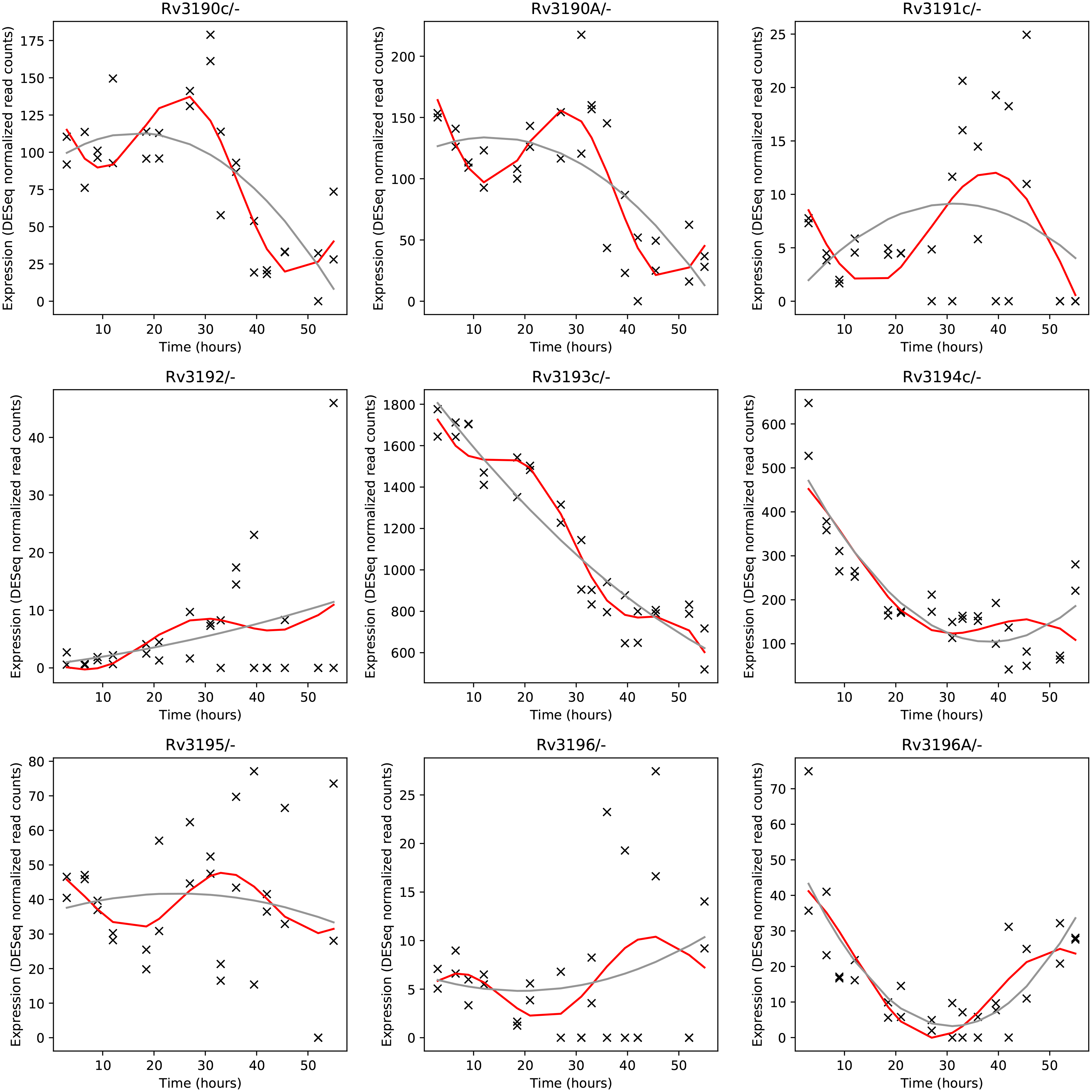
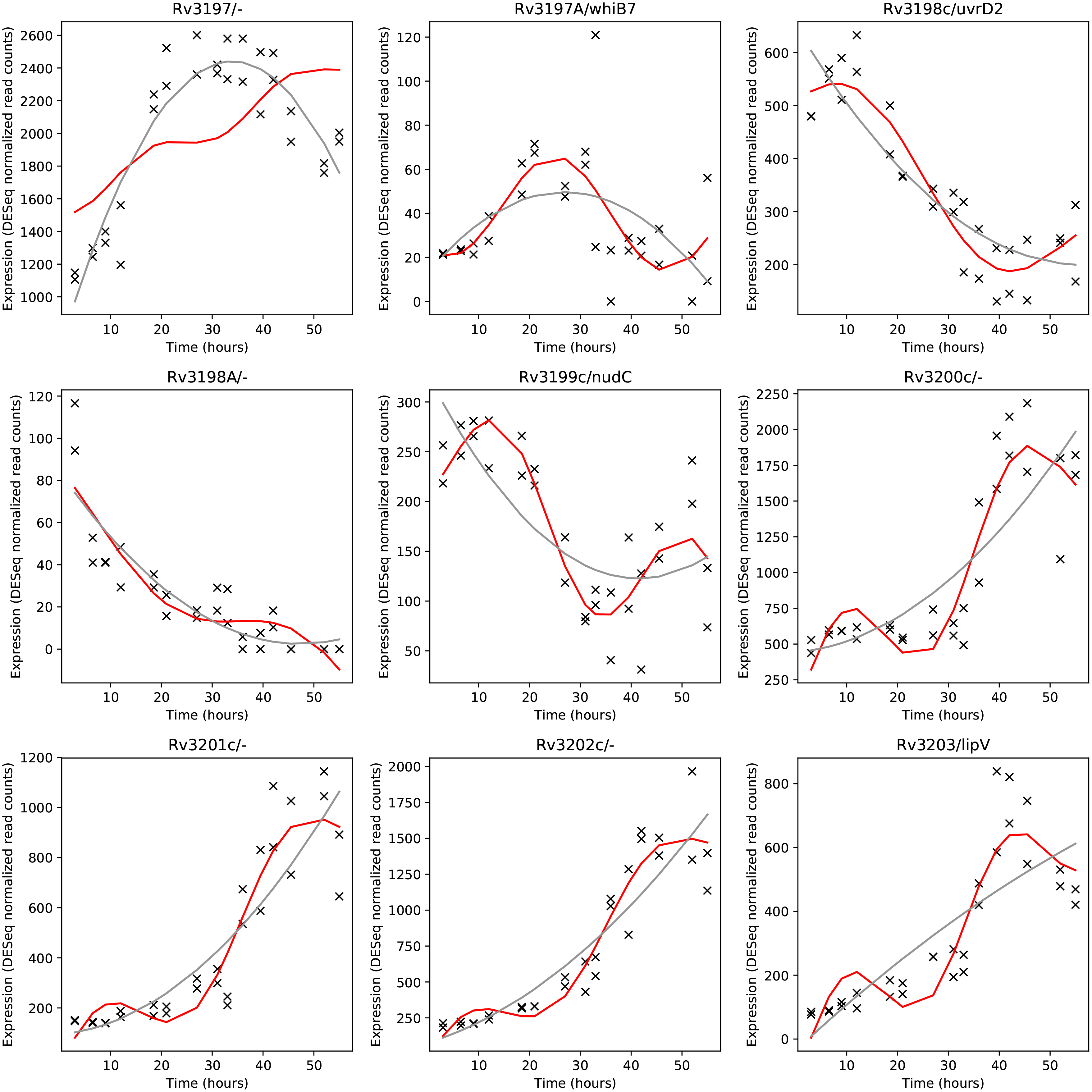
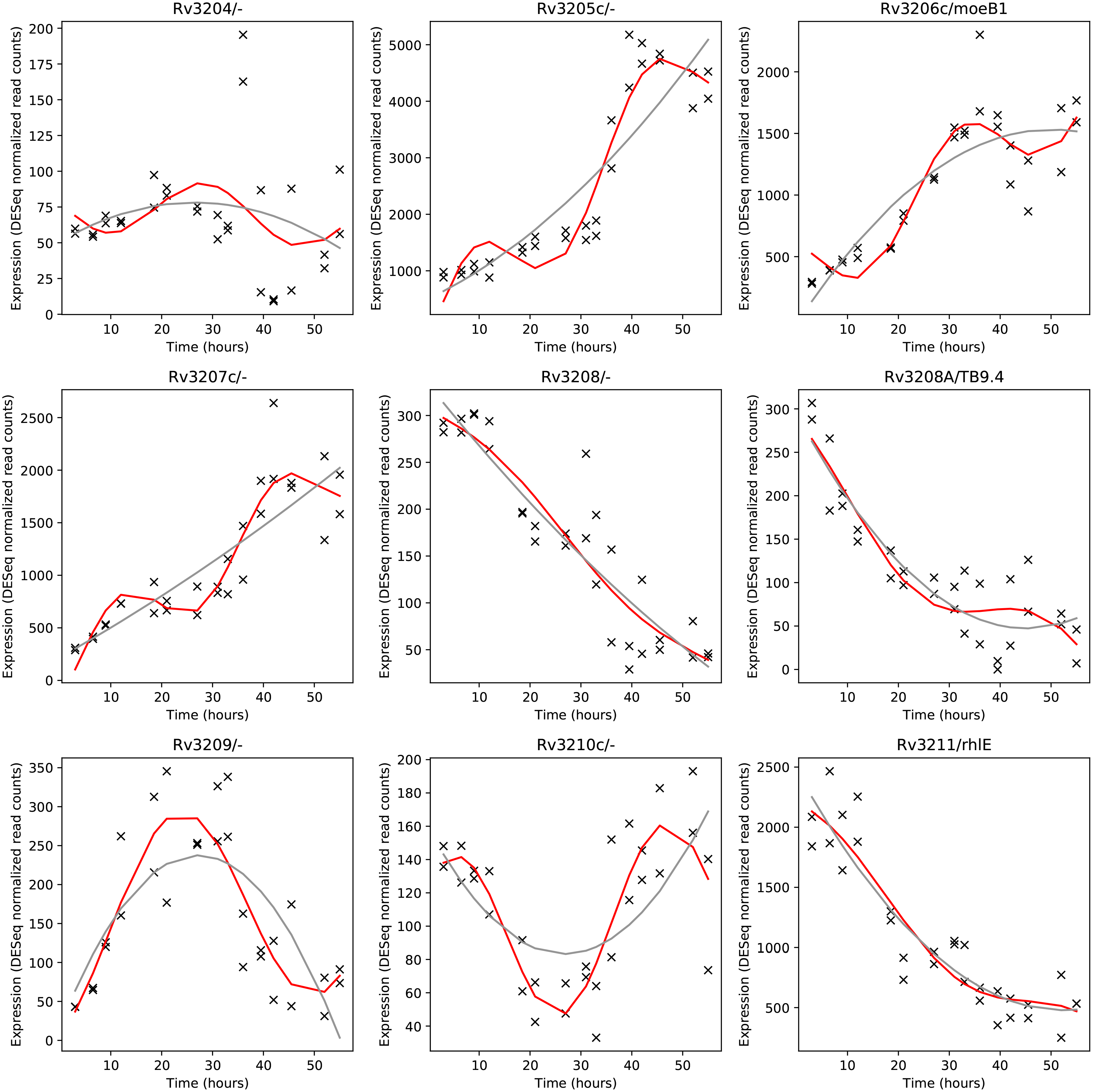
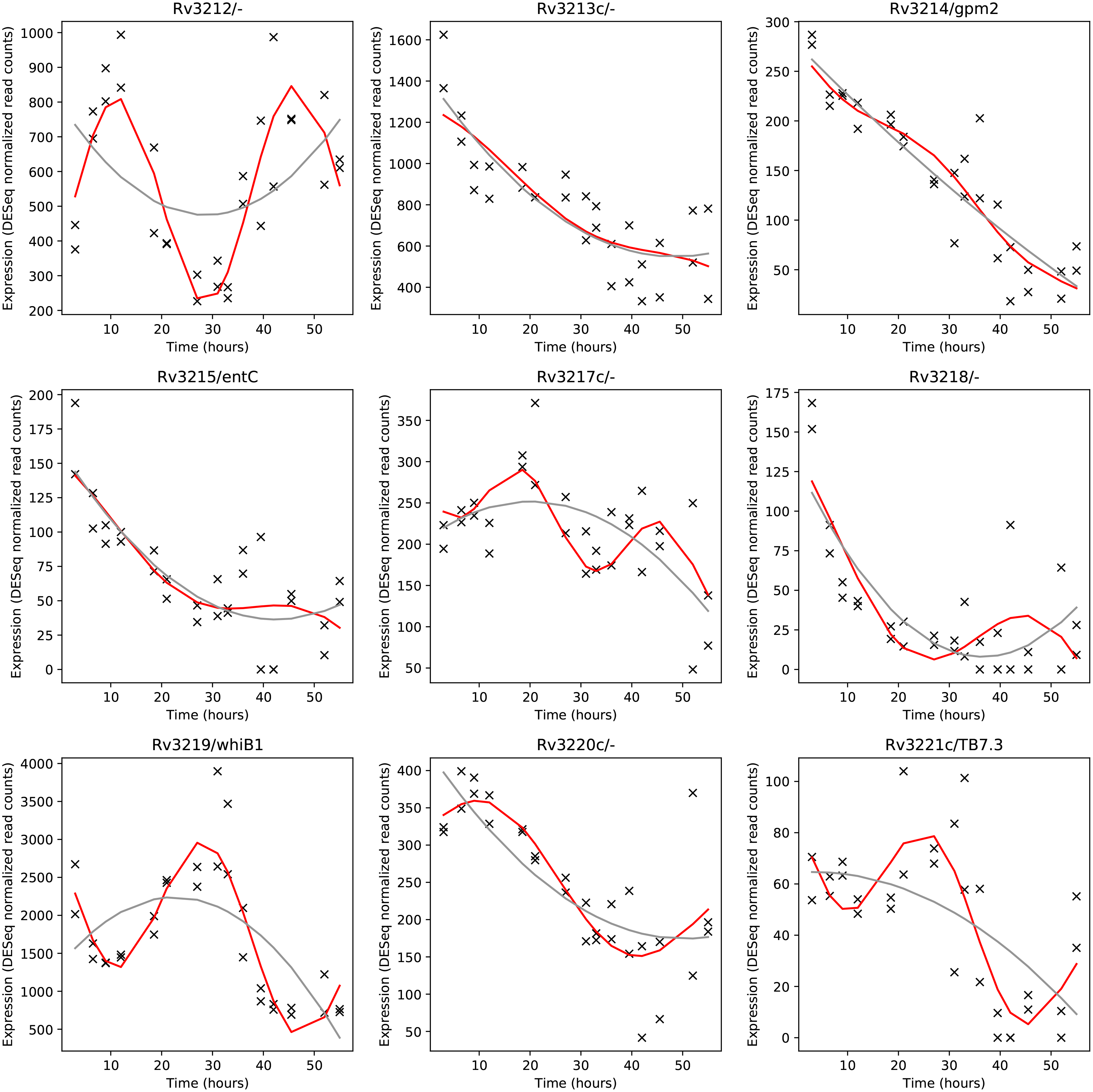
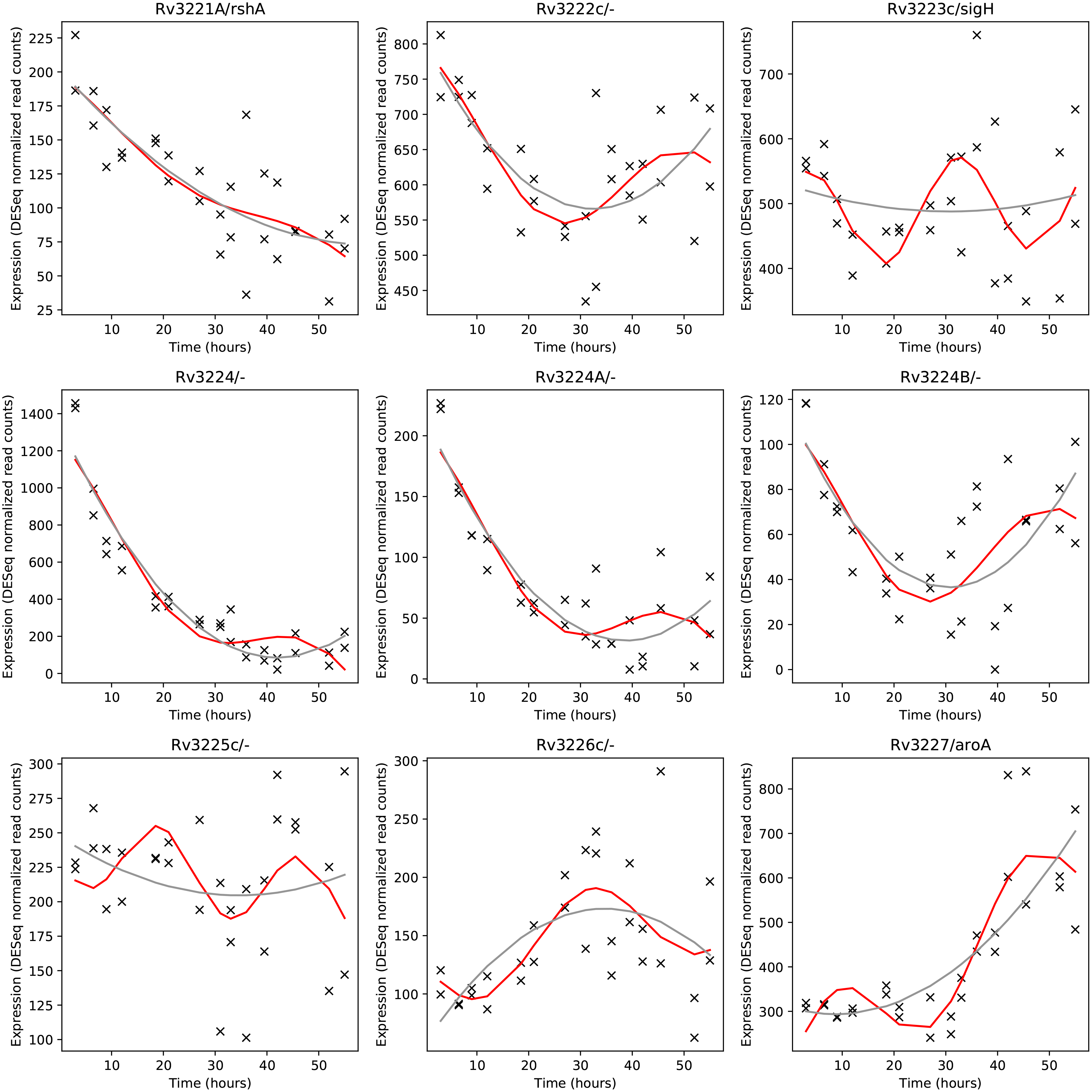
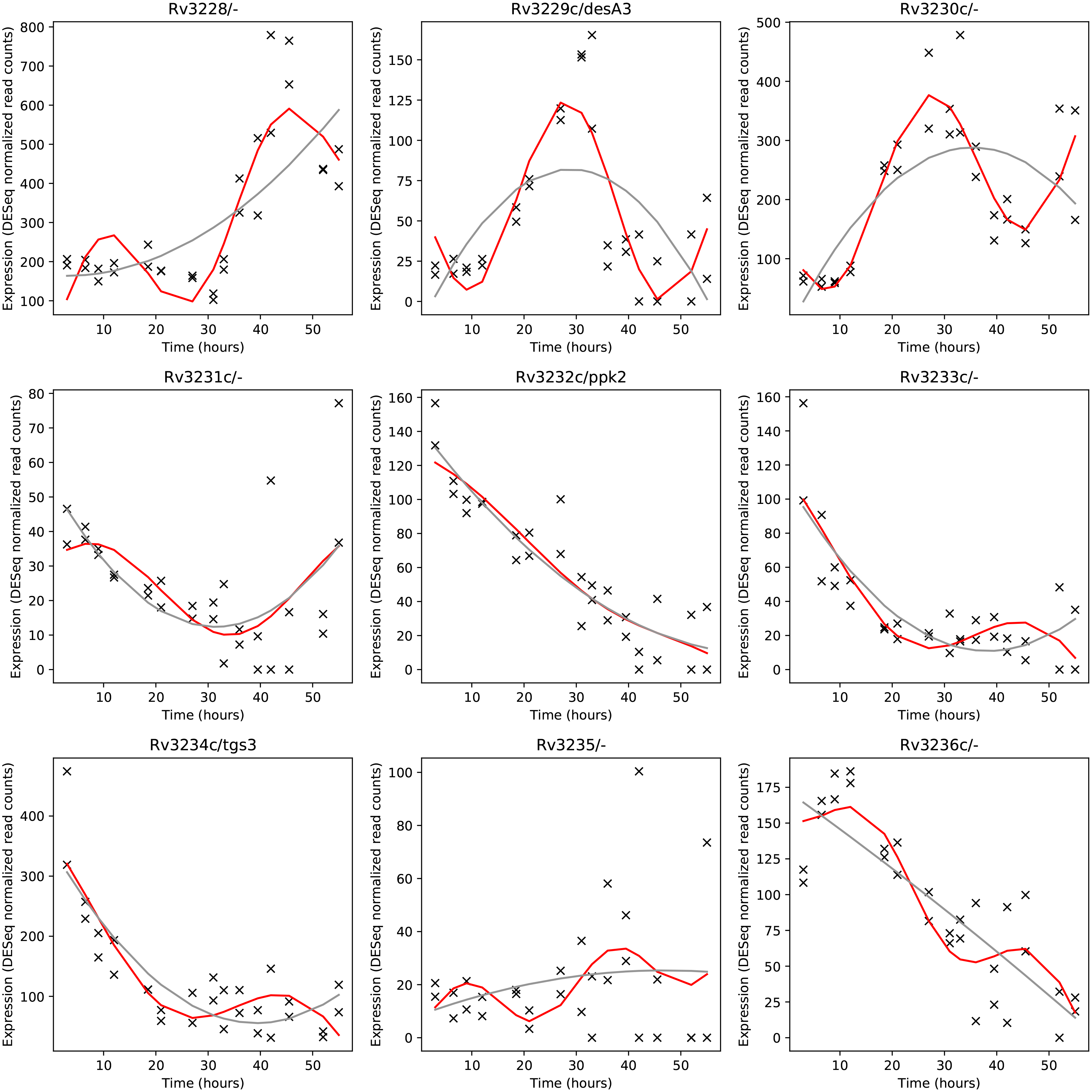
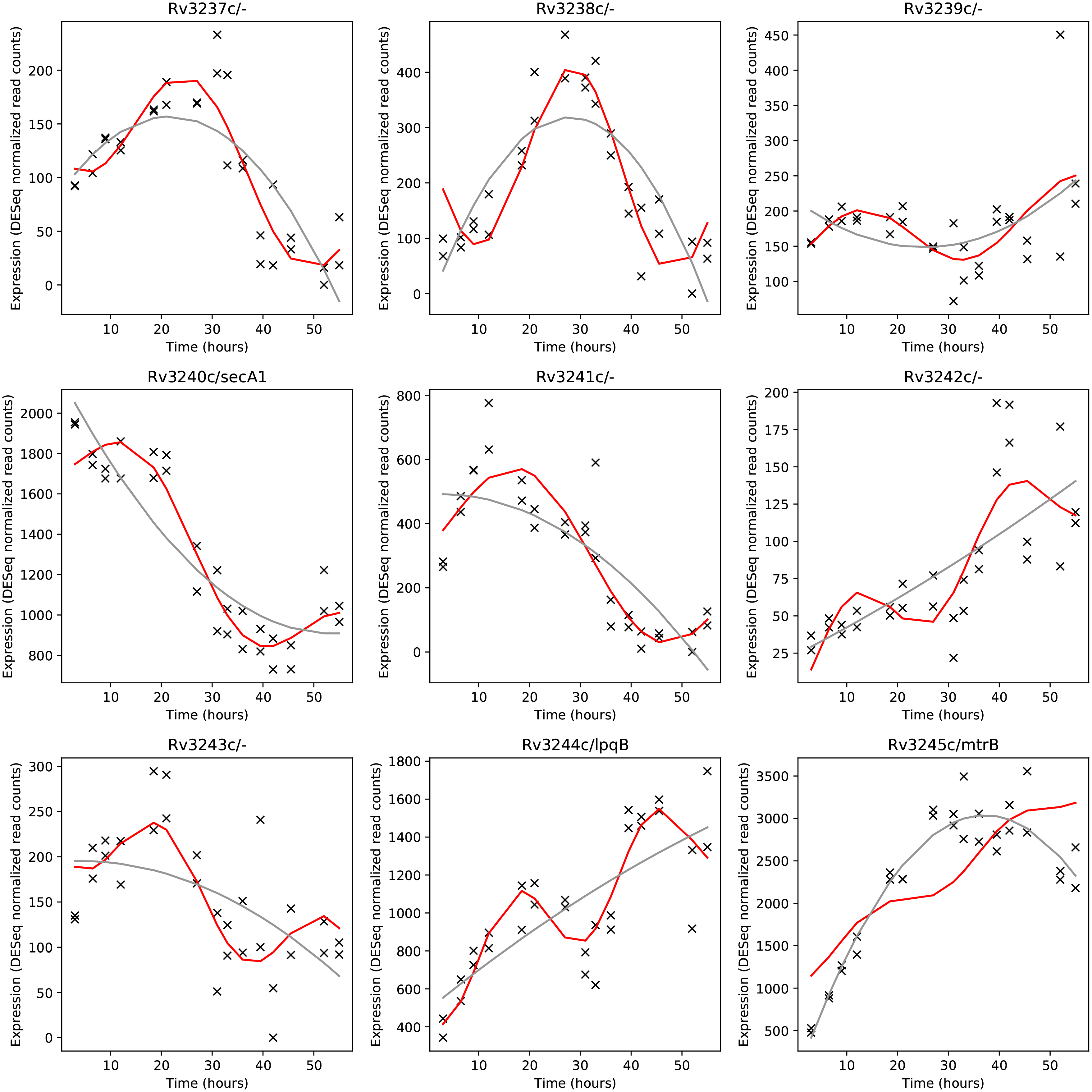
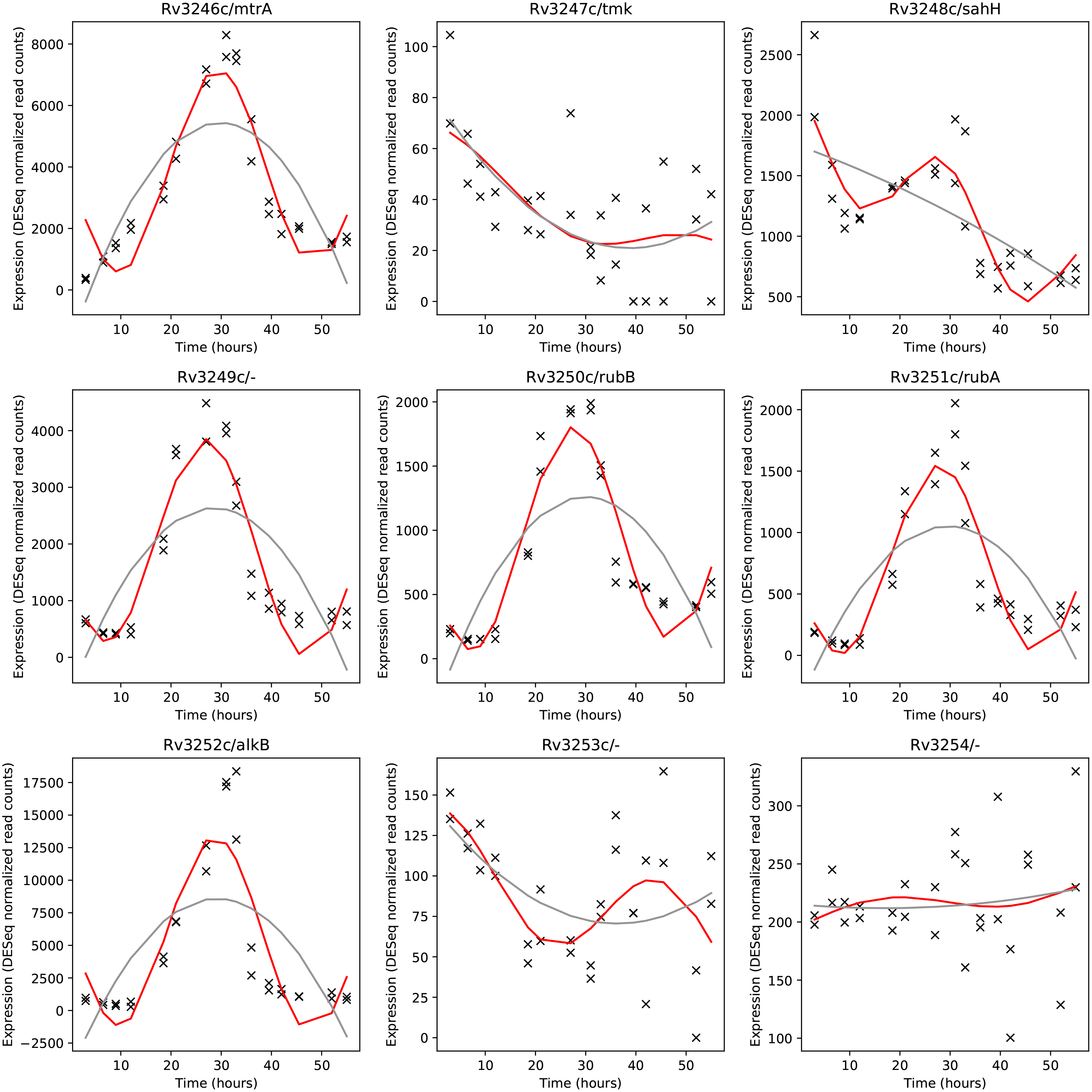
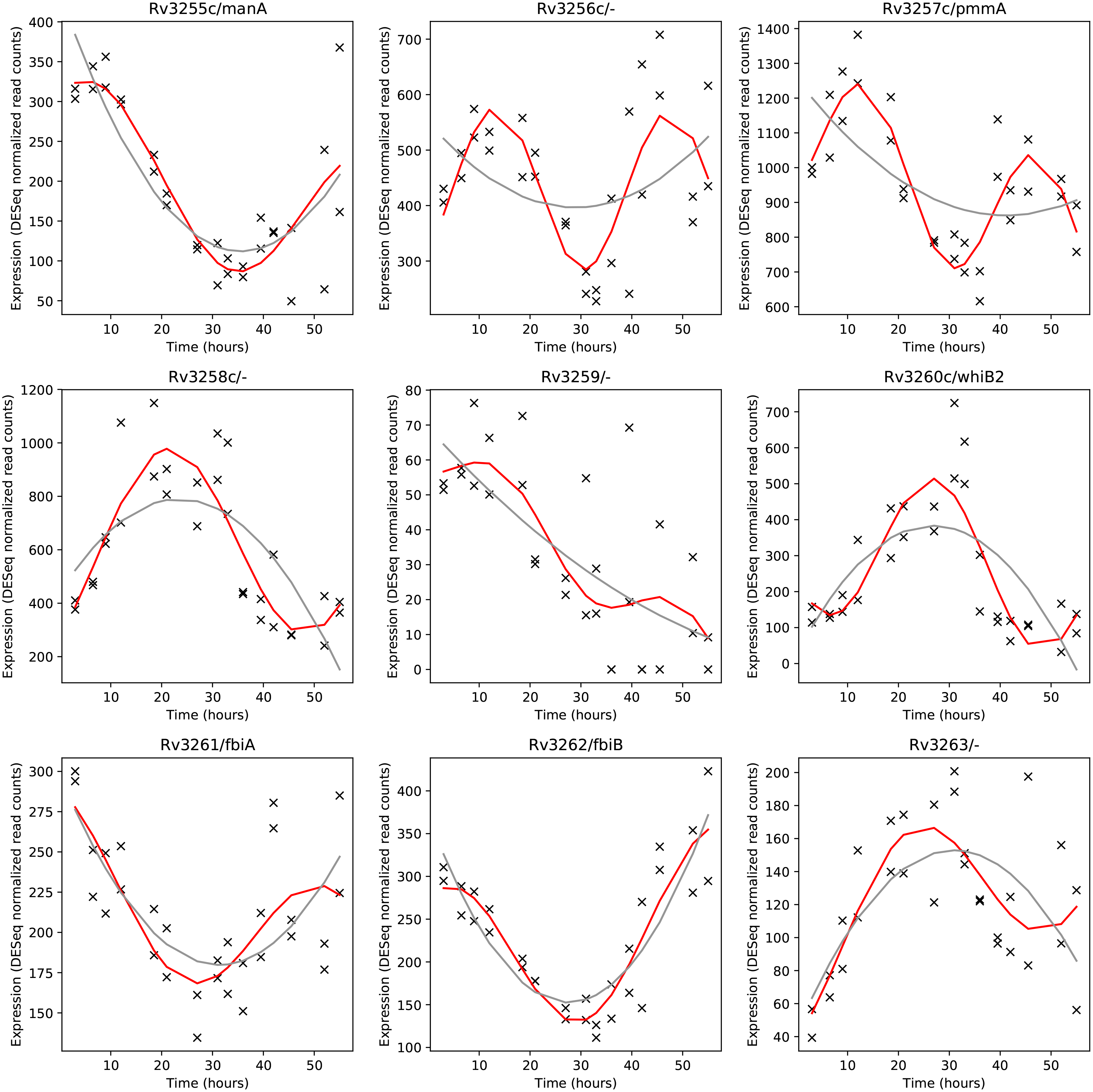
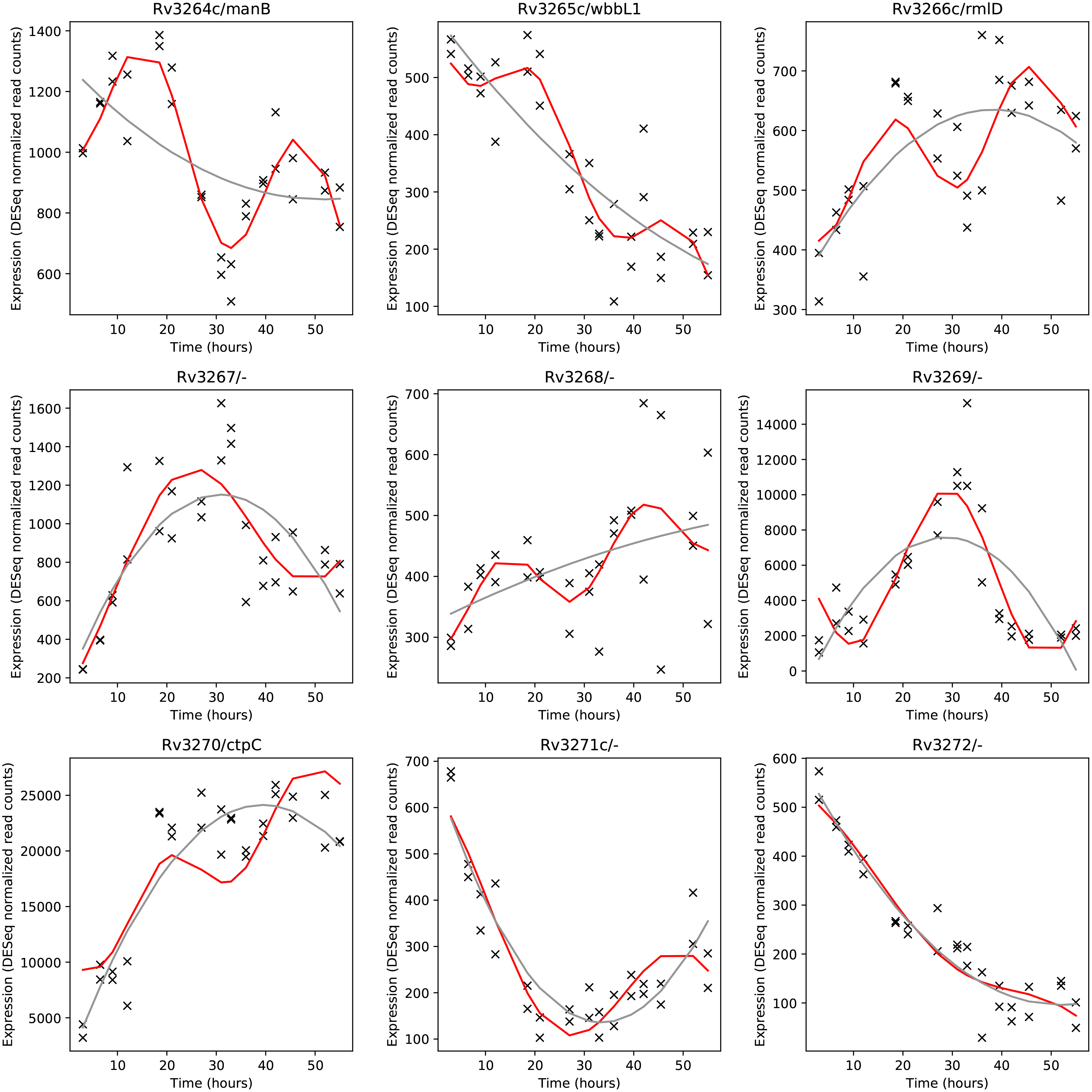
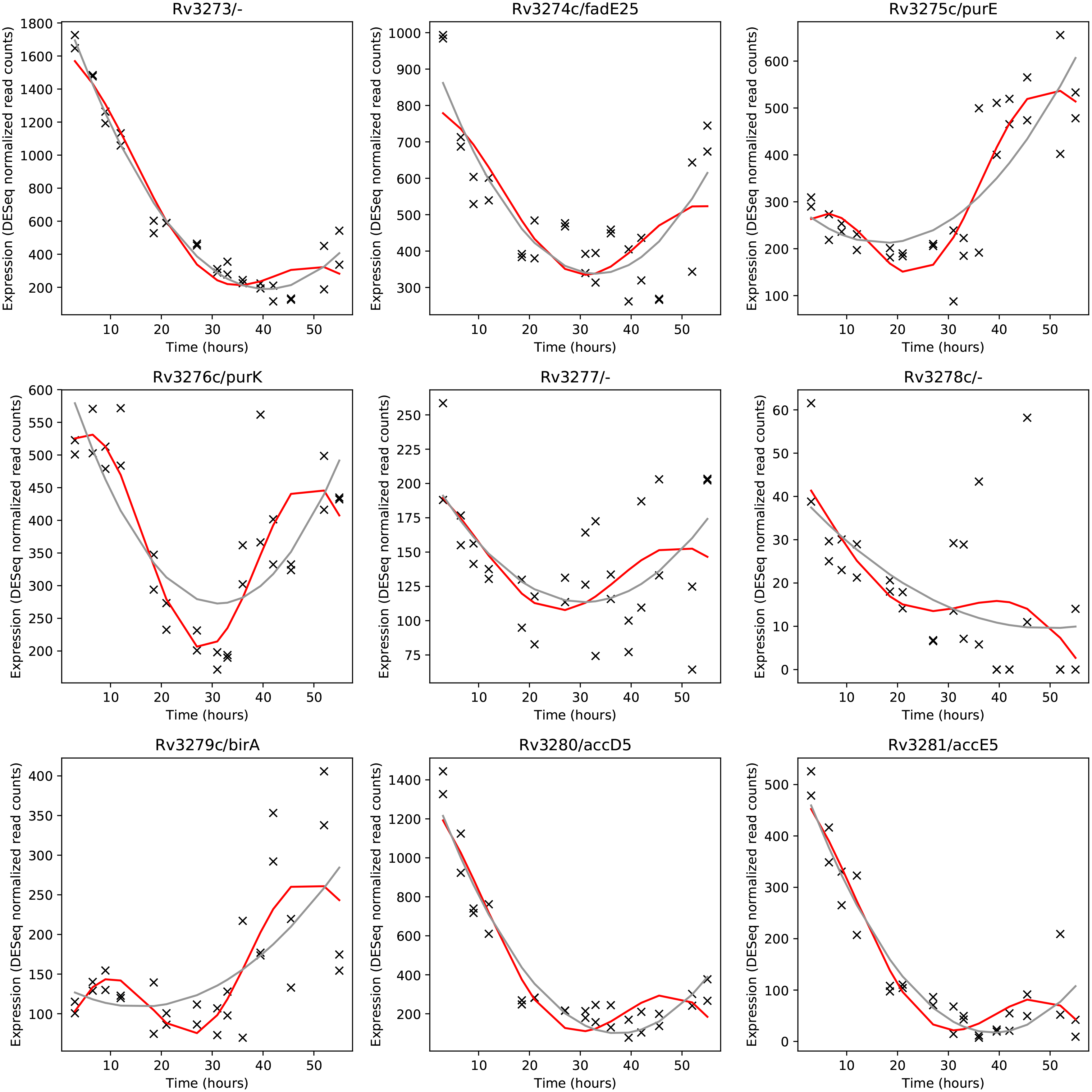
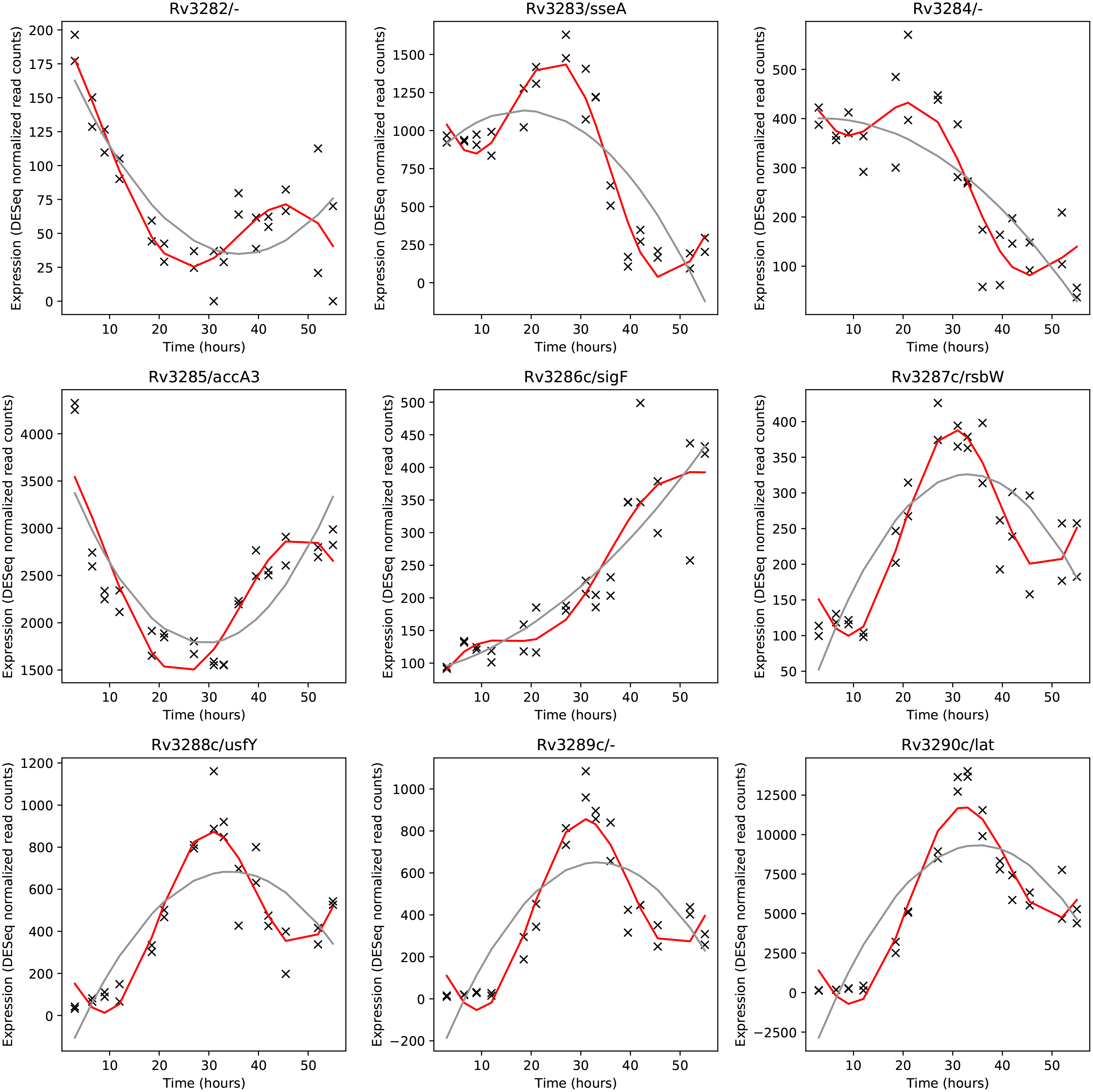
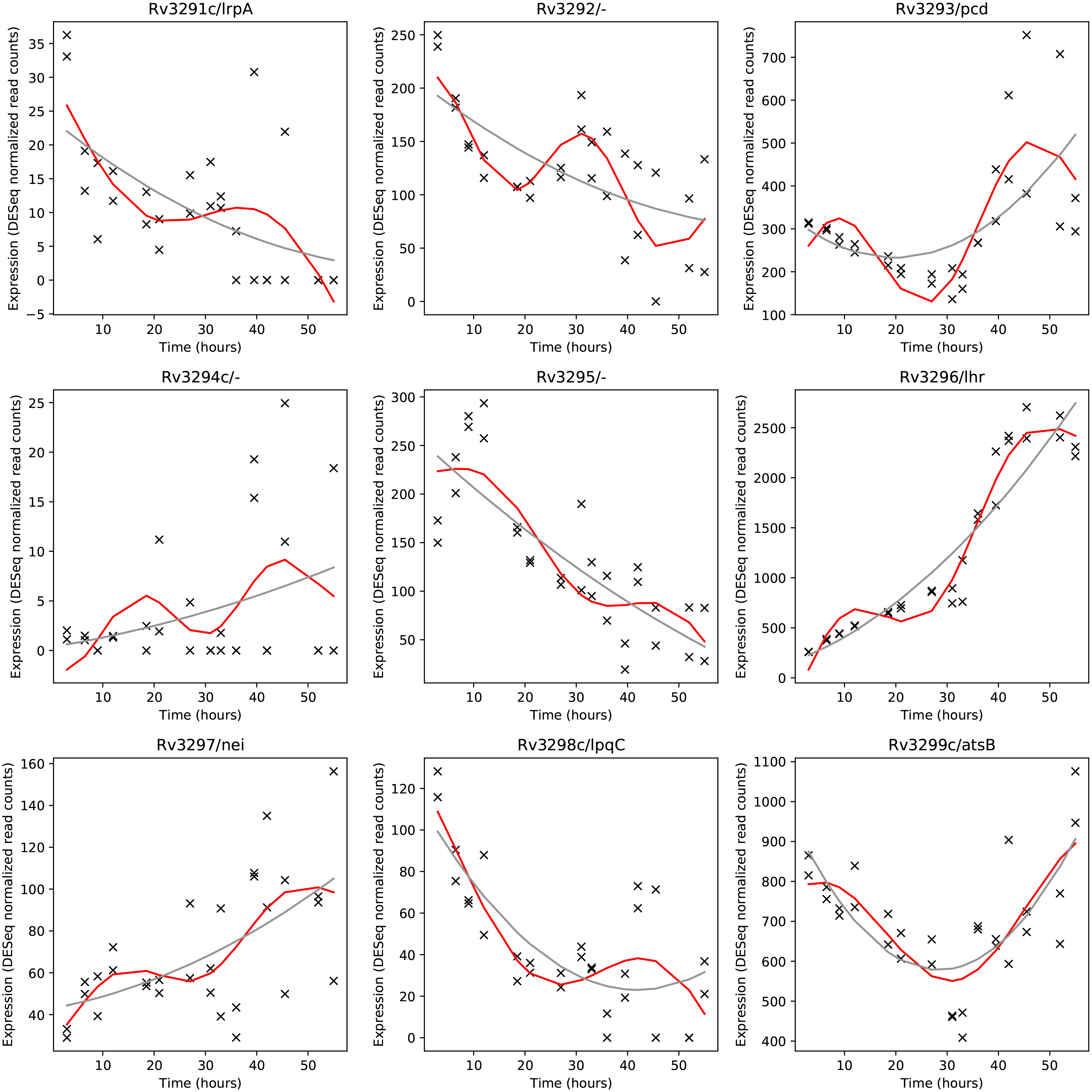
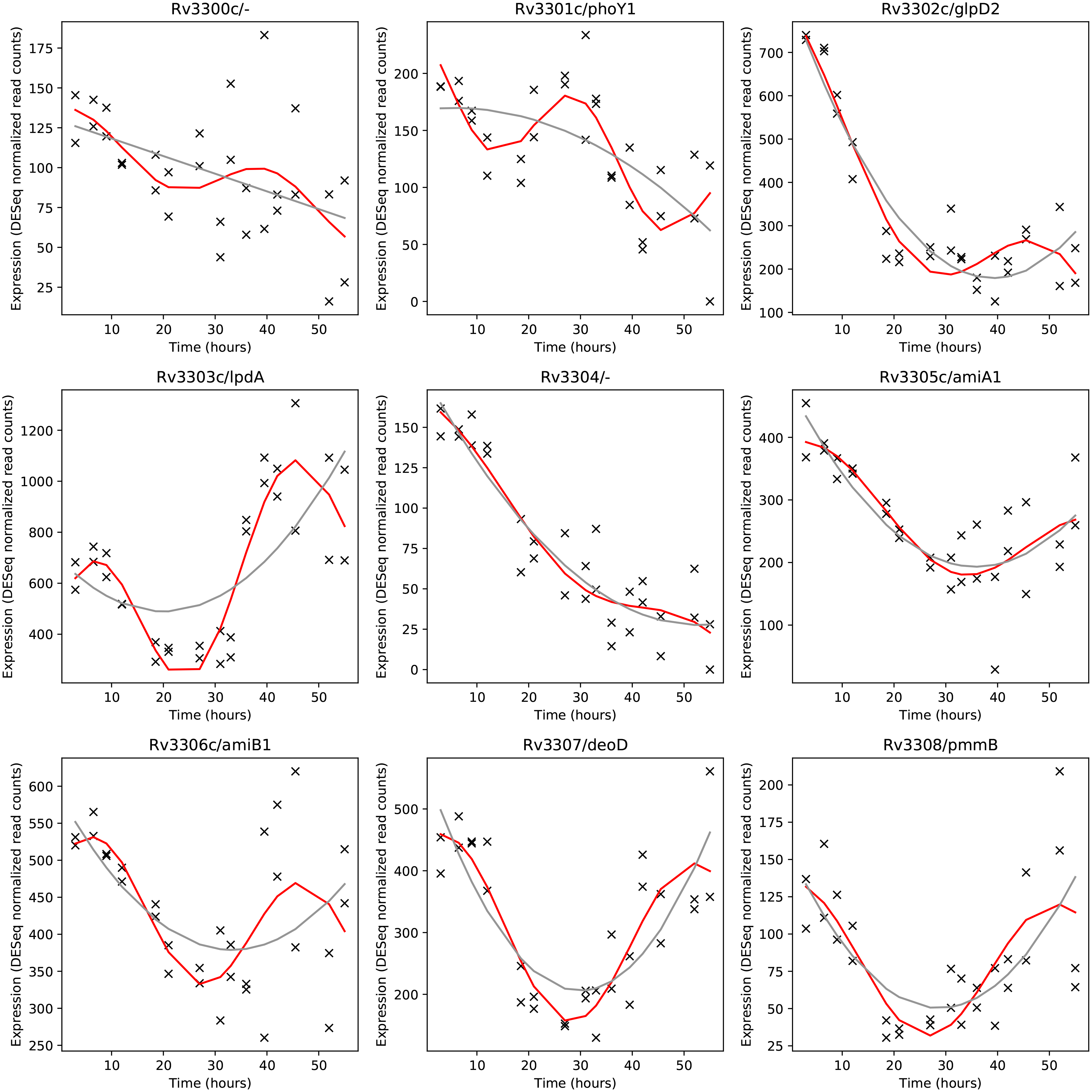
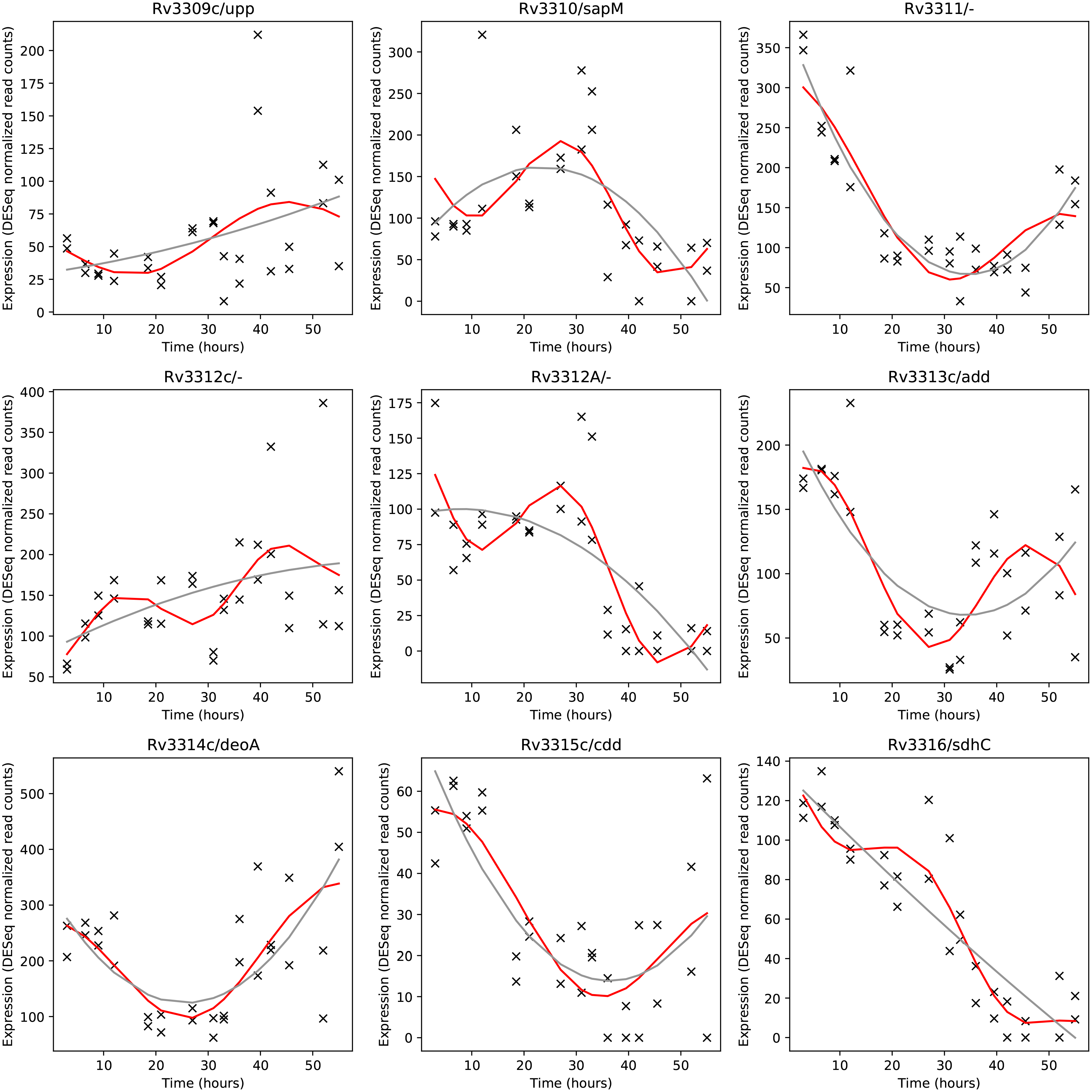
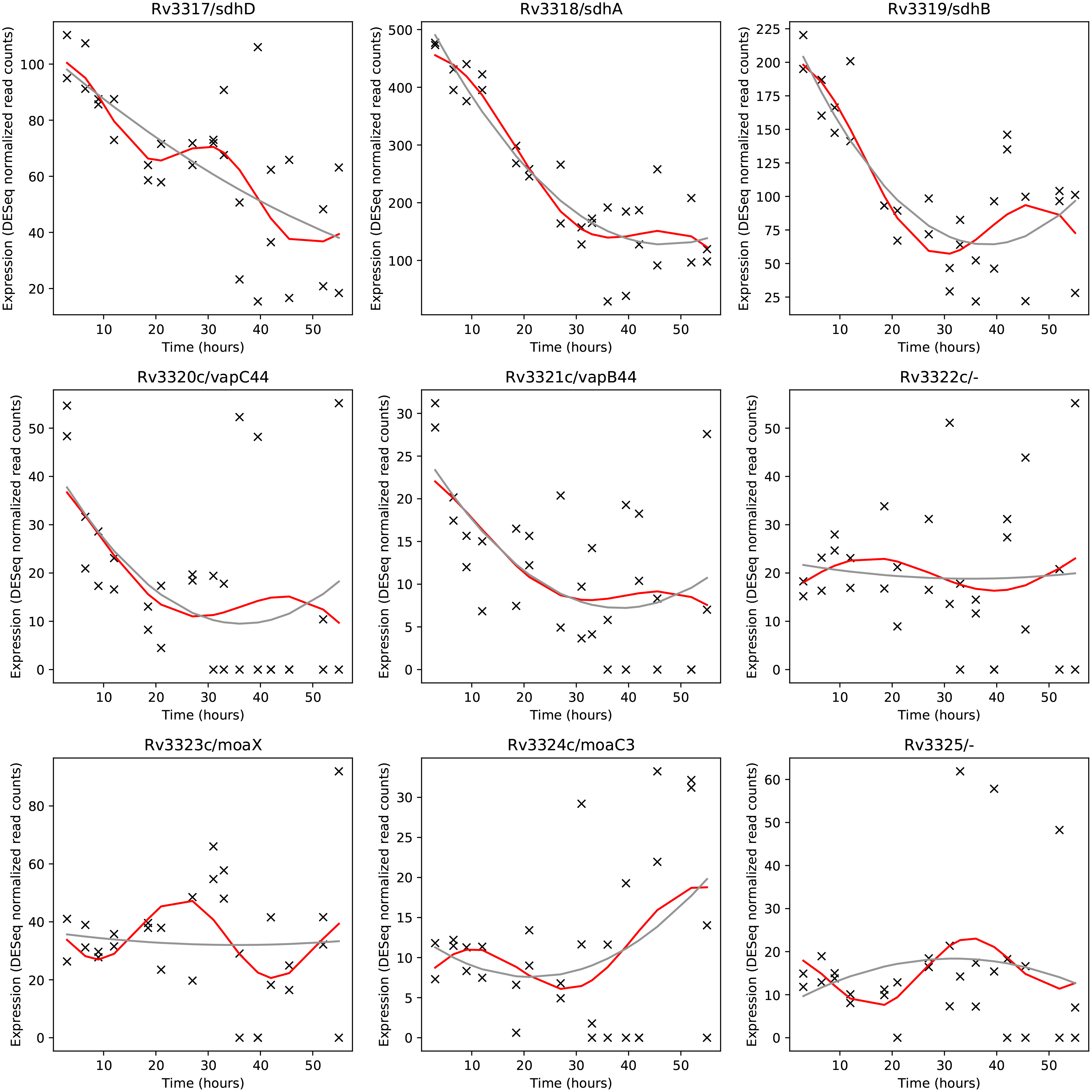
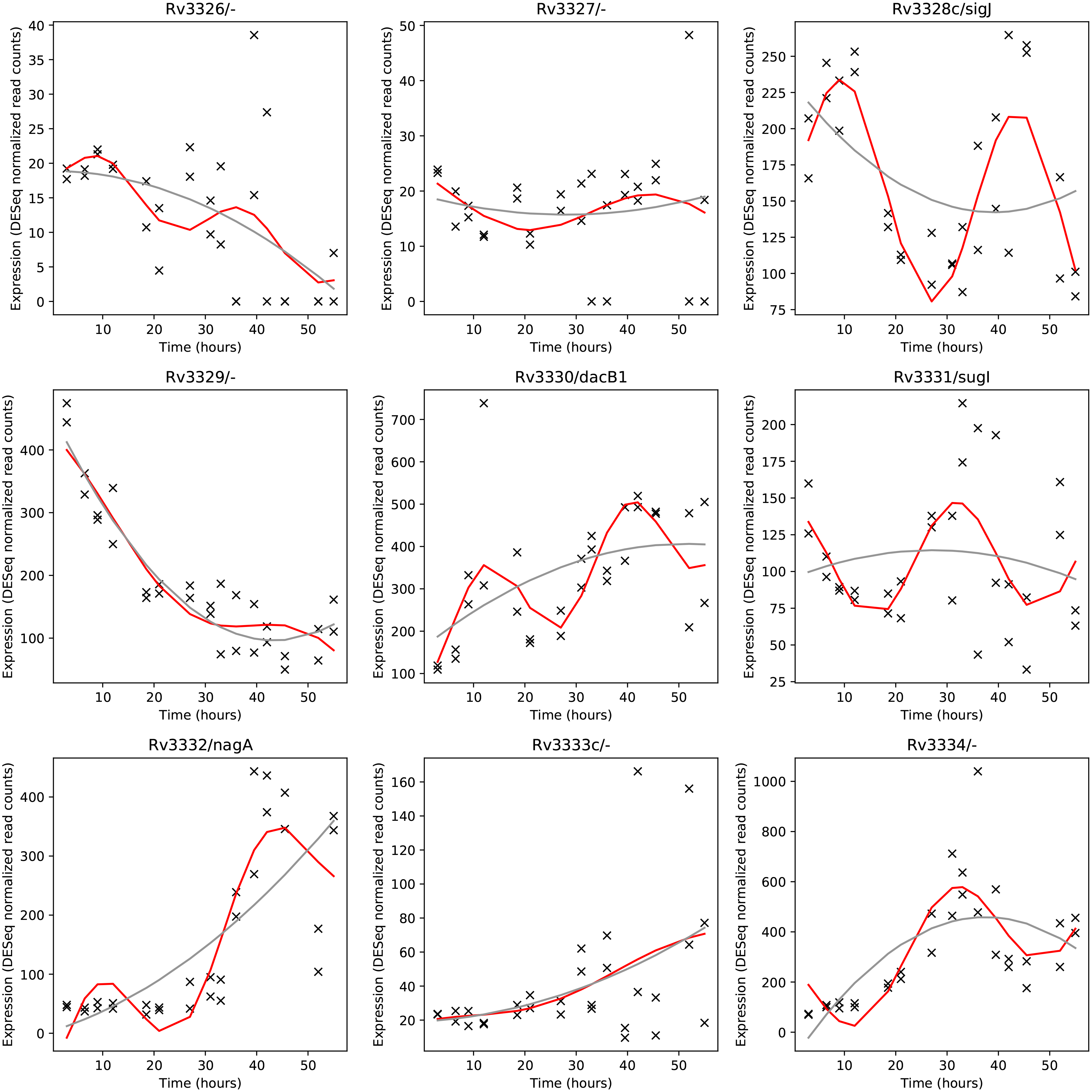
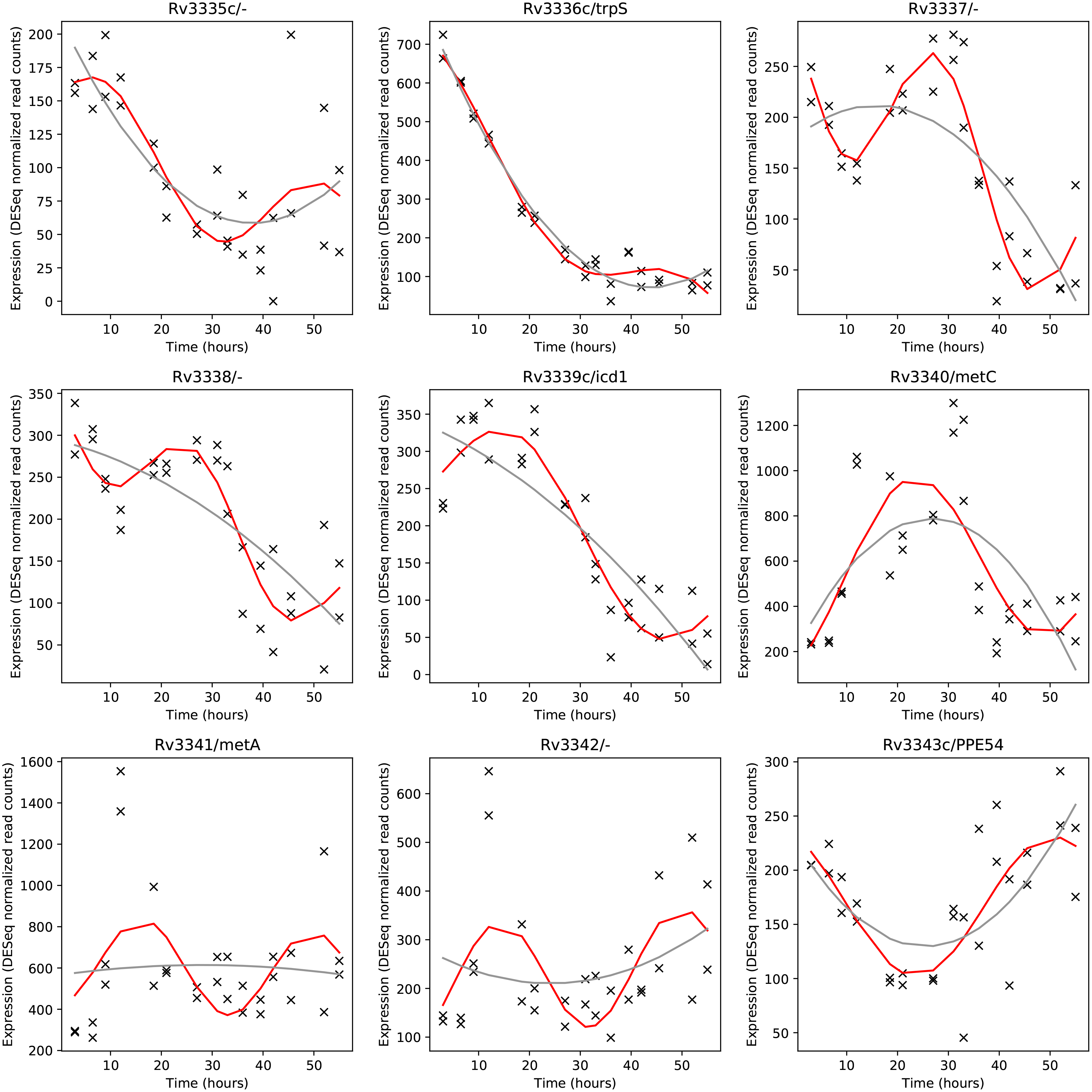
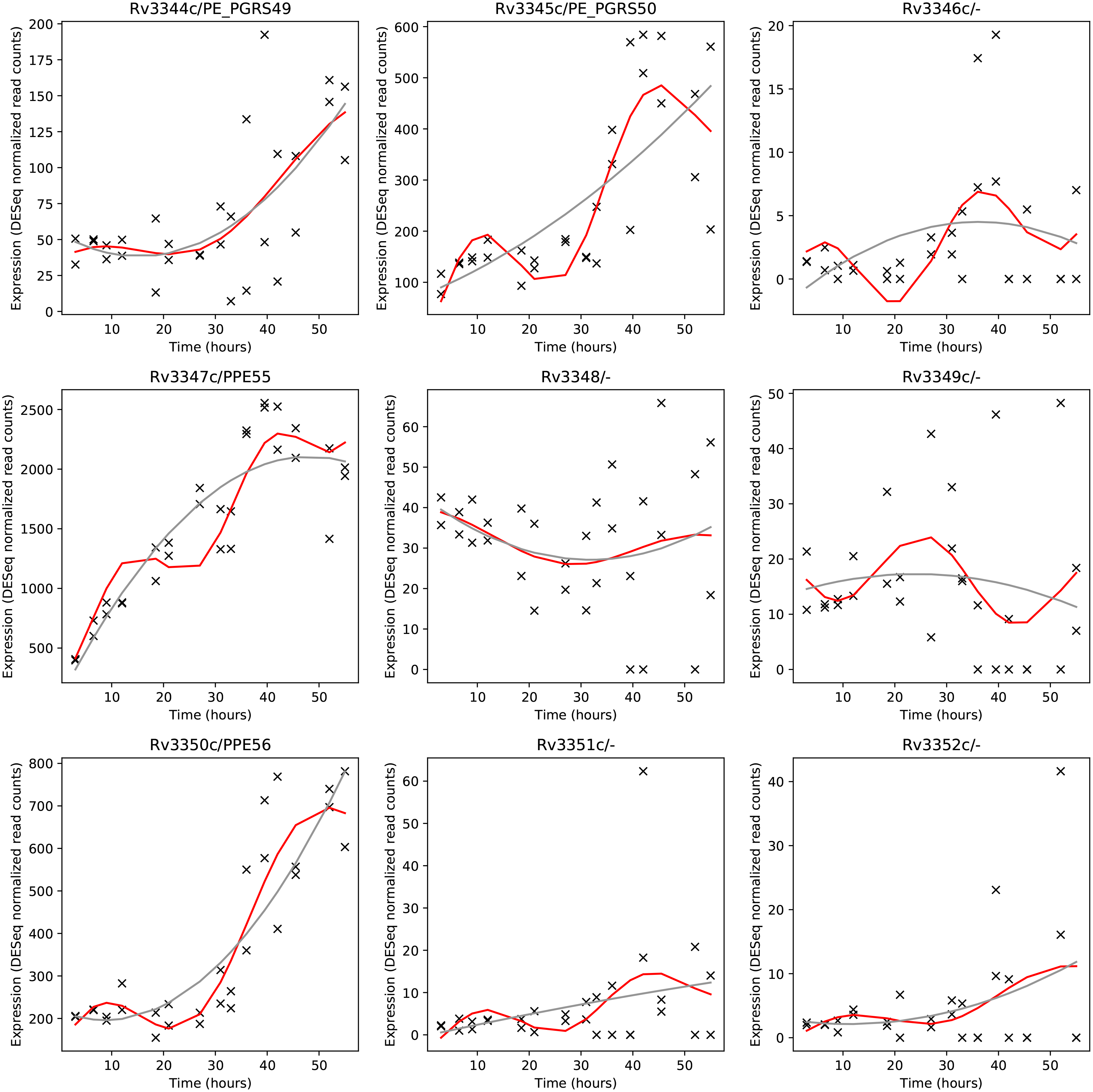
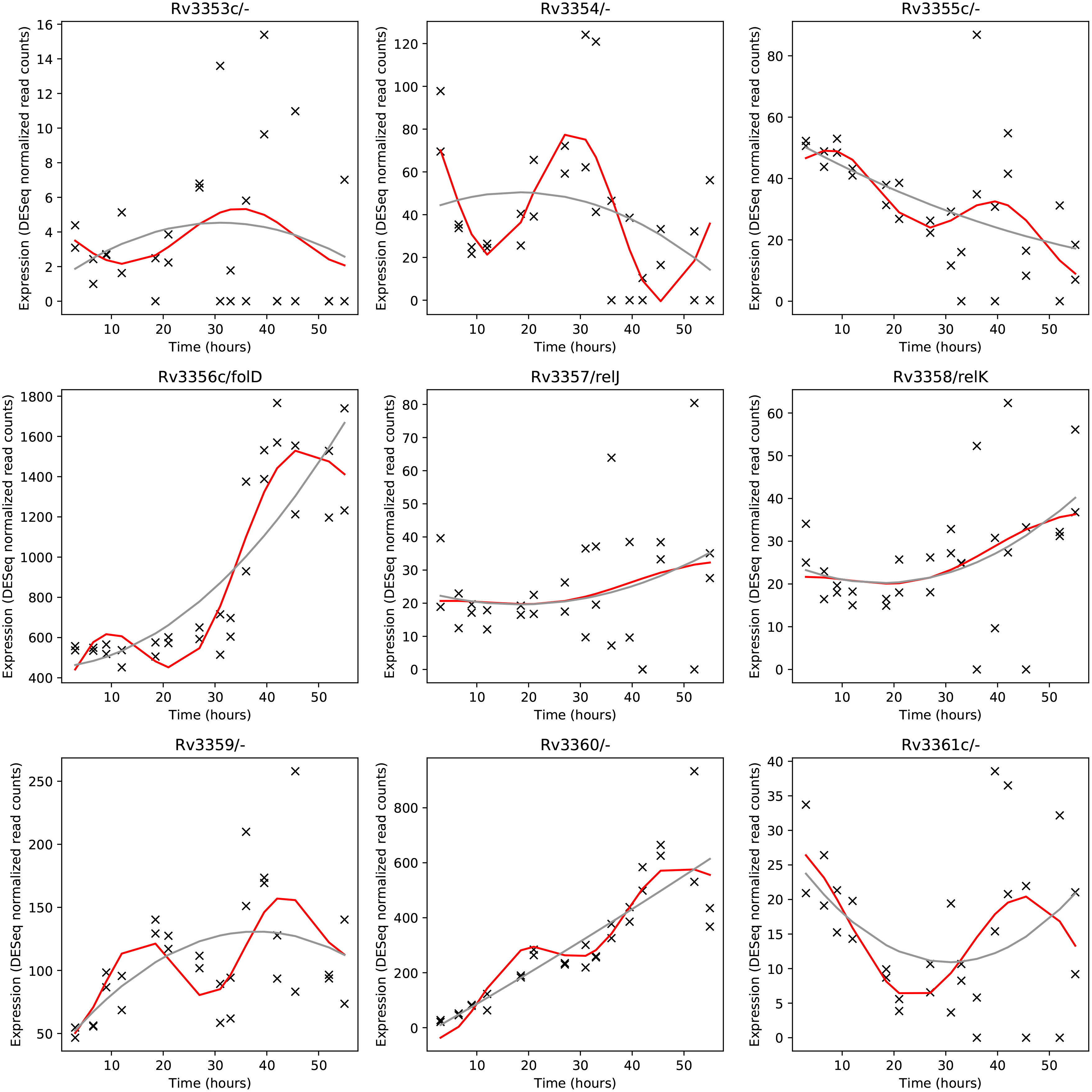
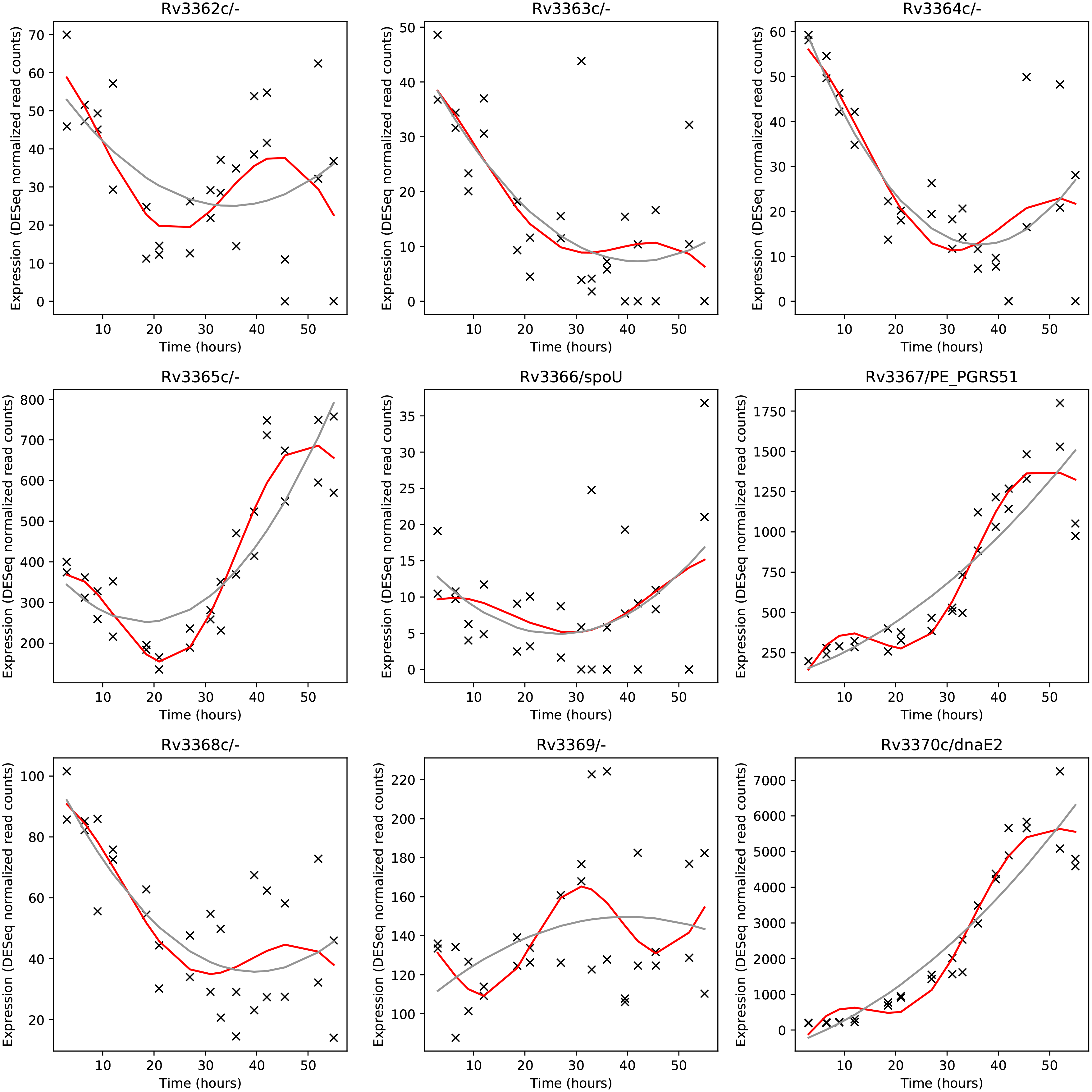
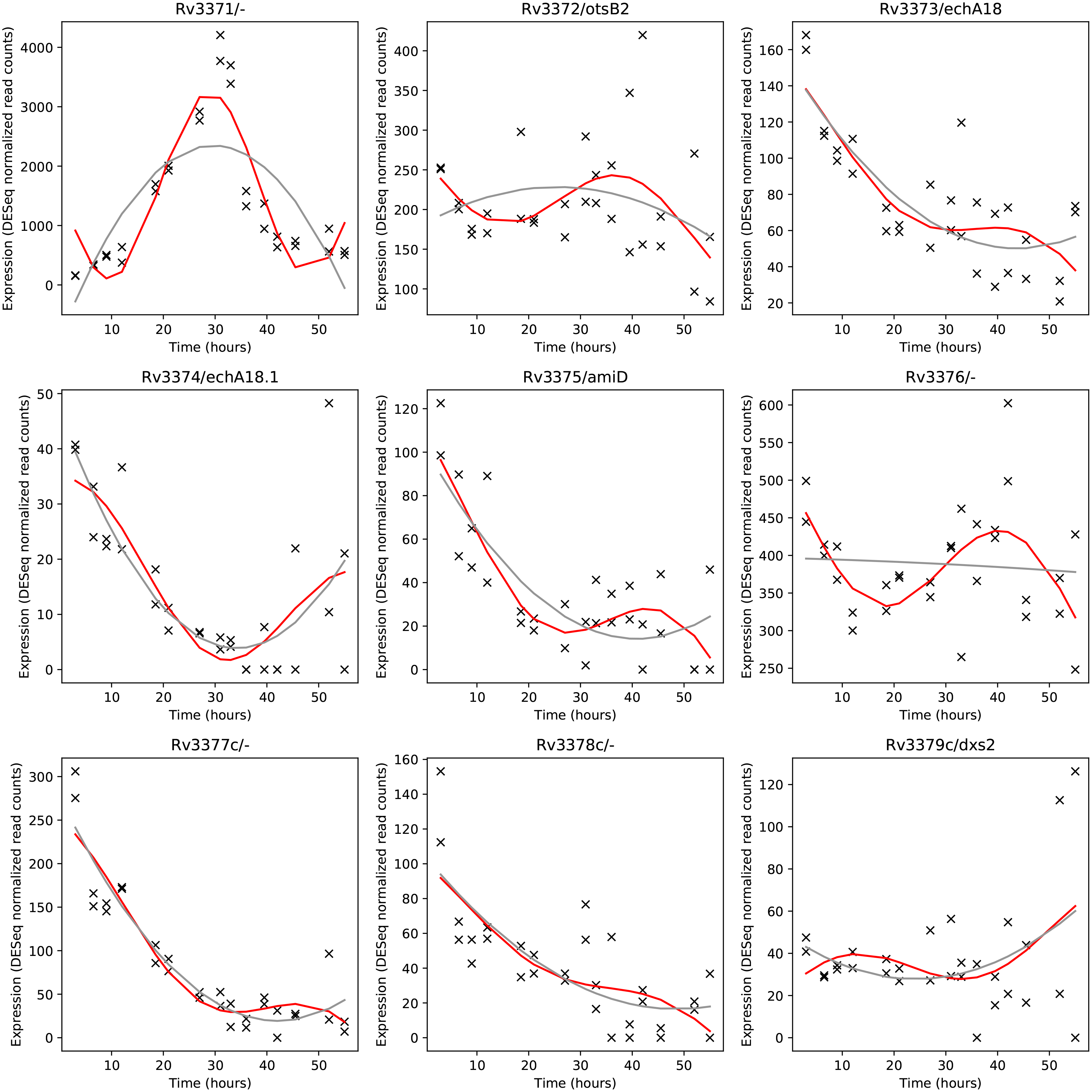
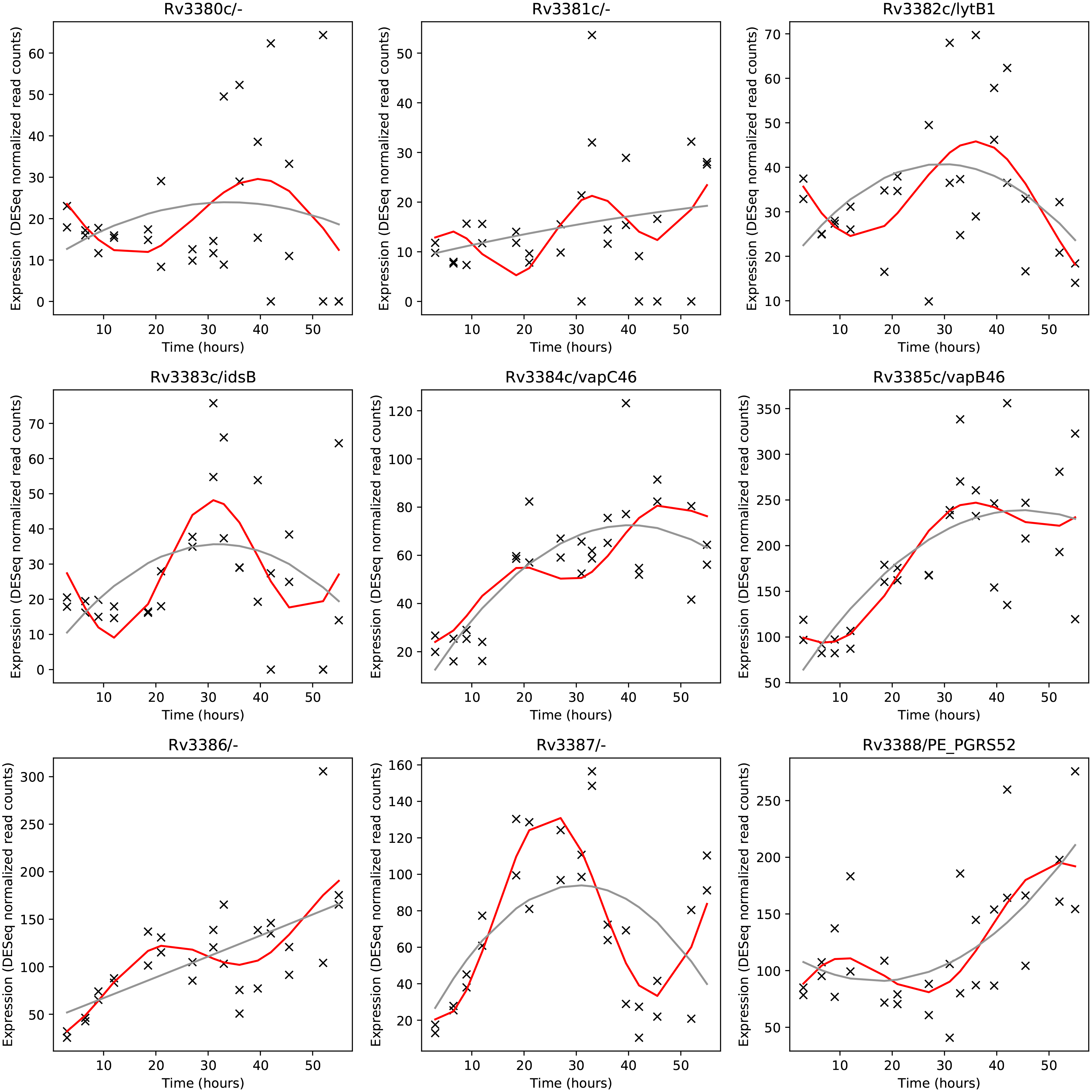
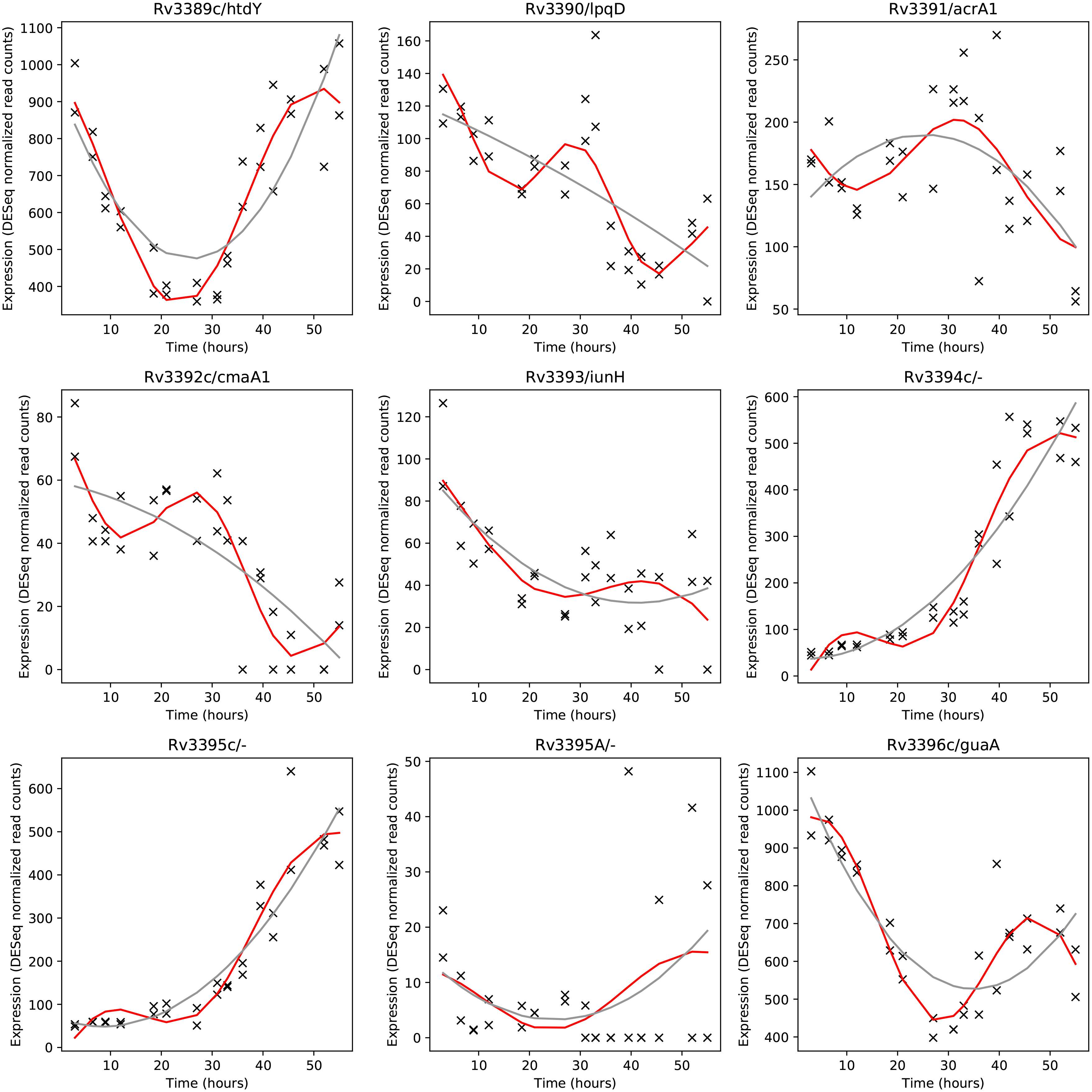
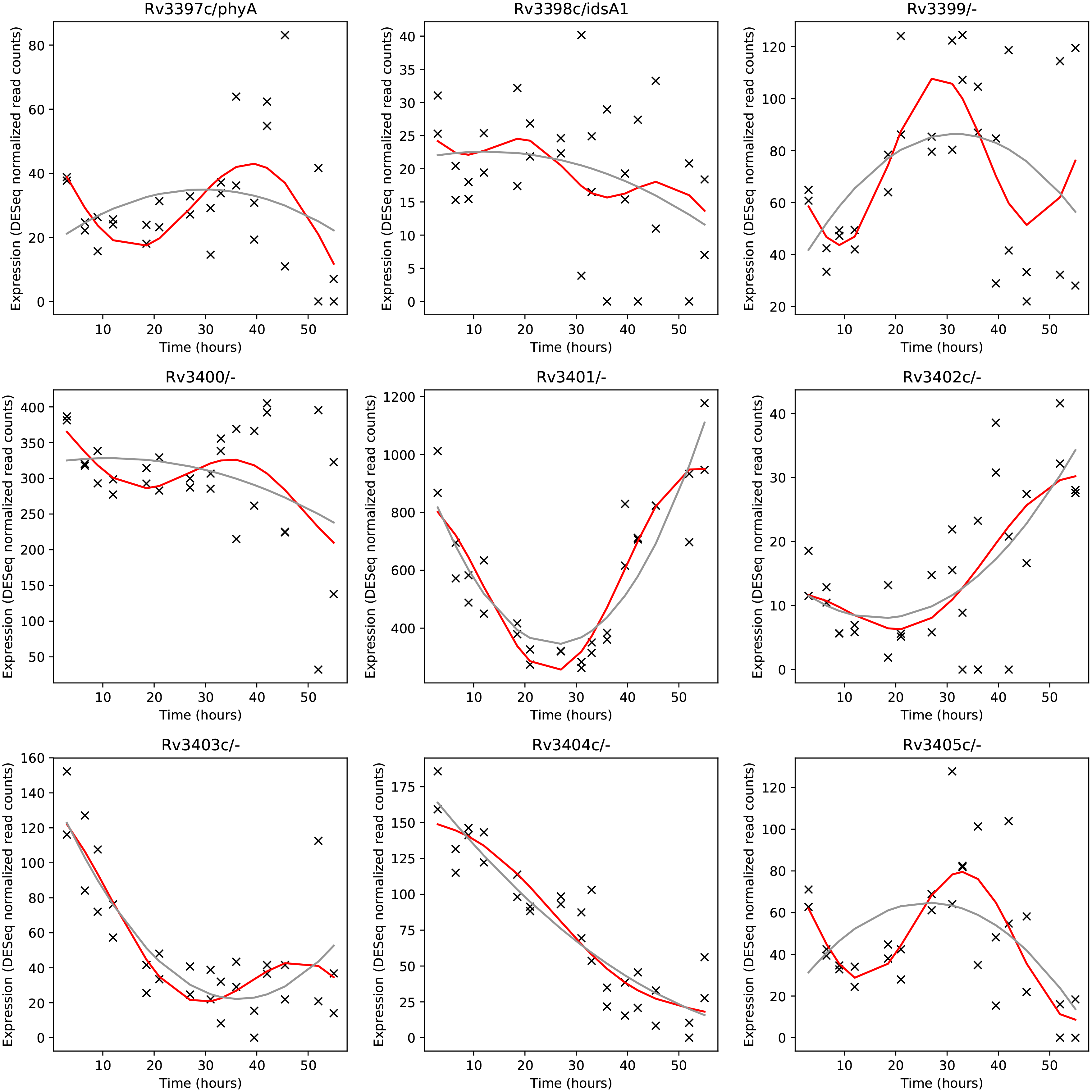
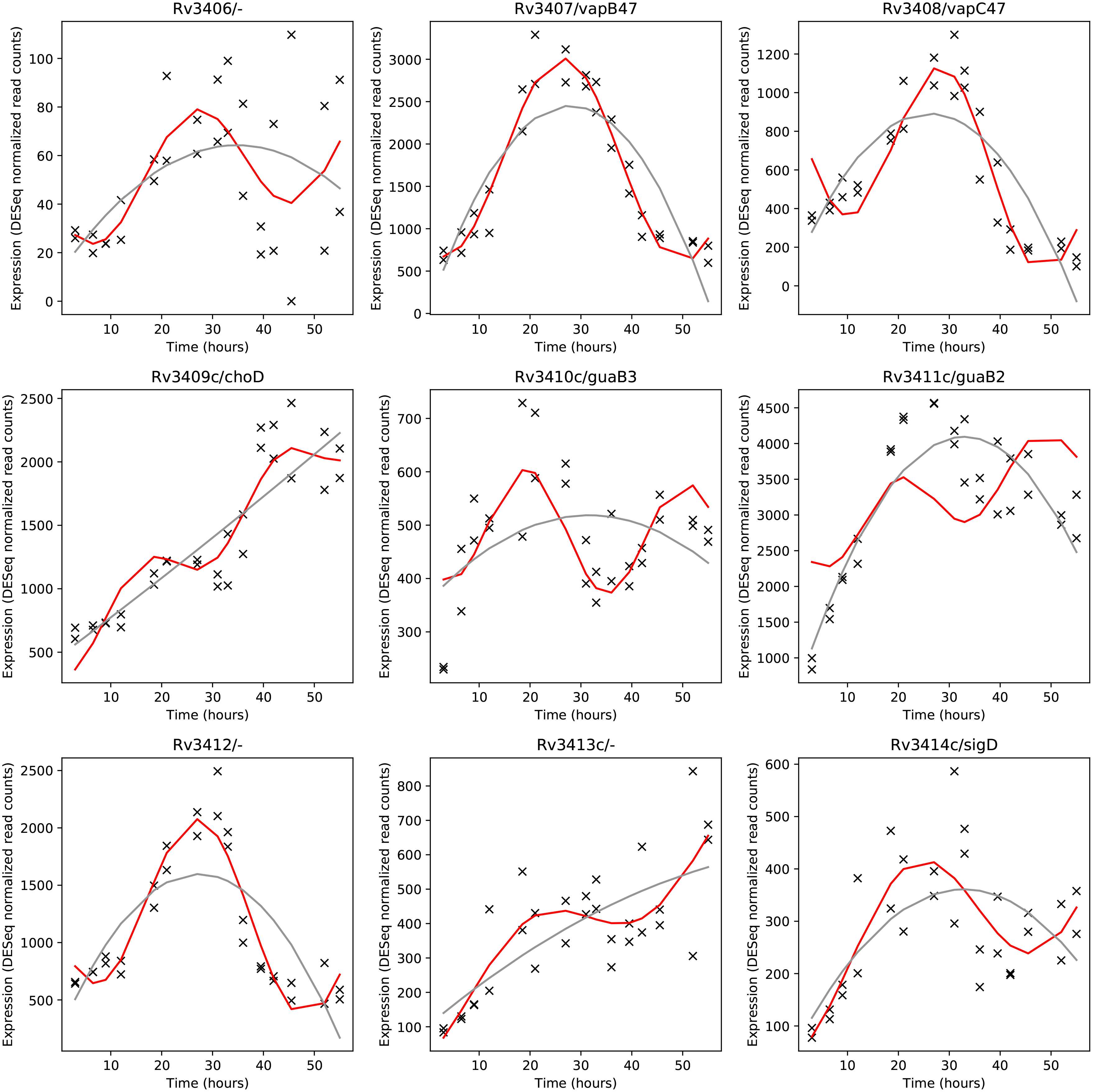
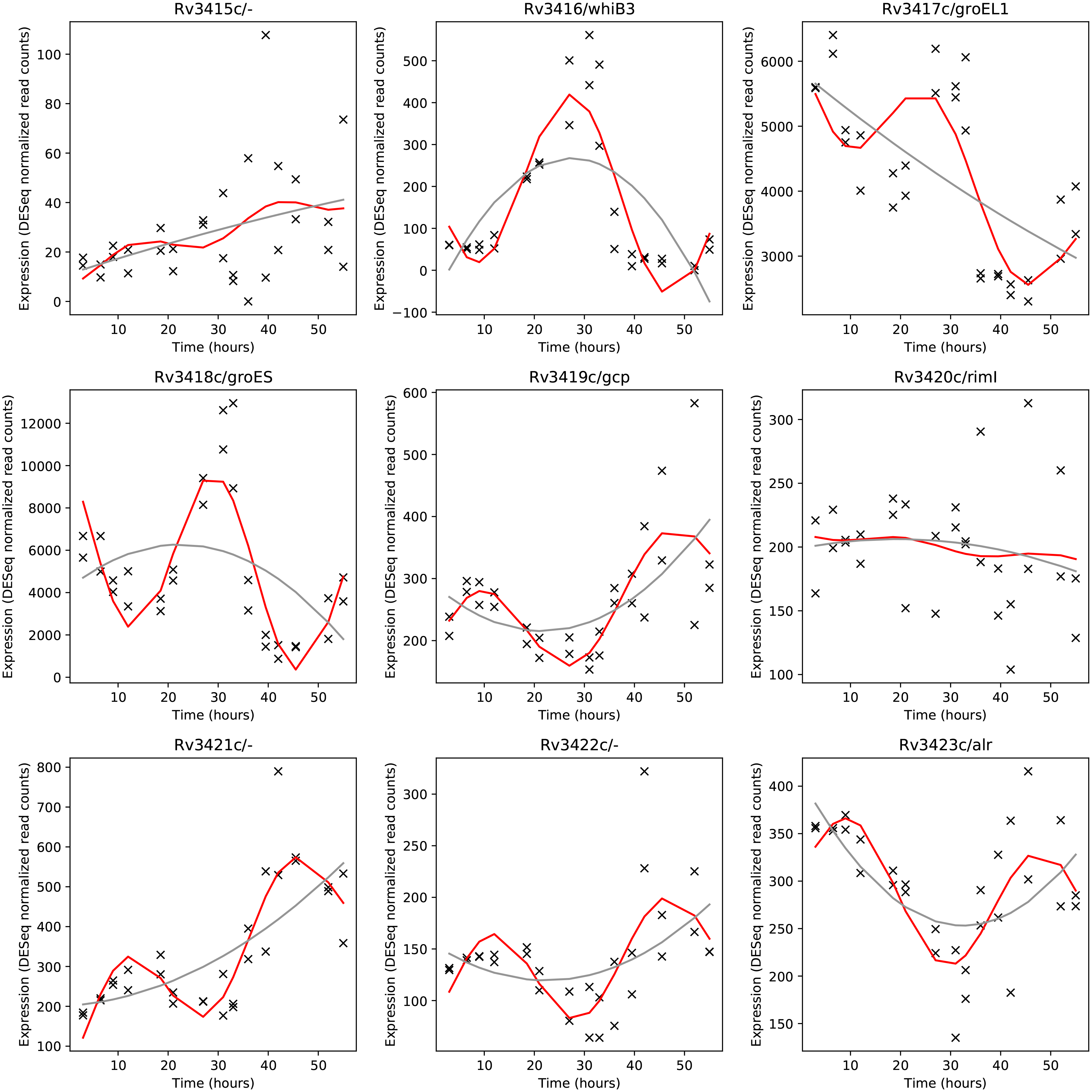
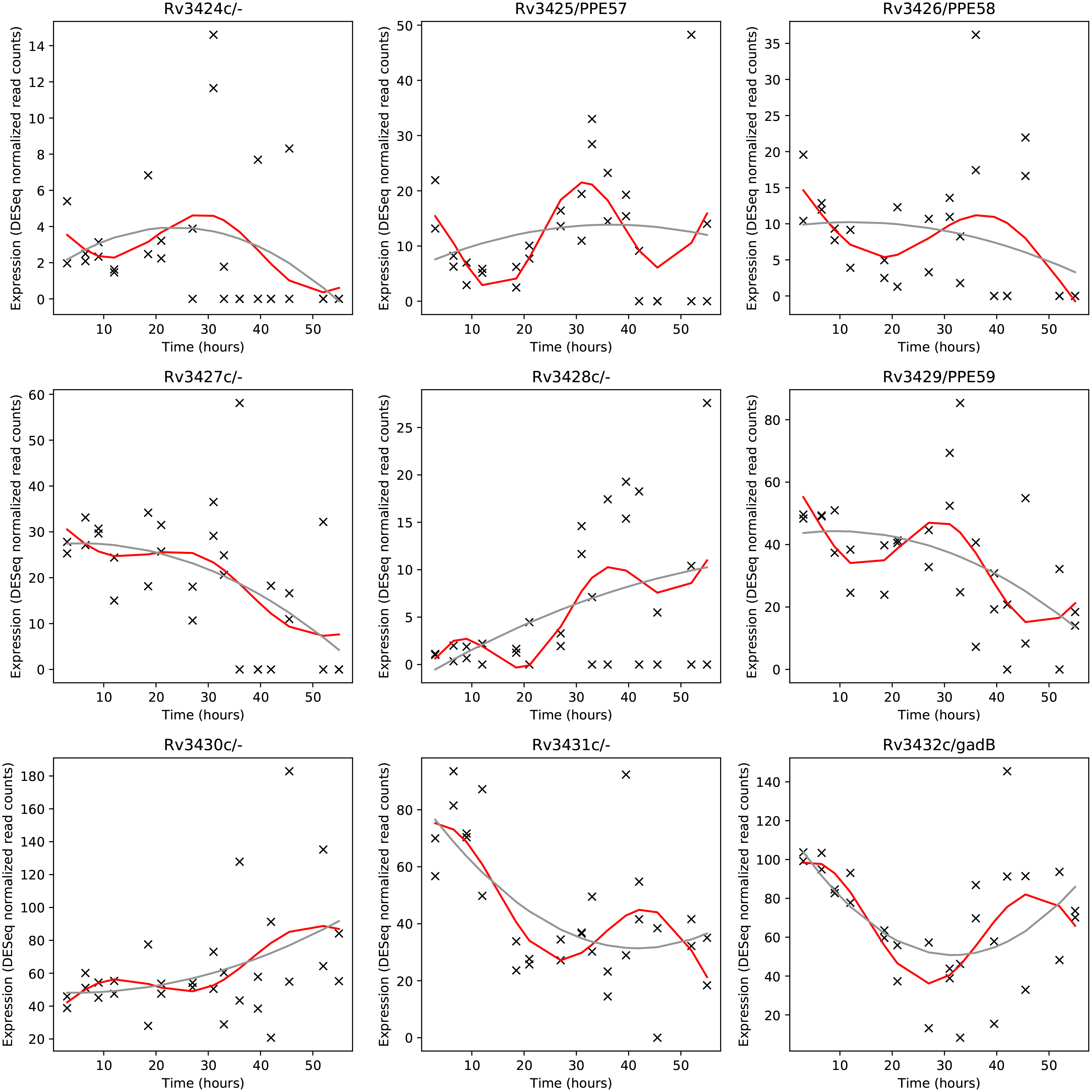
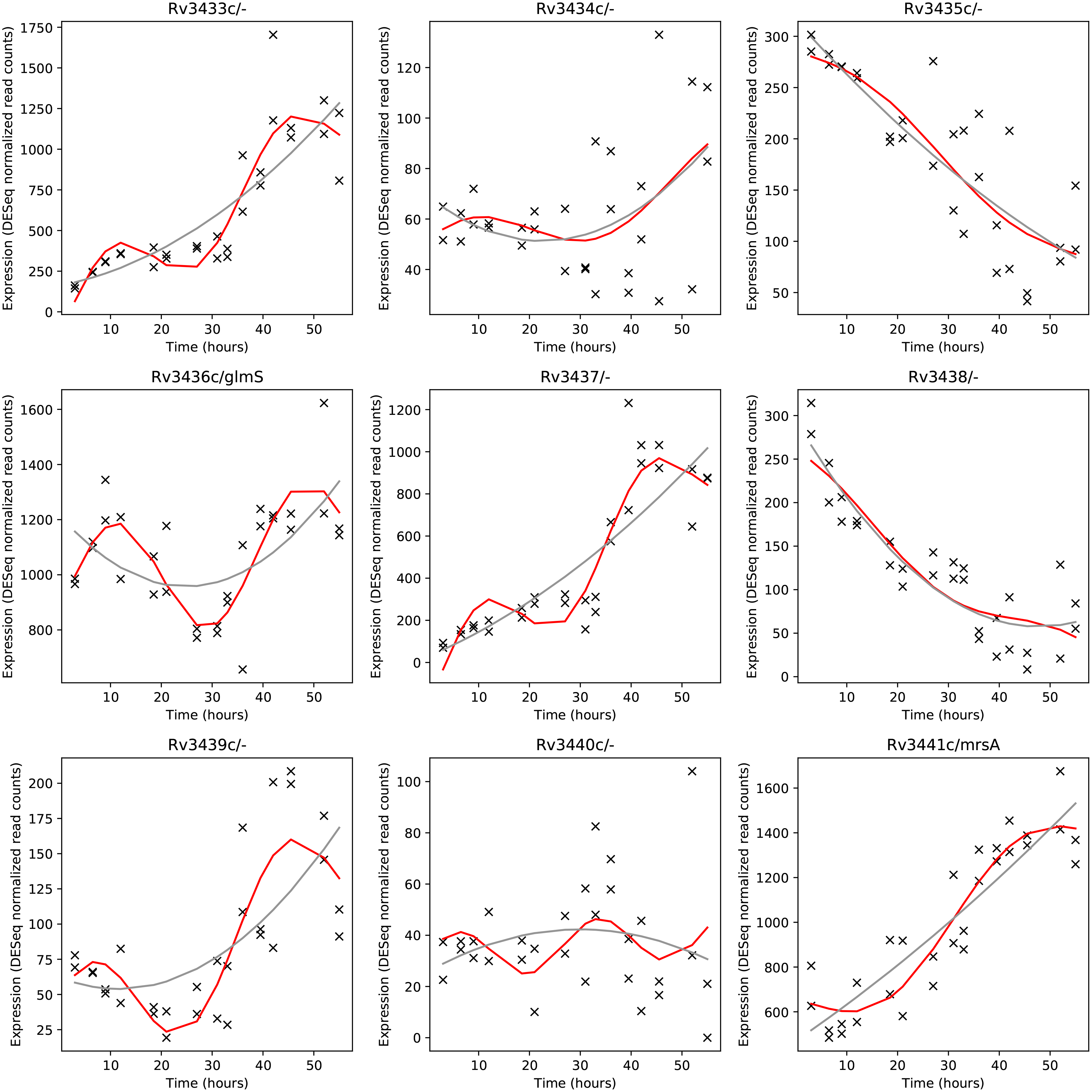
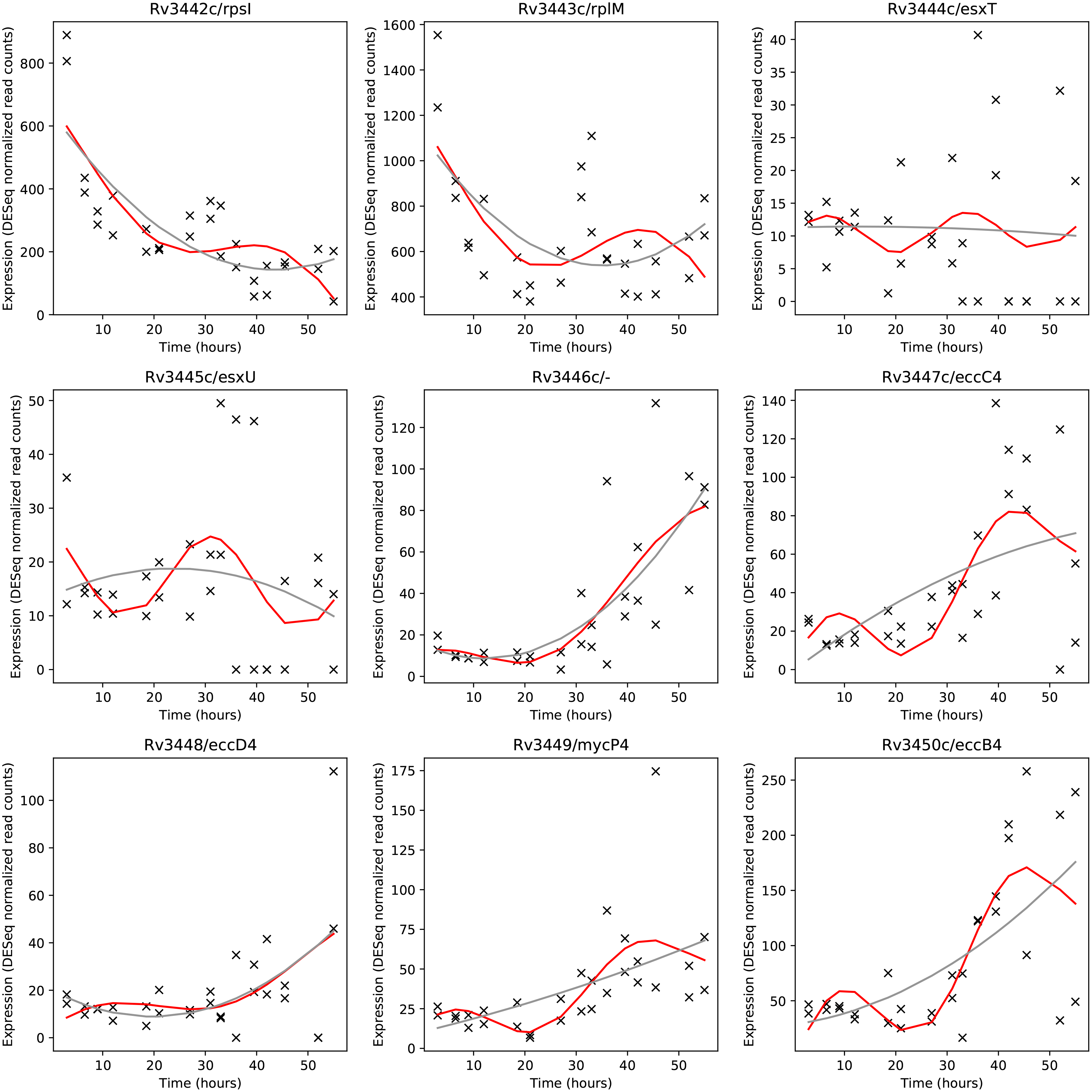
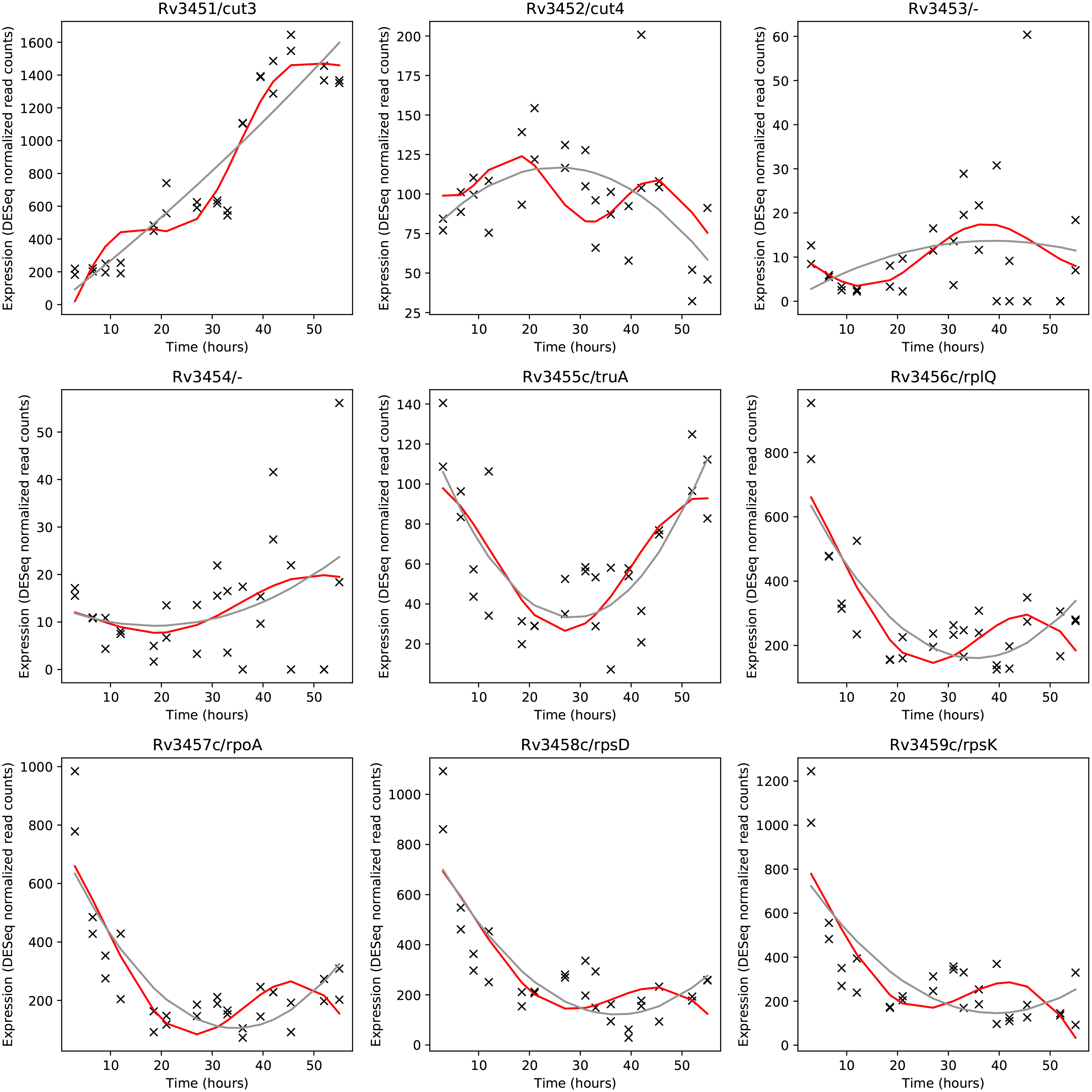
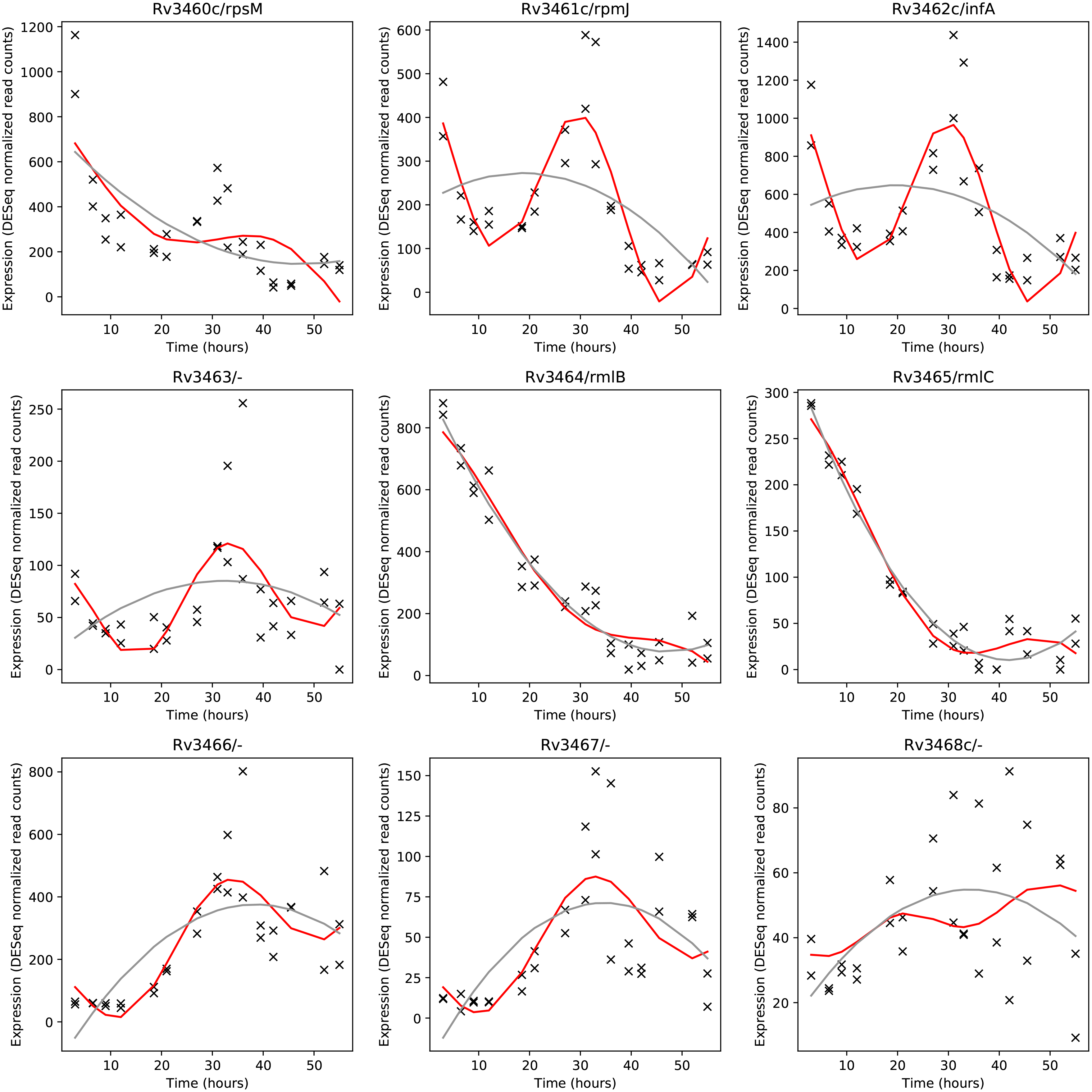
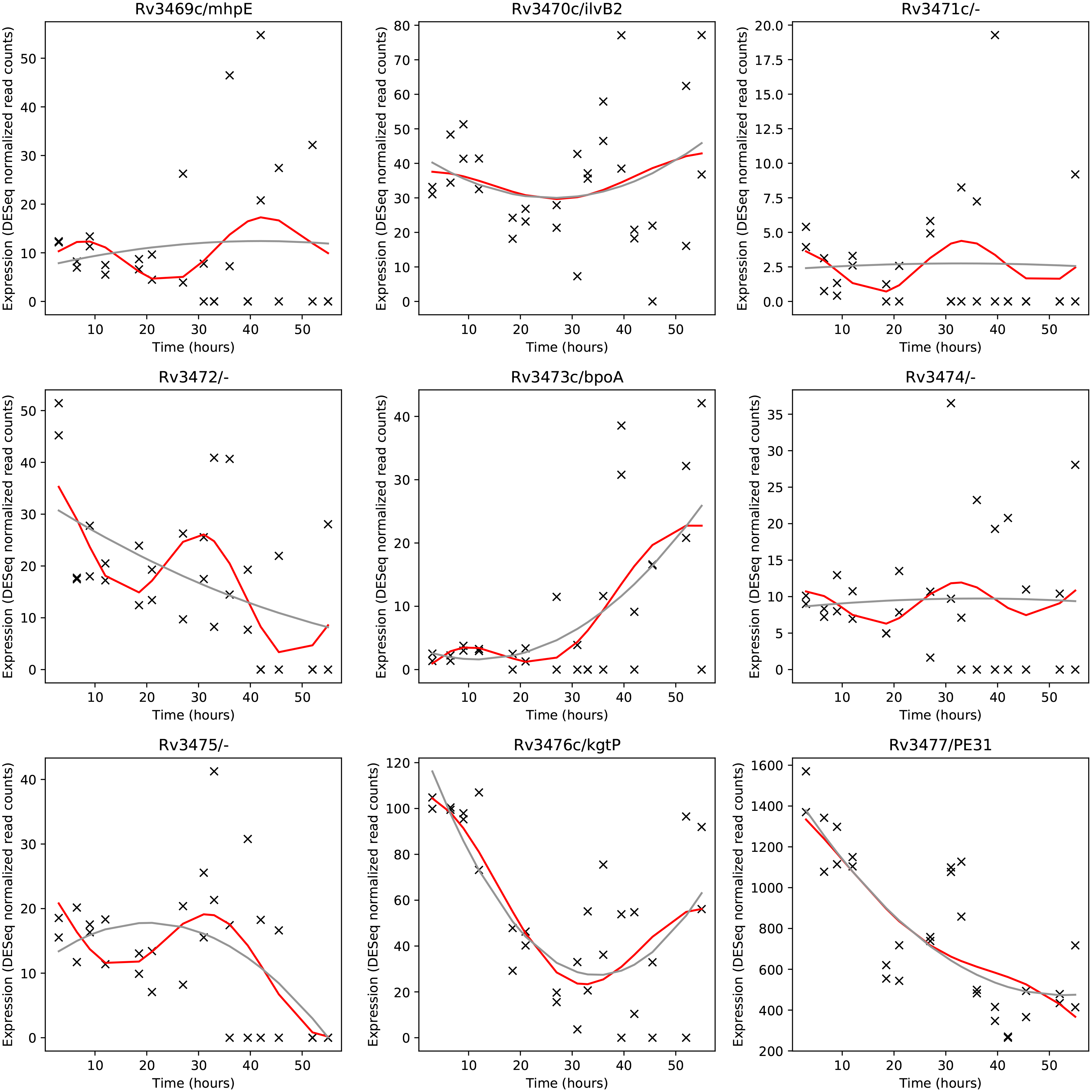
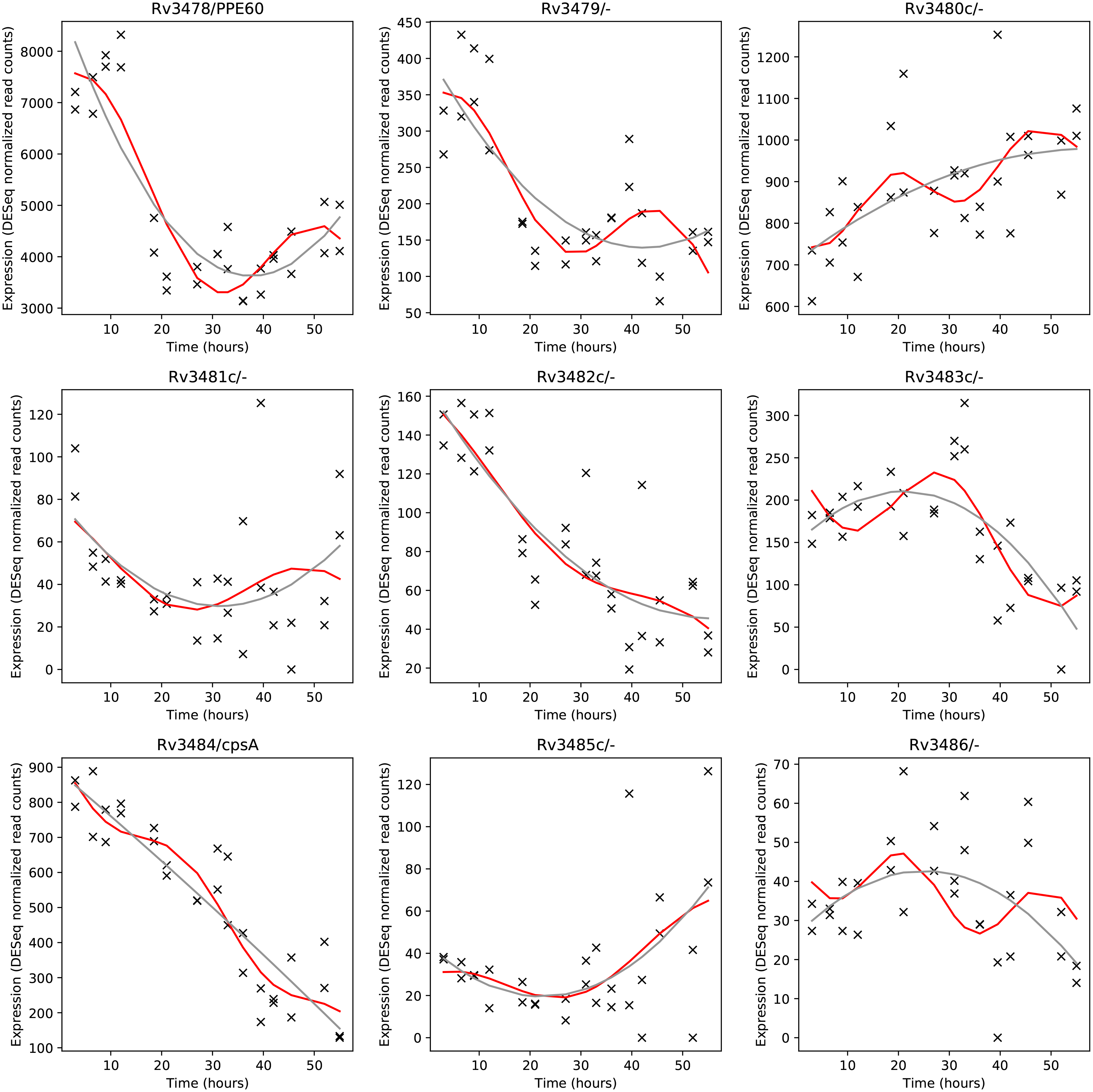
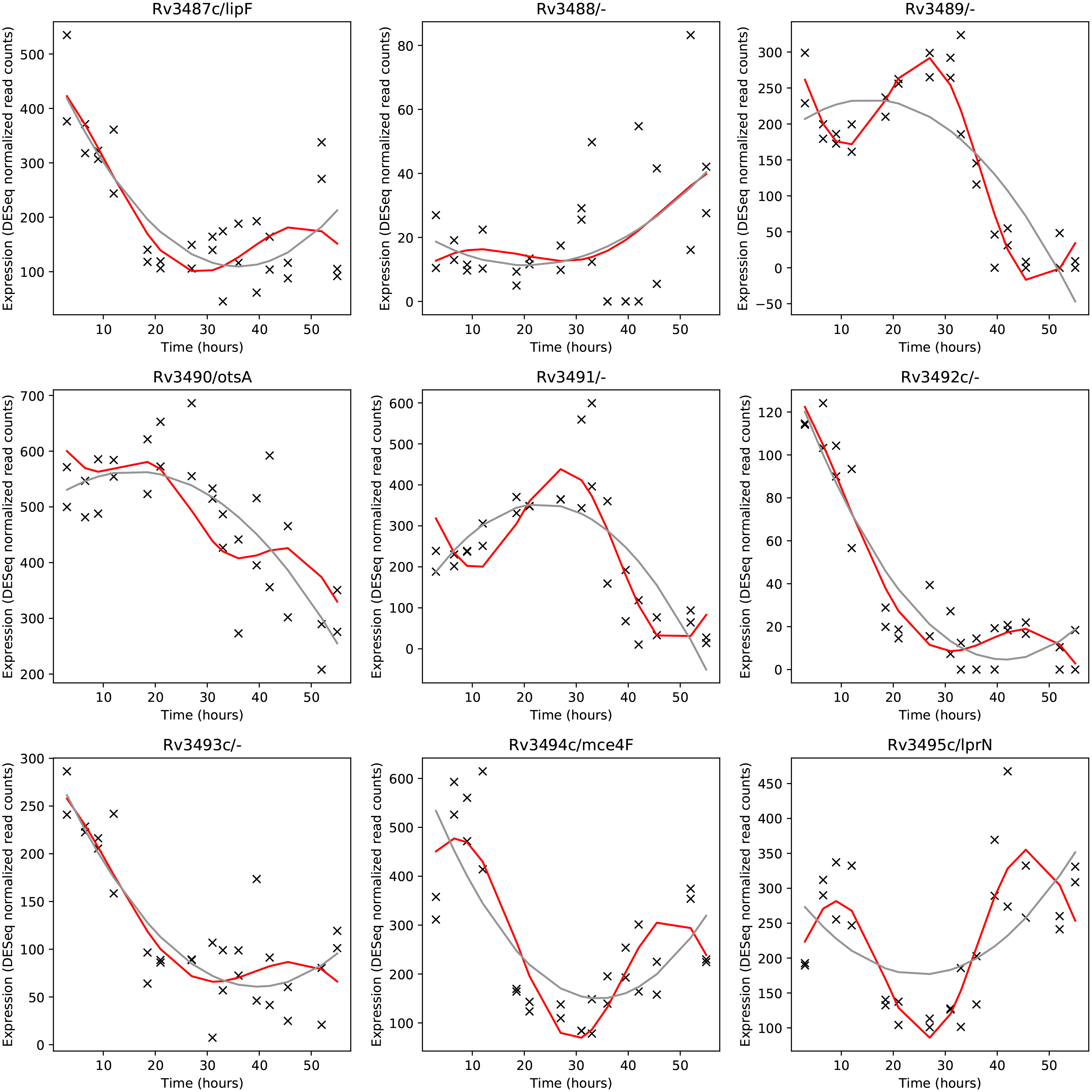
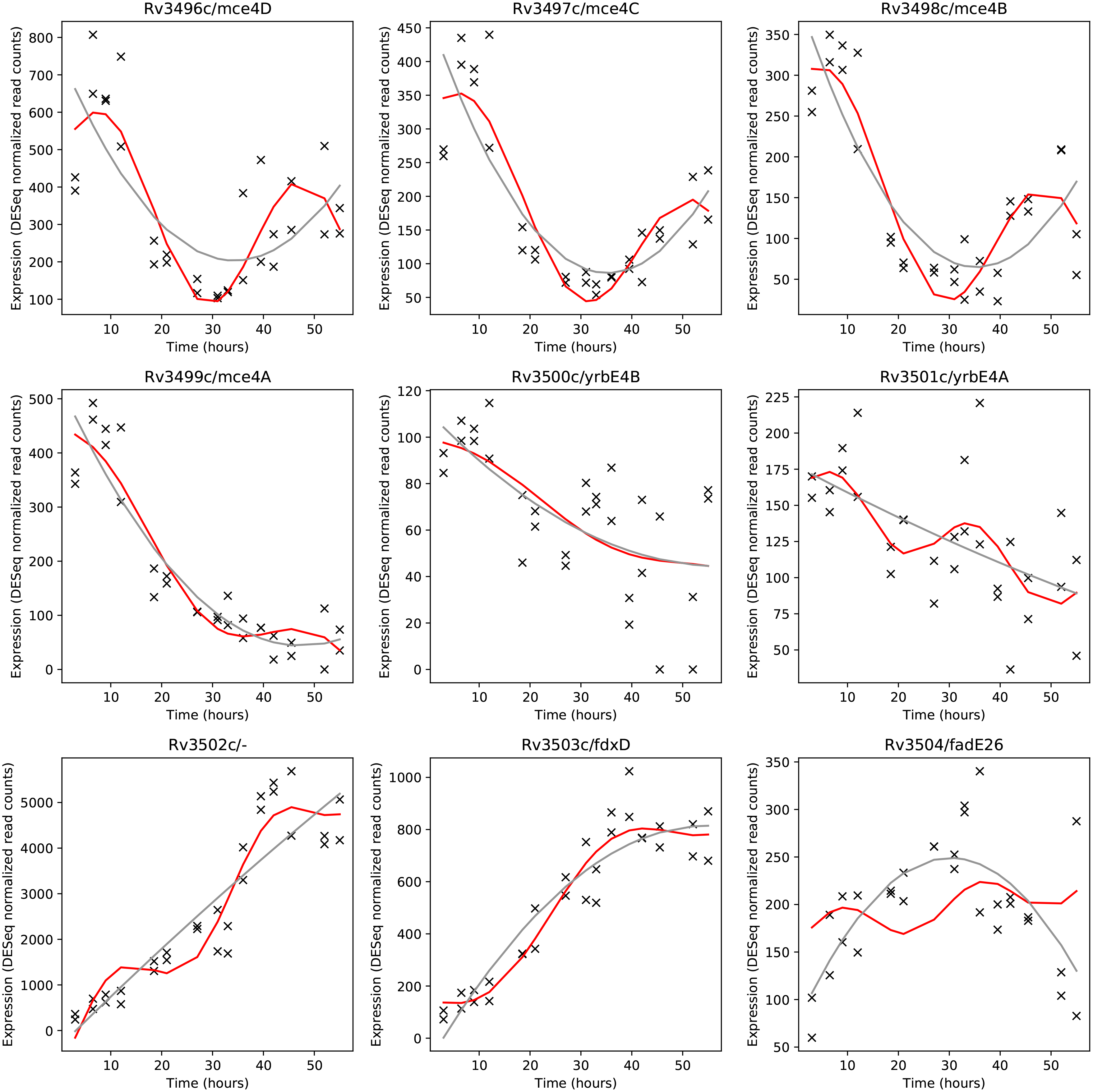
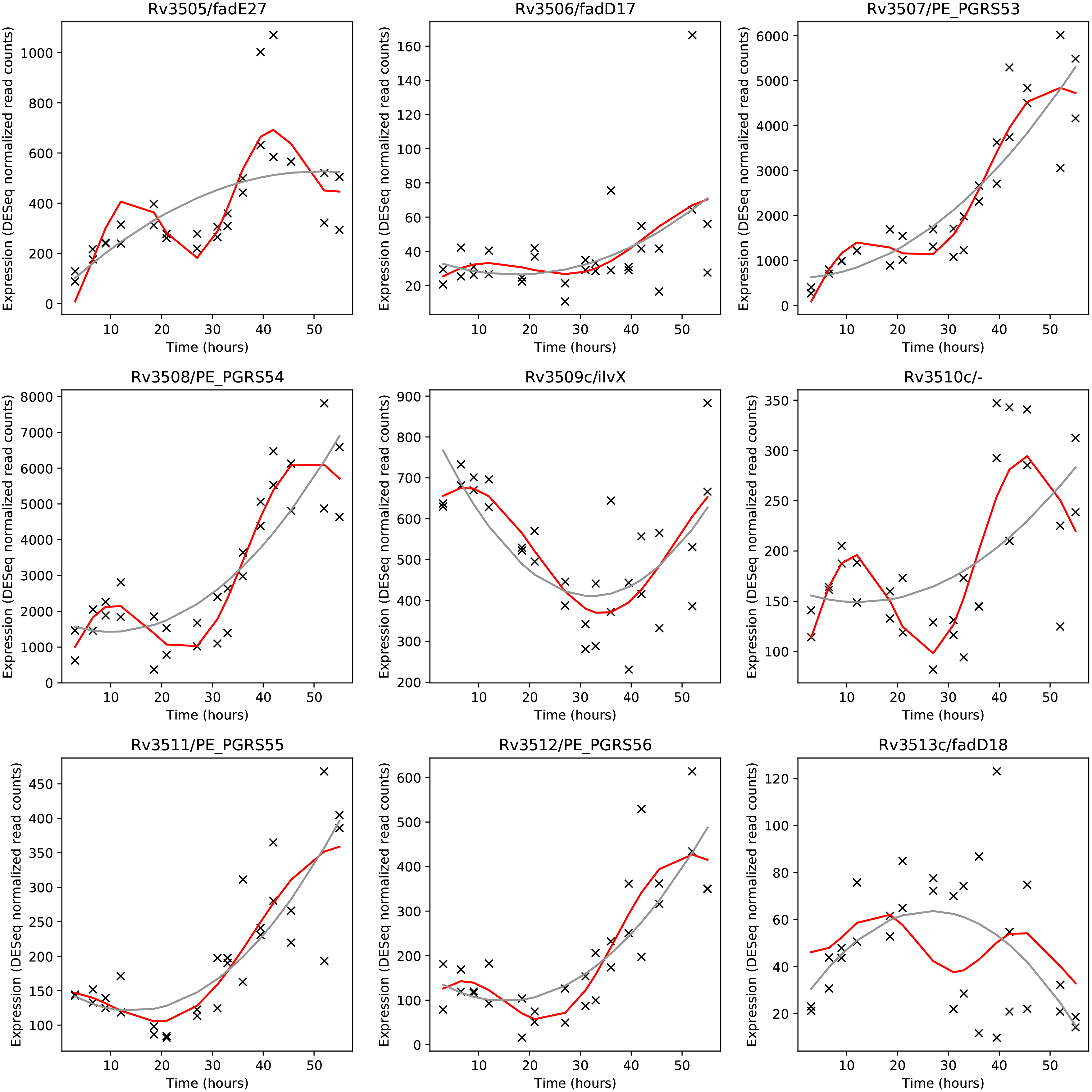
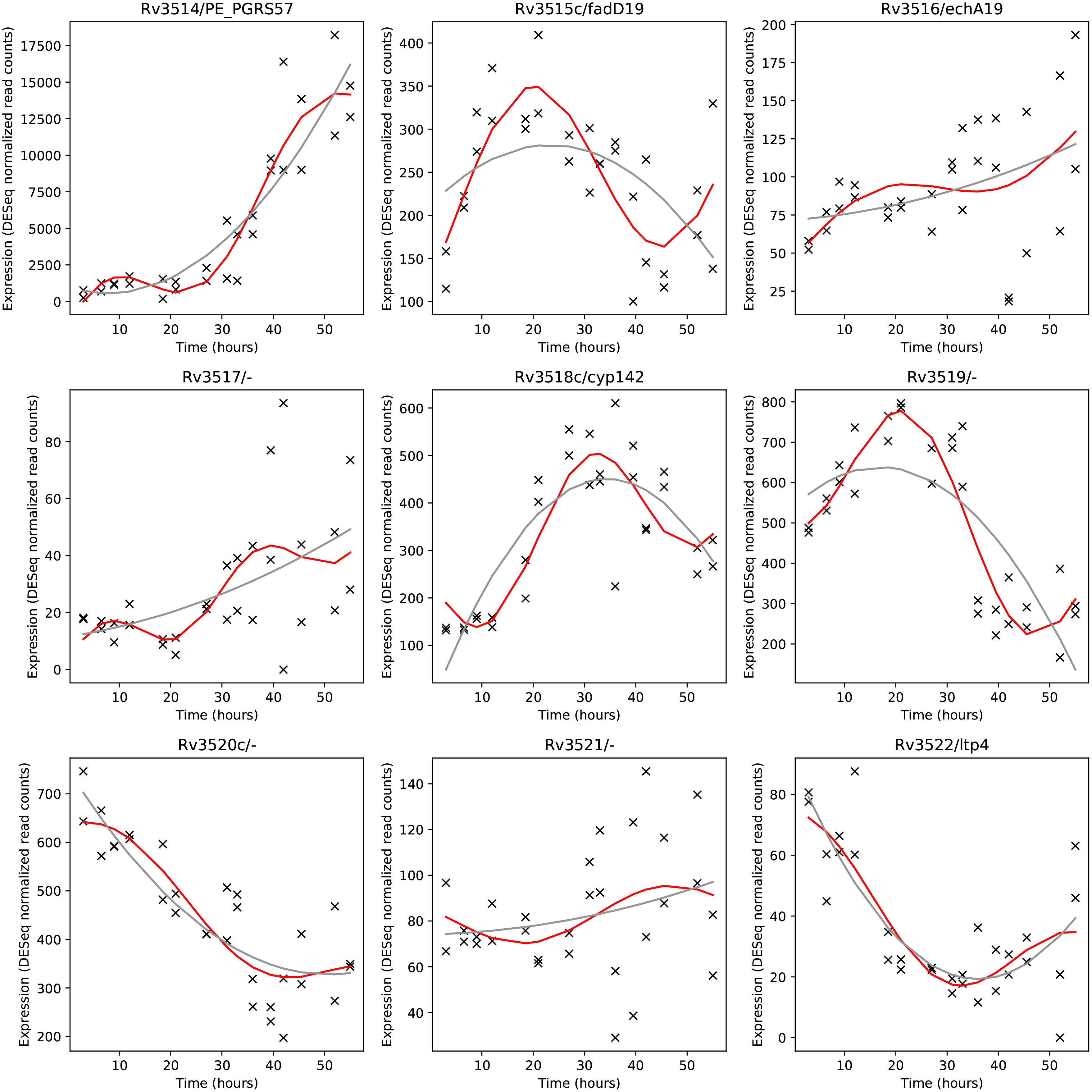
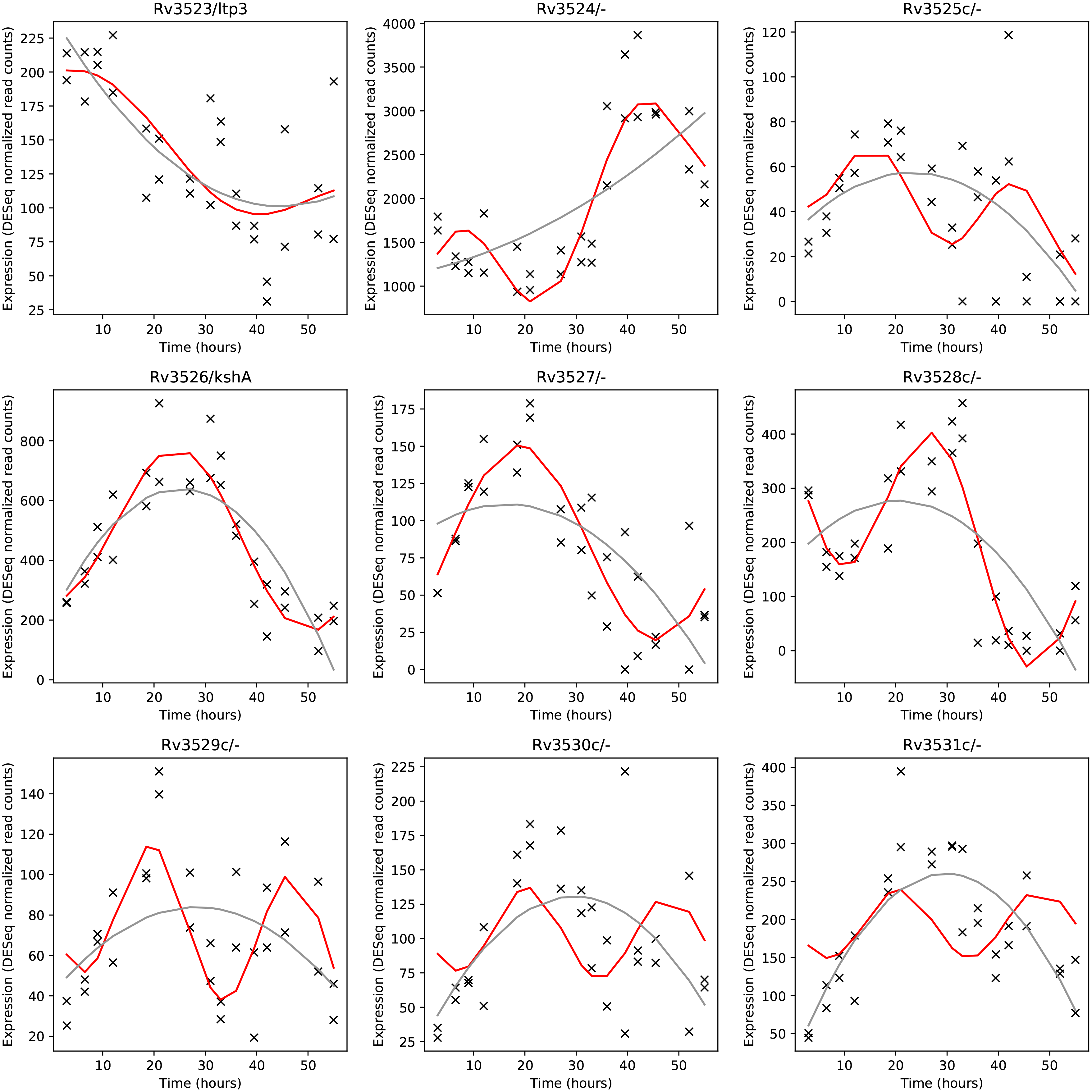
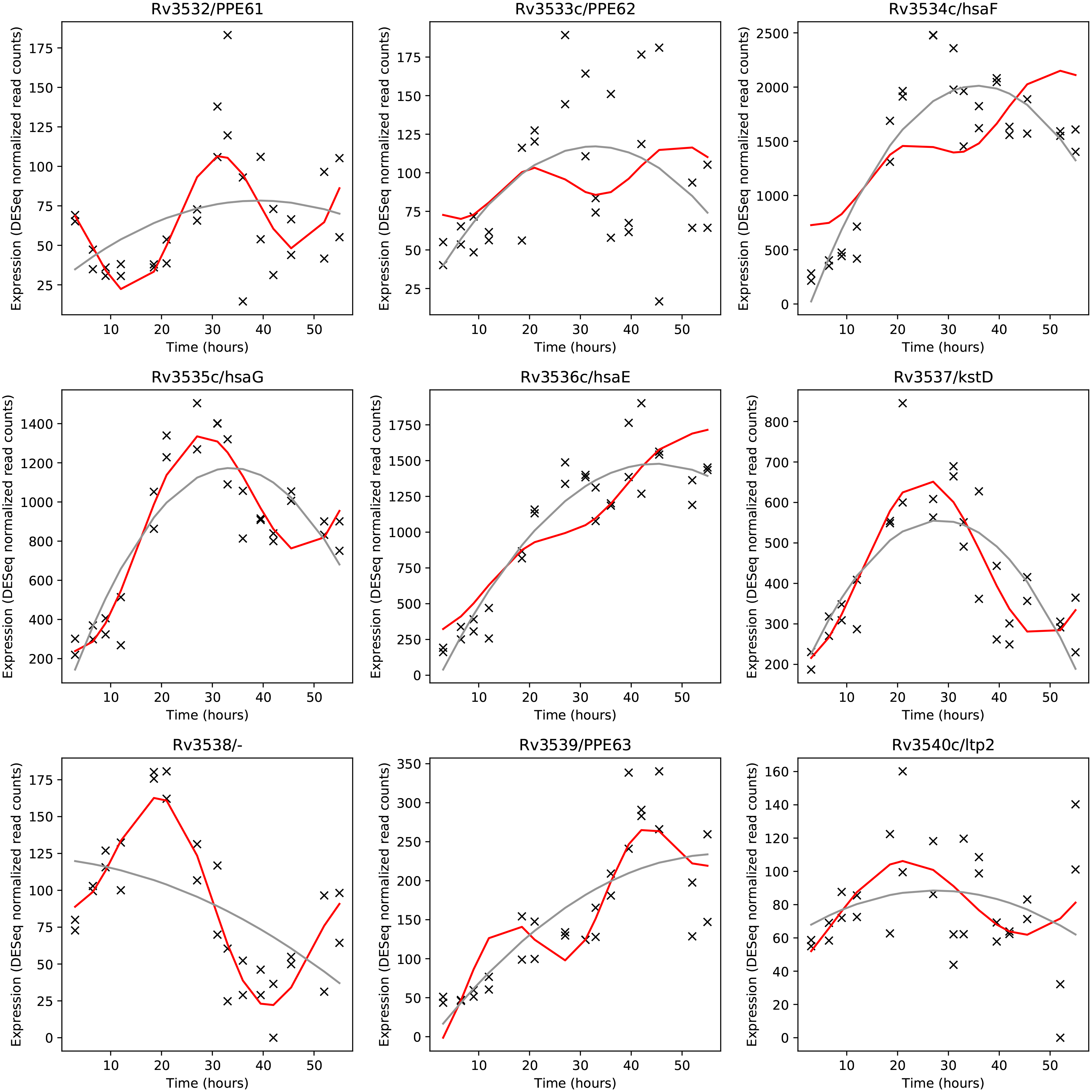
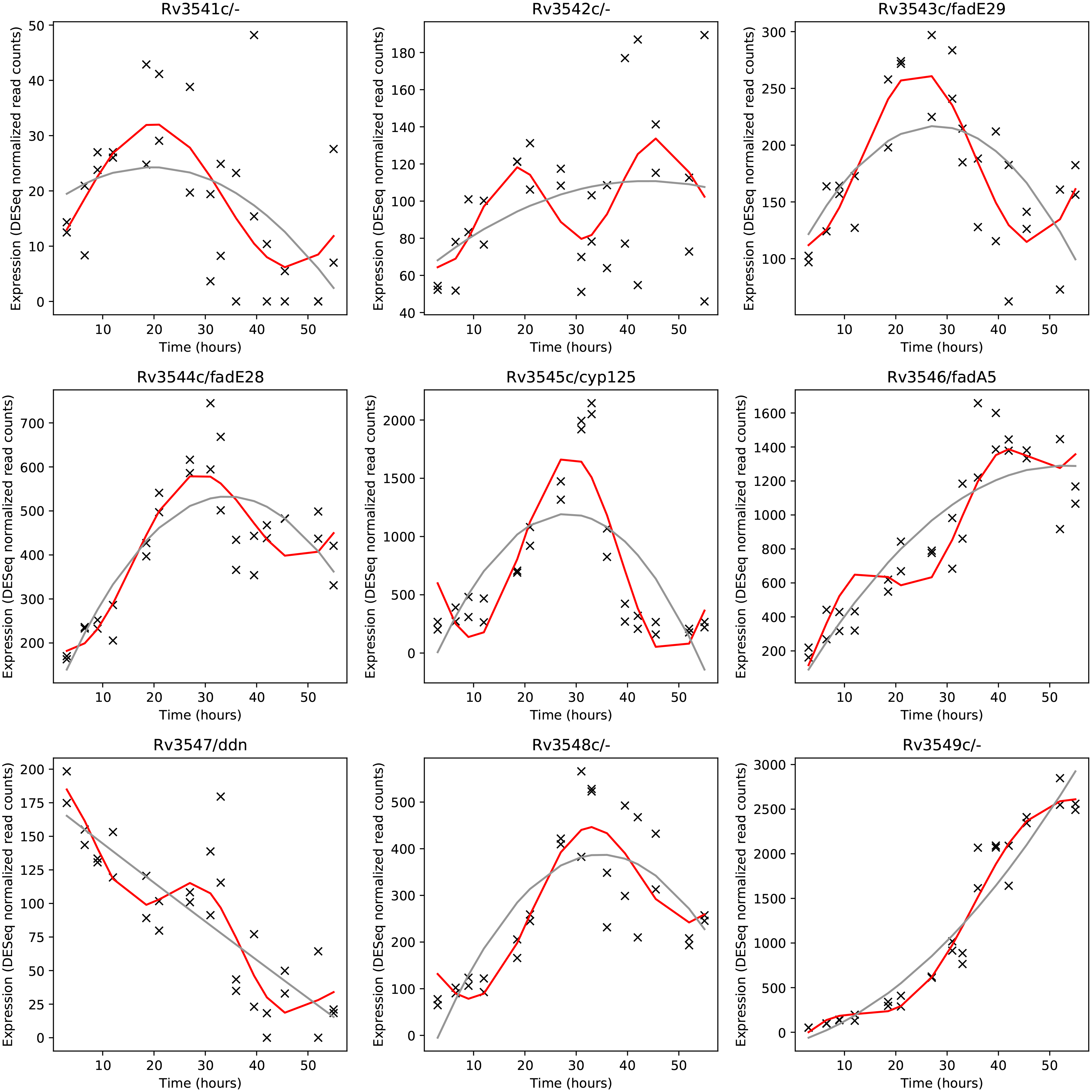
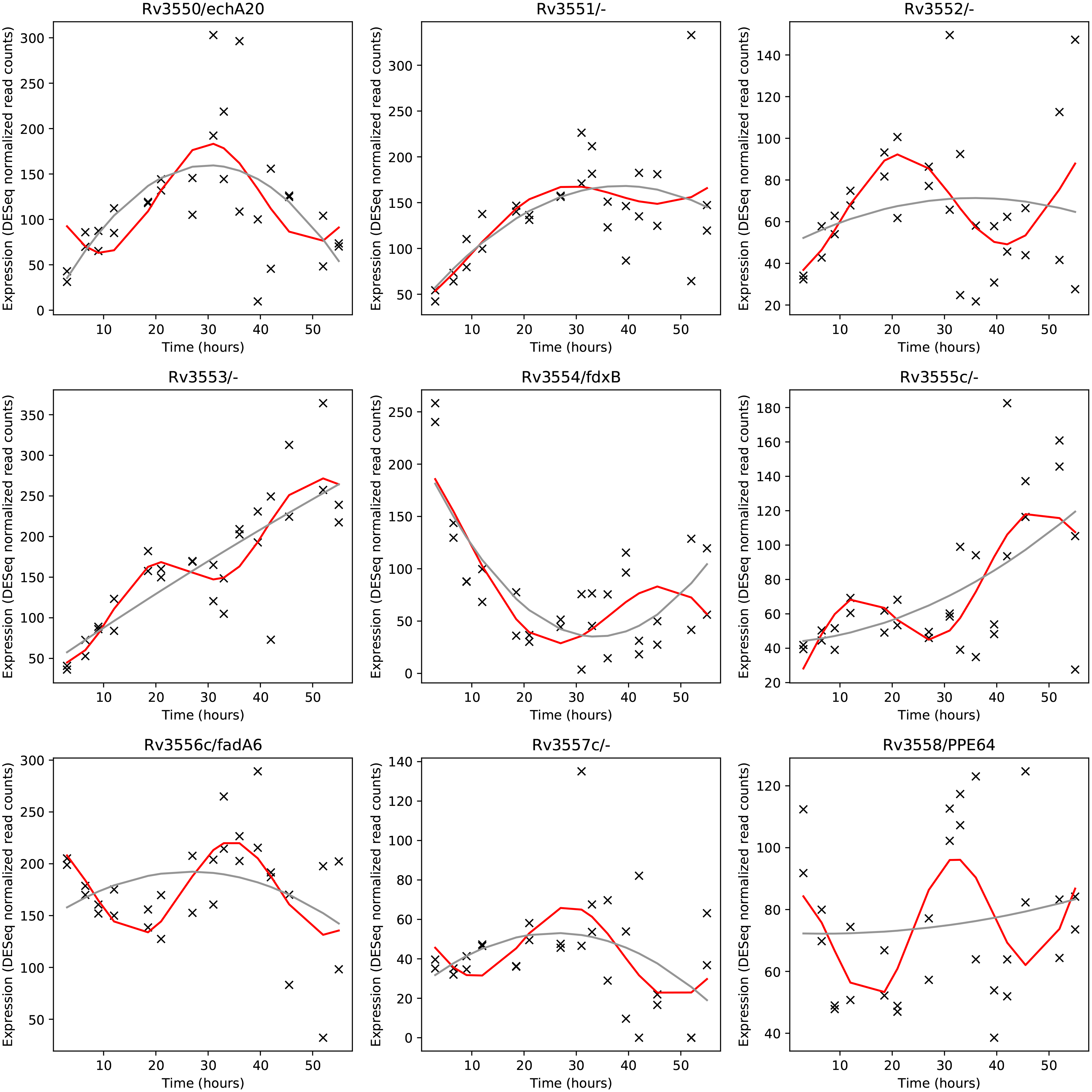
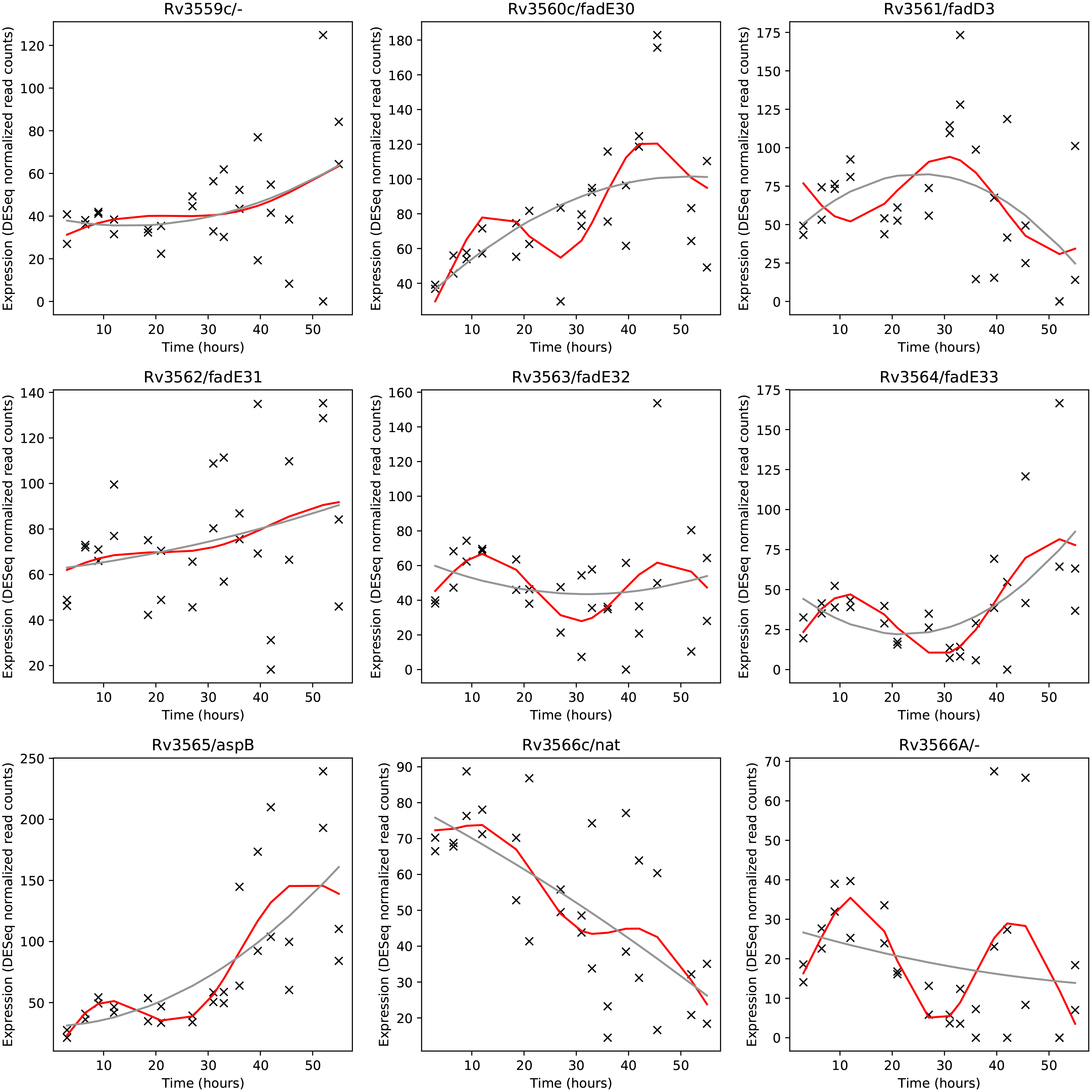
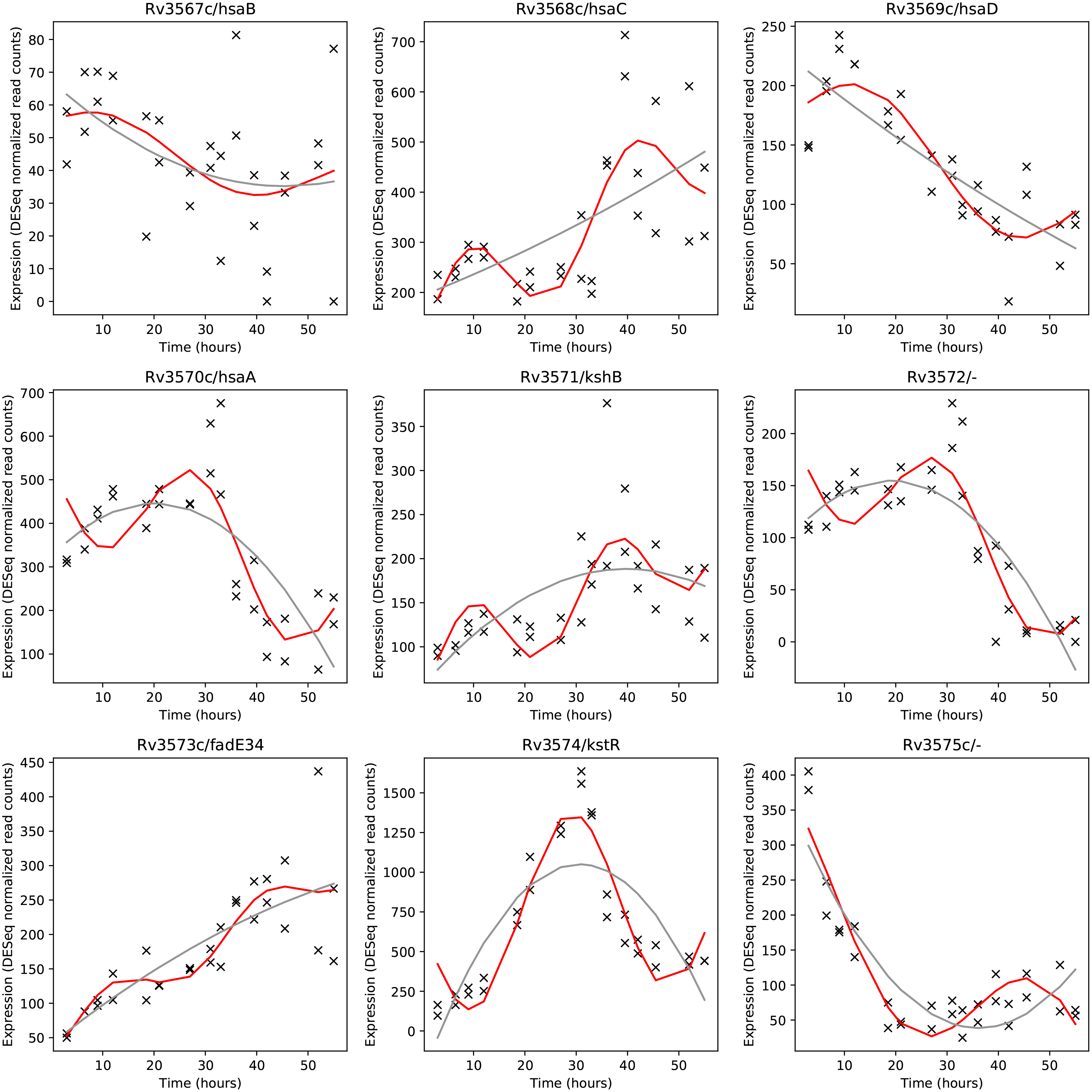
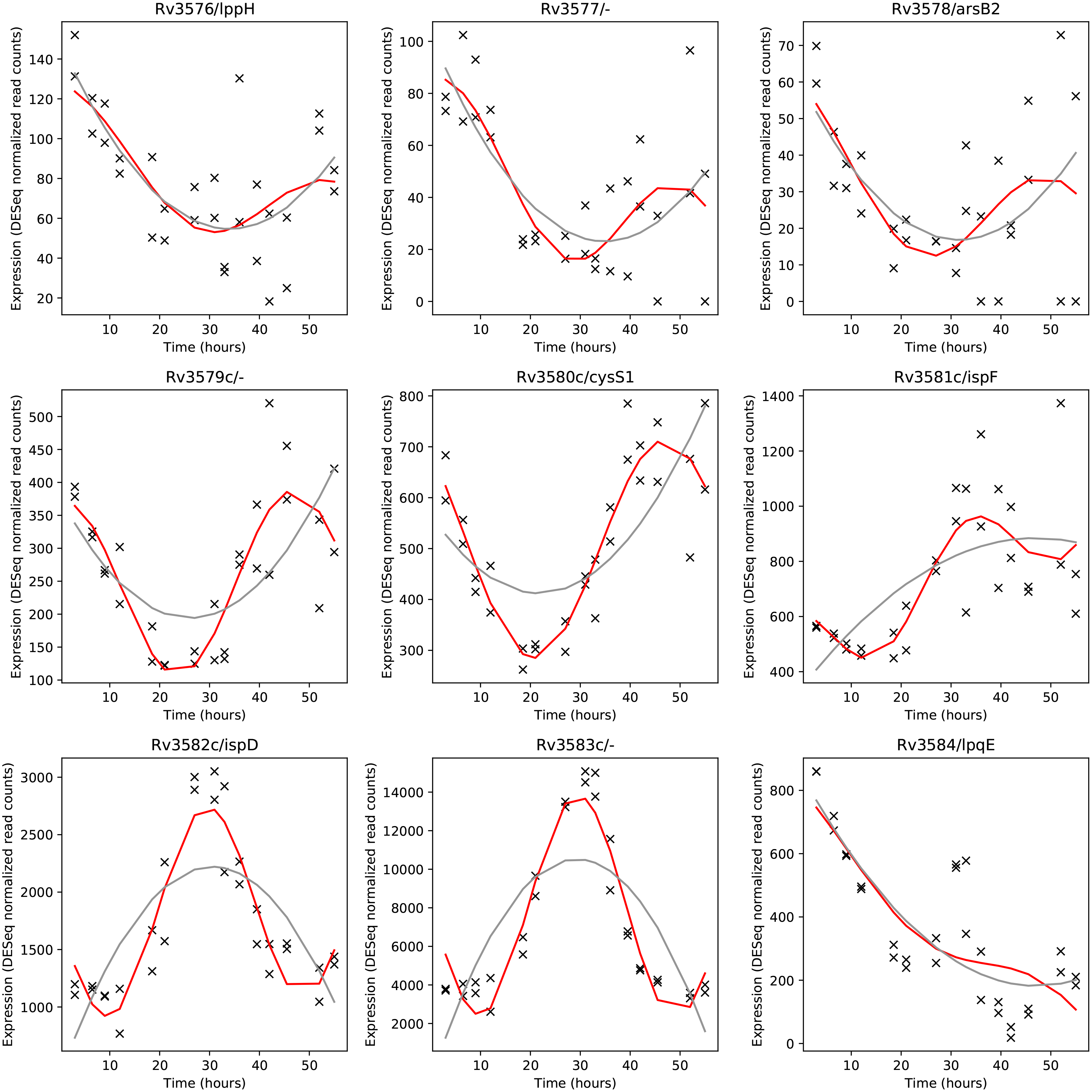
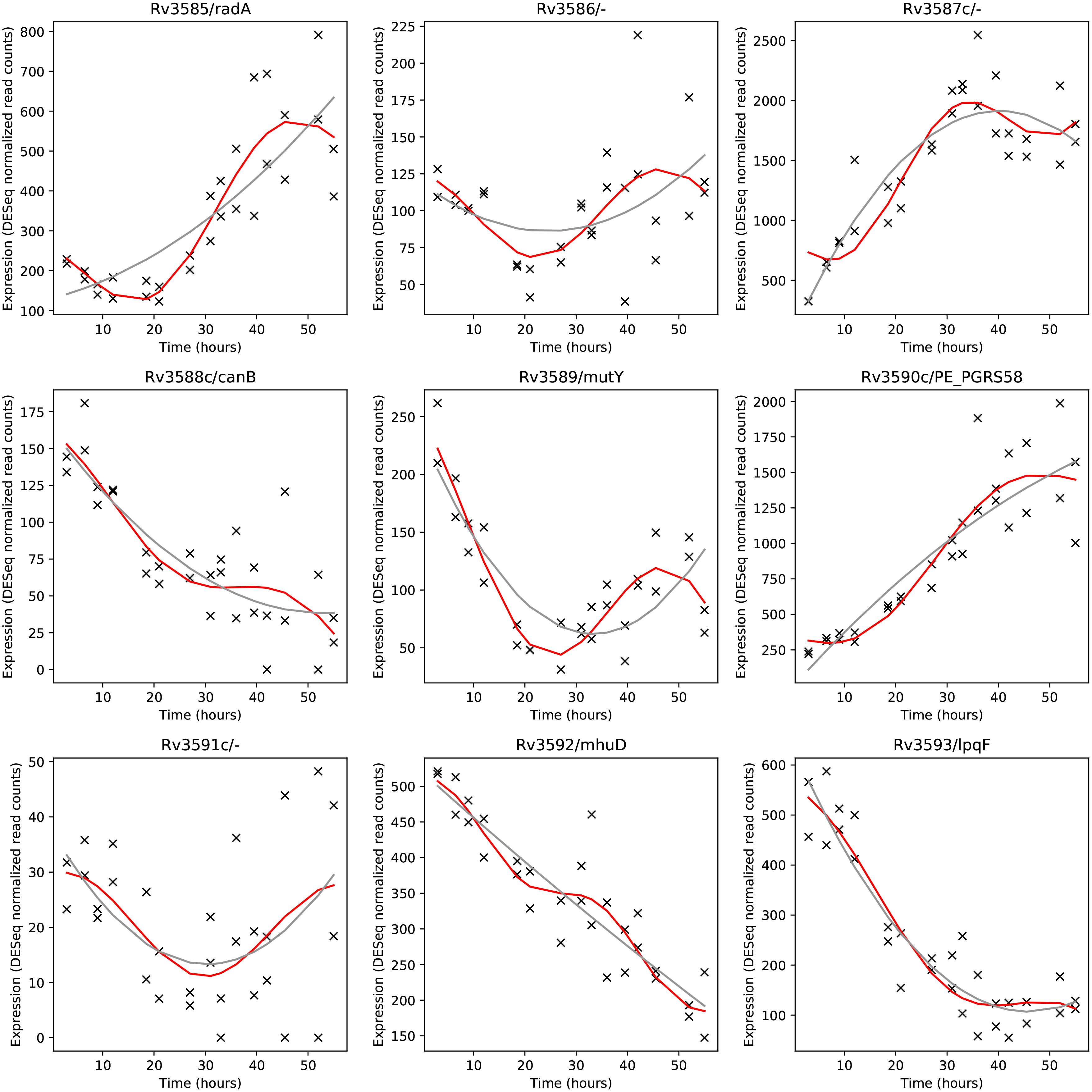
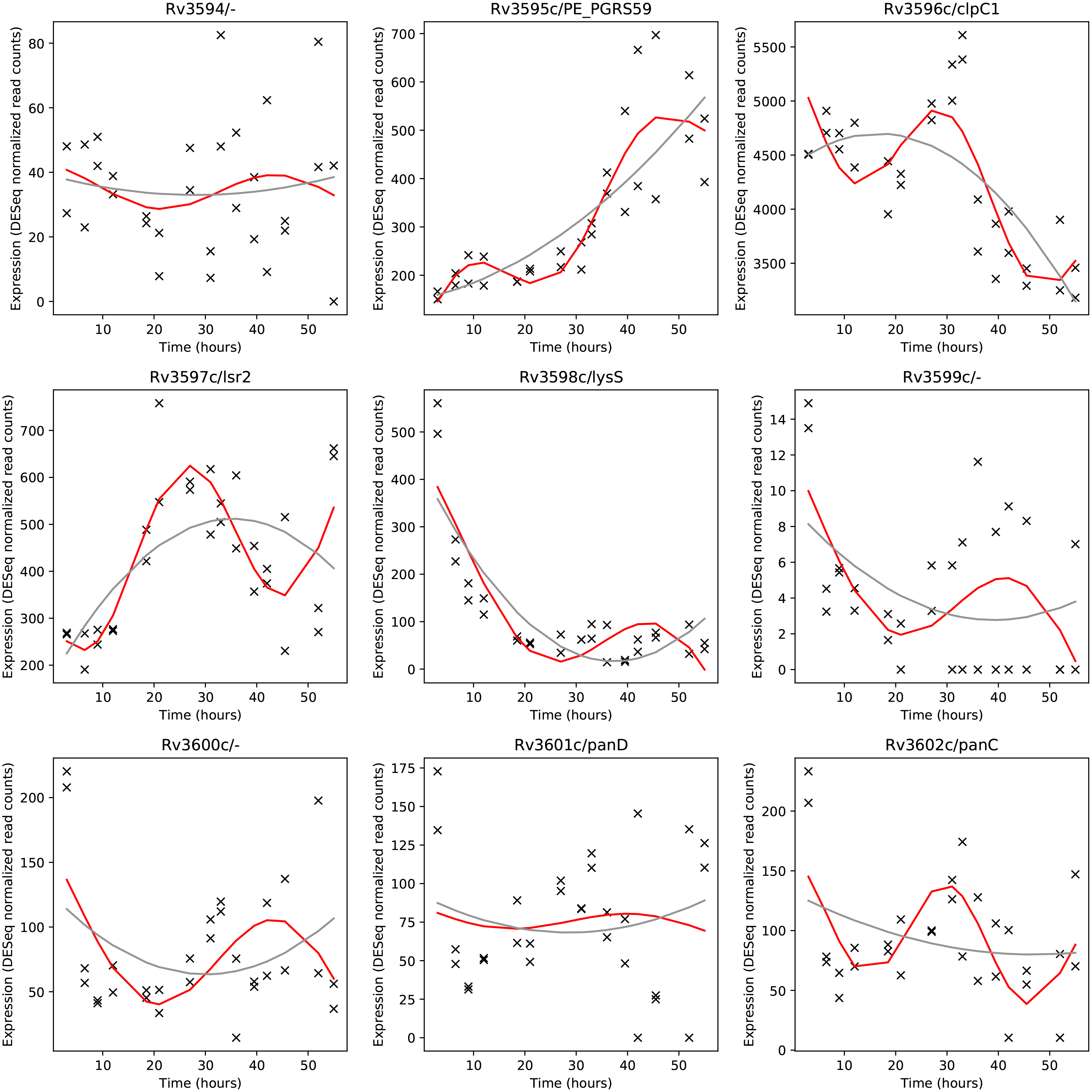
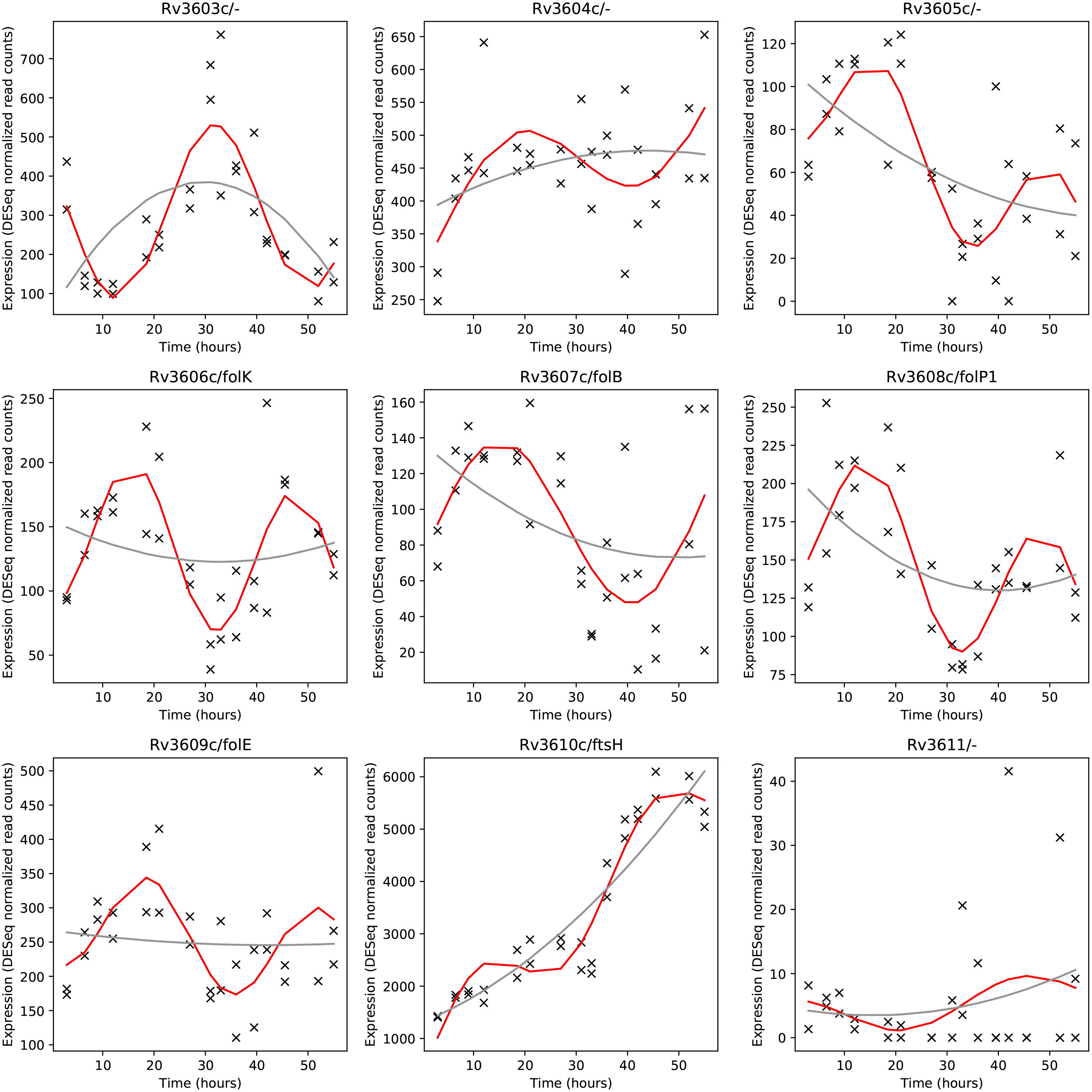
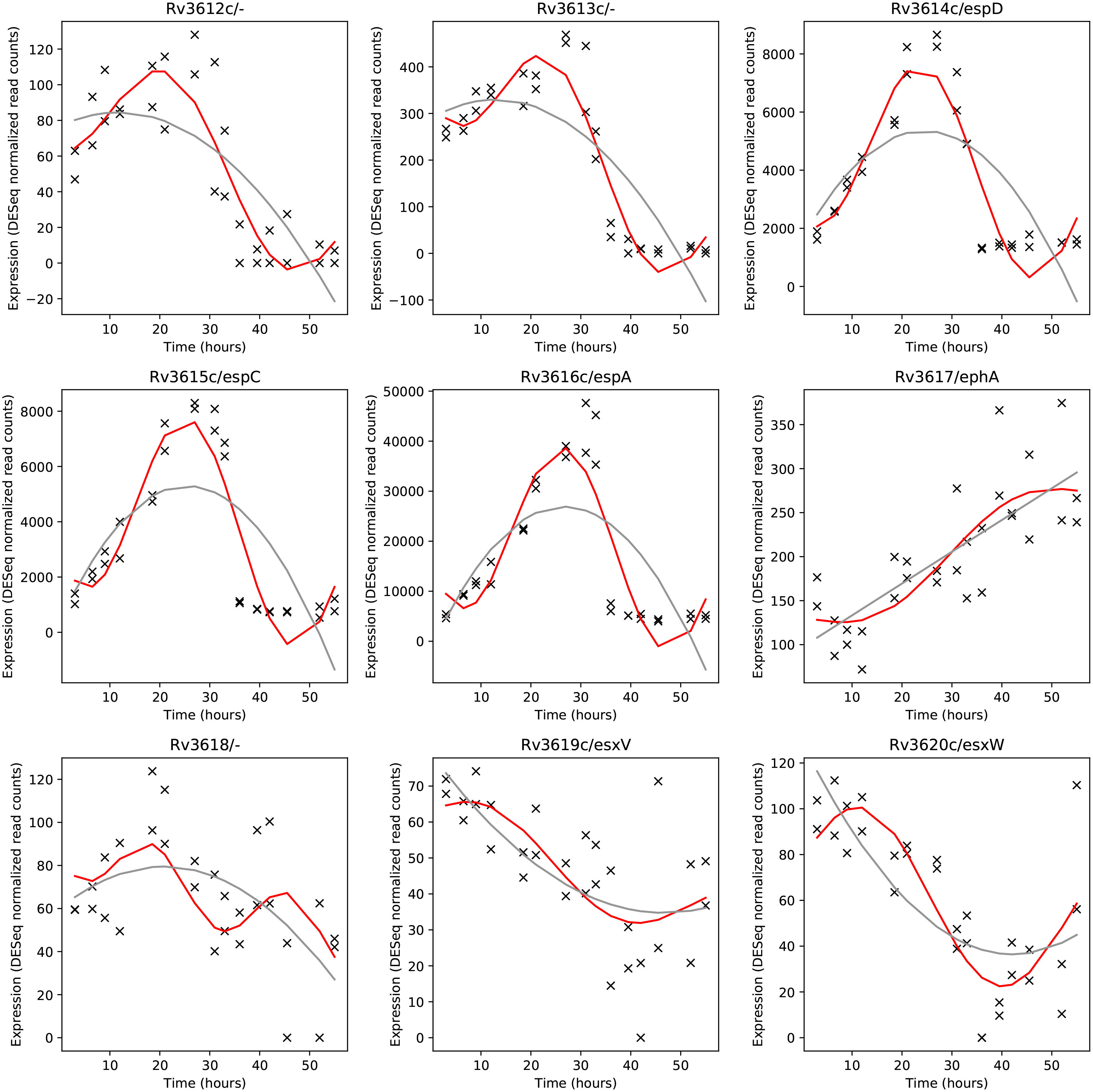
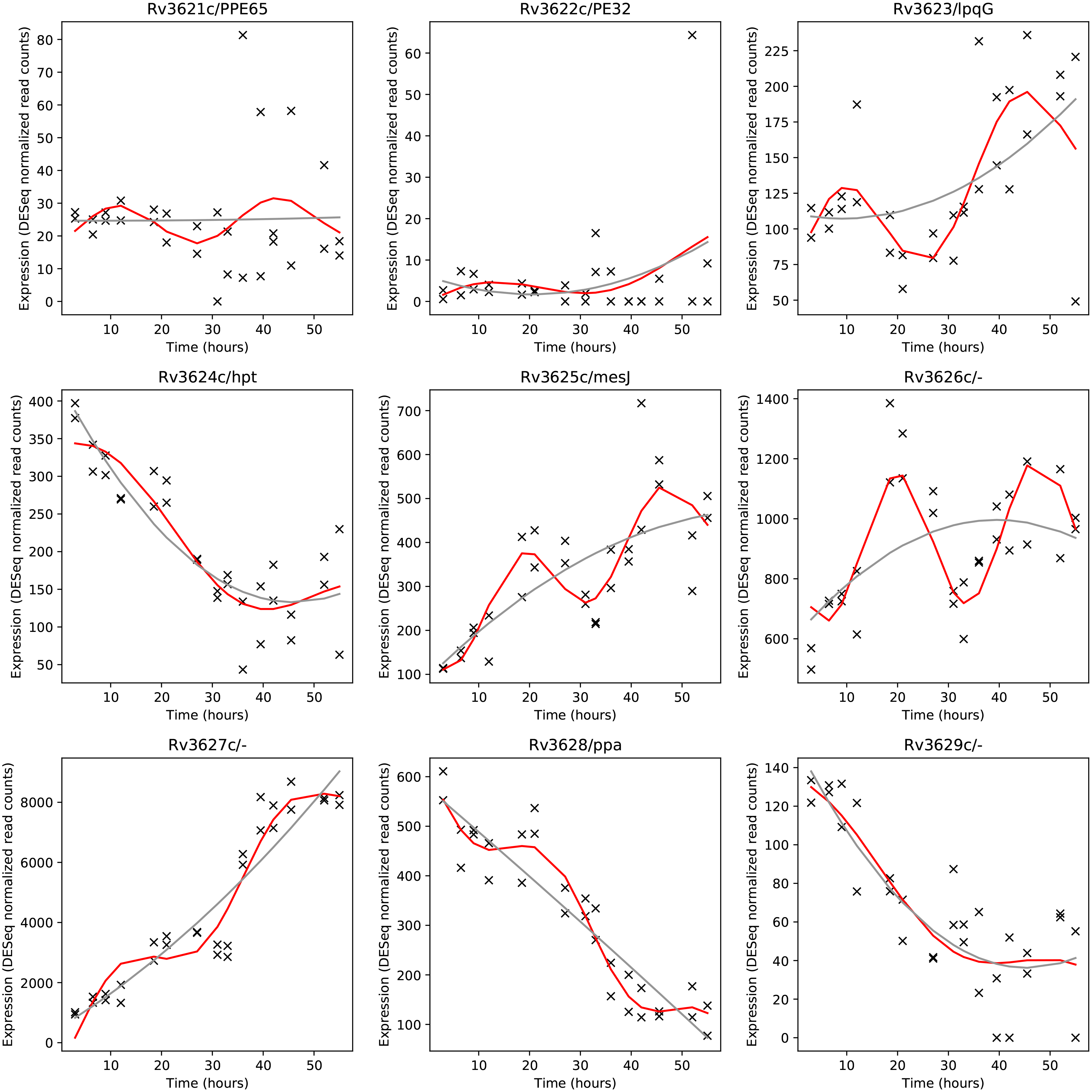
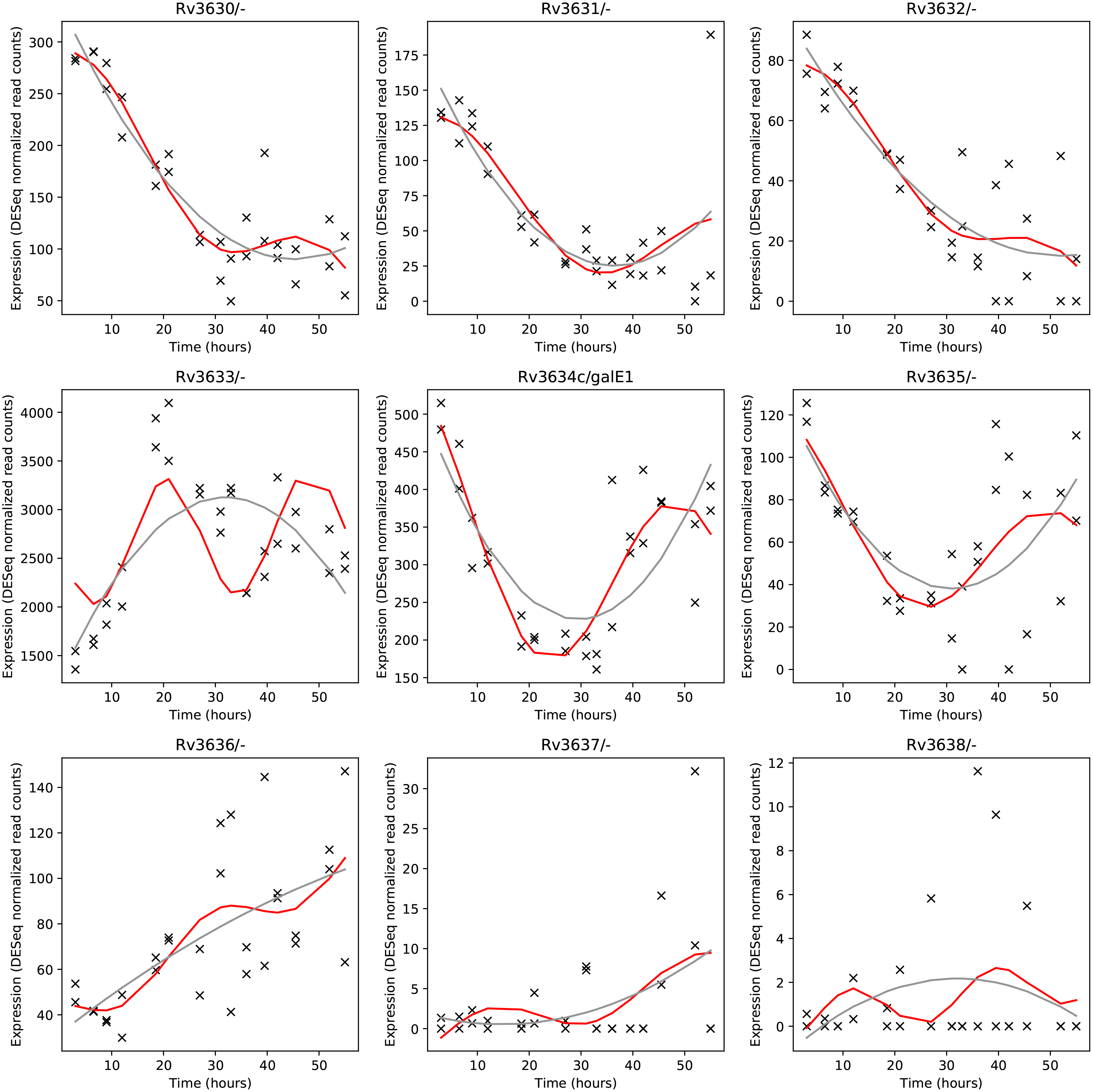
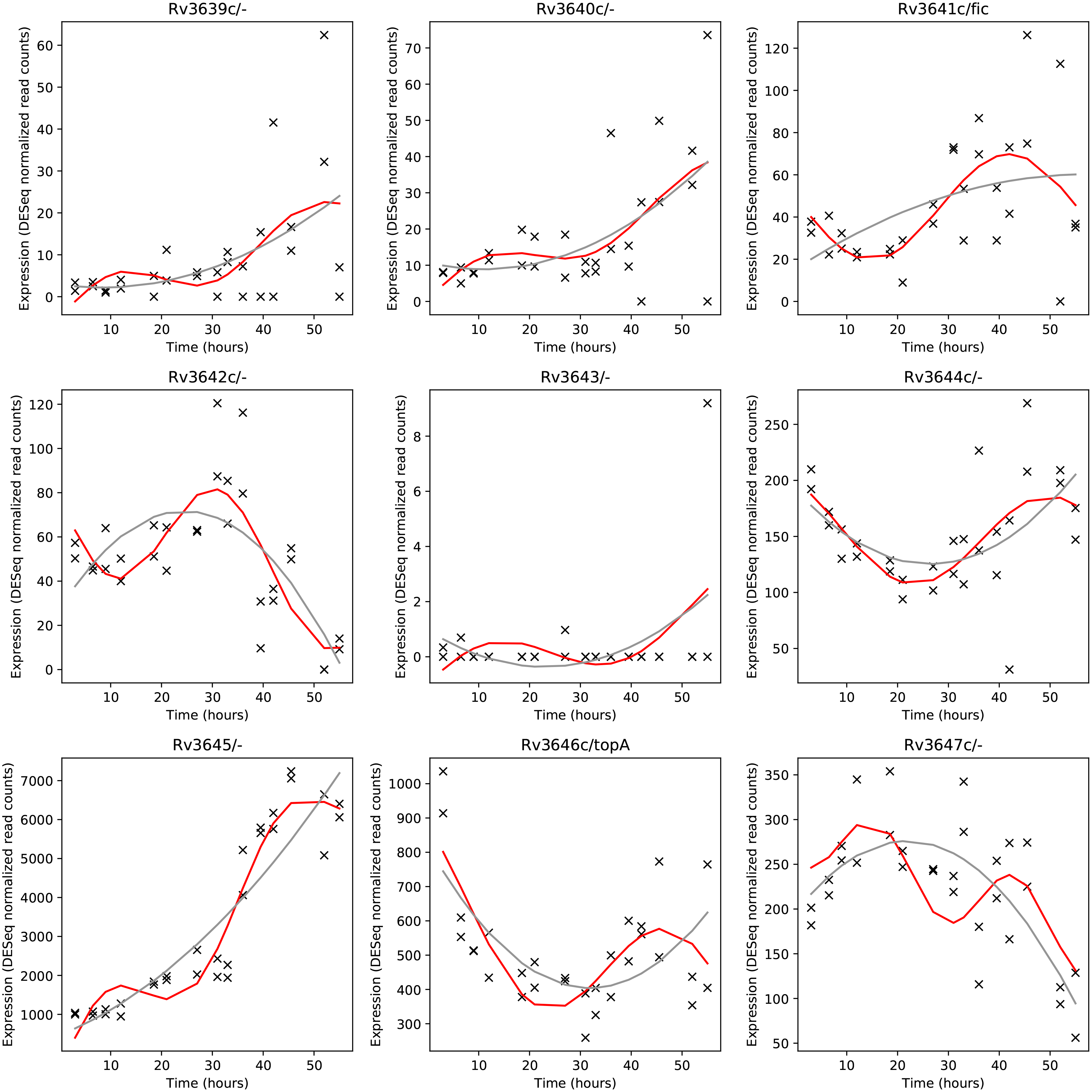
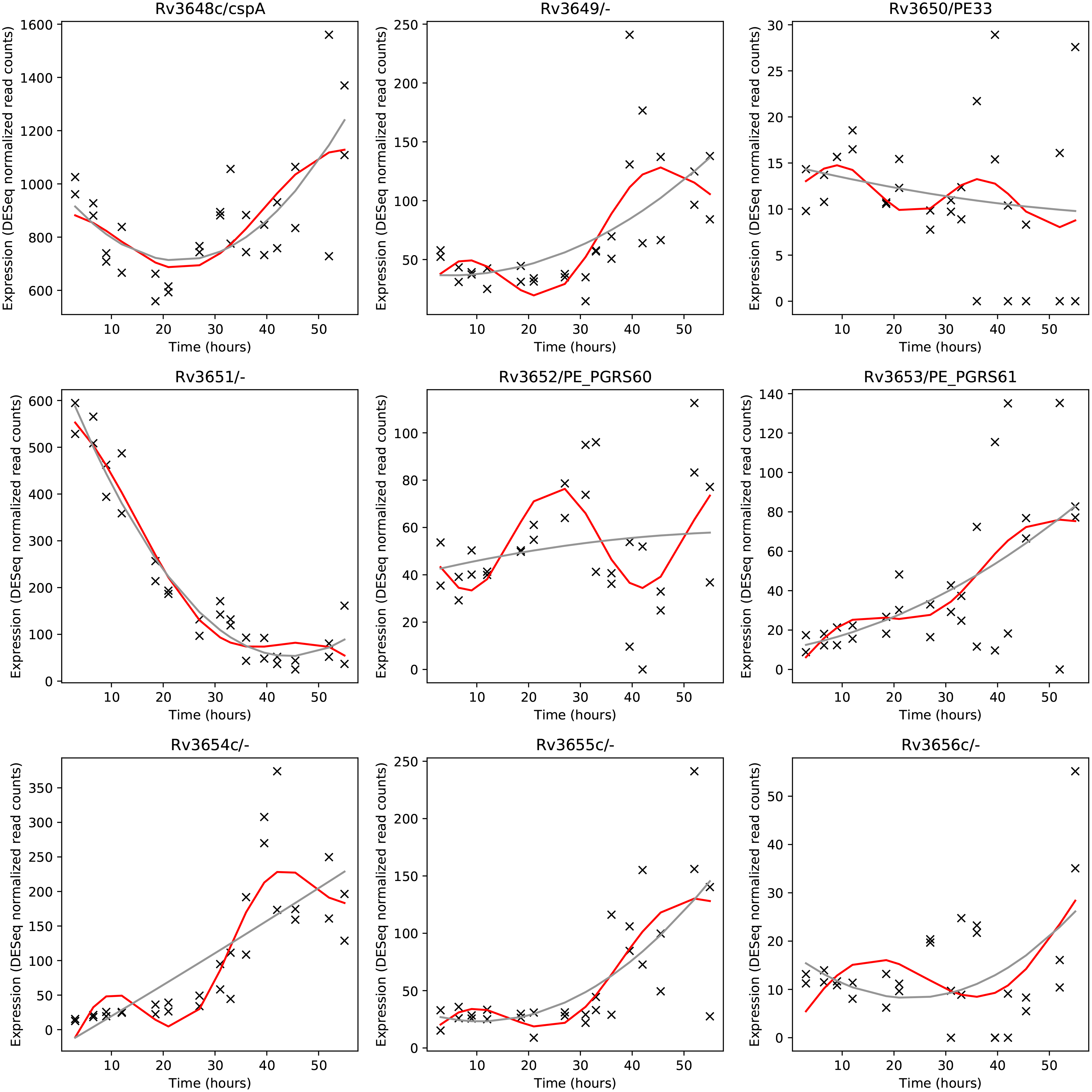
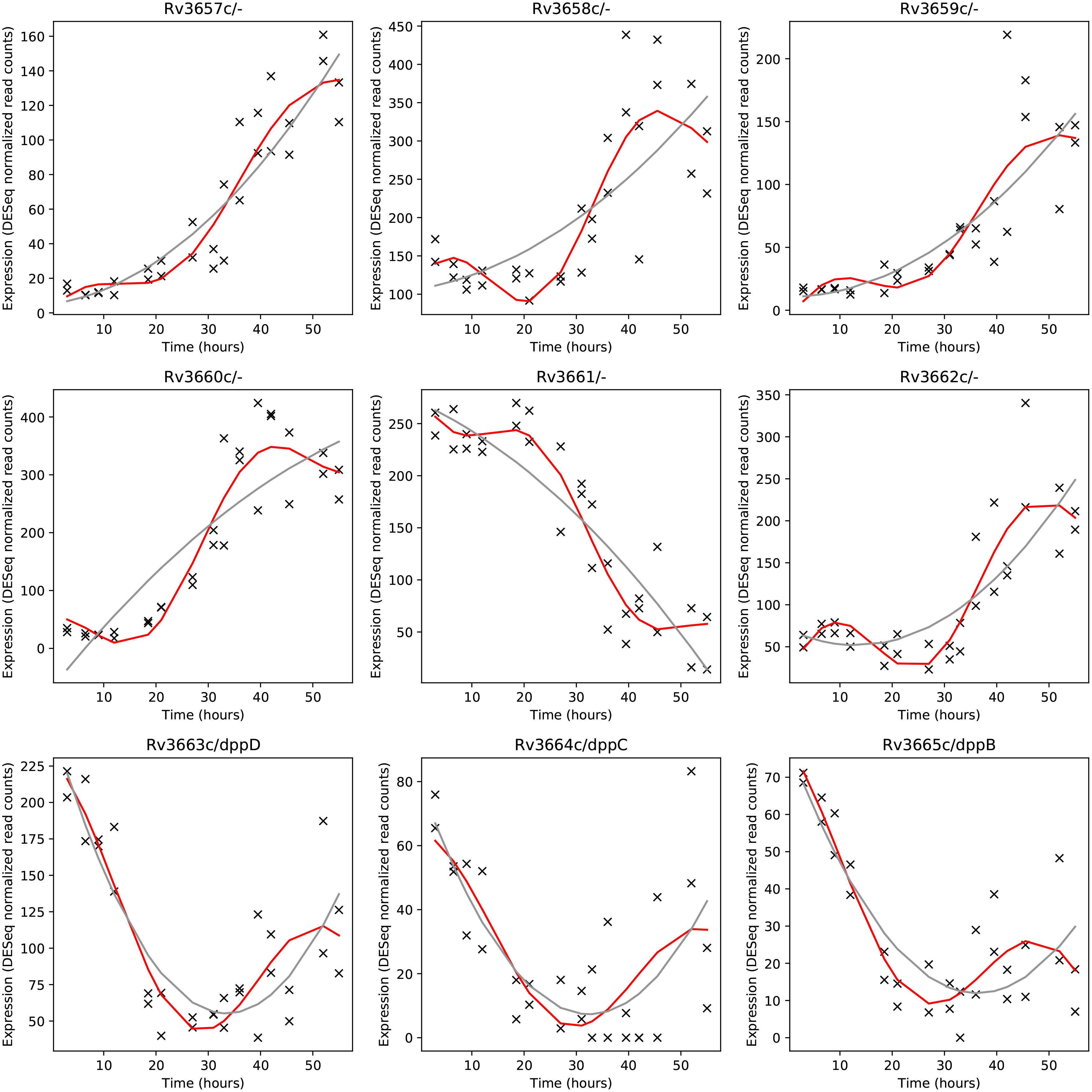
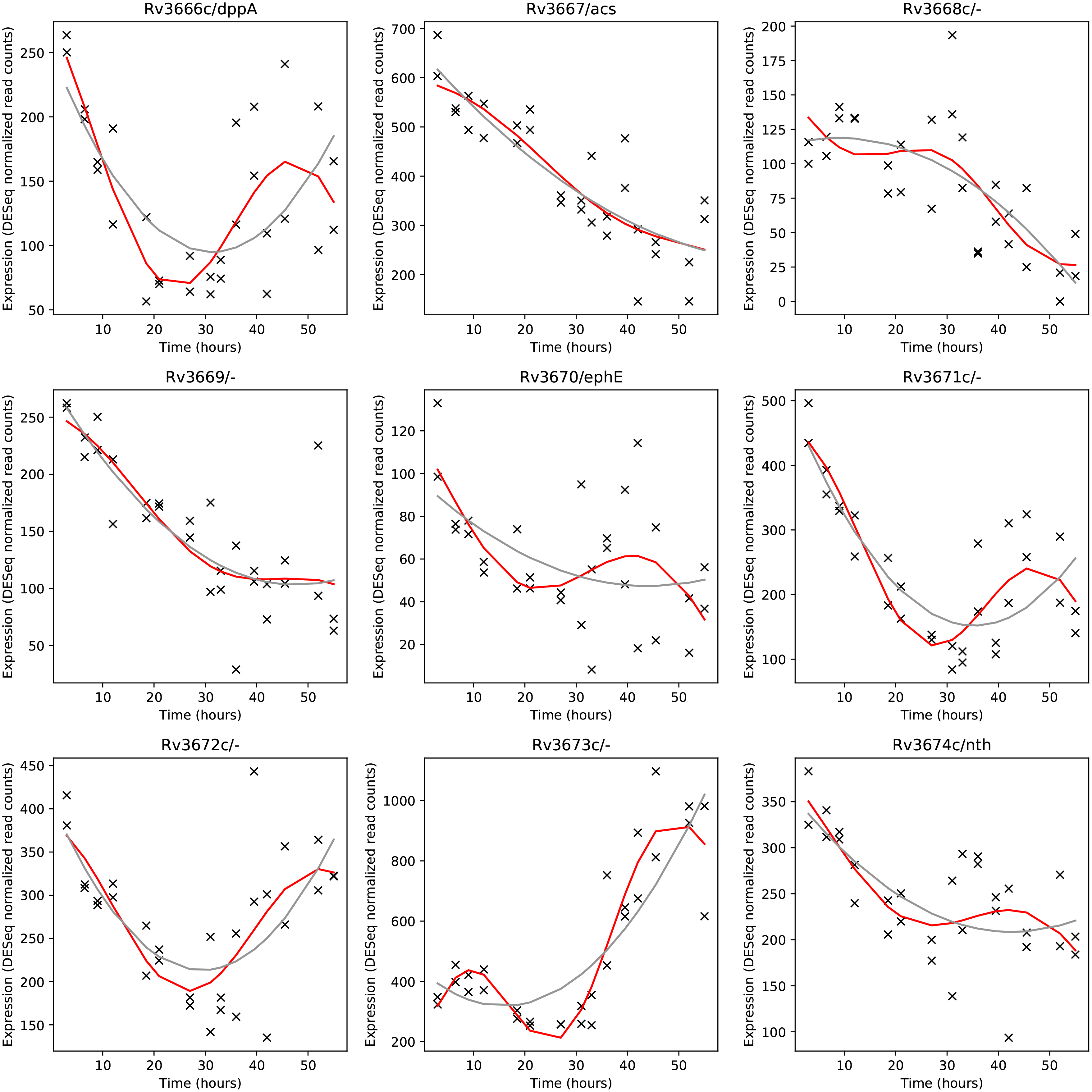
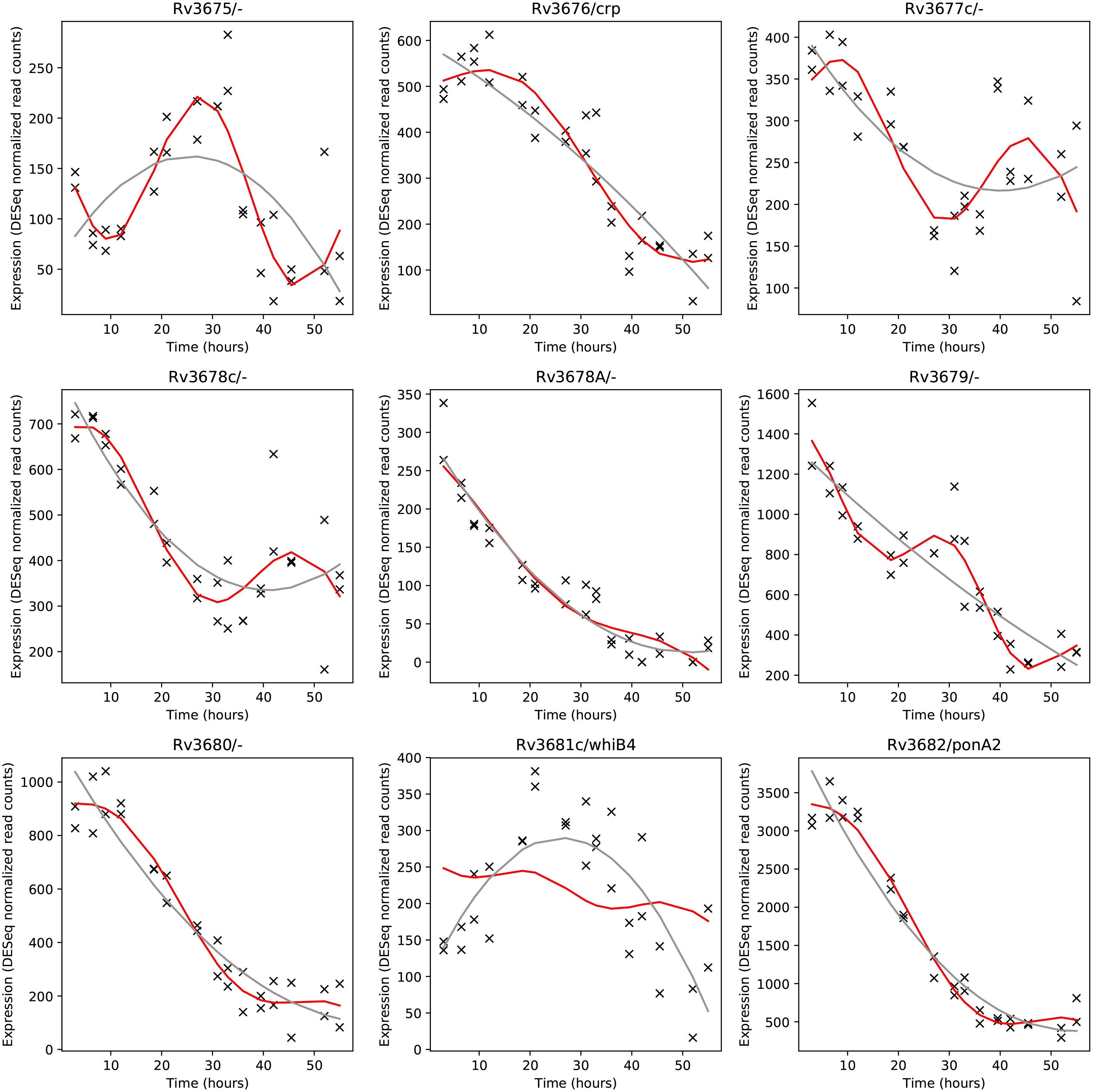
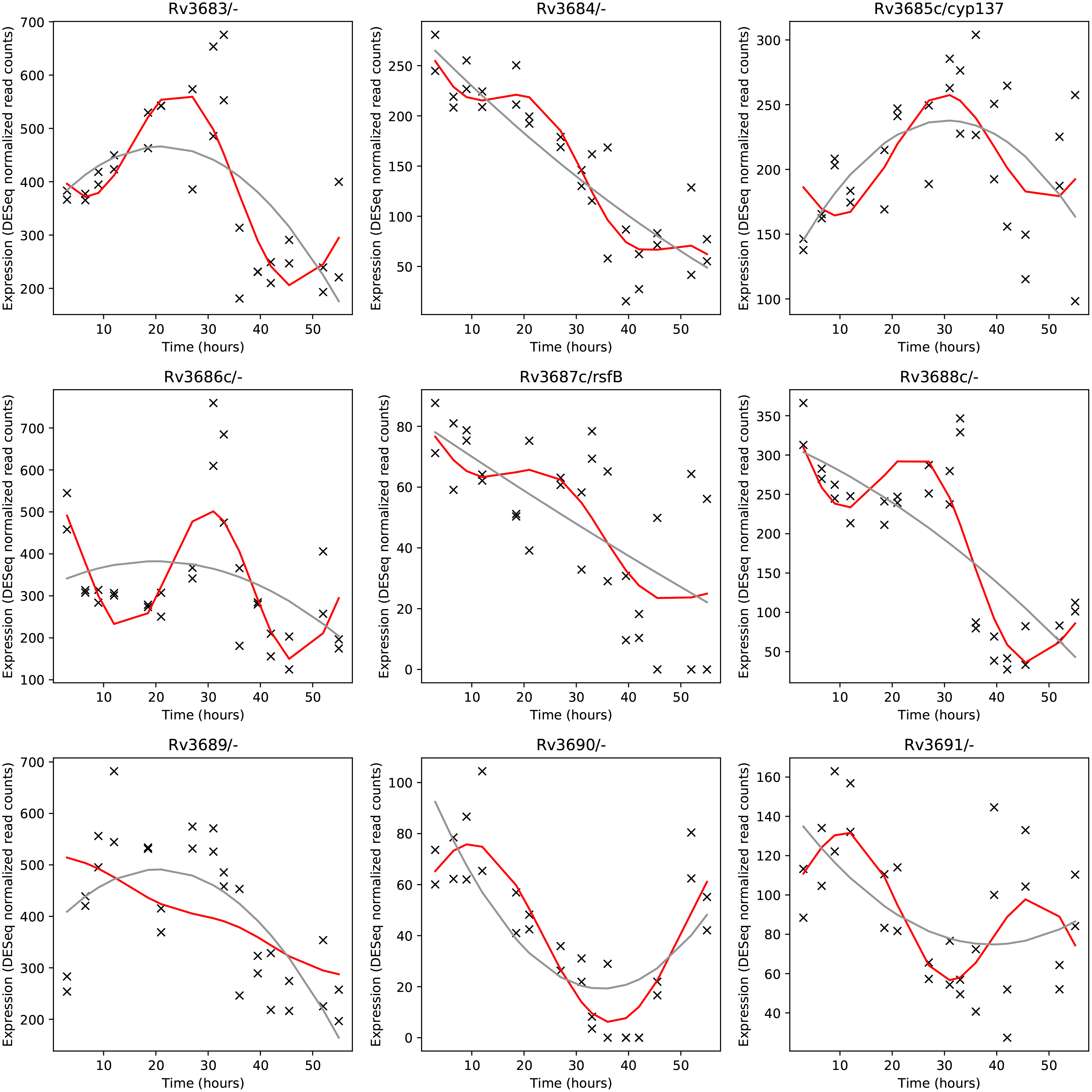
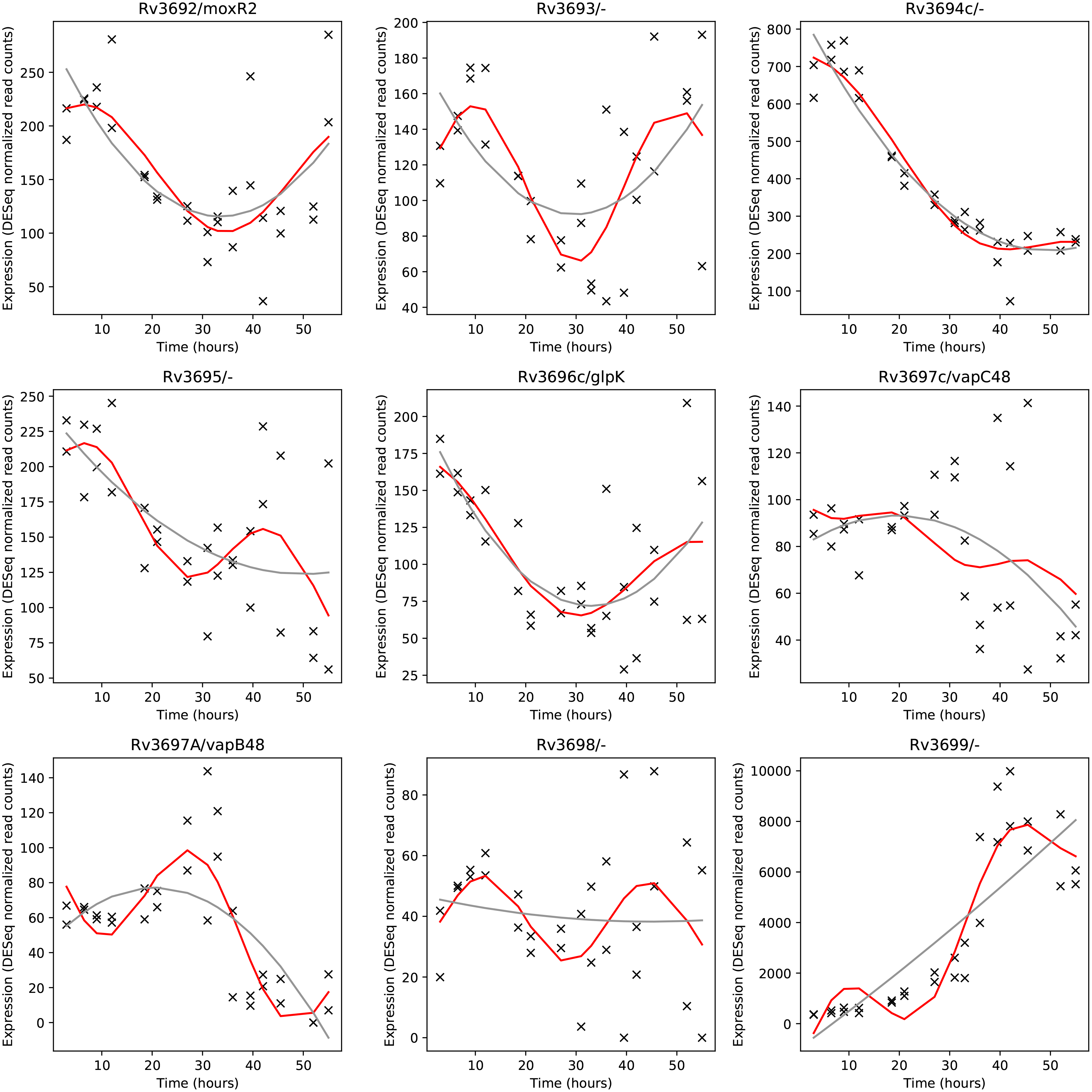
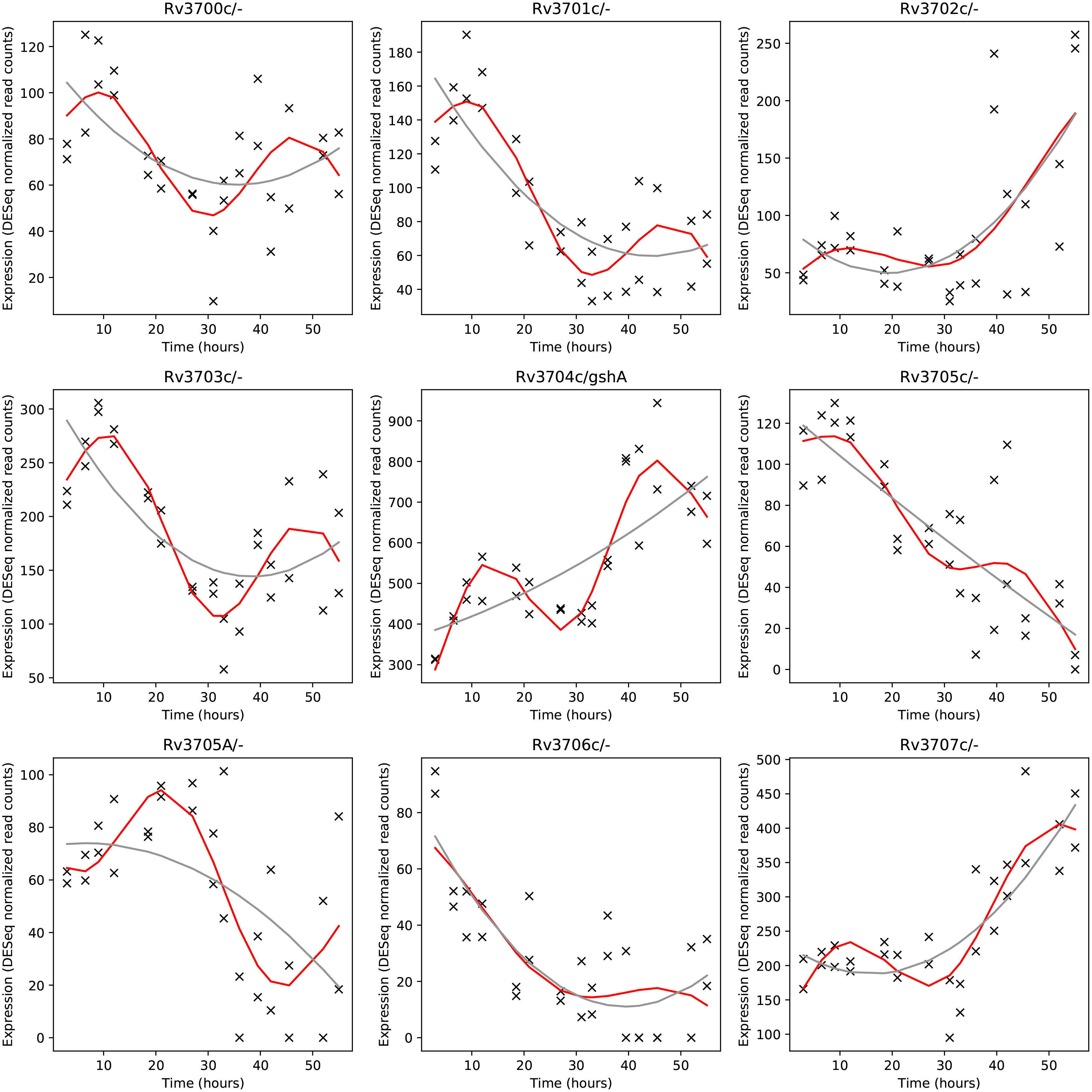
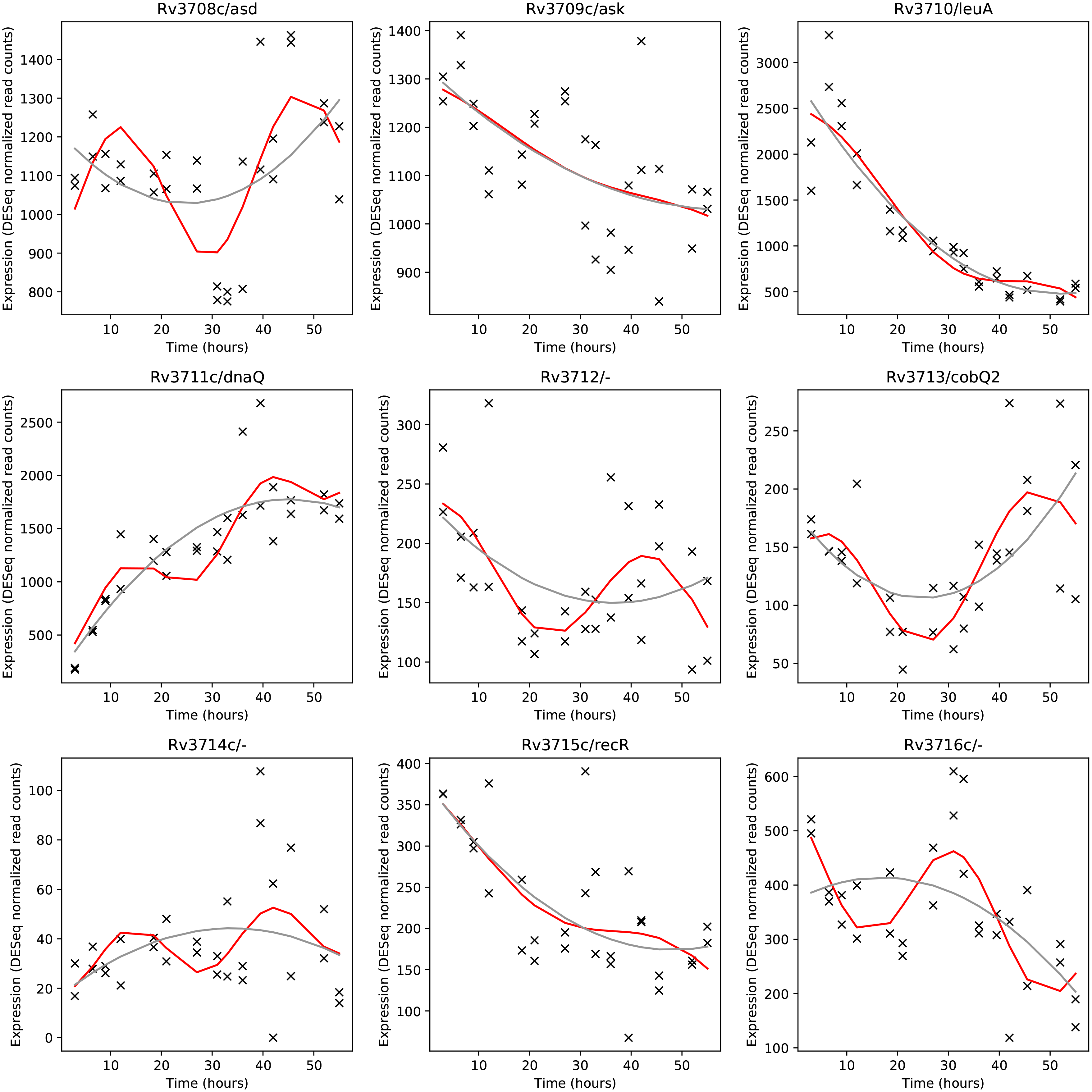
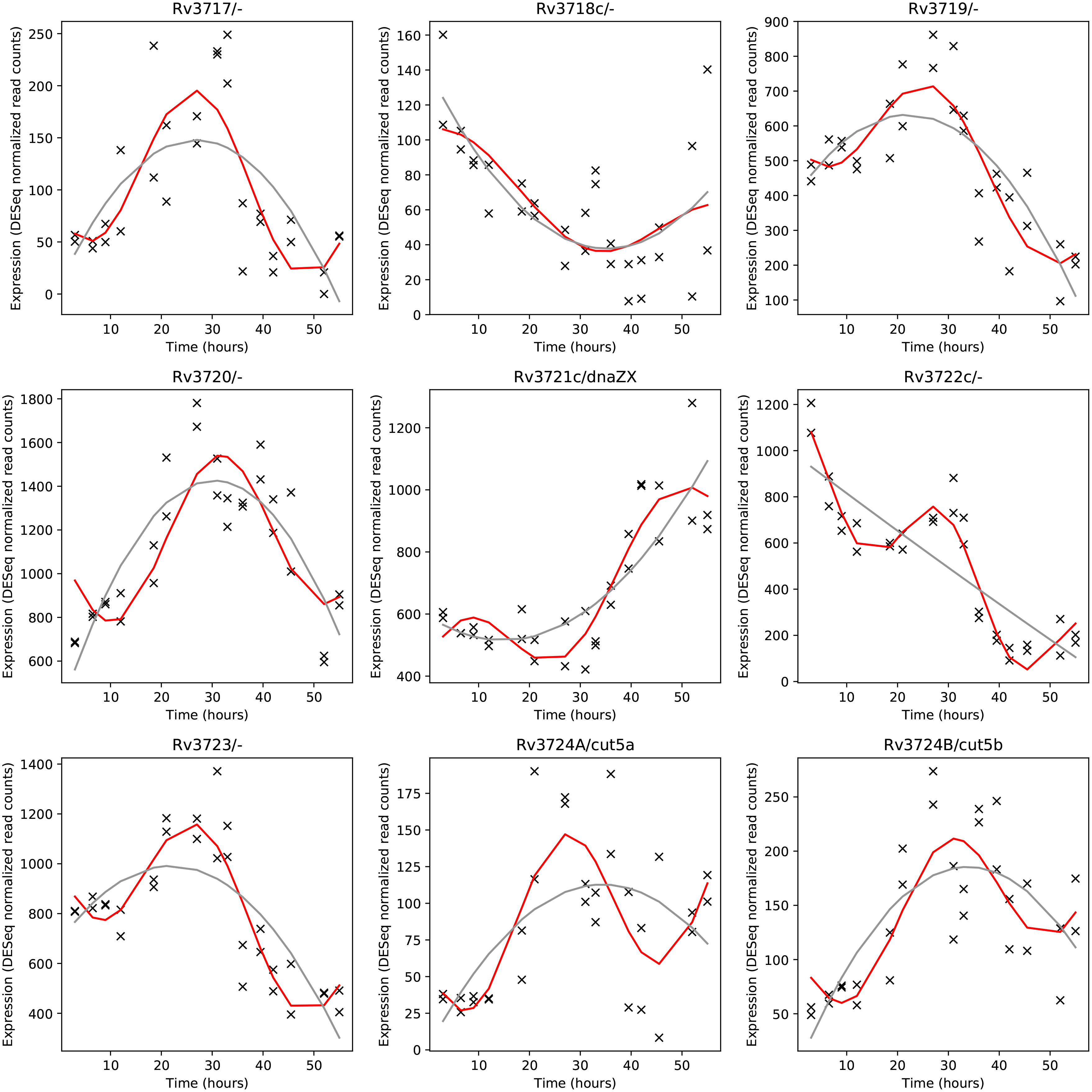
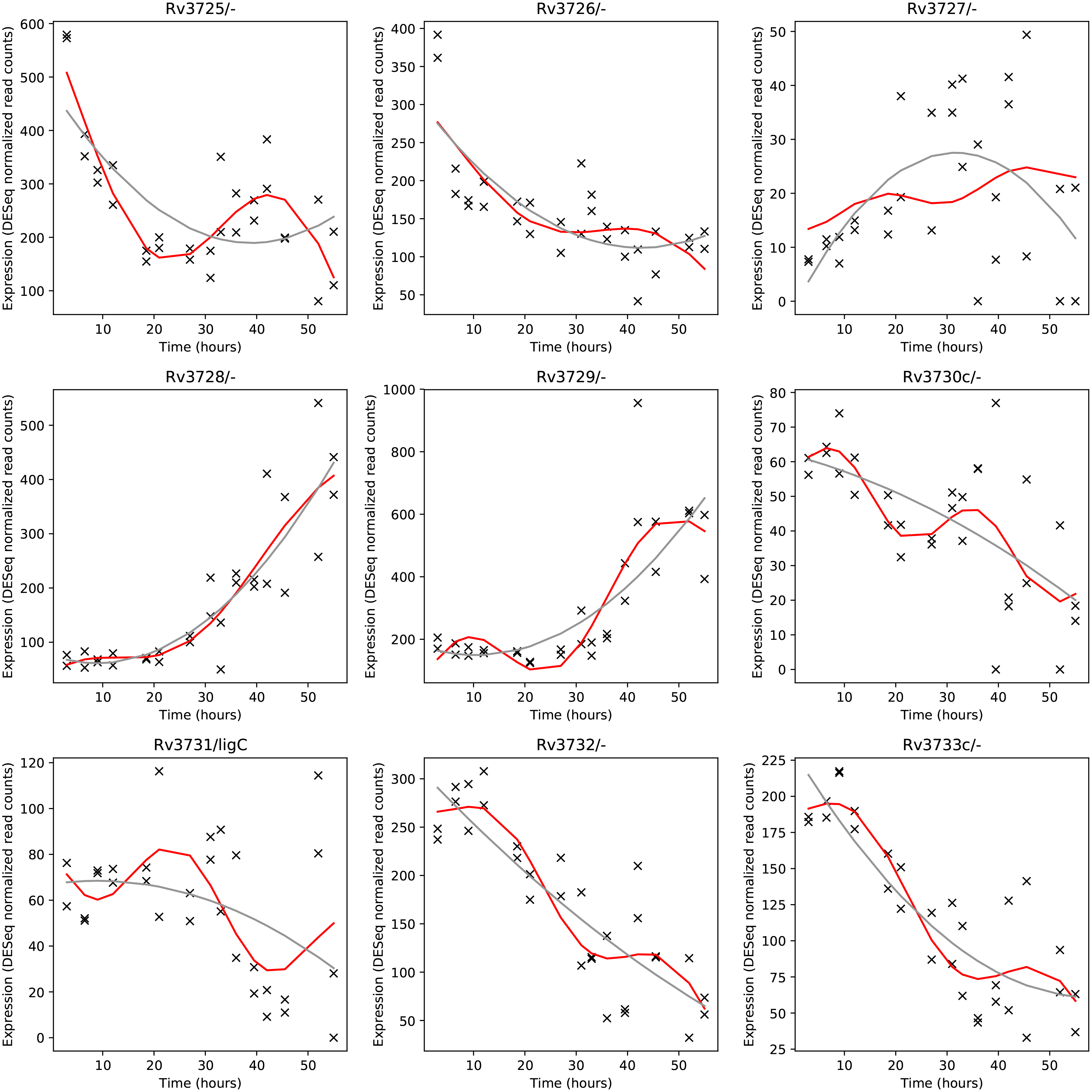
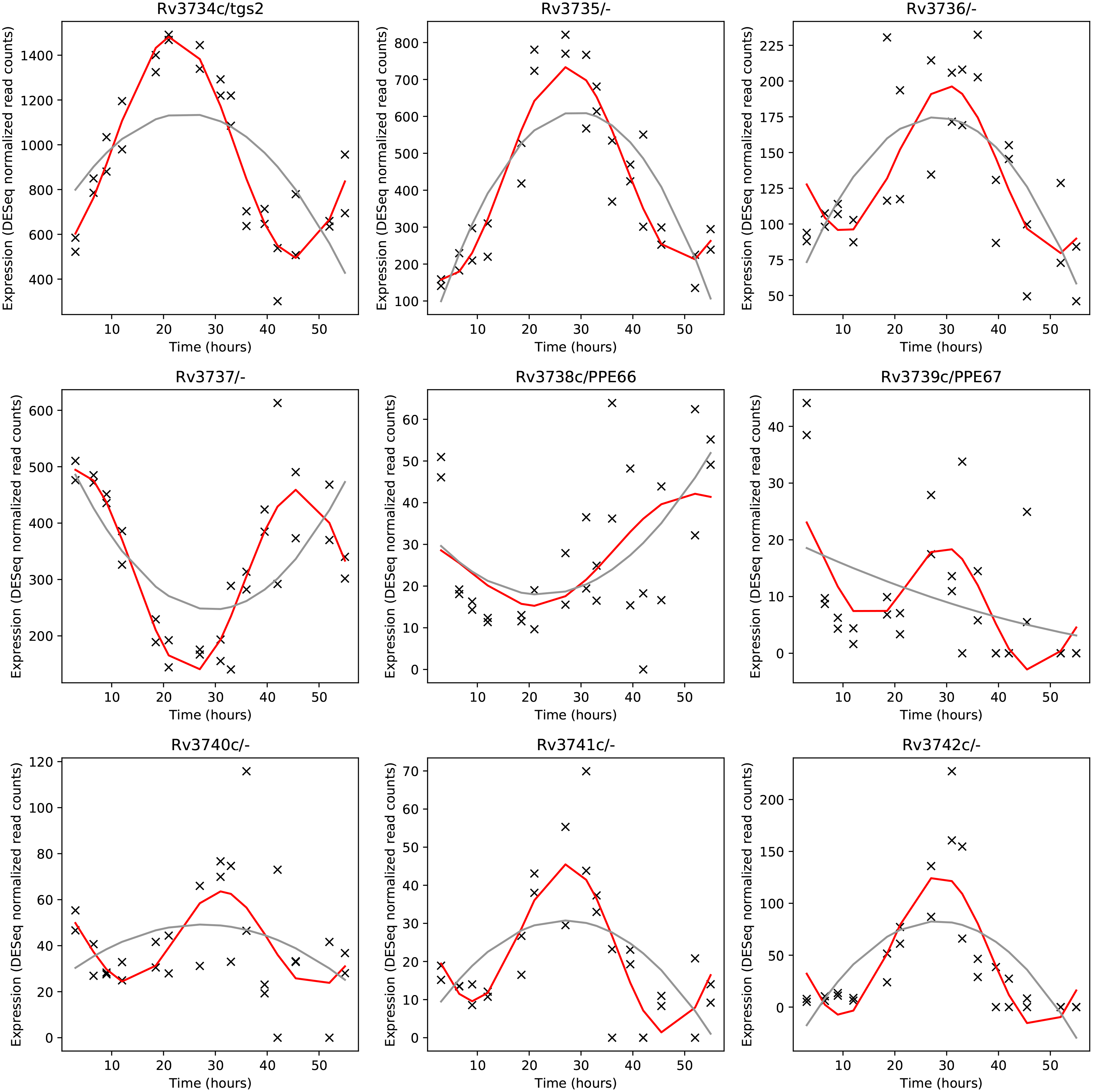
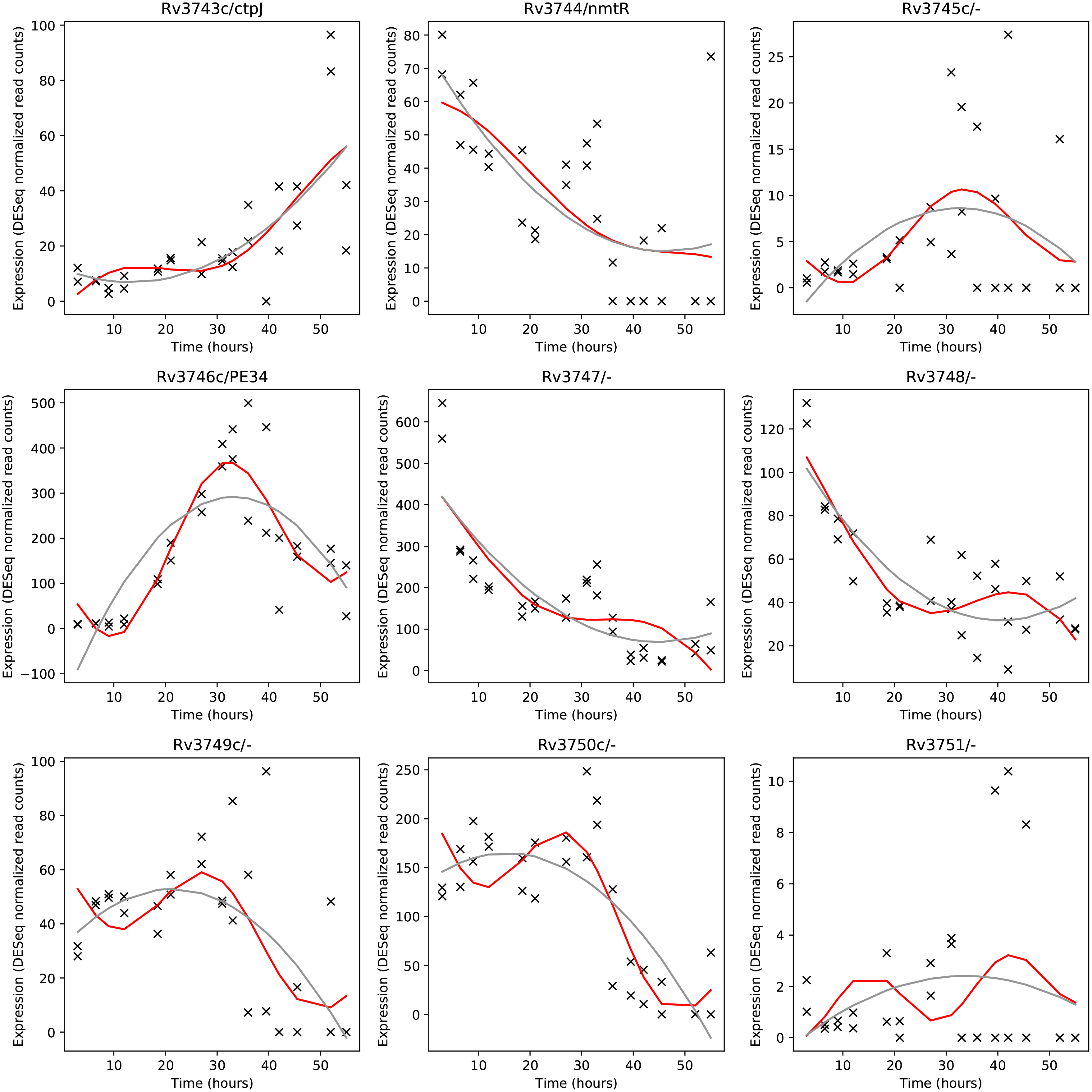
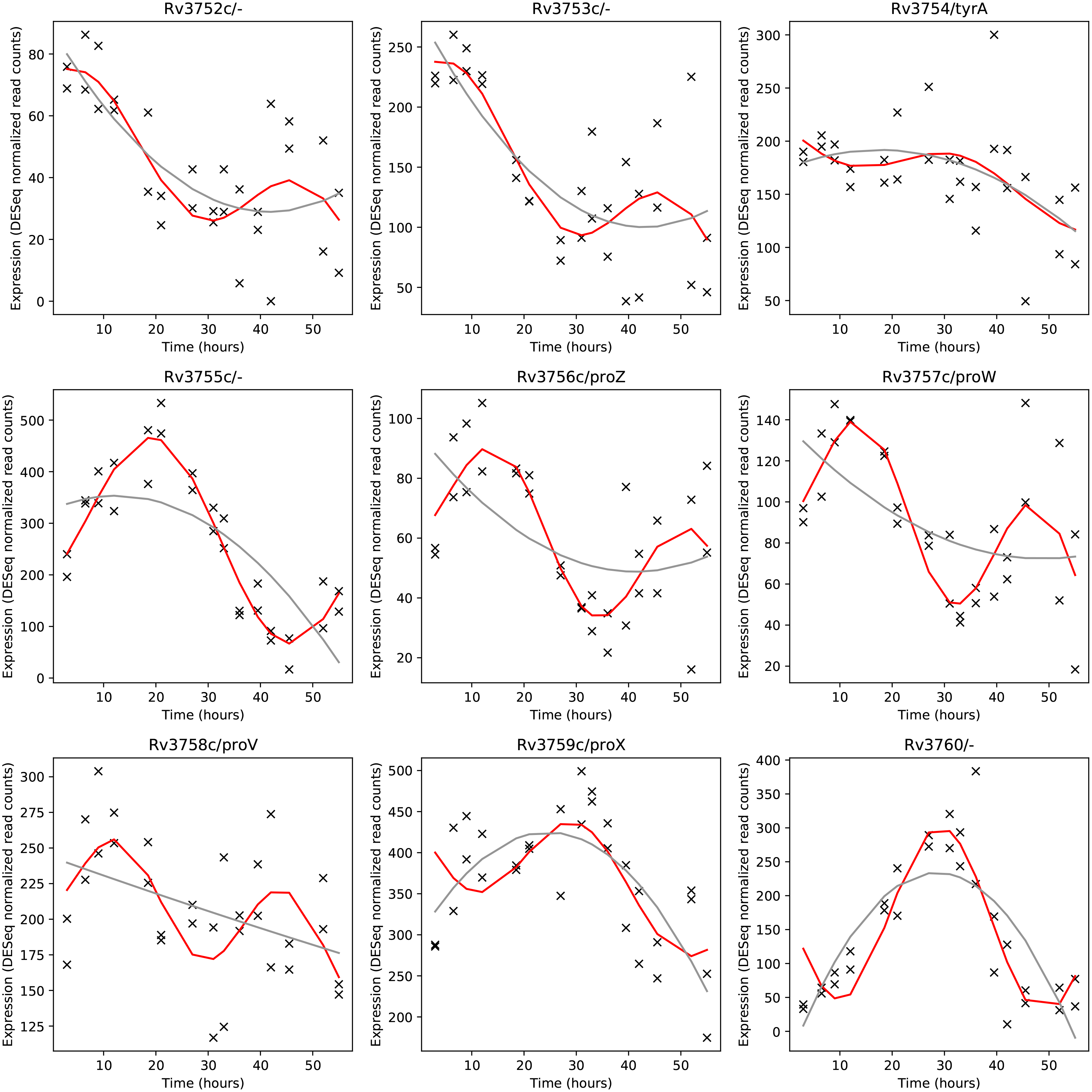
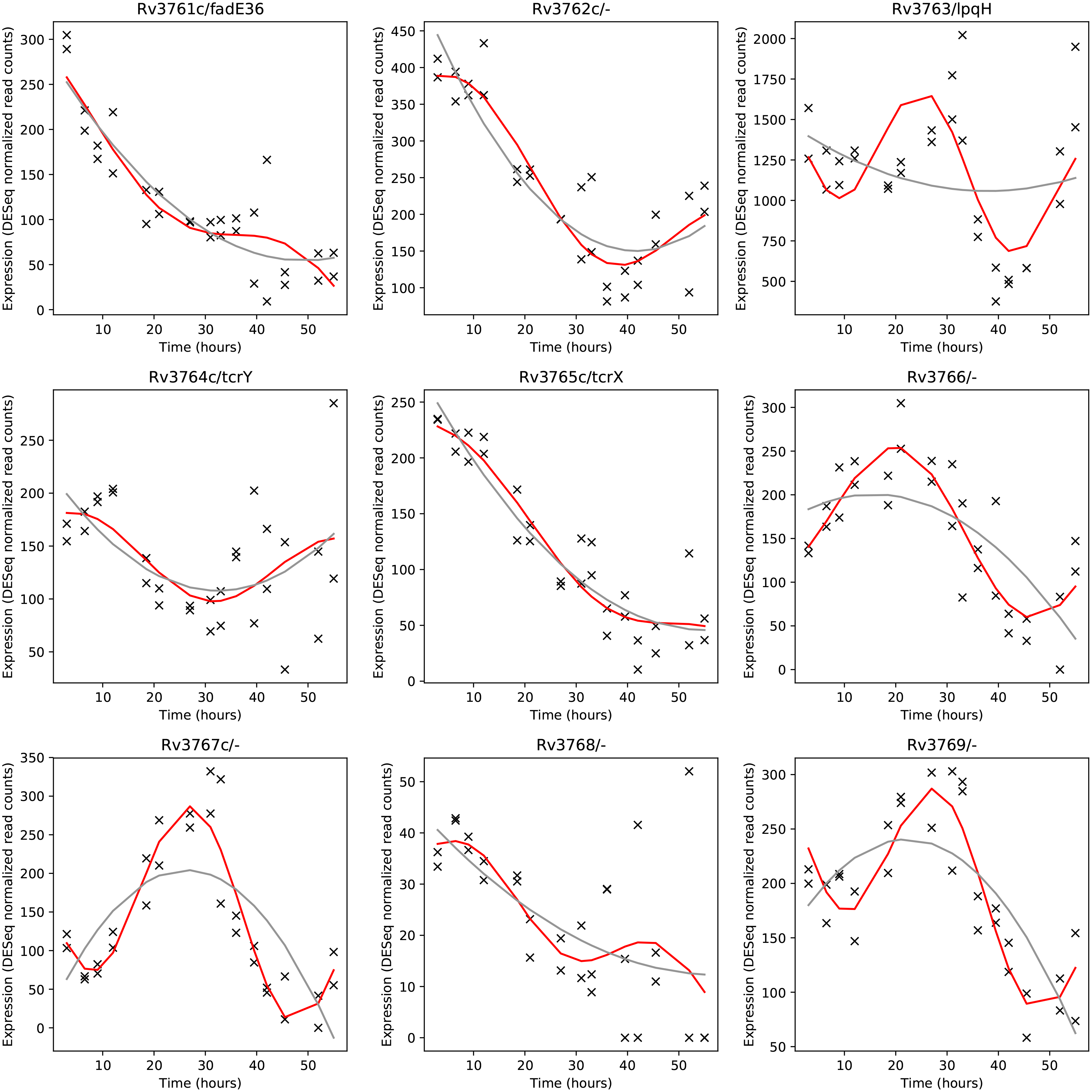
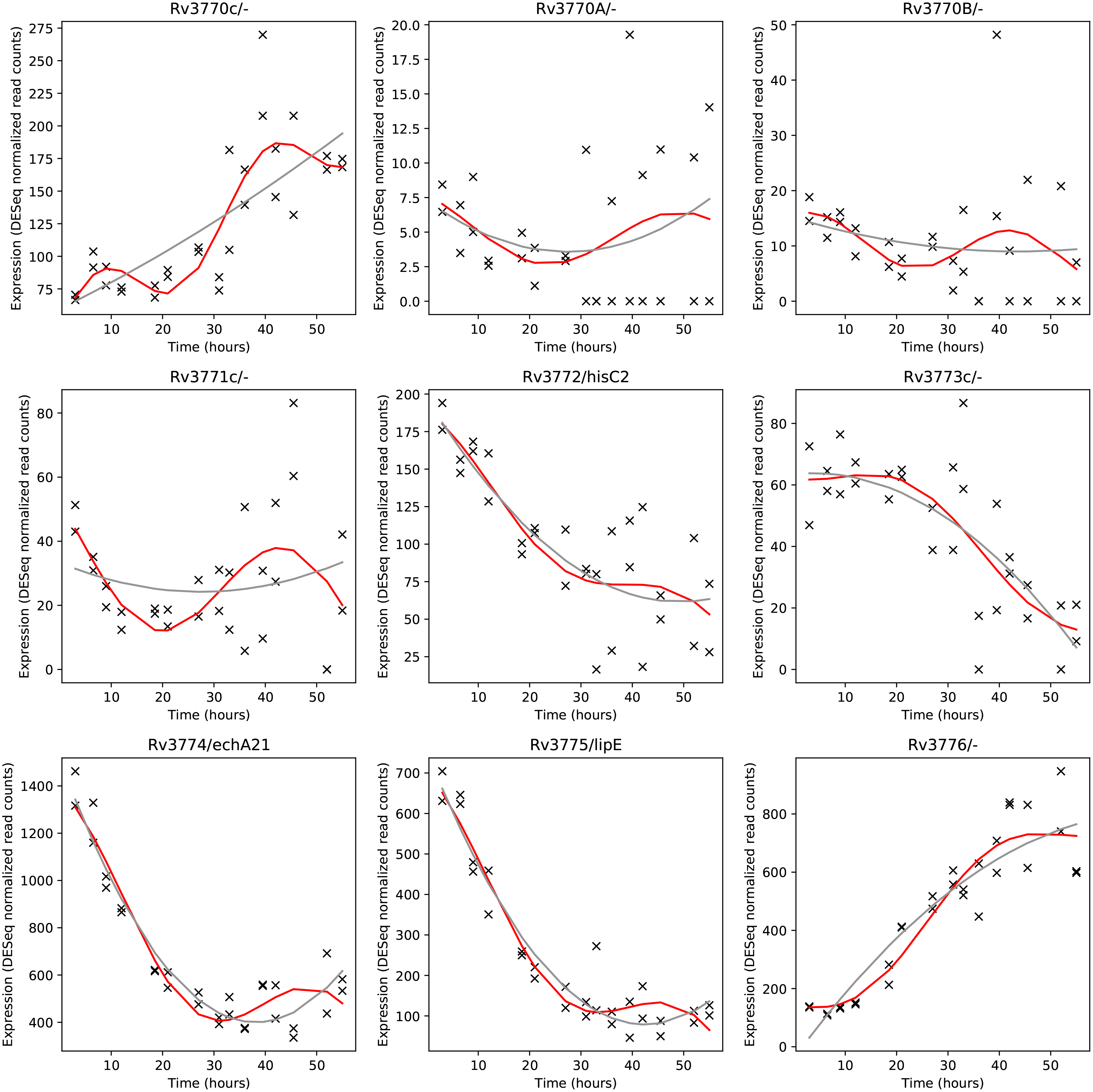
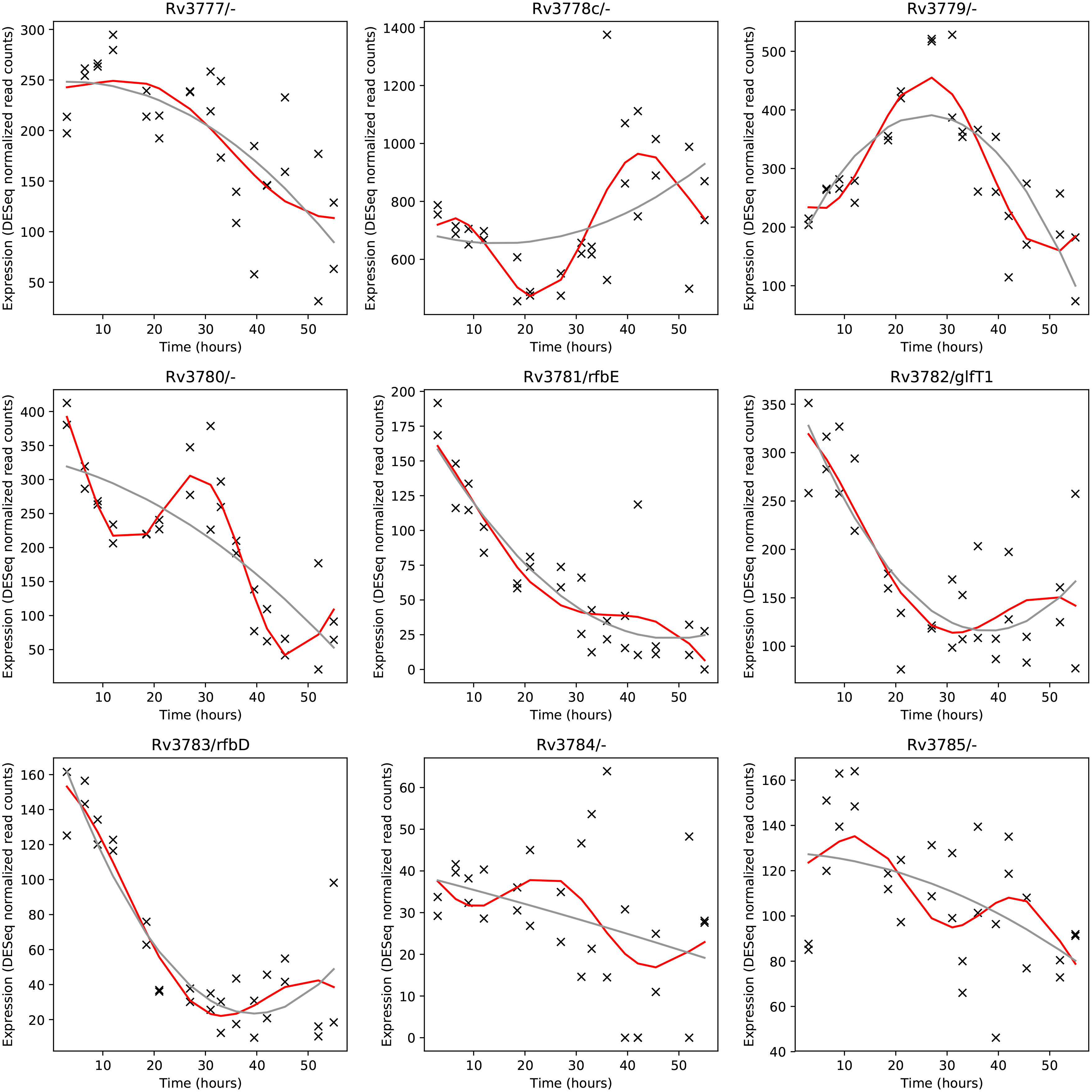
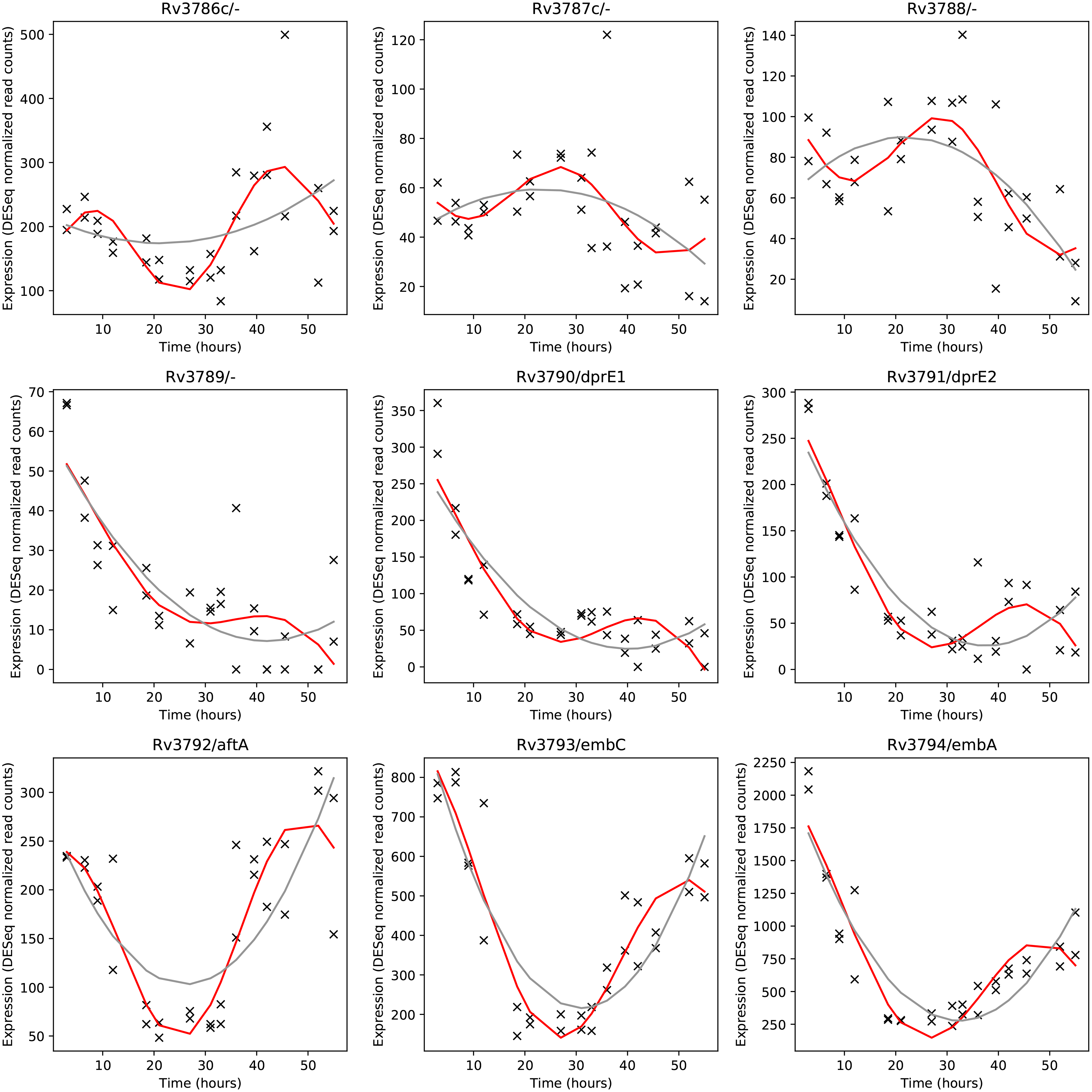
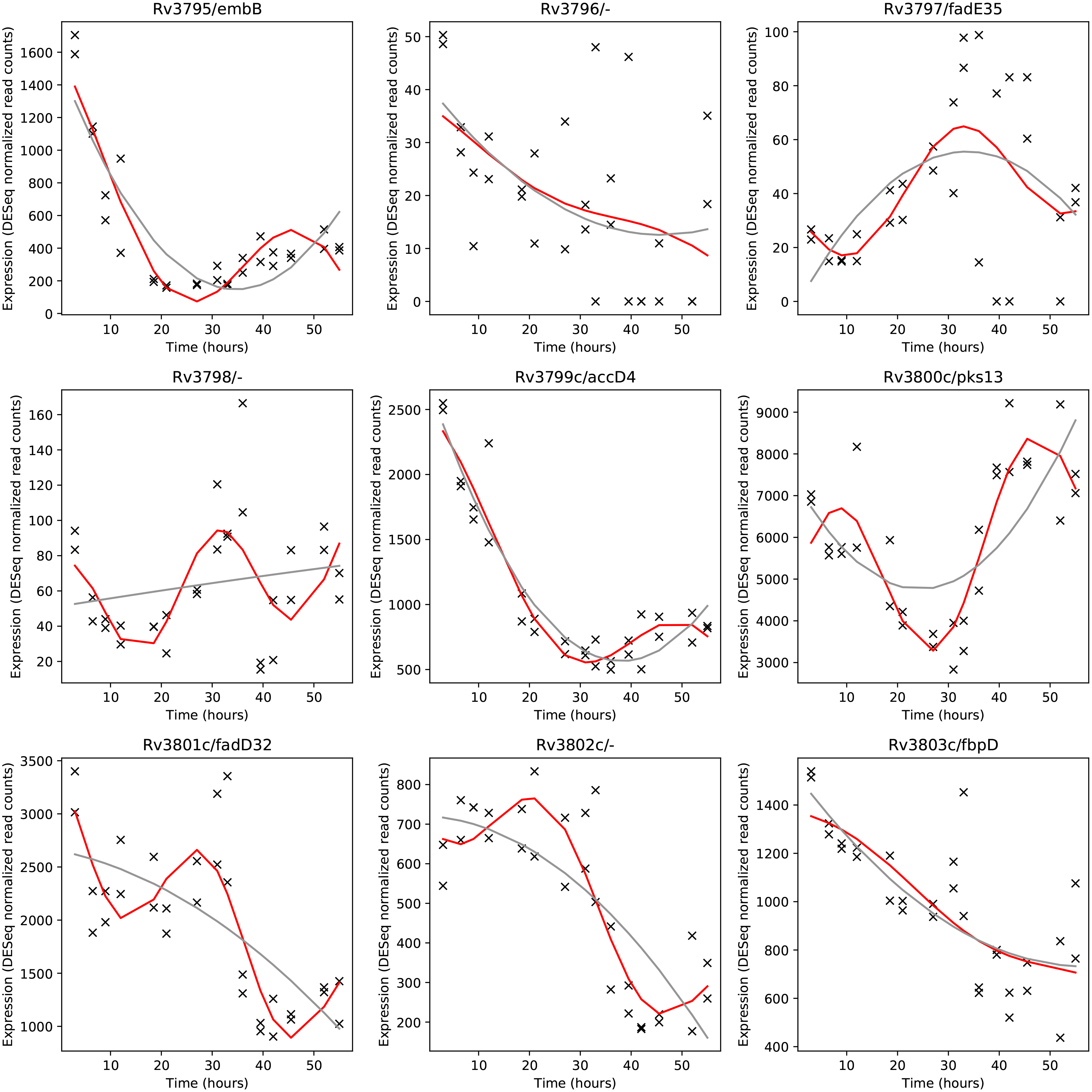
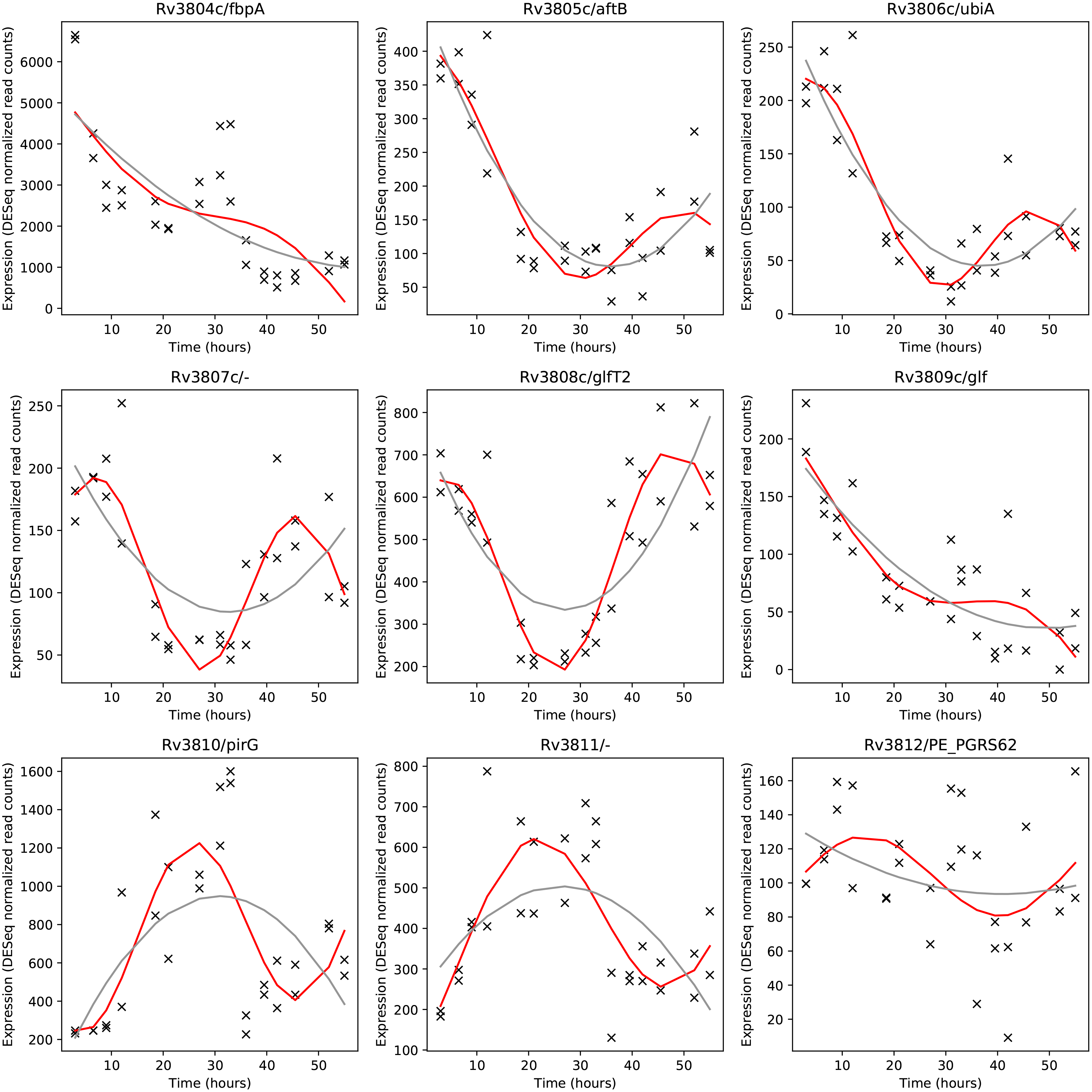
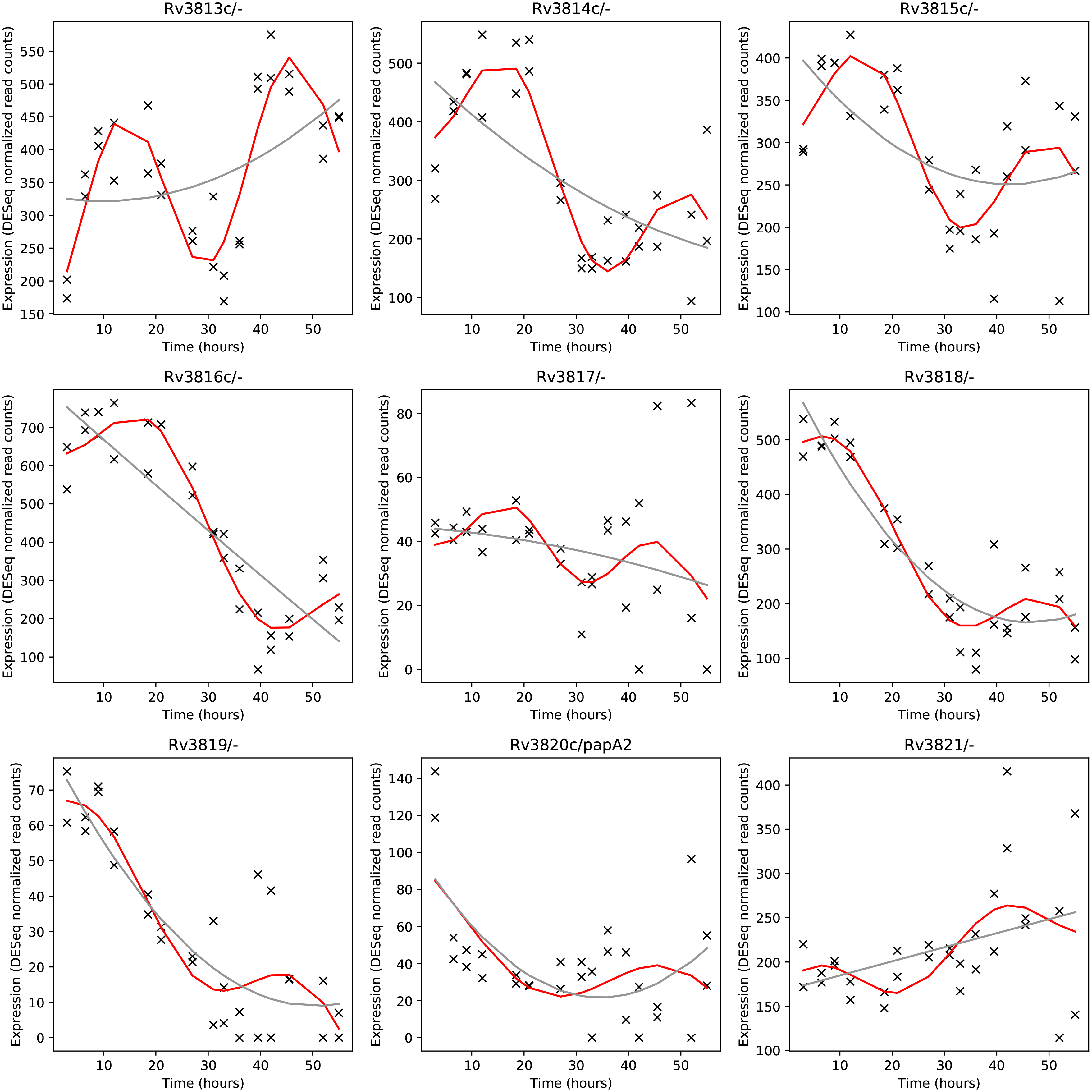
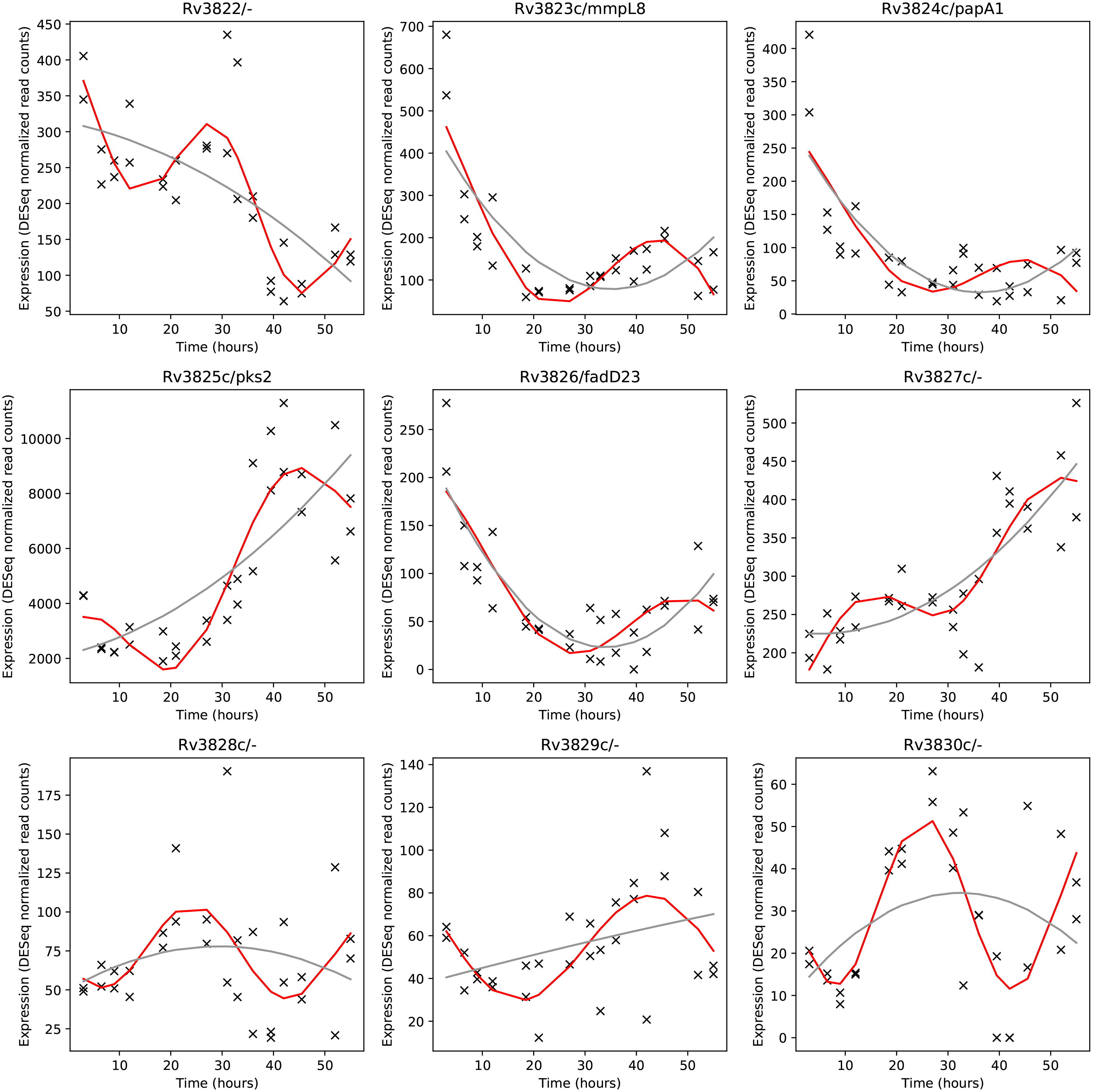
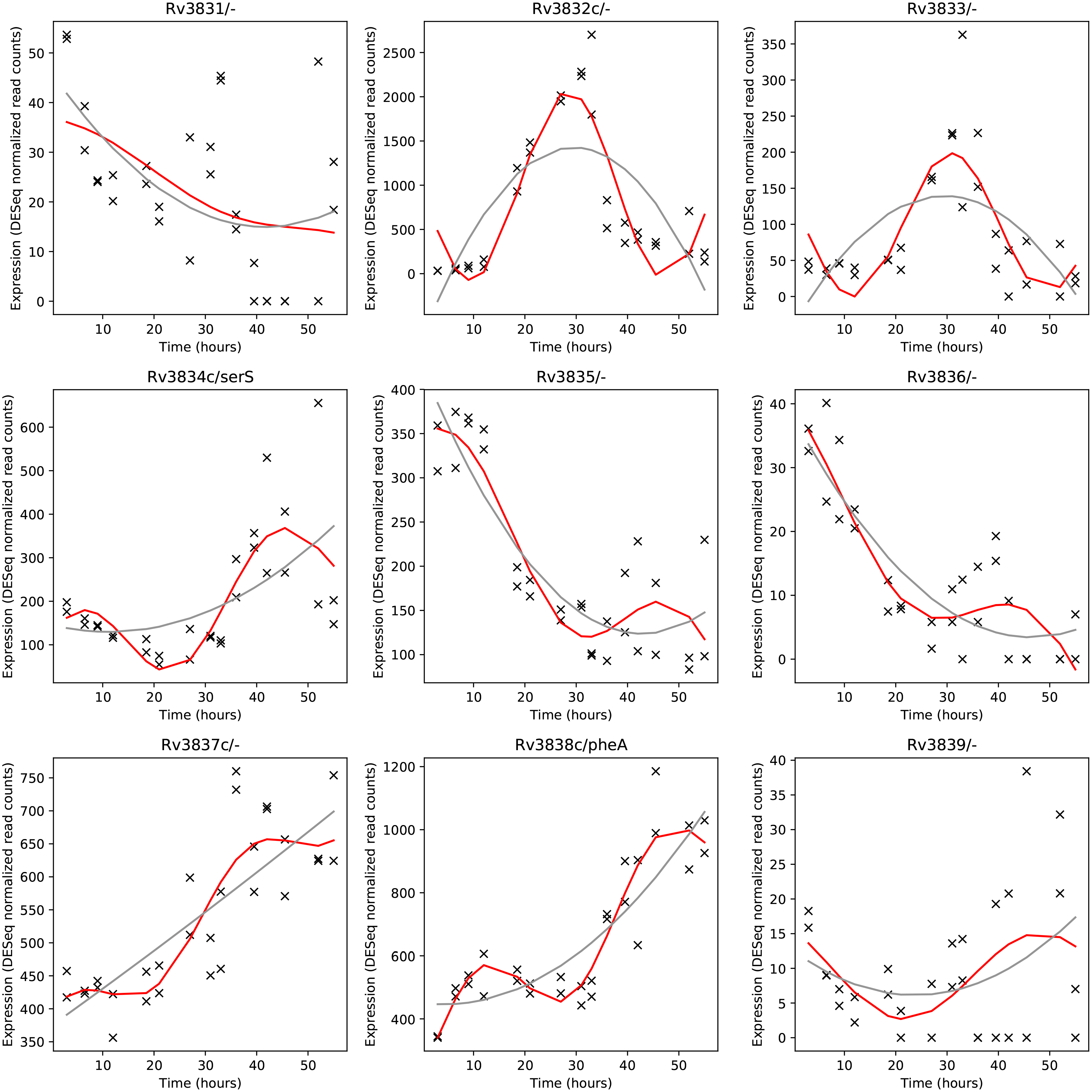
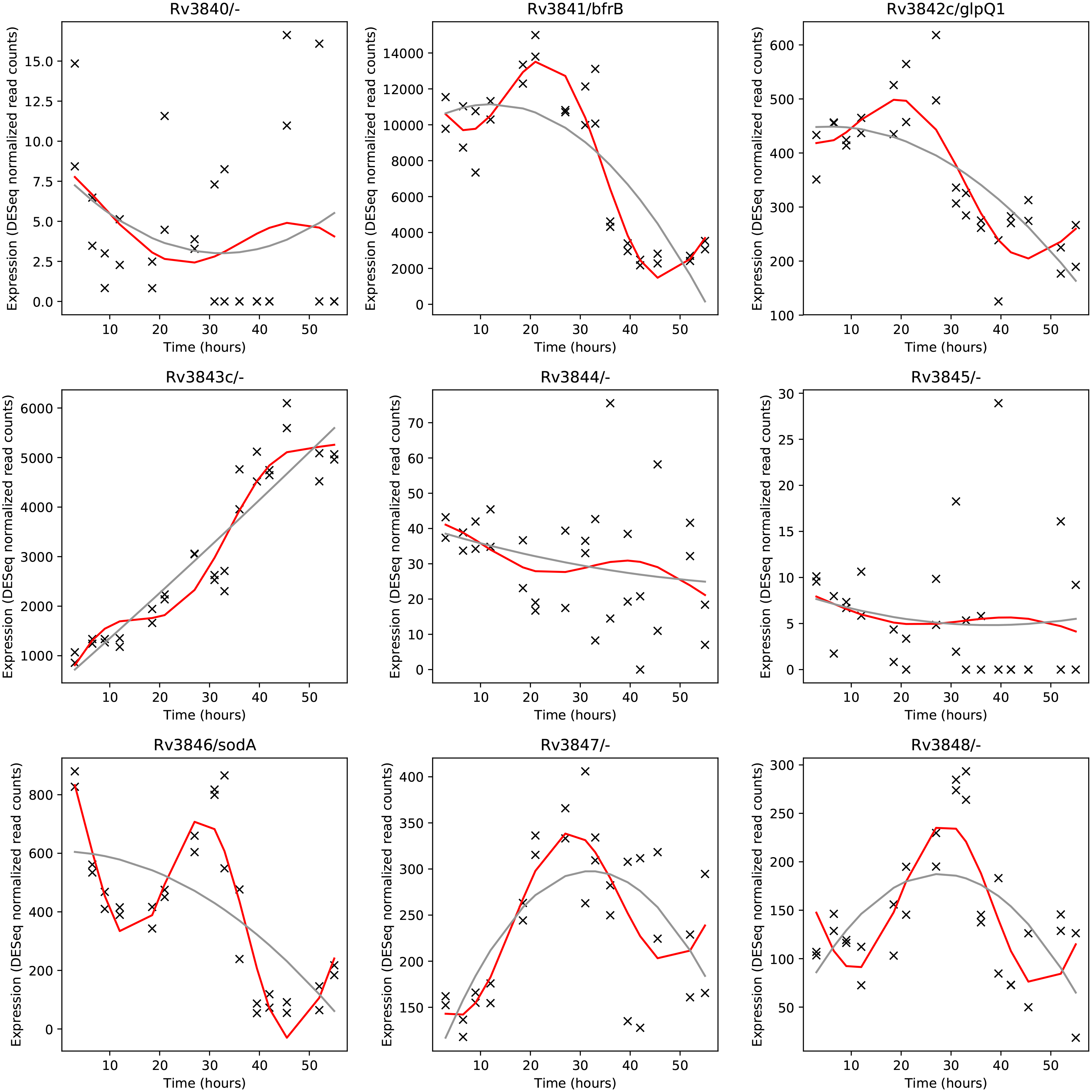
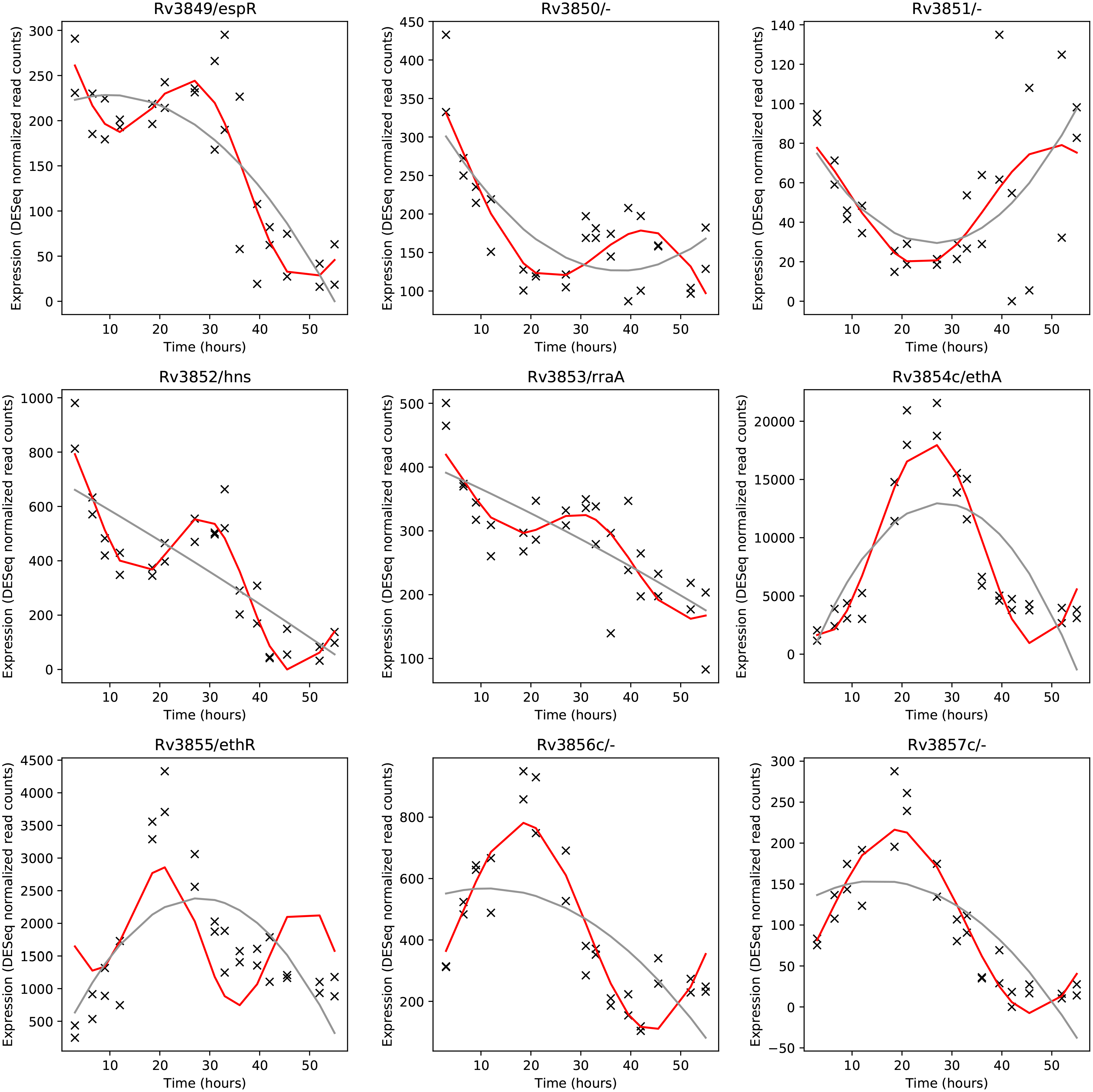
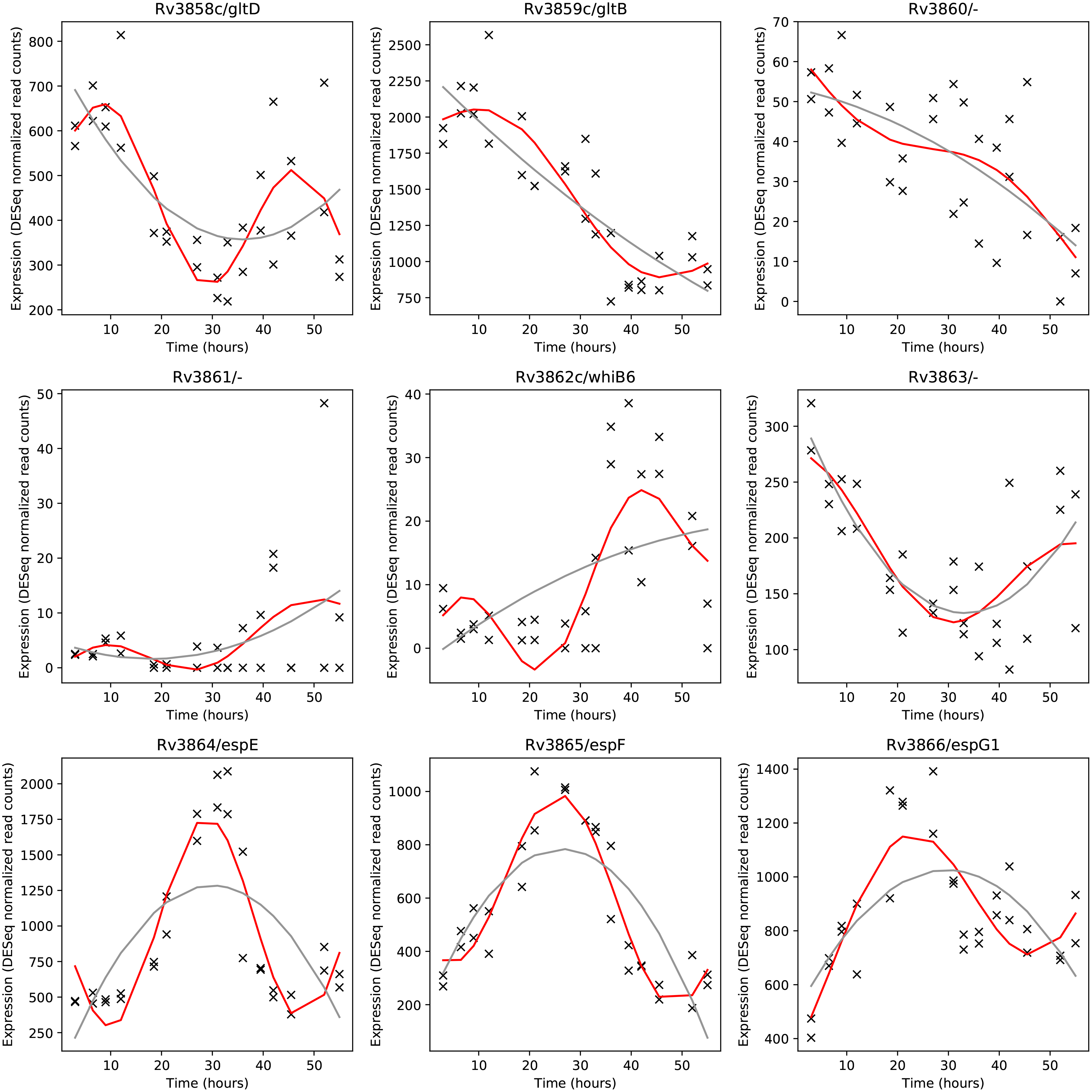
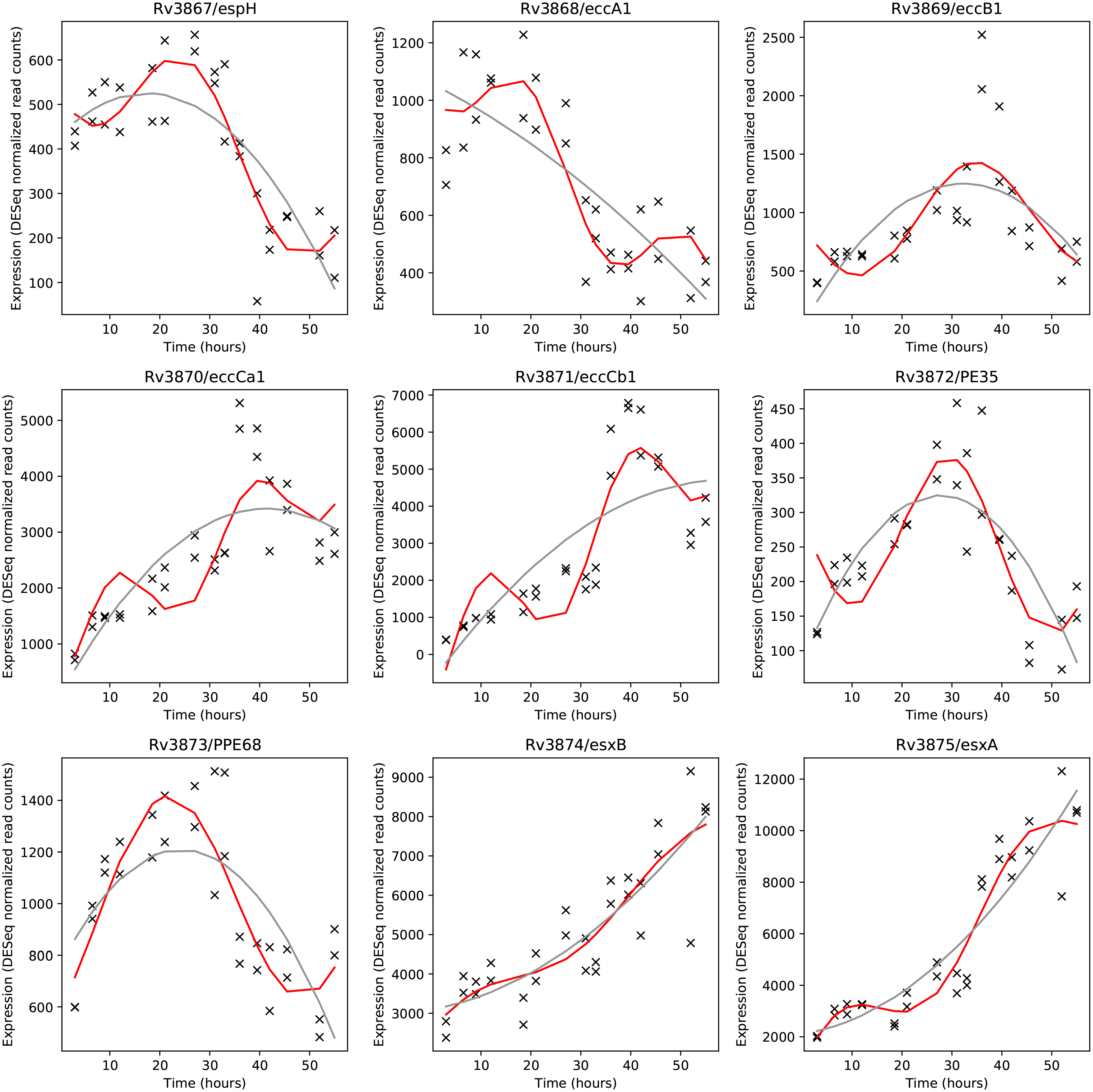
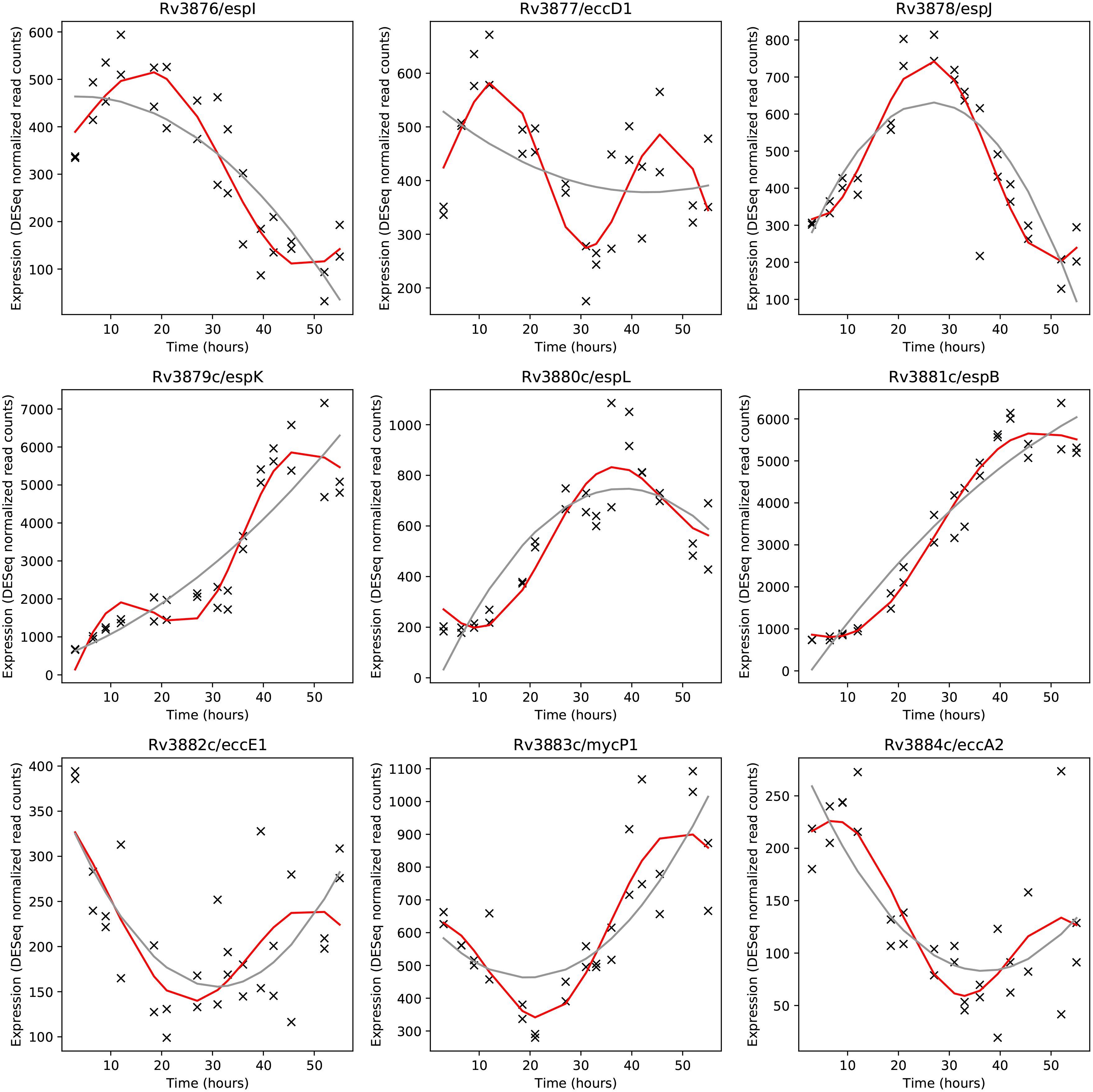
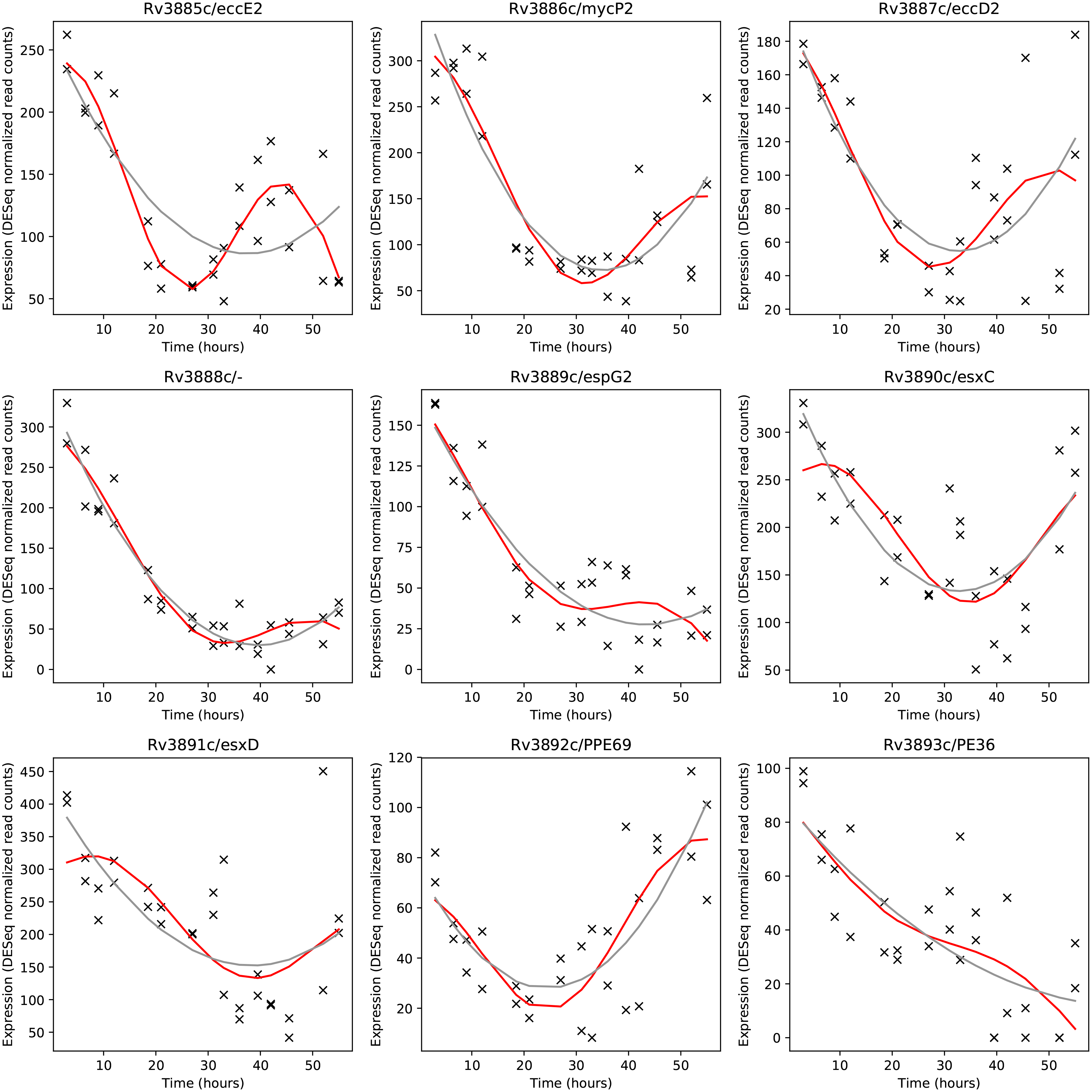
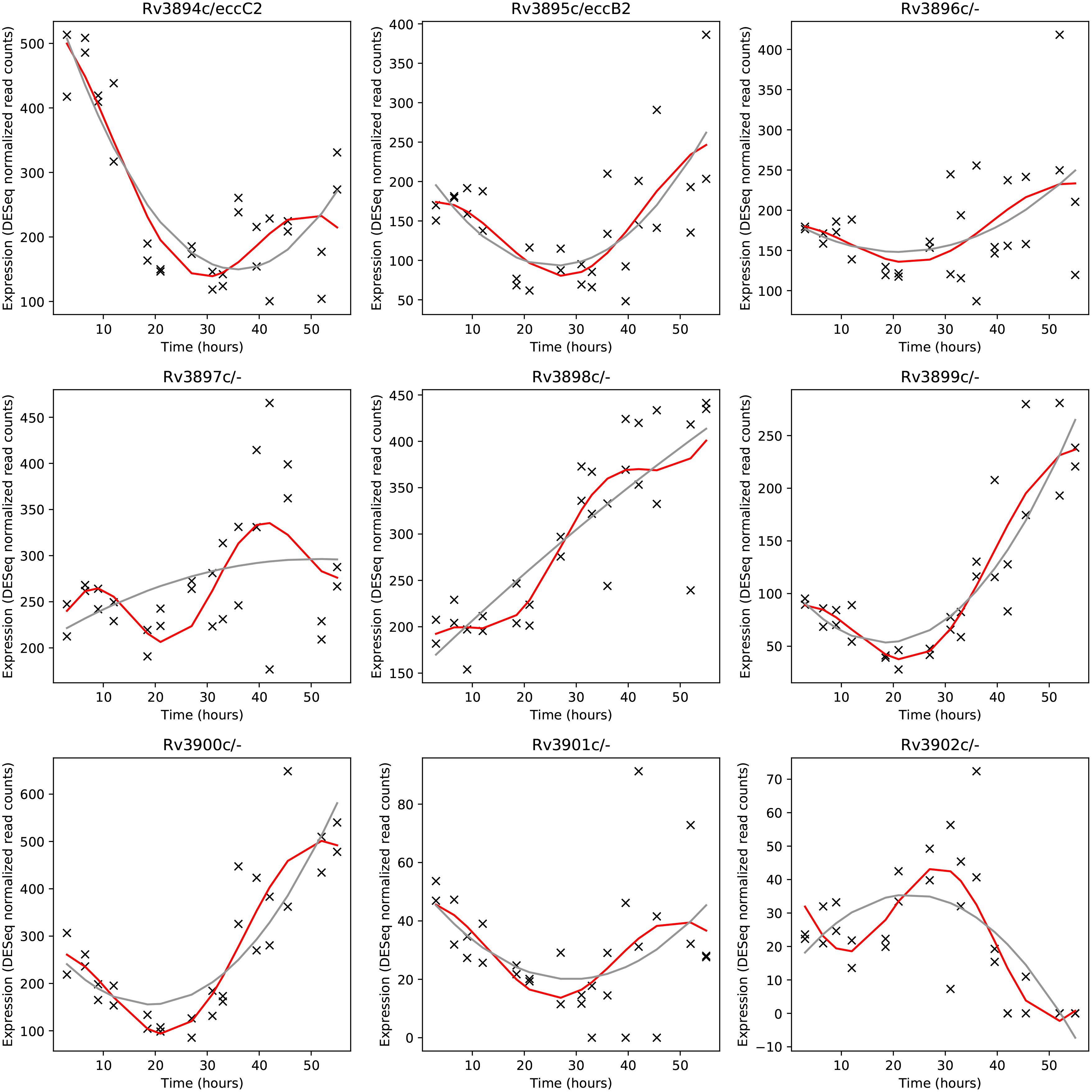
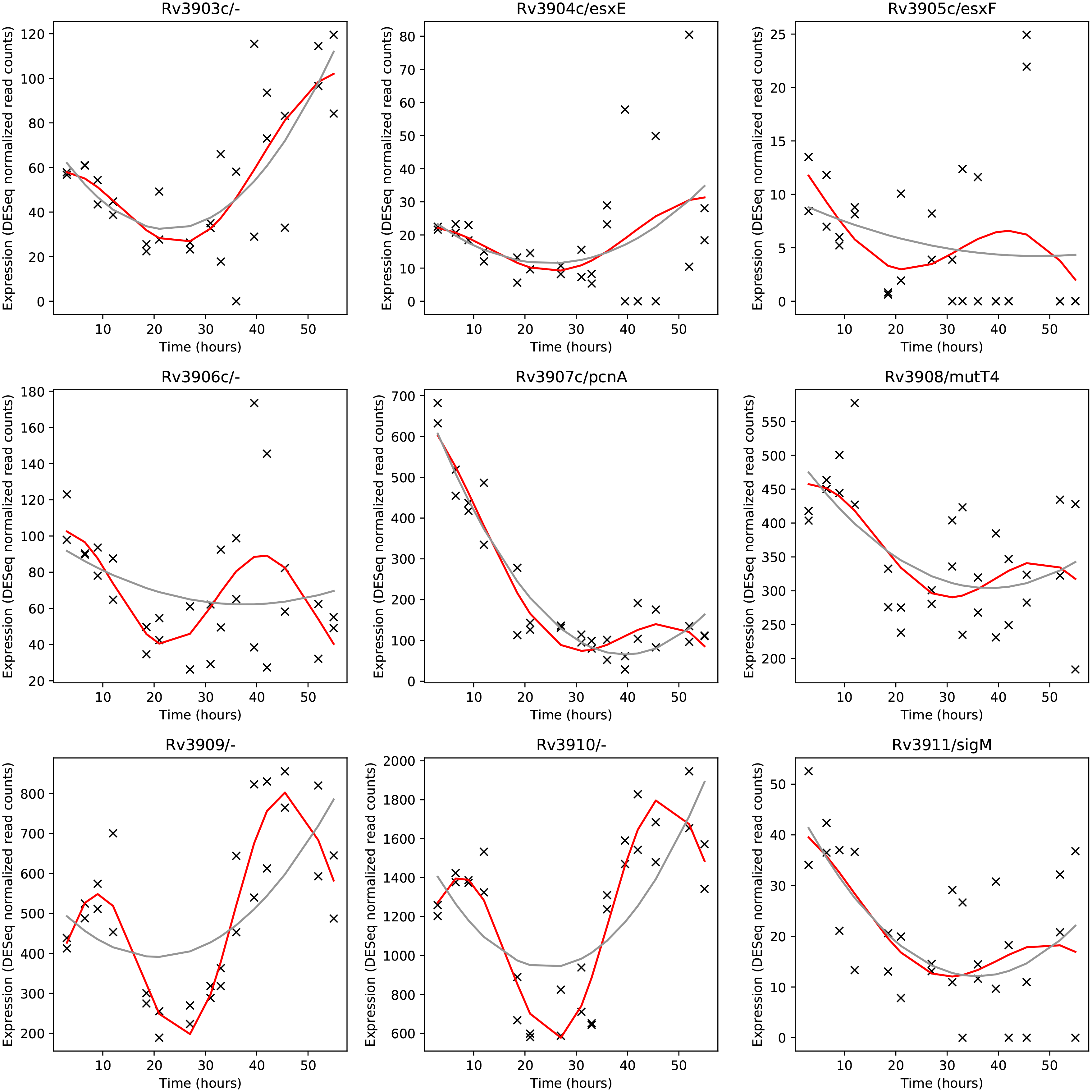
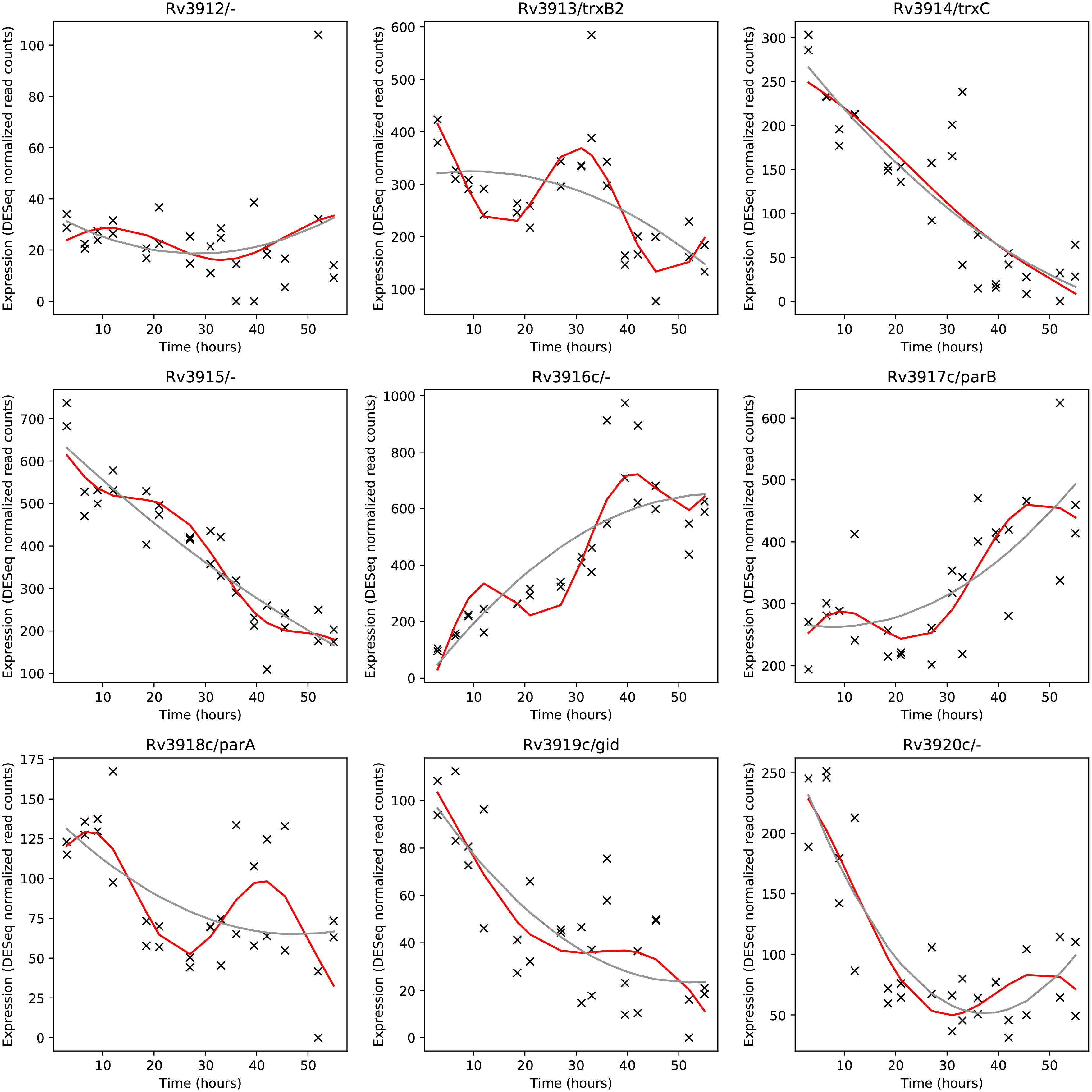
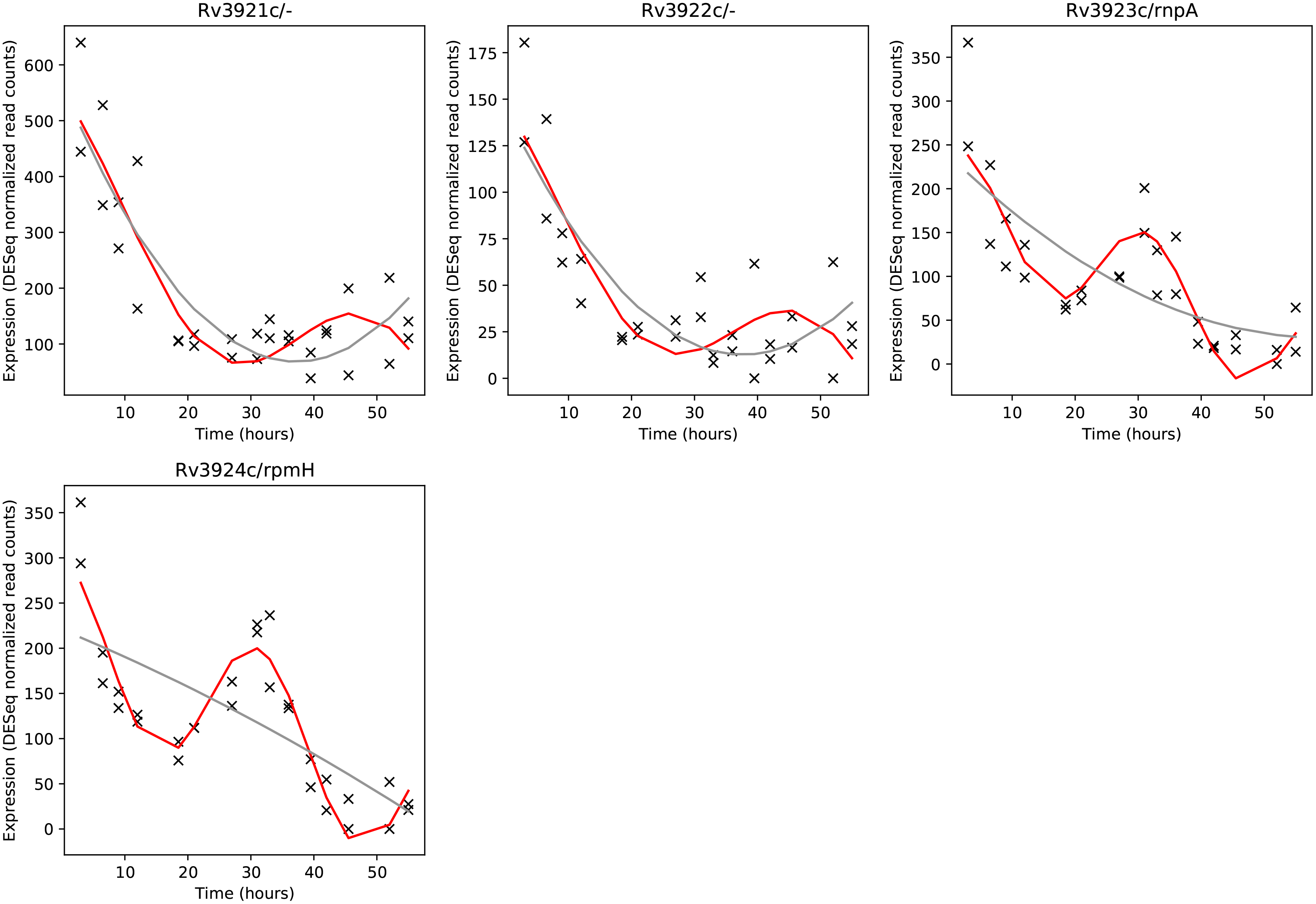
Sinusoidal fits (red) and quadratic fits (gray) to standard normalized DESeq read counts (count-mean/SD) for all genes in *Mtbcos*

**Fig S4:**
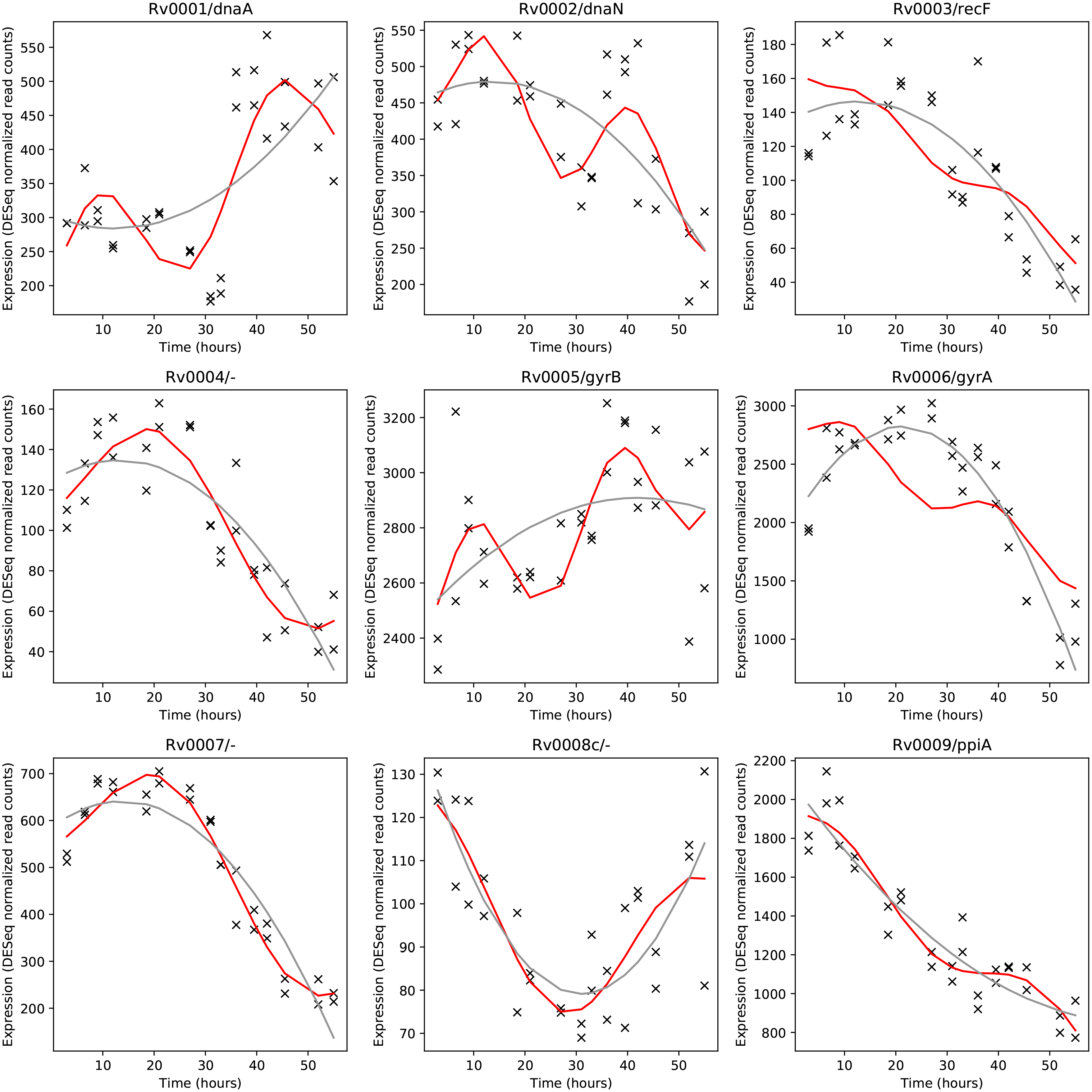
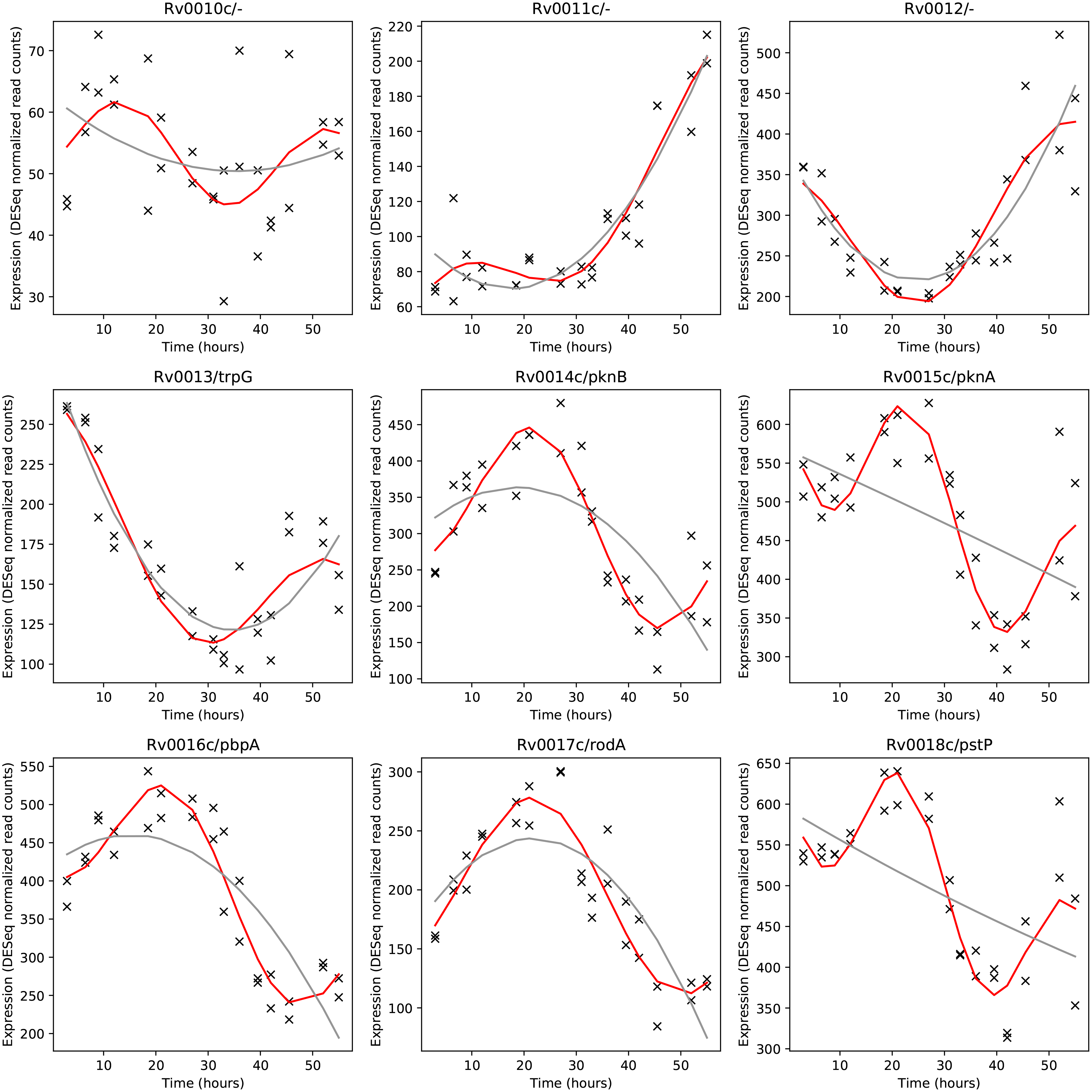
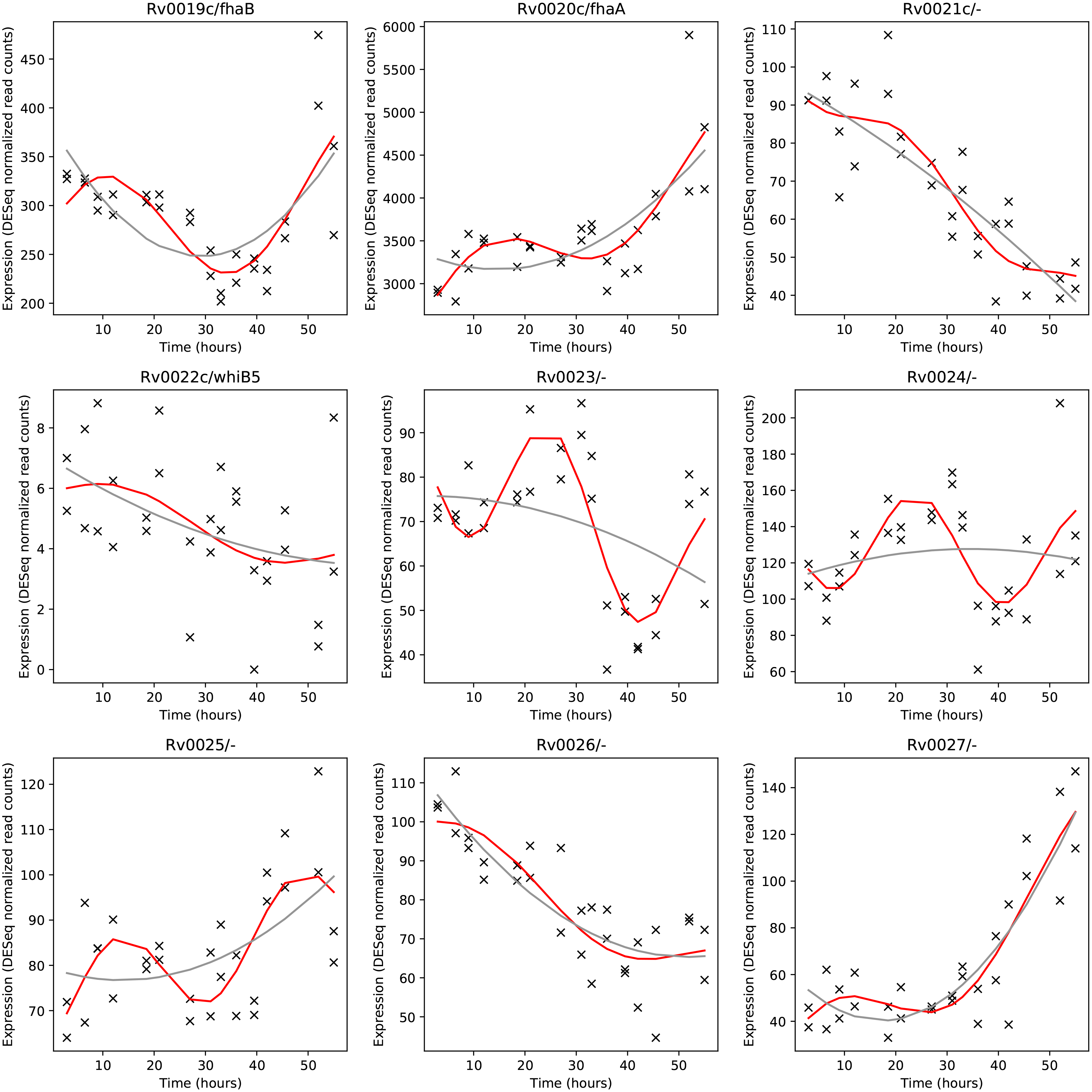
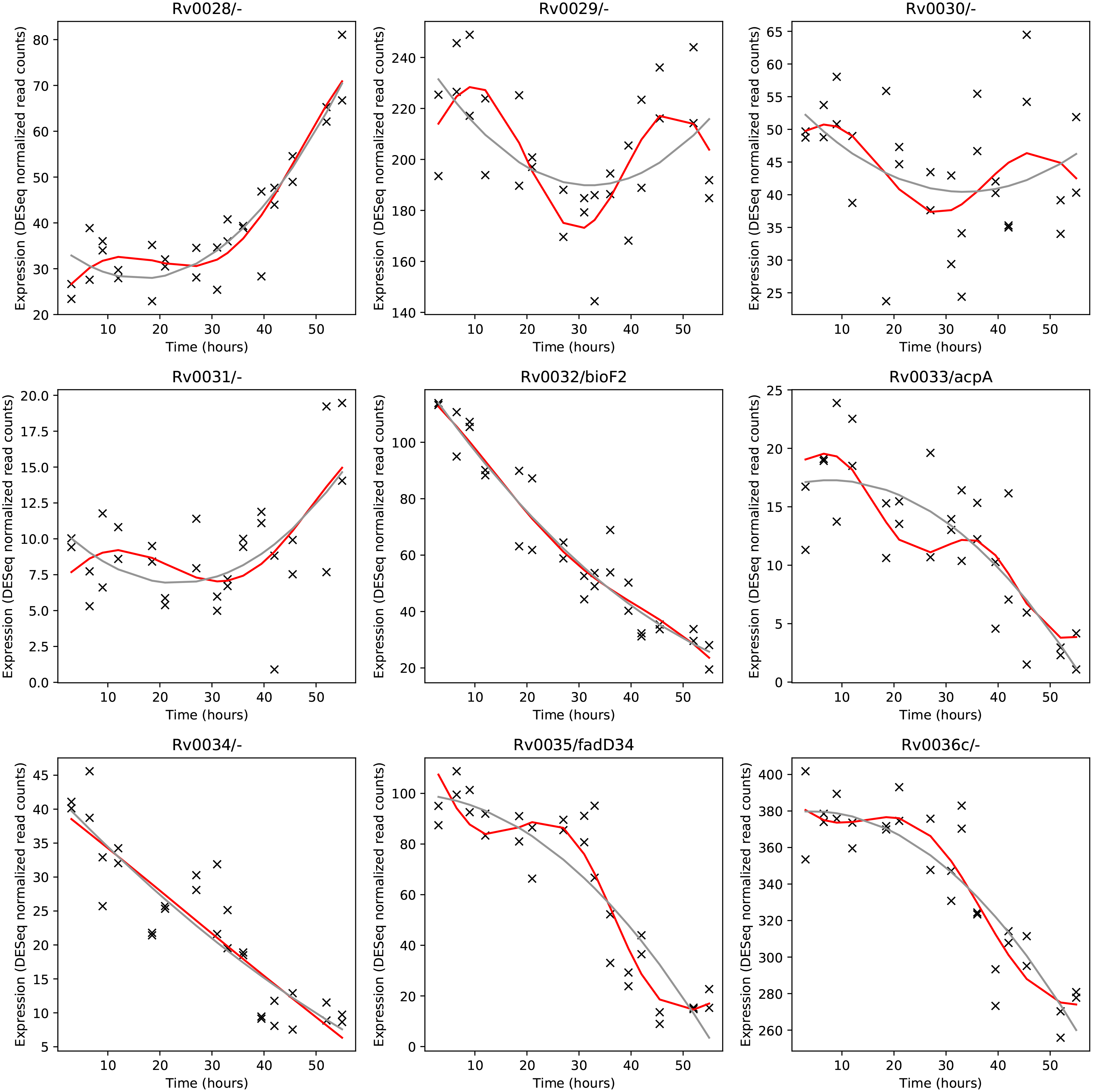
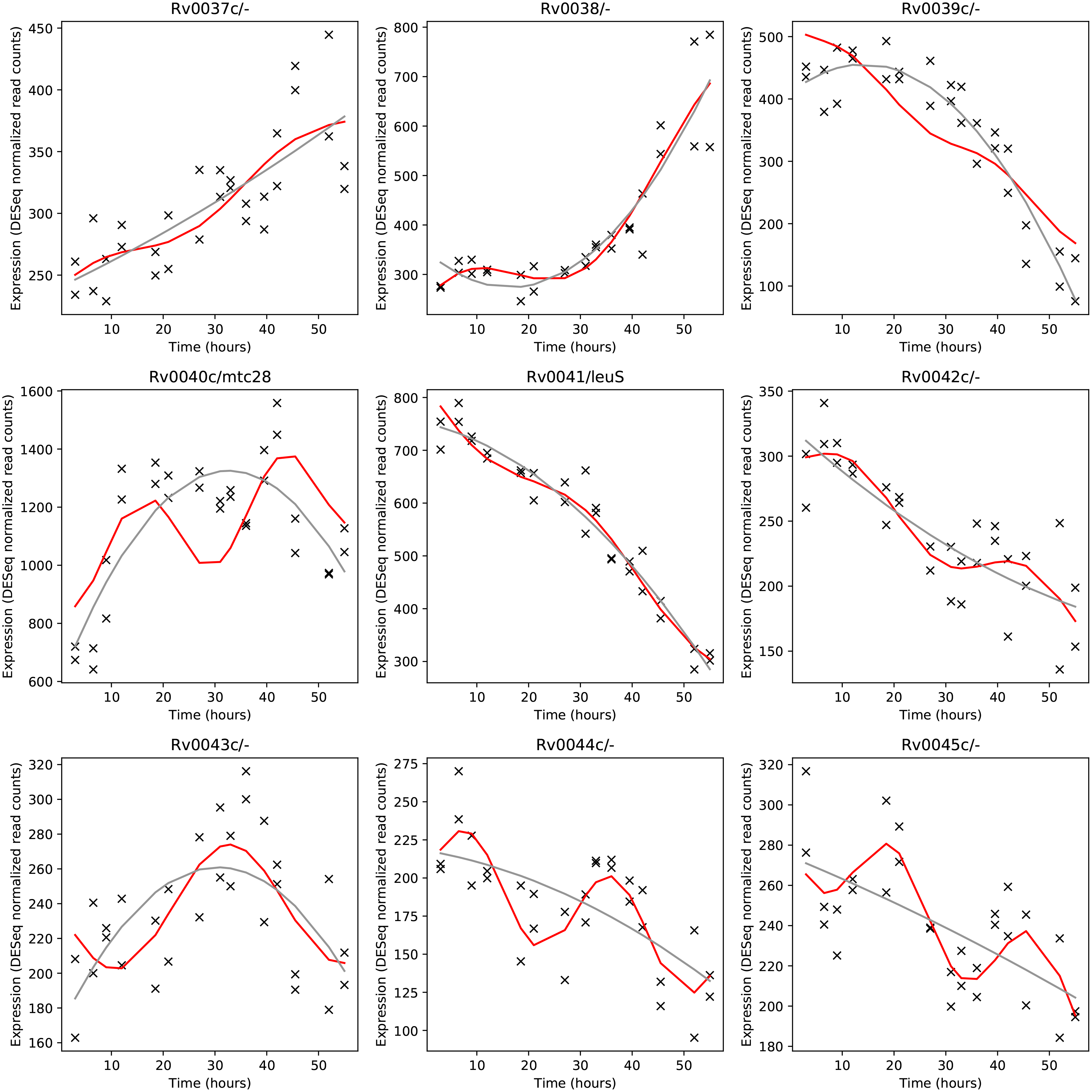
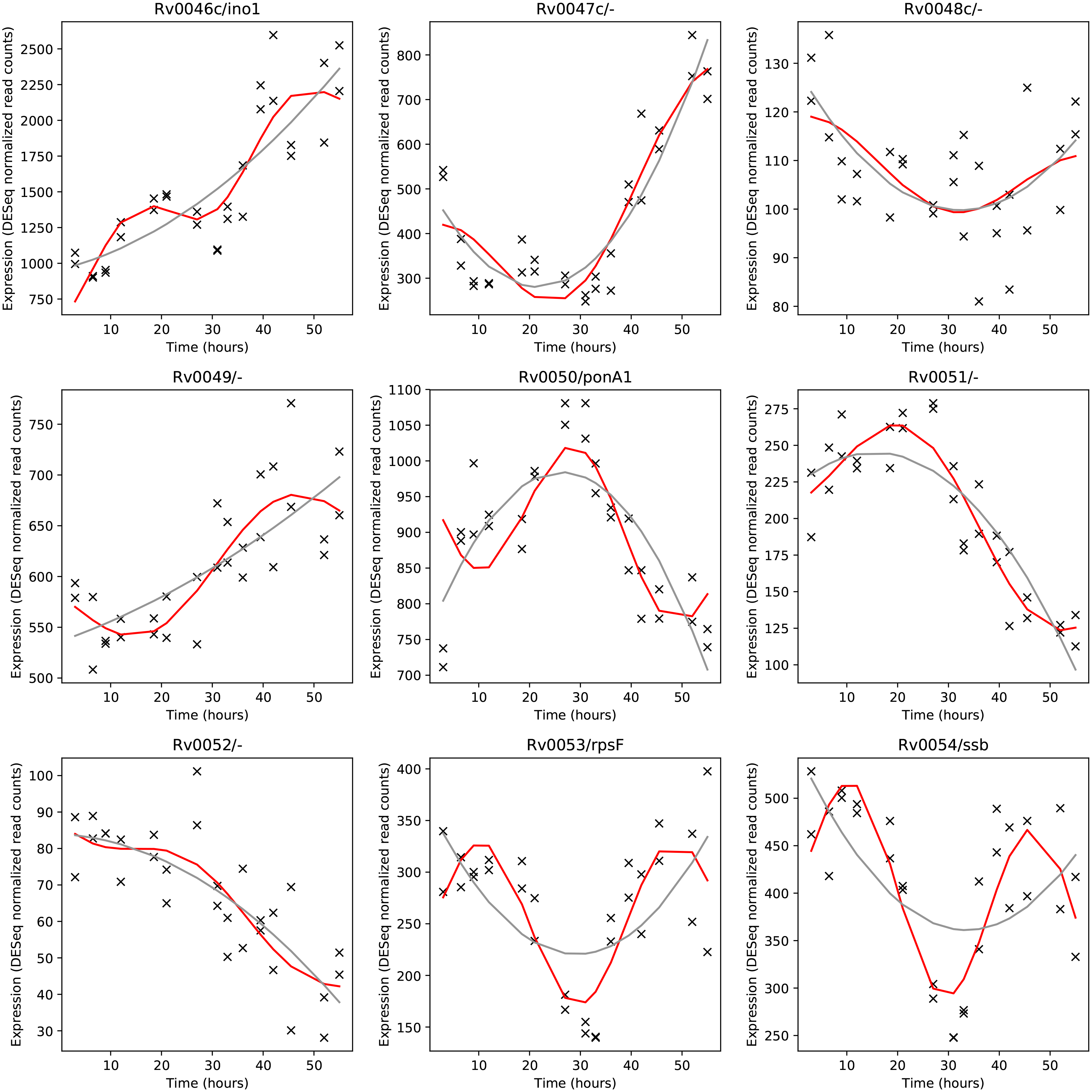
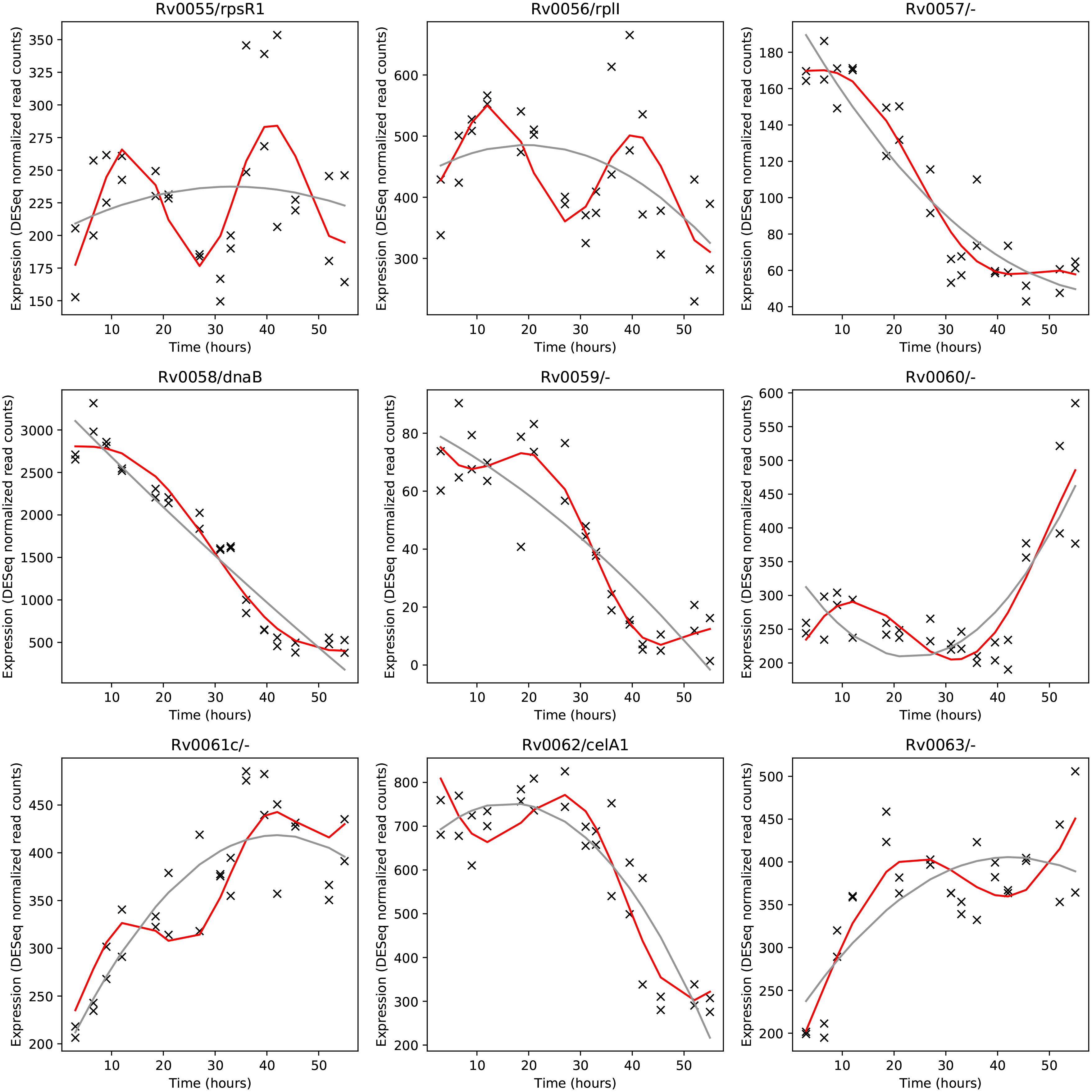
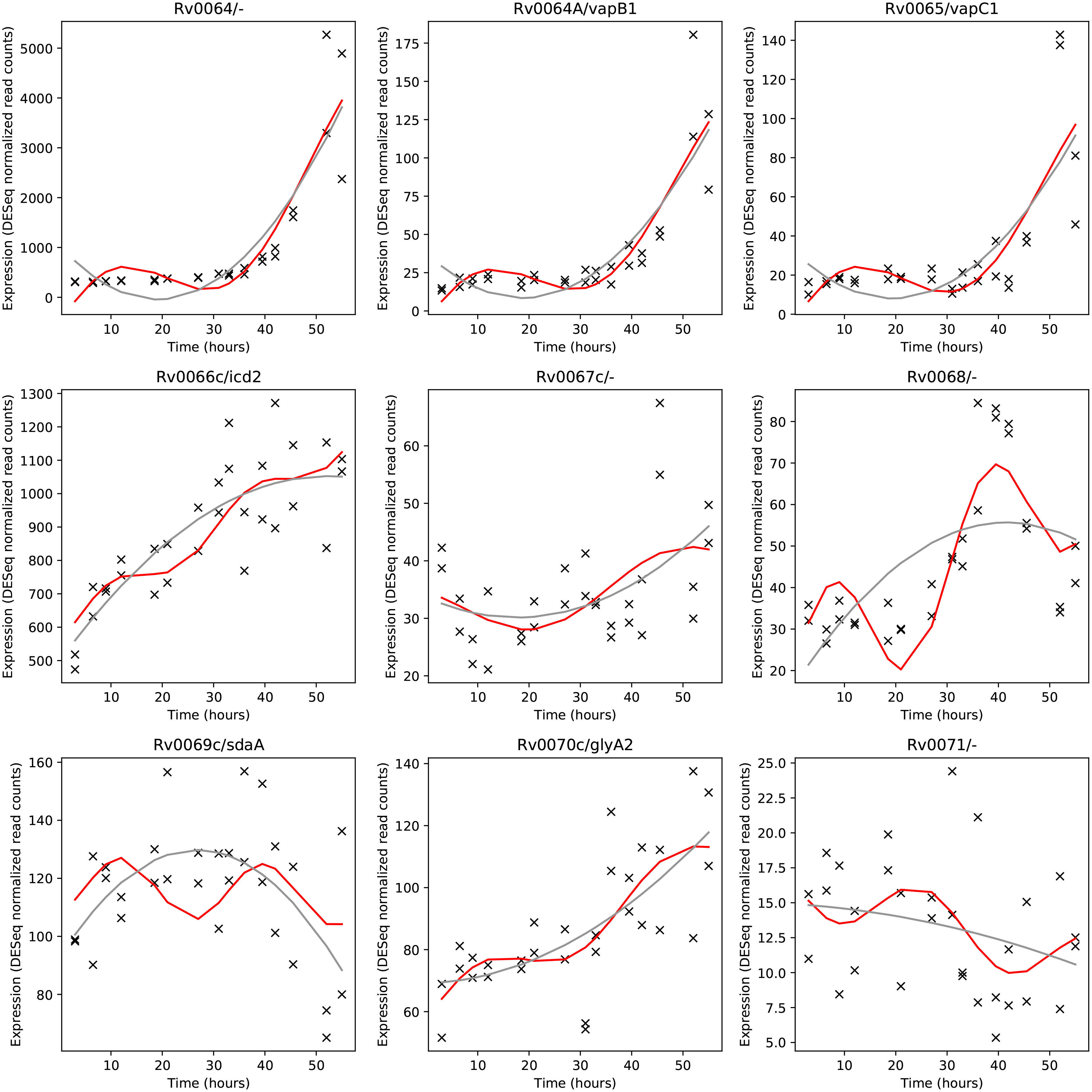
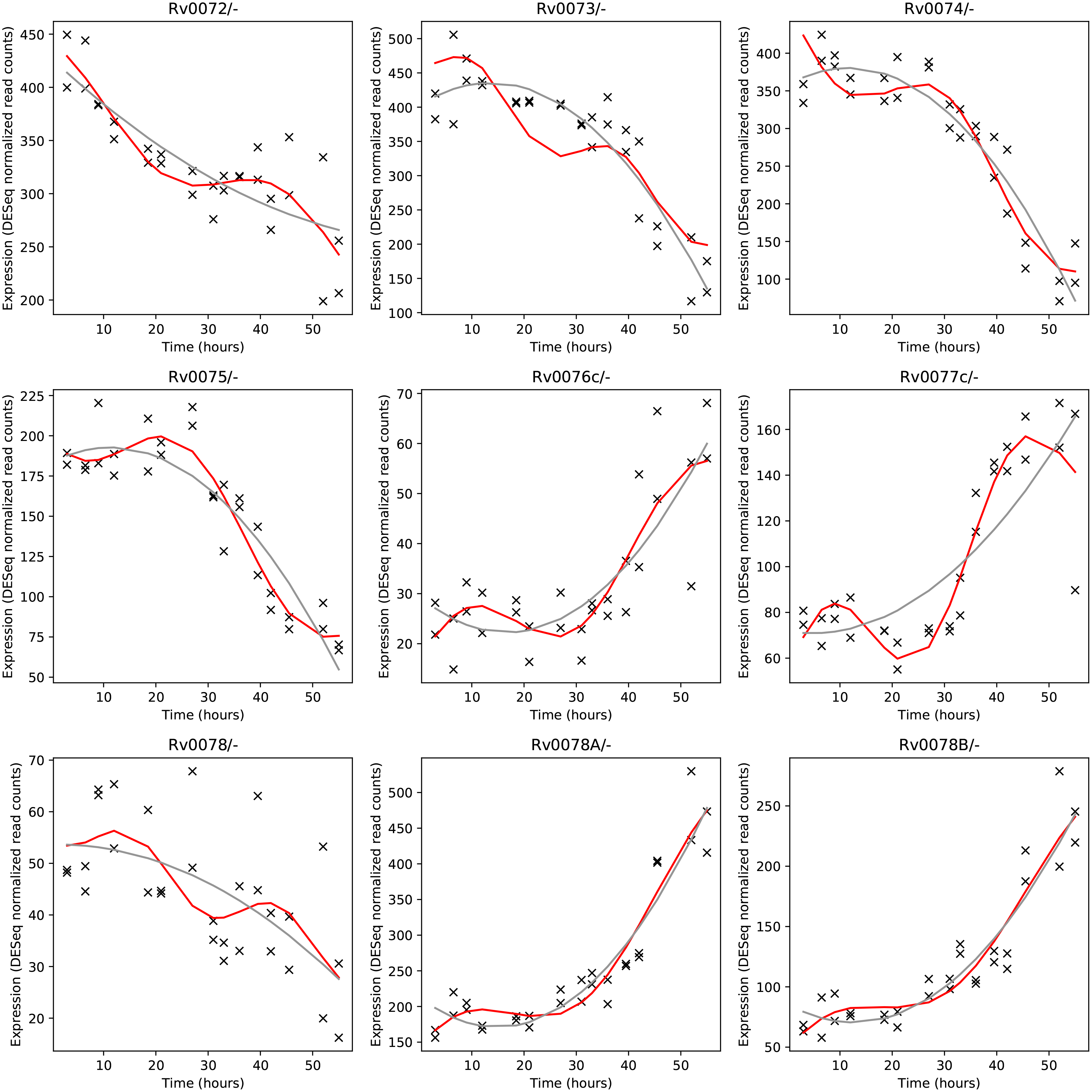
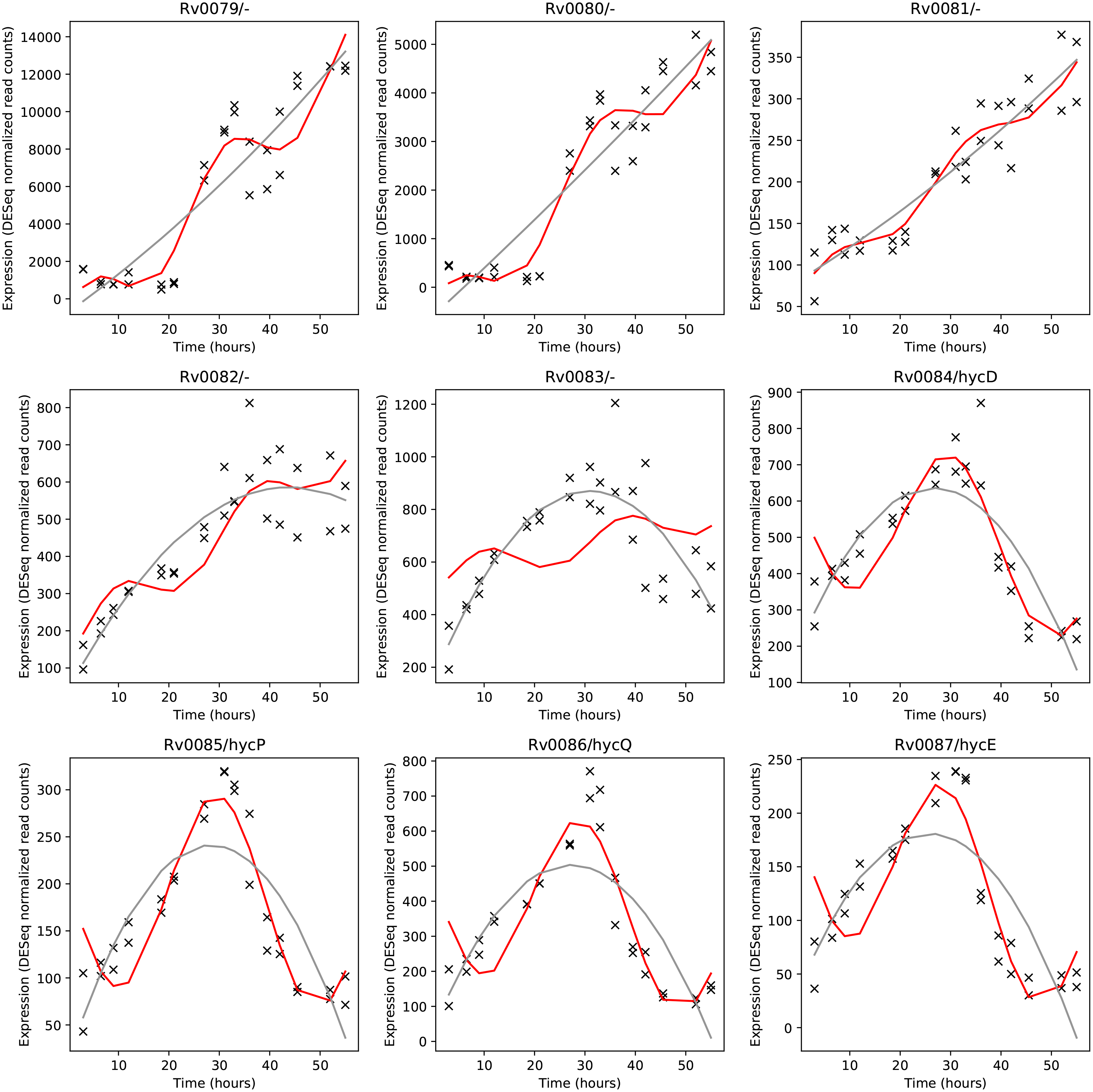
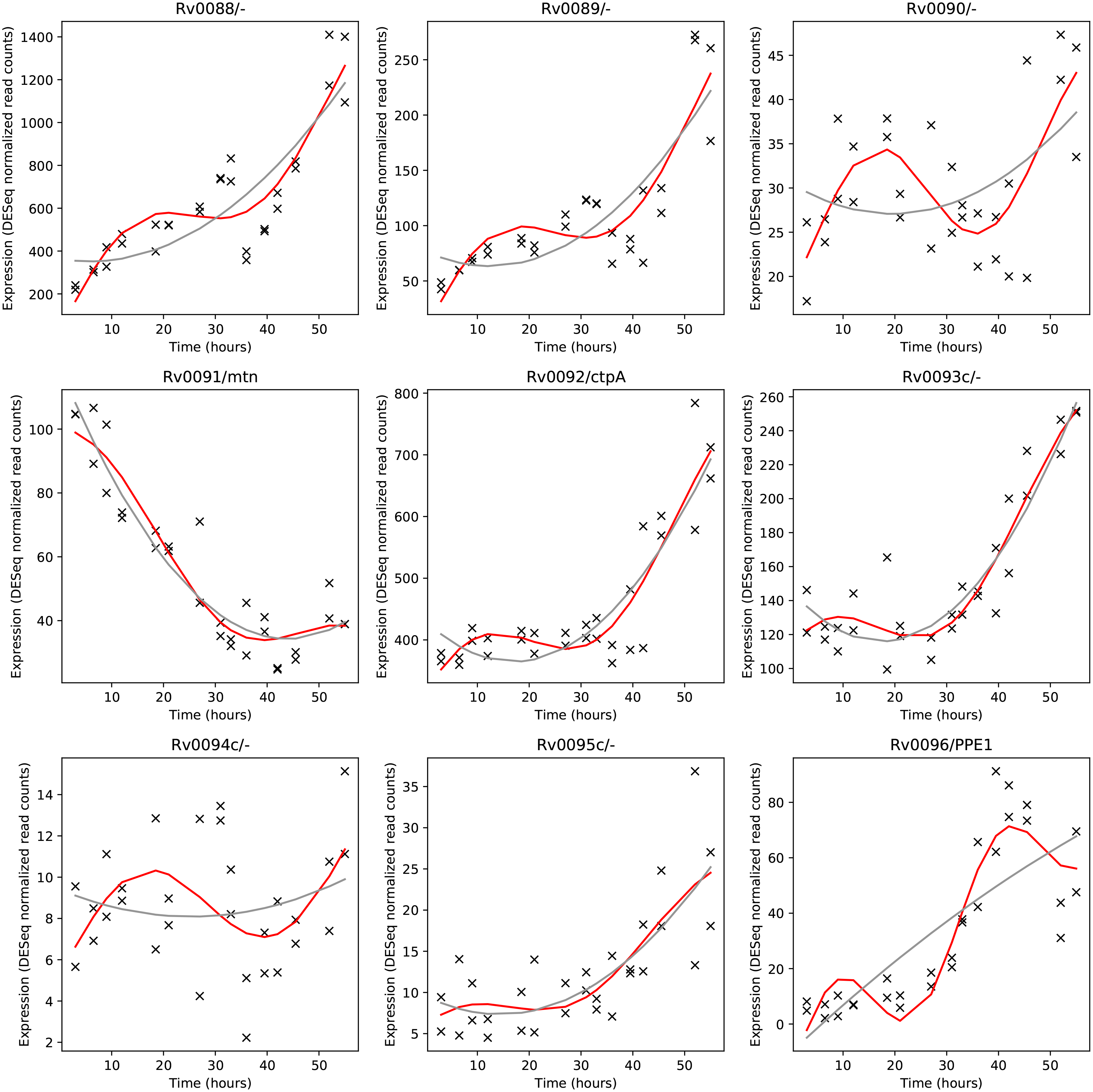
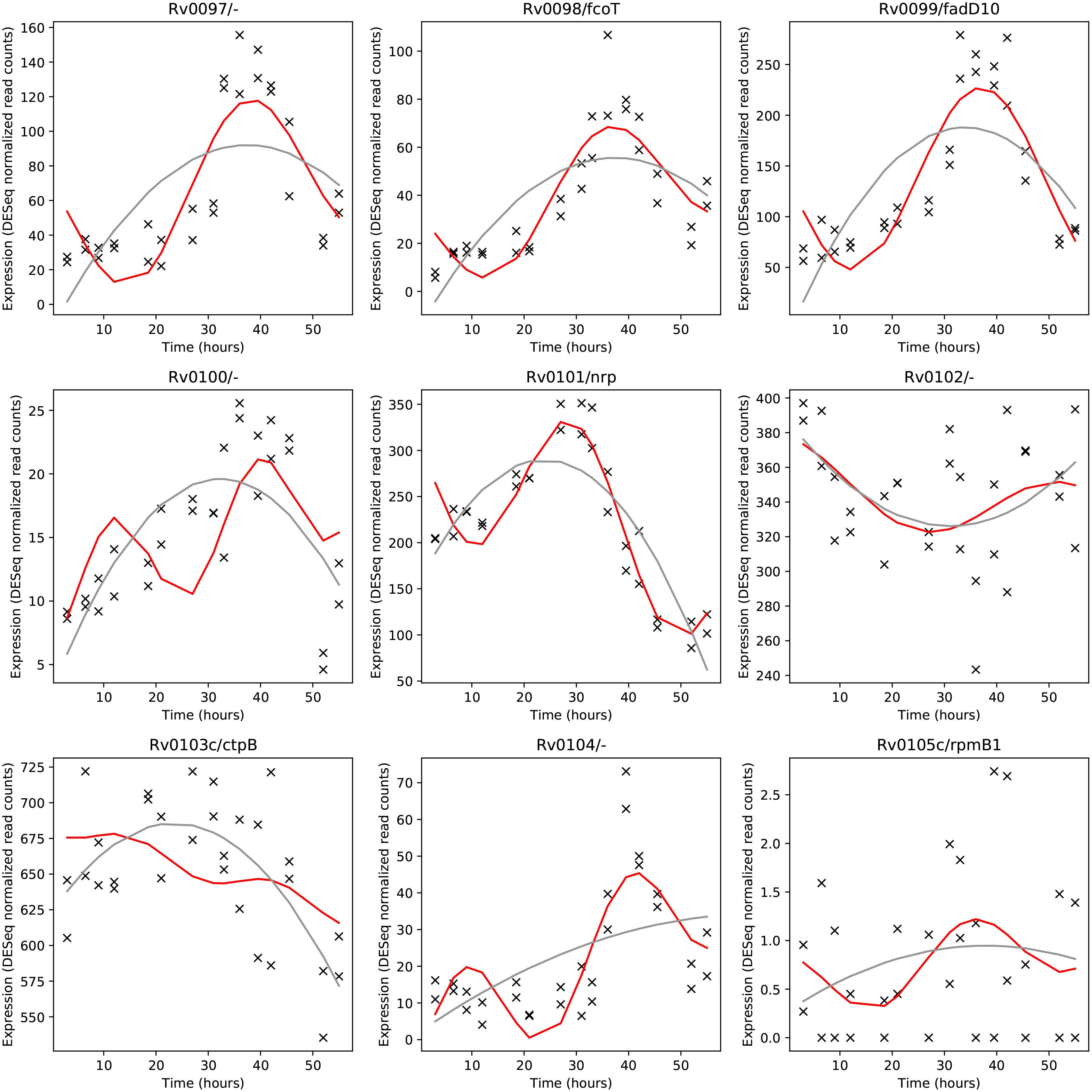
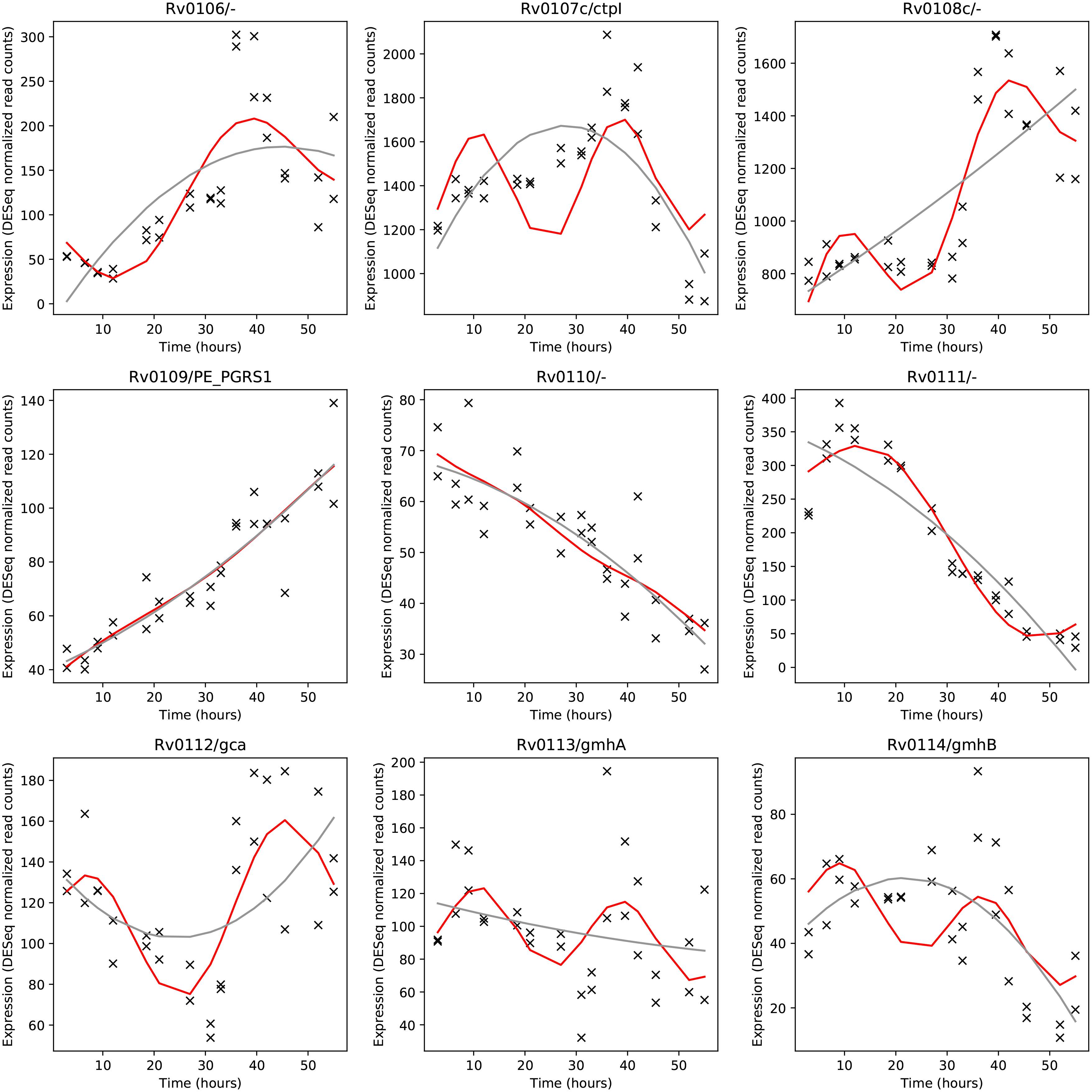
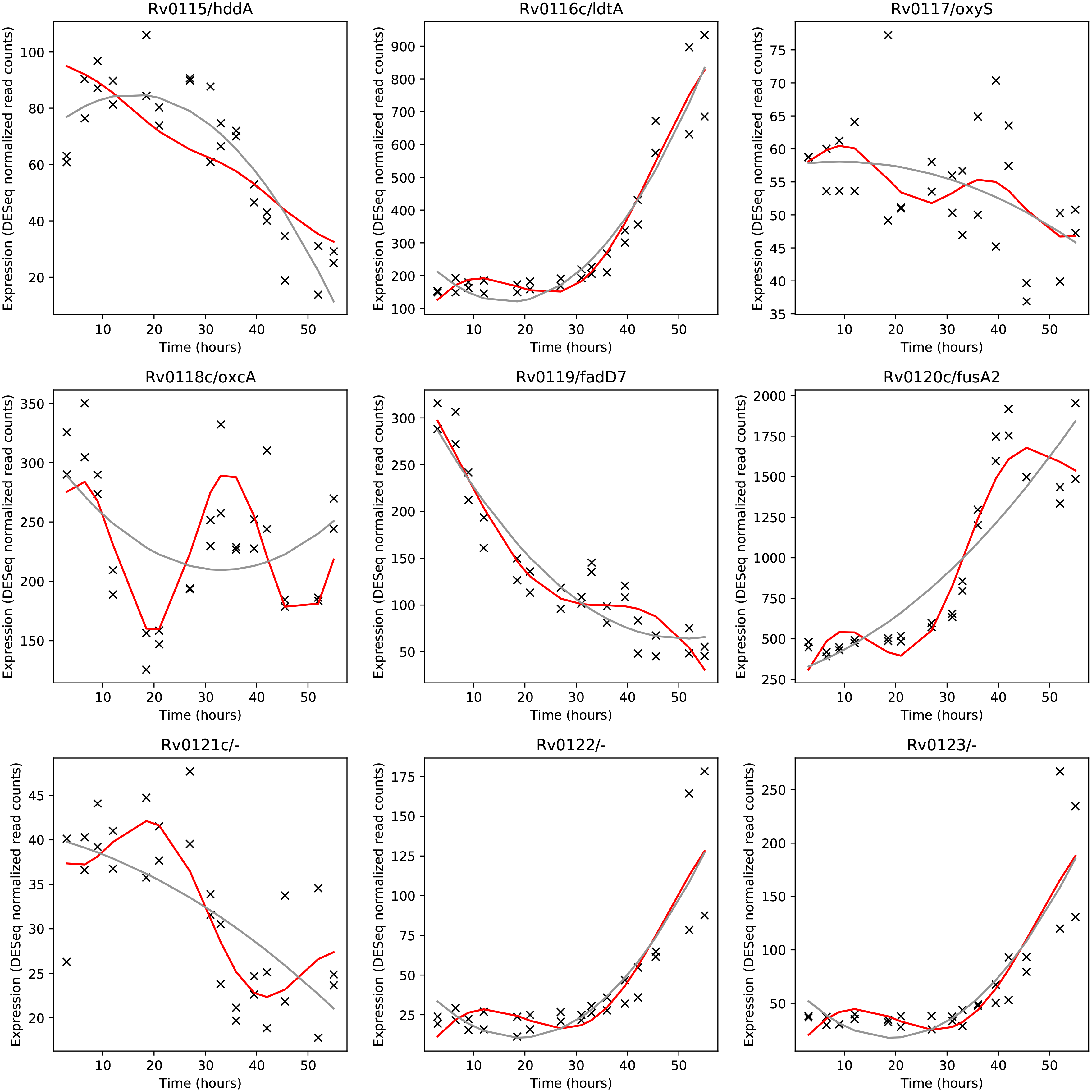
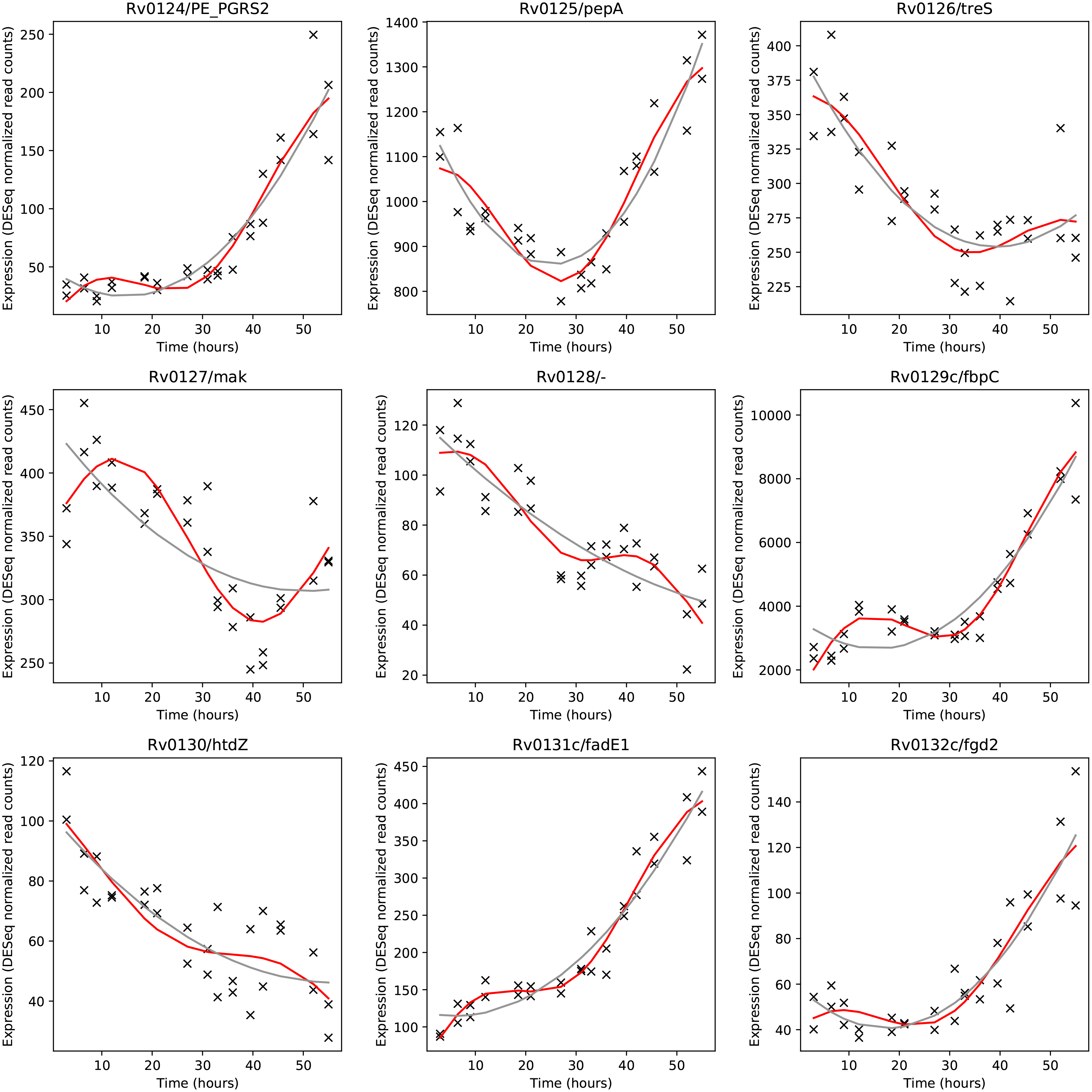
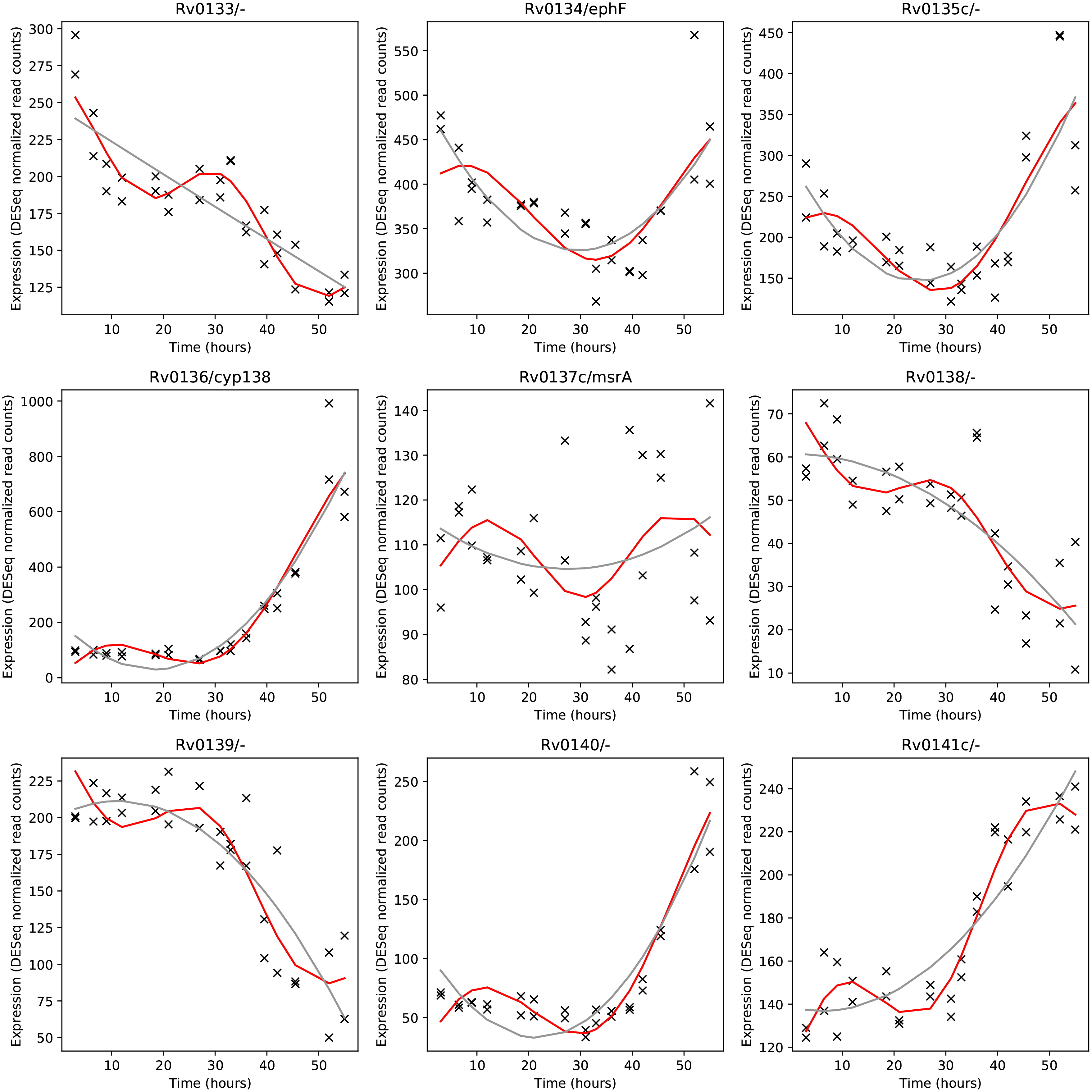
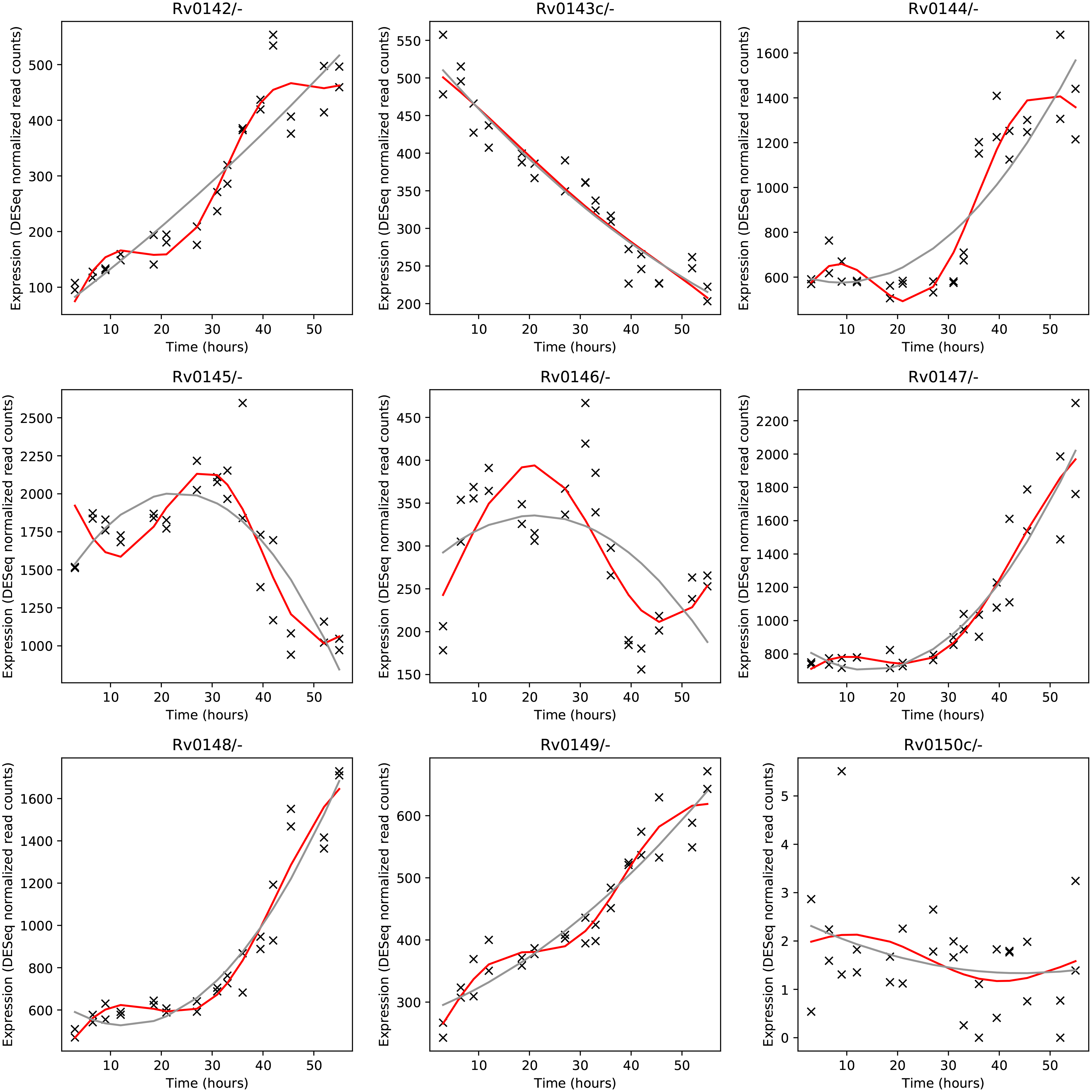
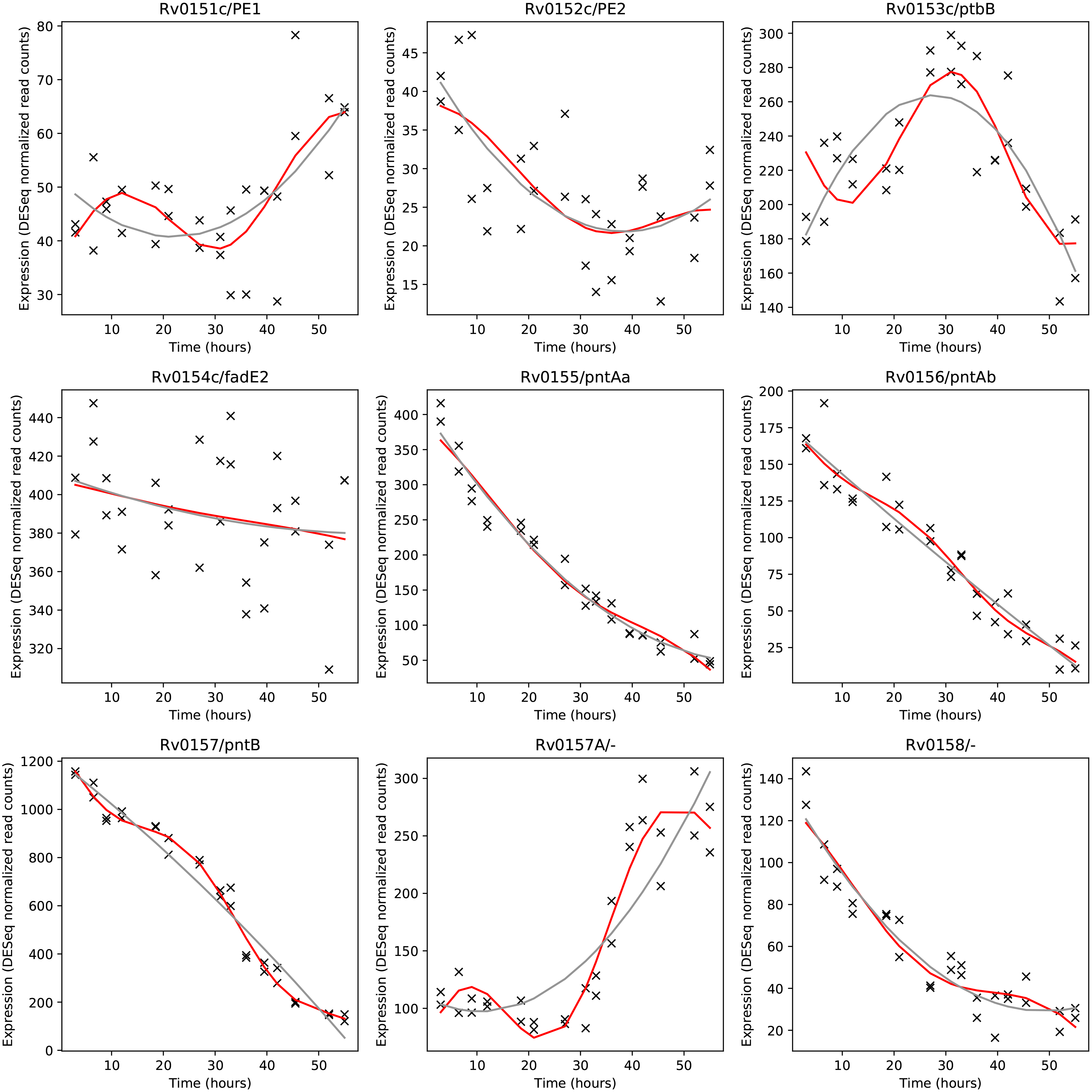
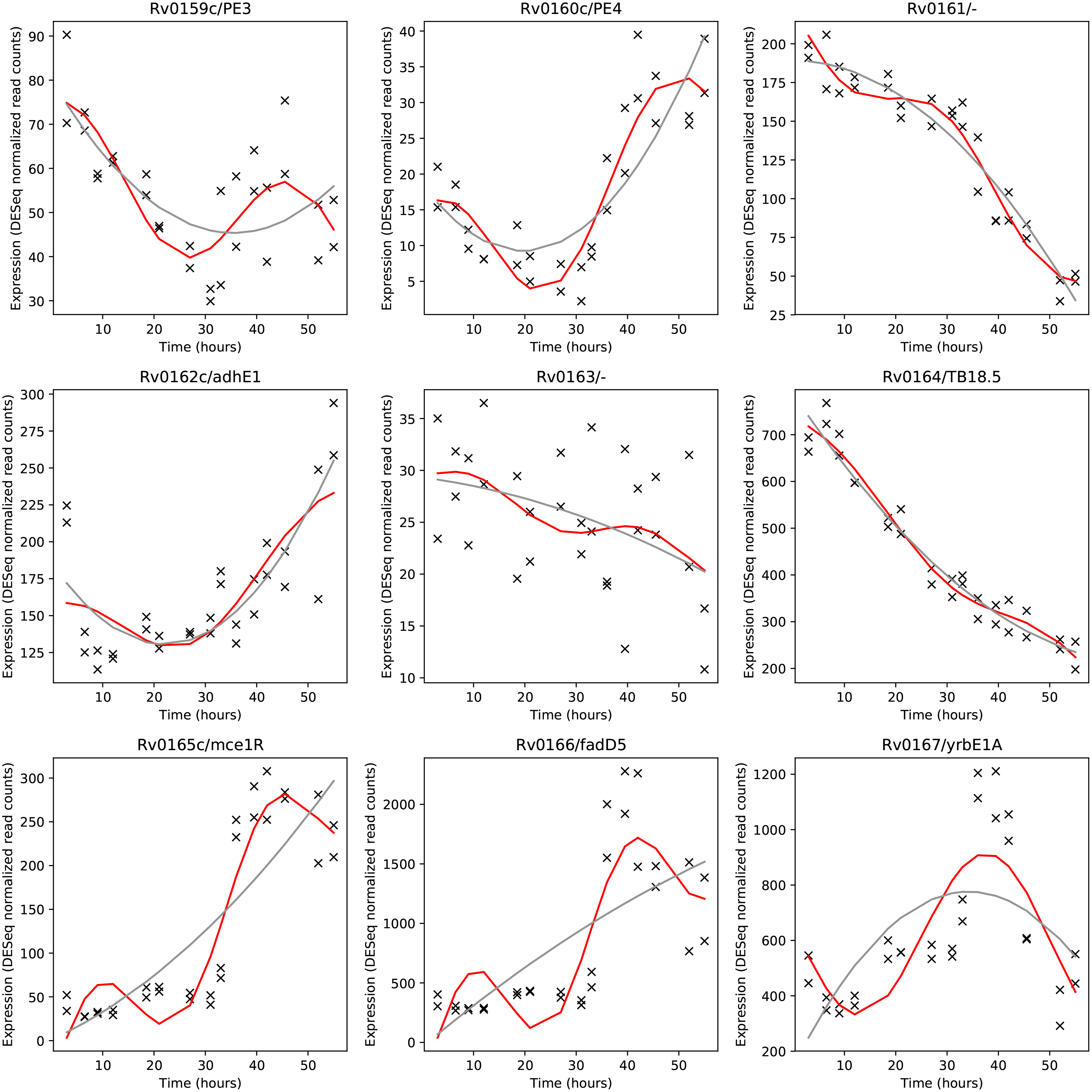
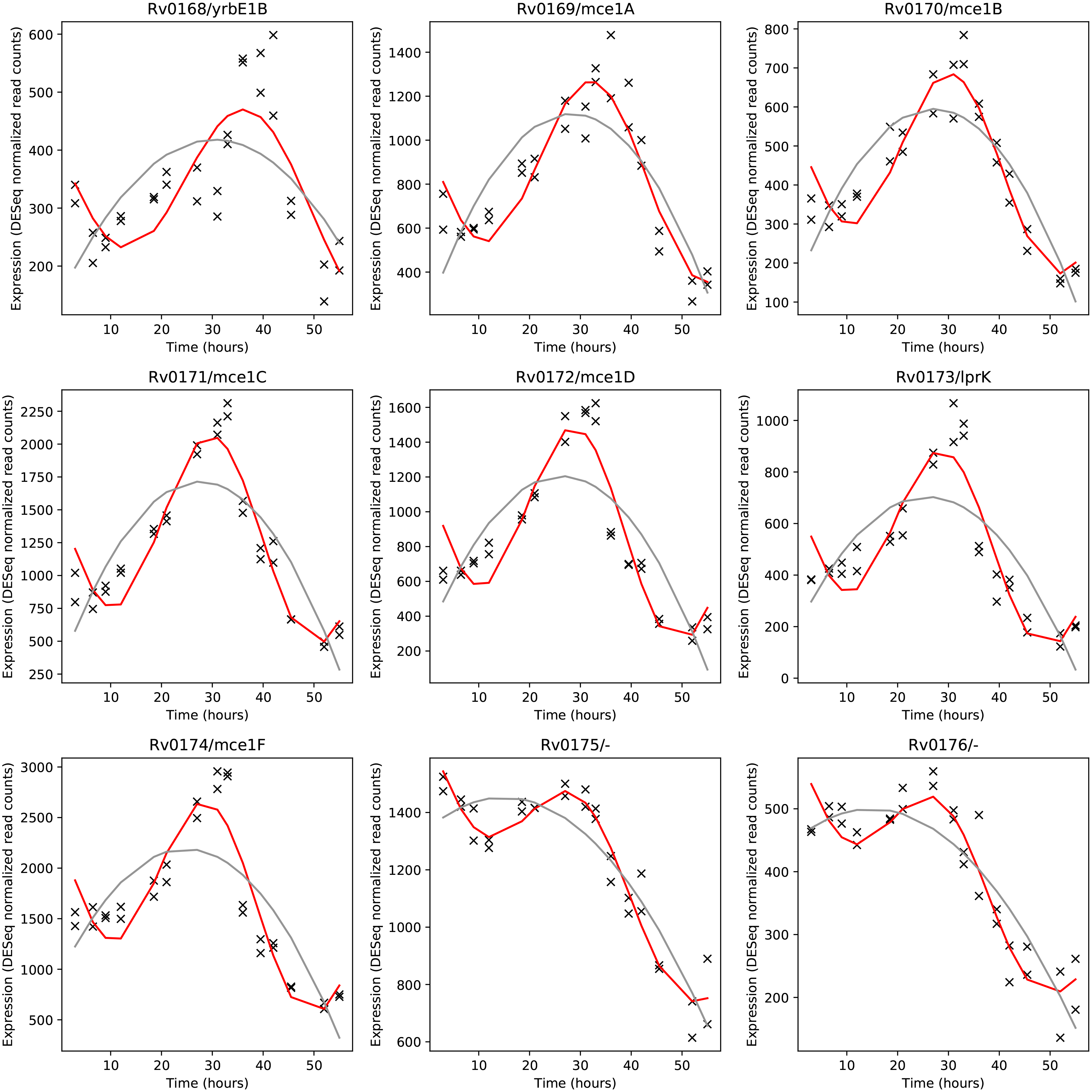
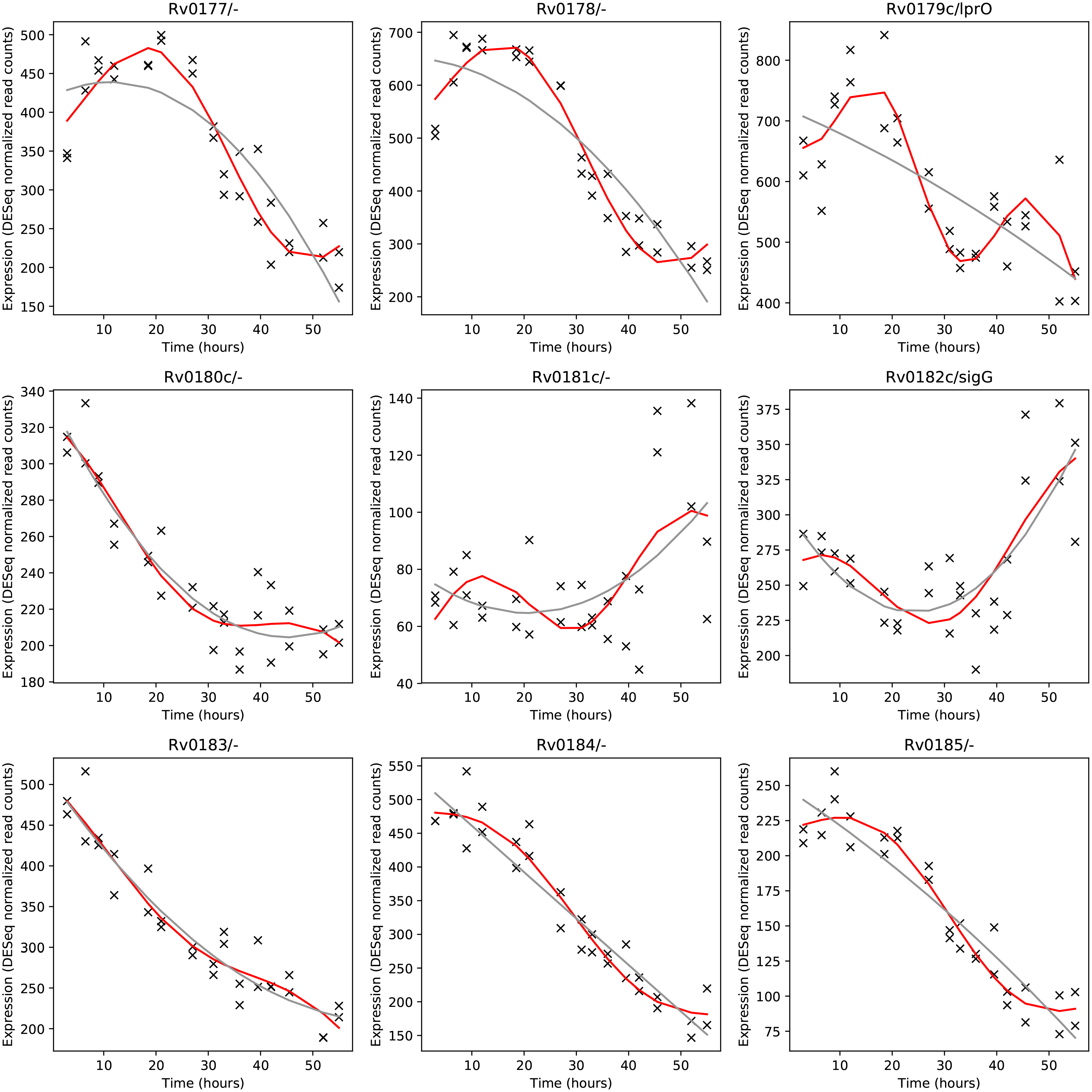
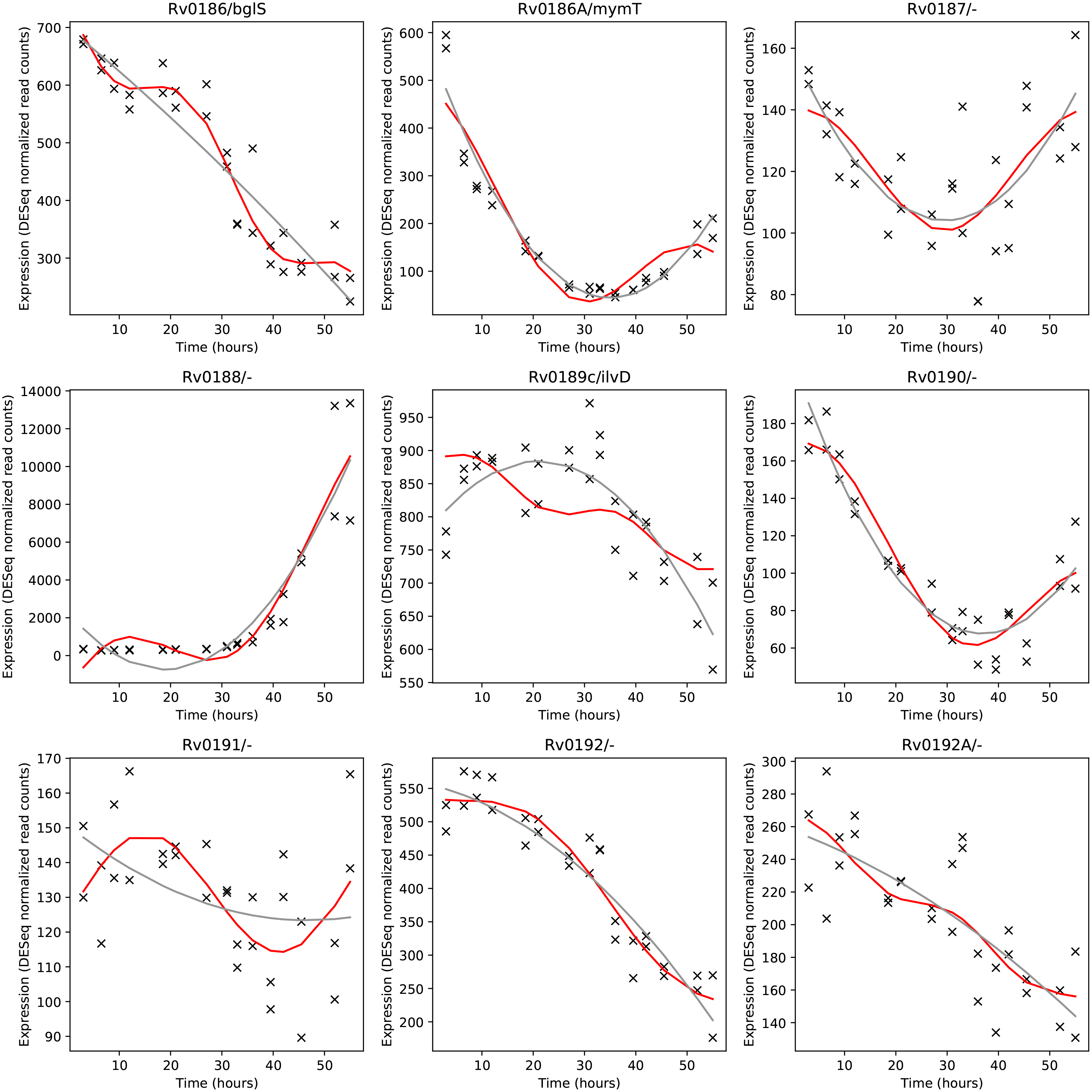
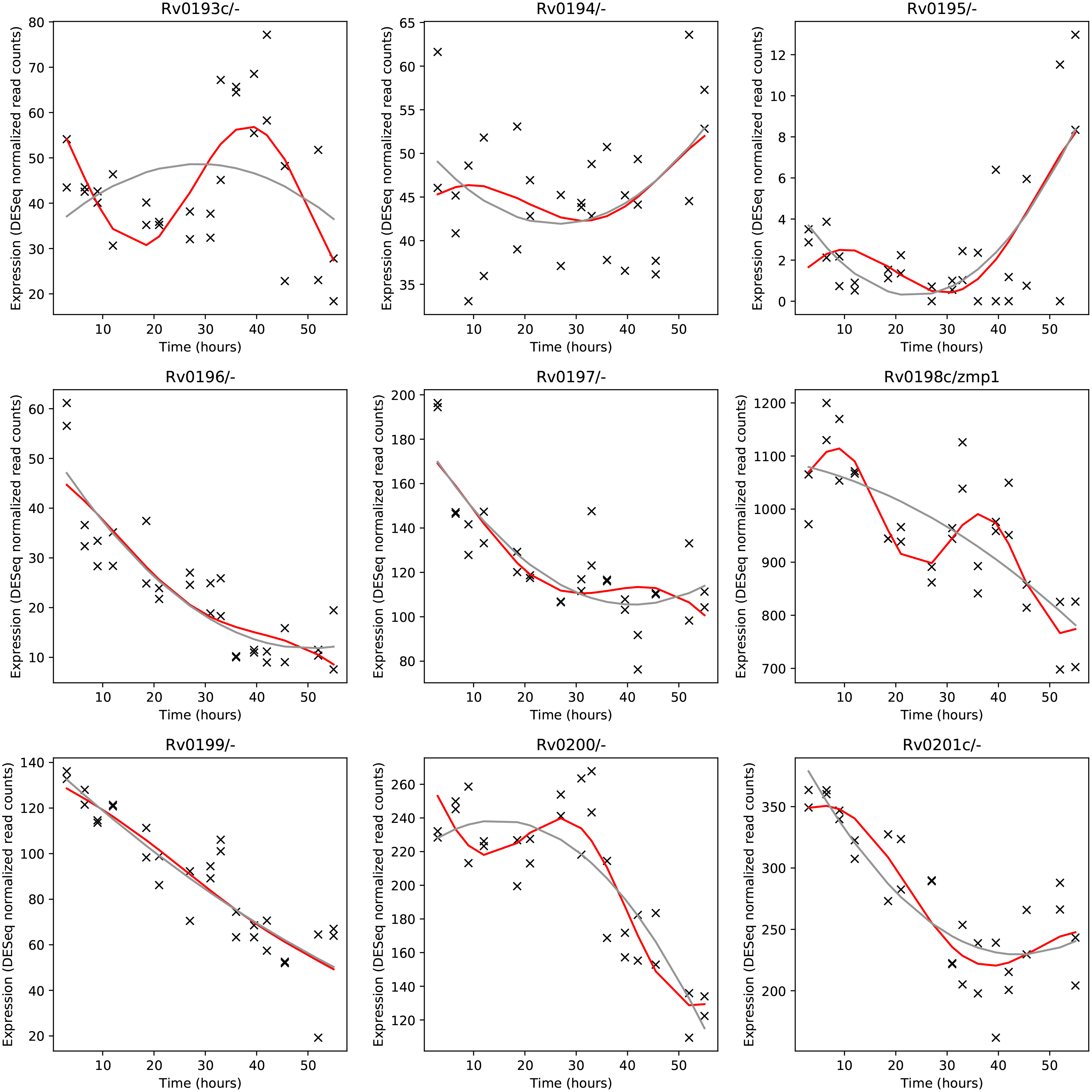
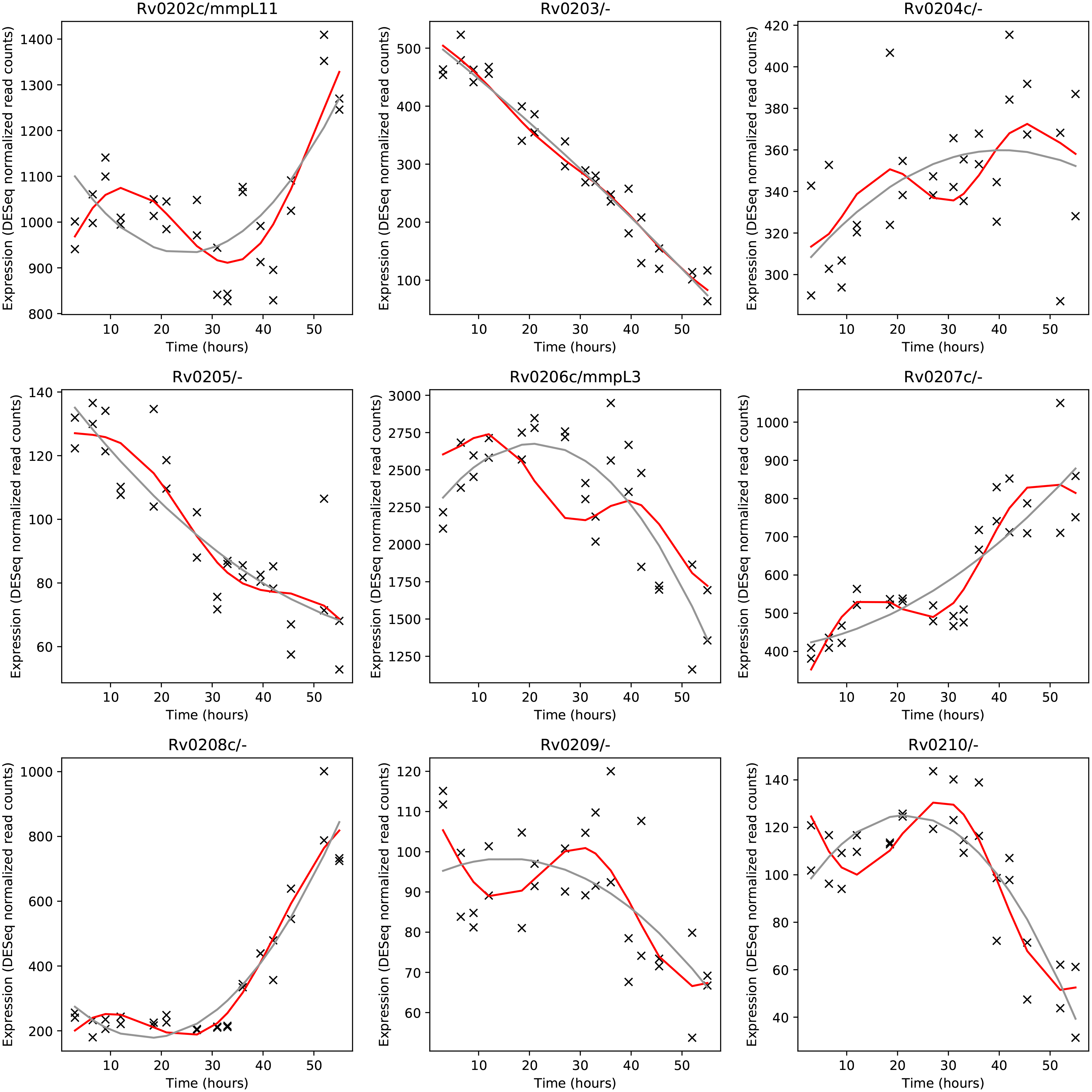
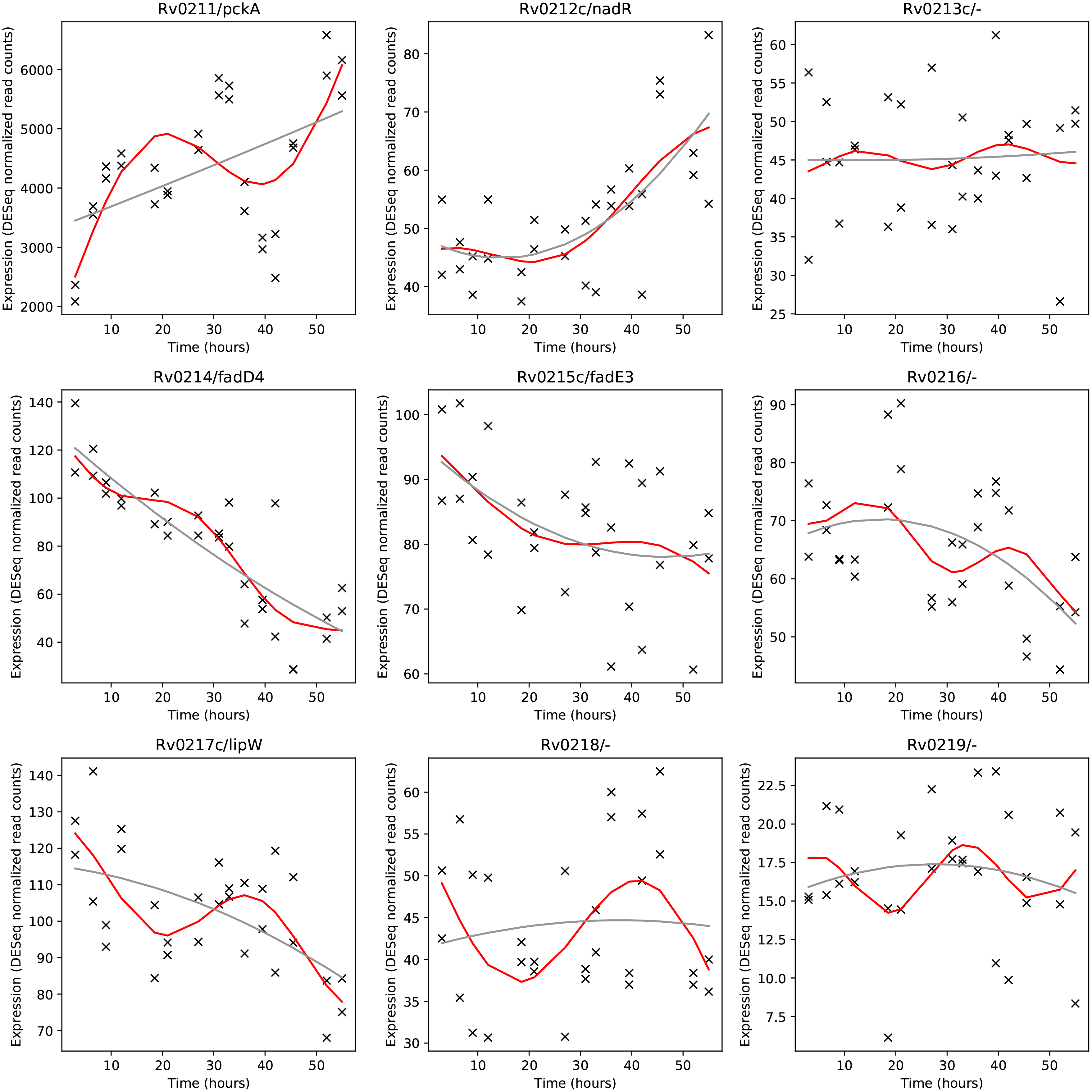
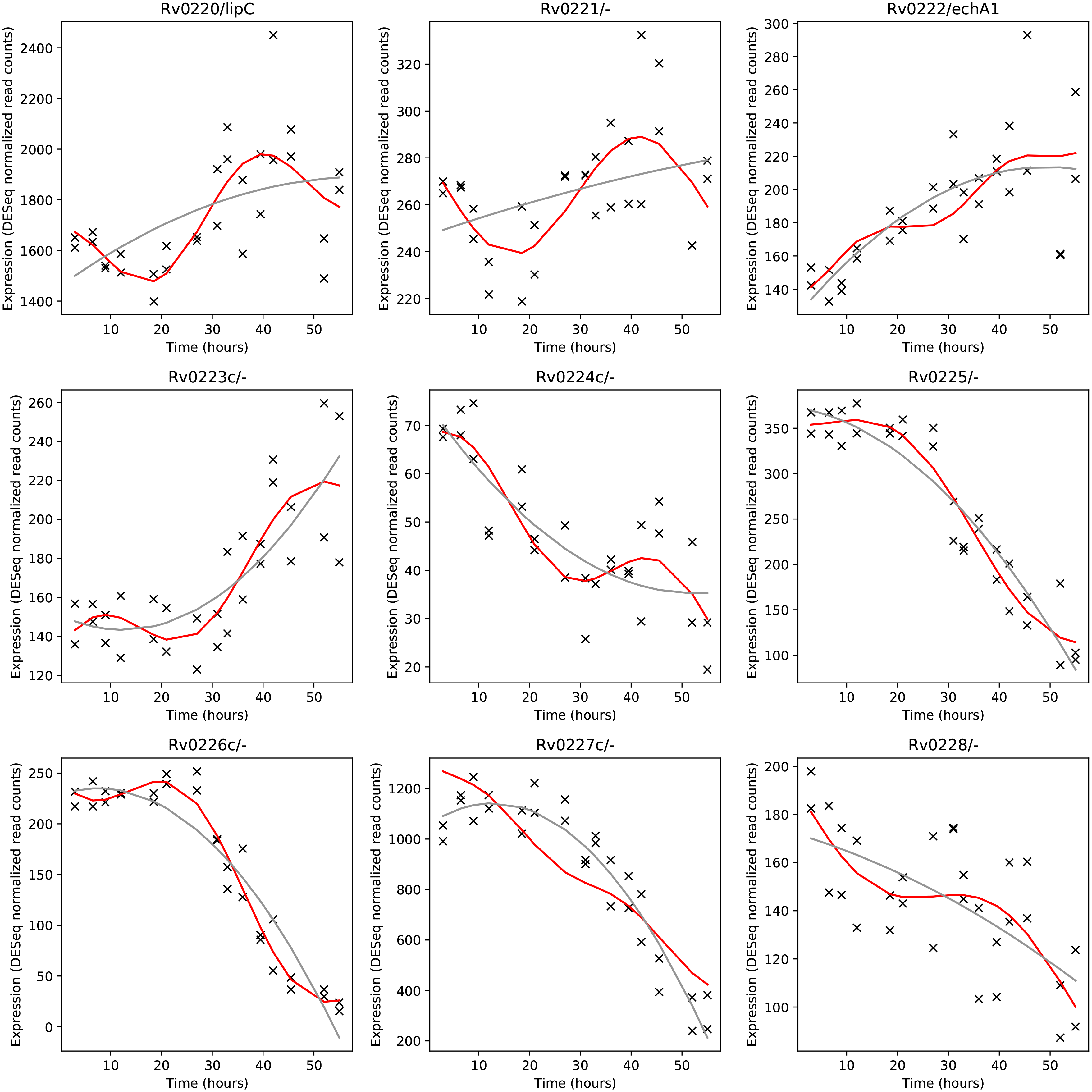
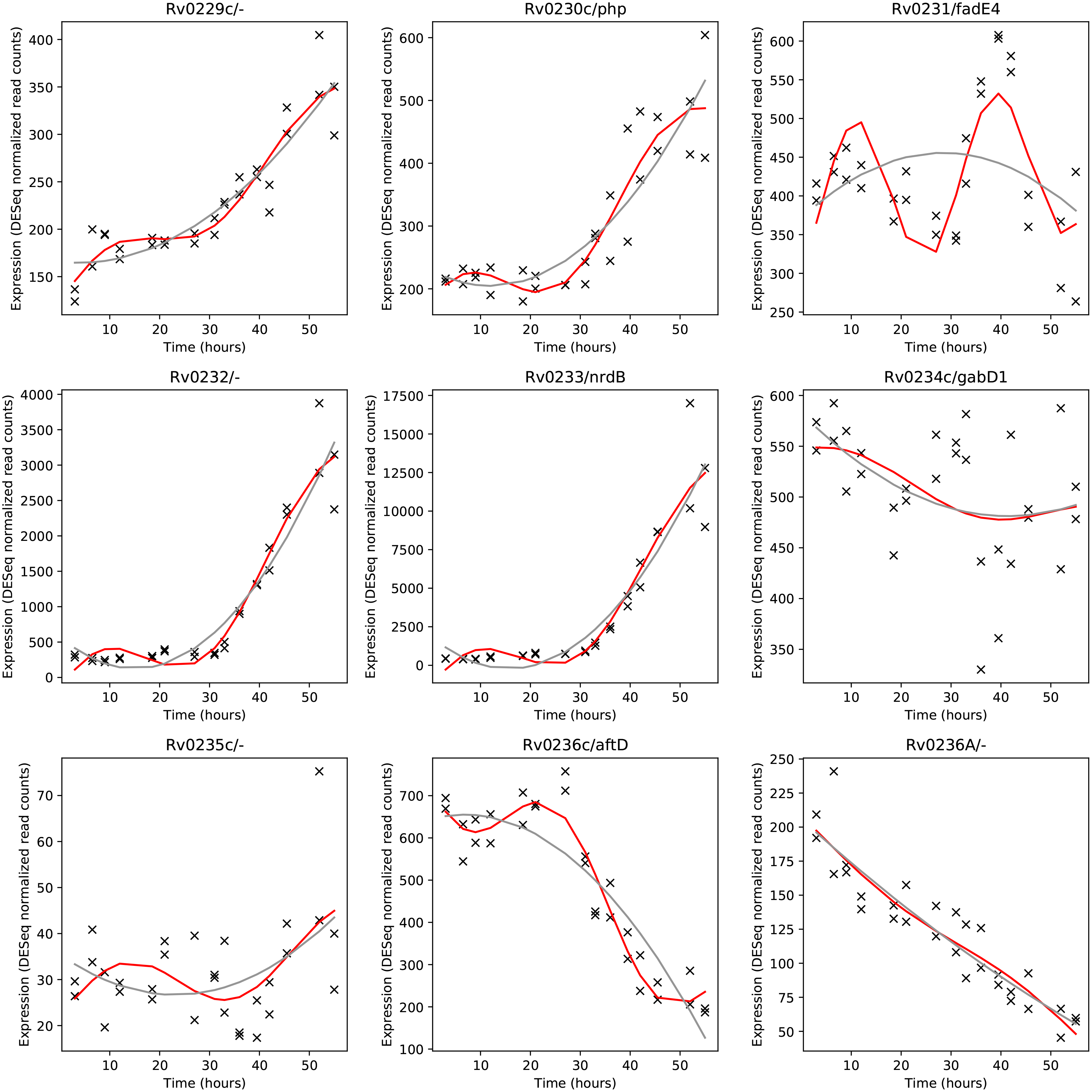
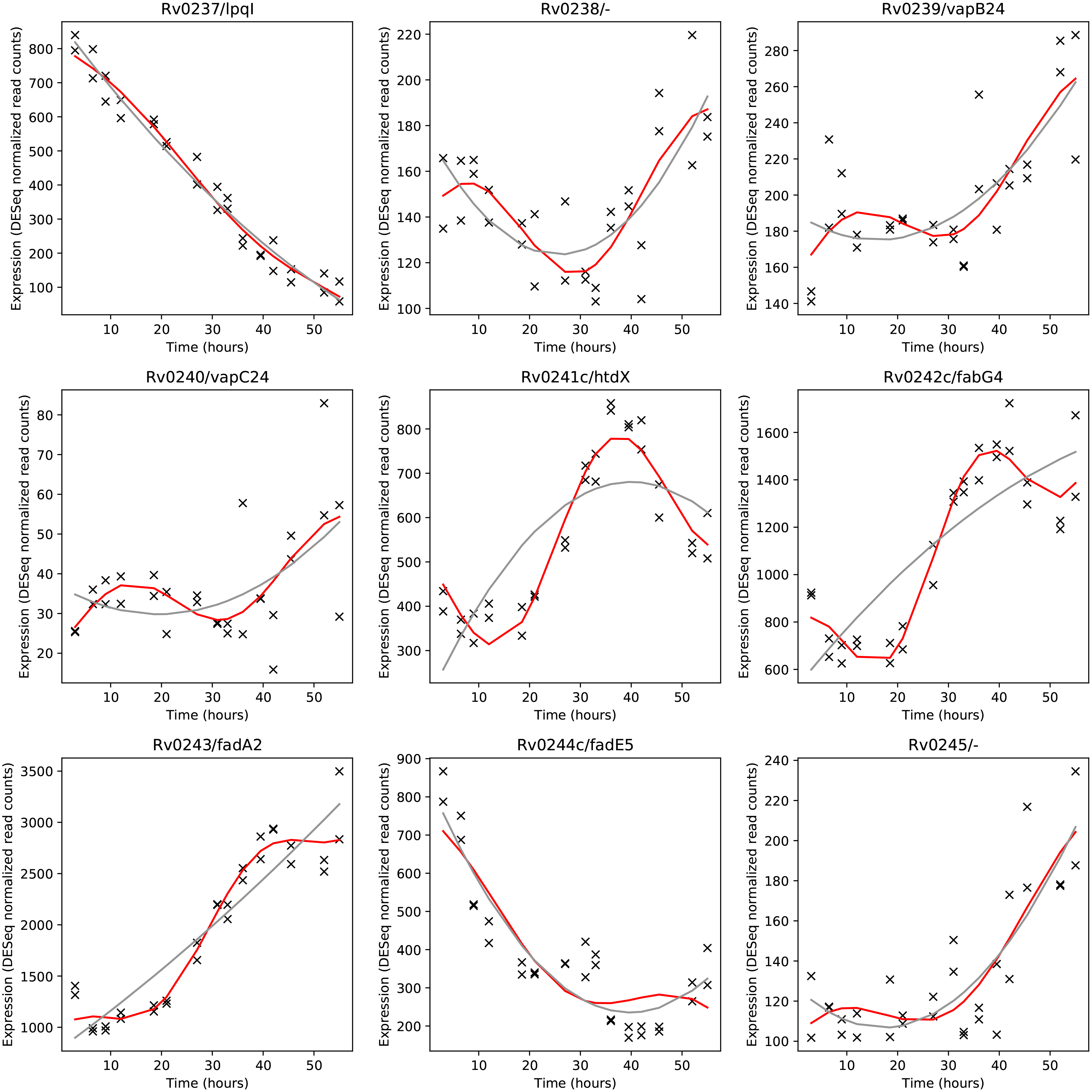
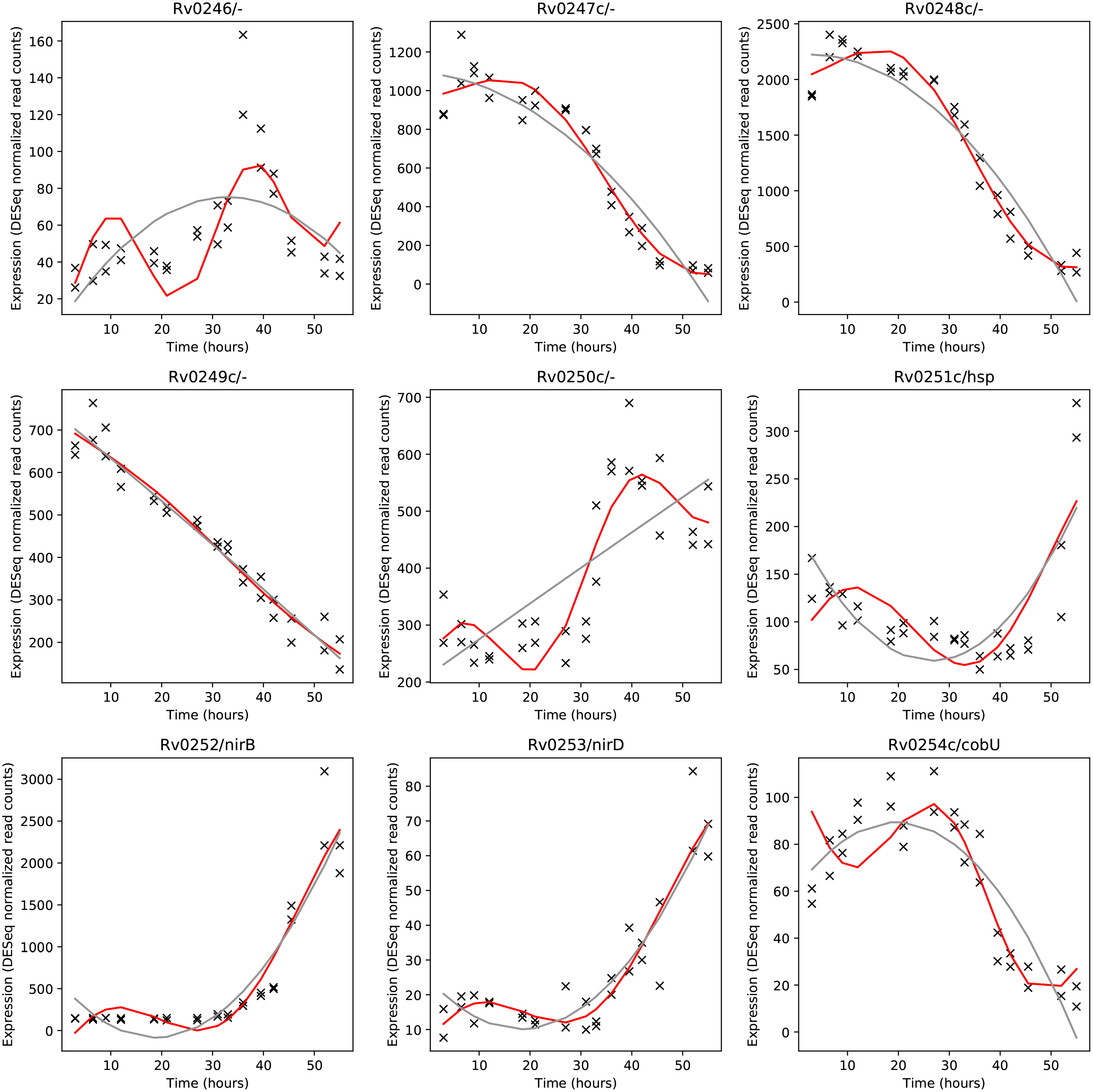
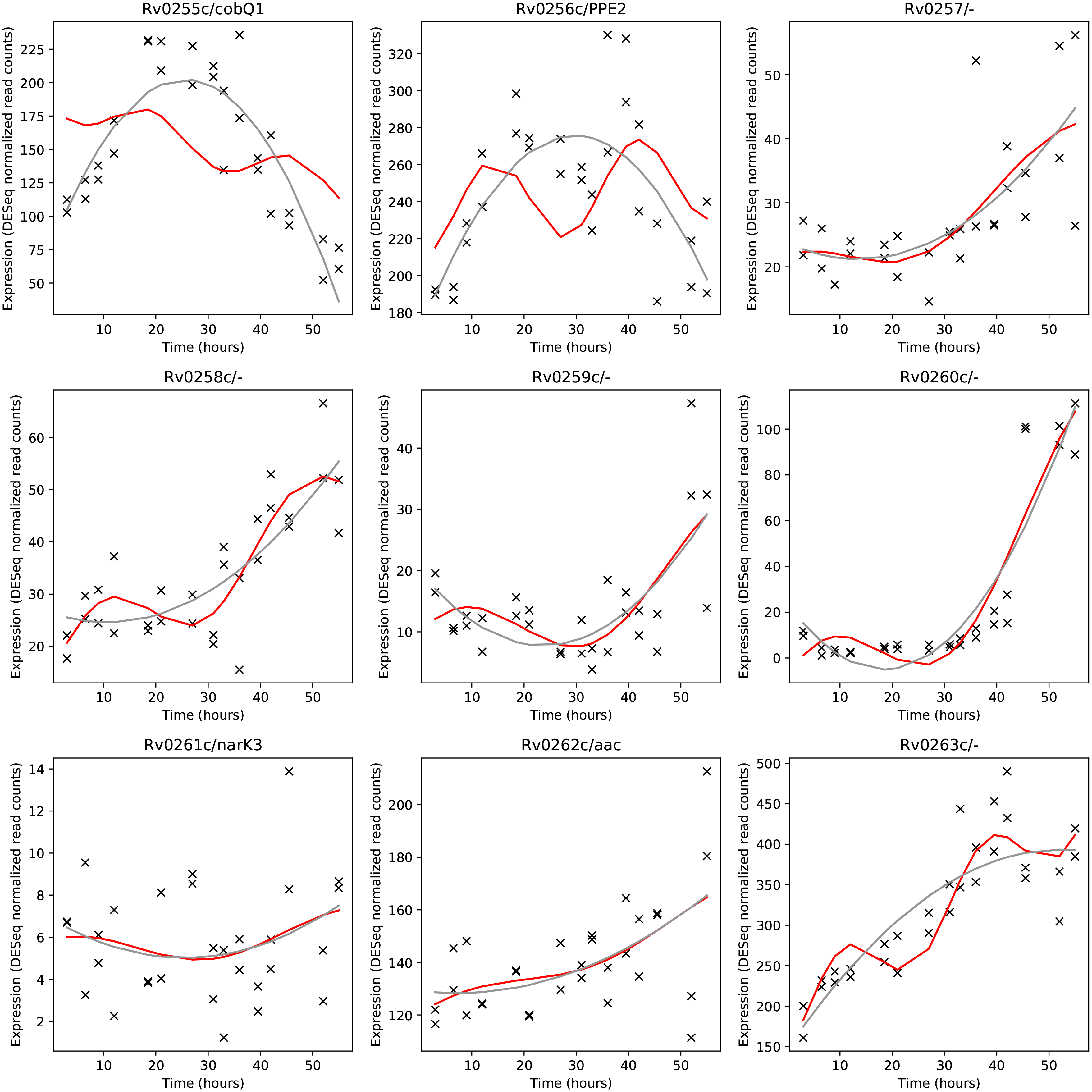
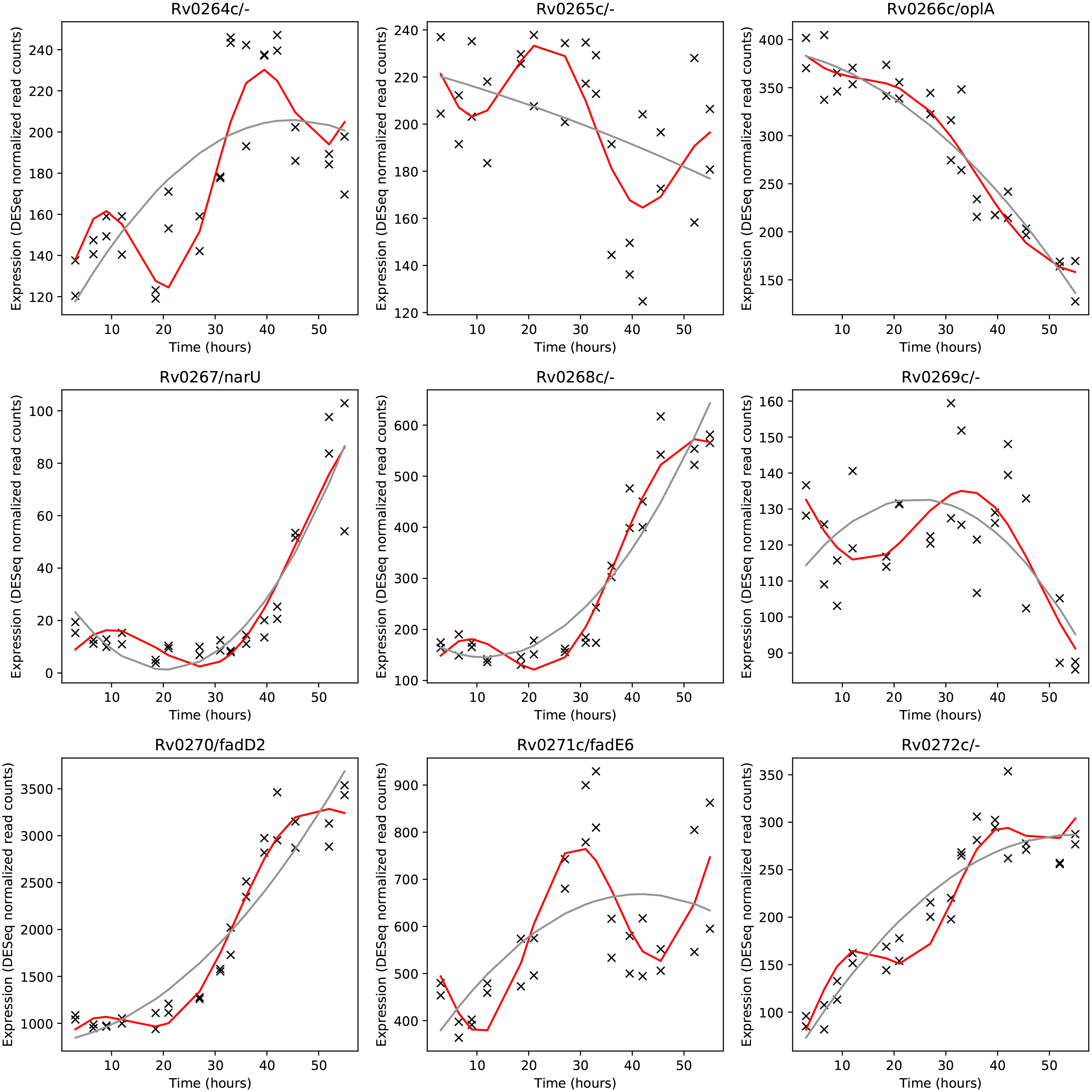
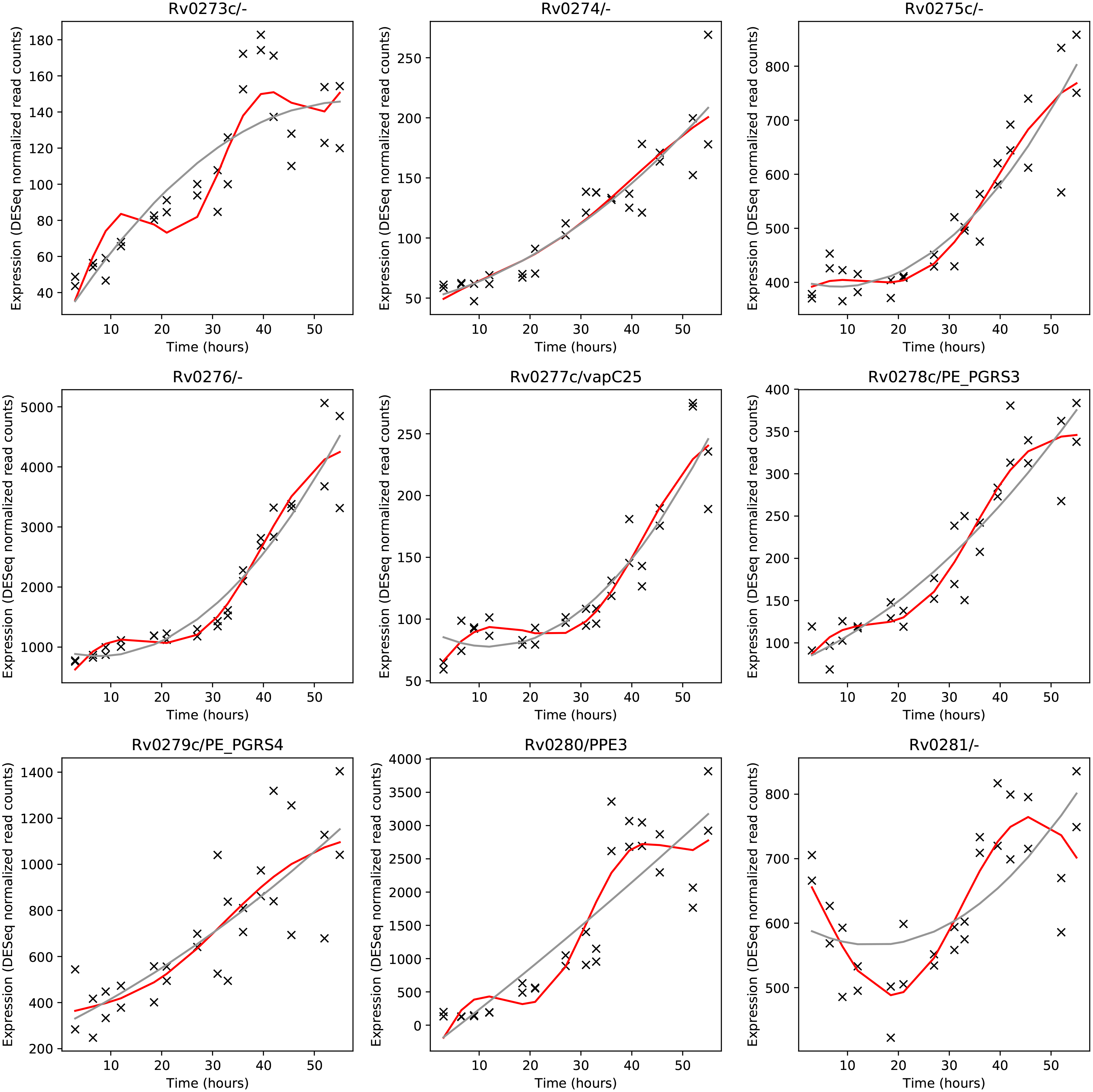
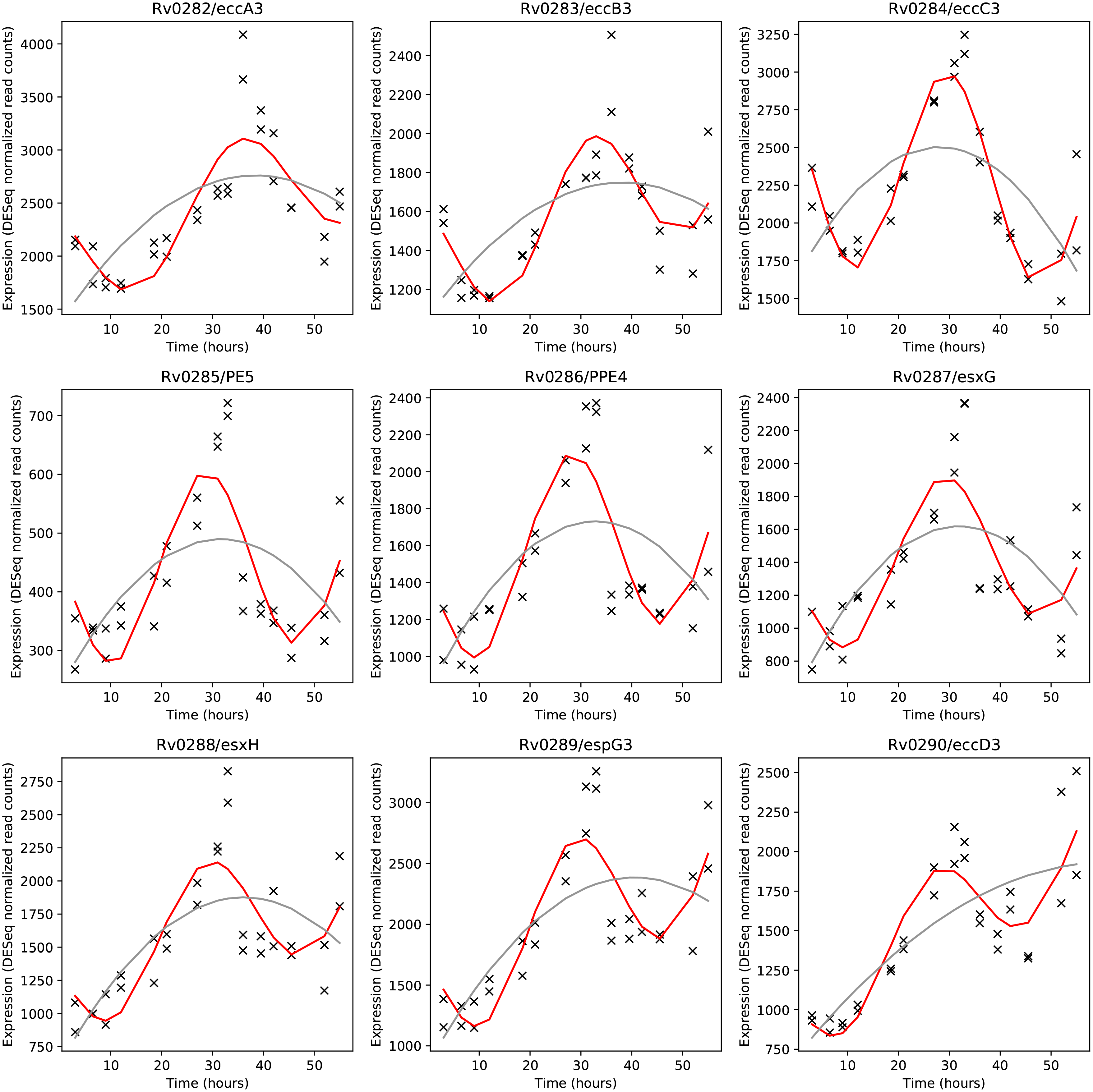
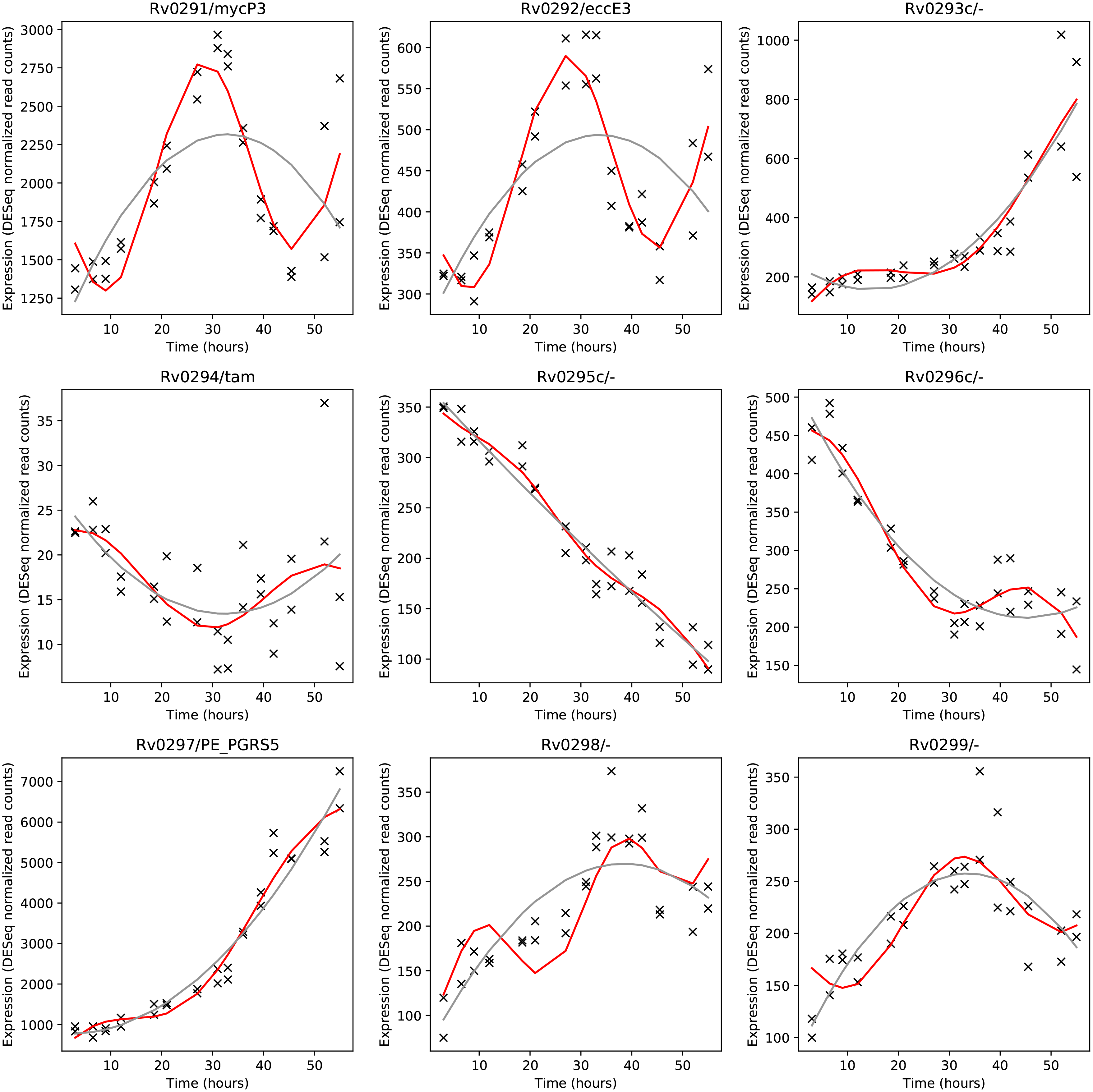
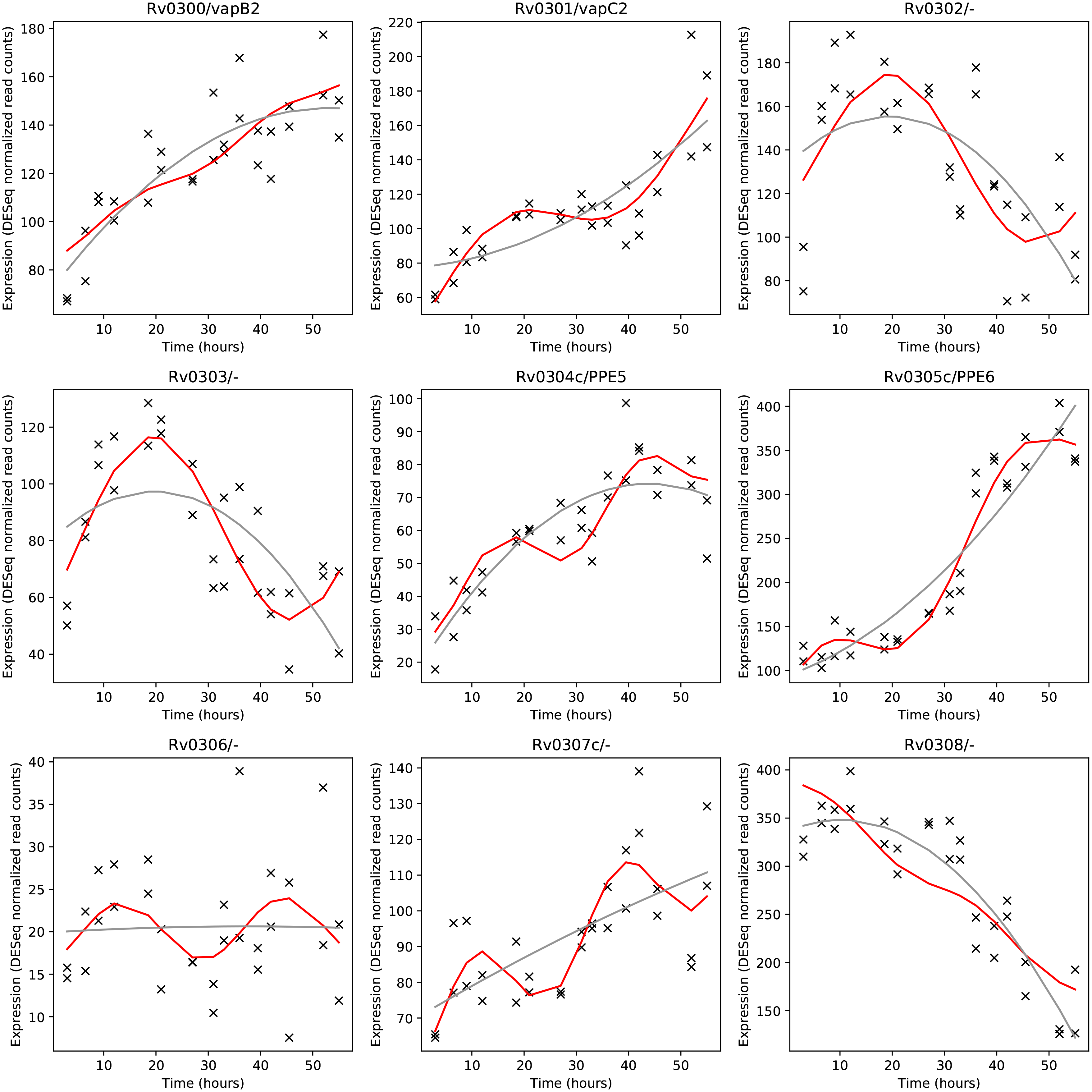
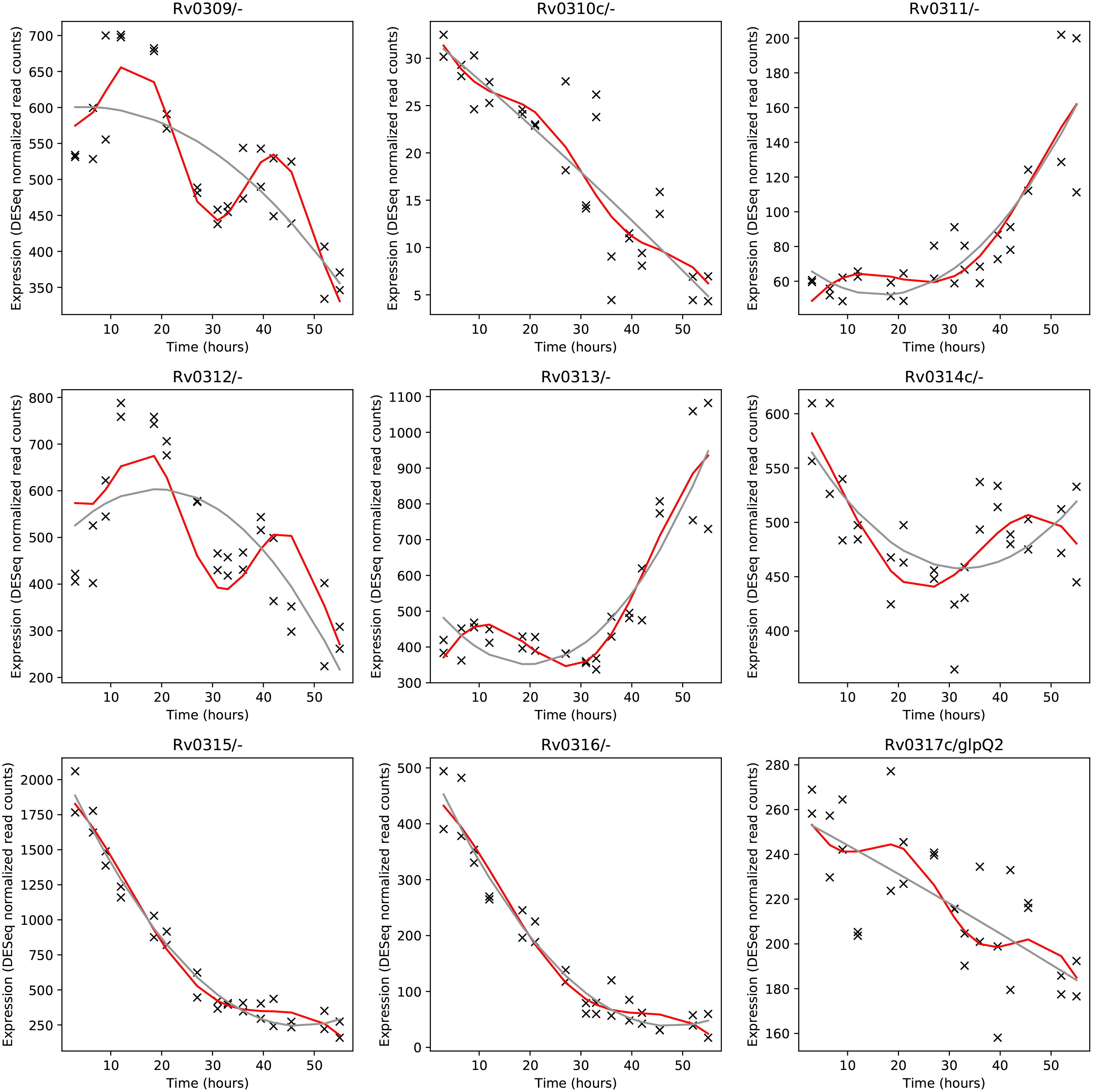
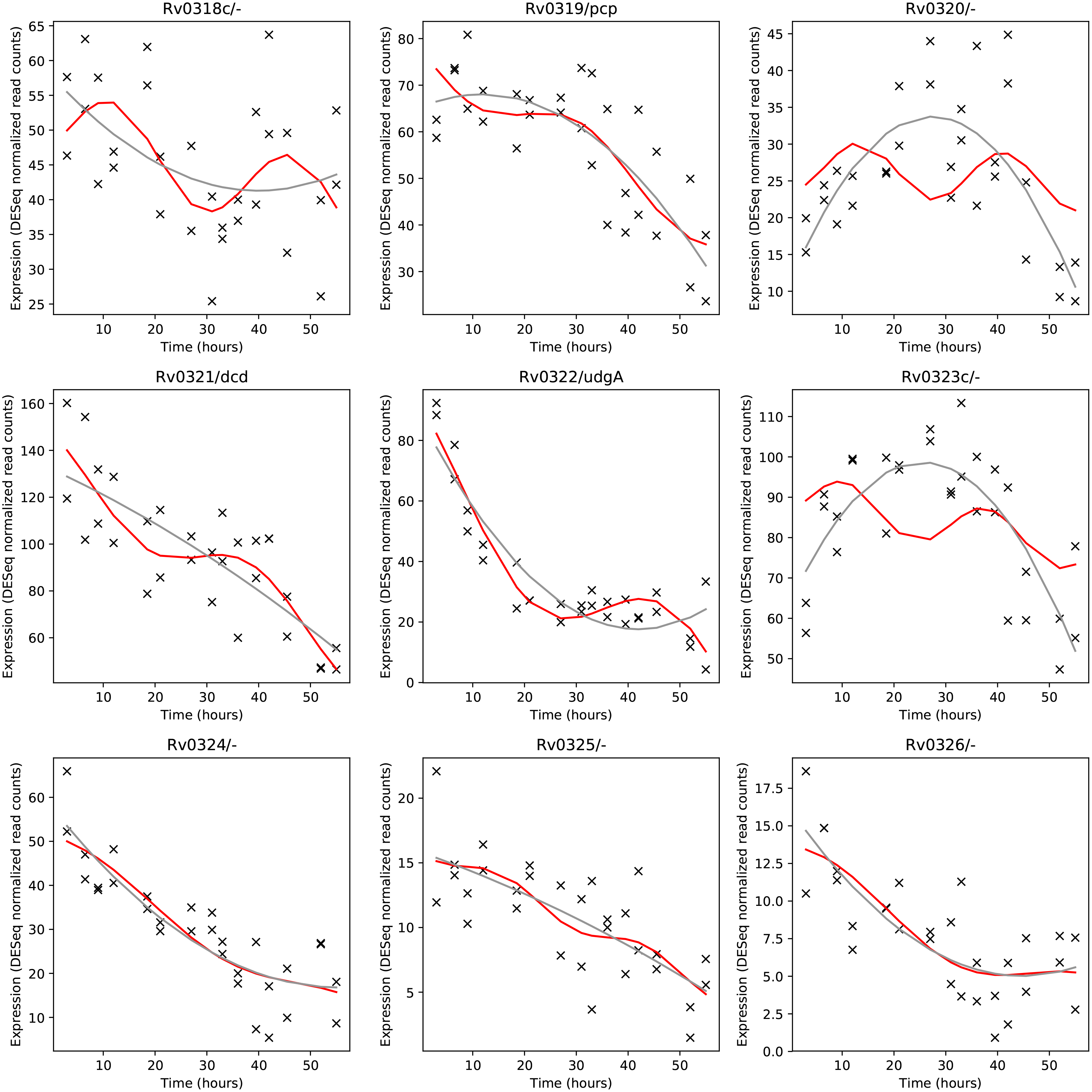
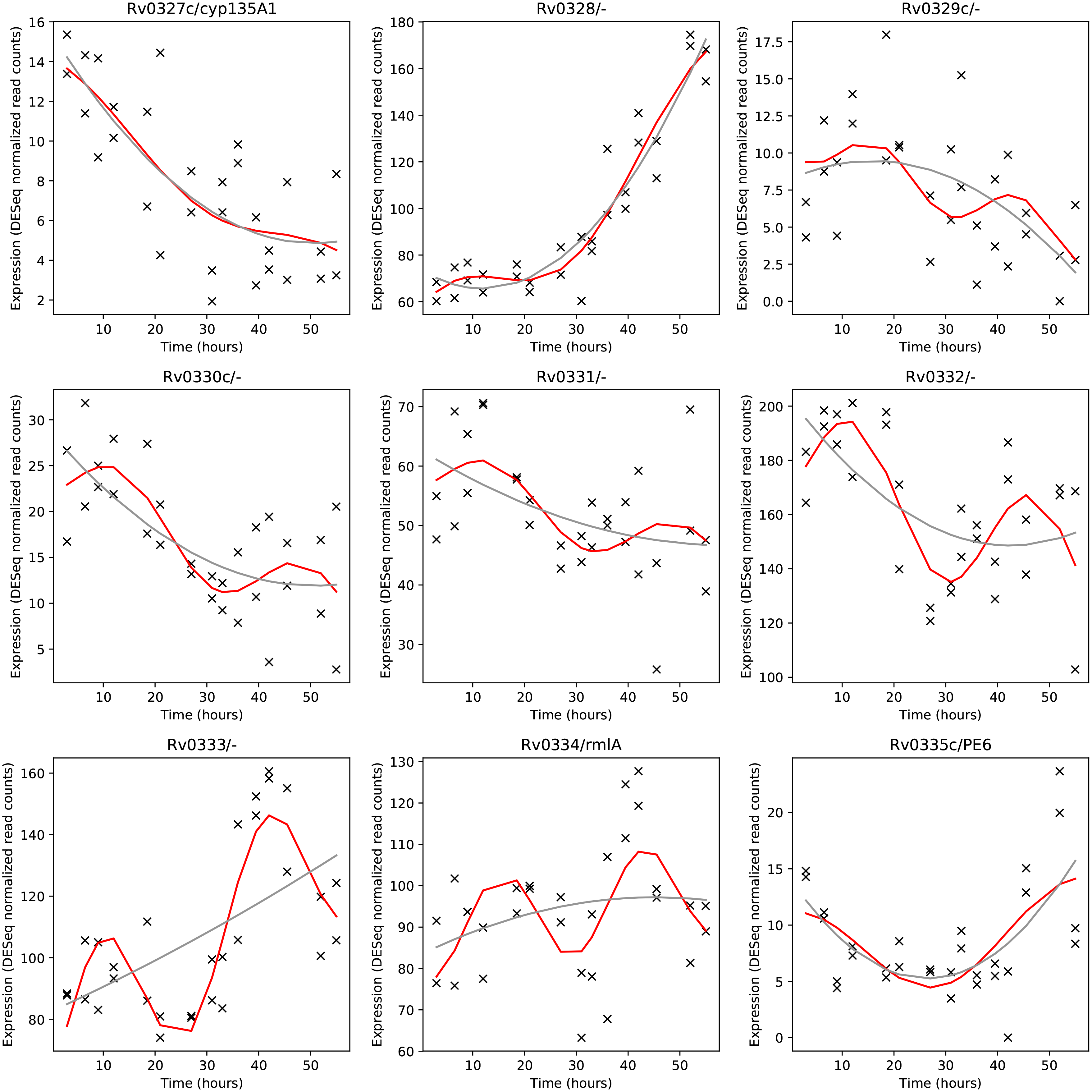
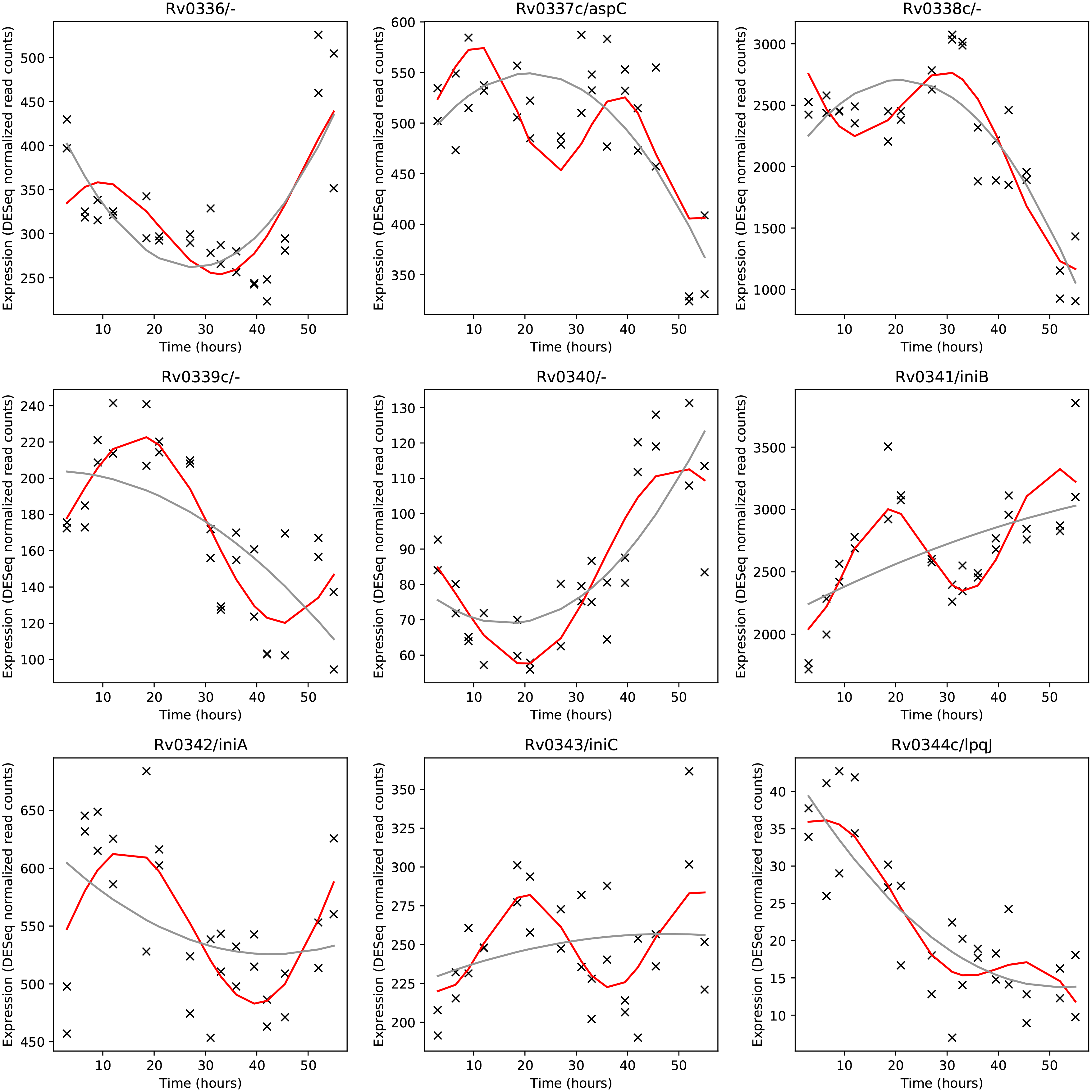
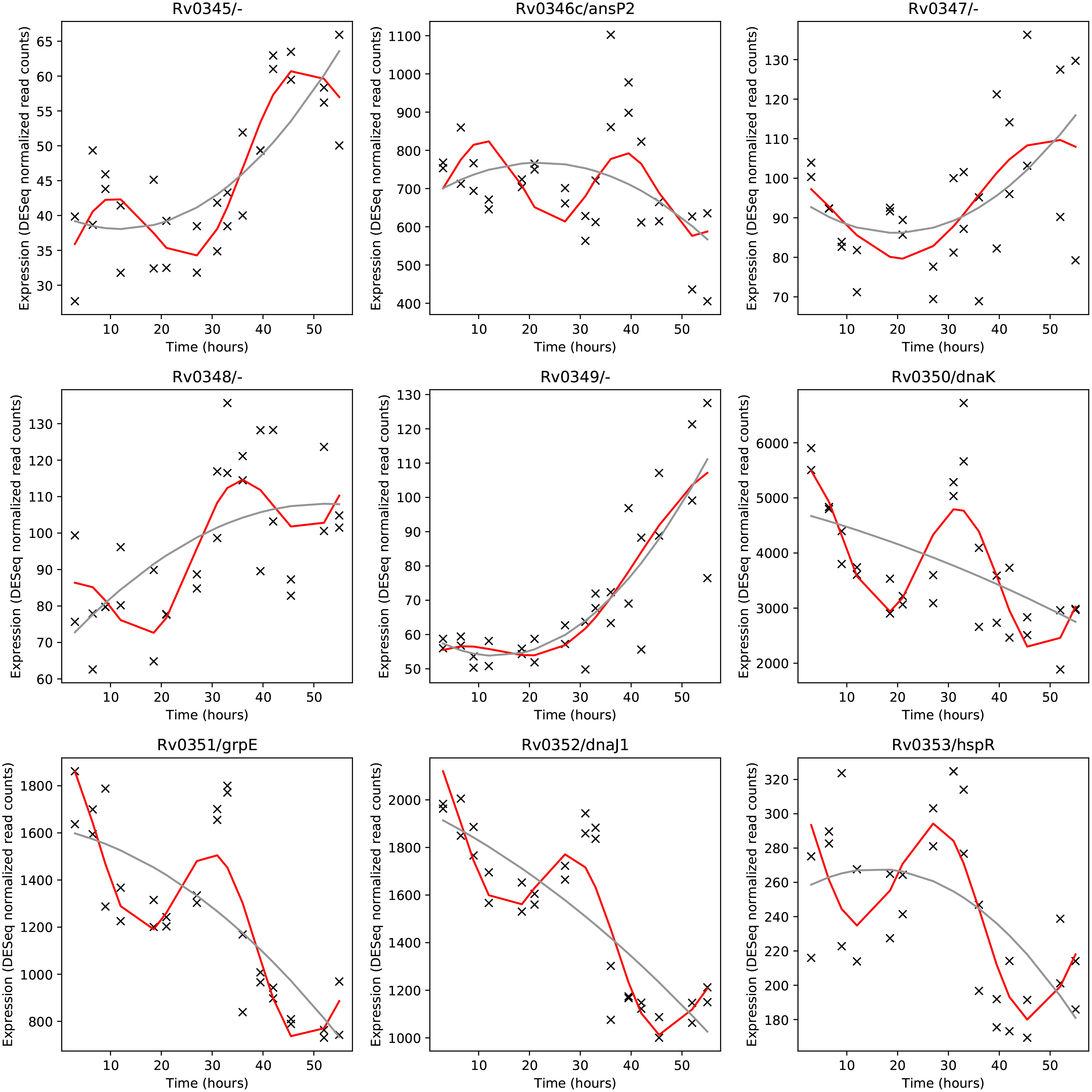
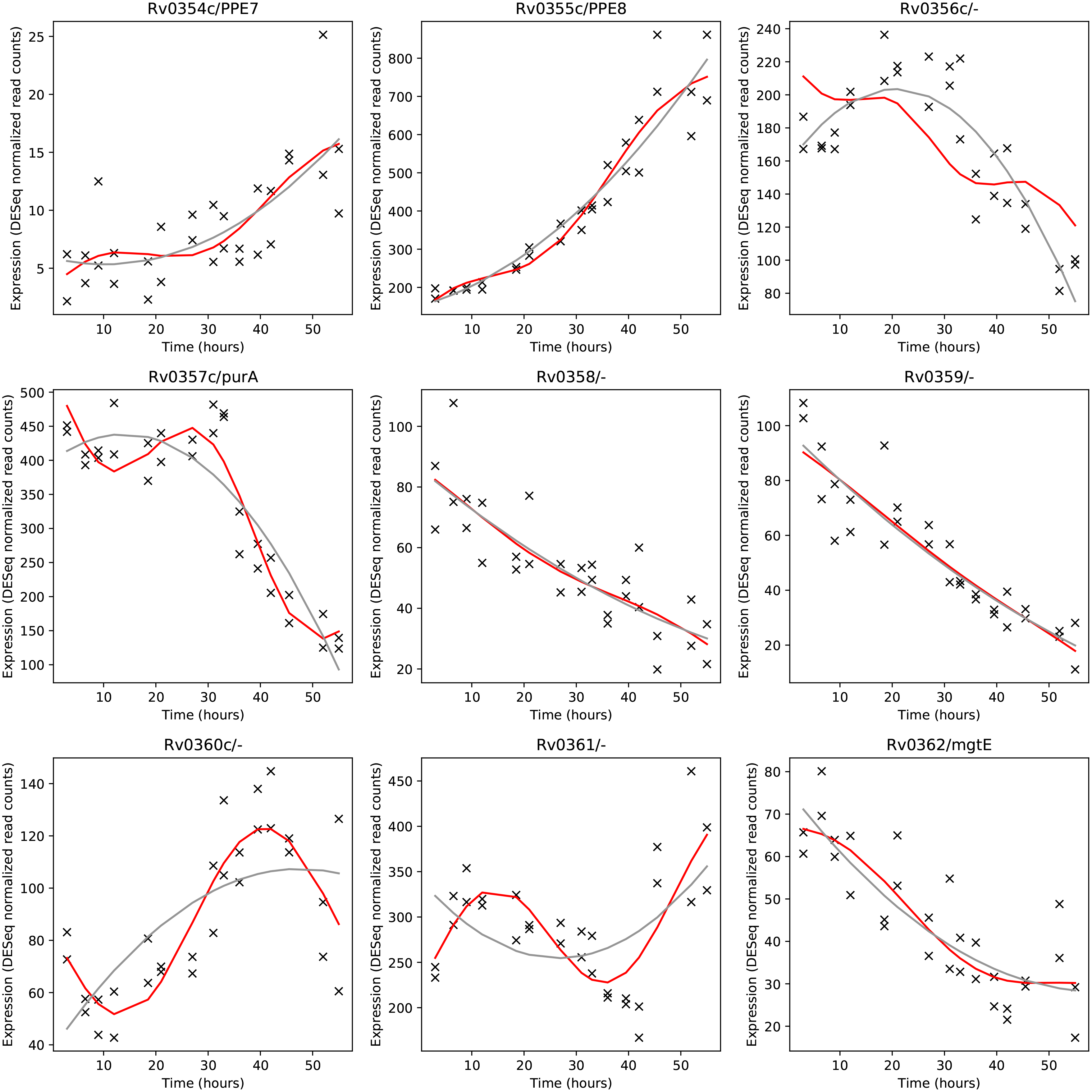
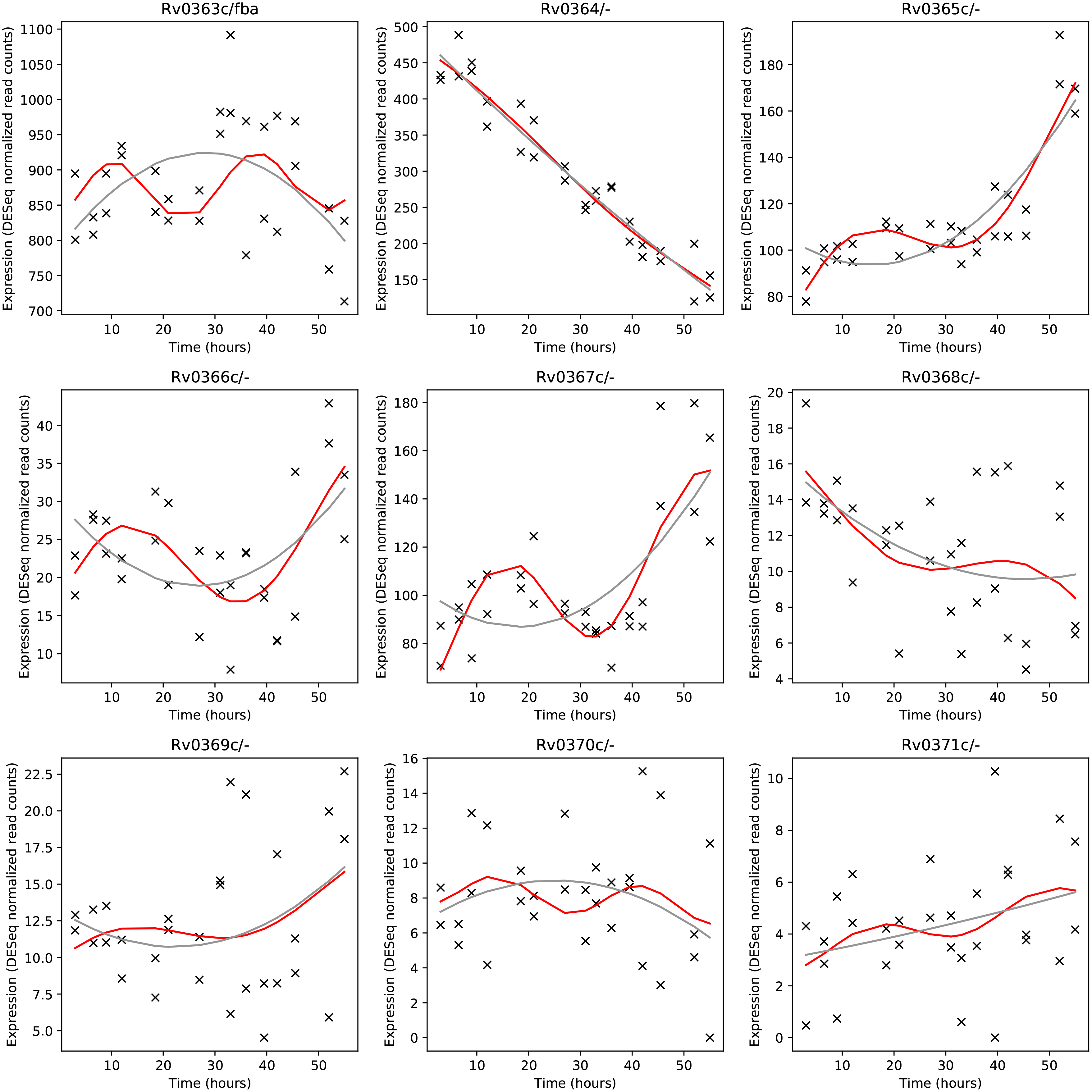
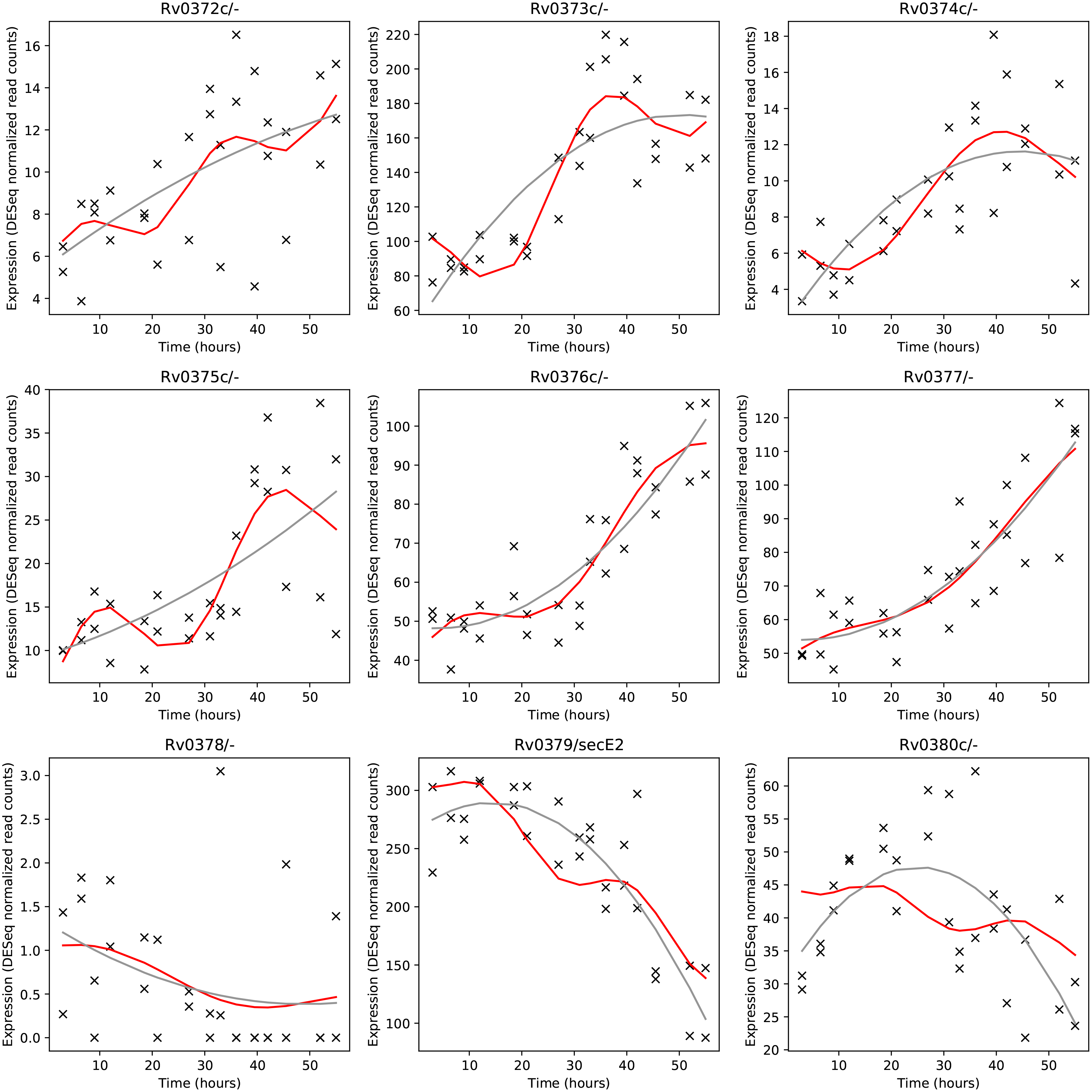
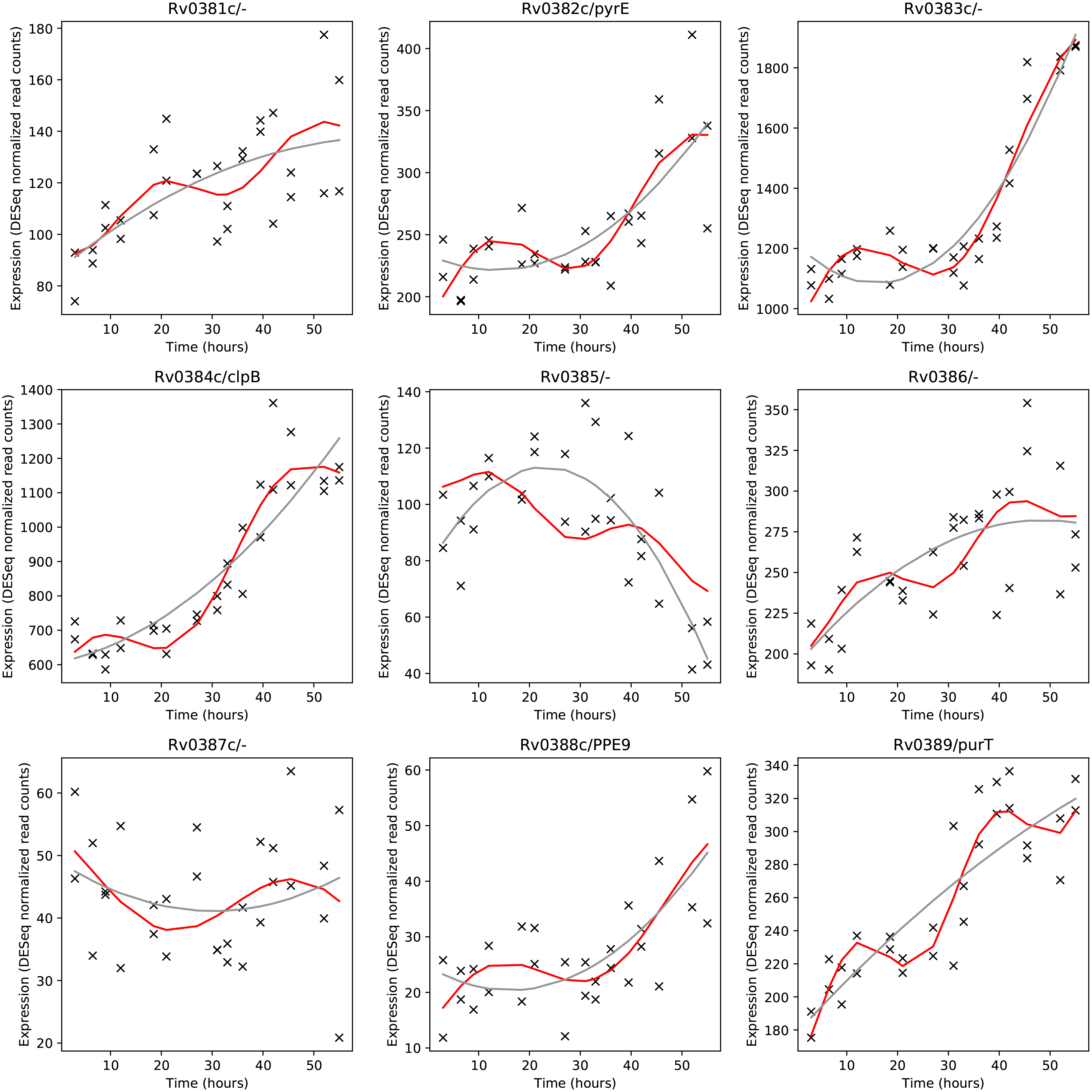
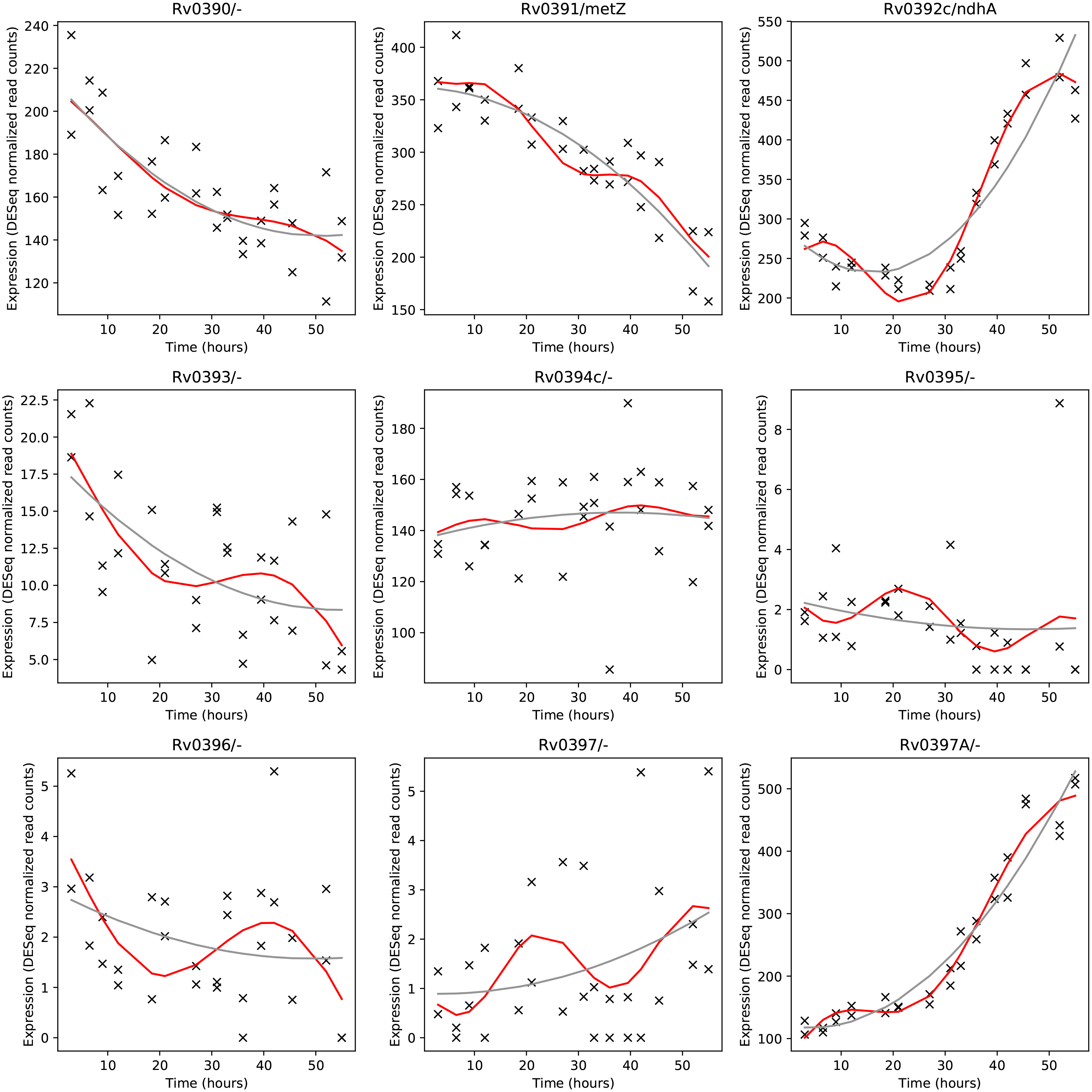
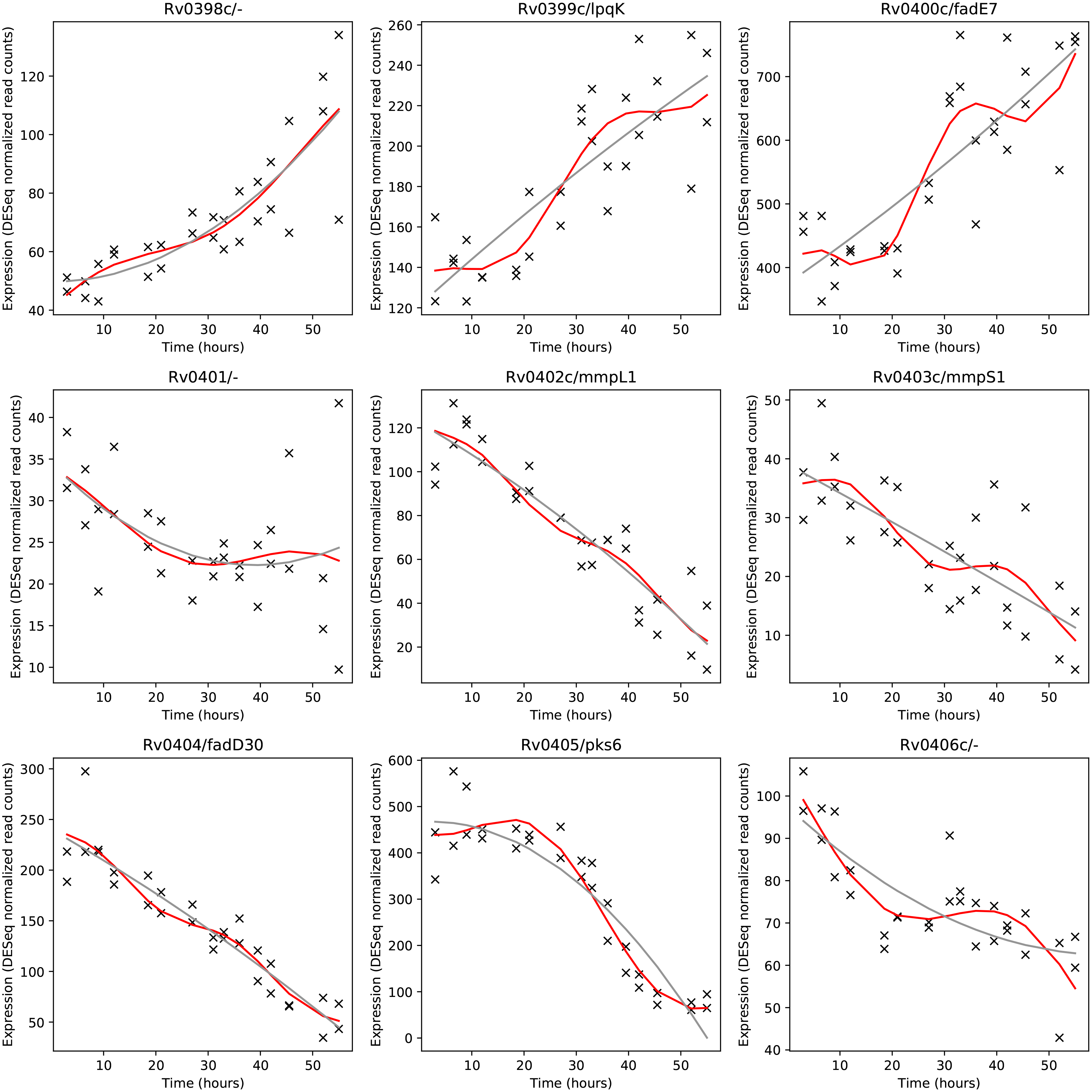
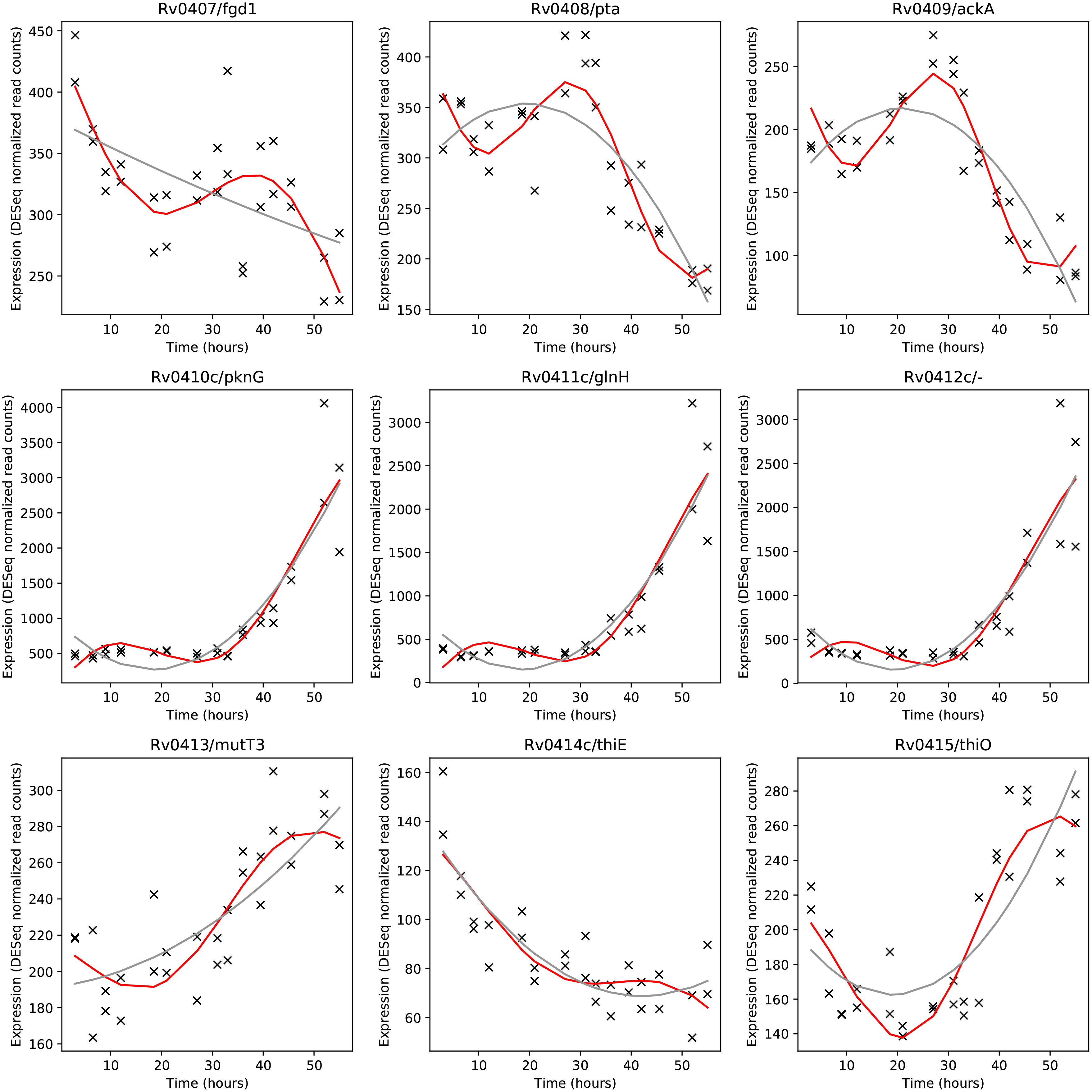
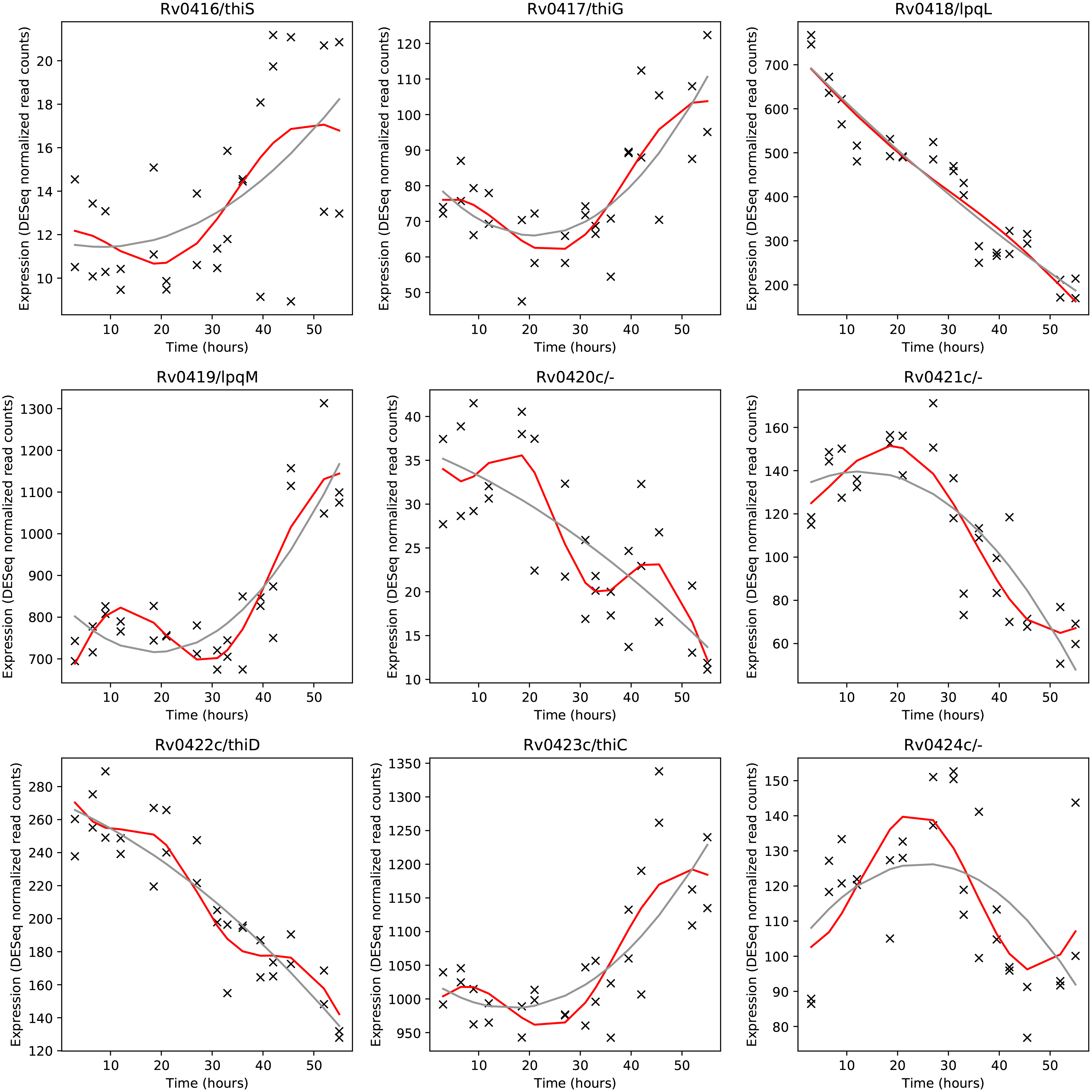
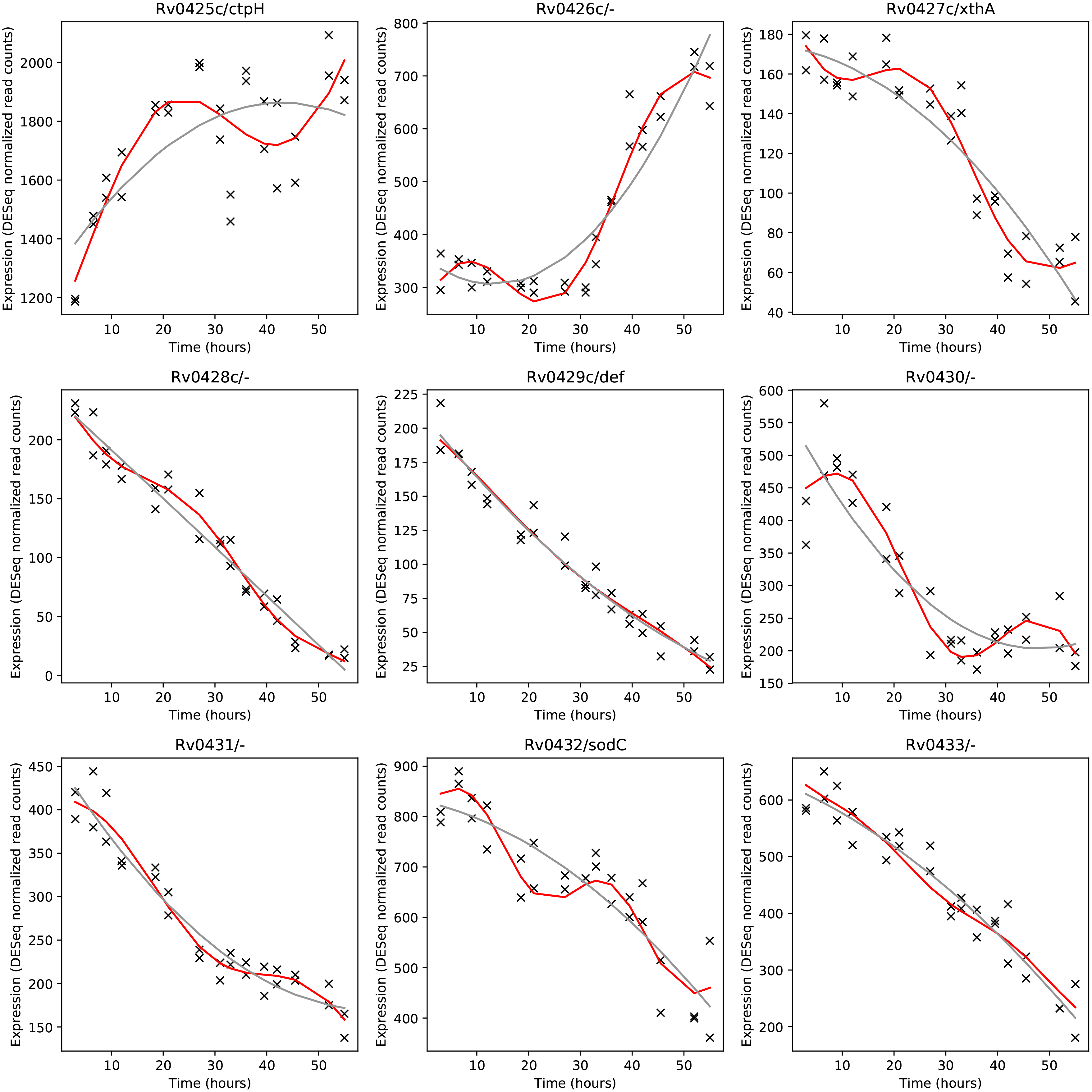
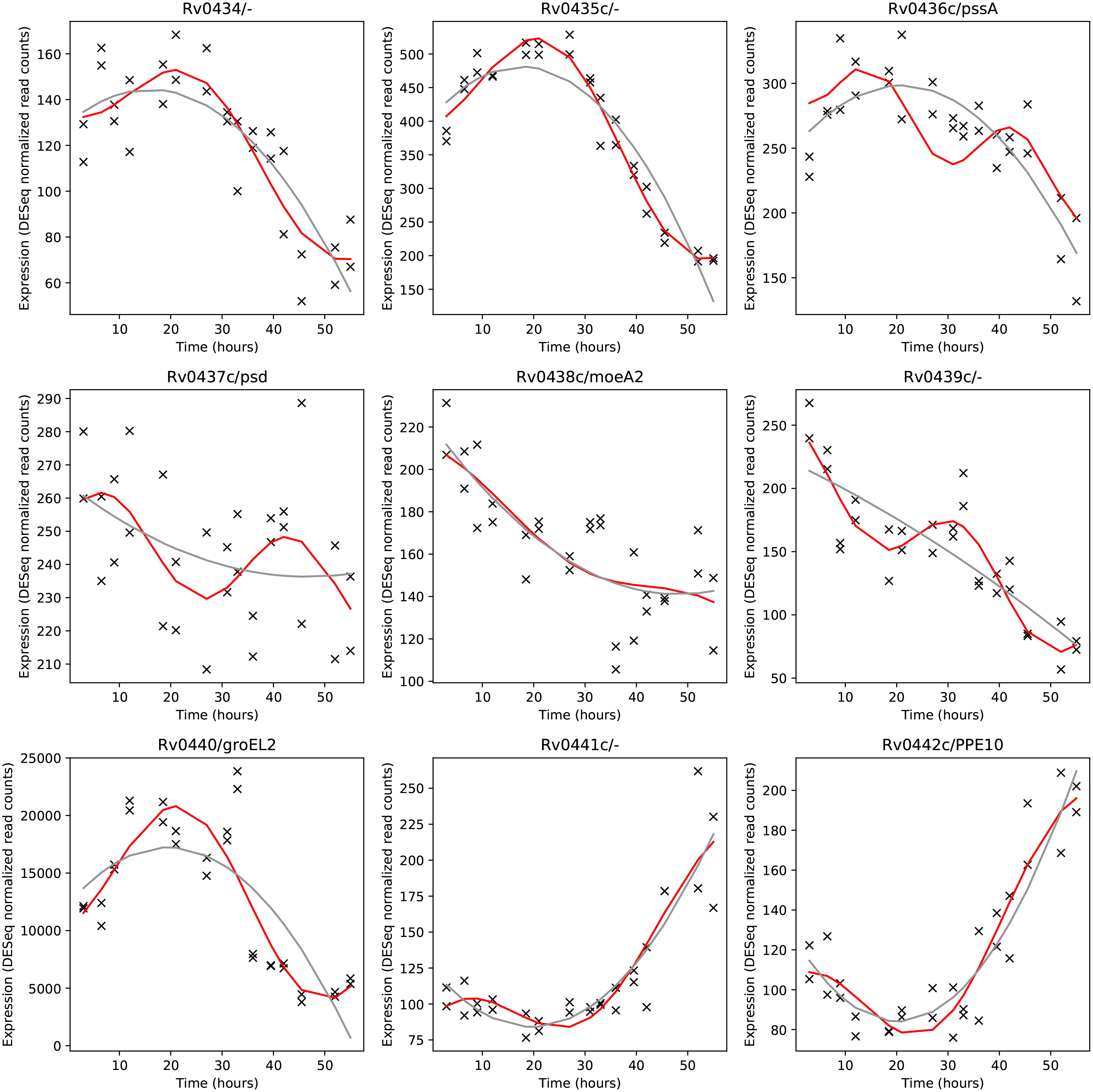
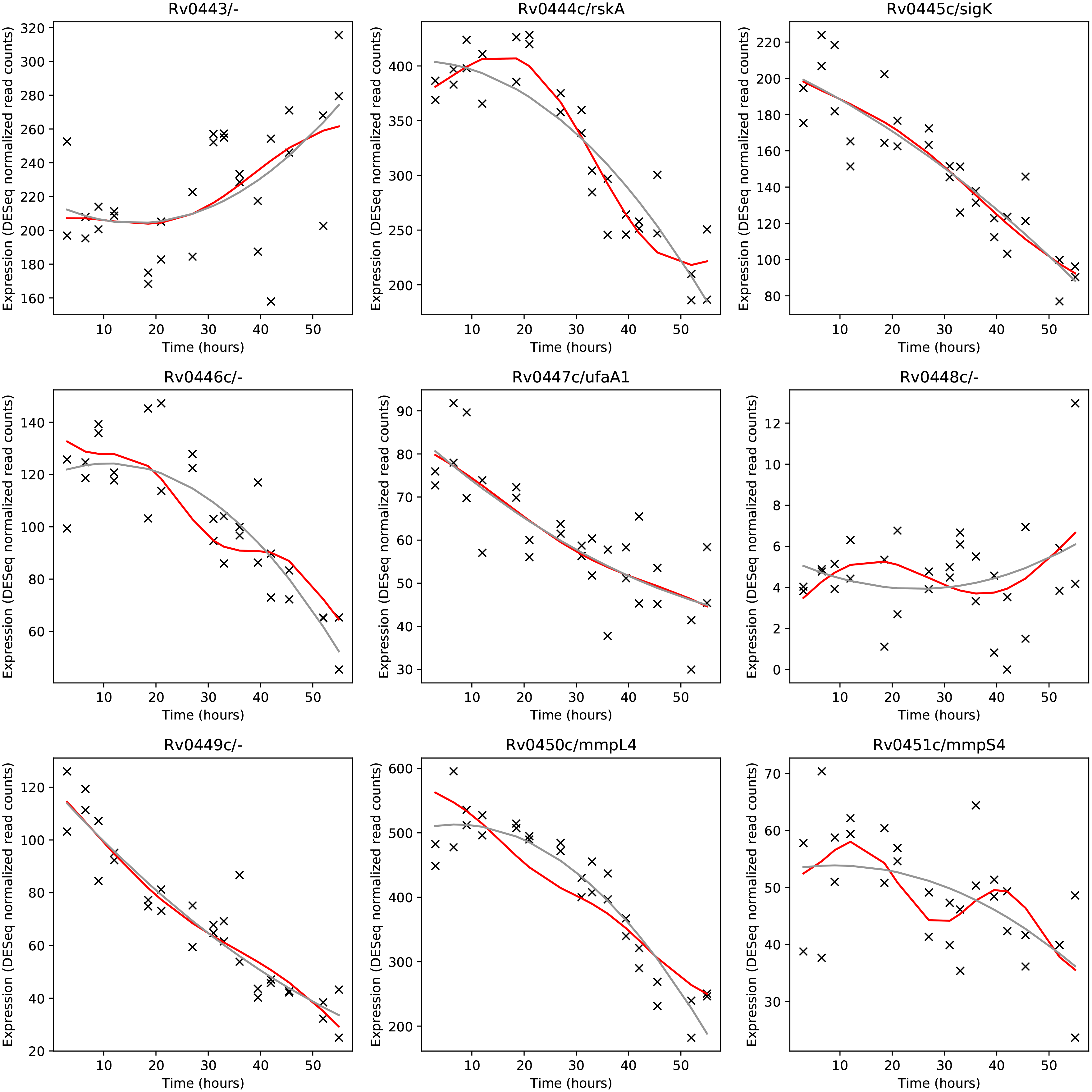
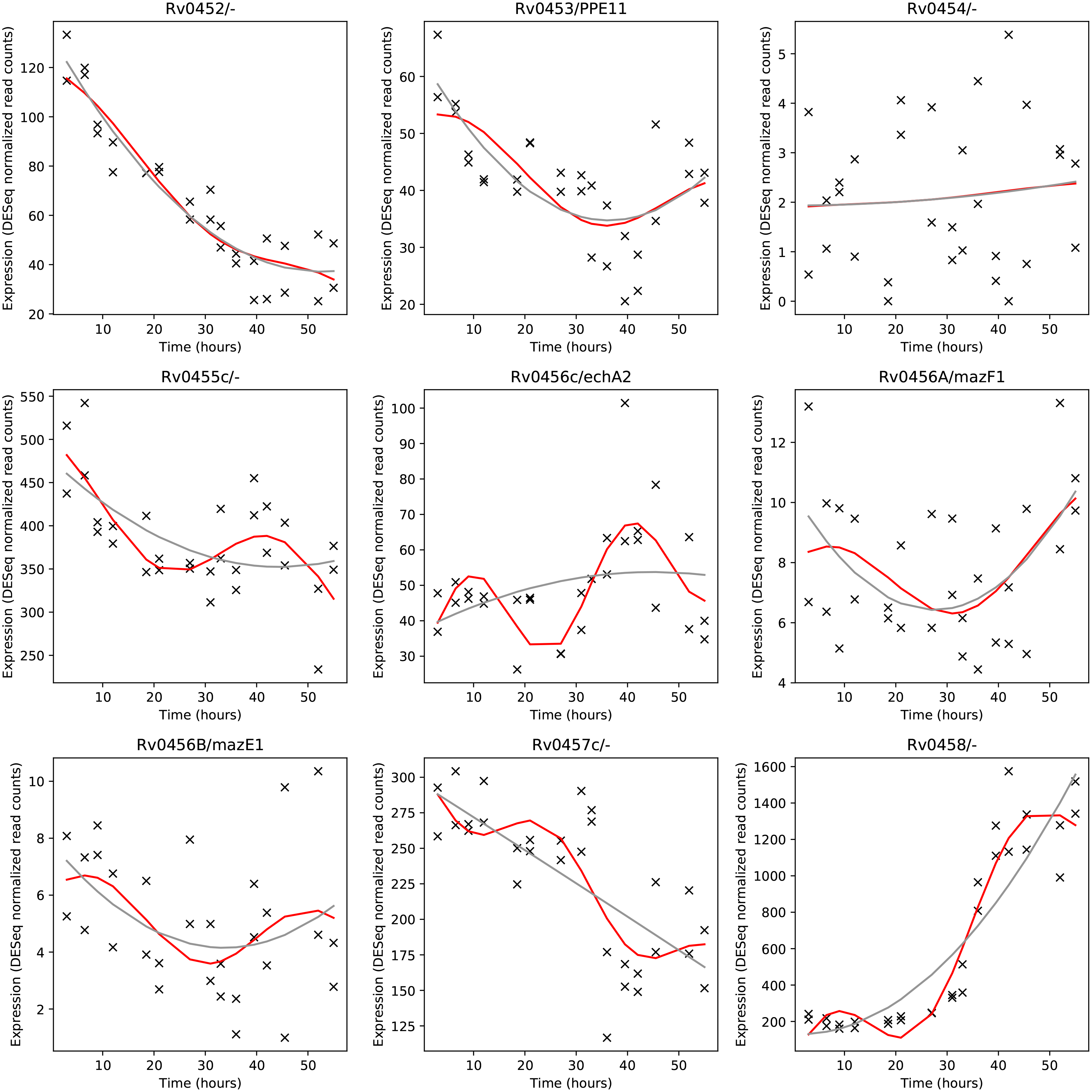
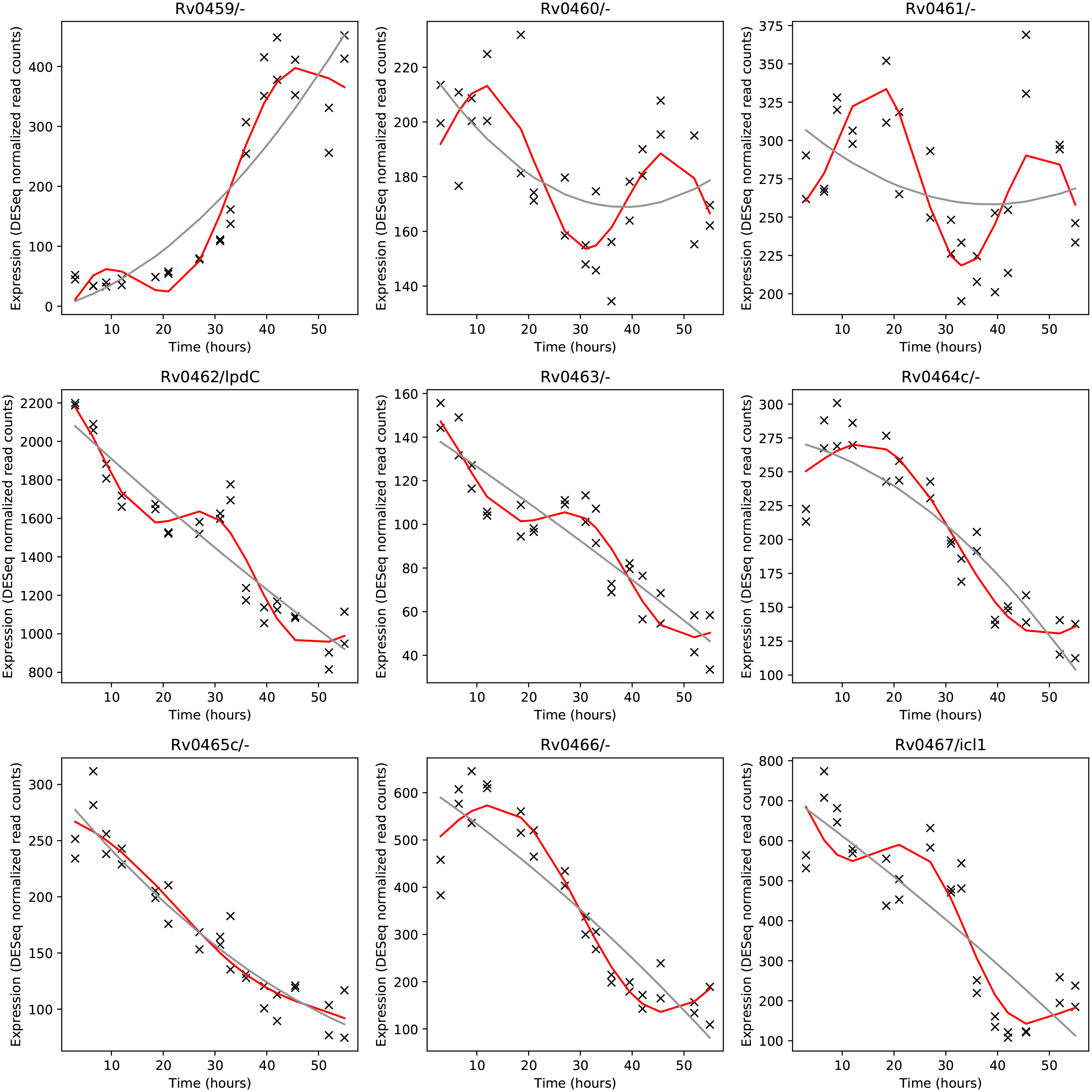
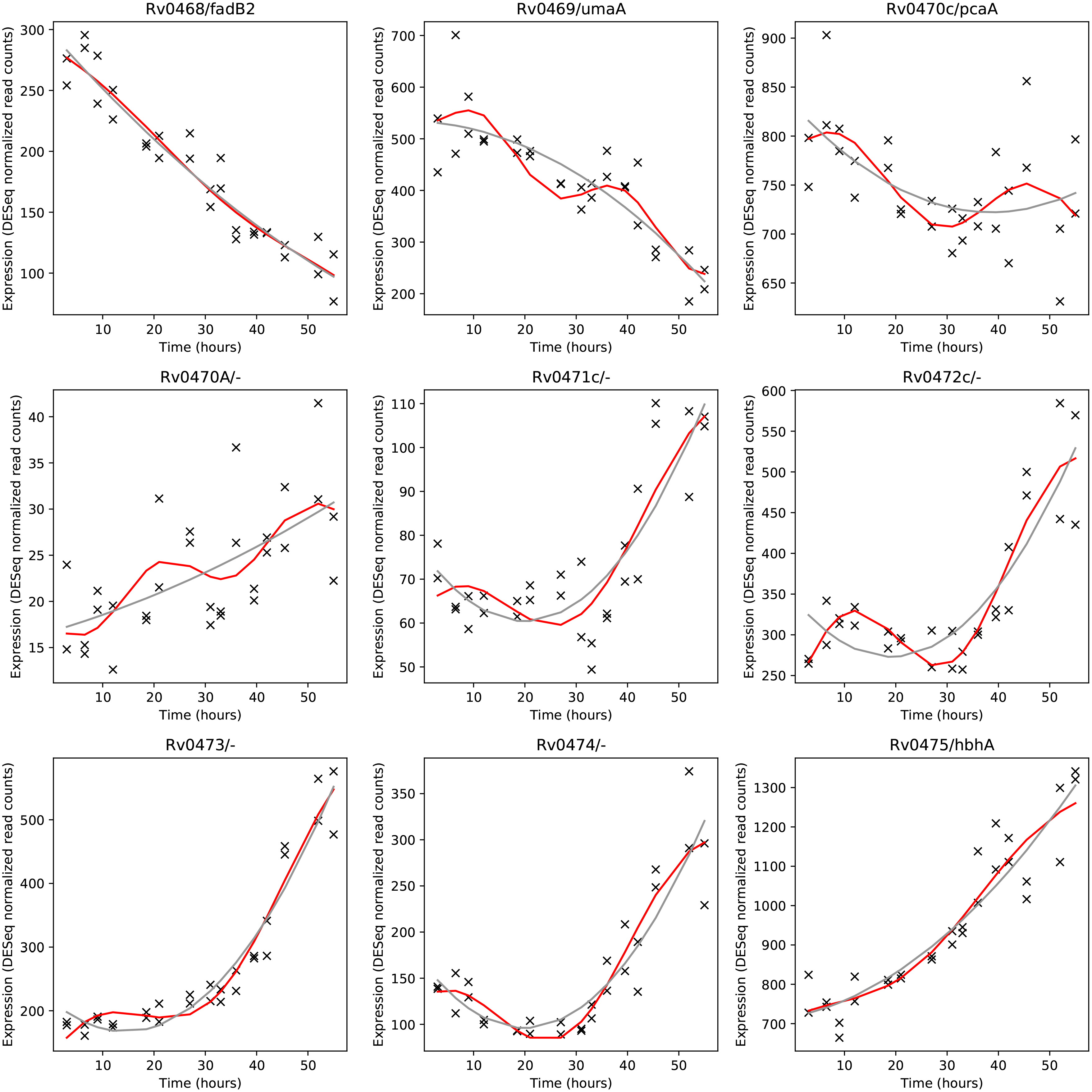
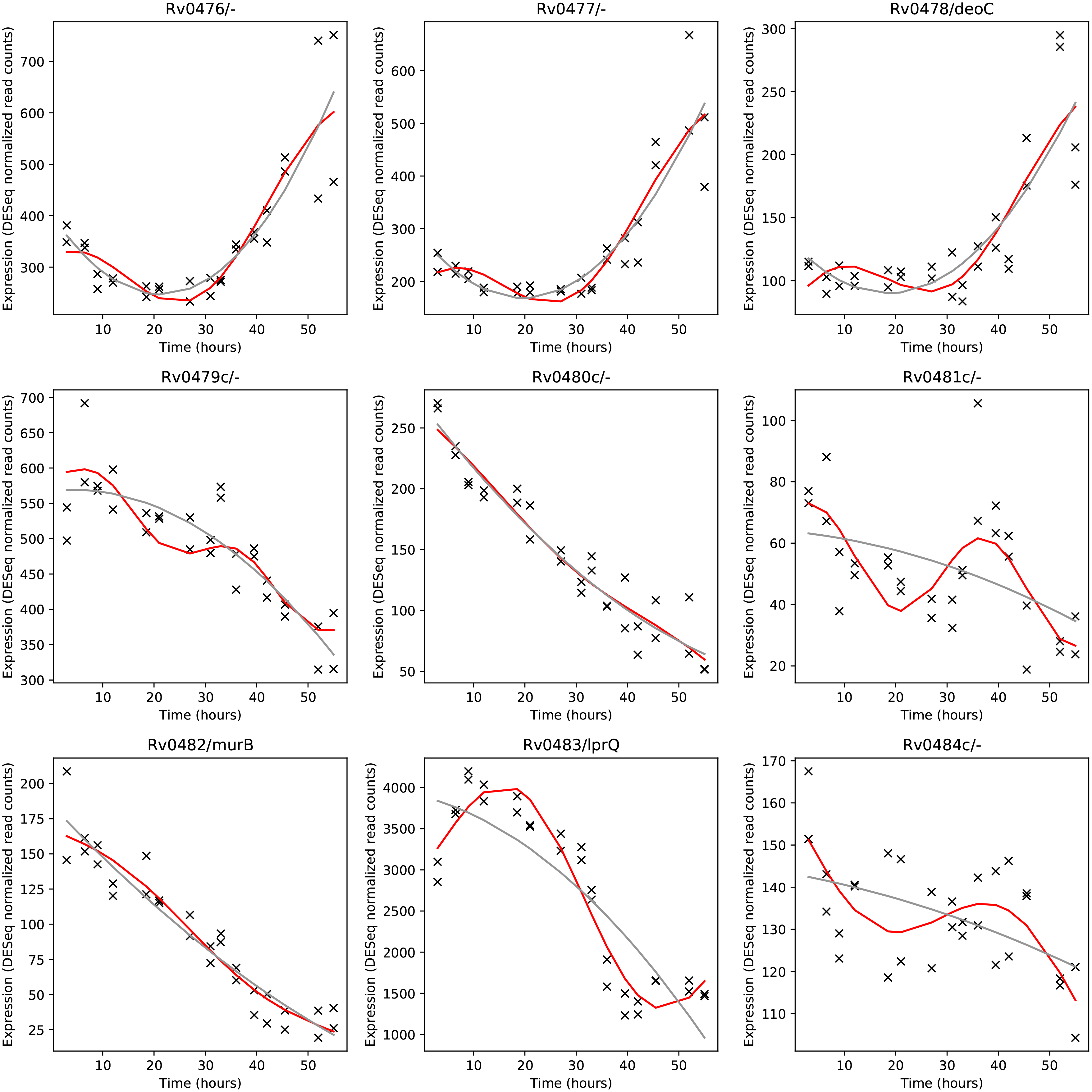
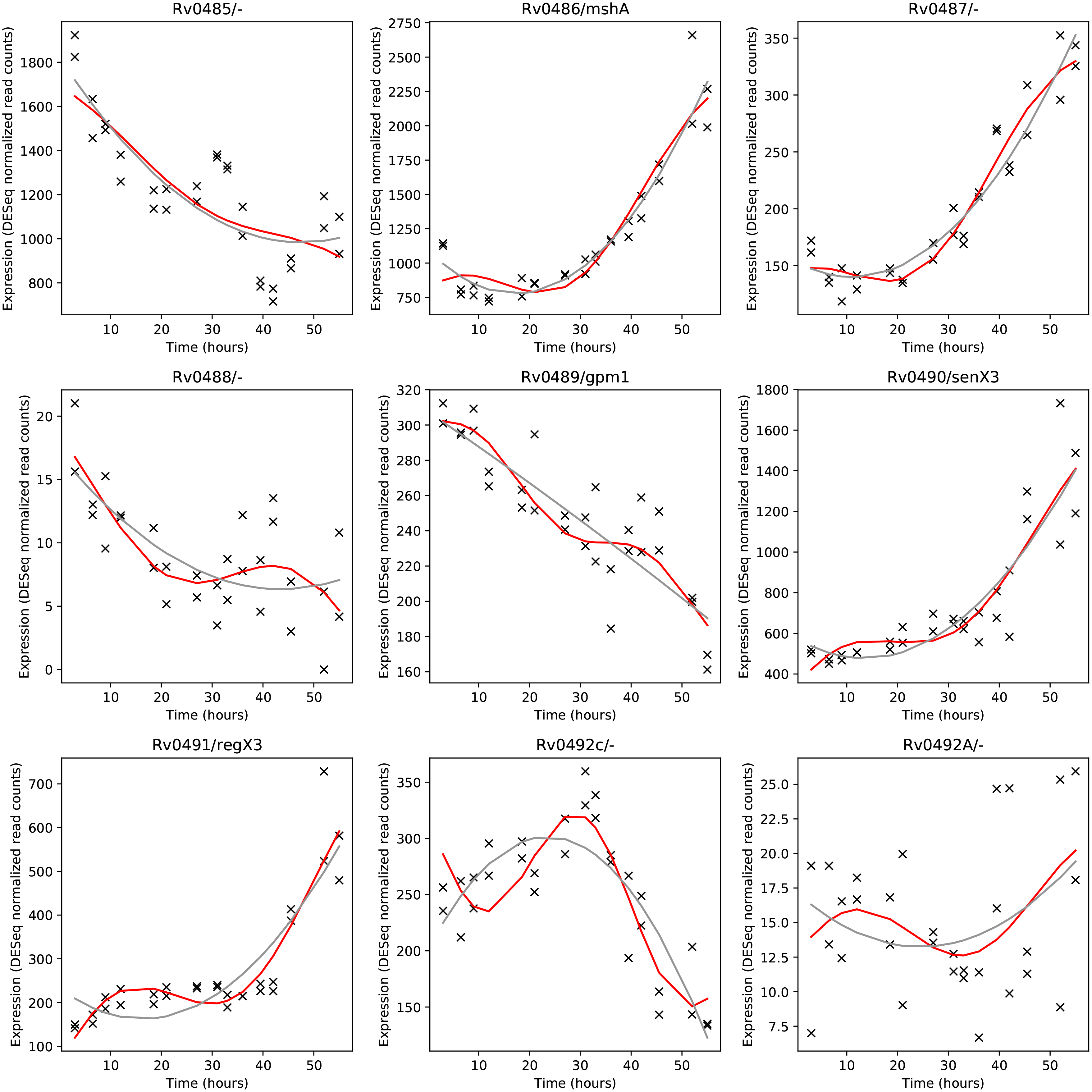
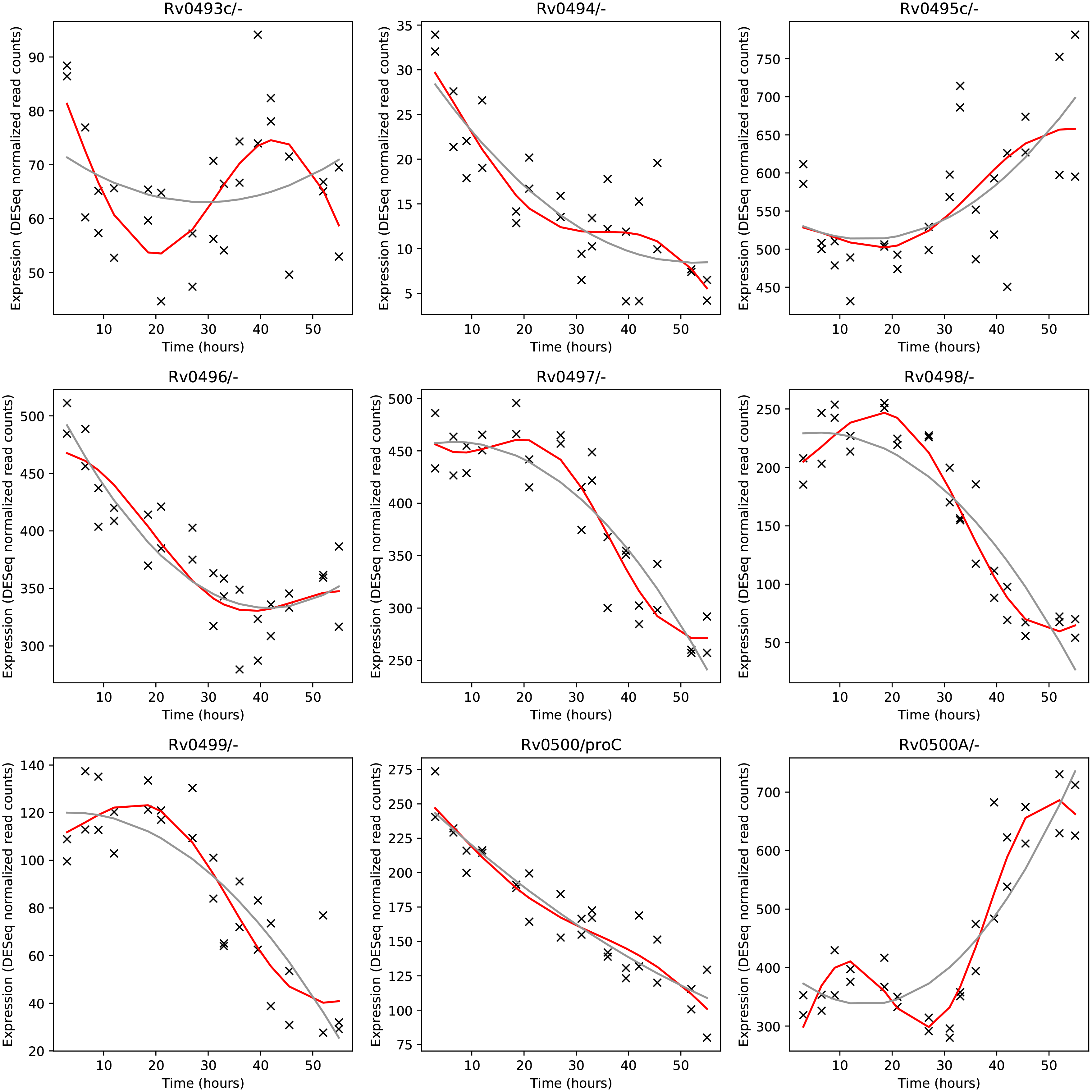
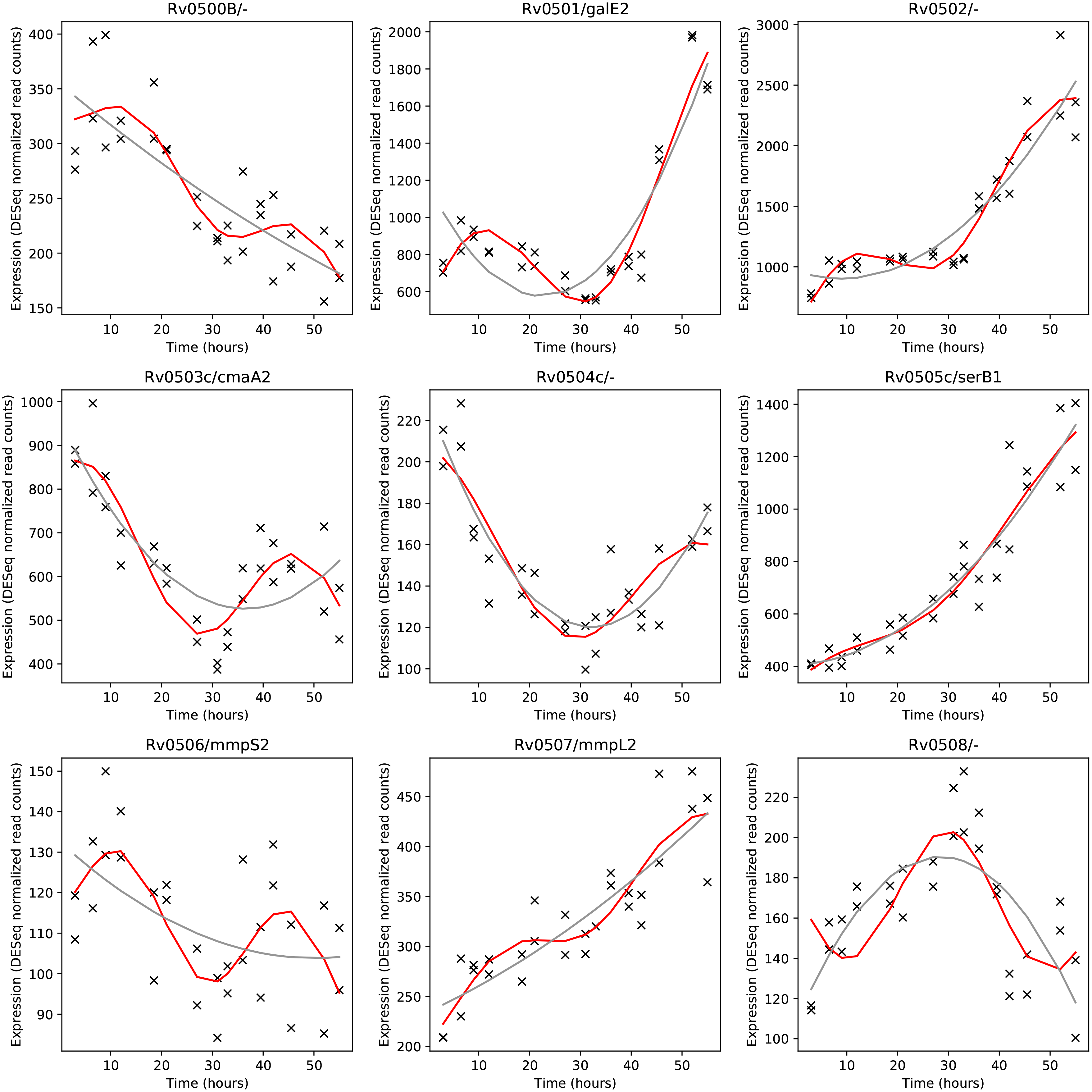
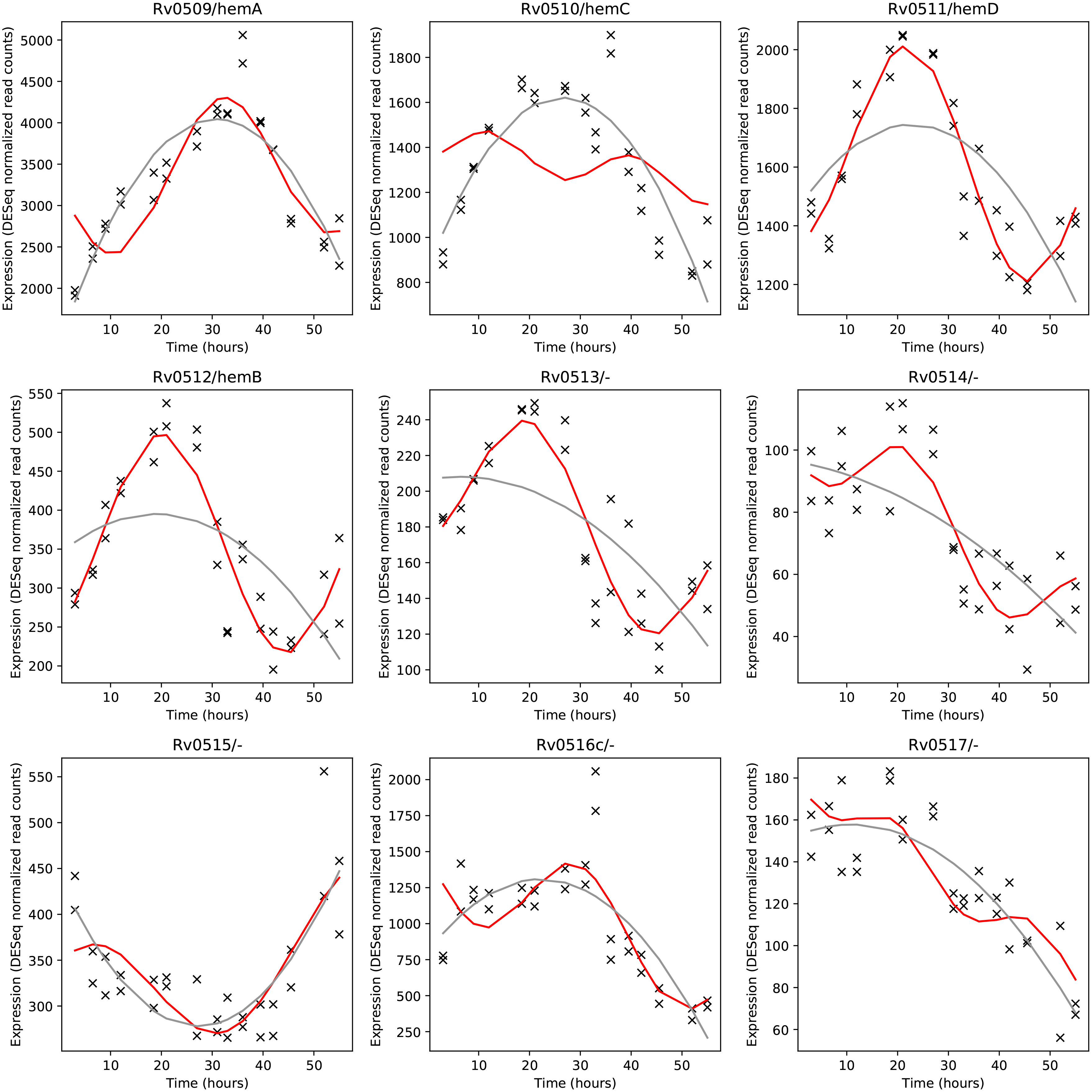
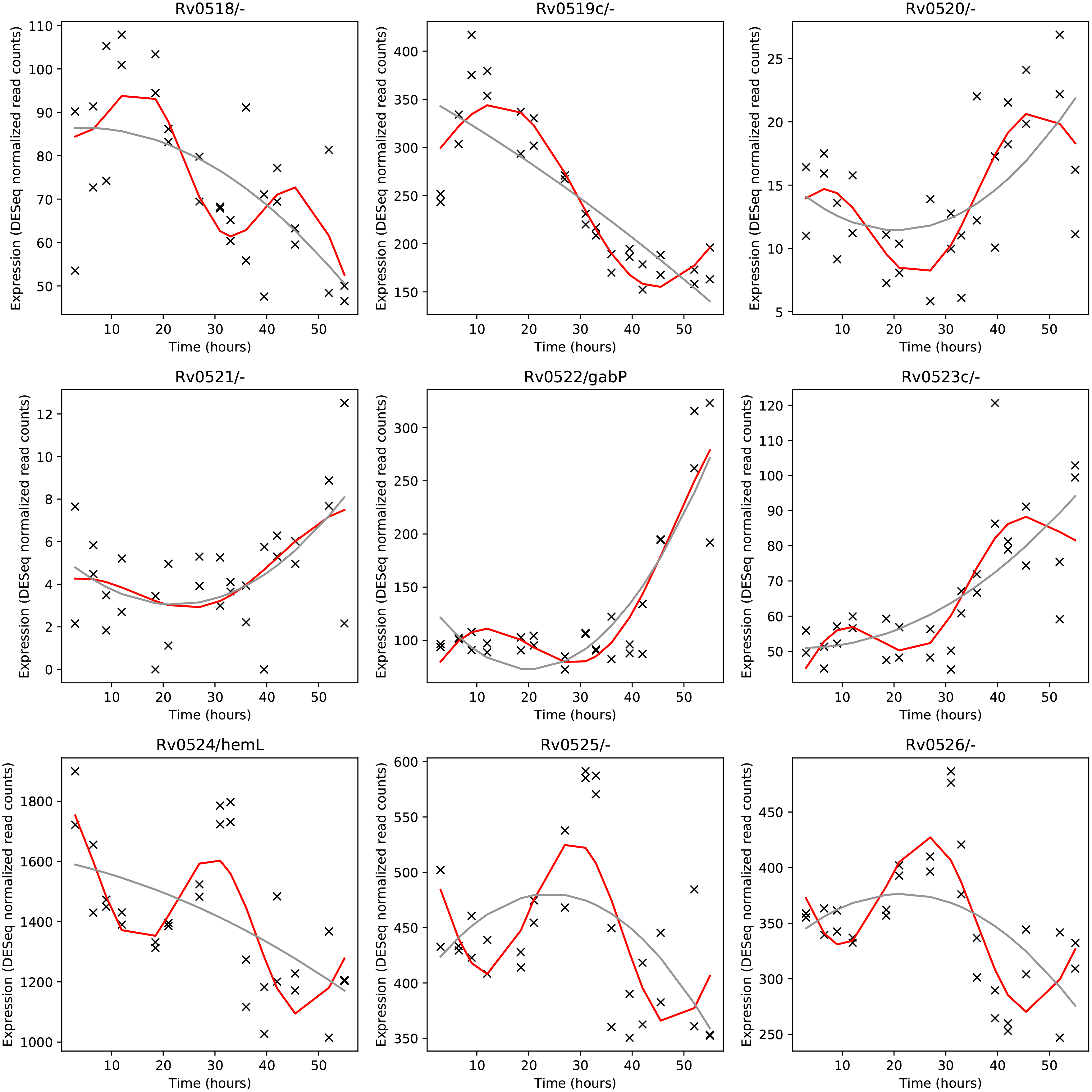
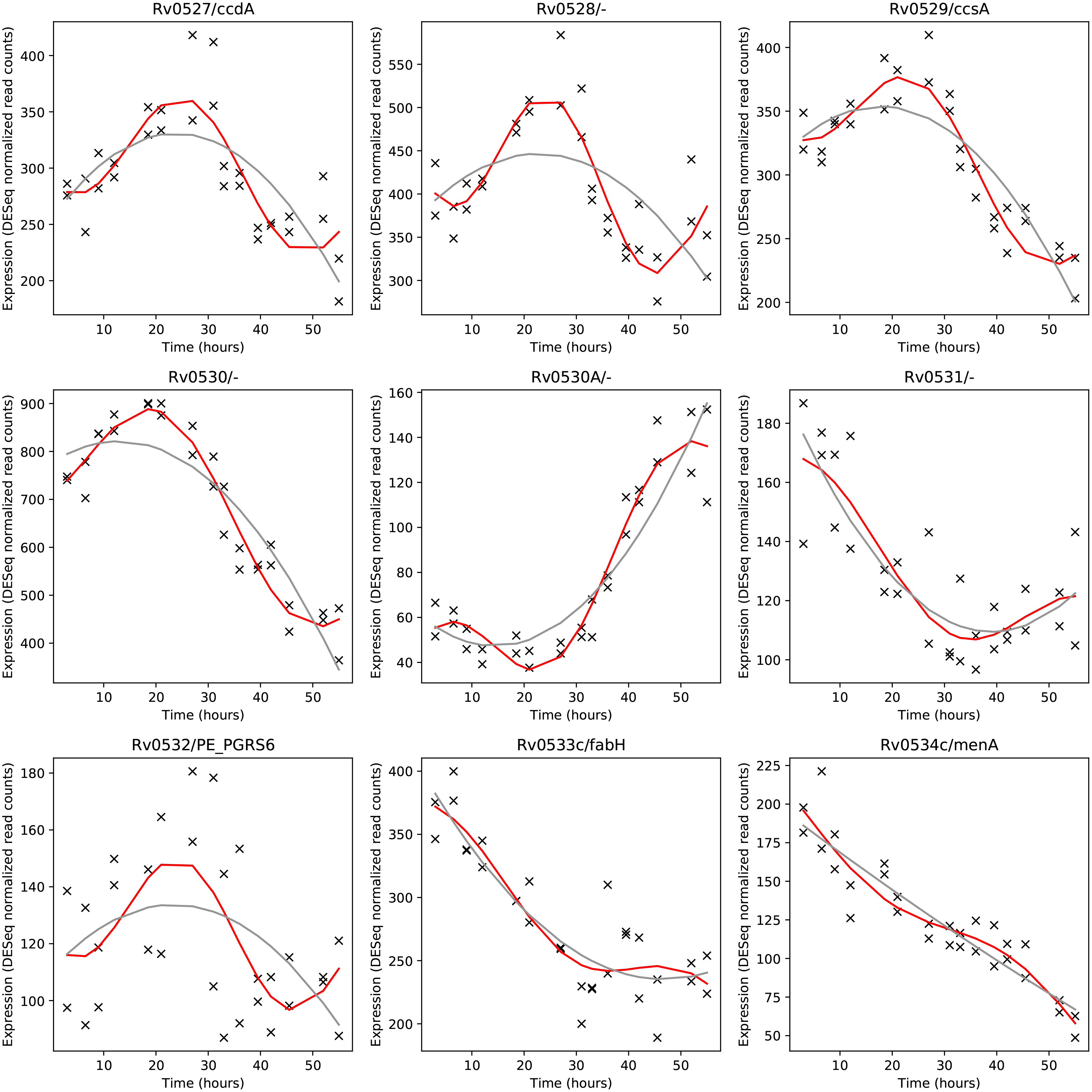
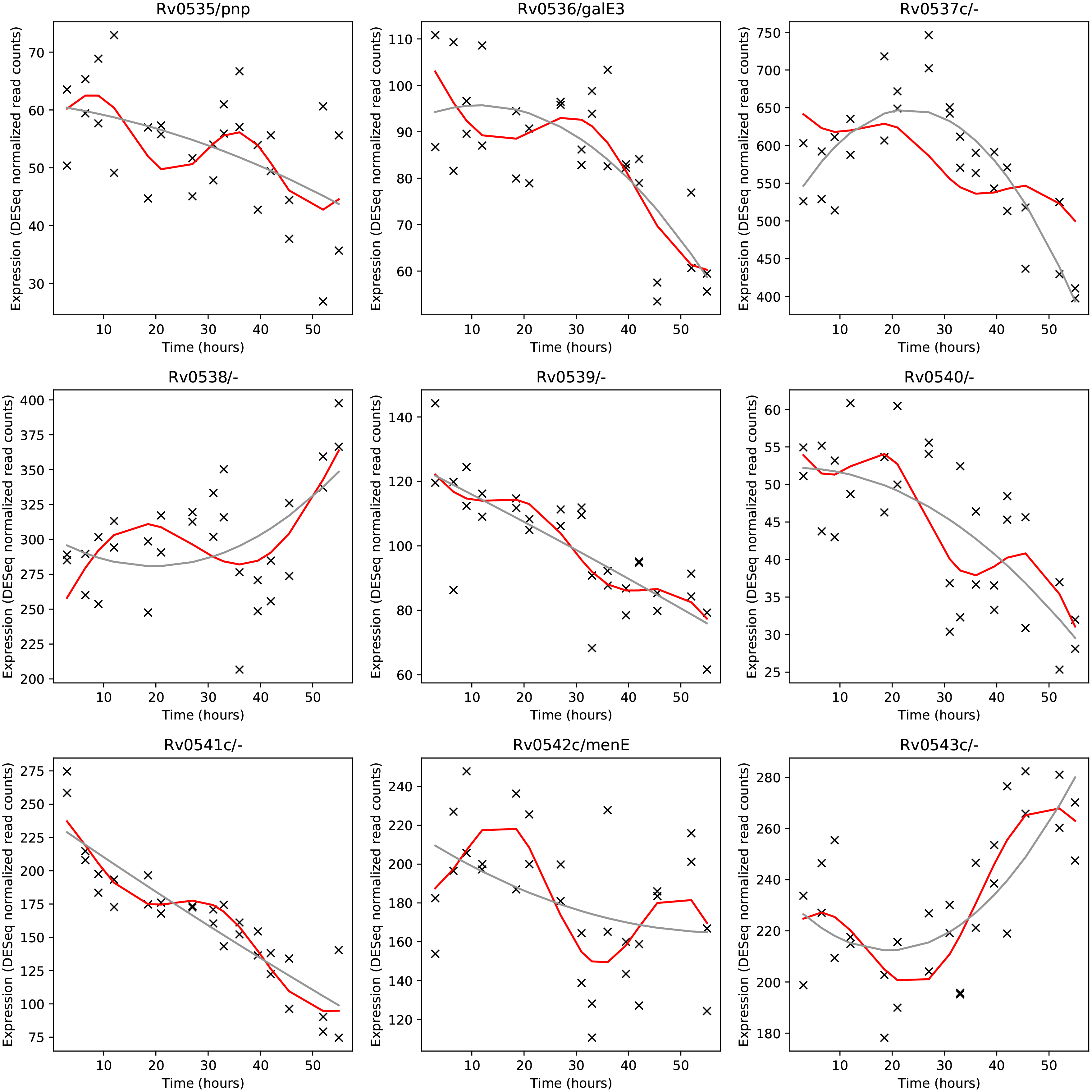
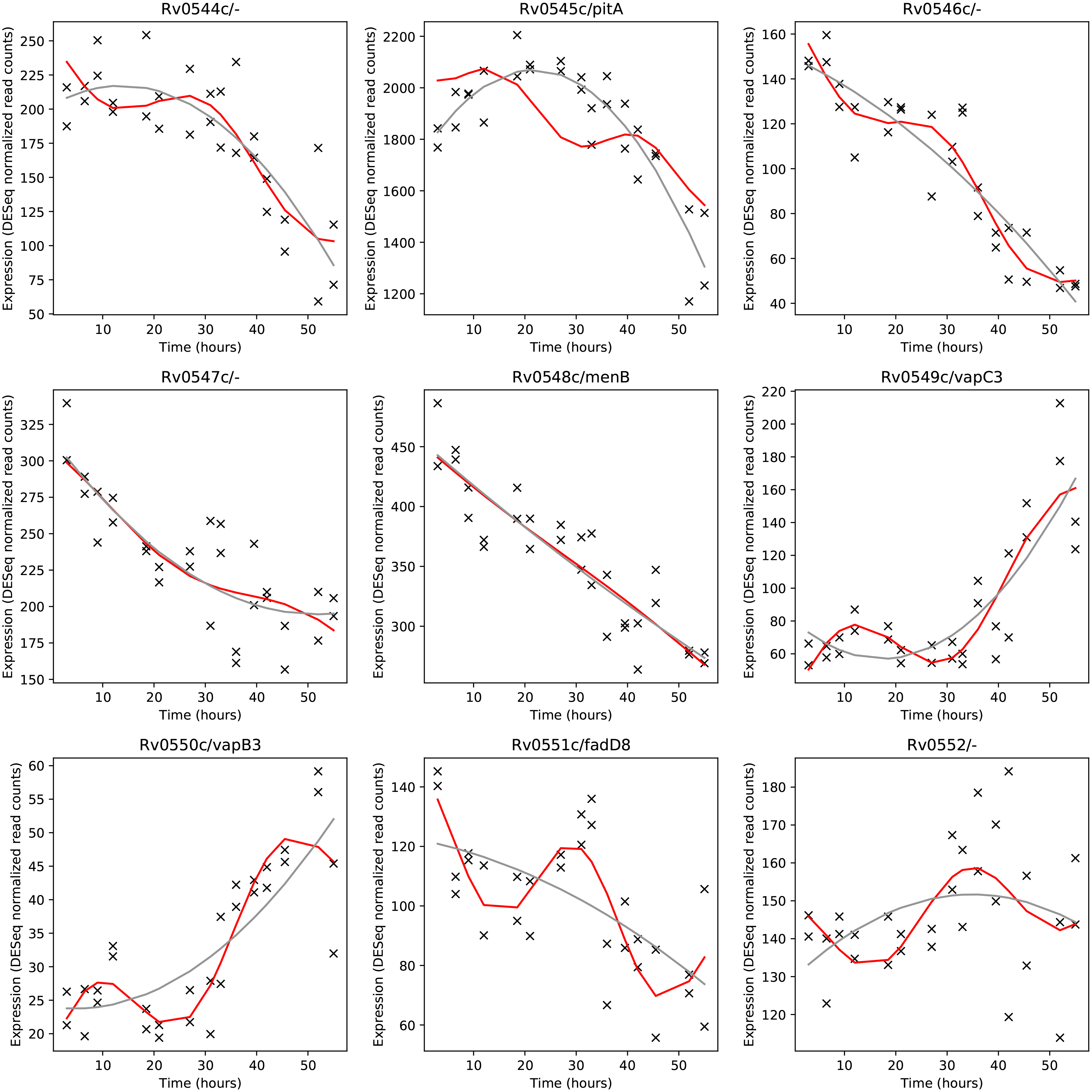
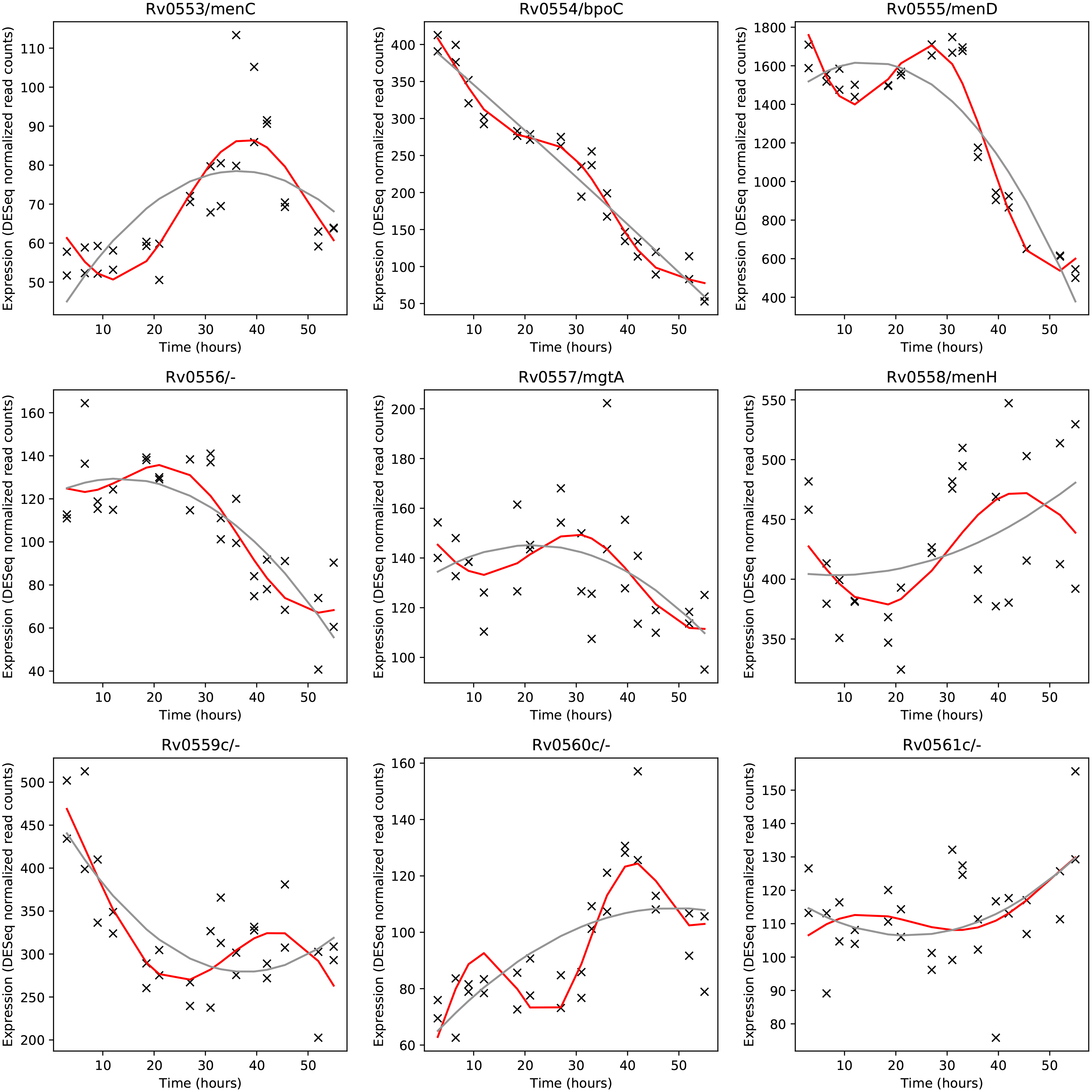
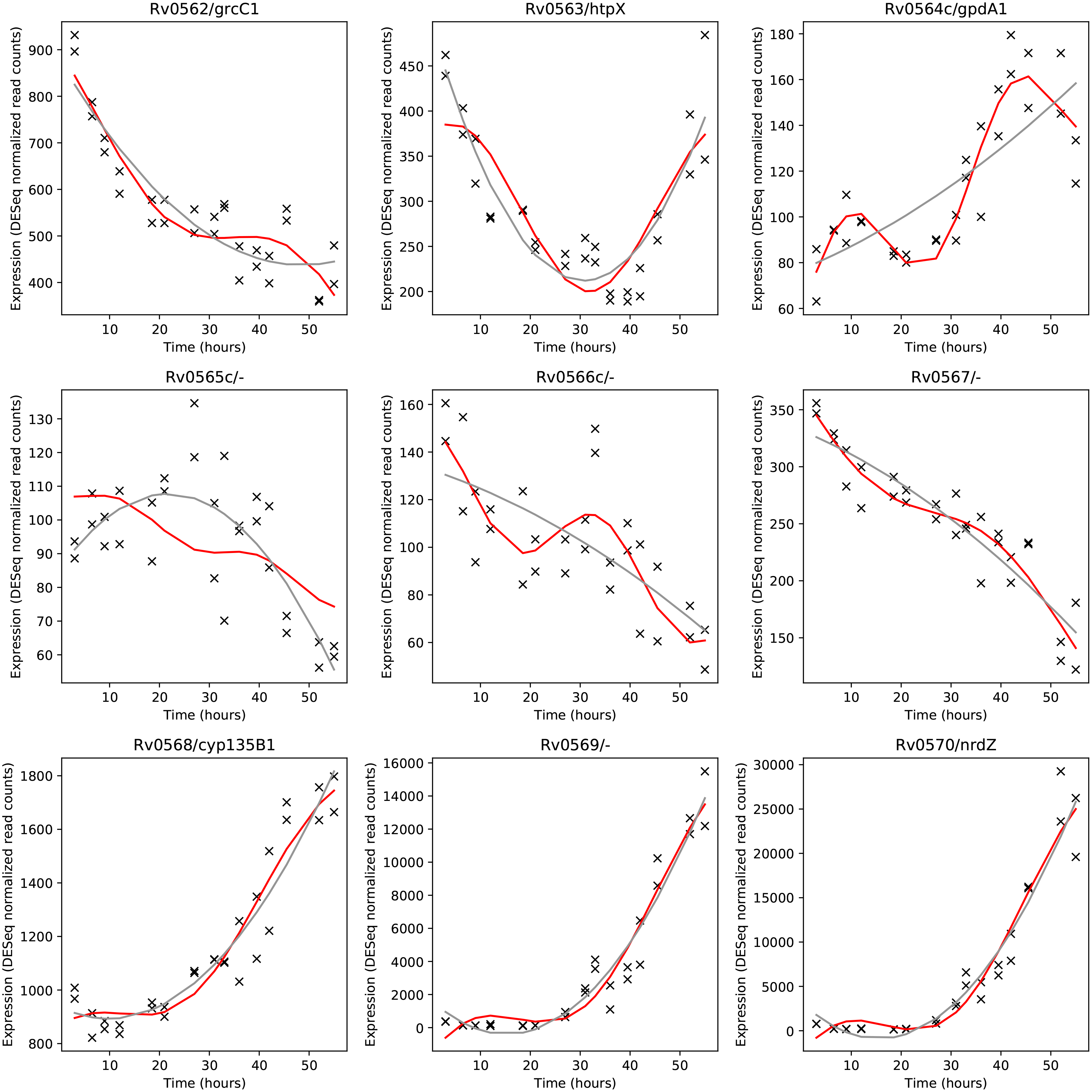
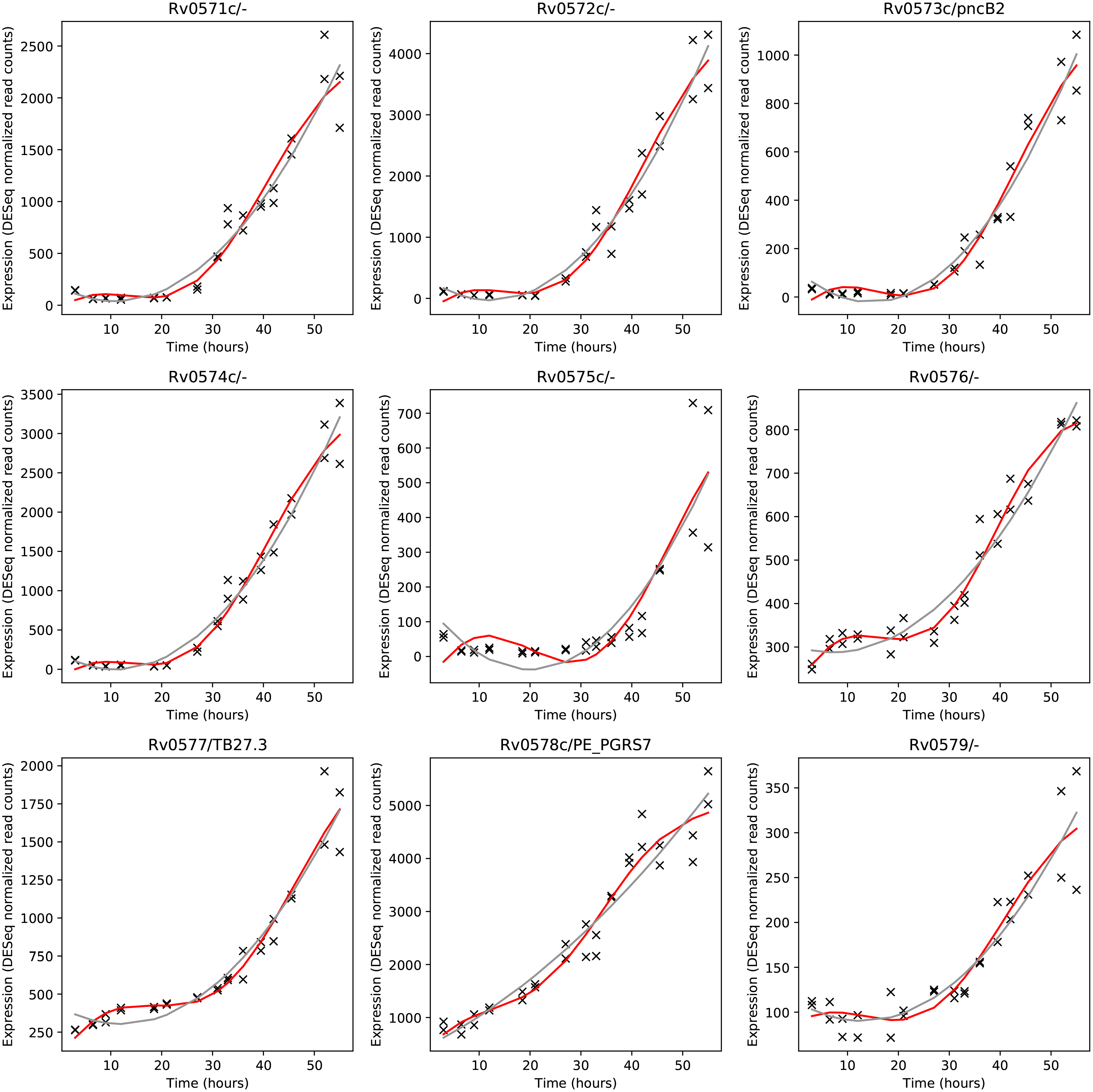
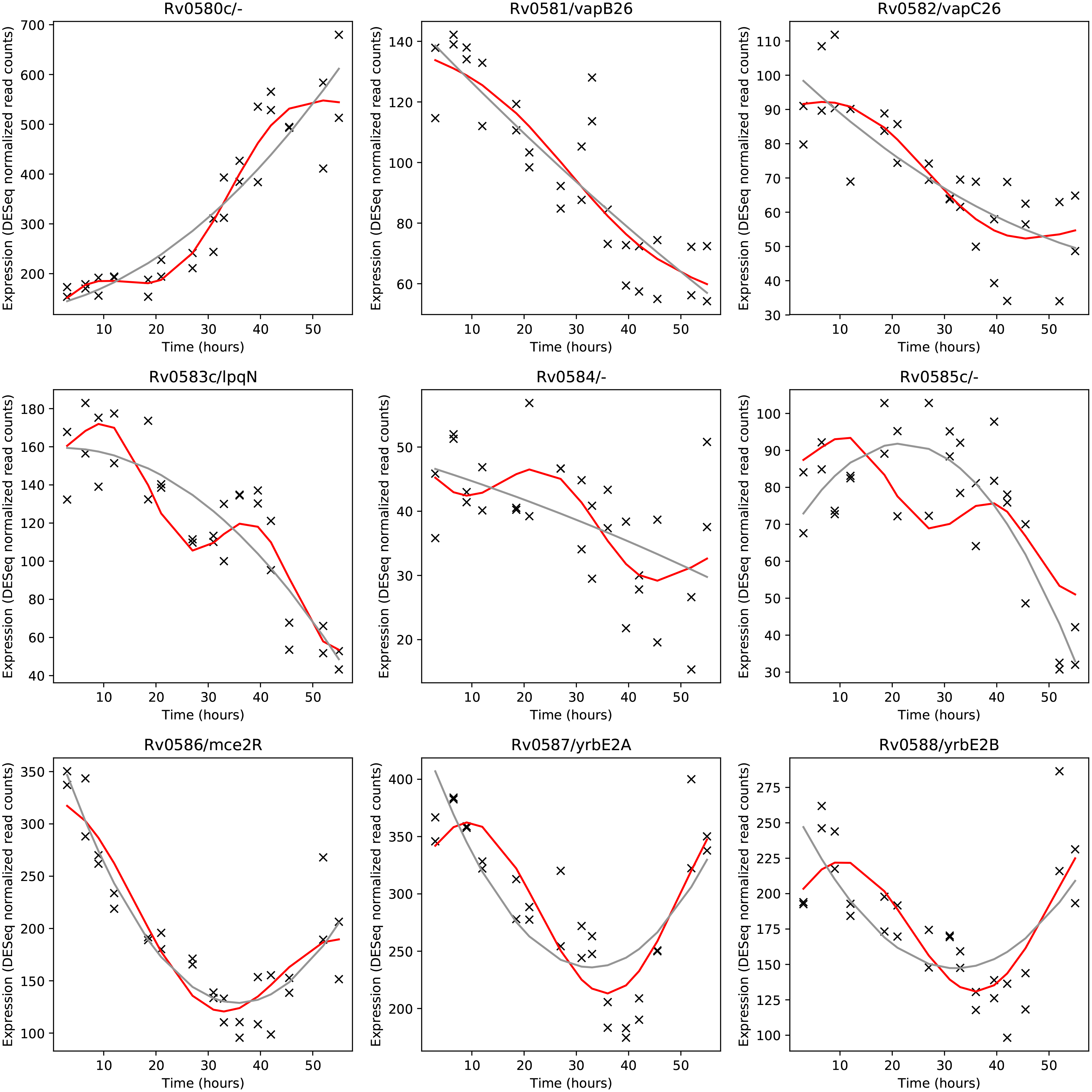
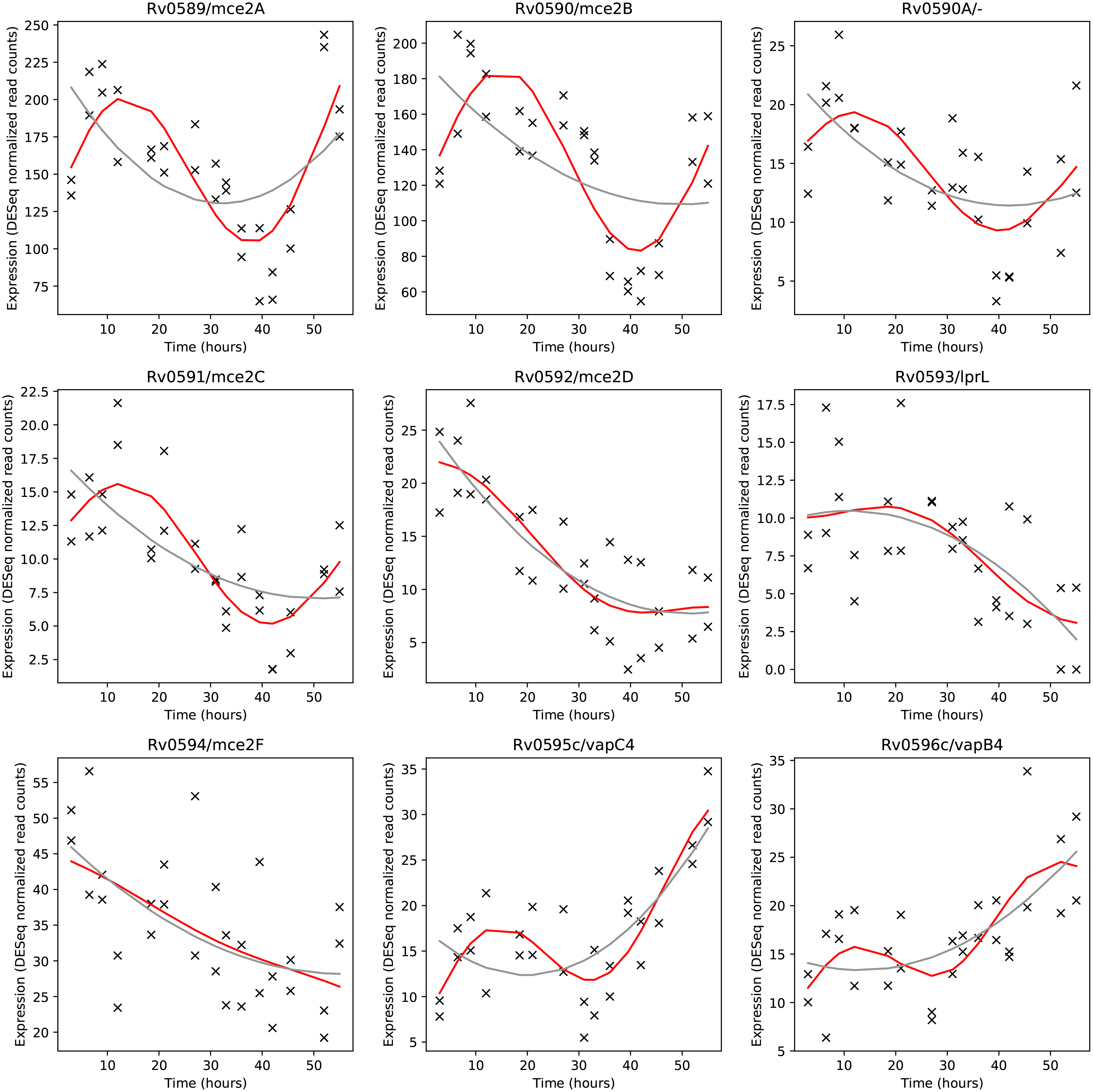
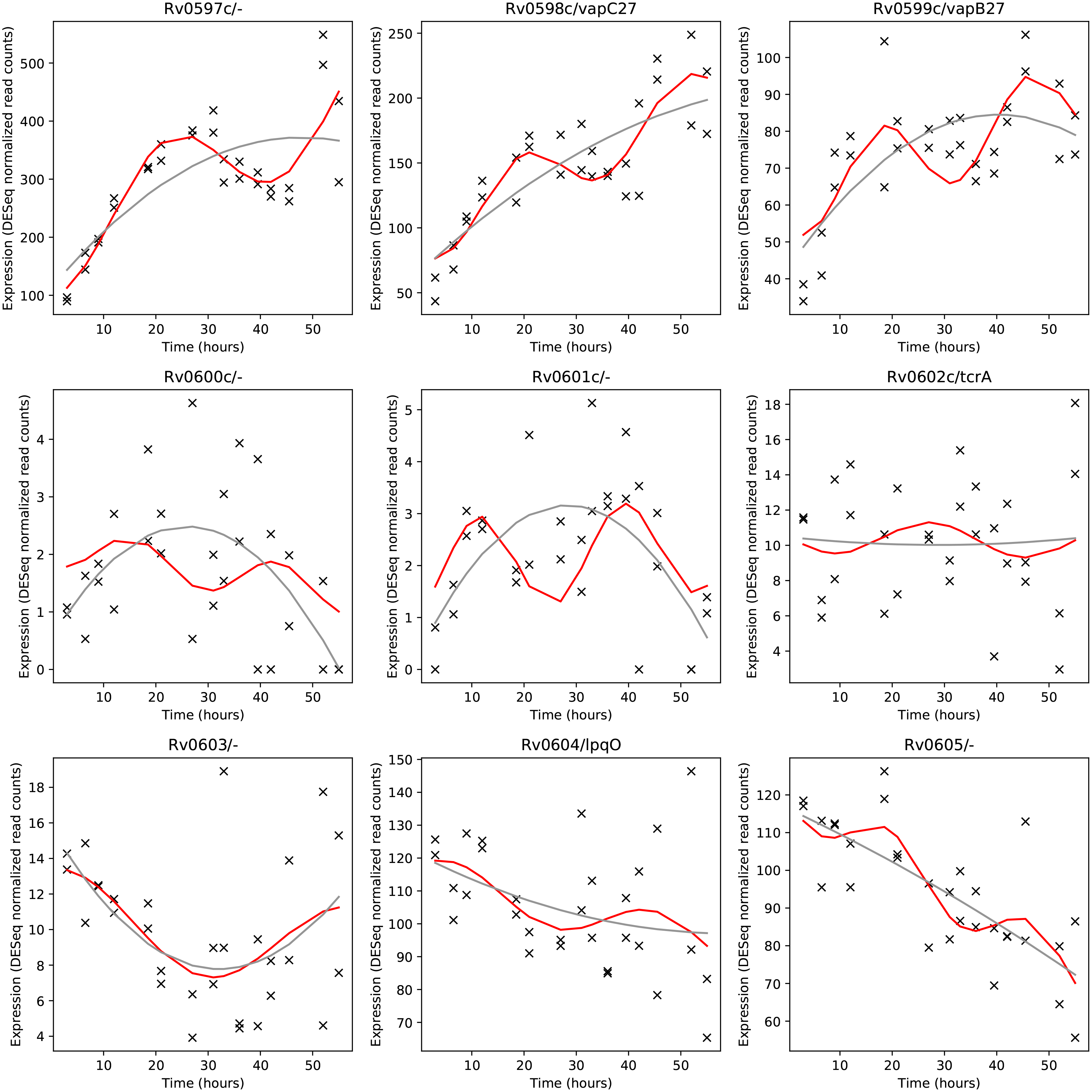
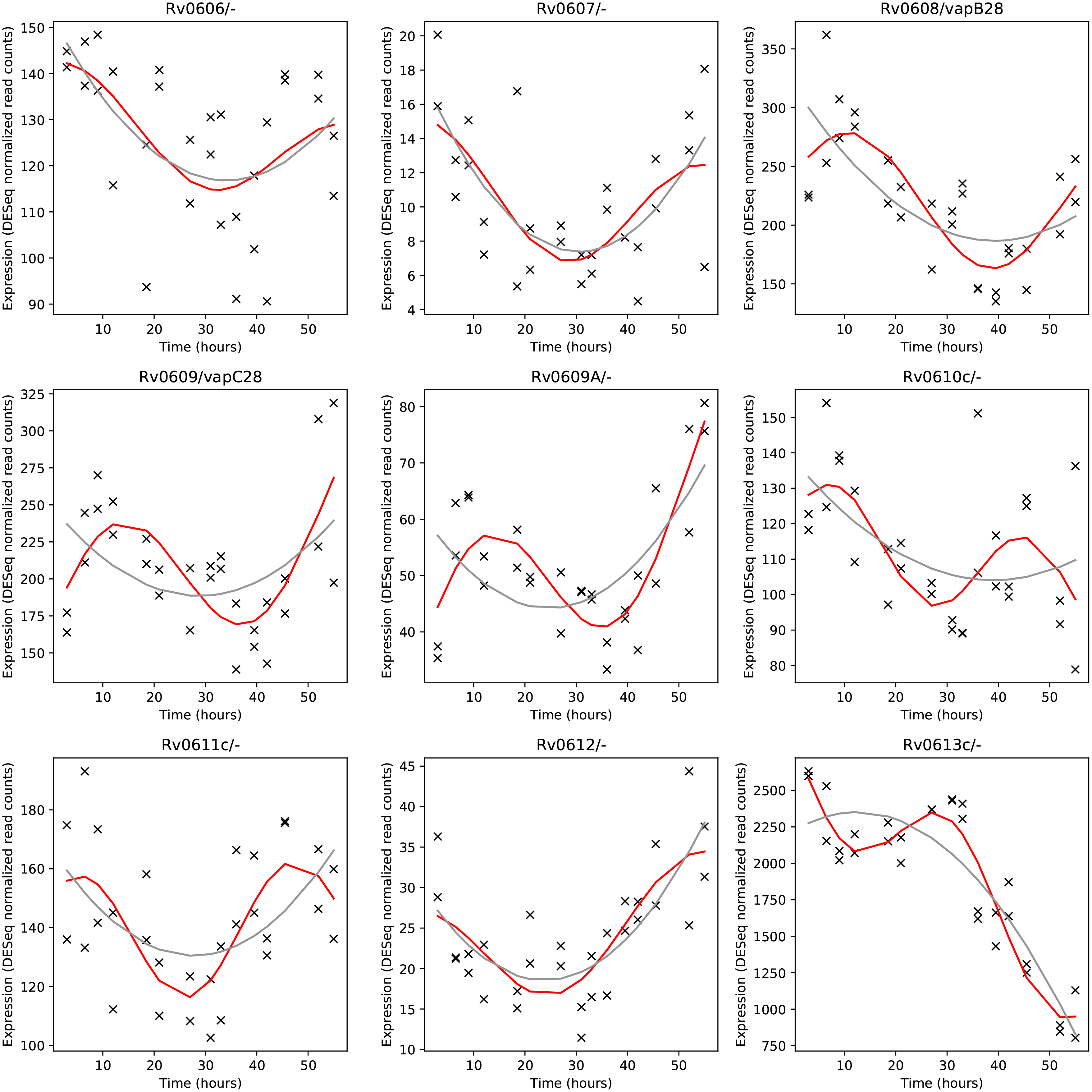
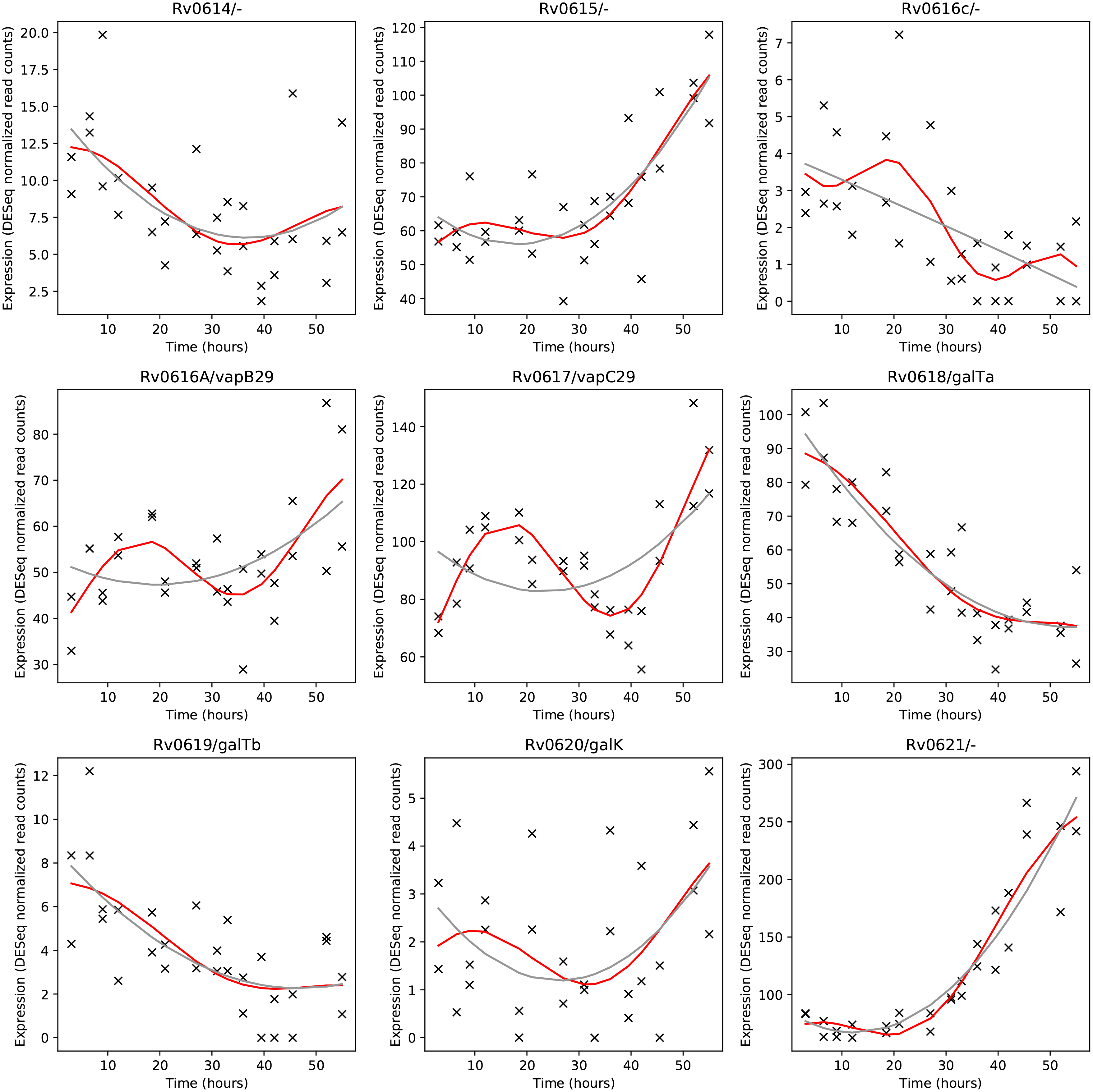
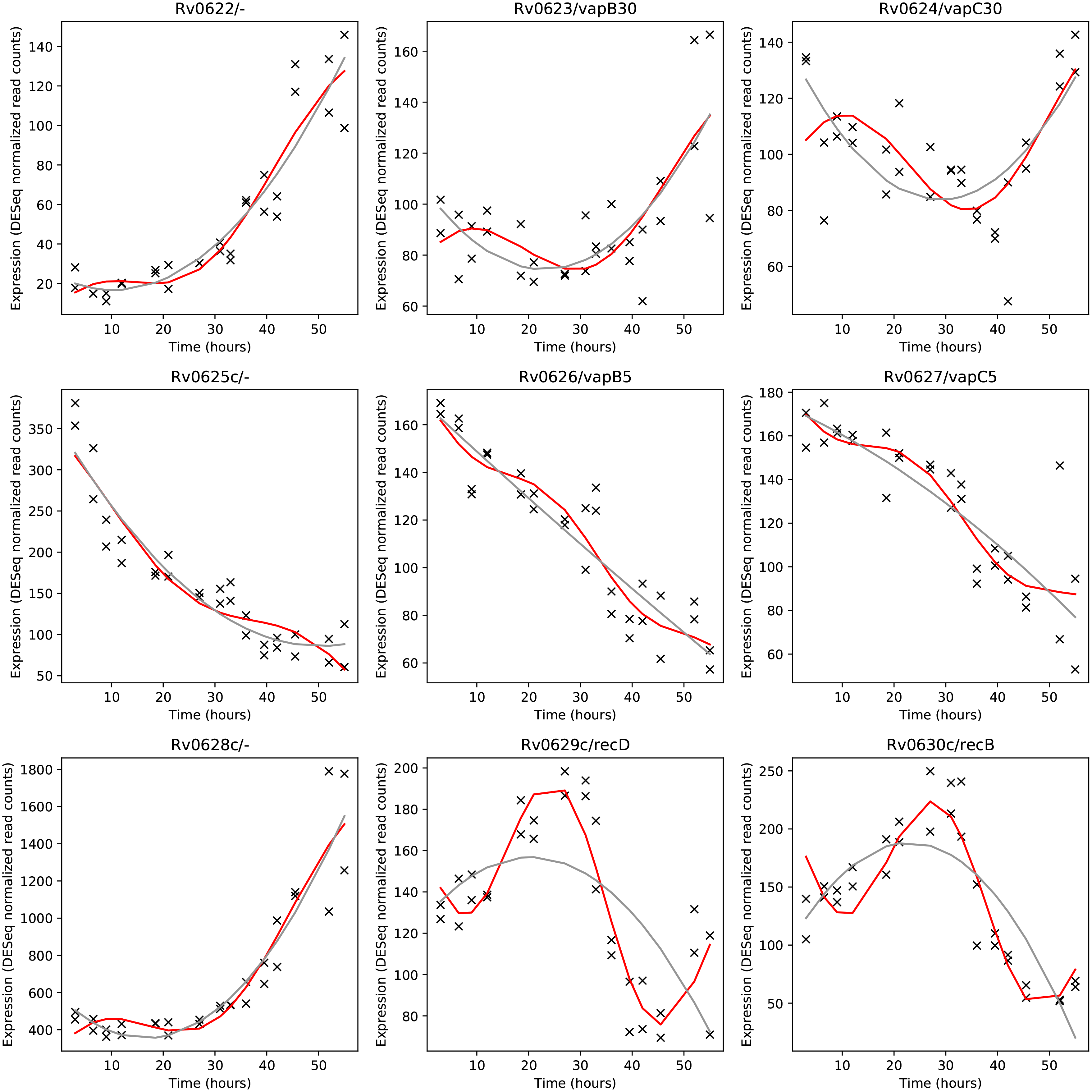
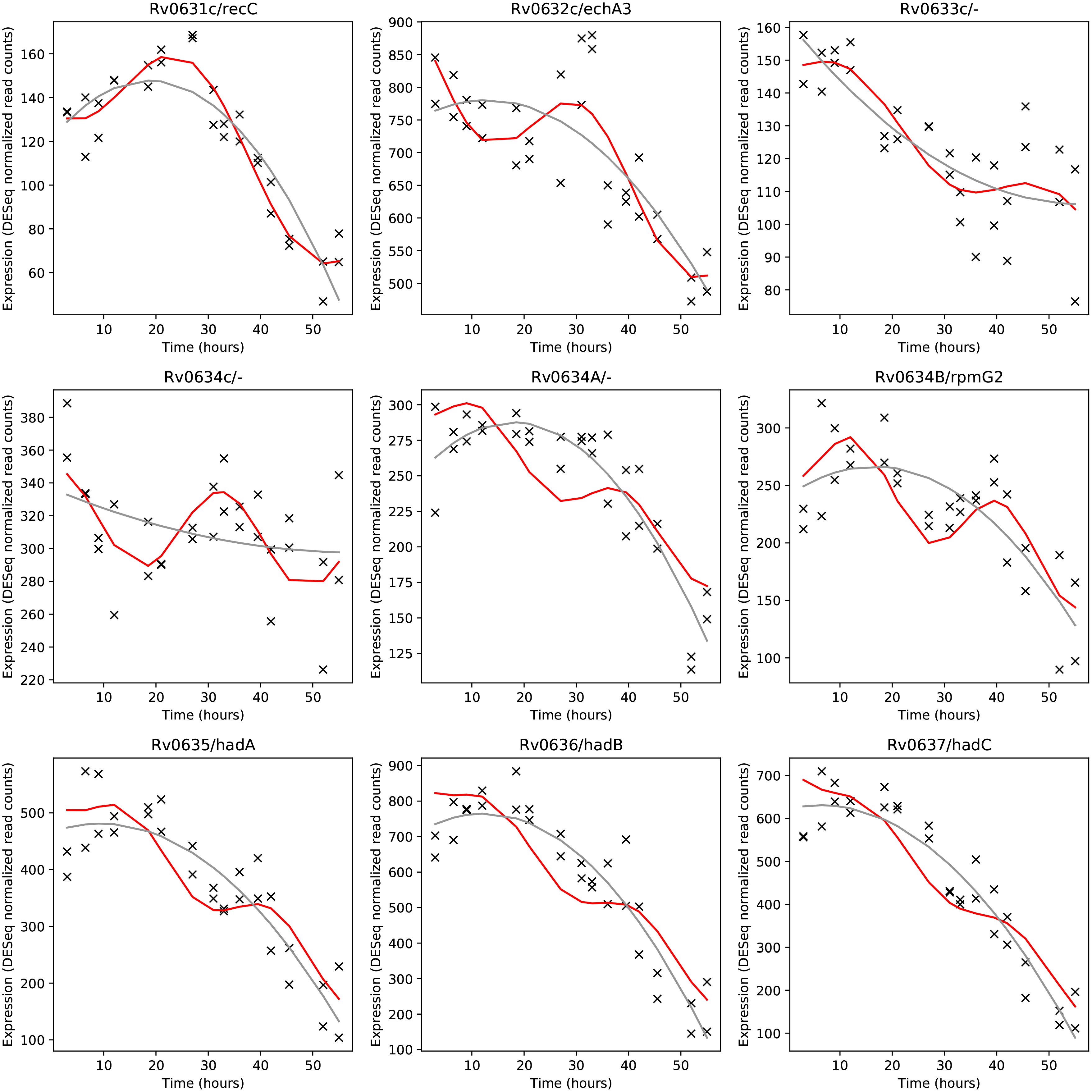
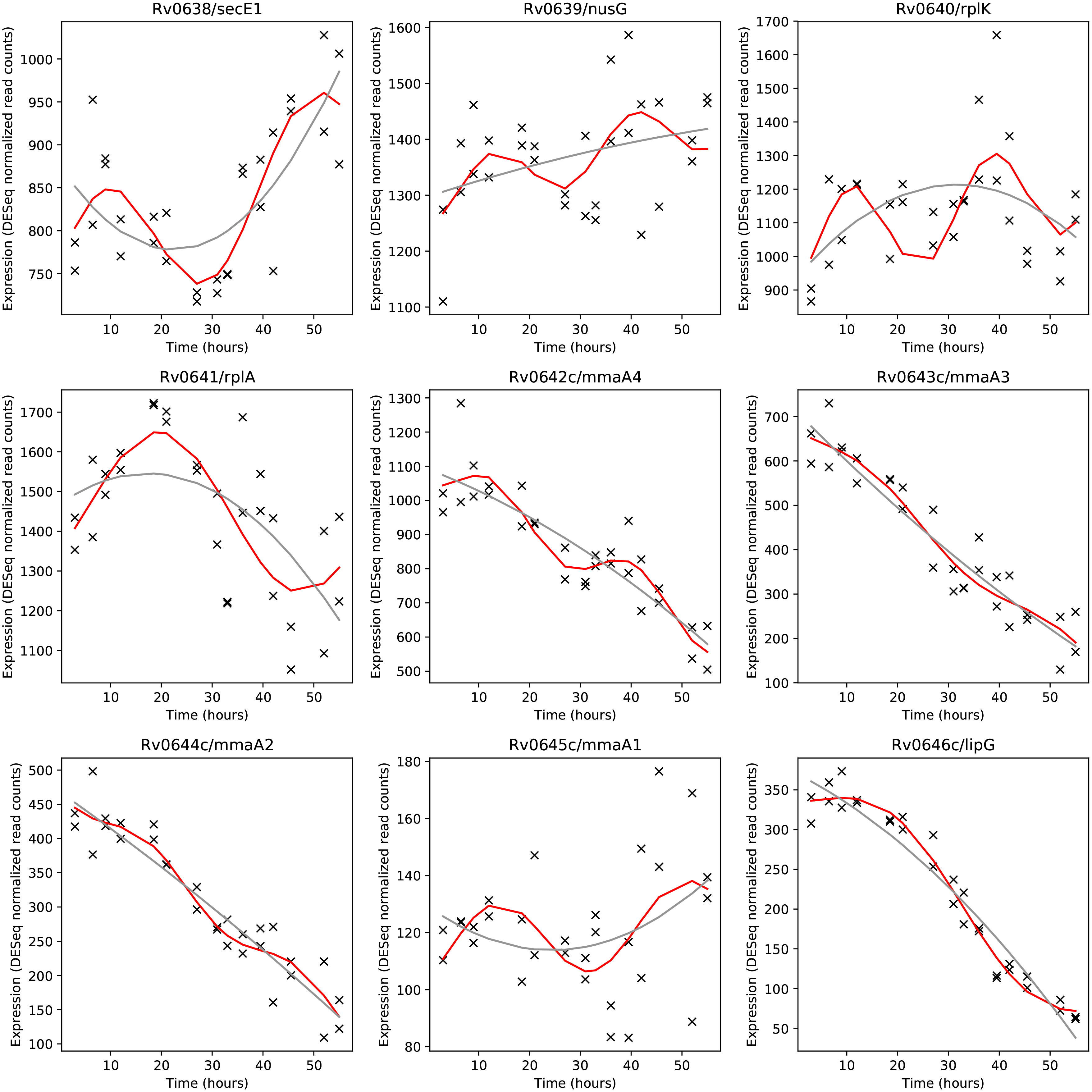
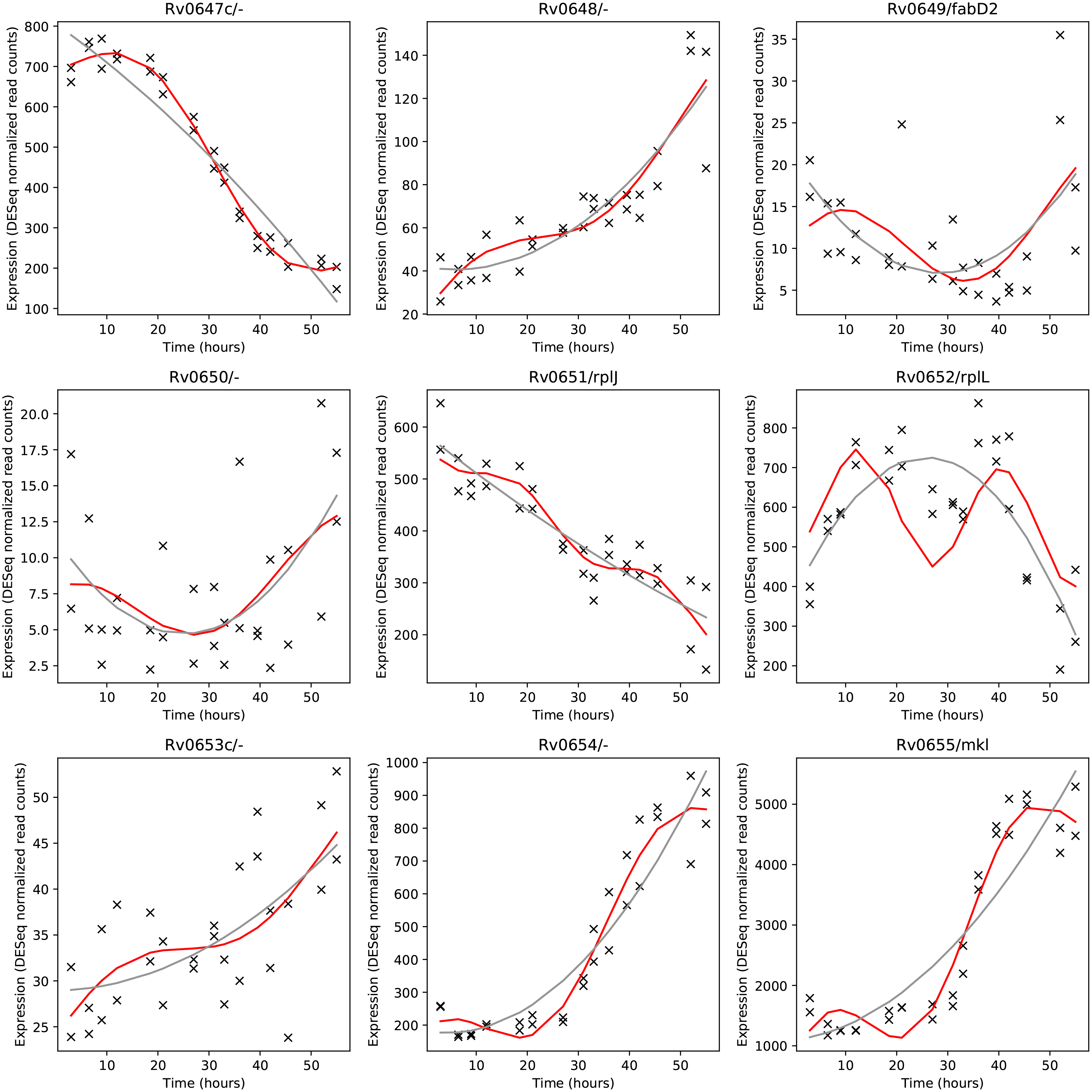
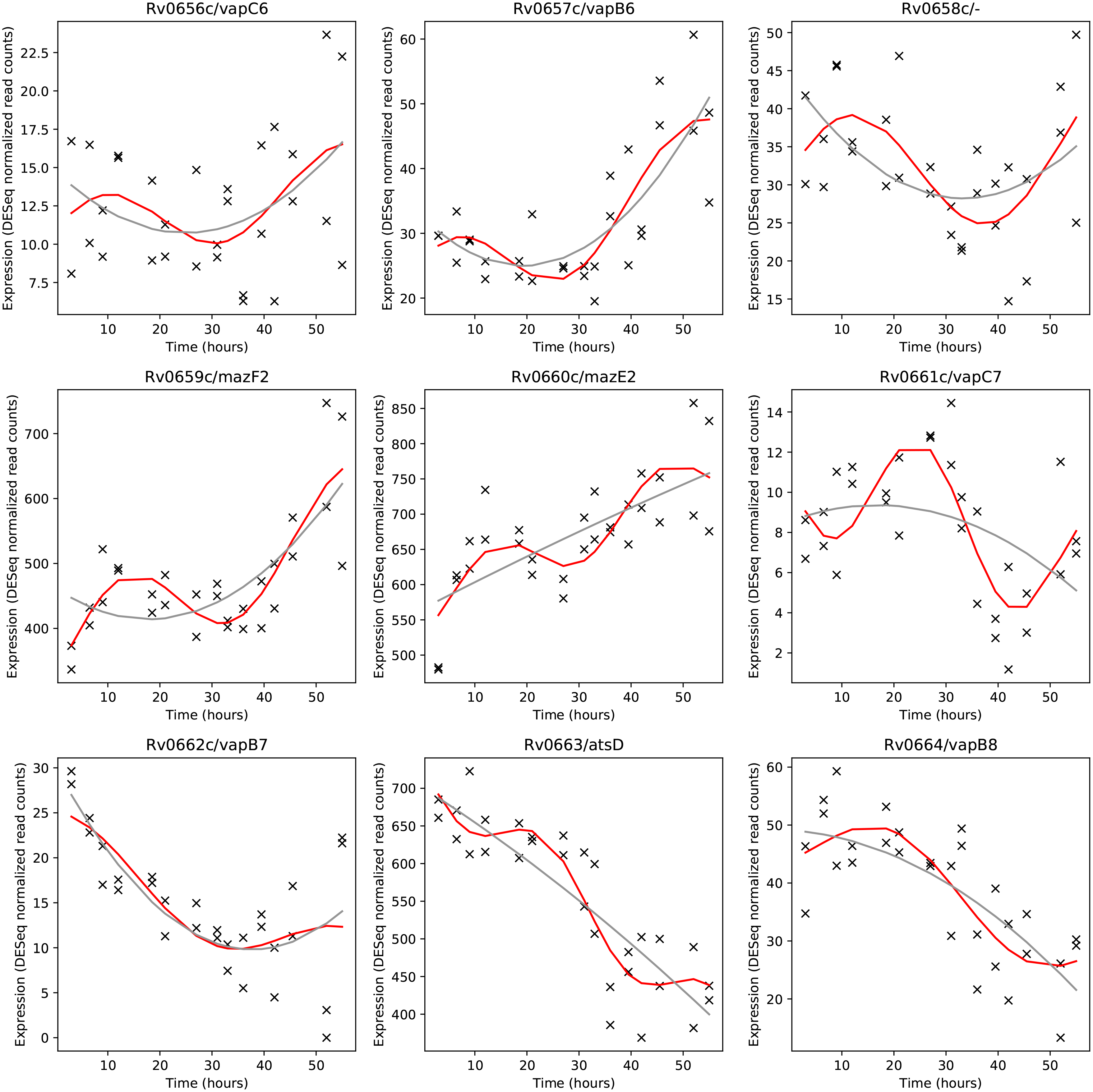
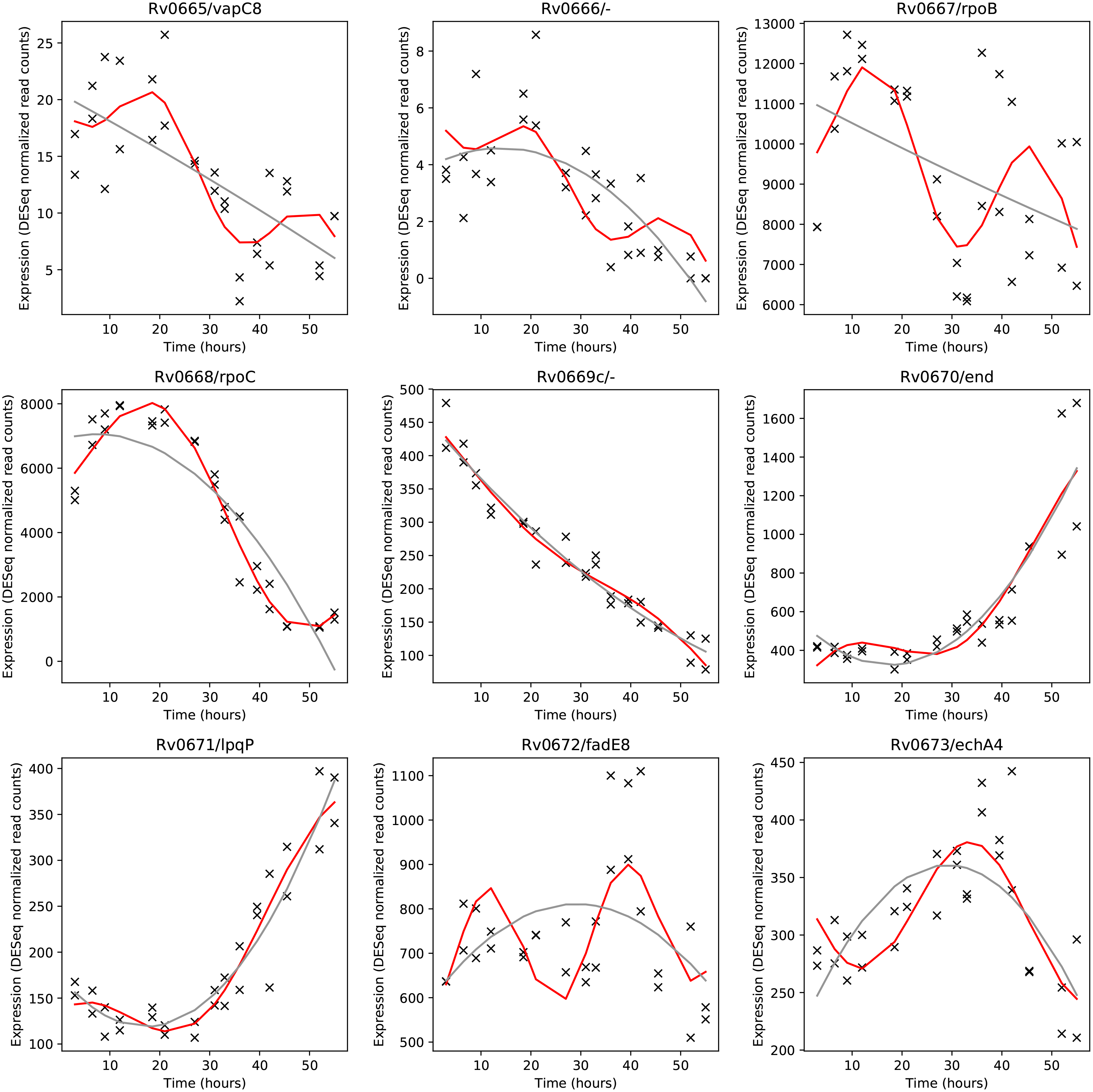
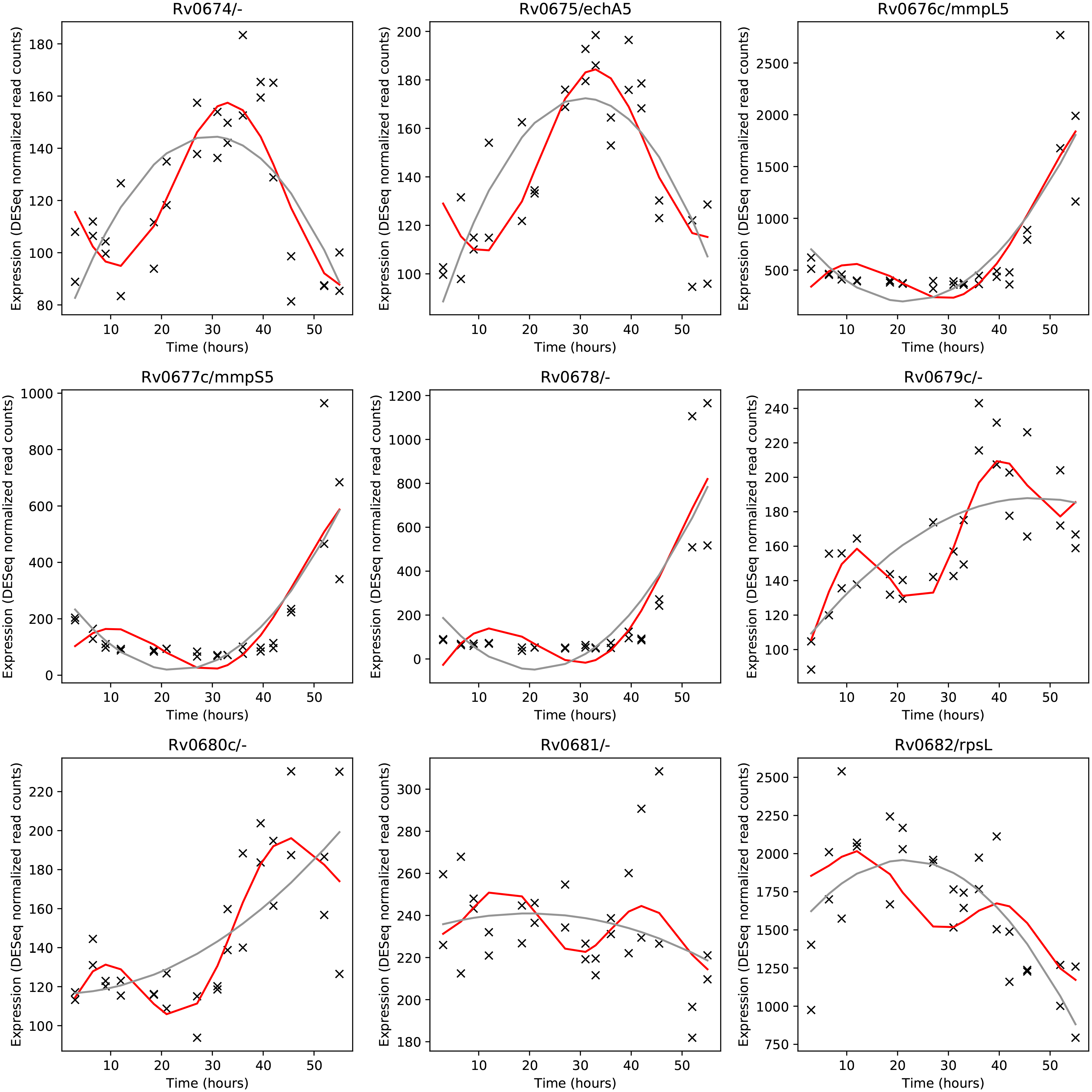
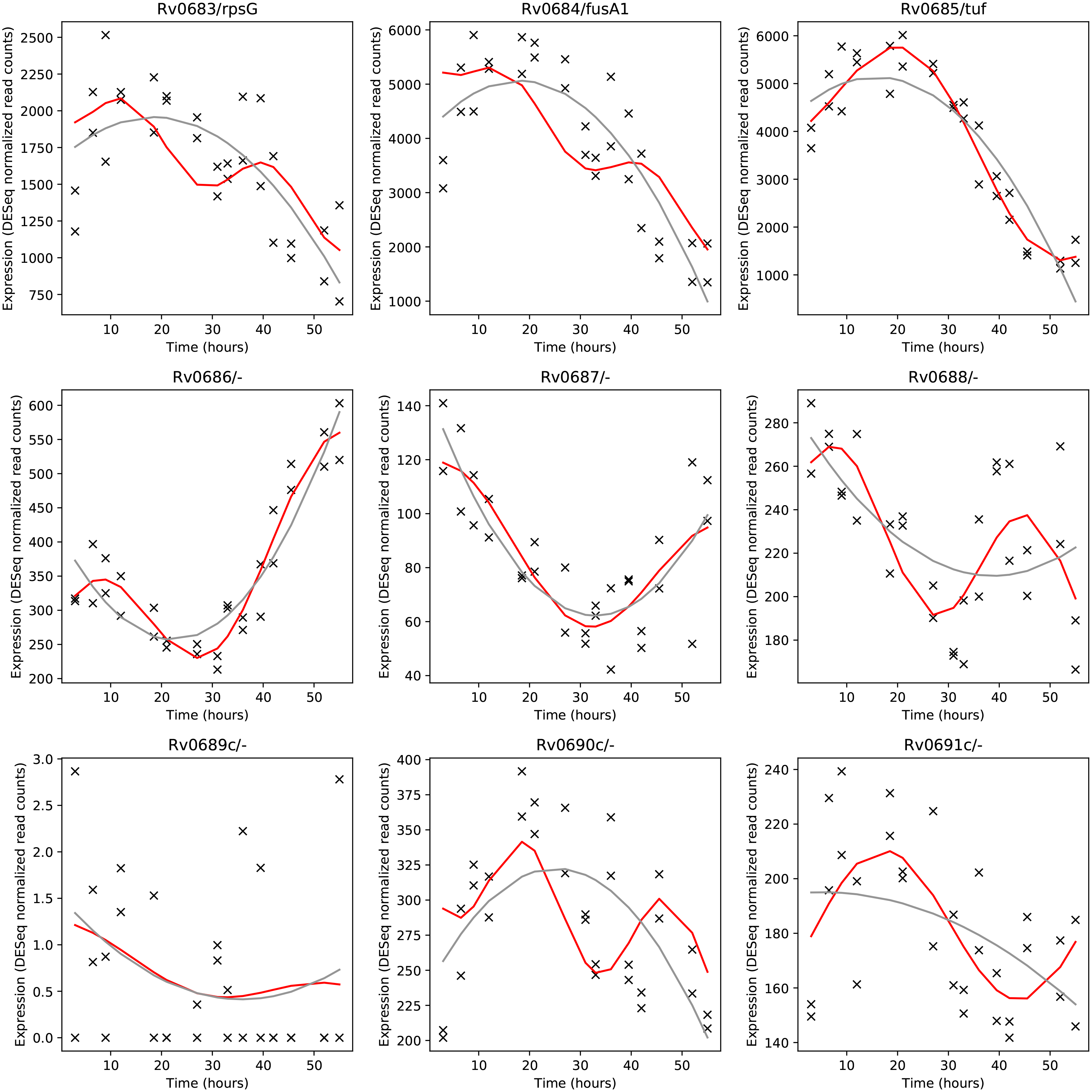
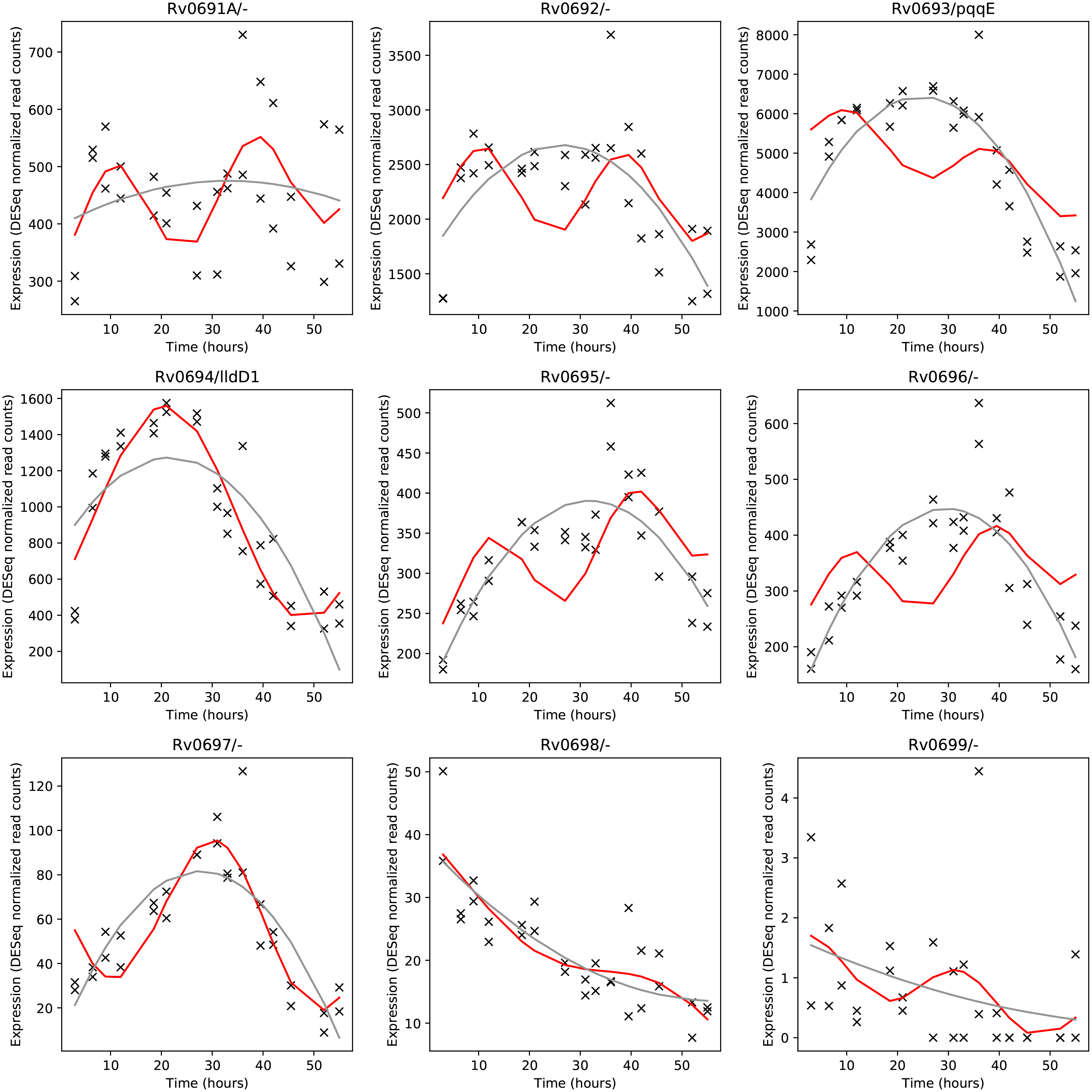
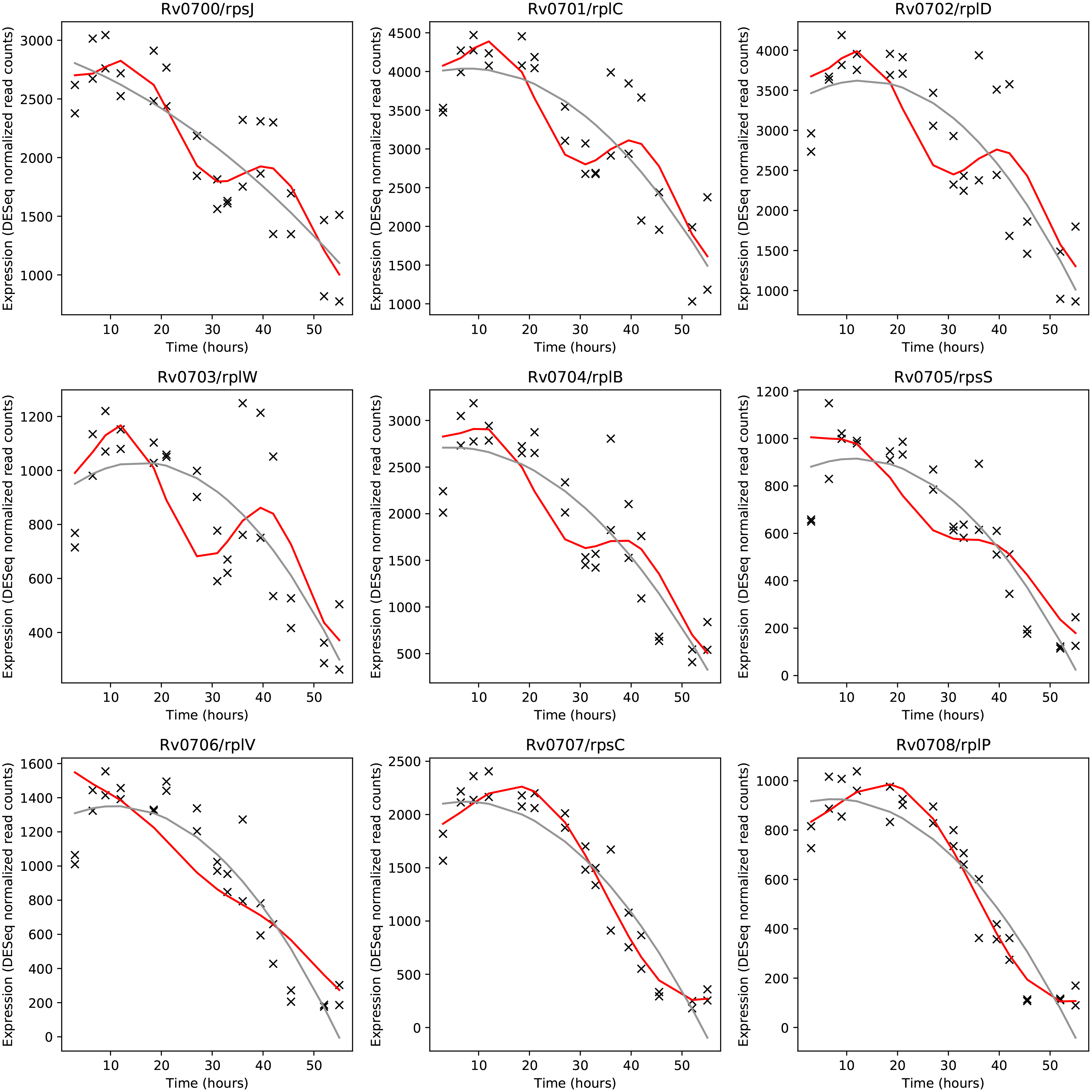
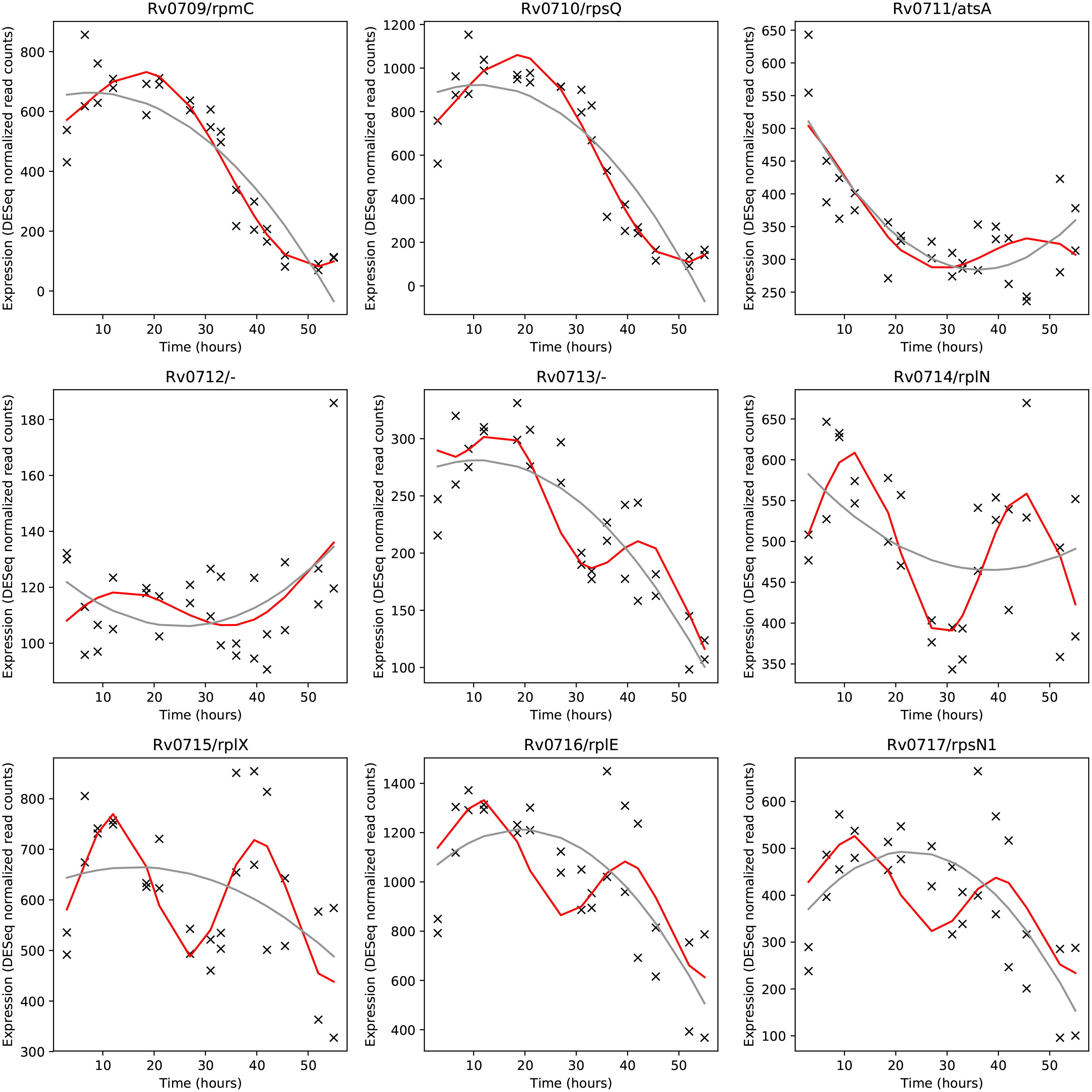
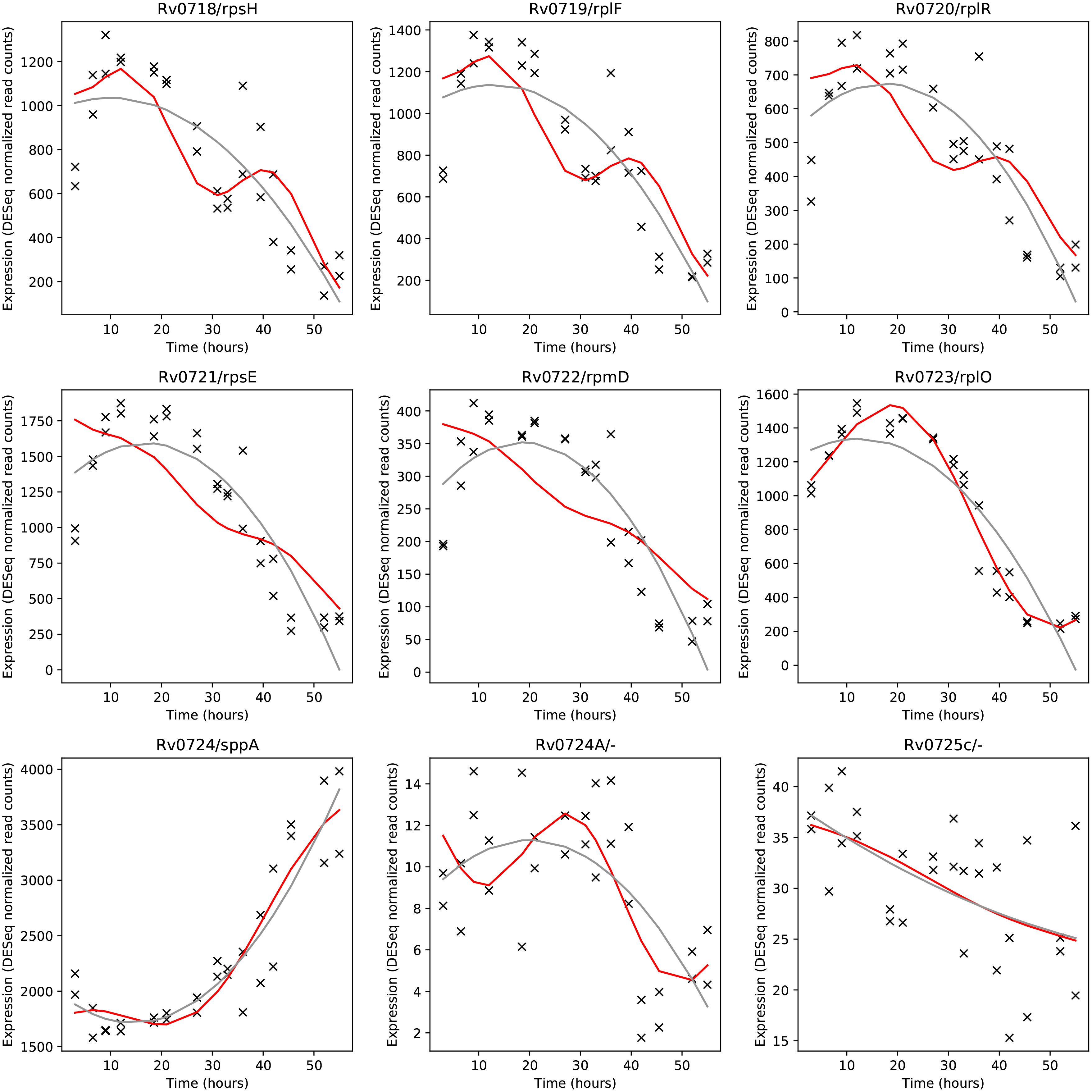
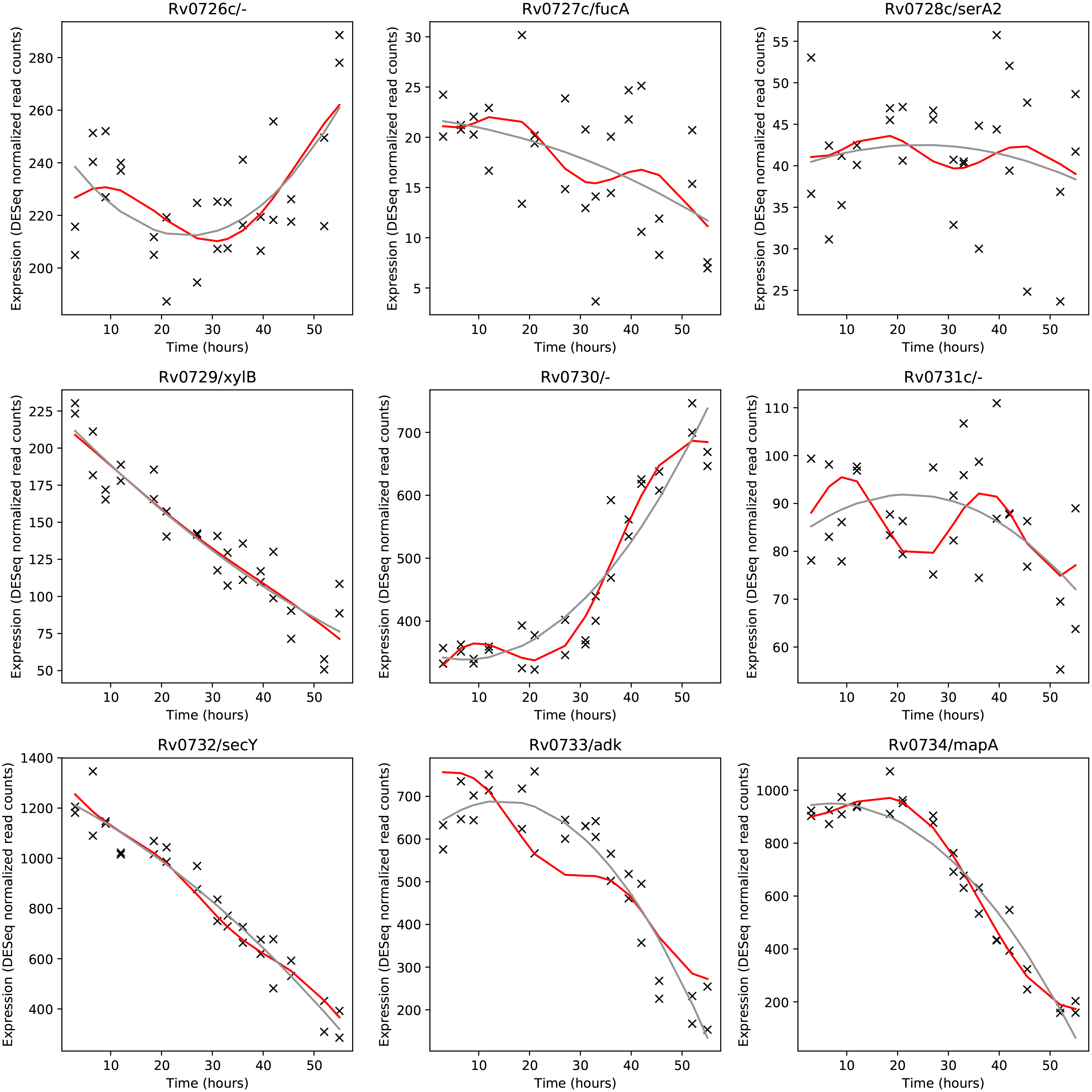
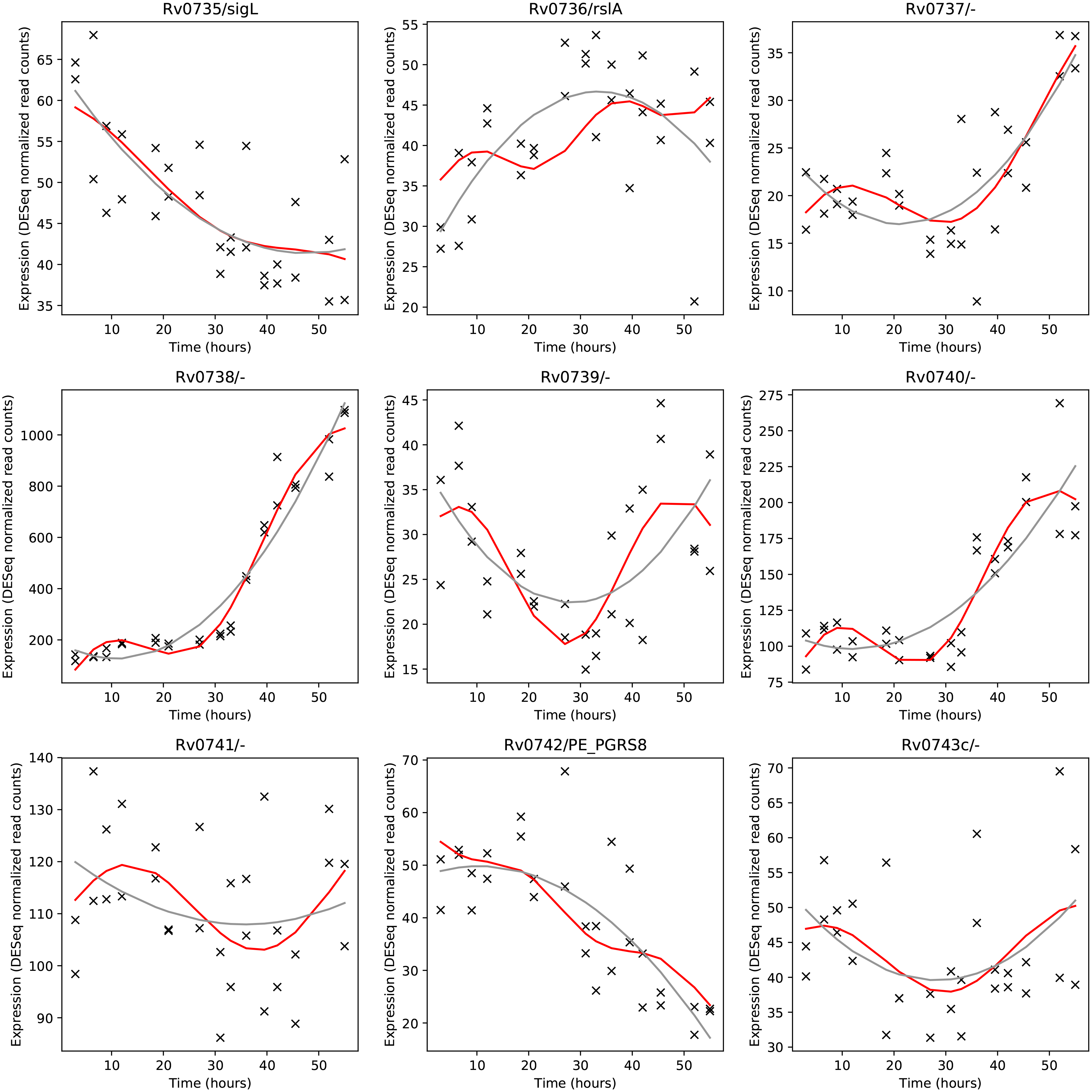
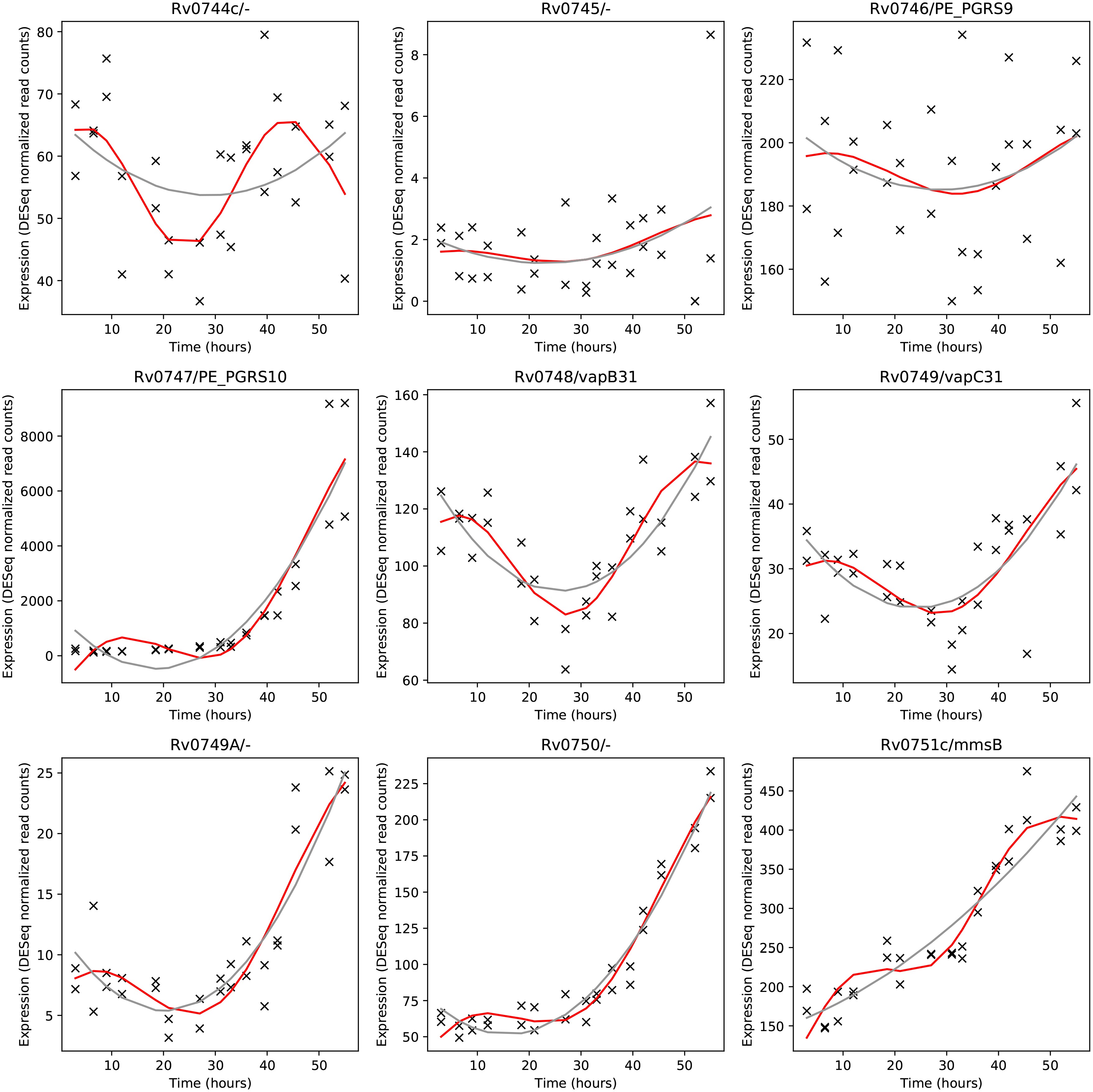
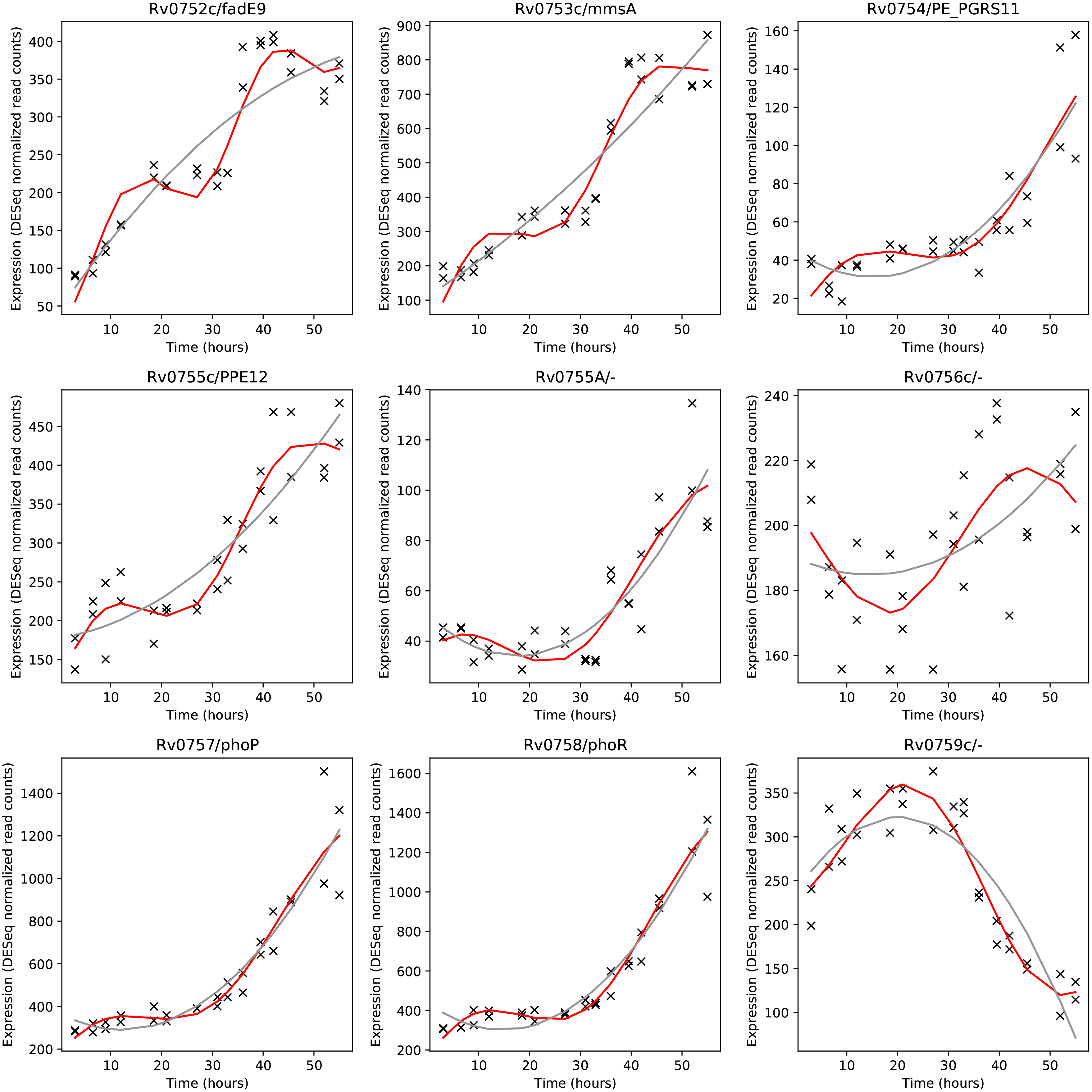
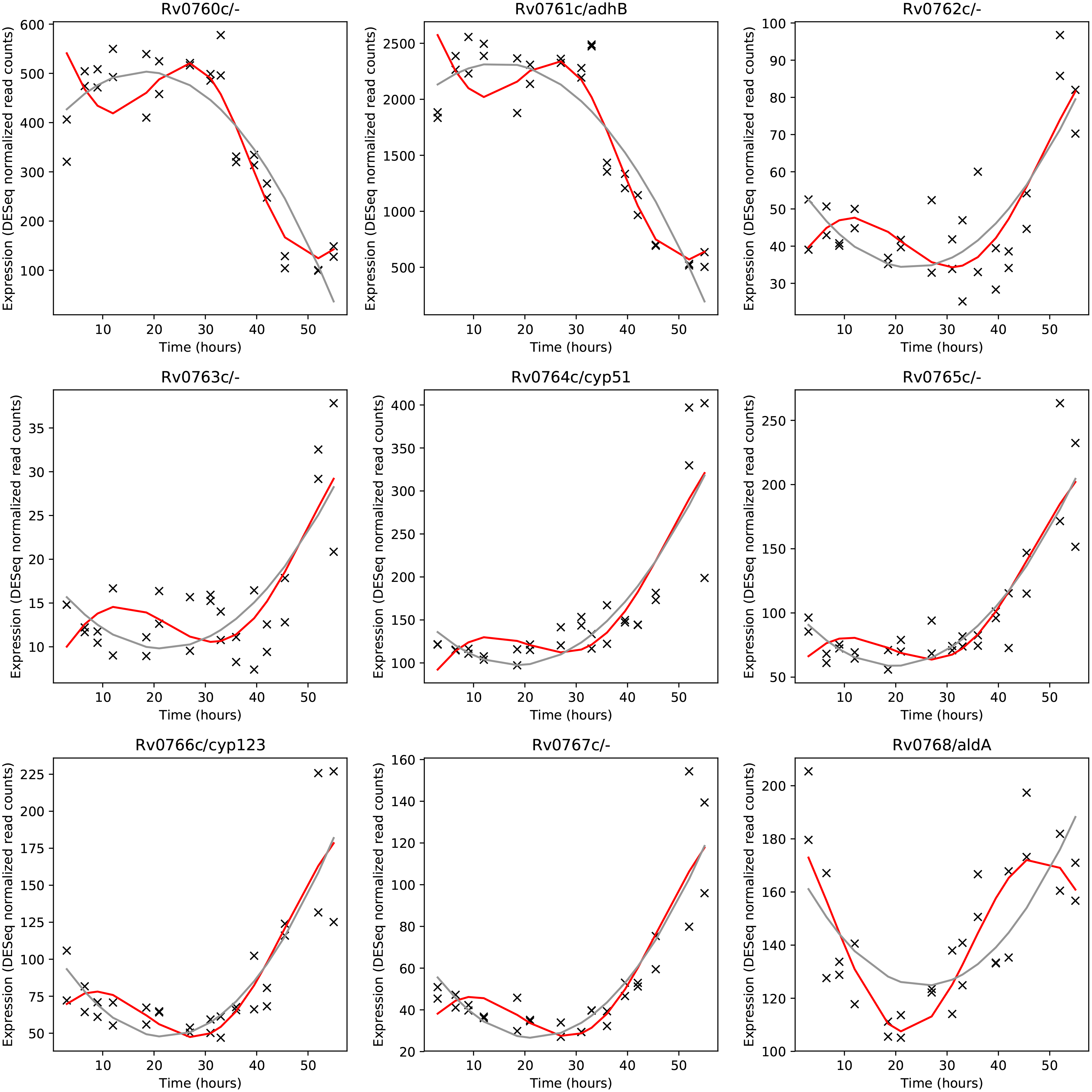
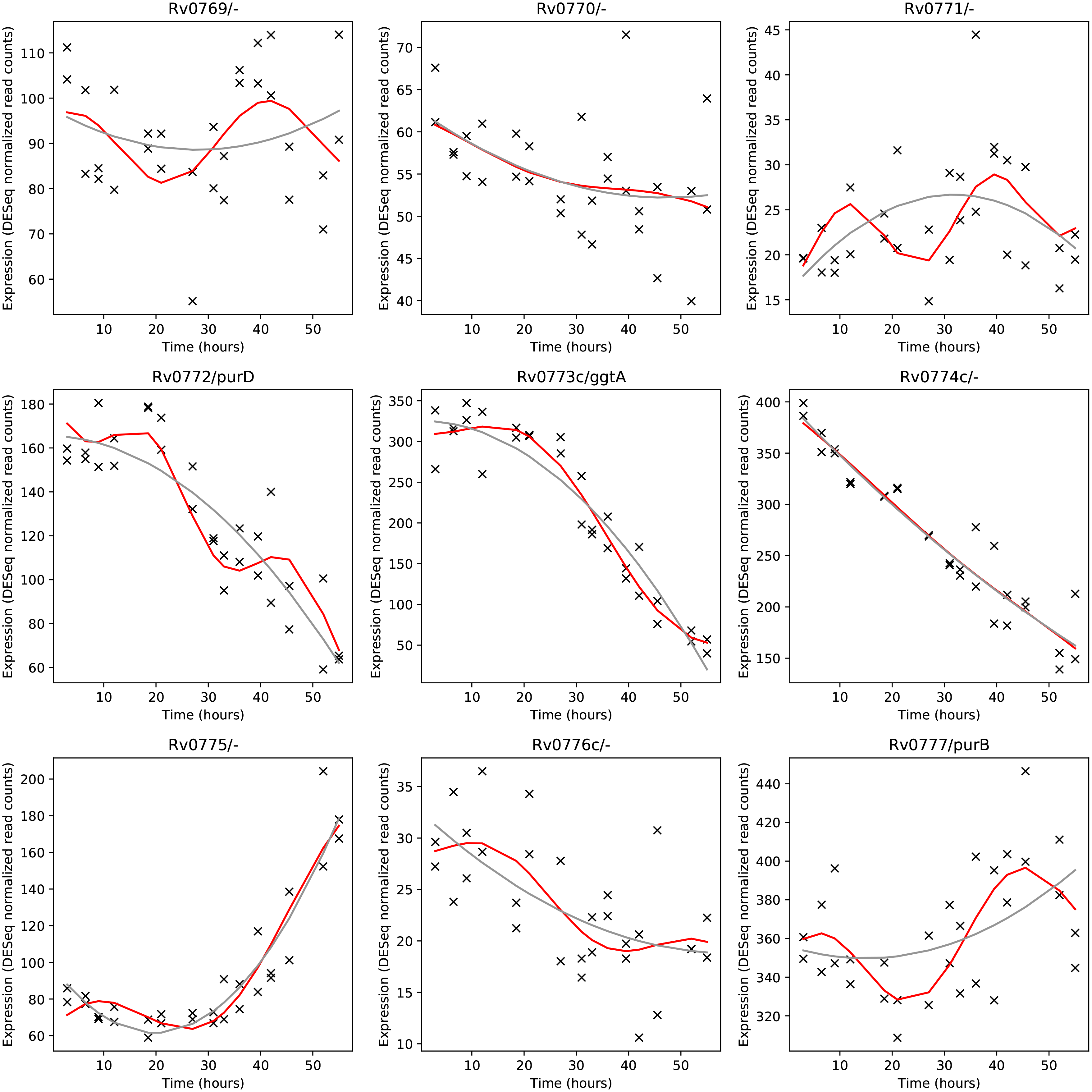
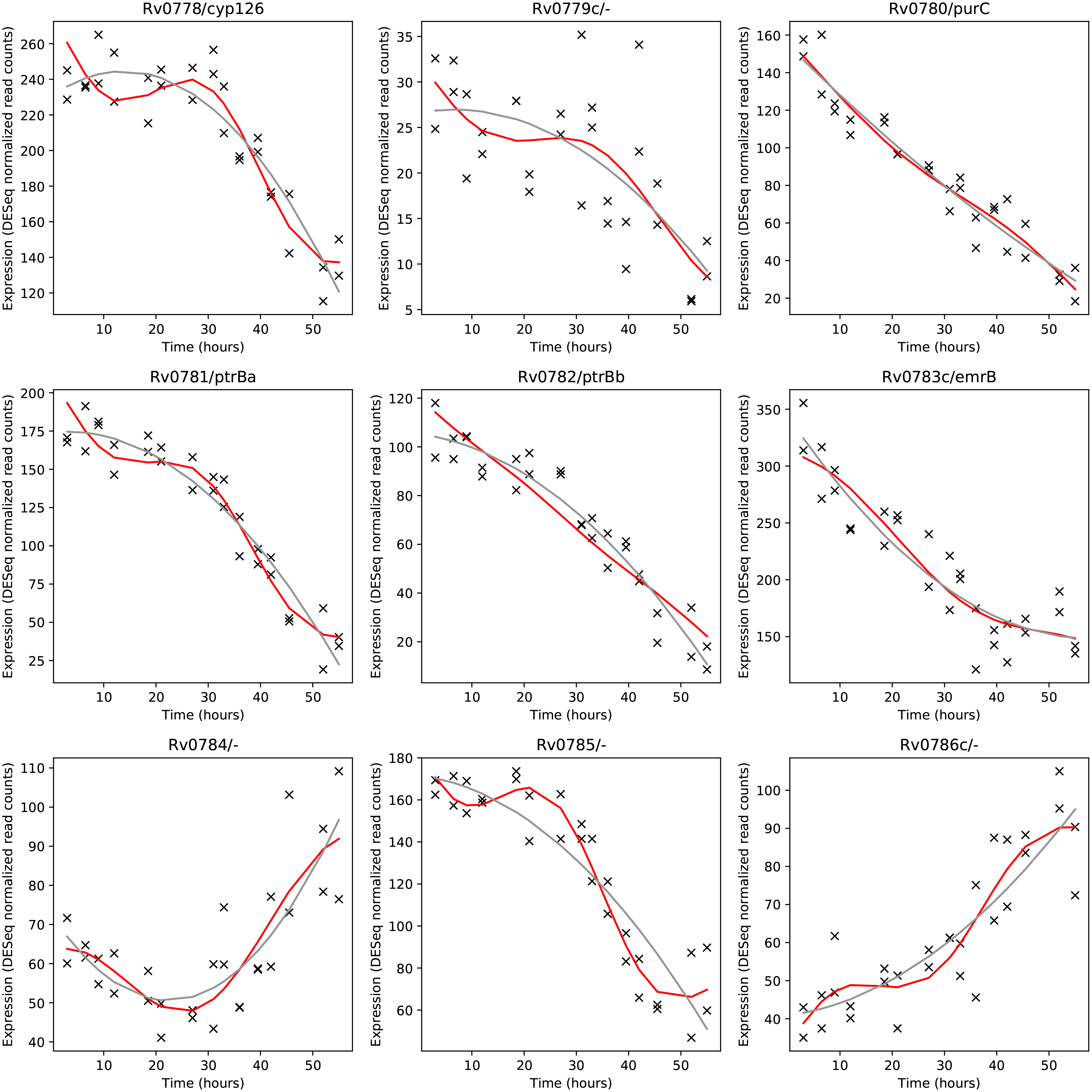
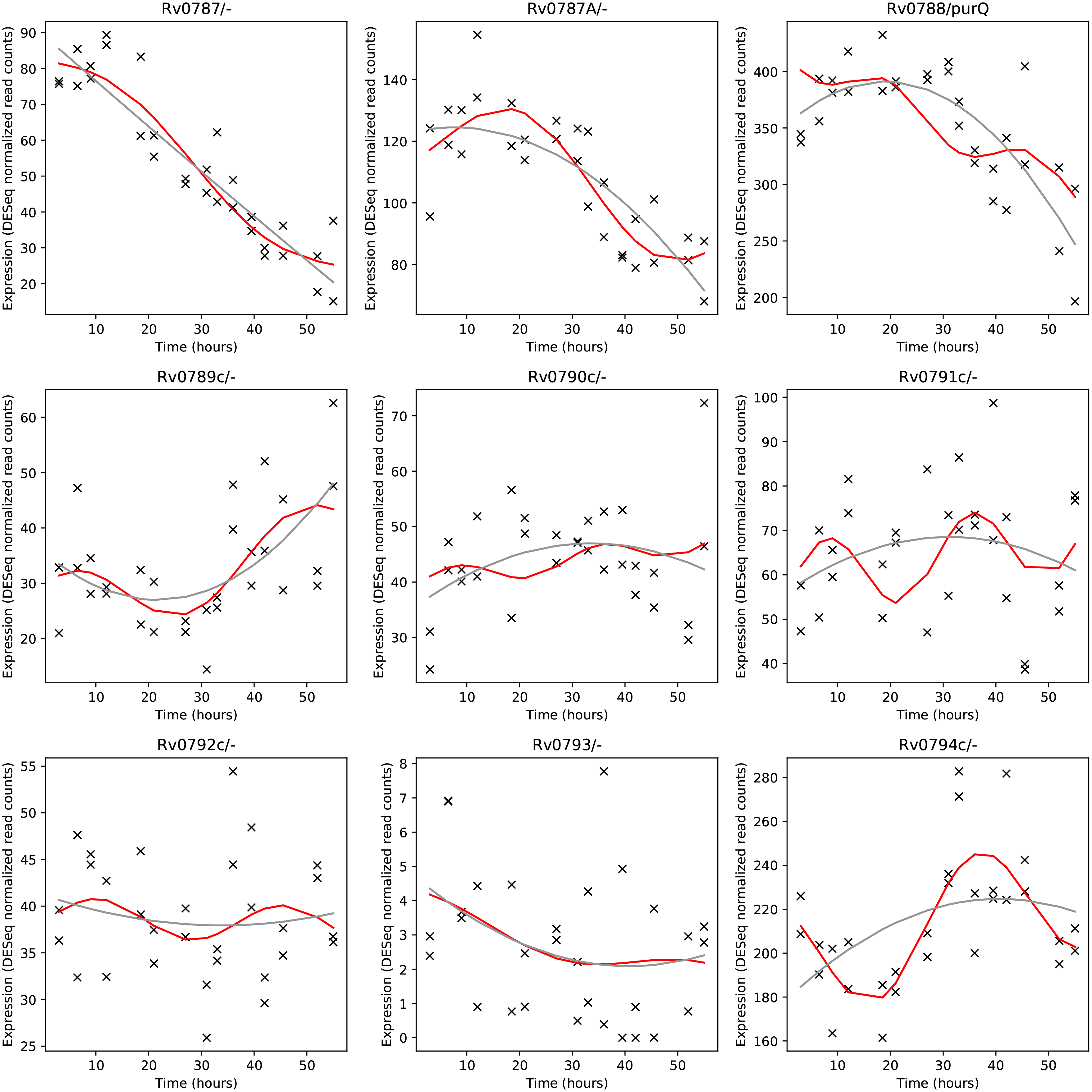
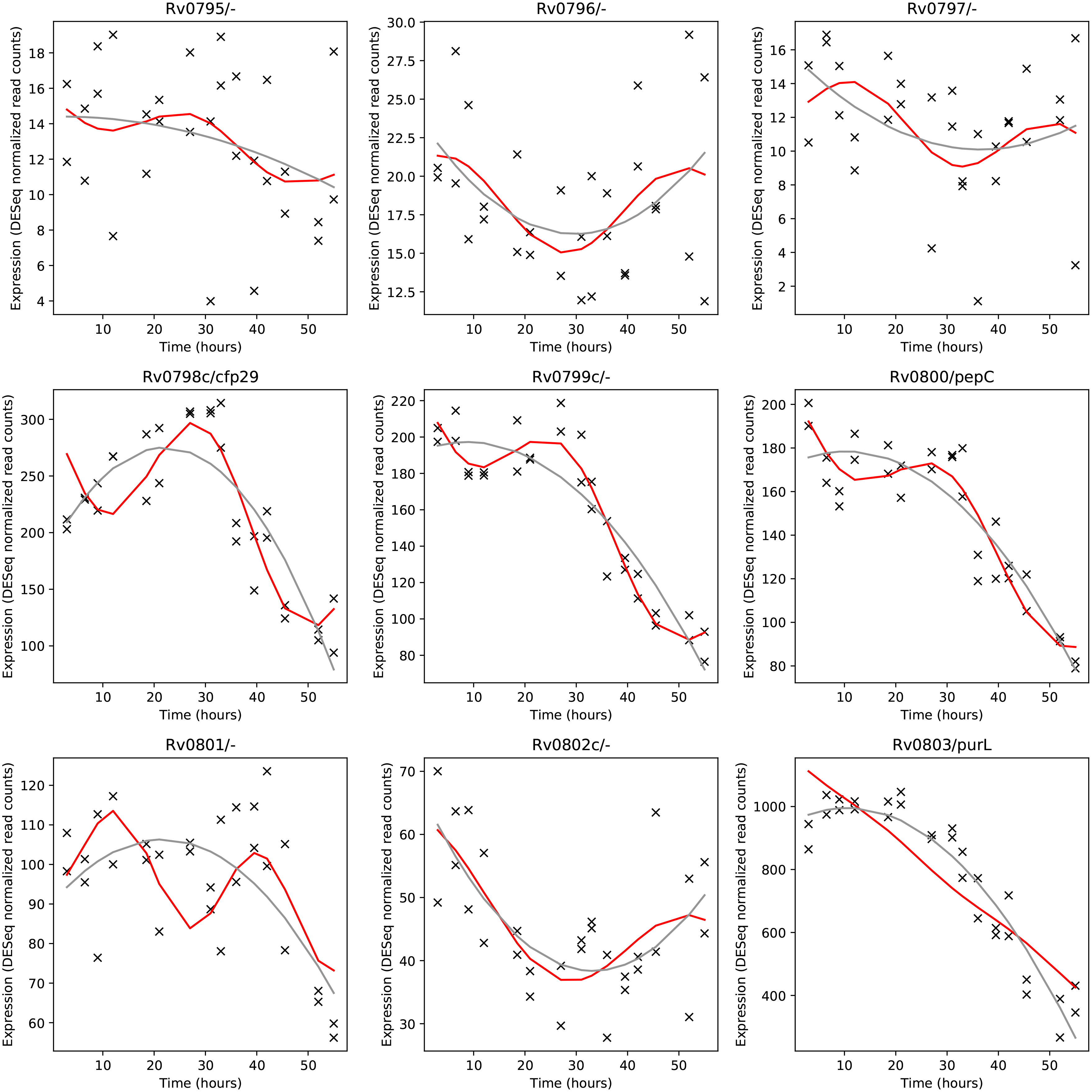
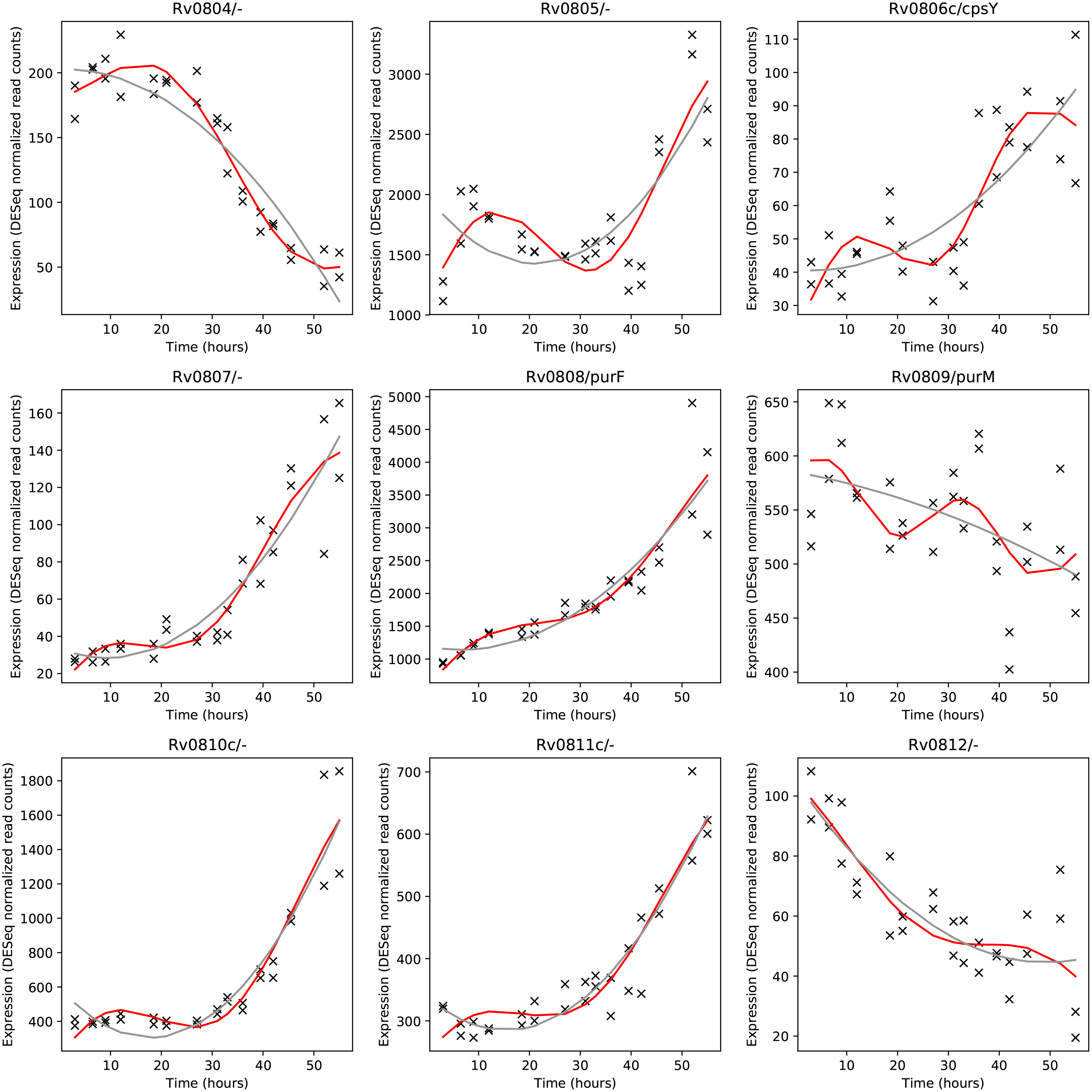
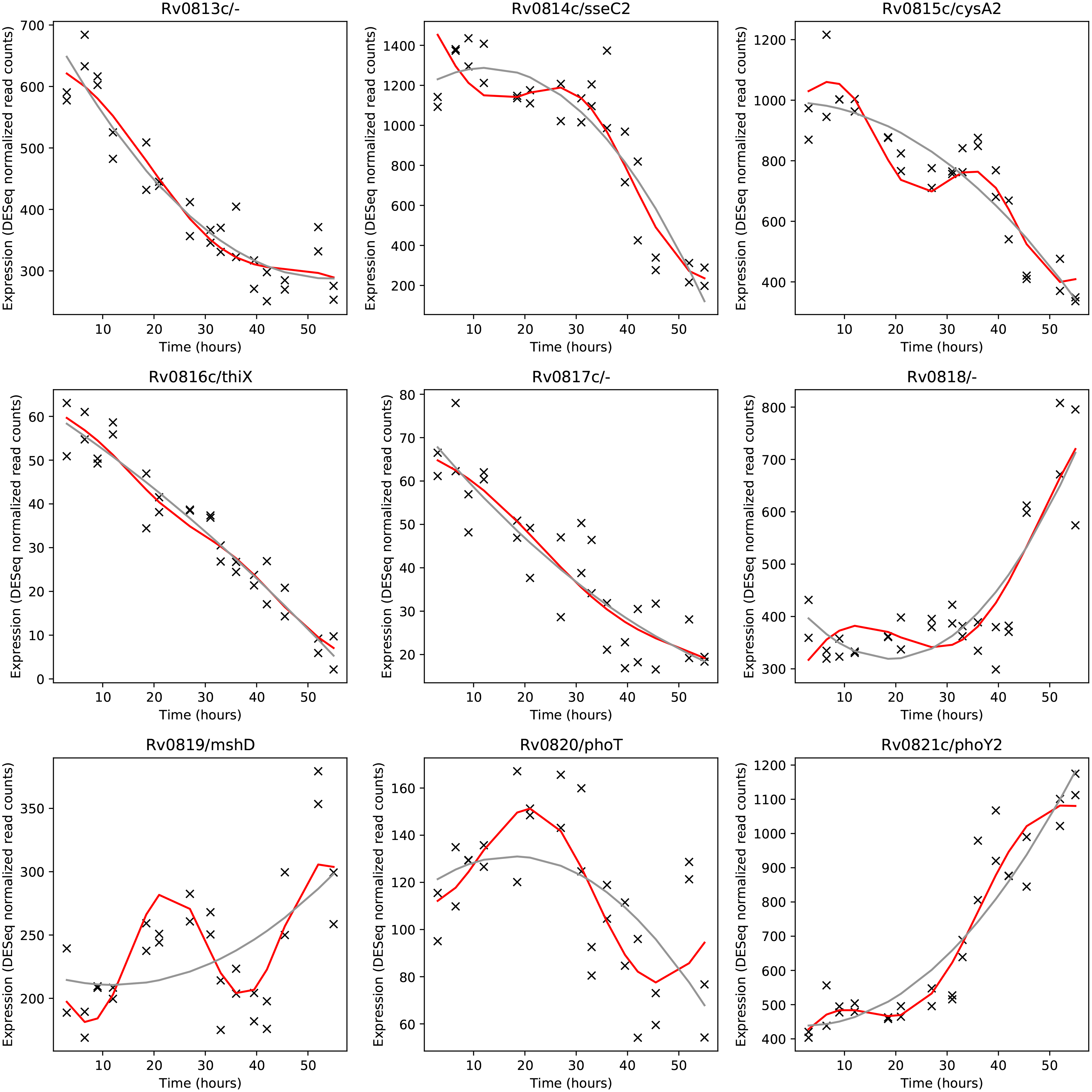
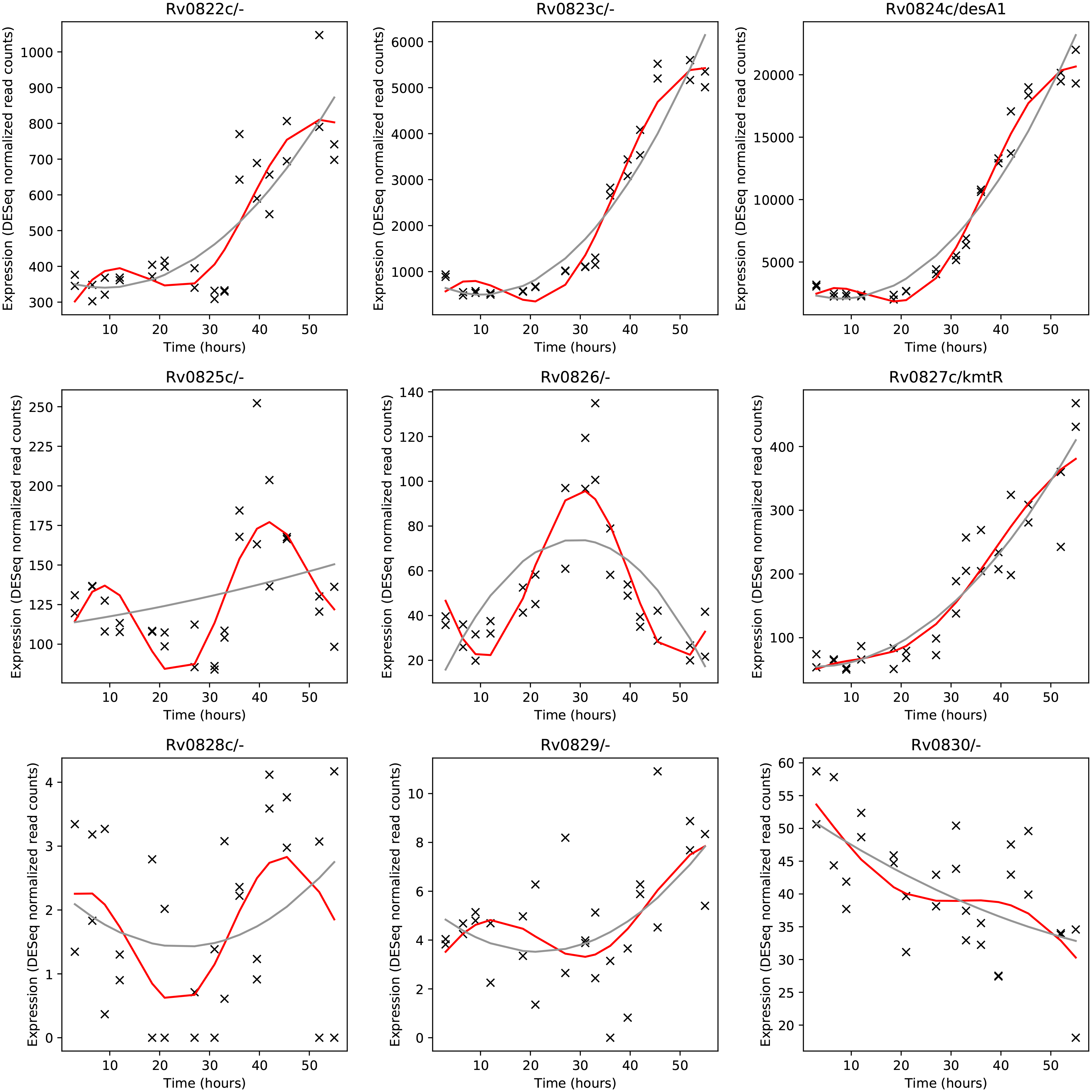
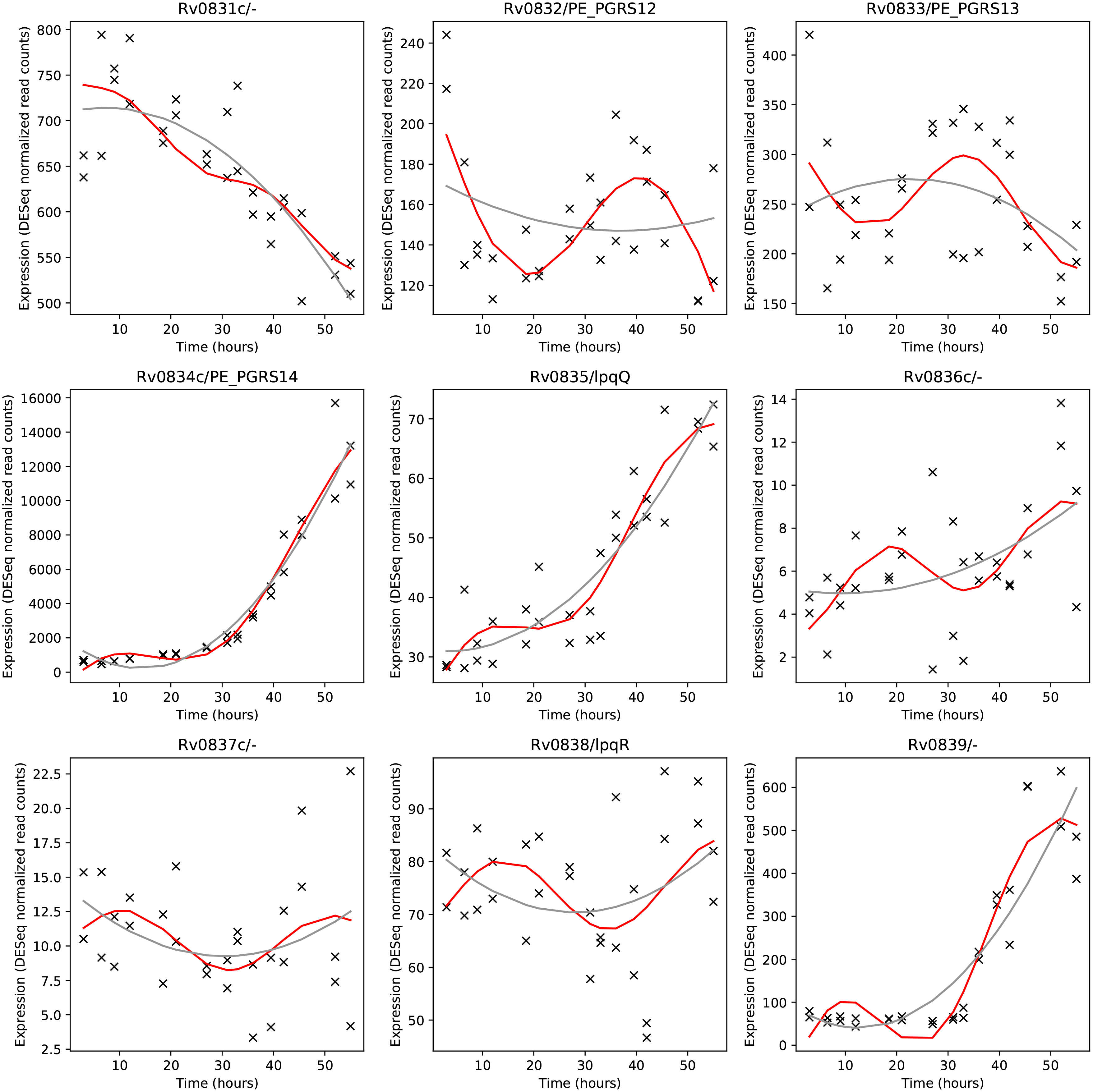
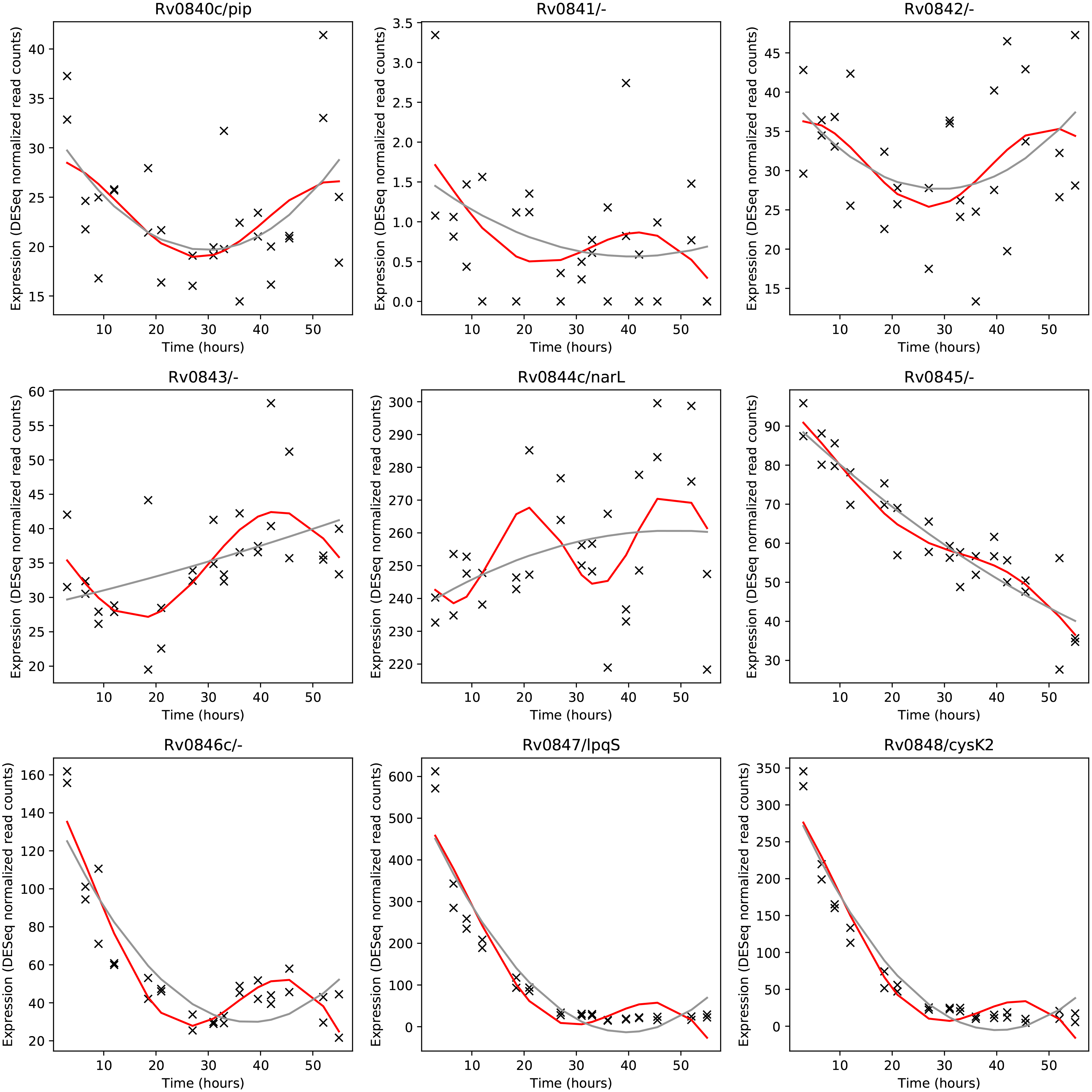
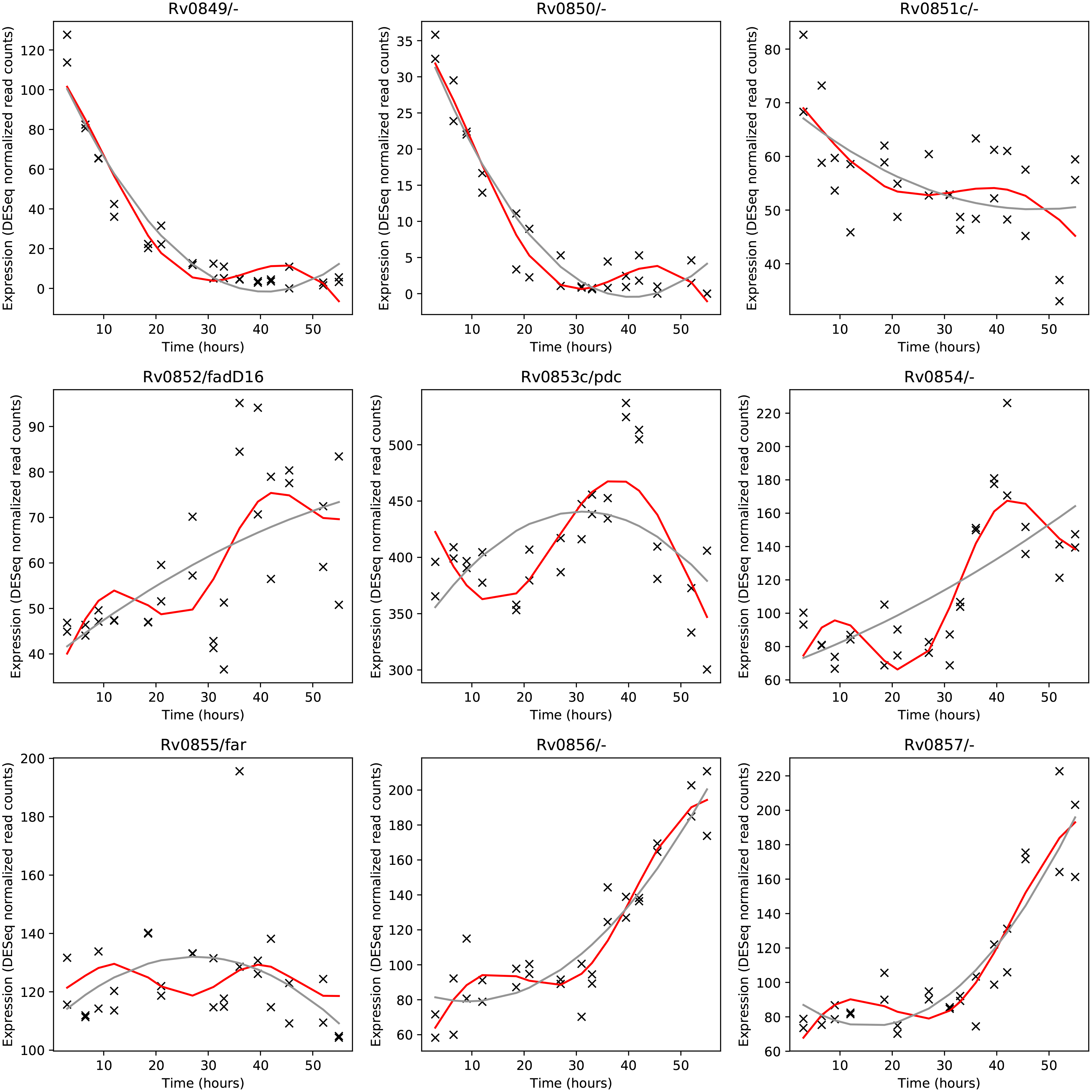
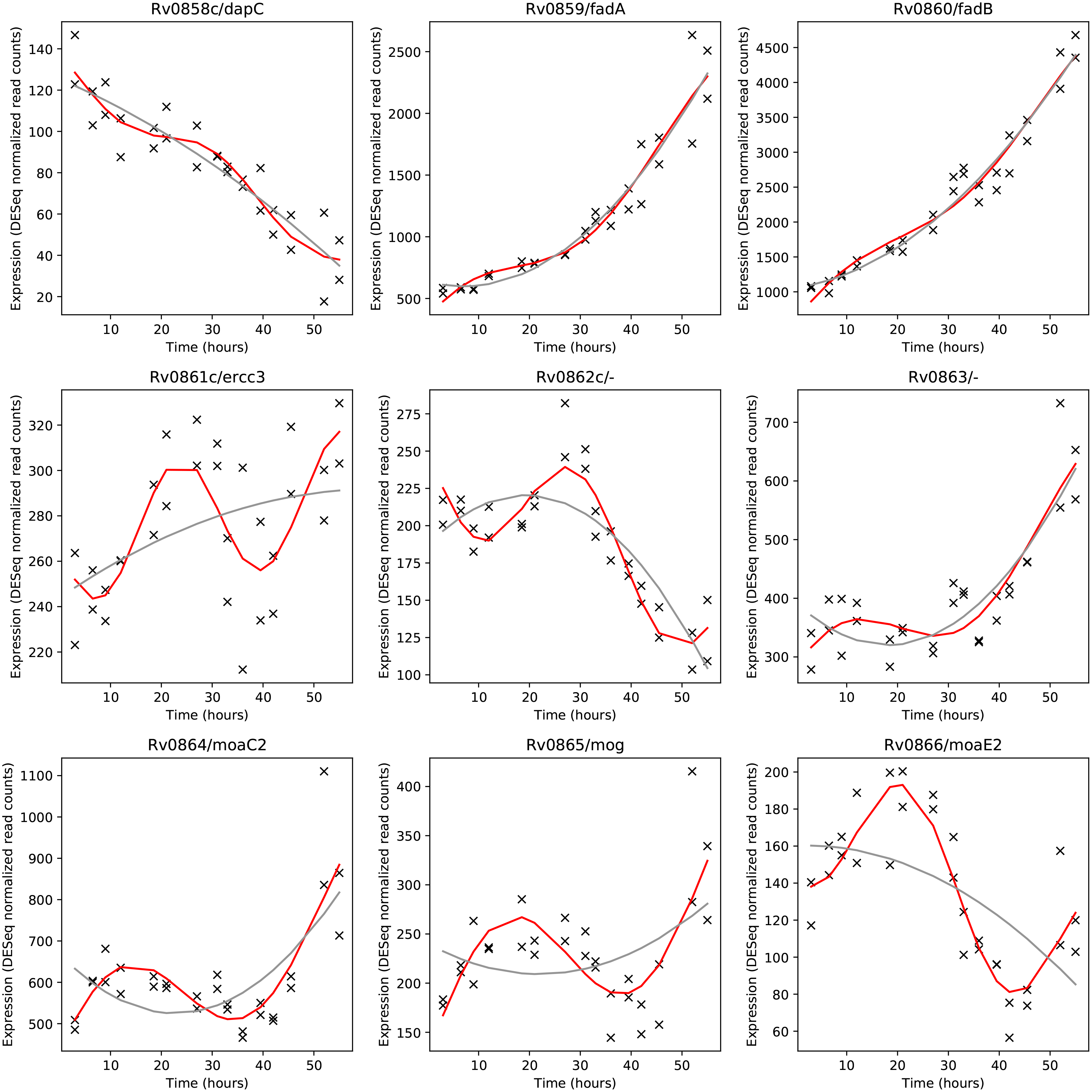
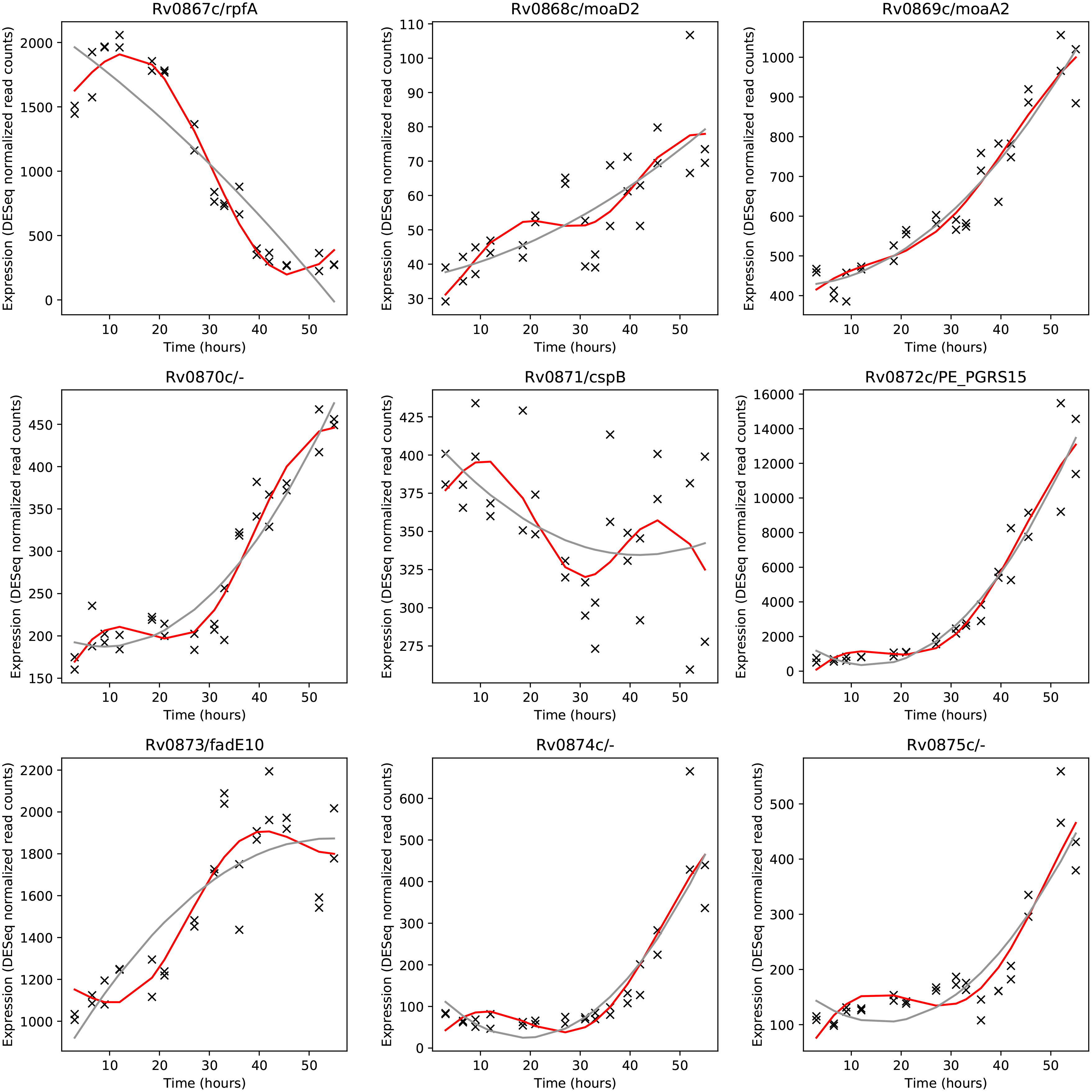
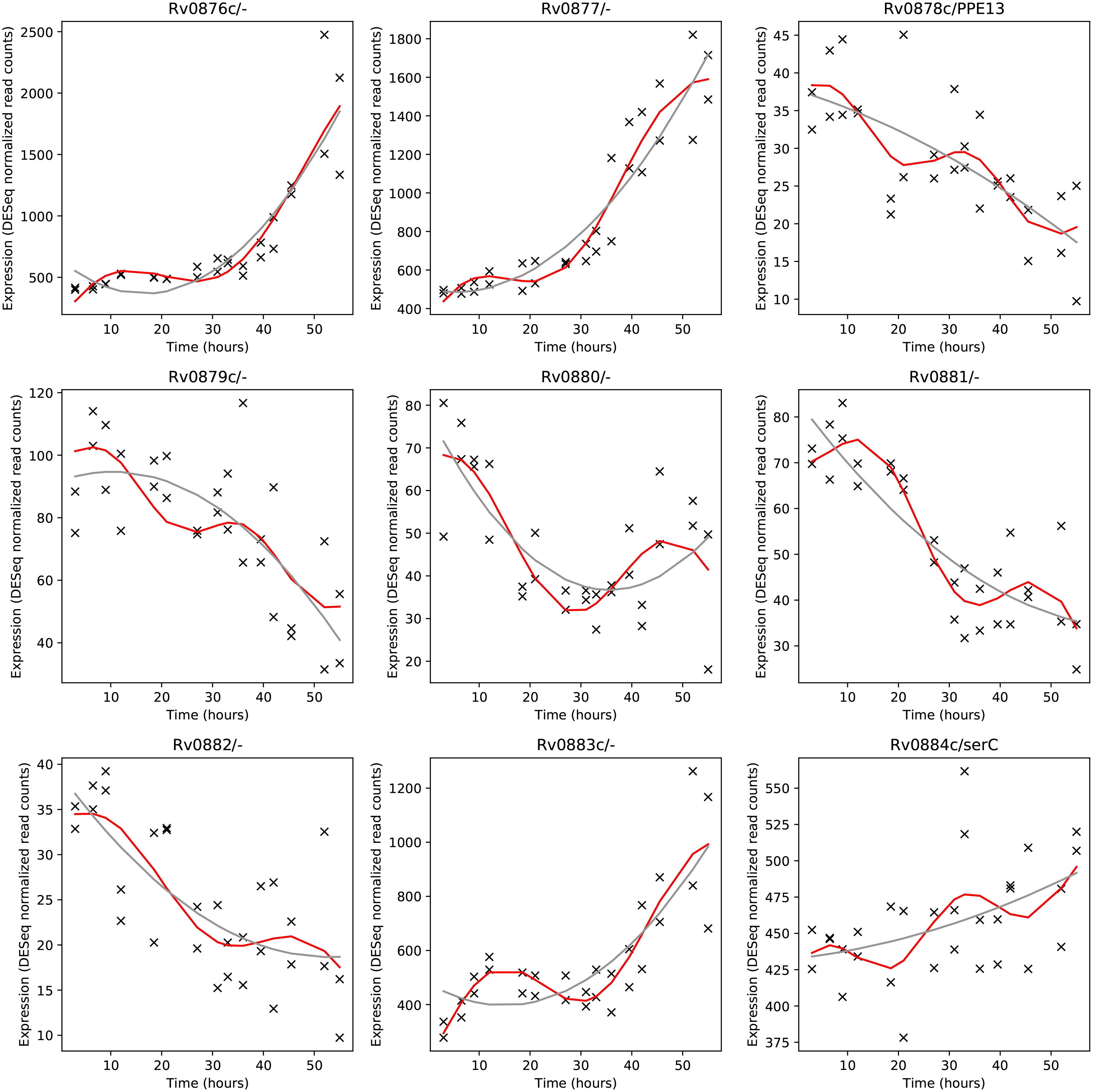
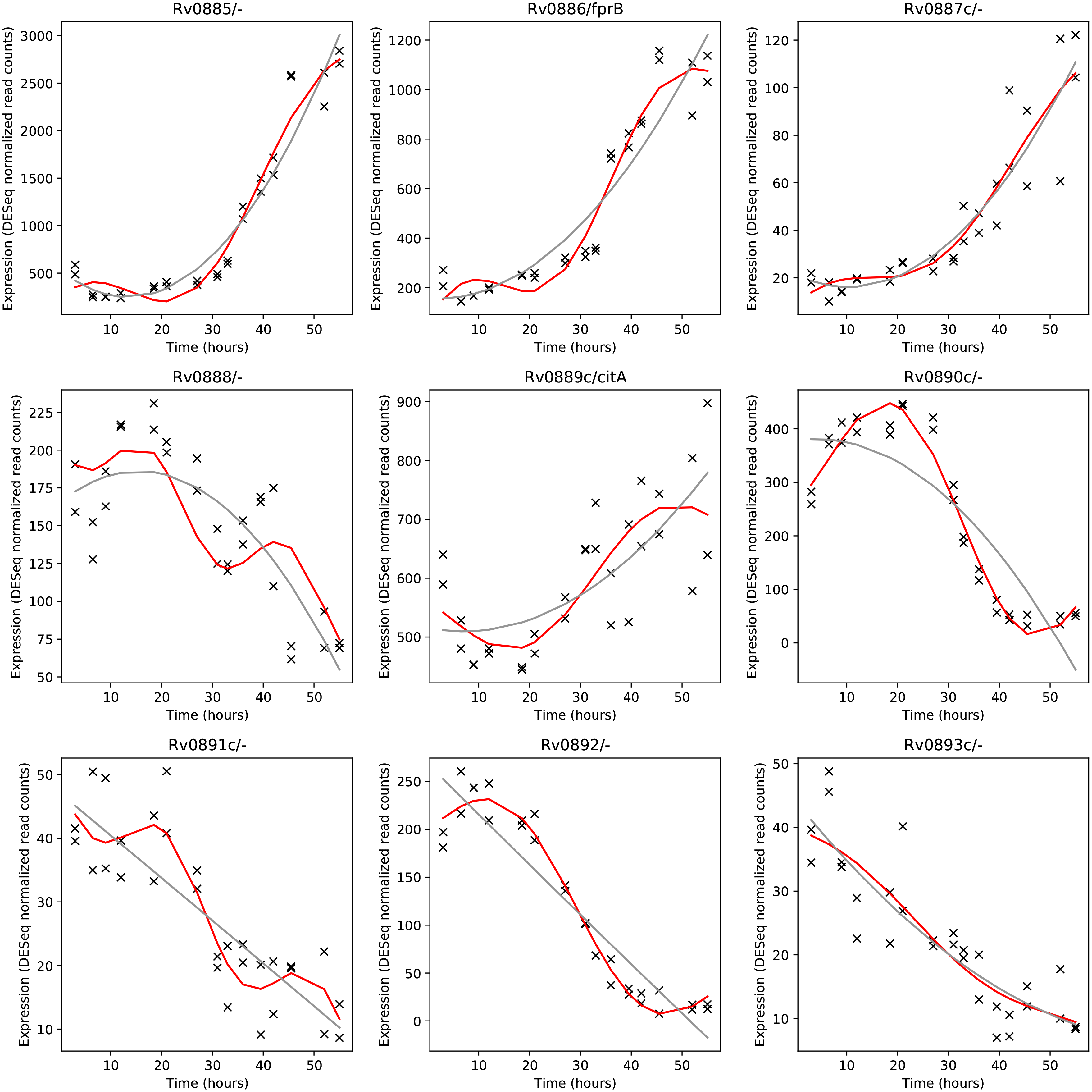
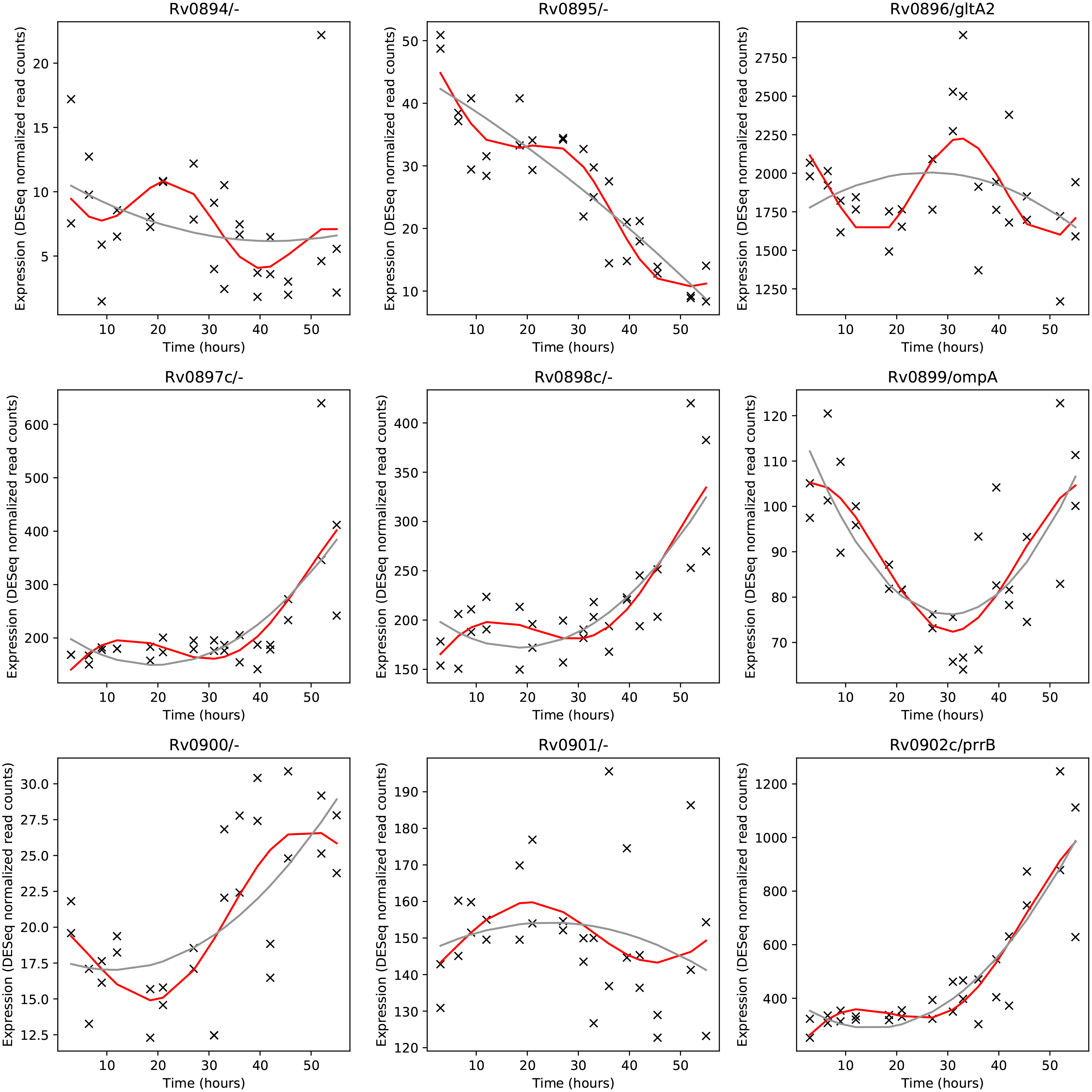
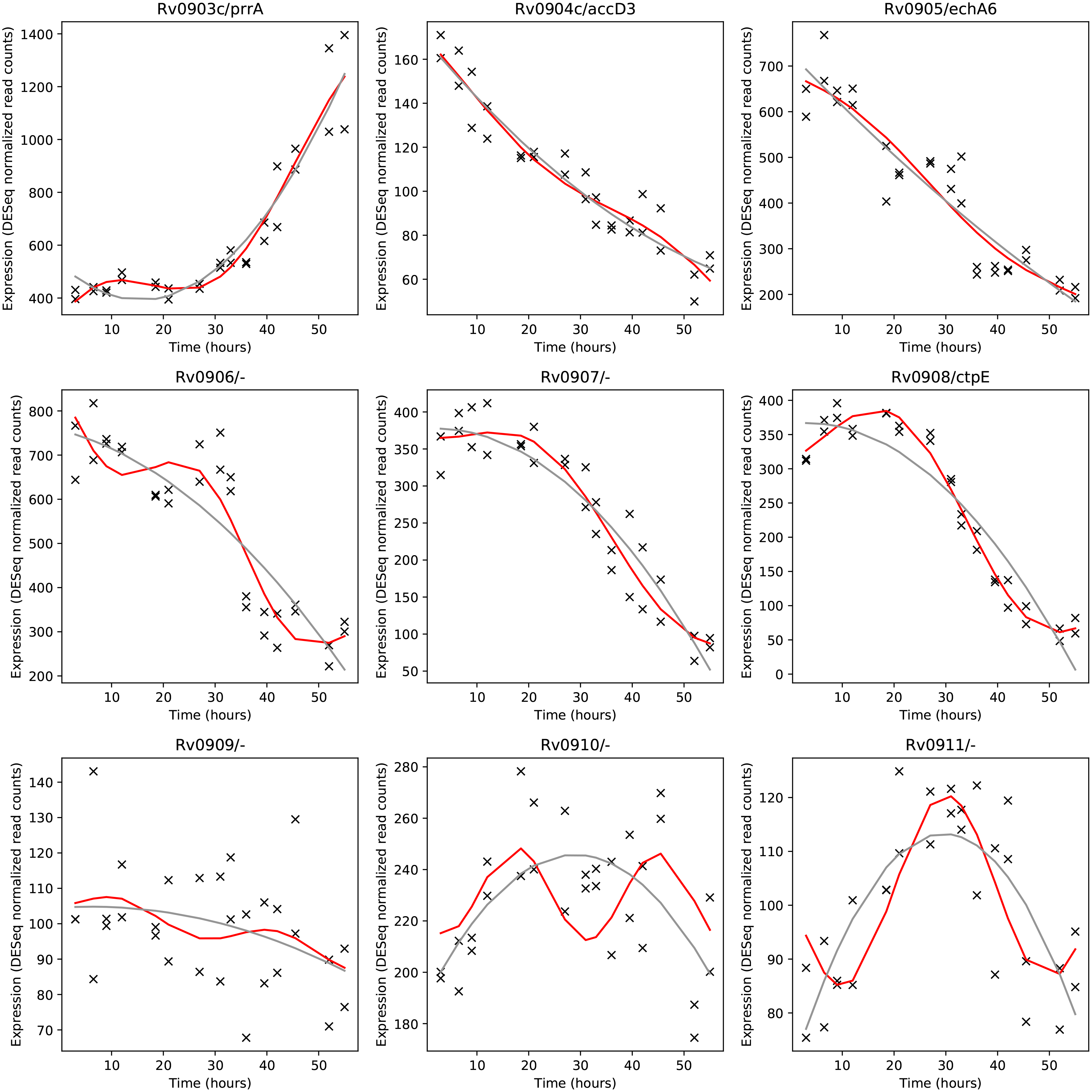
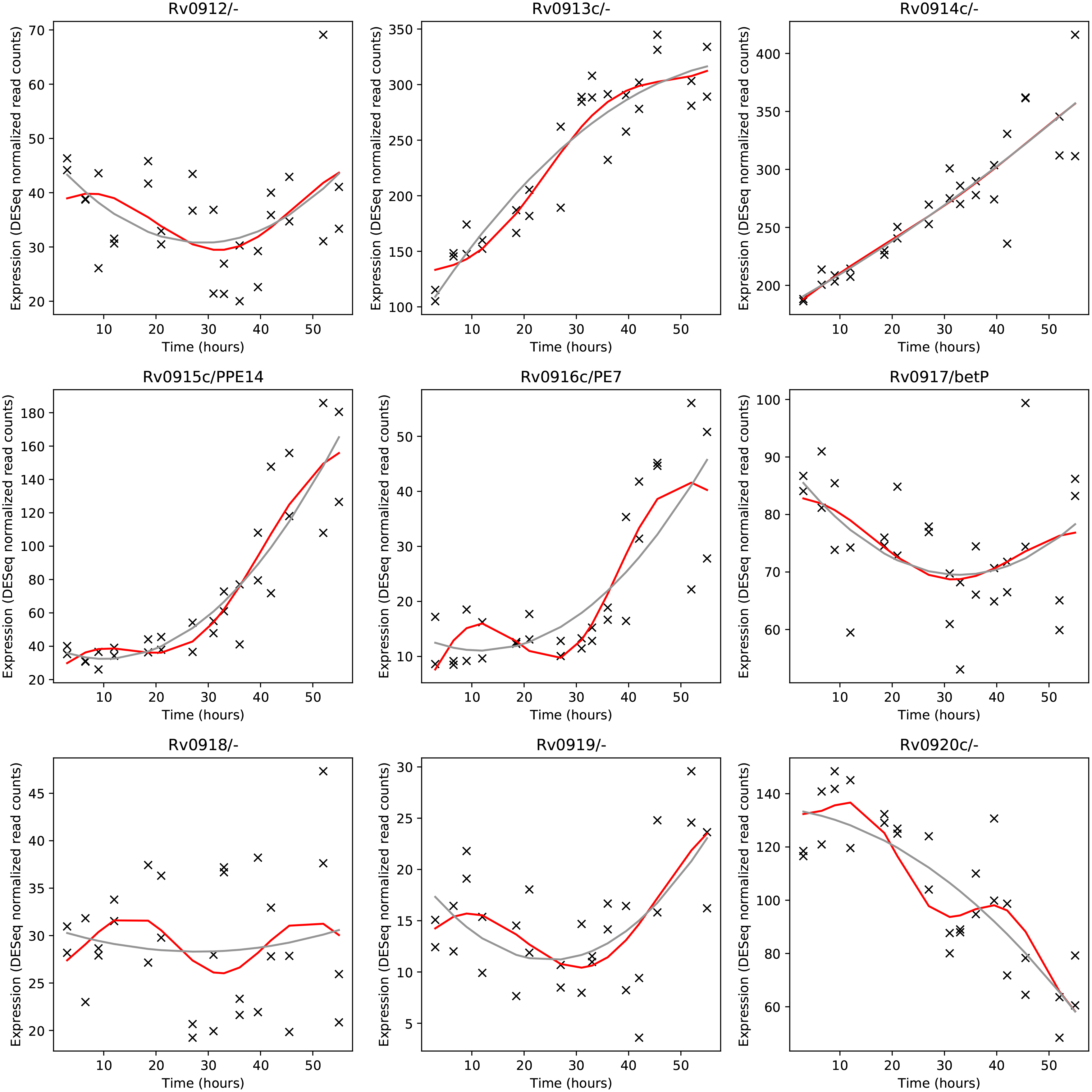
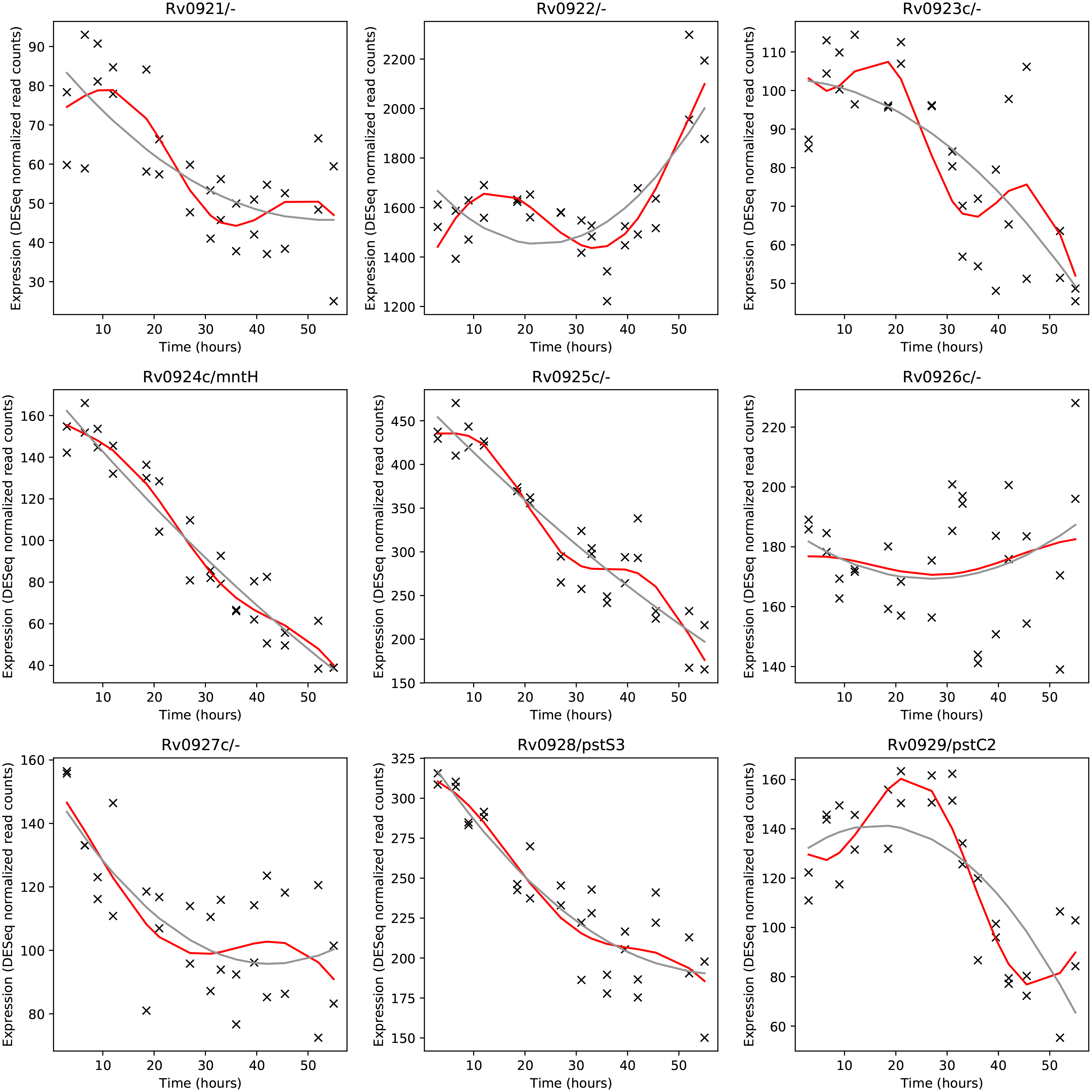
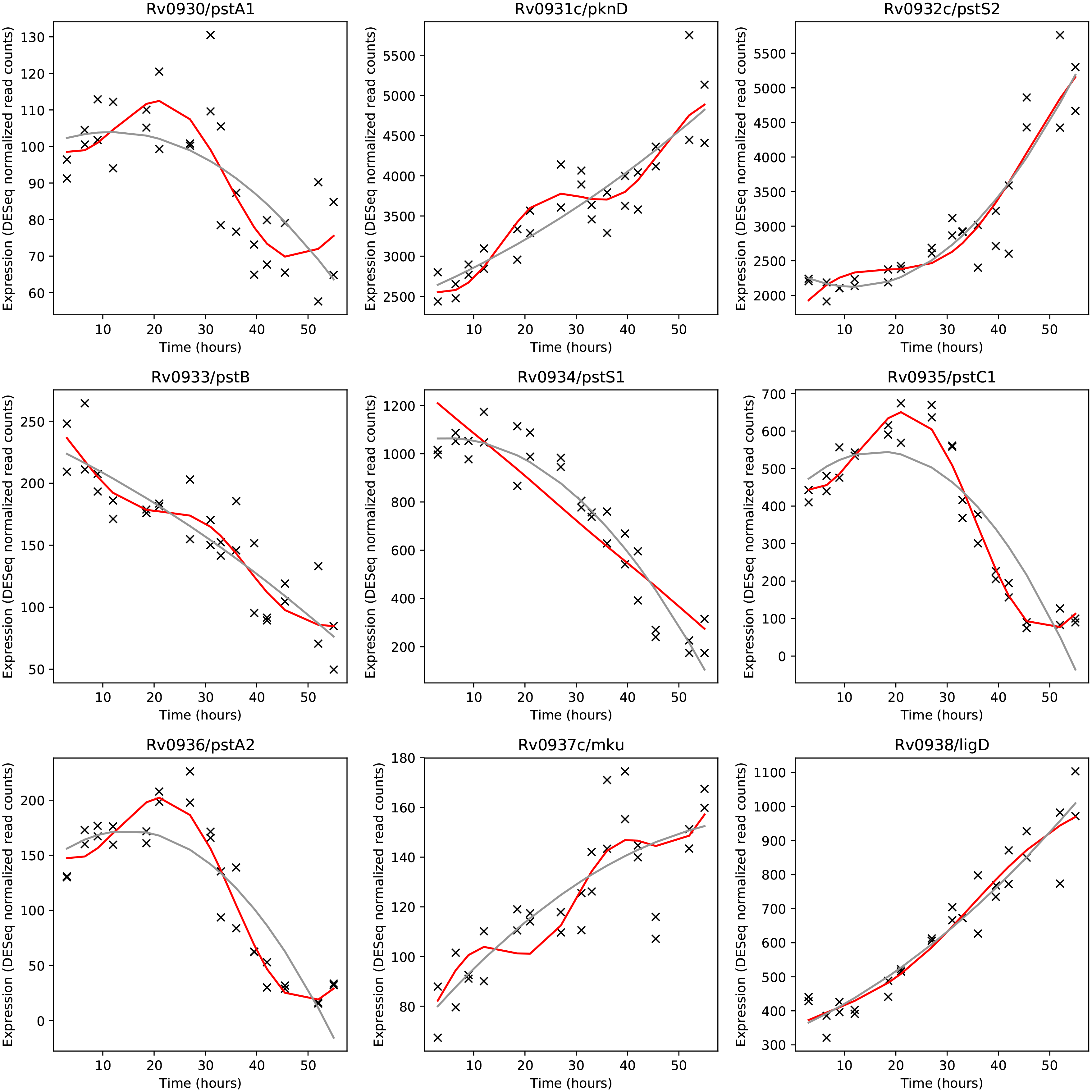
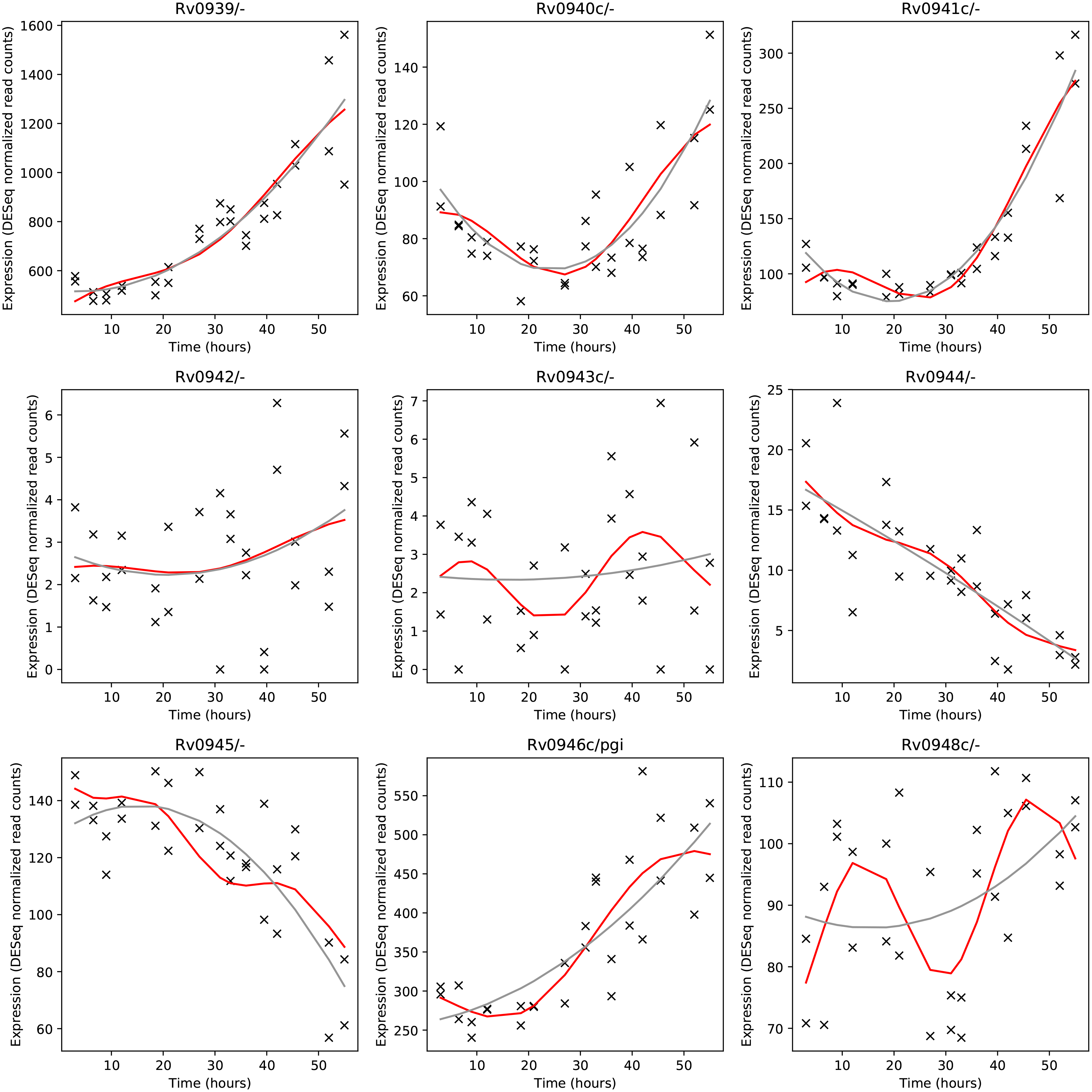
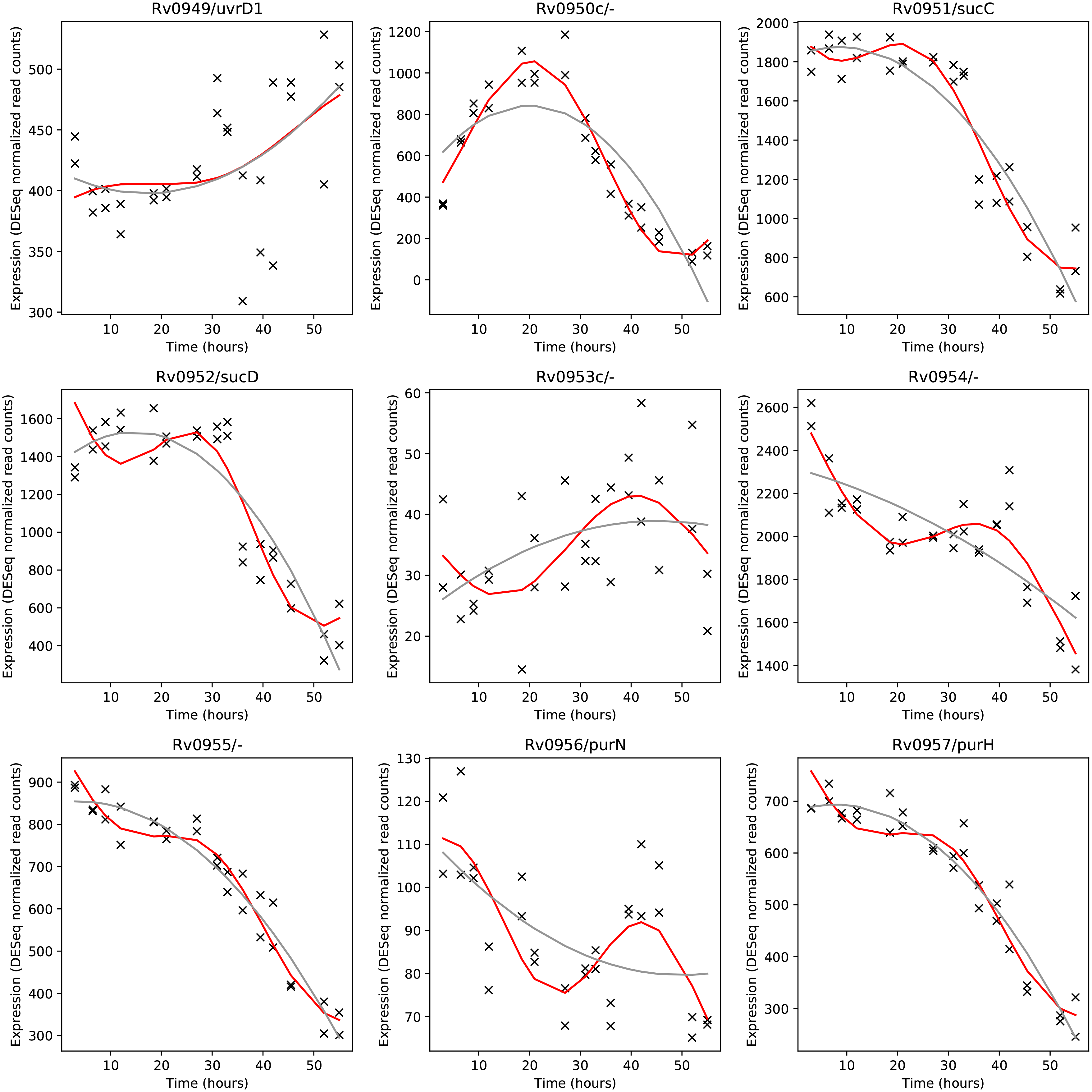
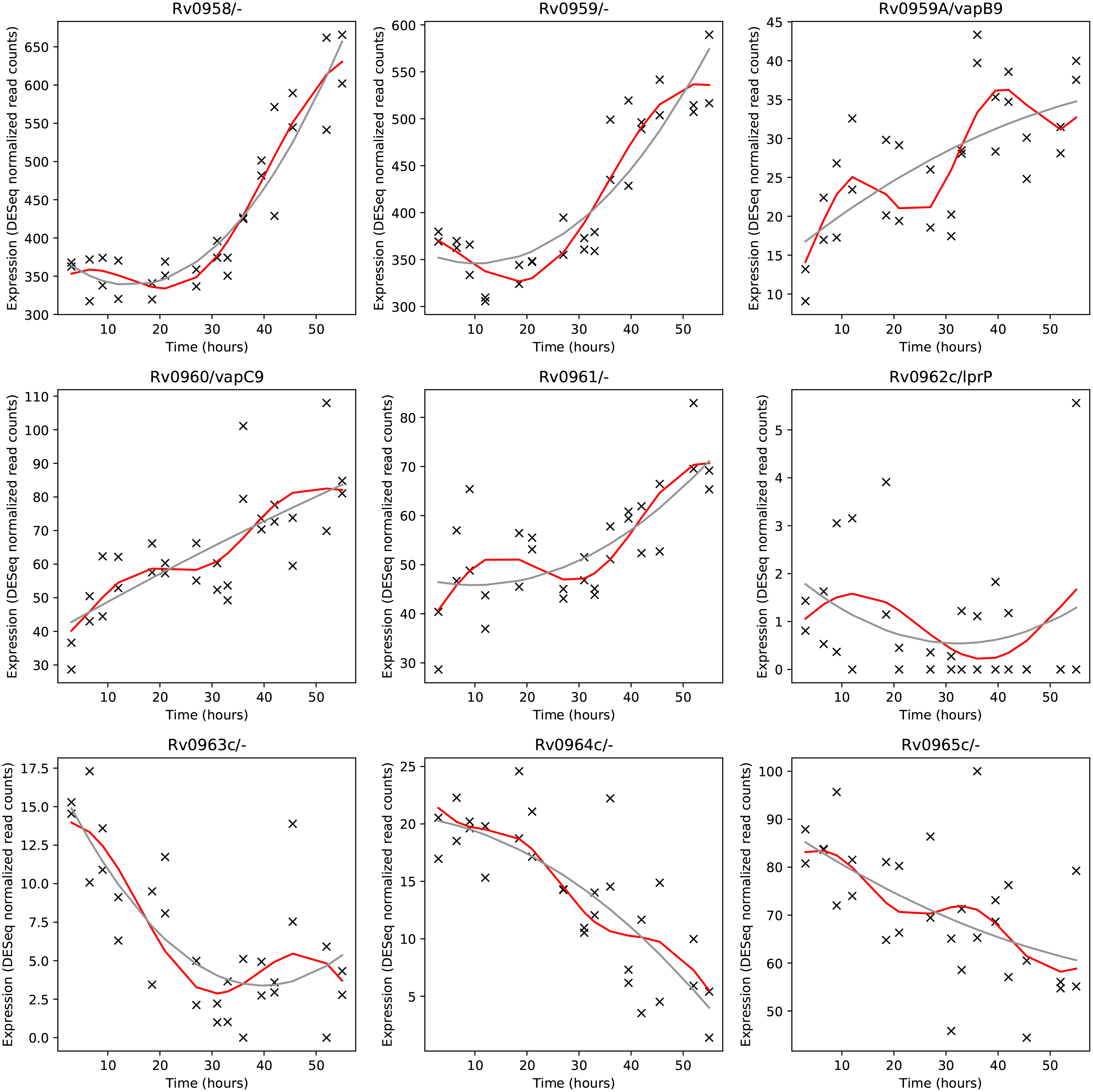
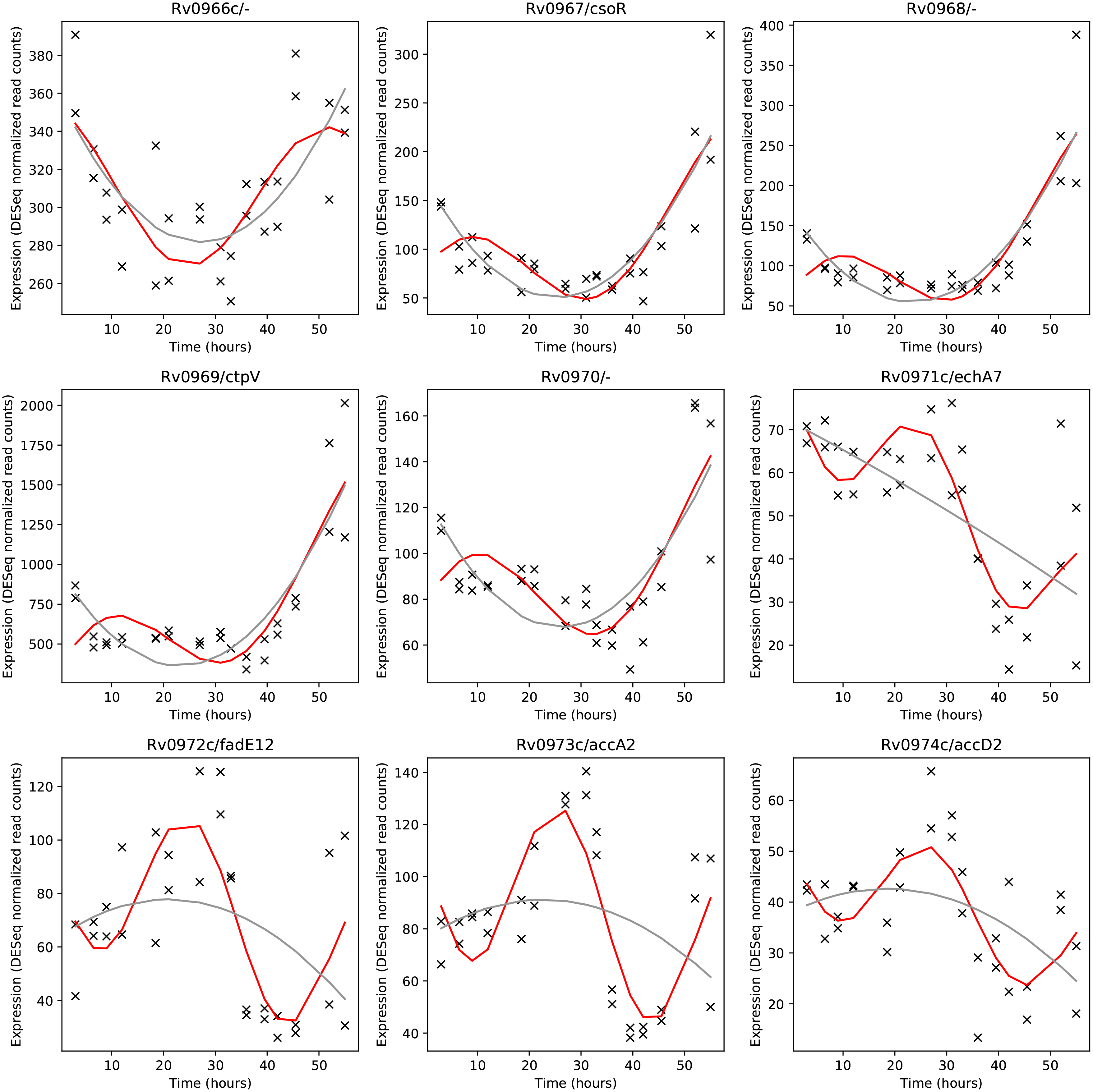
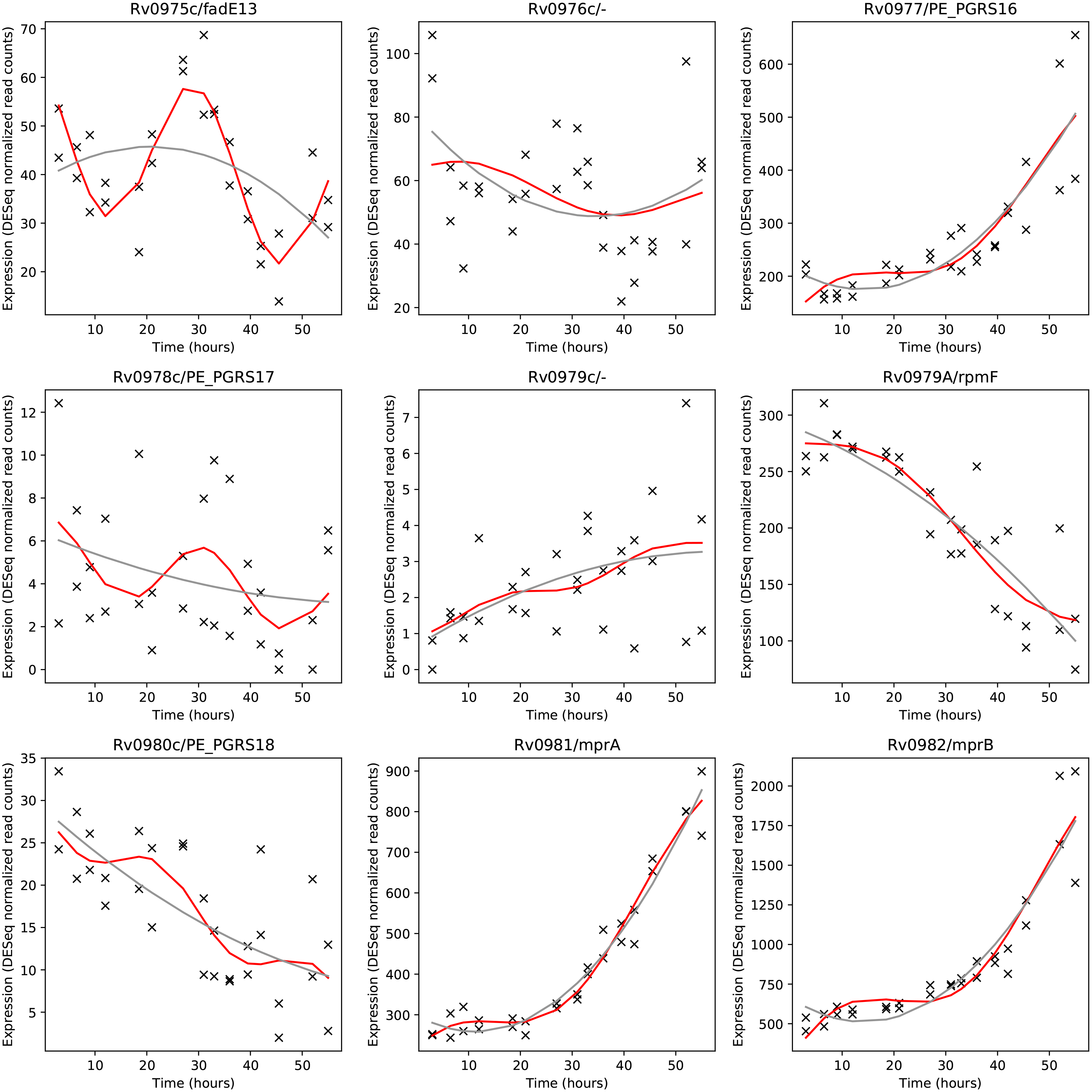
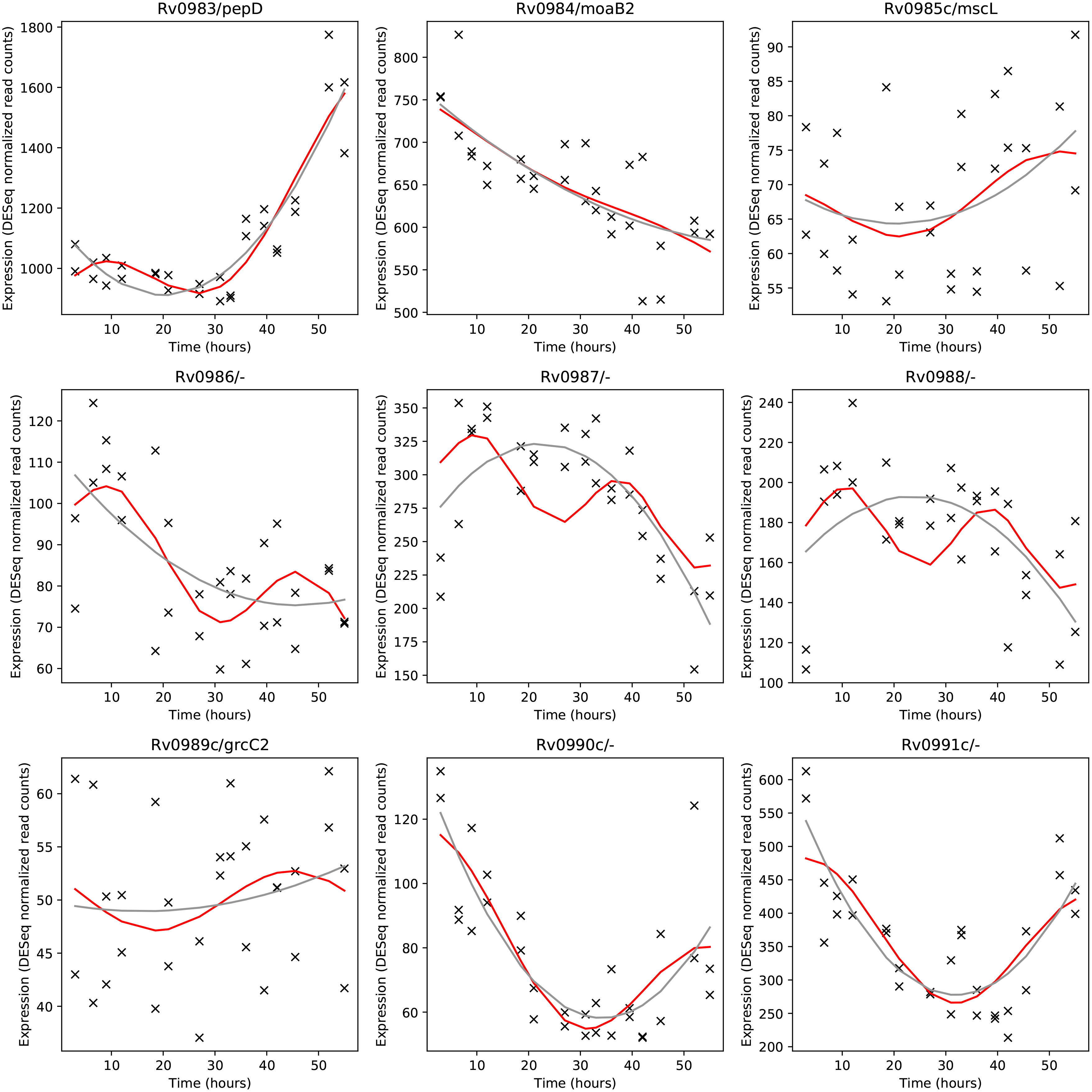
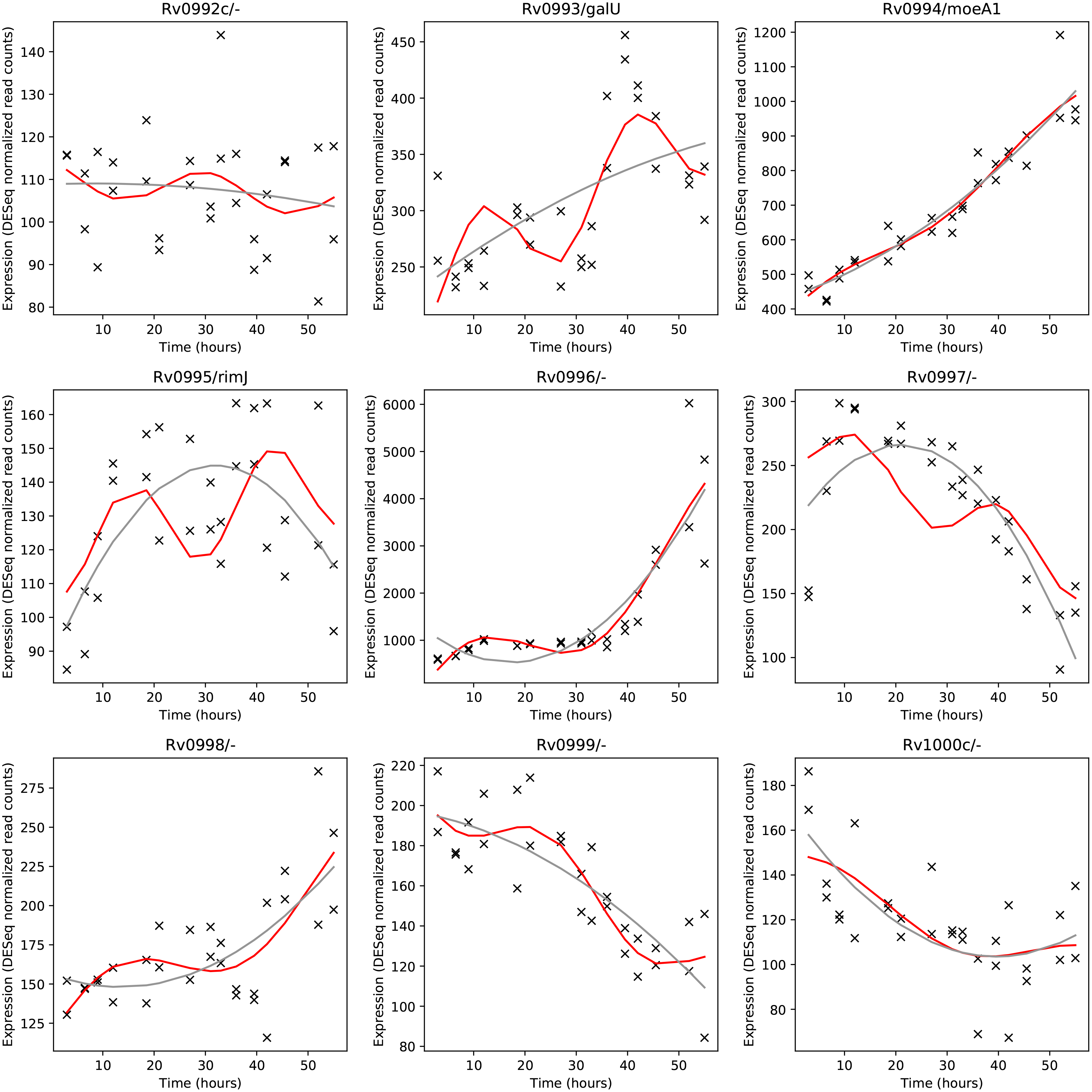
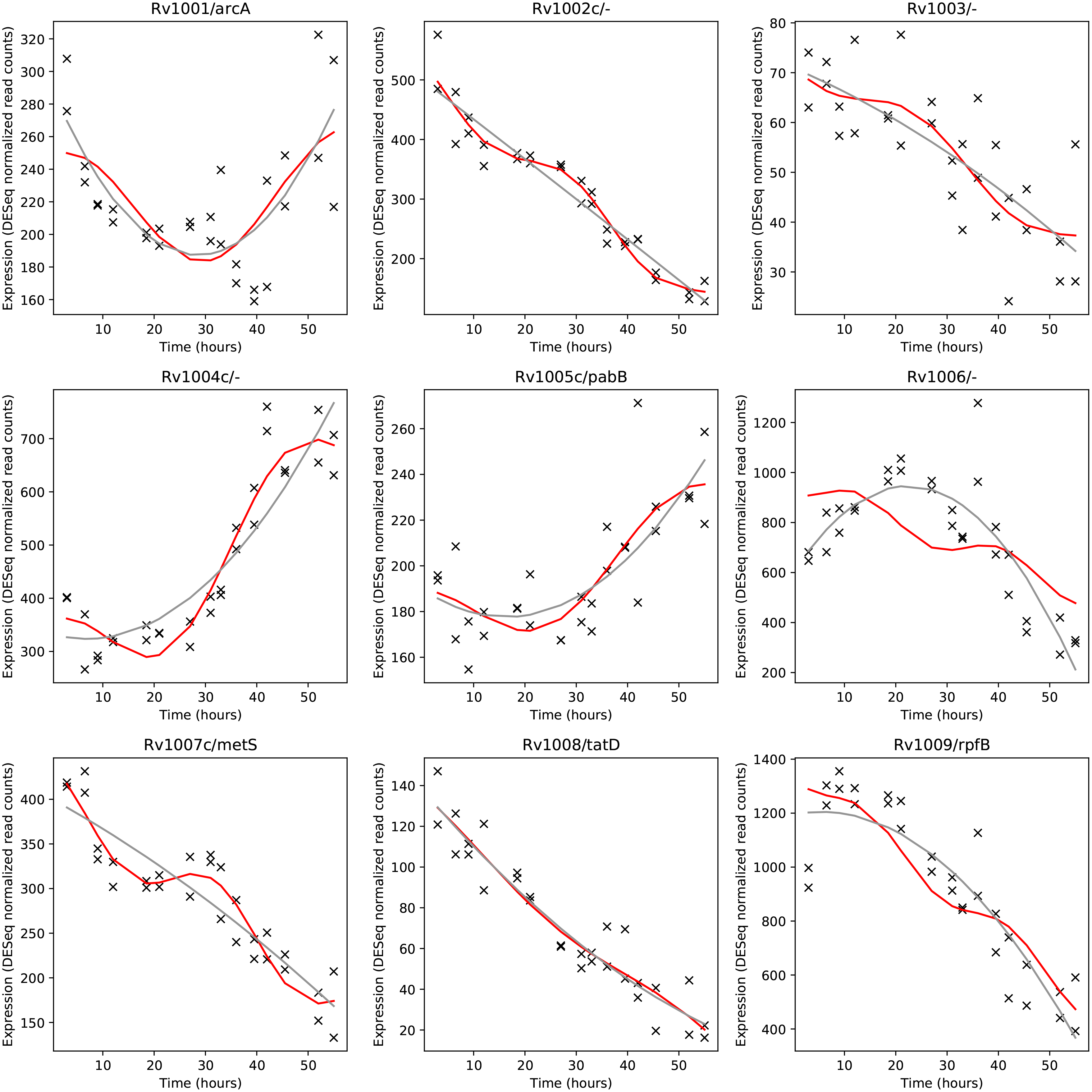
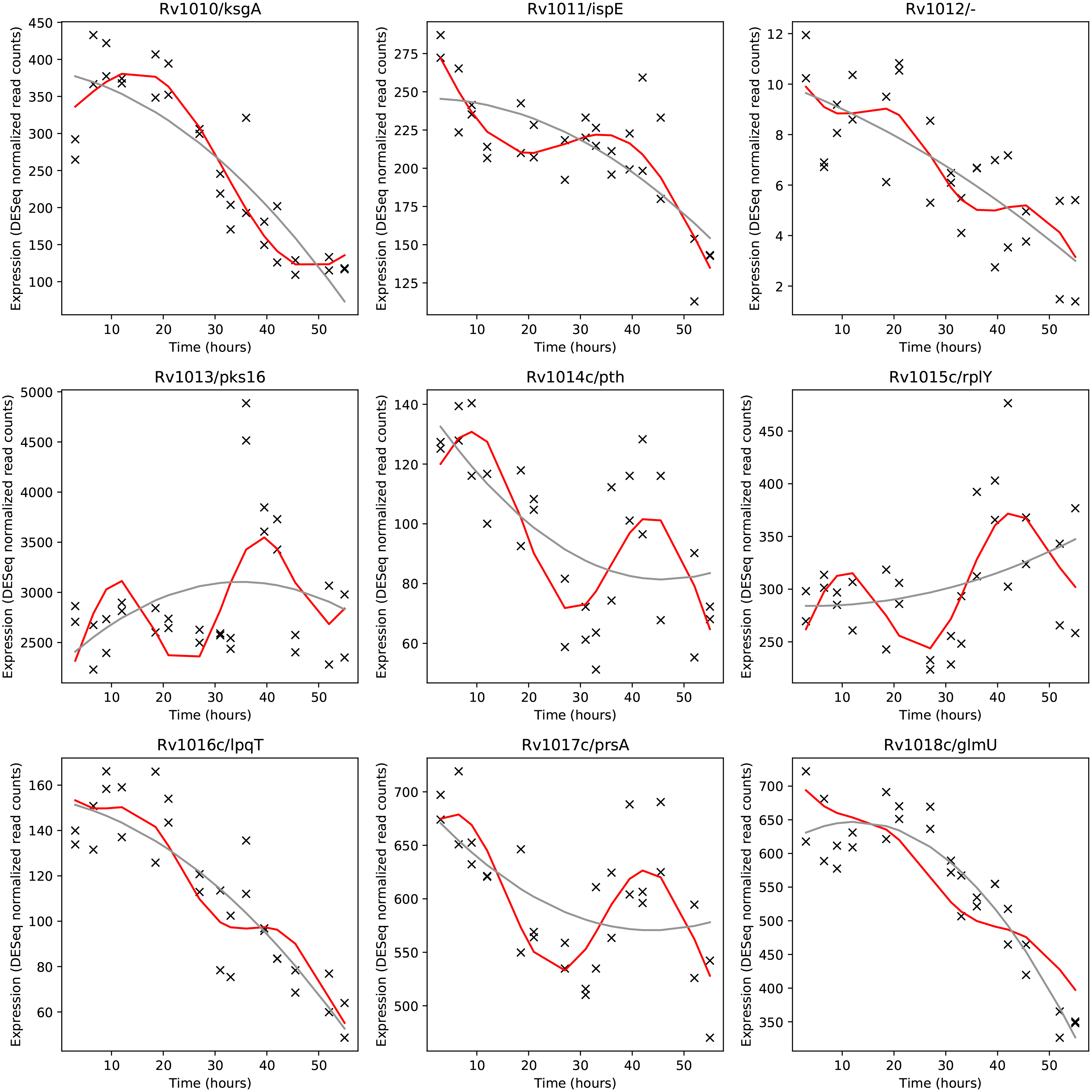
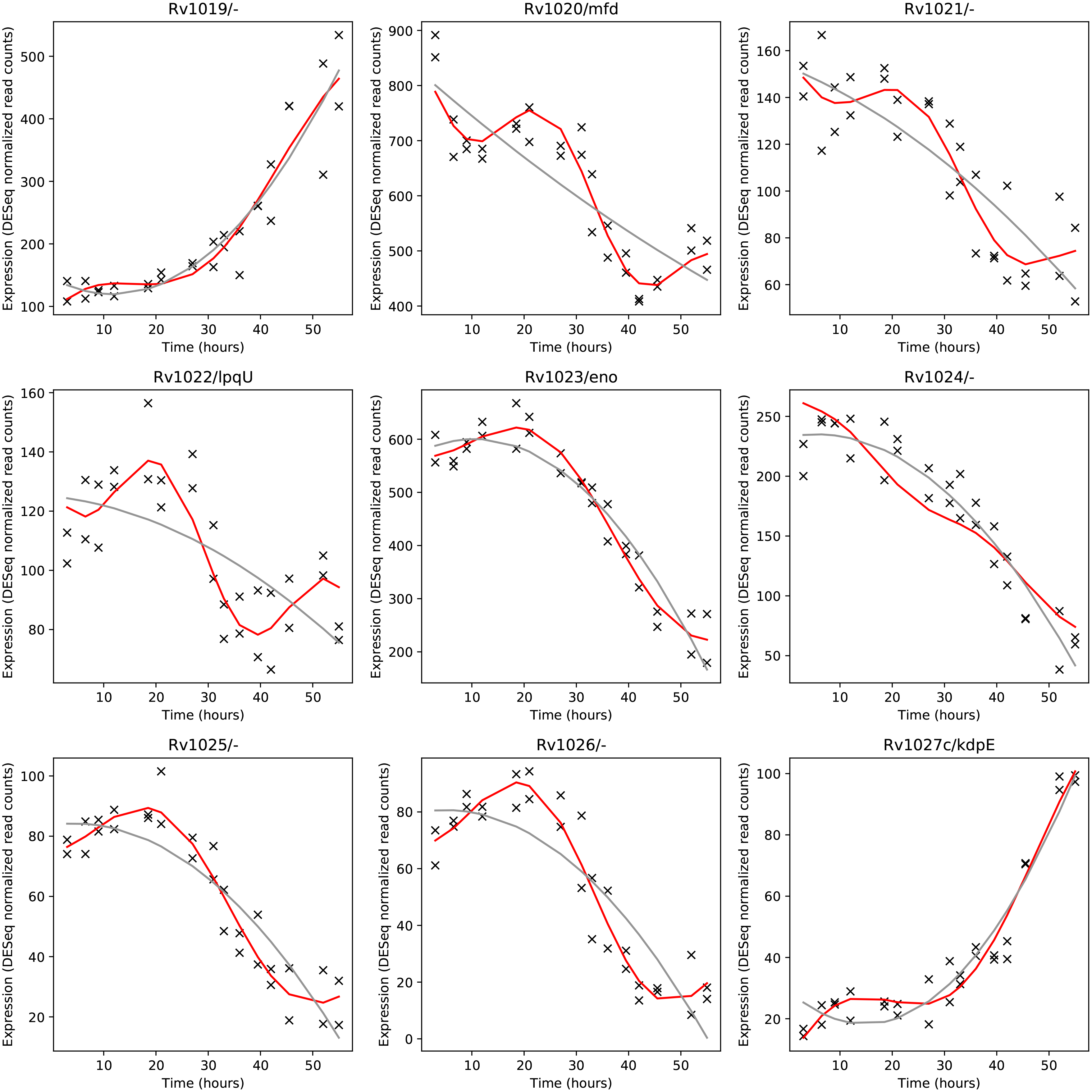
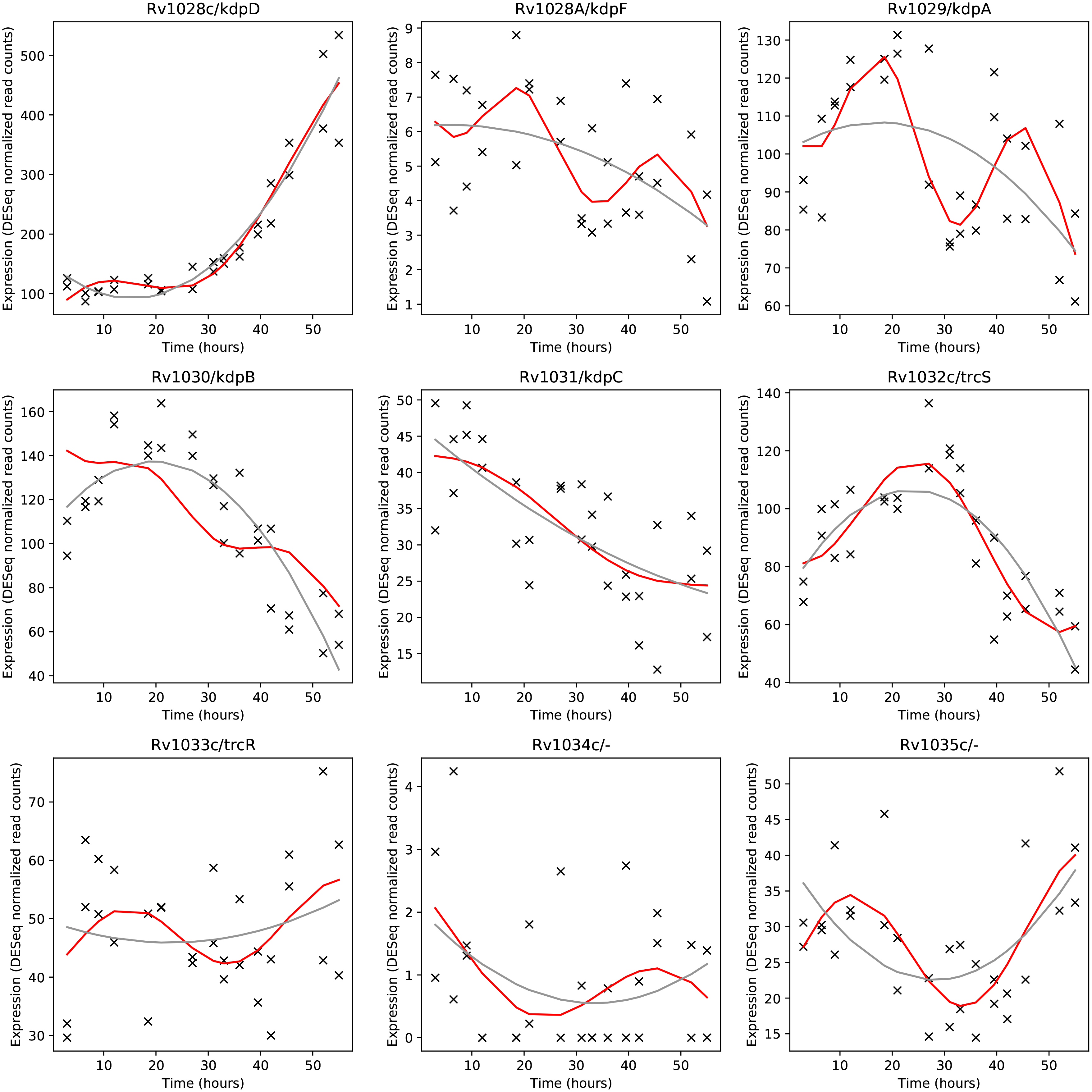
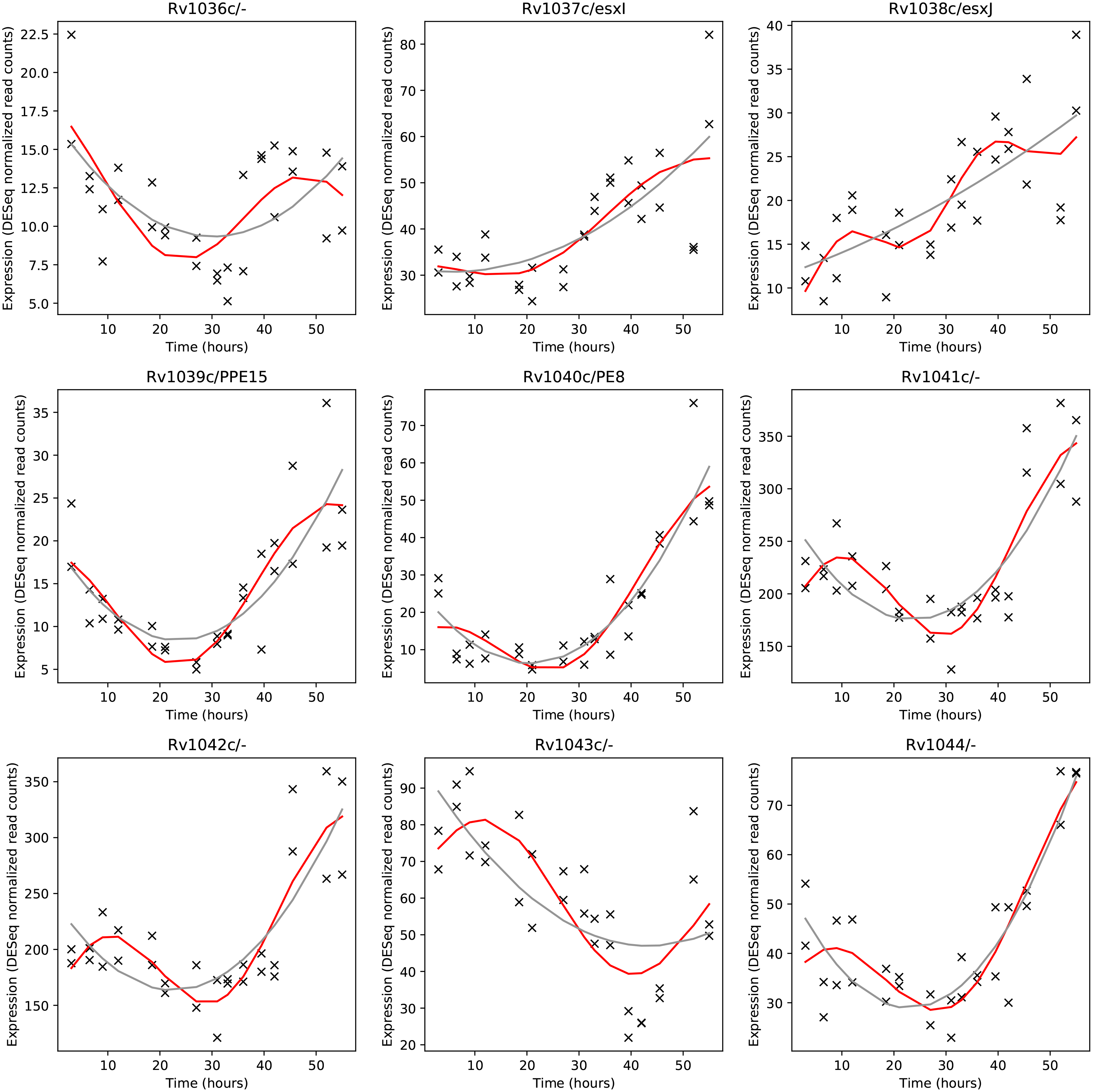
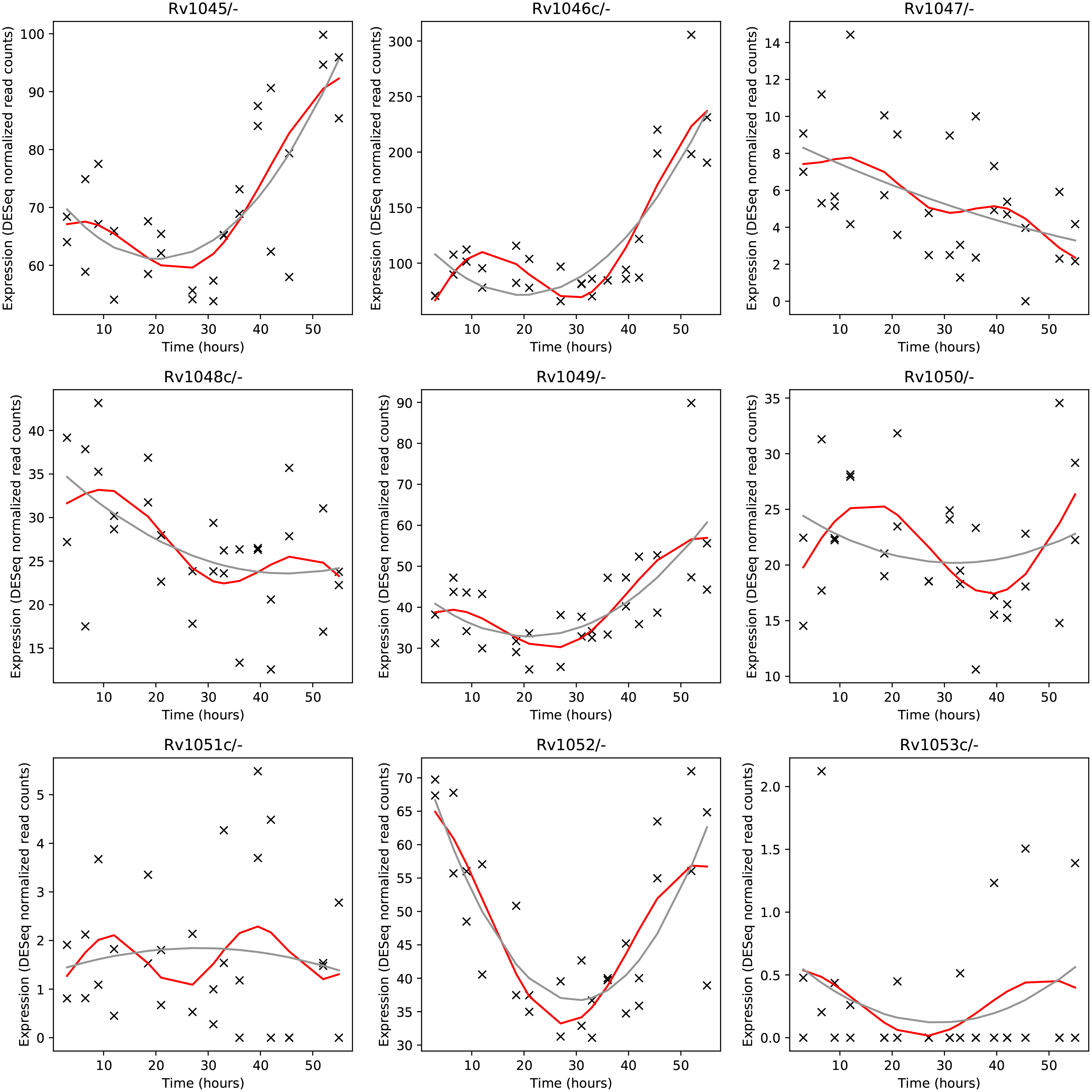
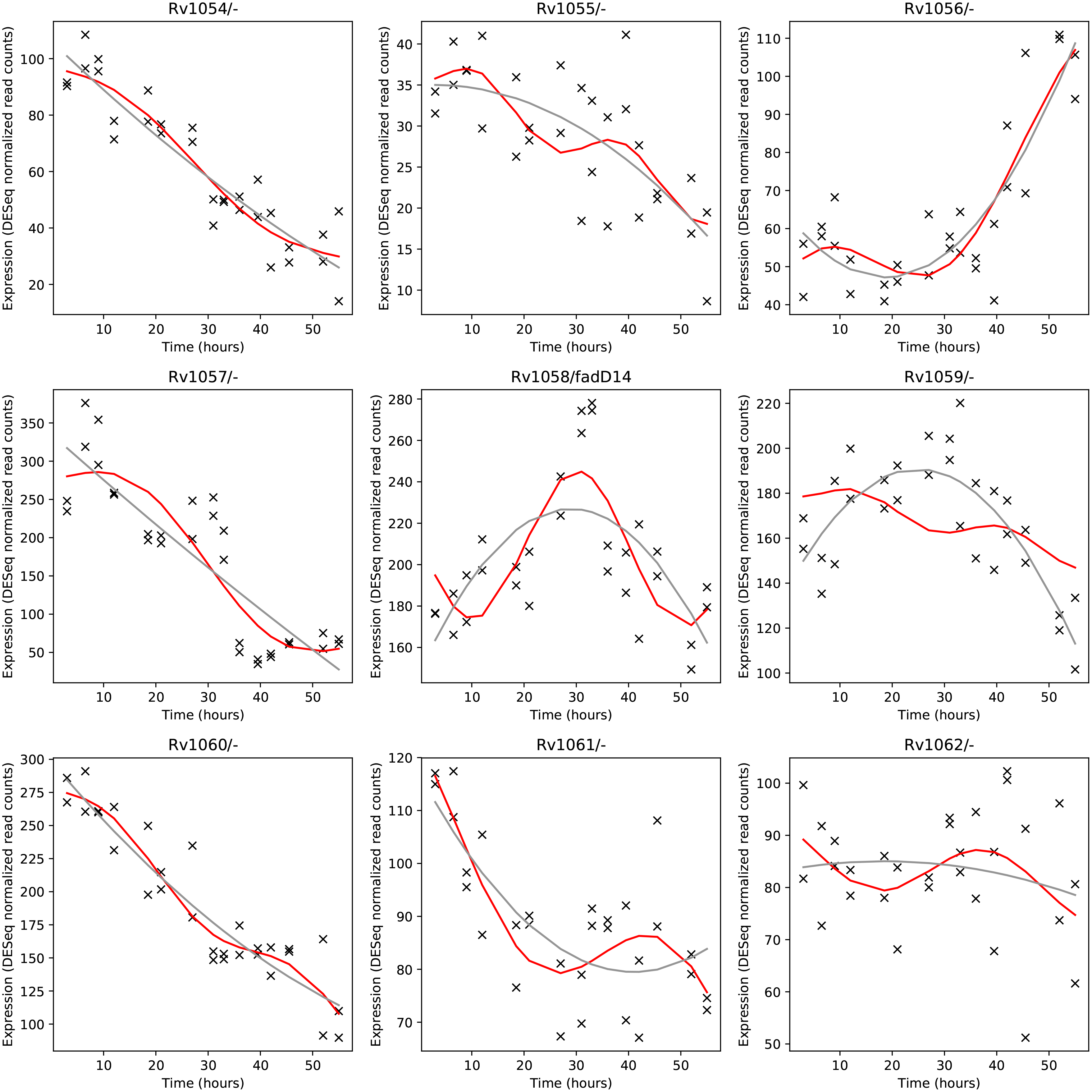
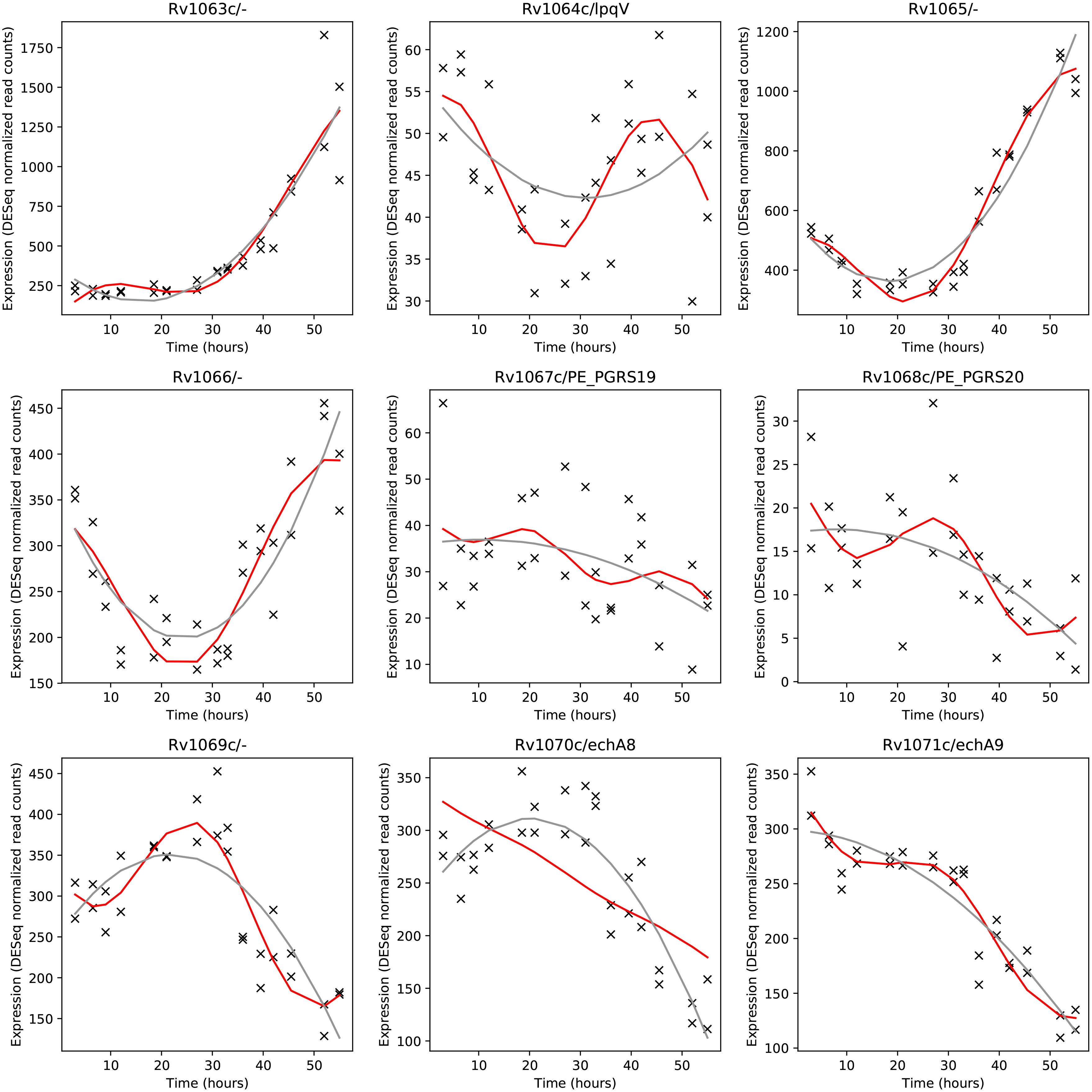
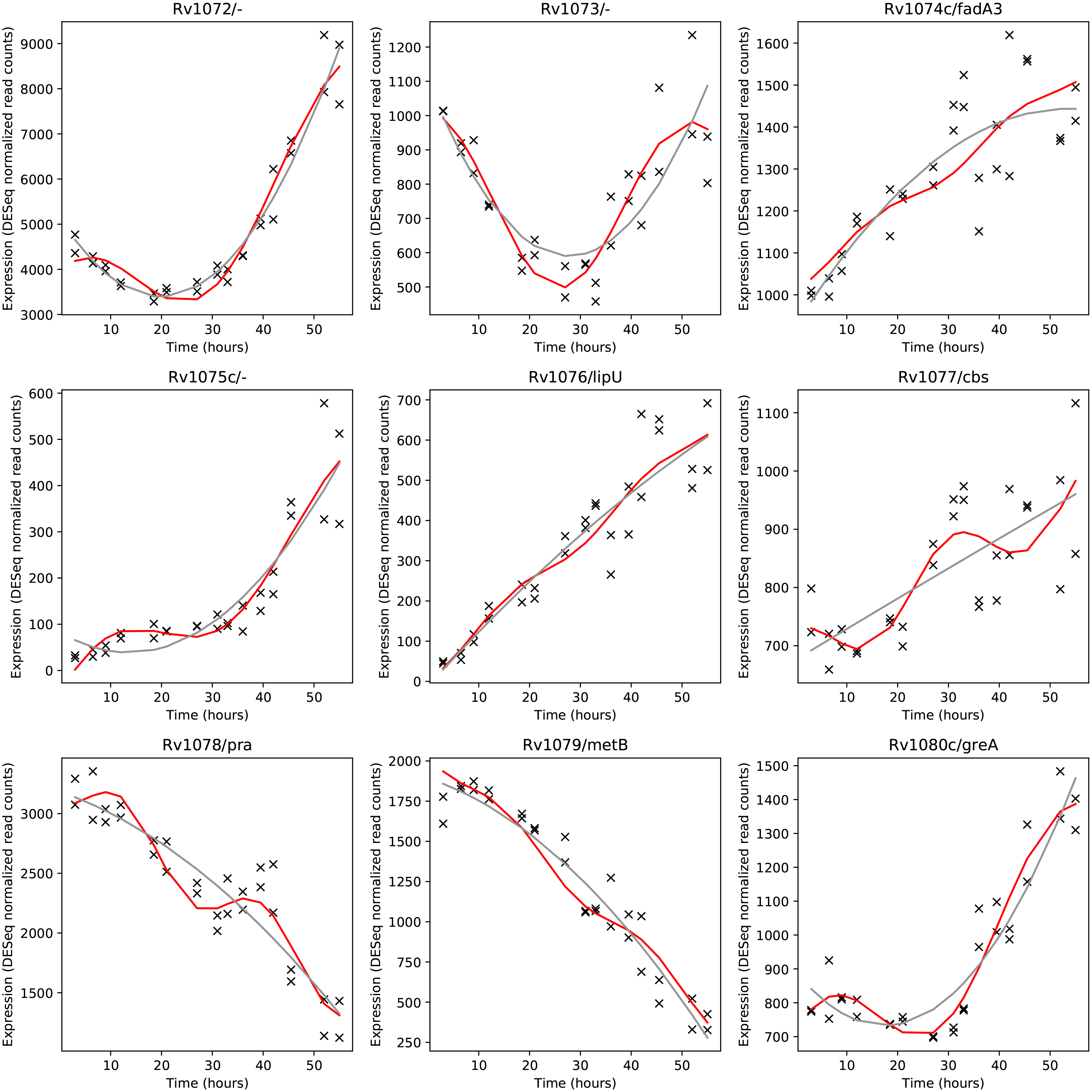
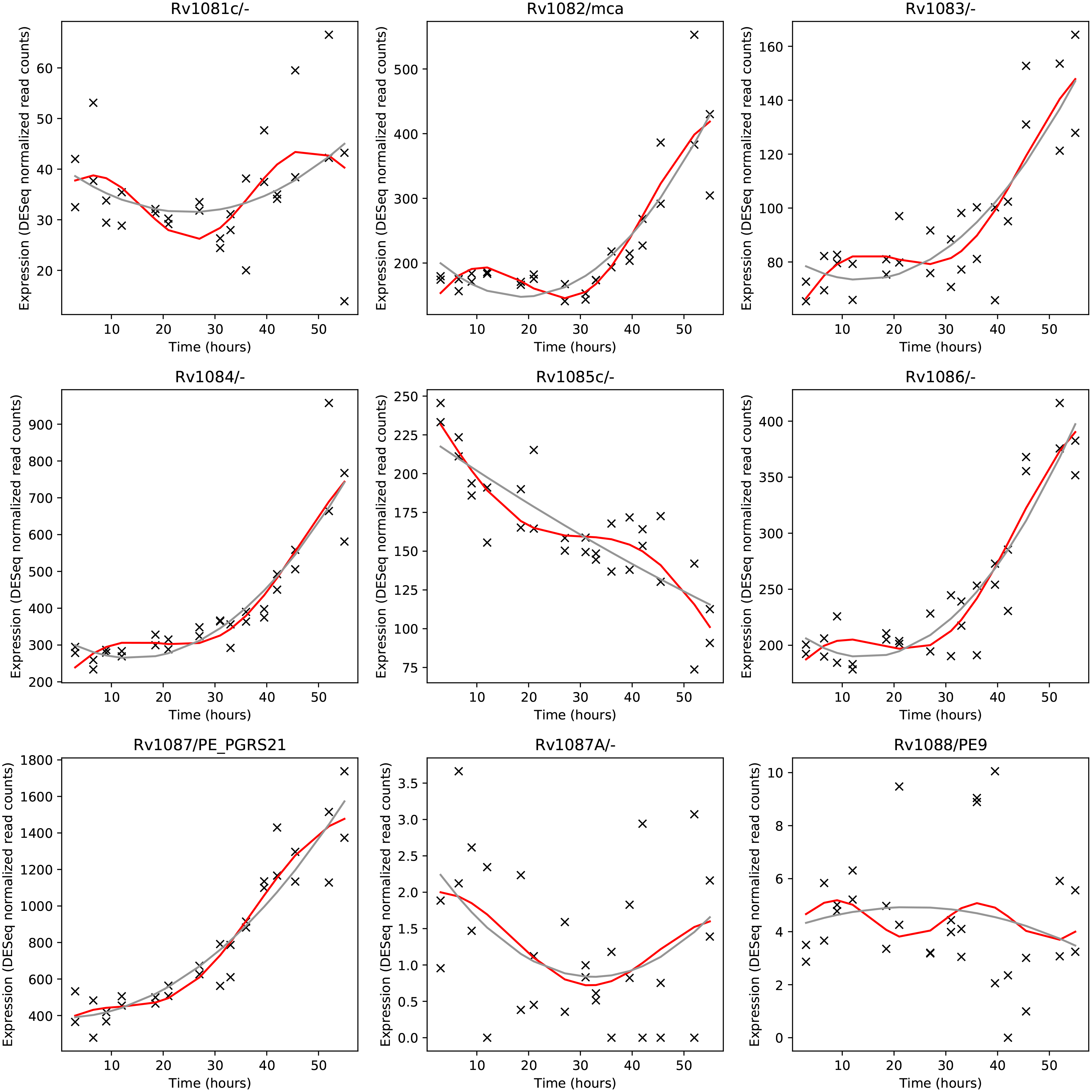
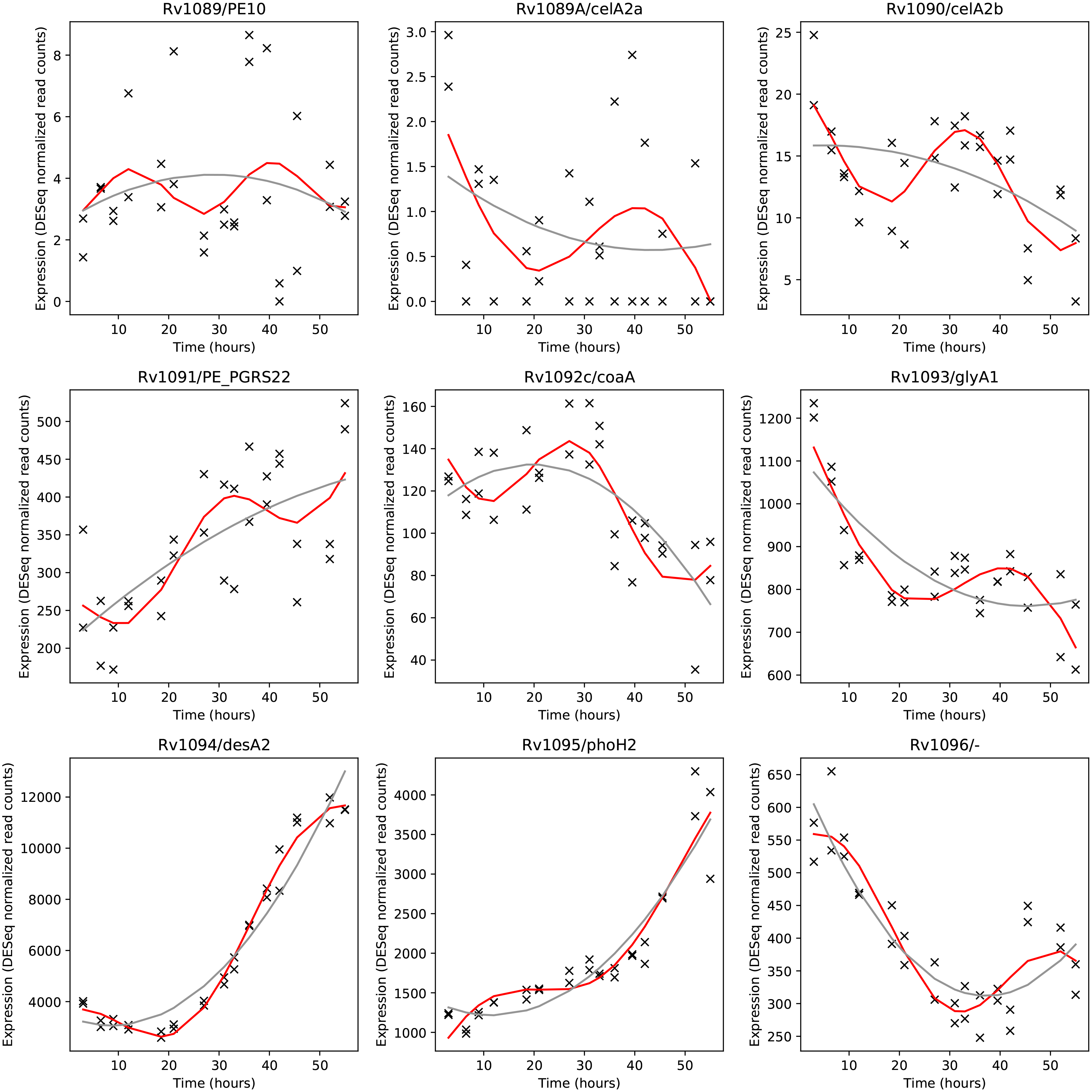
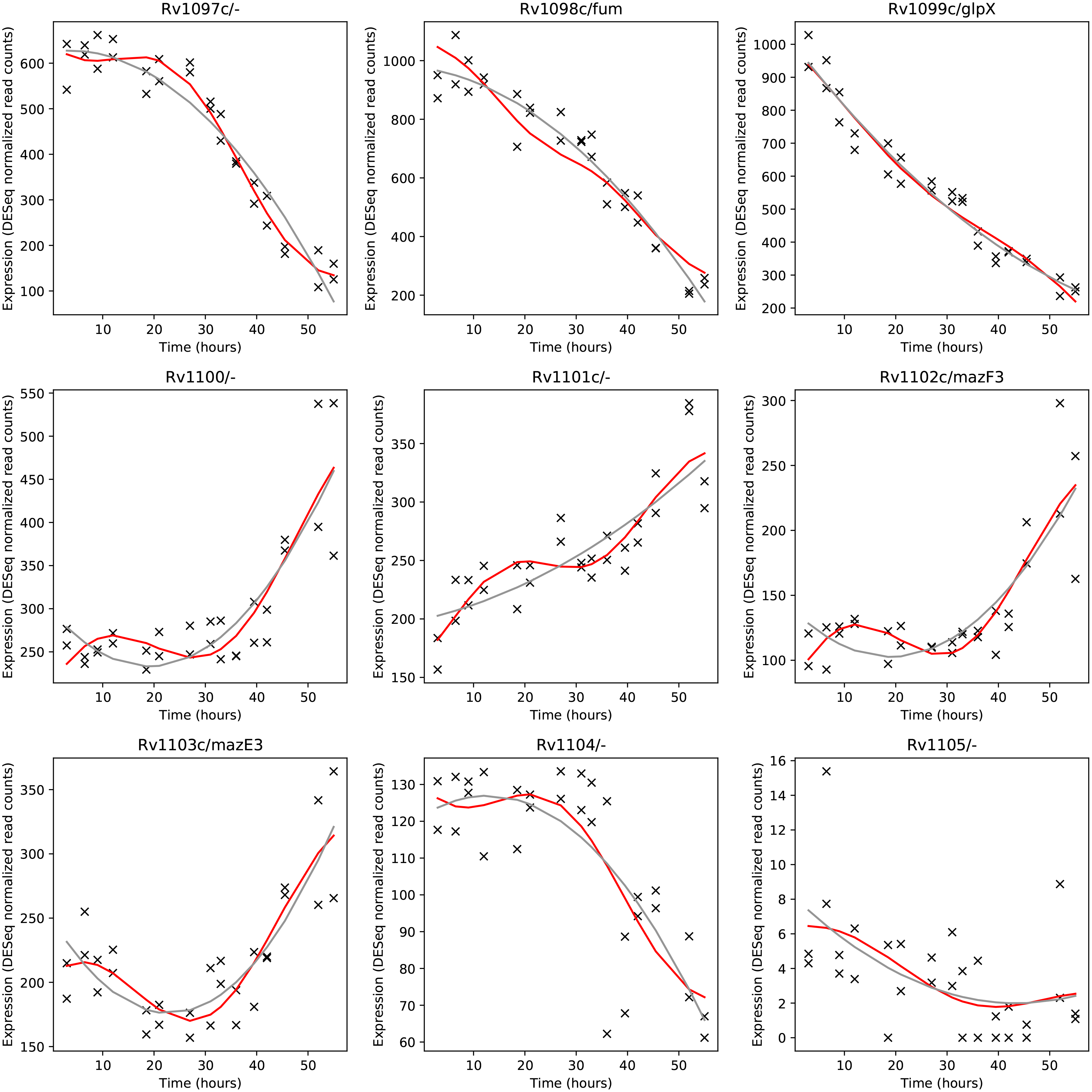
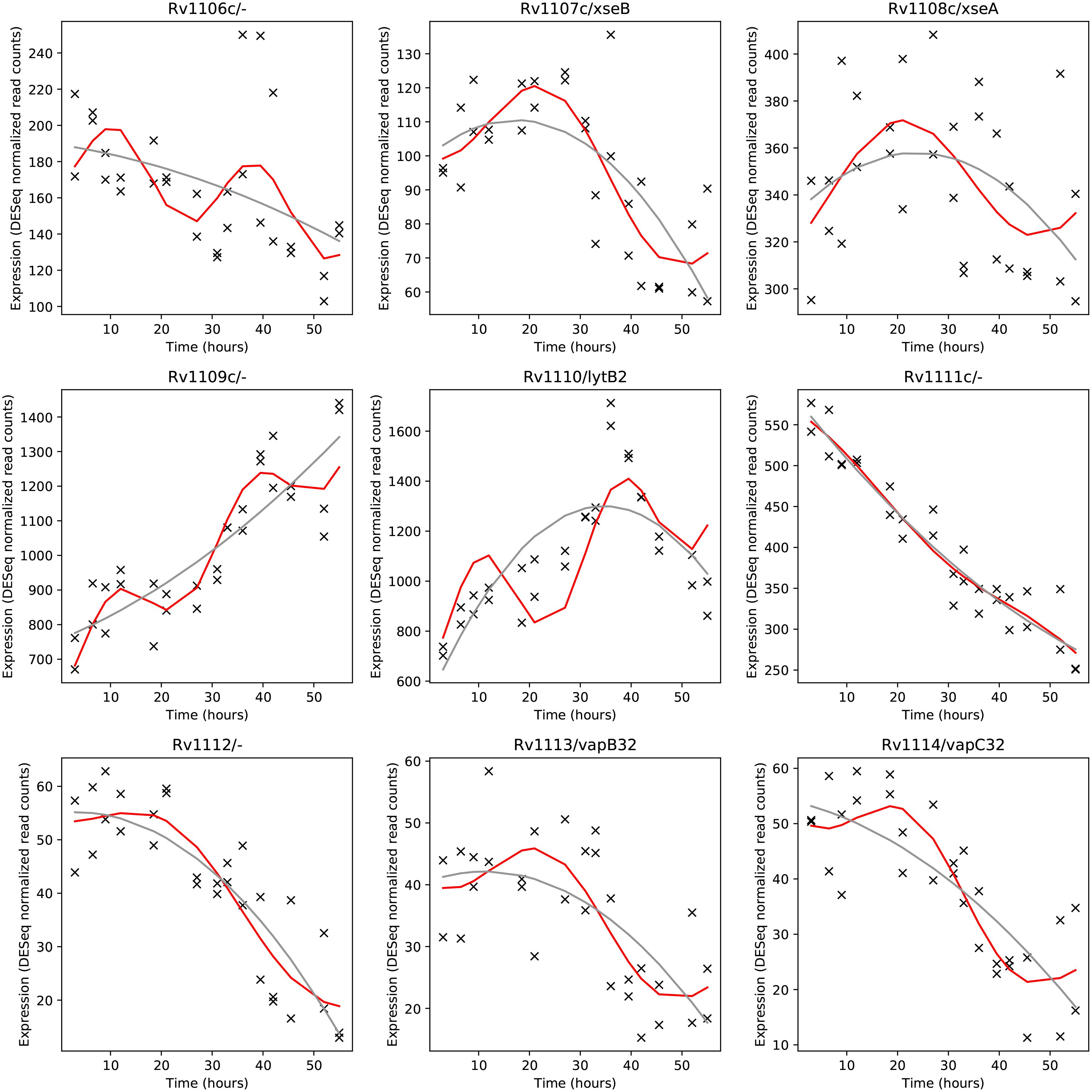
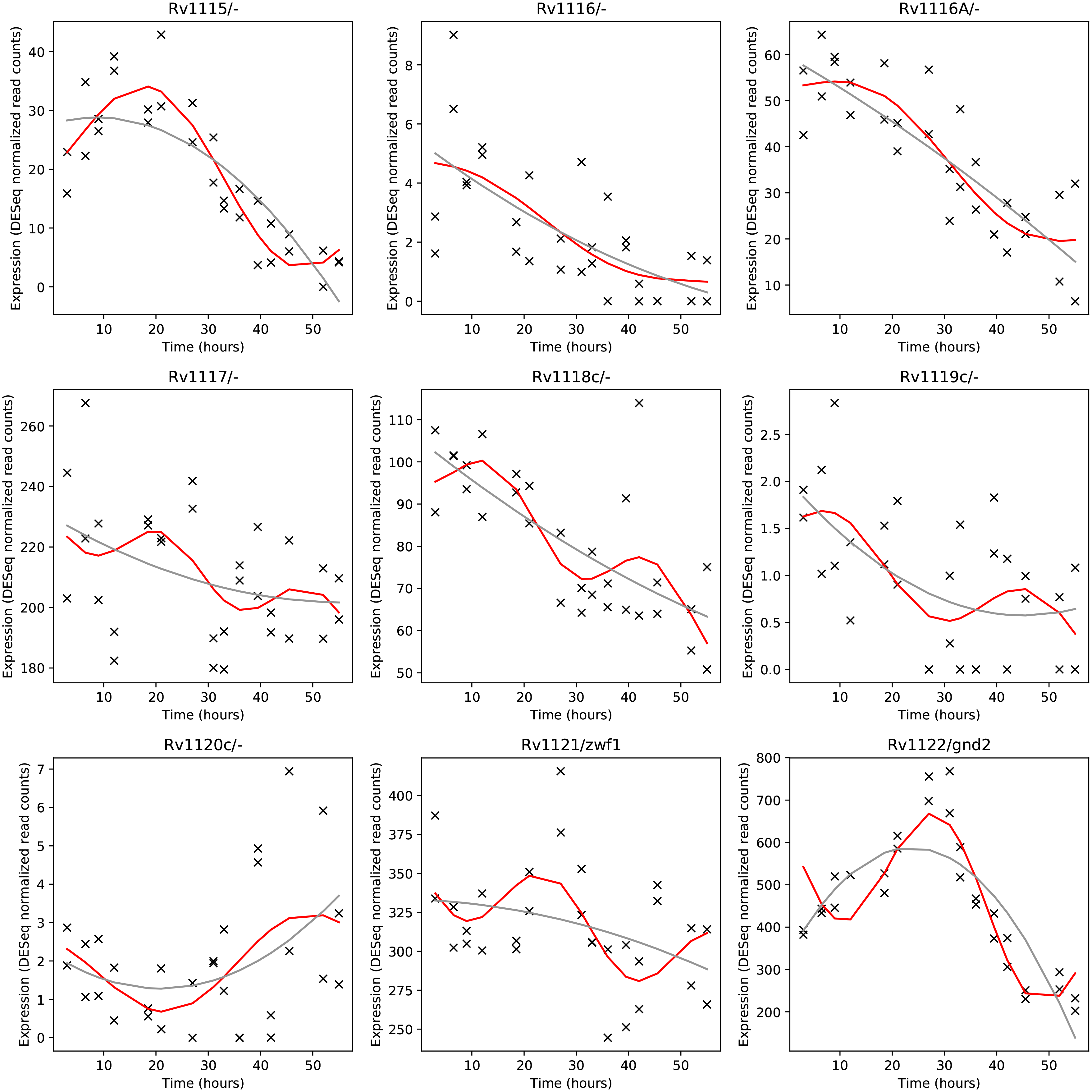
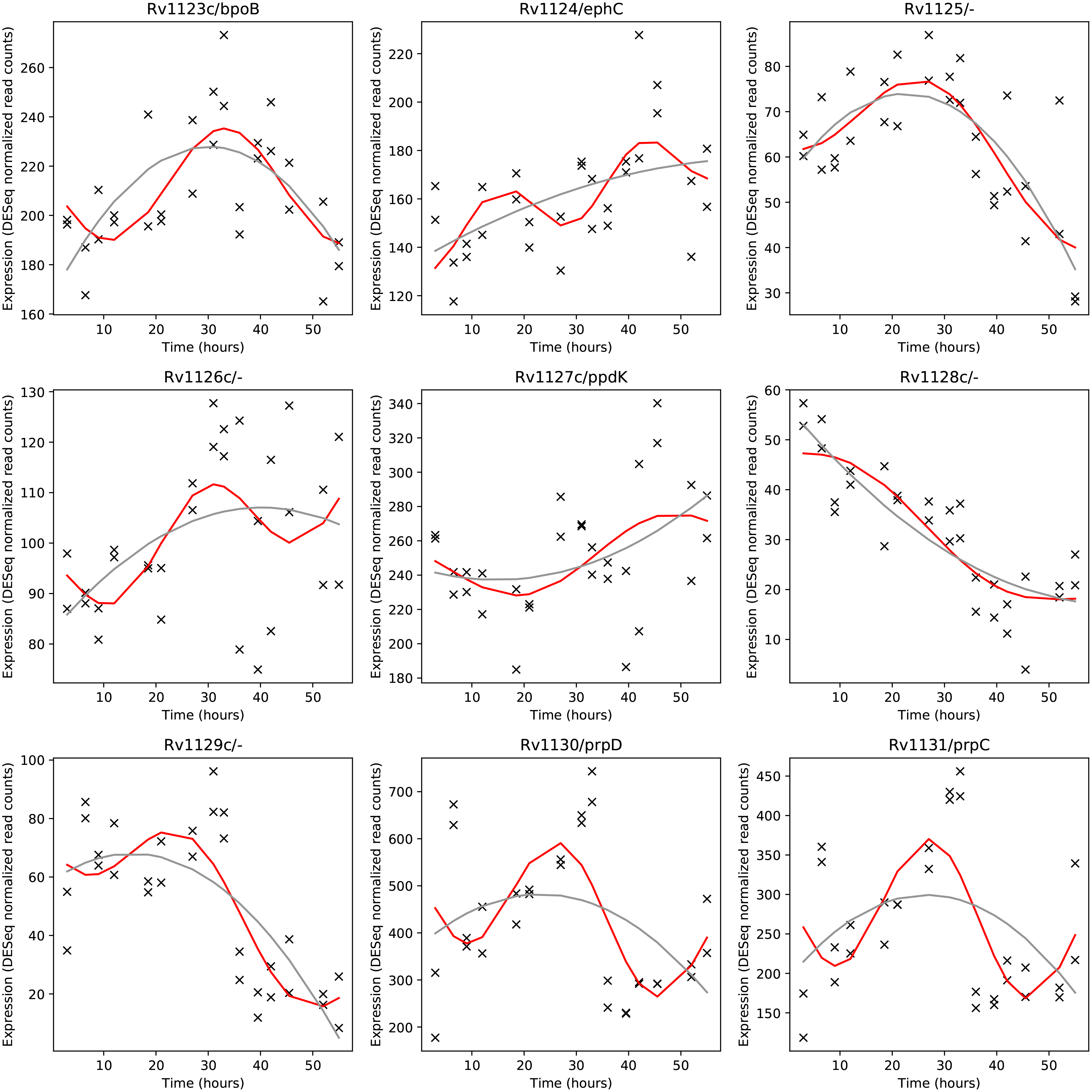
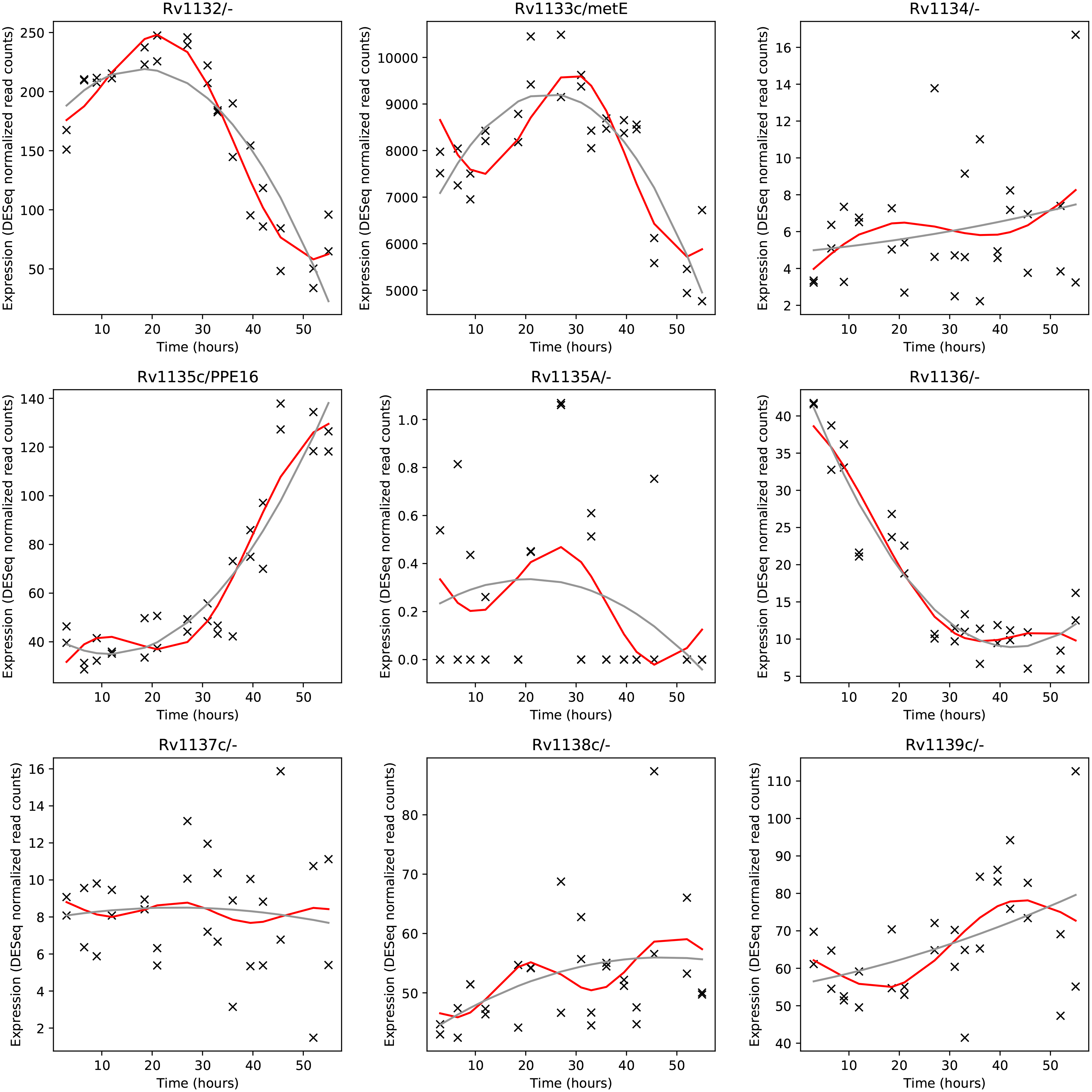
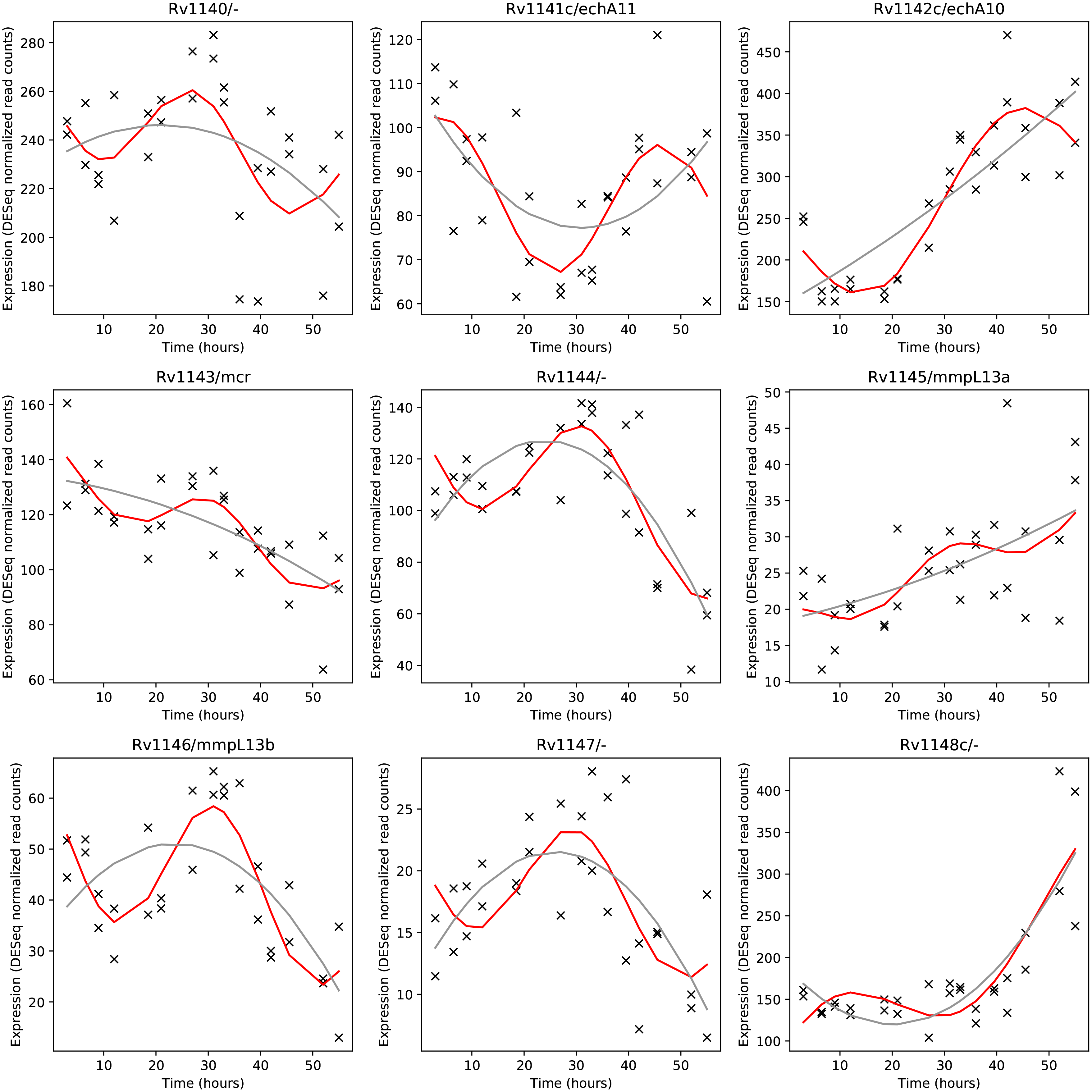
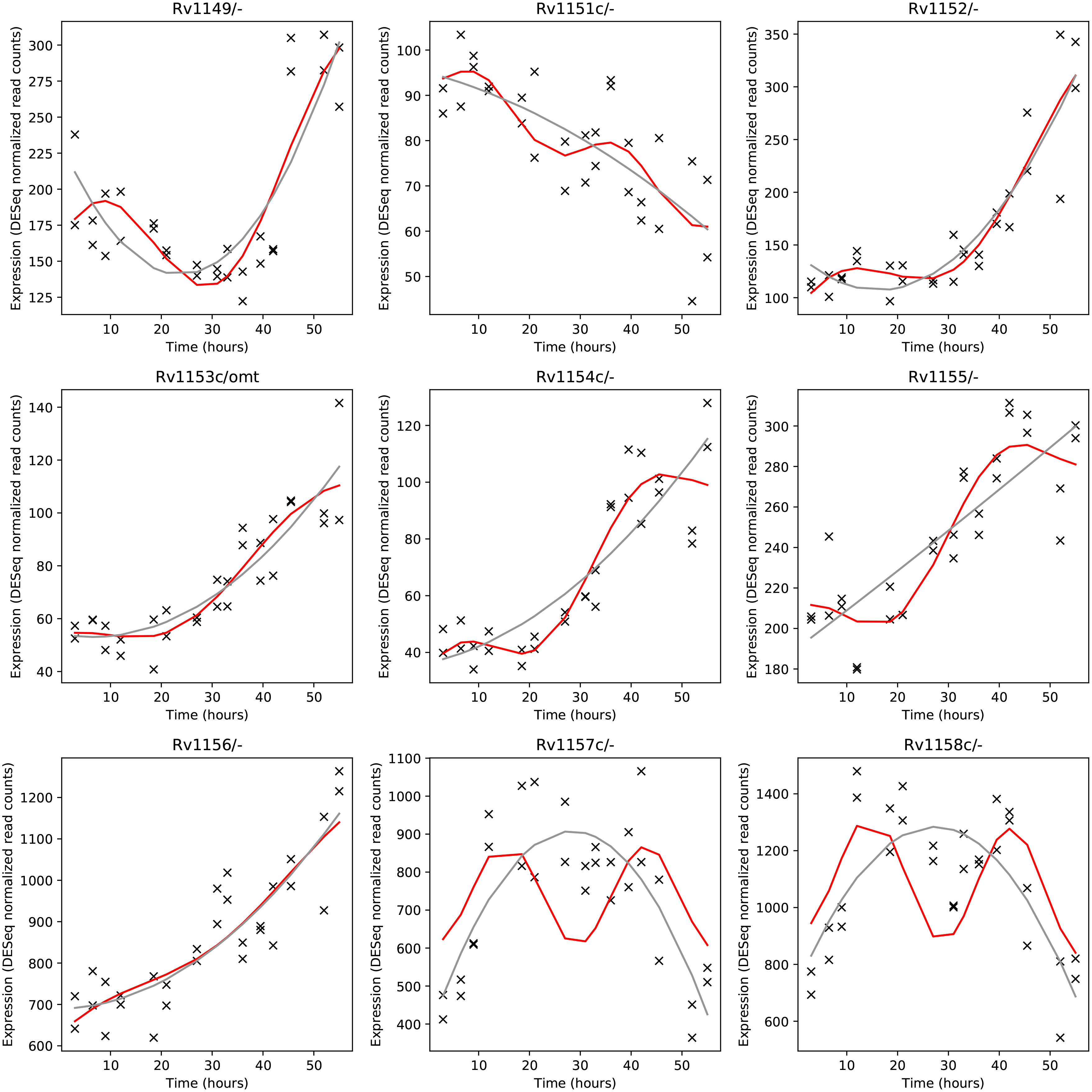
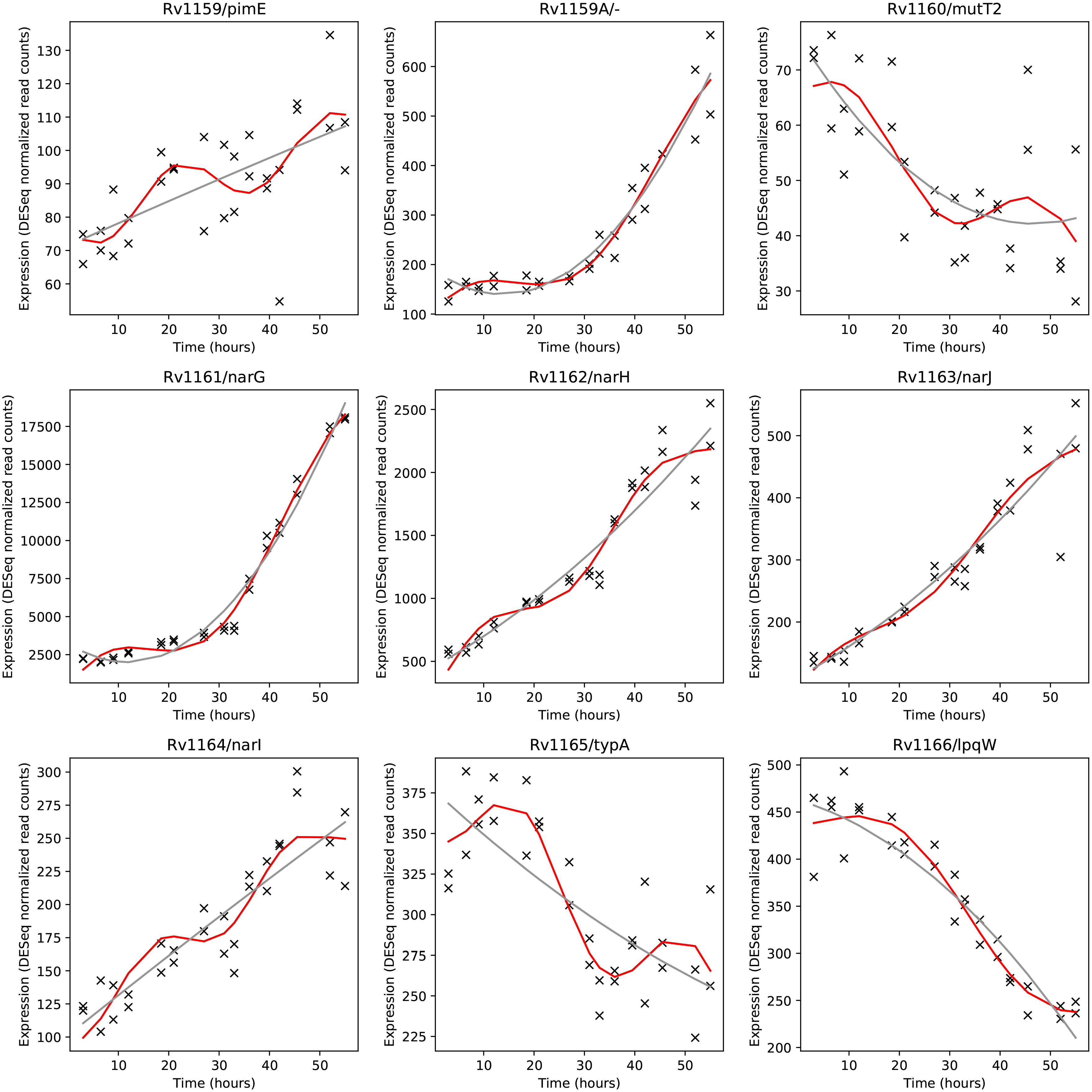
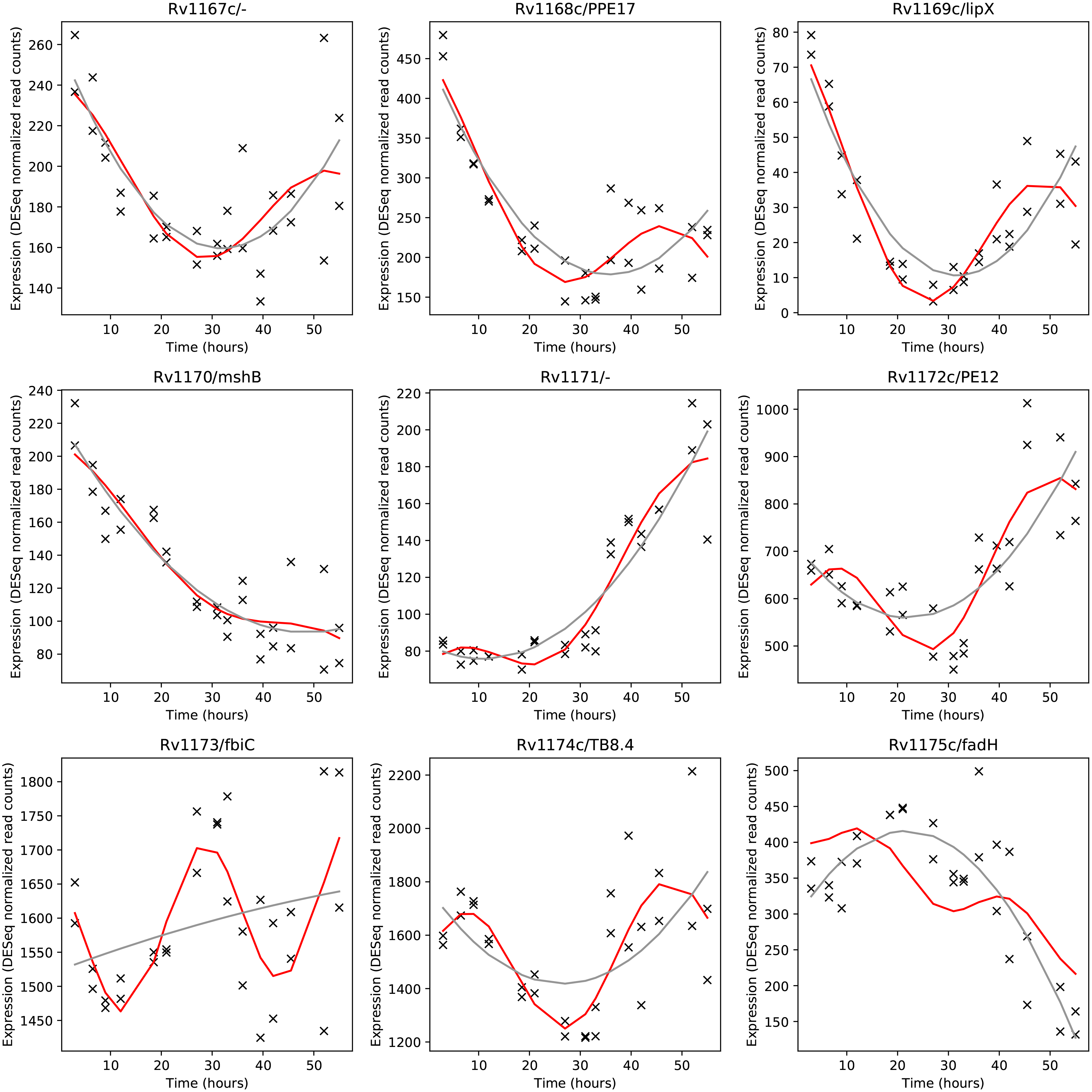
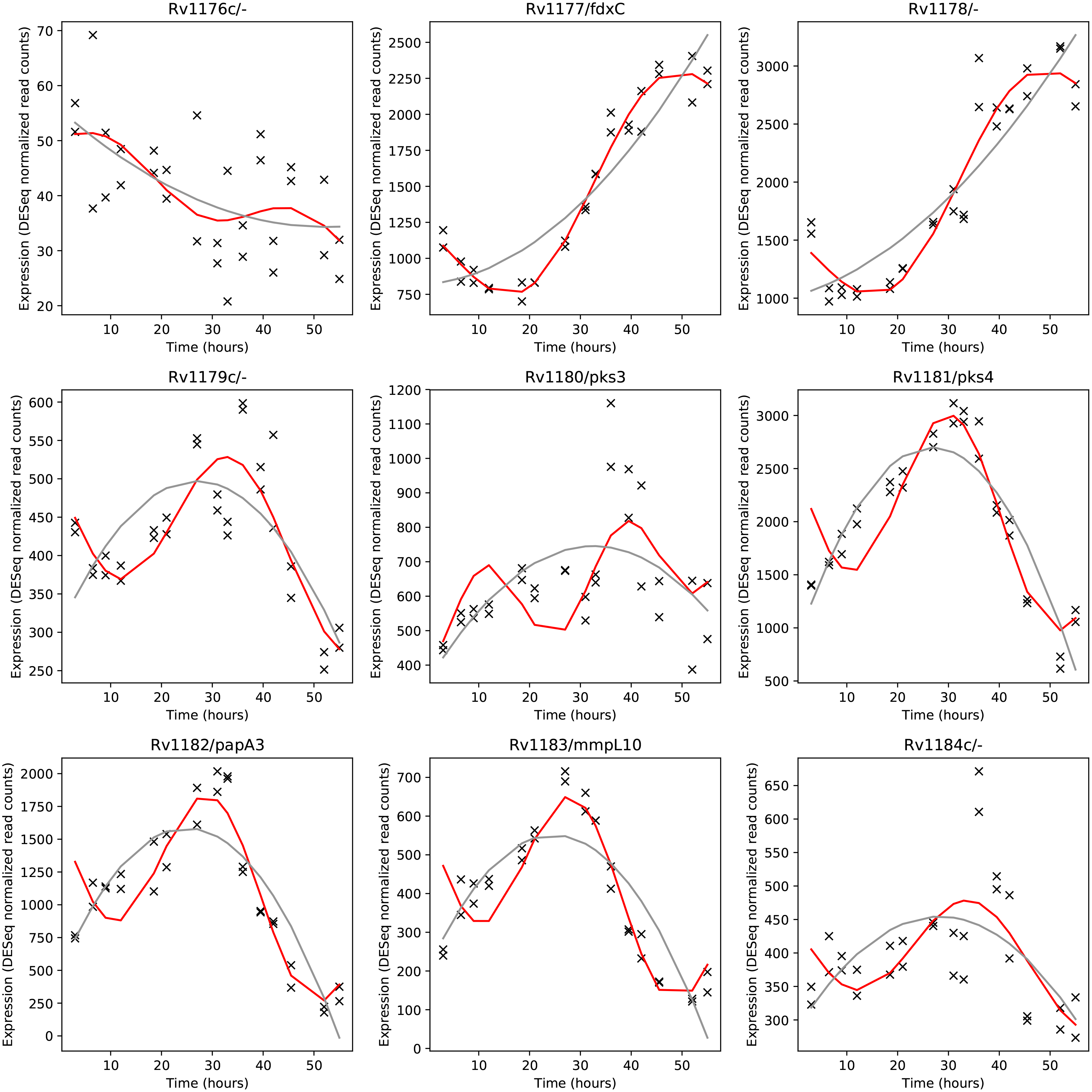
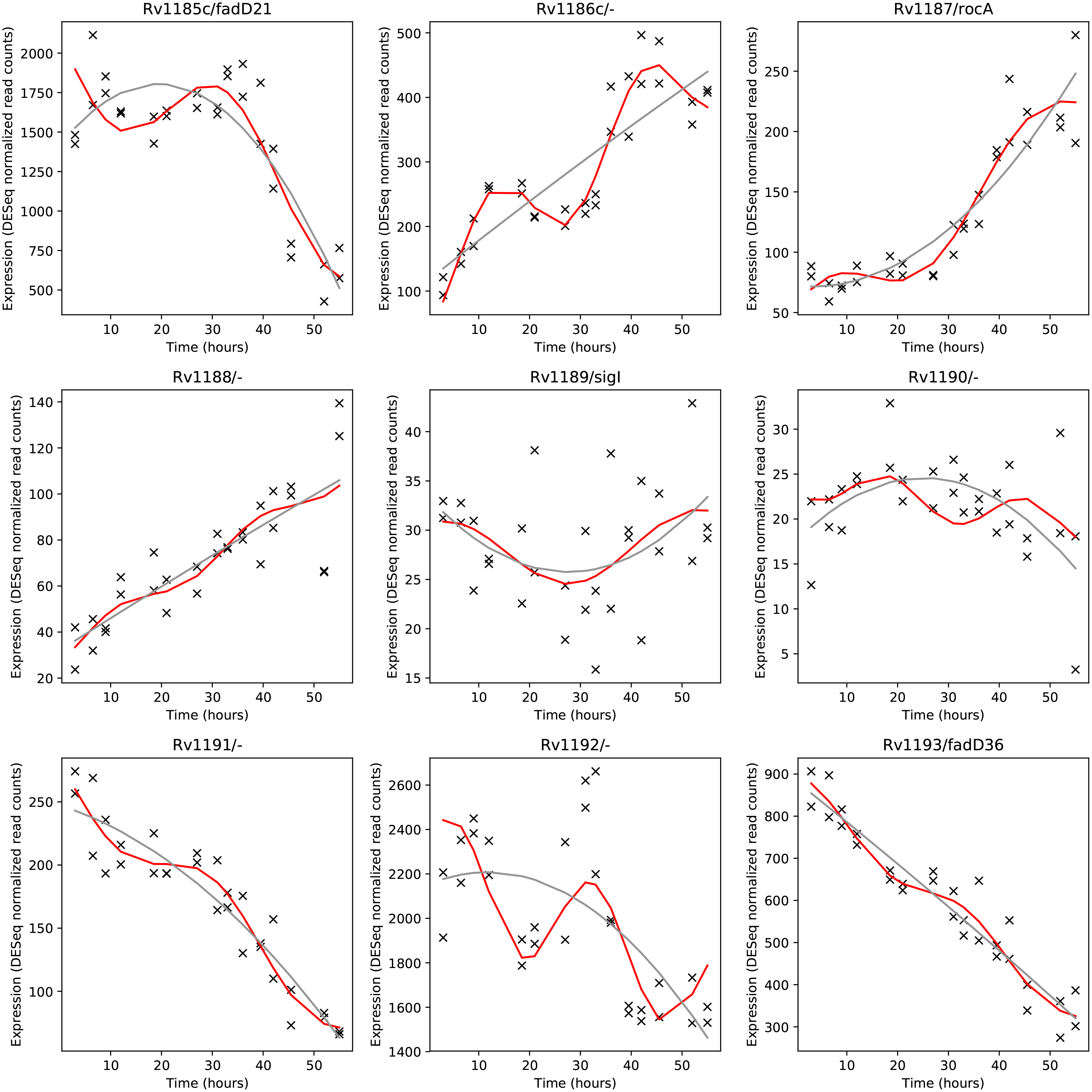
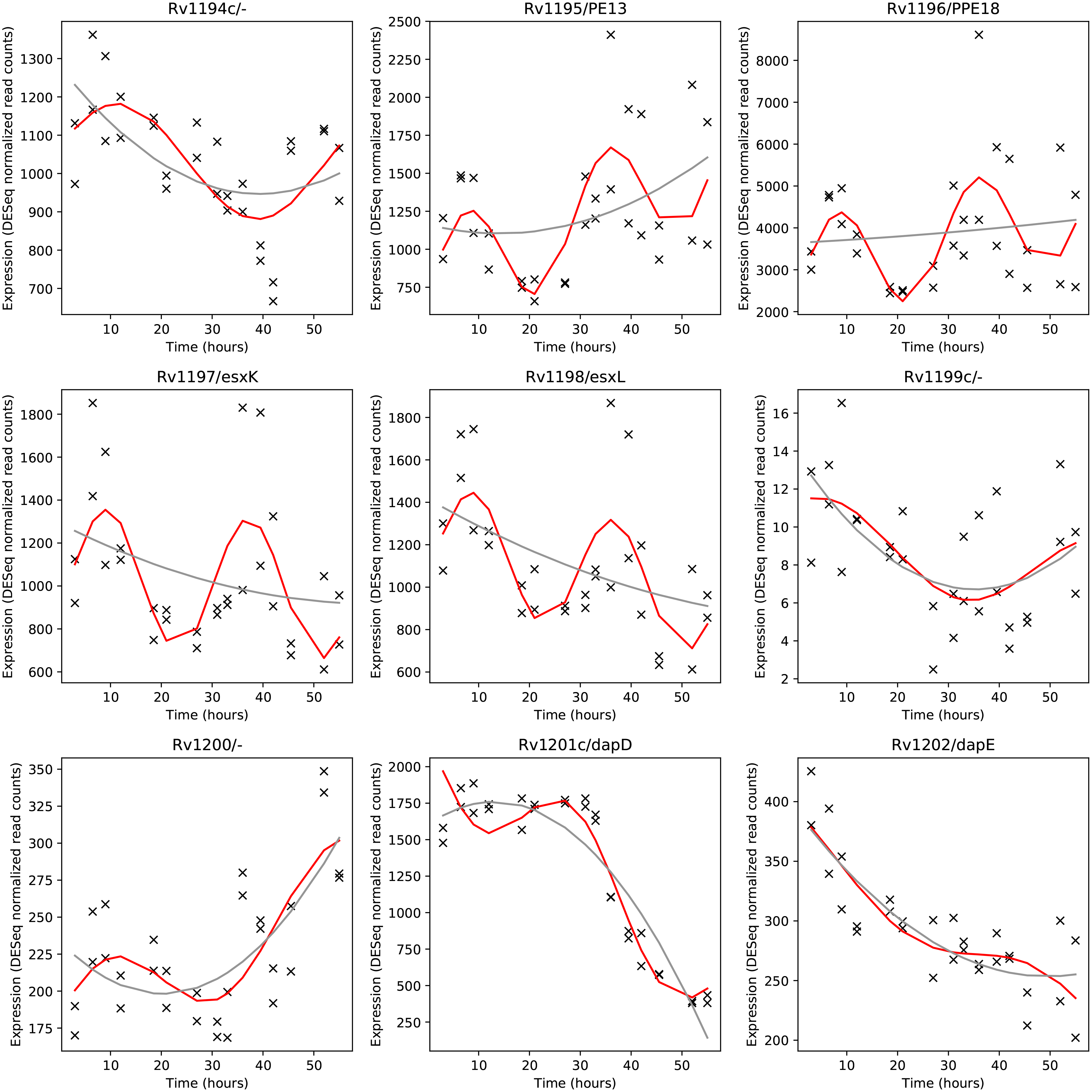
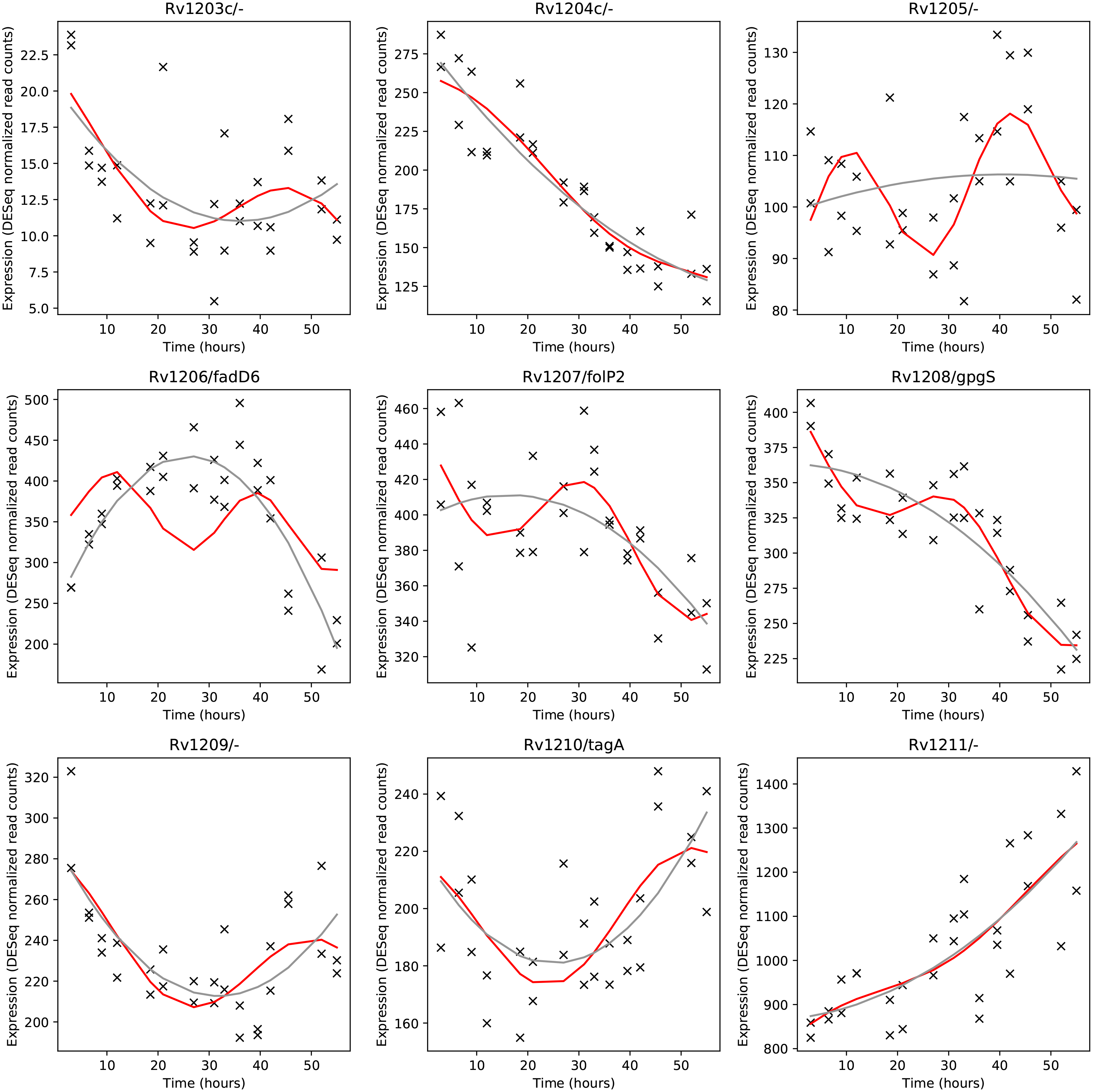
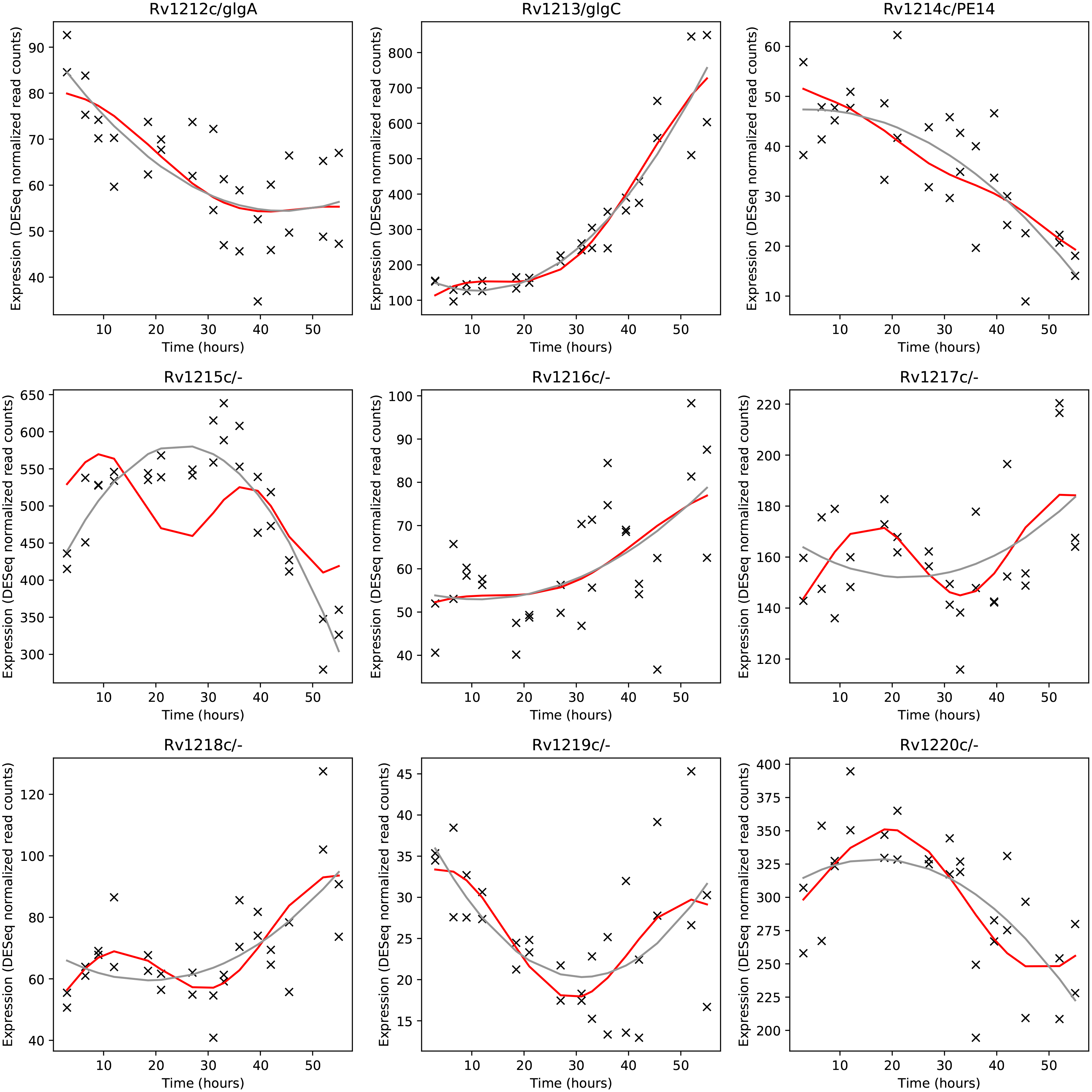
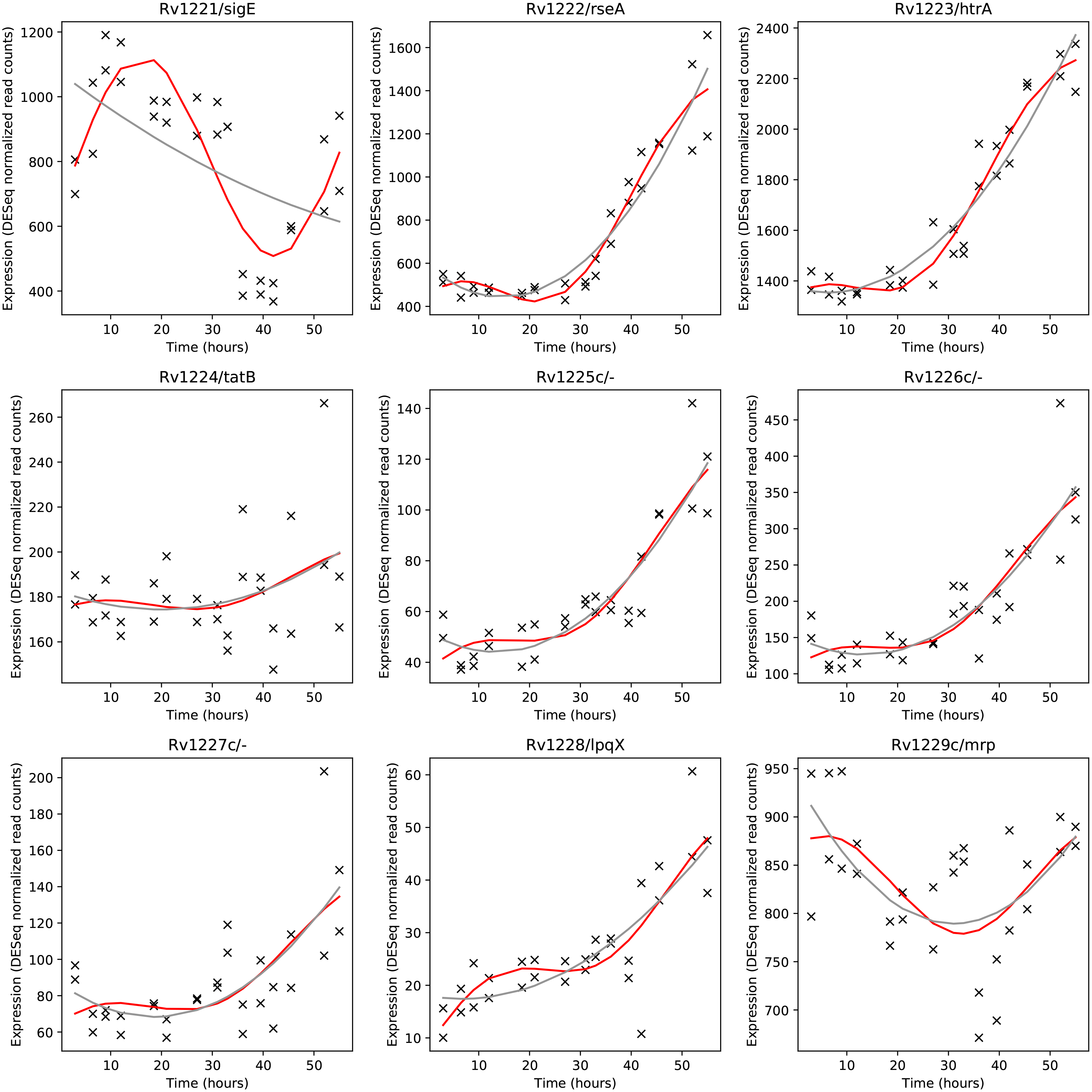
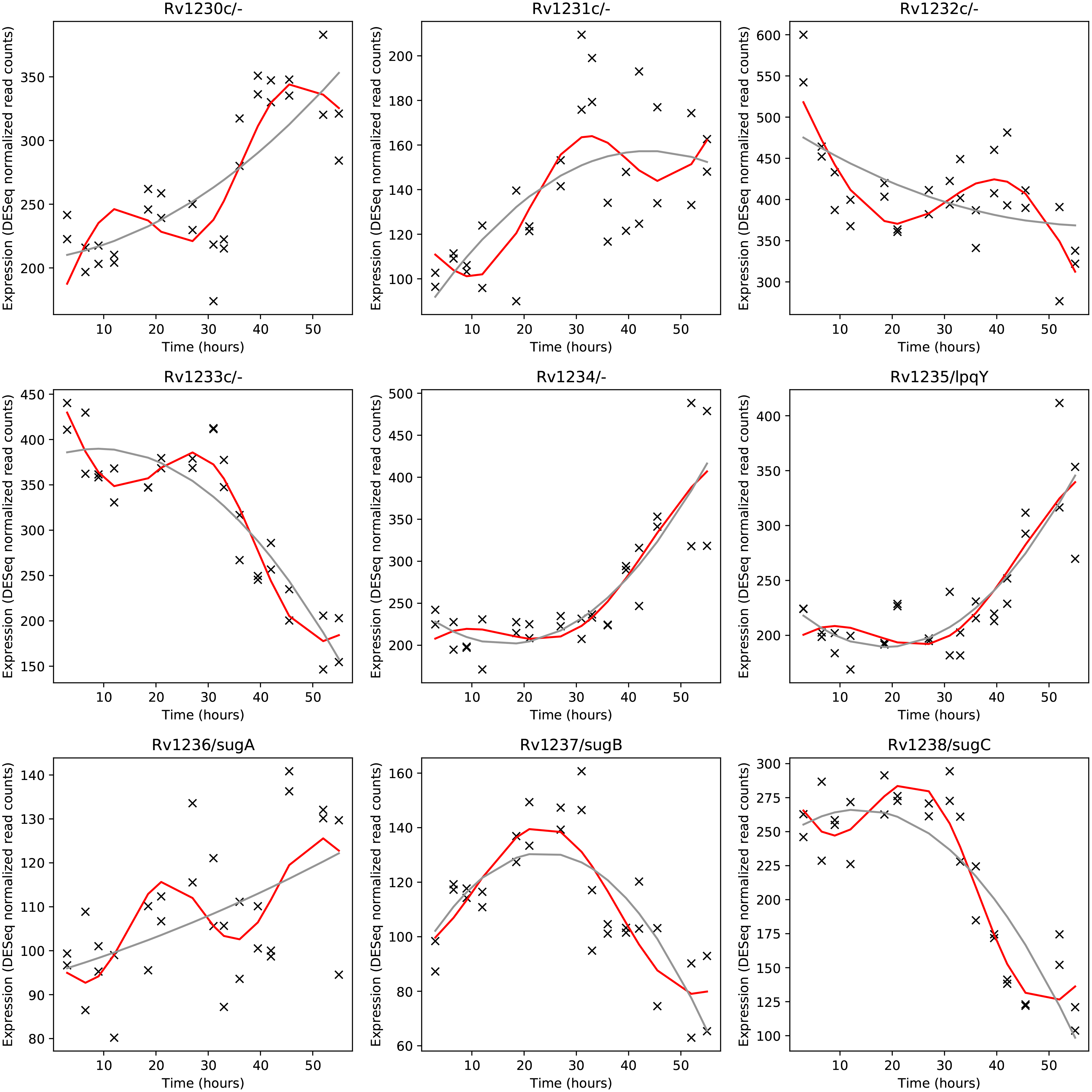
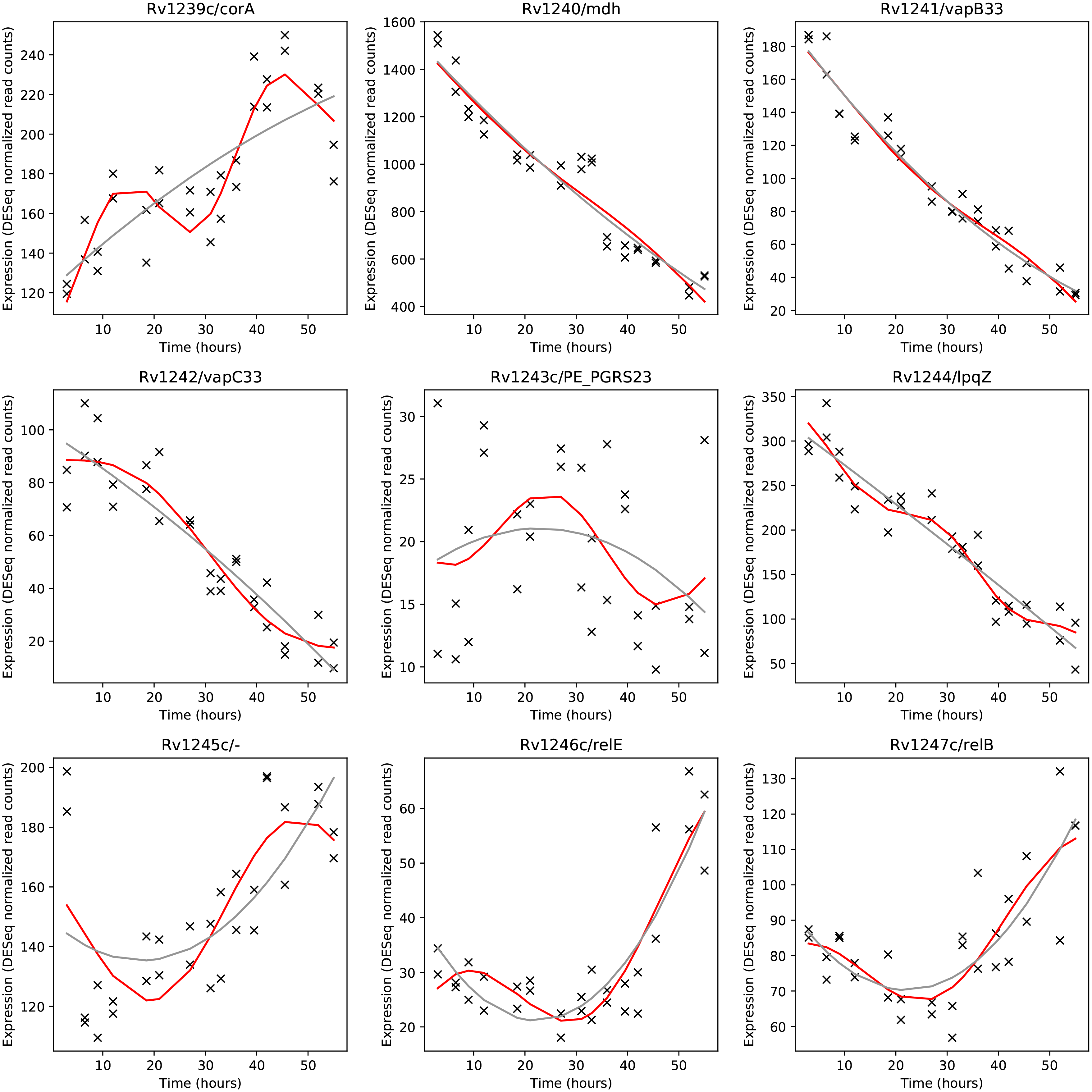
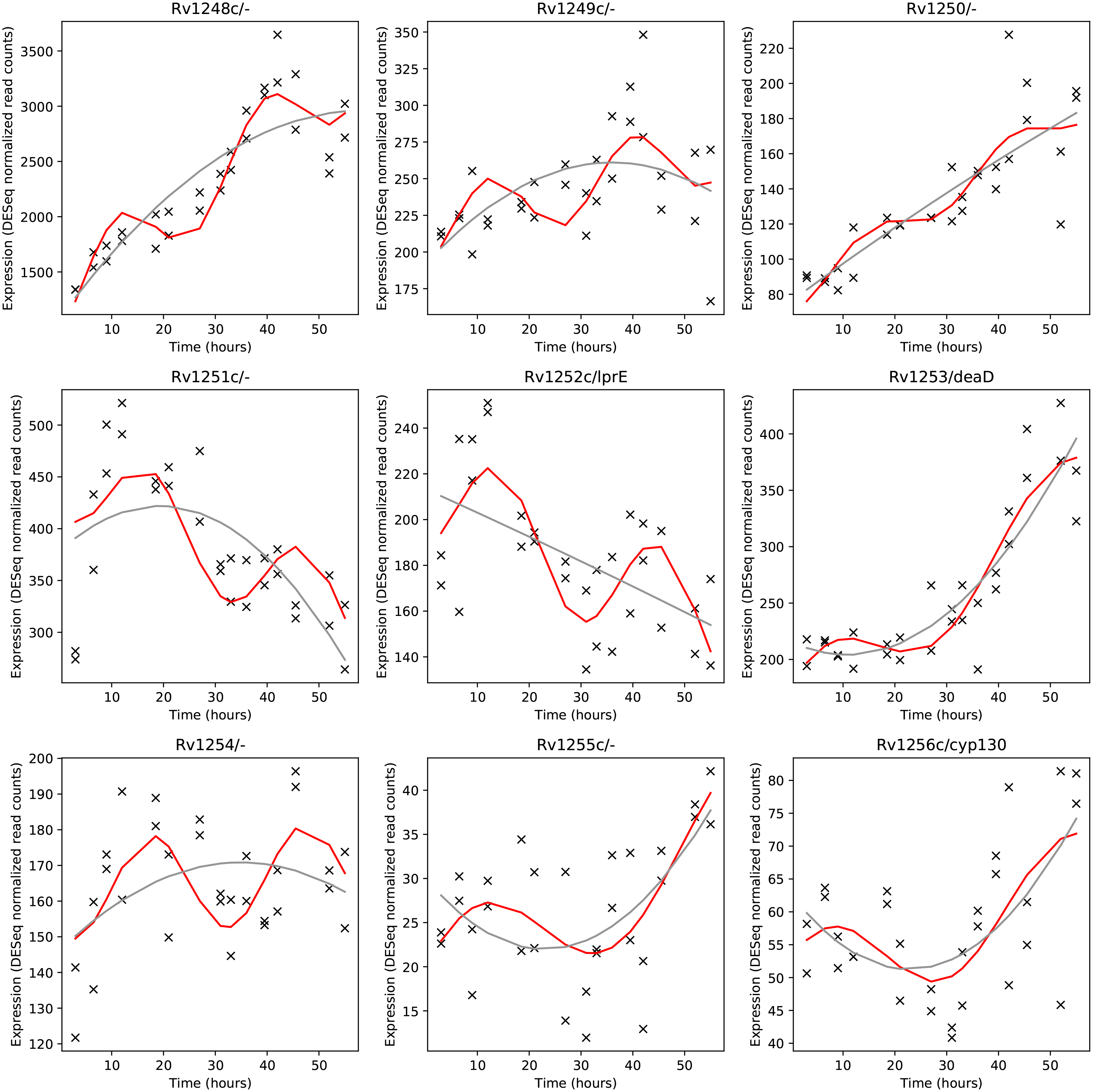
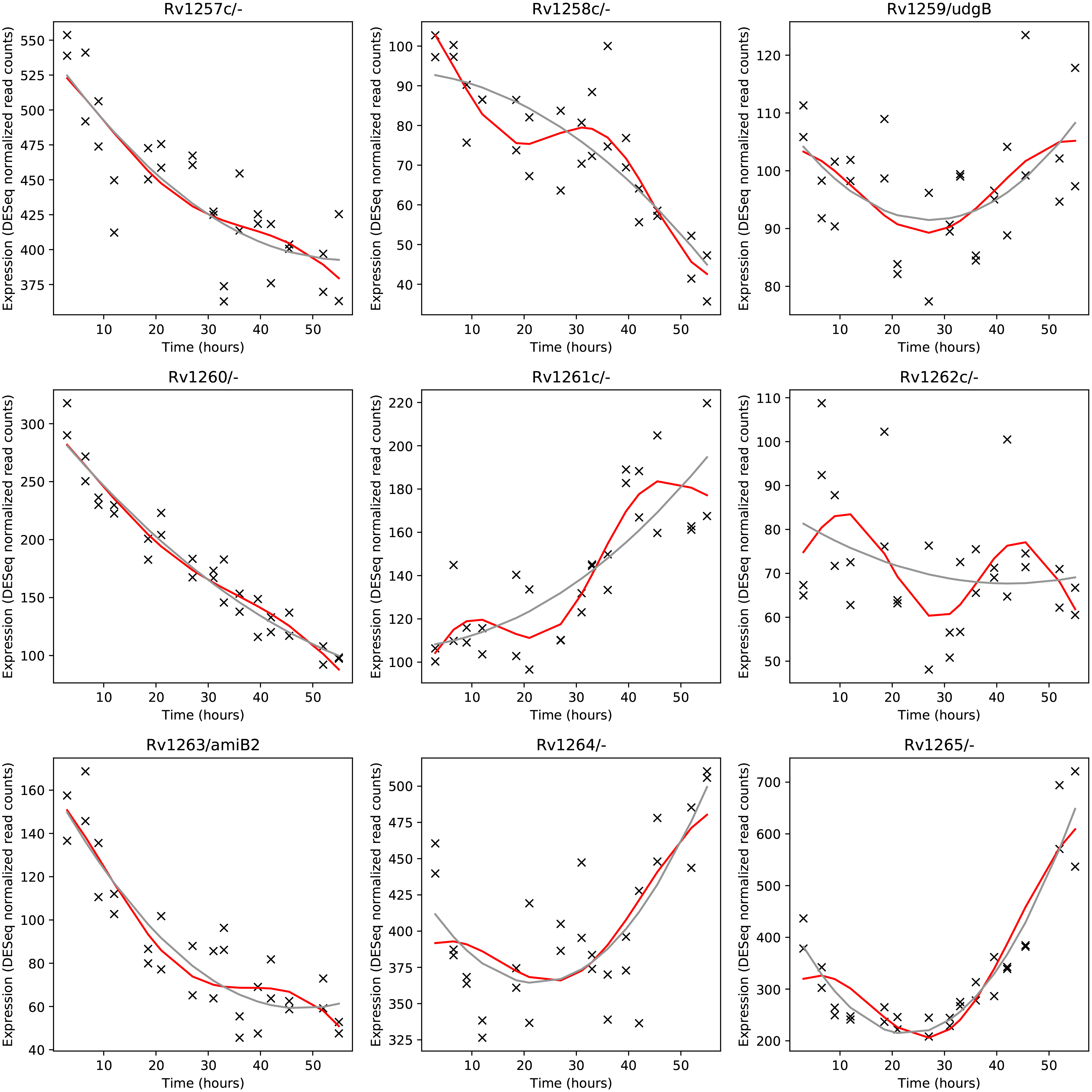
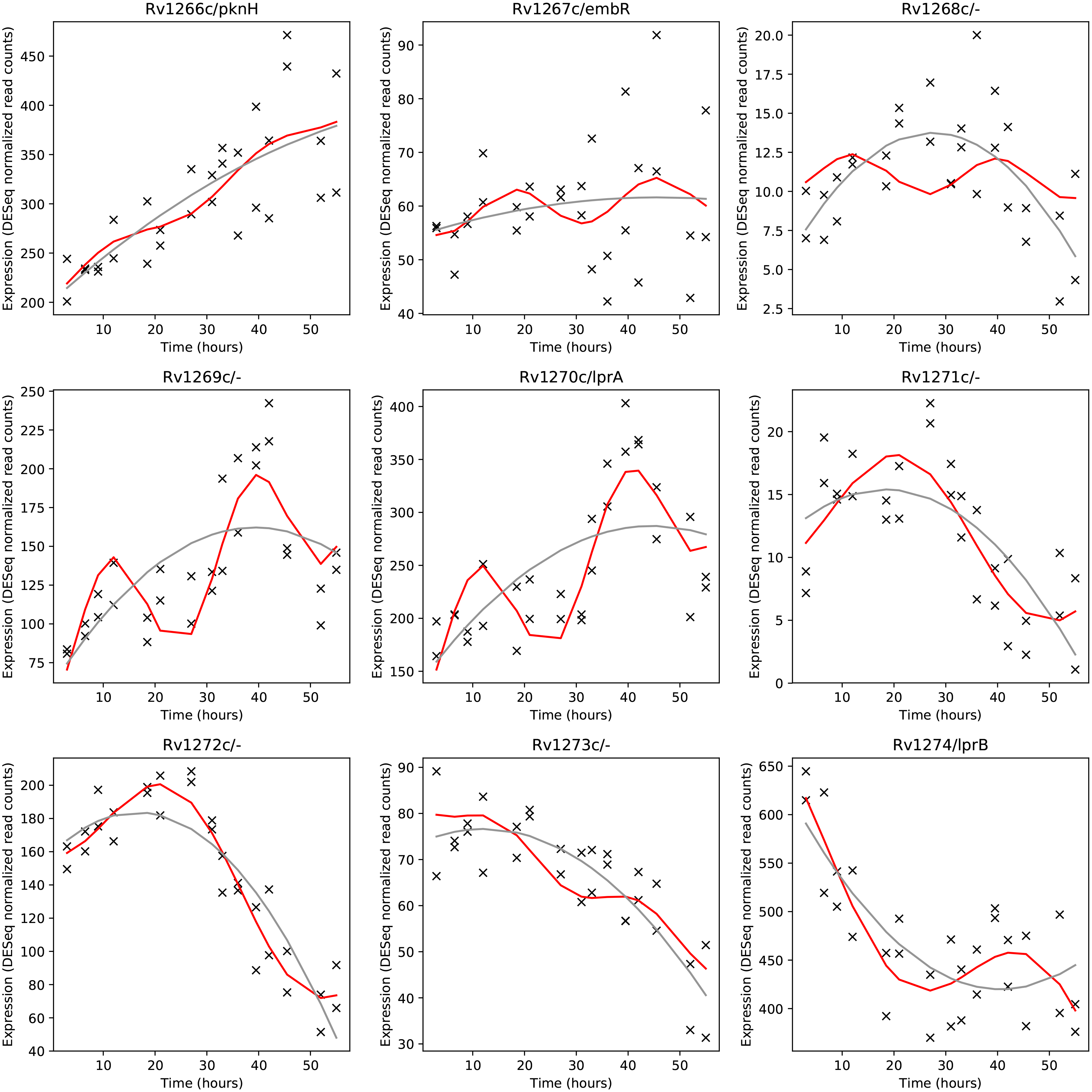
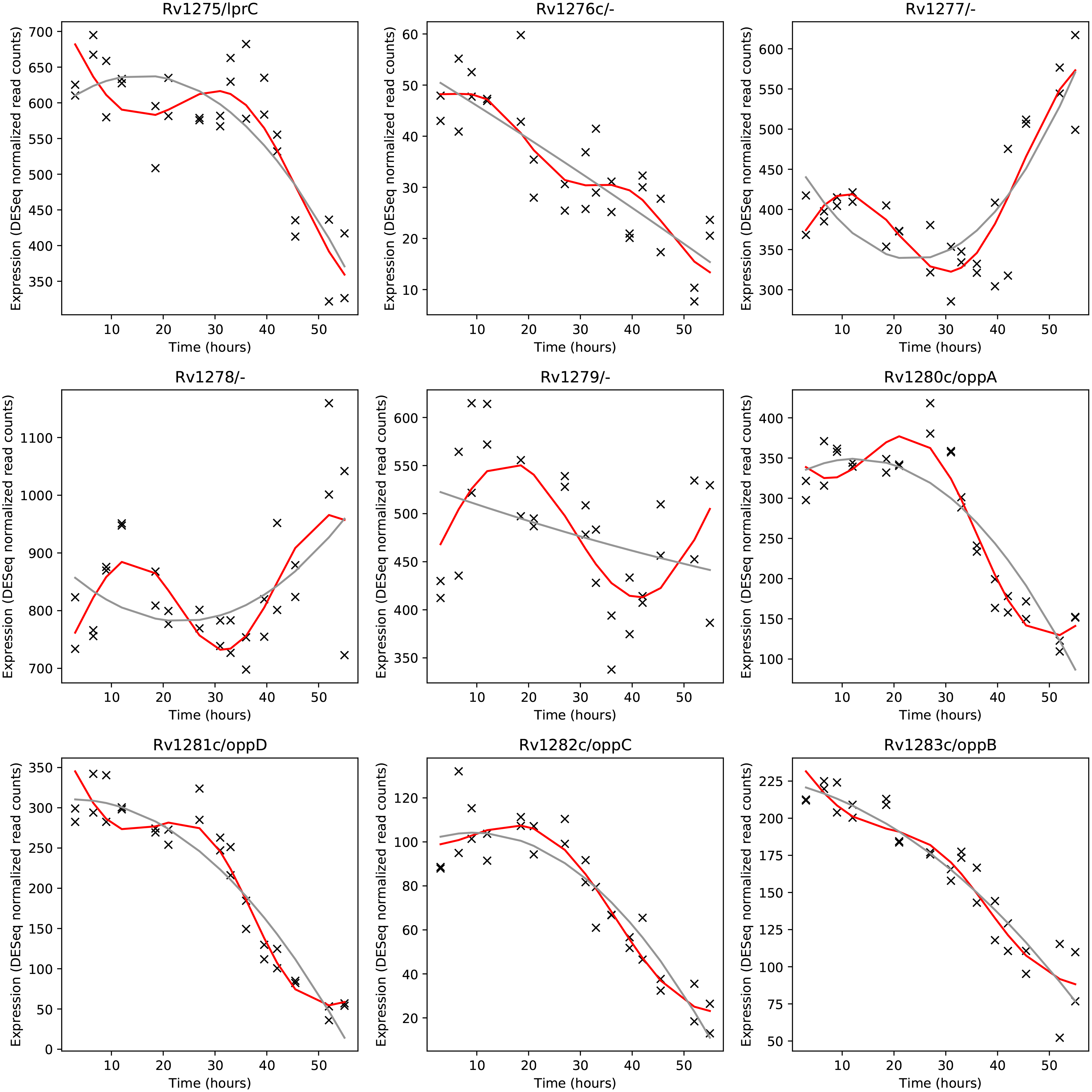
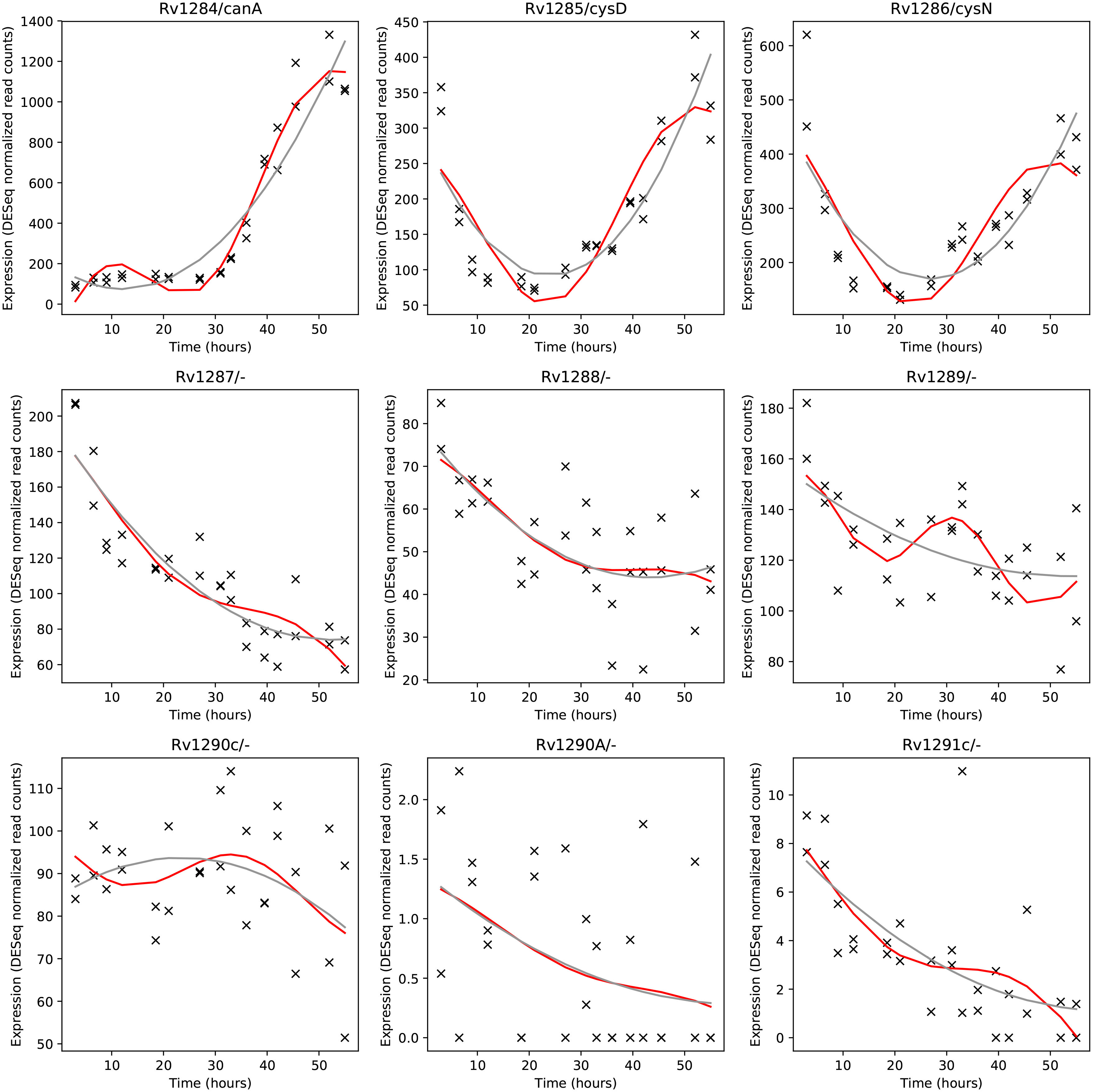
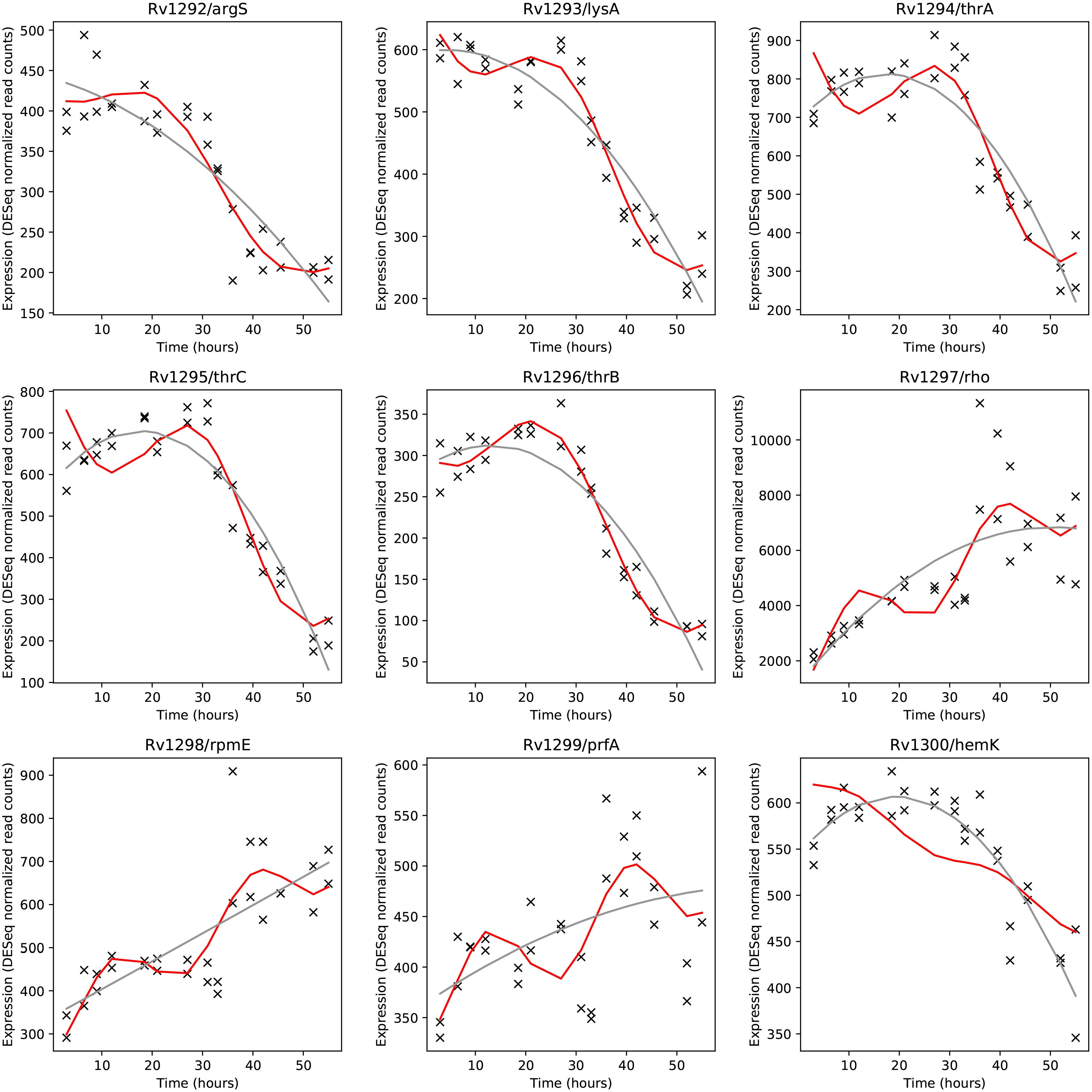
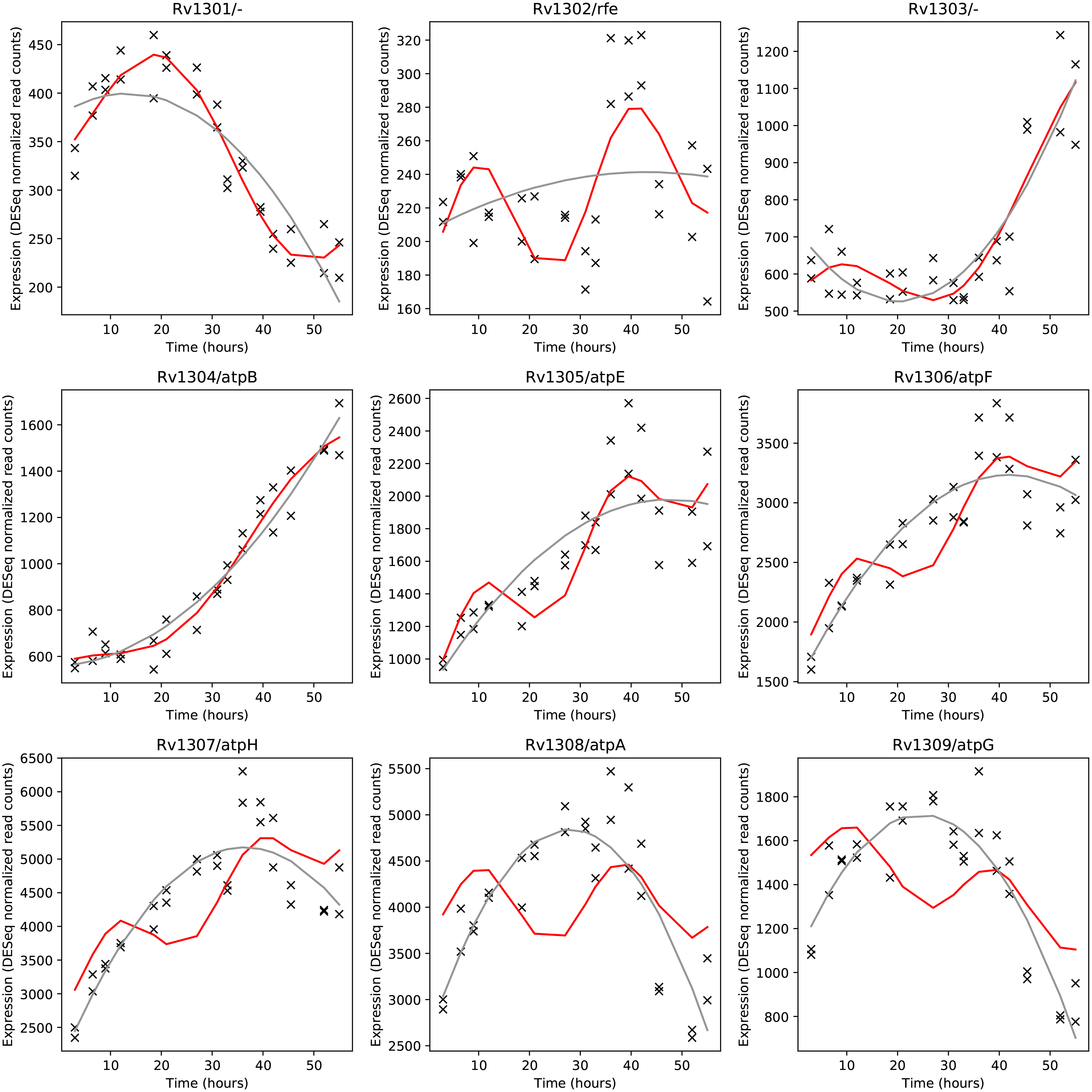
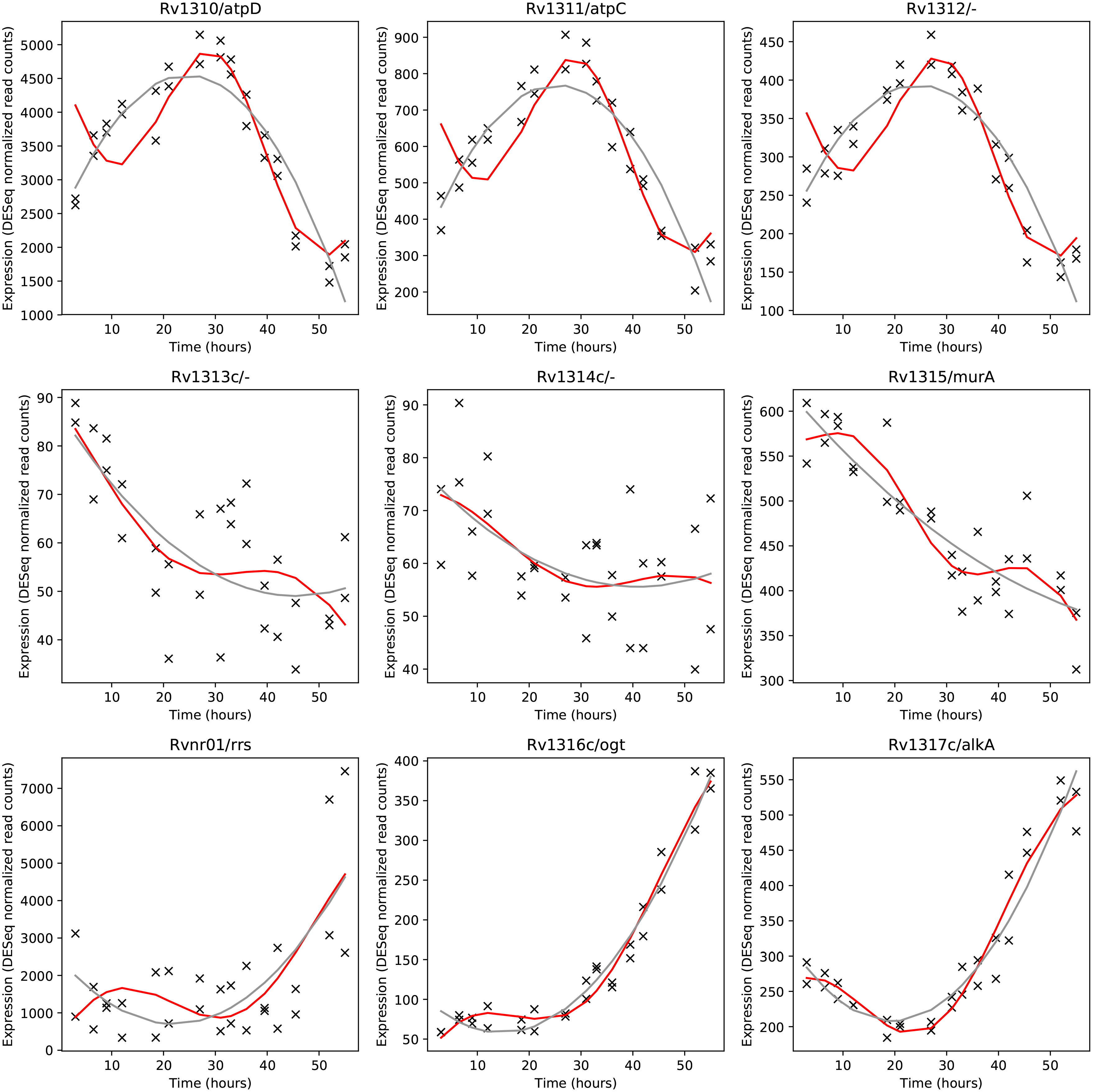
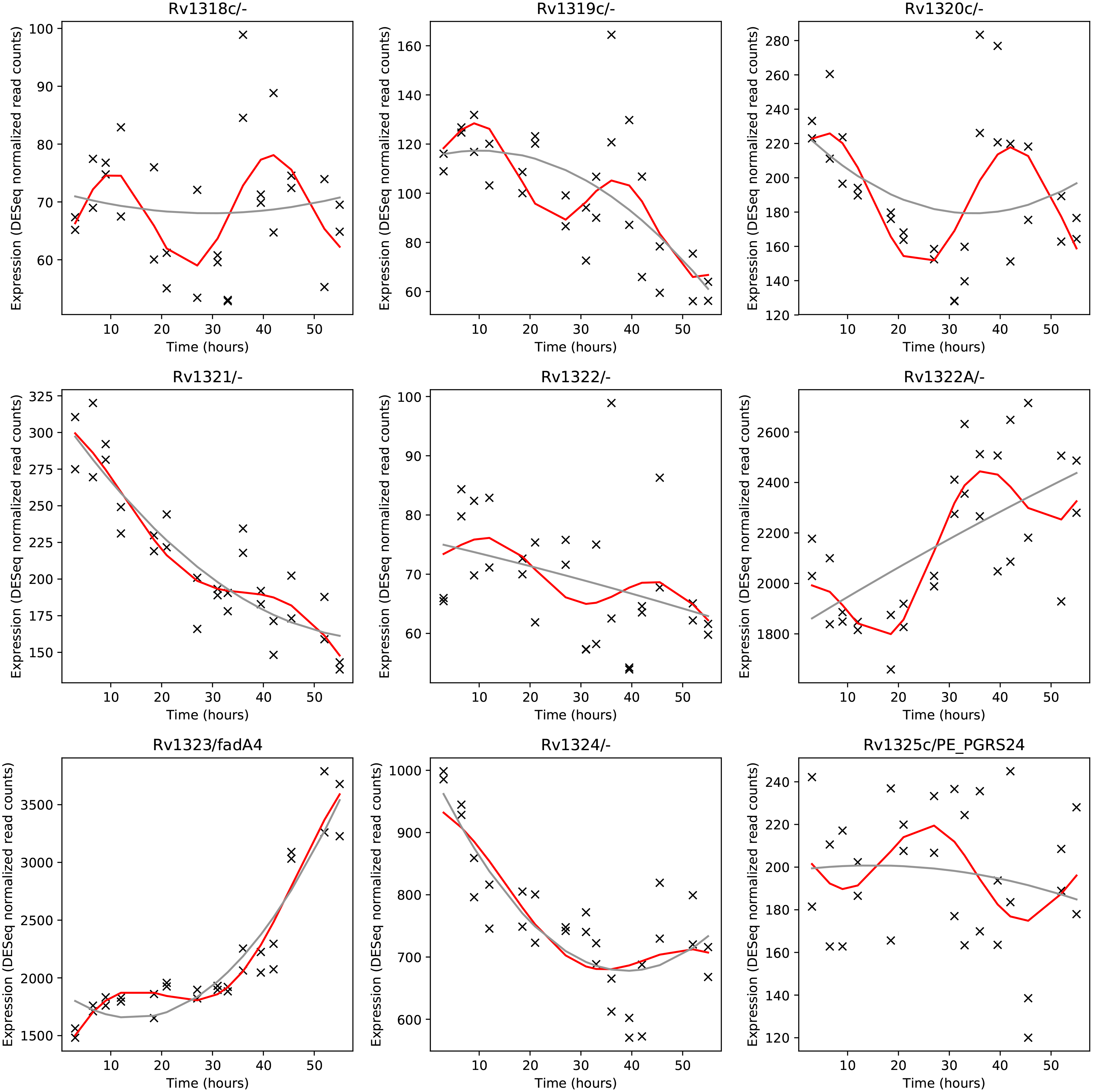
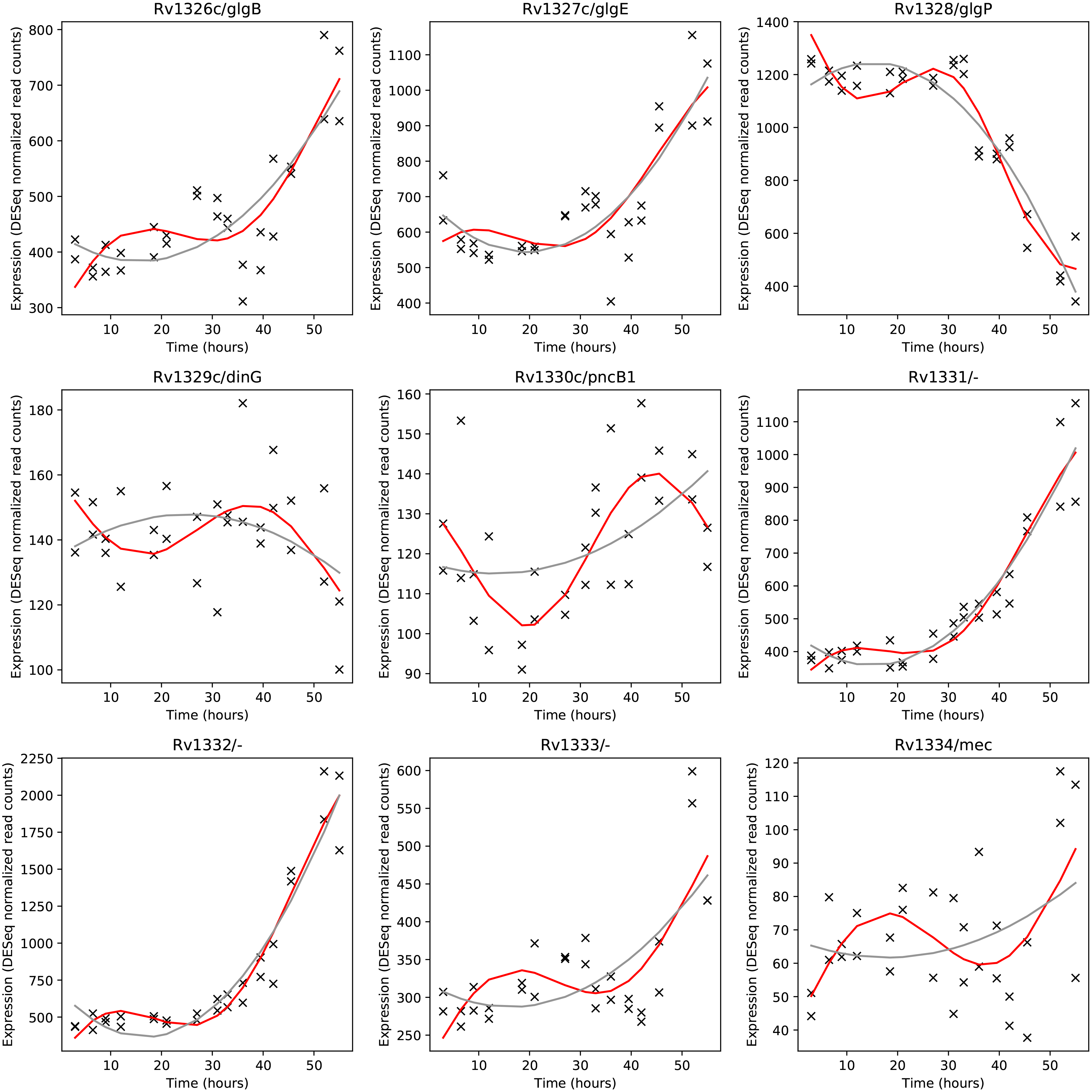
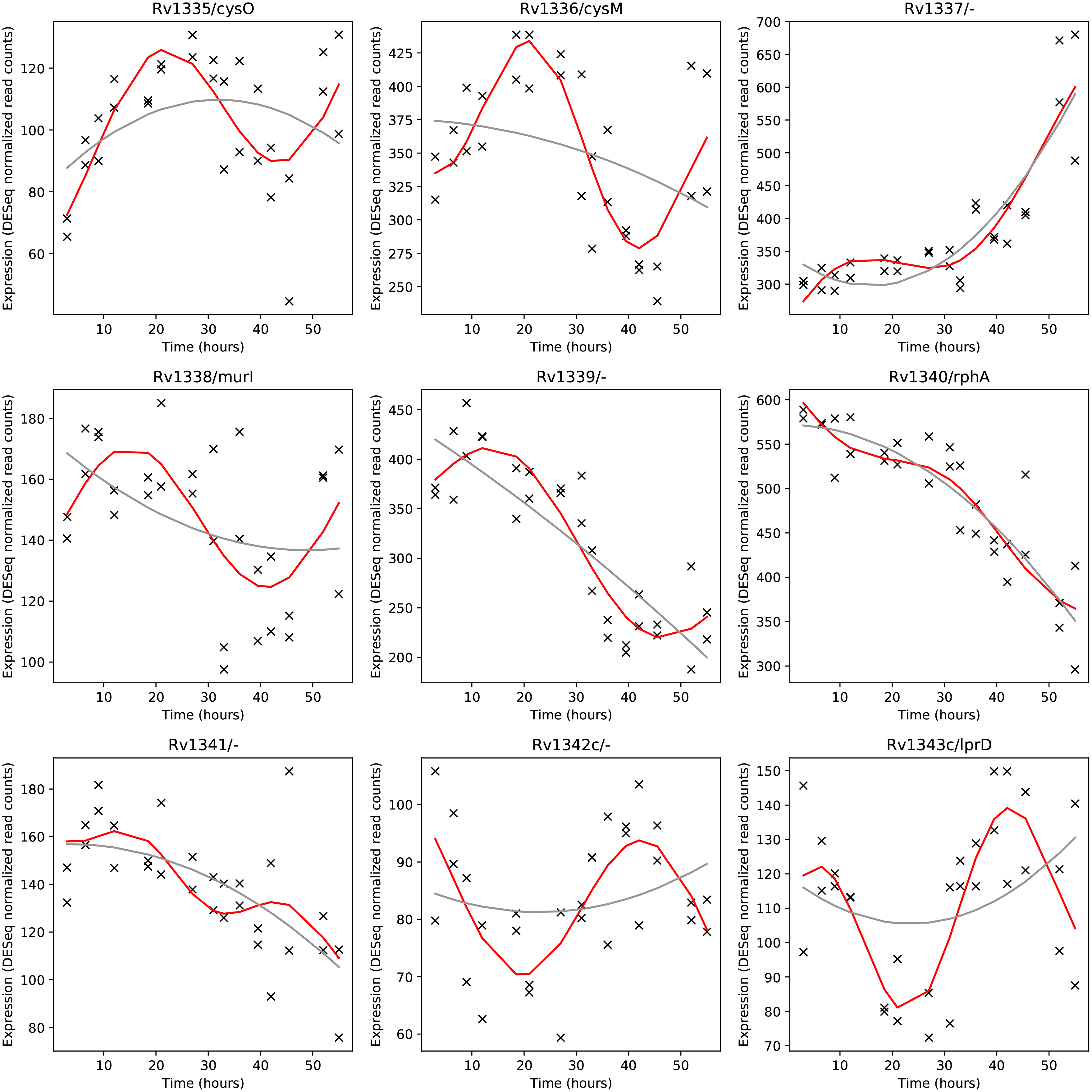
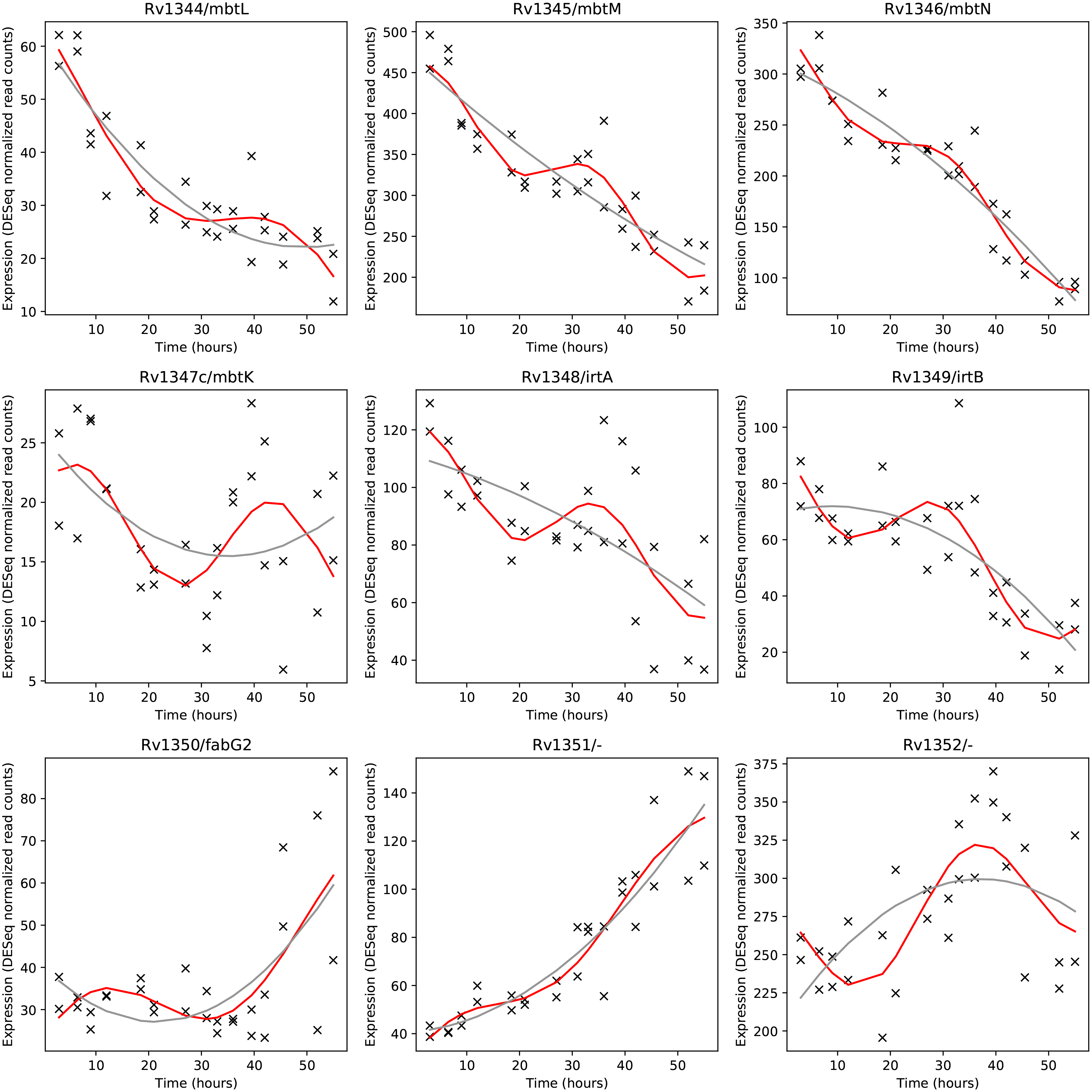
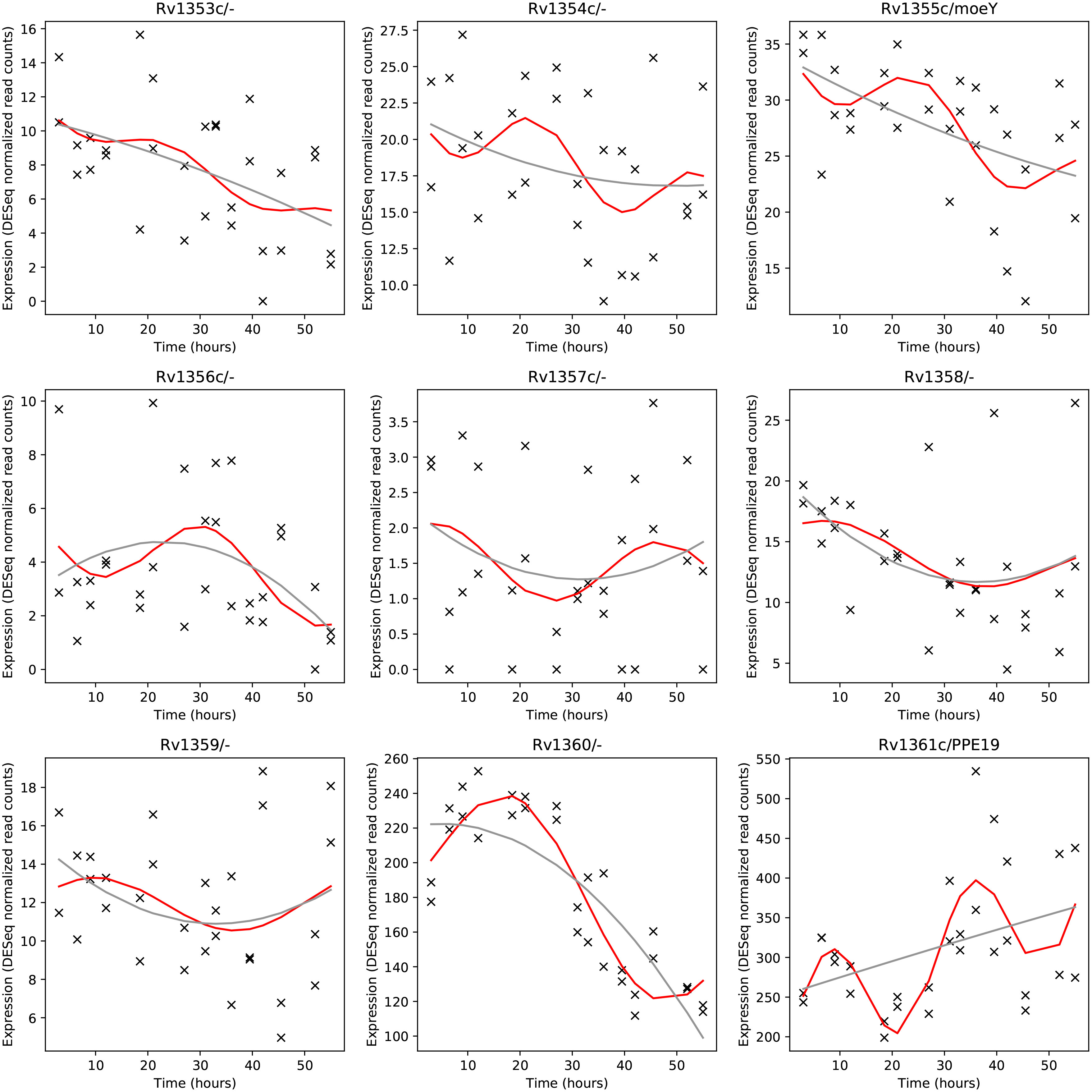
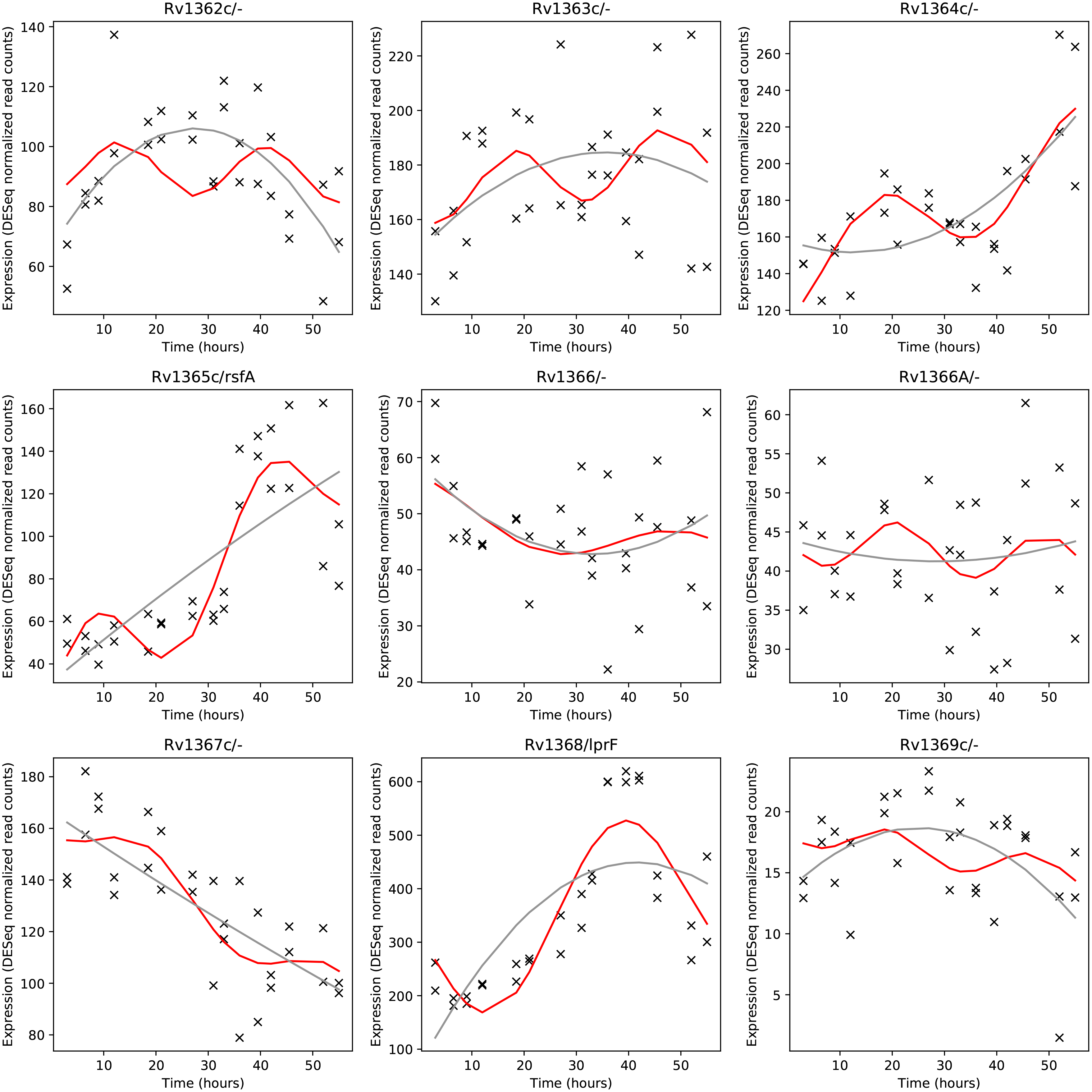
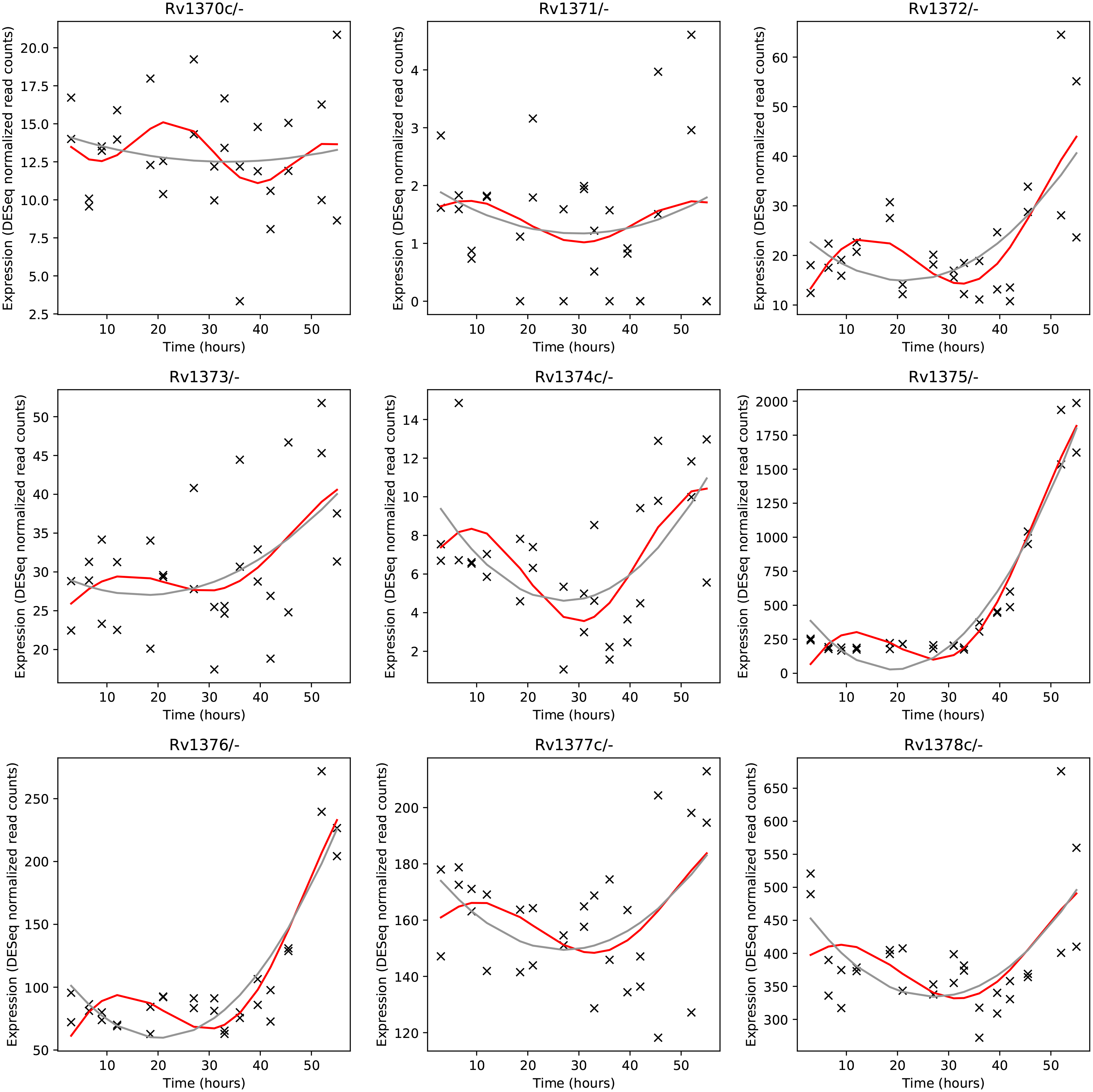
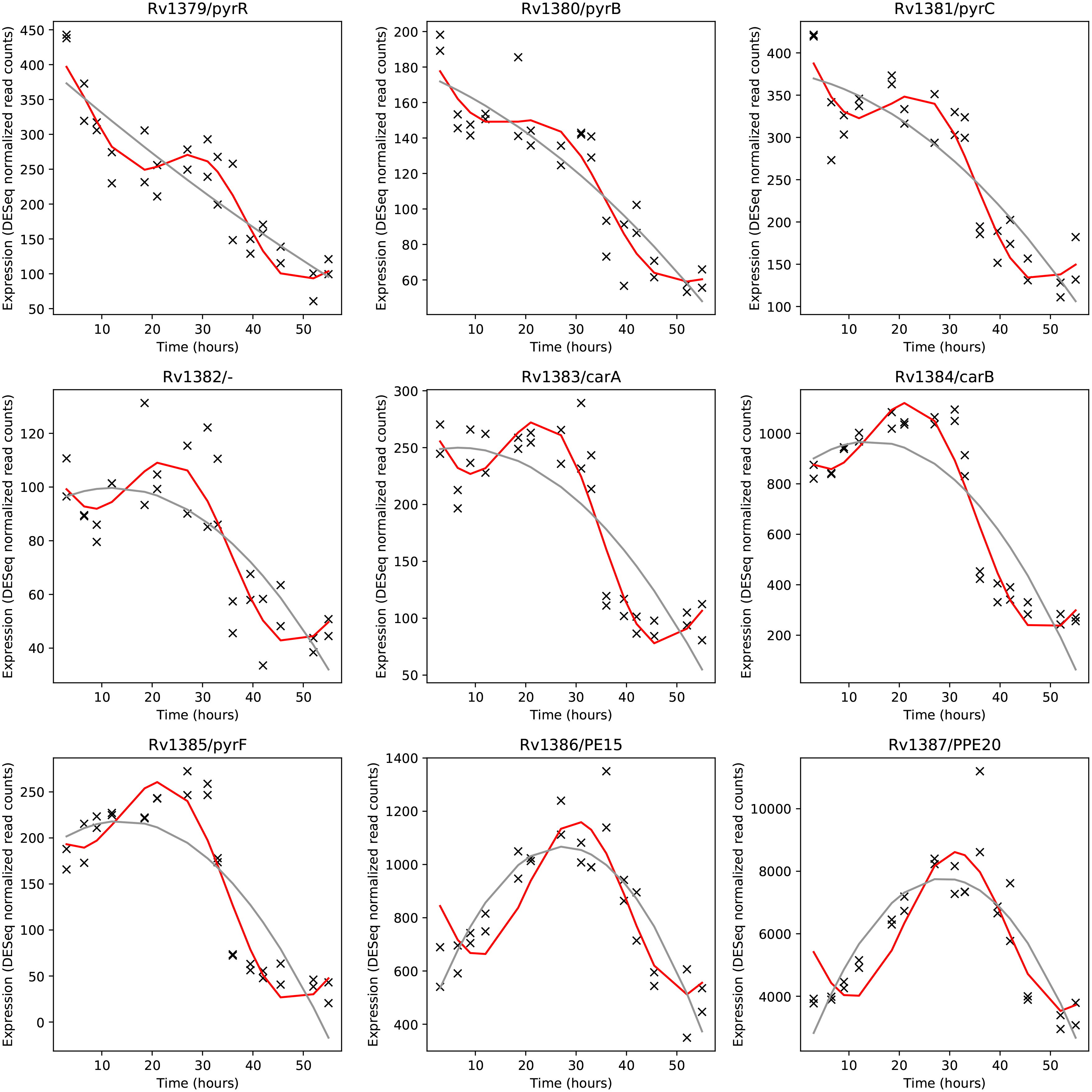
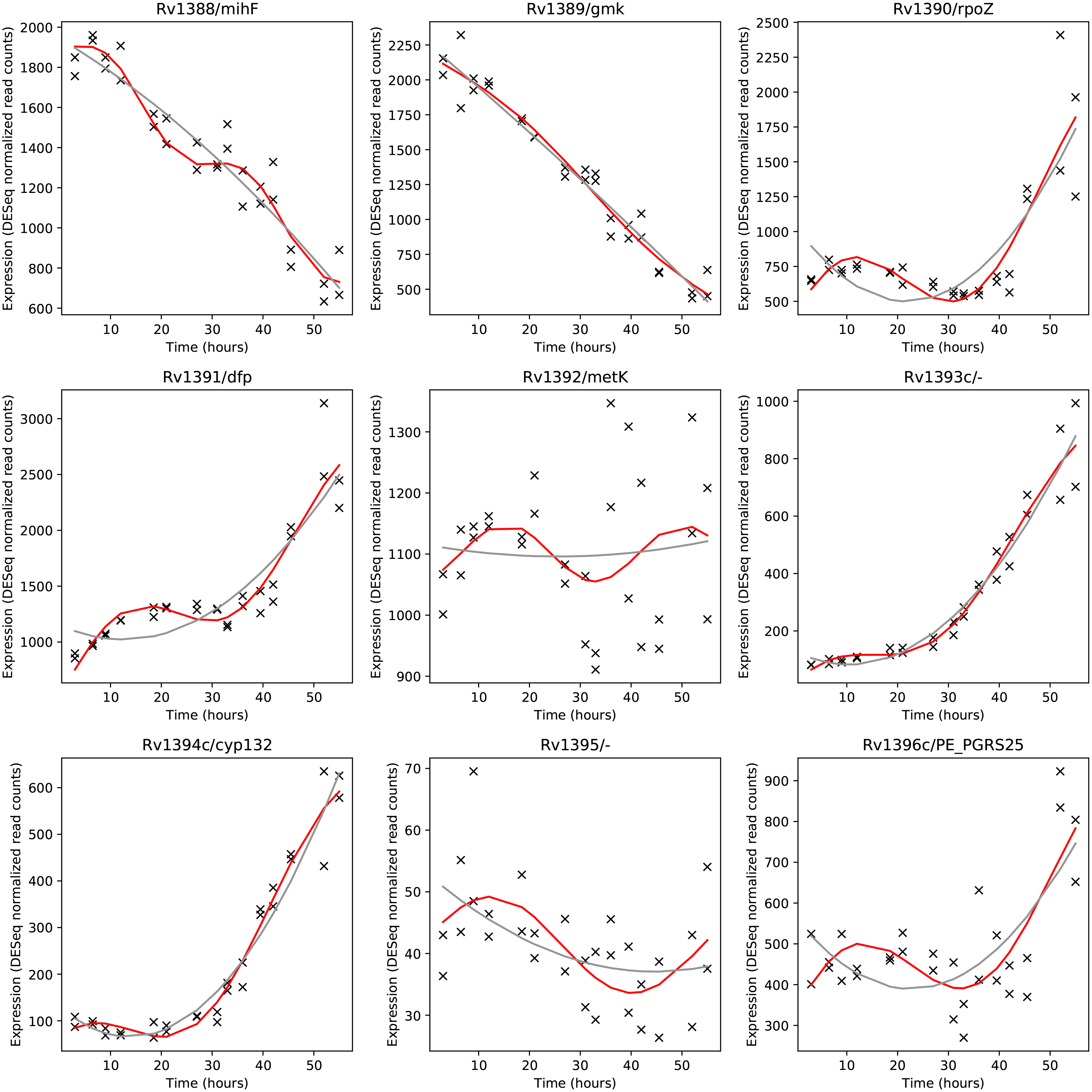
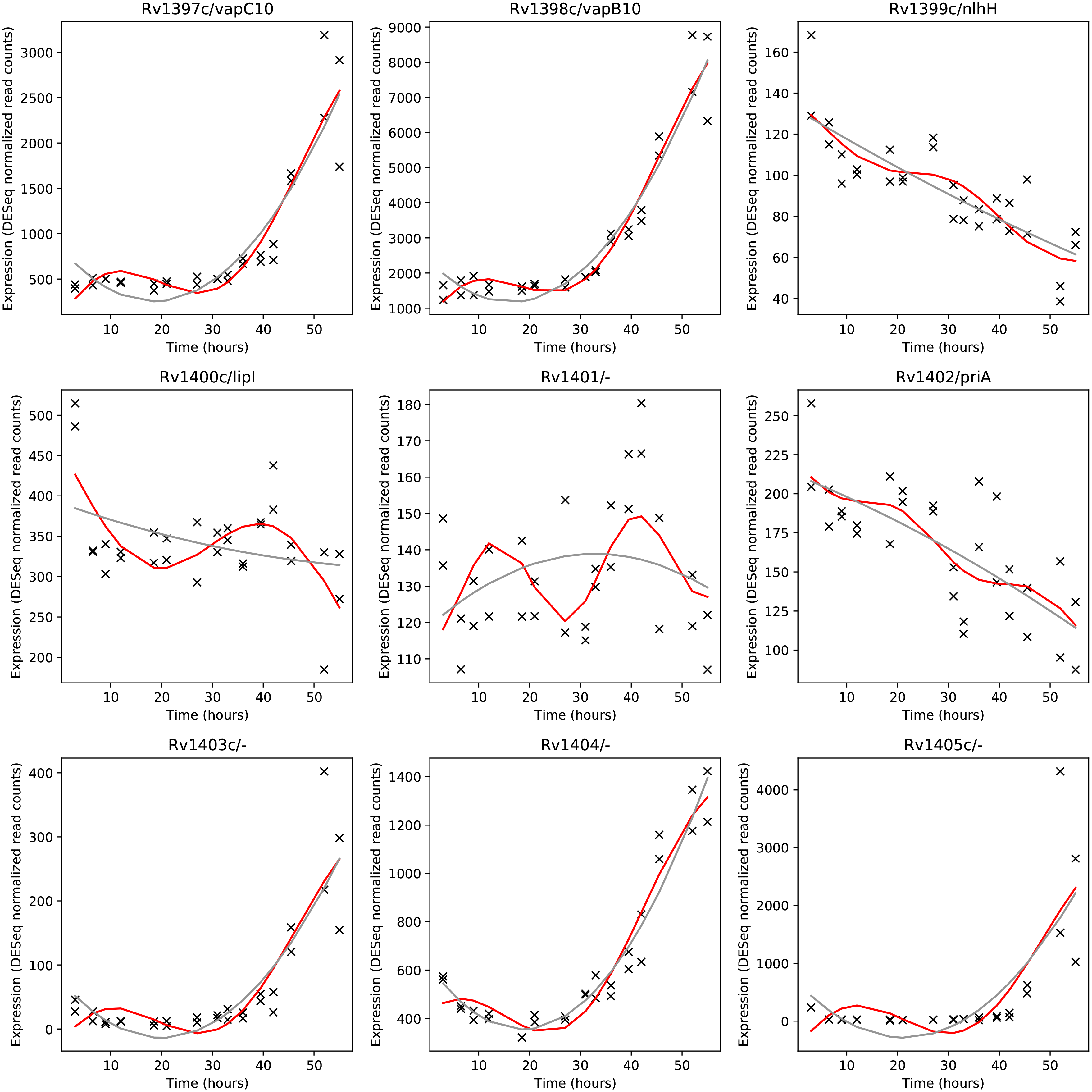
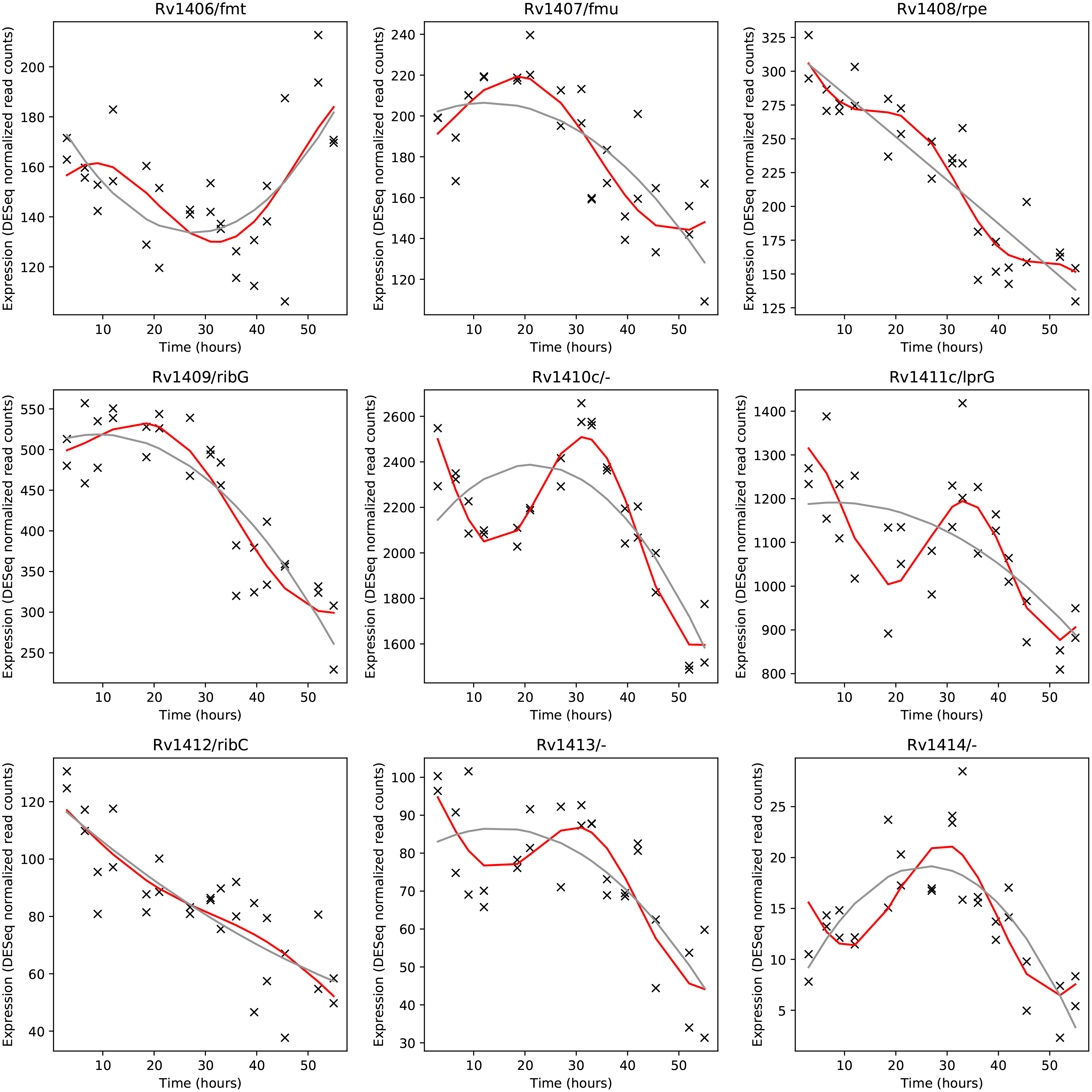
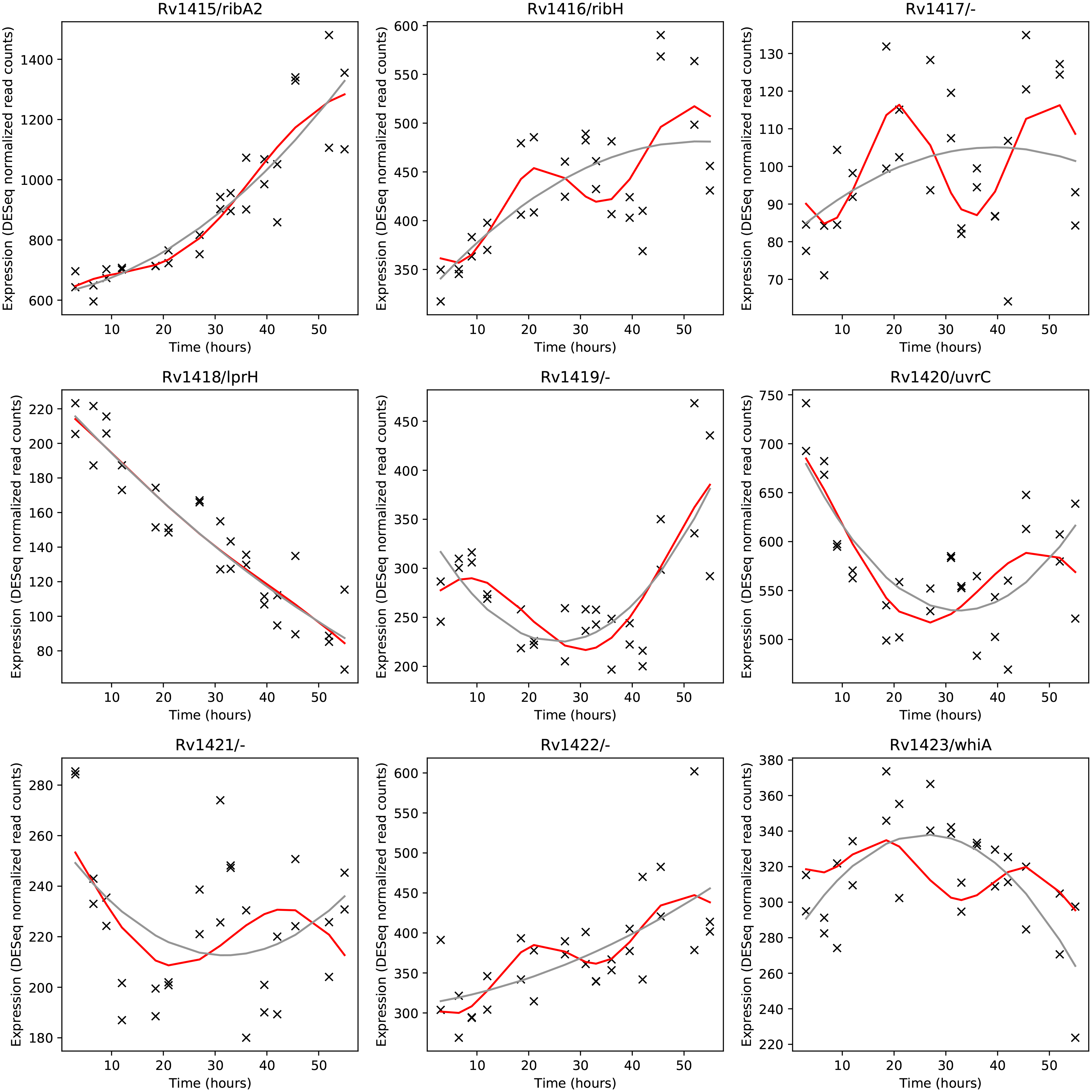
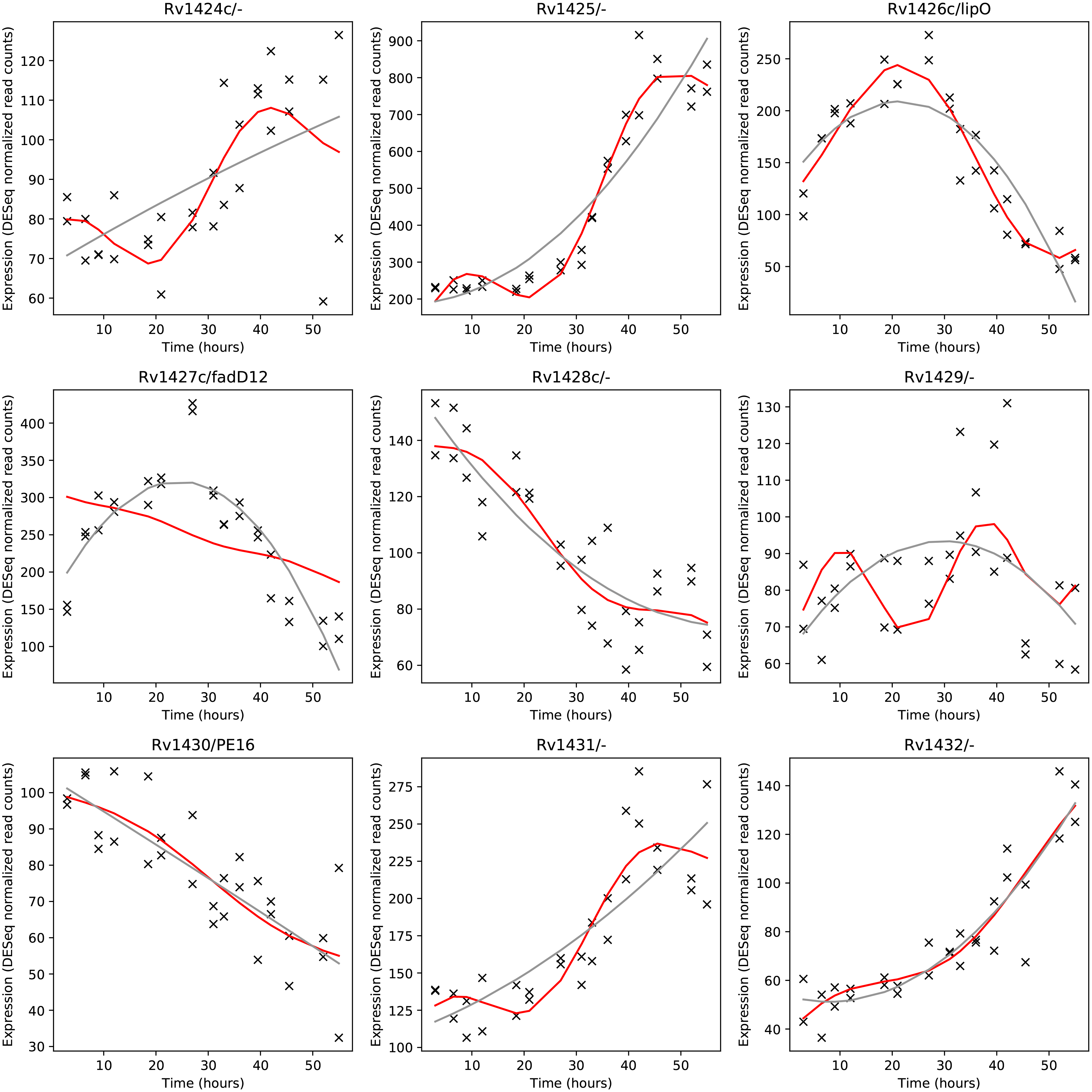
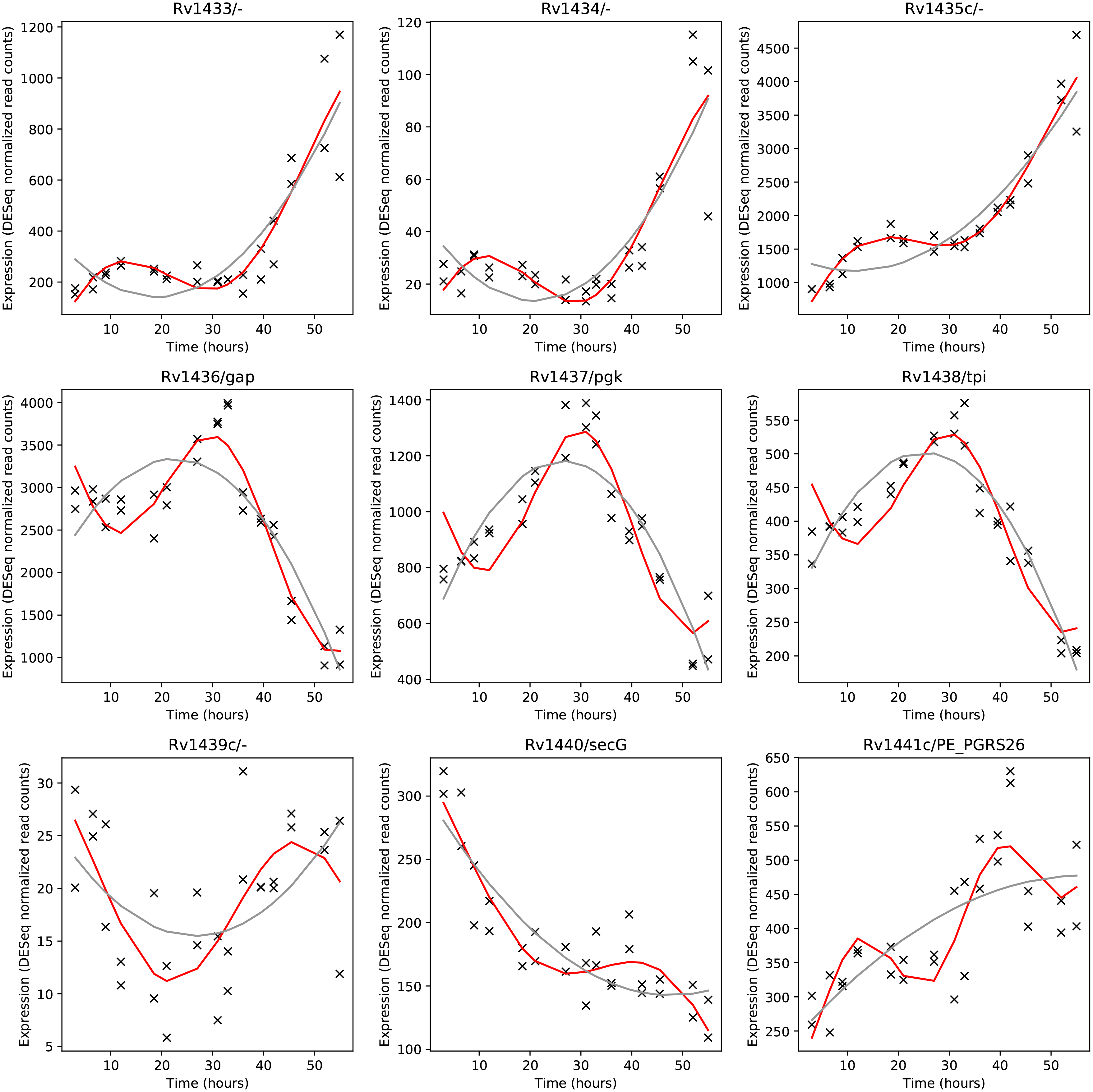
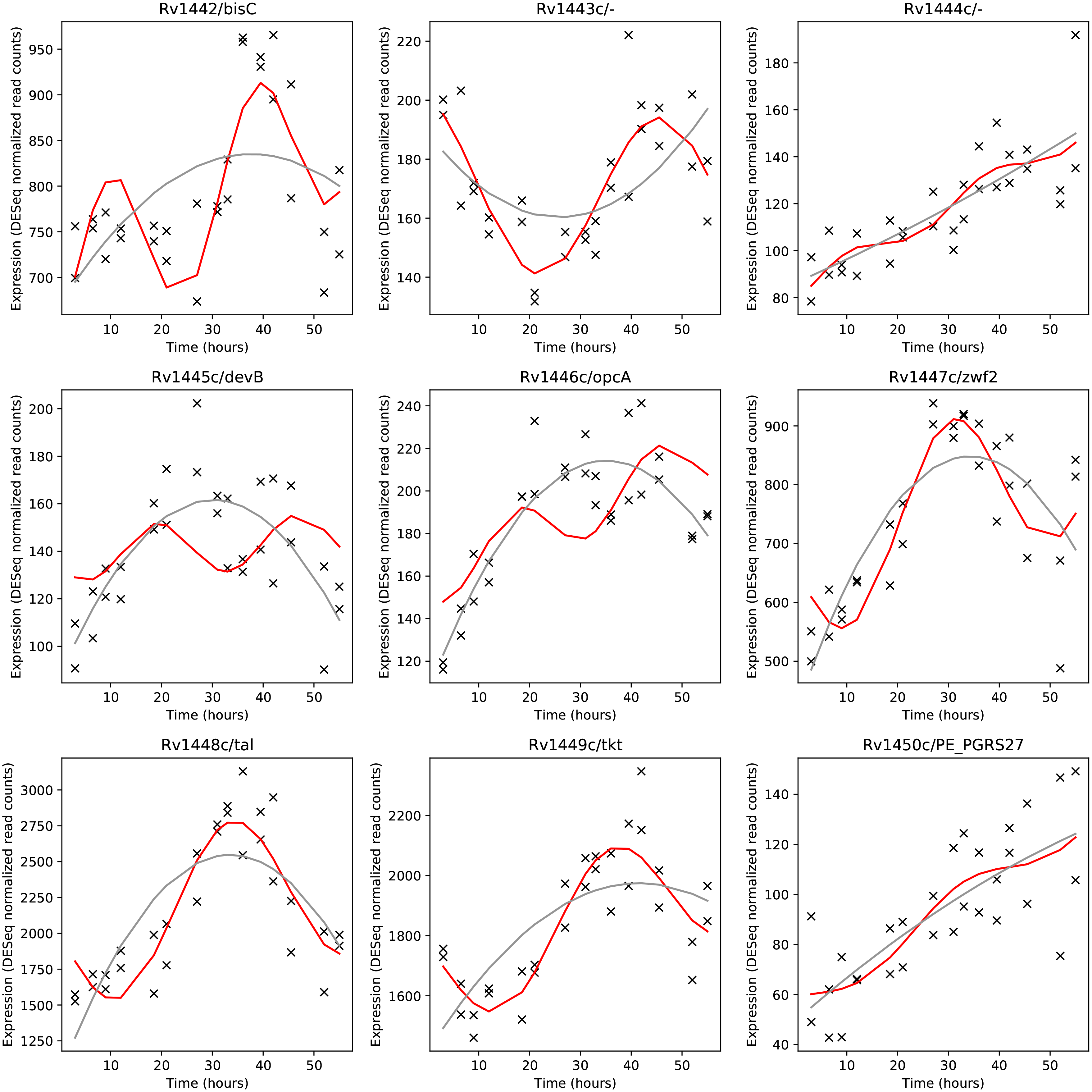
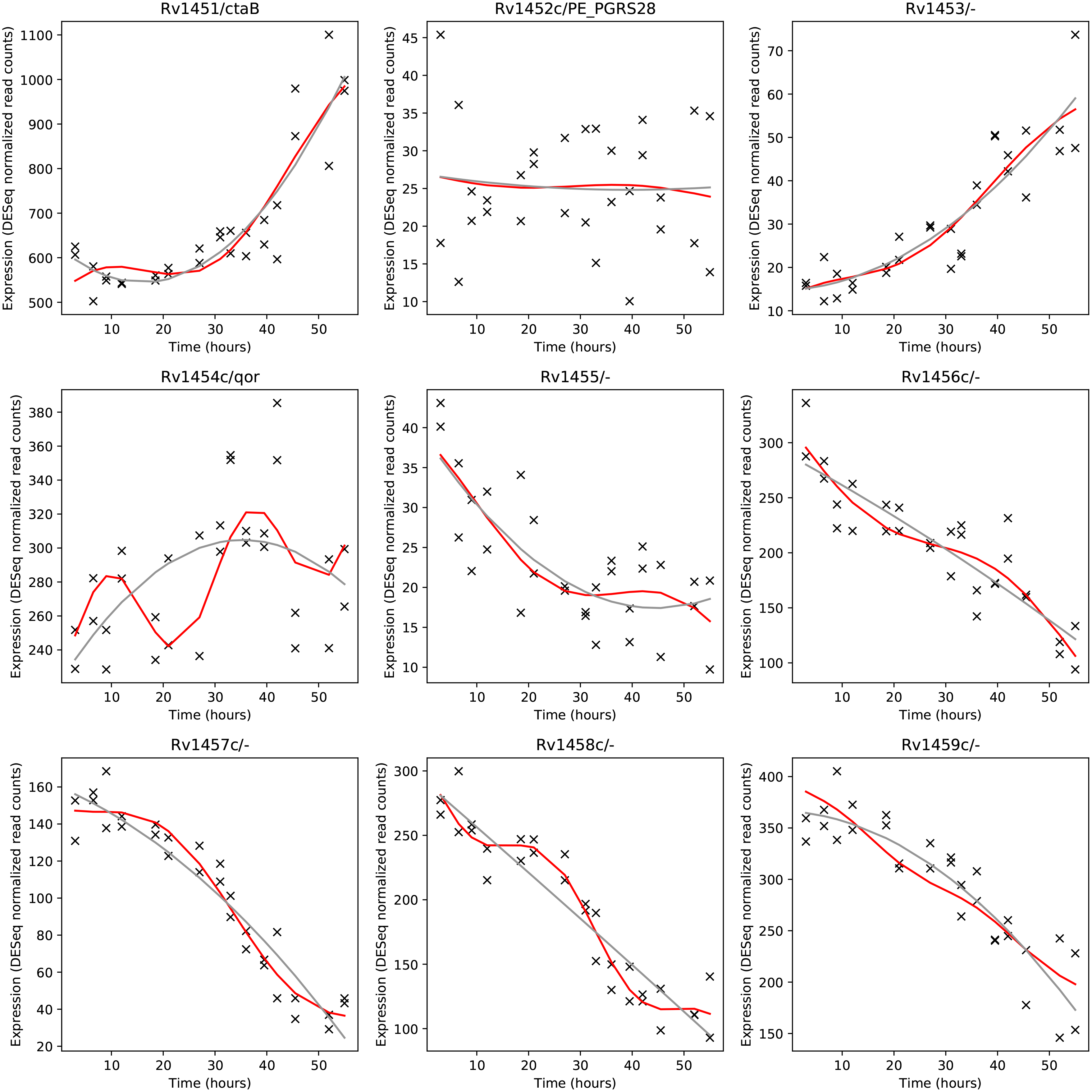
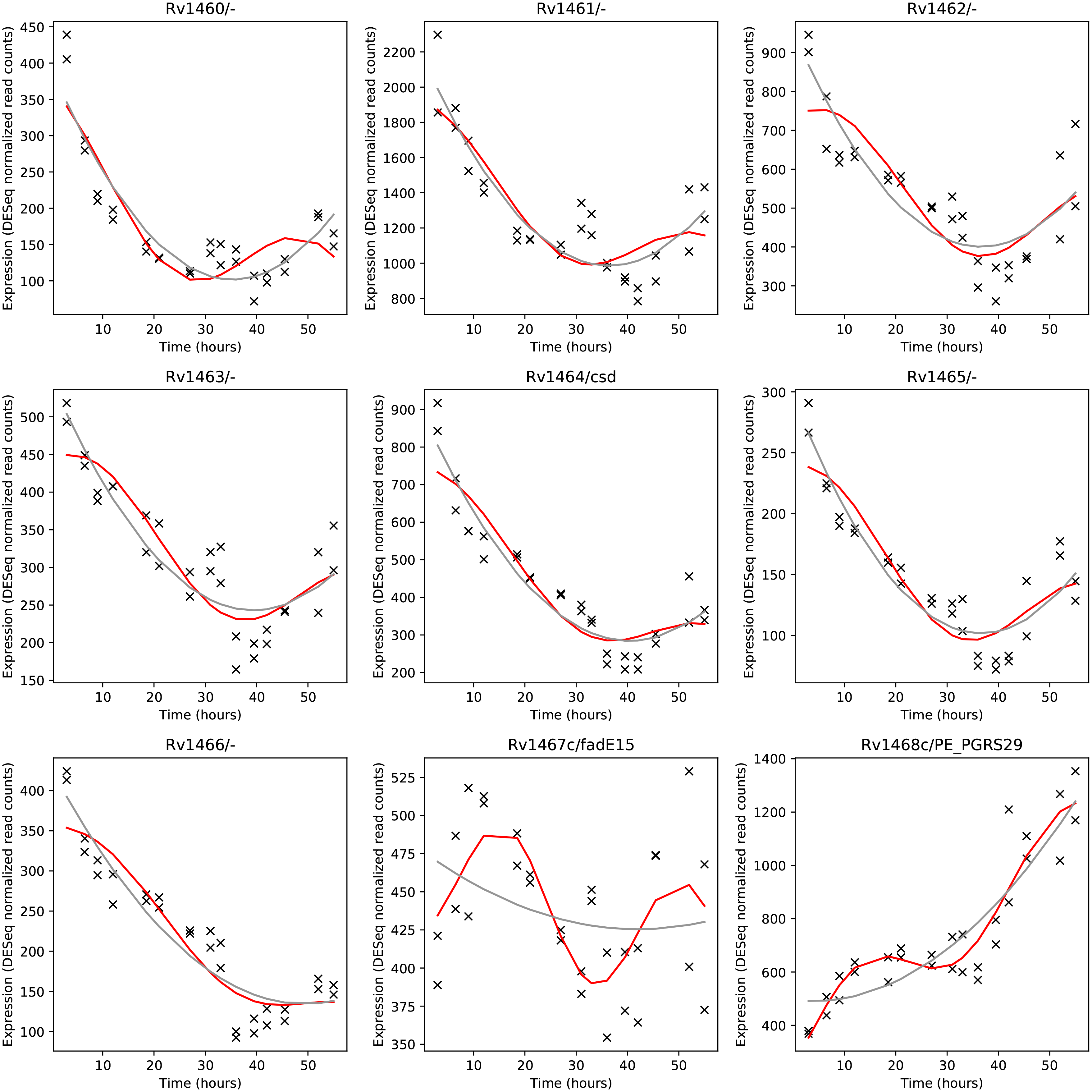
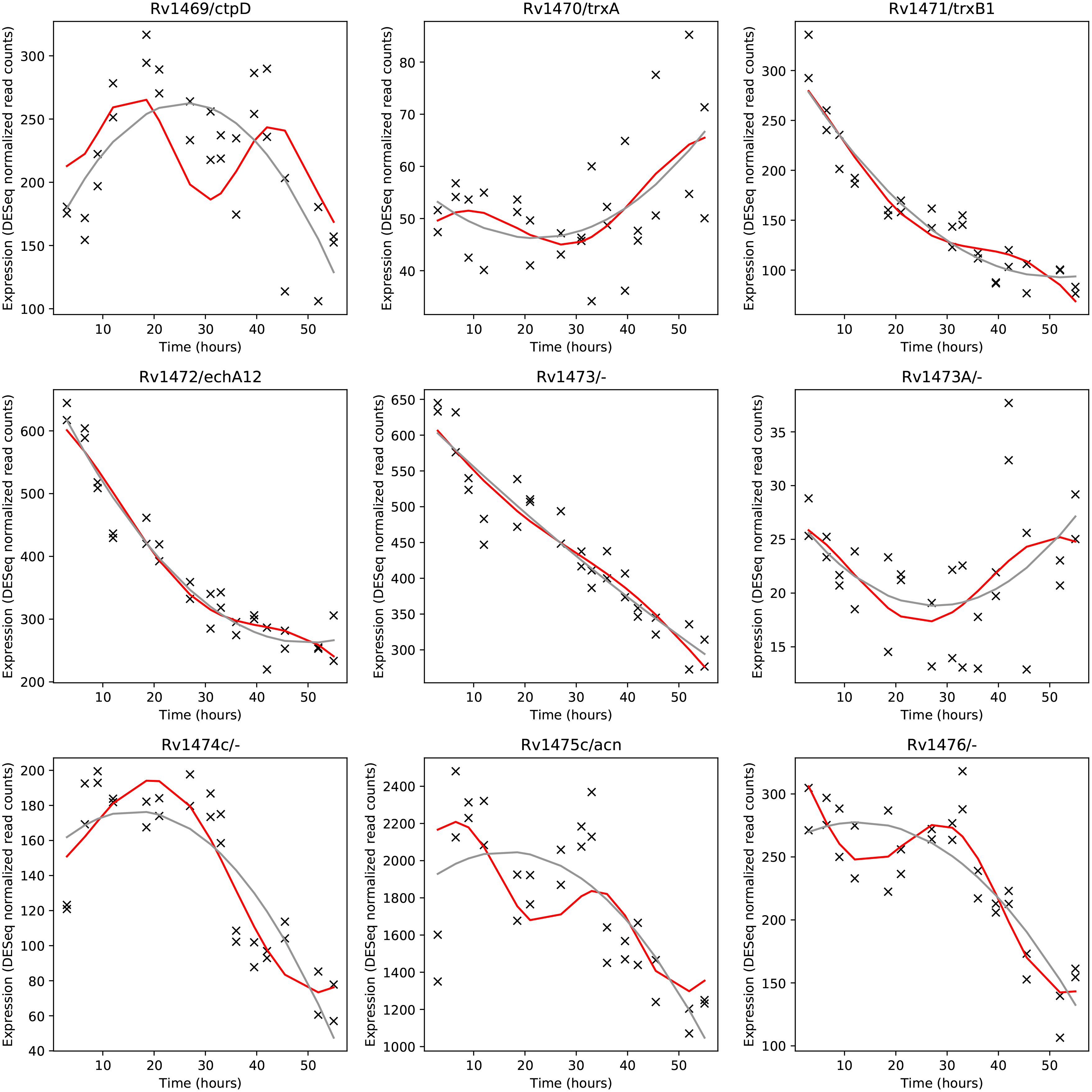
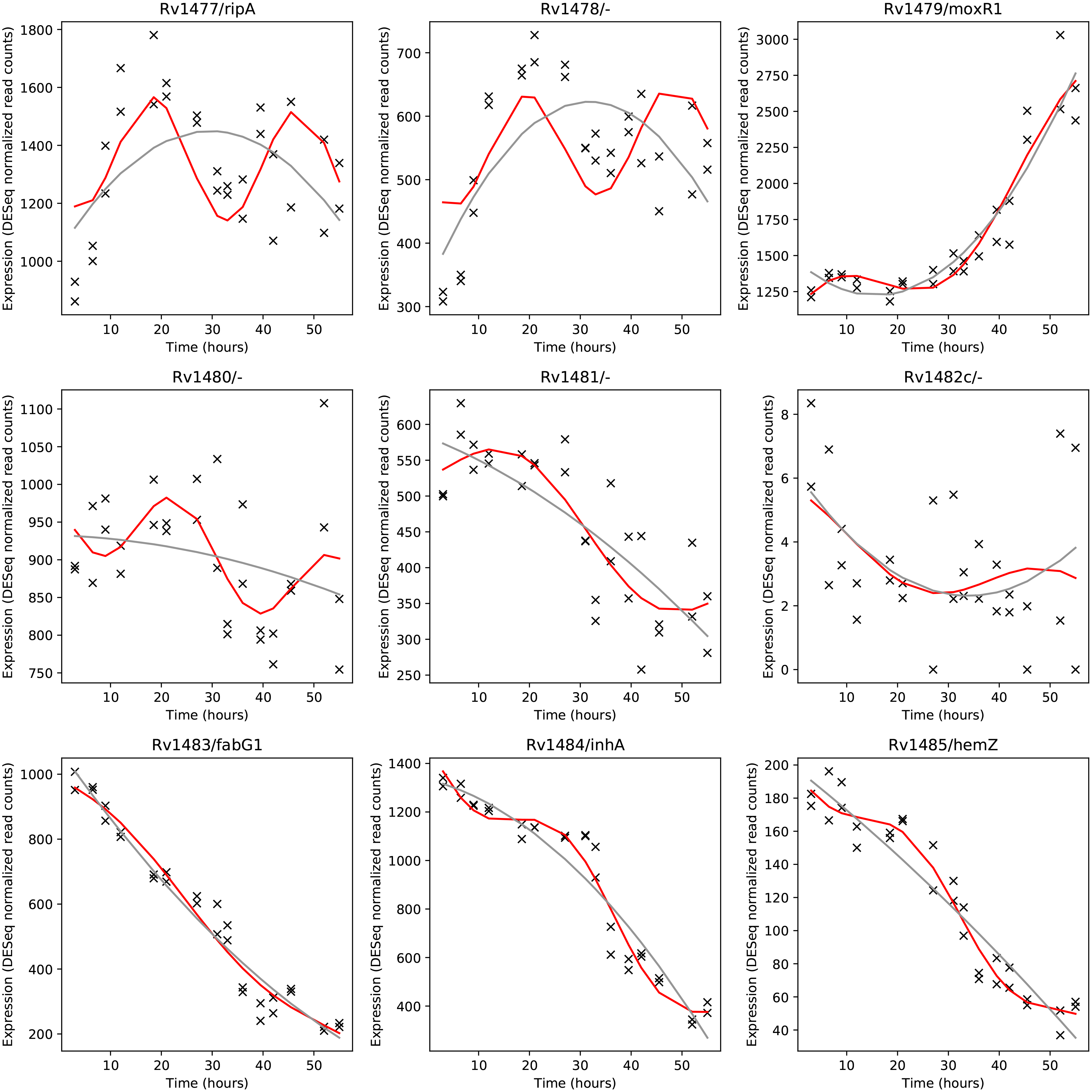
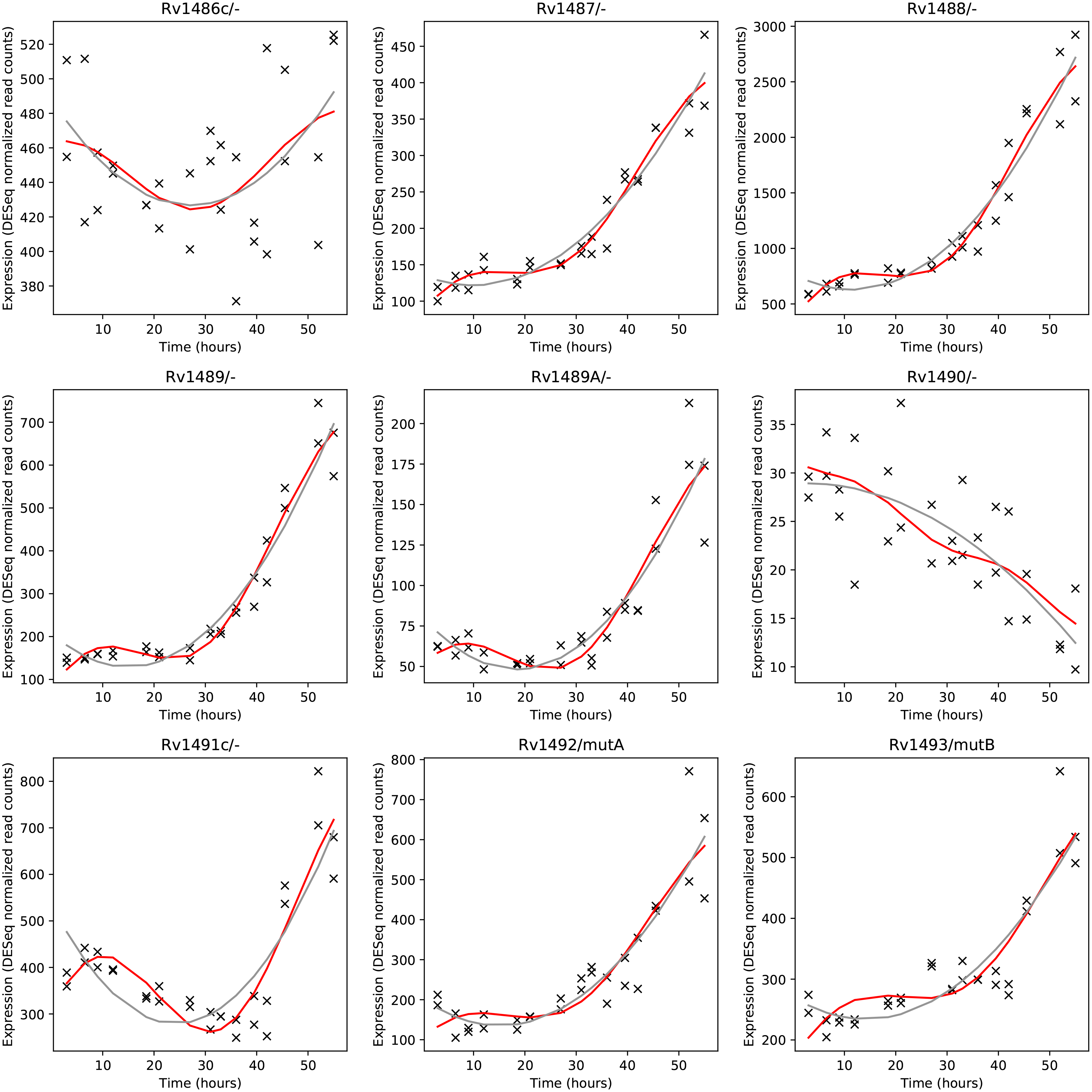
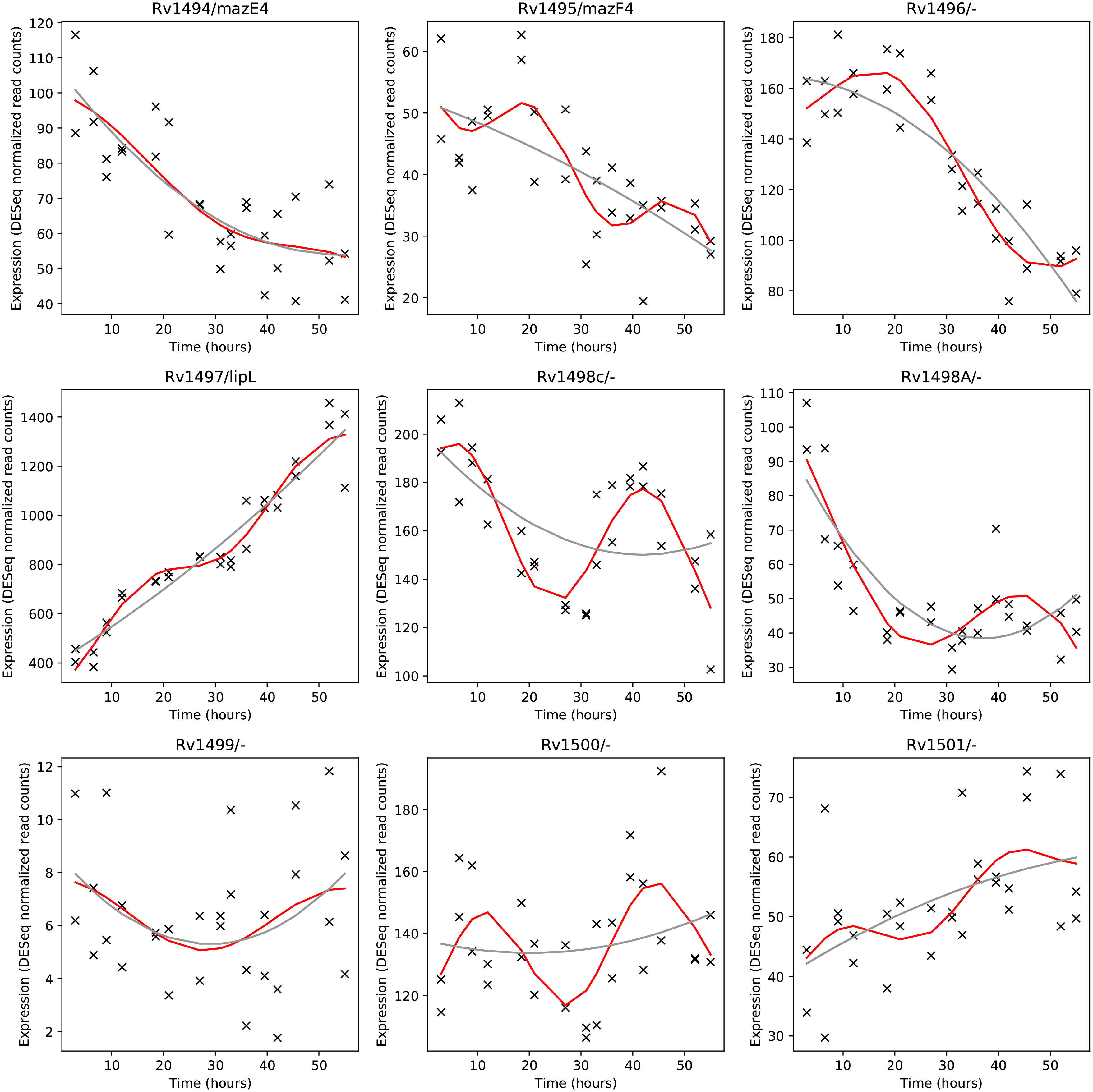
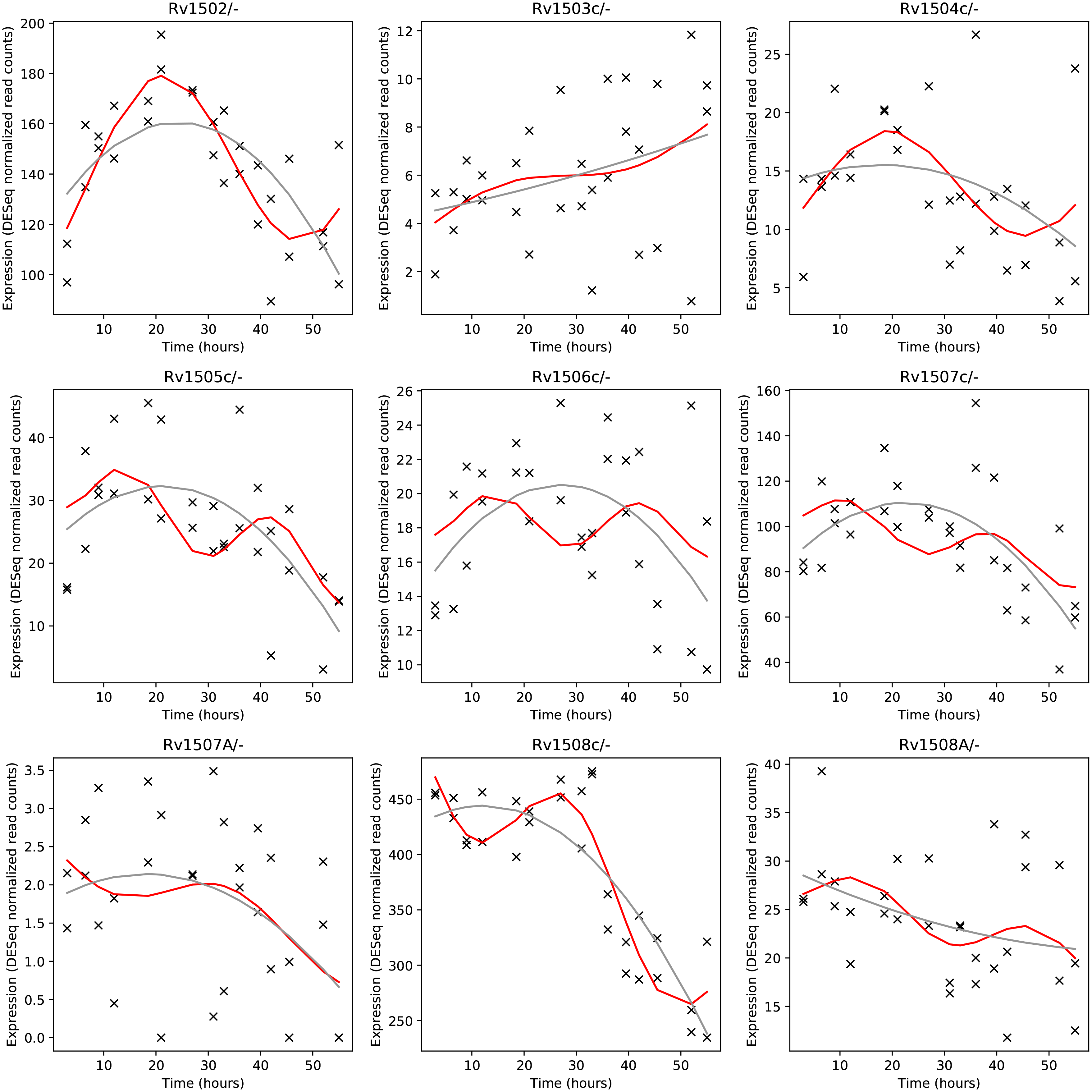
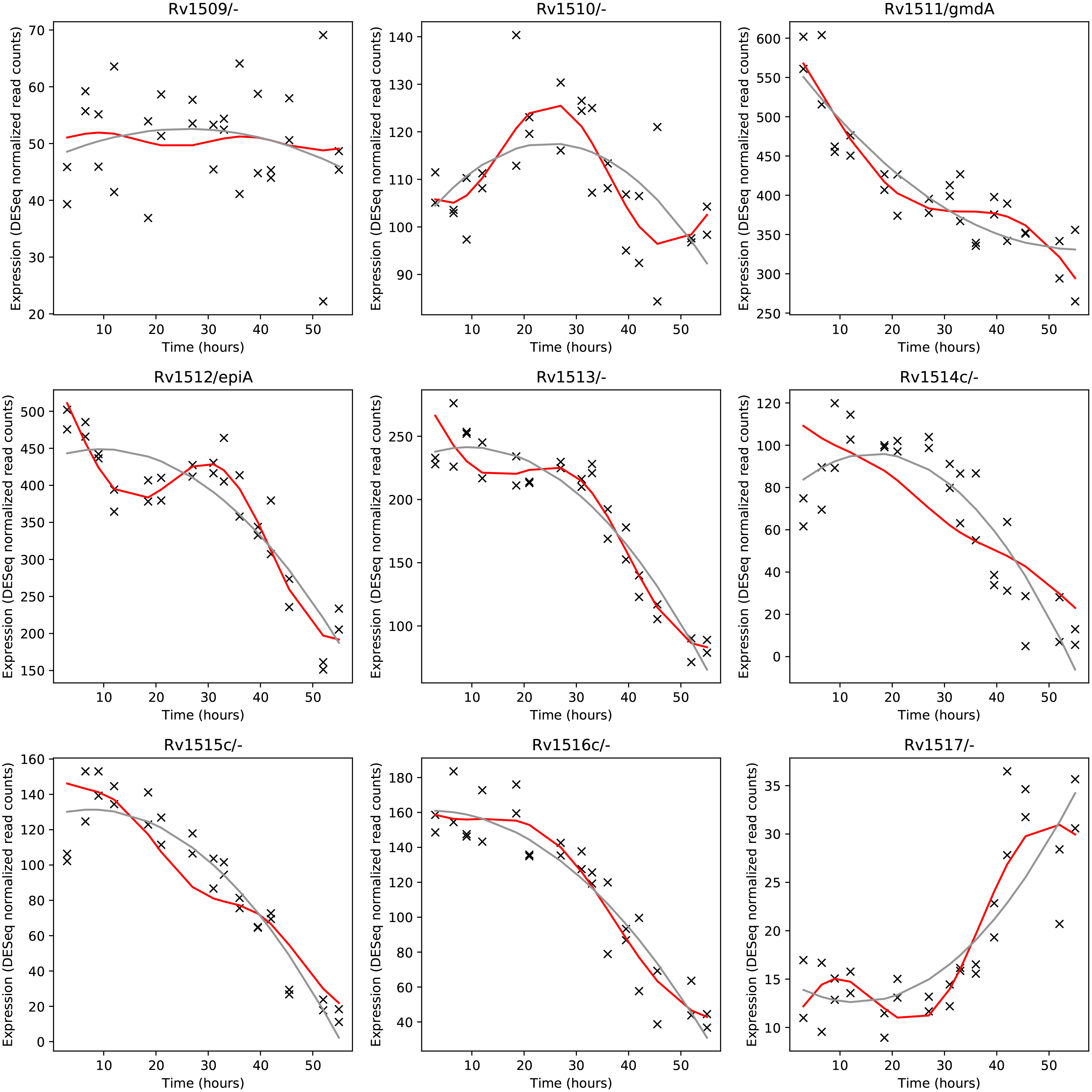
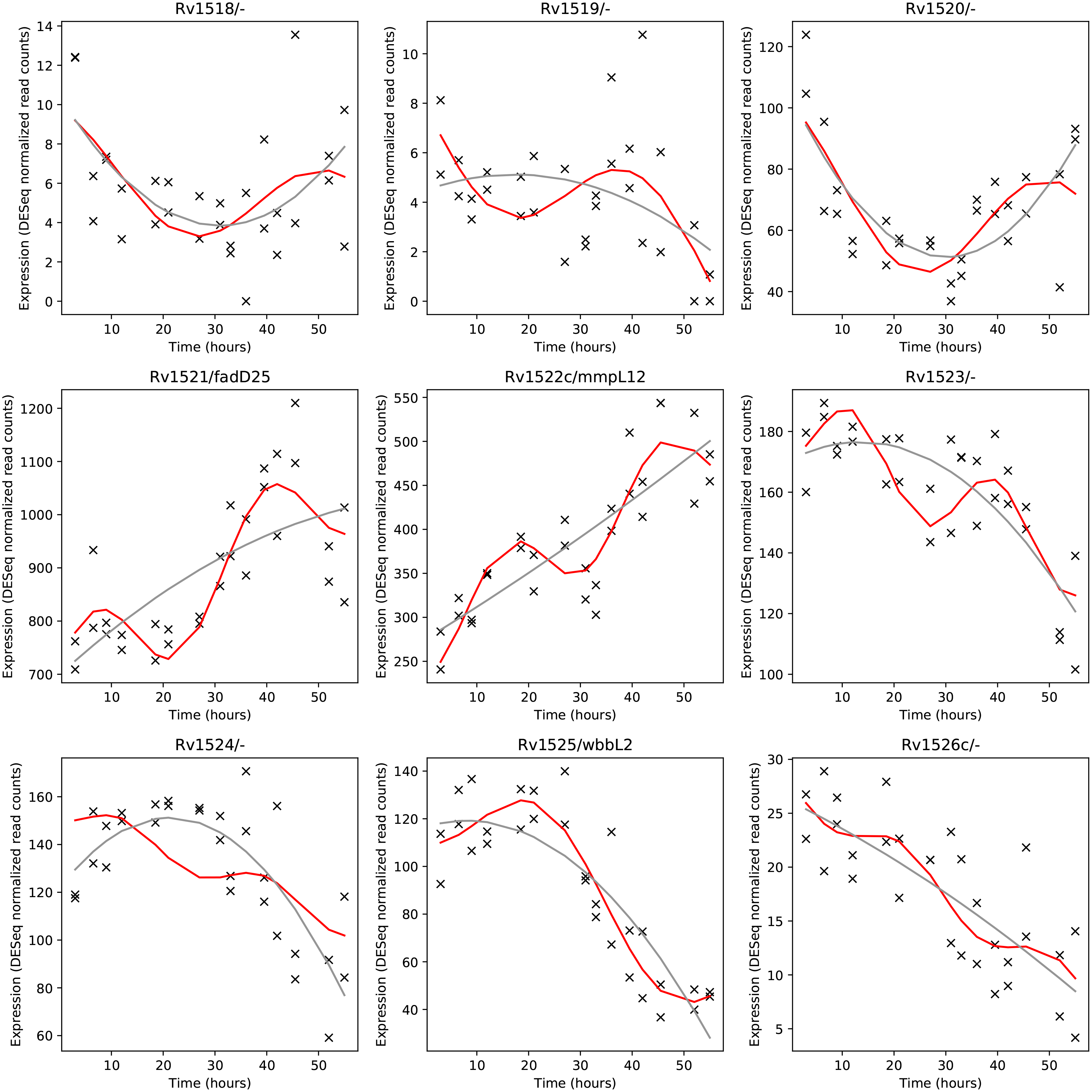
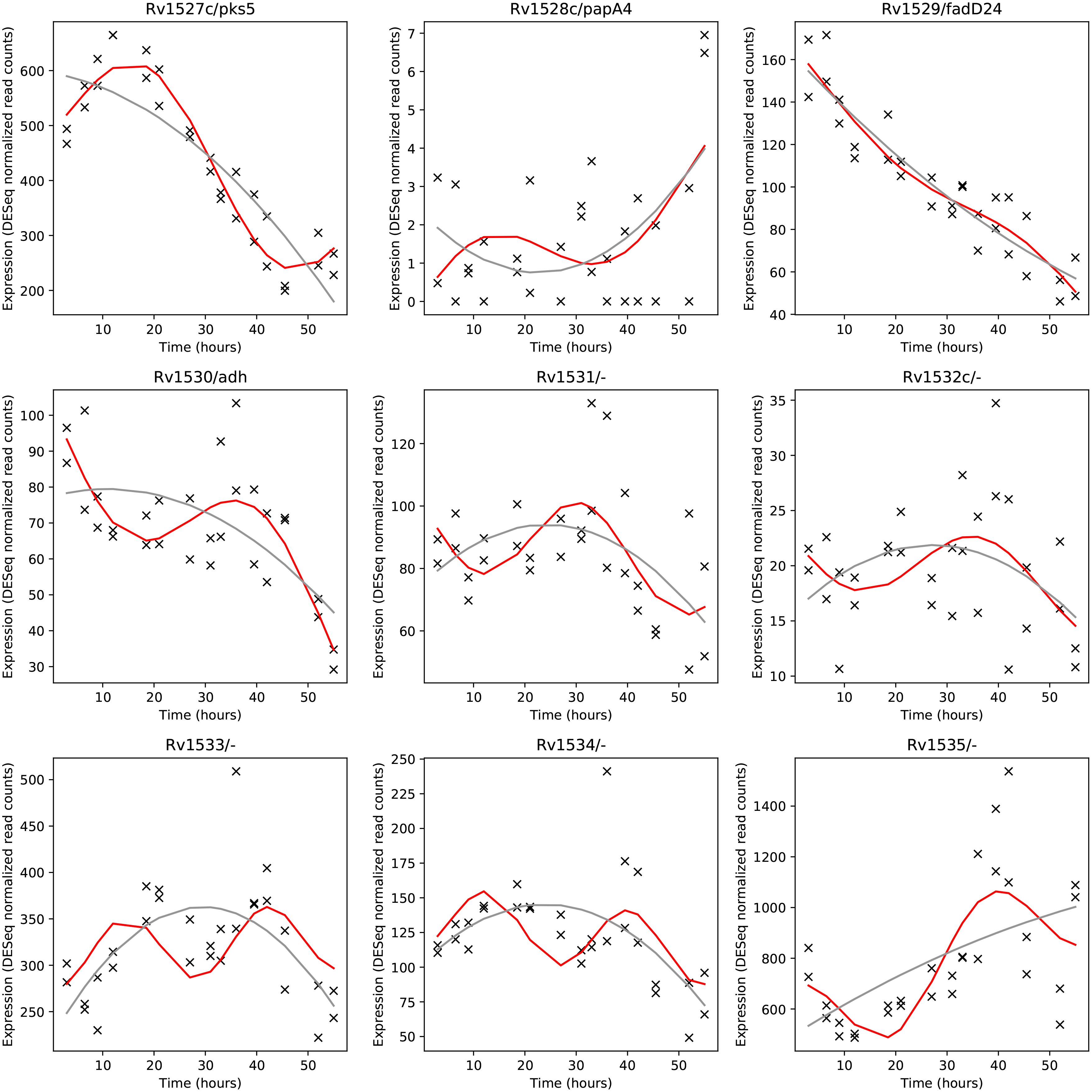
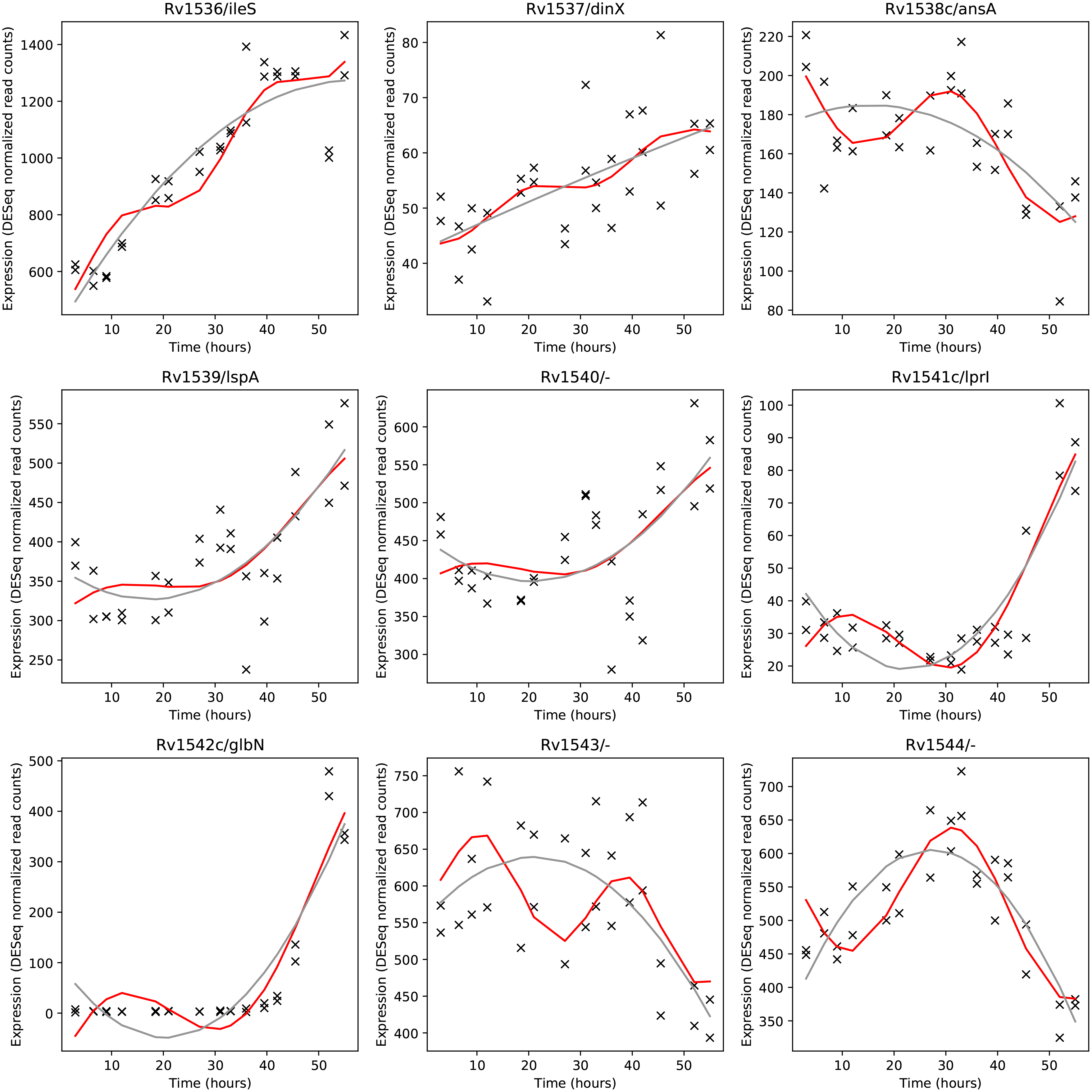
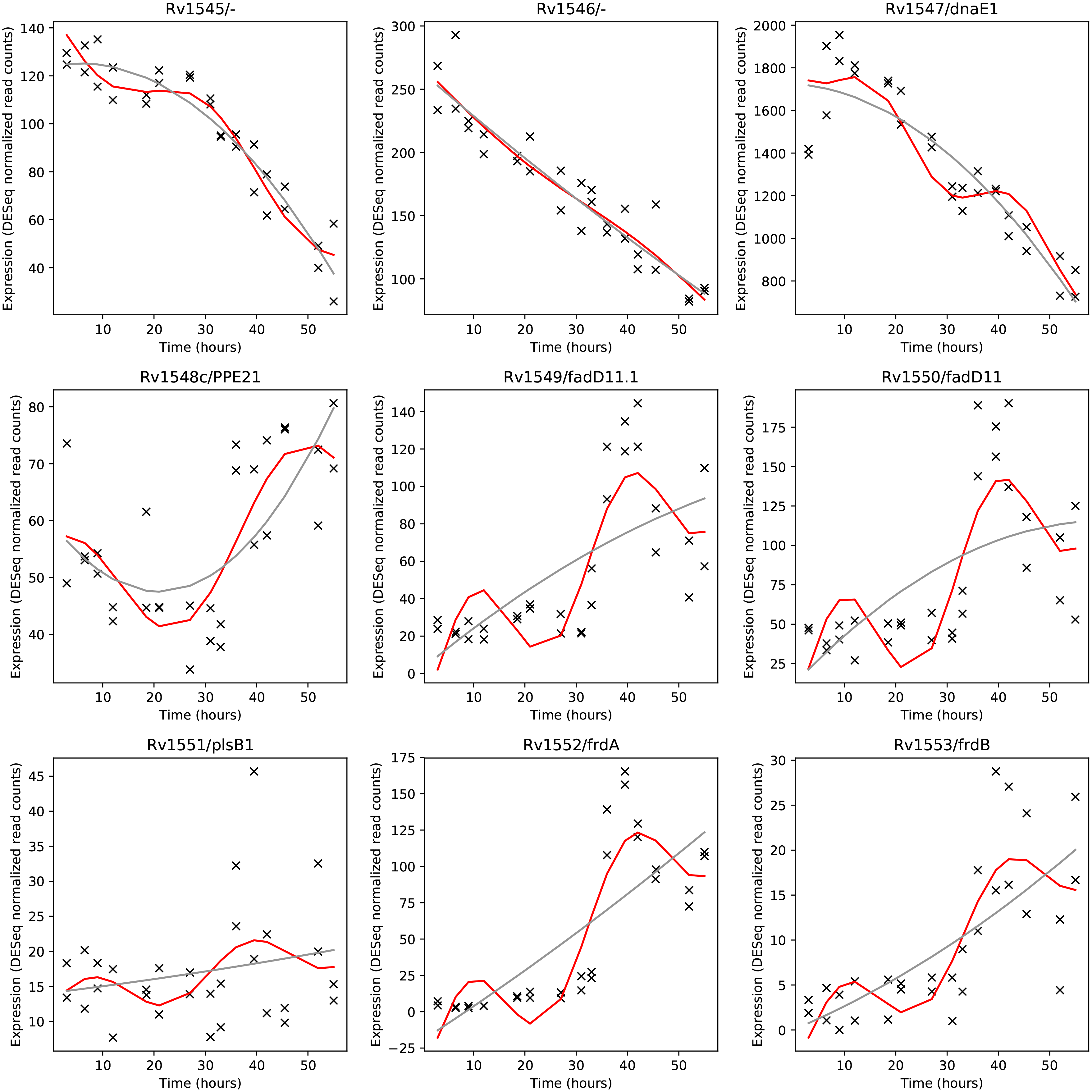
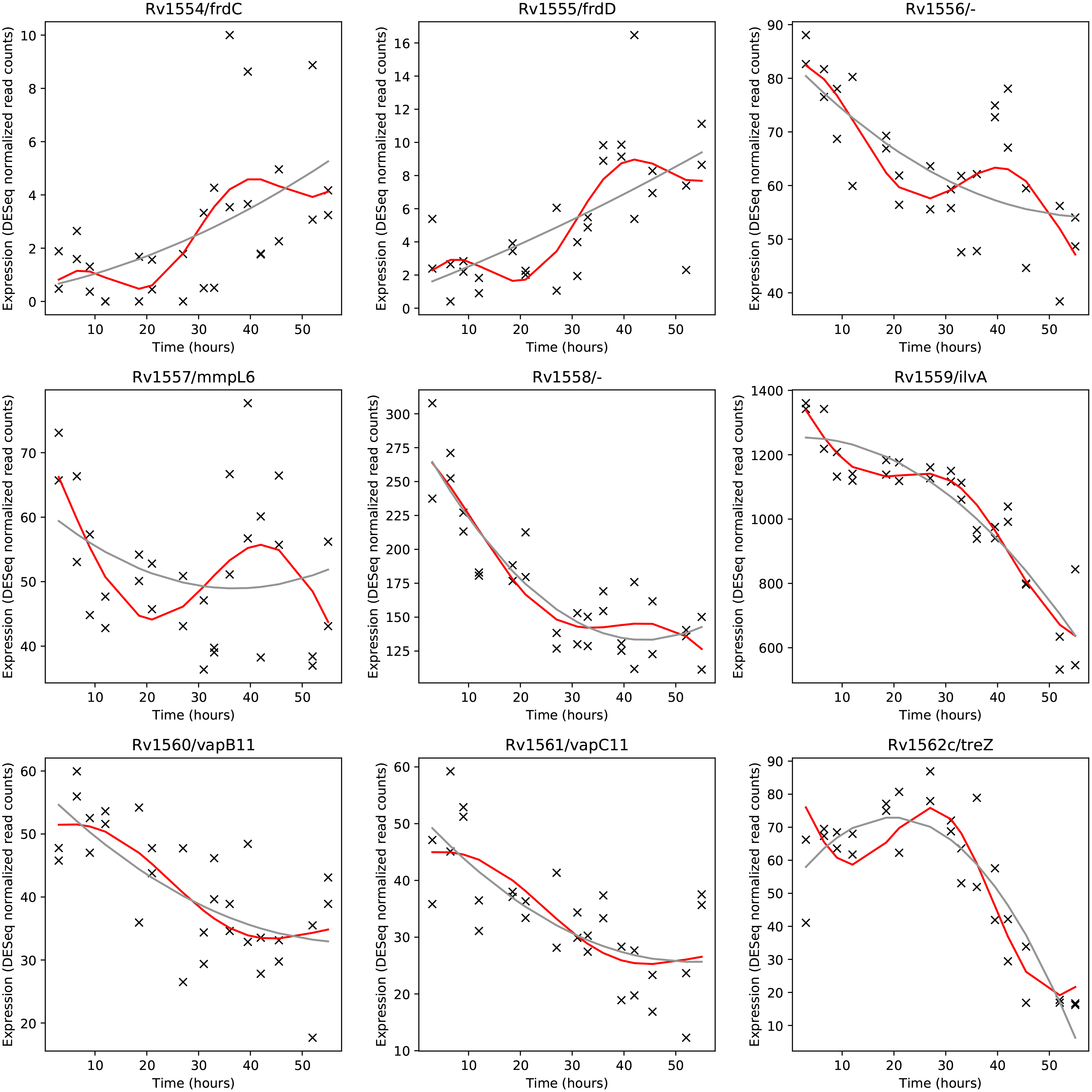
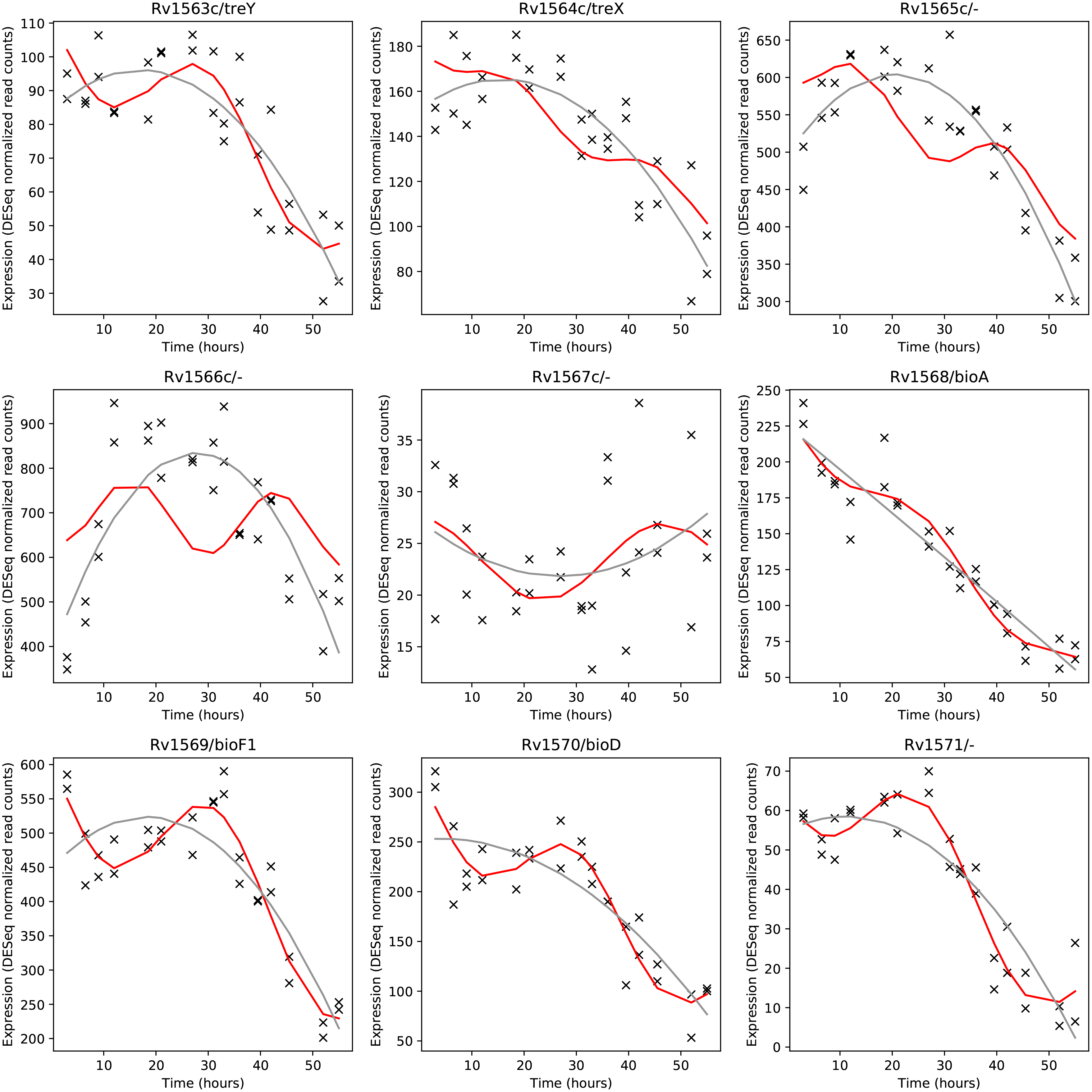
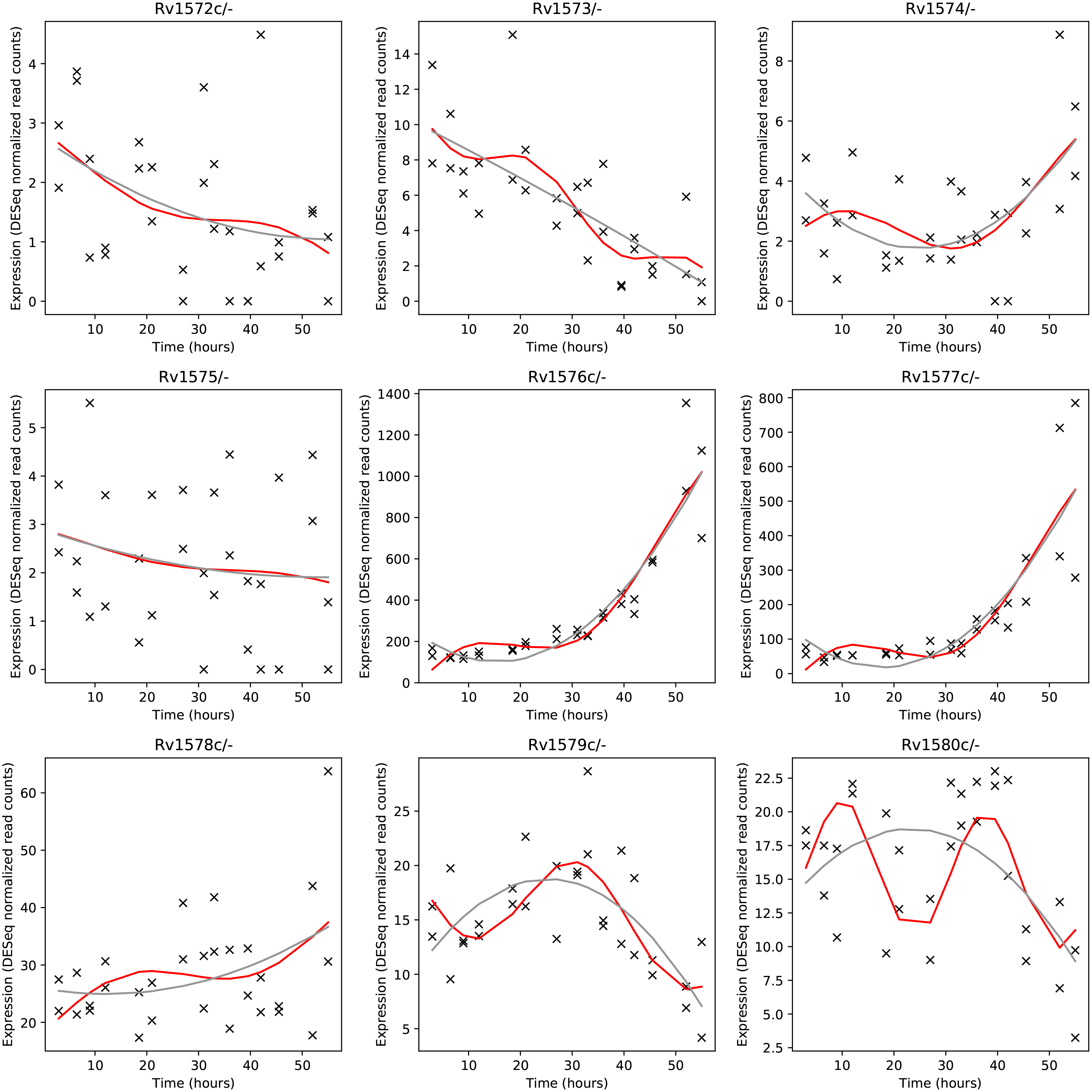
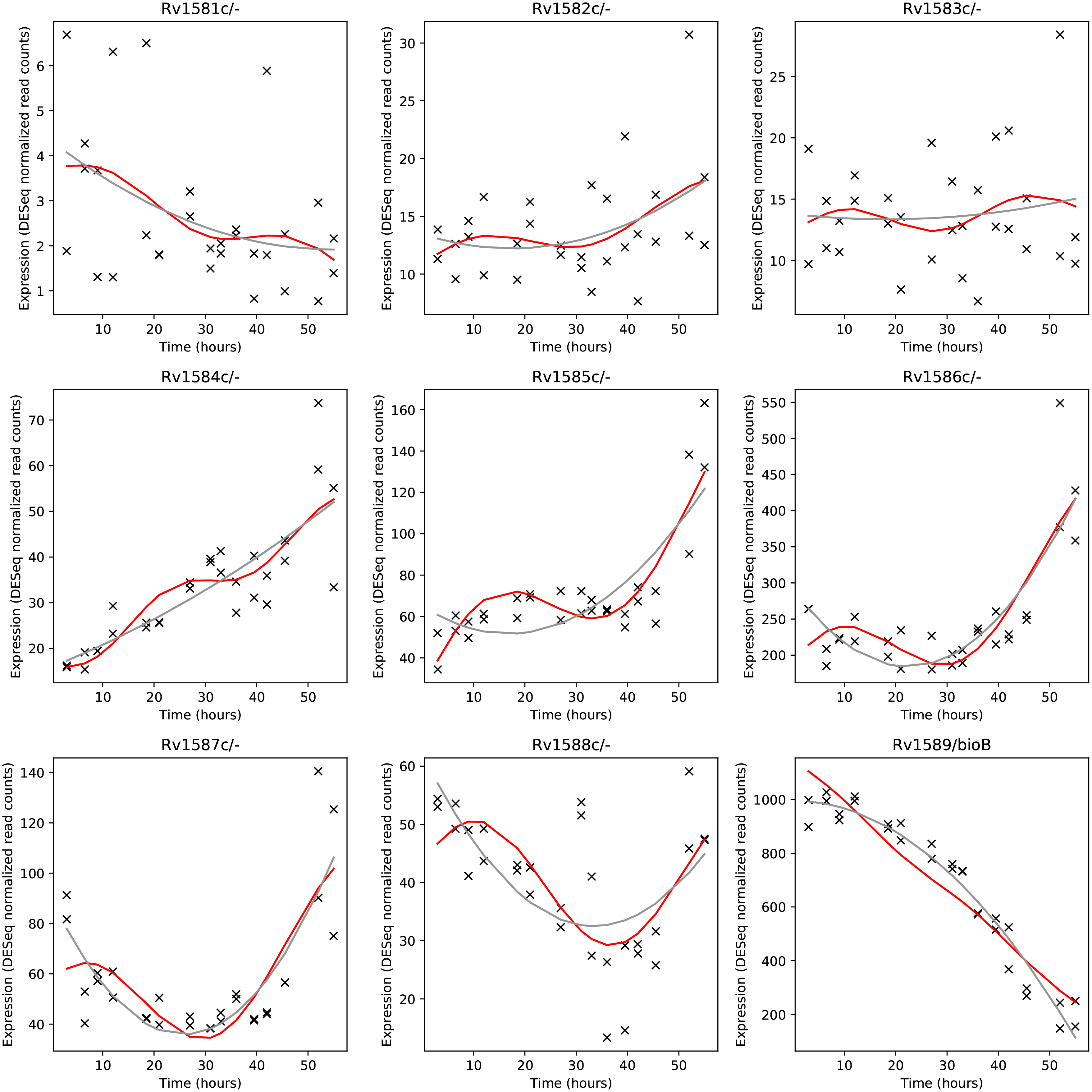
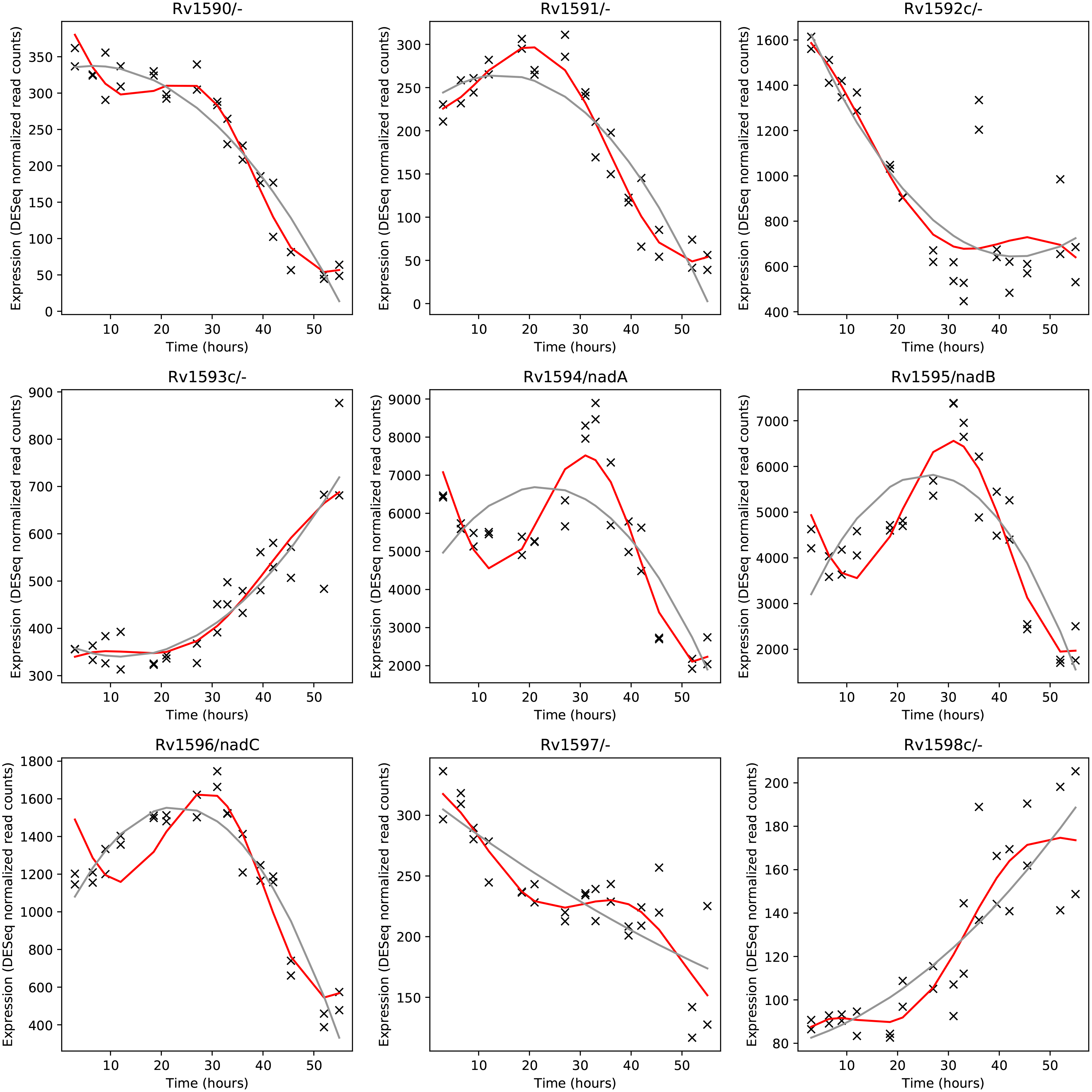
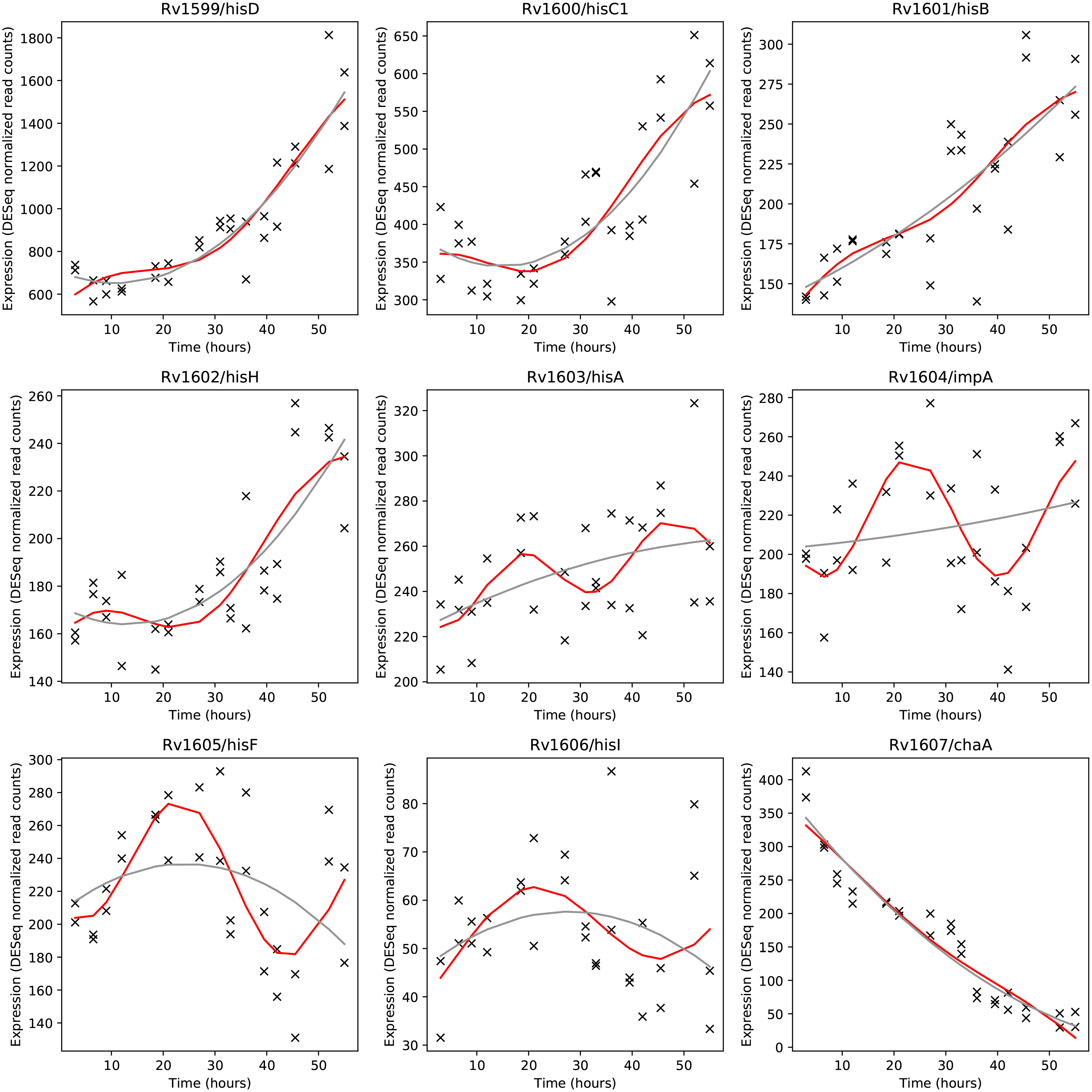
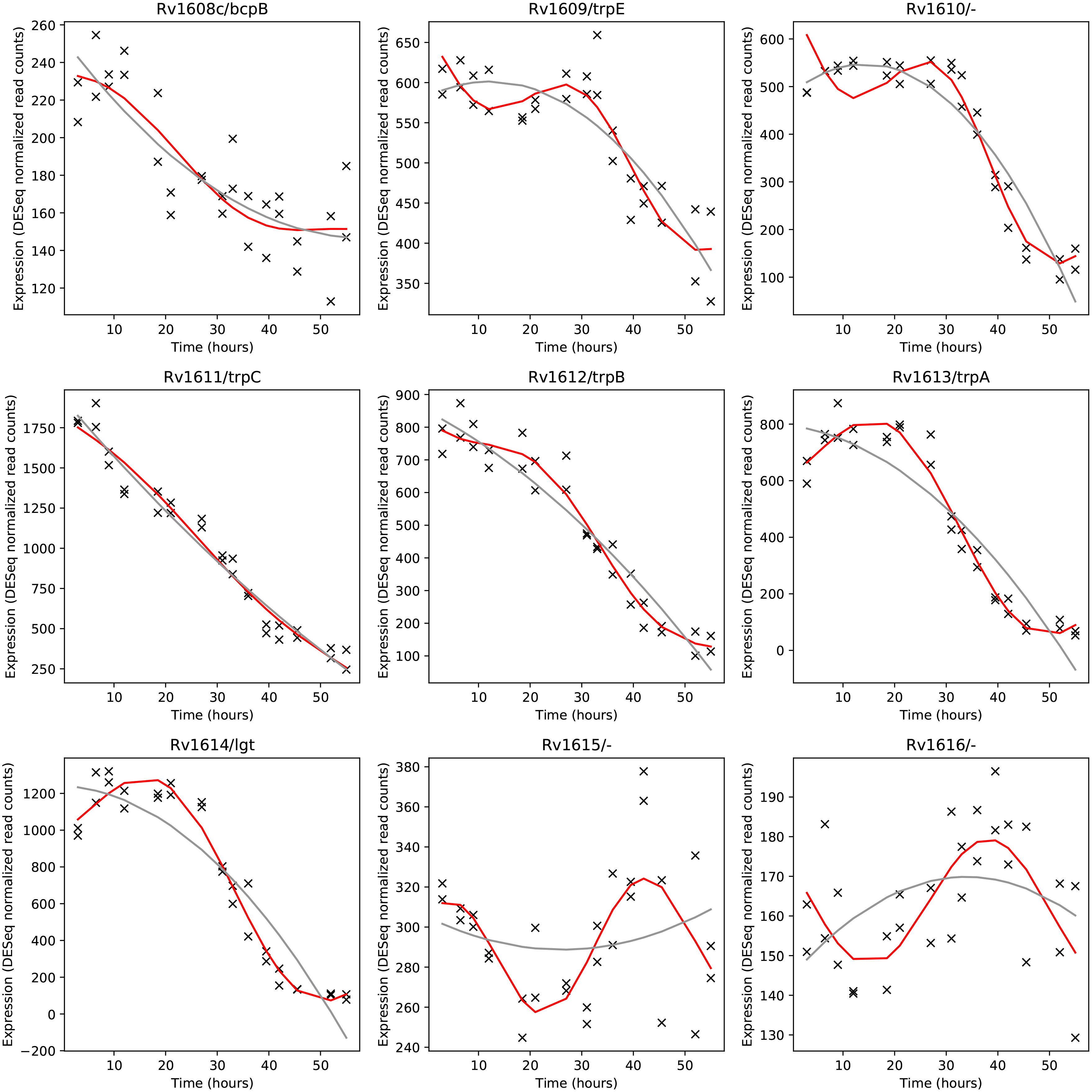
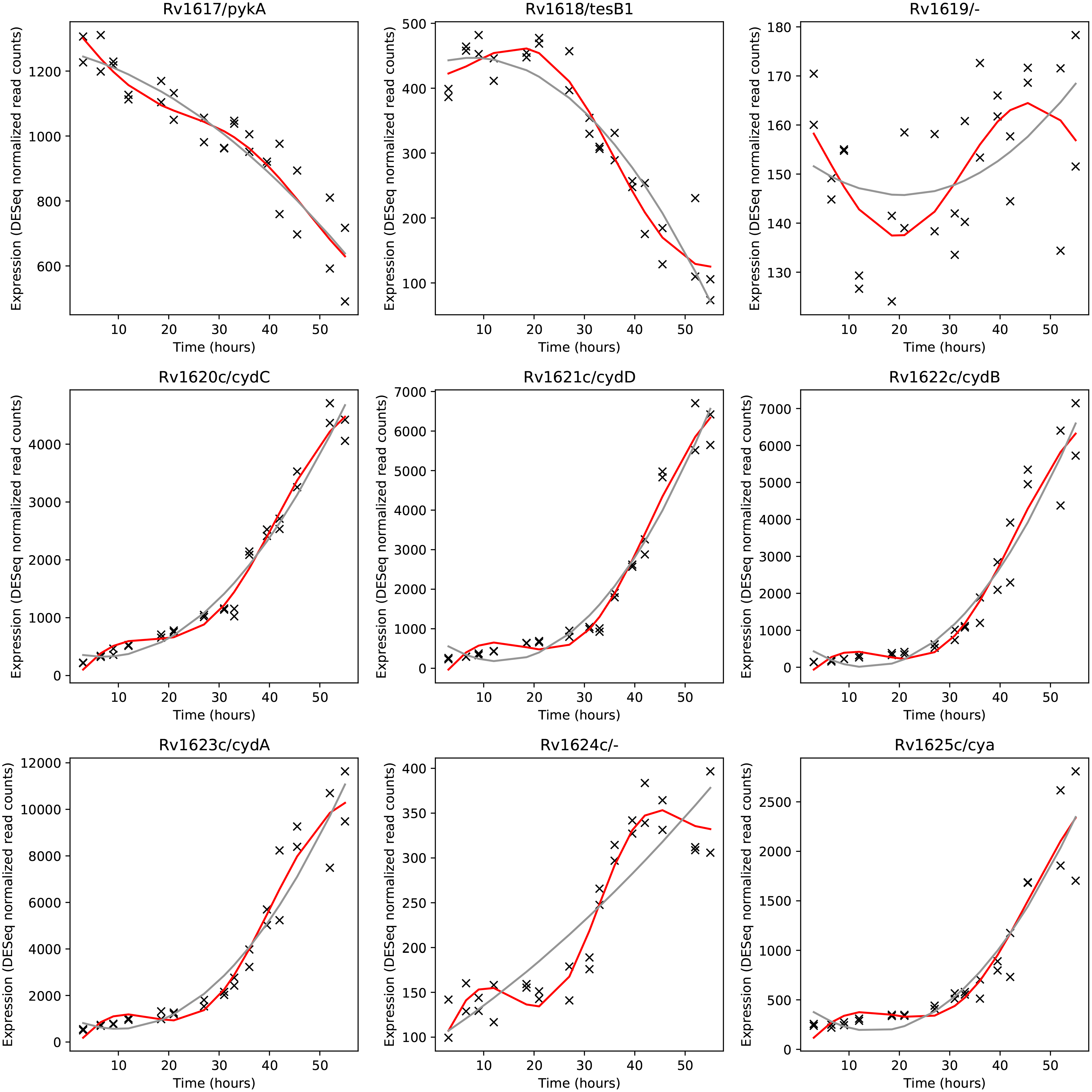
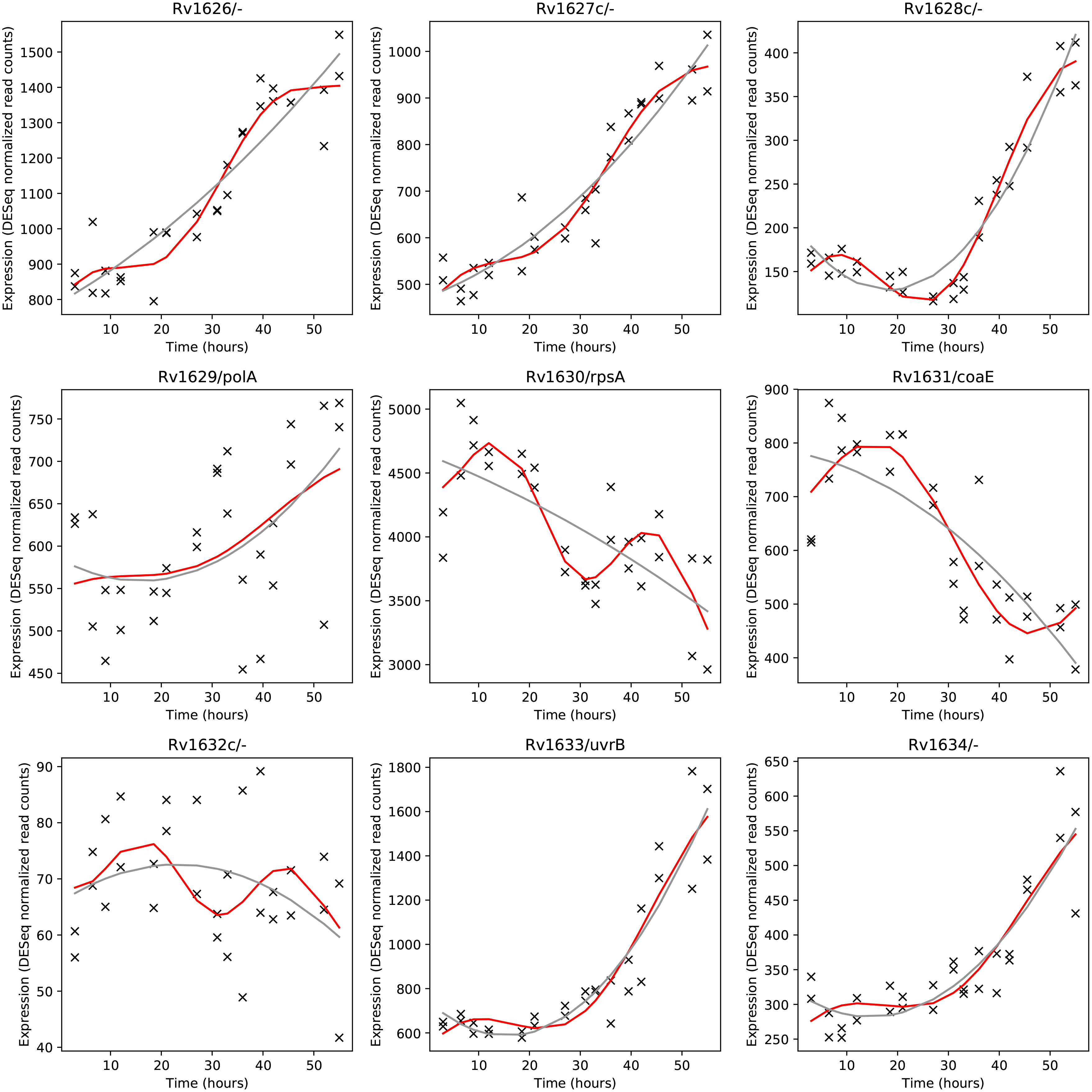
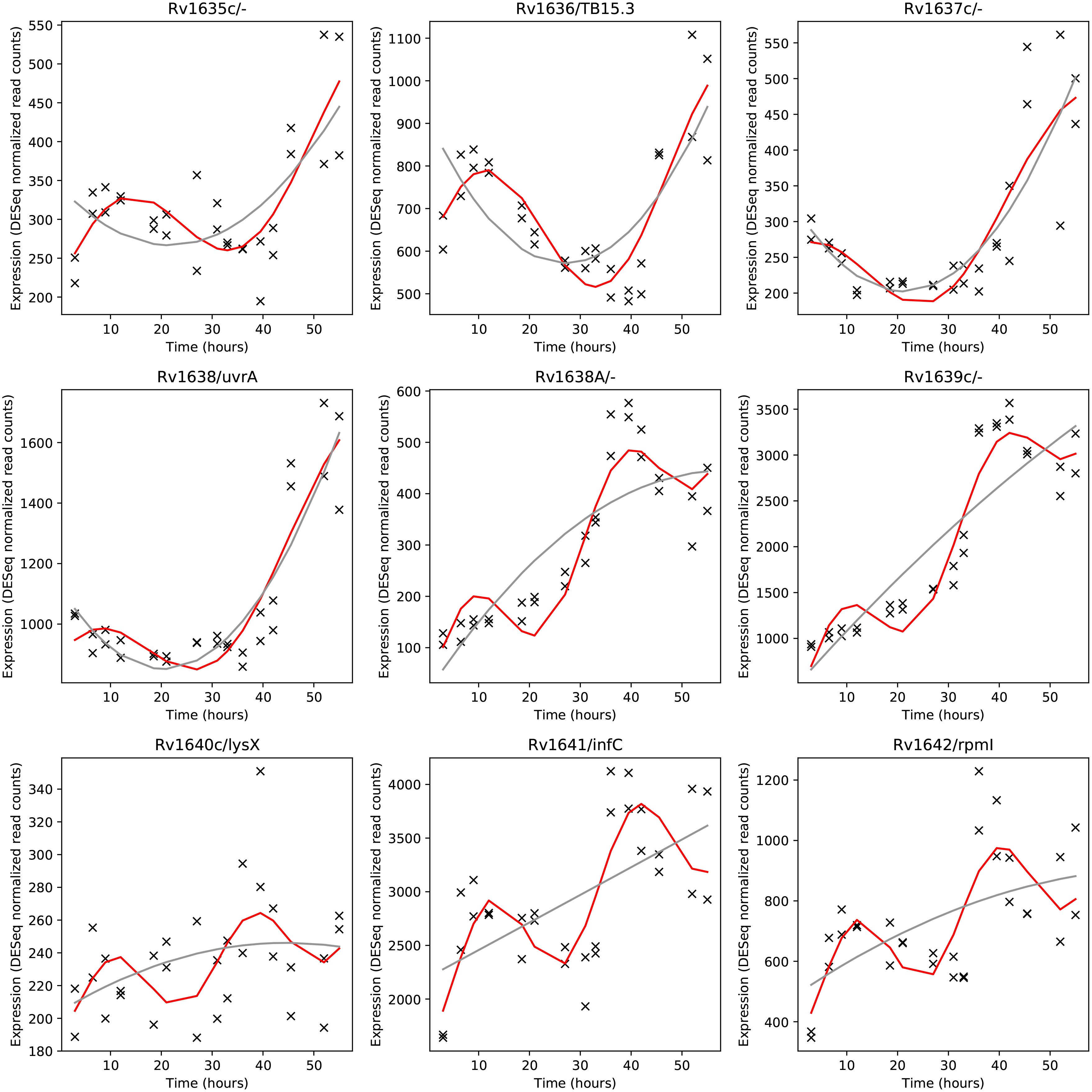
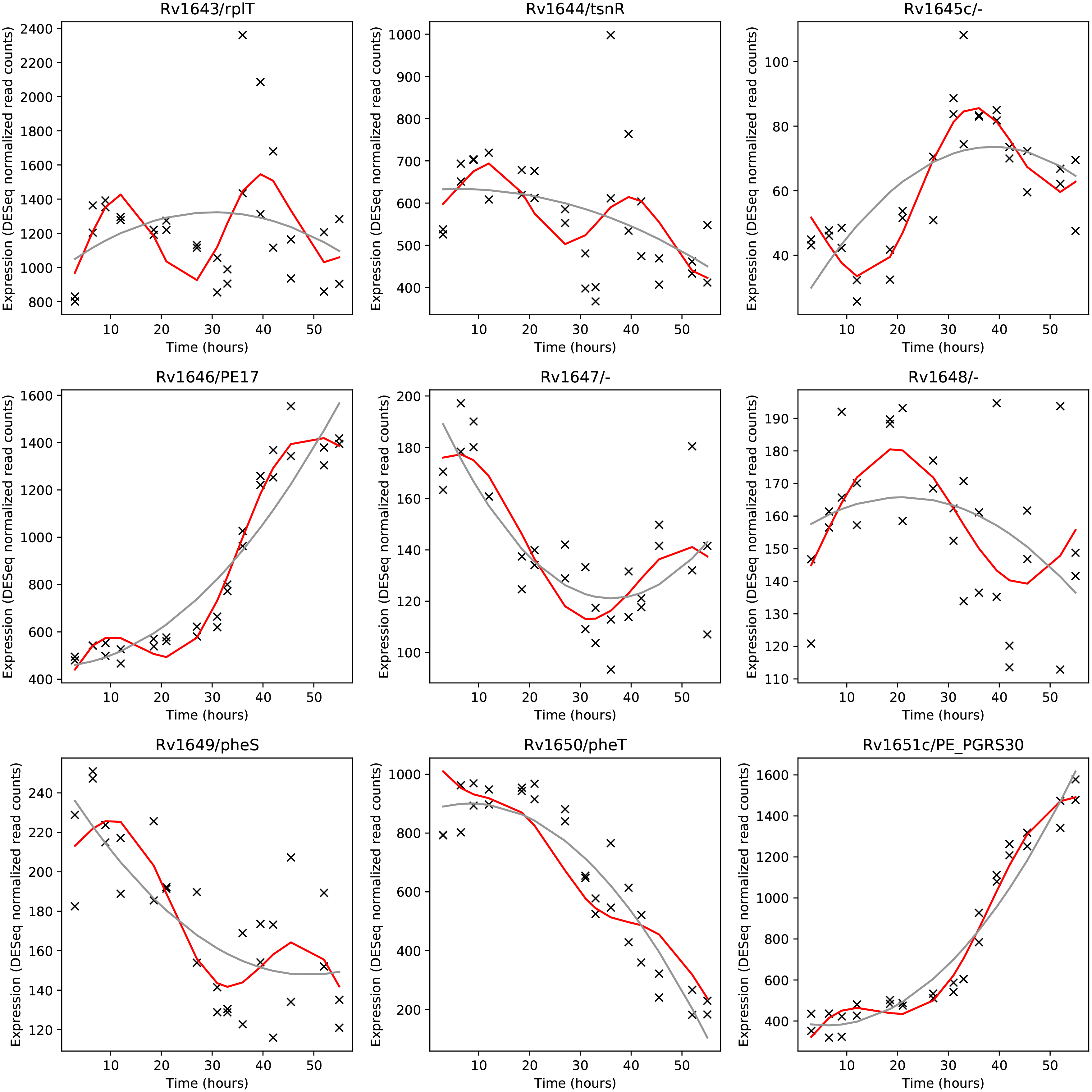
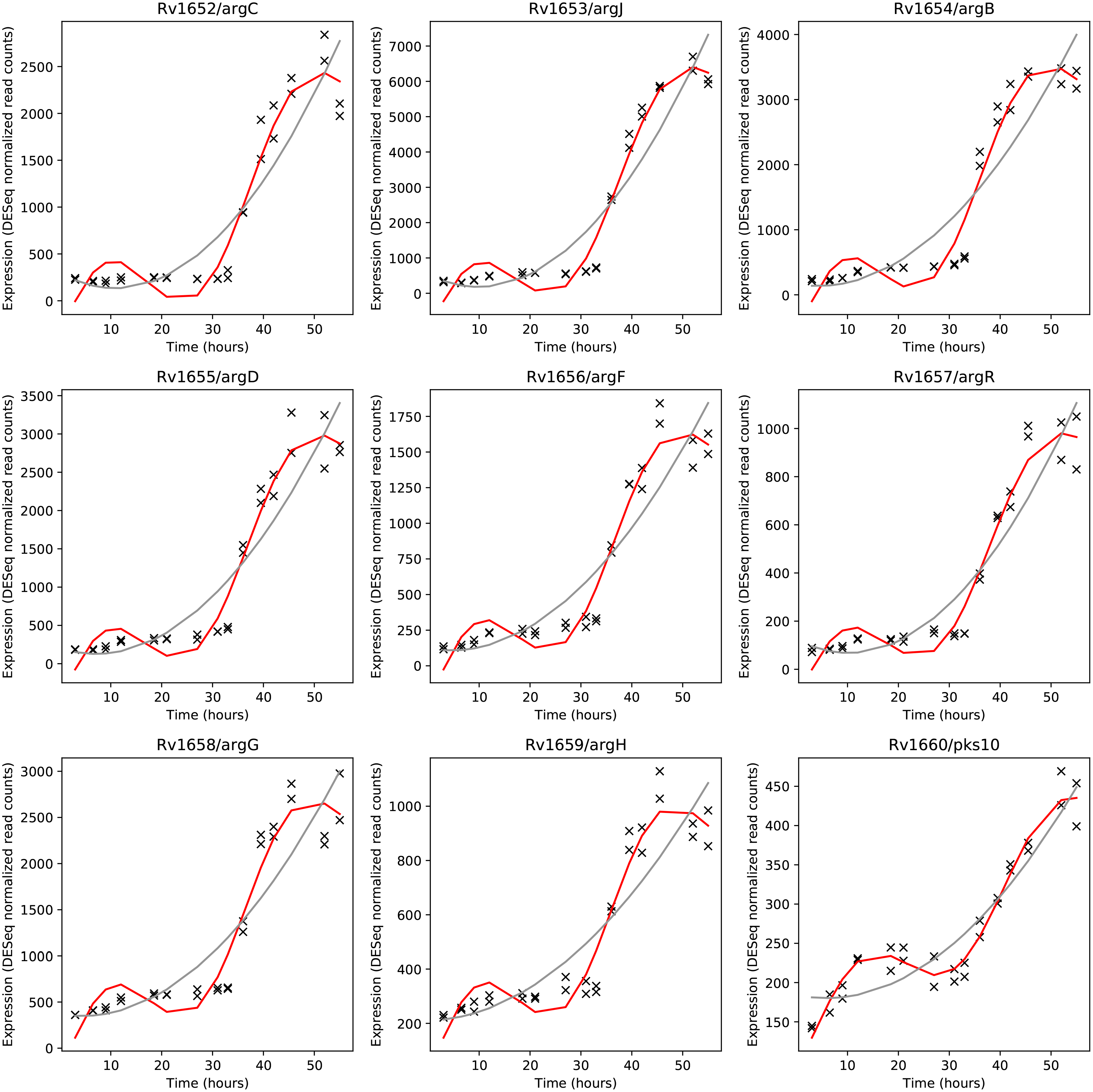
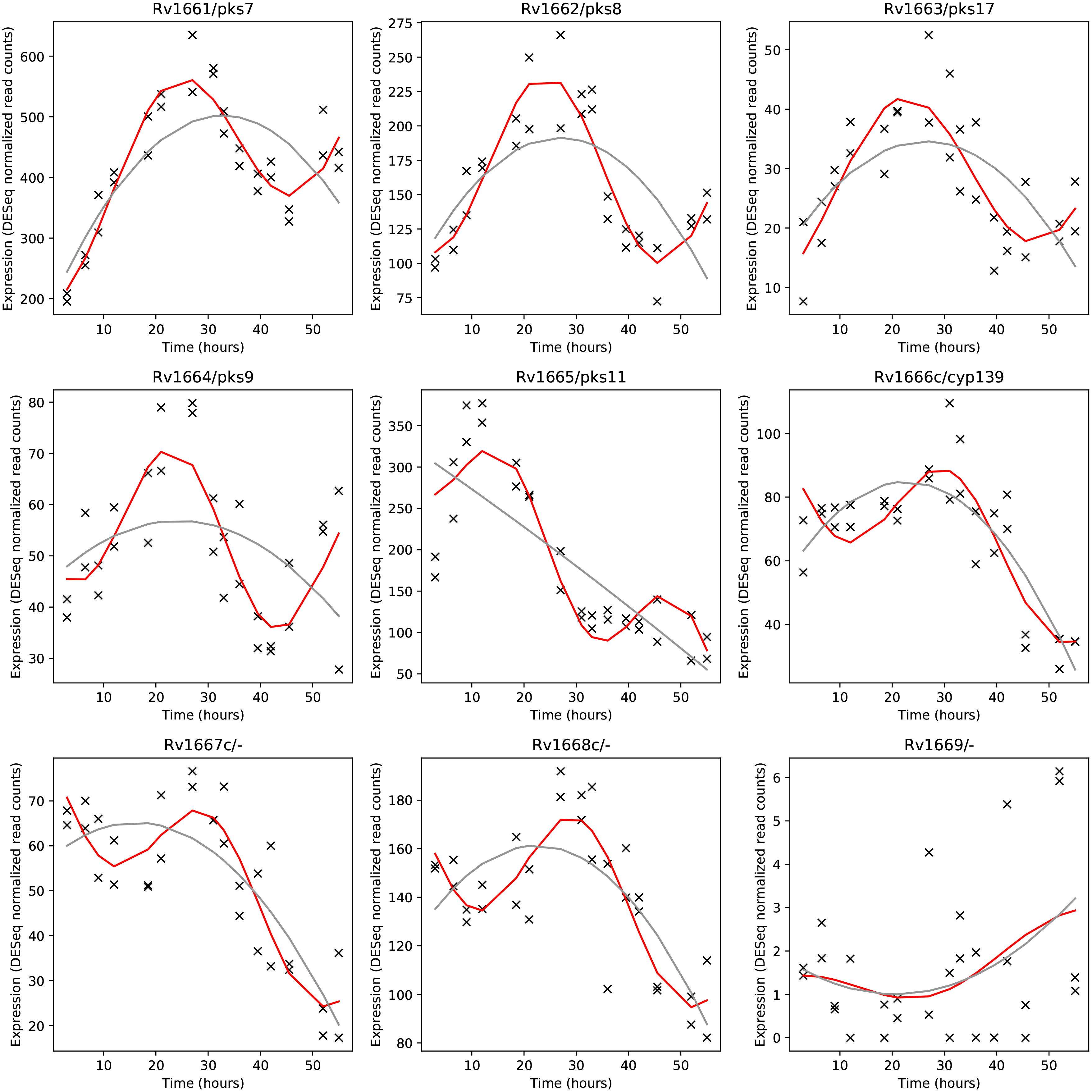
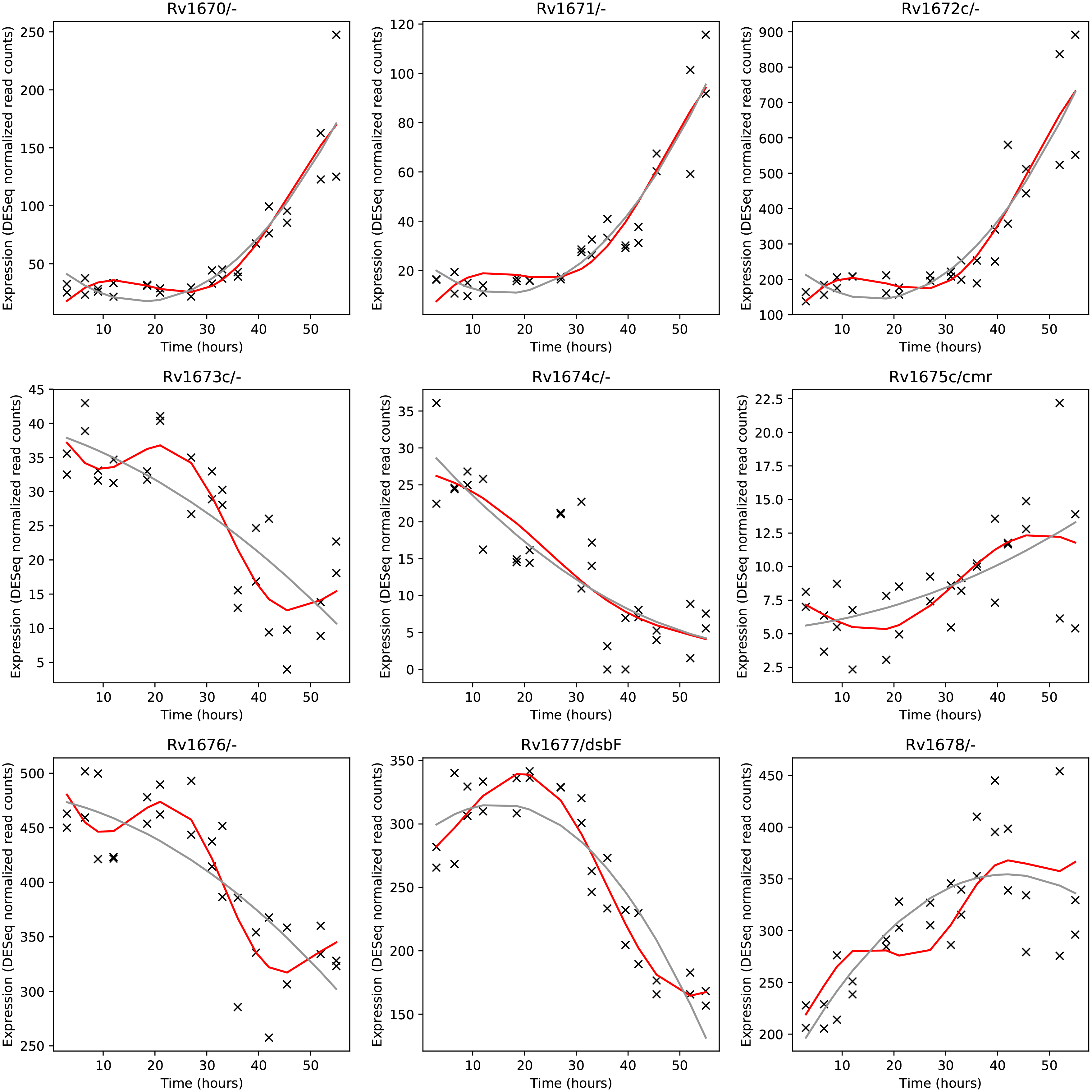
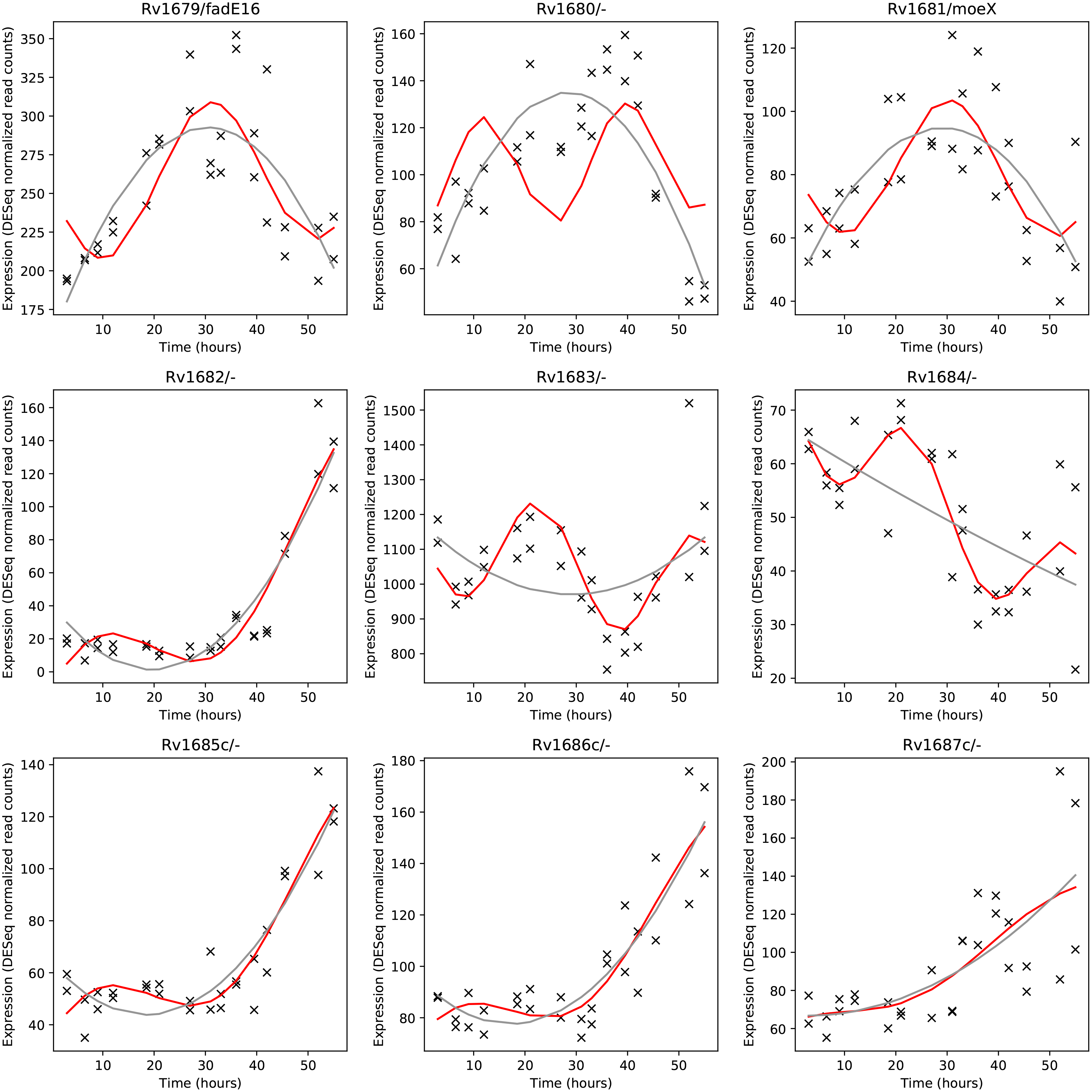
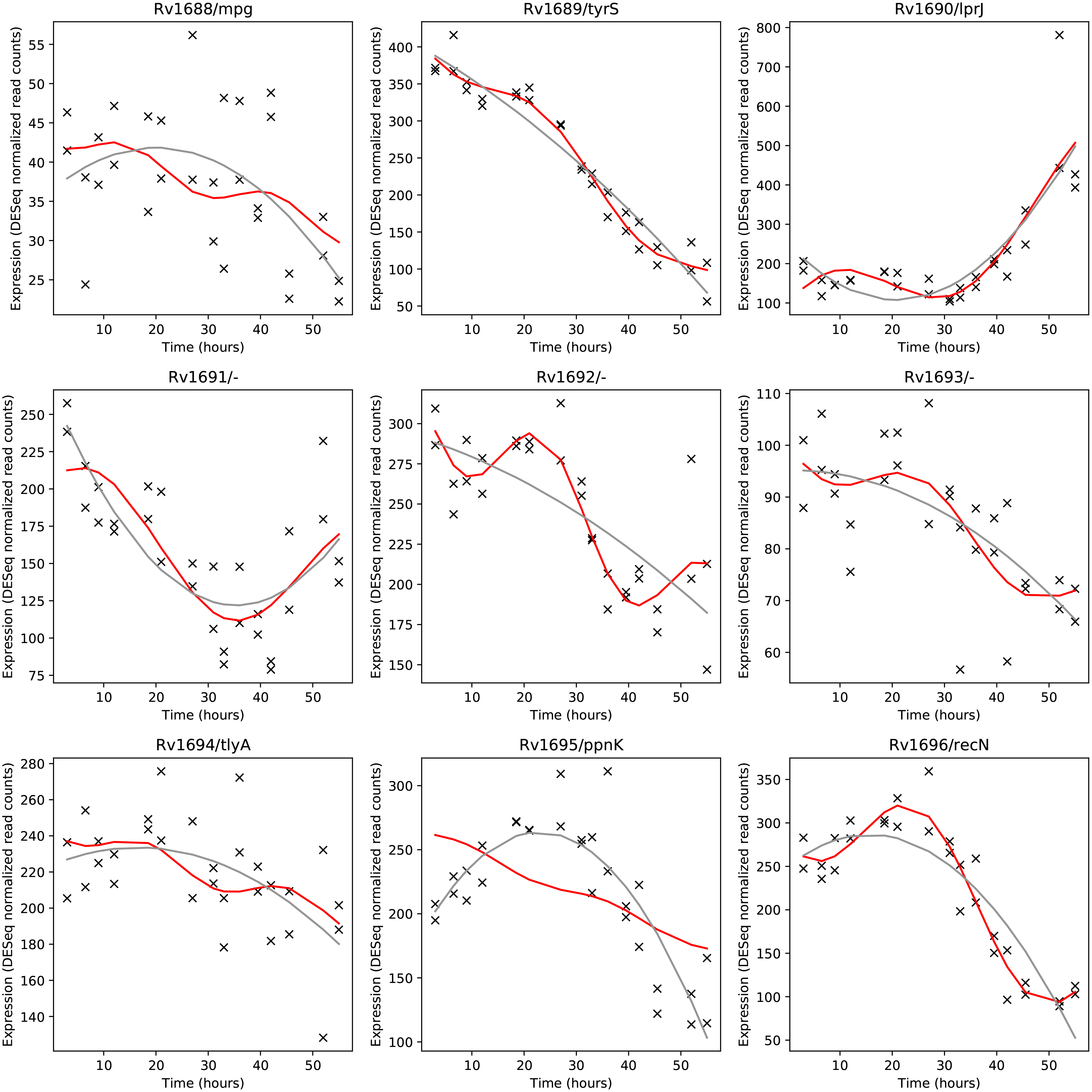
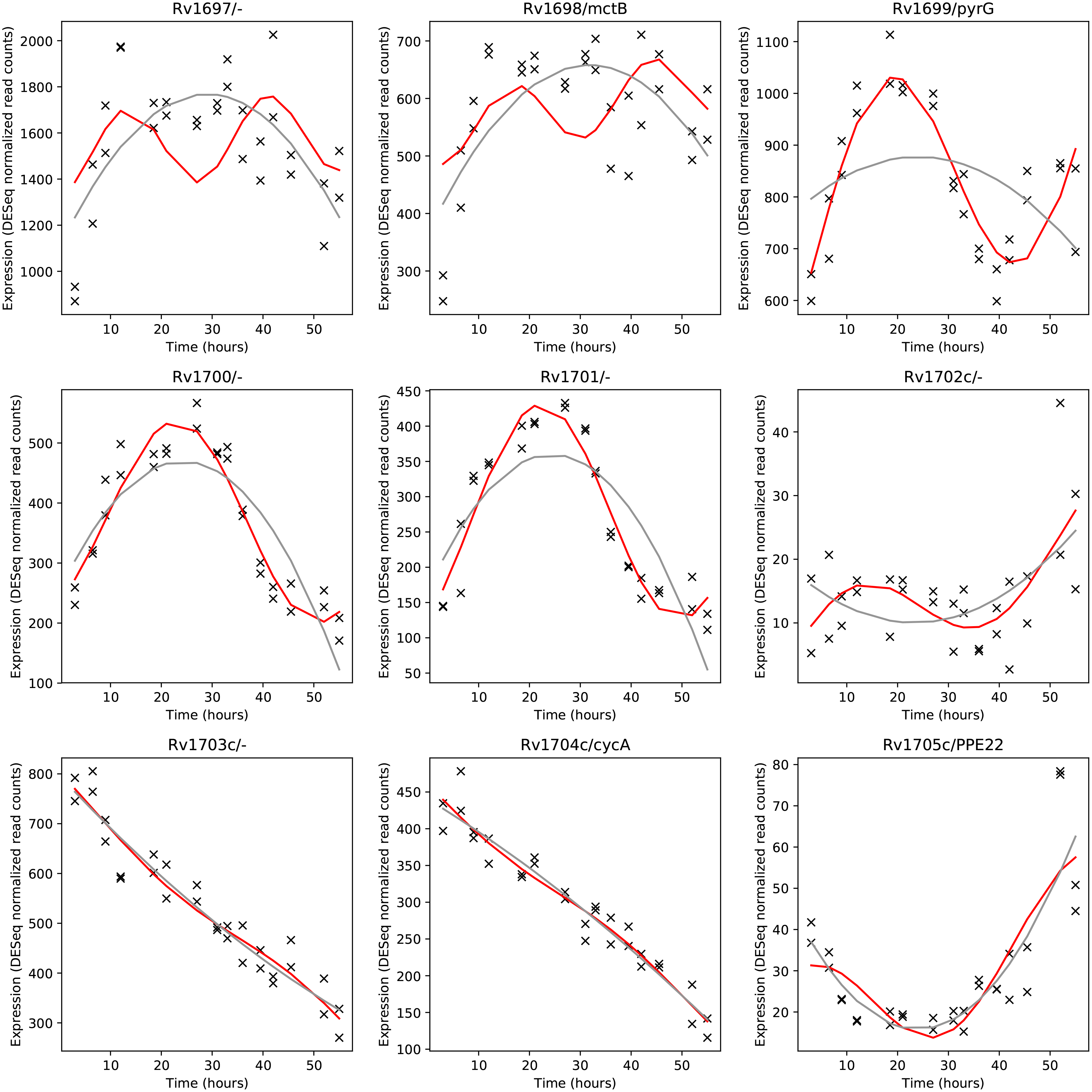
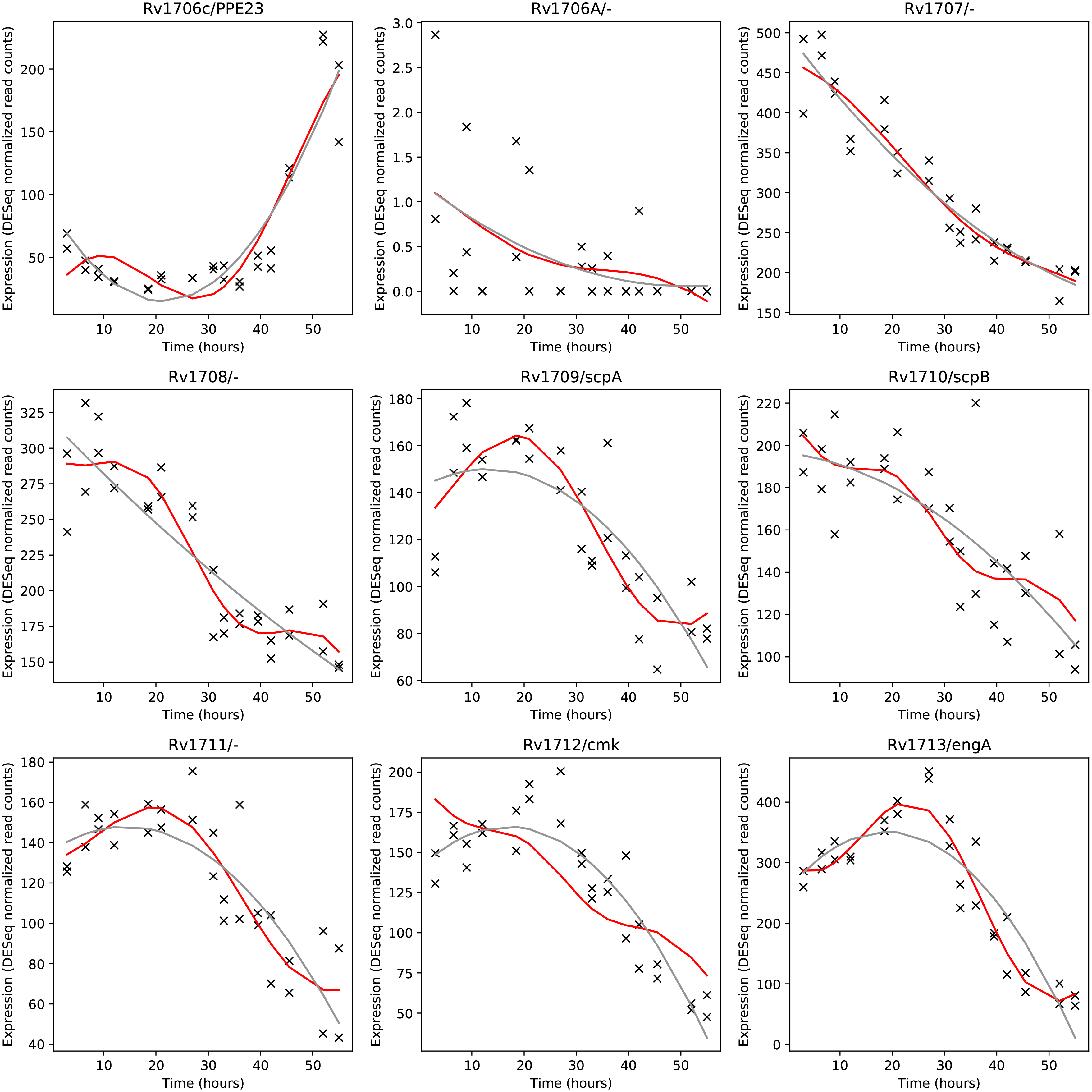
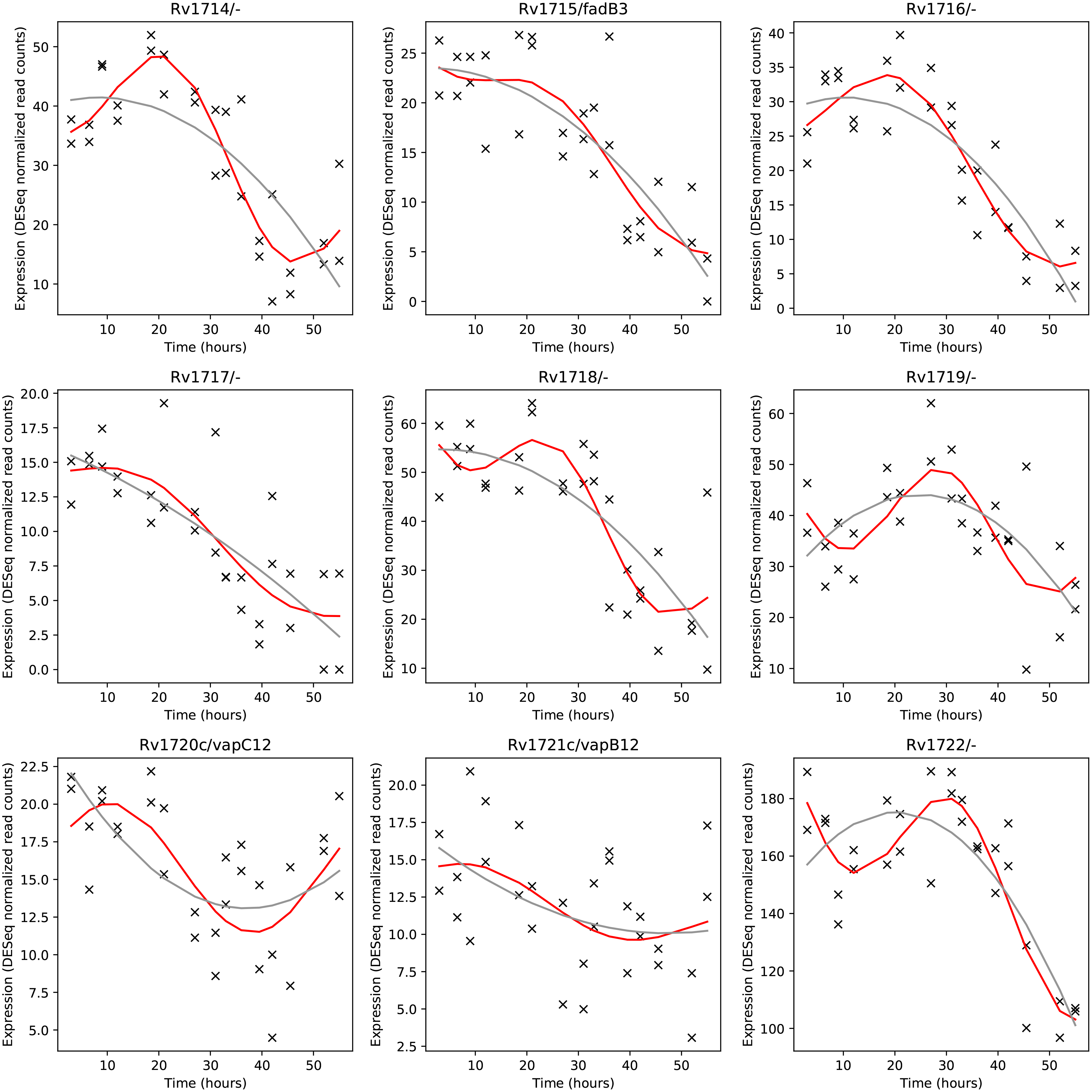
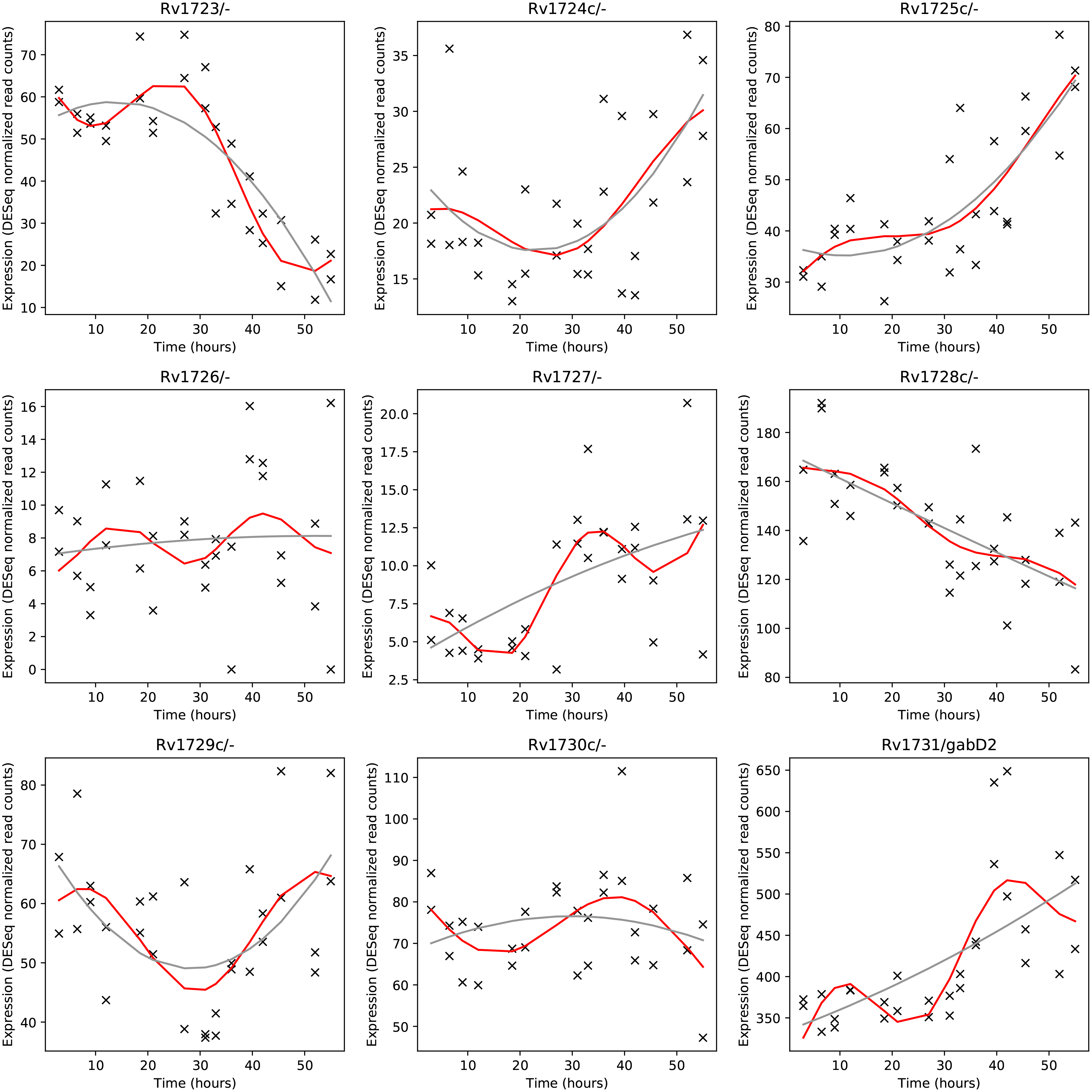
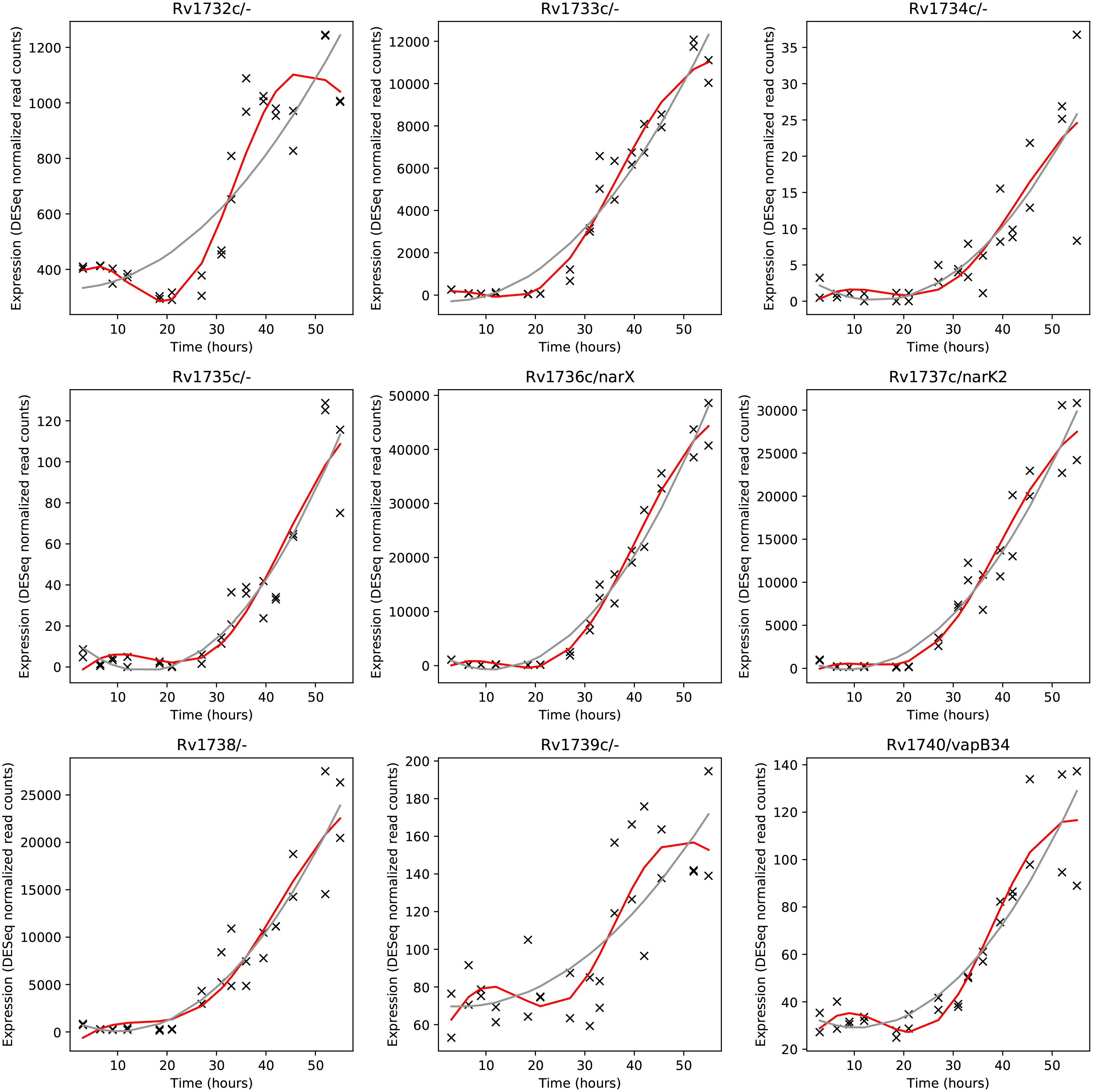
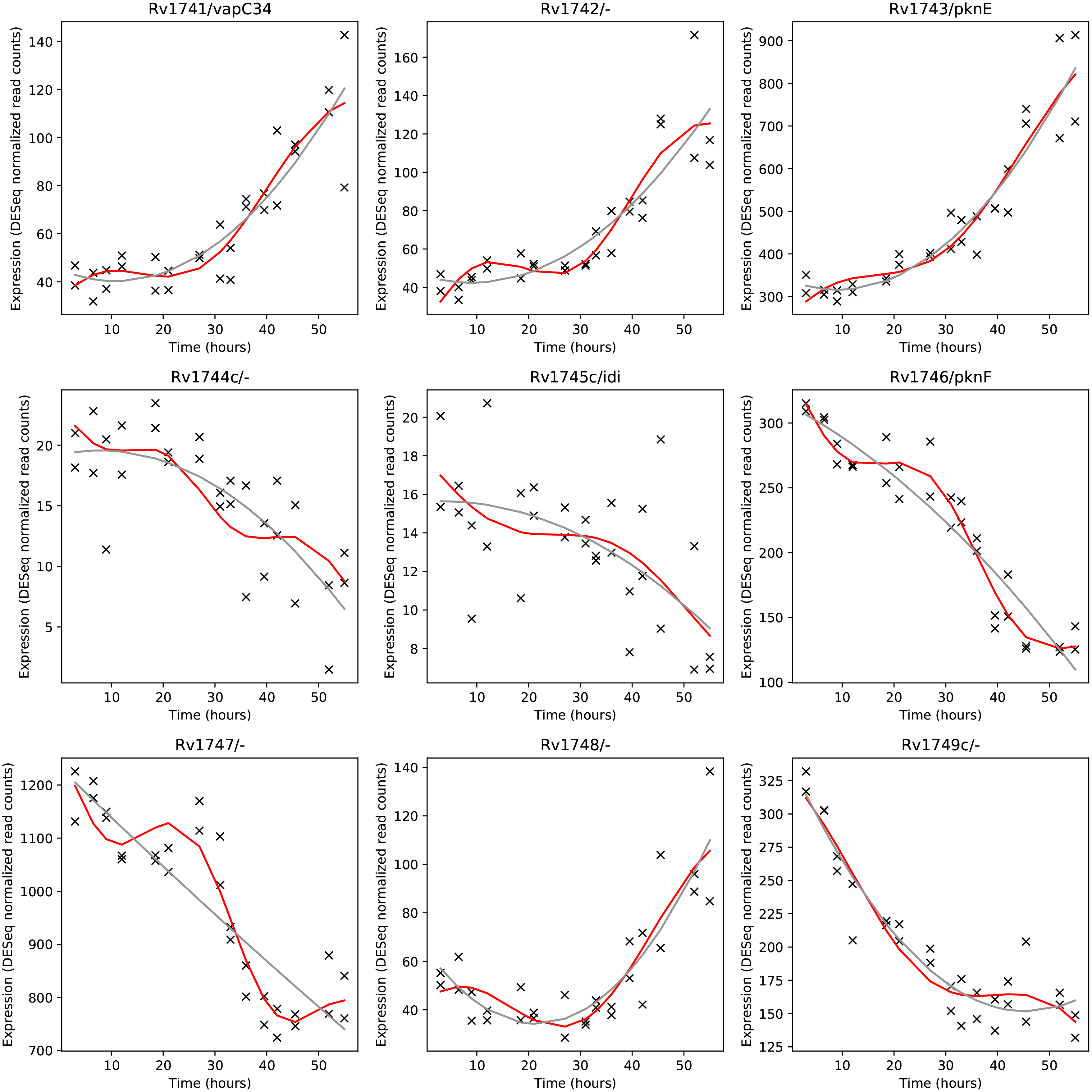
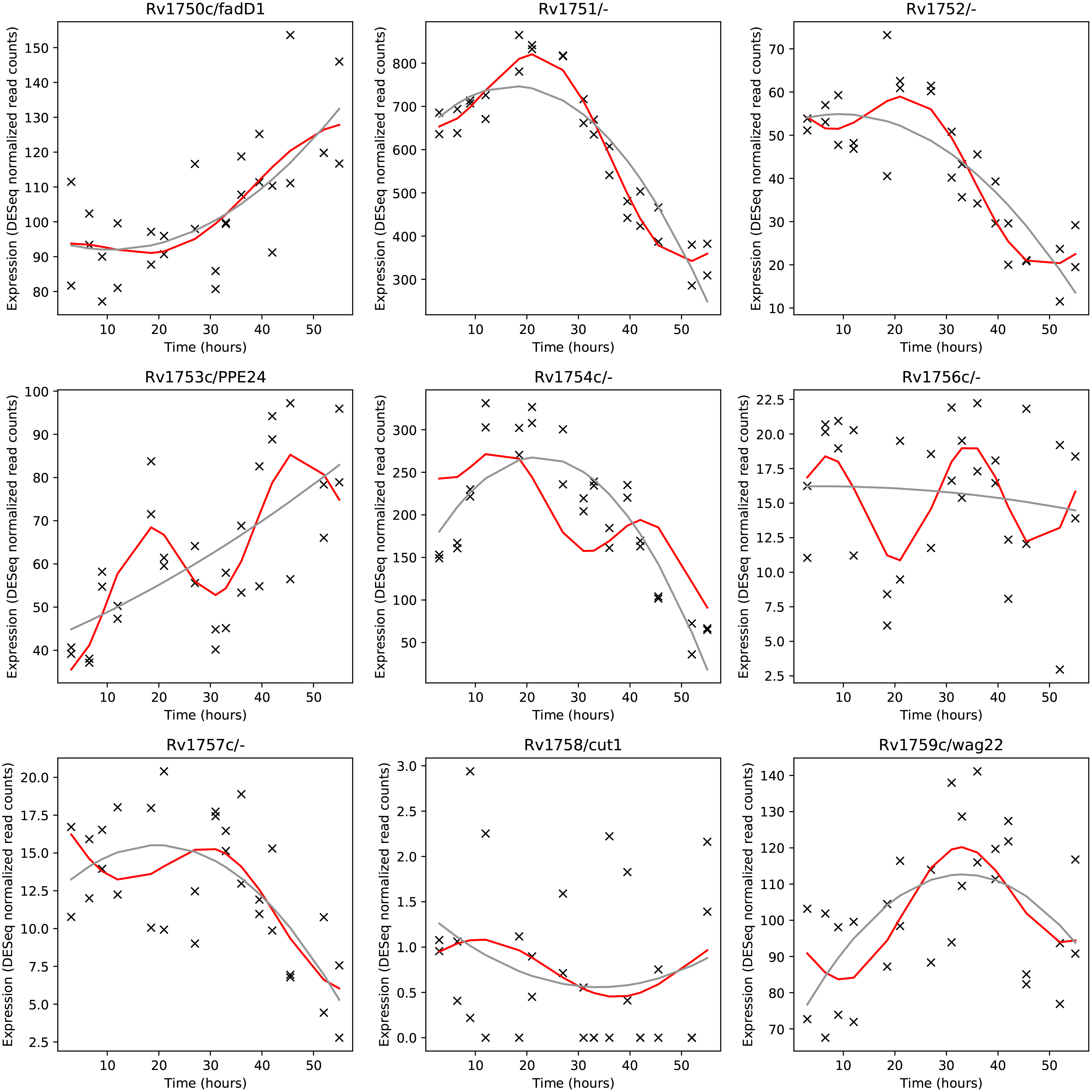
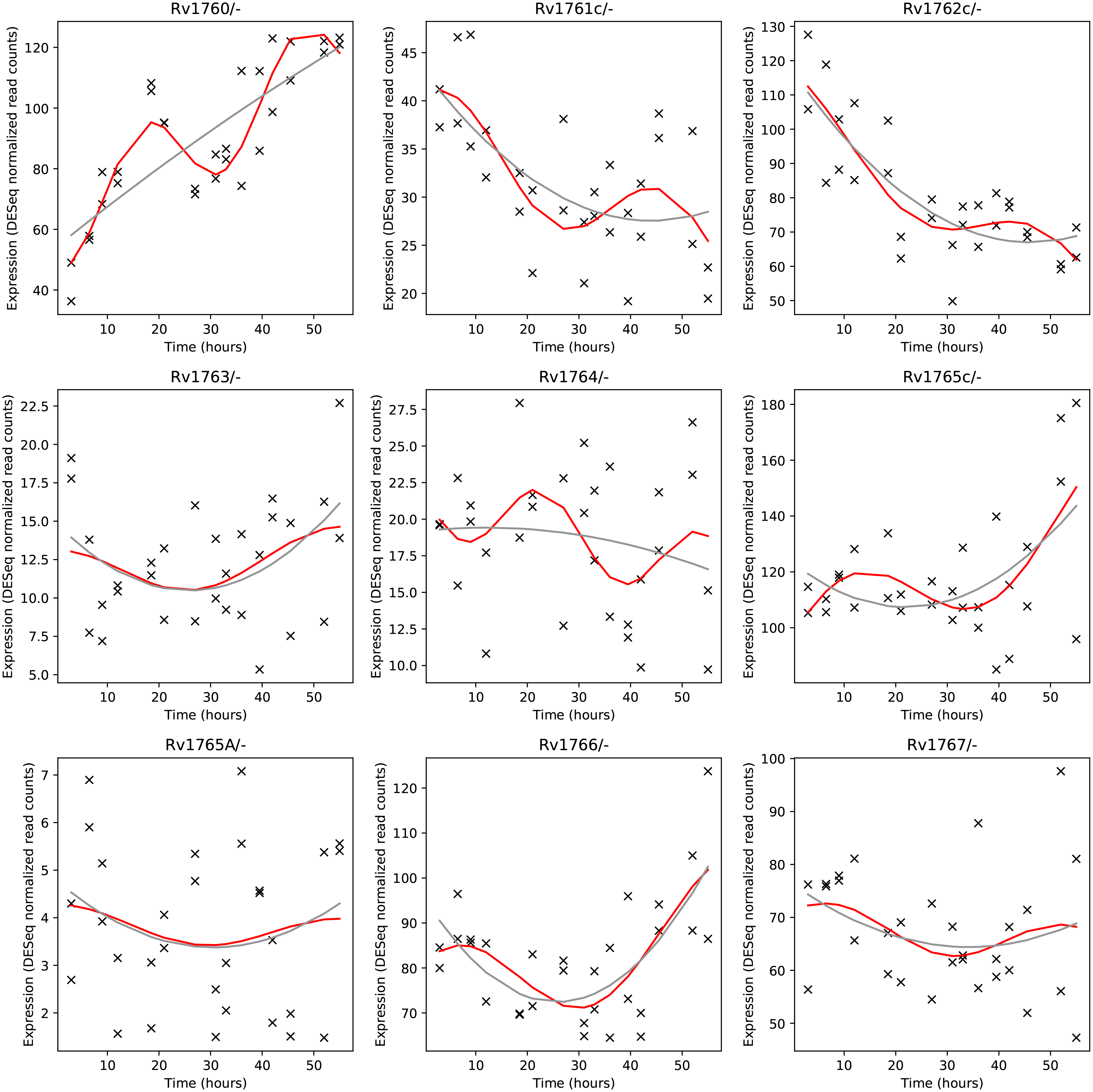
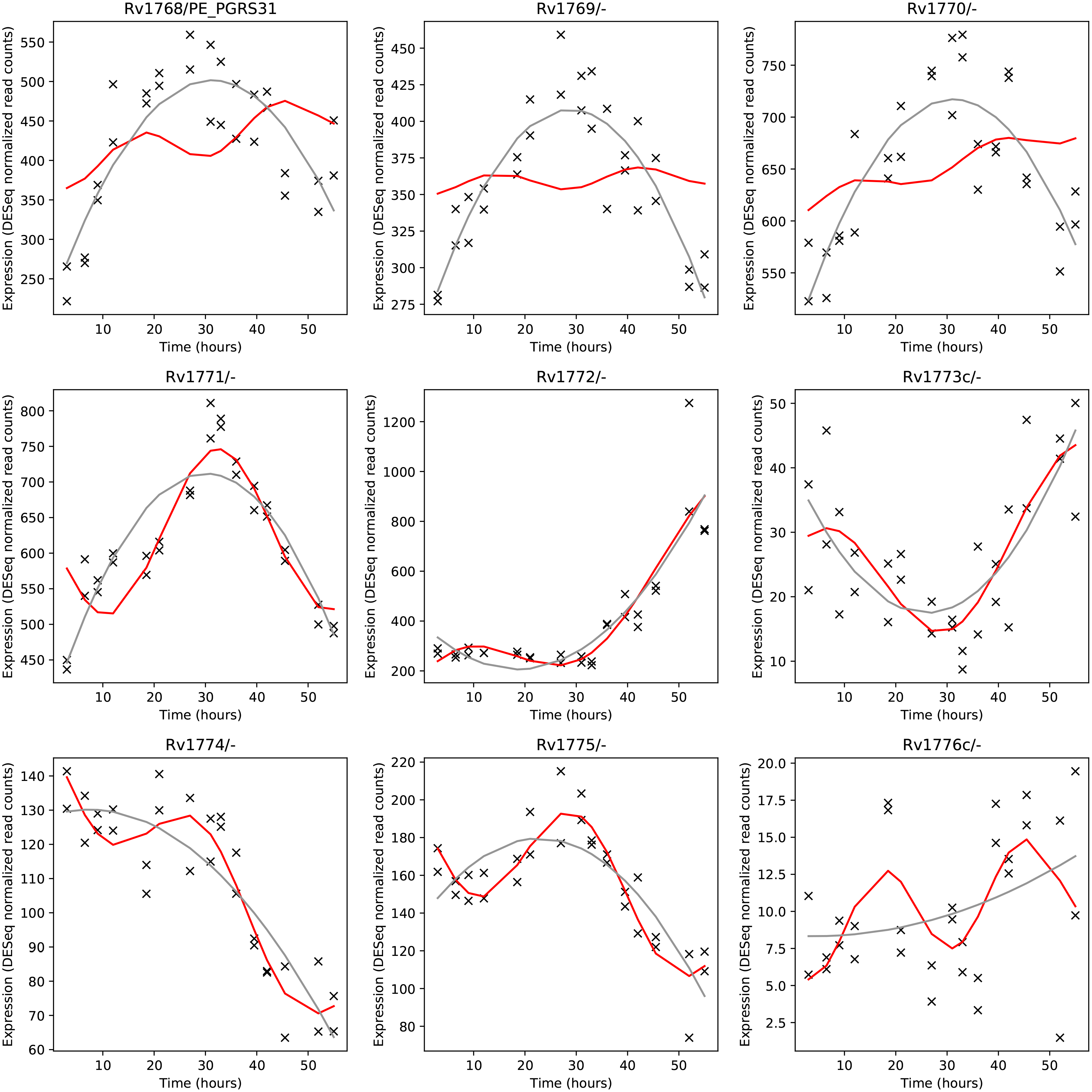
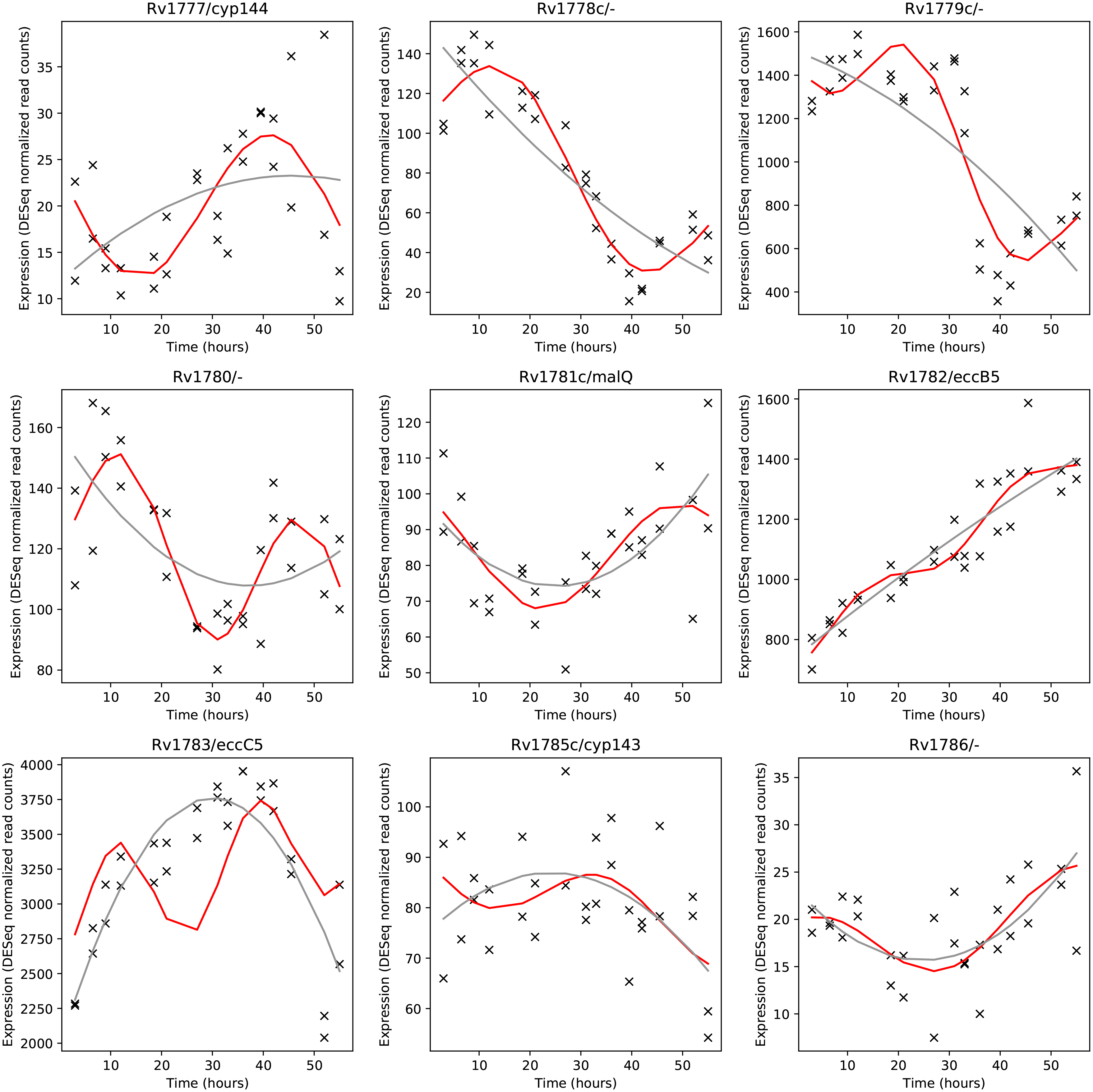
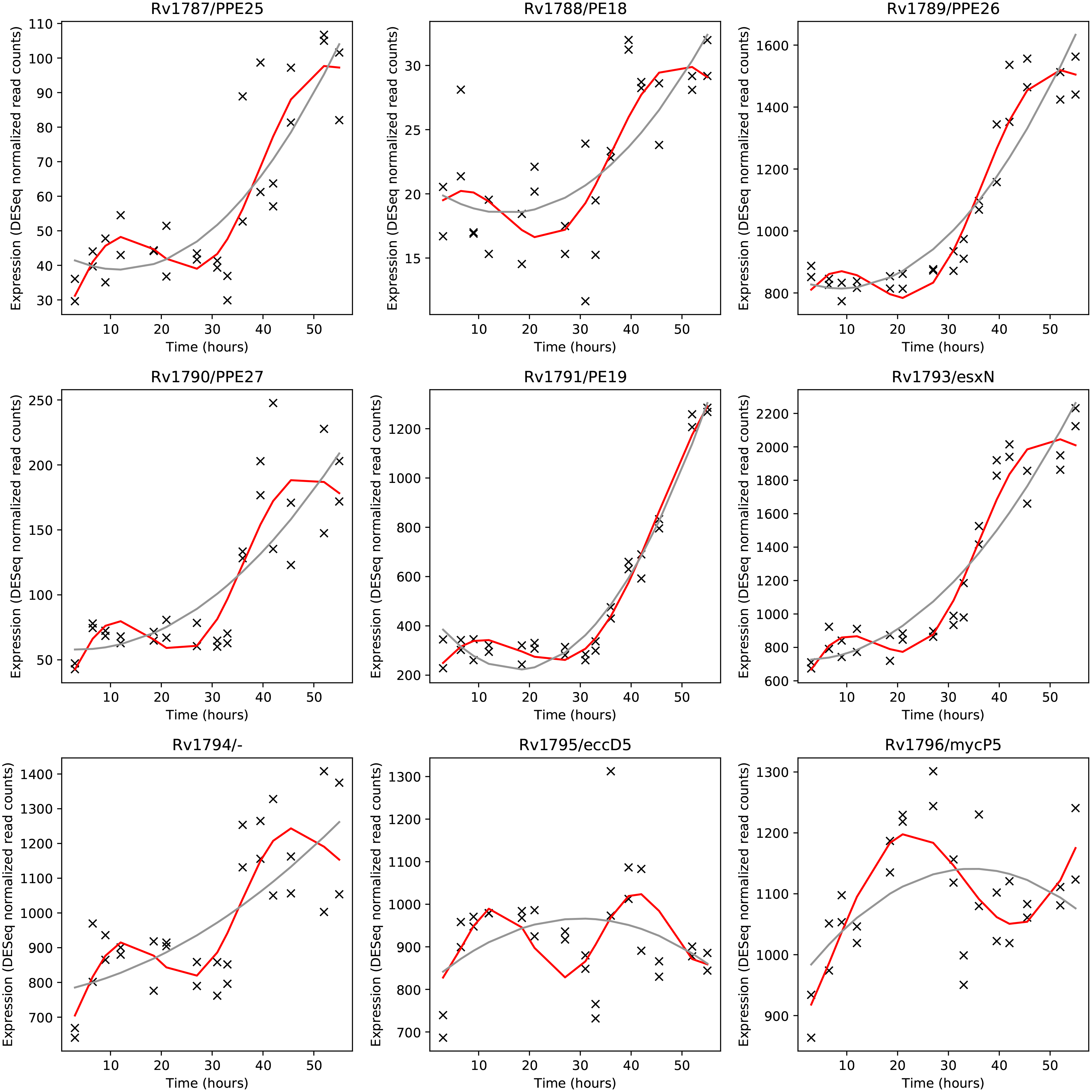
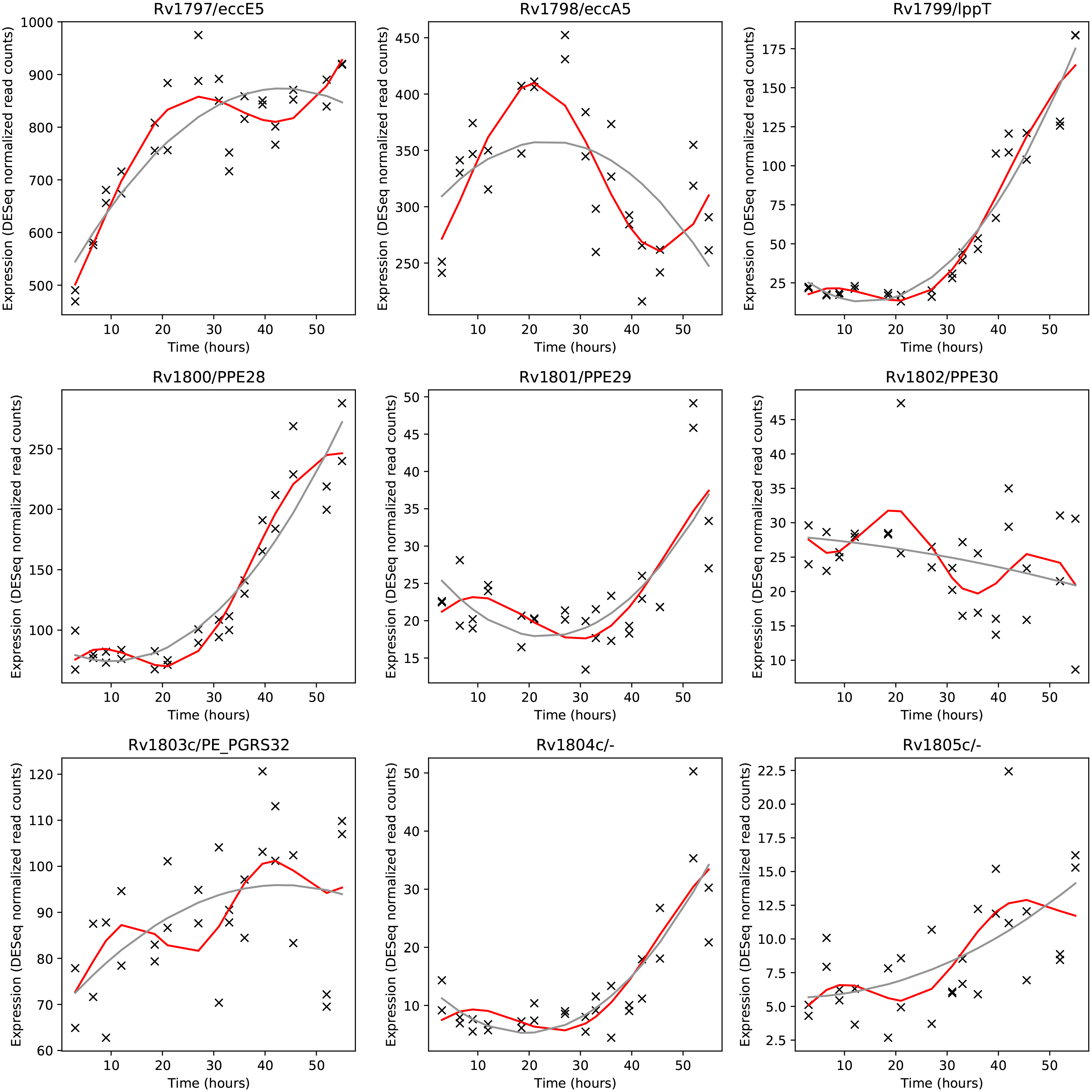
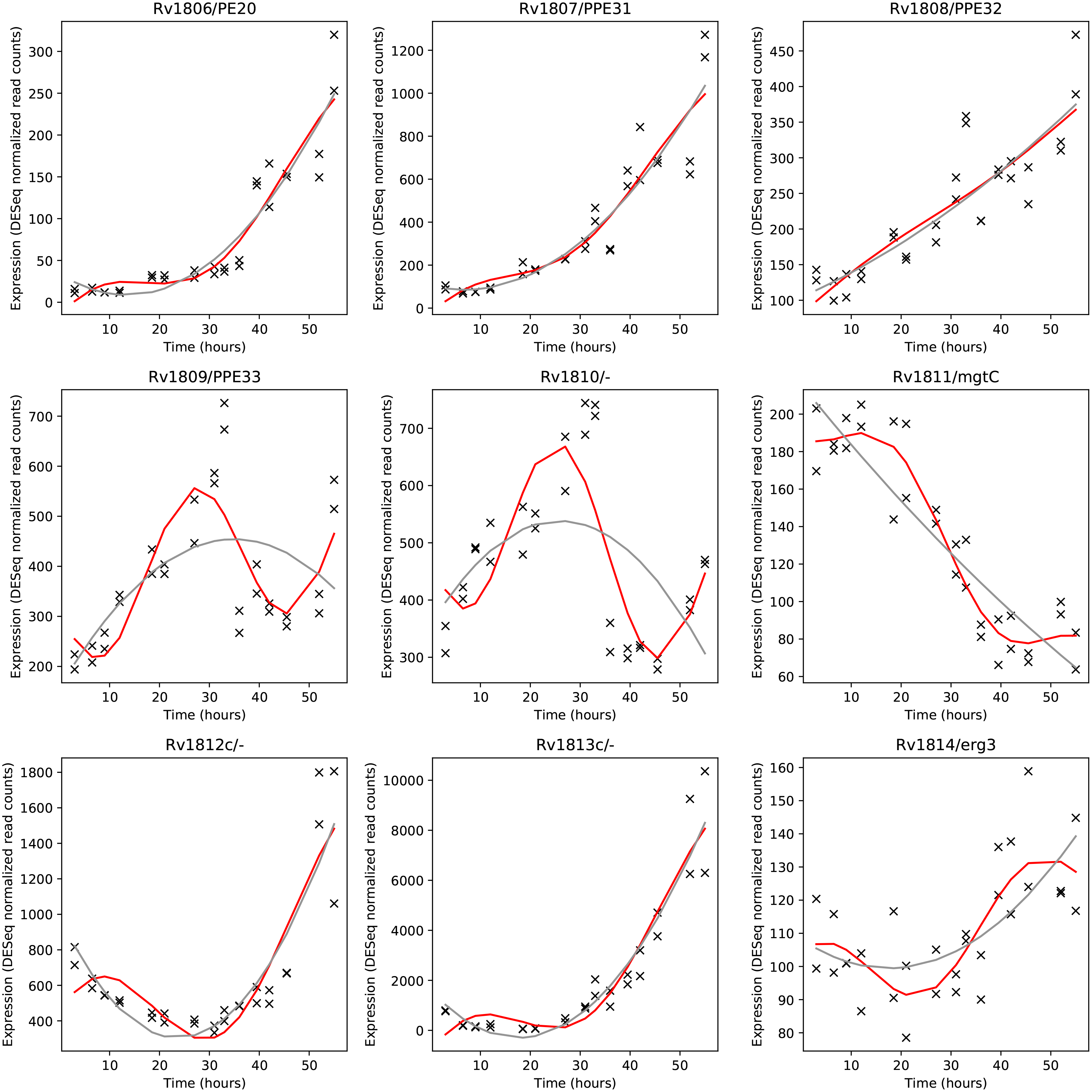
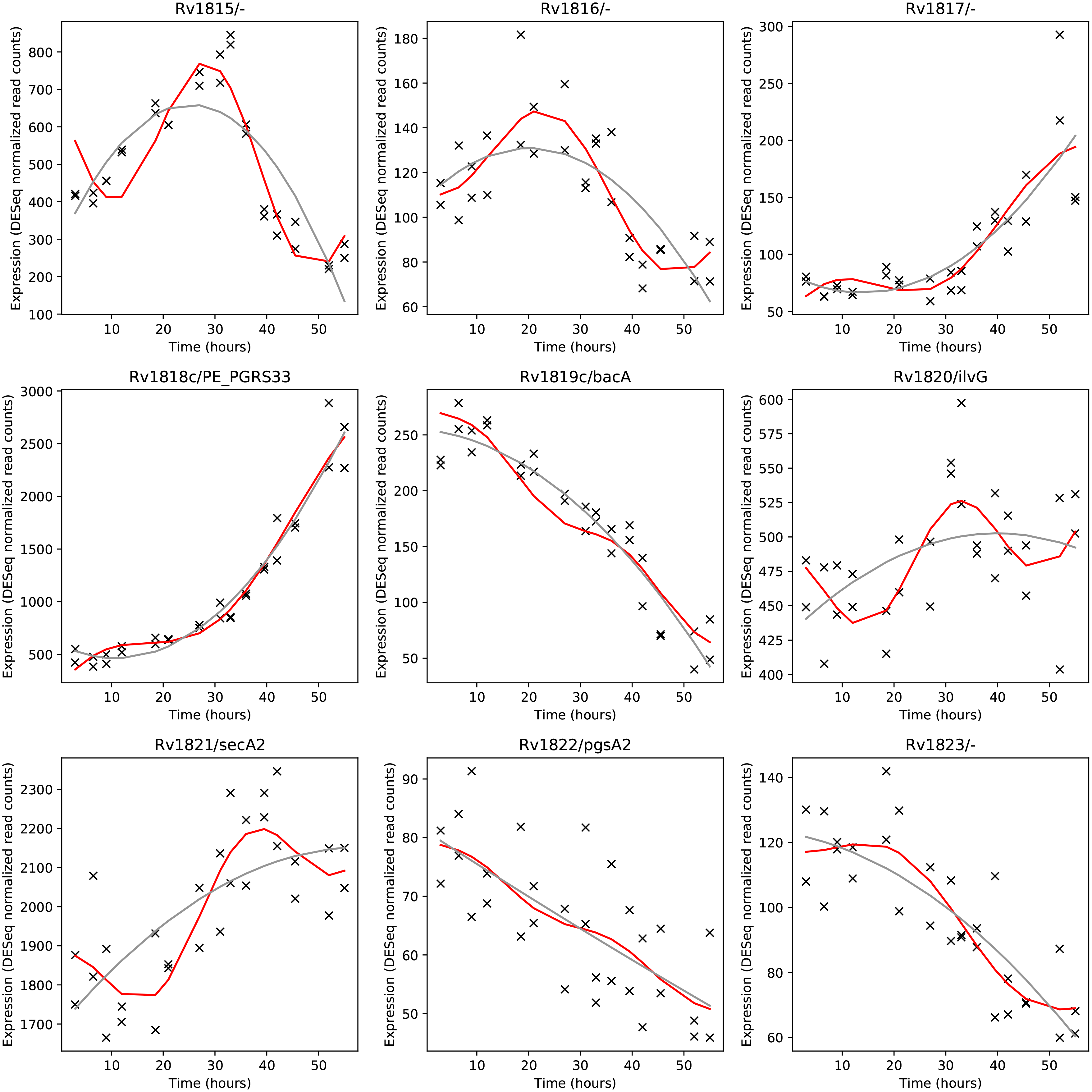
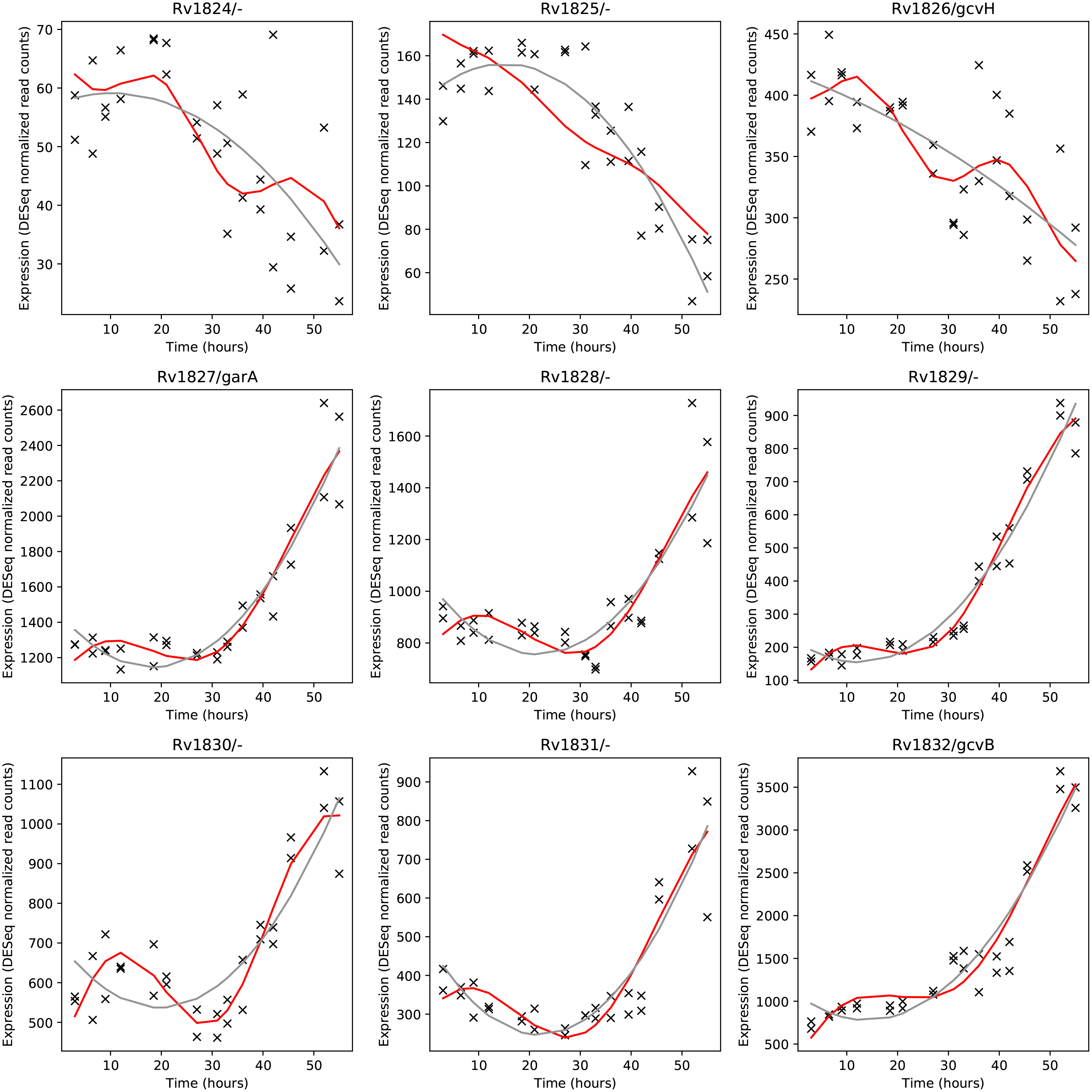
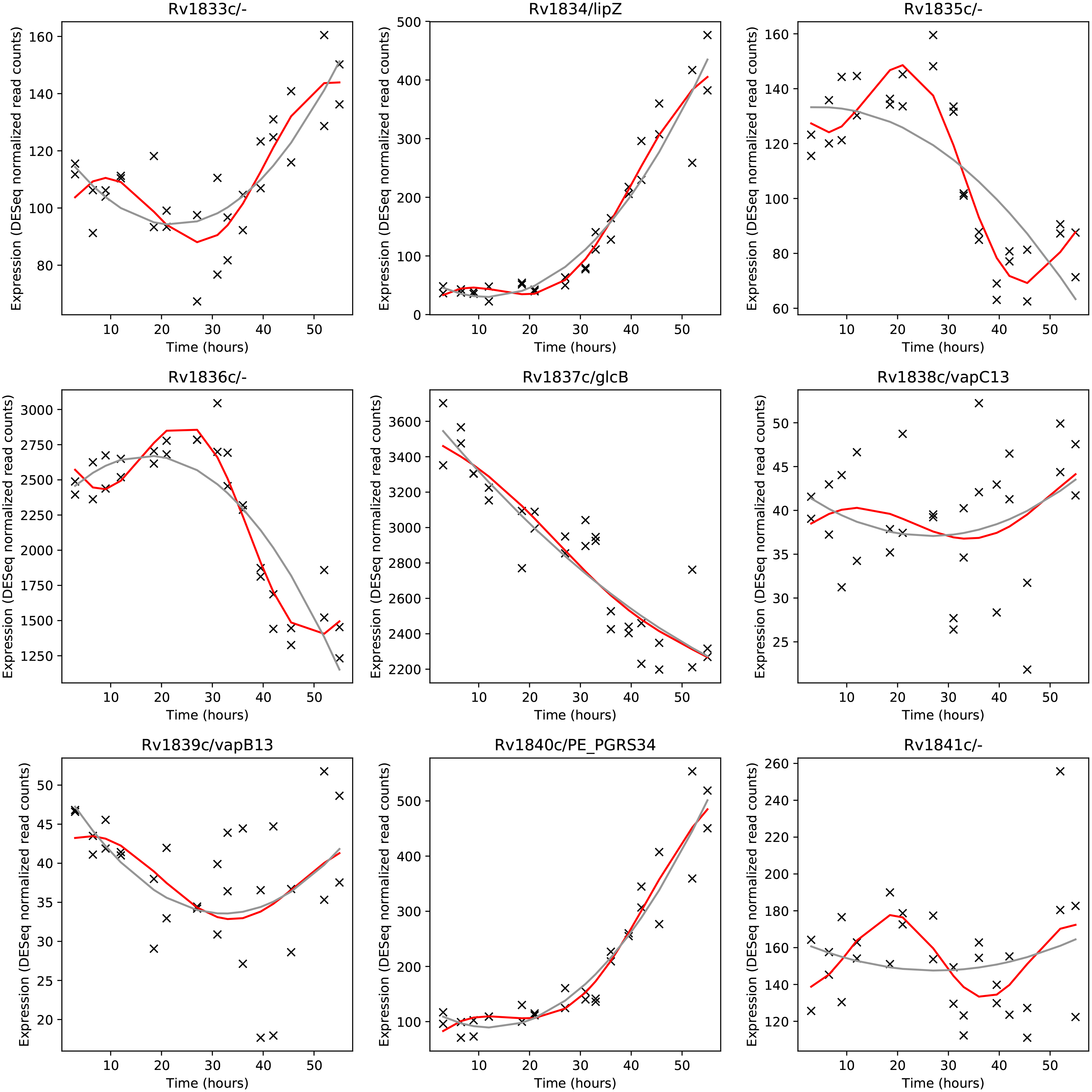
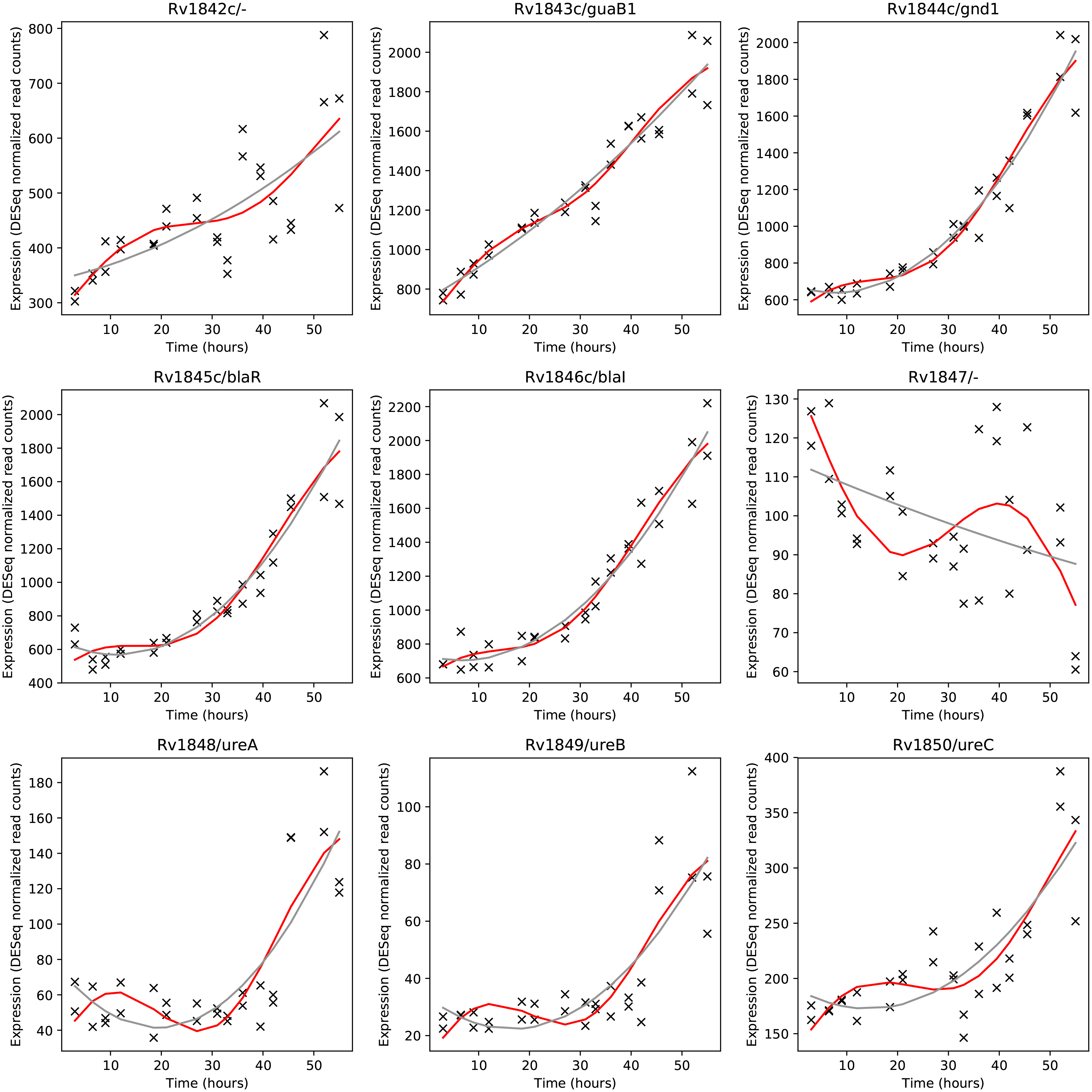
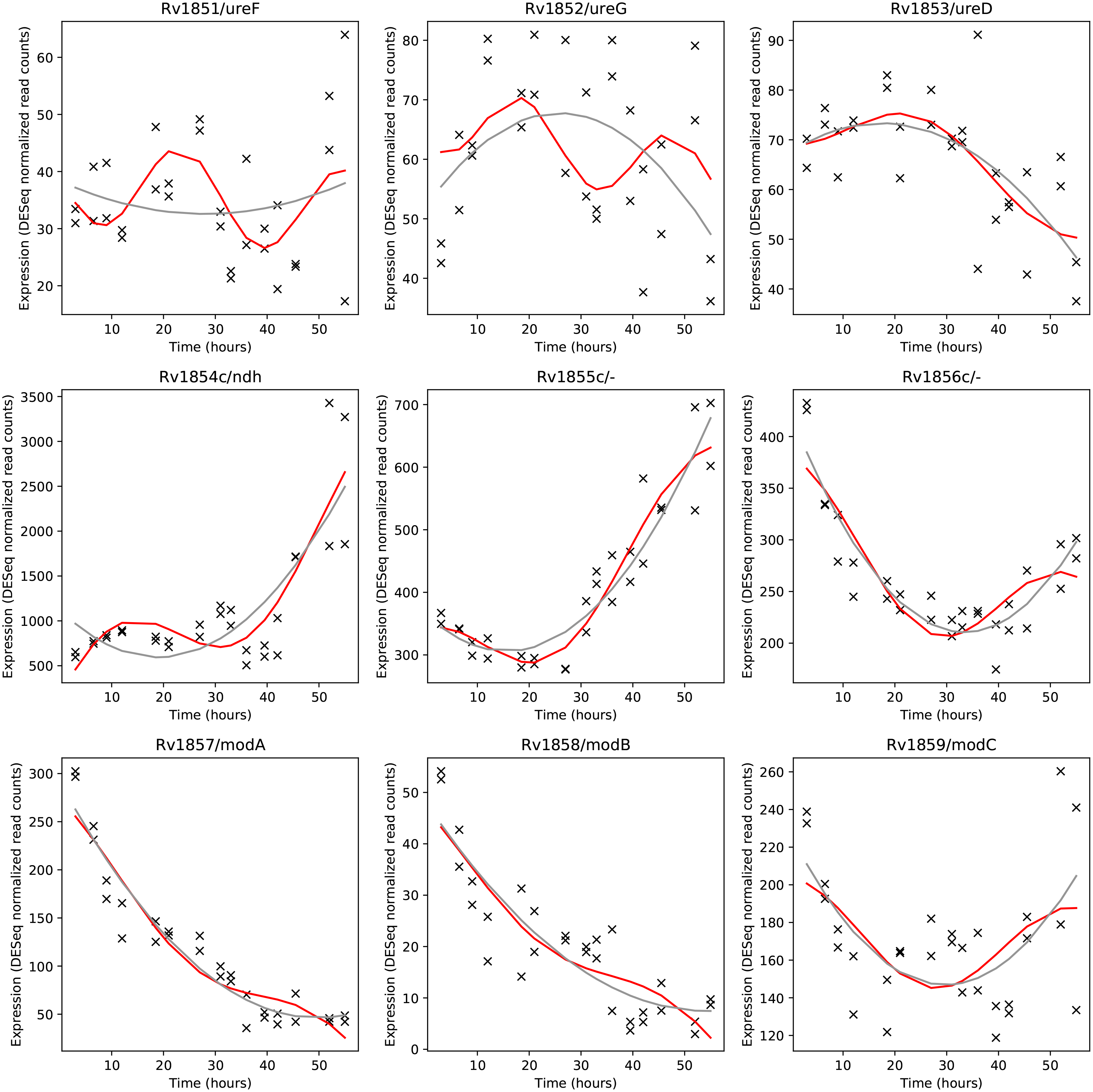
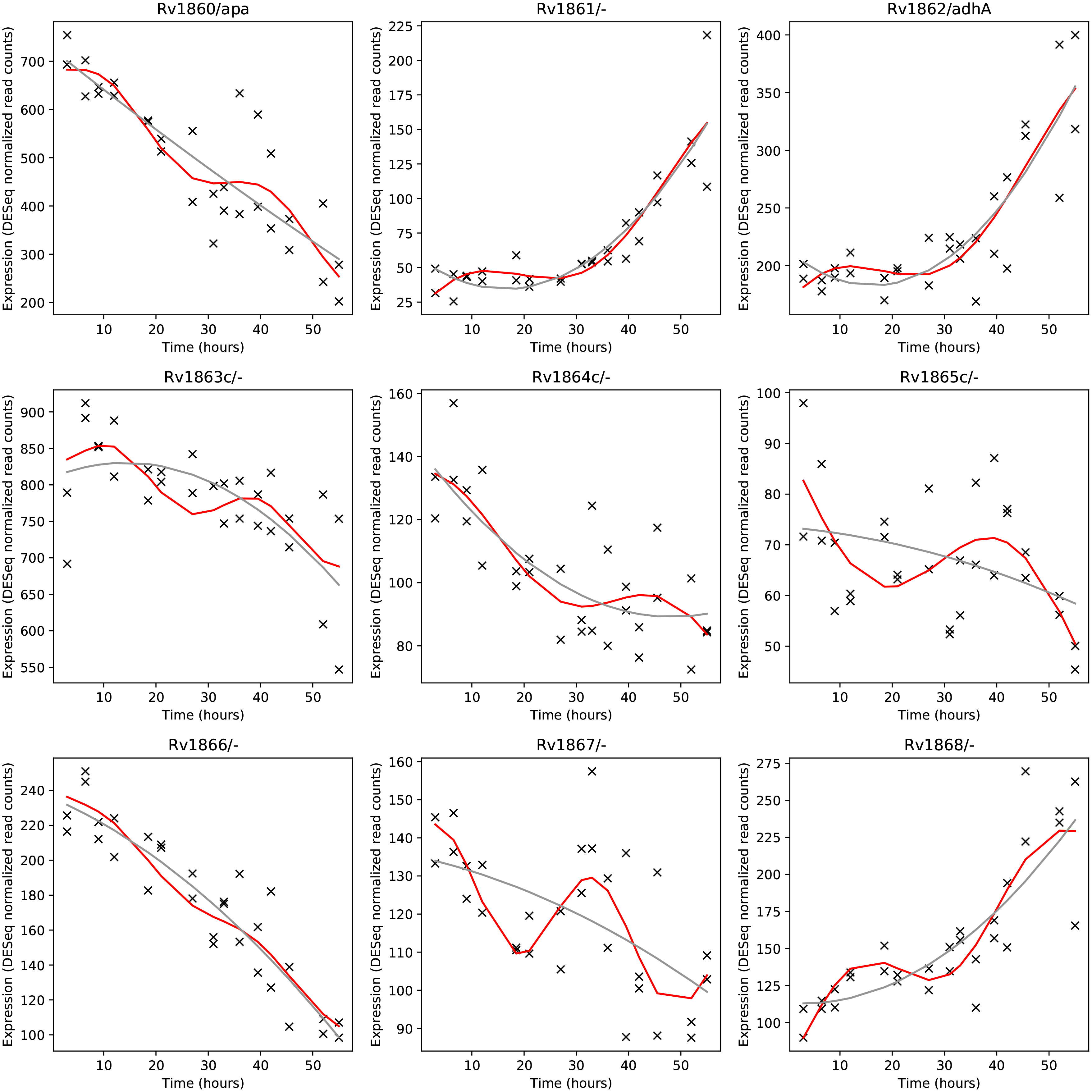
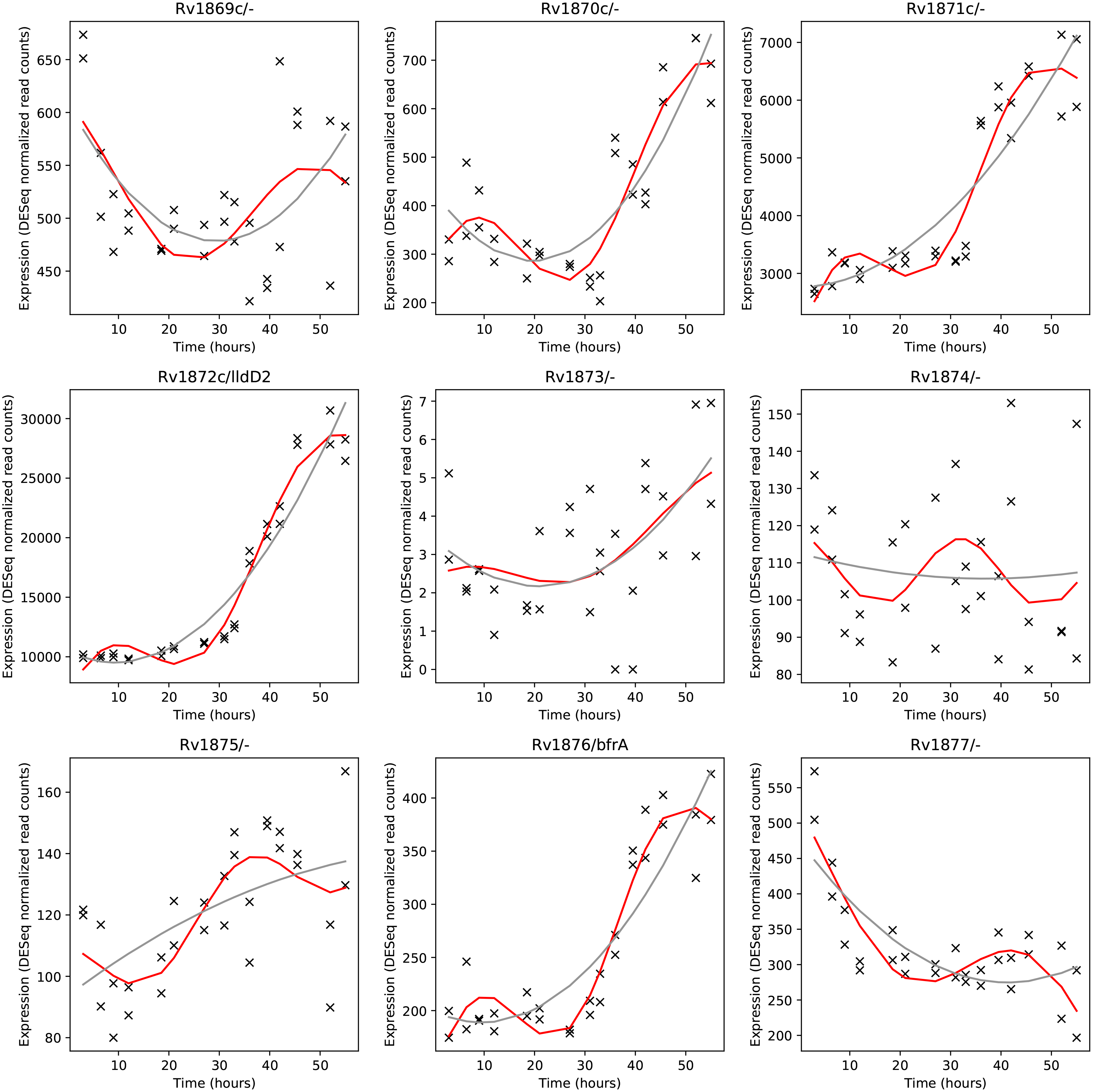
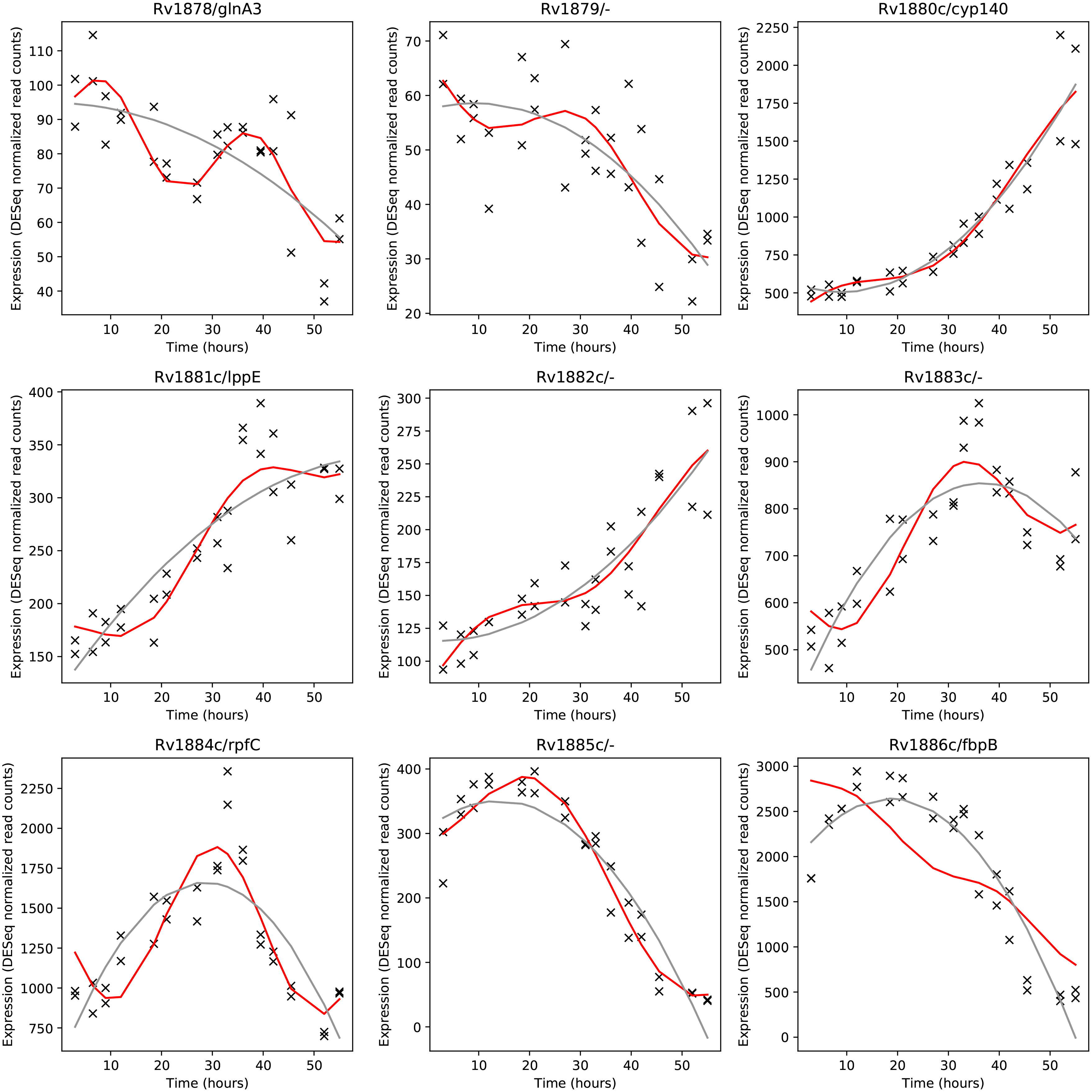
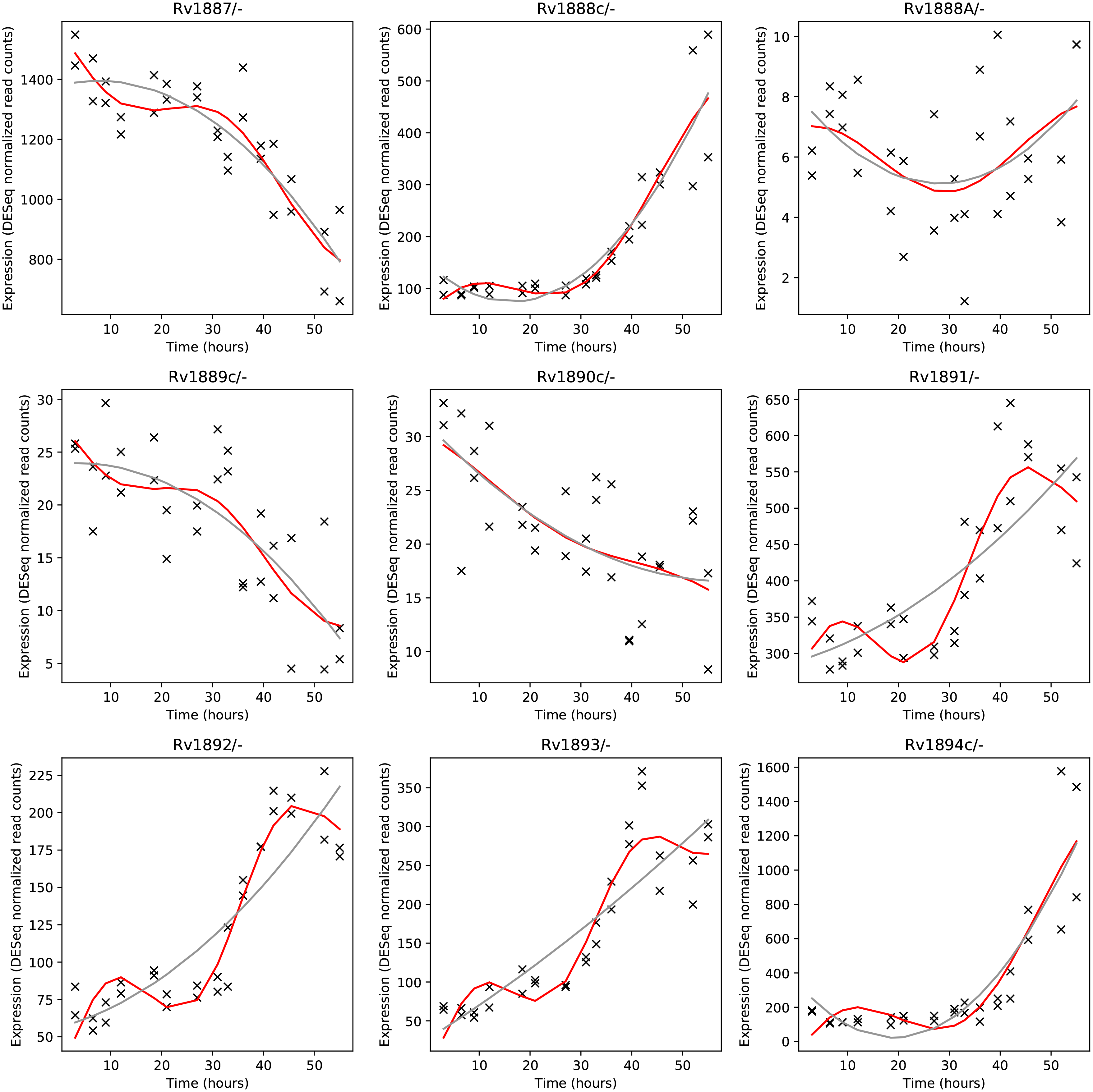
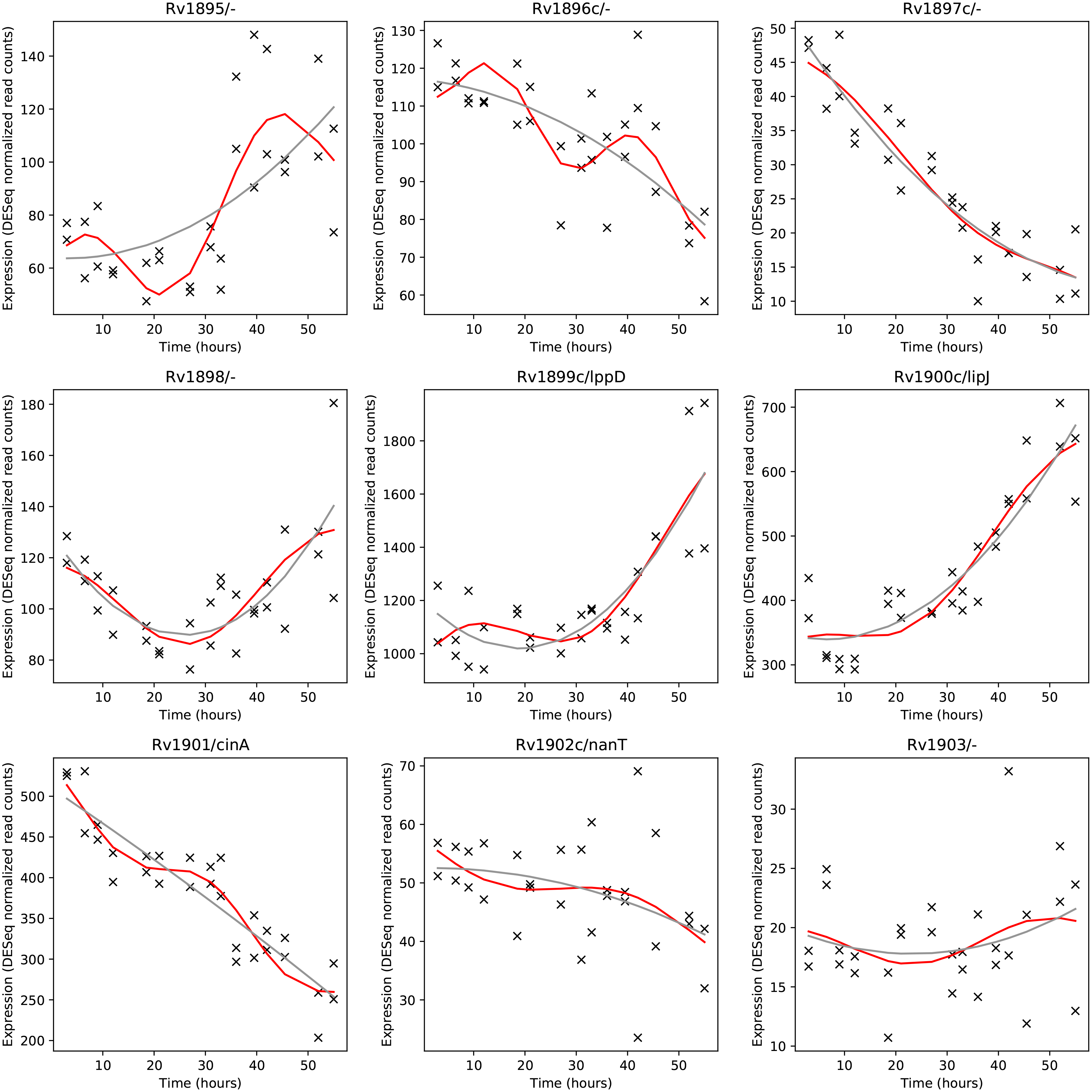
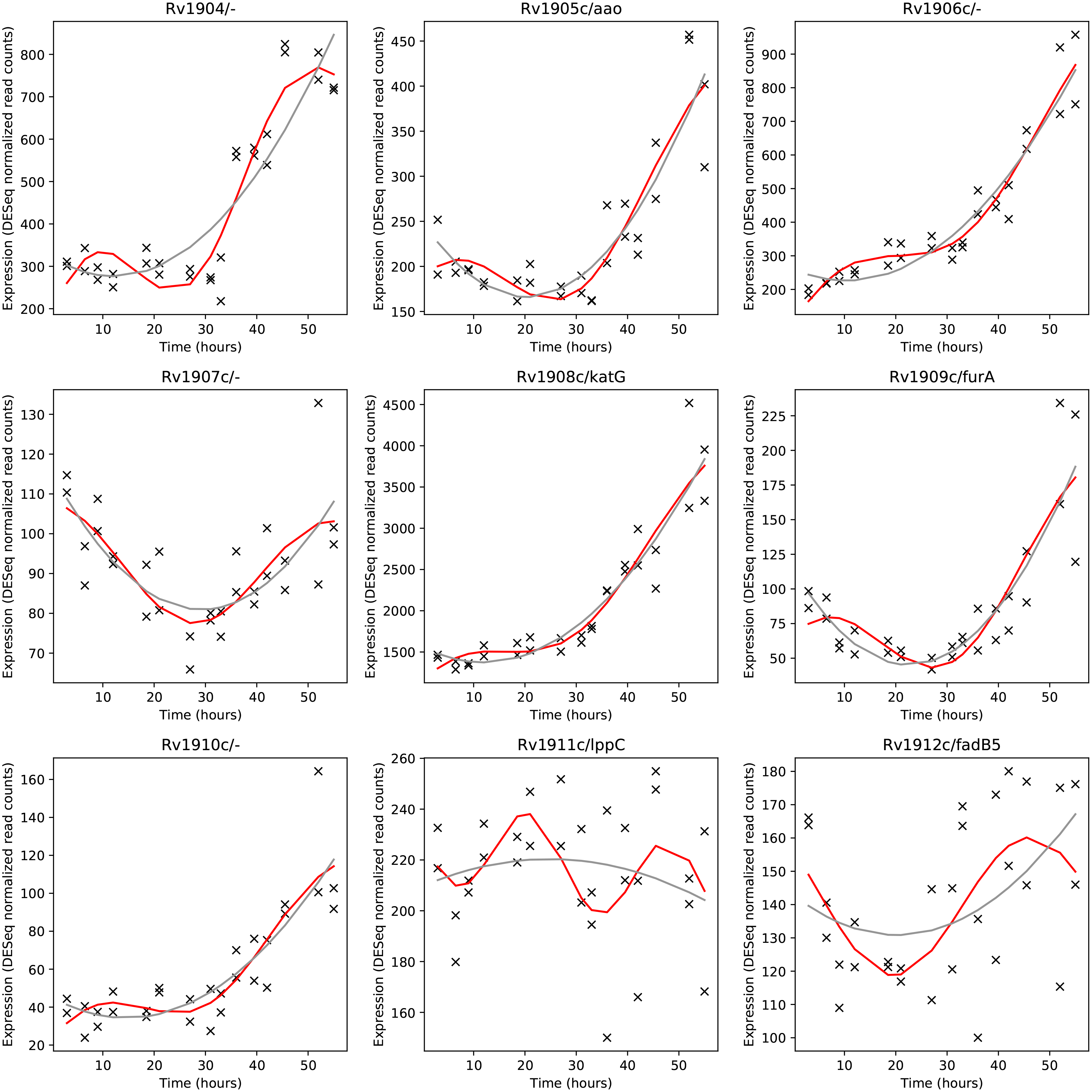
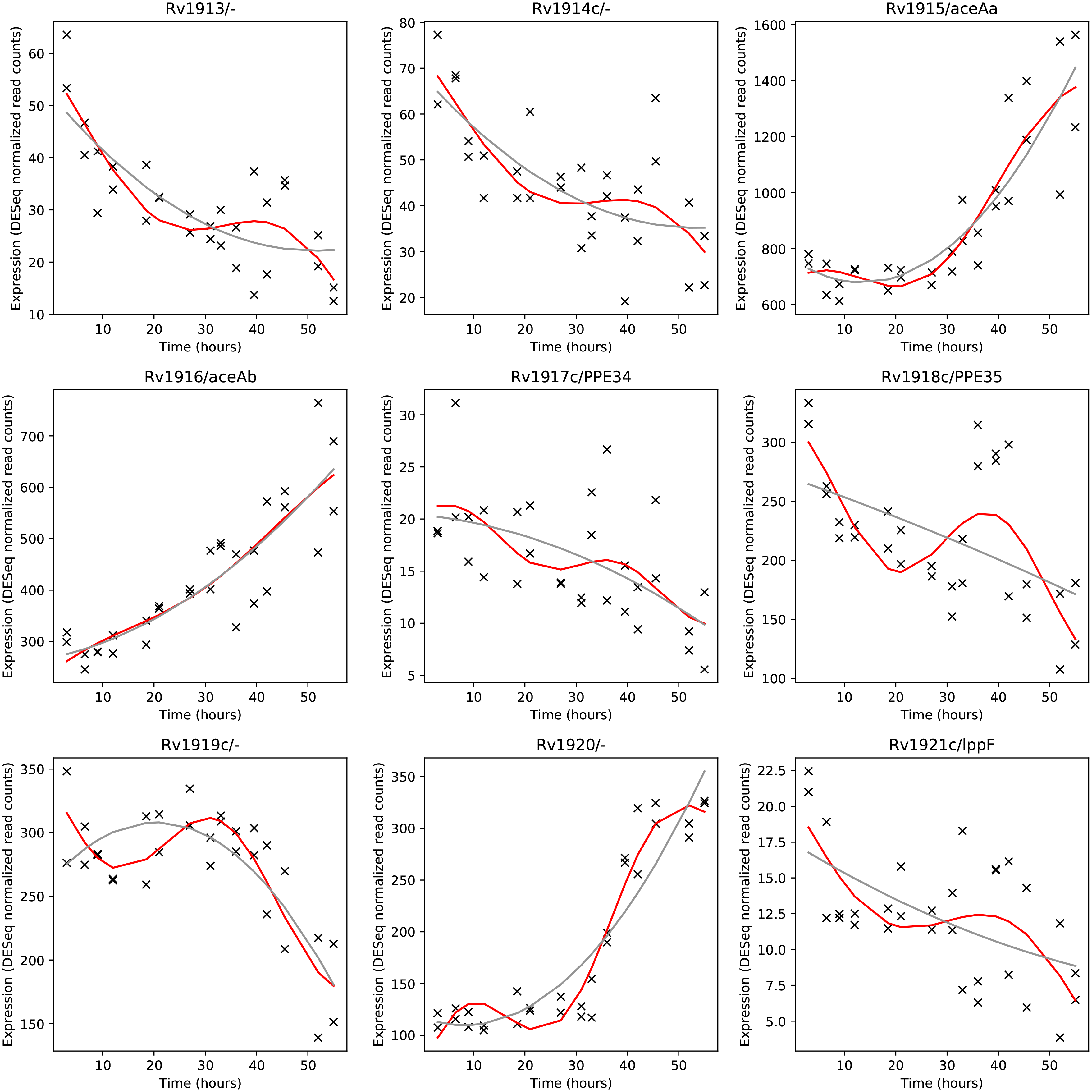
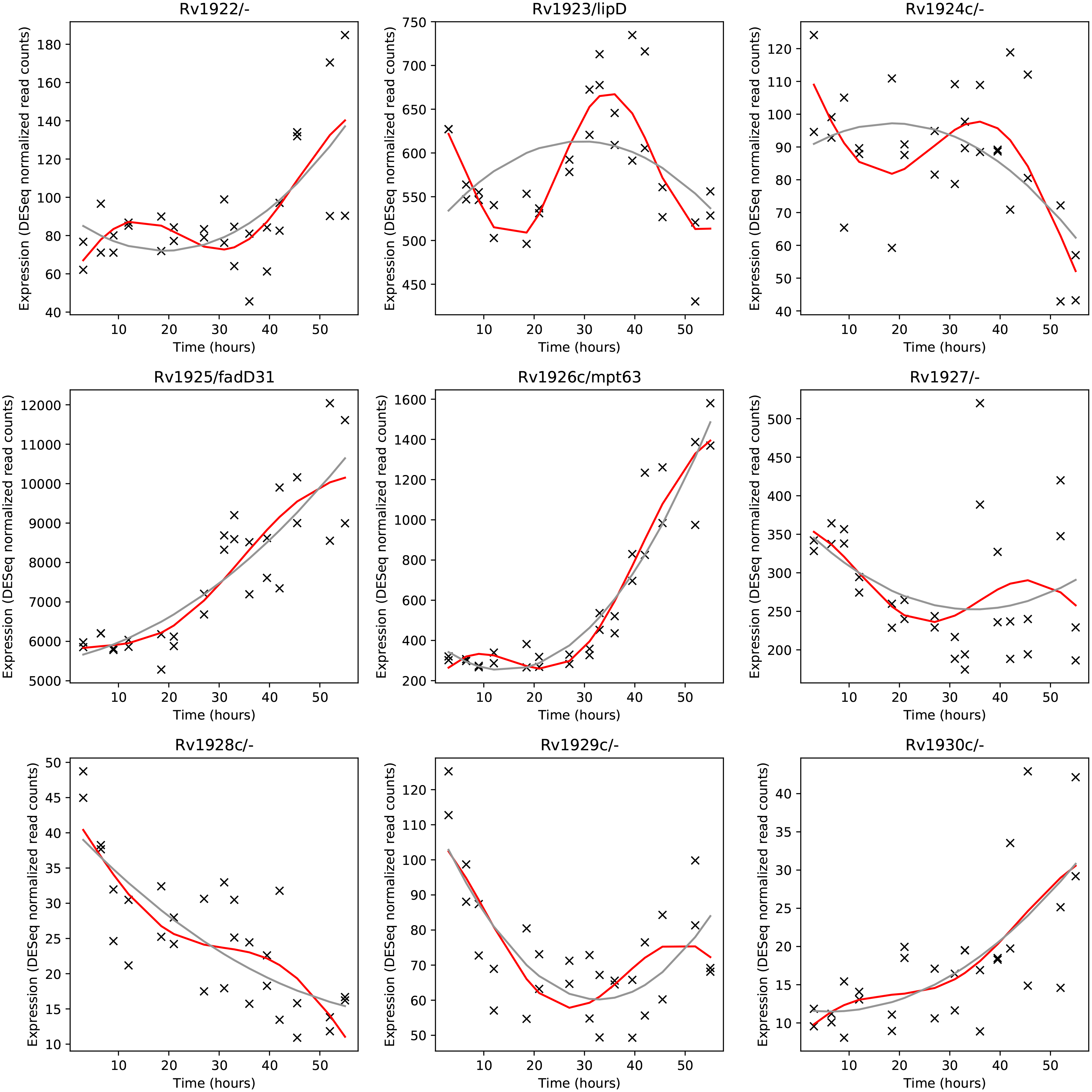
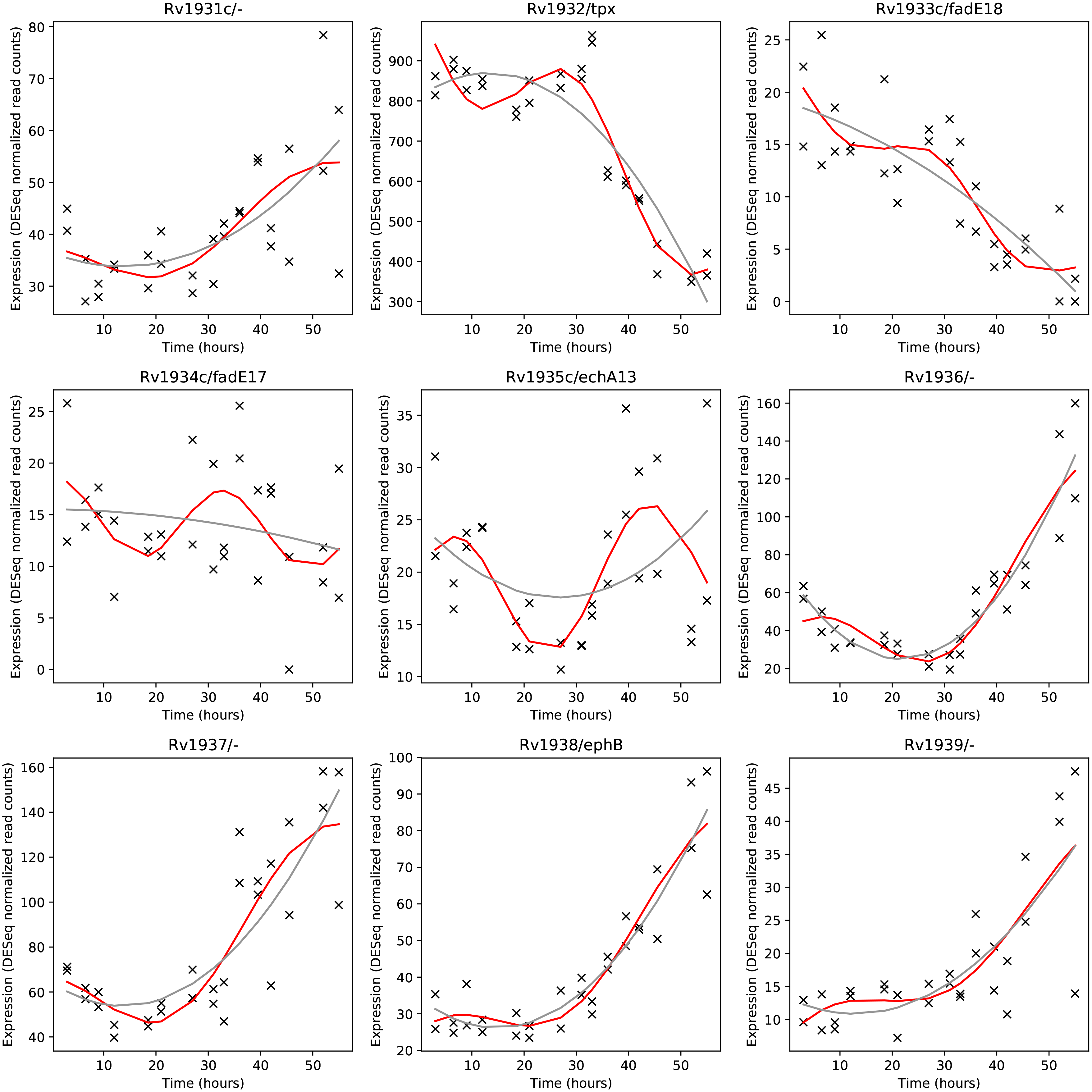
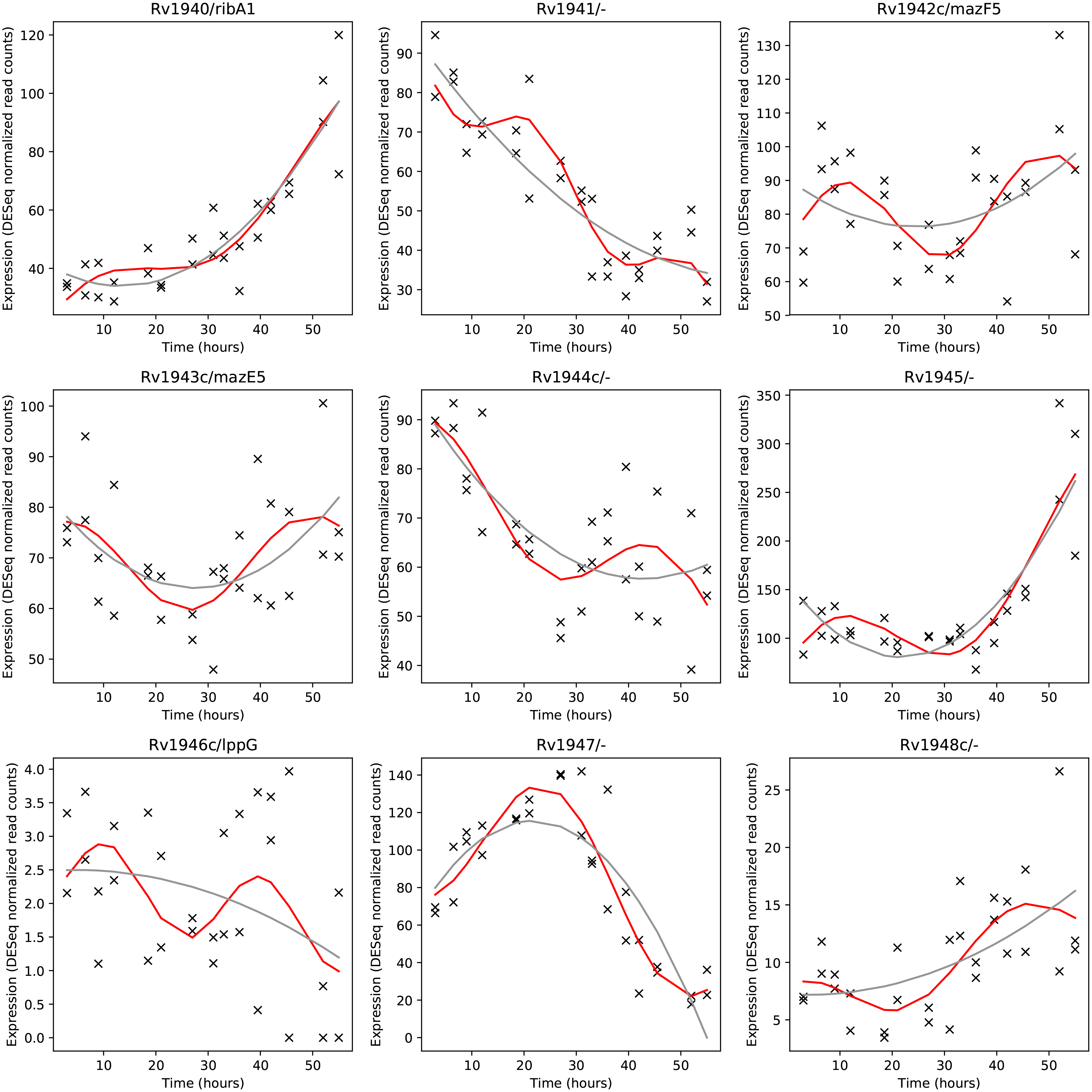
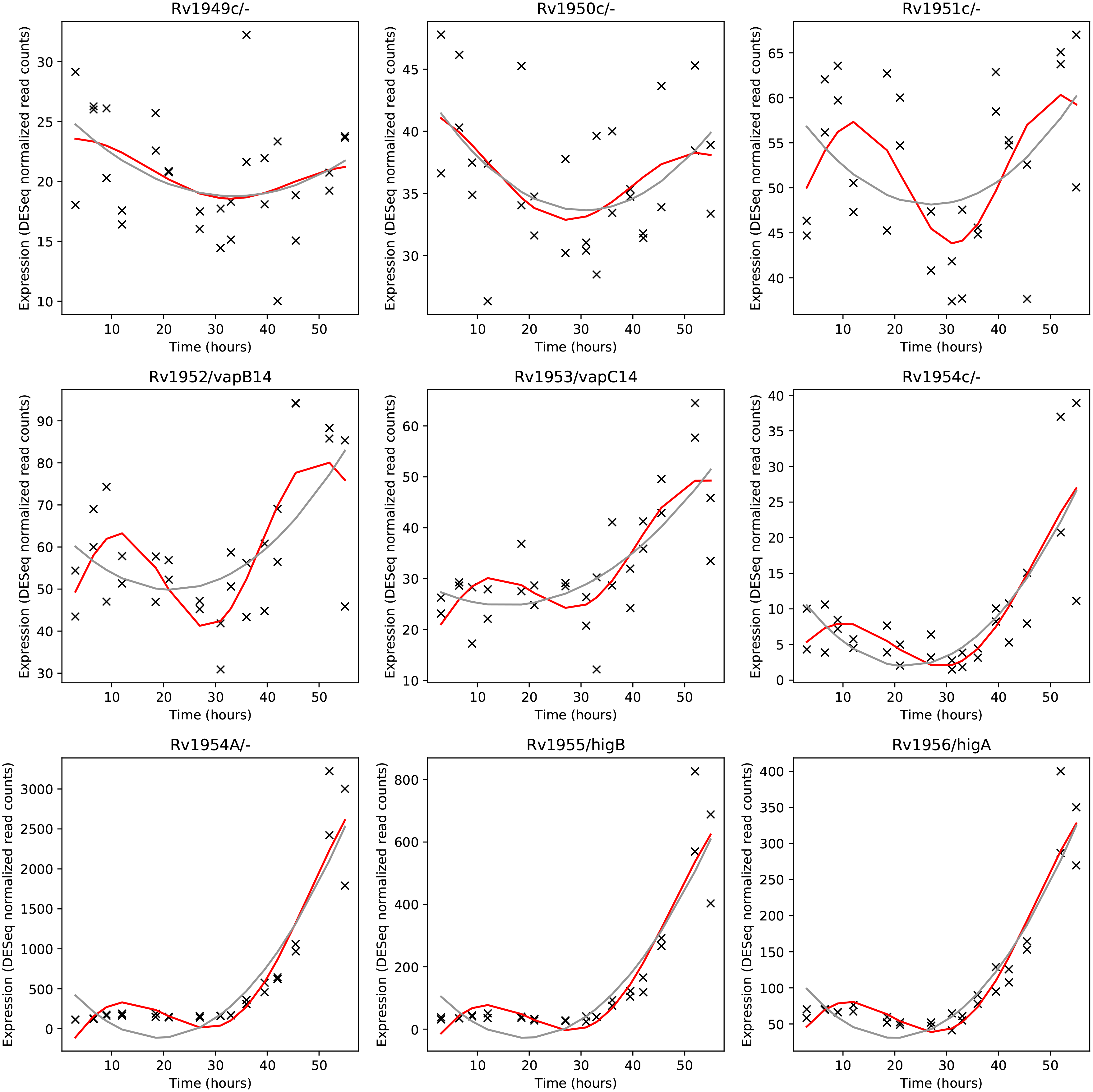
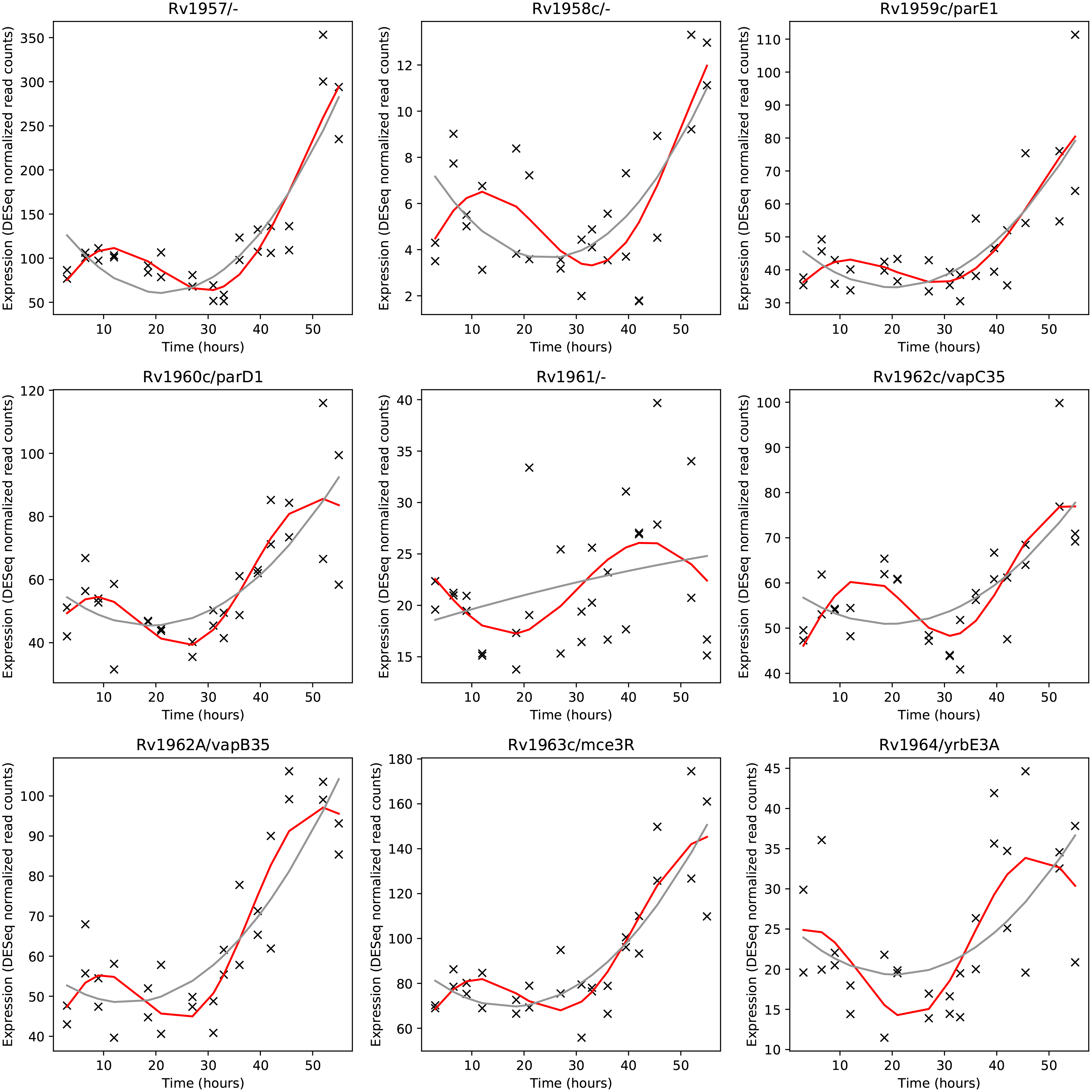
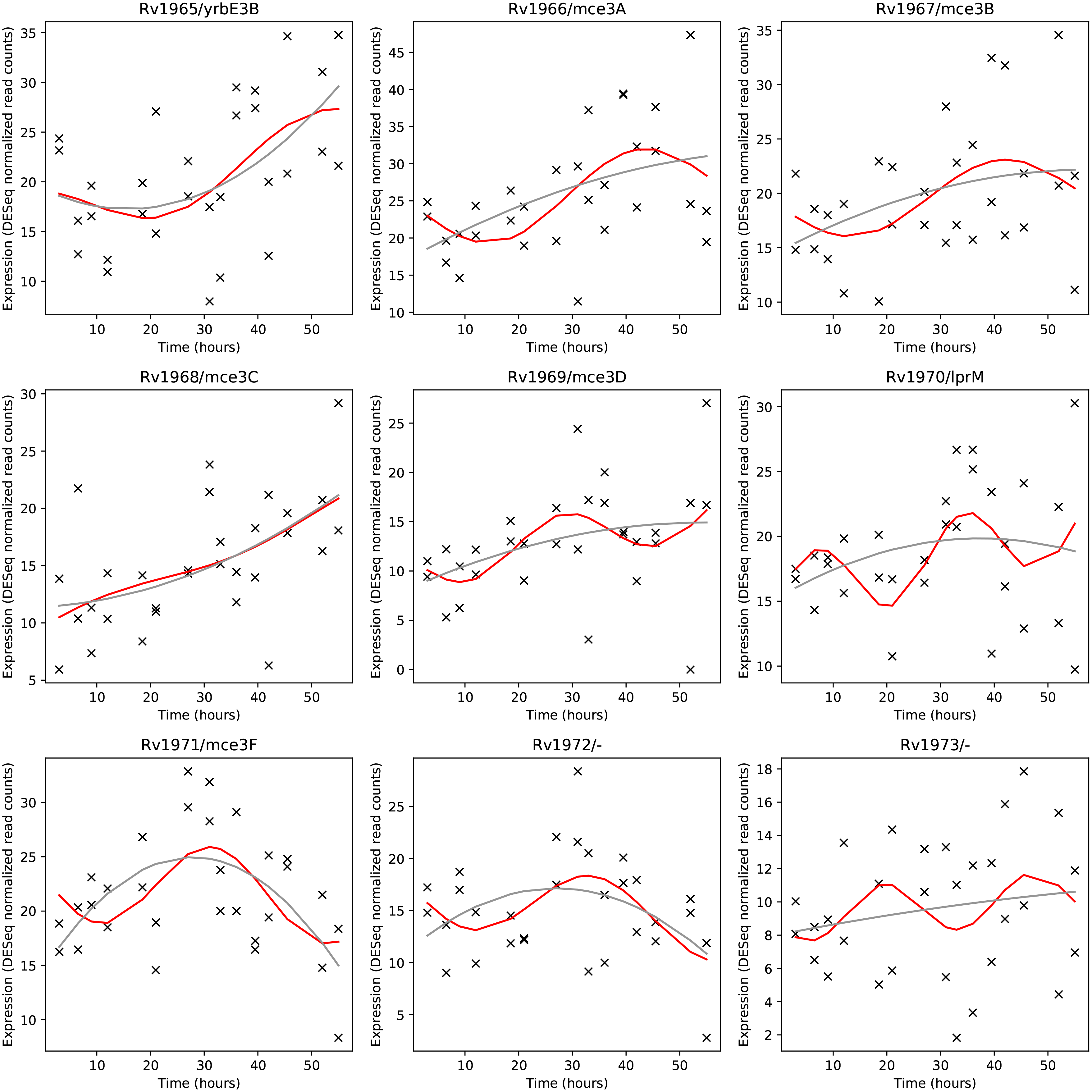
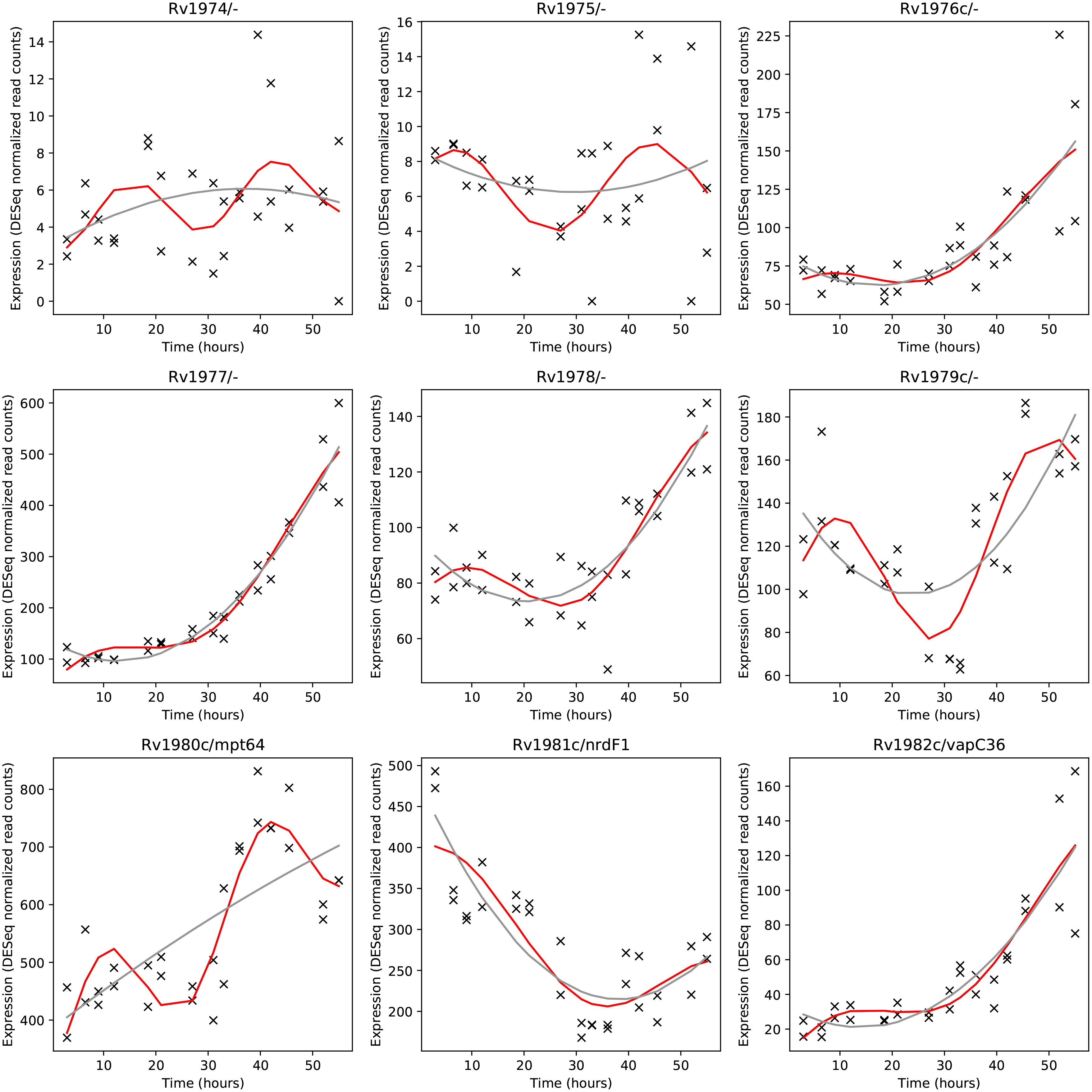
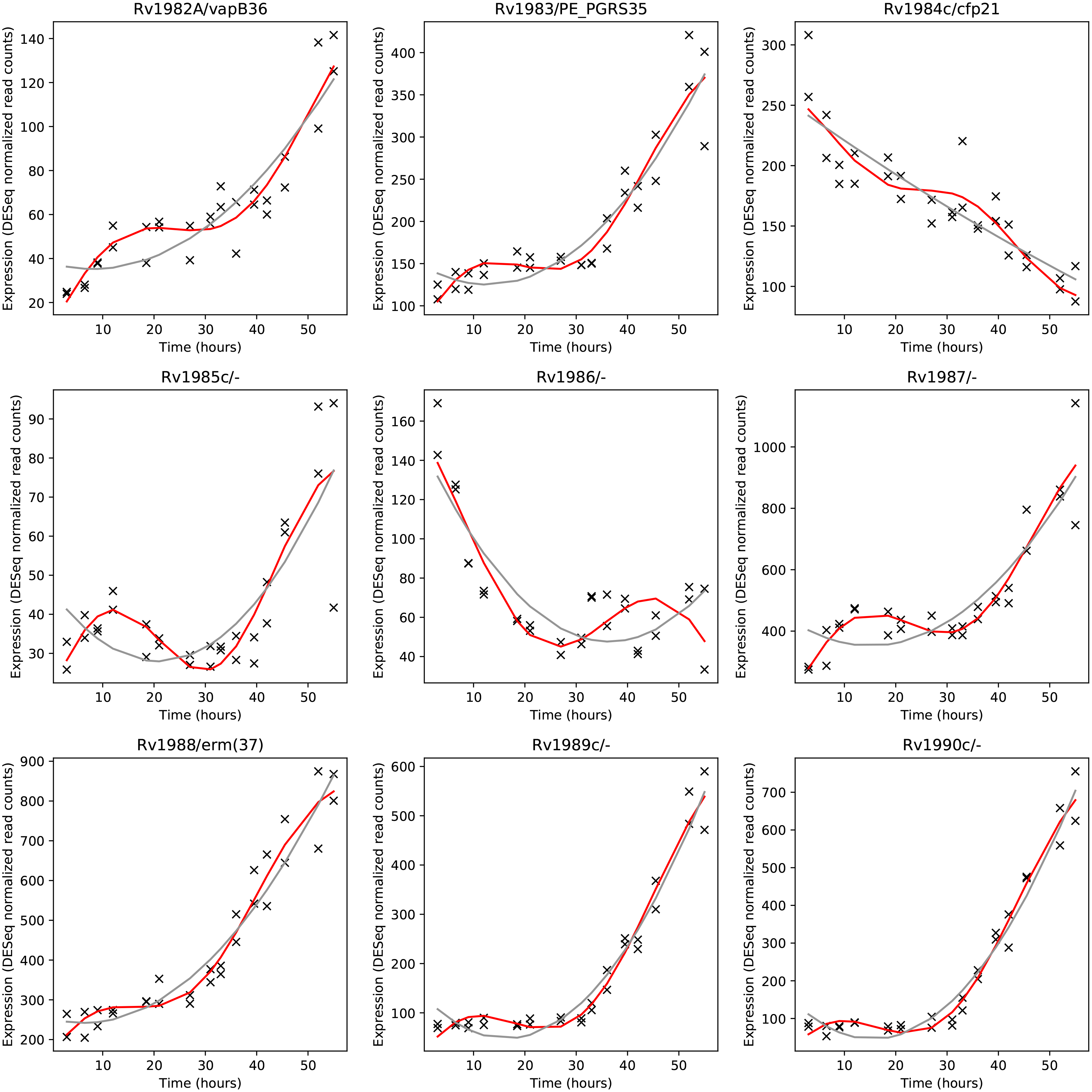
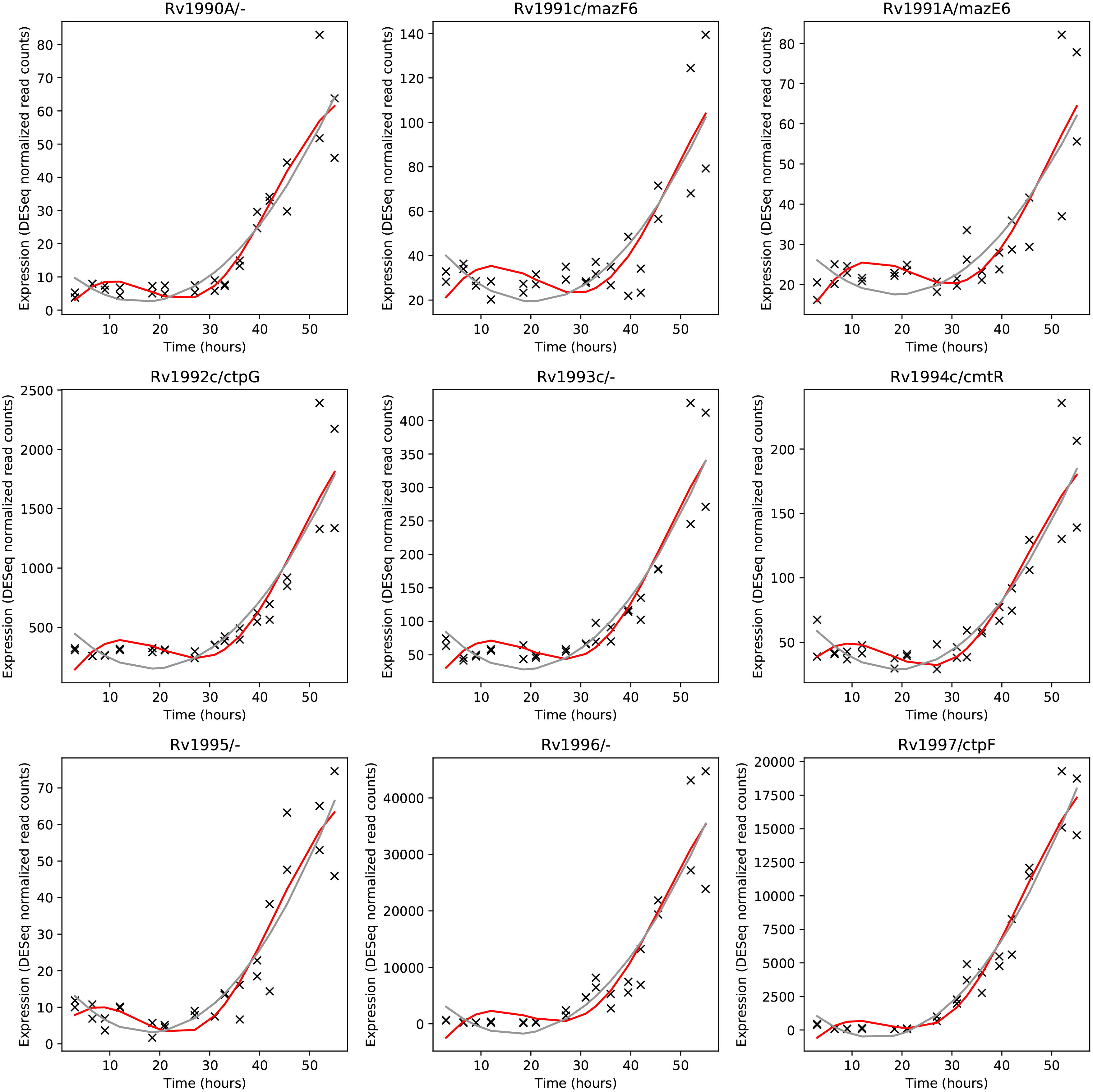
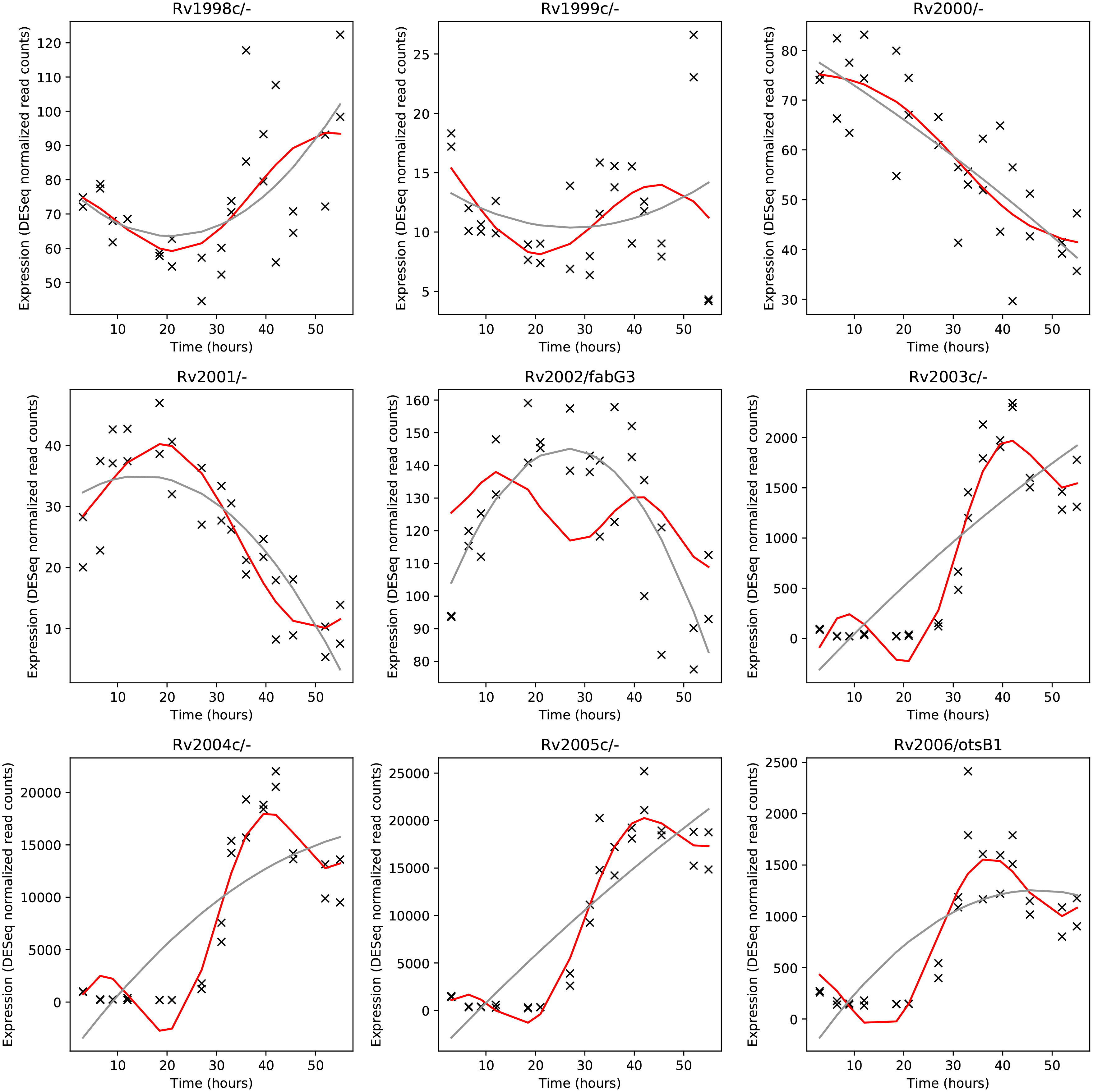
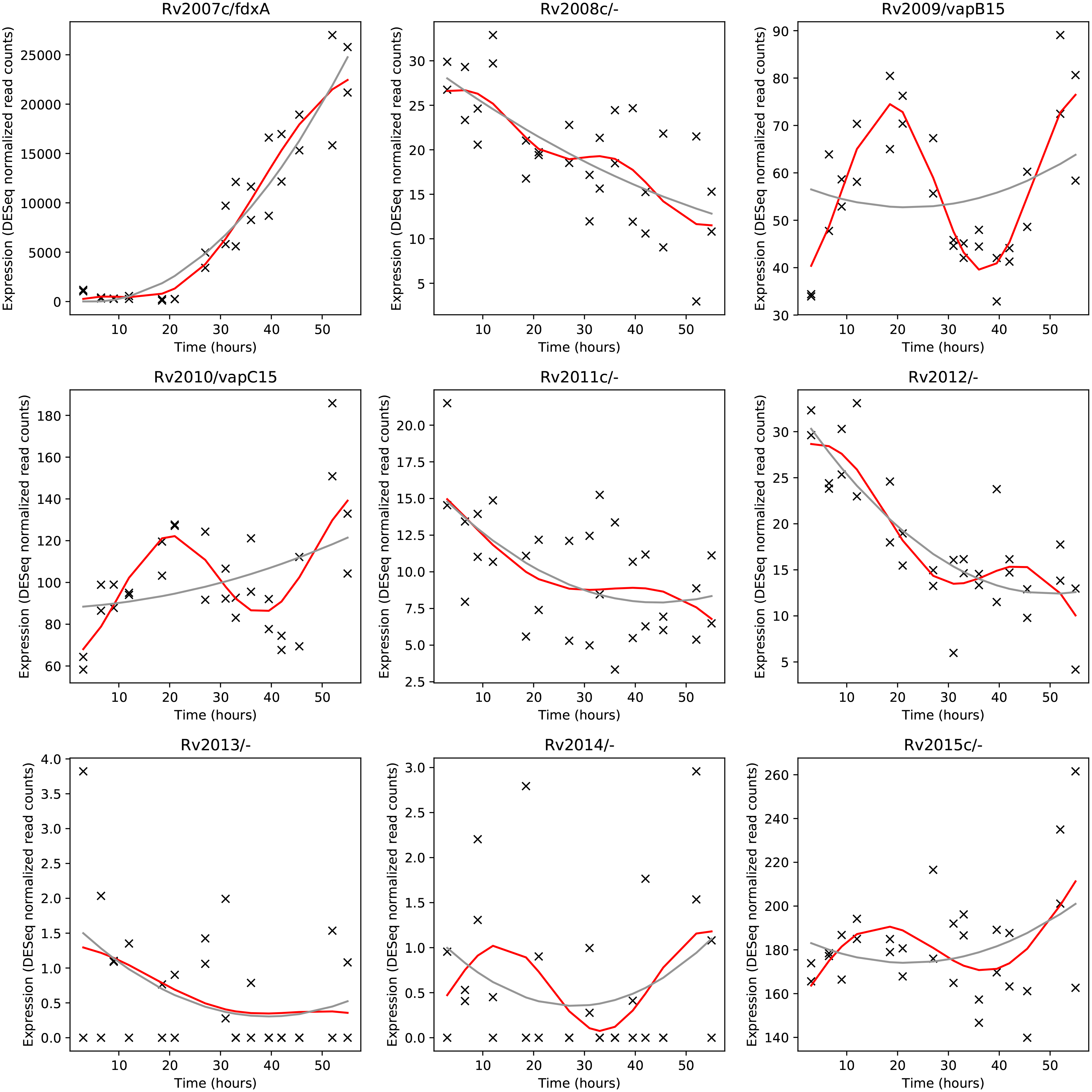
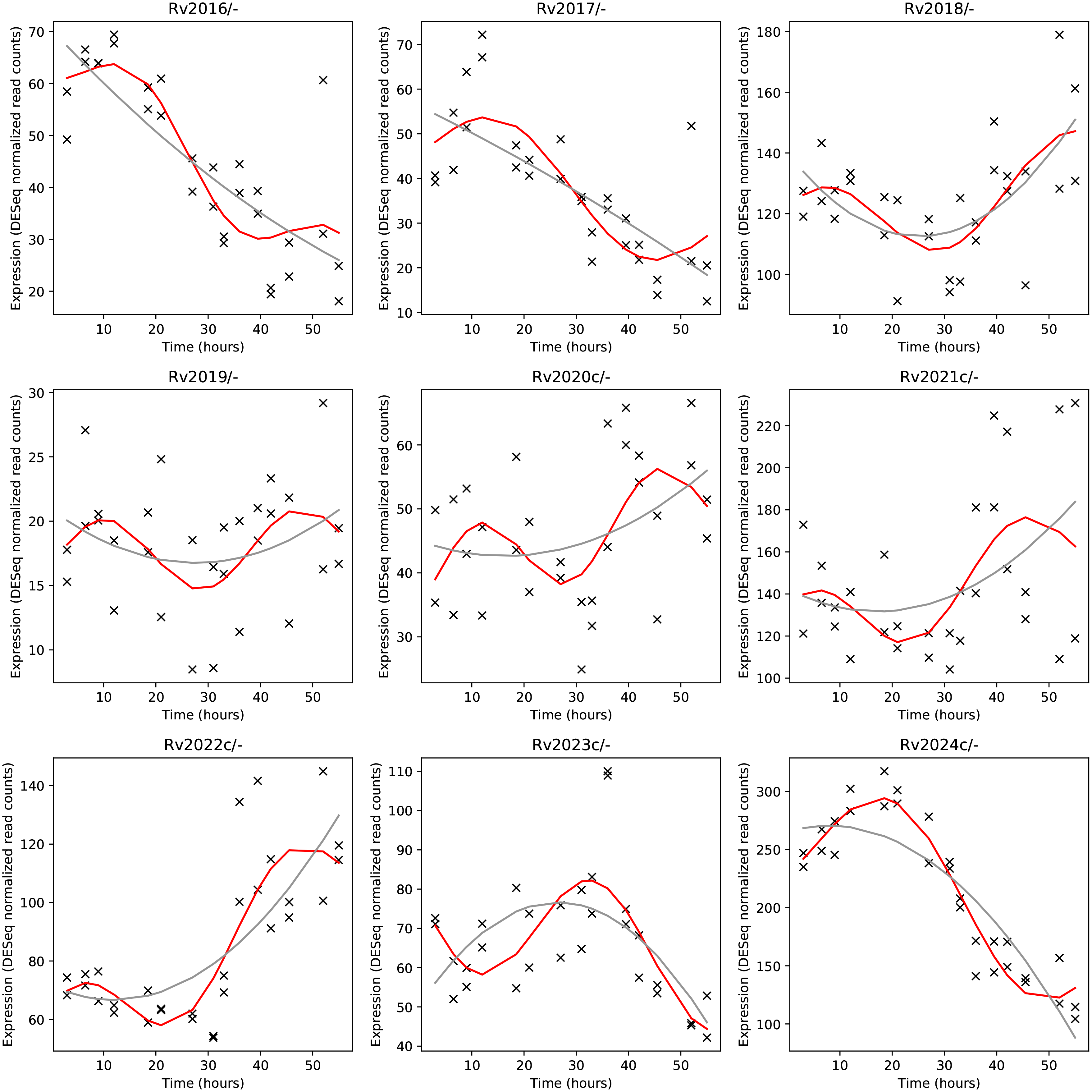
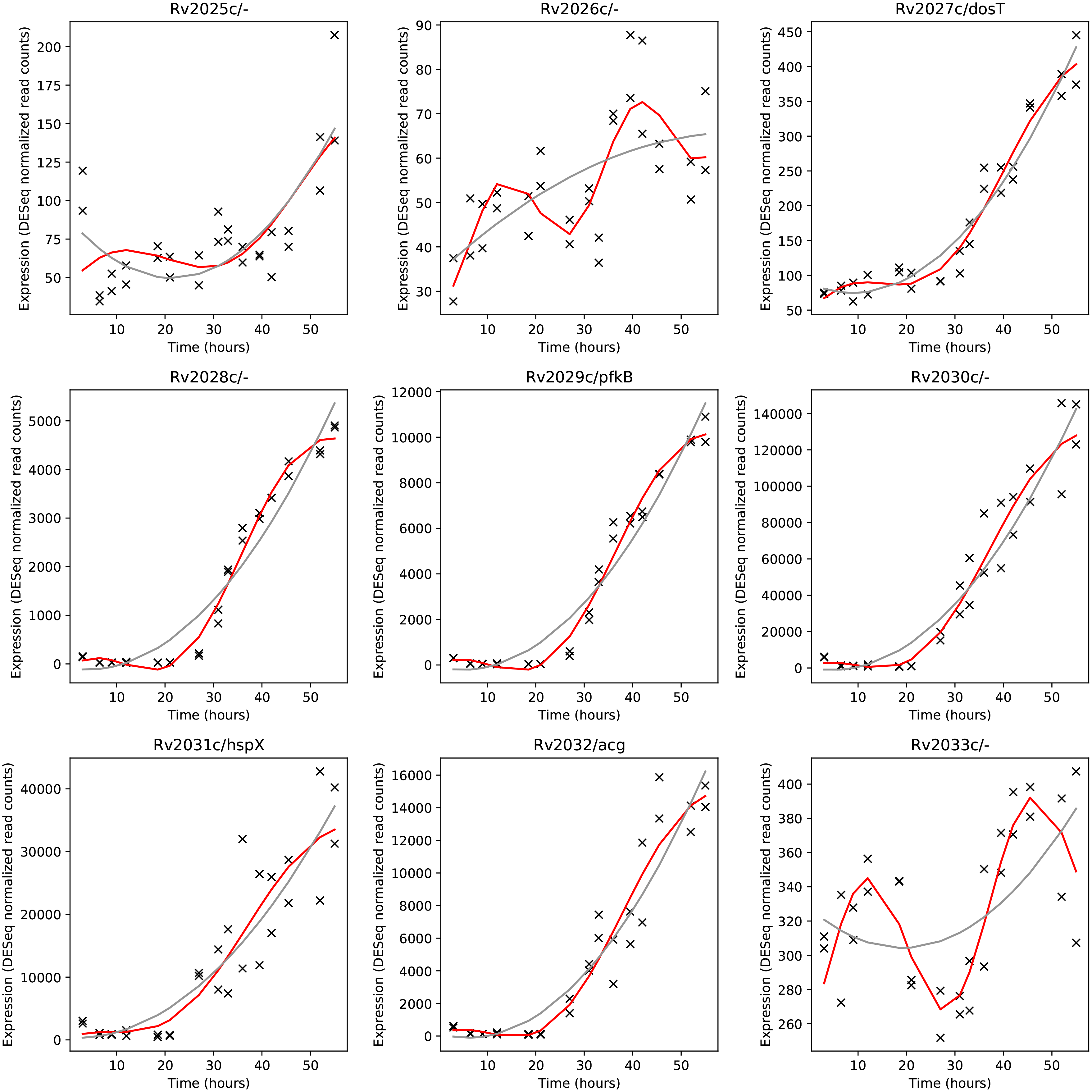
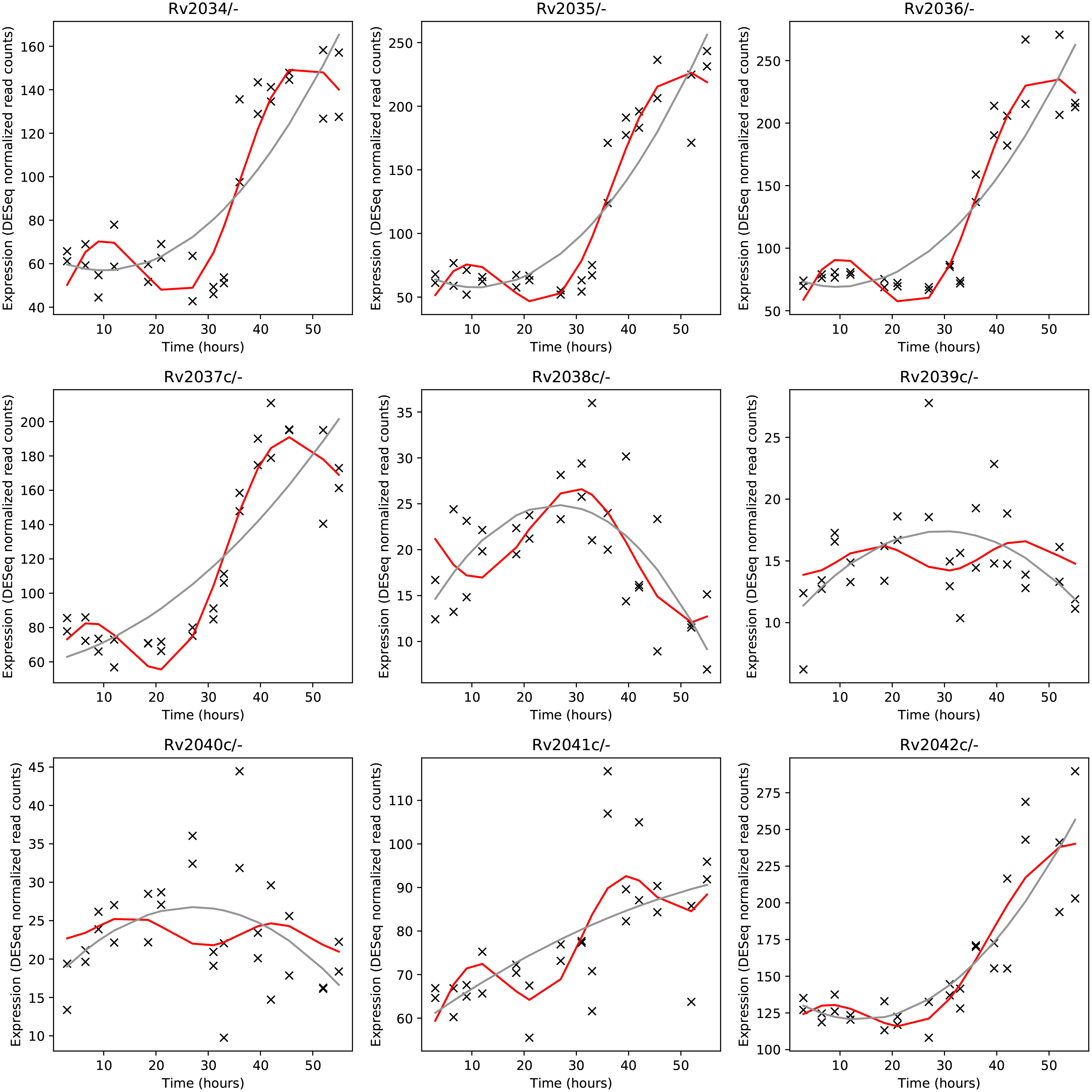
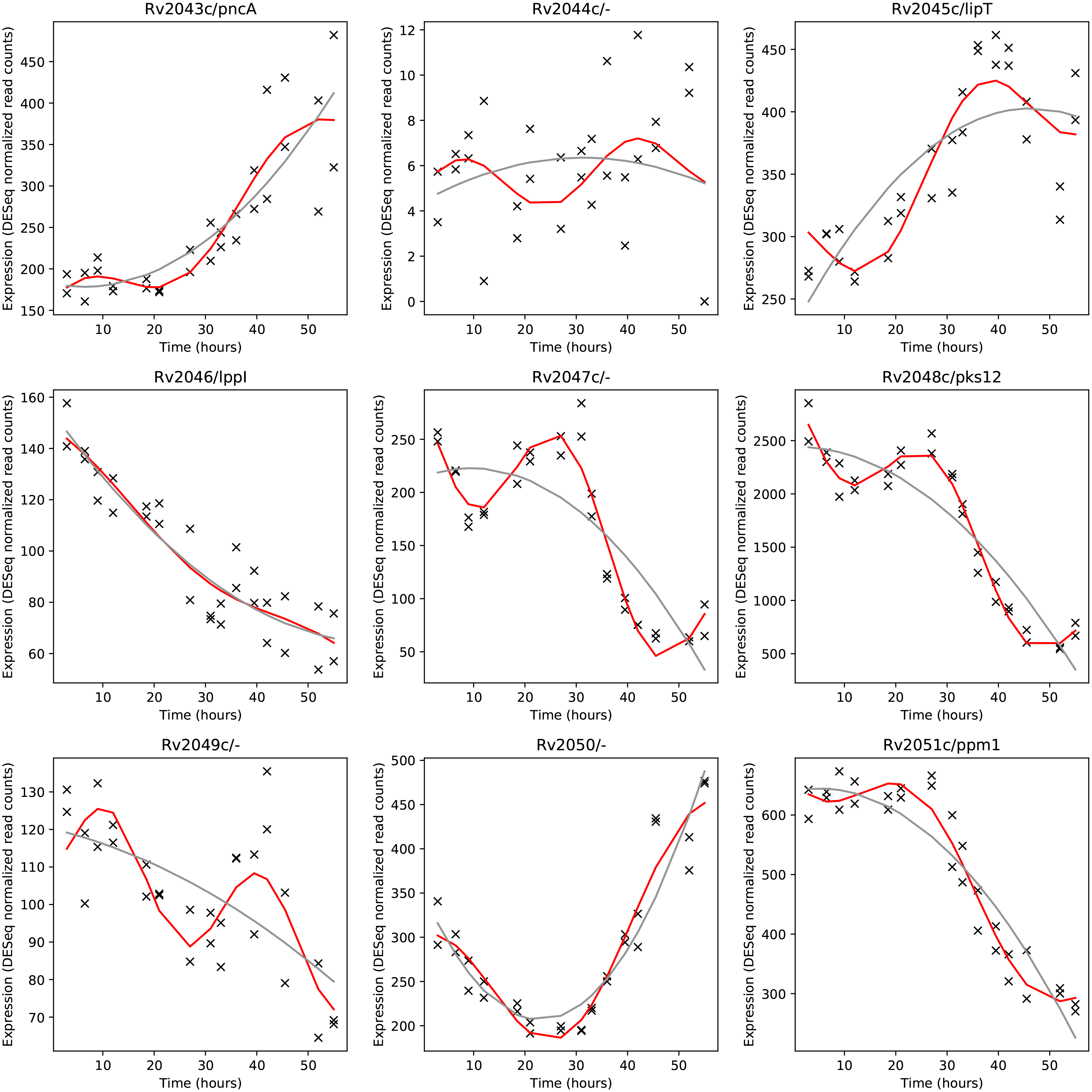
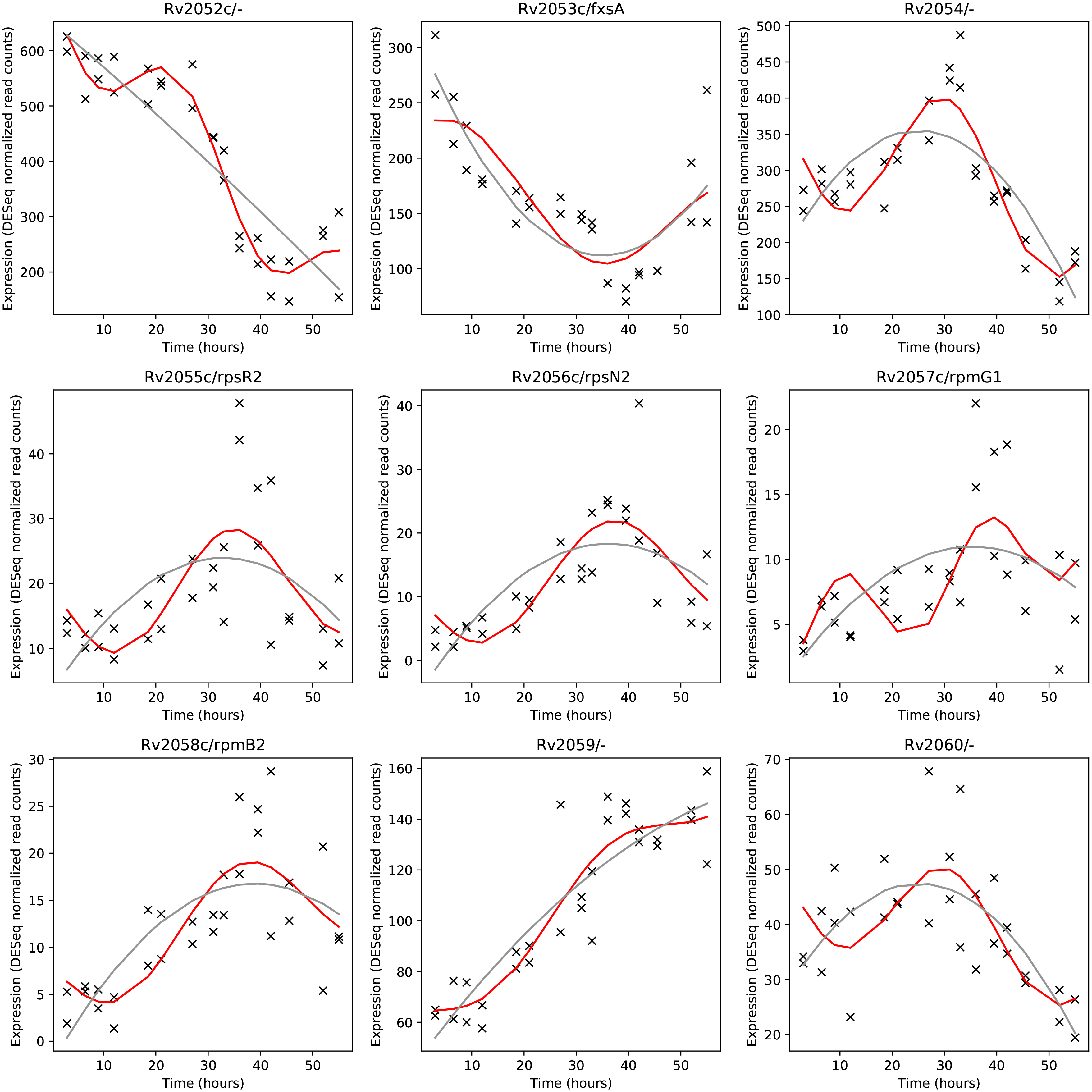
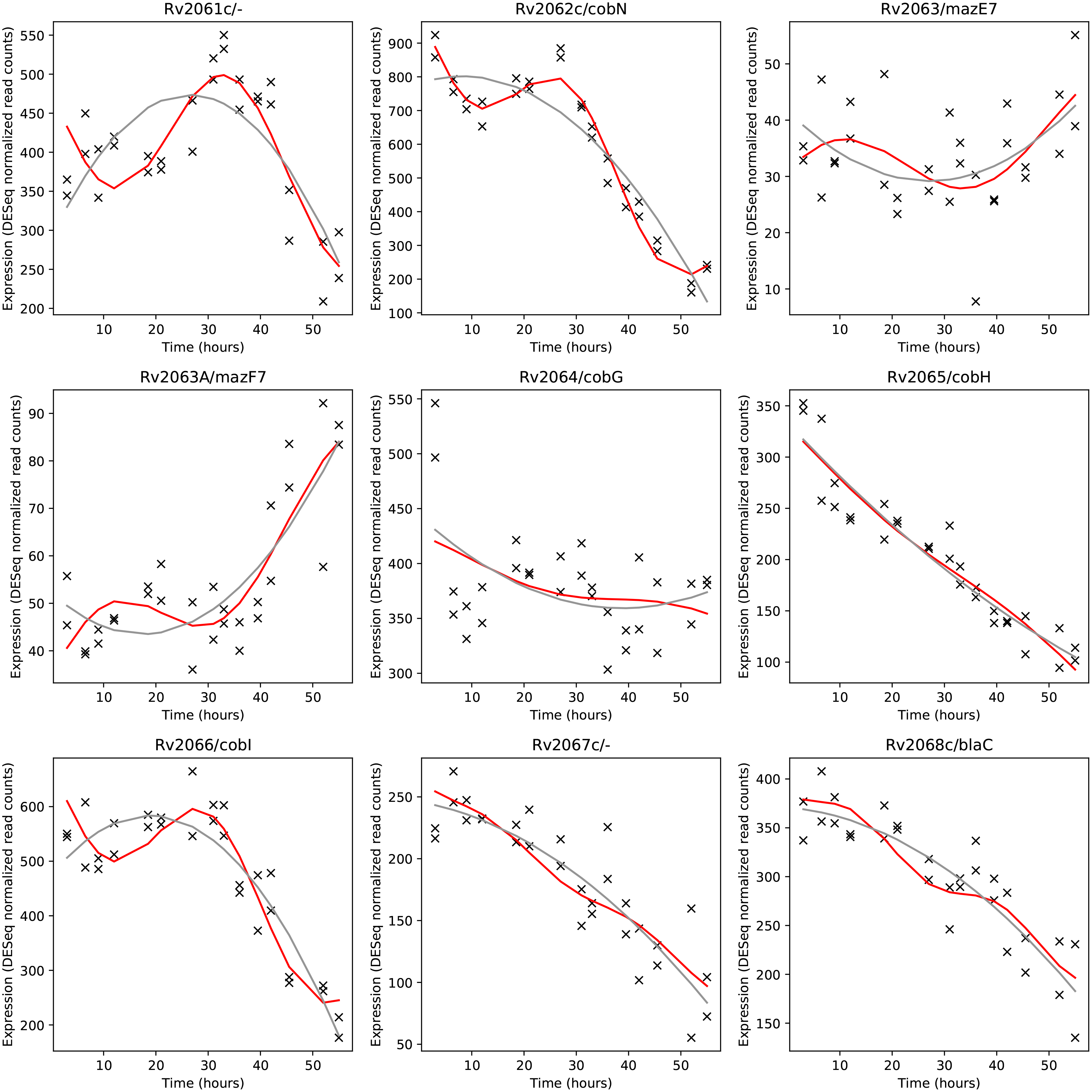
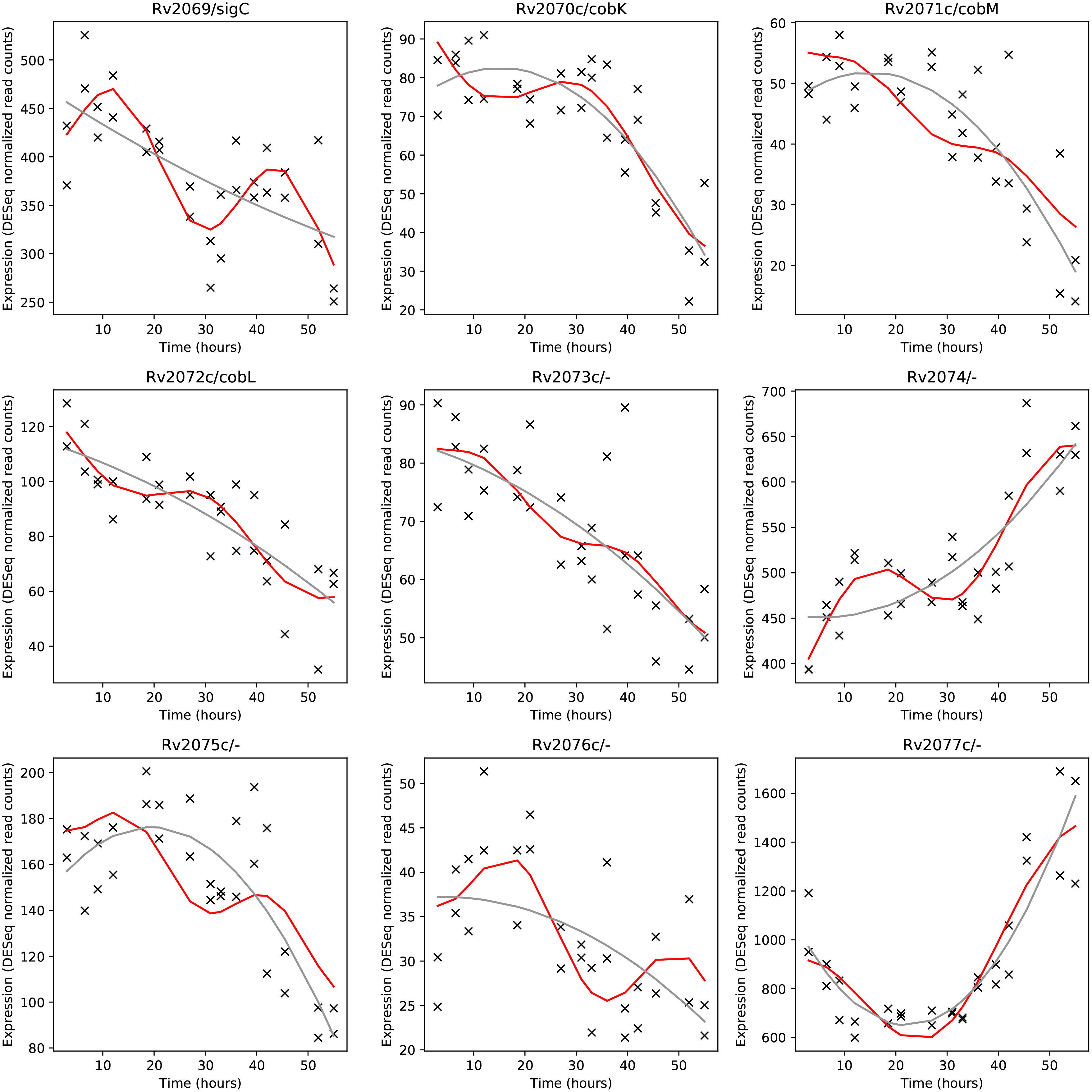
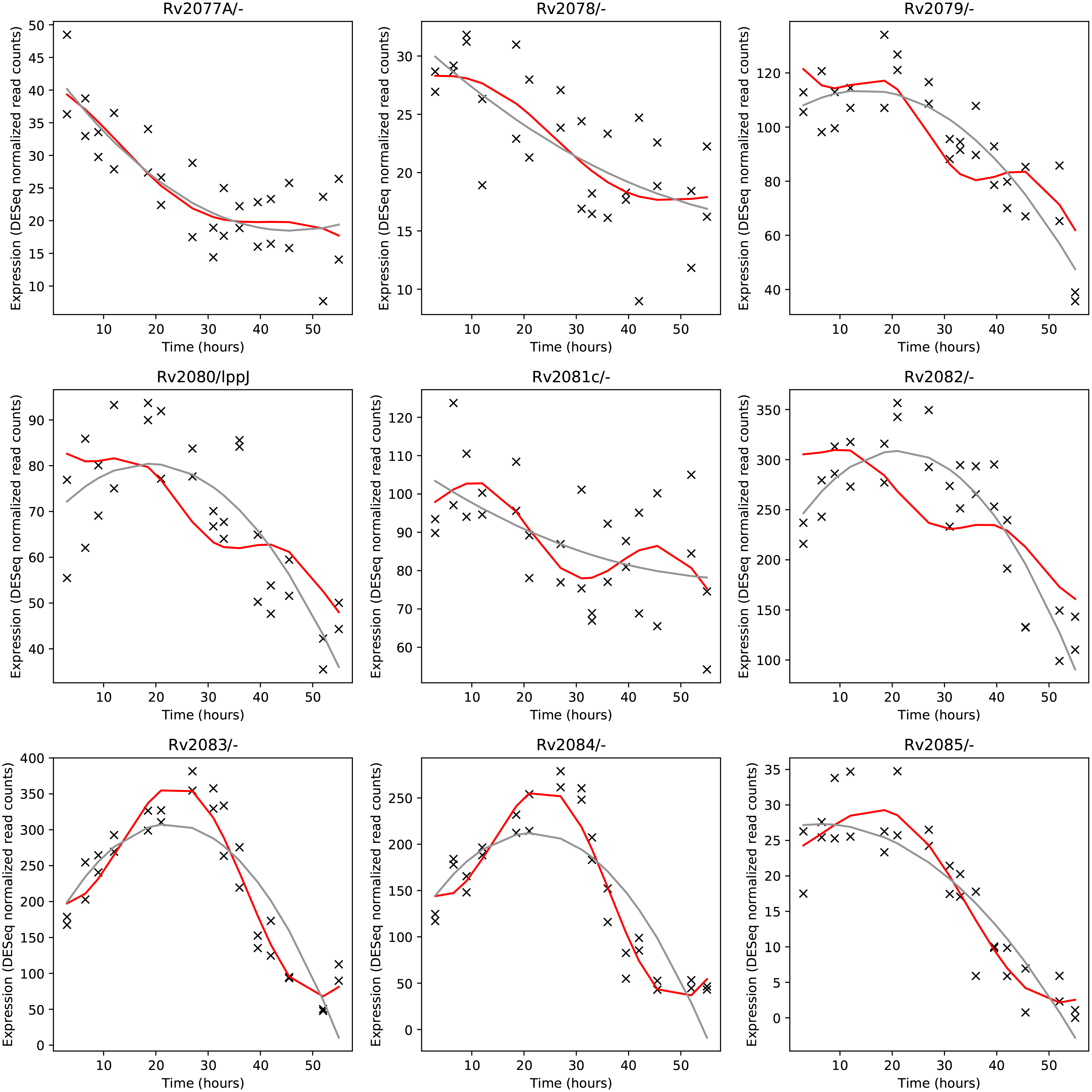
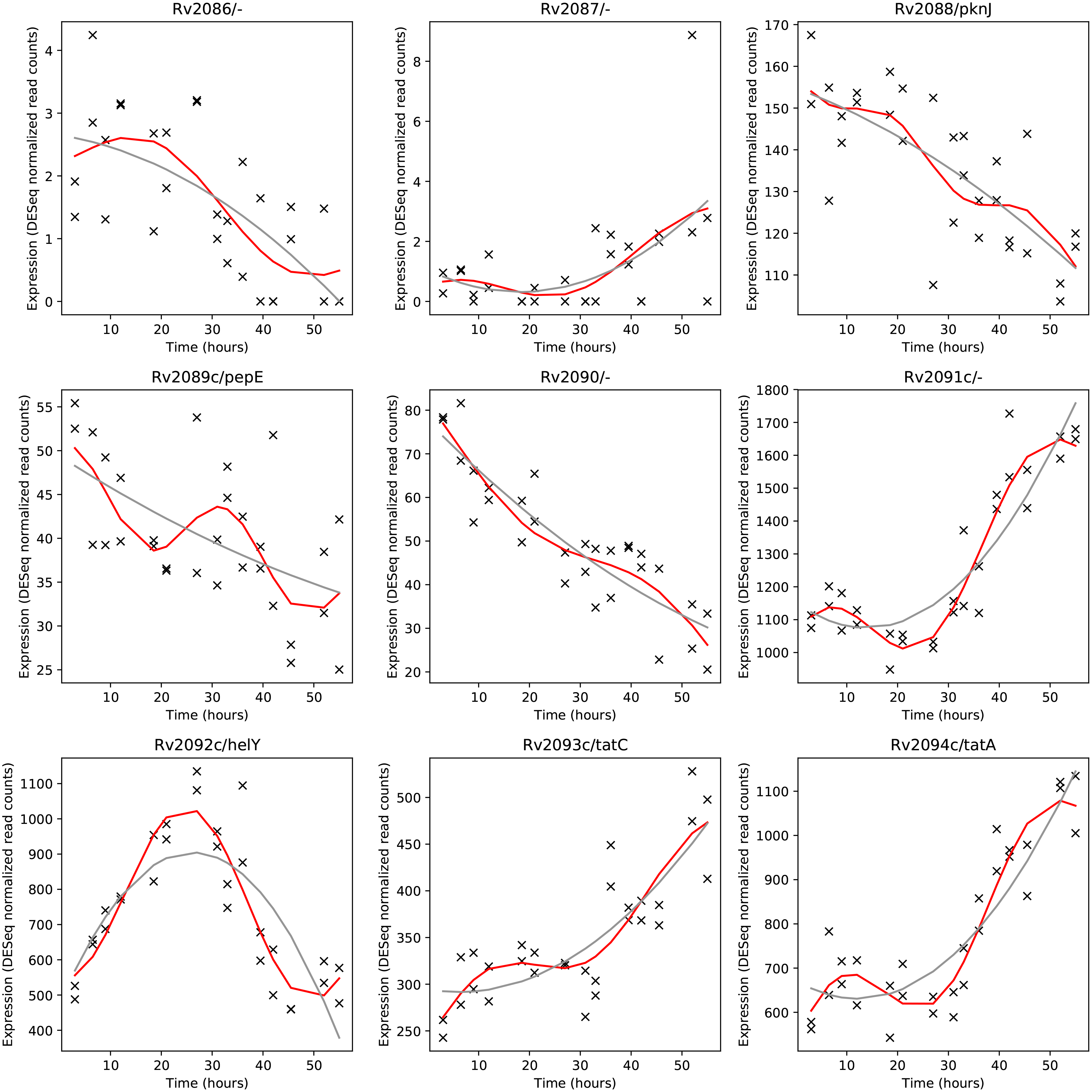
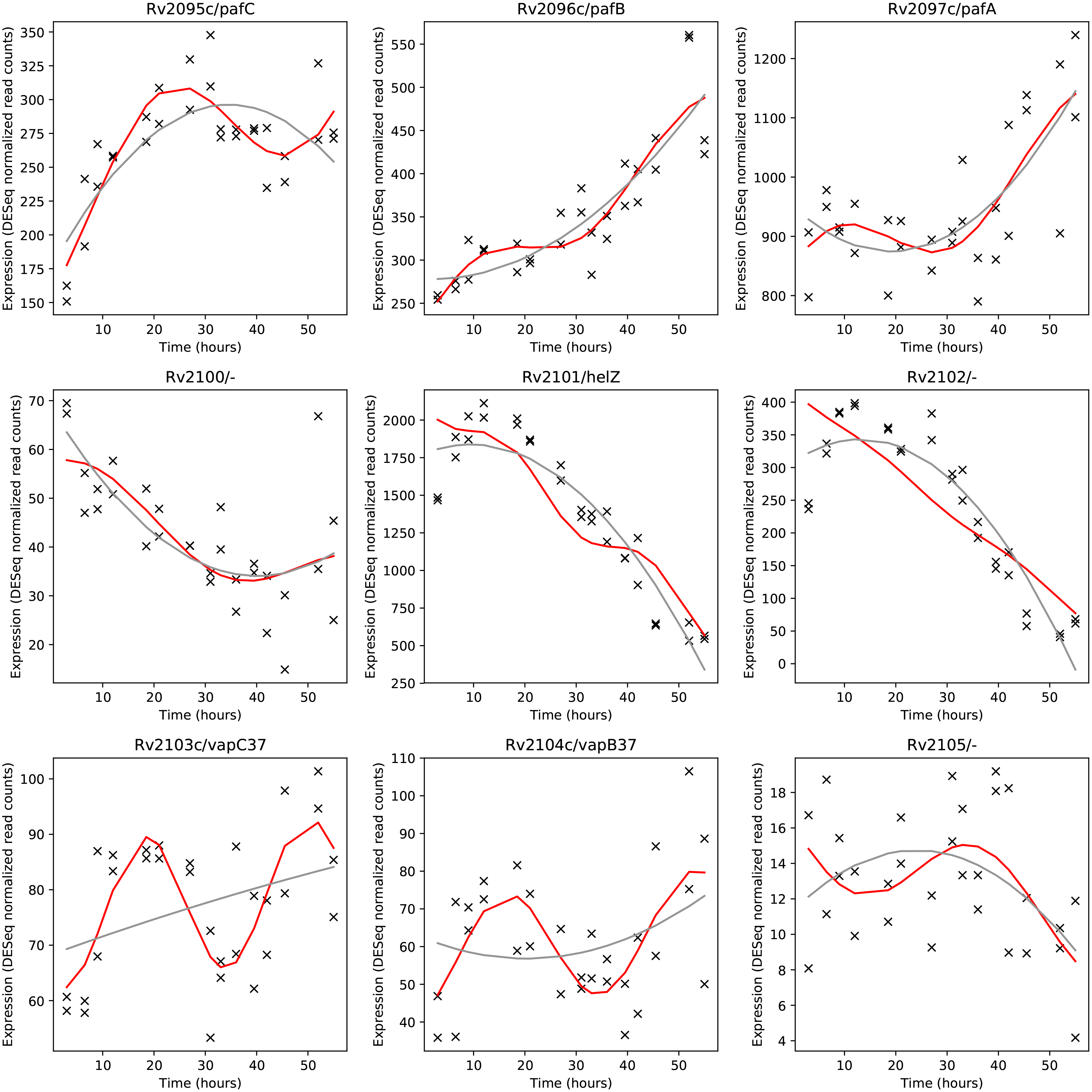
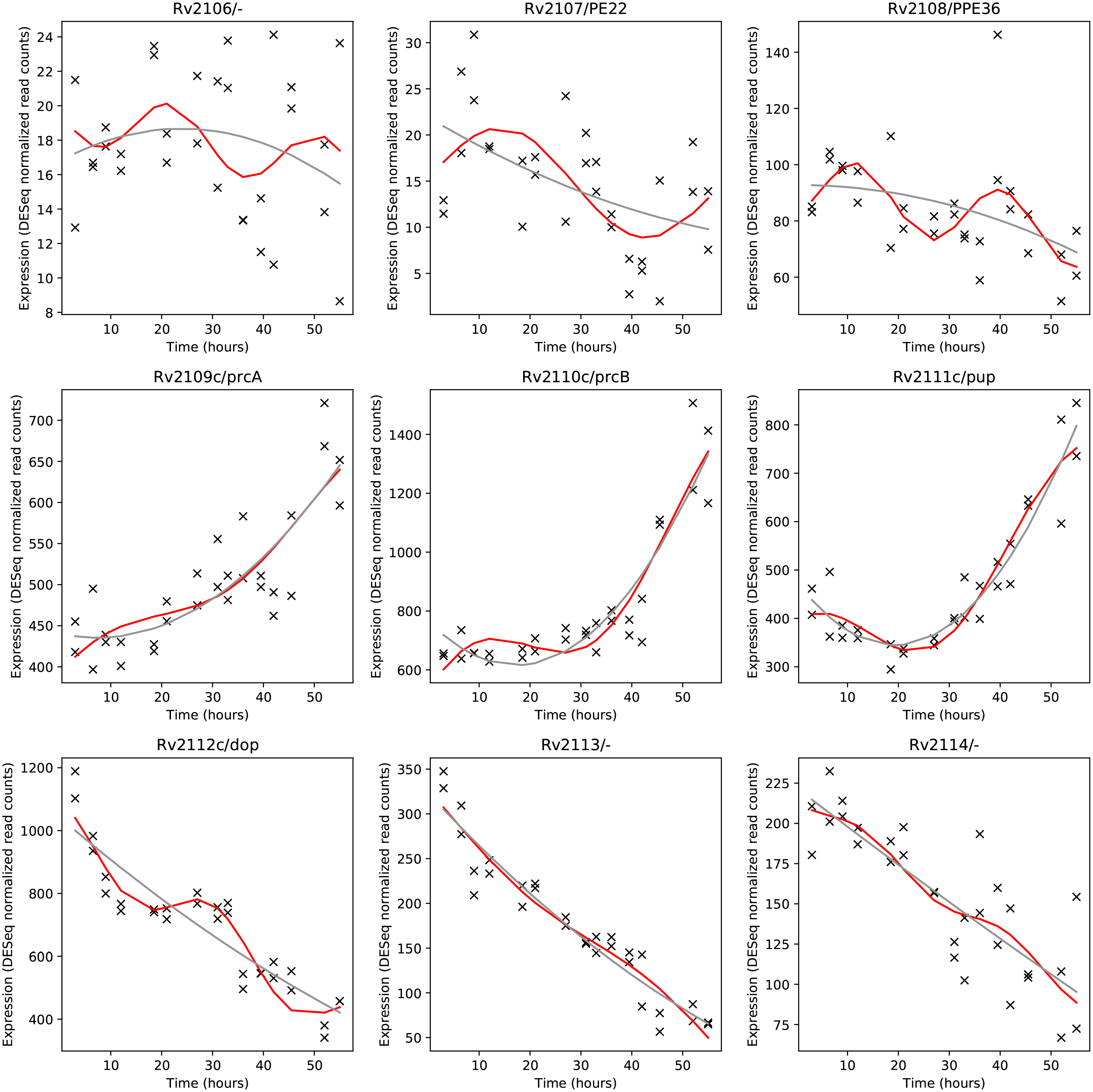
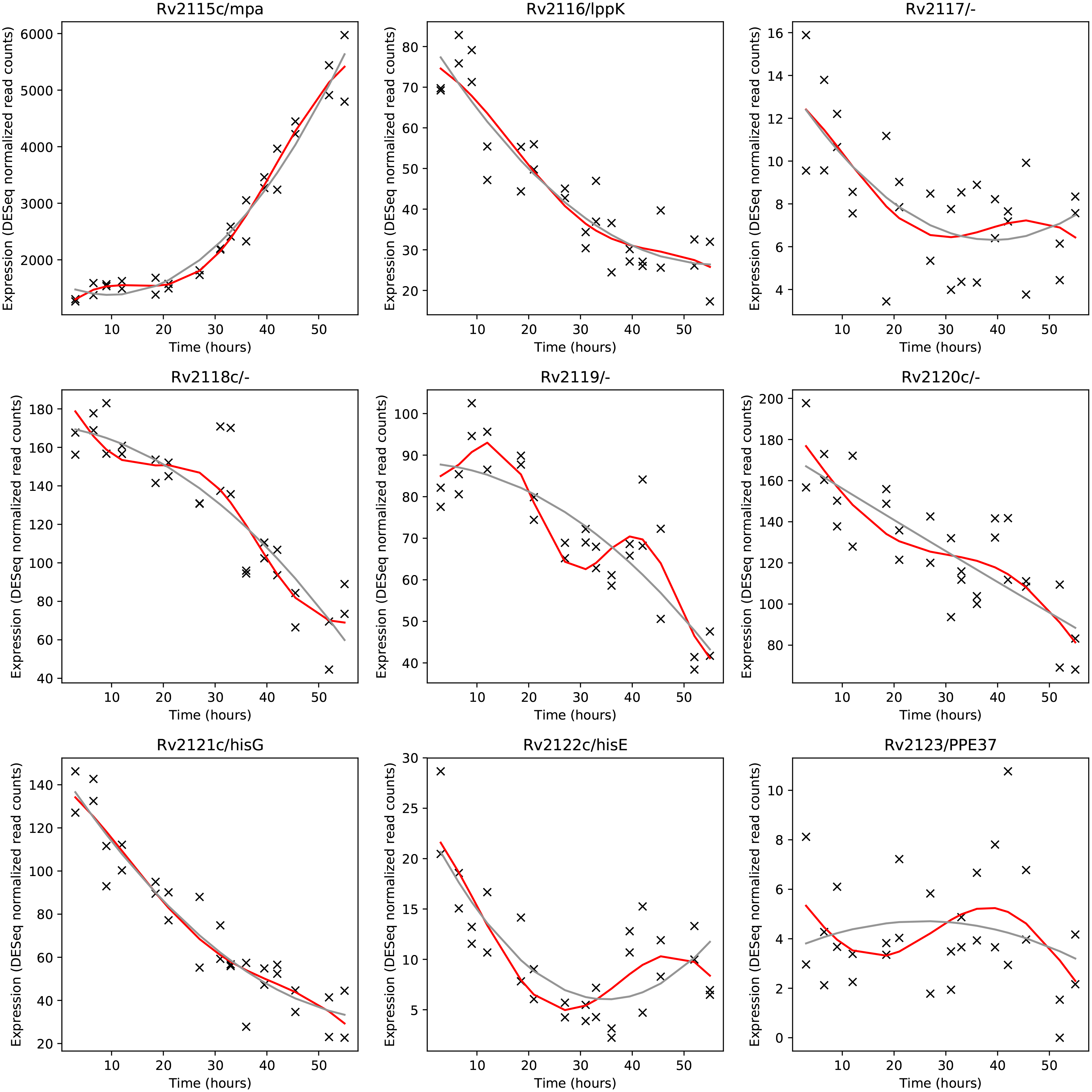
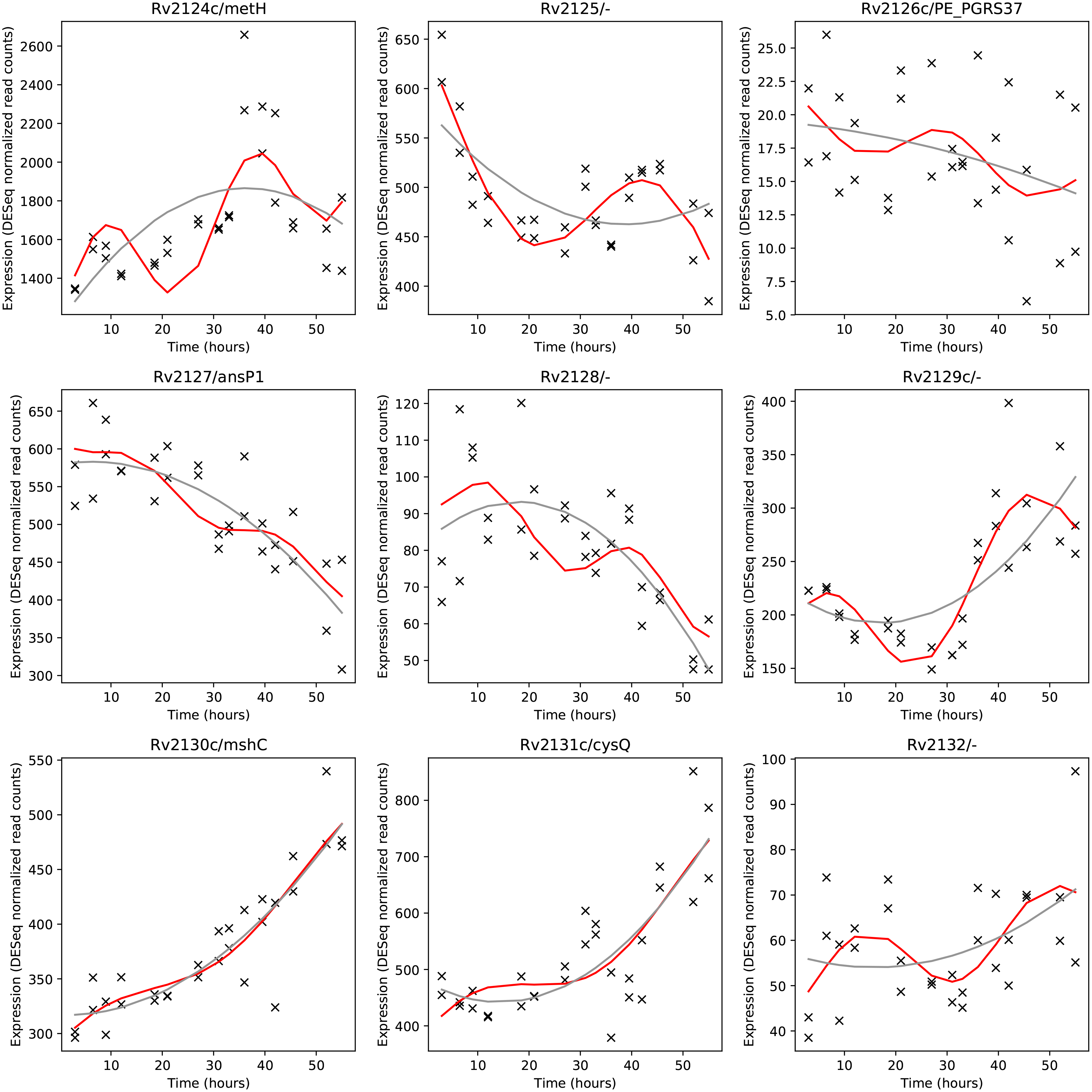
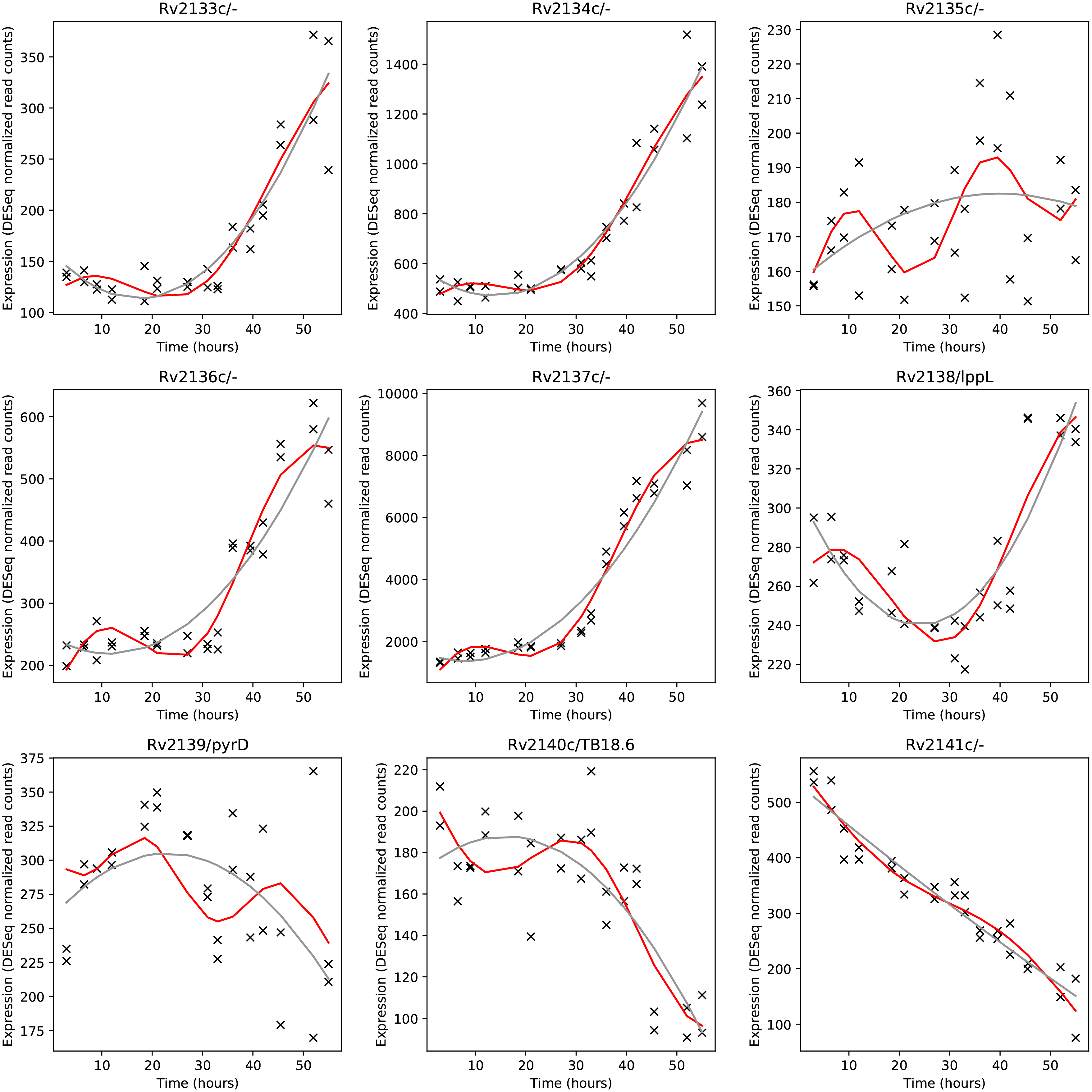
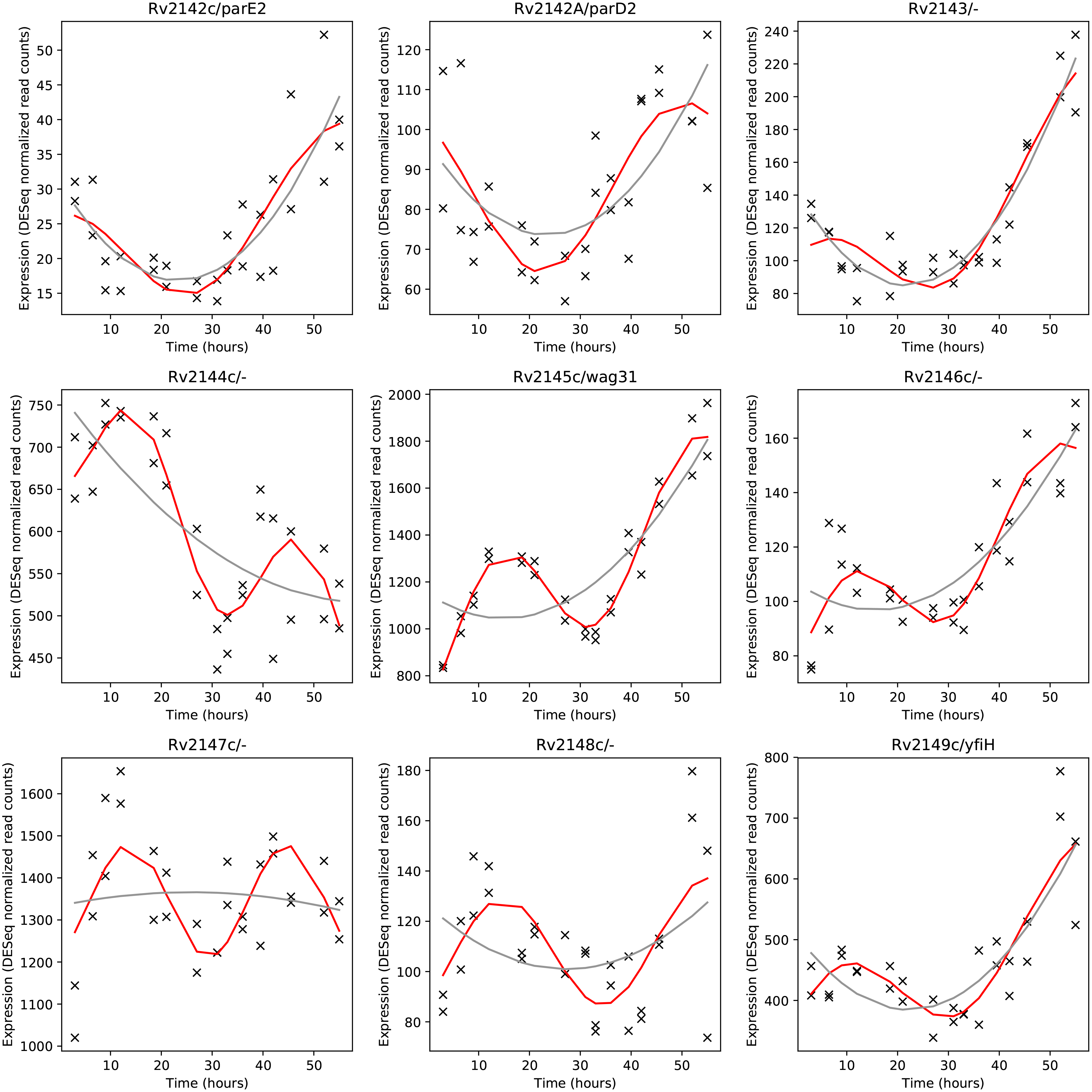
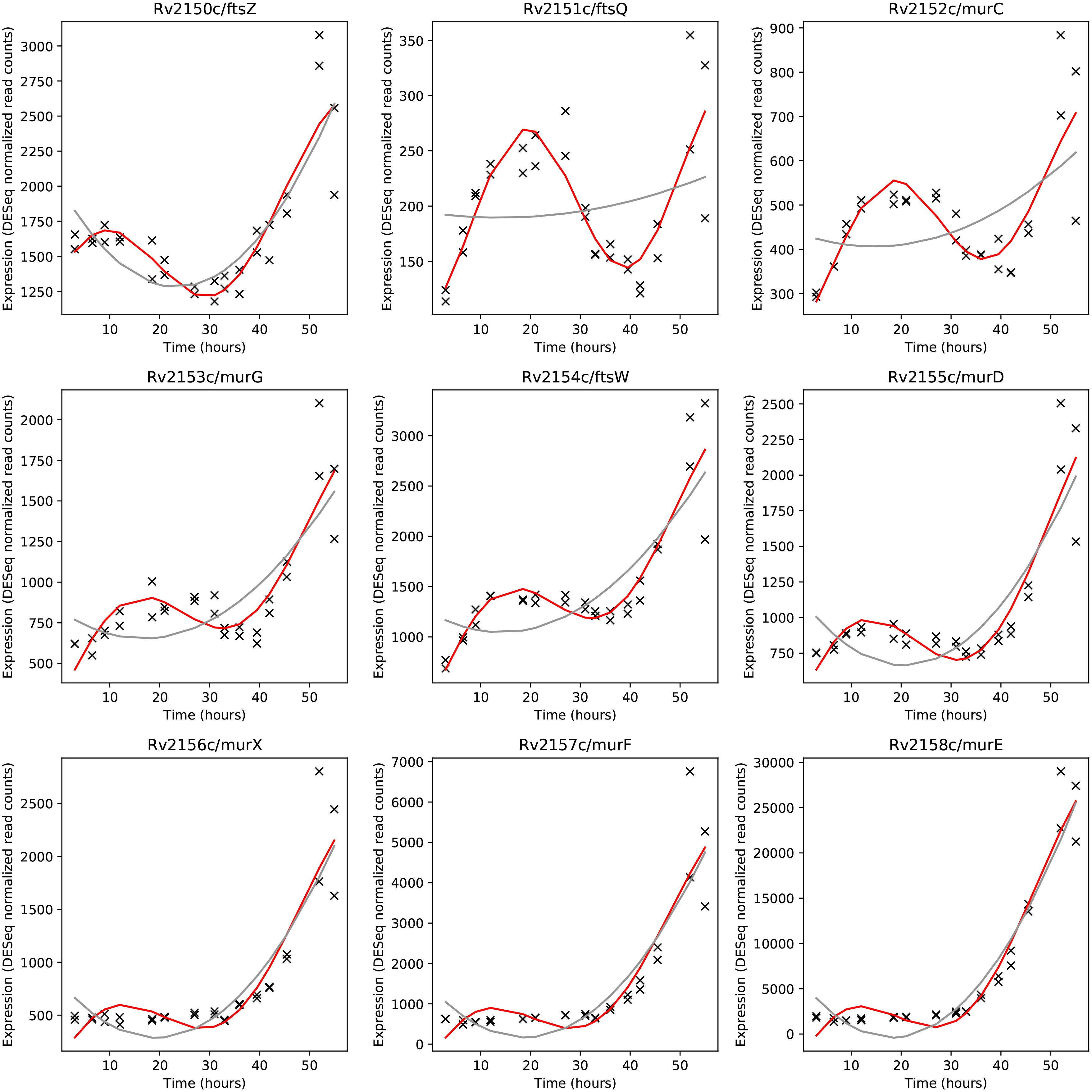
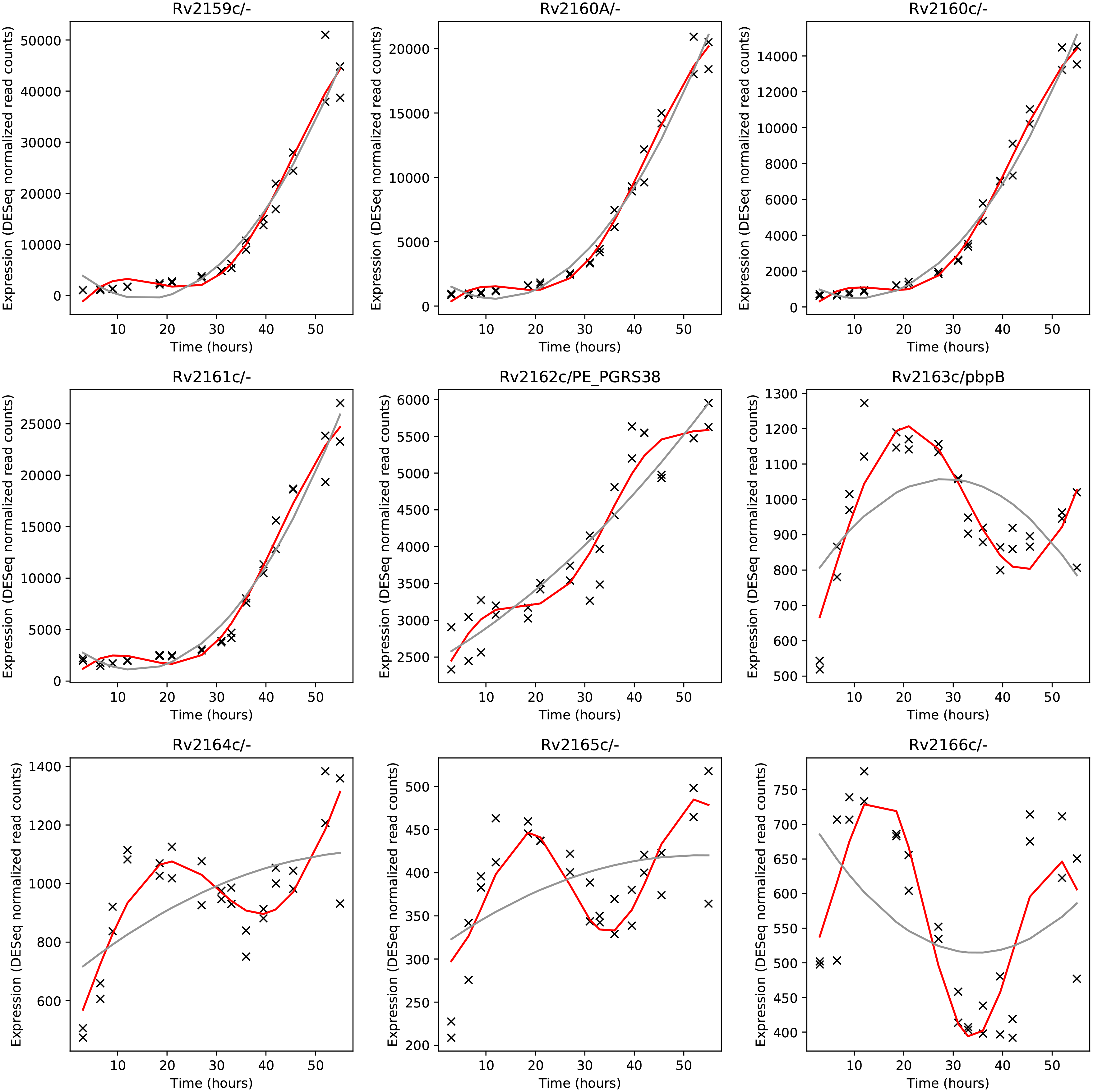
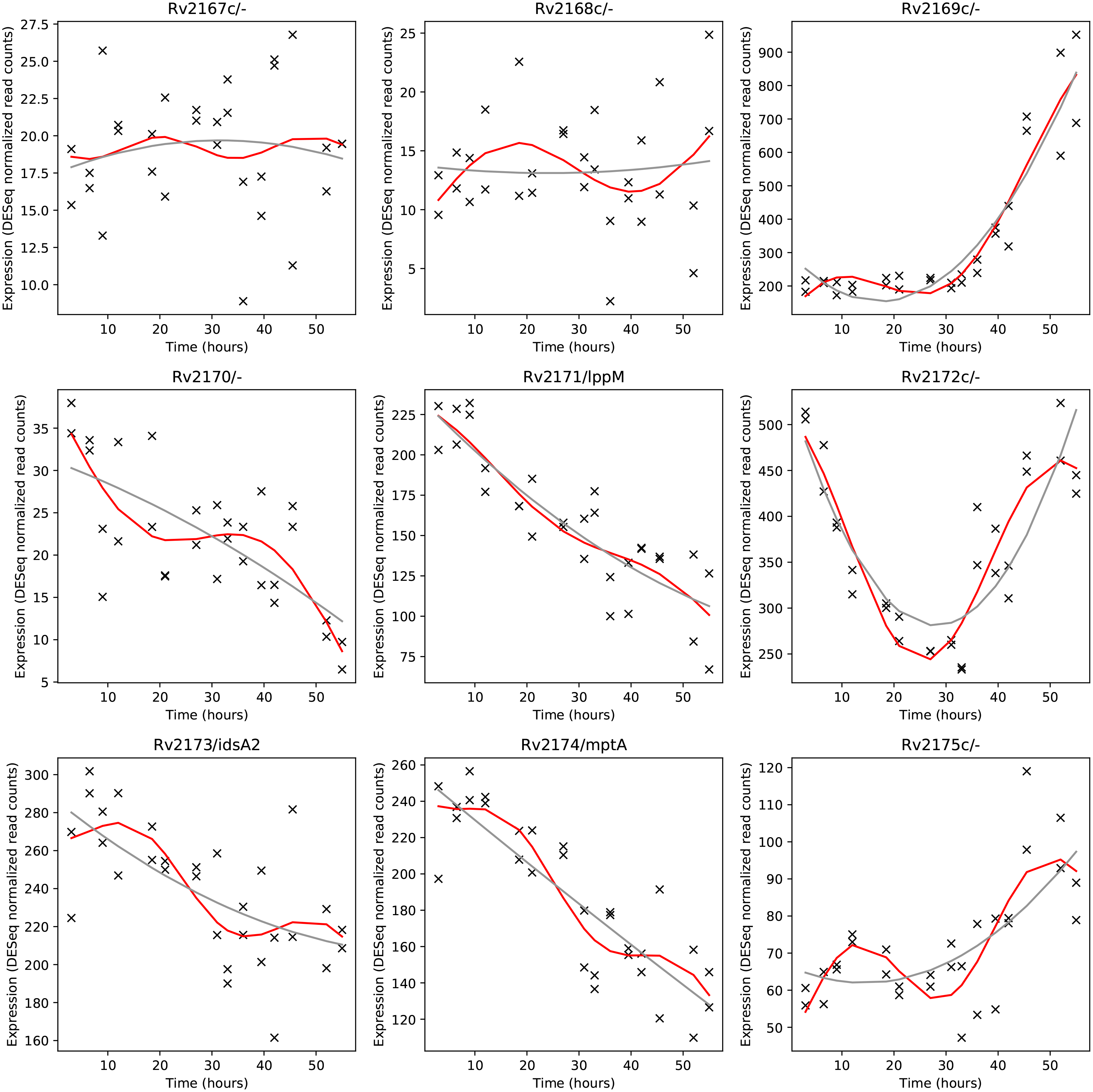
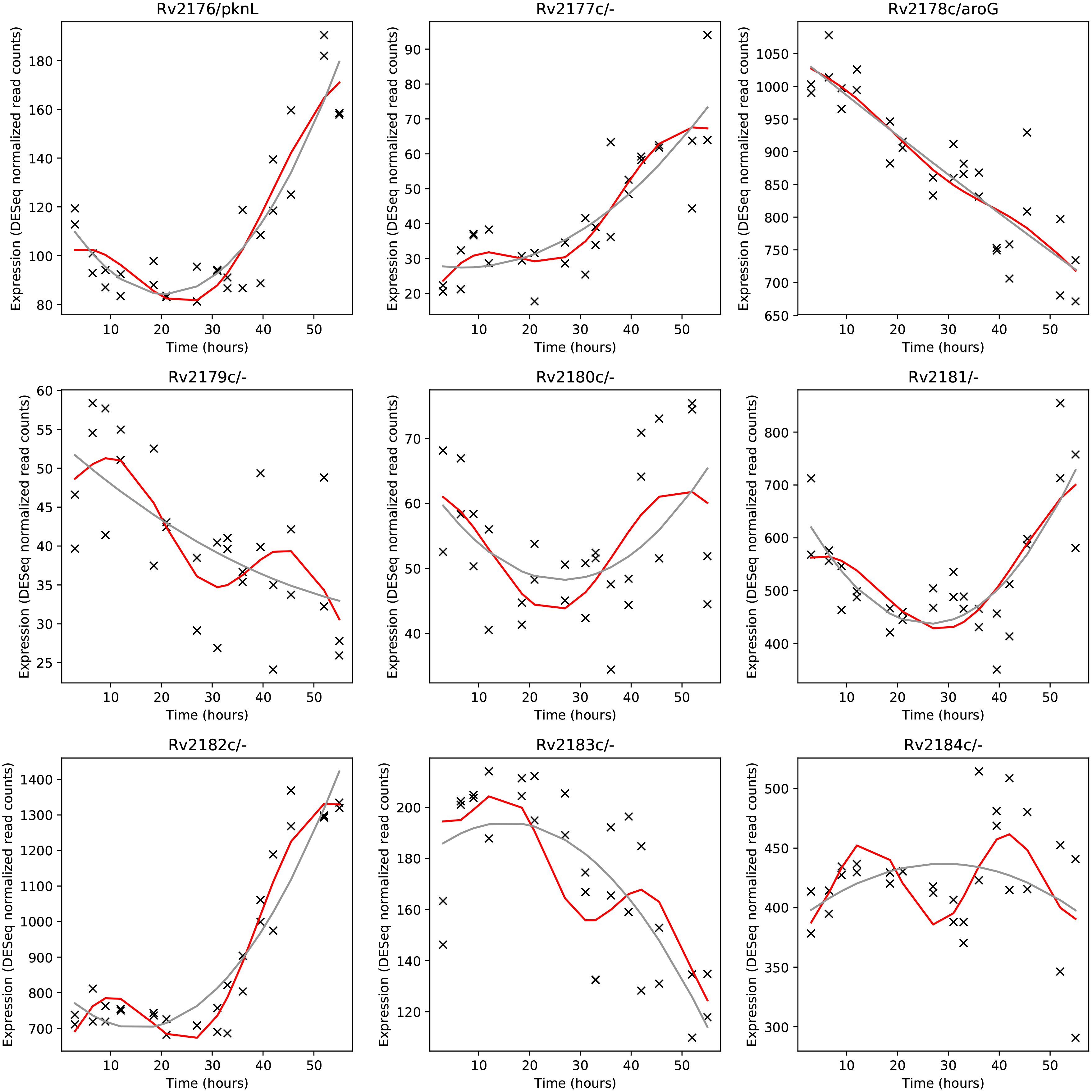
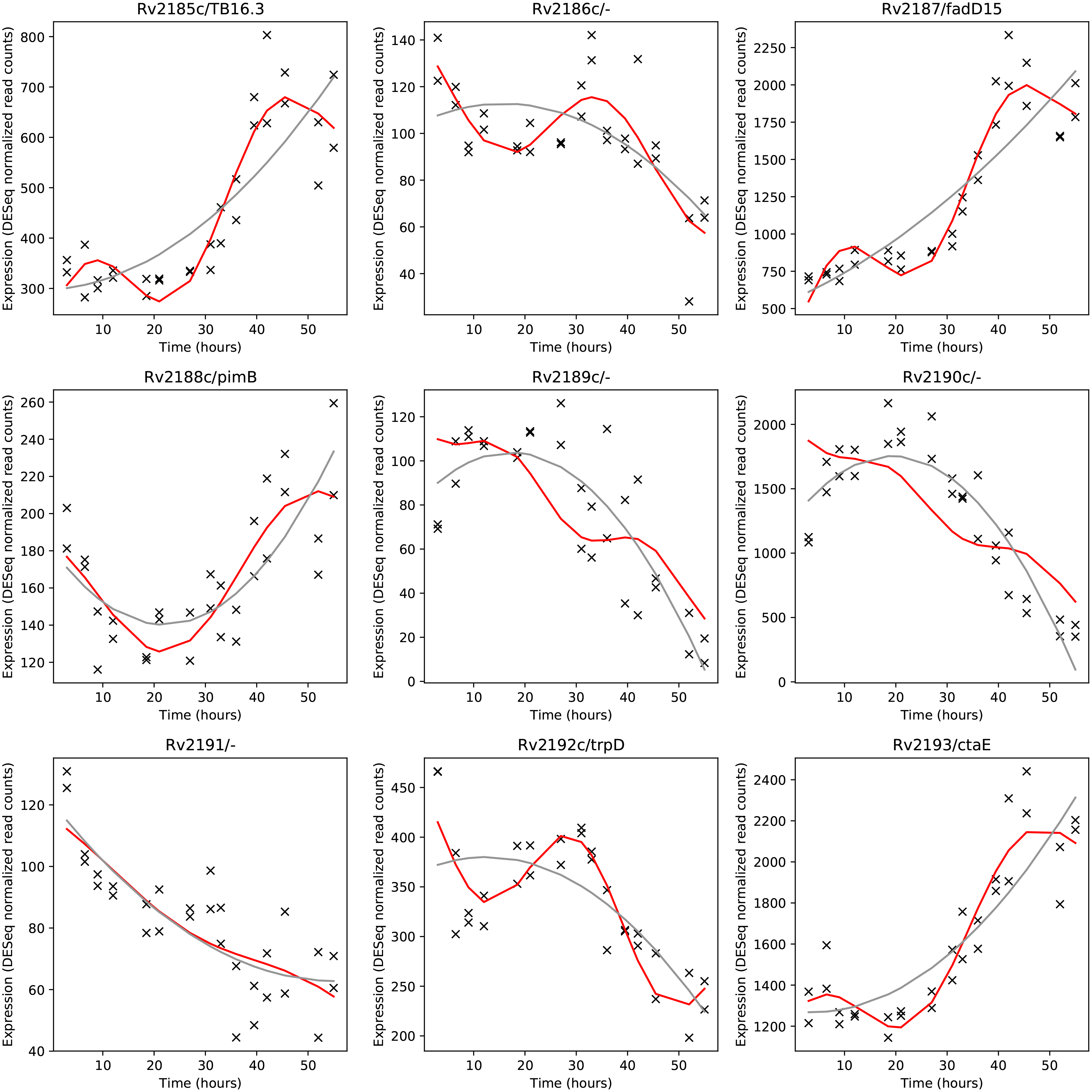
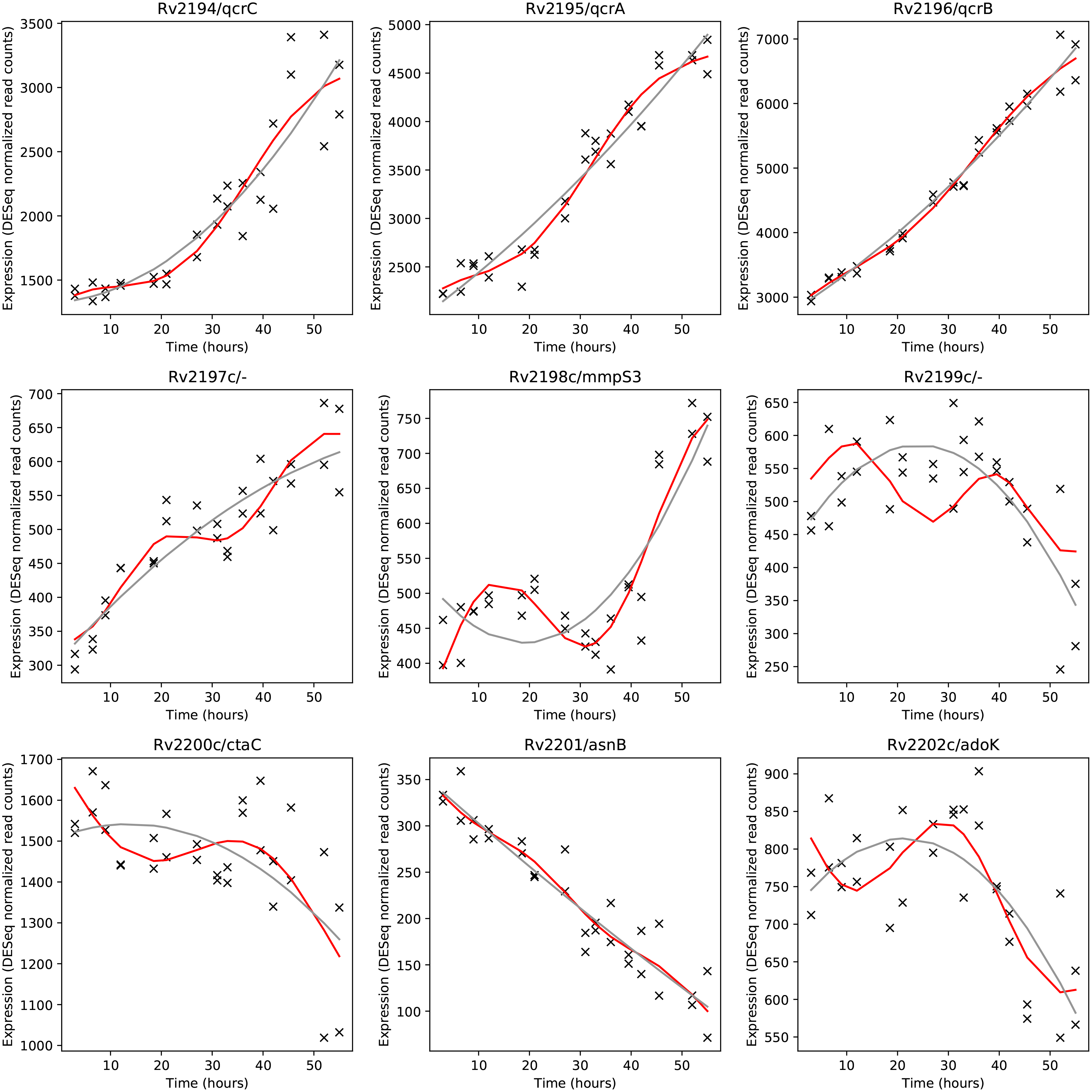
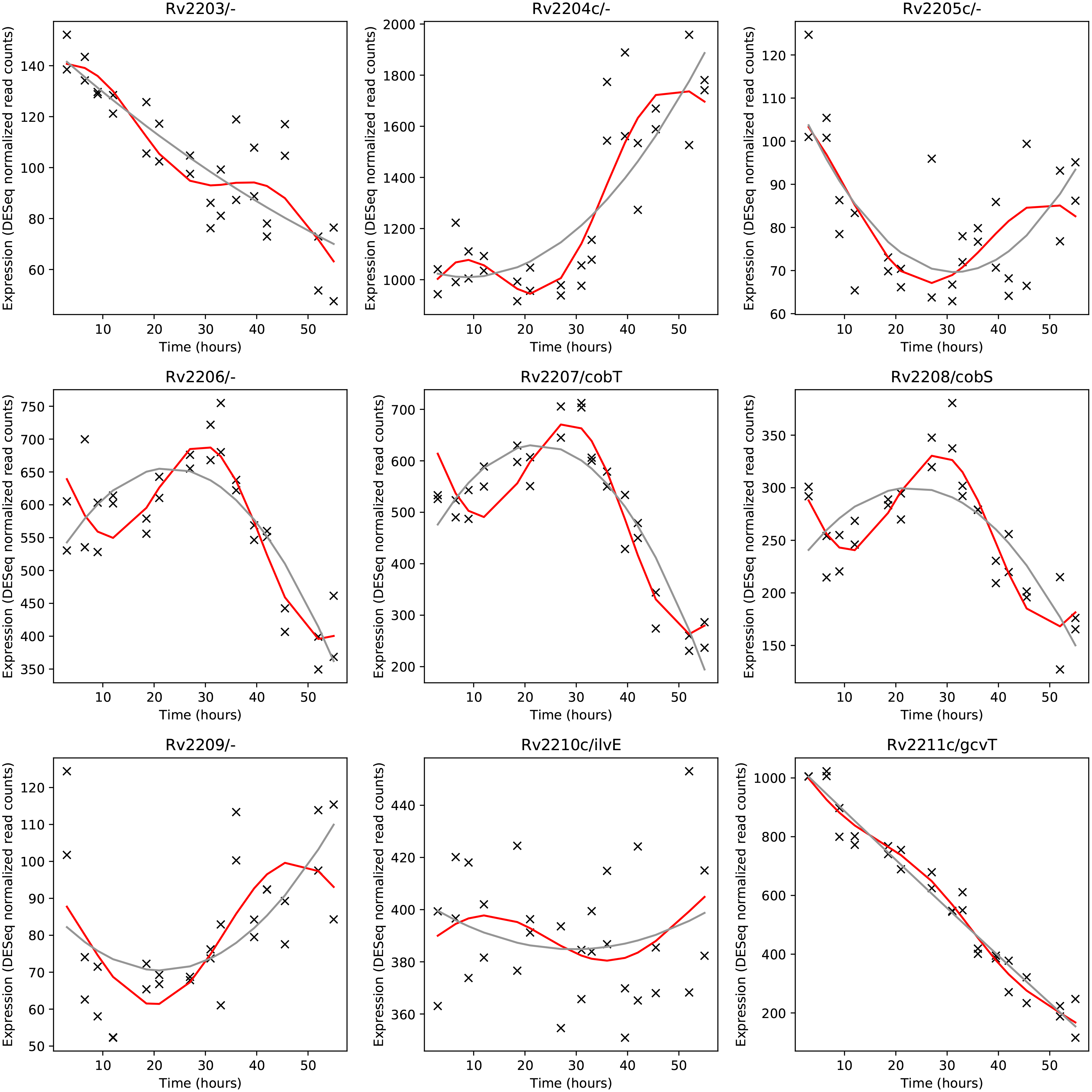
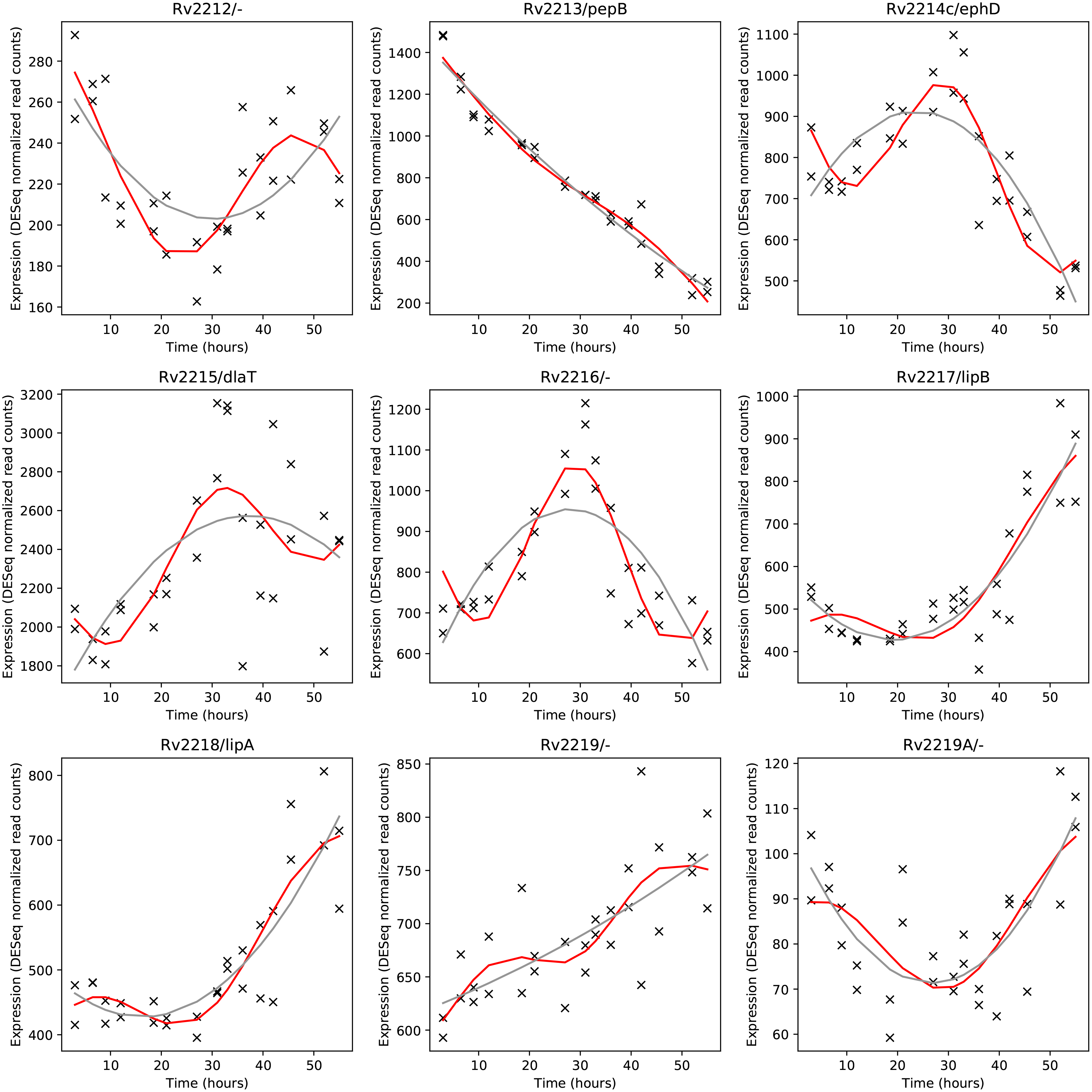
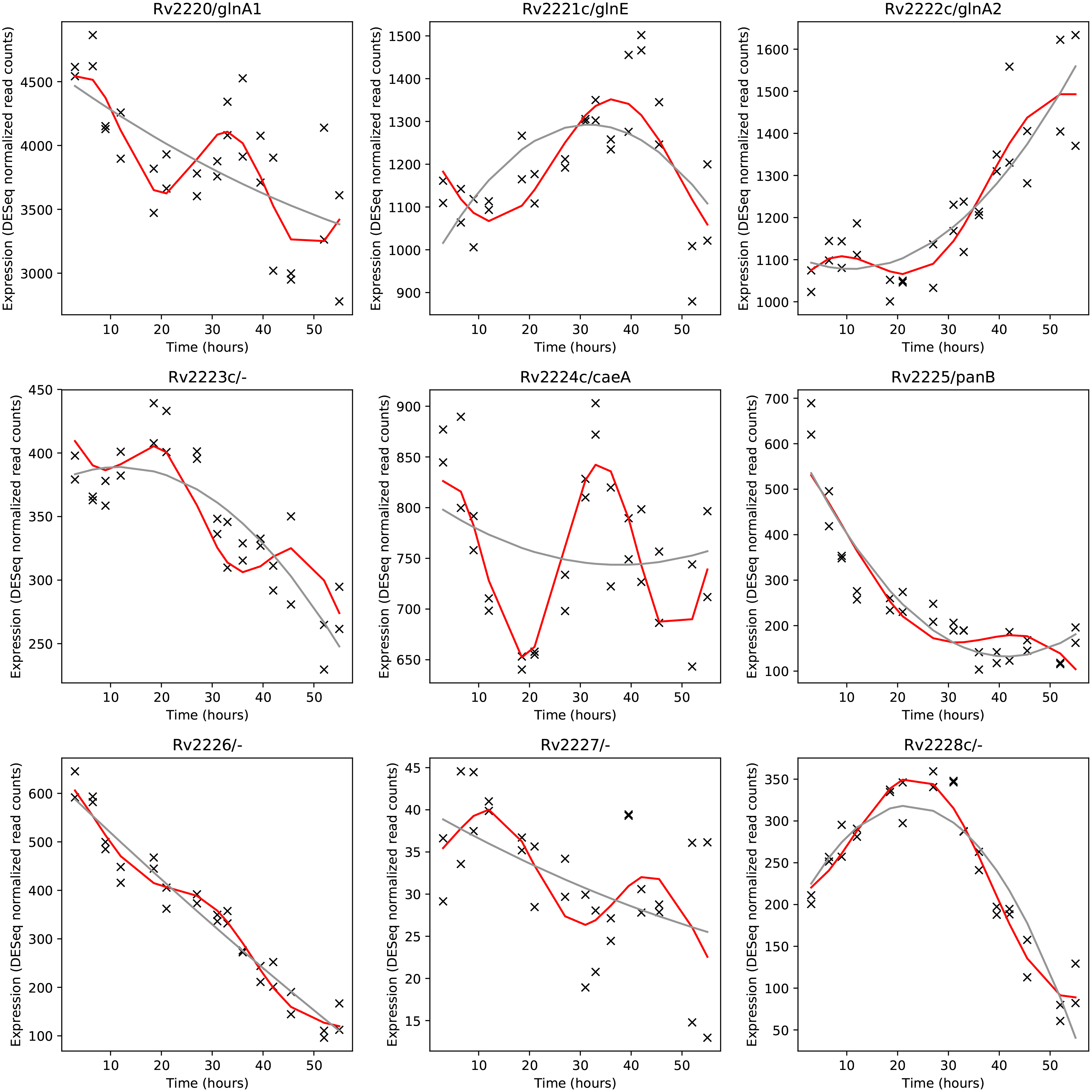
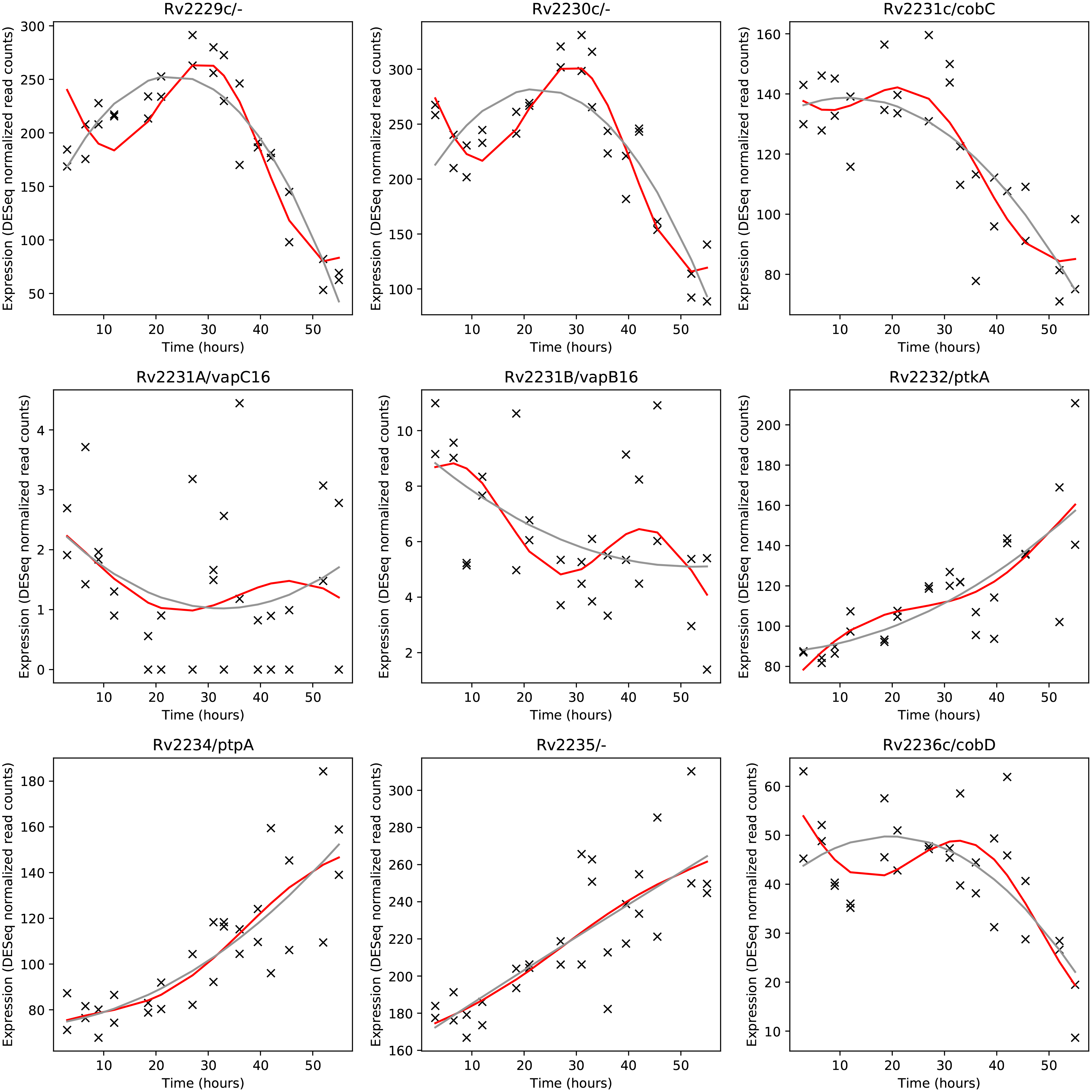
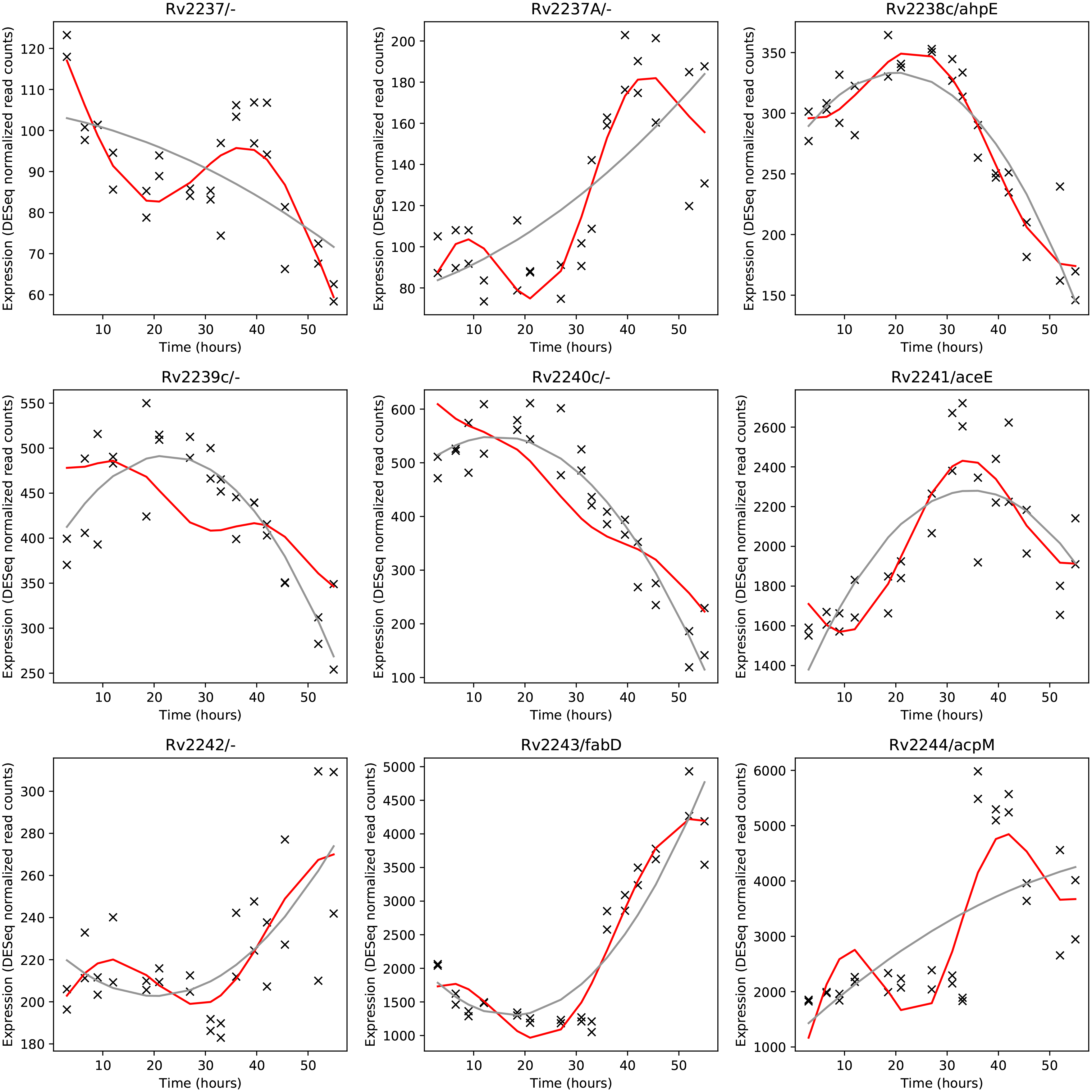
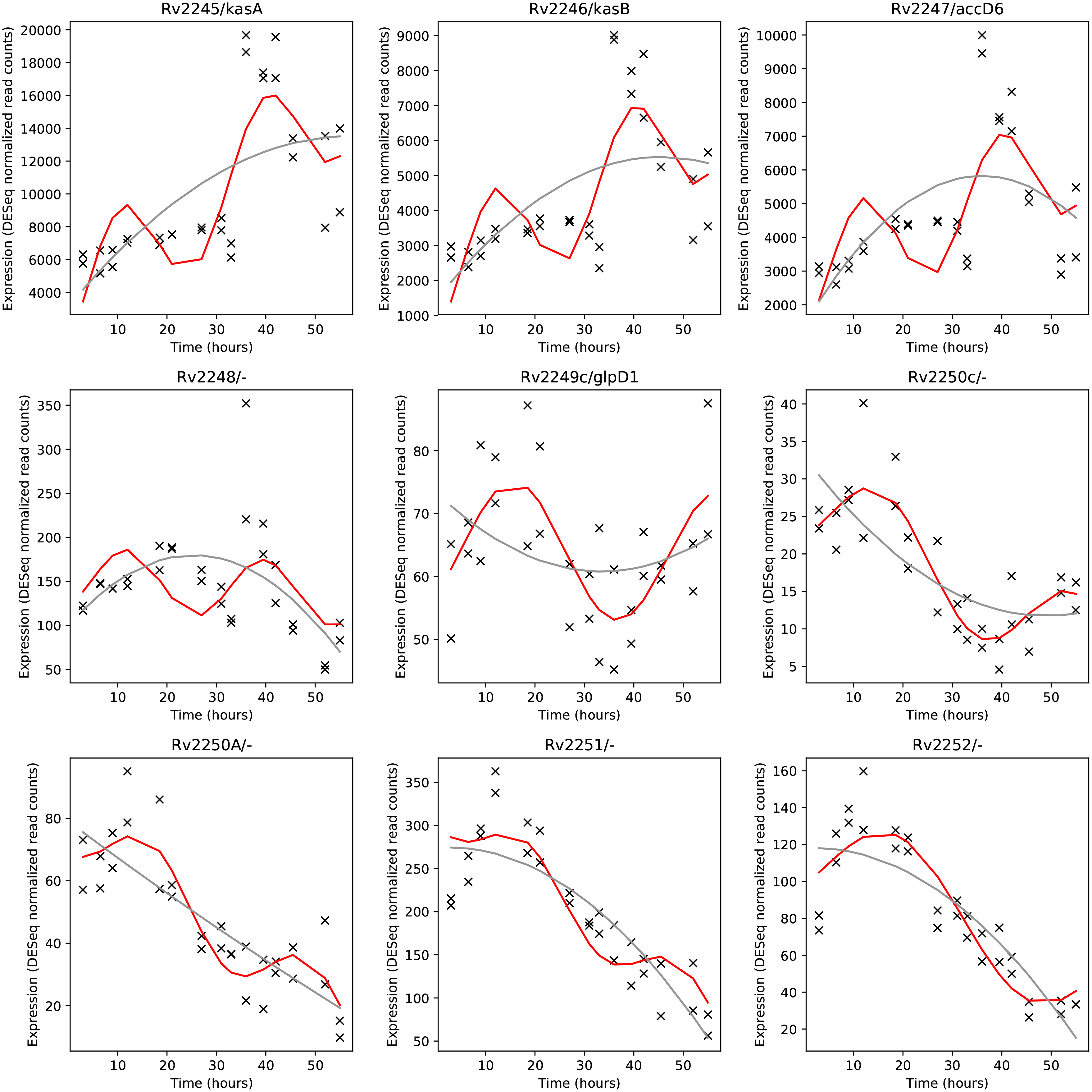
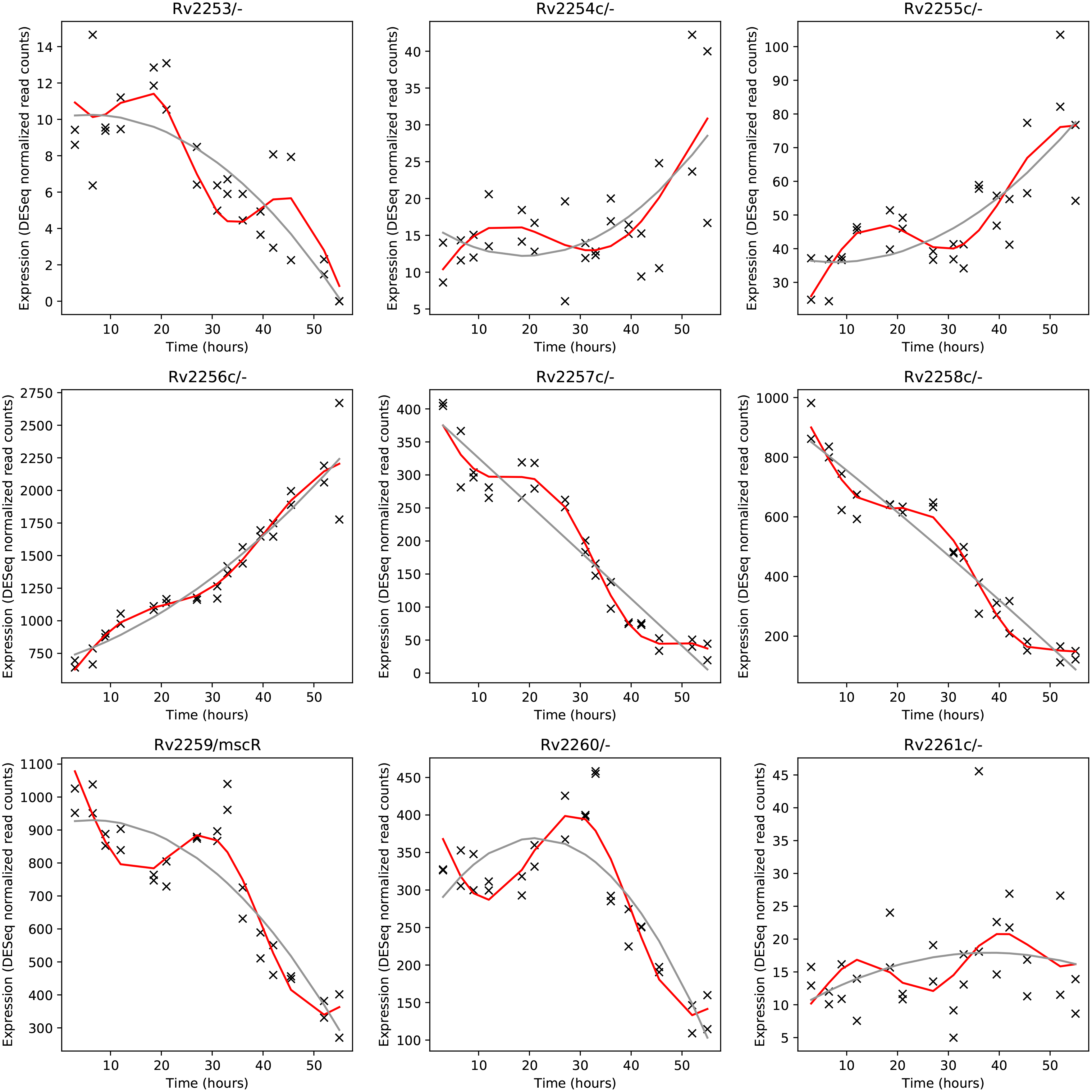
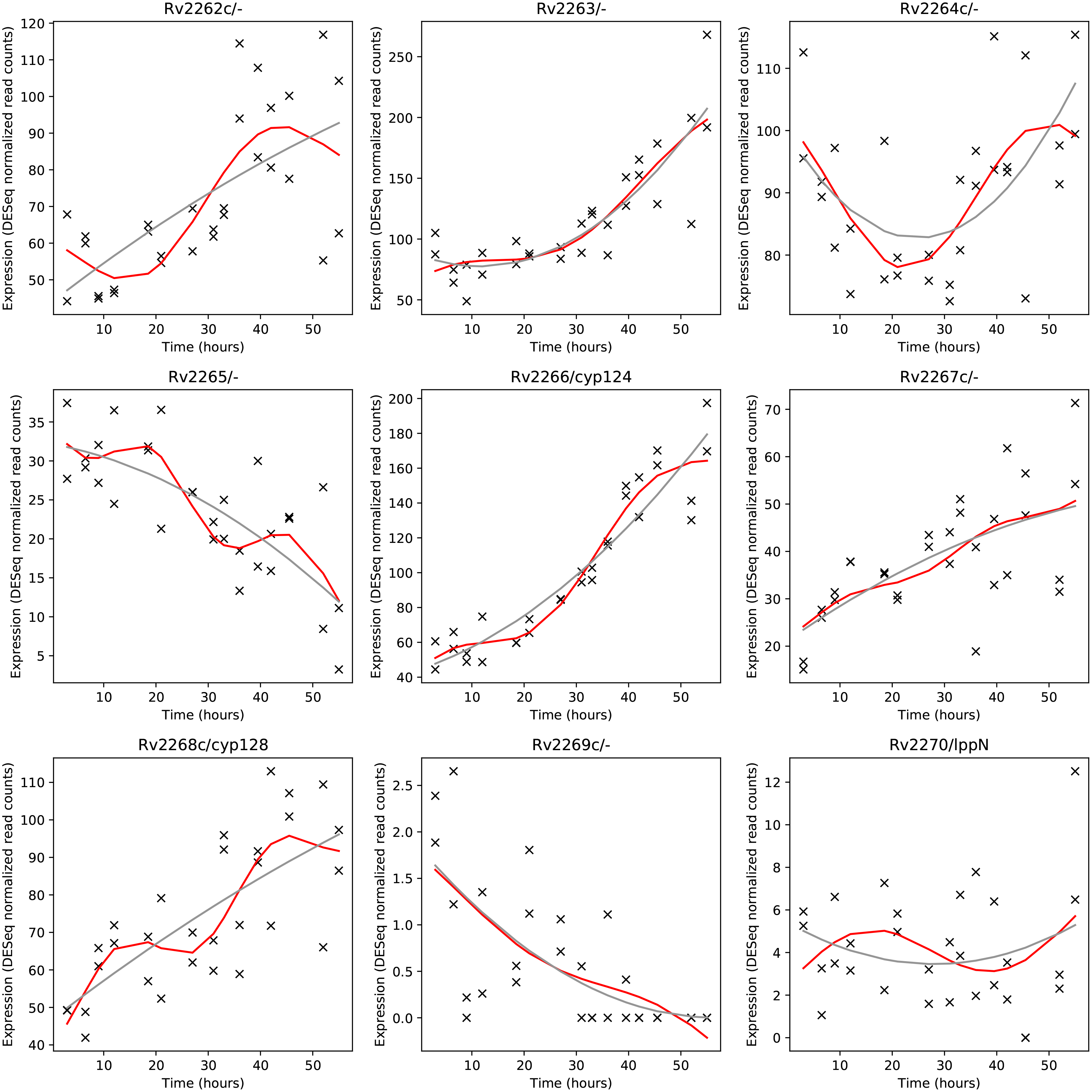
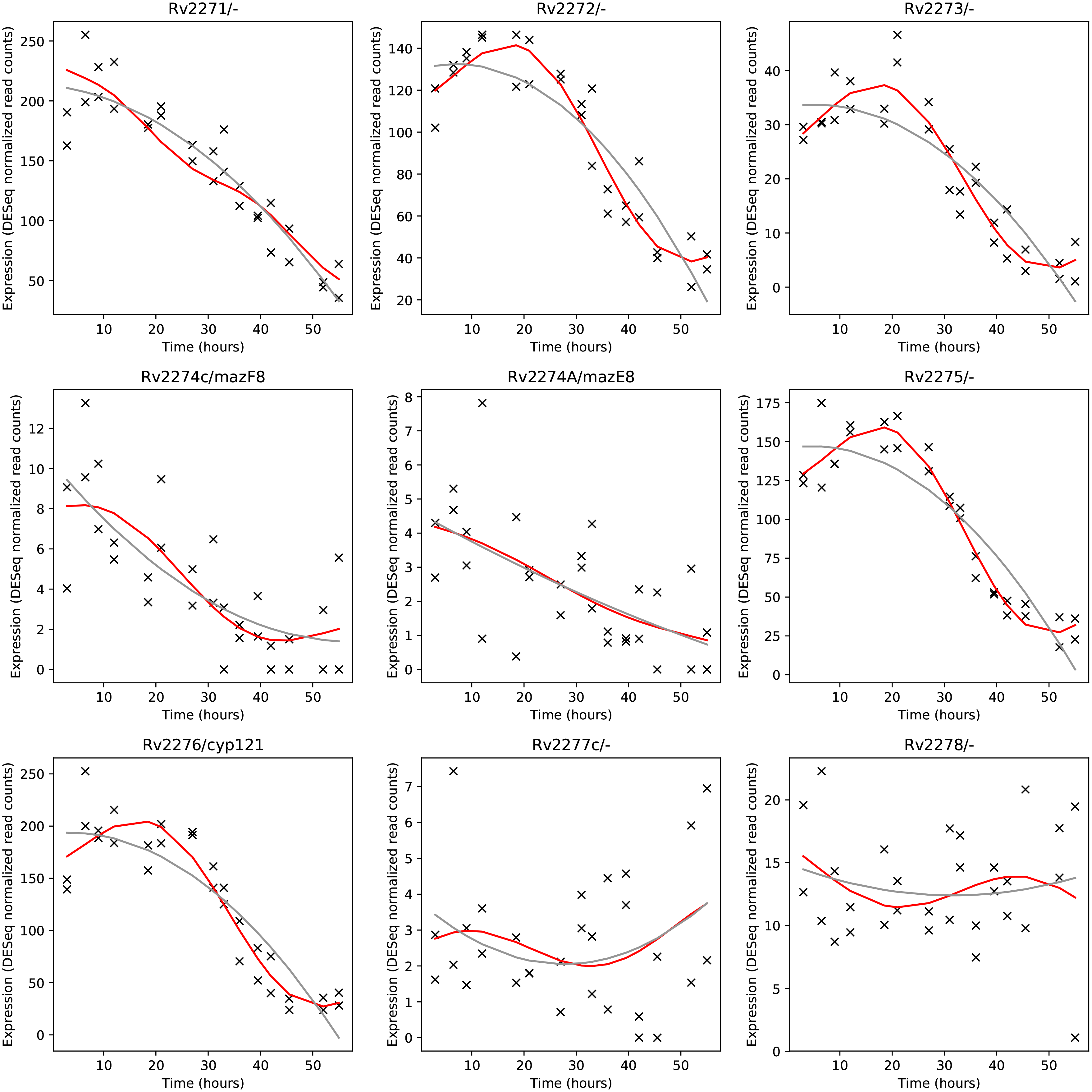
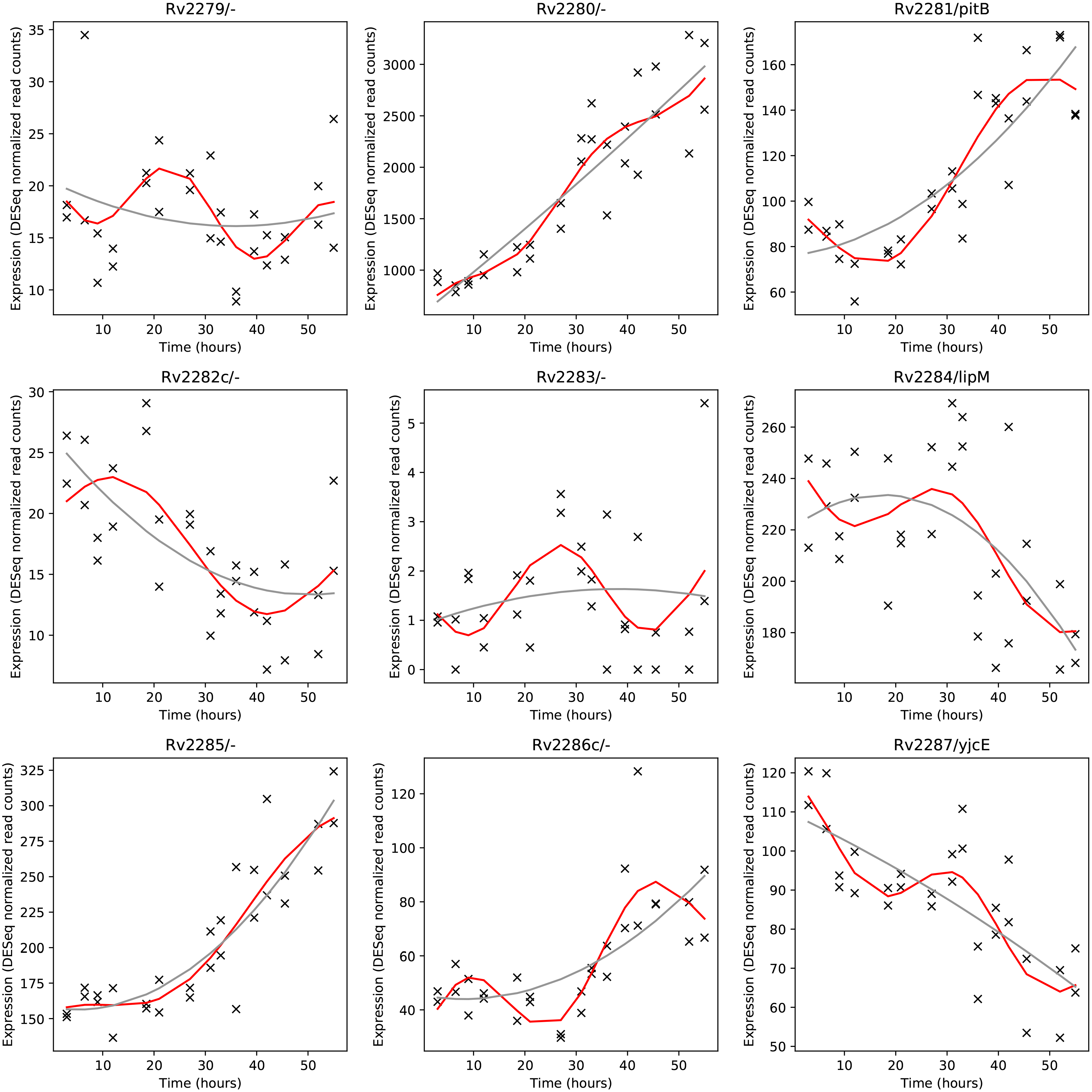
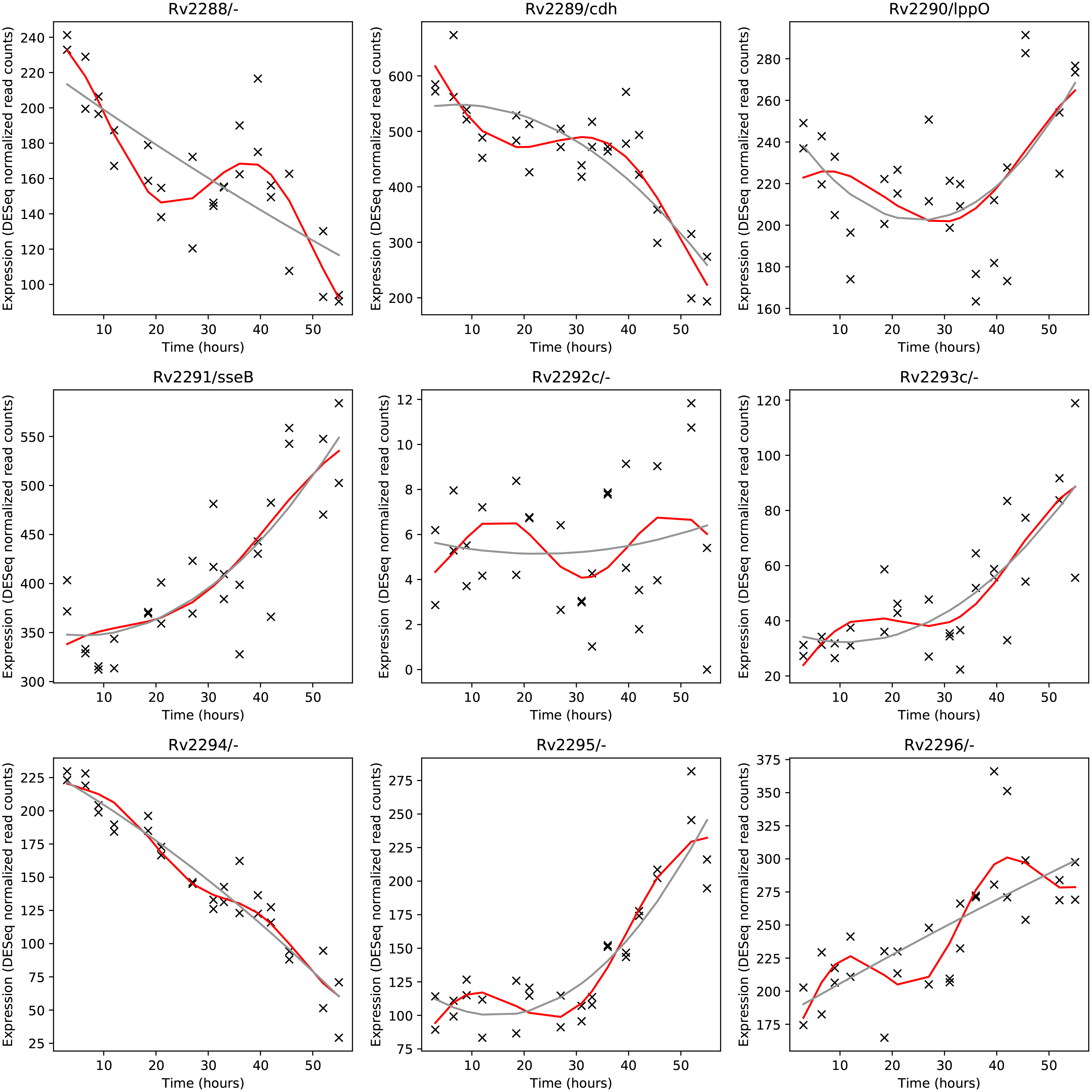
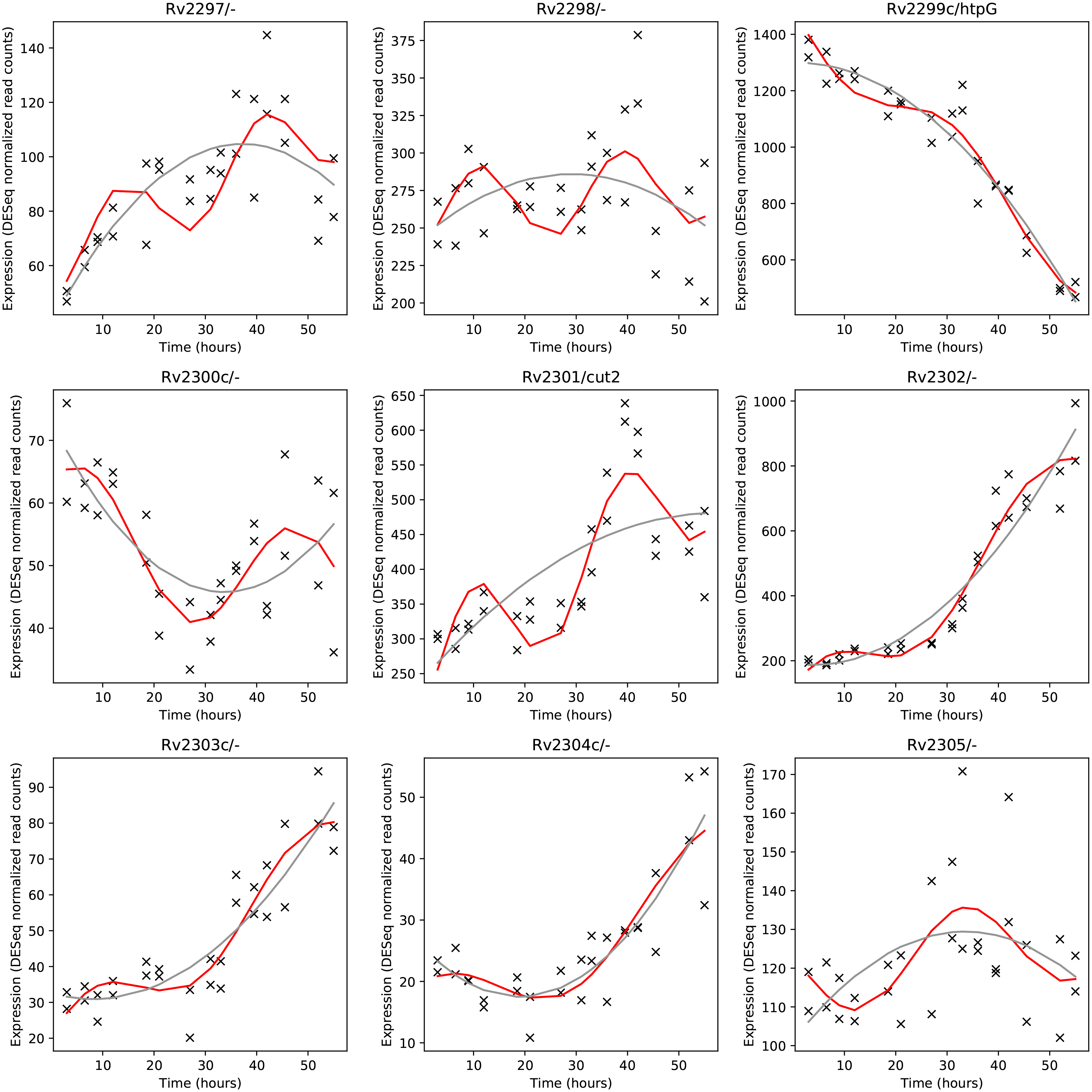
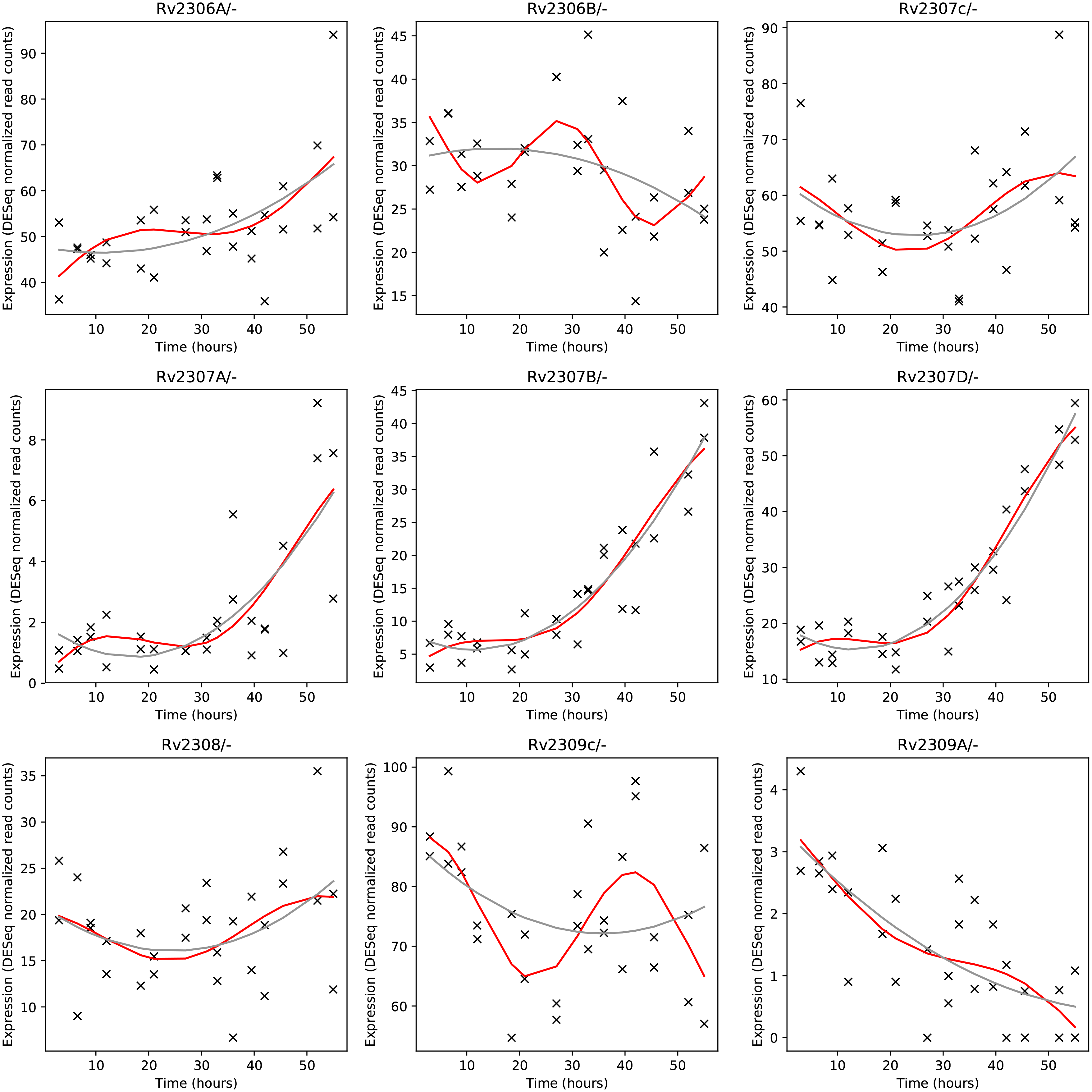
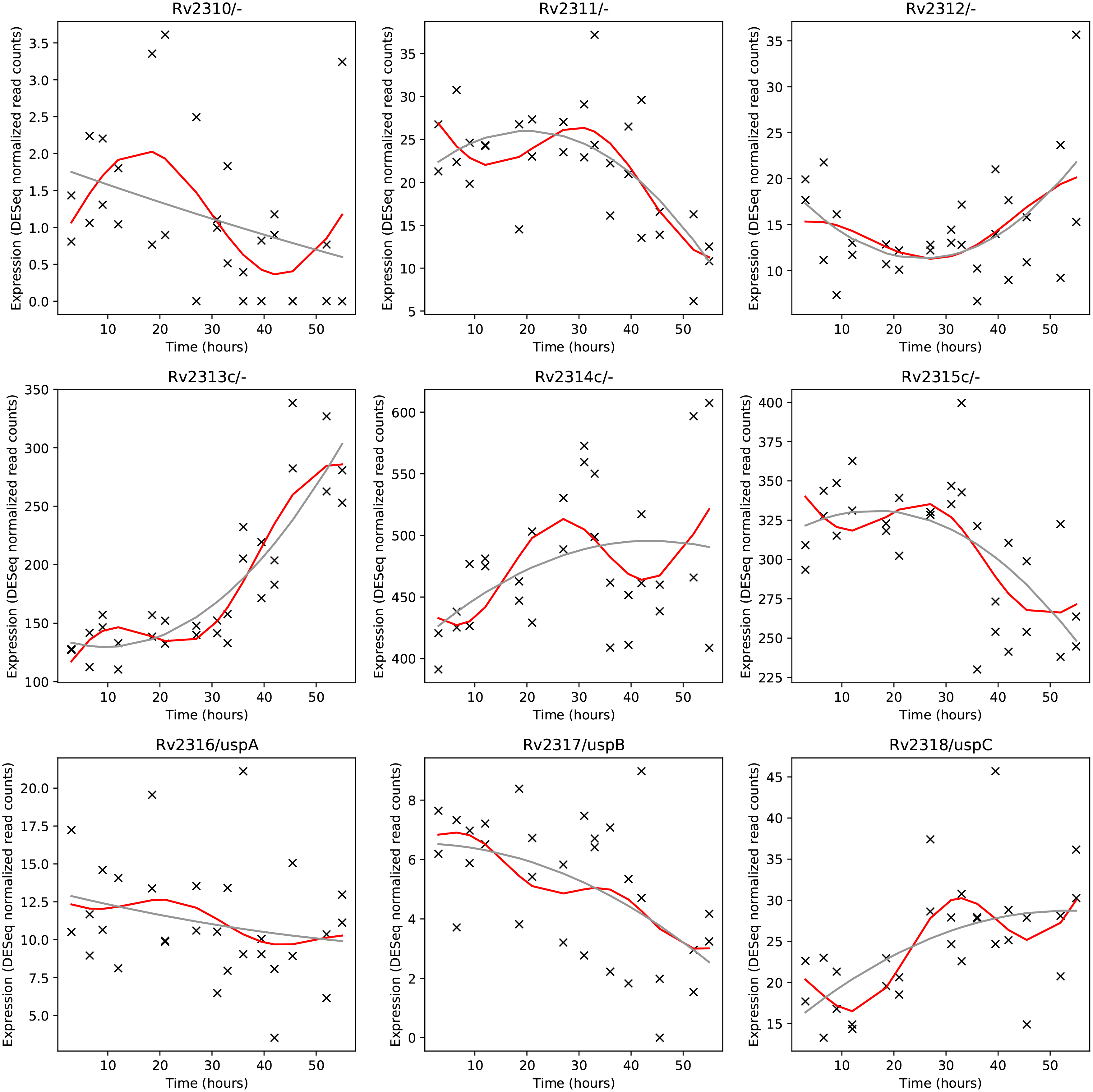
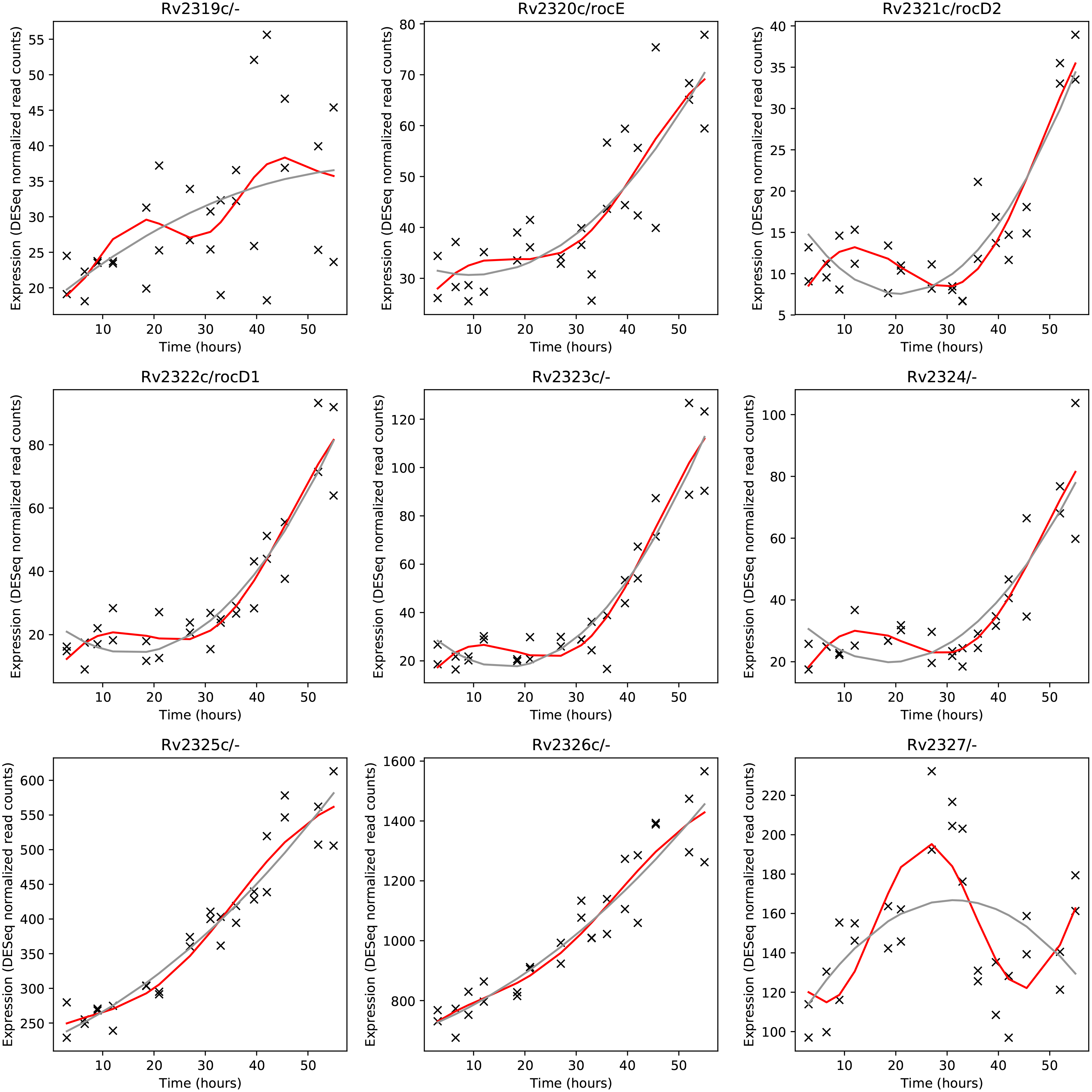
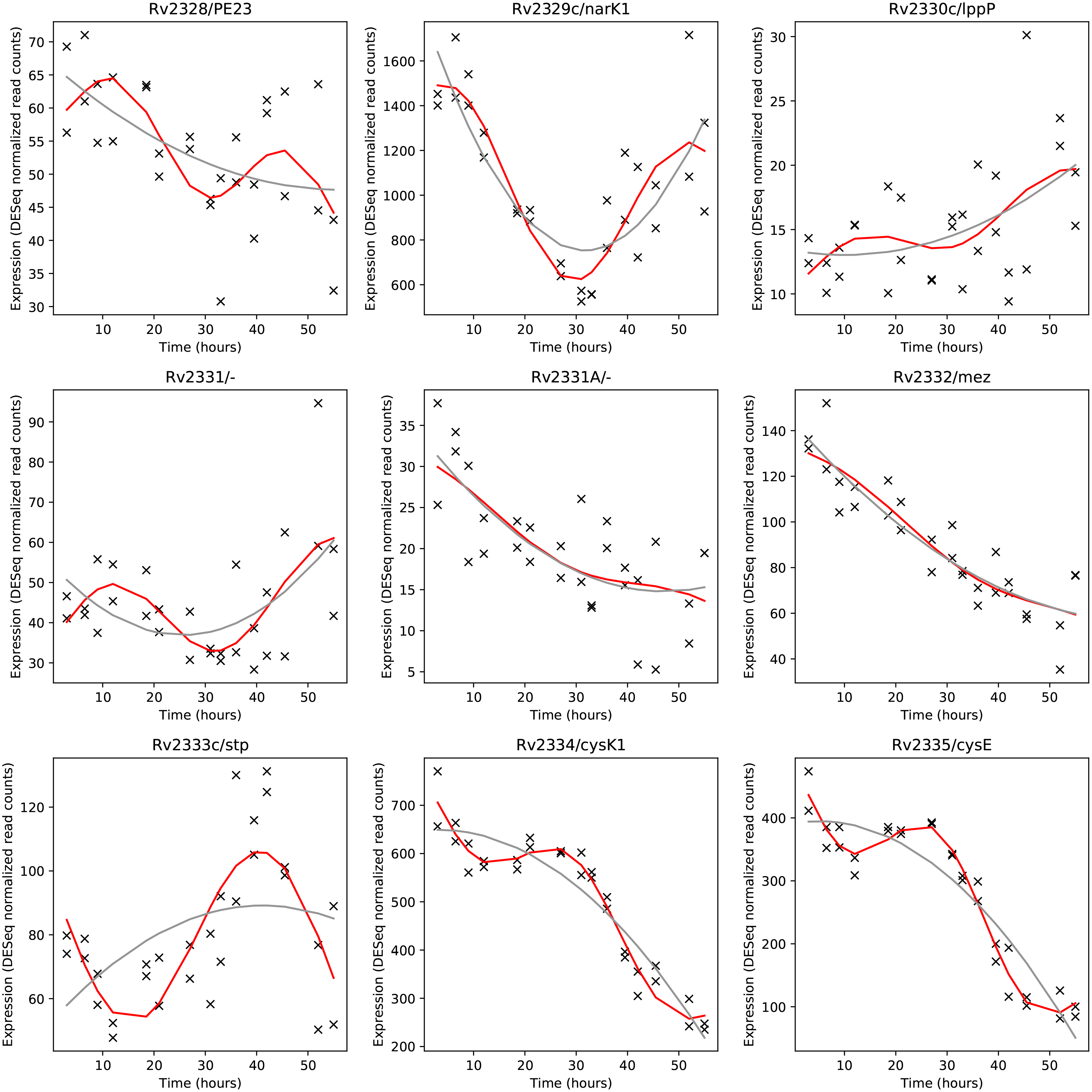
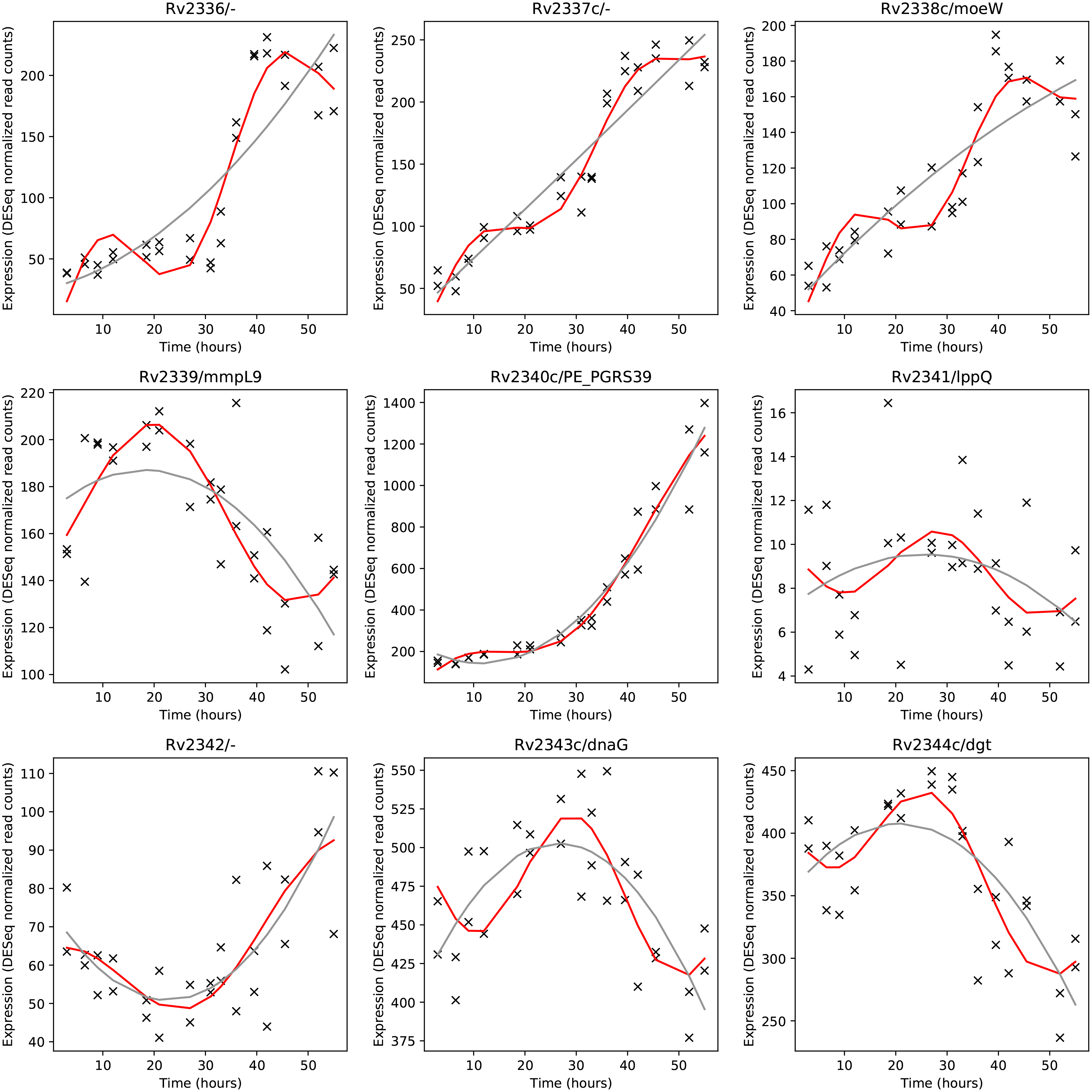
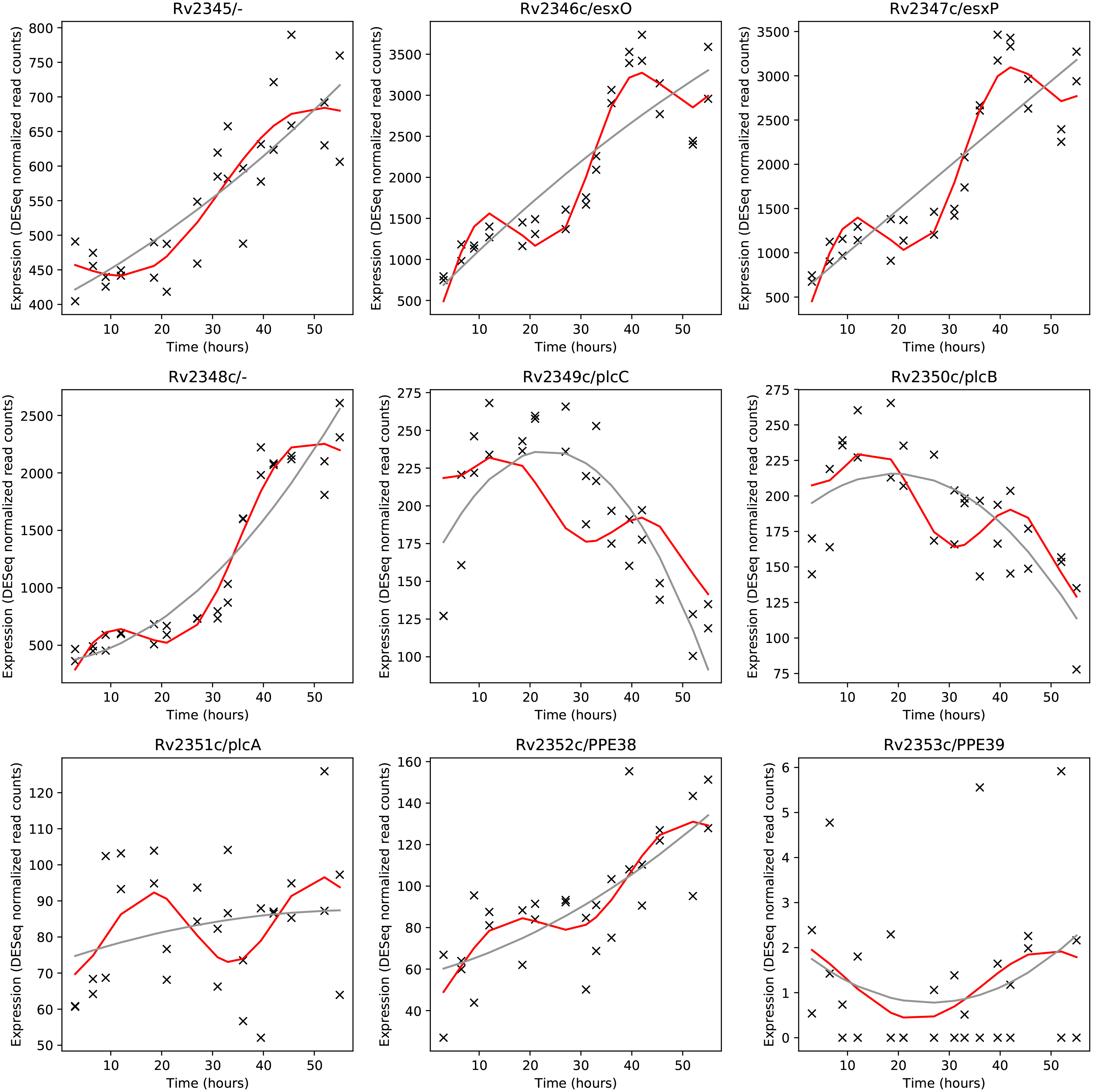
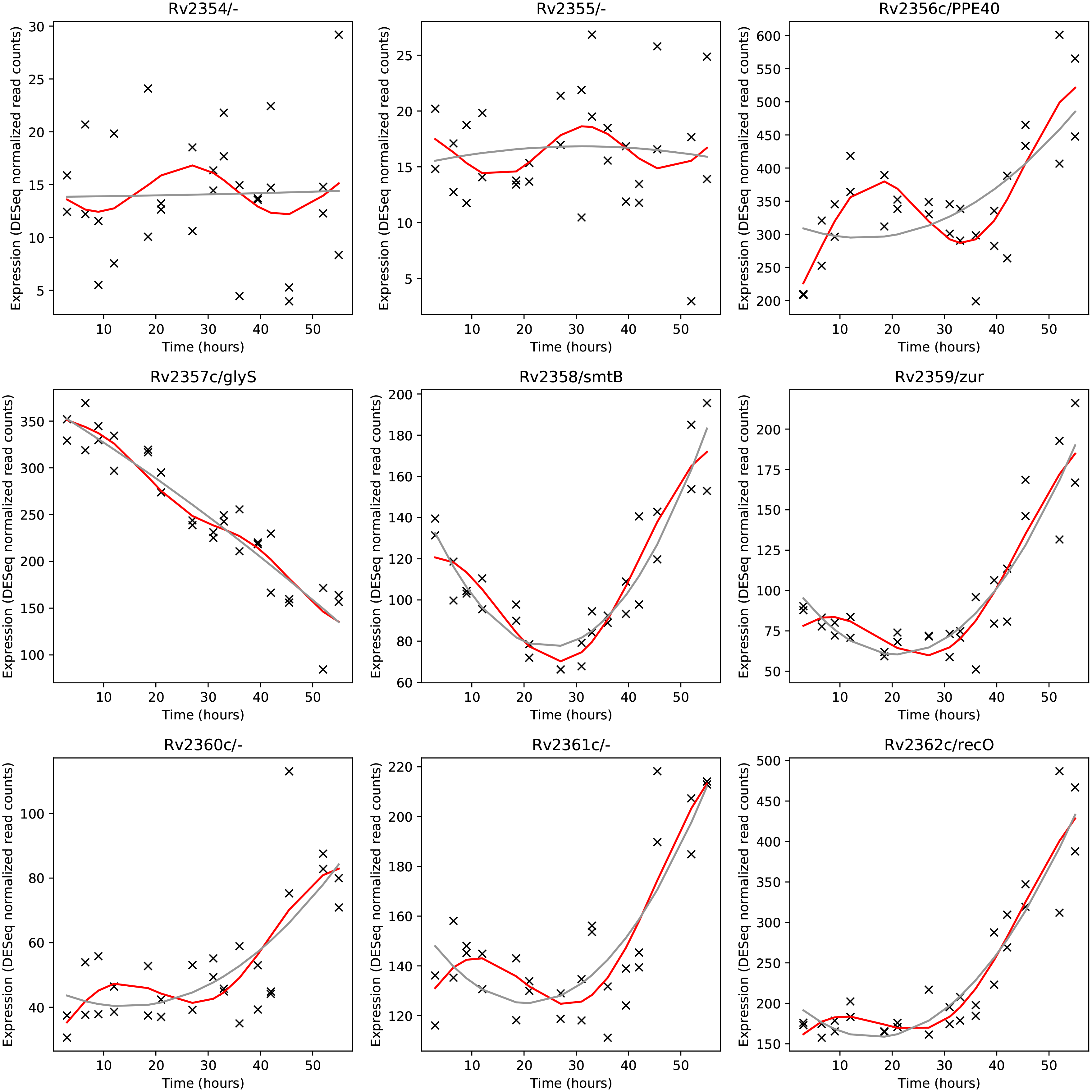
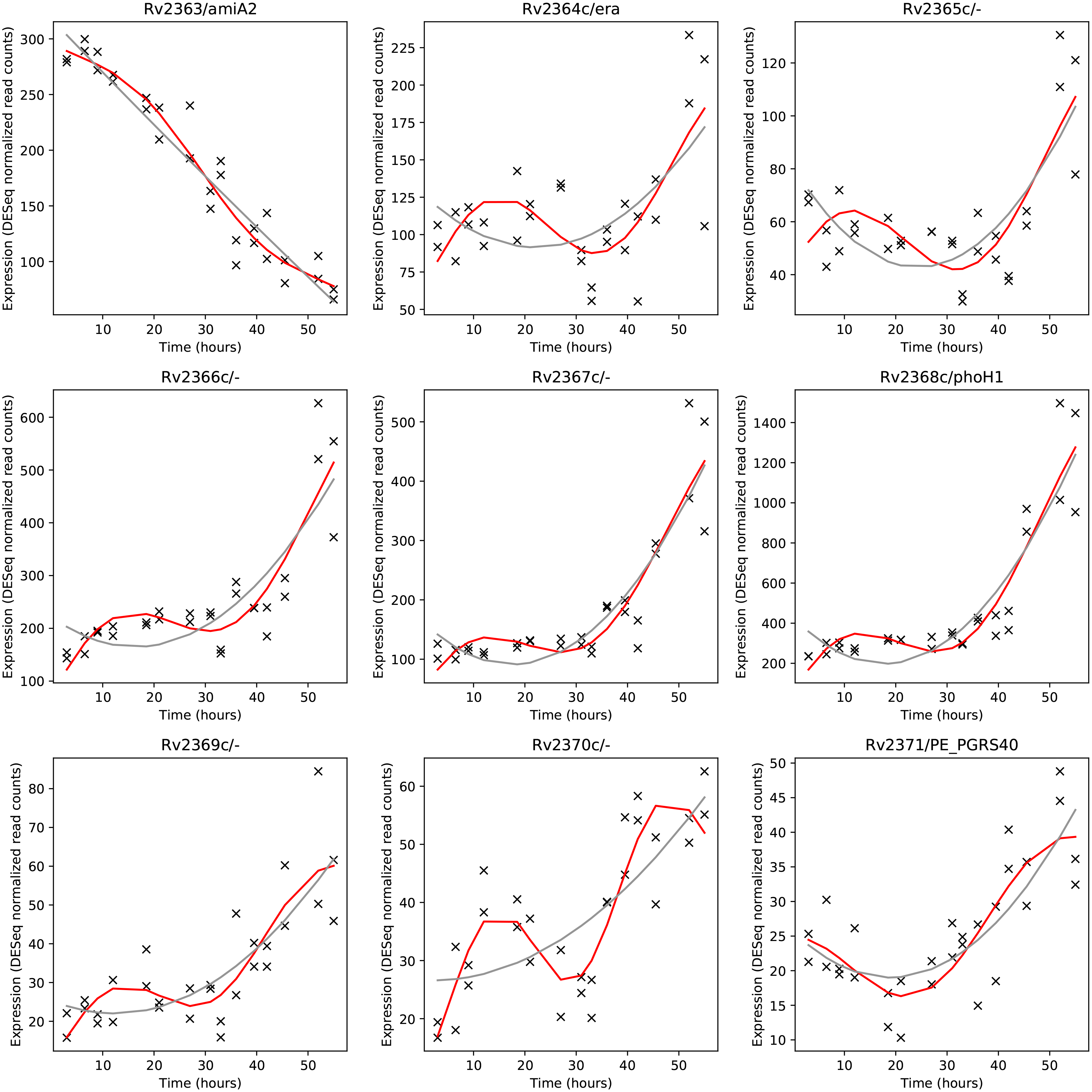
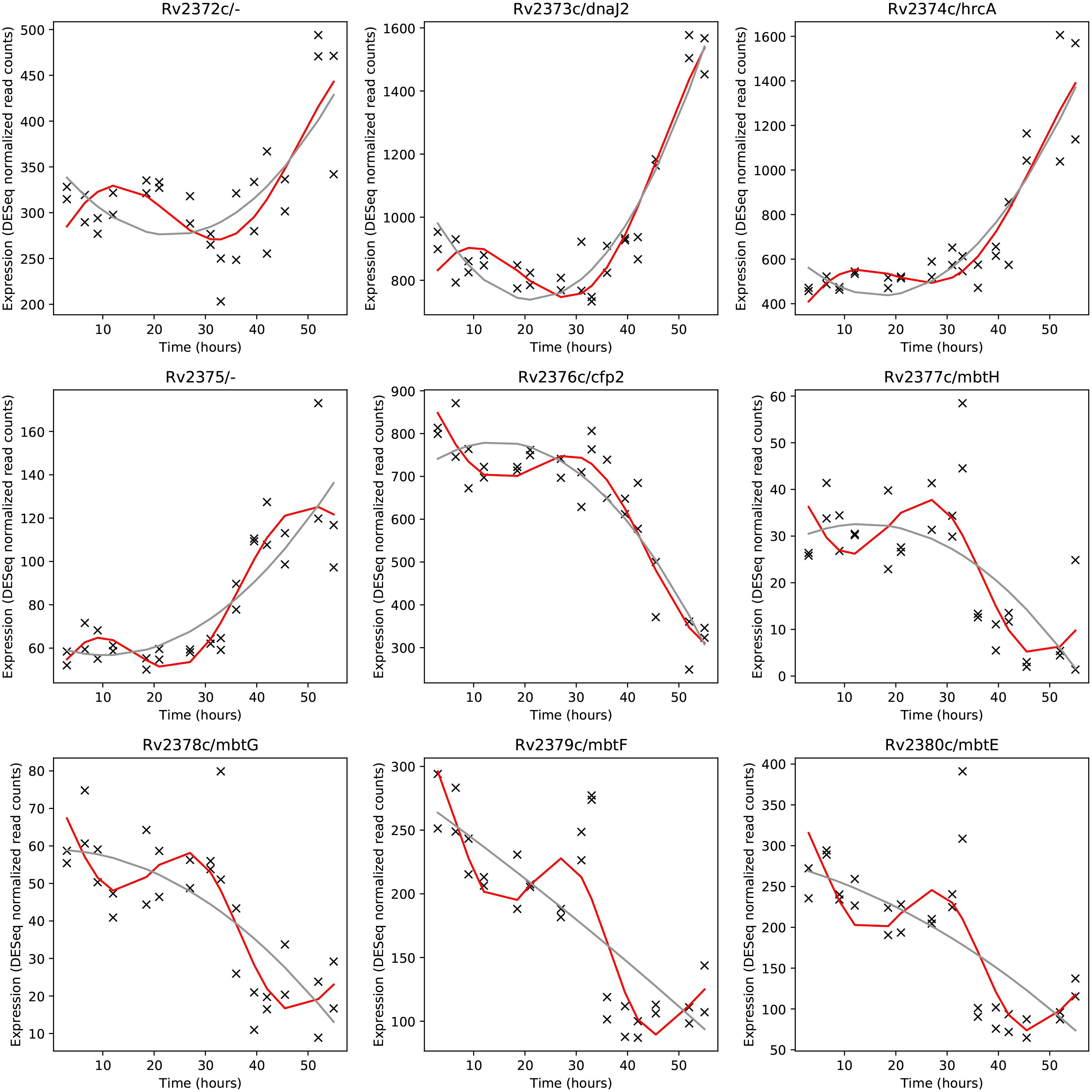
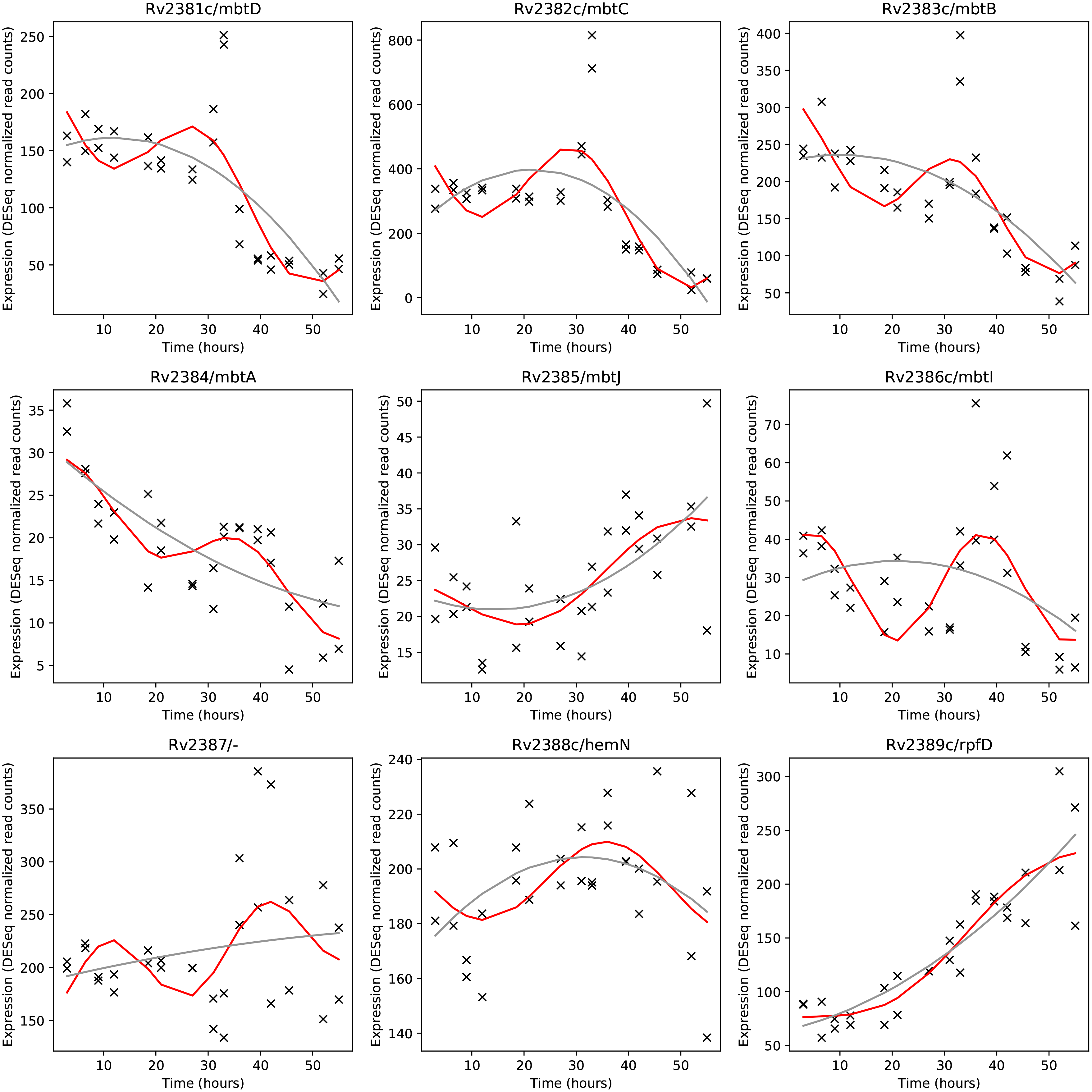
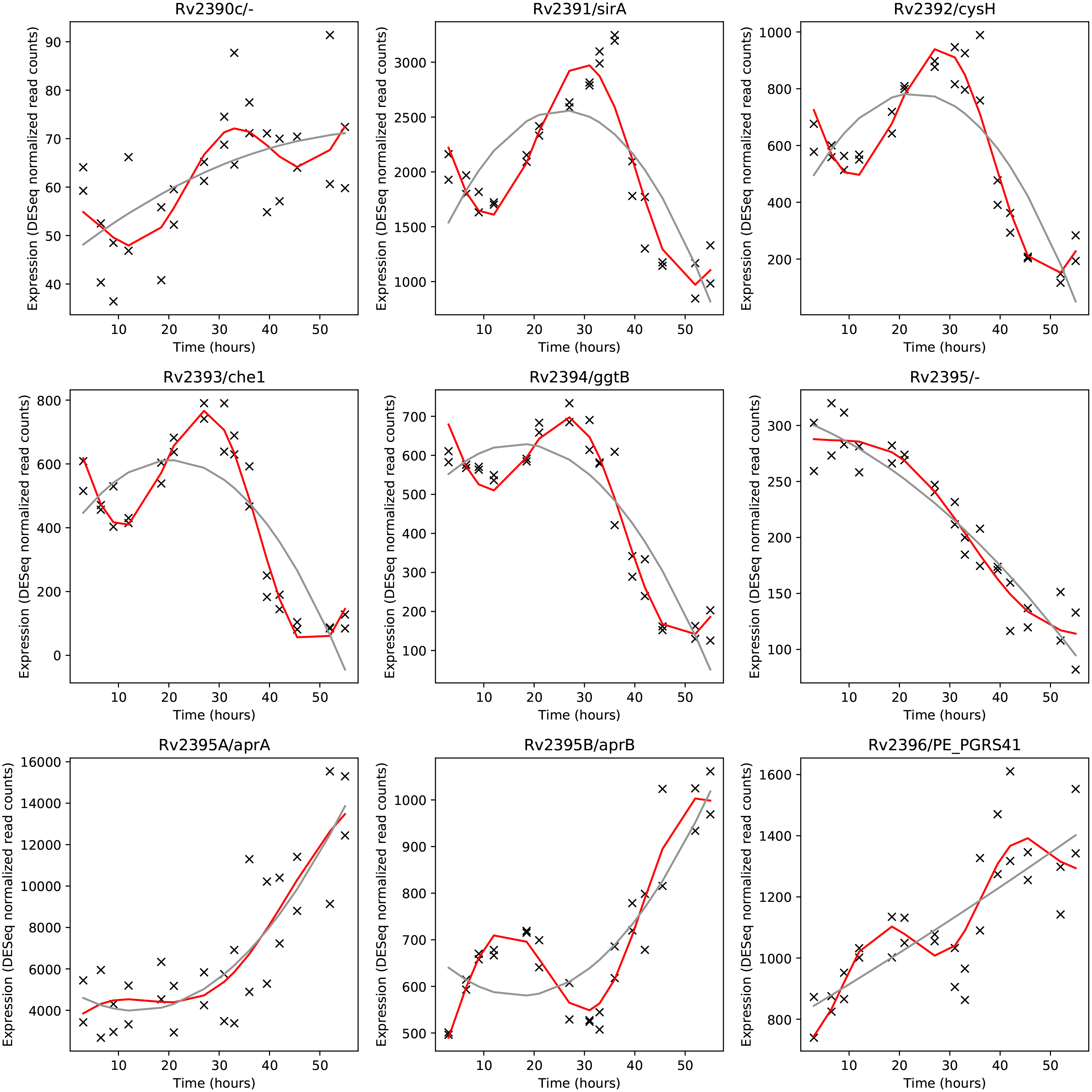
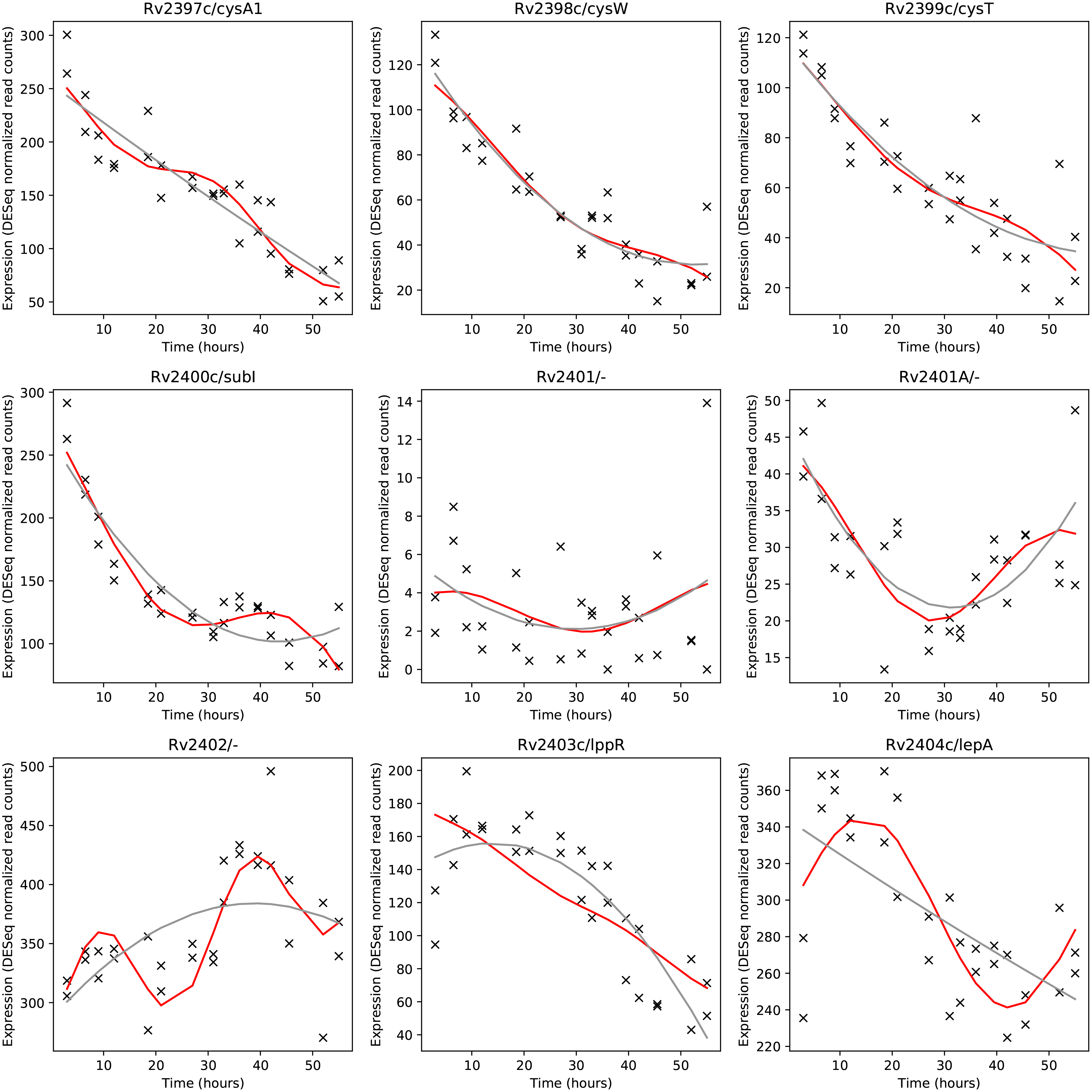
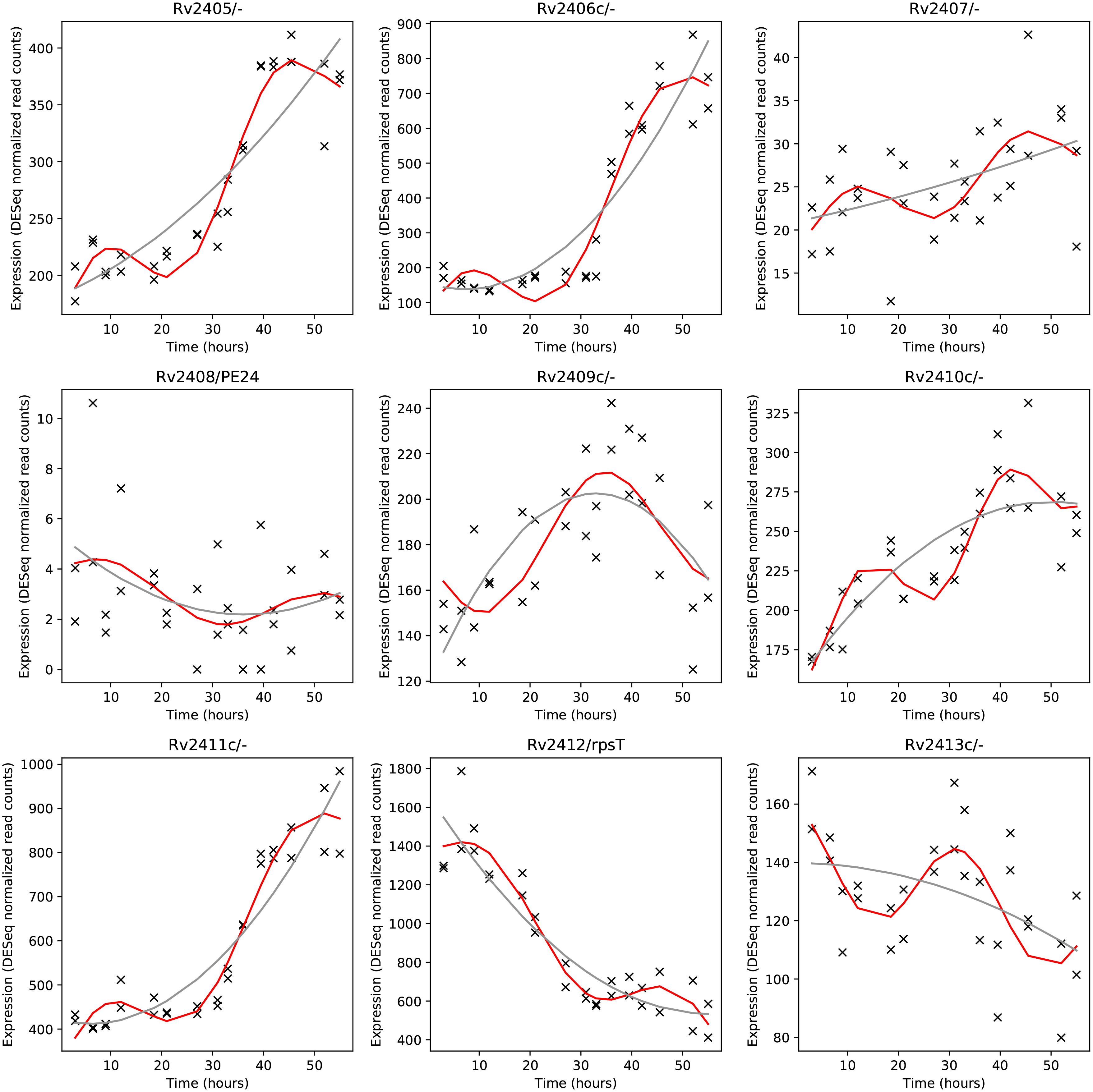
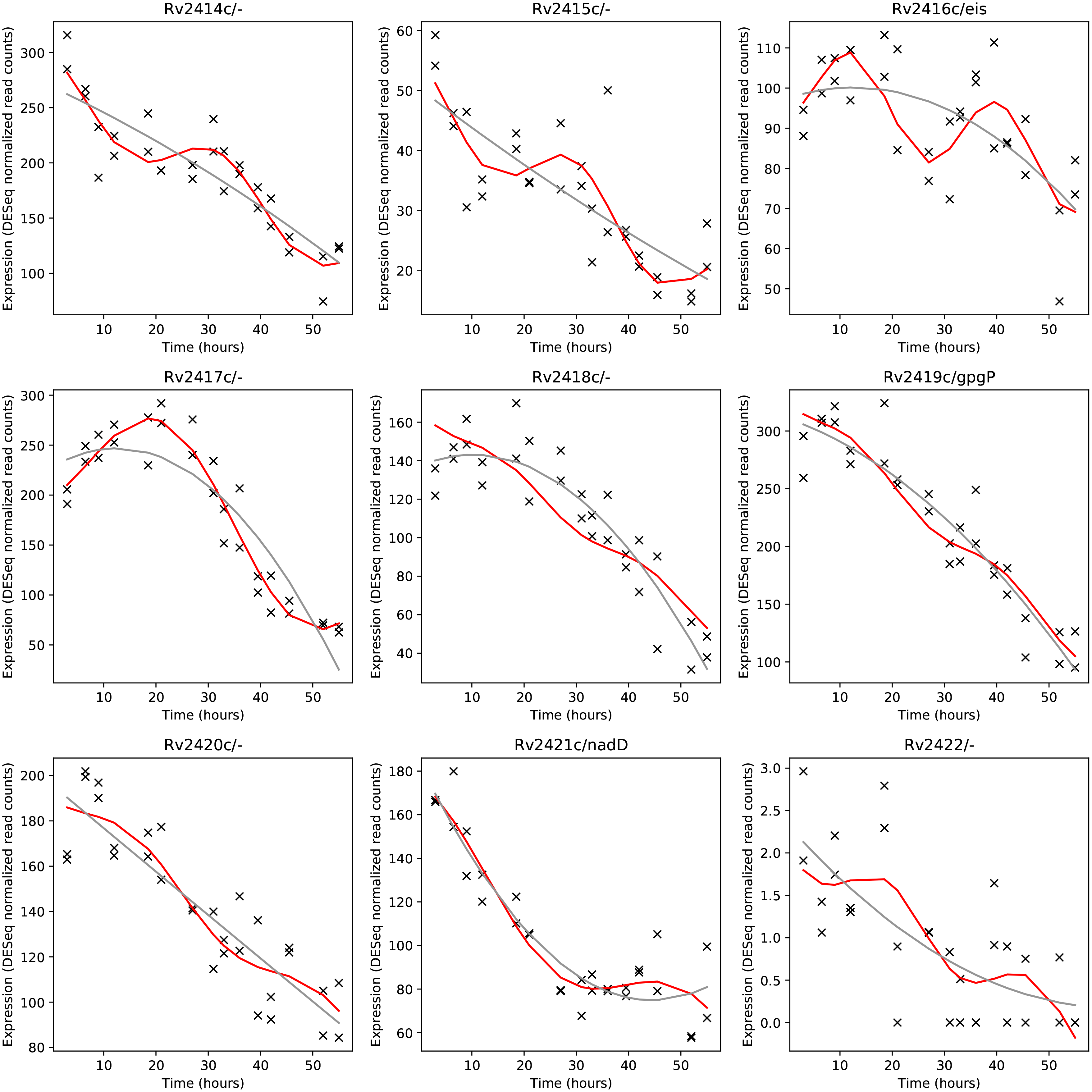
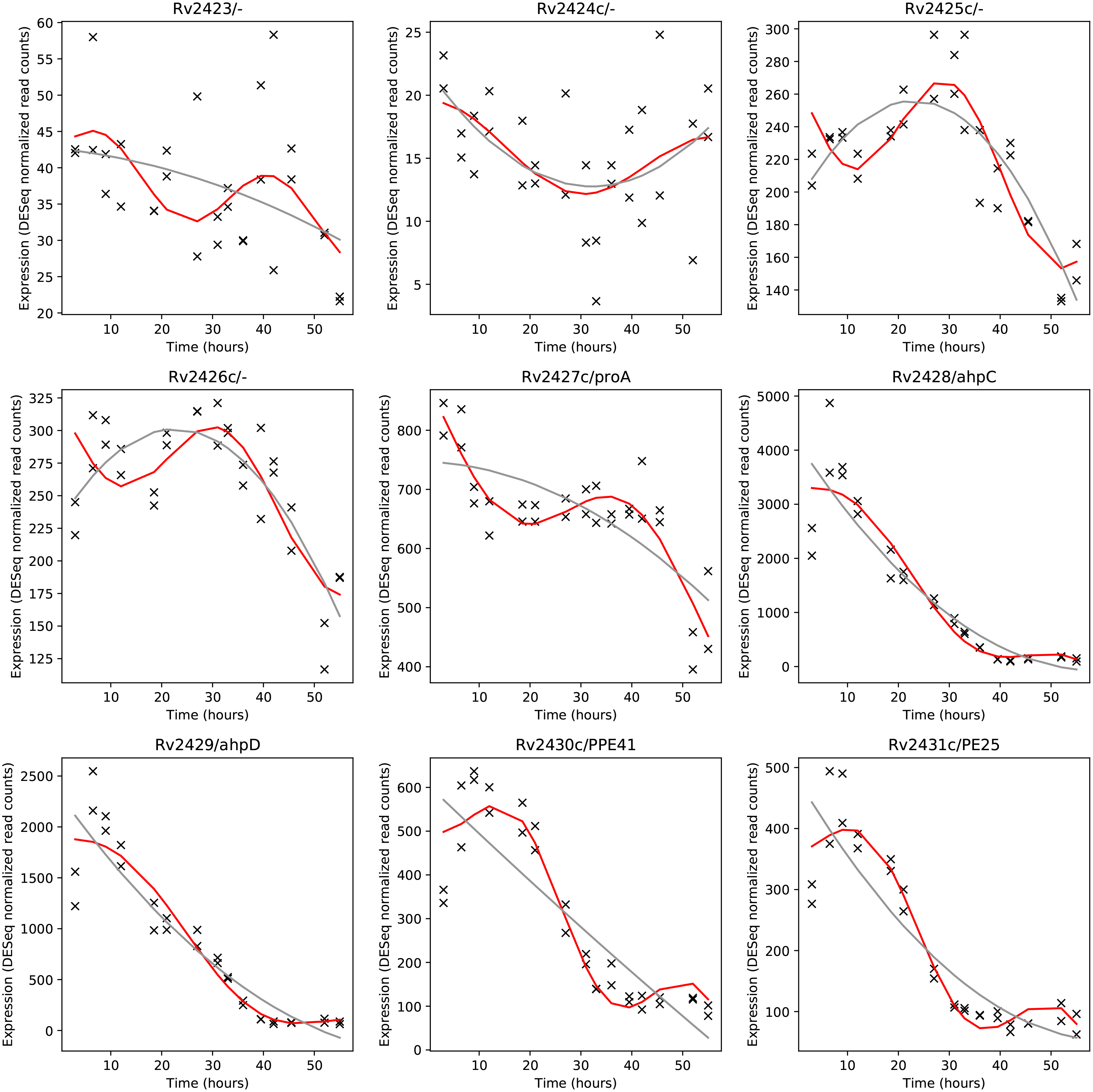
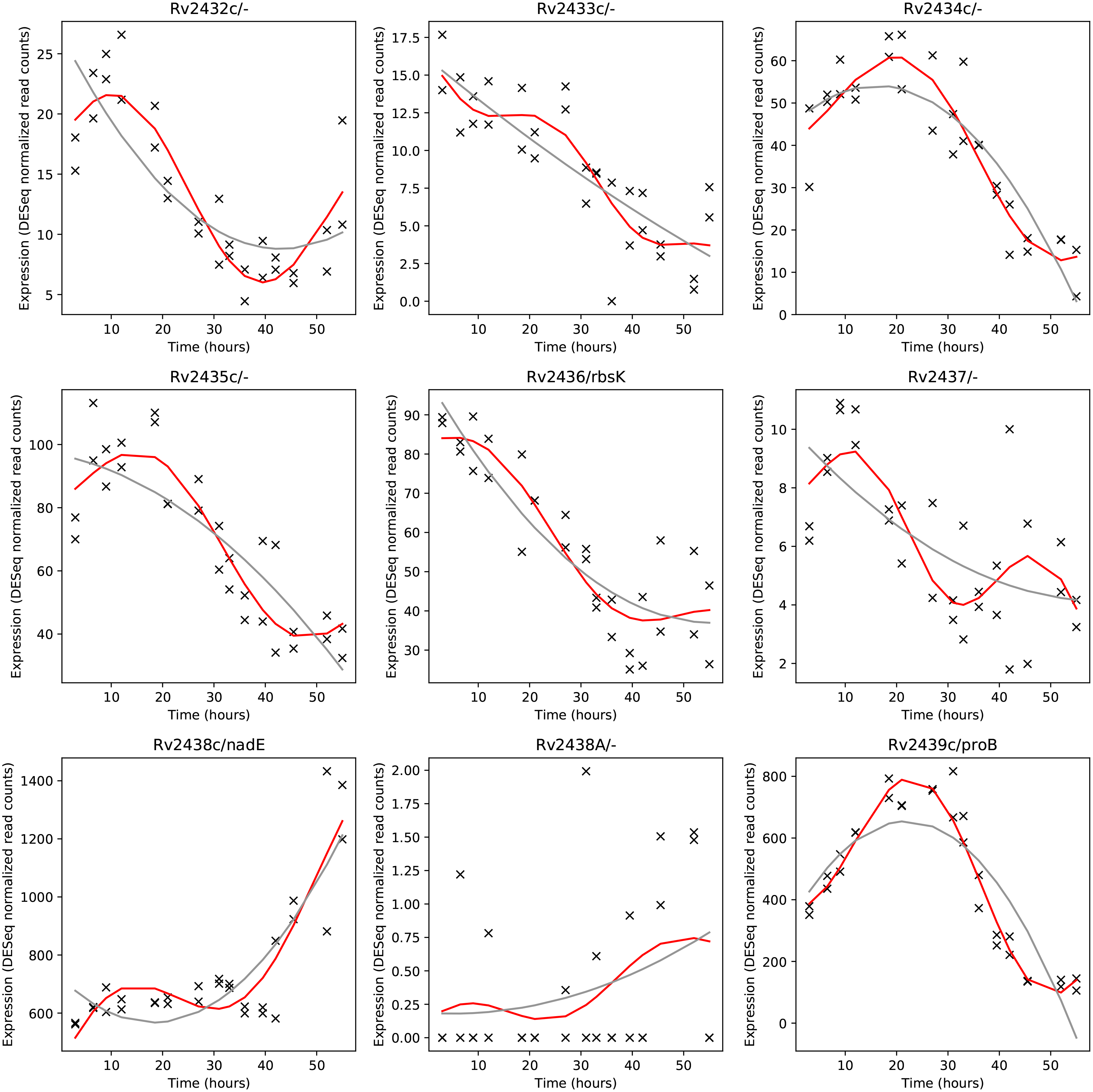
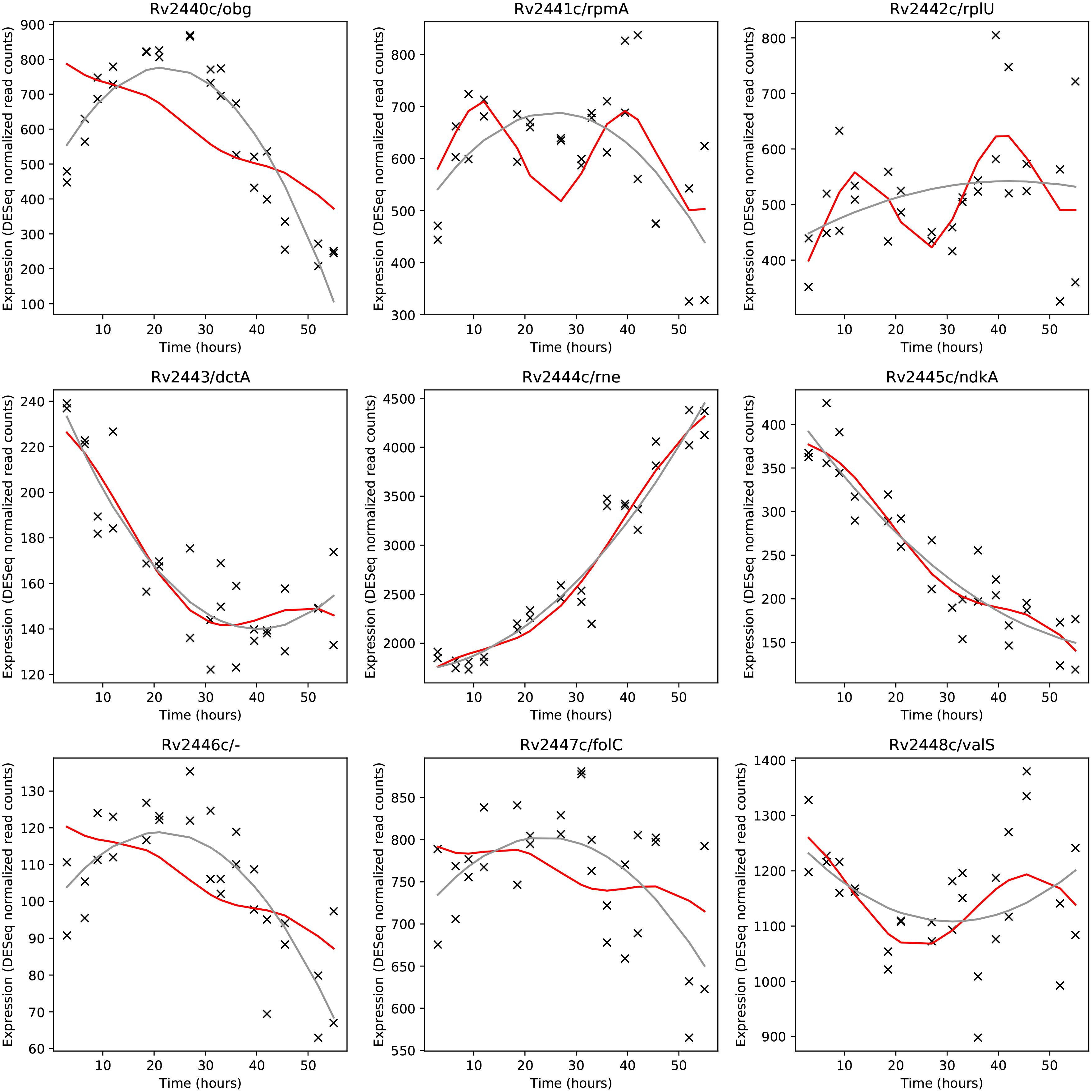
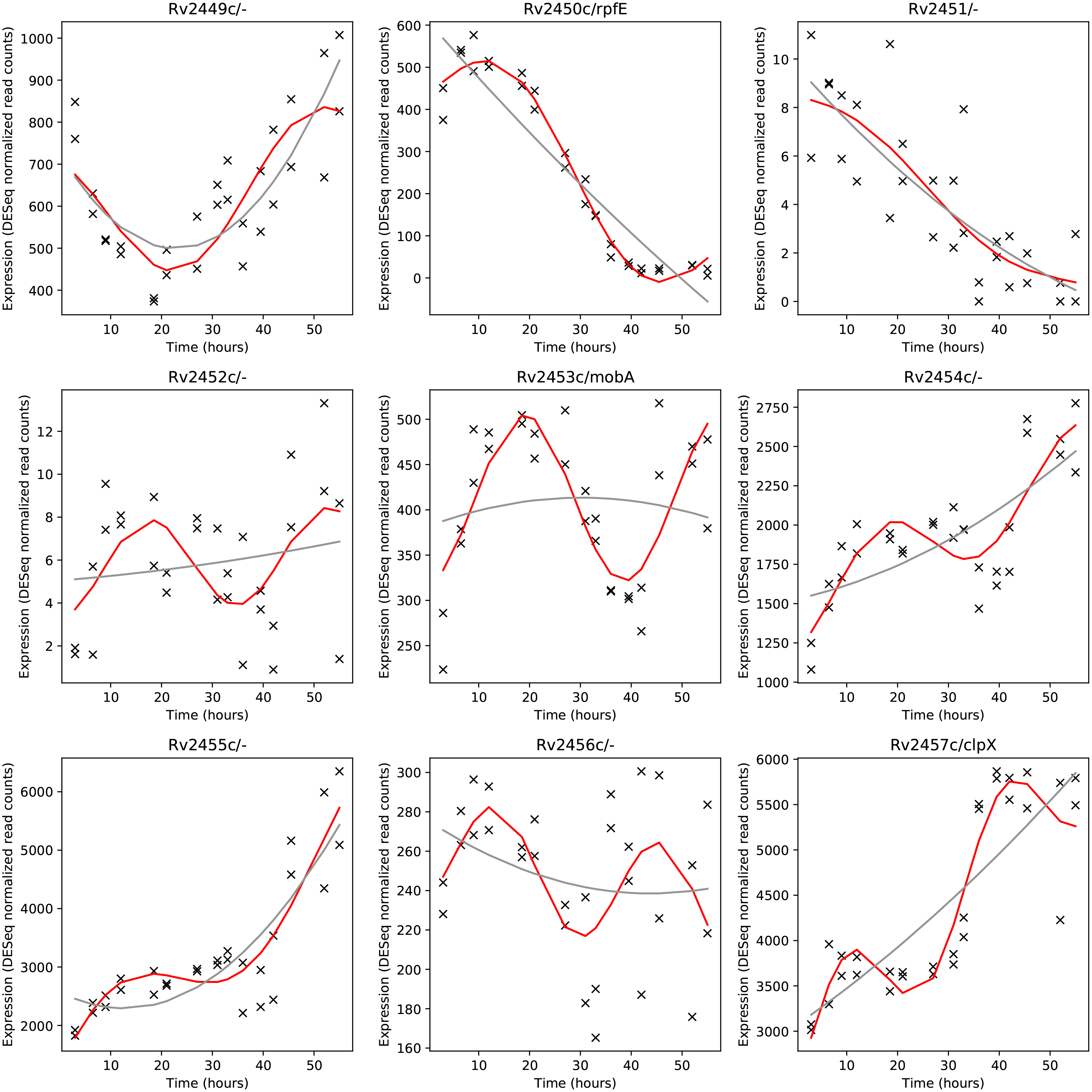
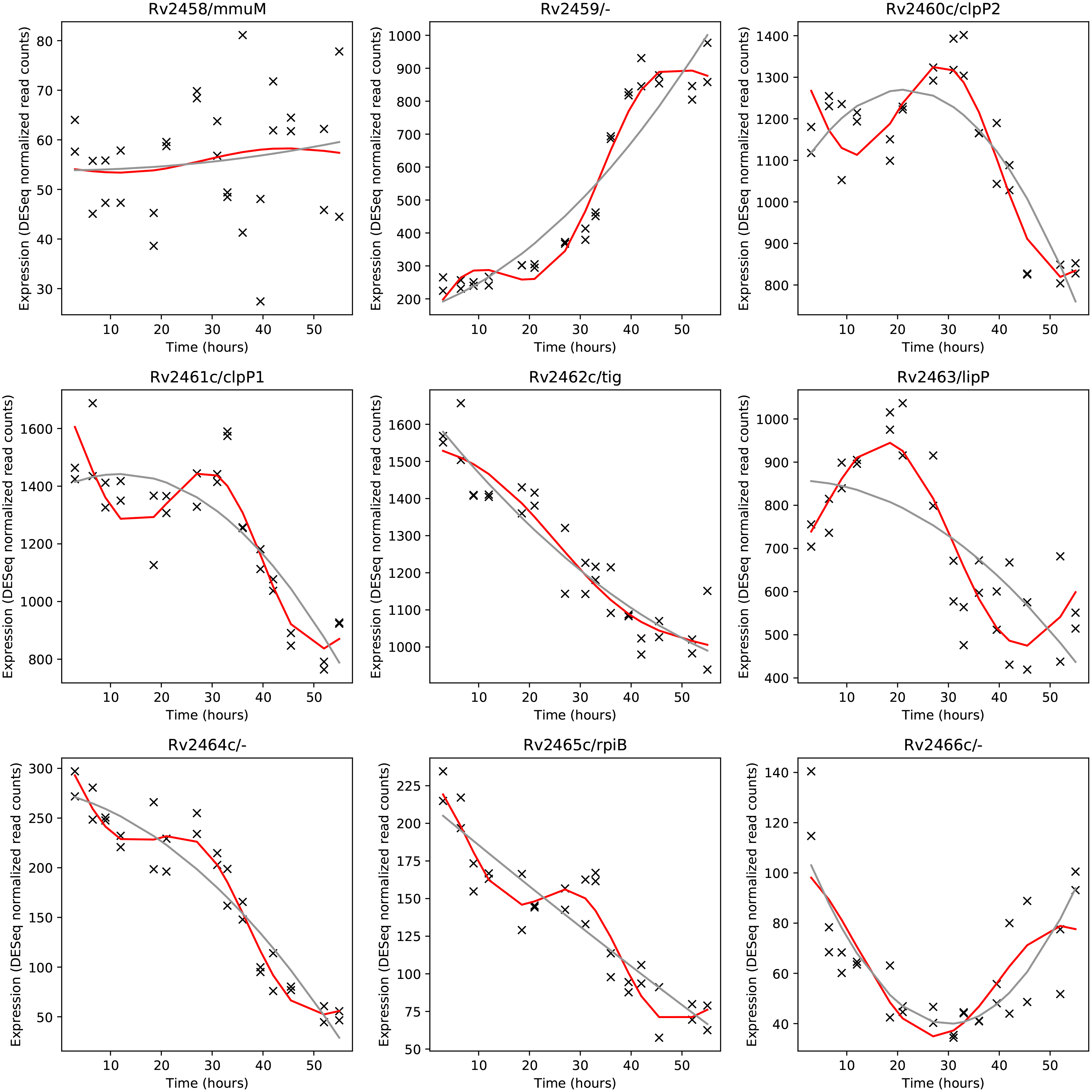
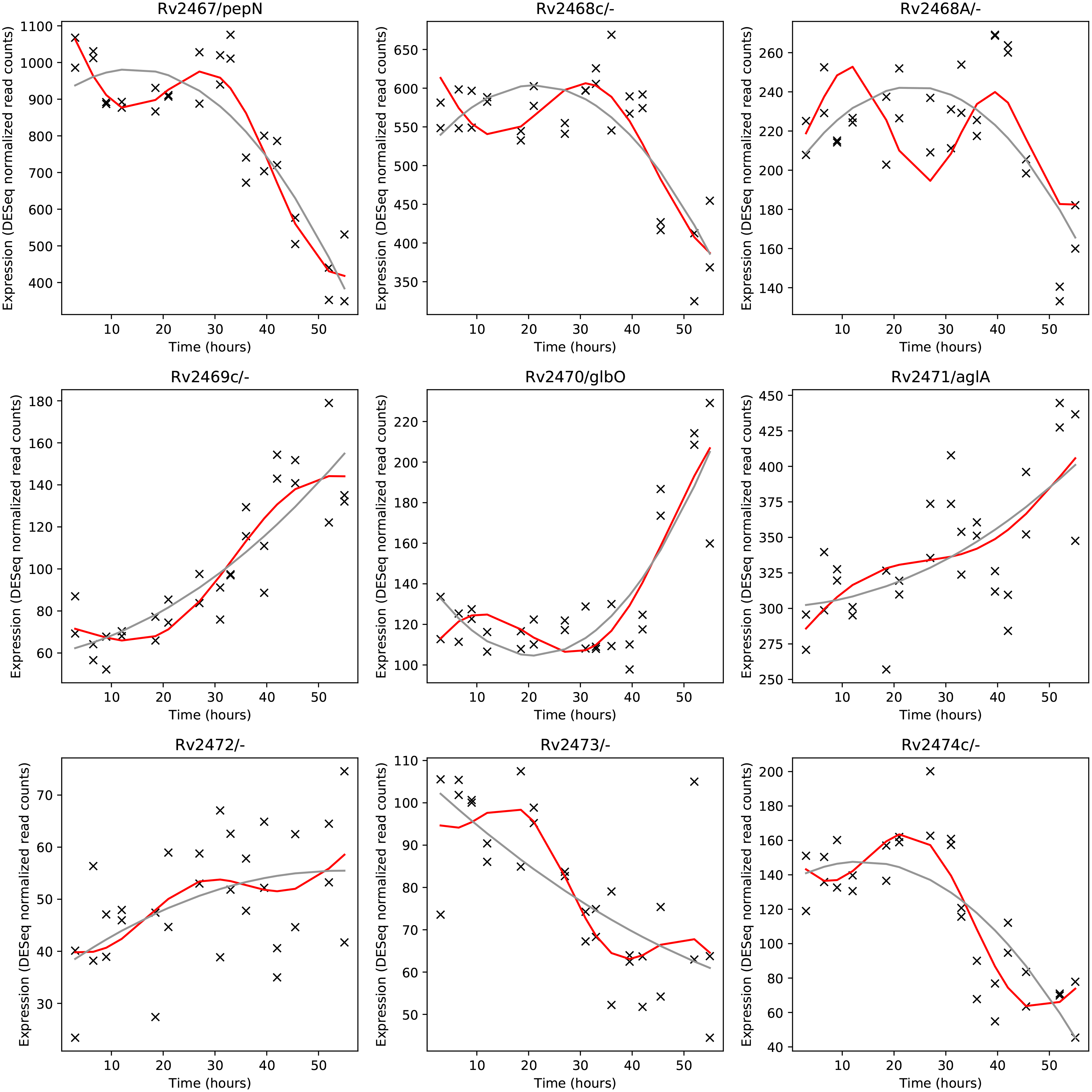
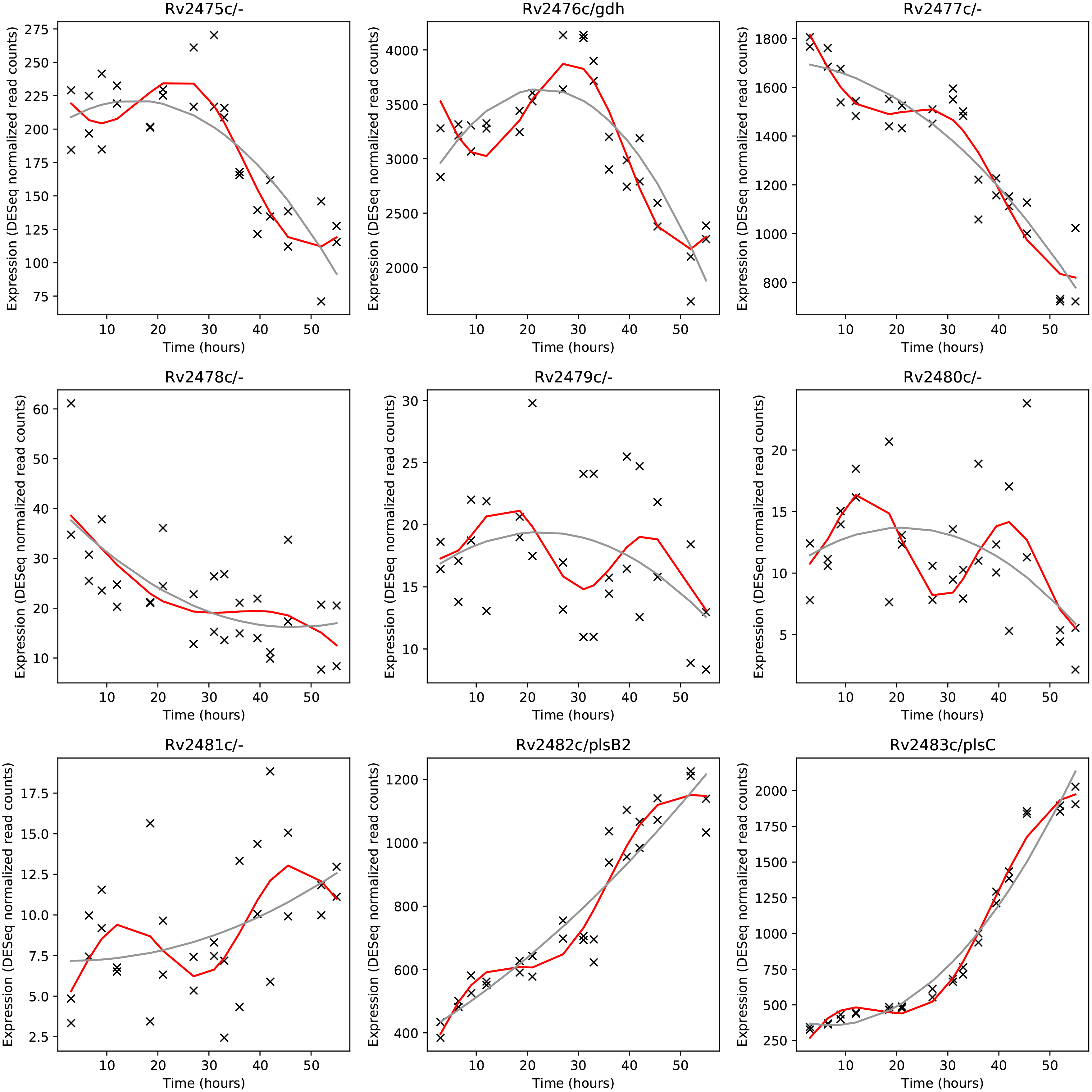
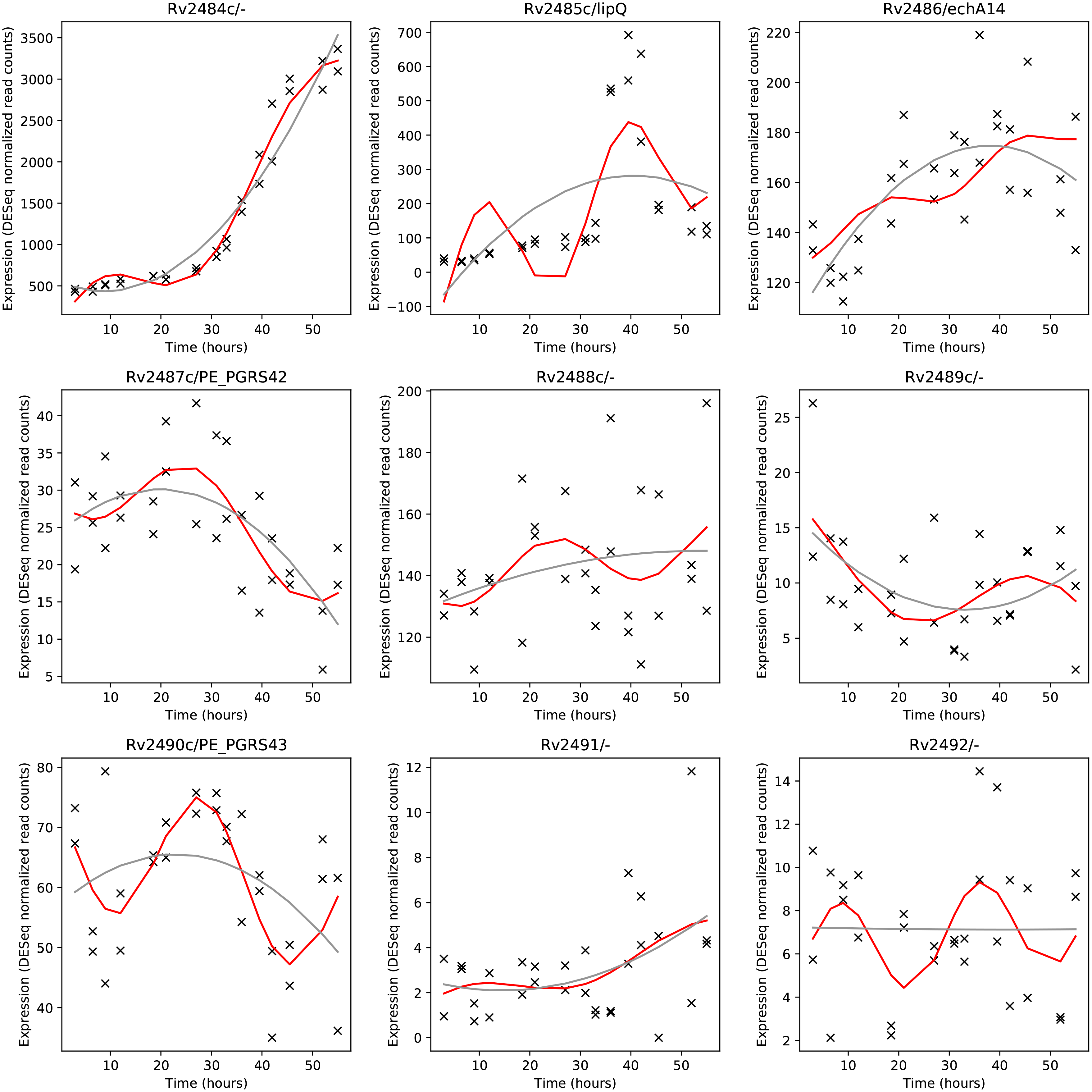
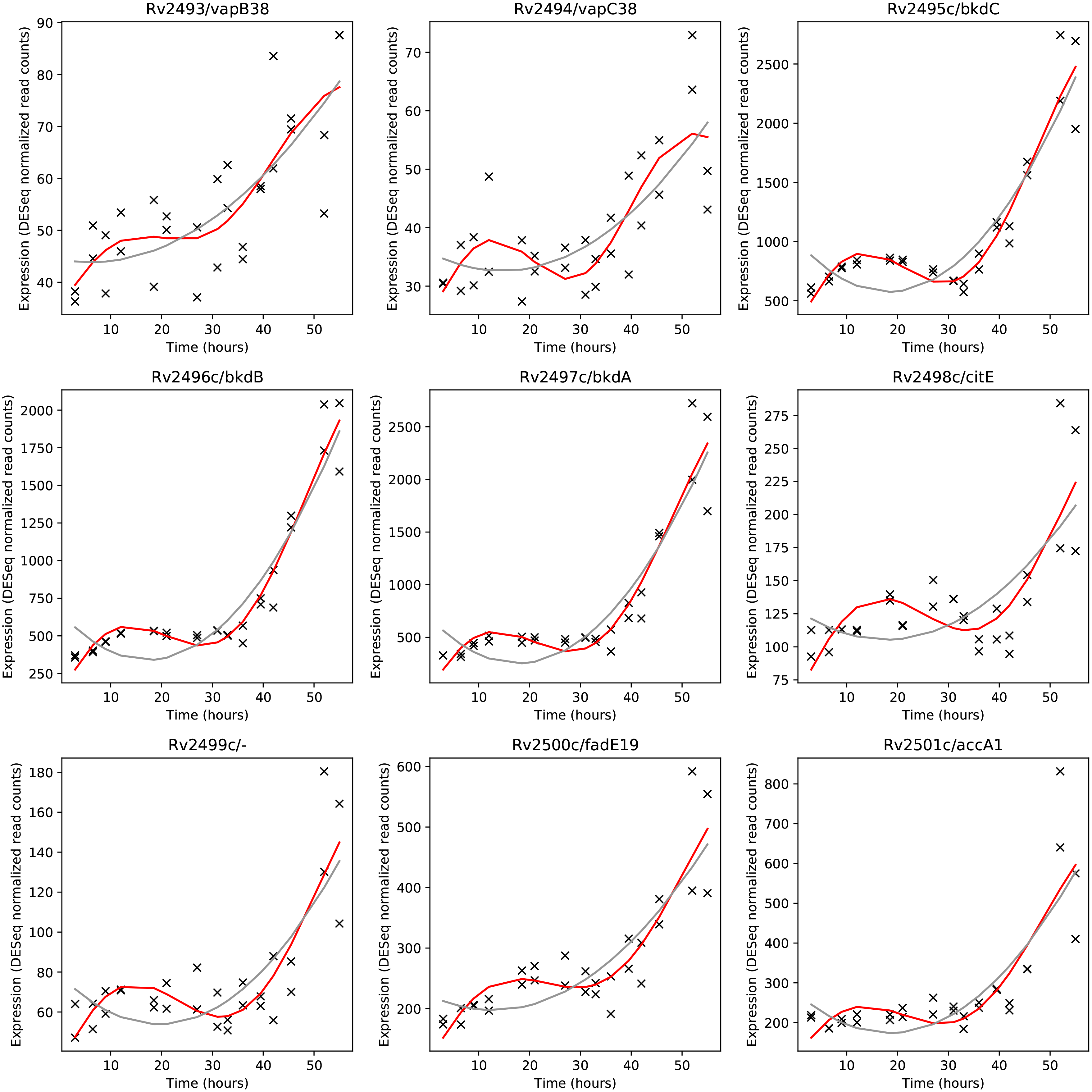
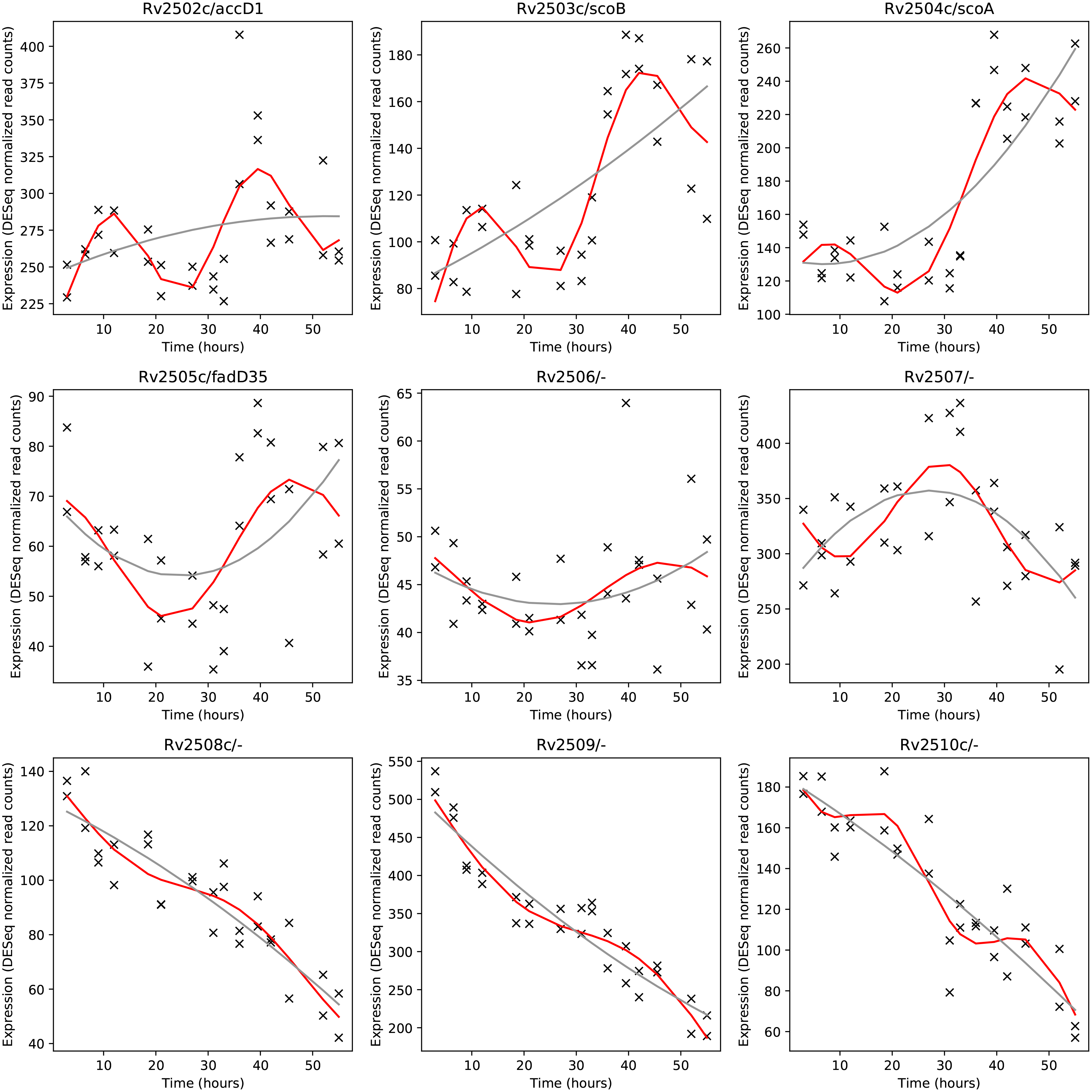
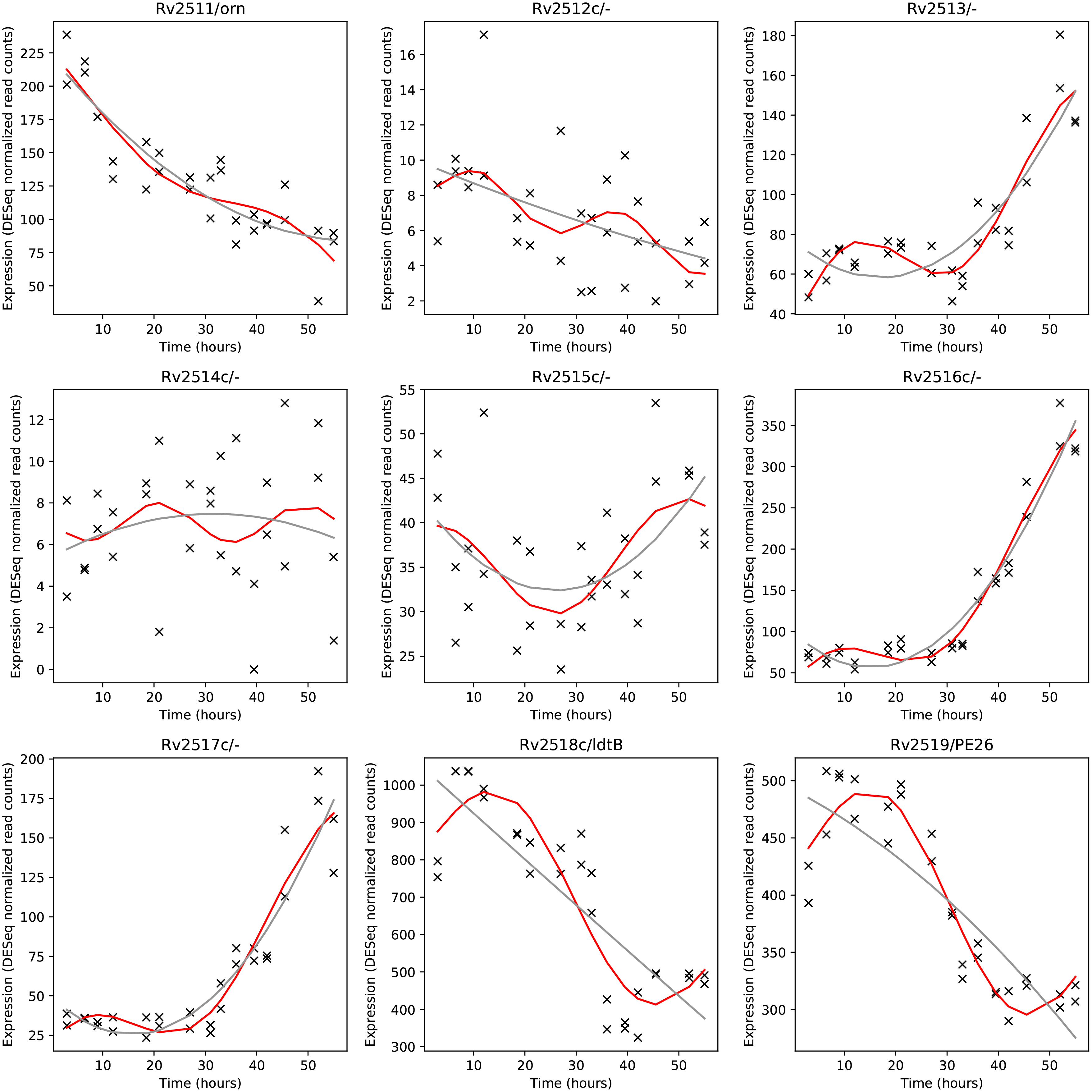
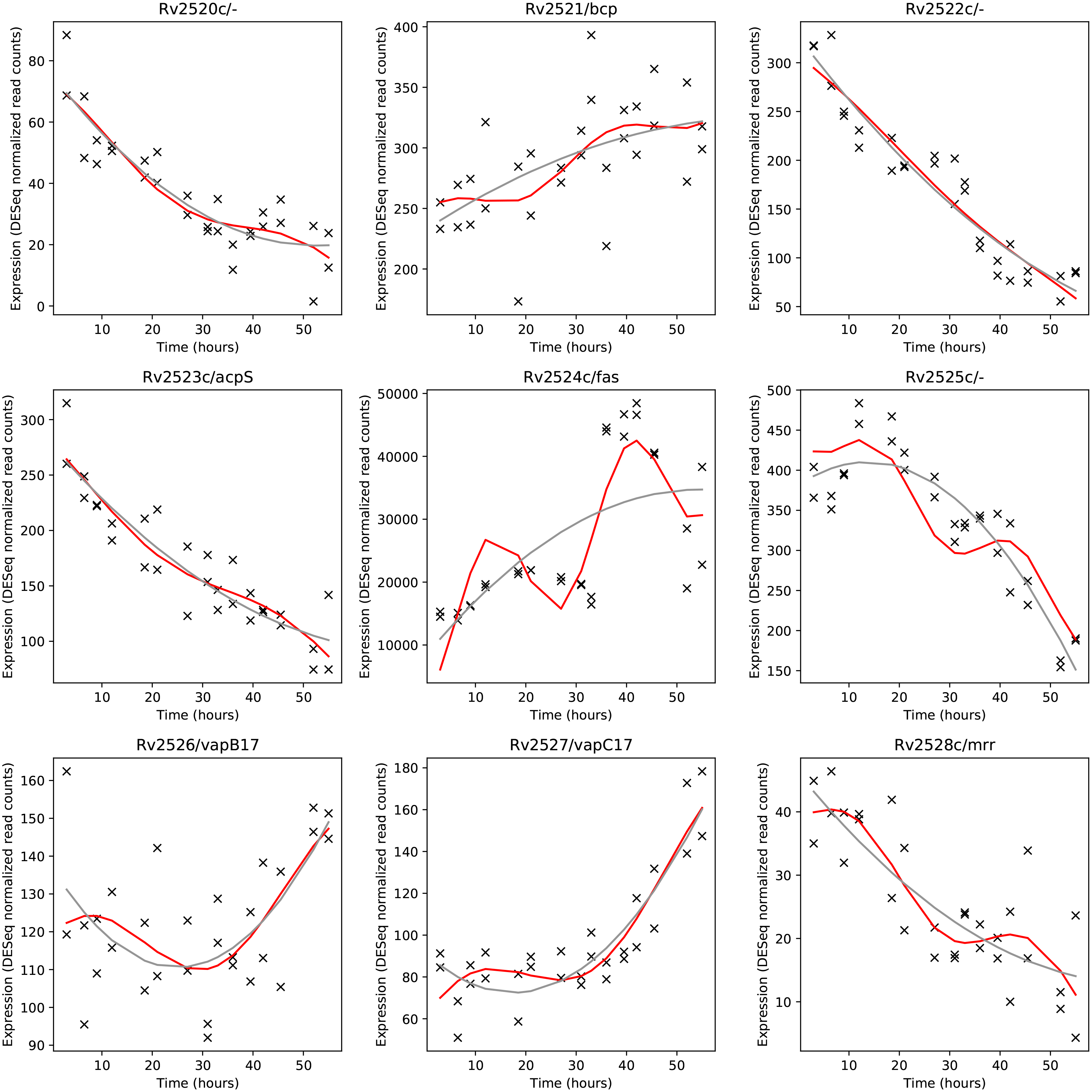
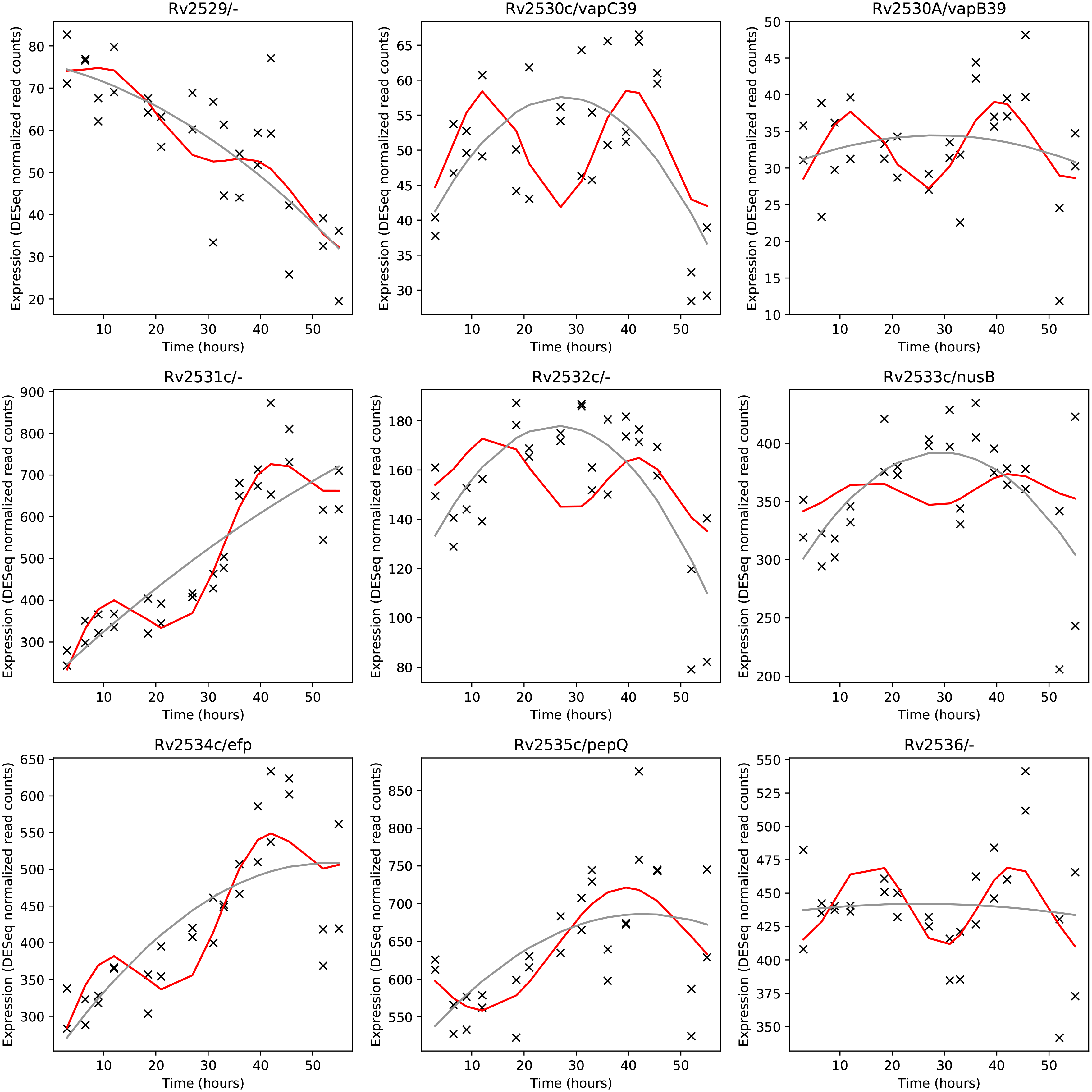
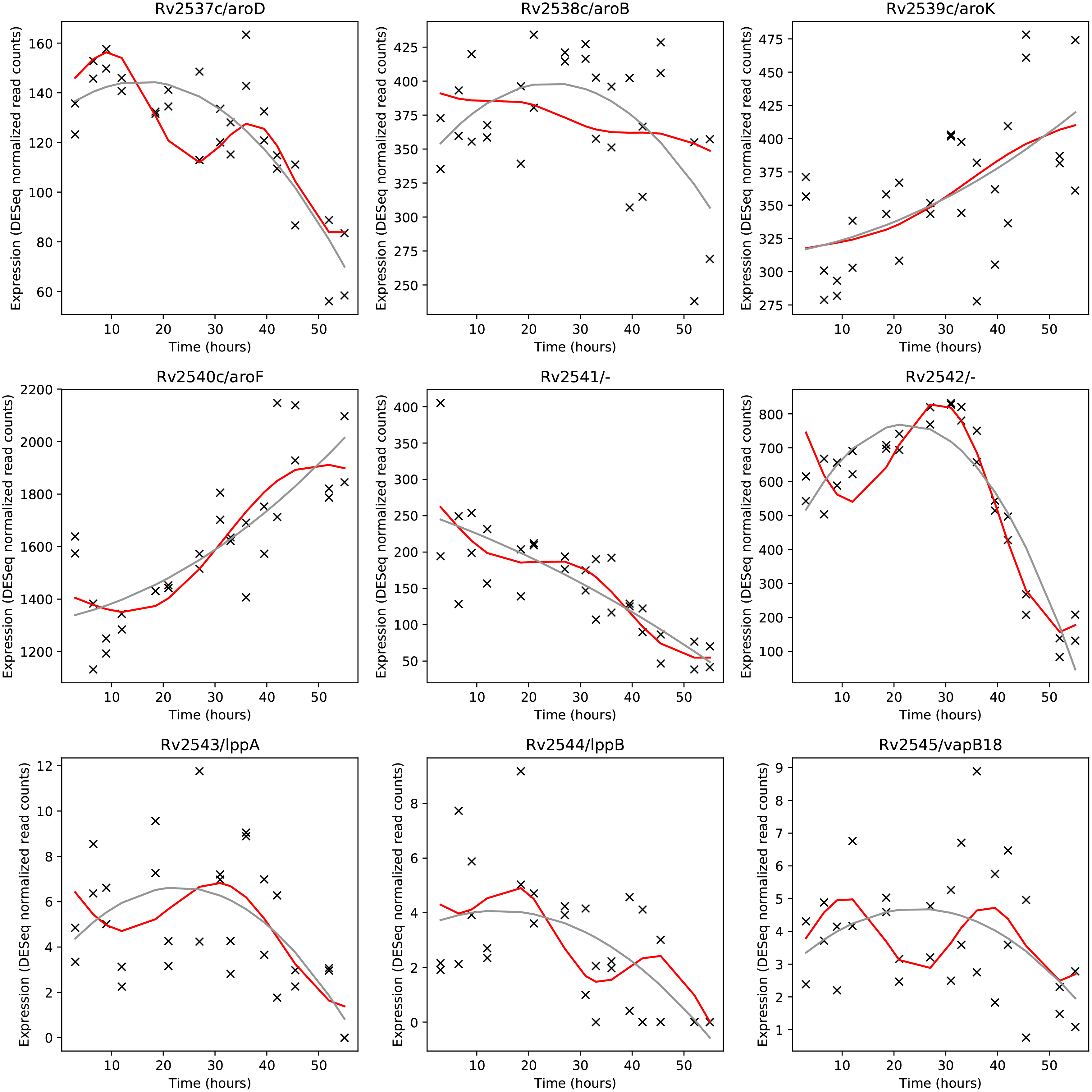
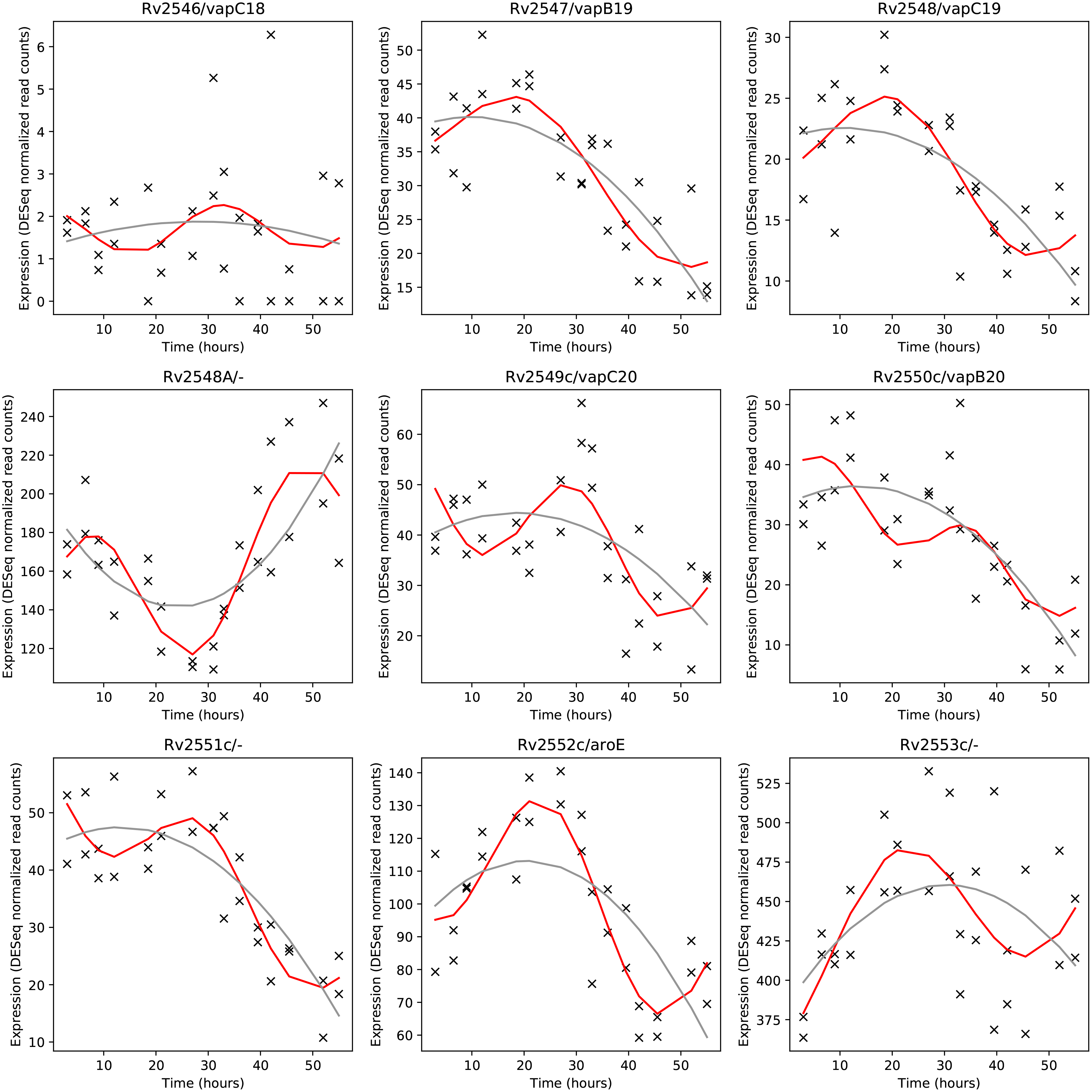
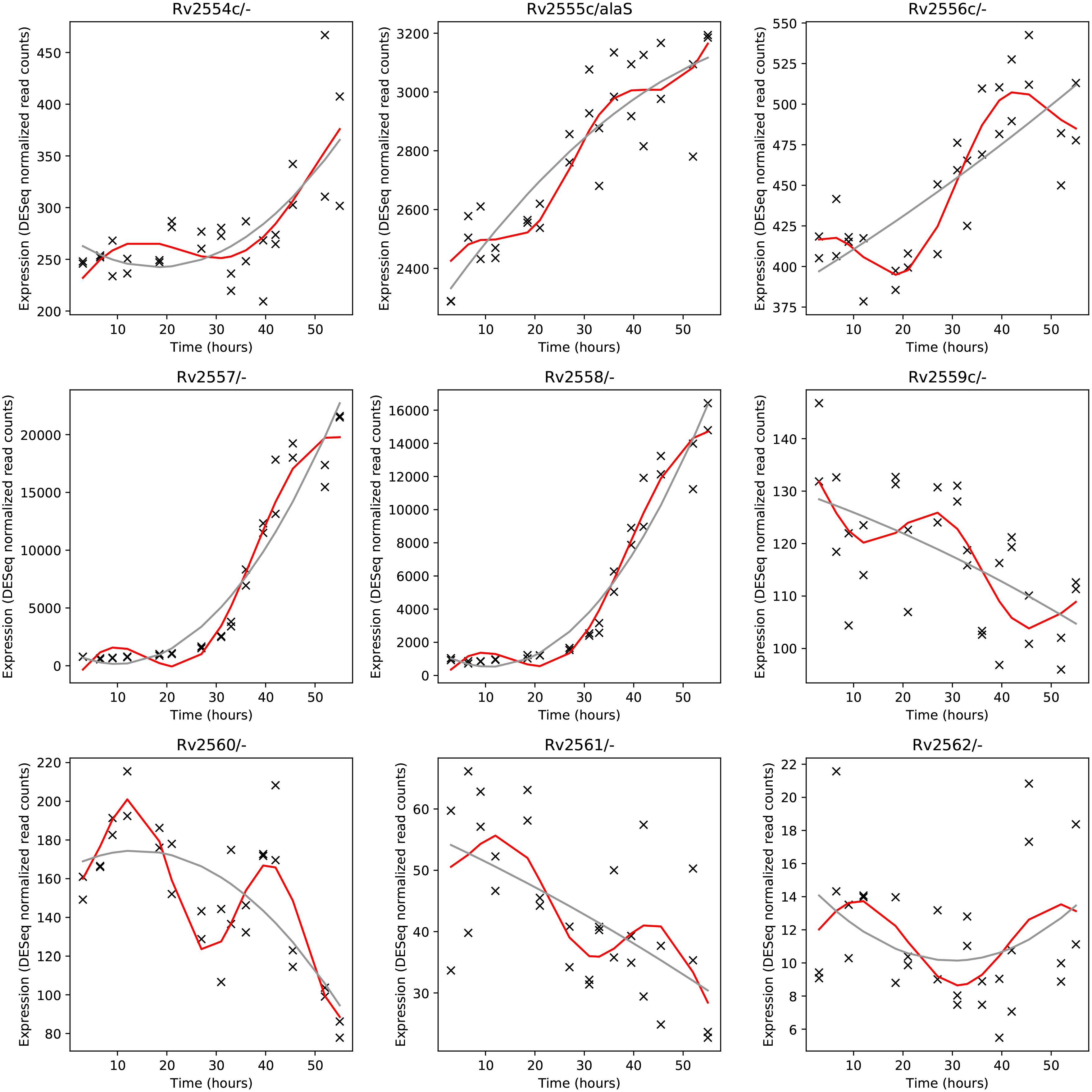
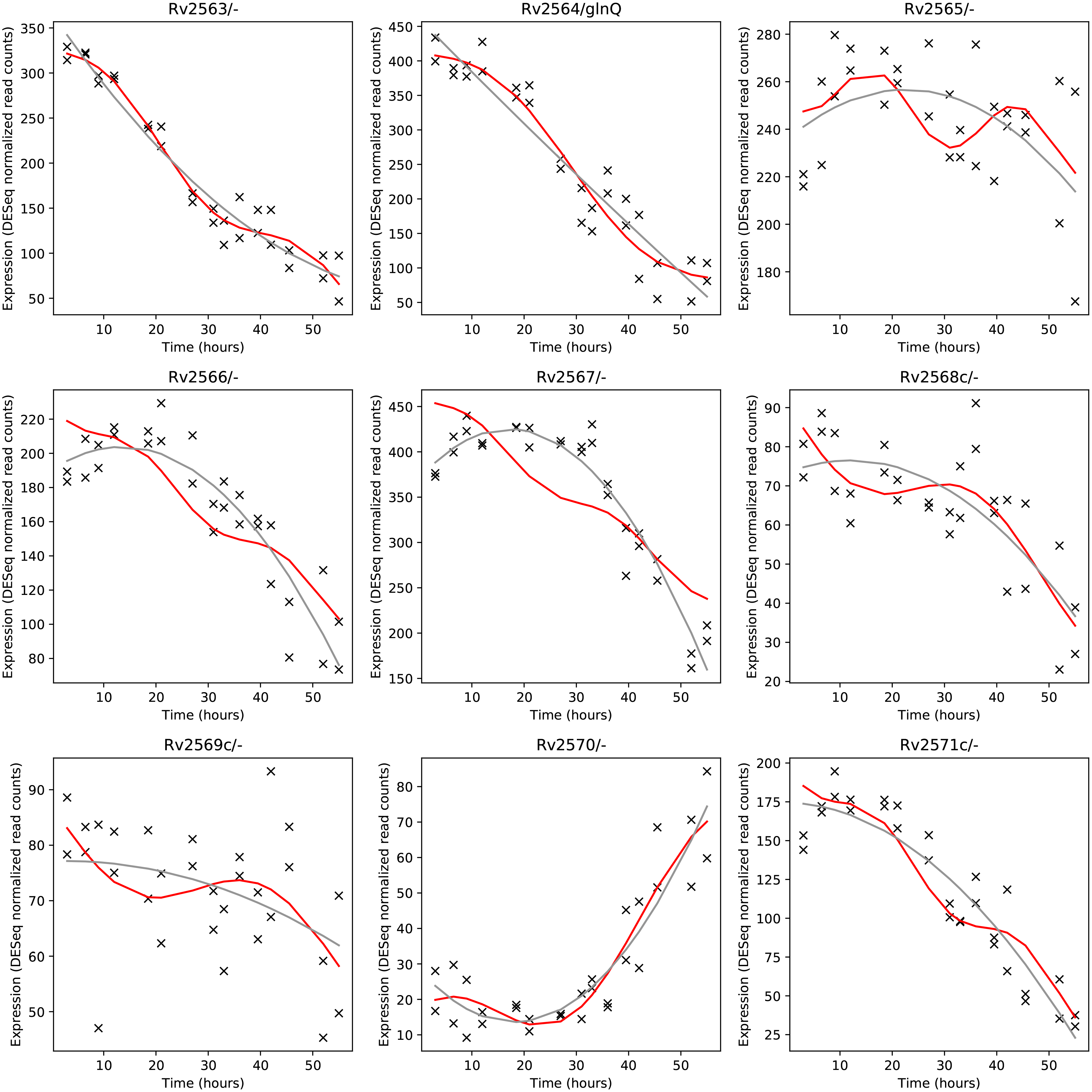
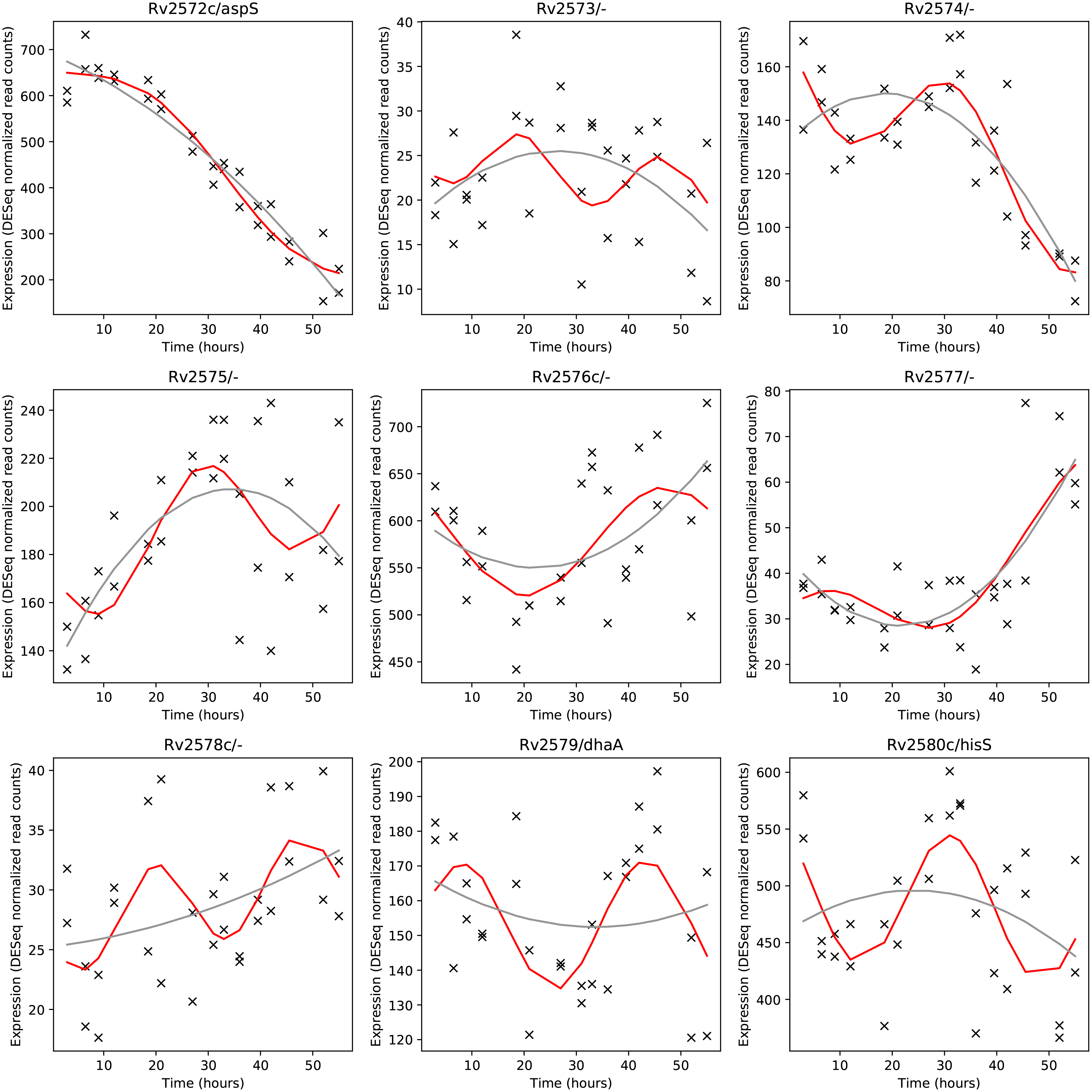
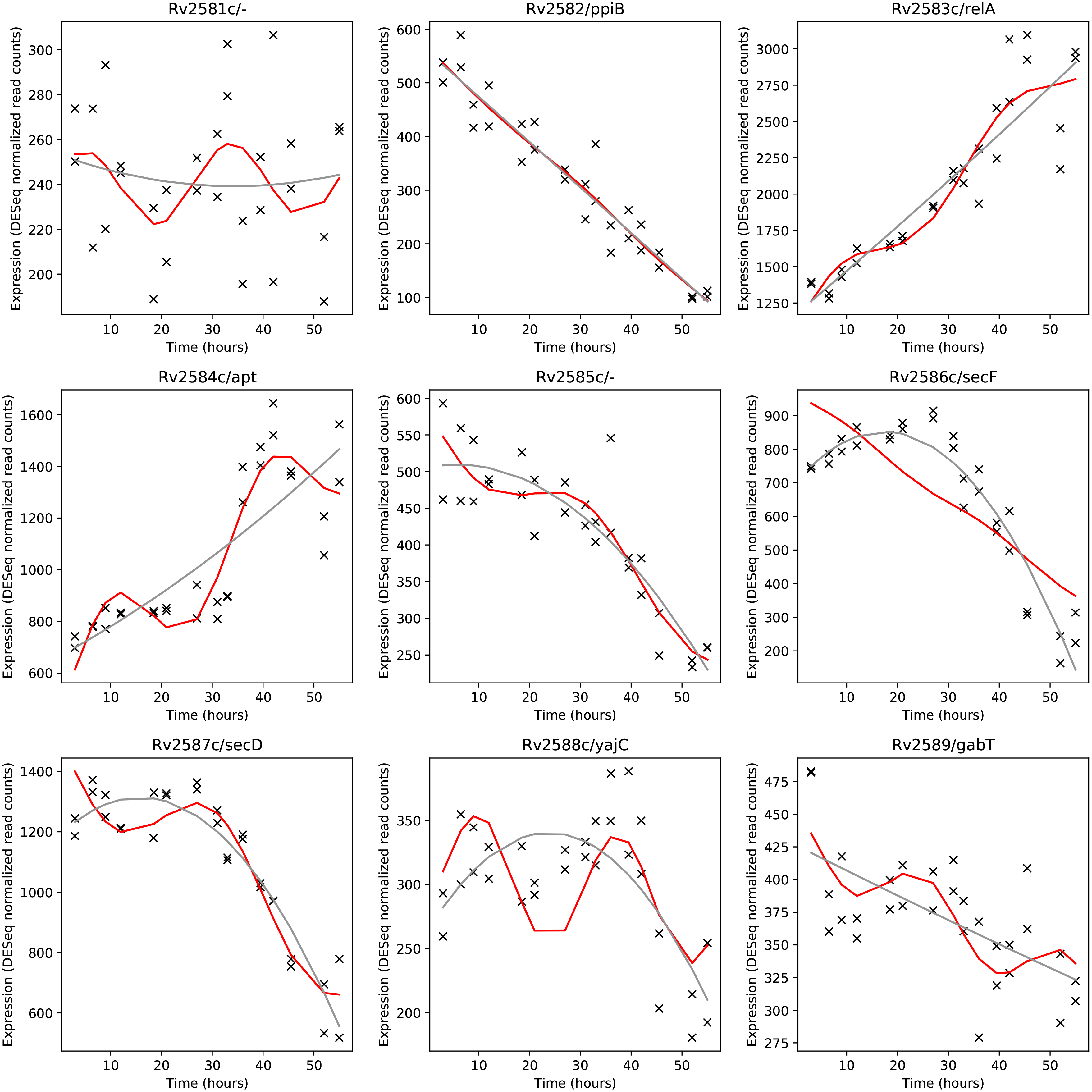
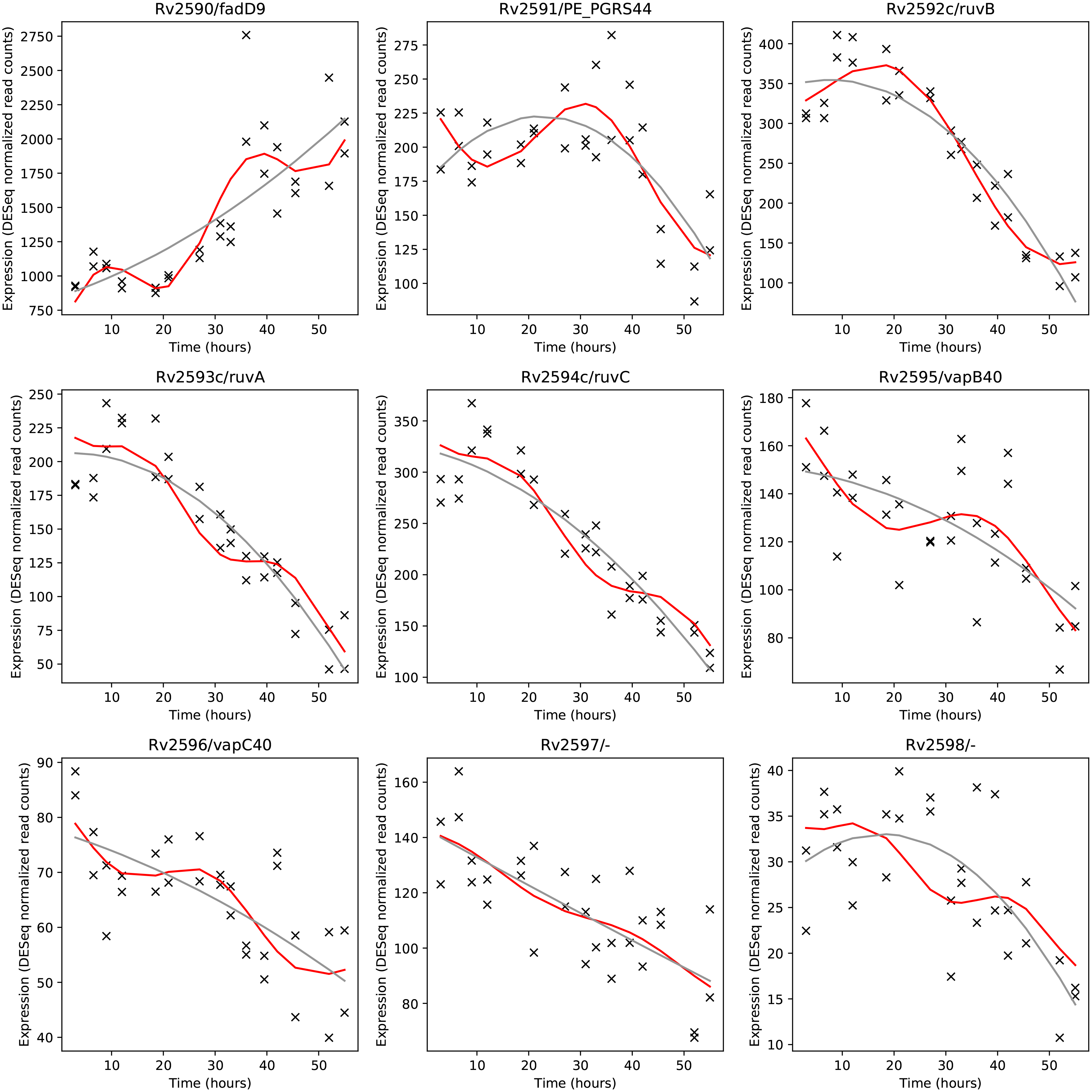
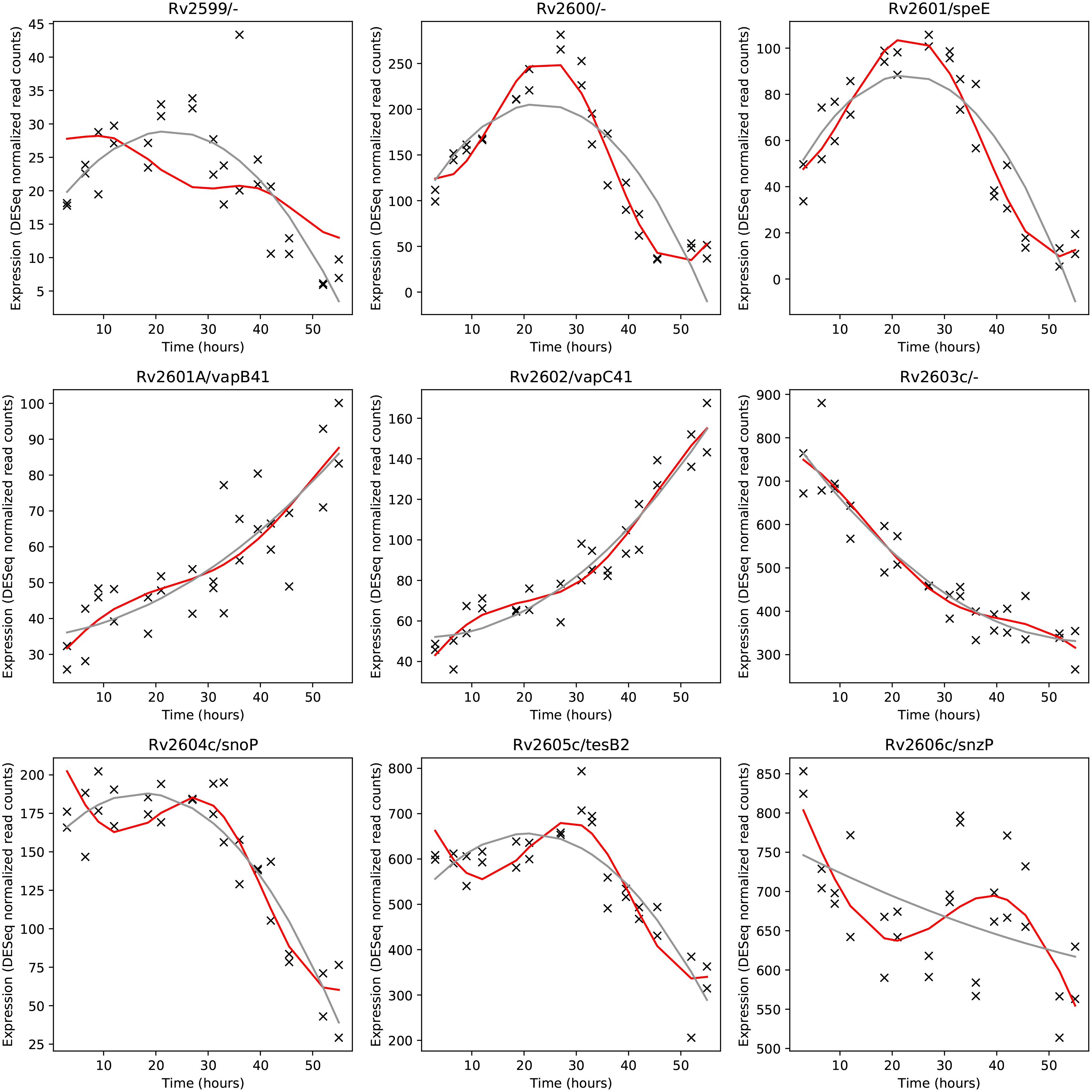
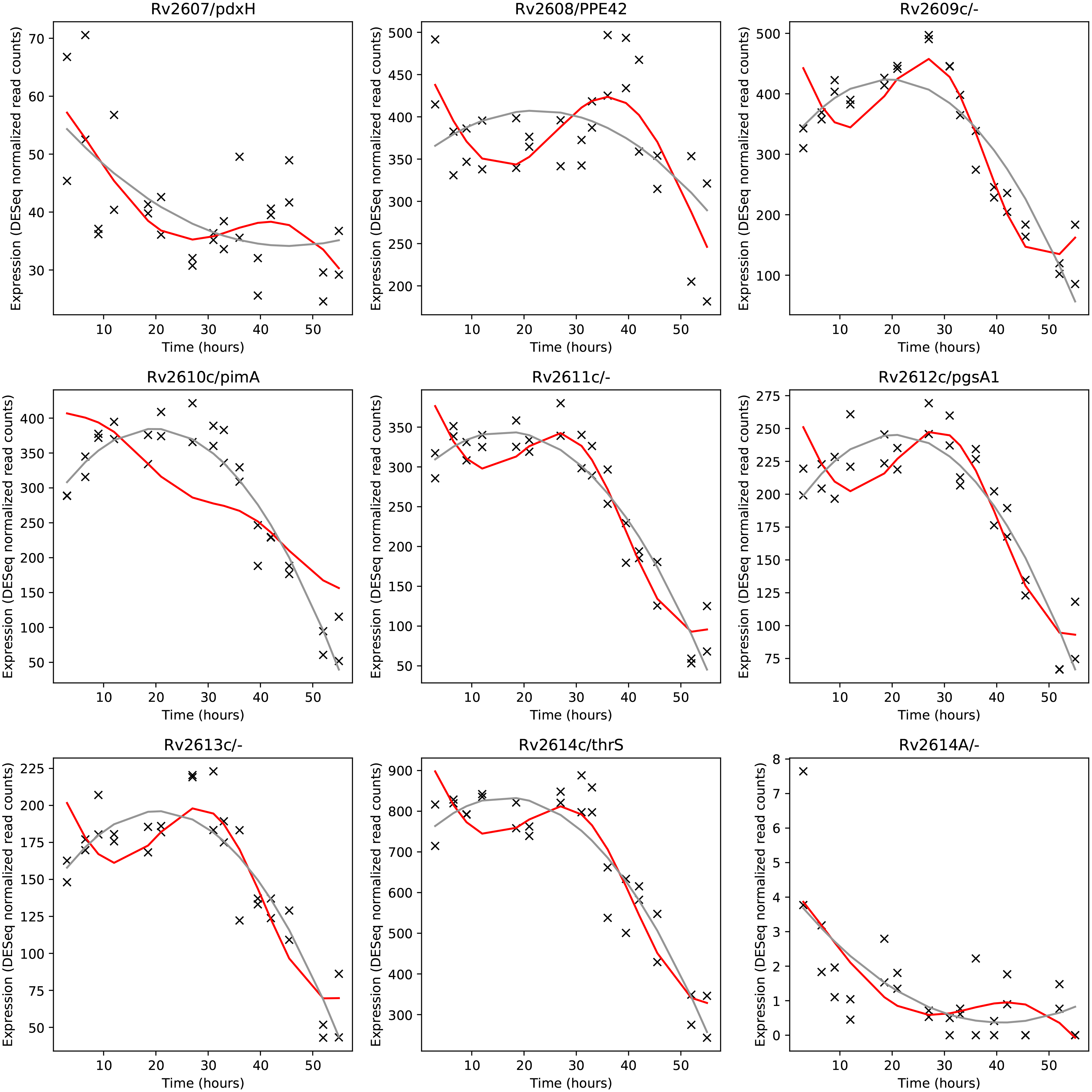
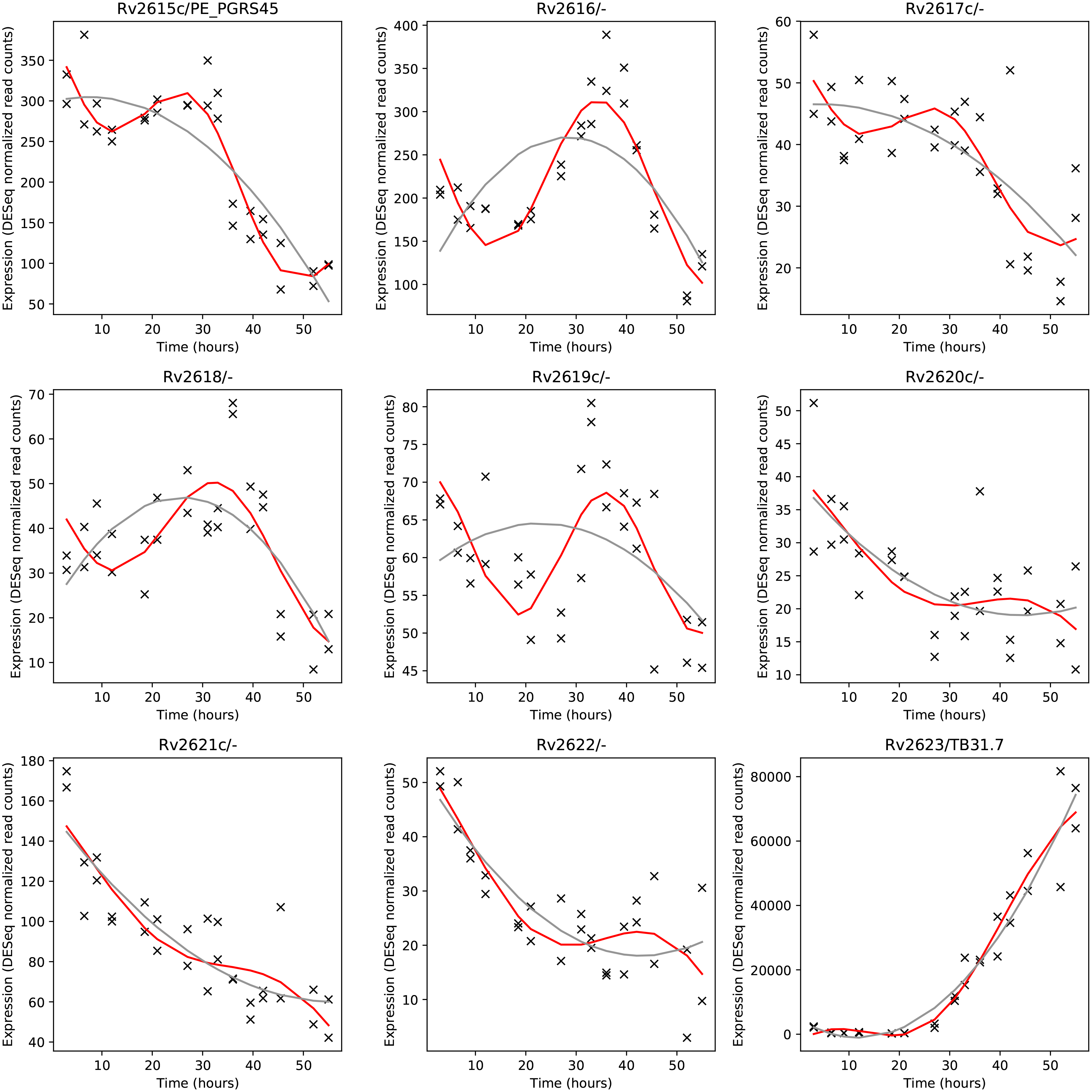
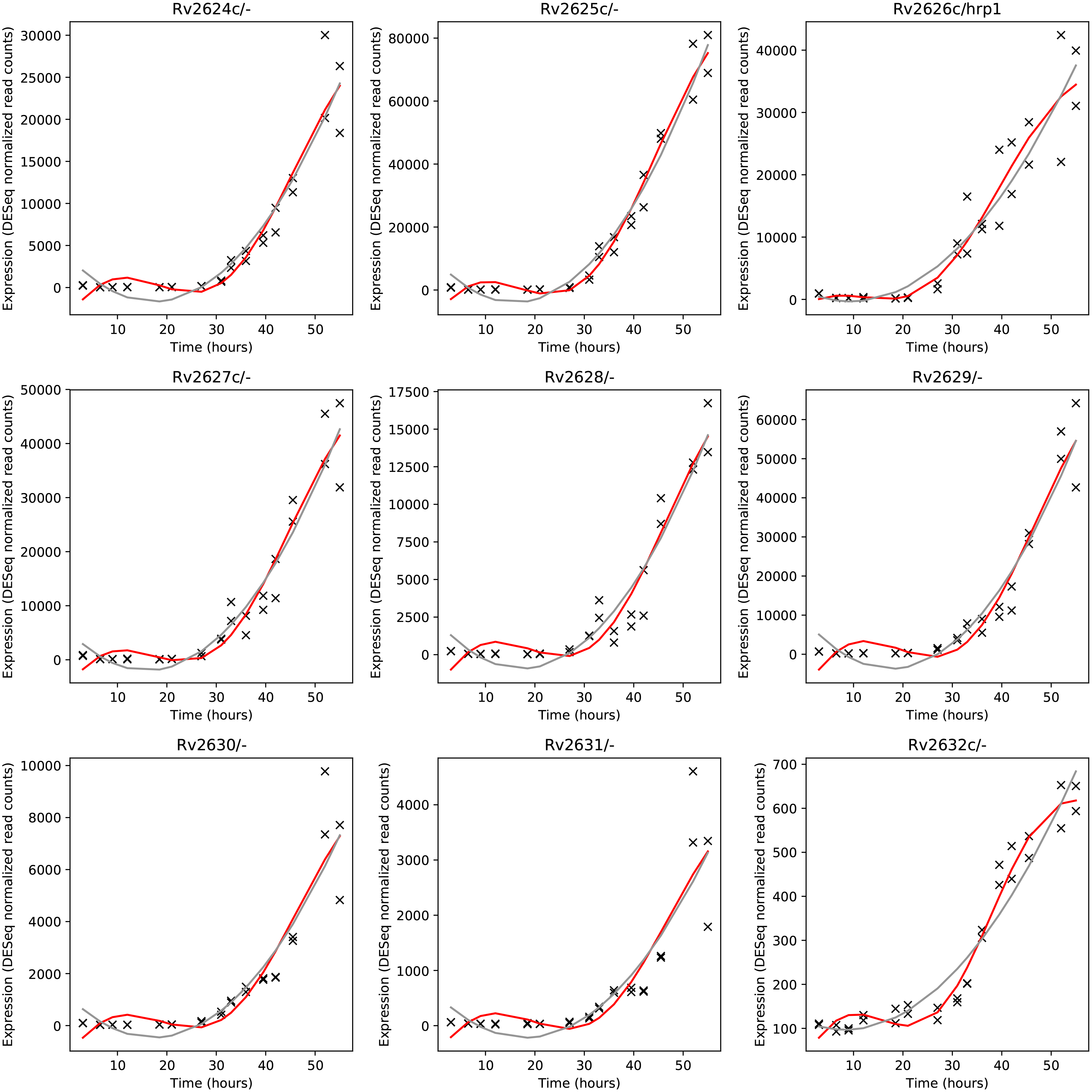
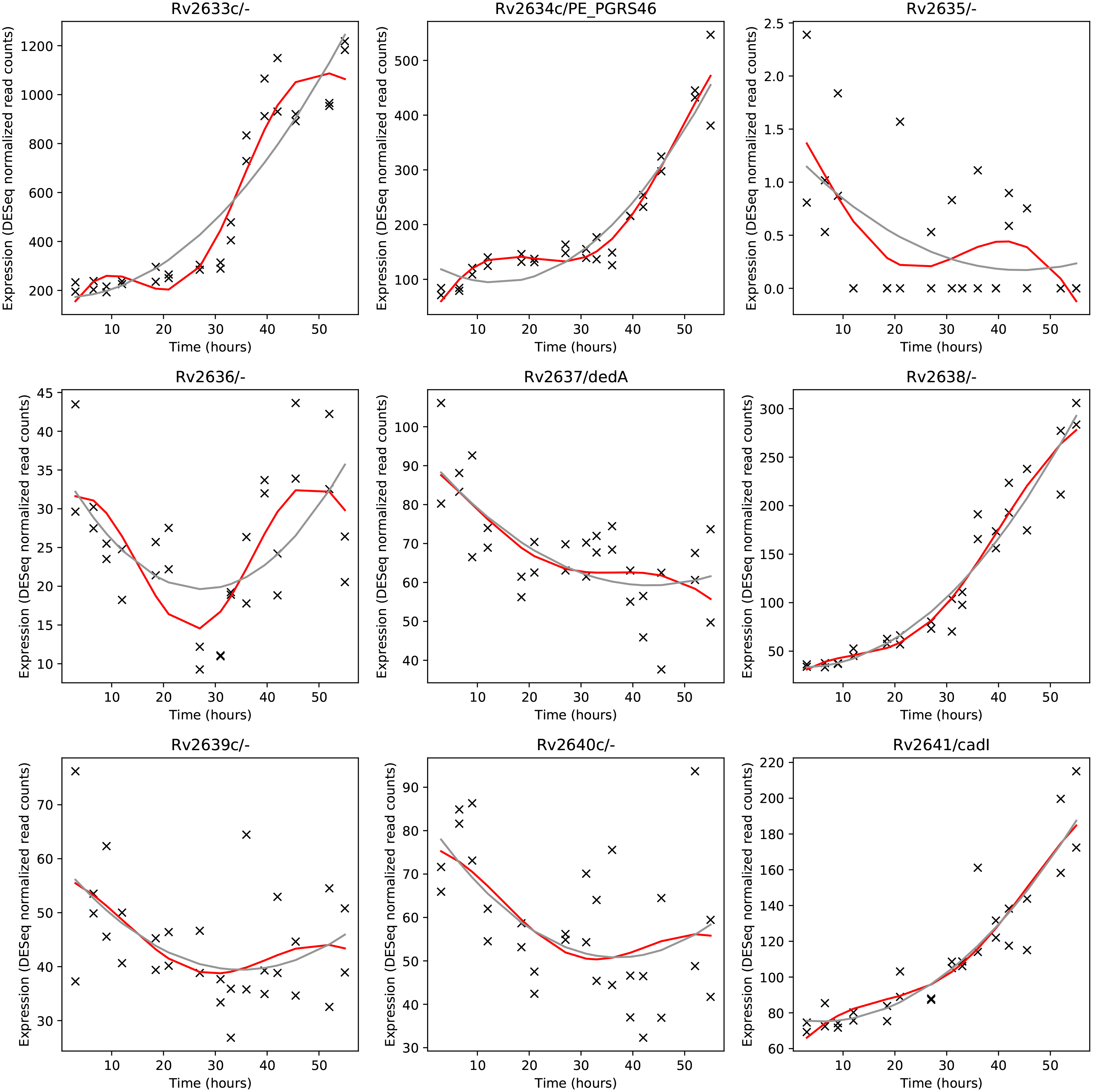
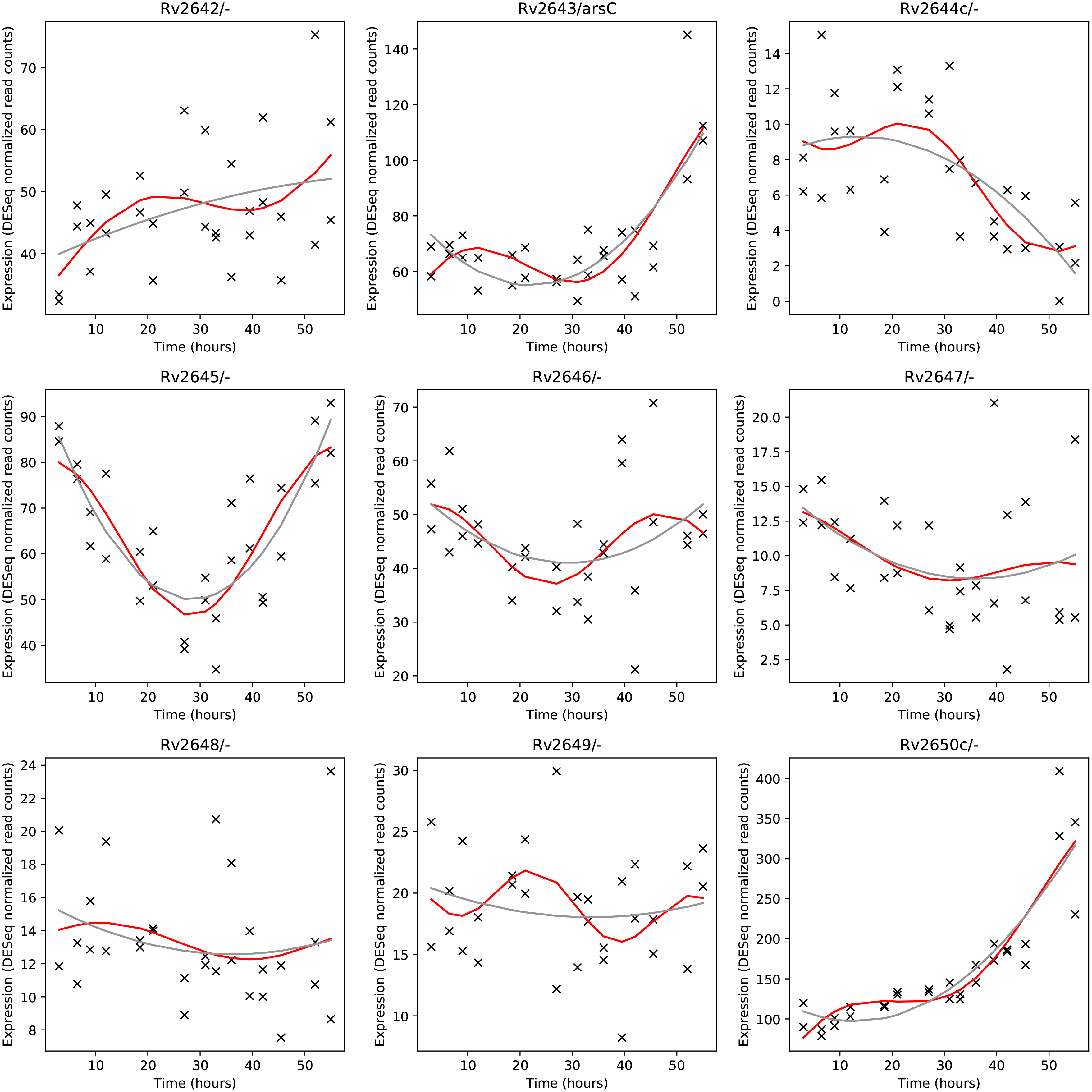
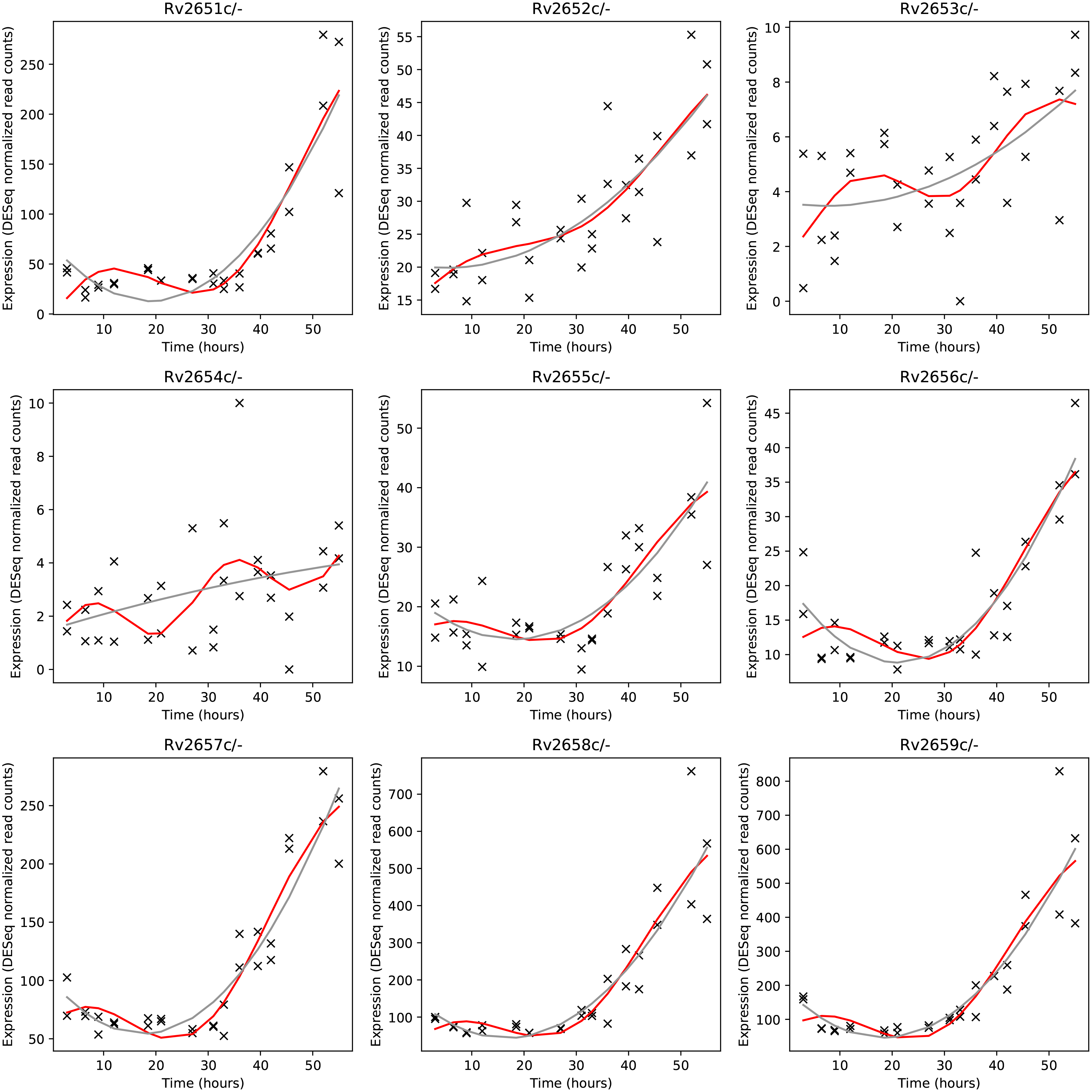
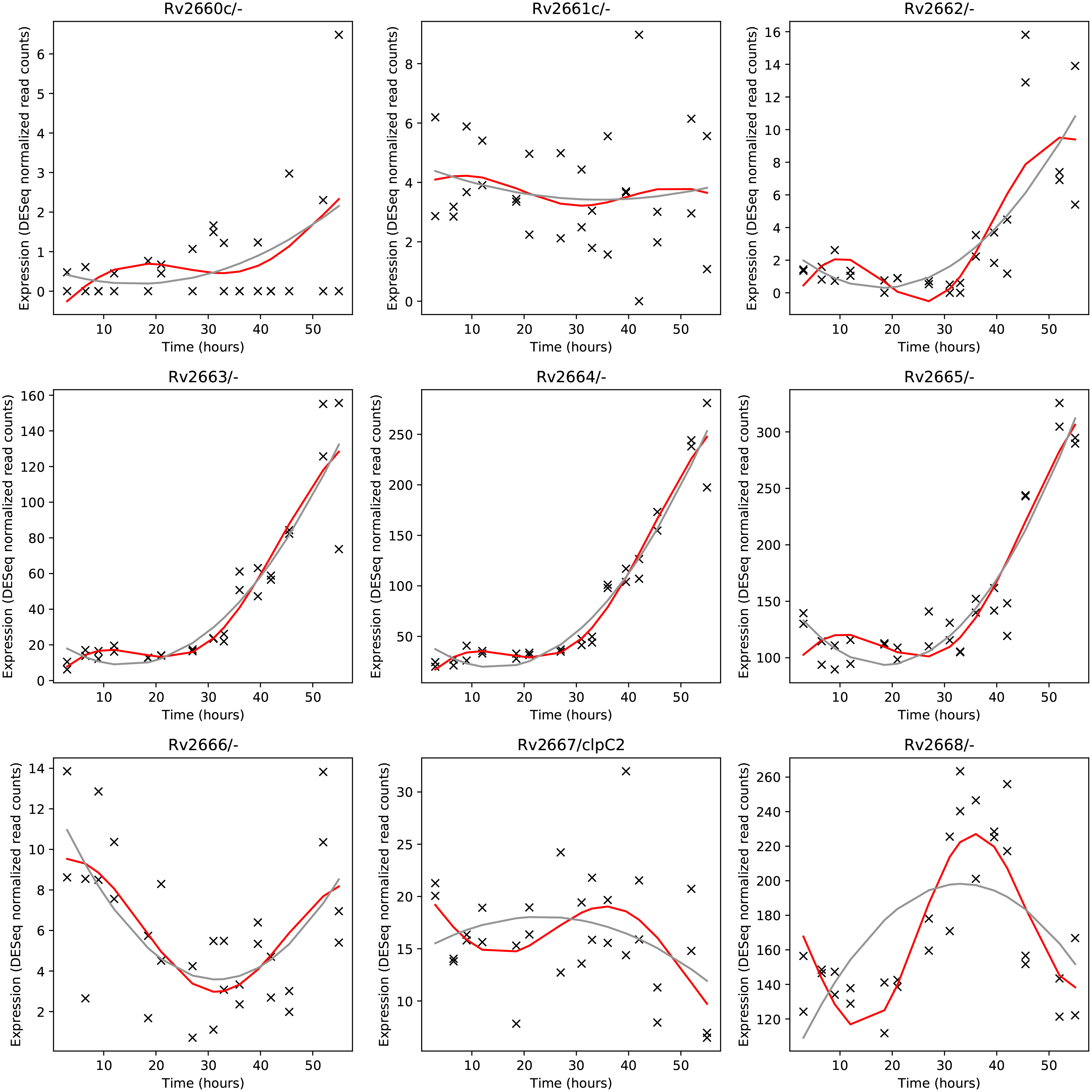
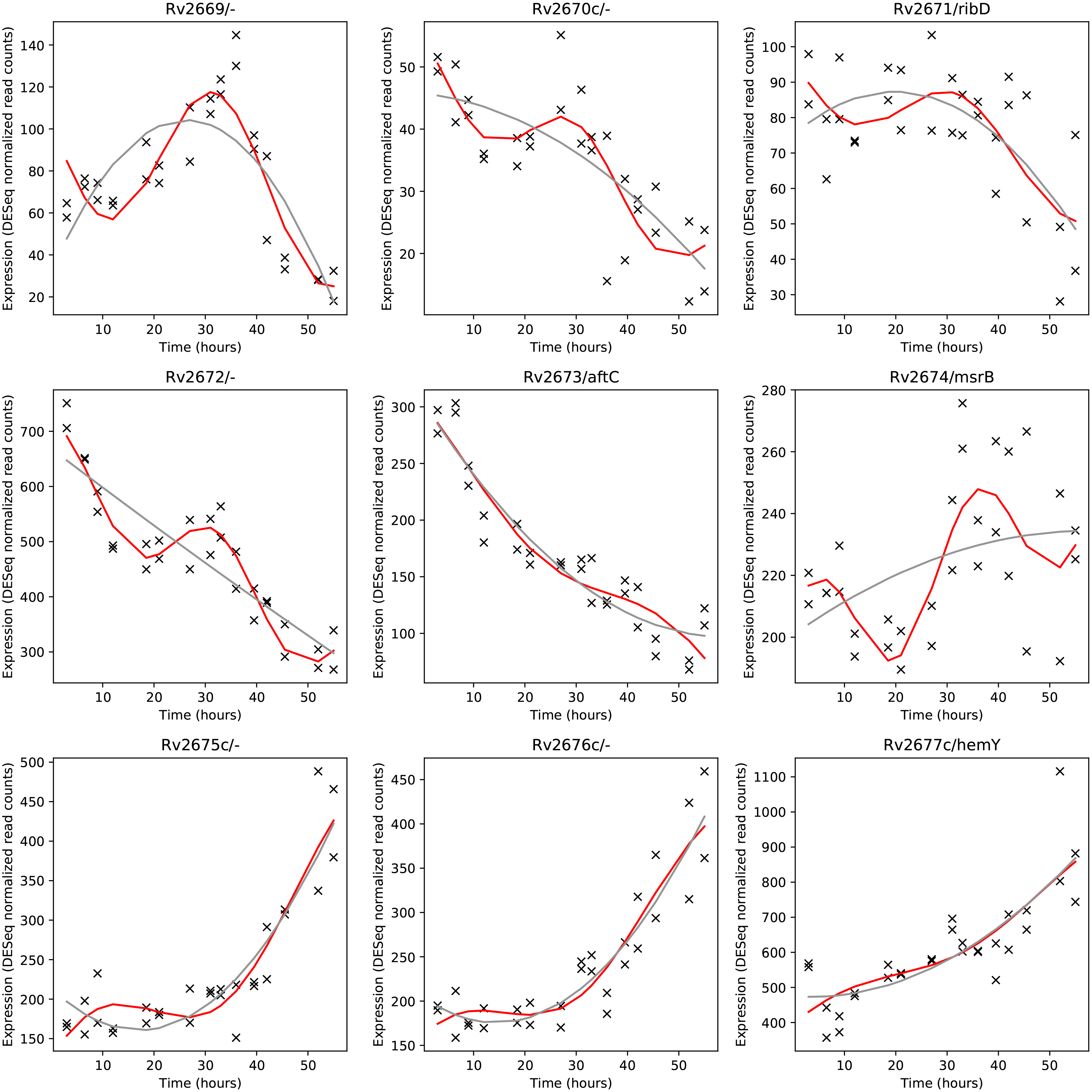
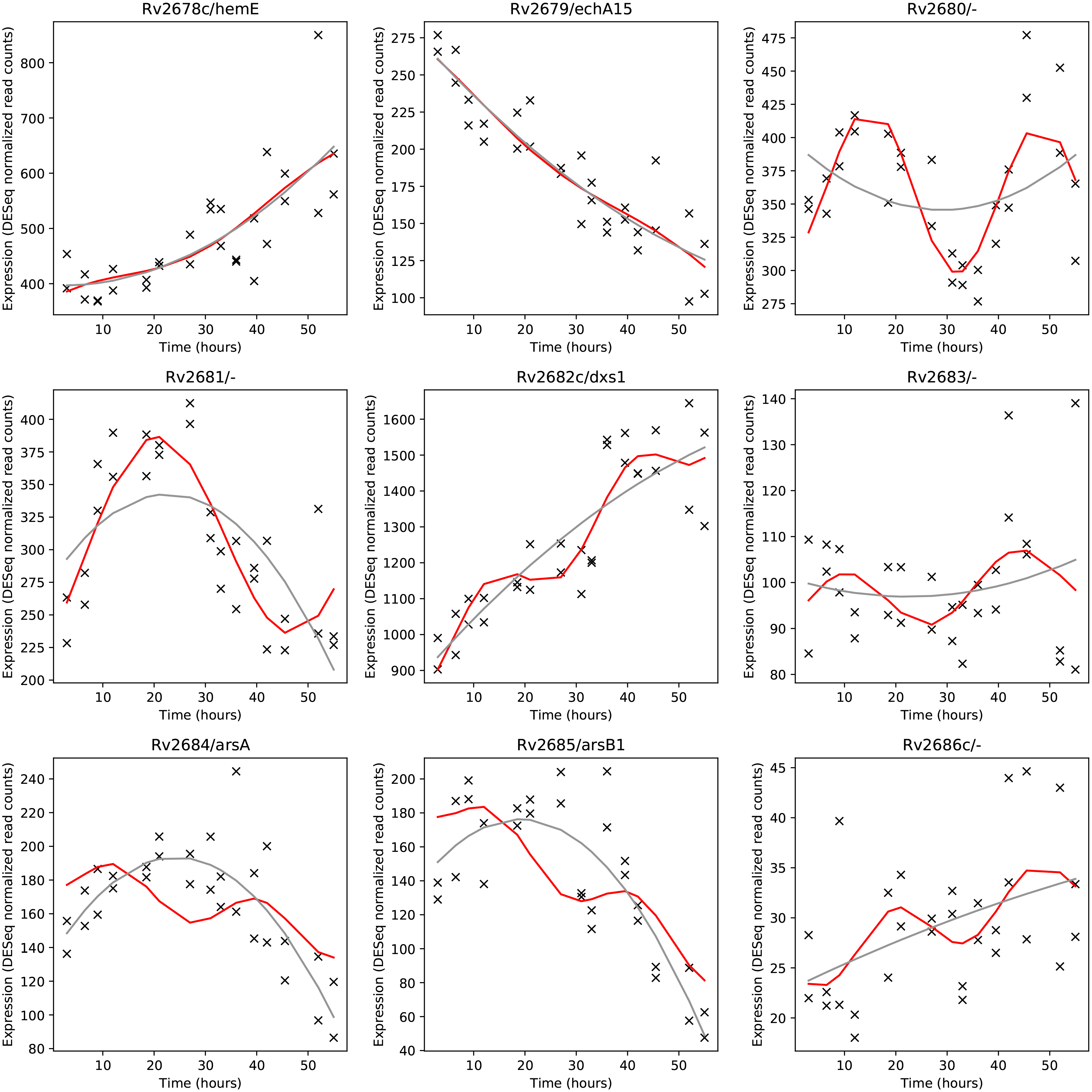
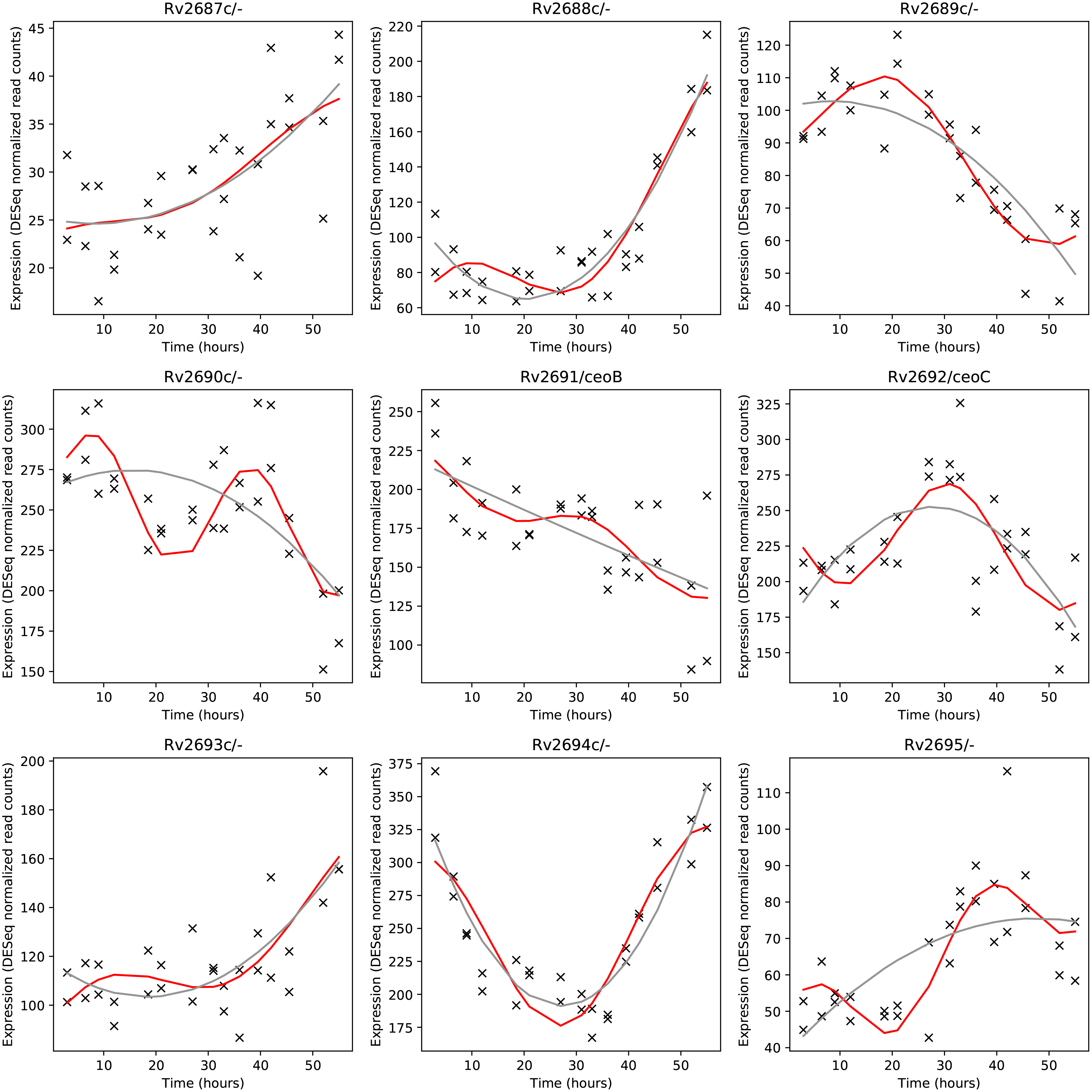
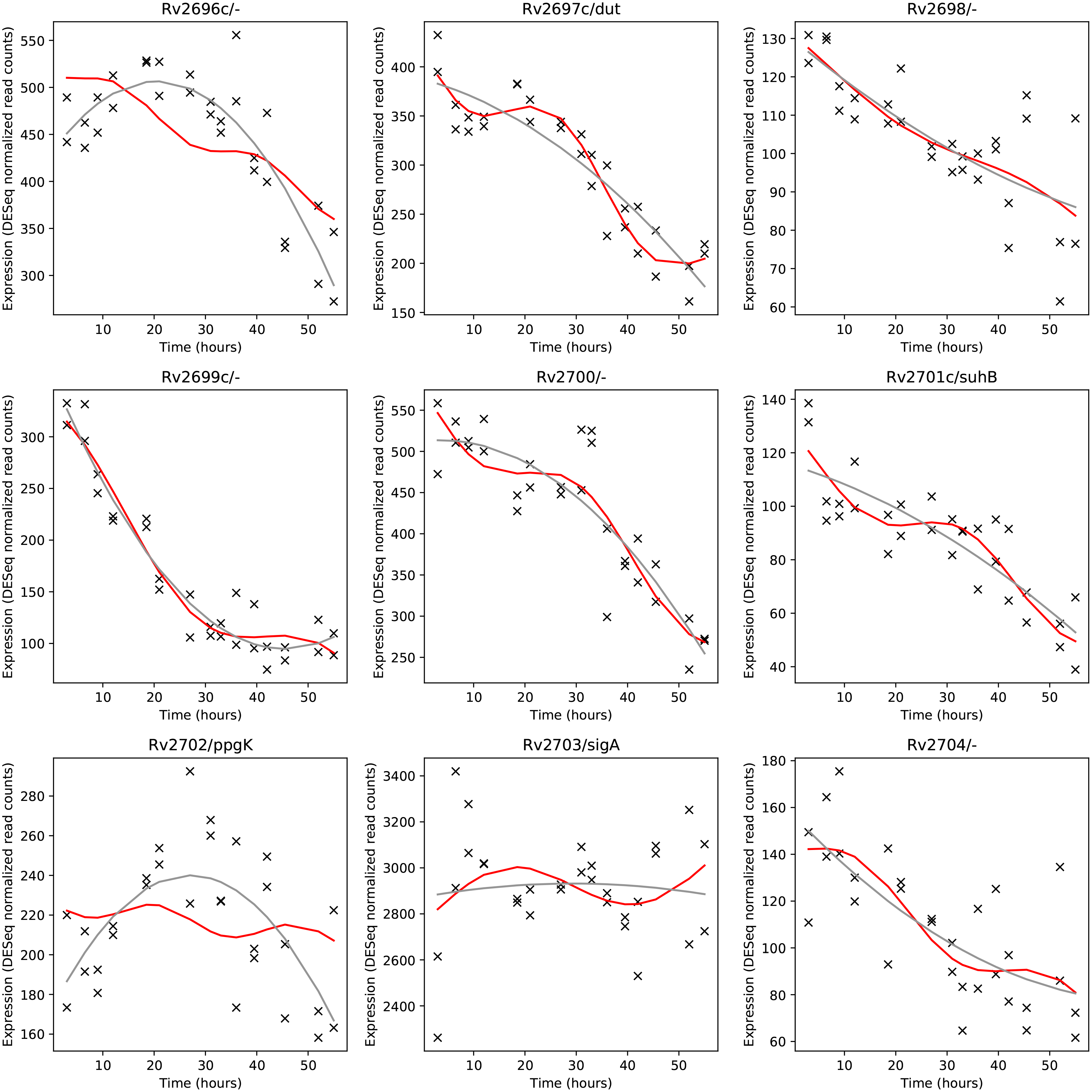
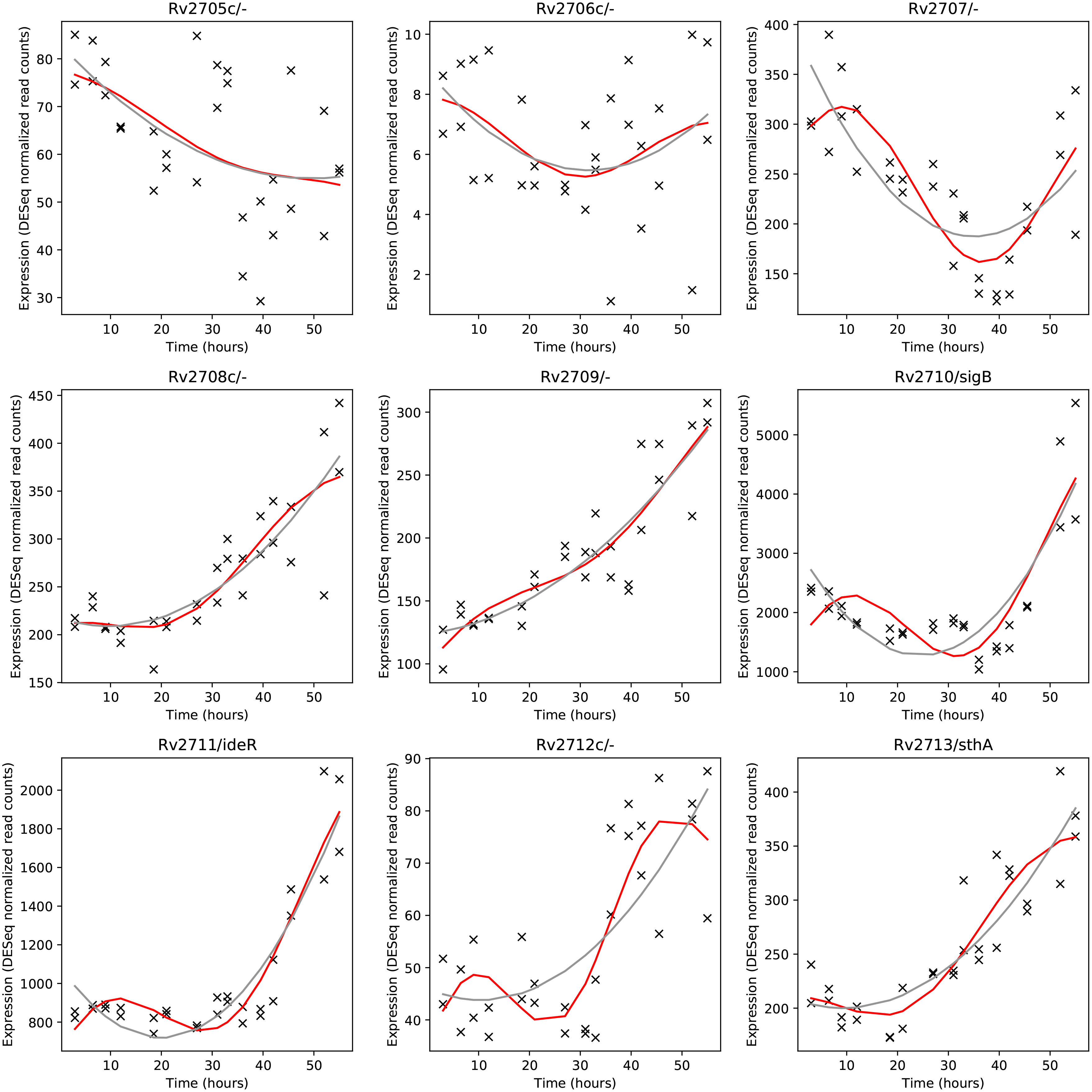
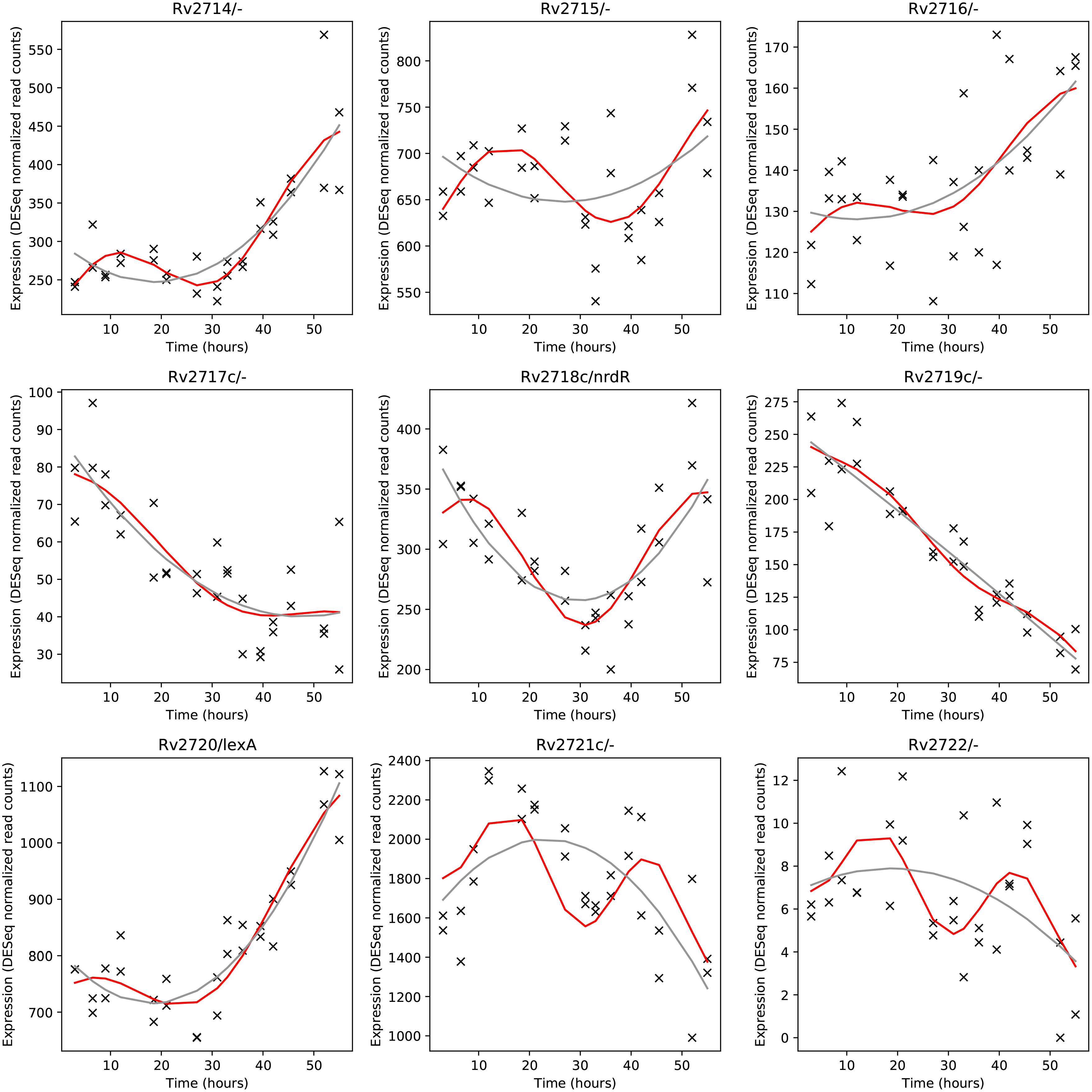
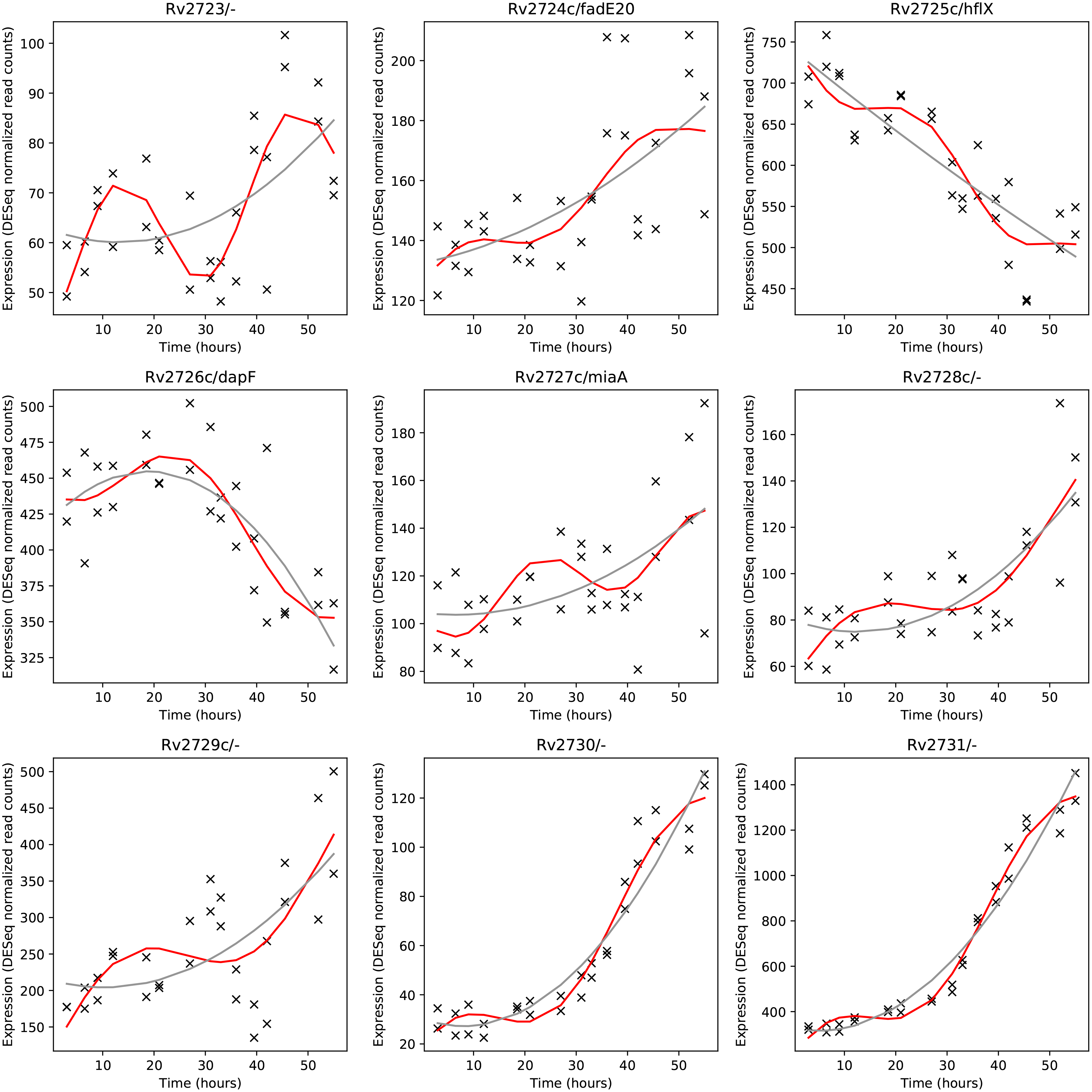
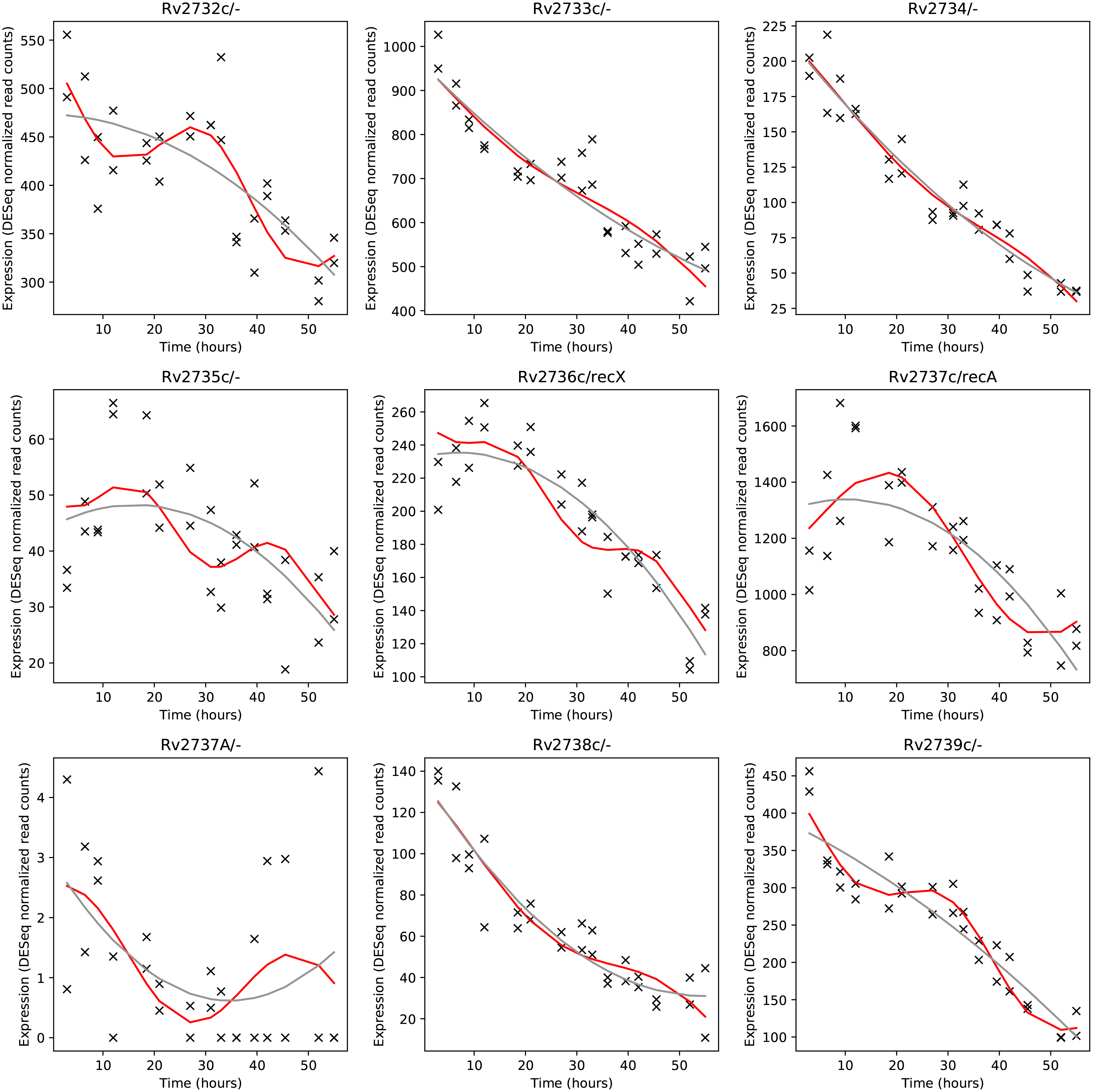
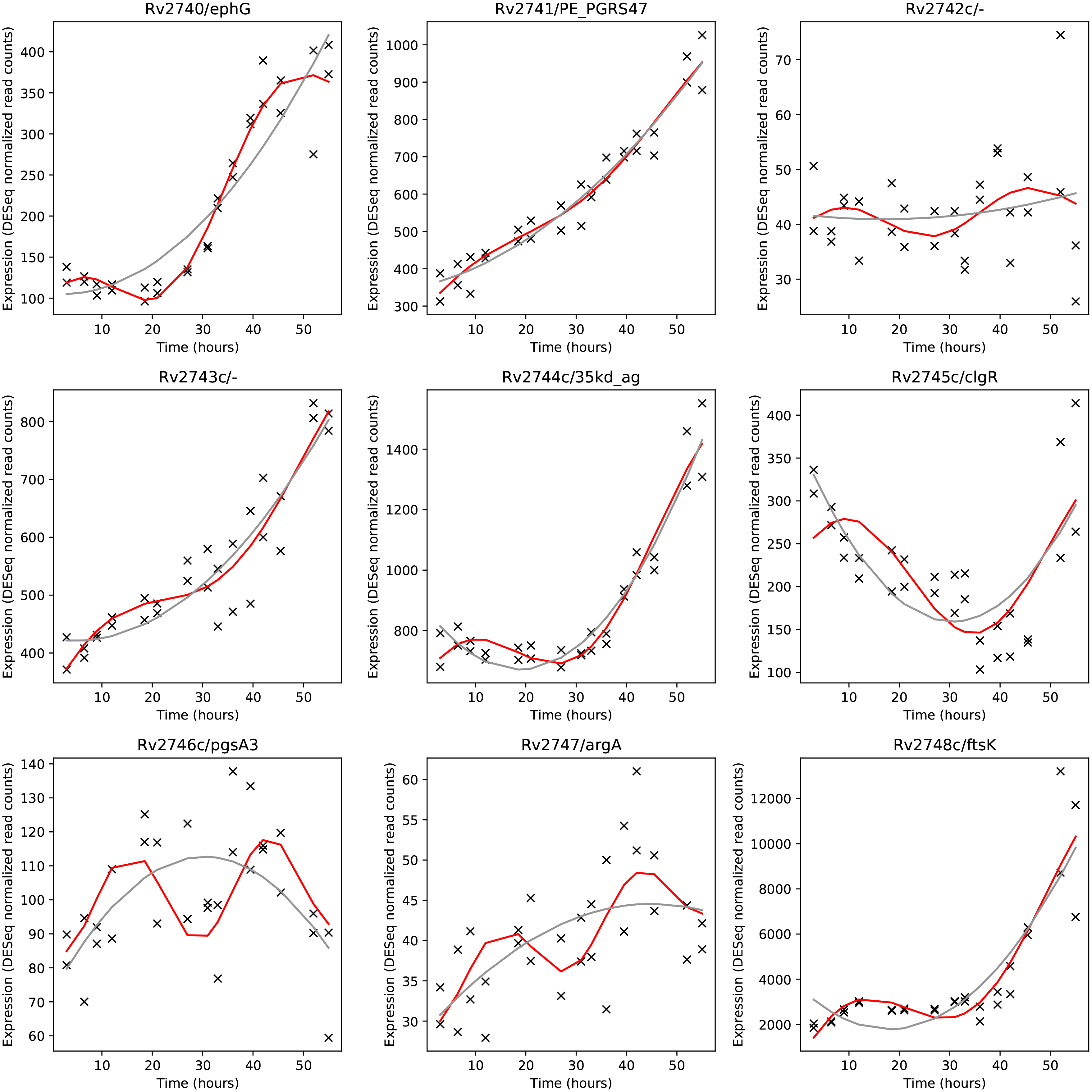
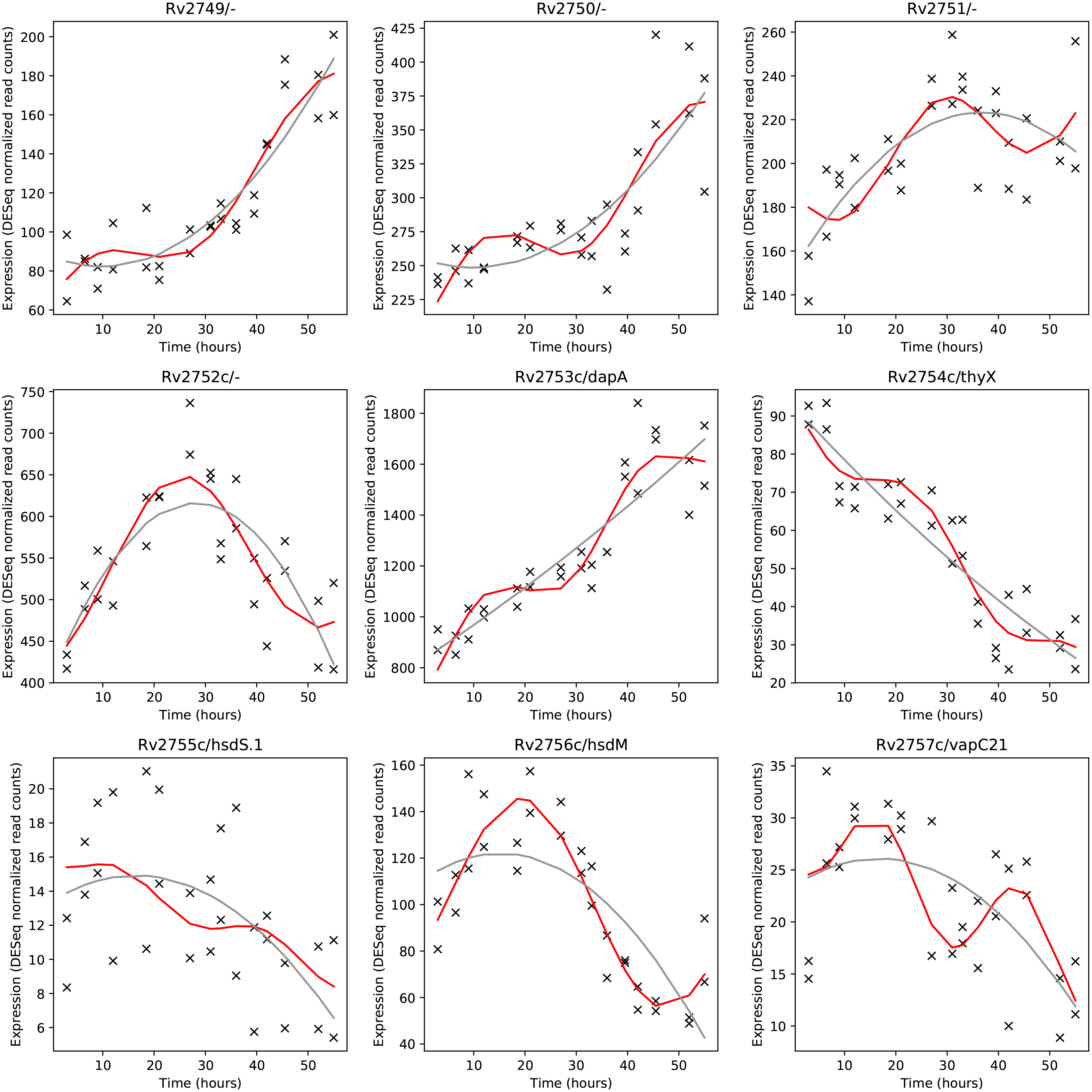
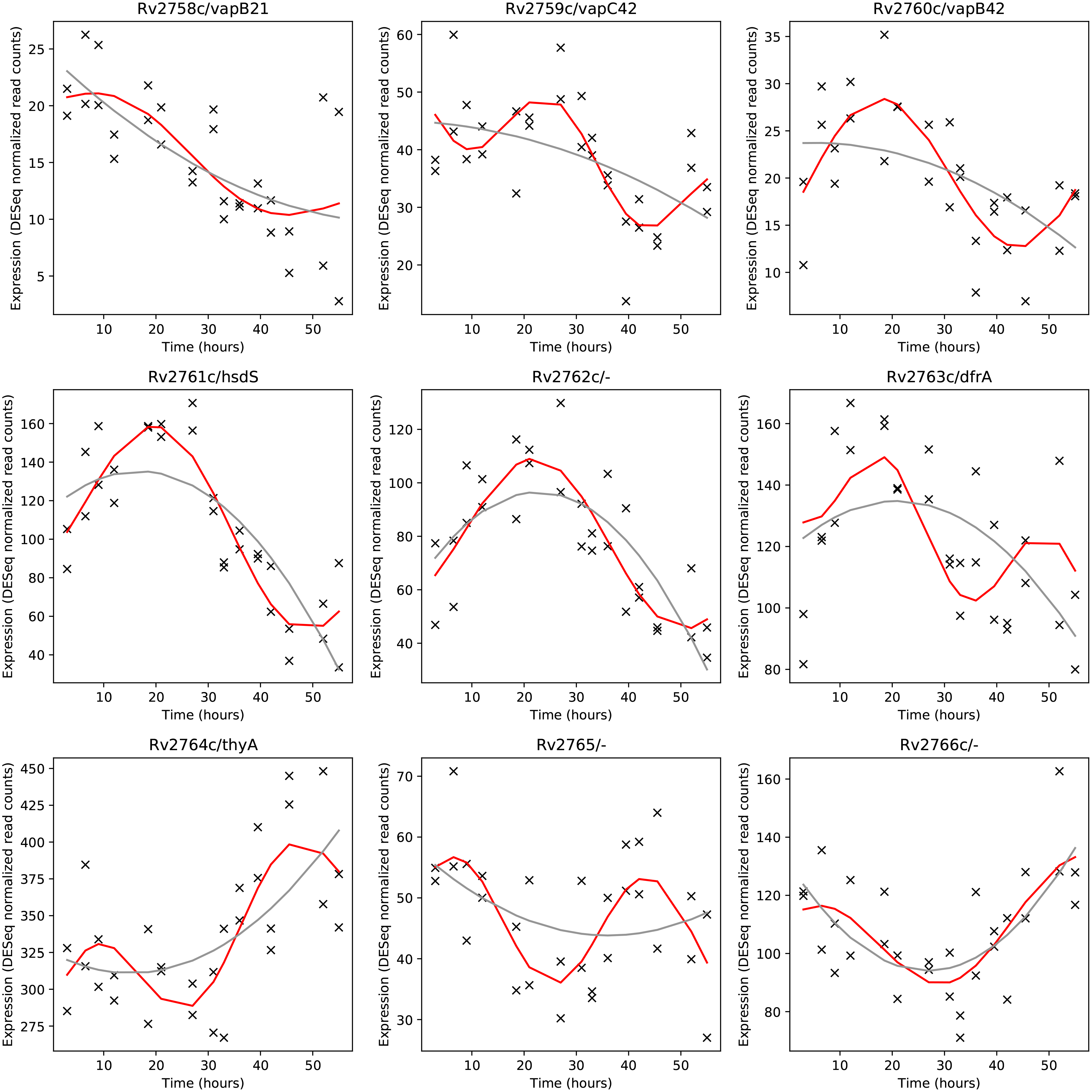
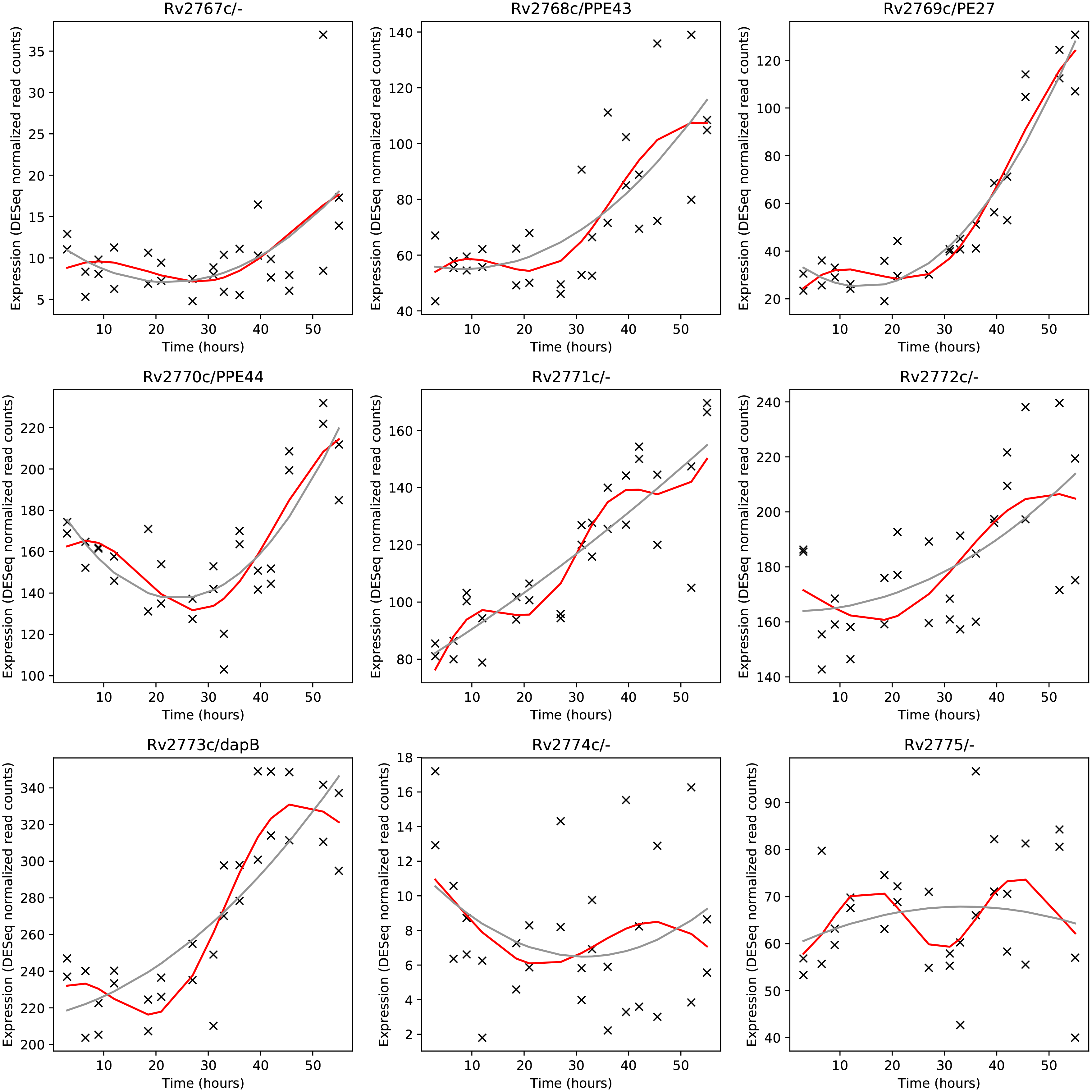
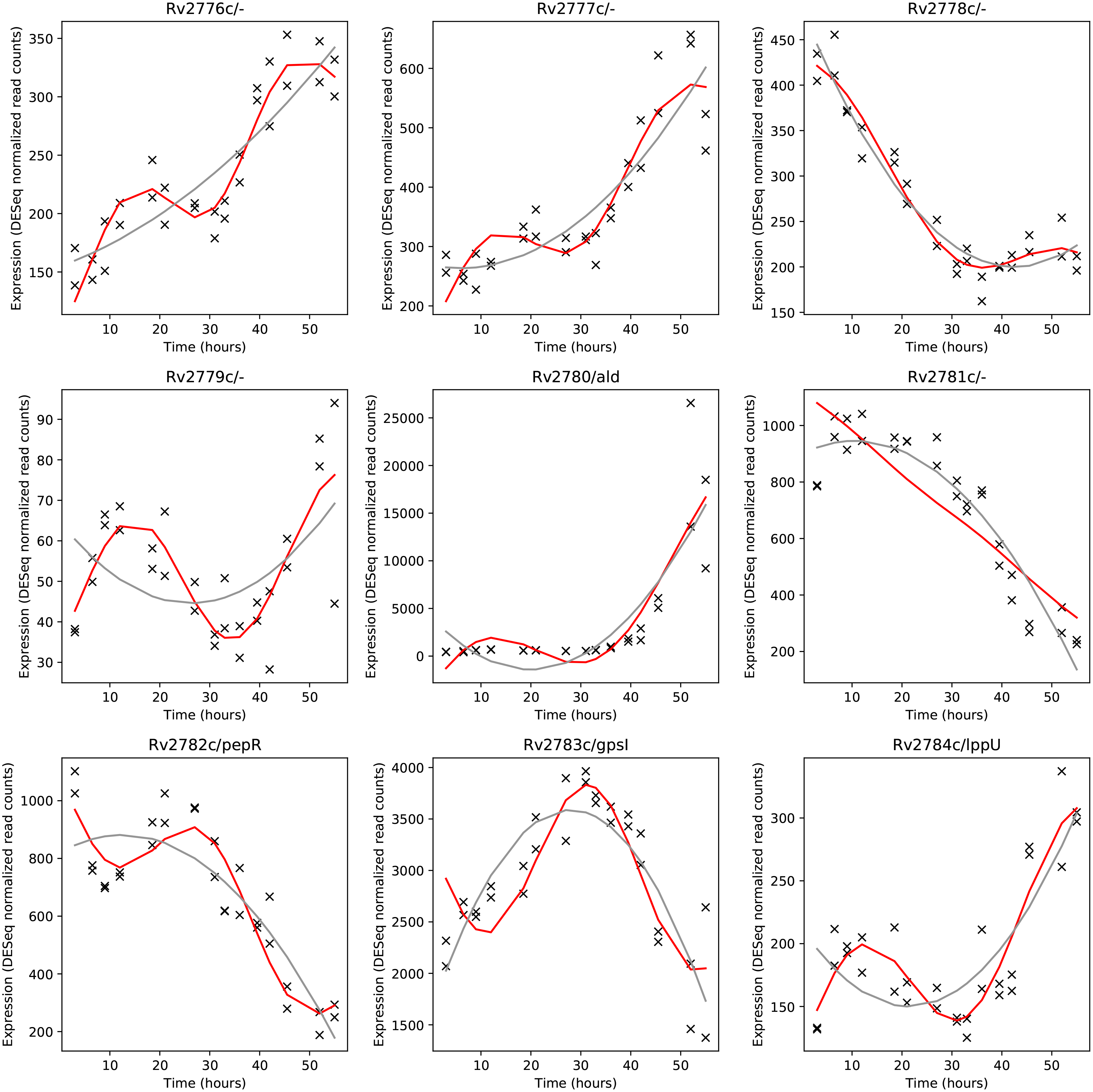
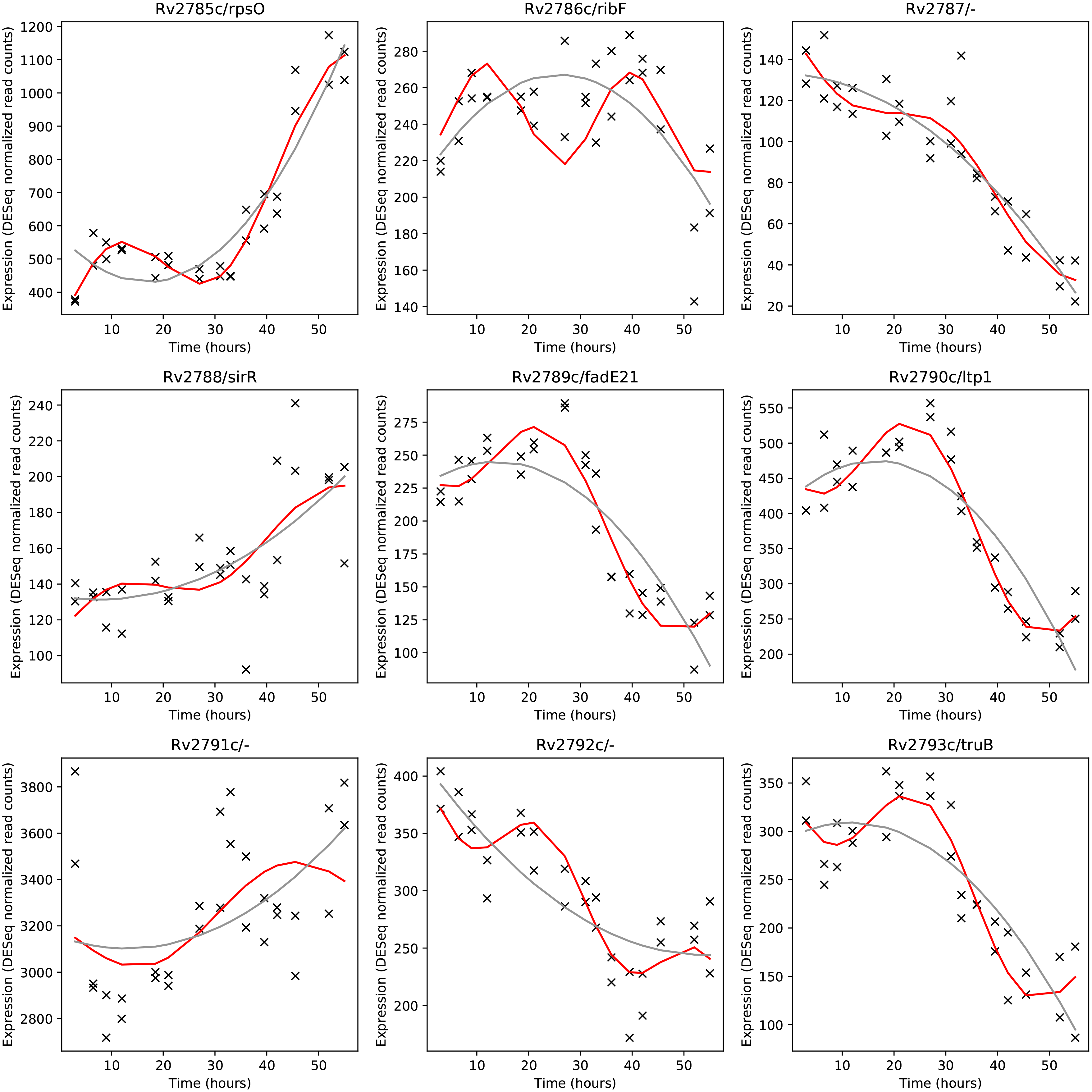
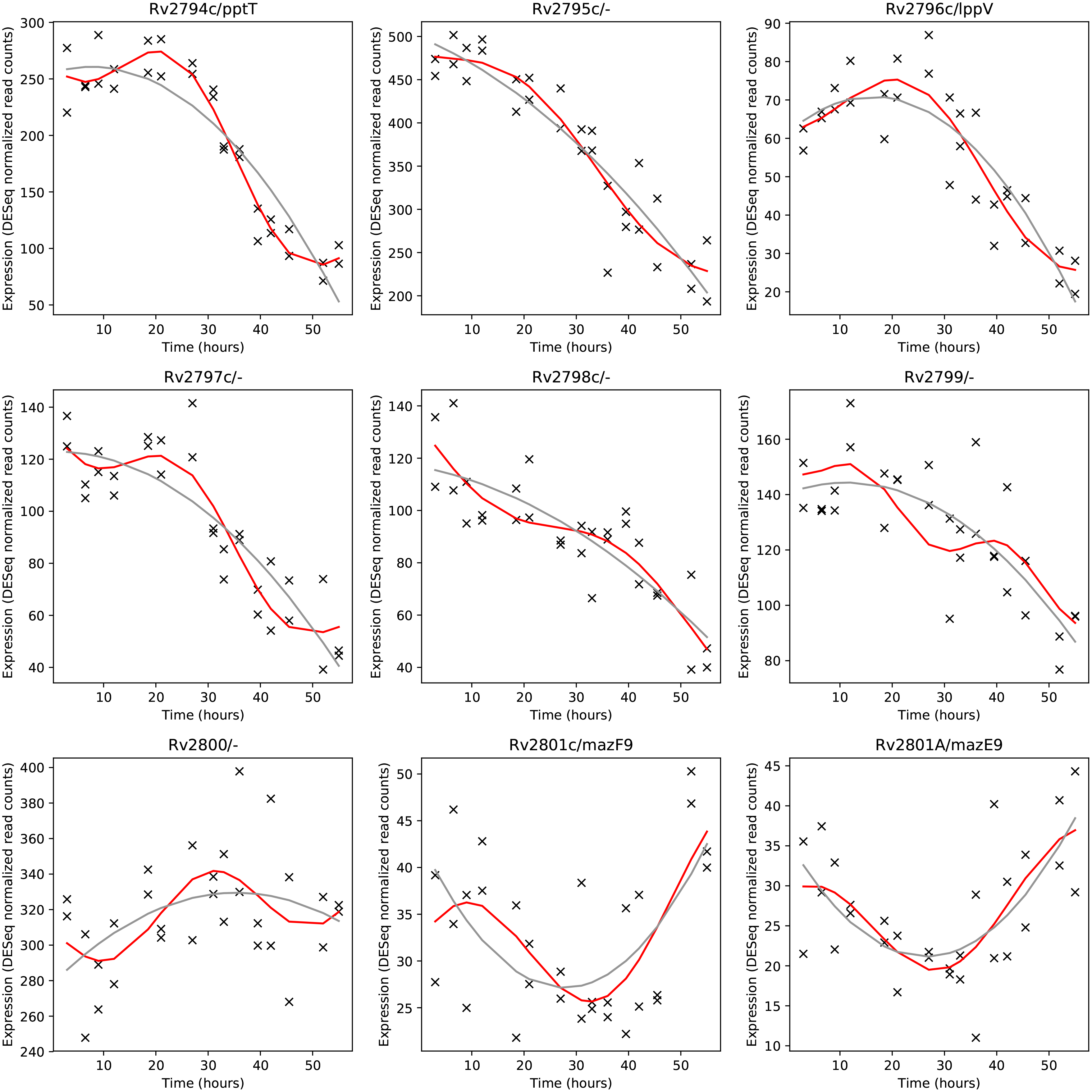
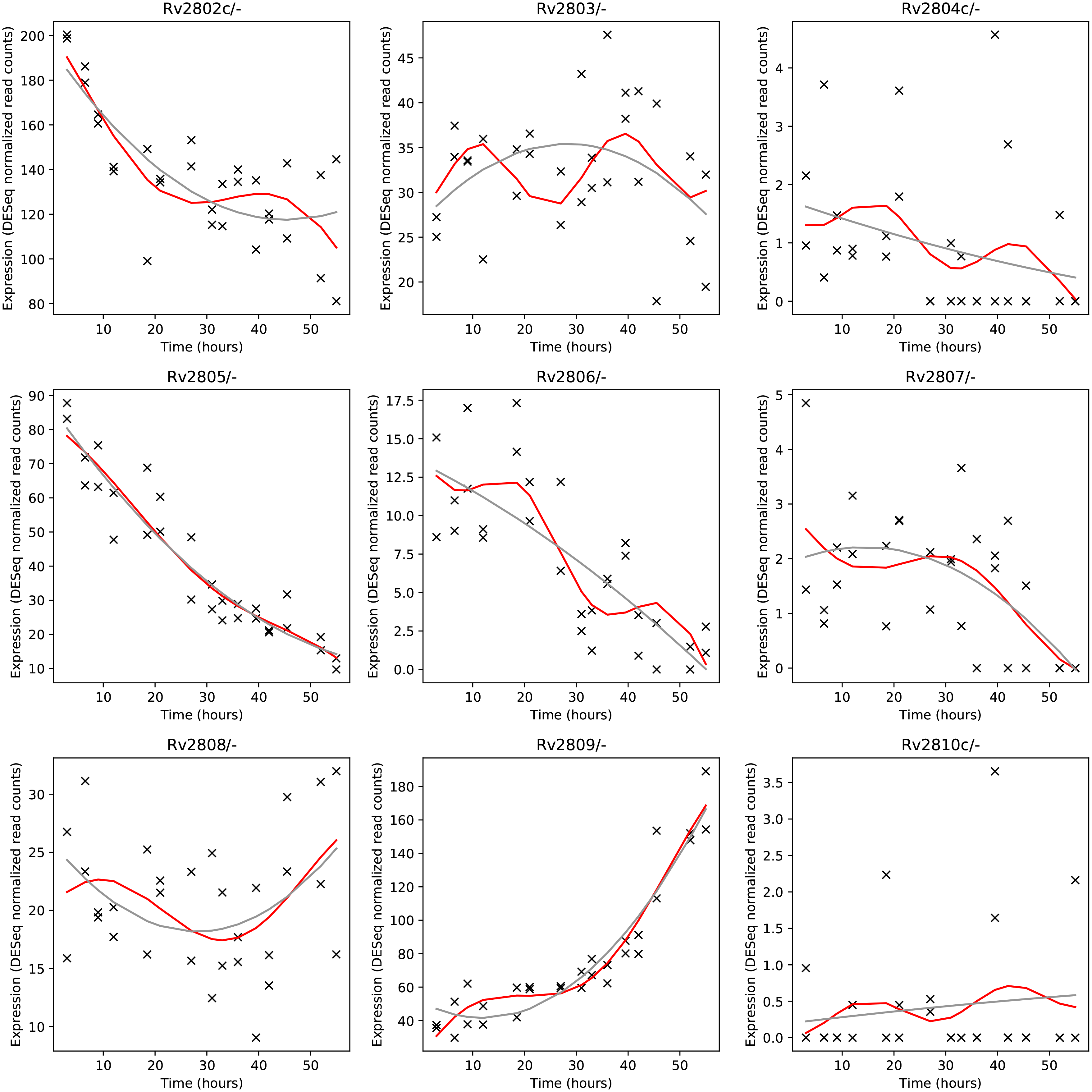
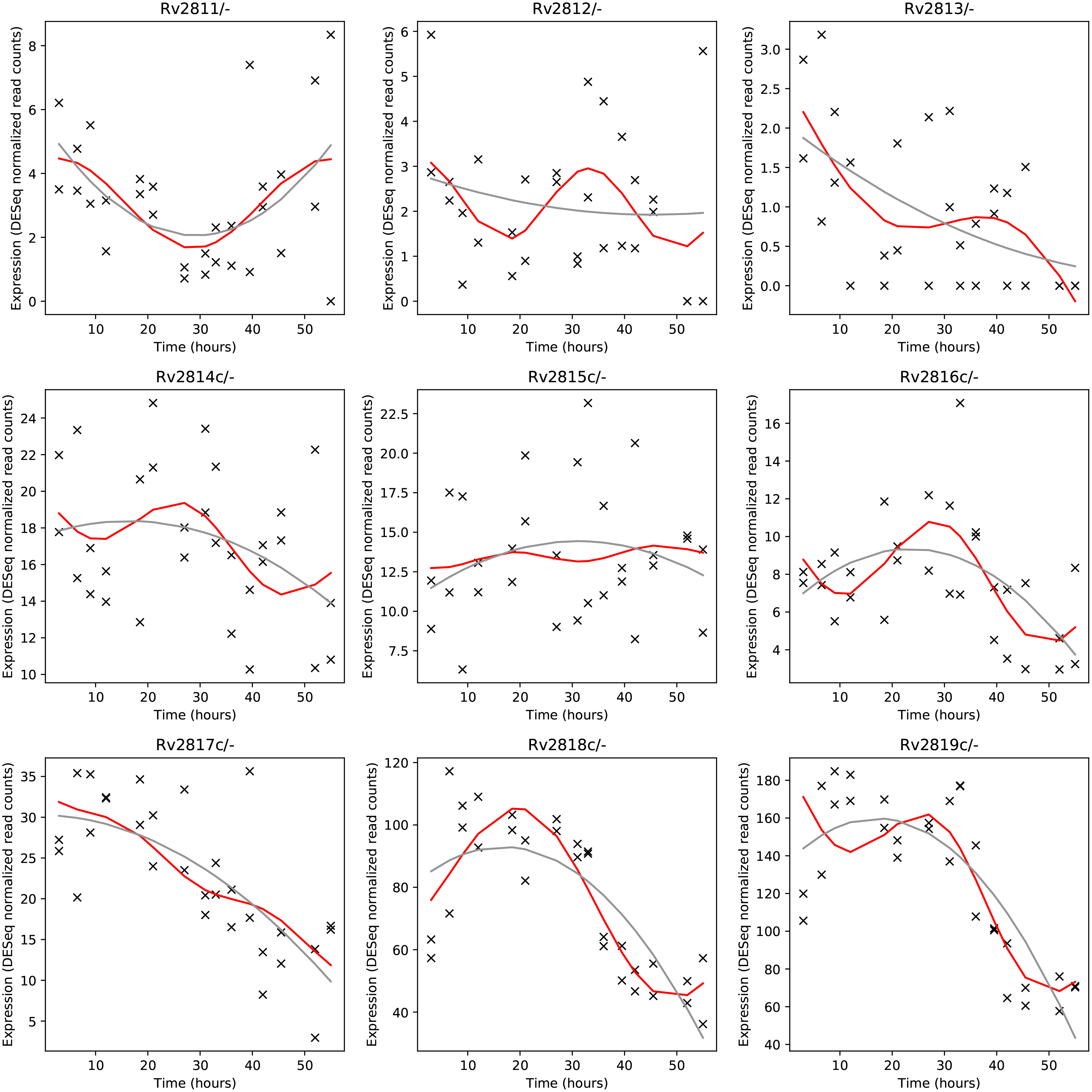
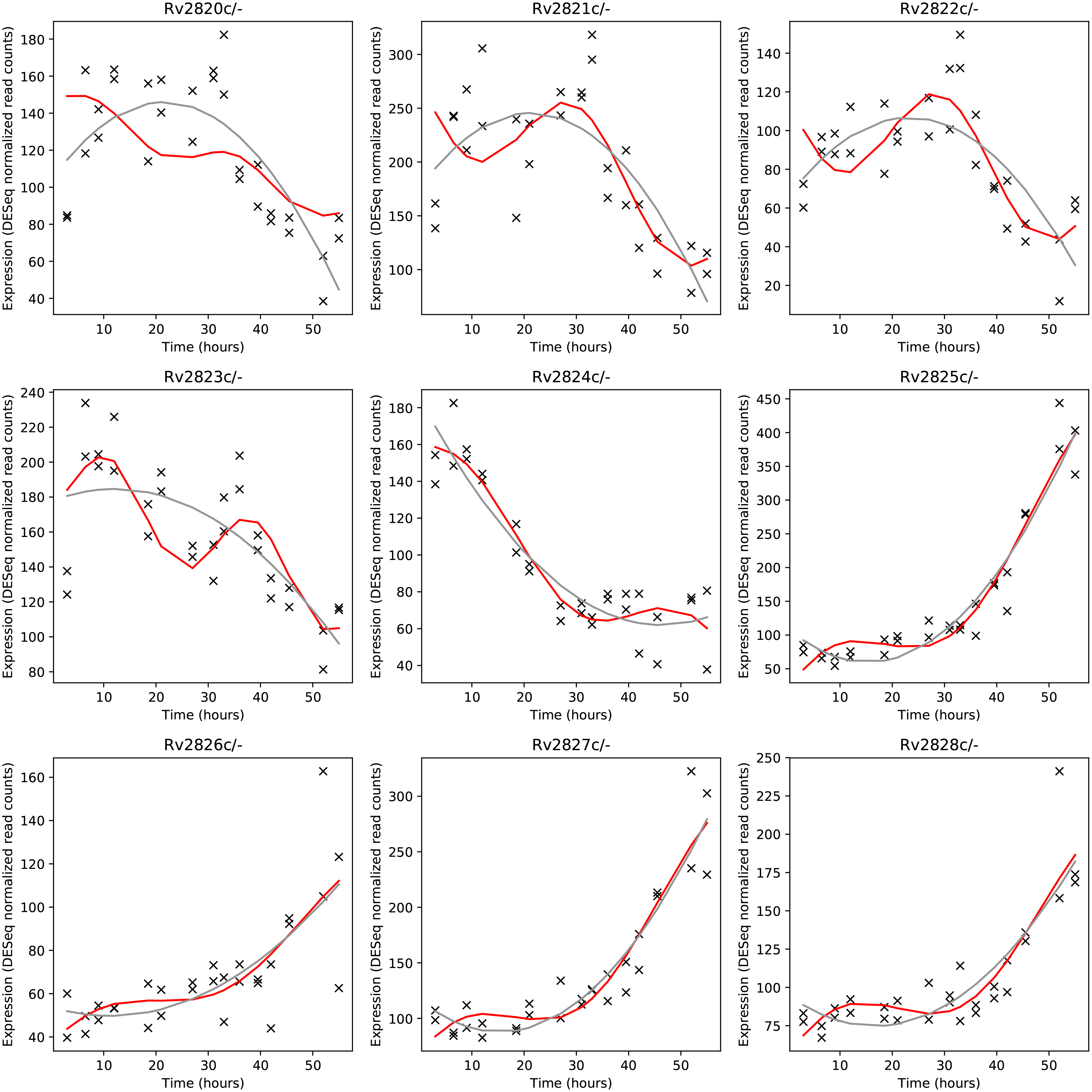
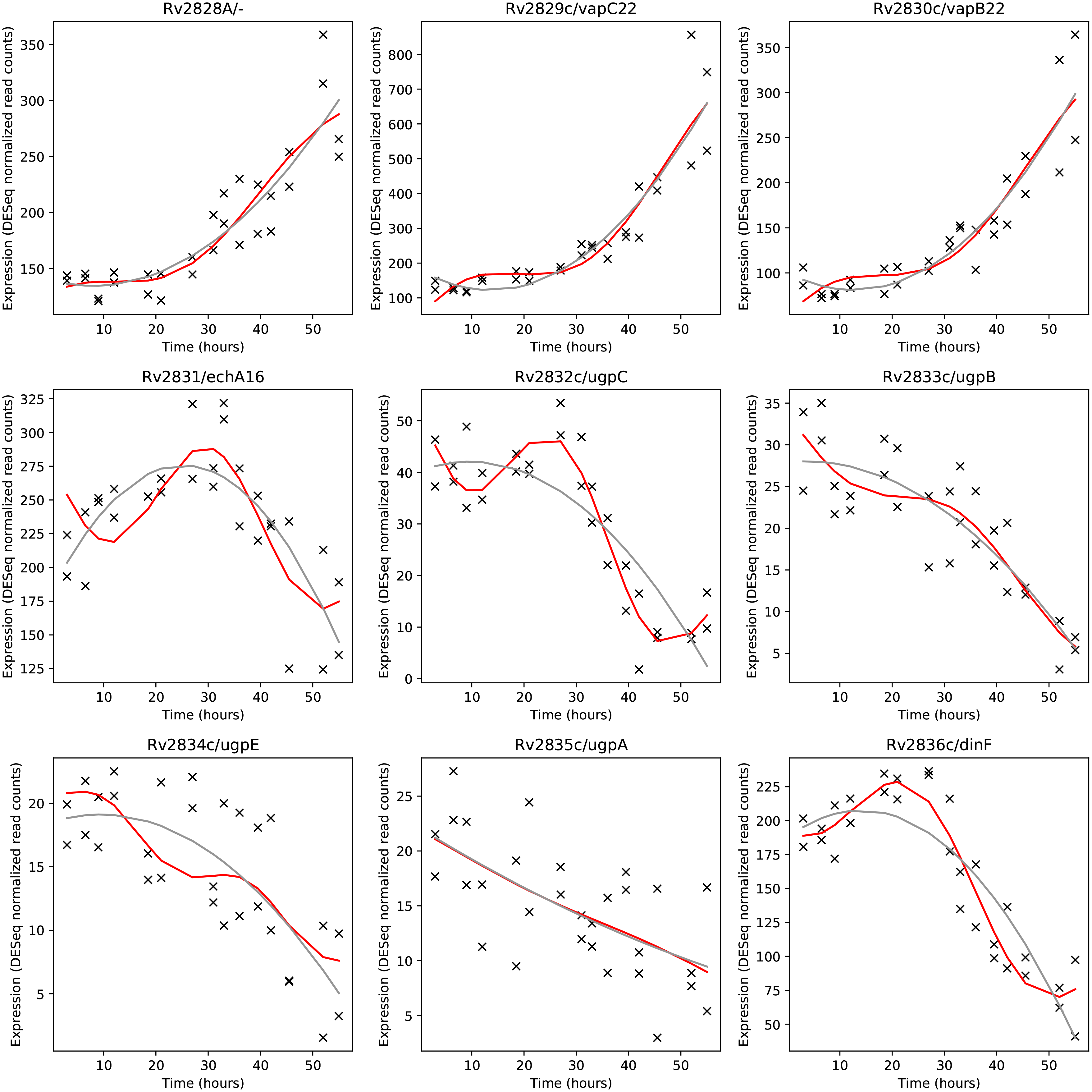
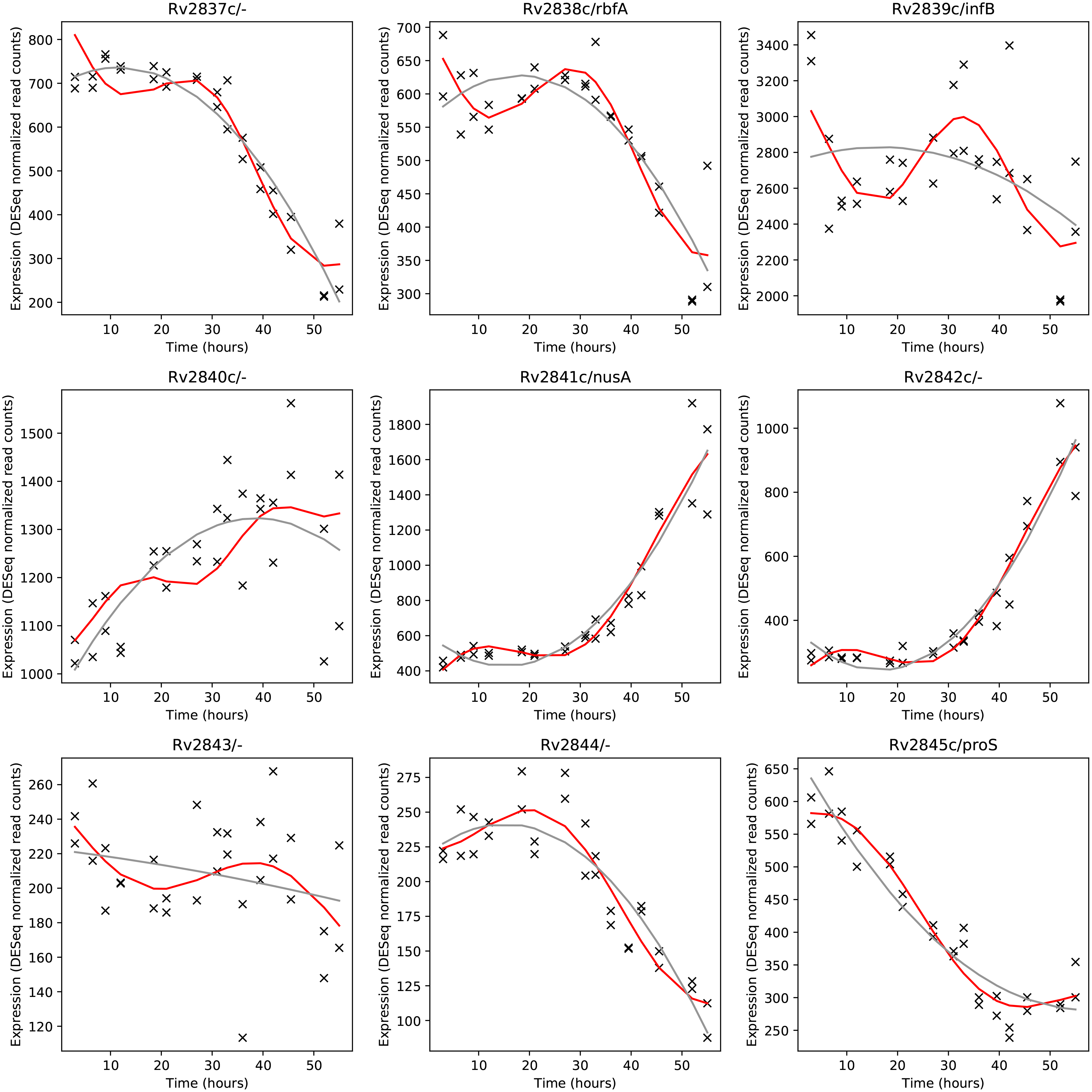
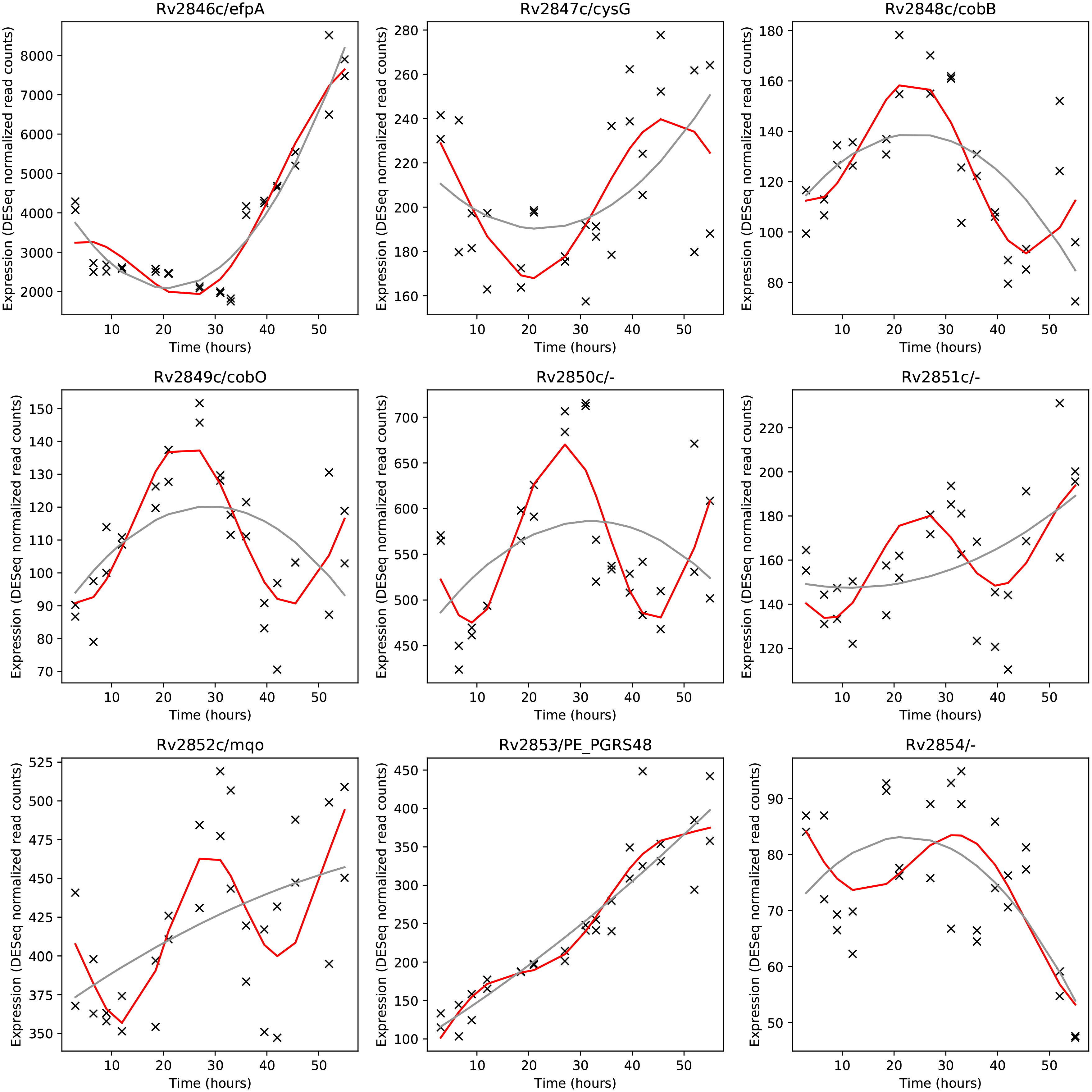
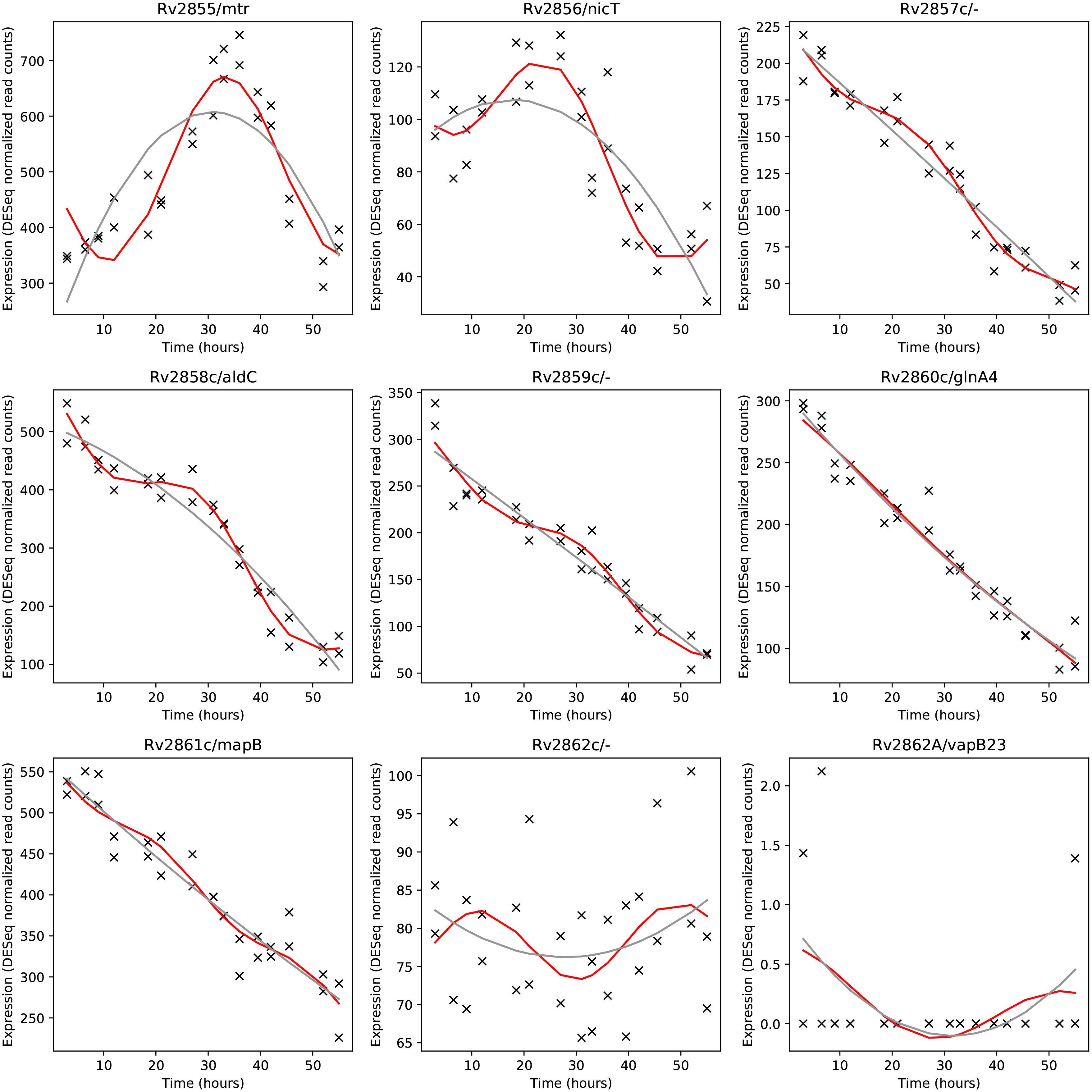
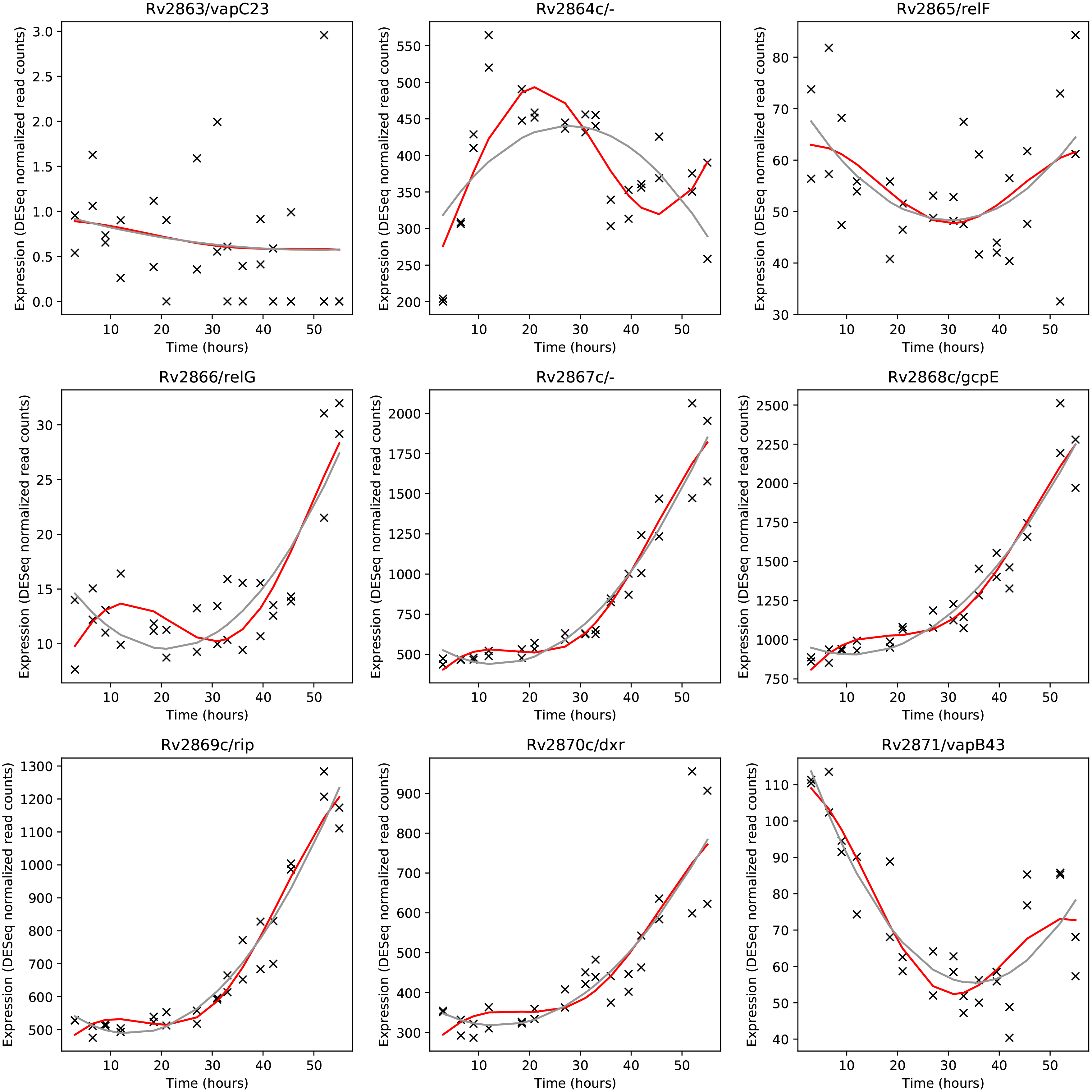
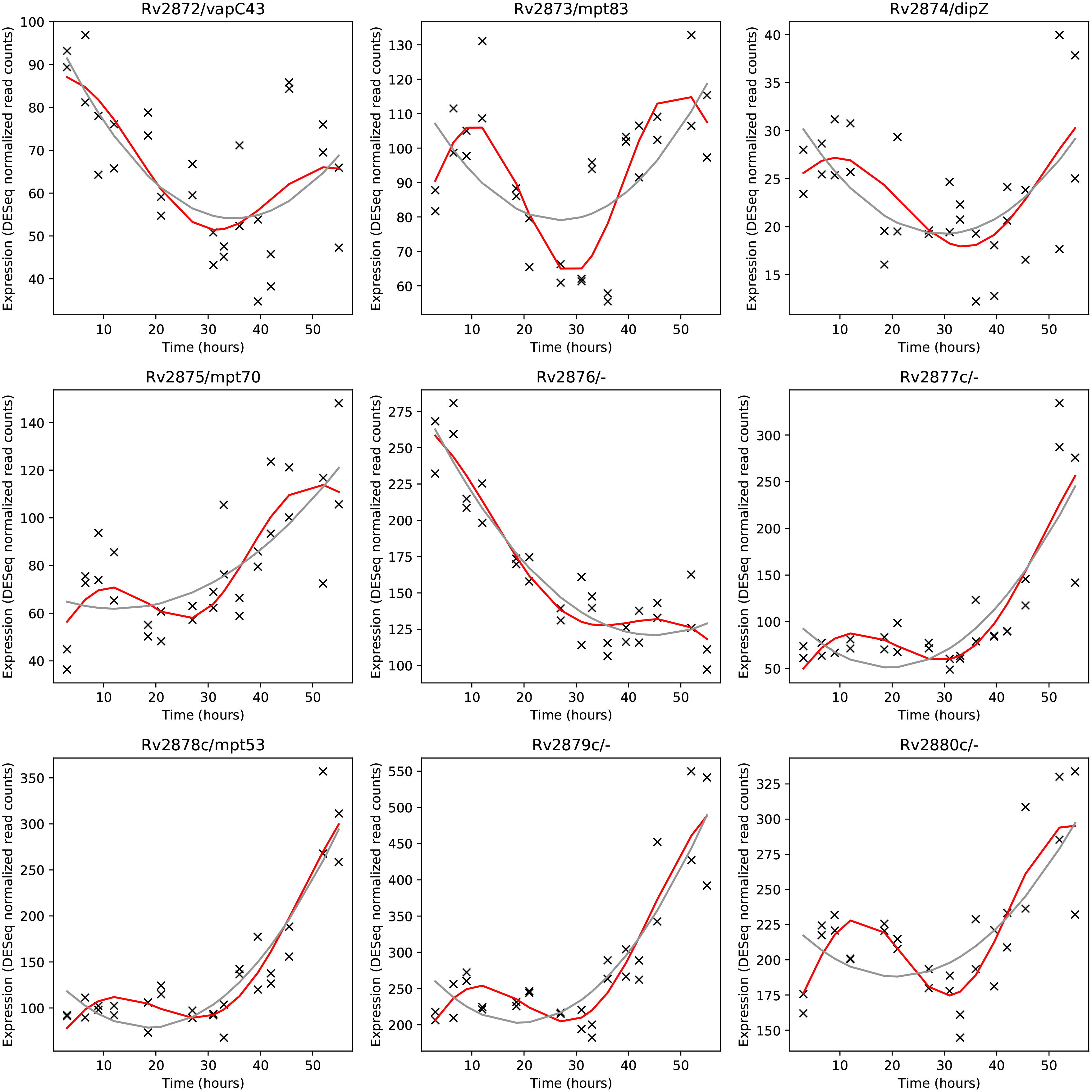
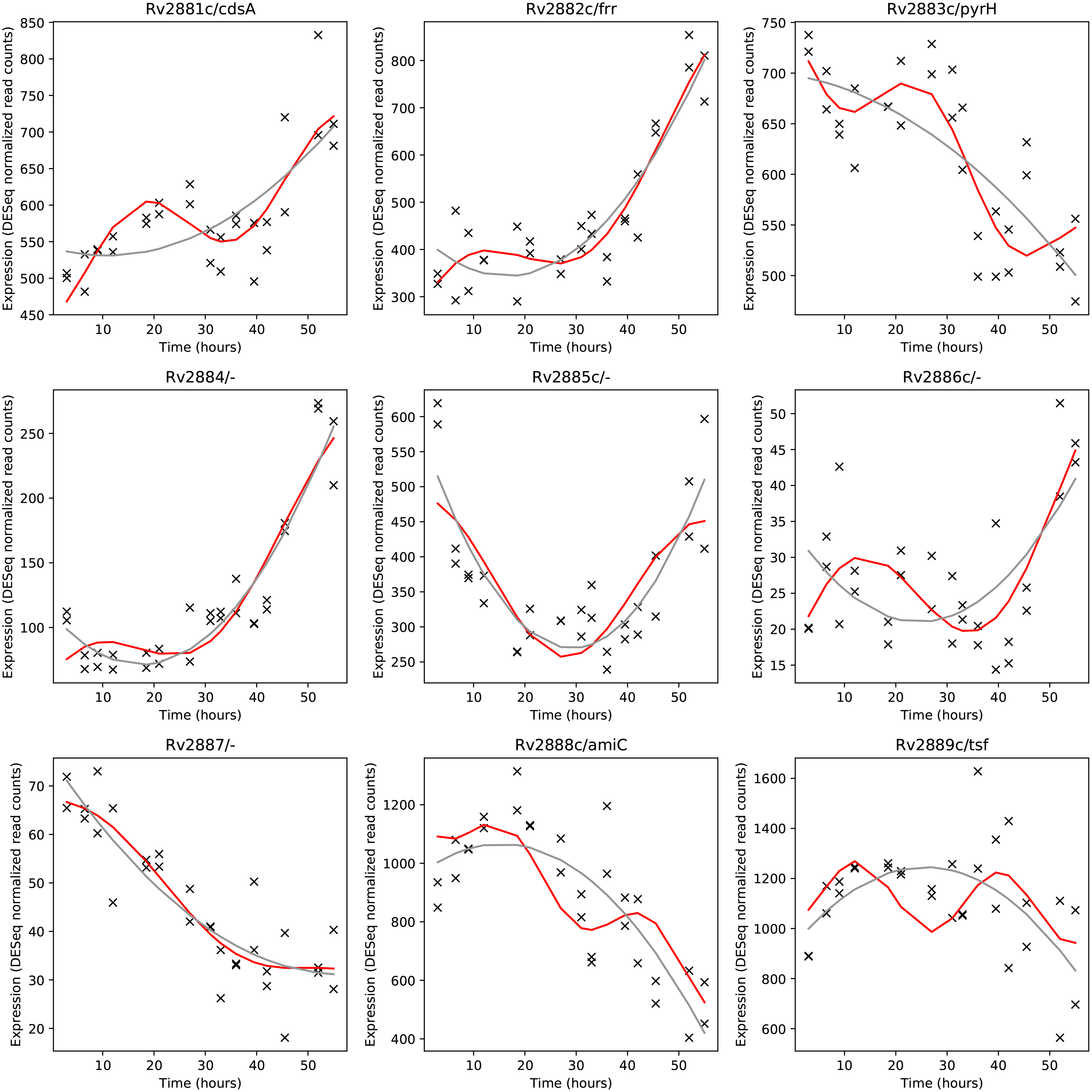
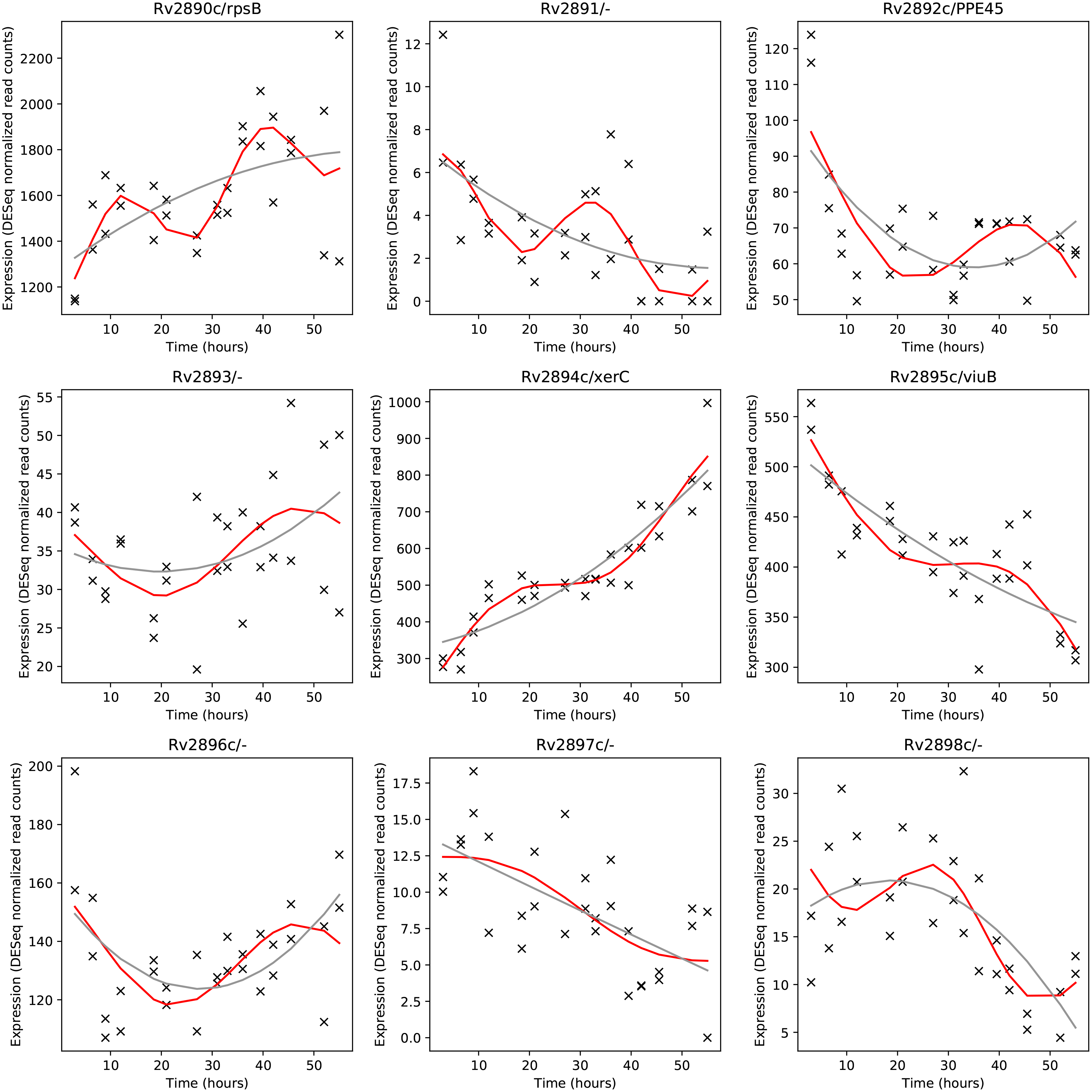
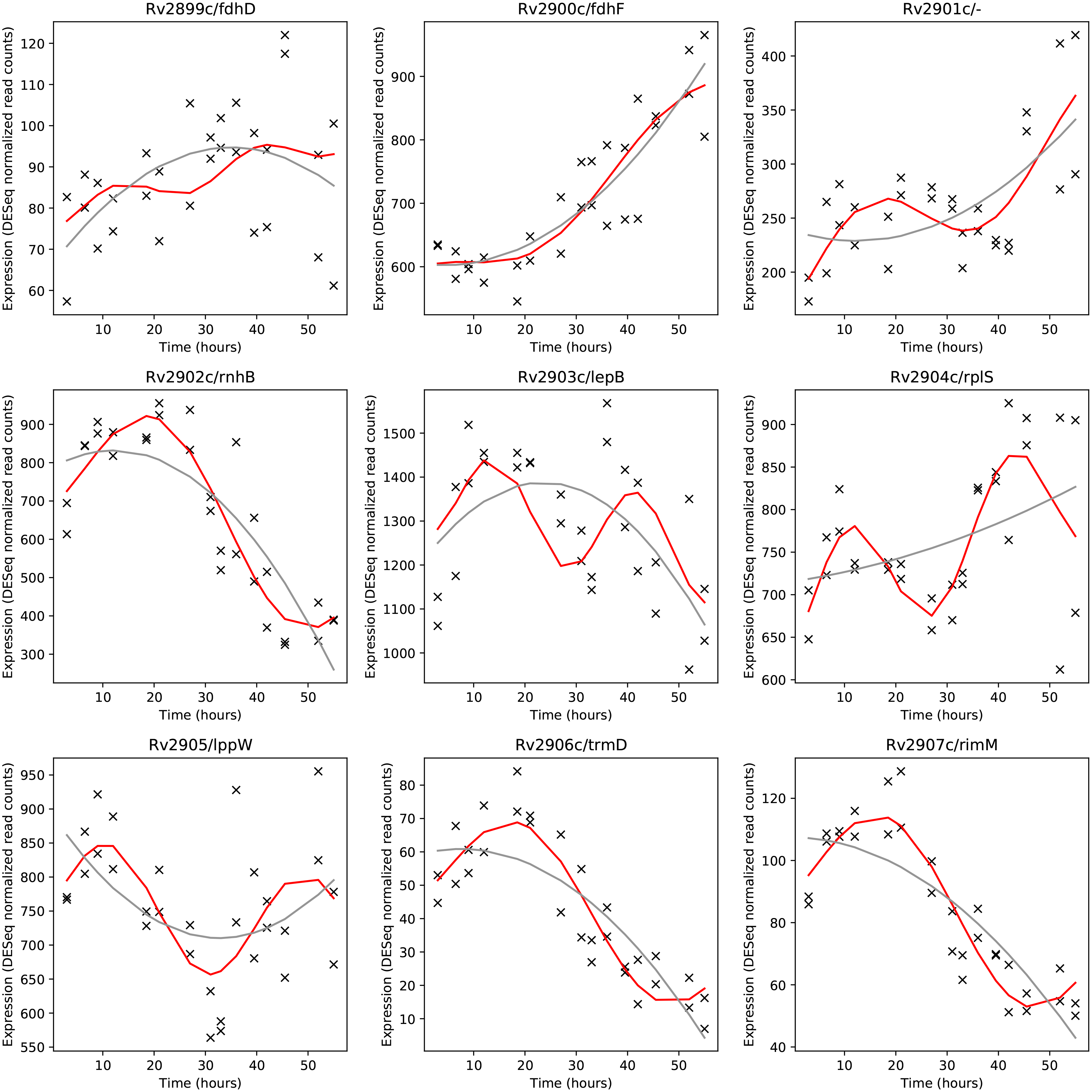
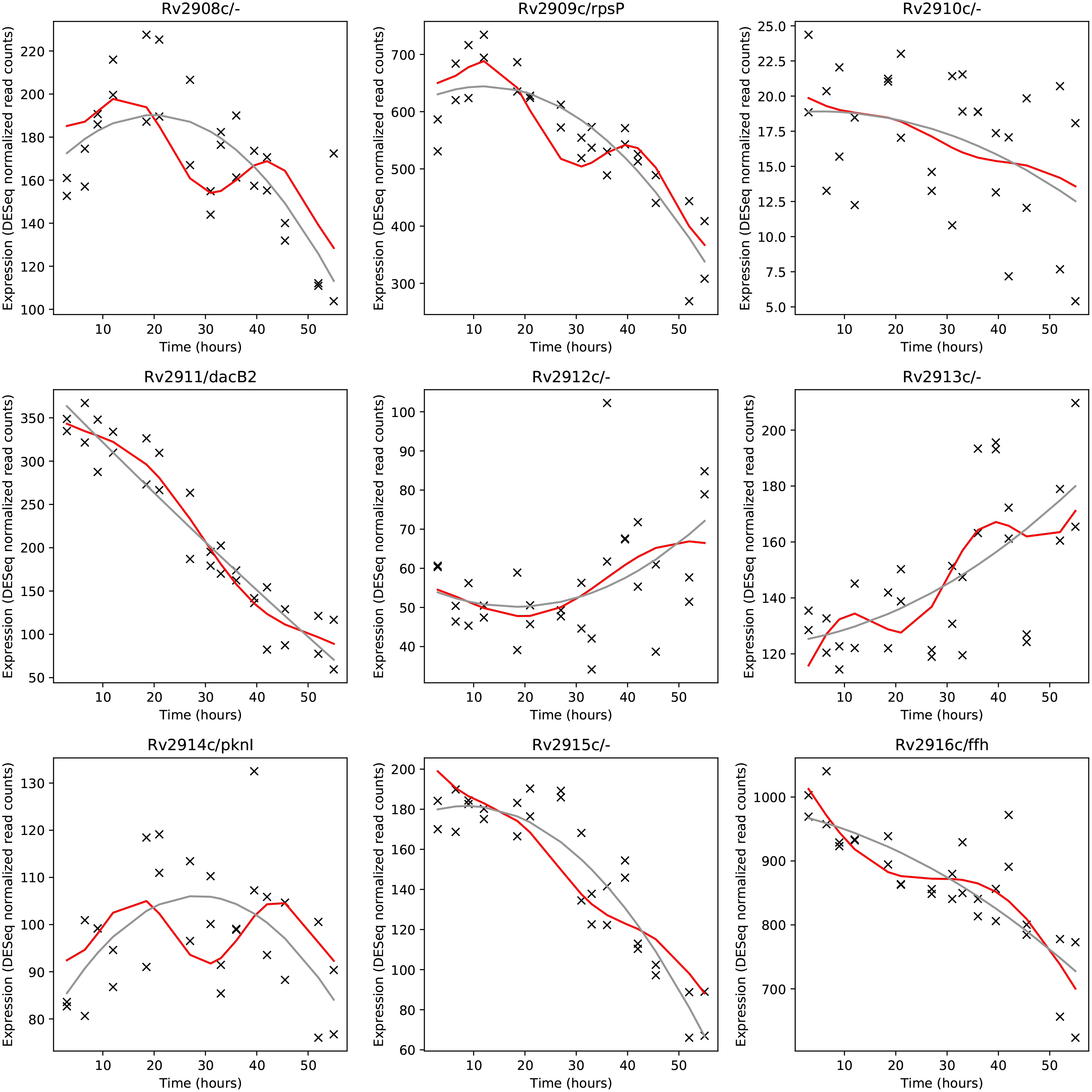
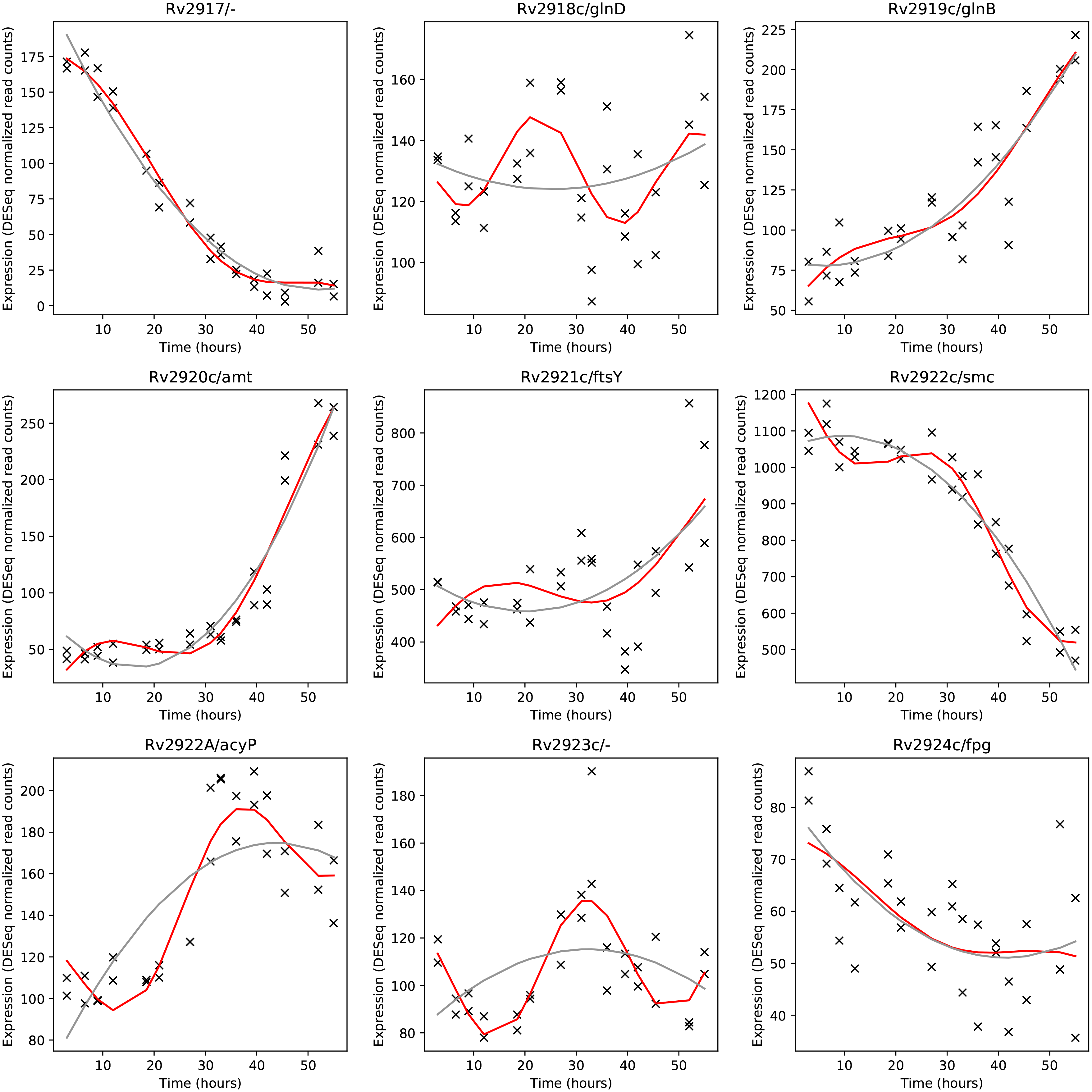
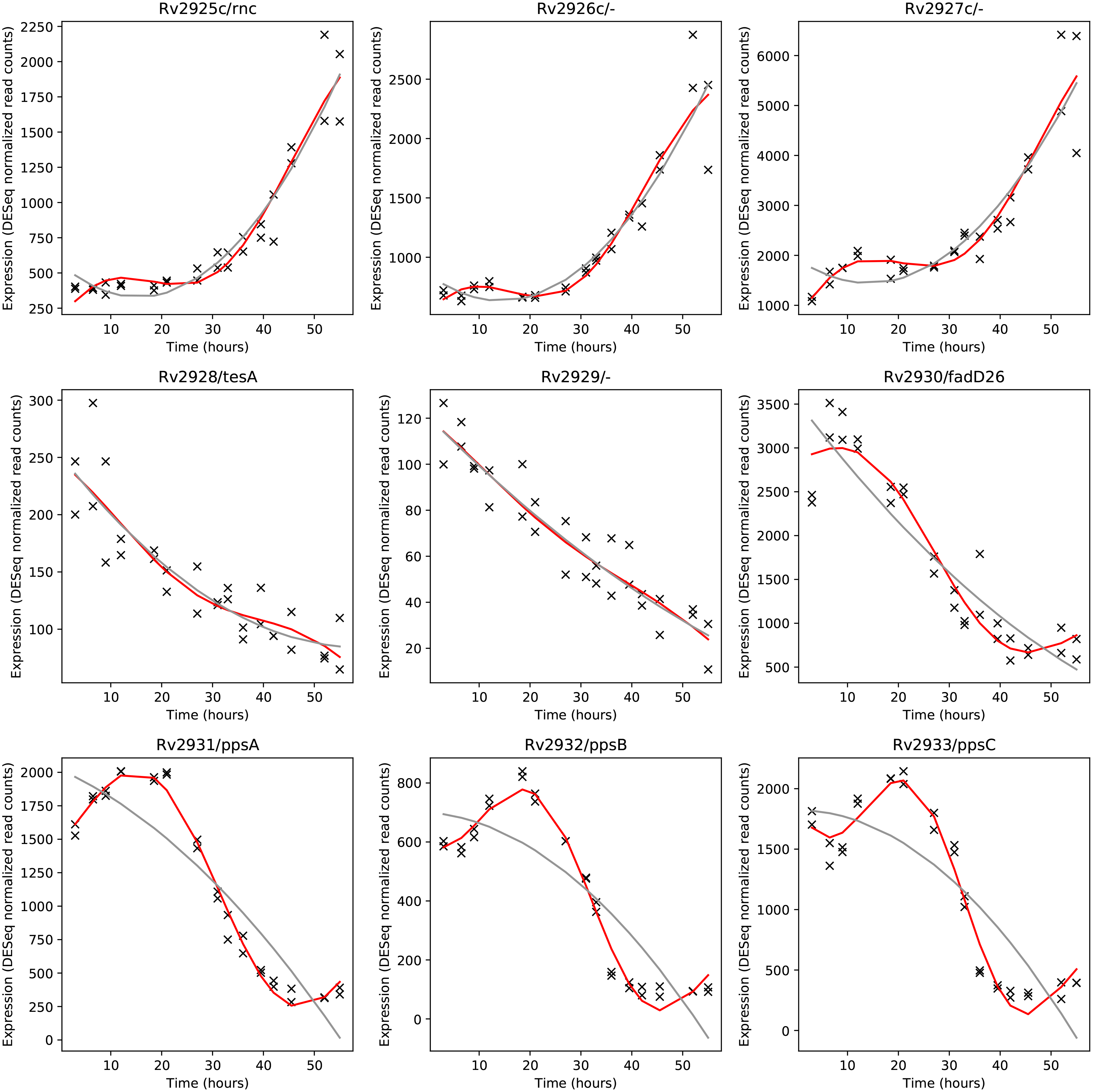
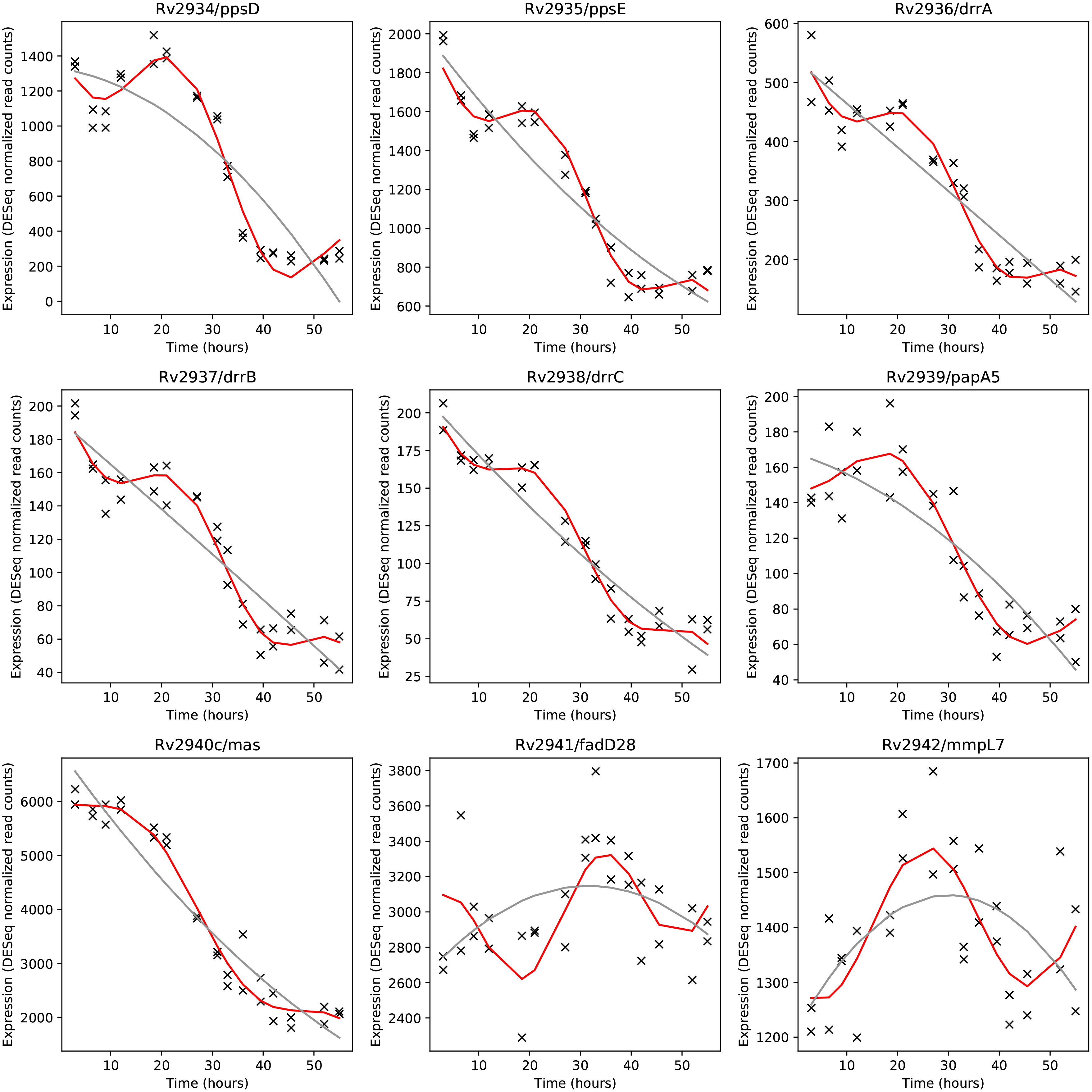
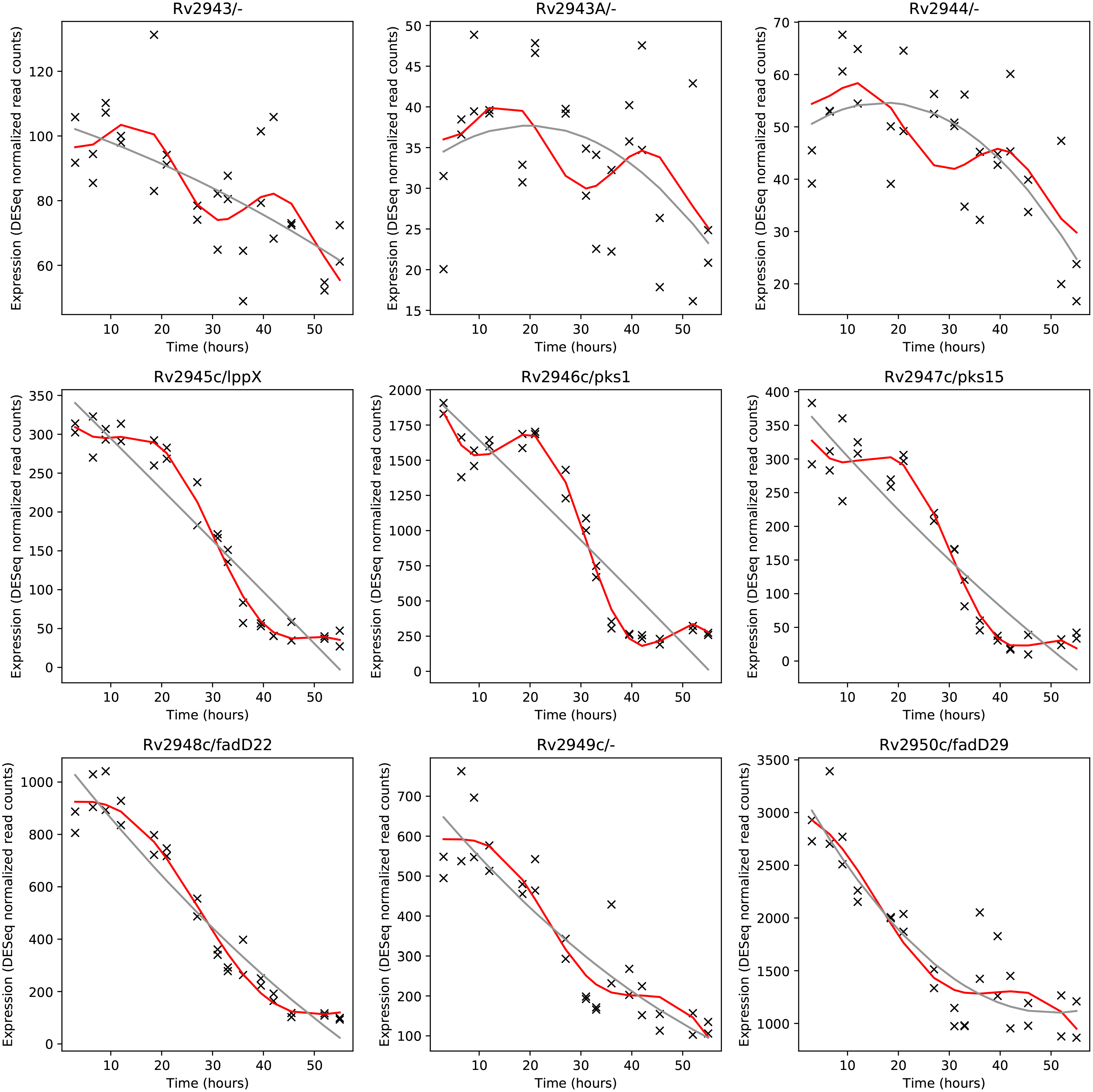
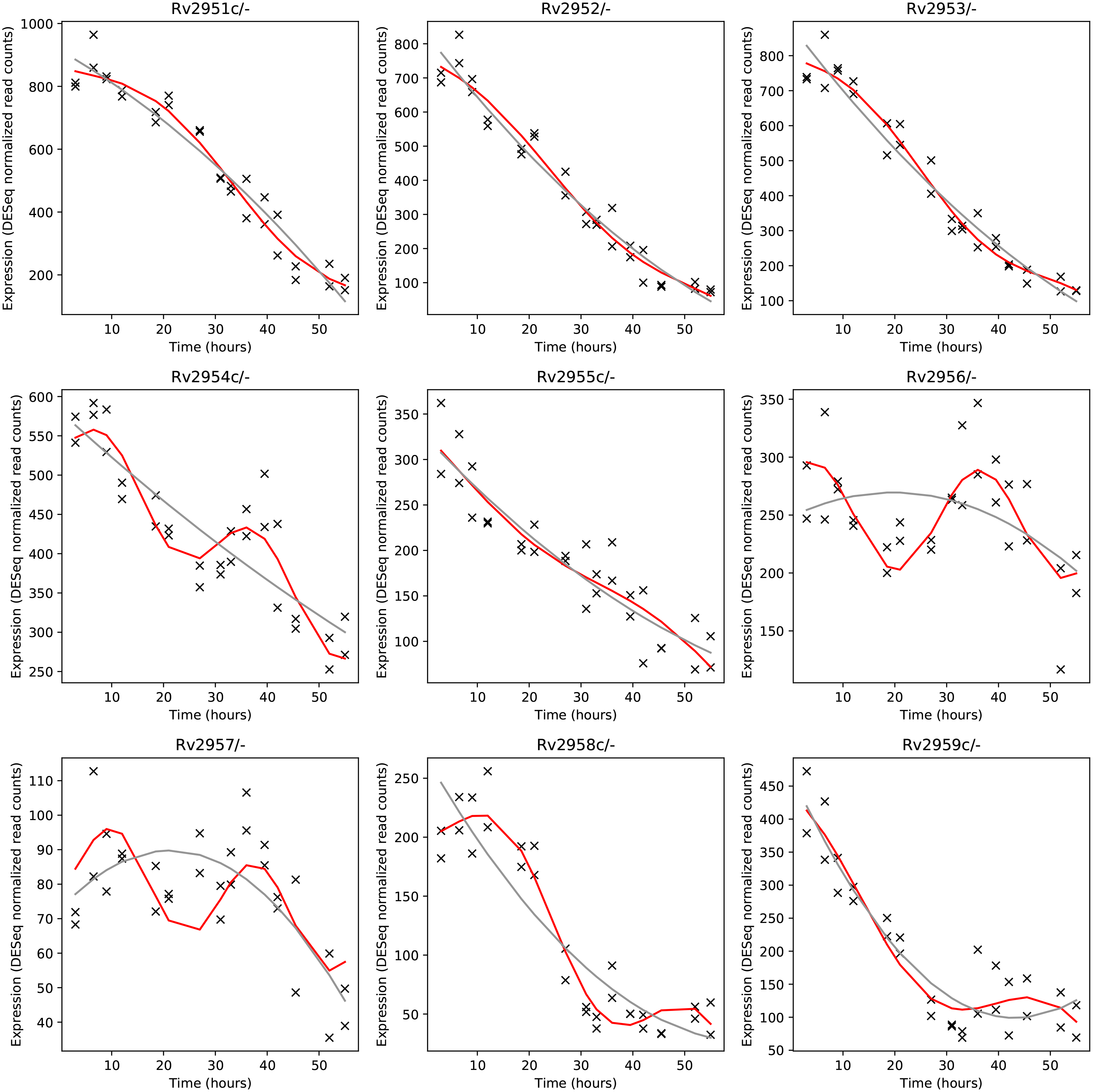
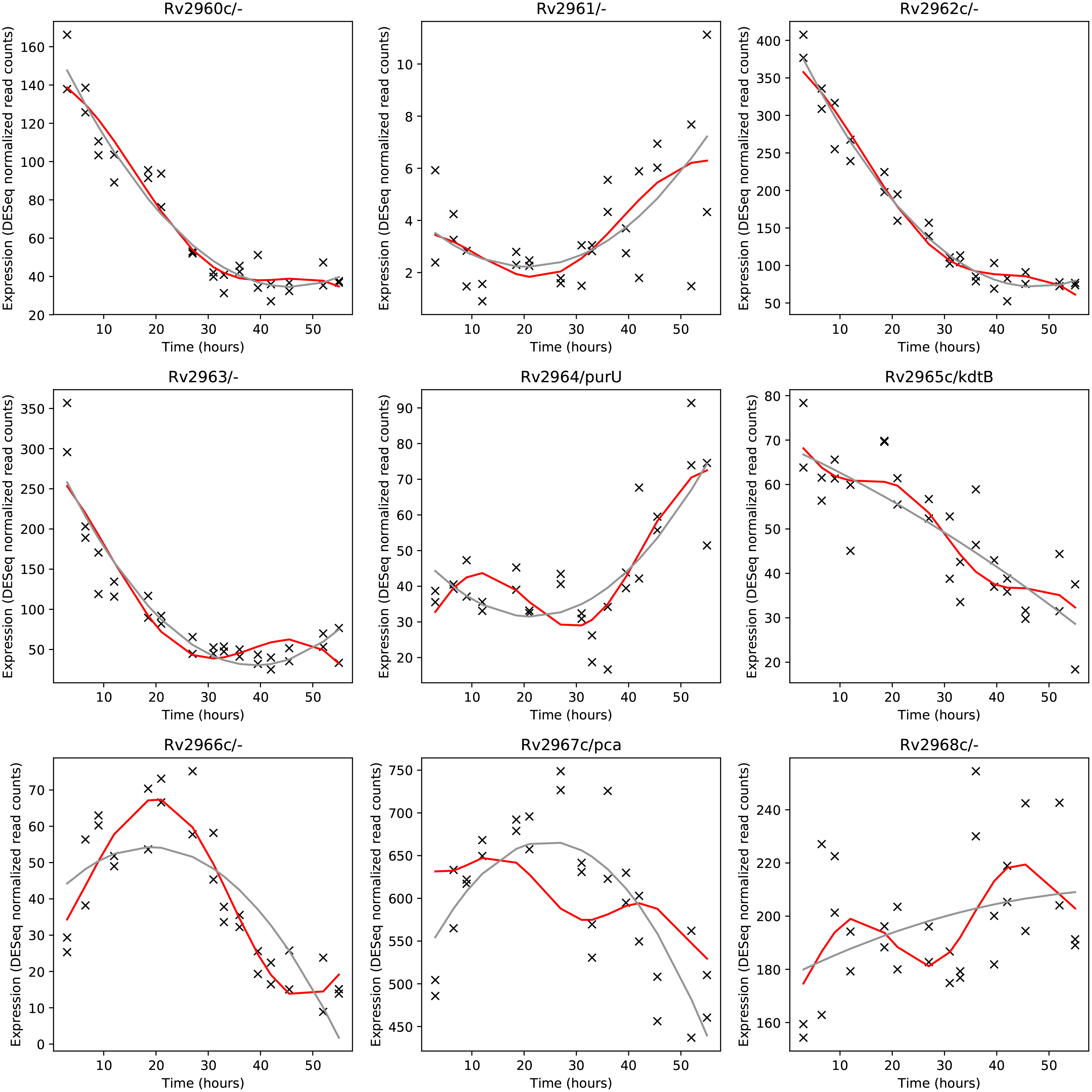
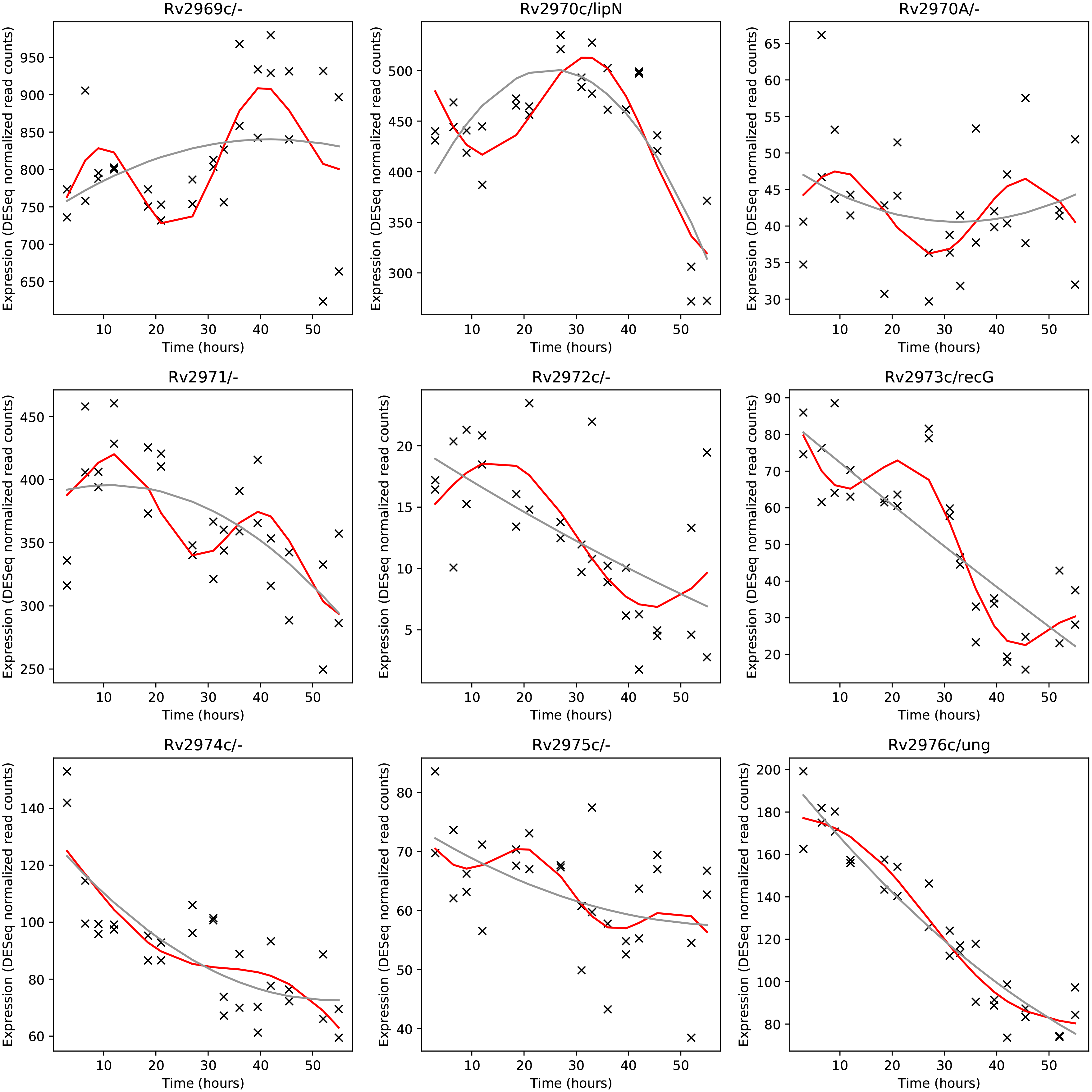
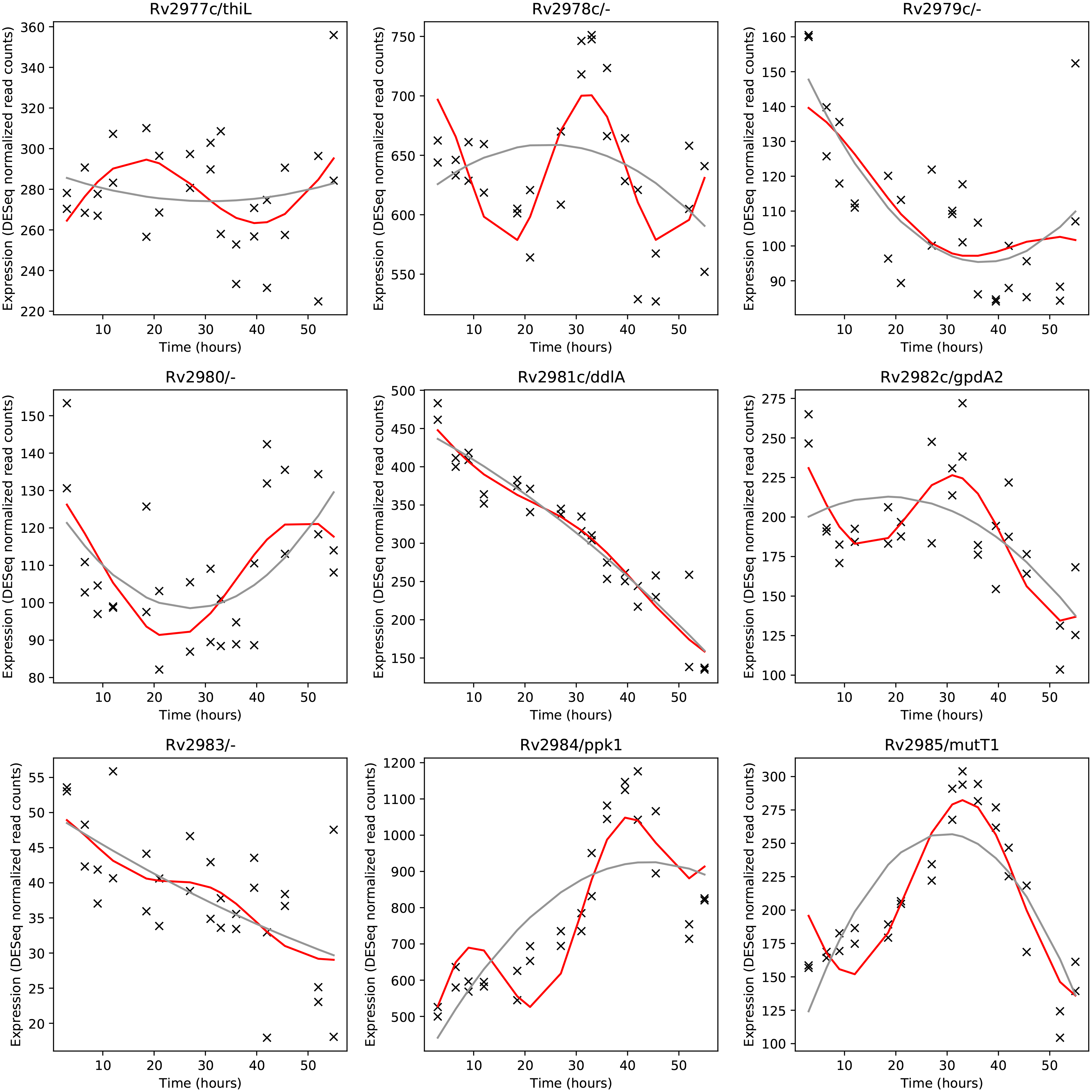
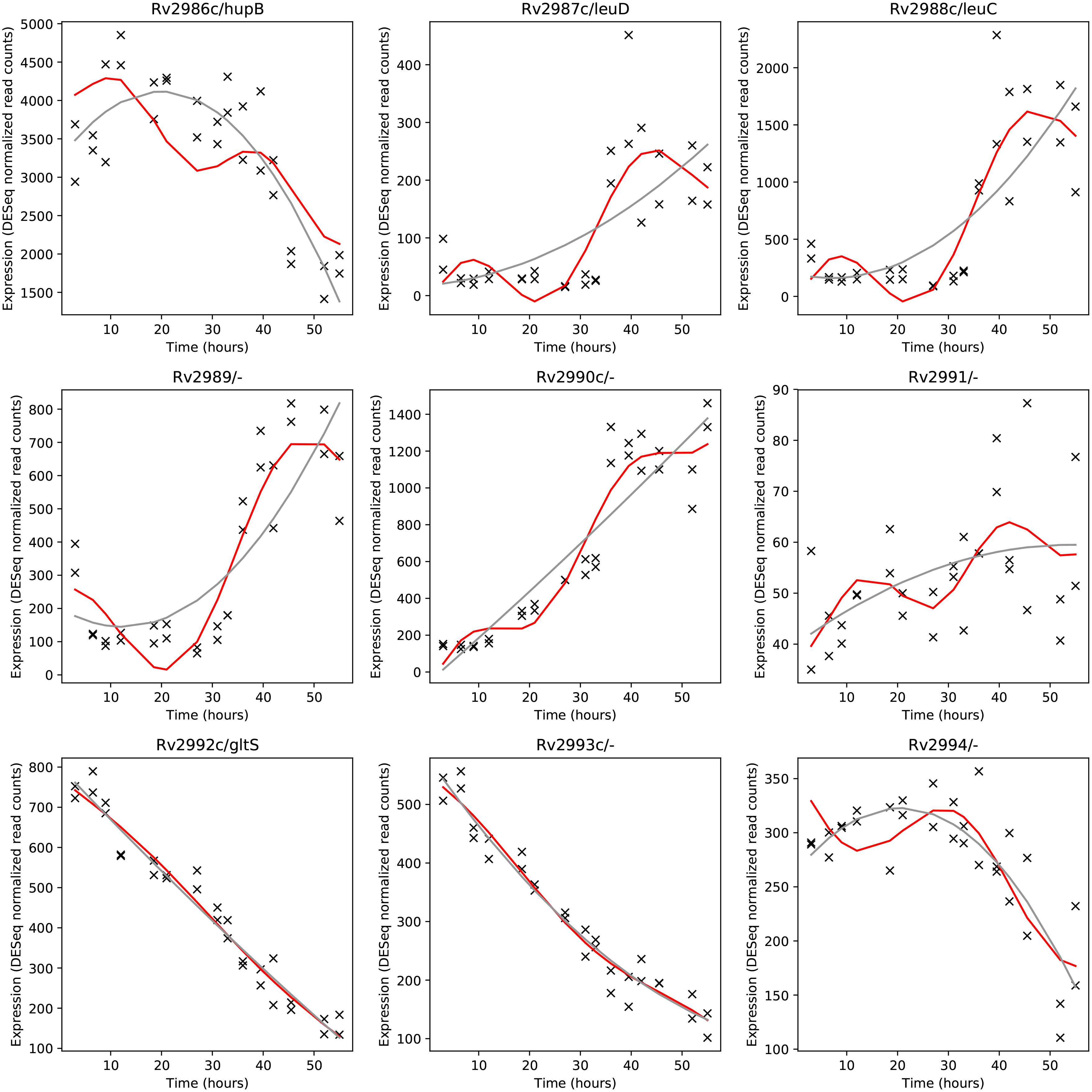
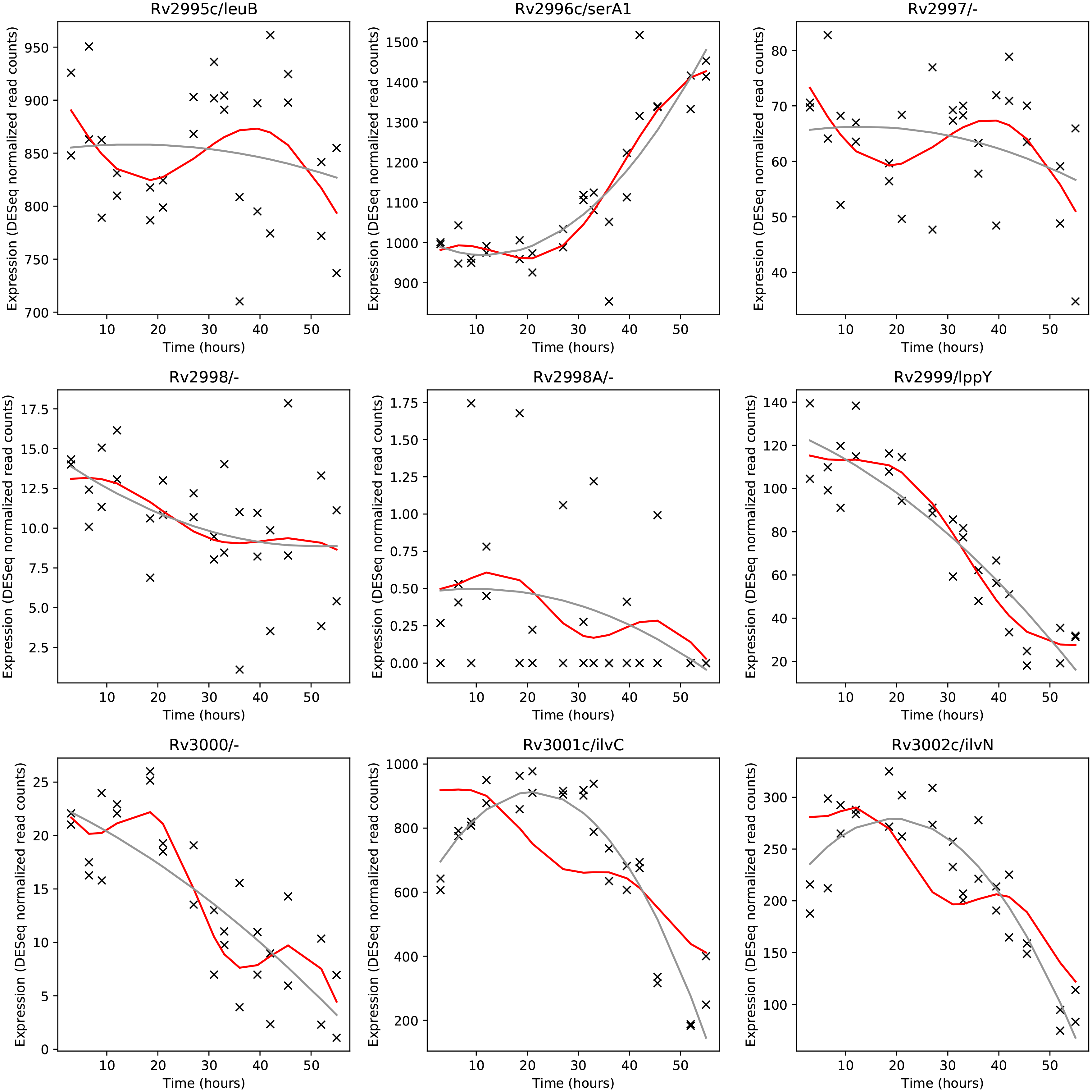
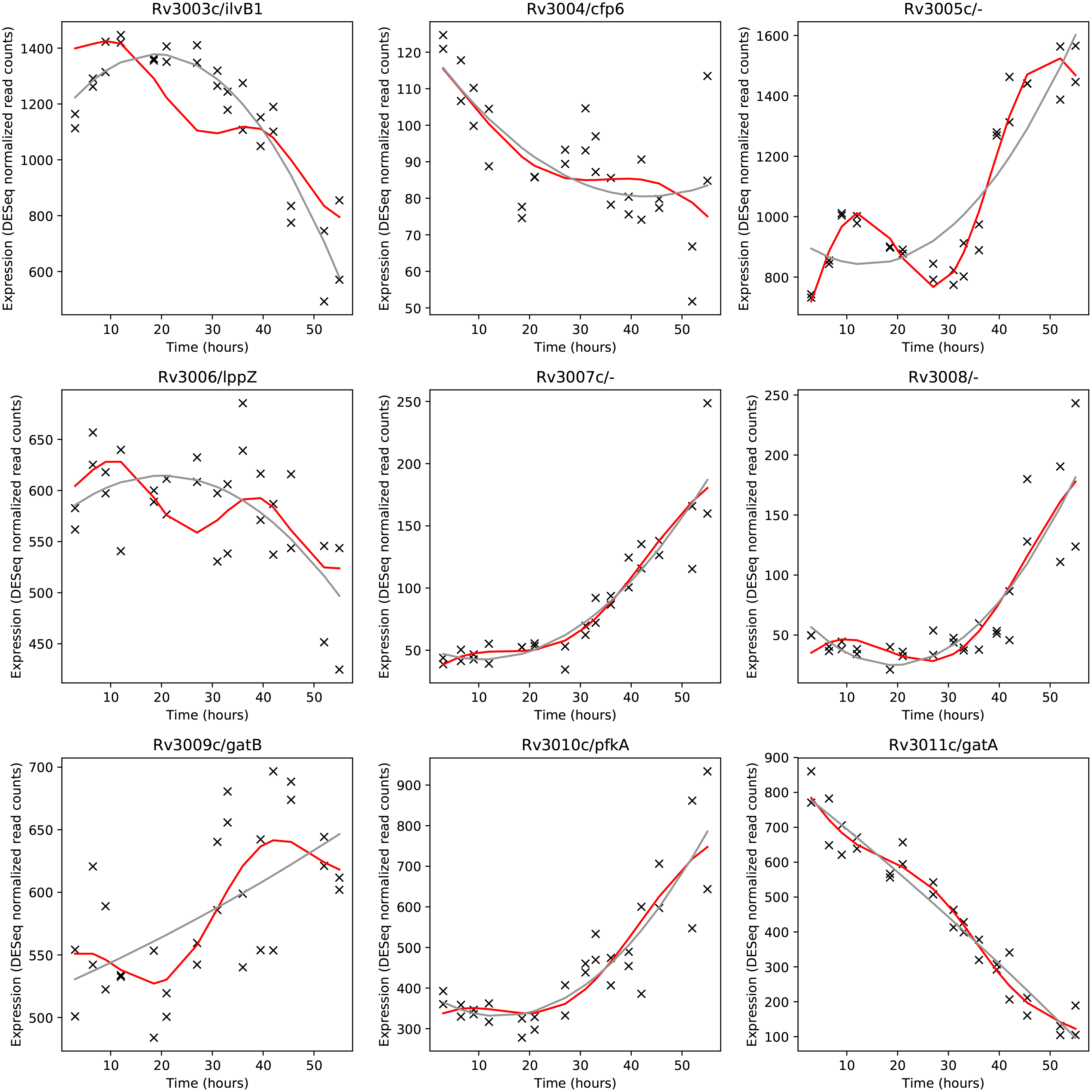
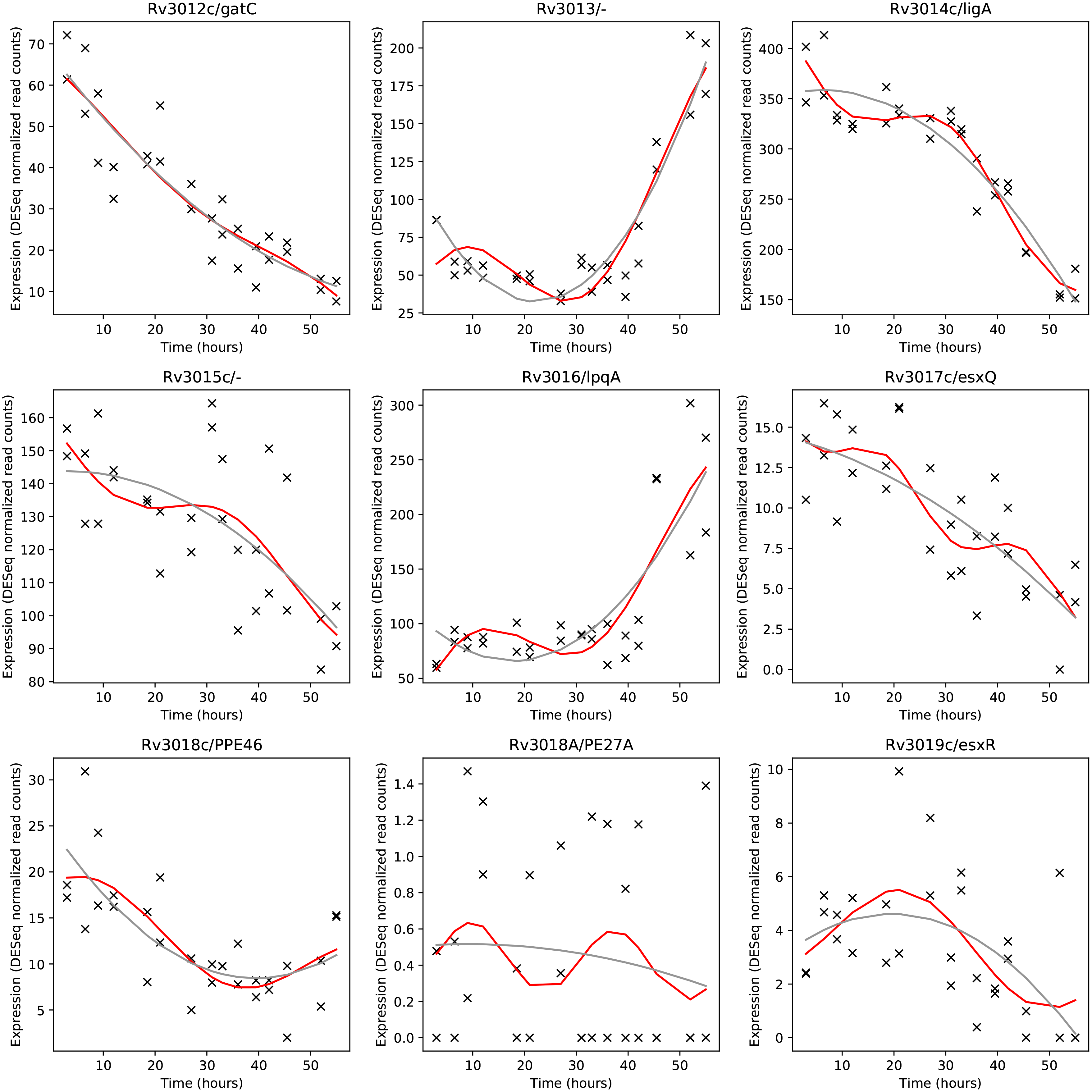
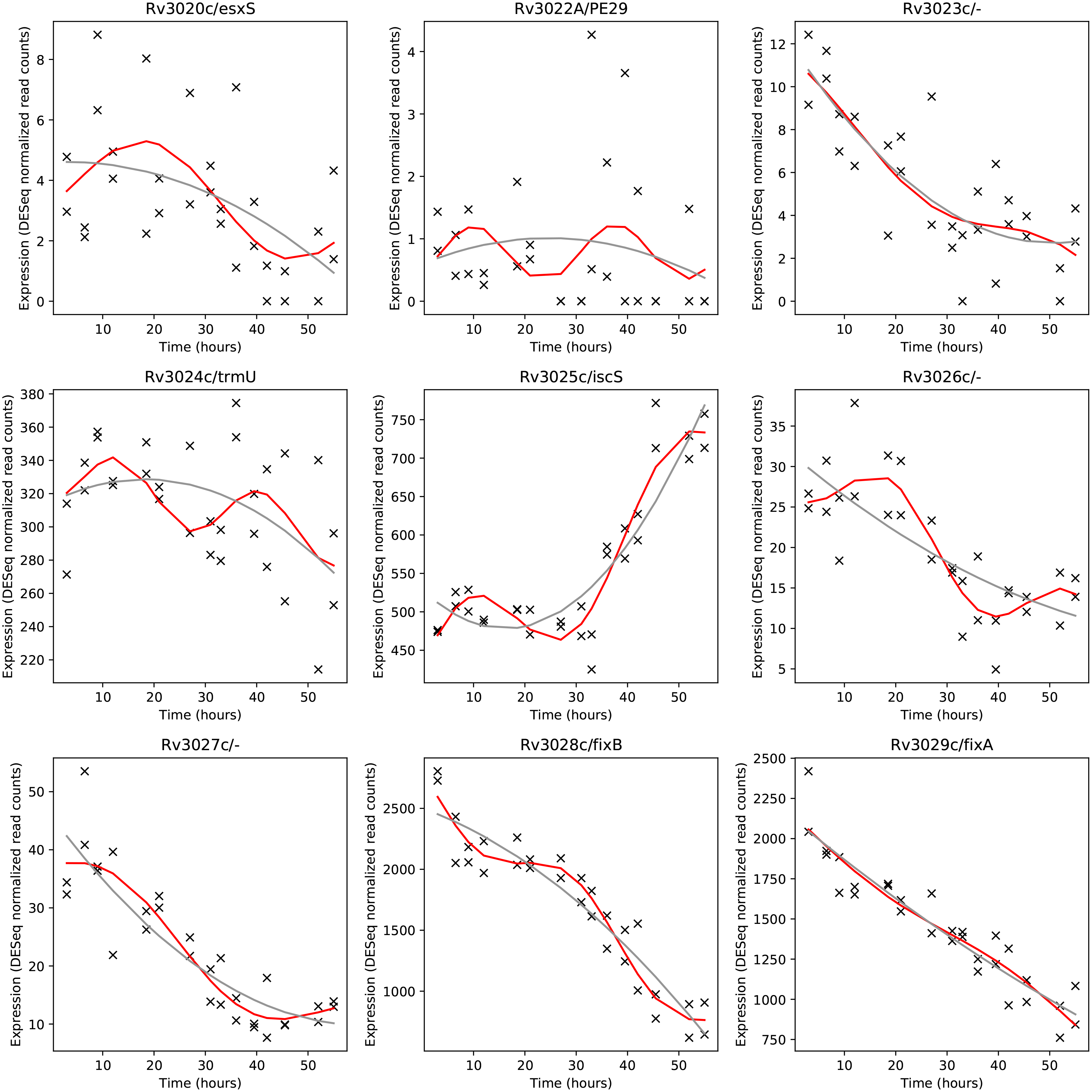
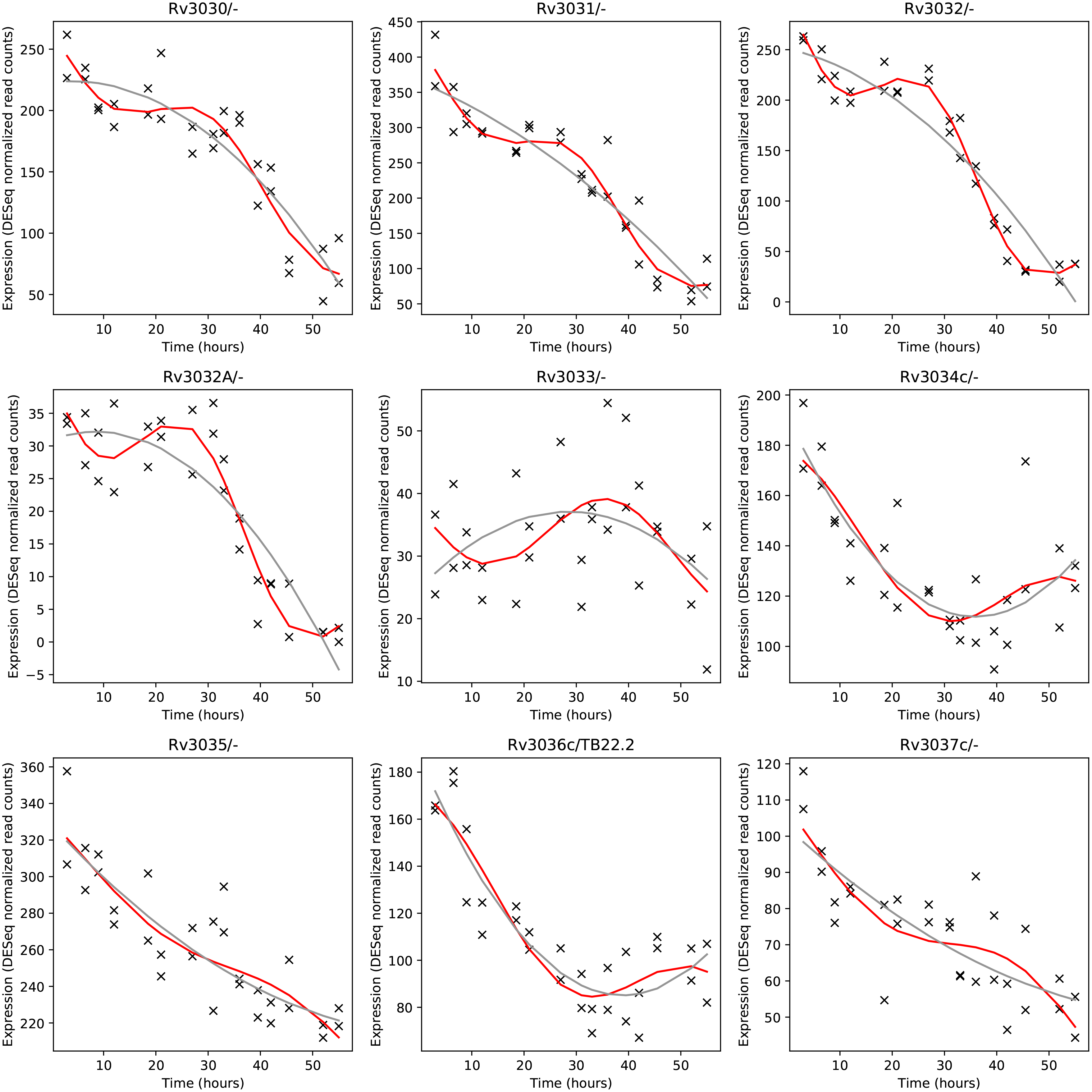
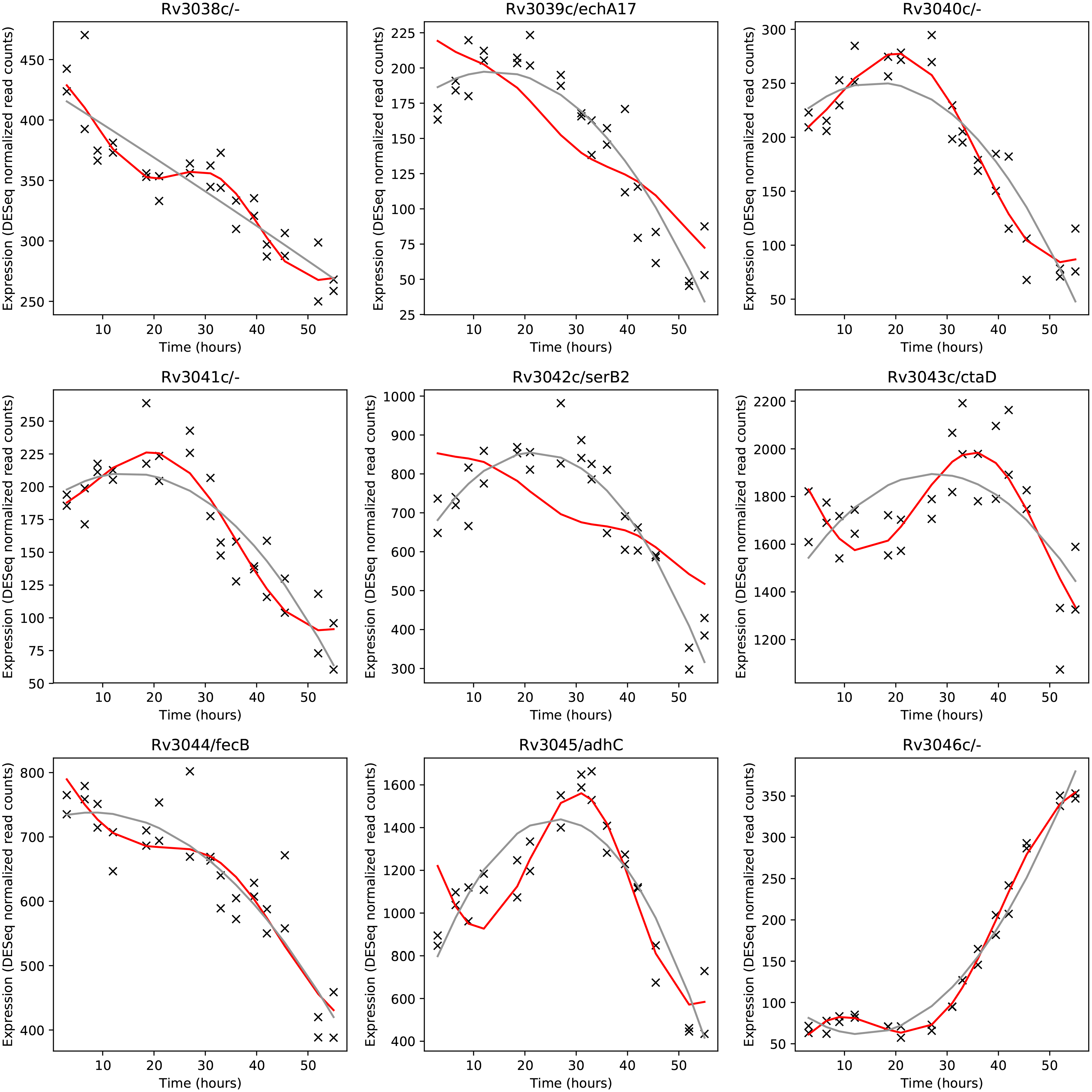
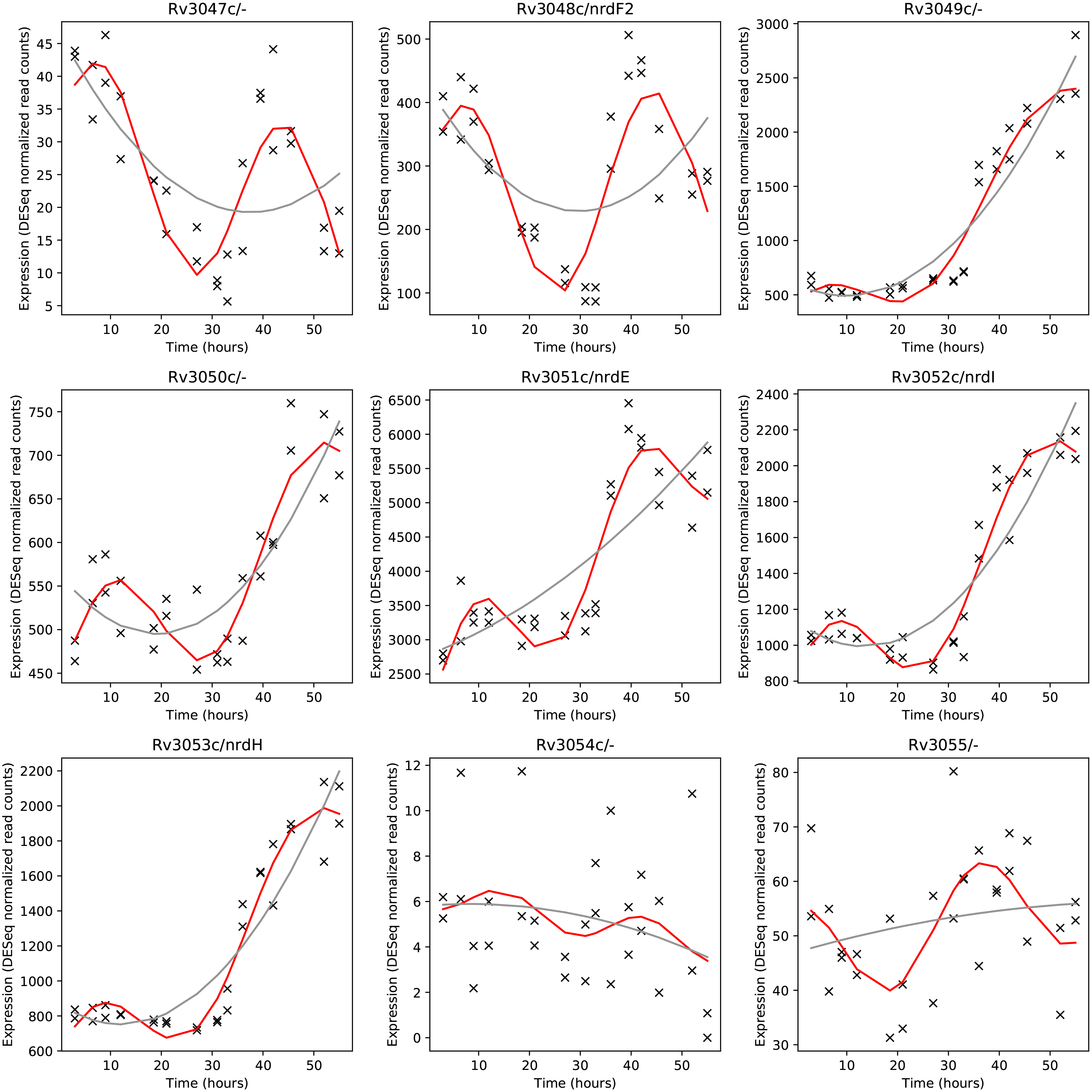
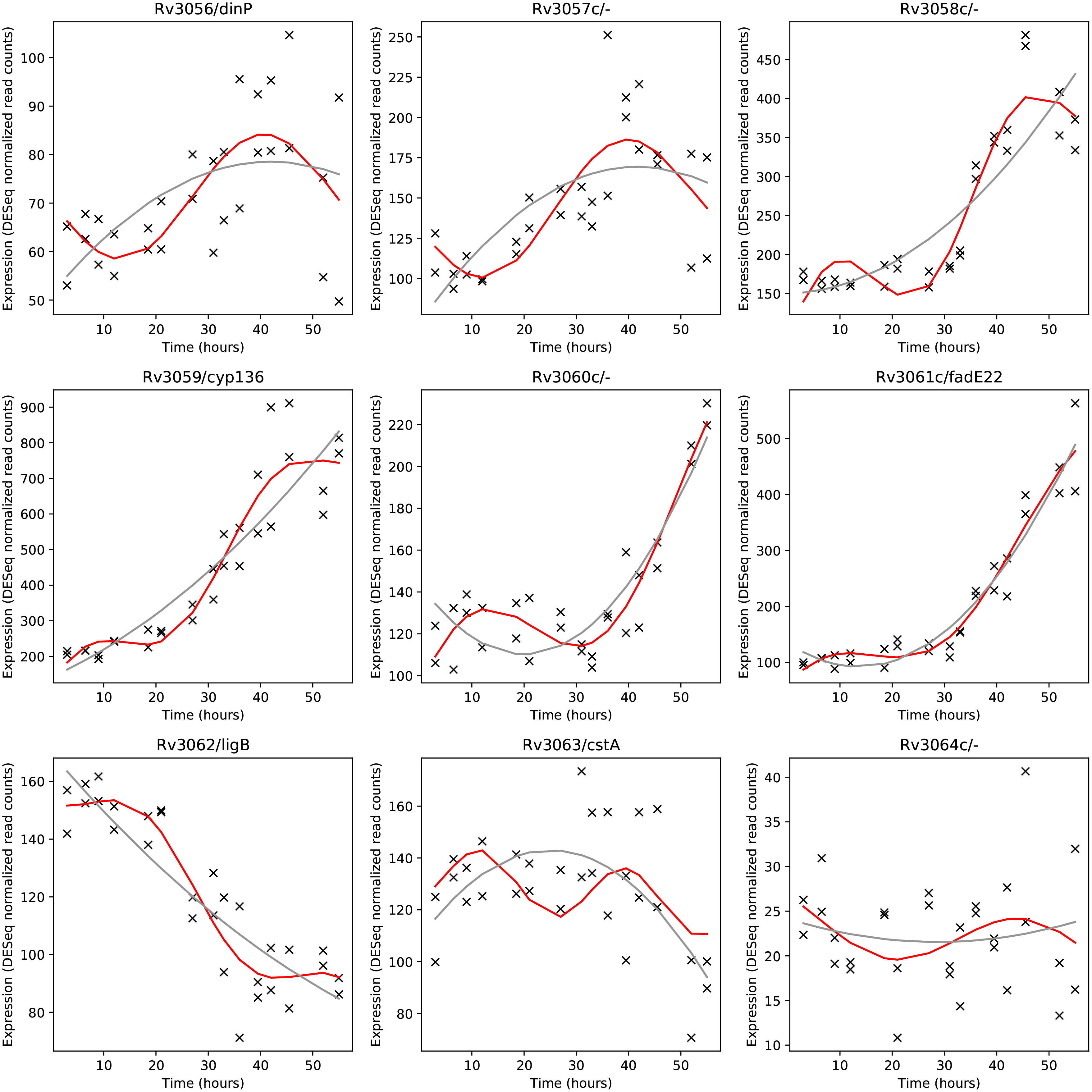
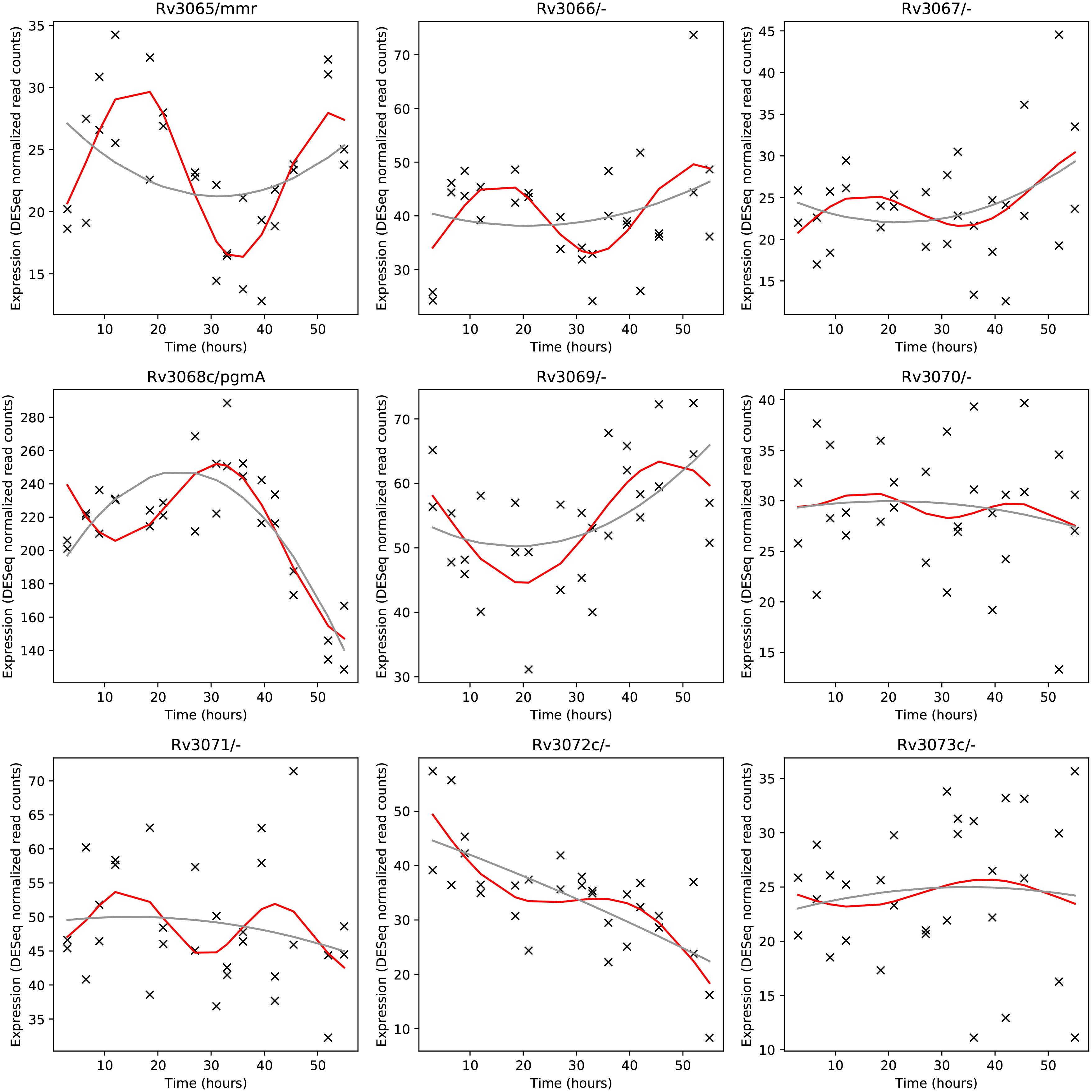
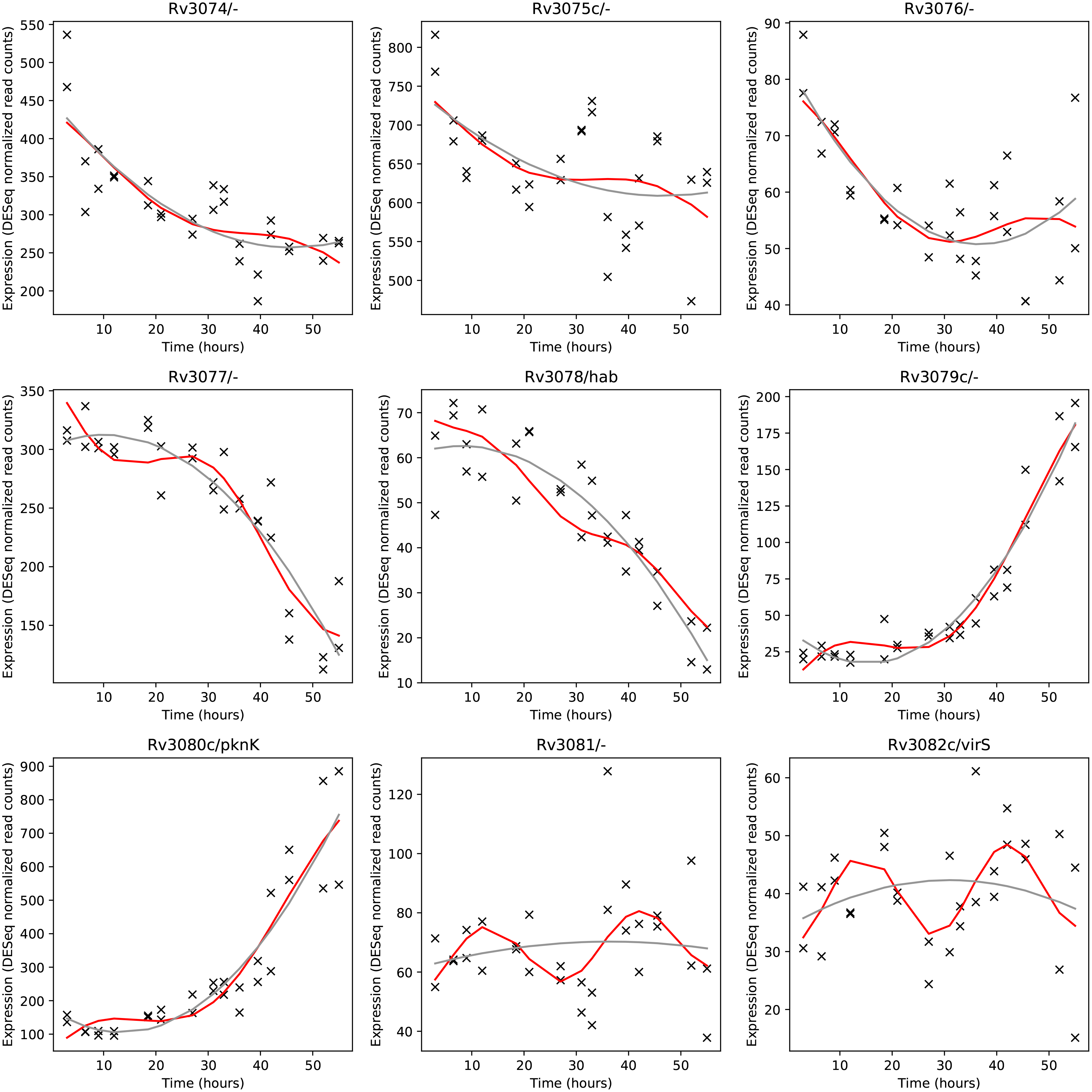
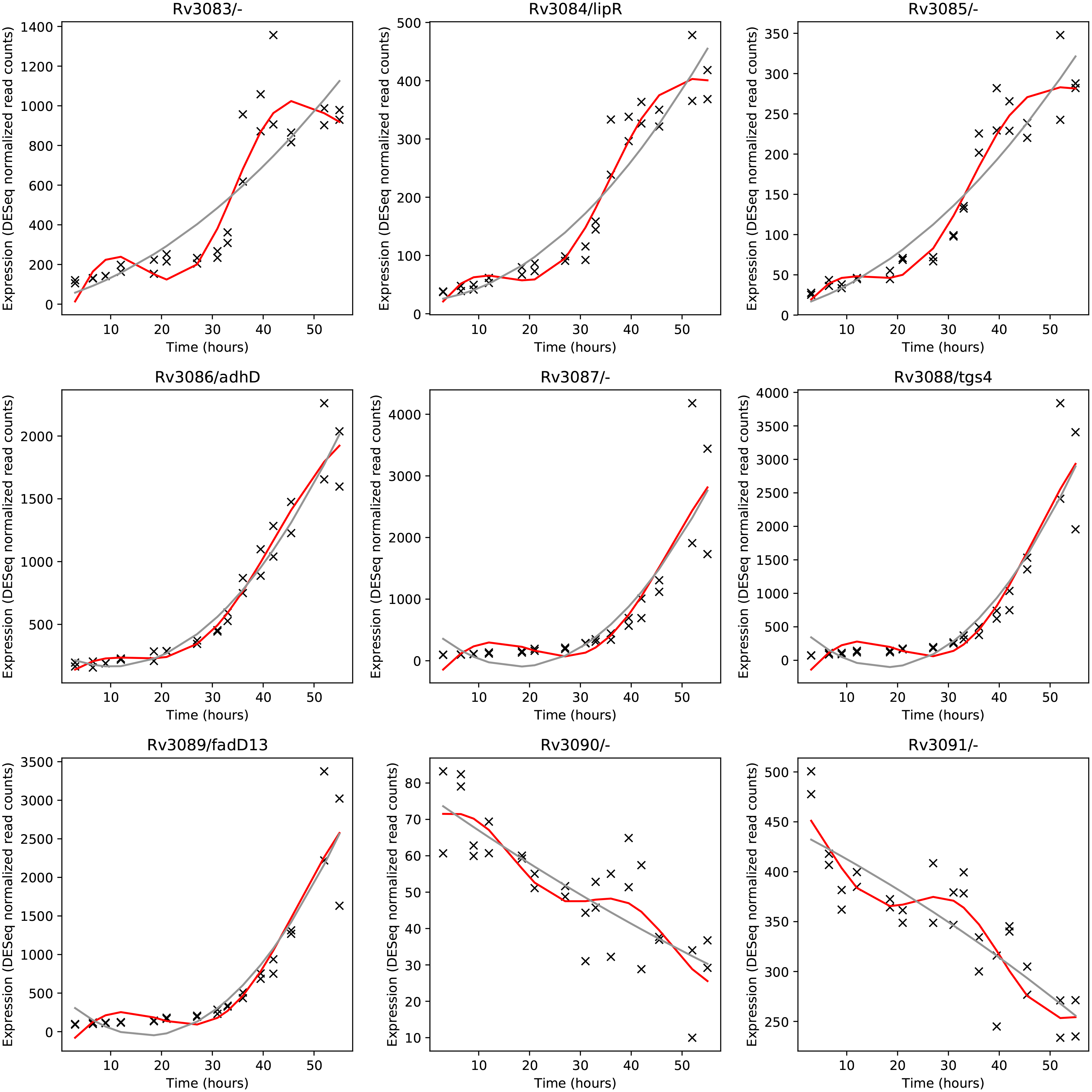
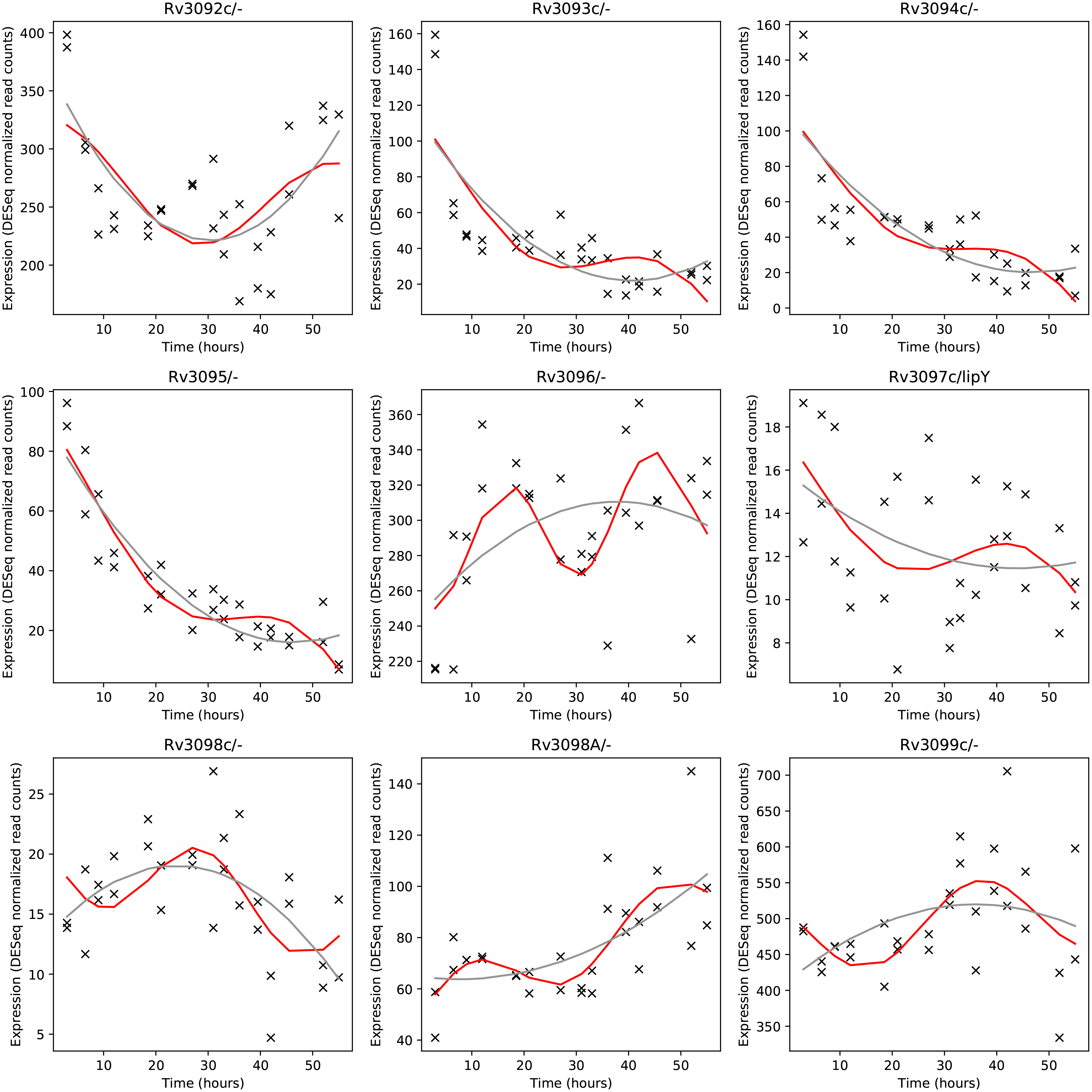
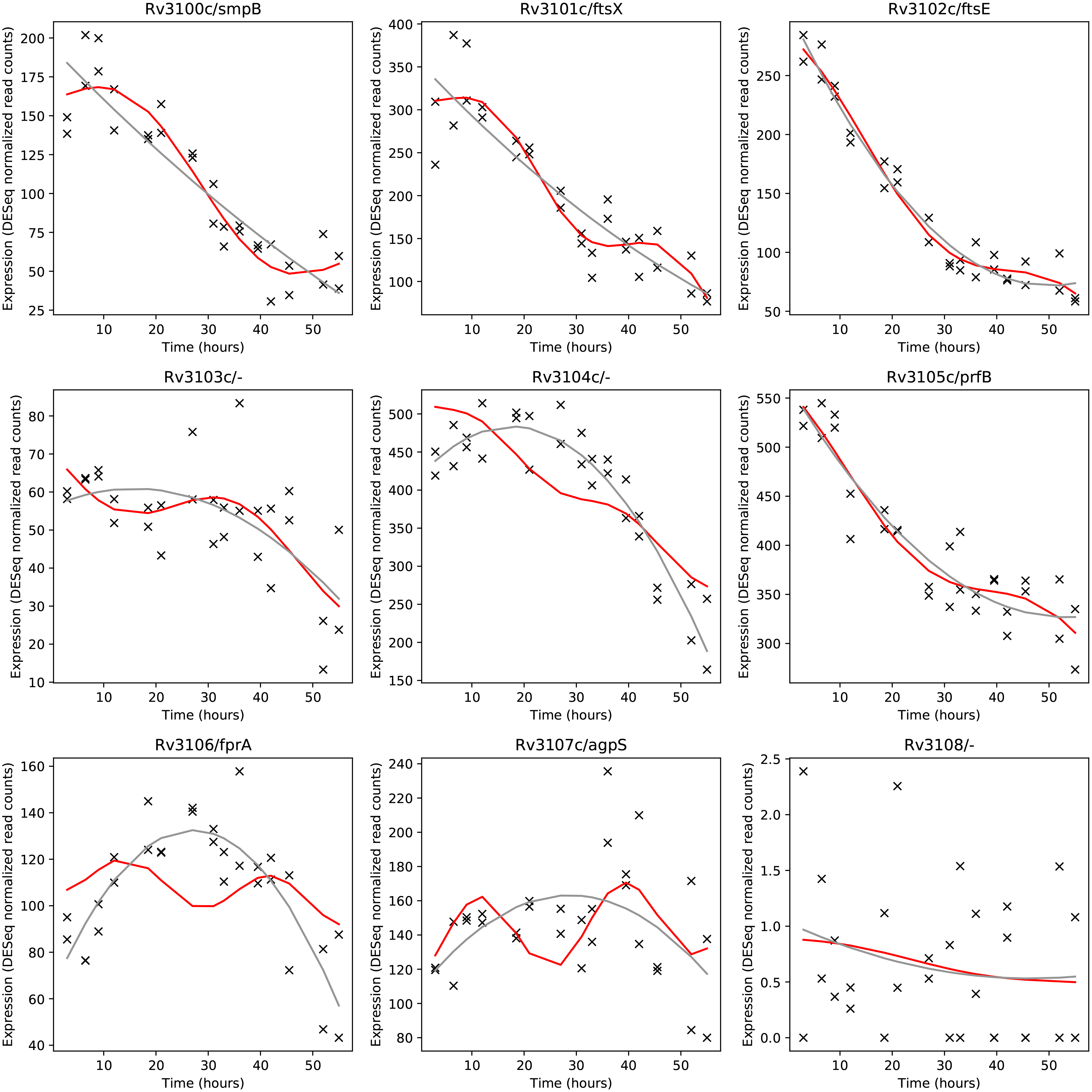
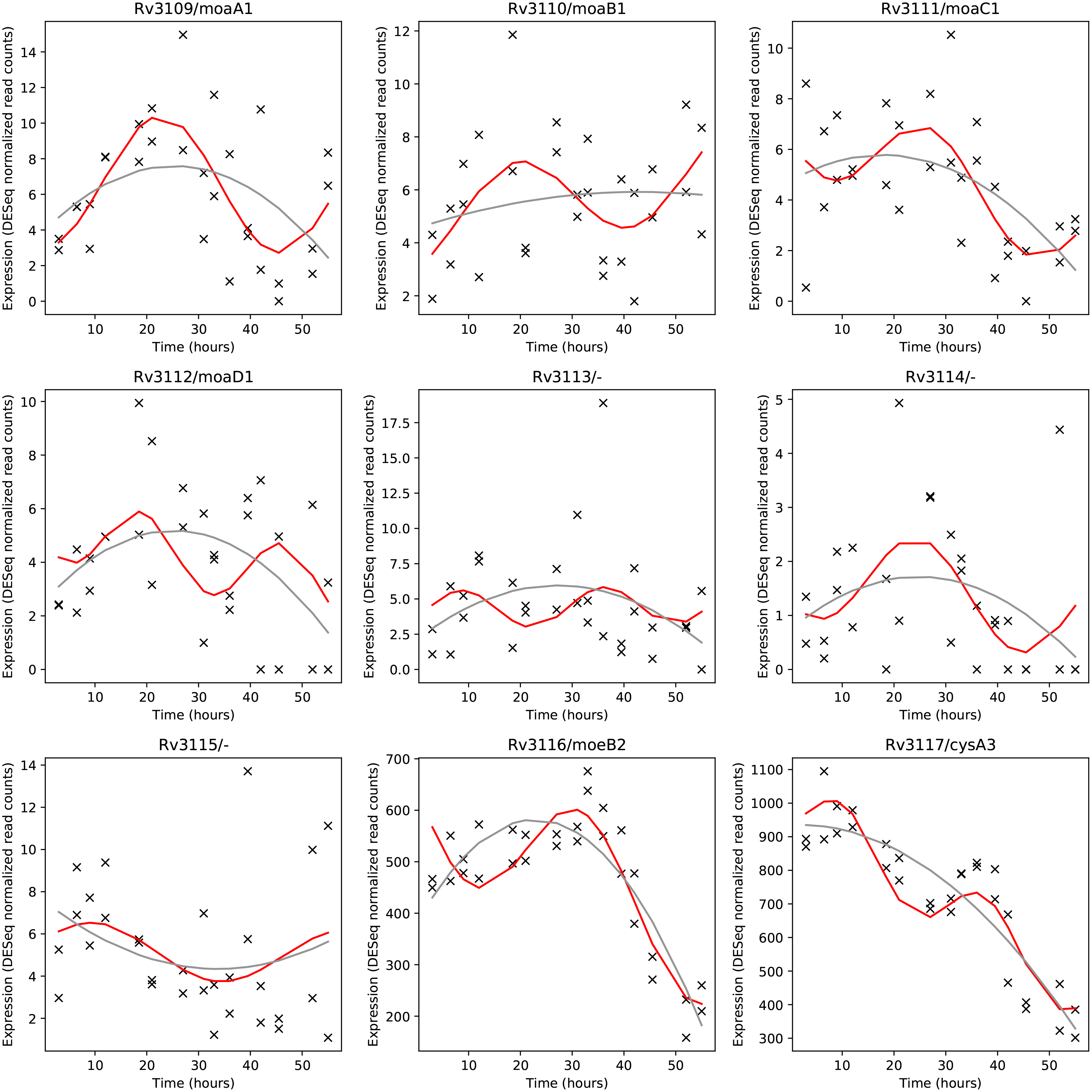
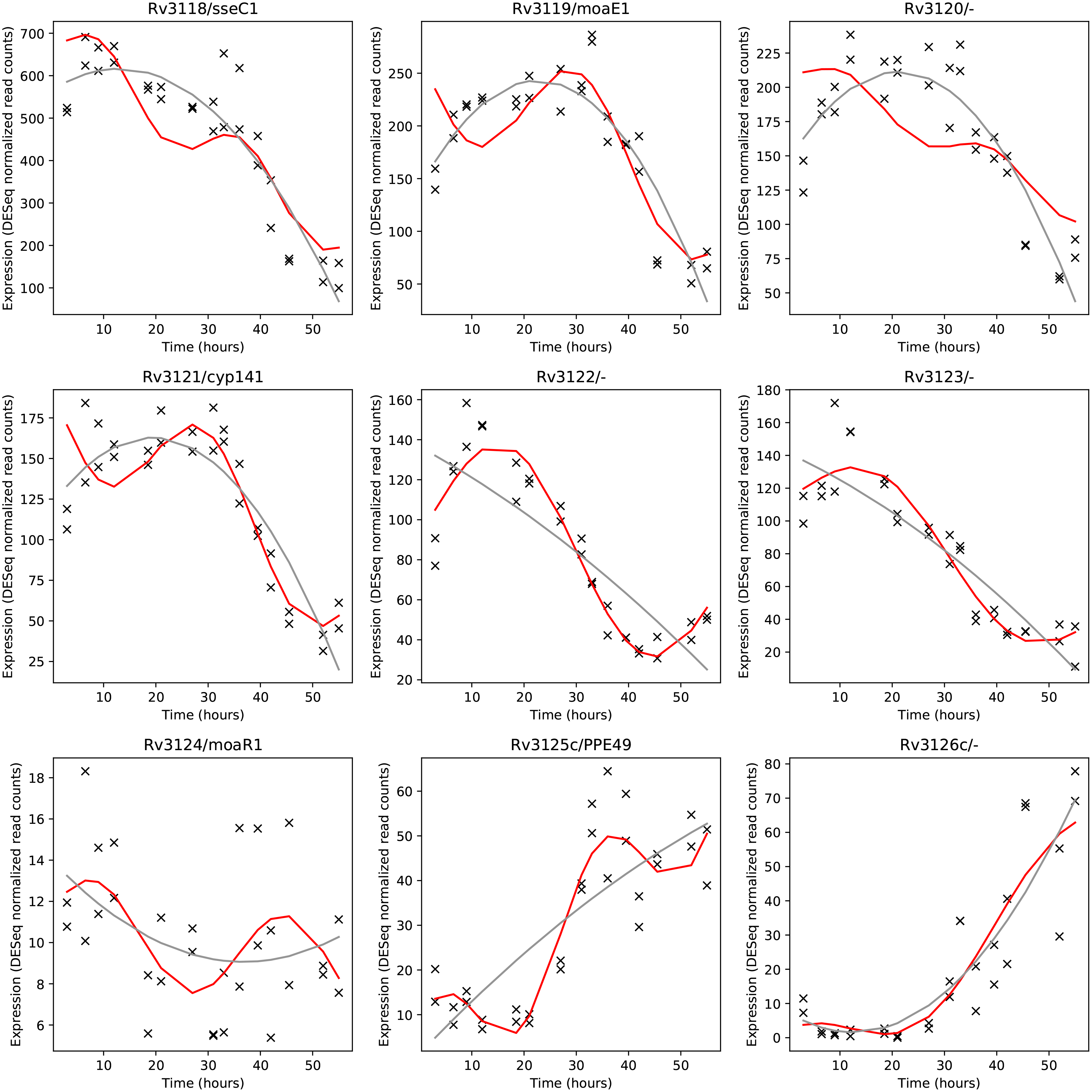
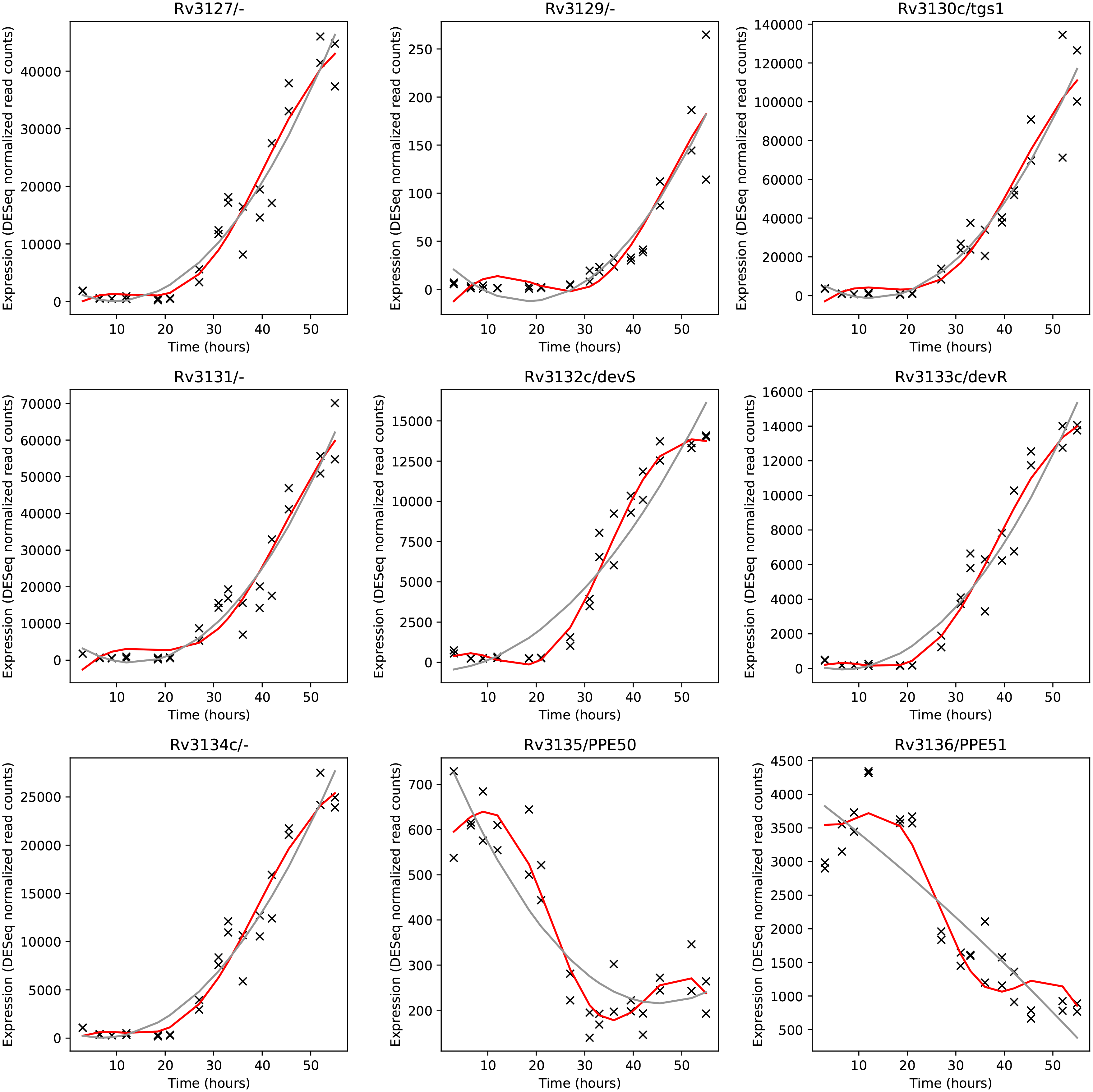
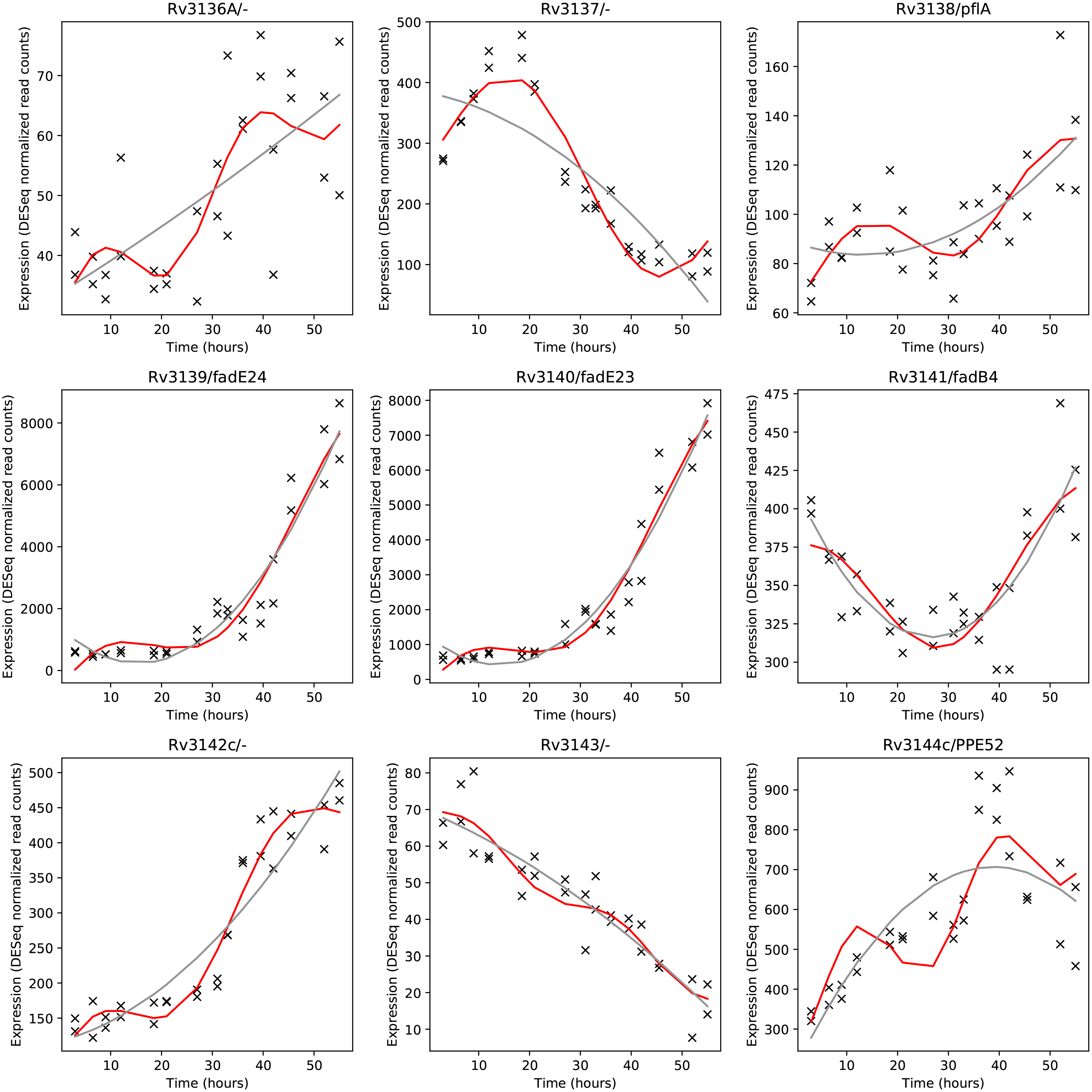
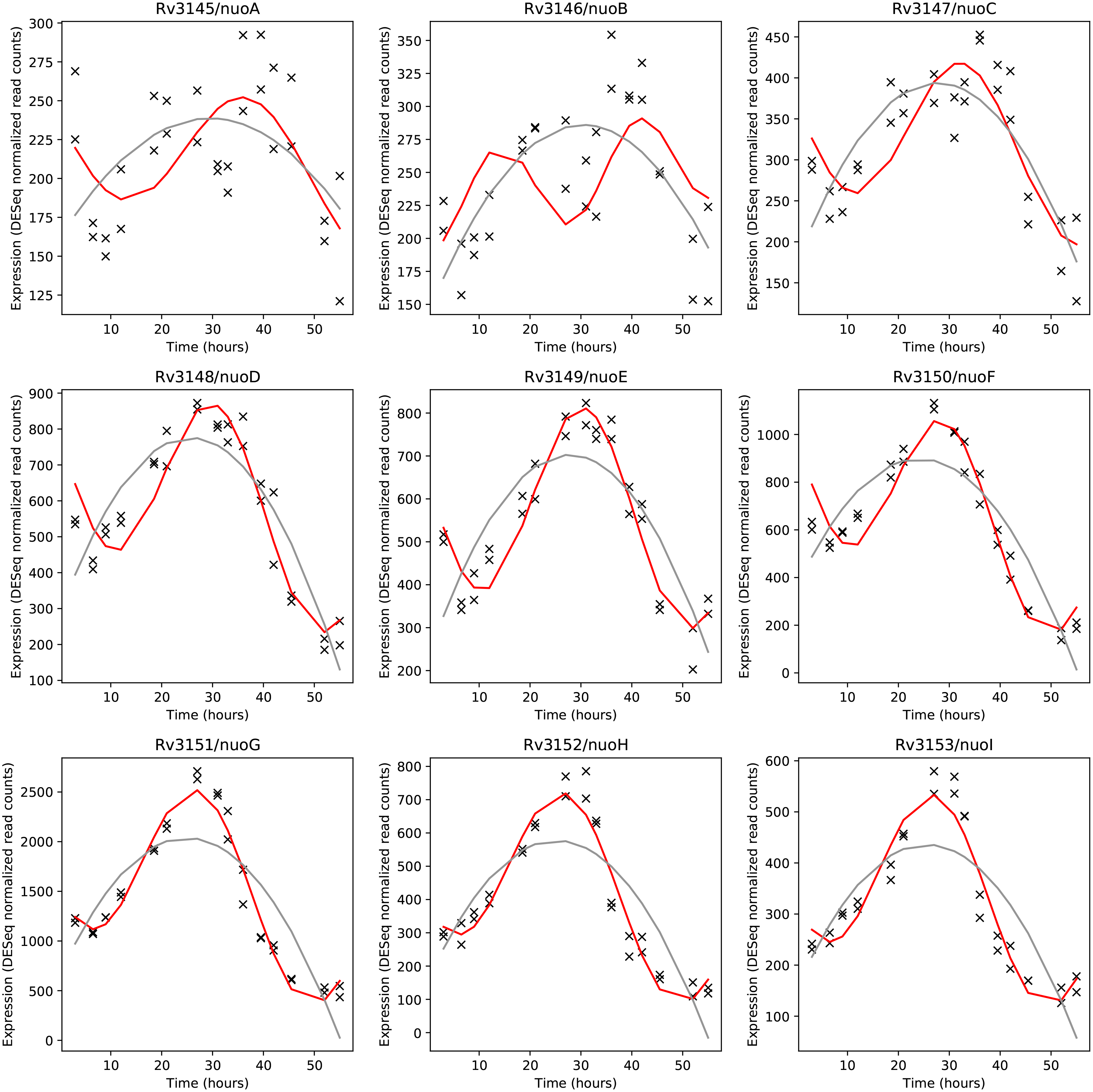
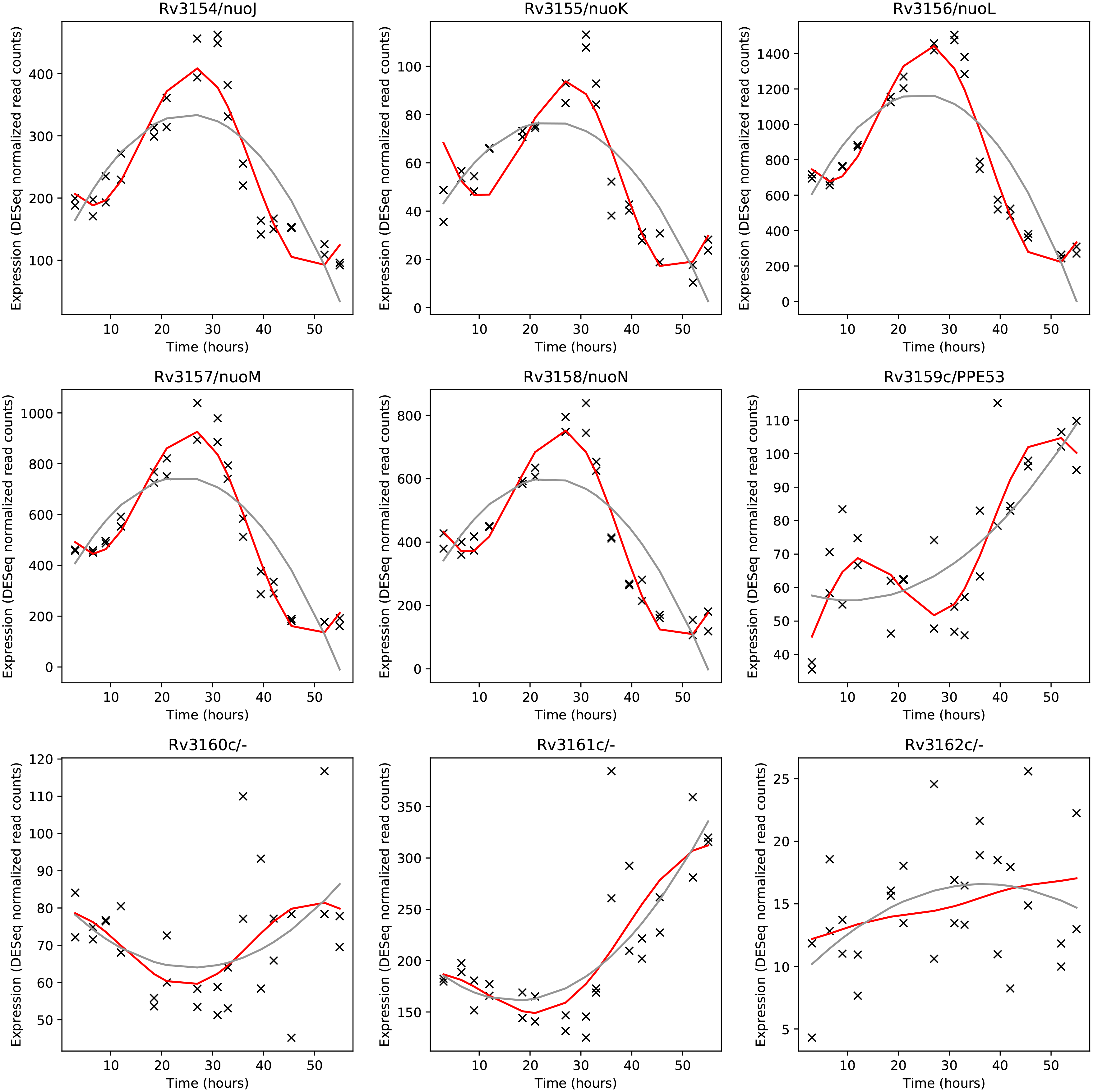
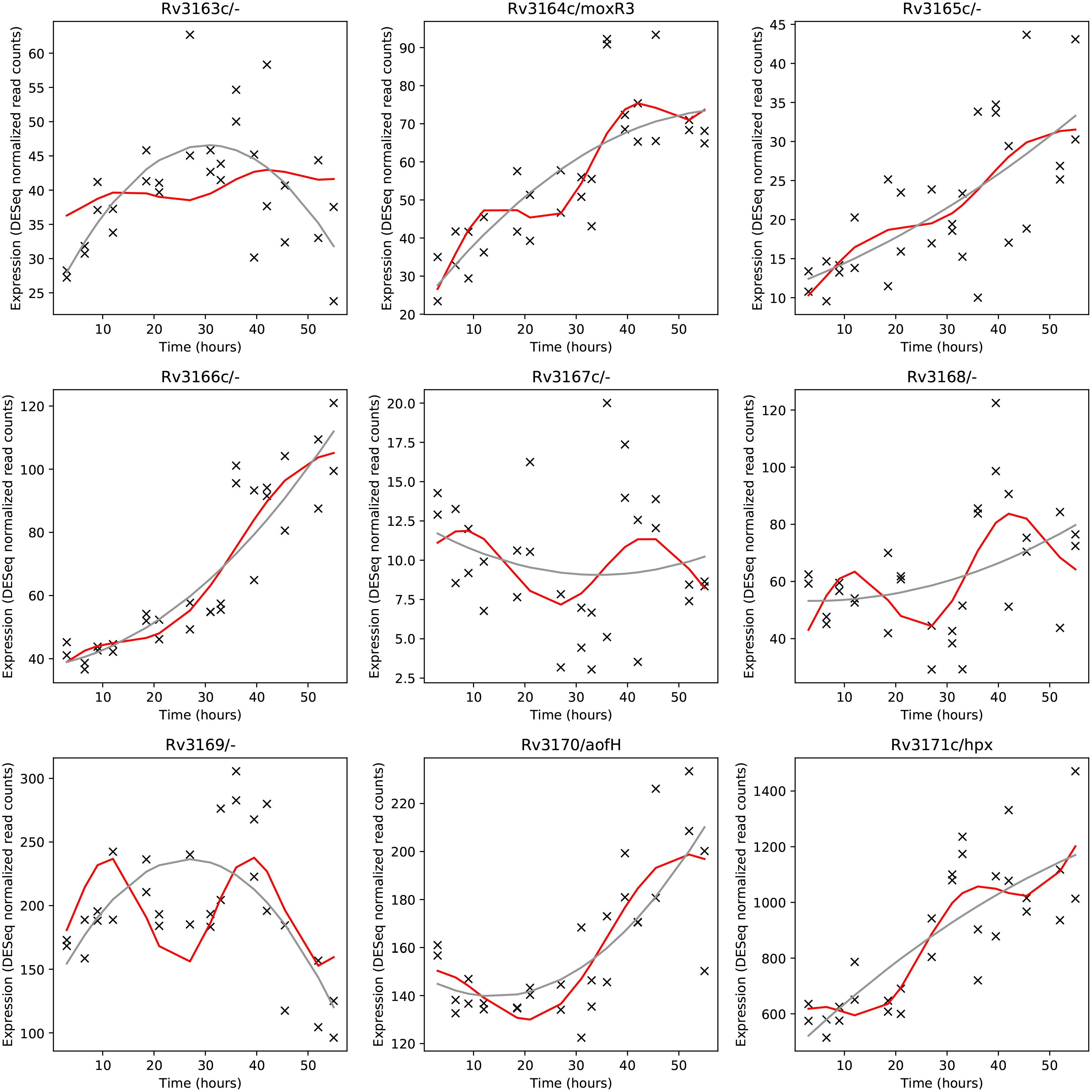
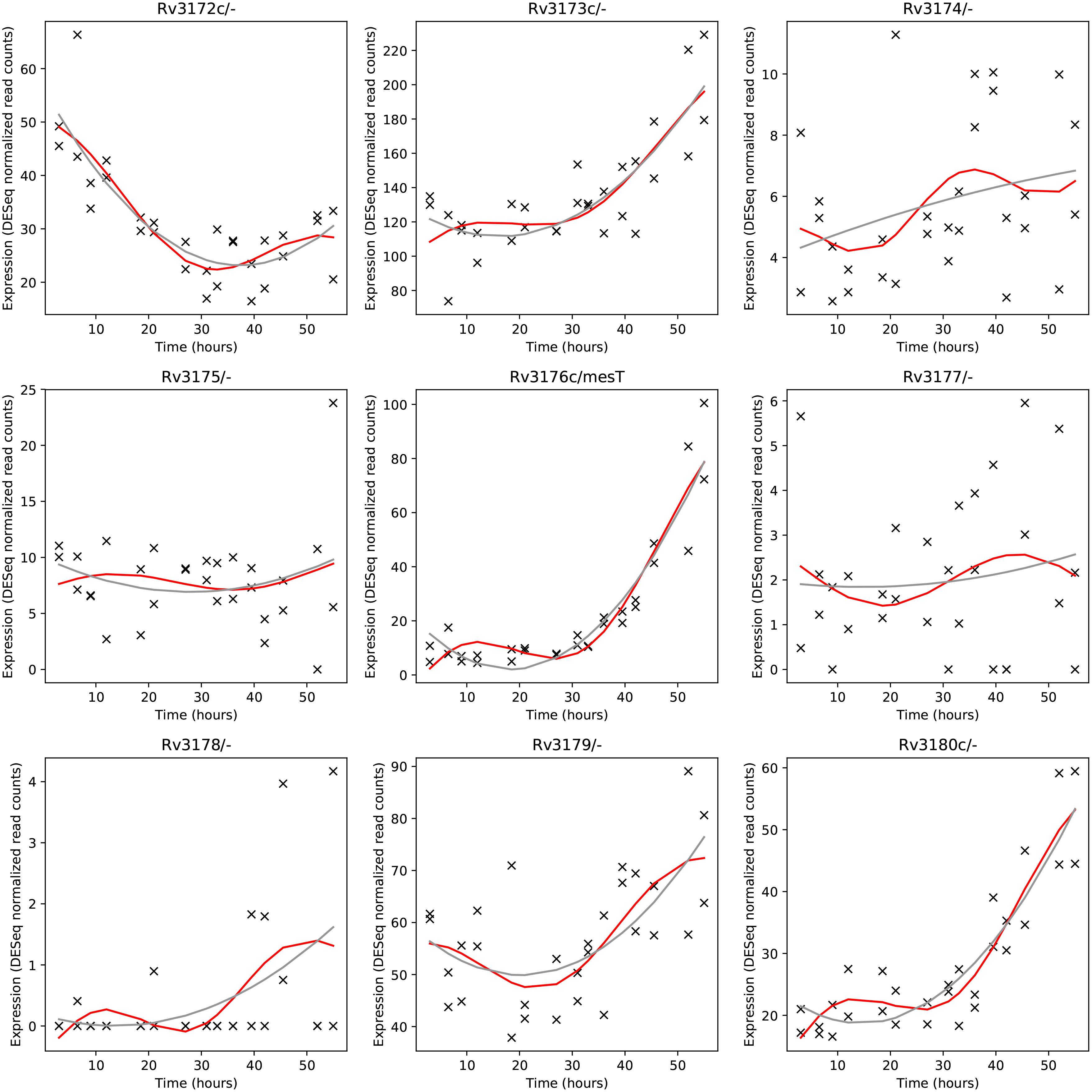
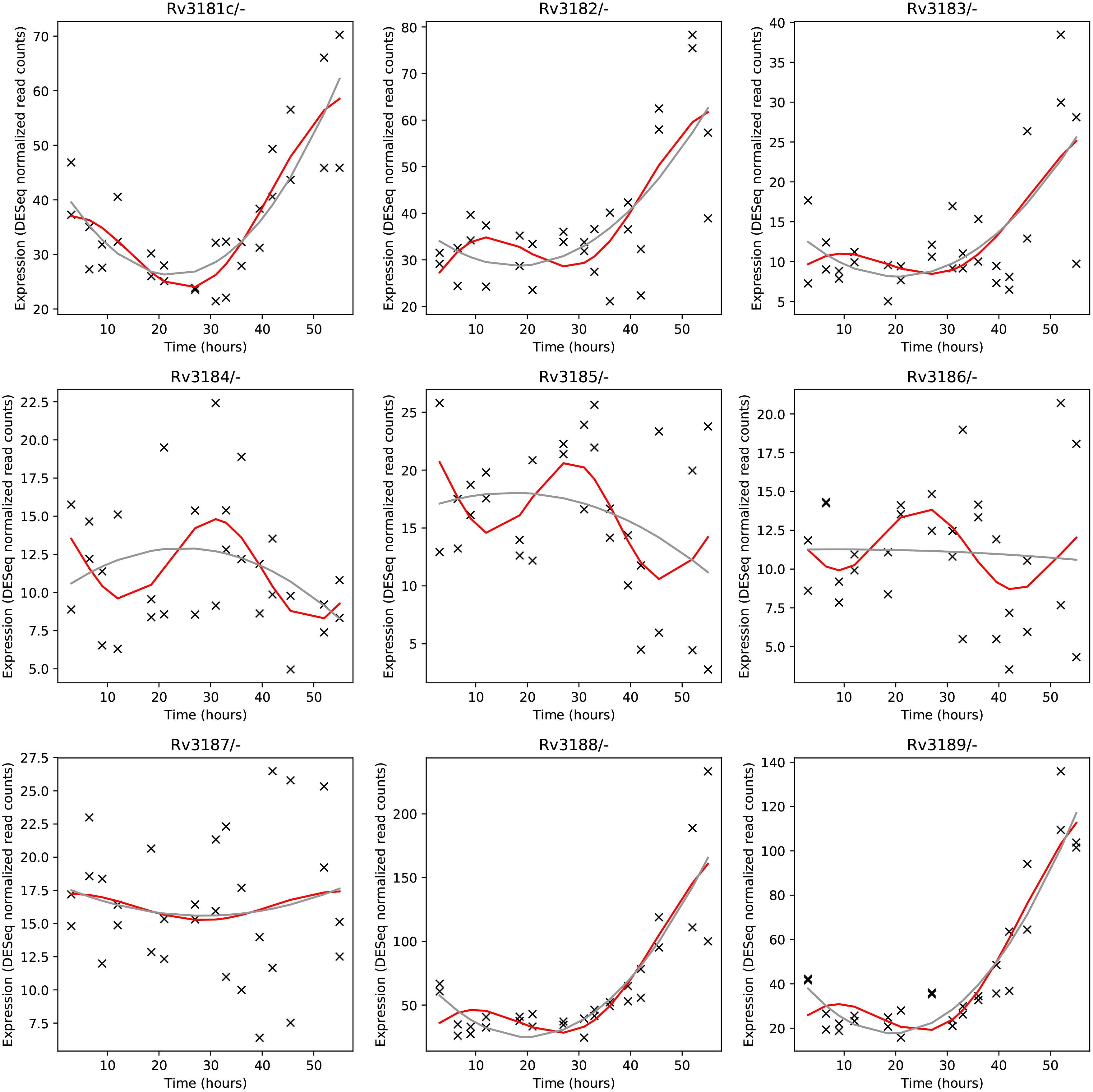
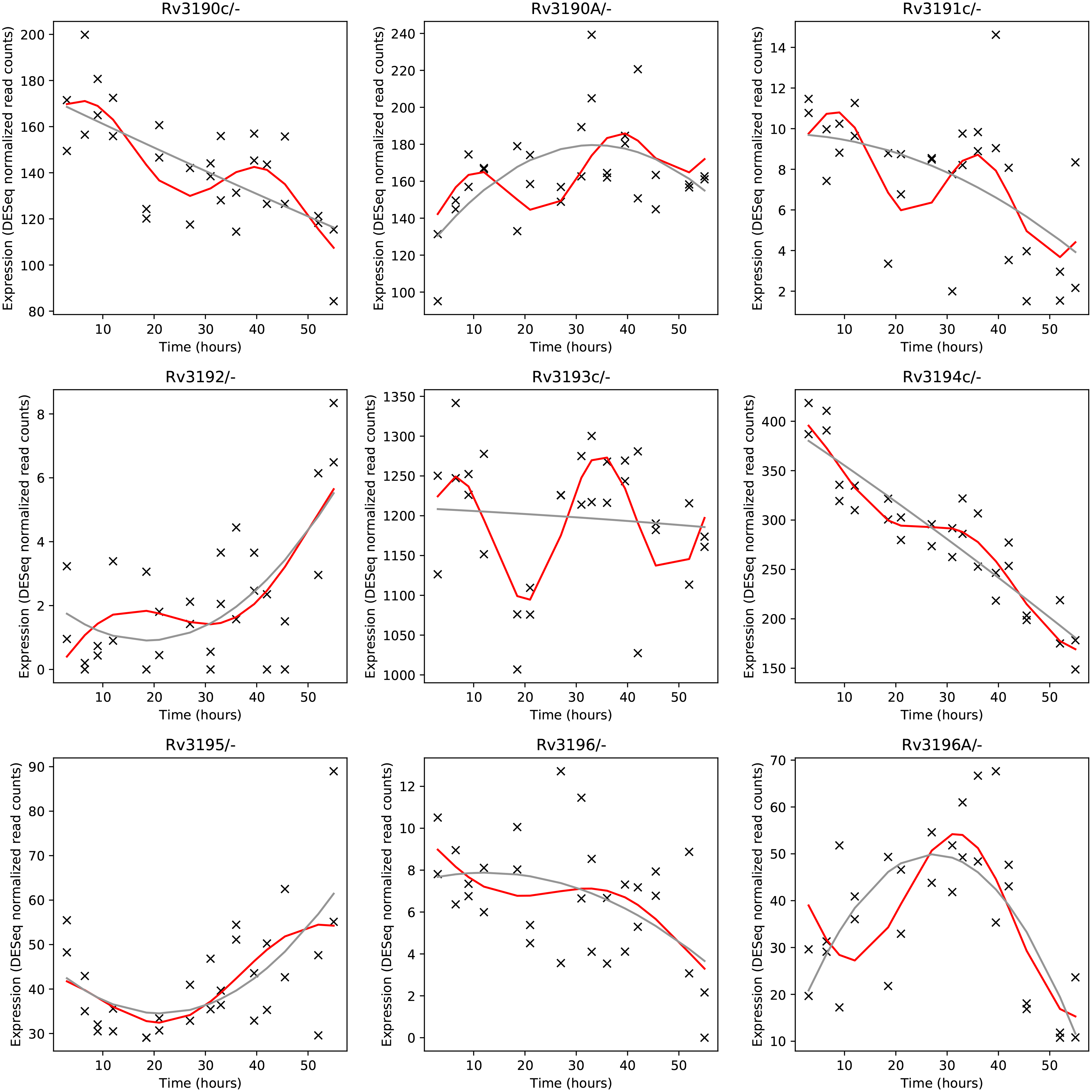
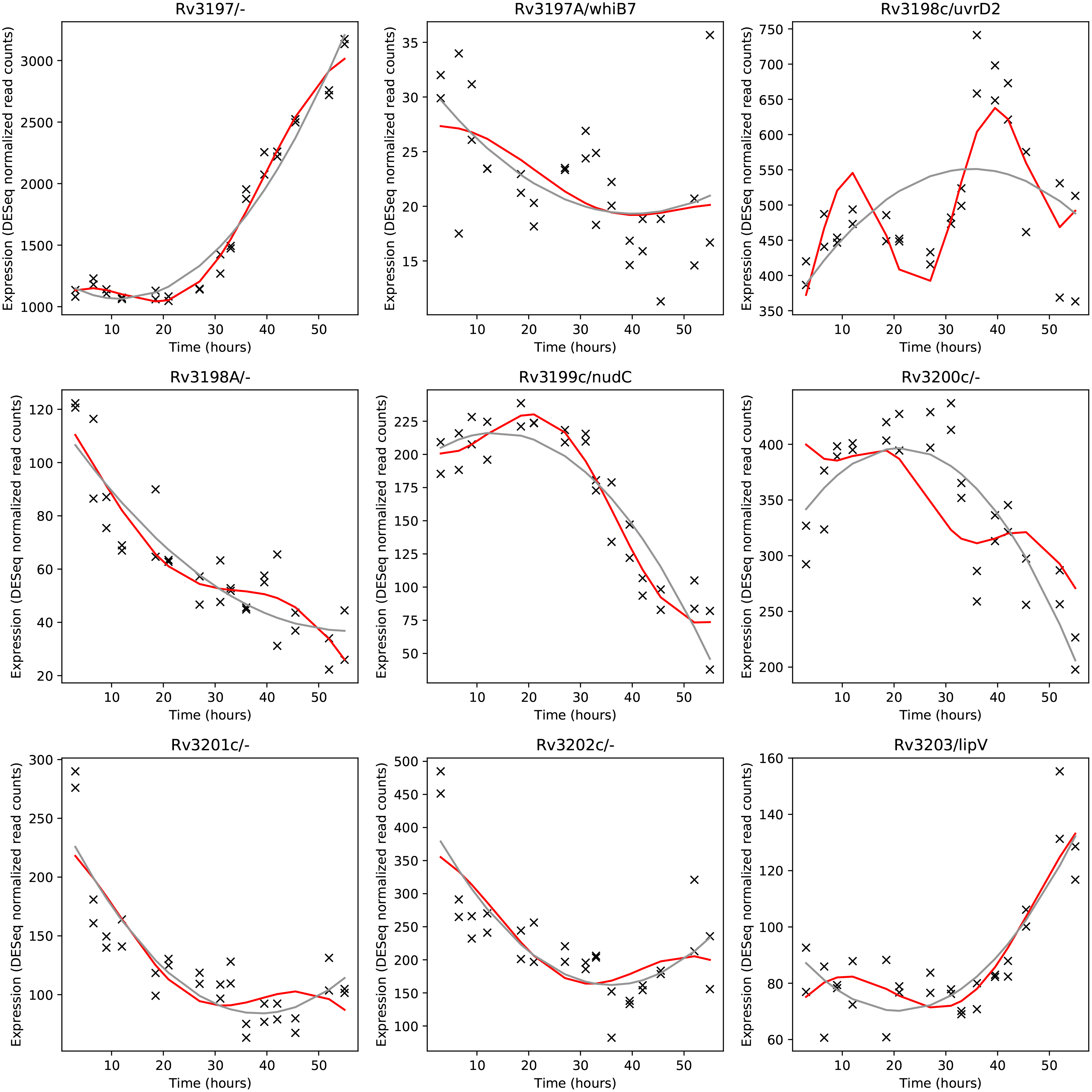
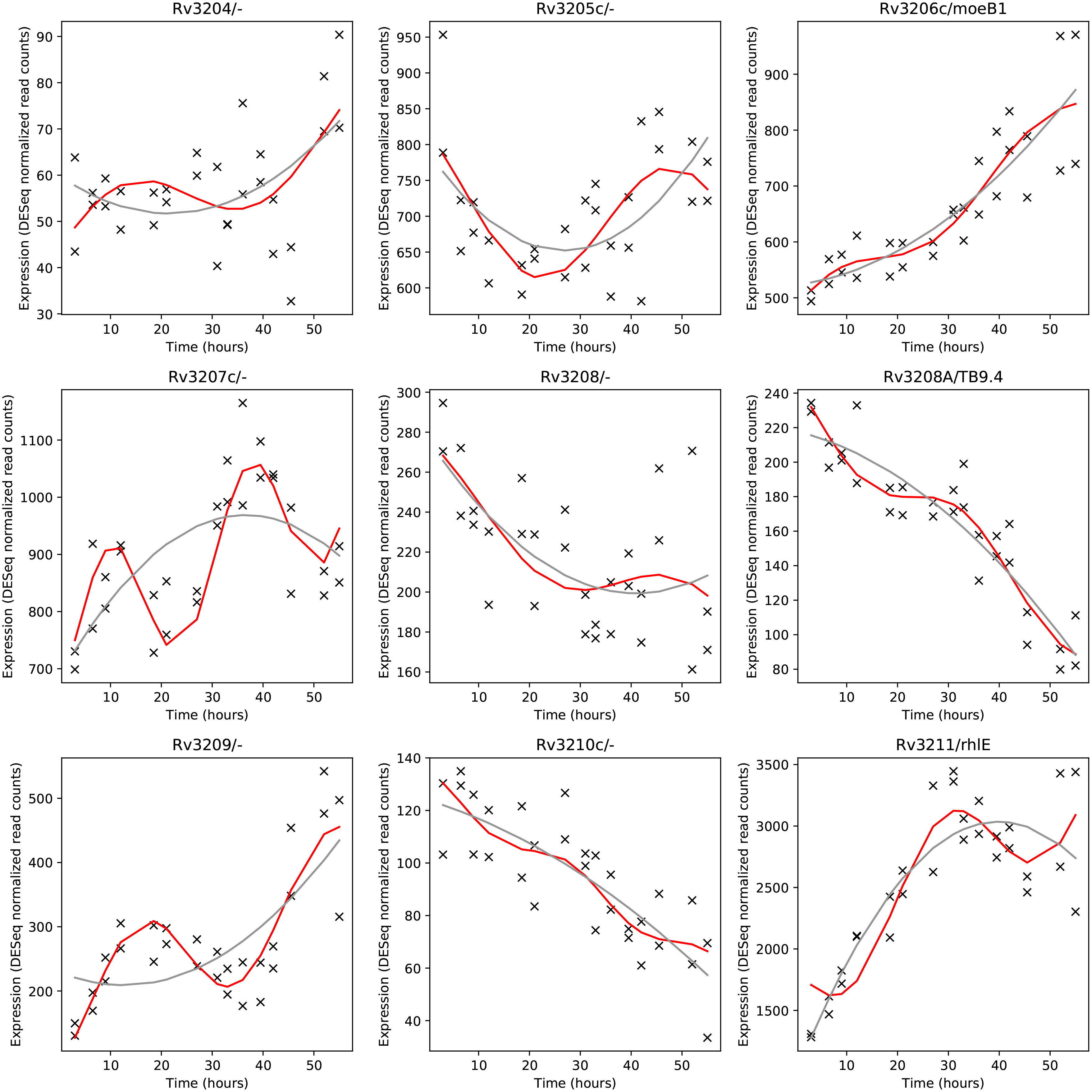
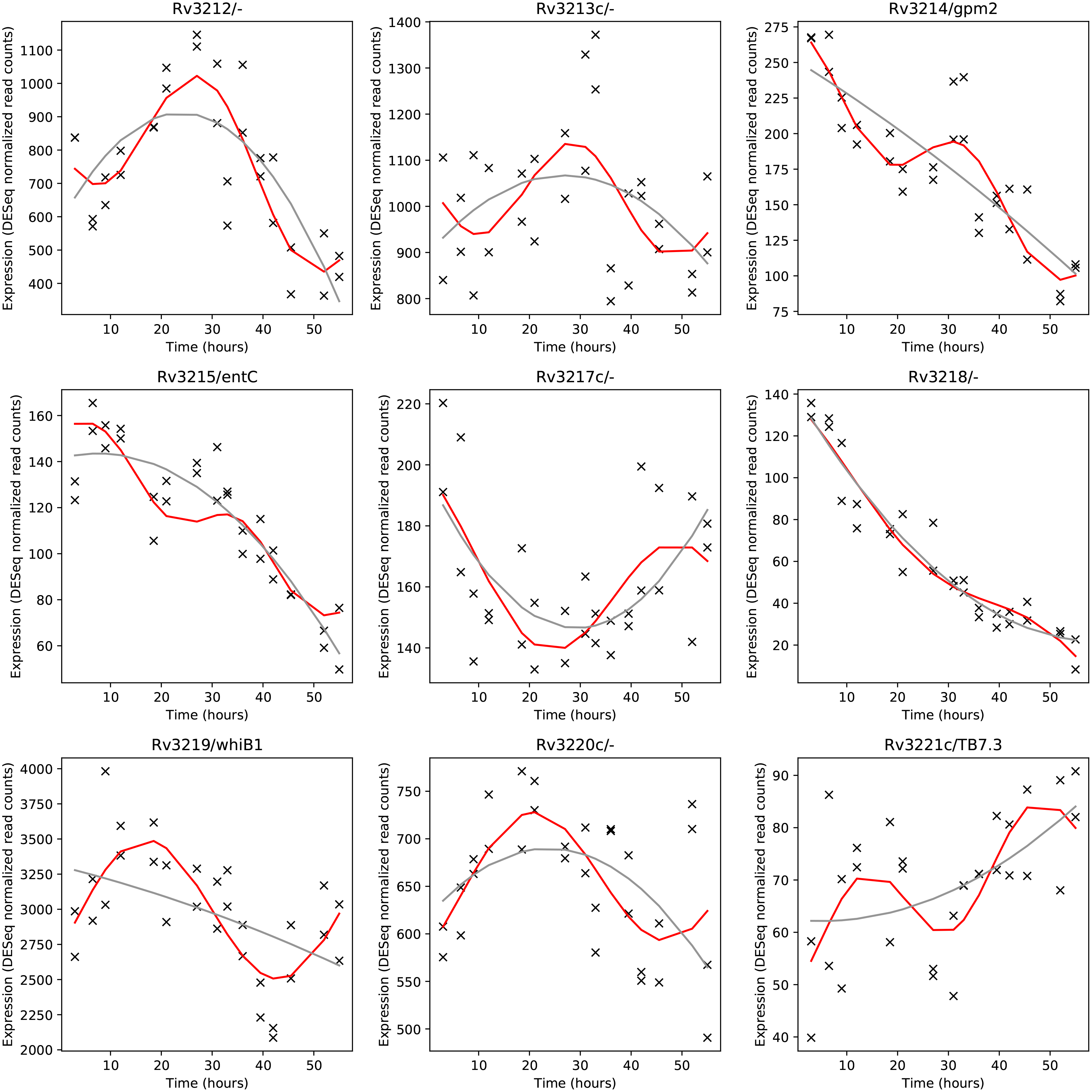
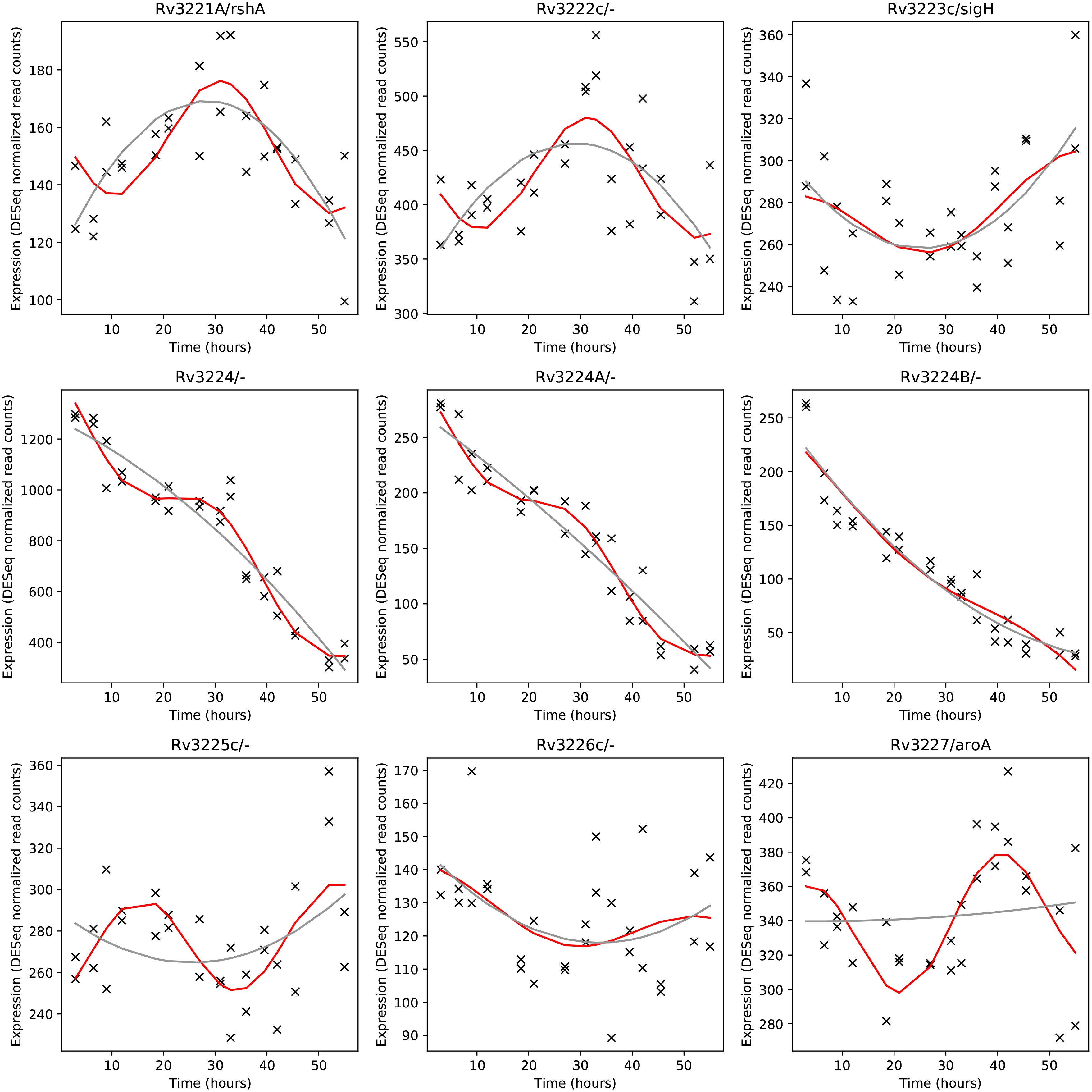
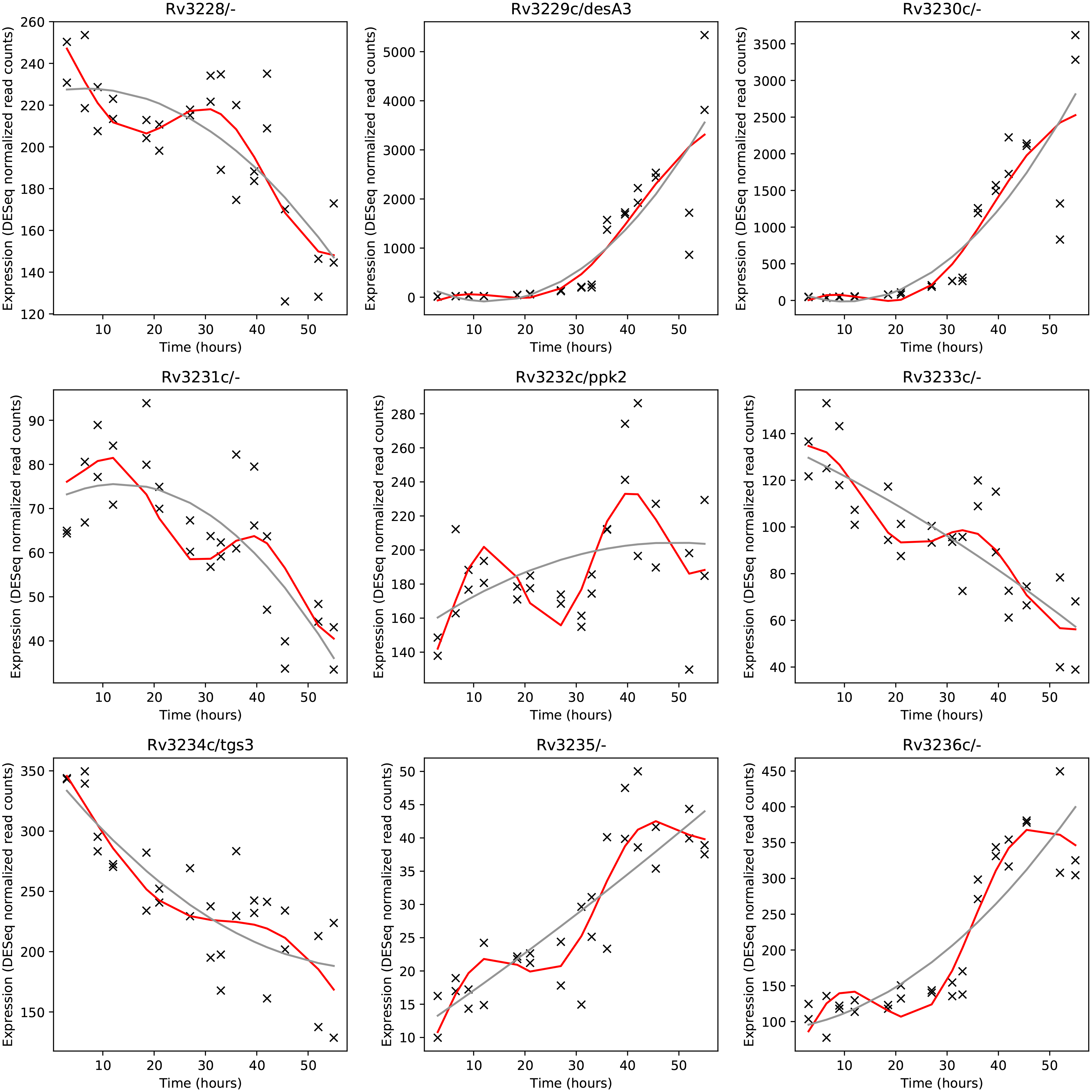
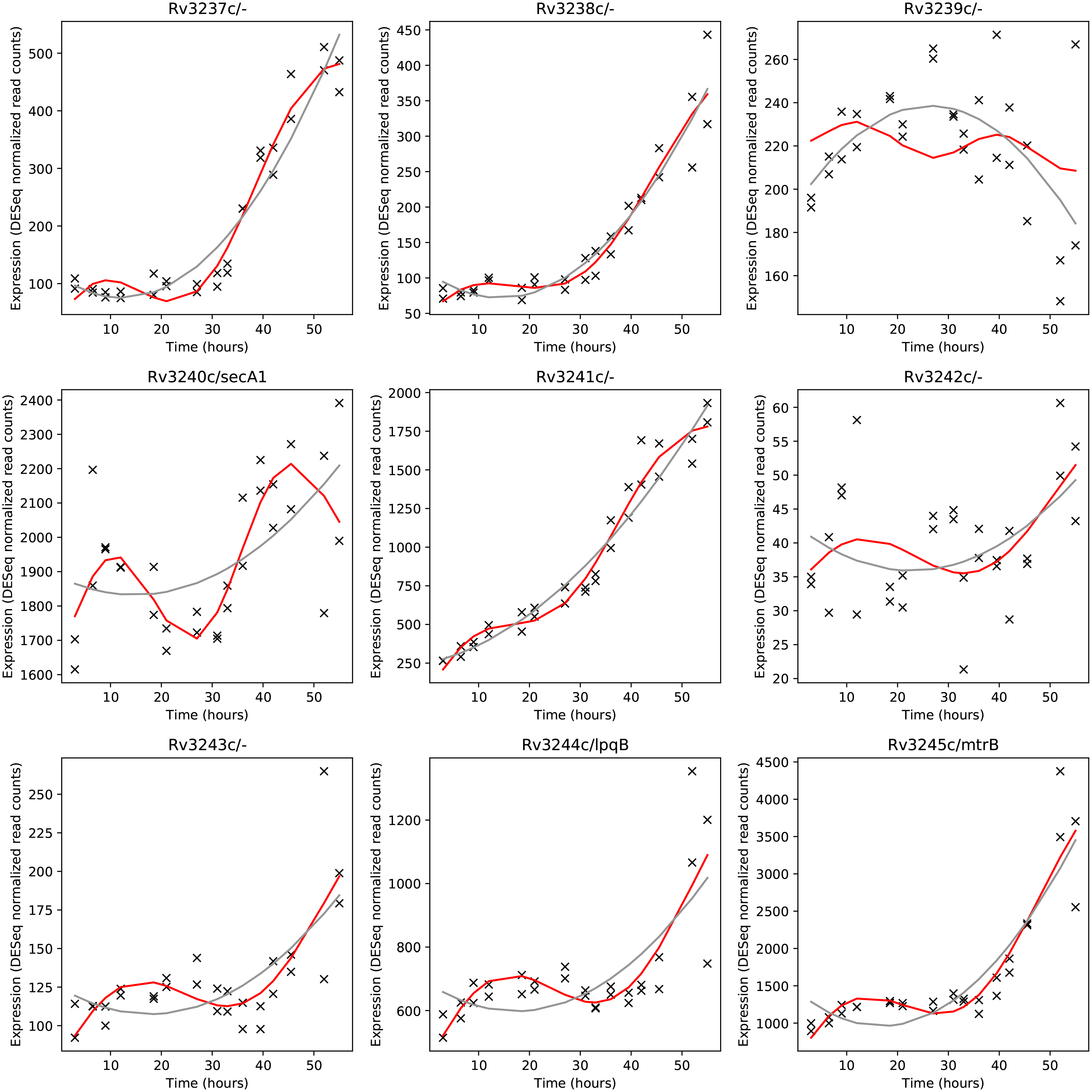
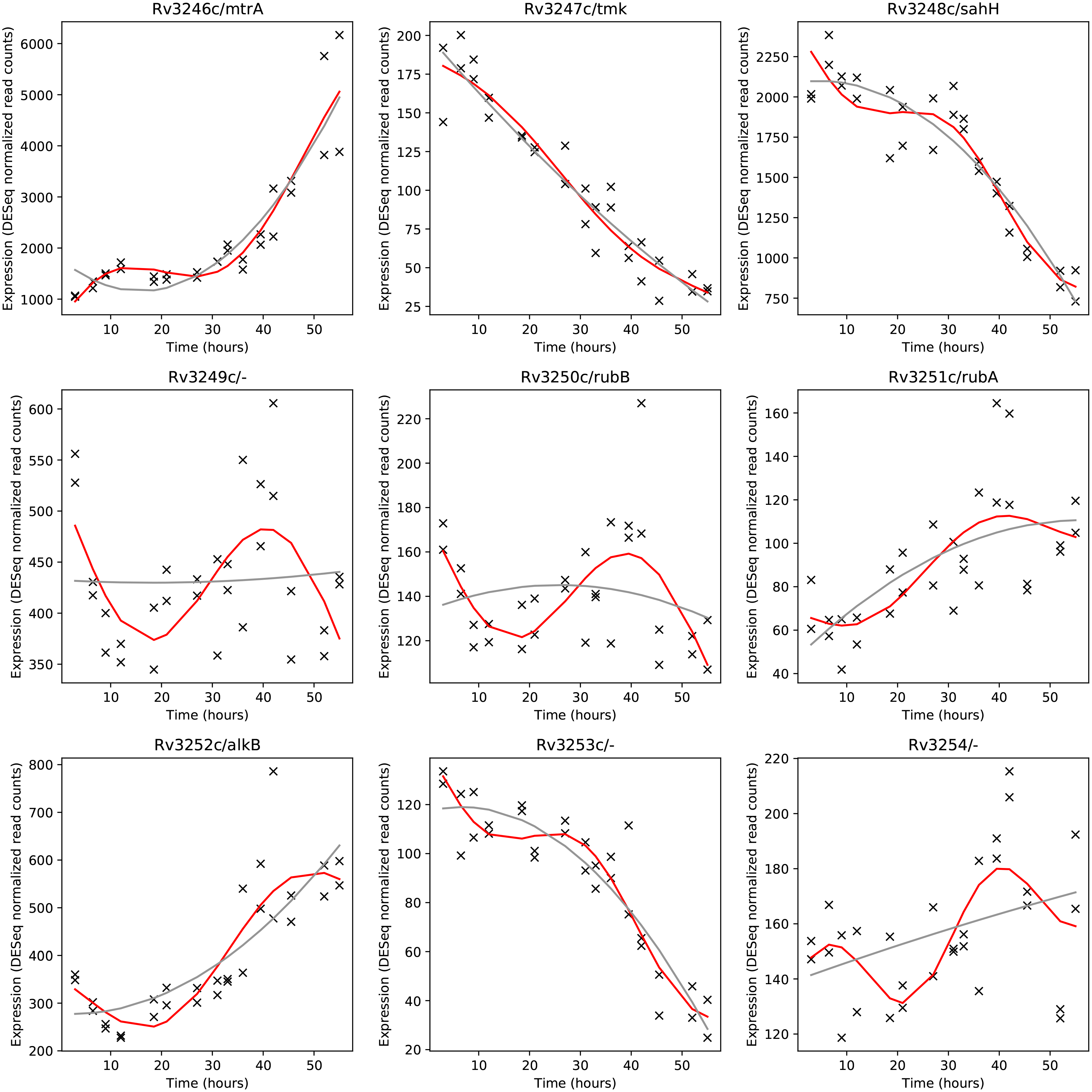
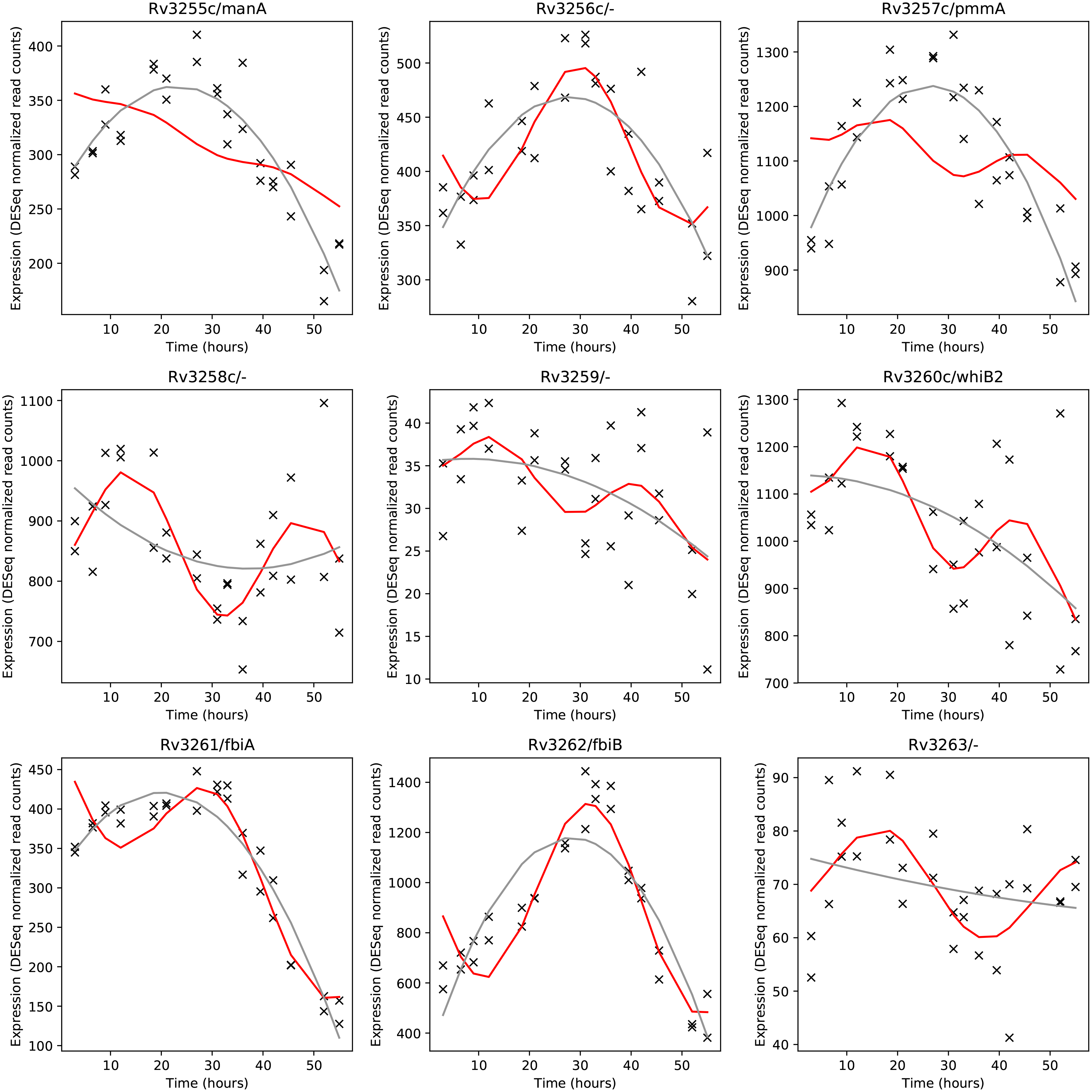
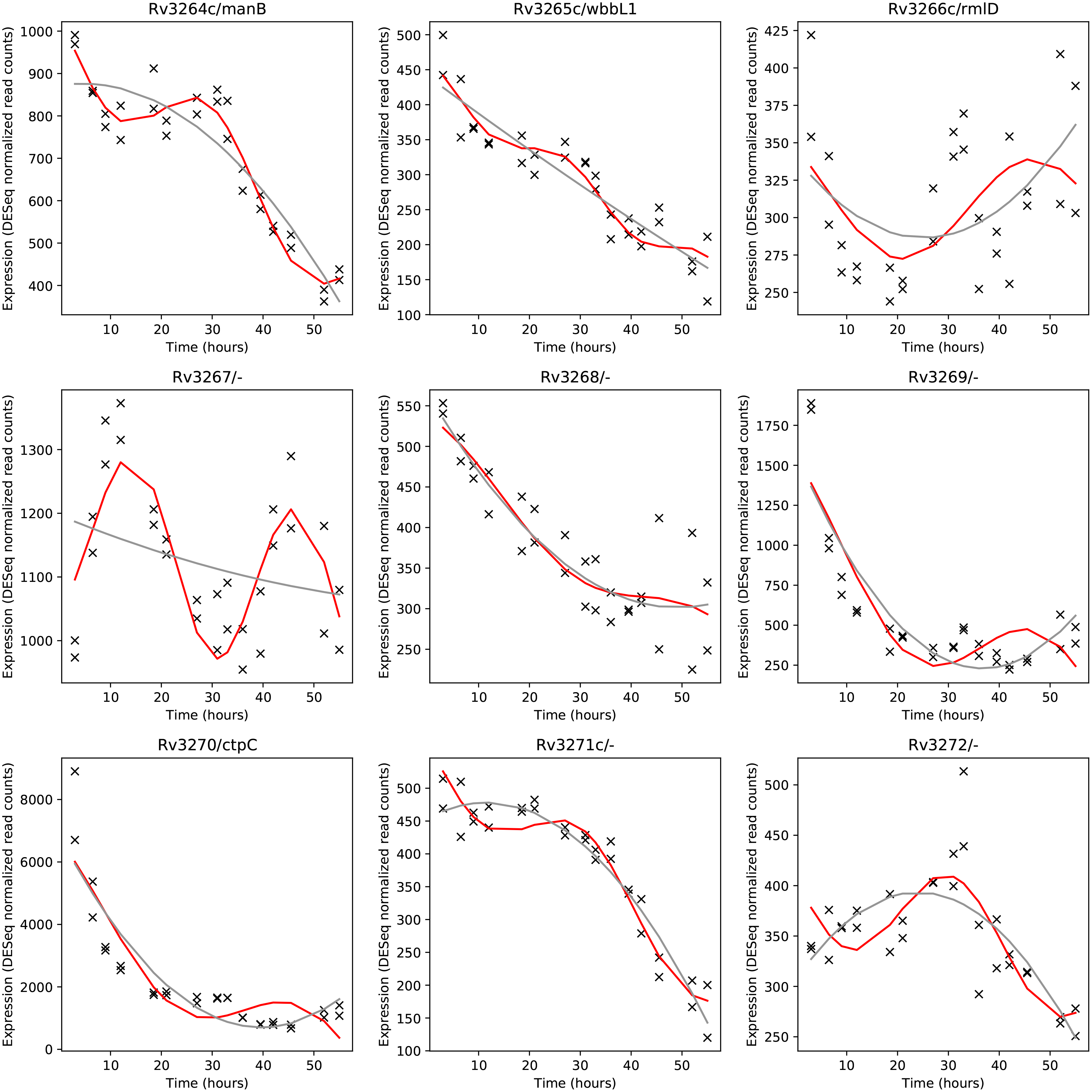
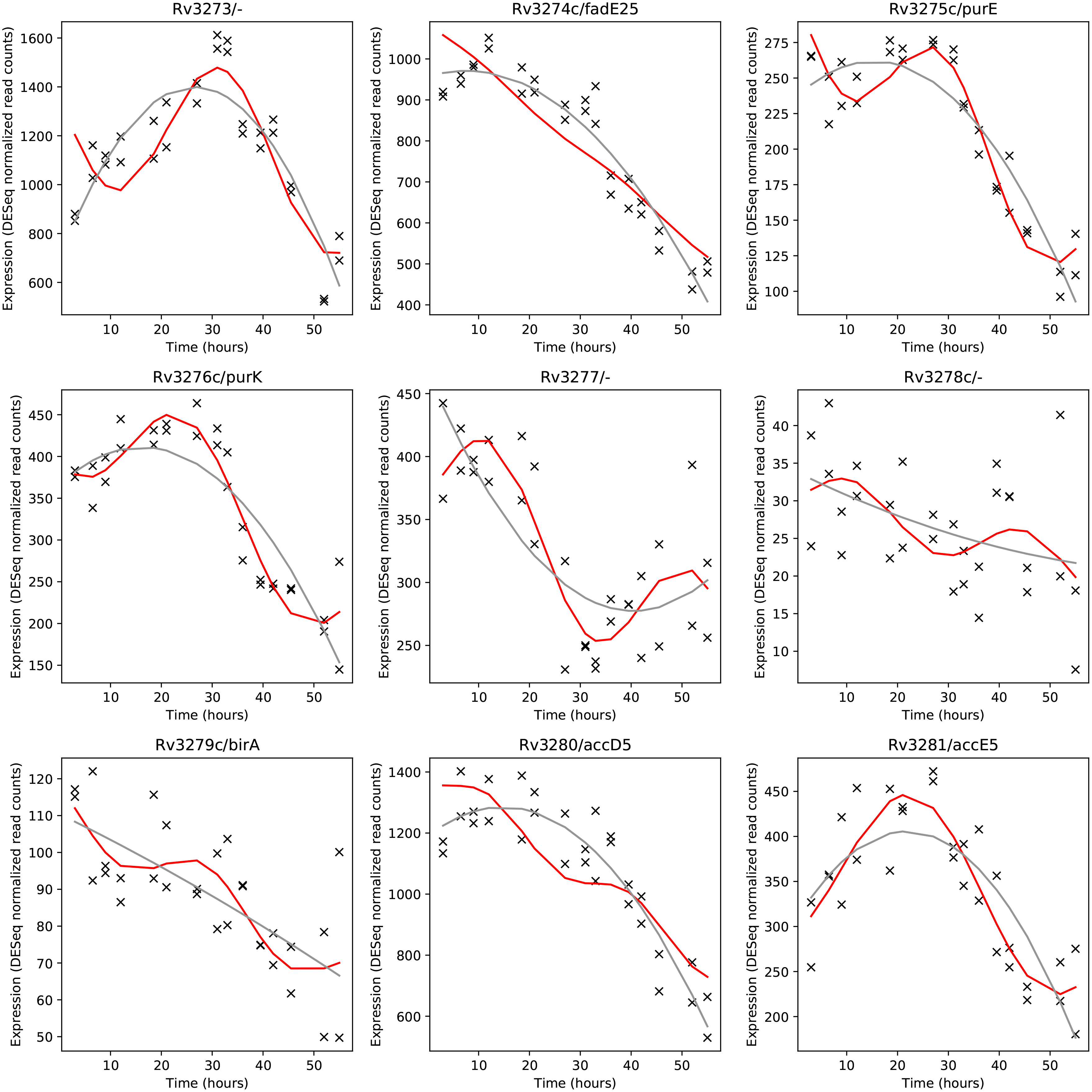
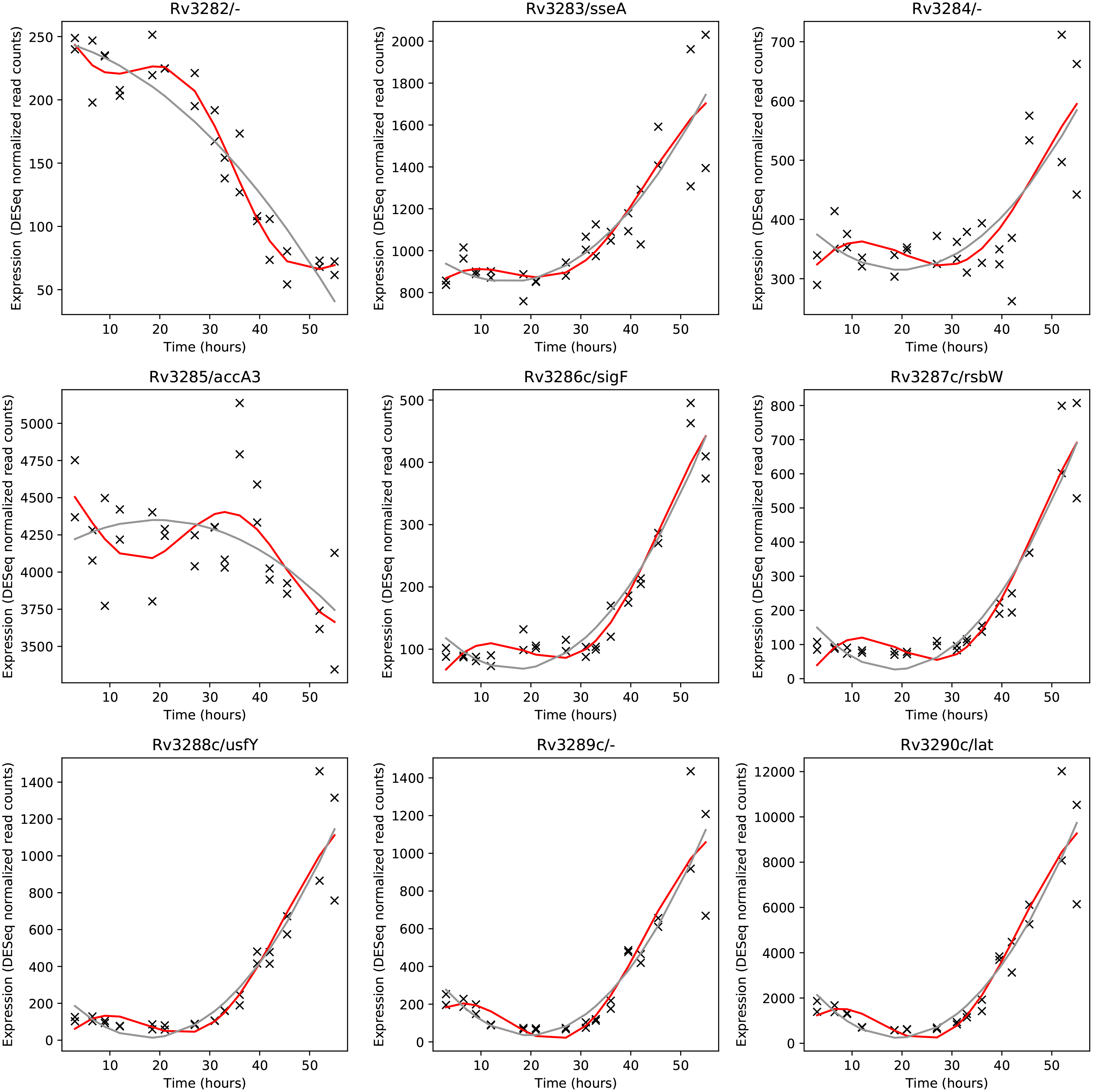
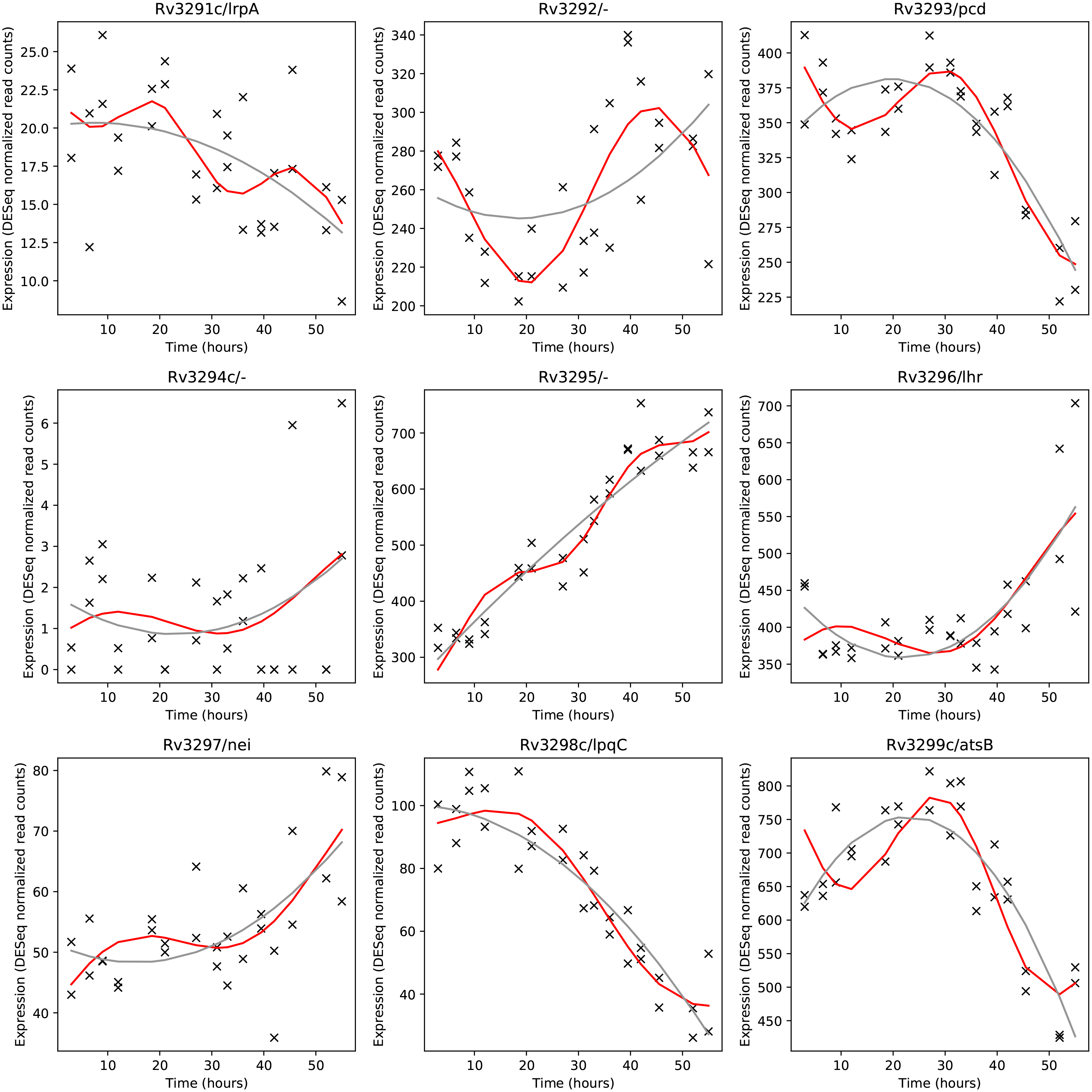
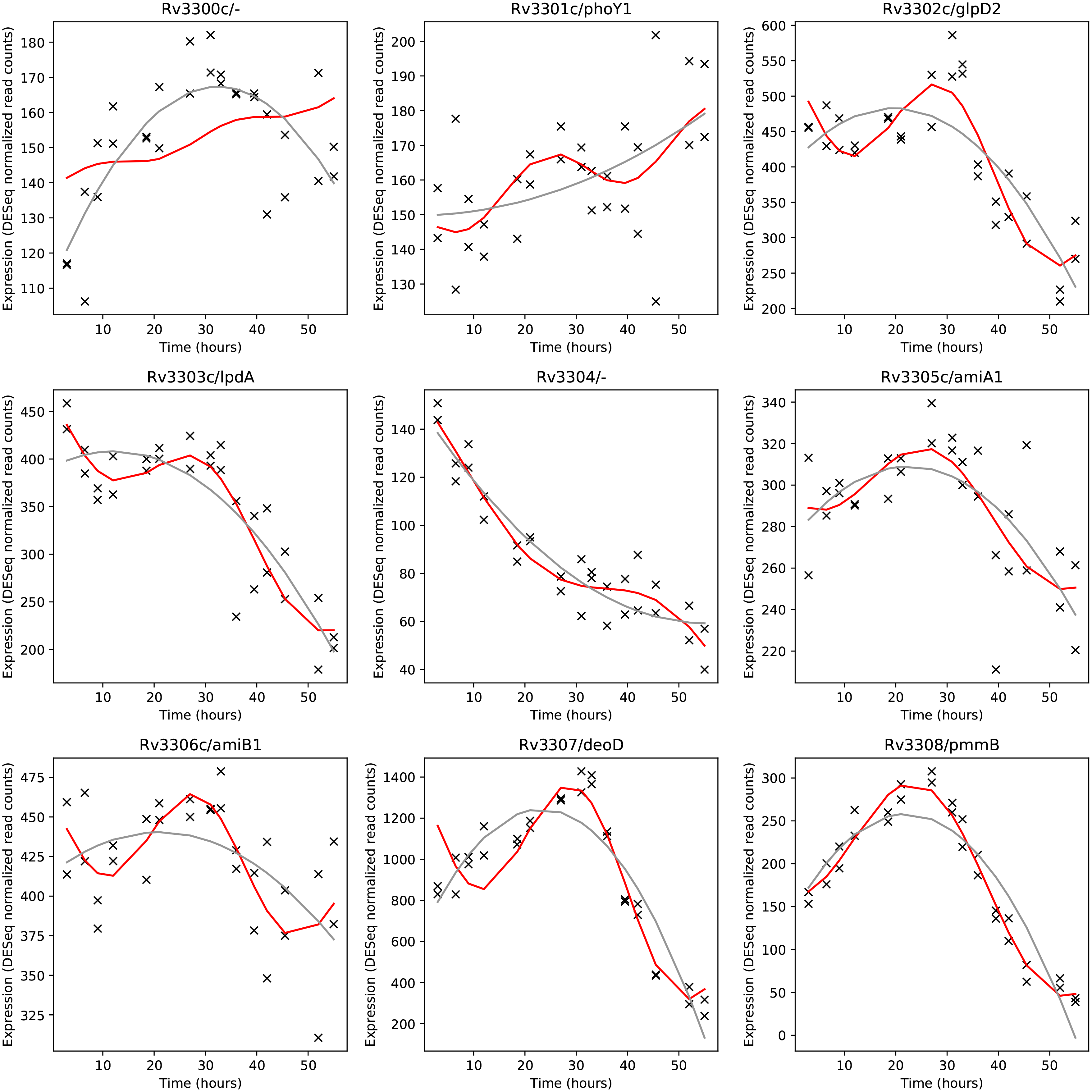
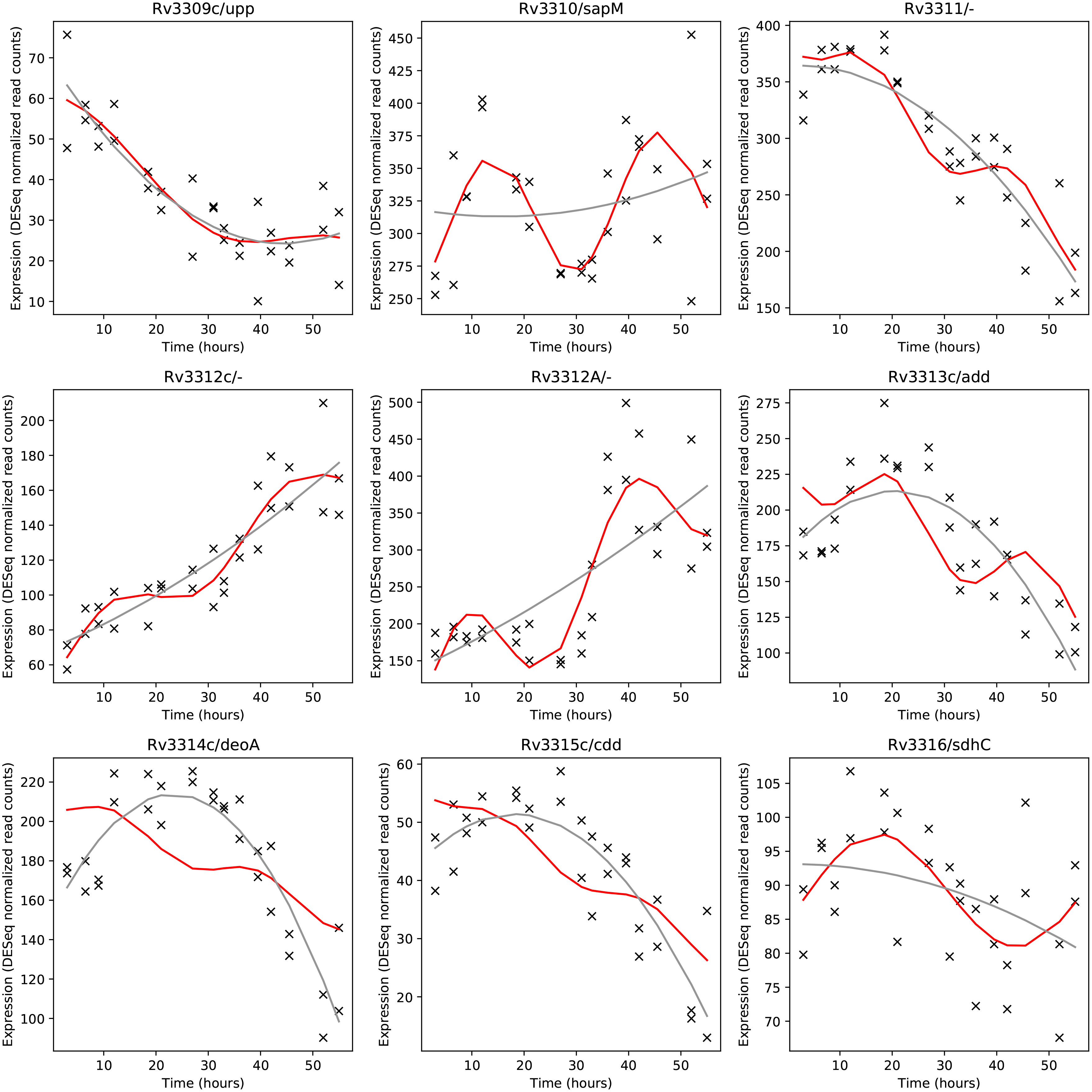
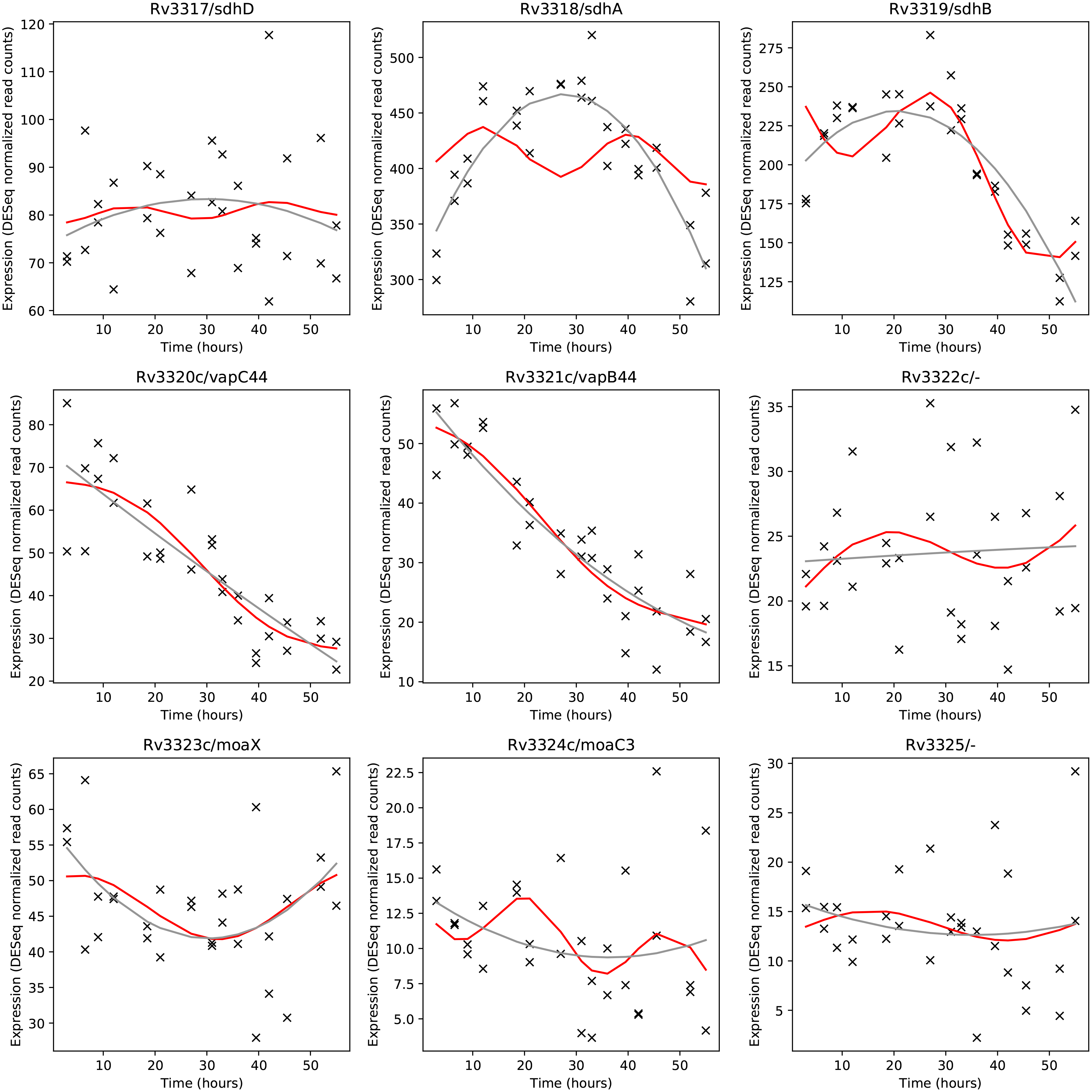
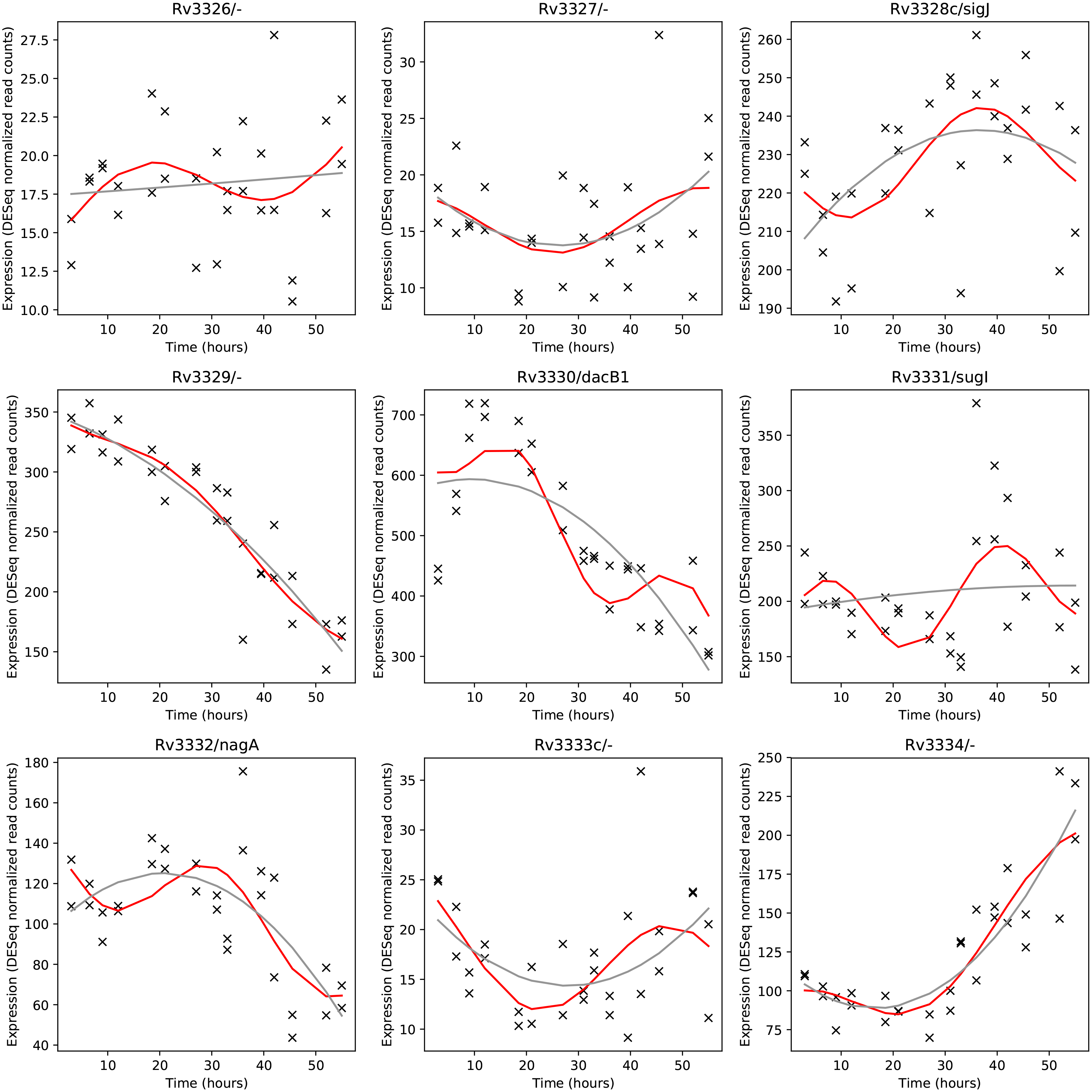
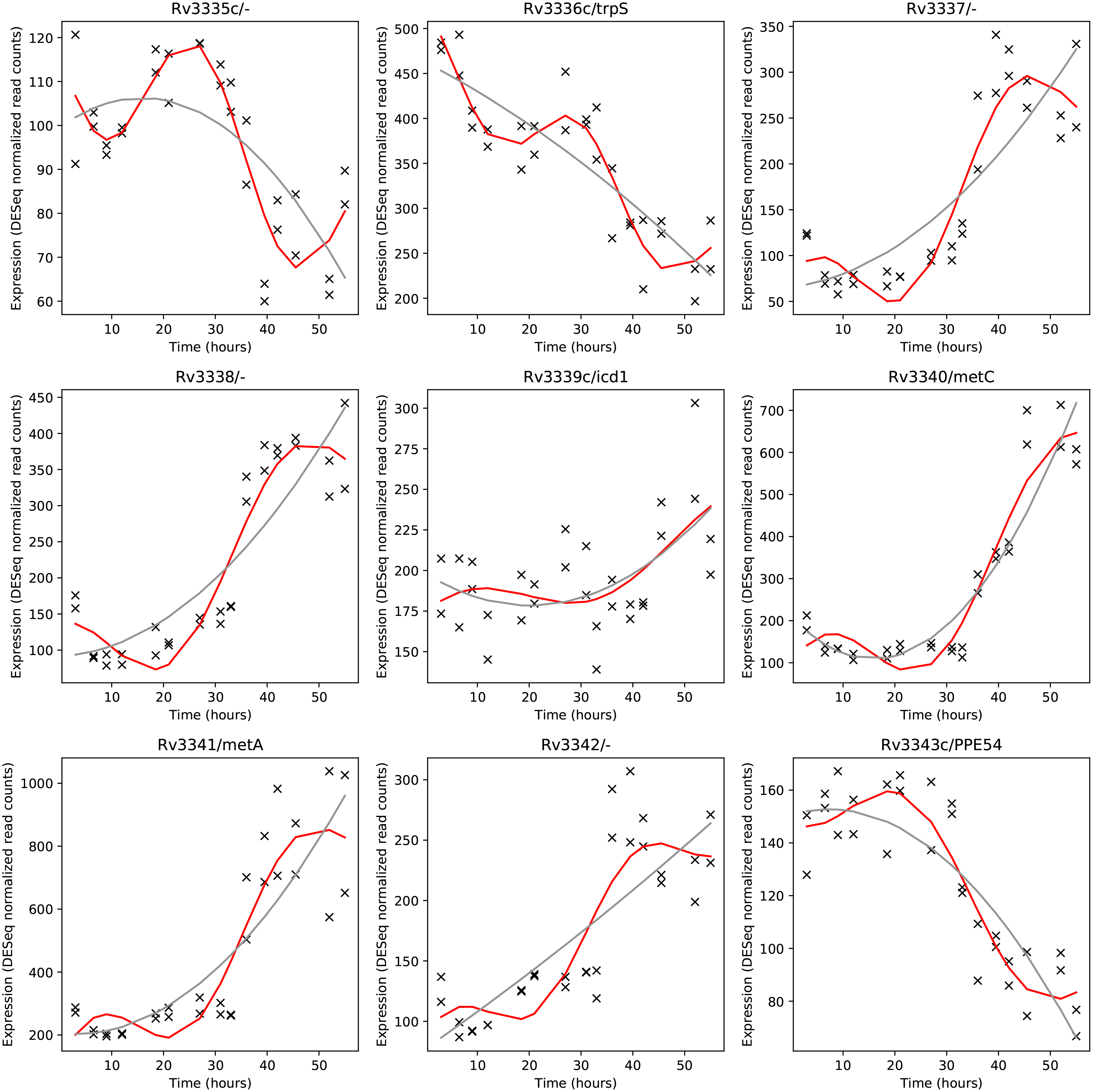
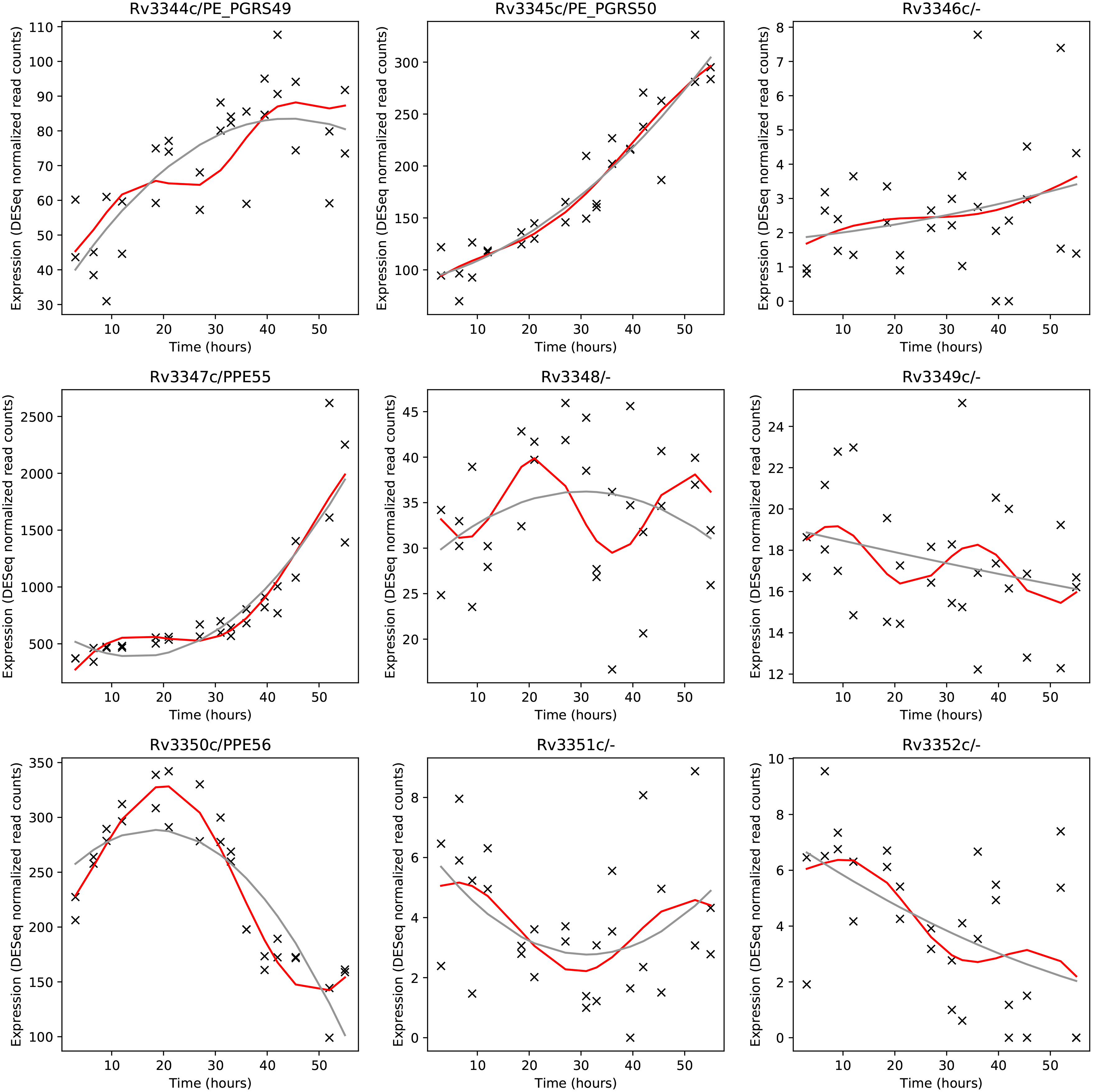
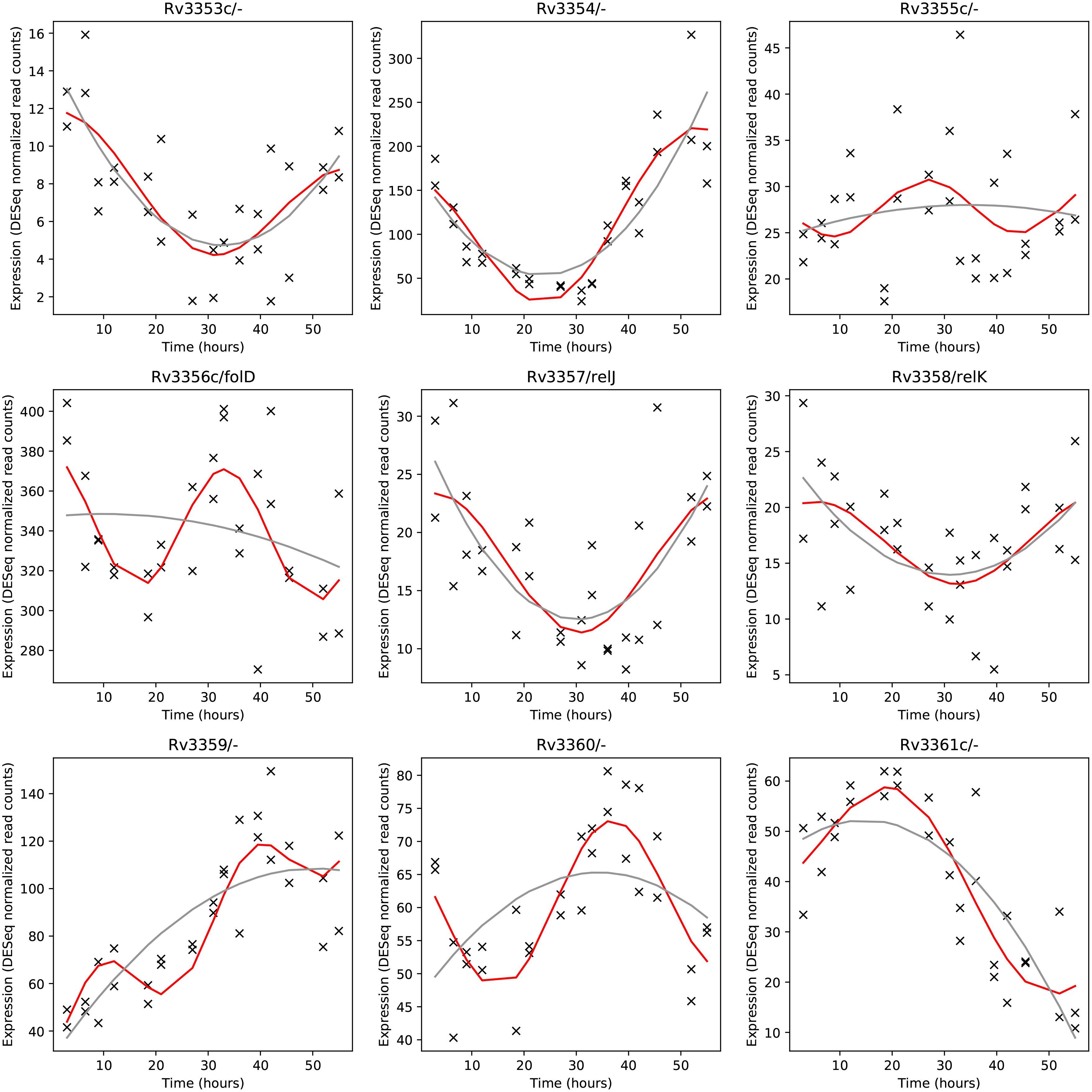
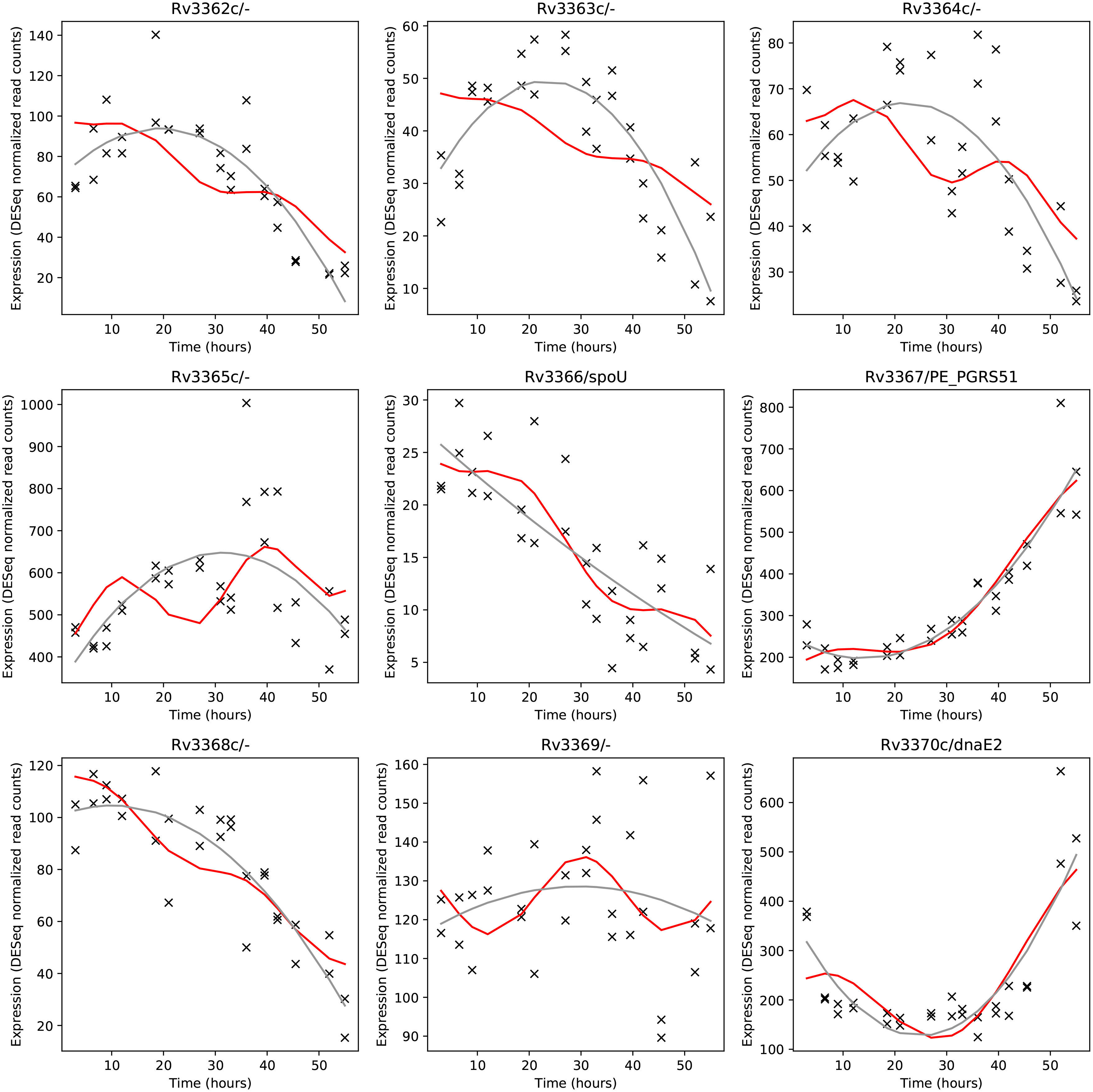
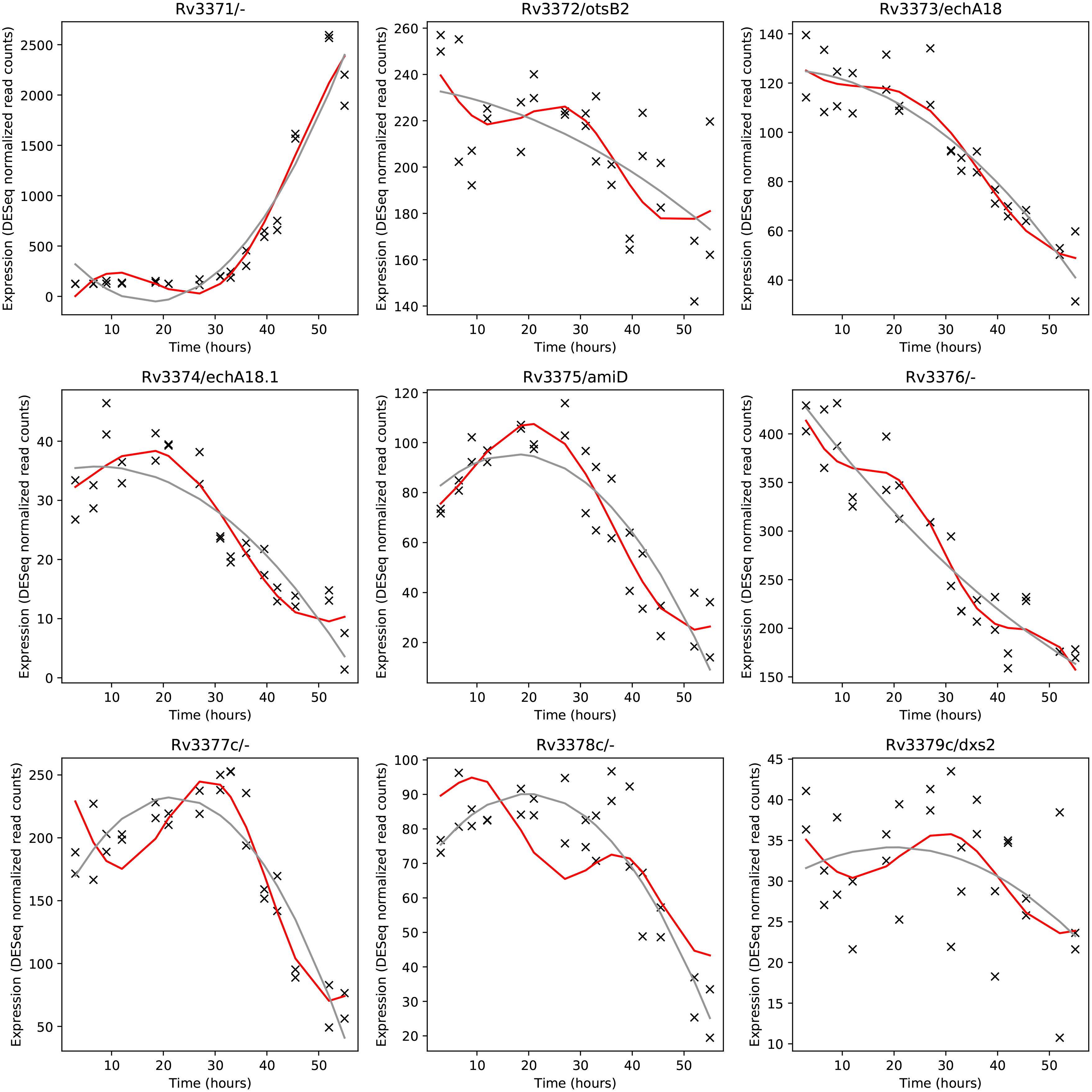
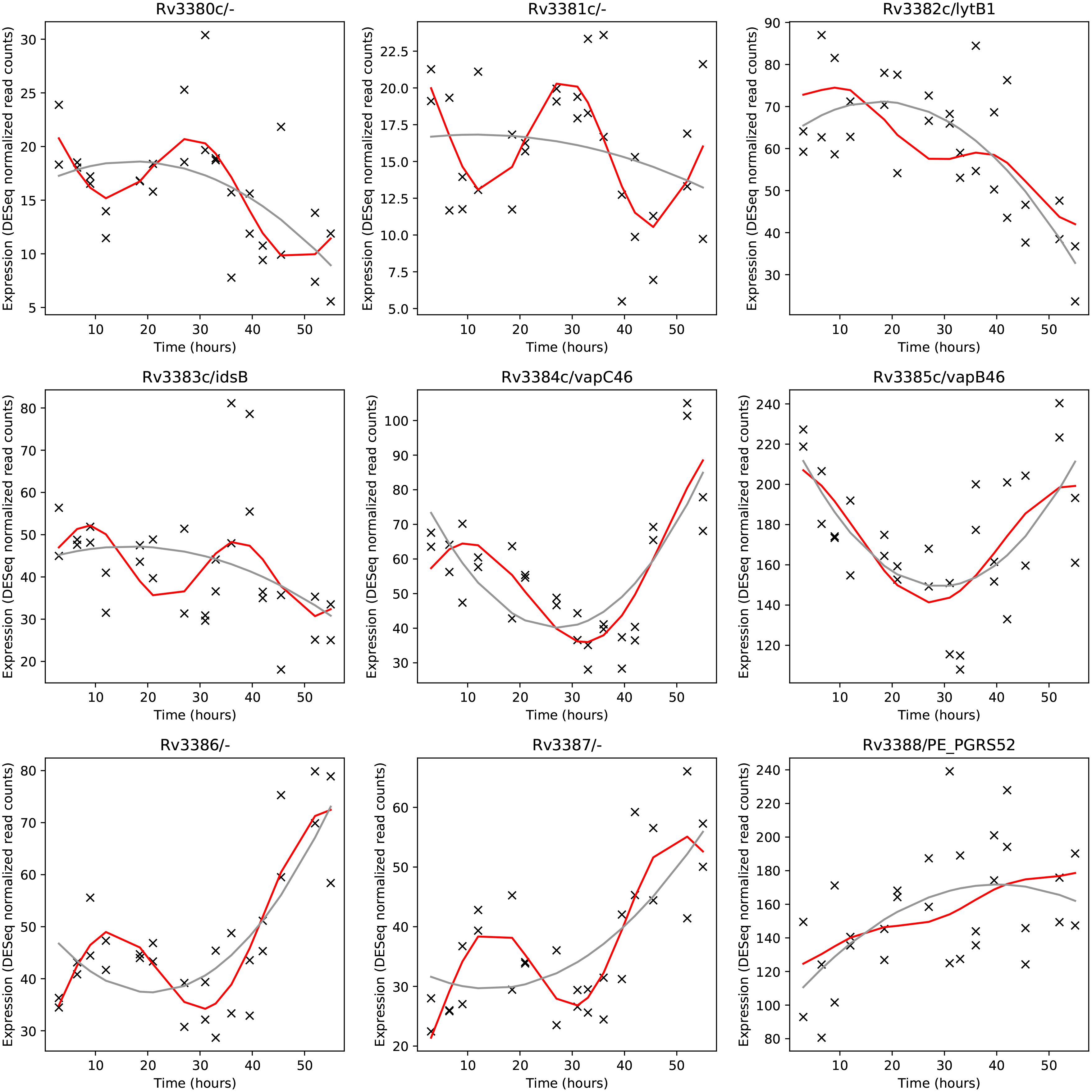
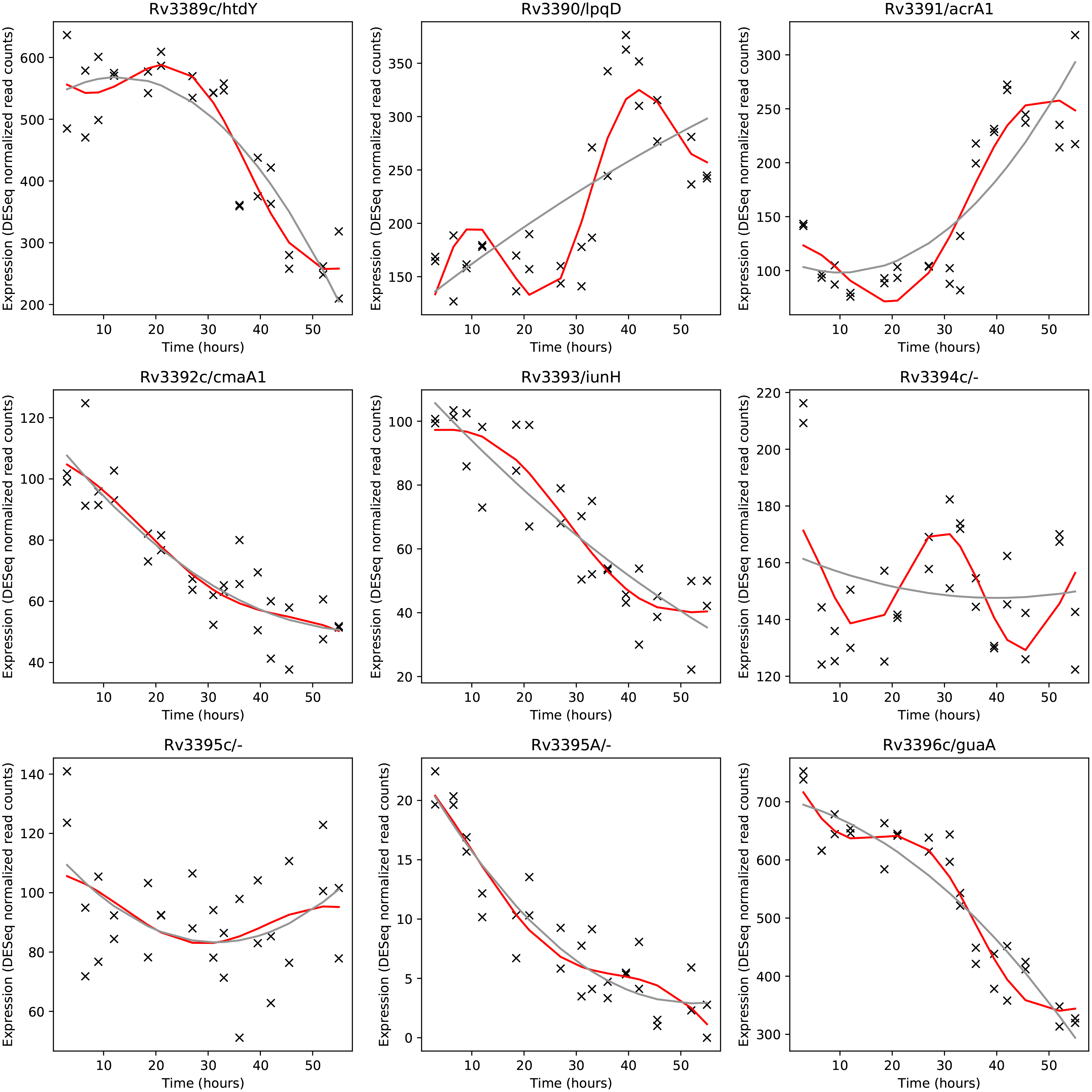
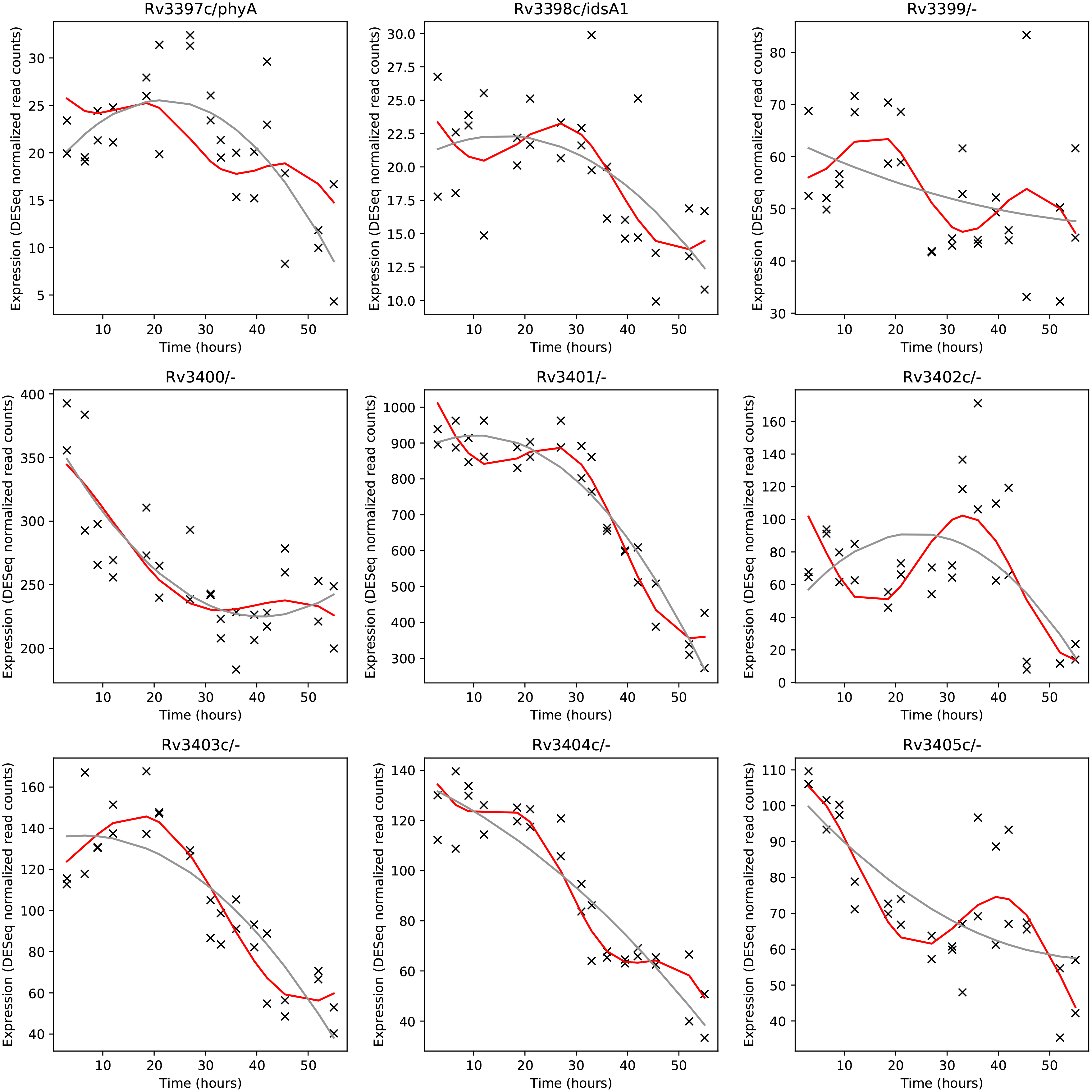
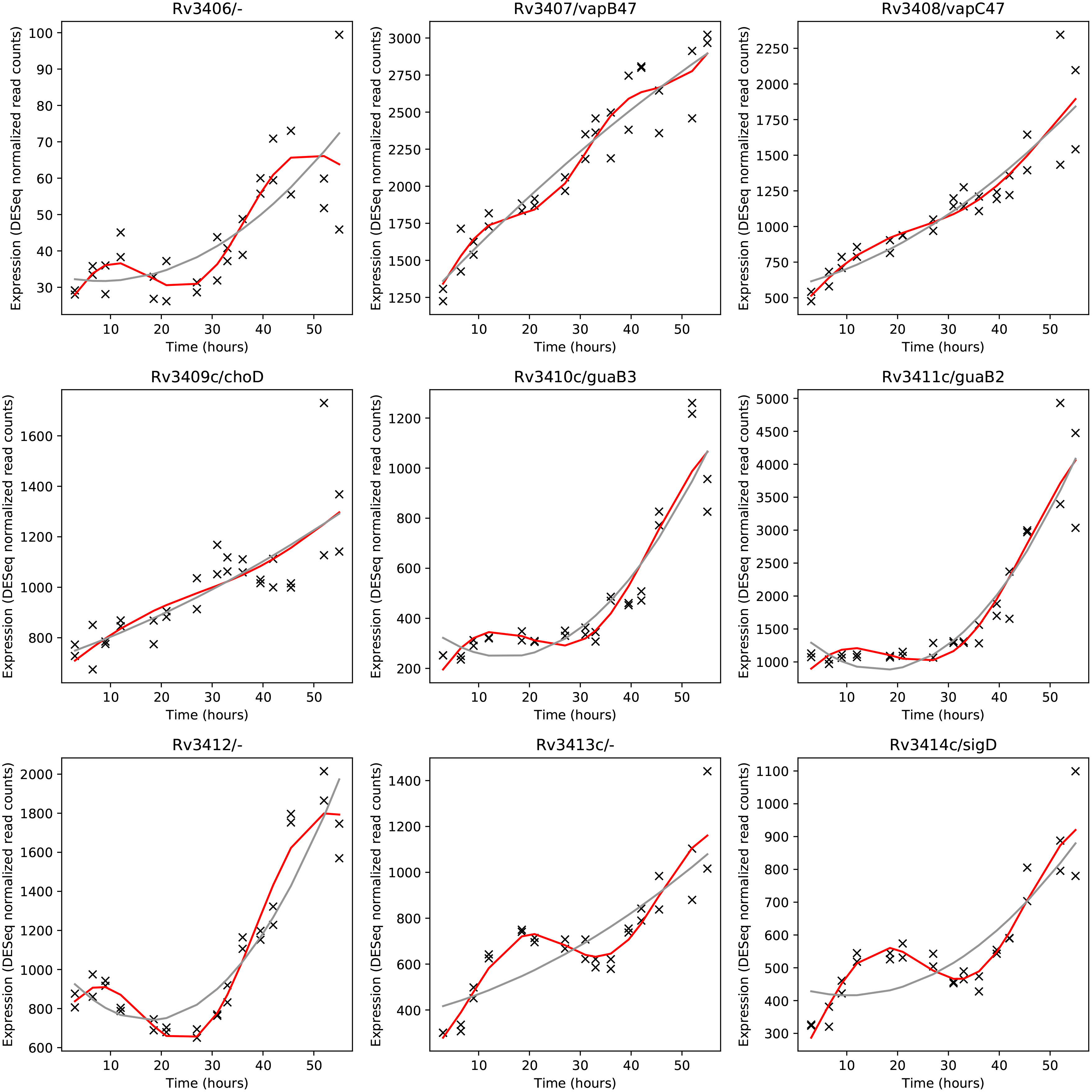
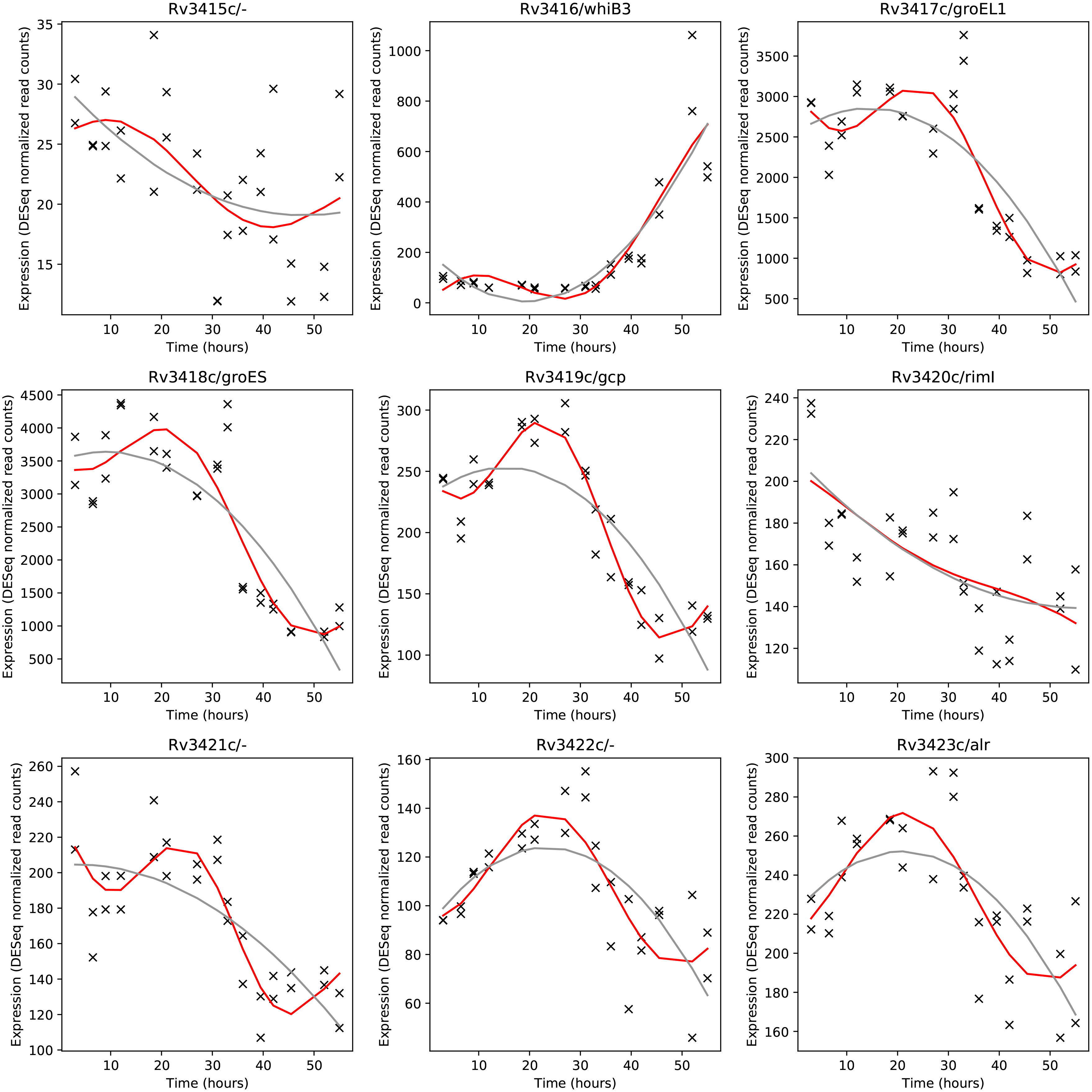
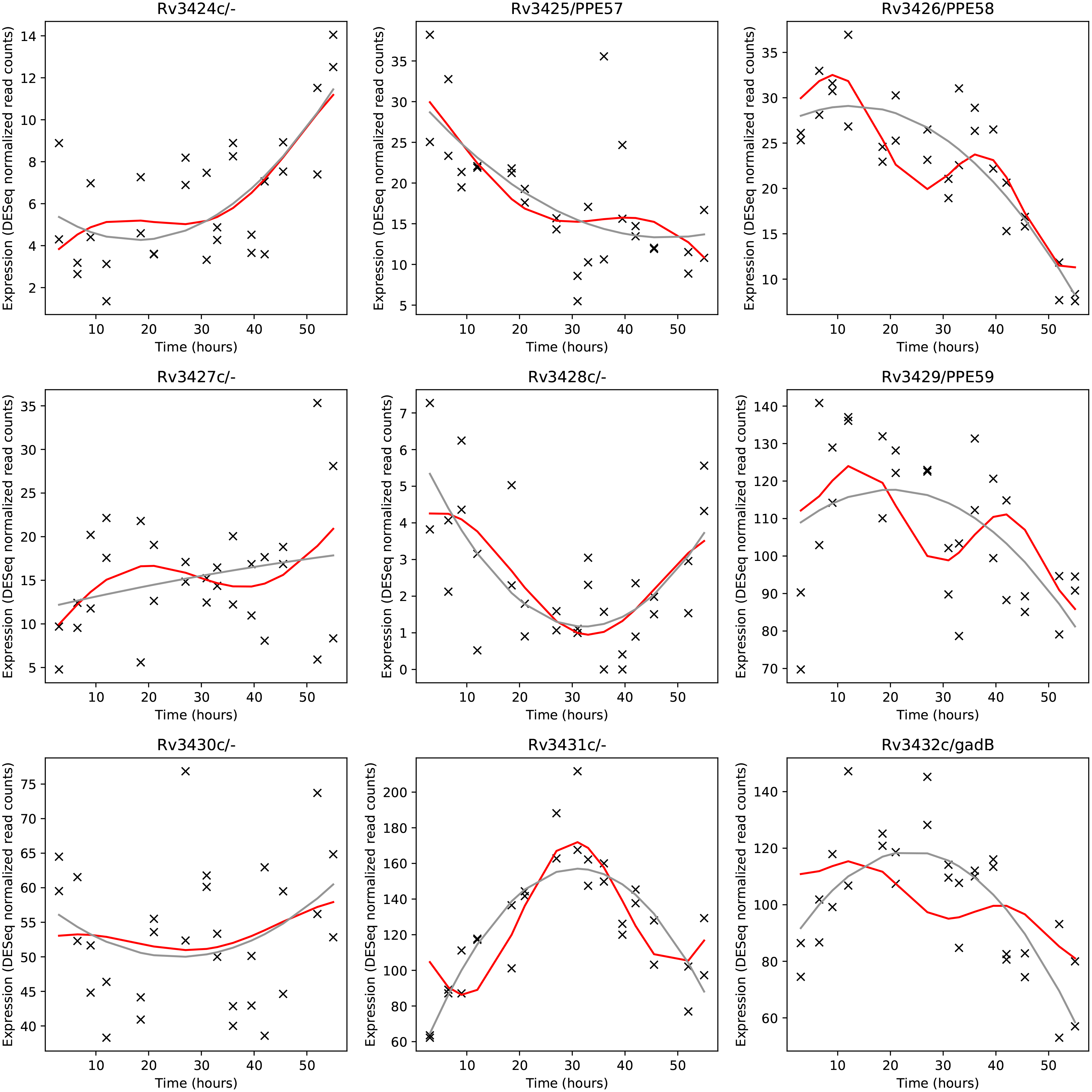
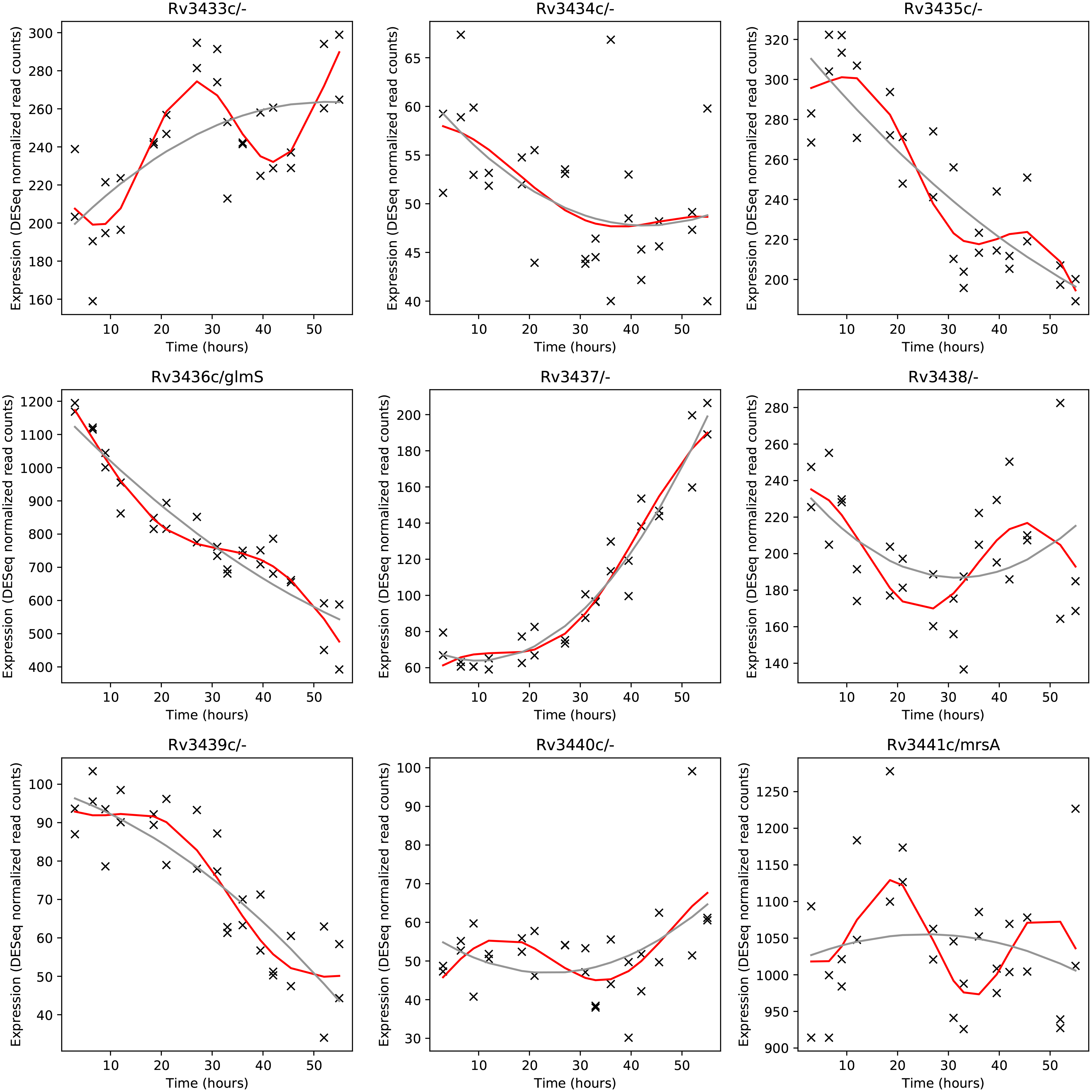
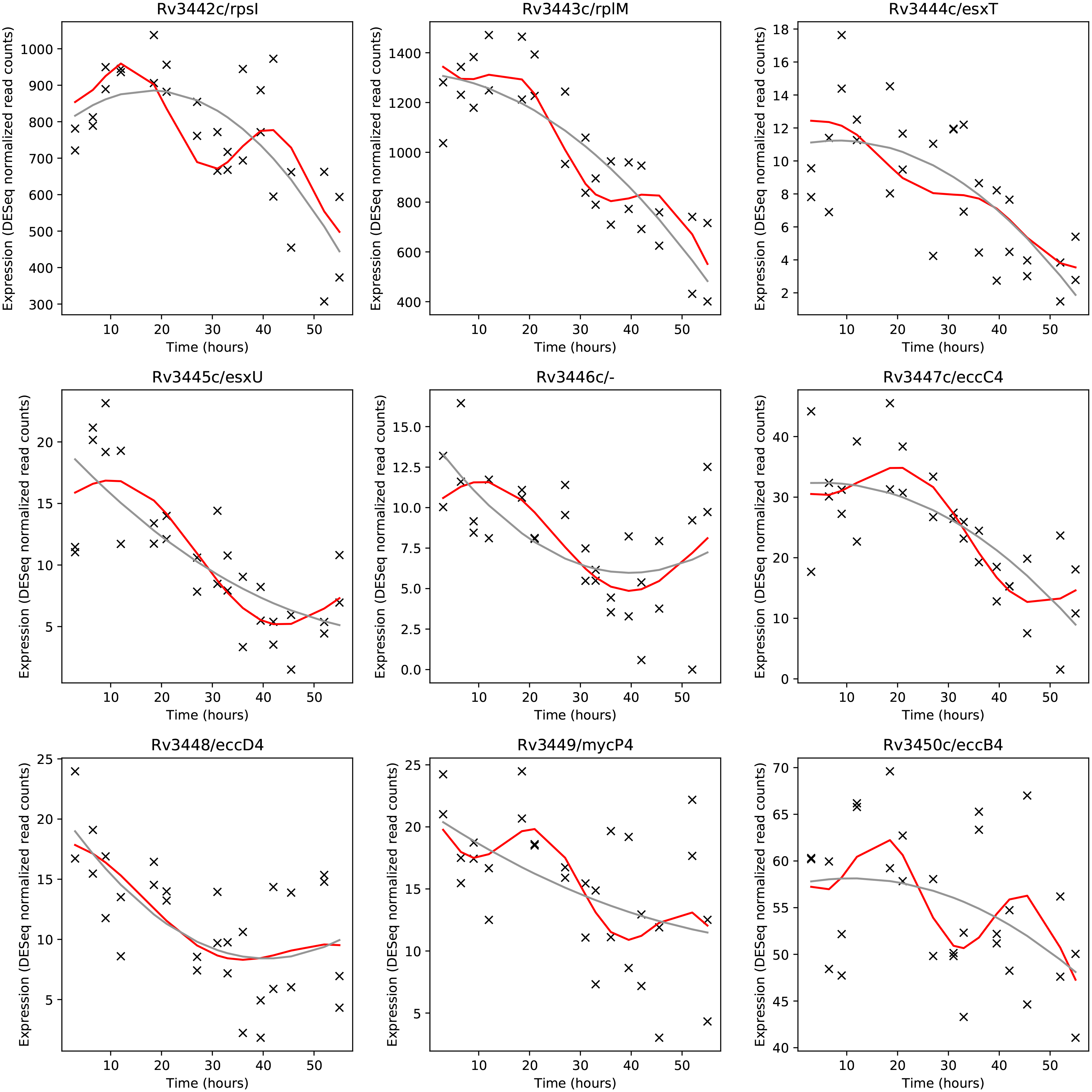
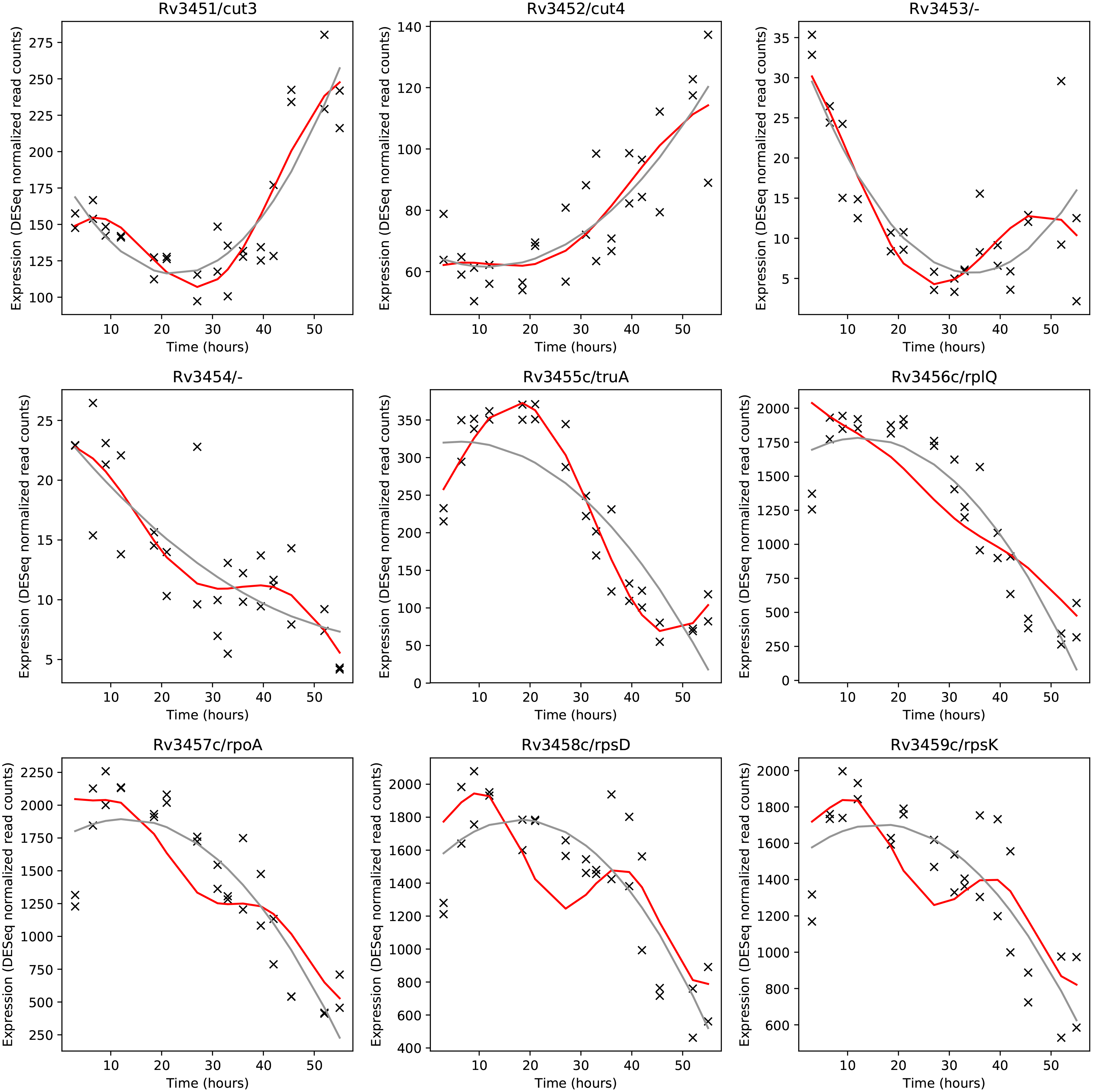
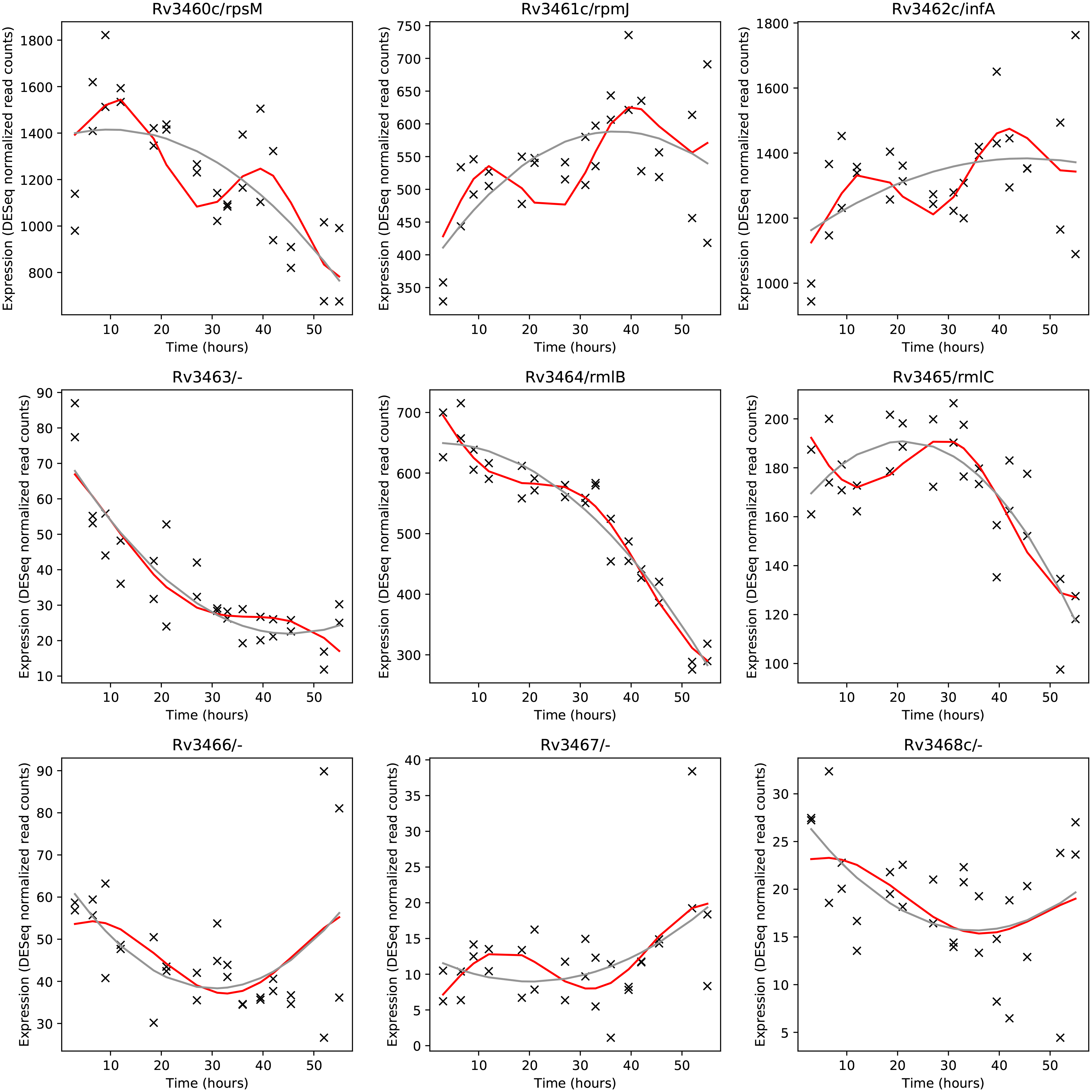
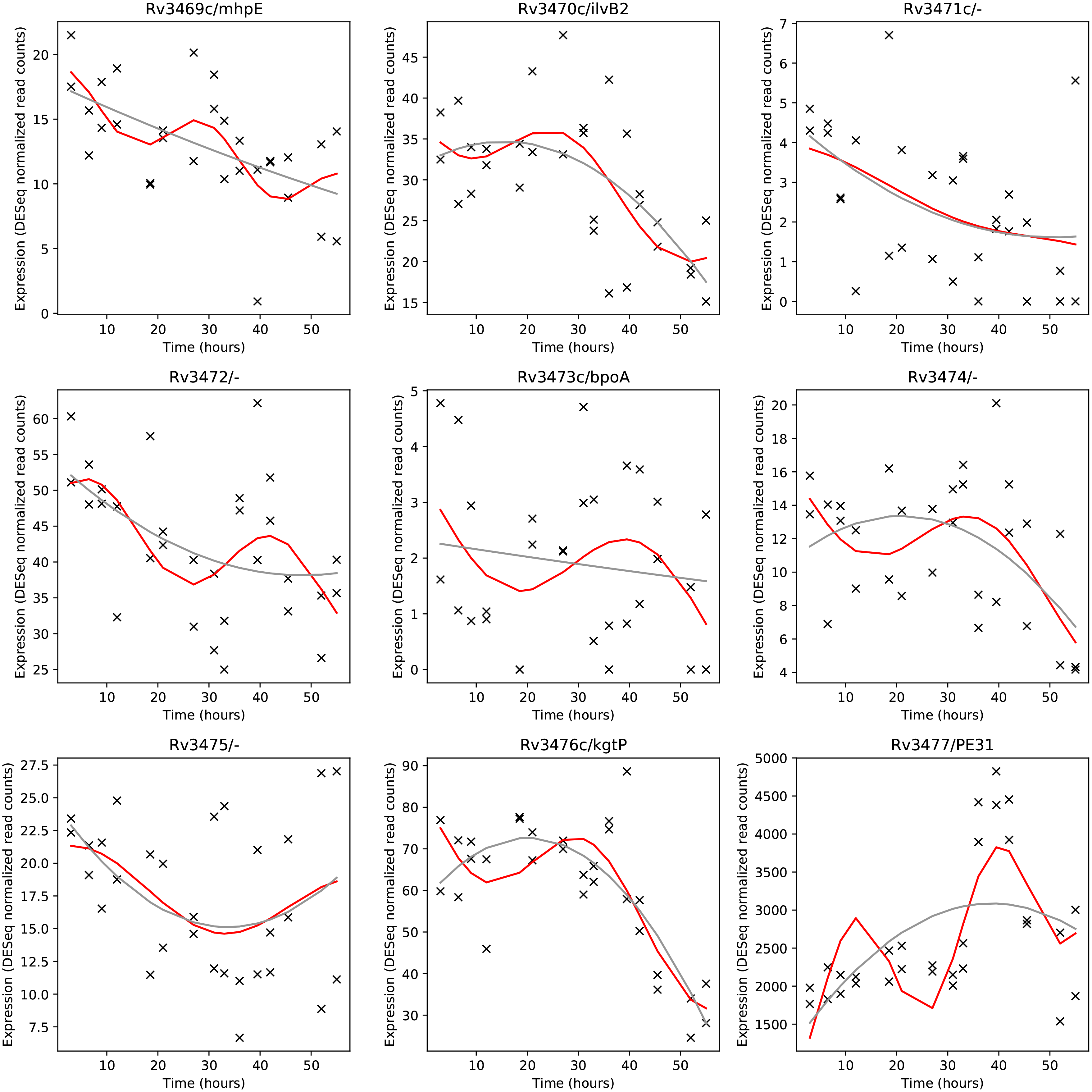
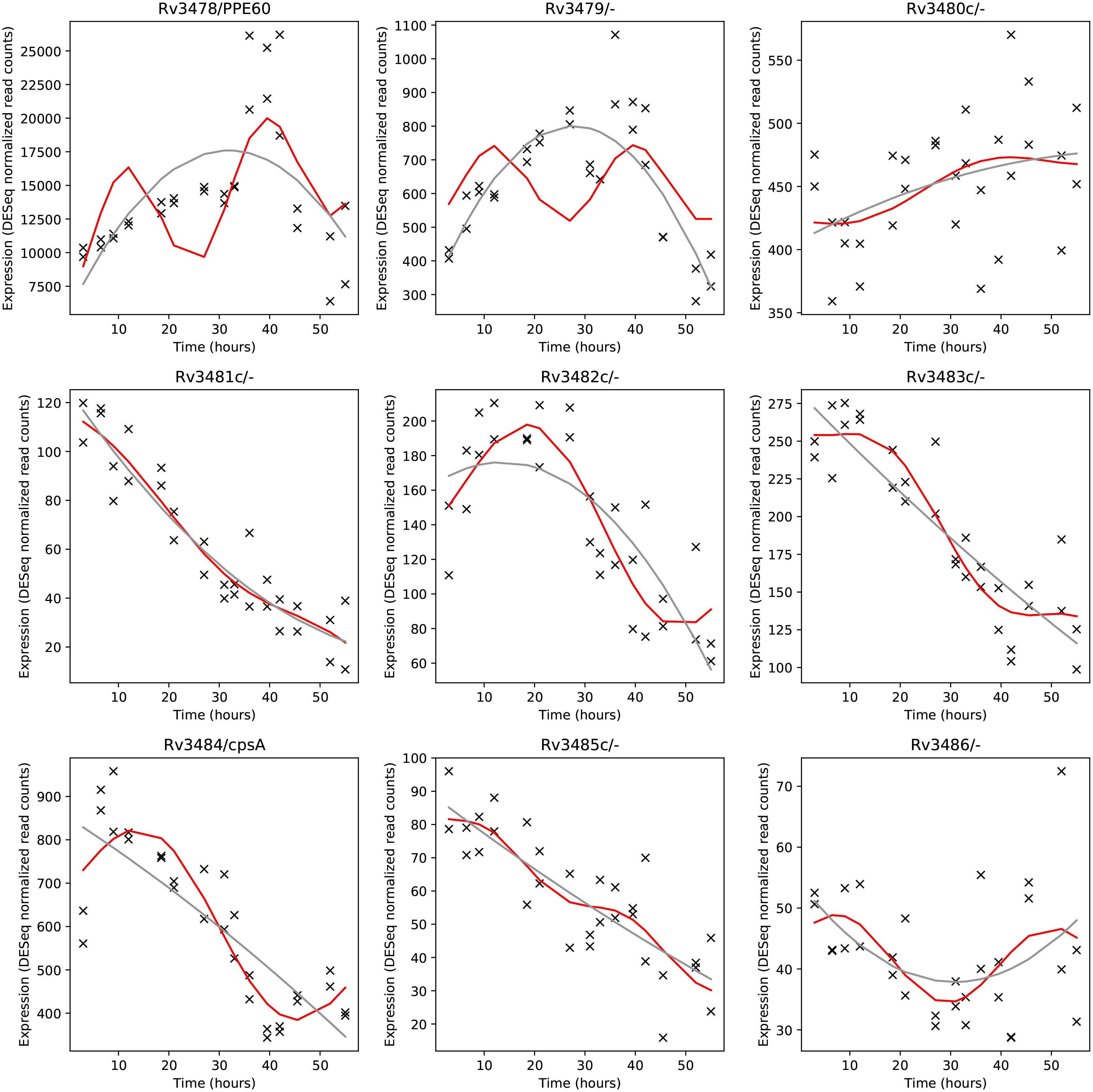
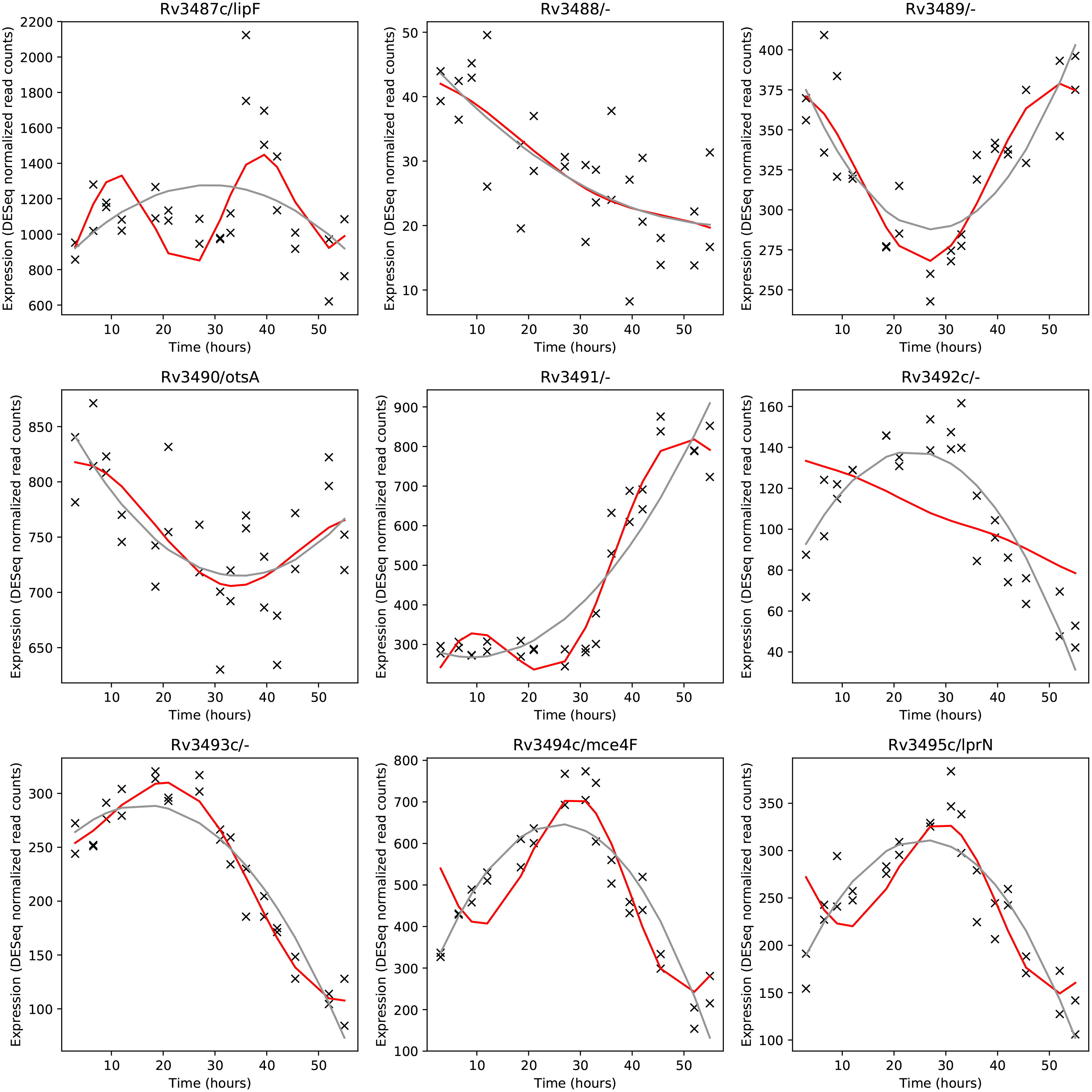
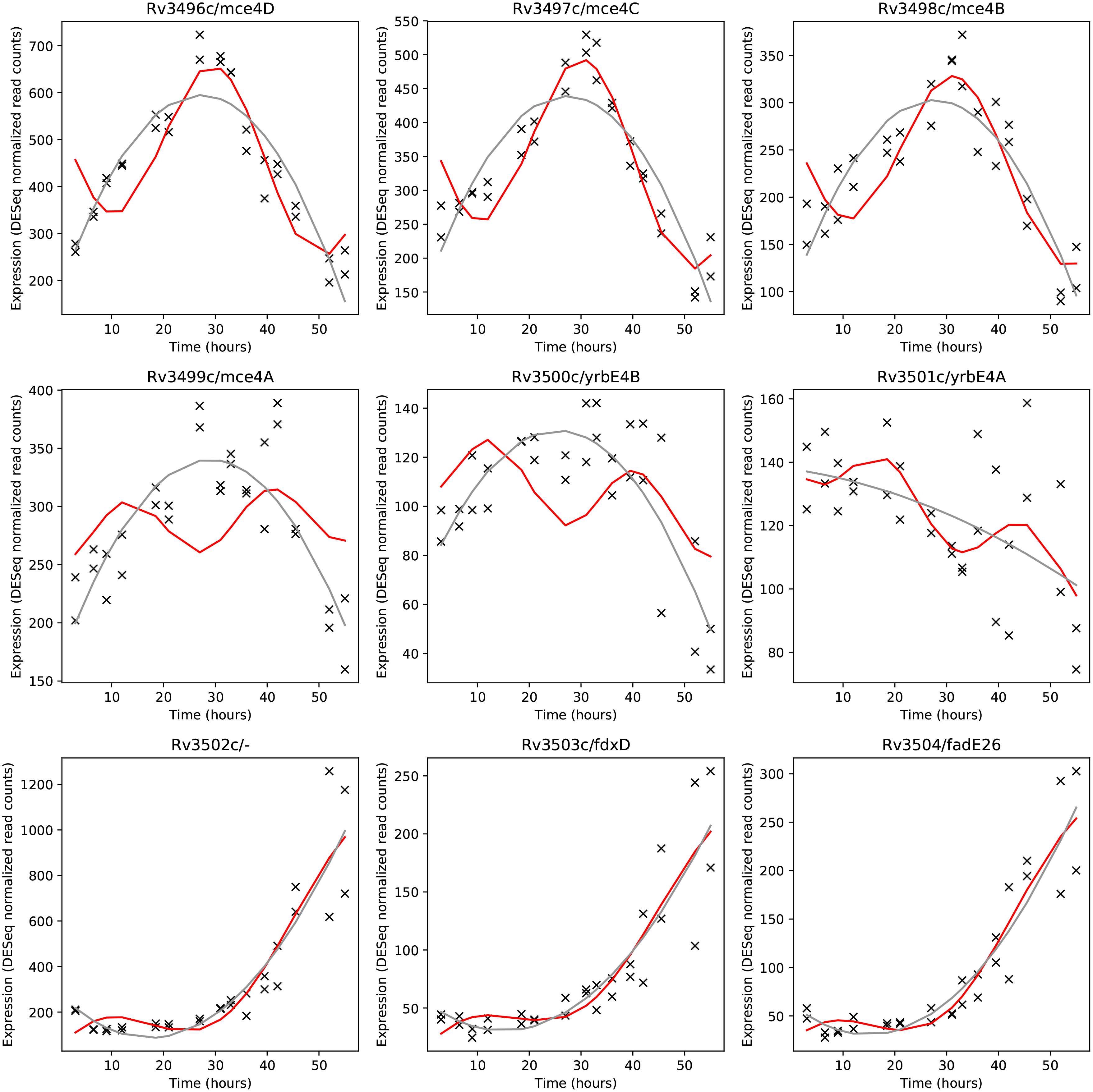
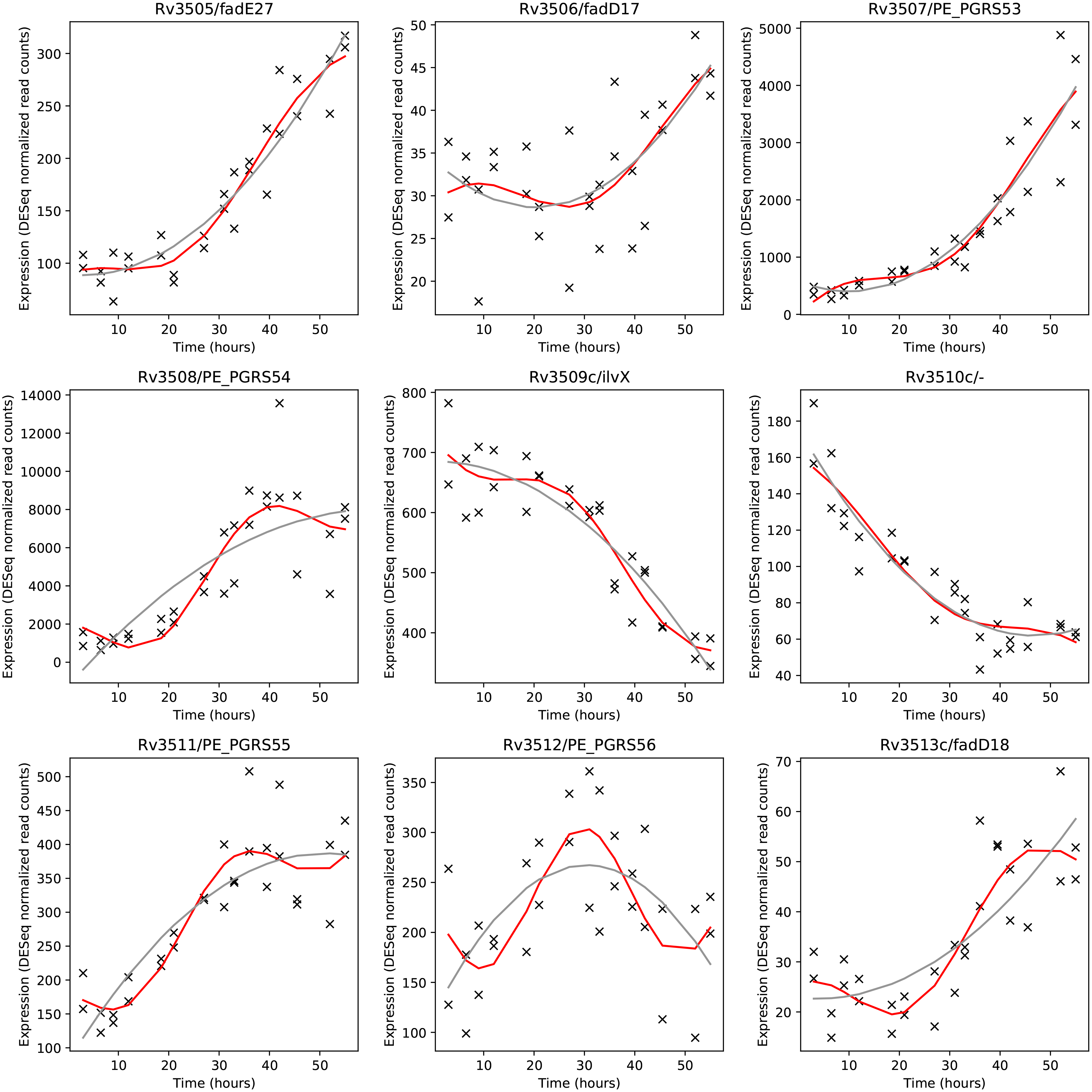
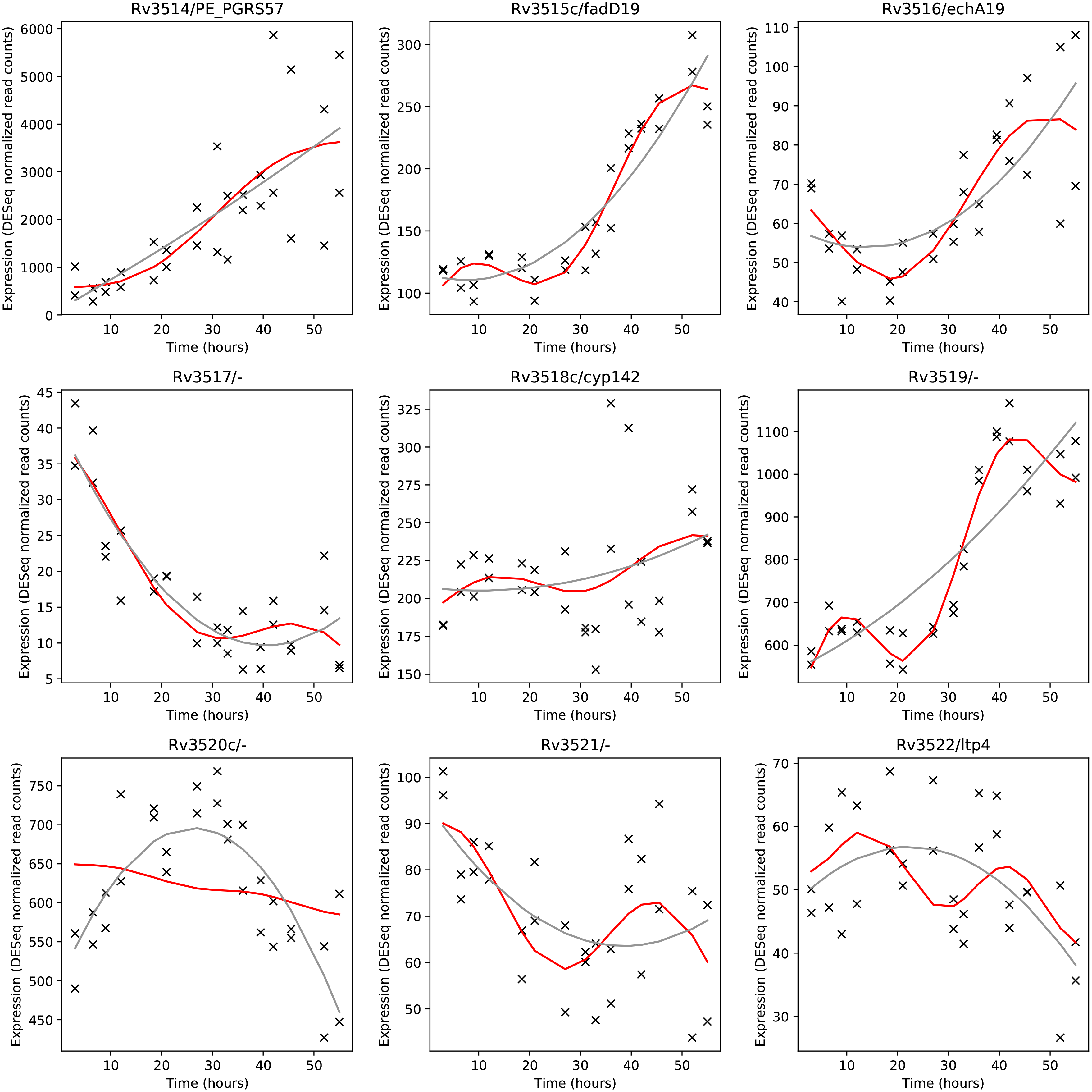
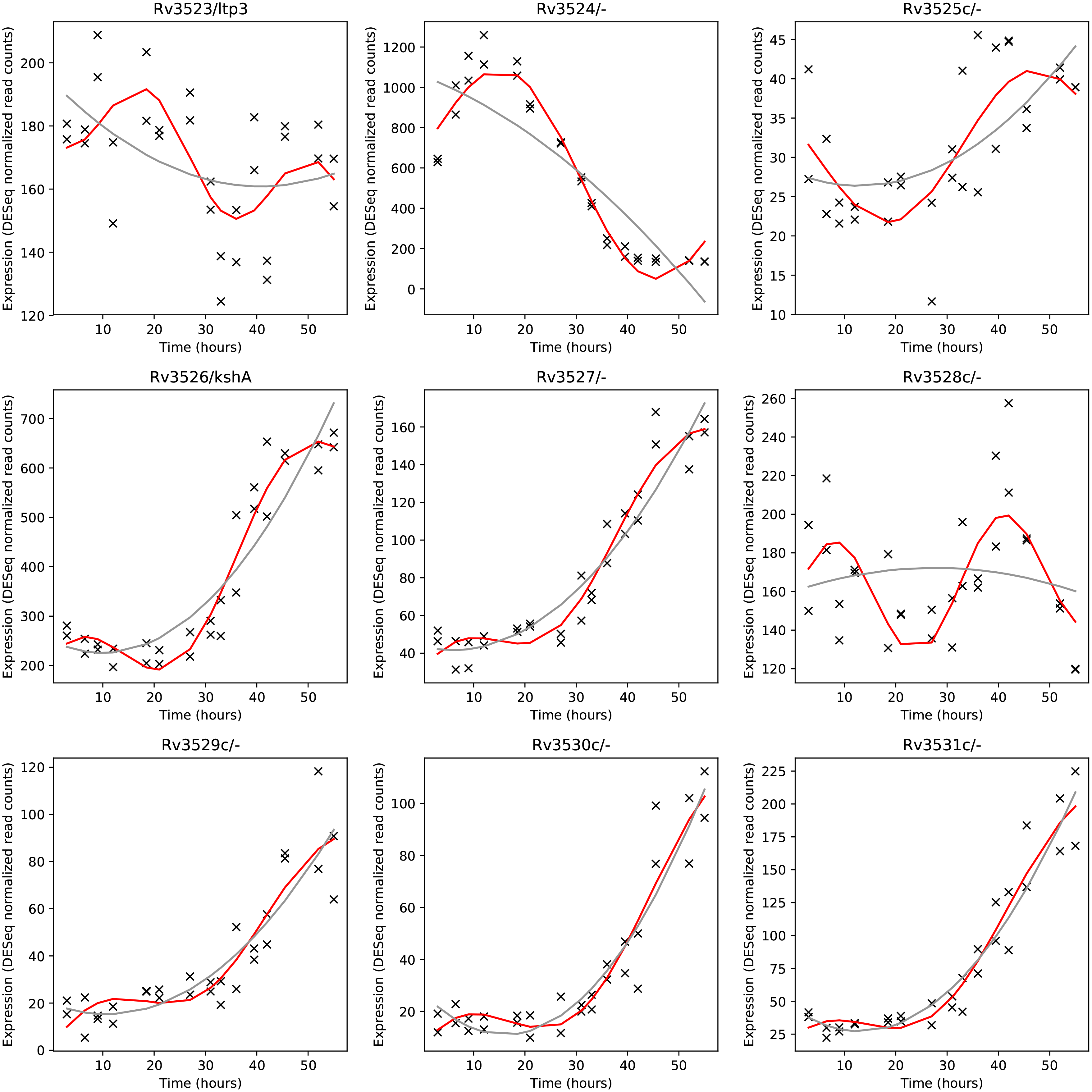
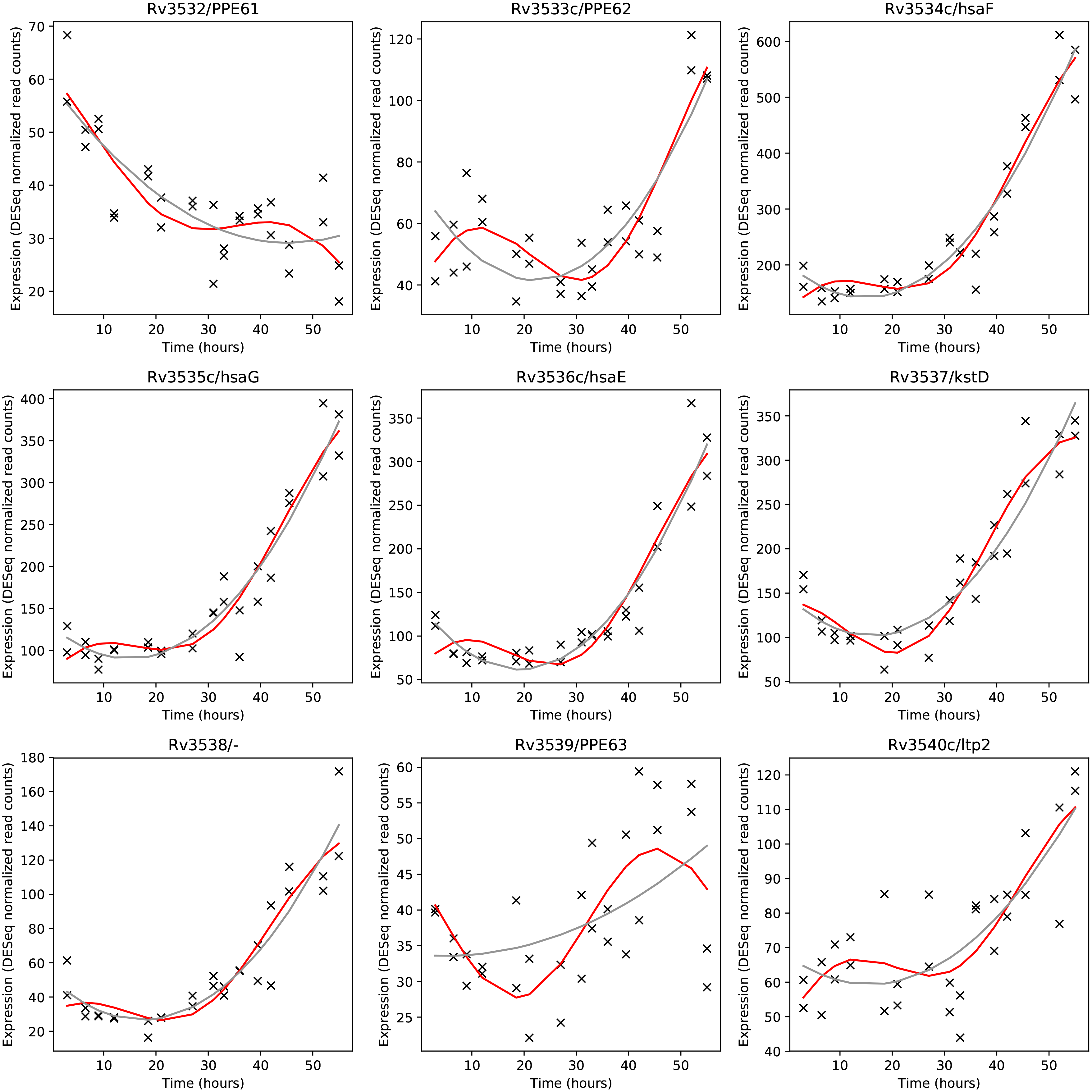
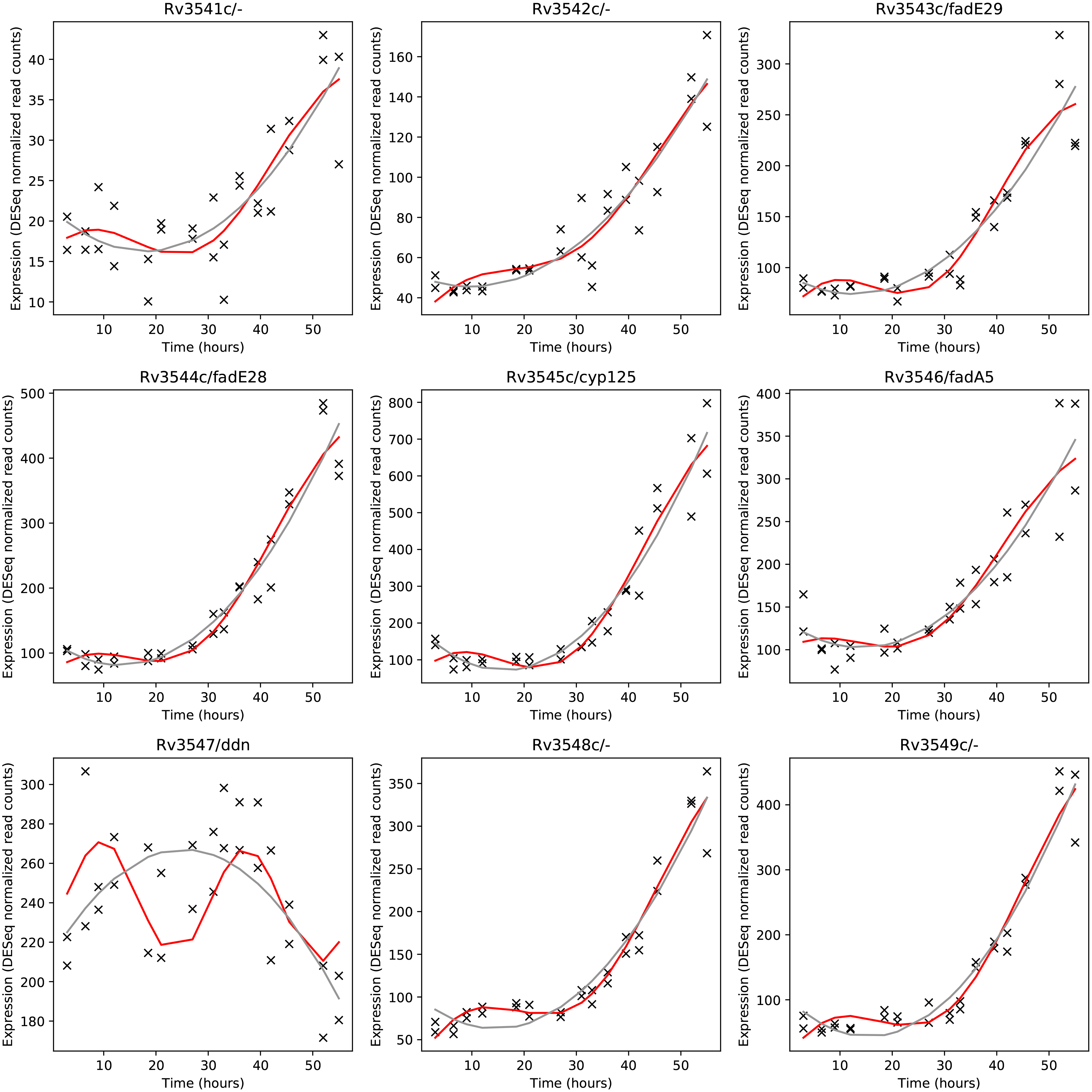
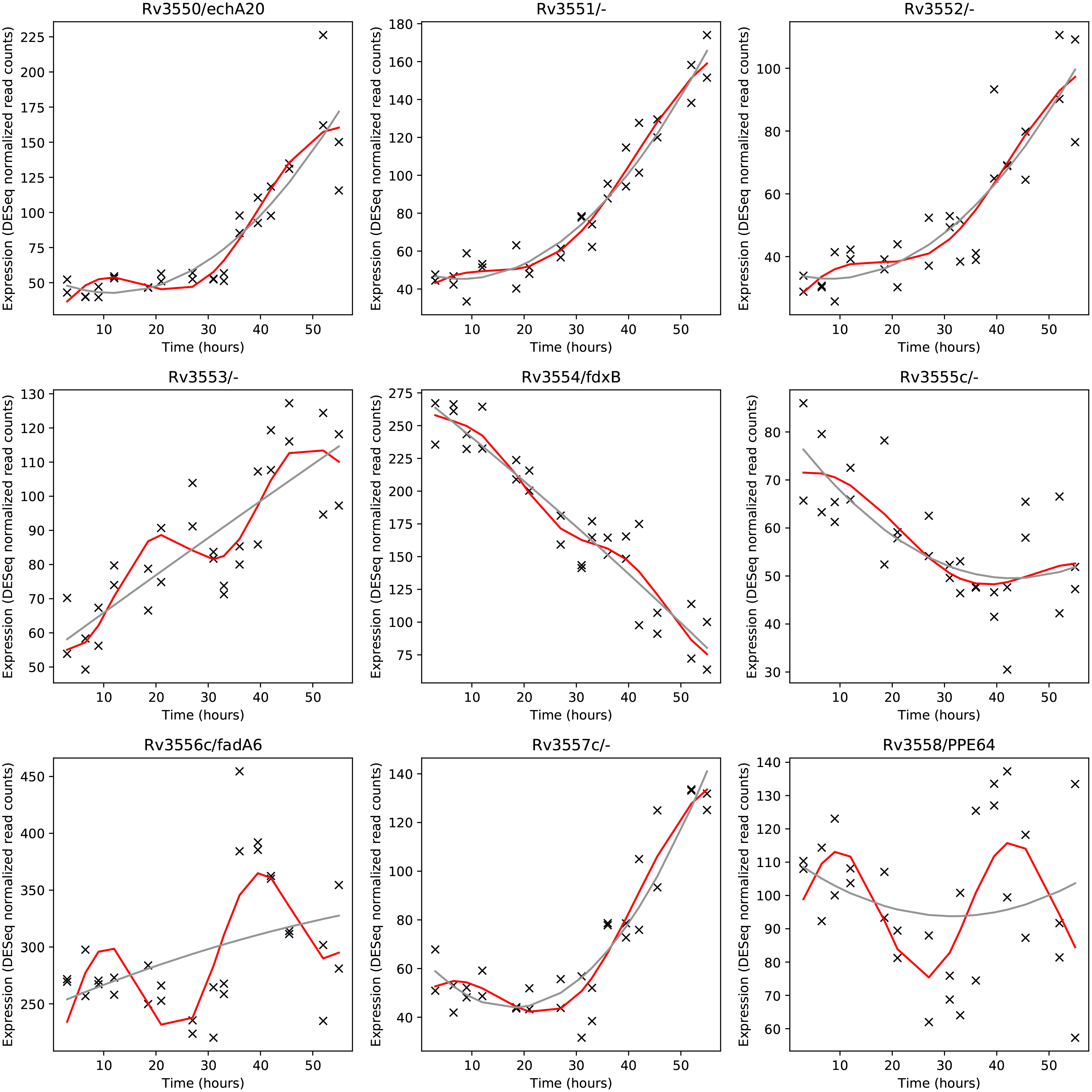
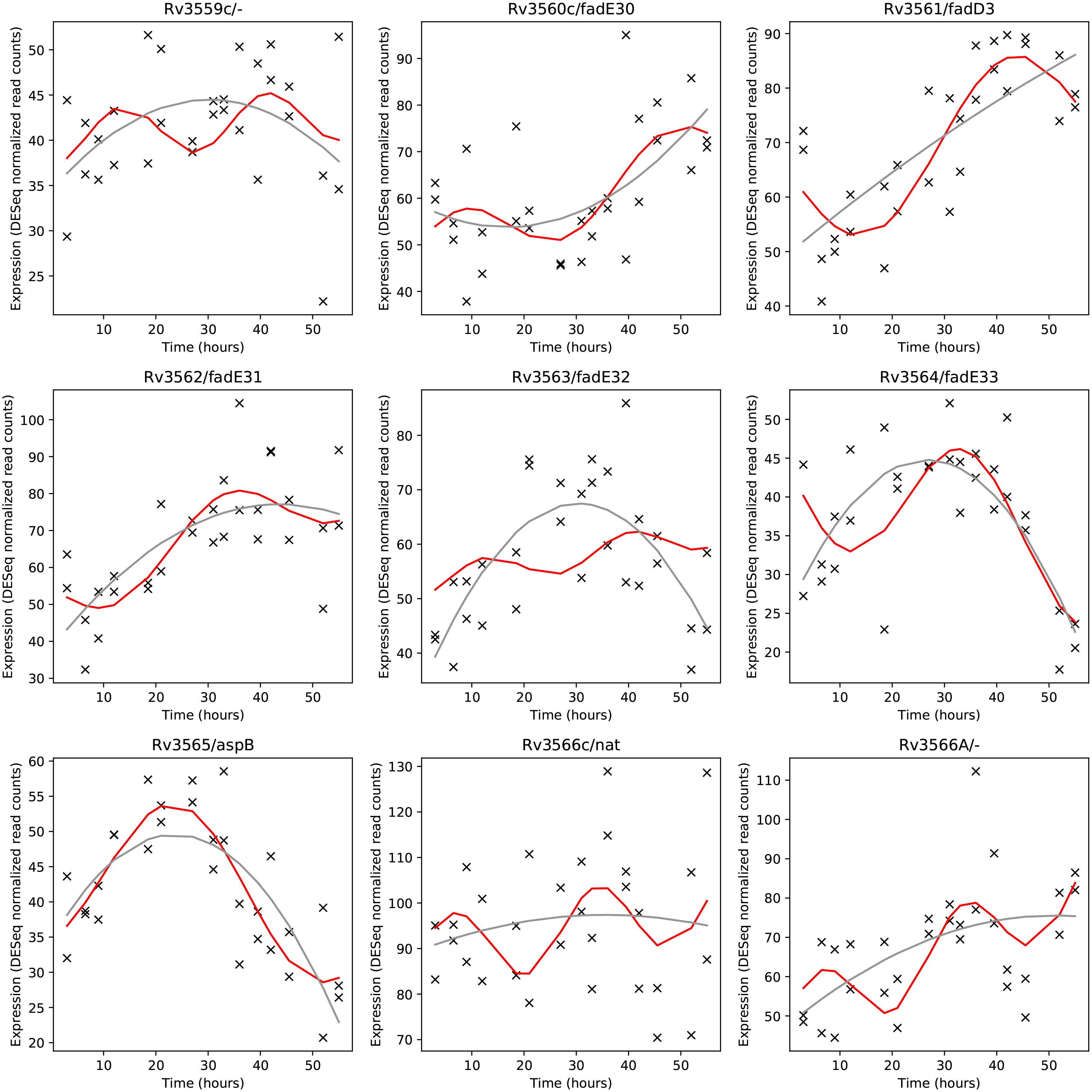
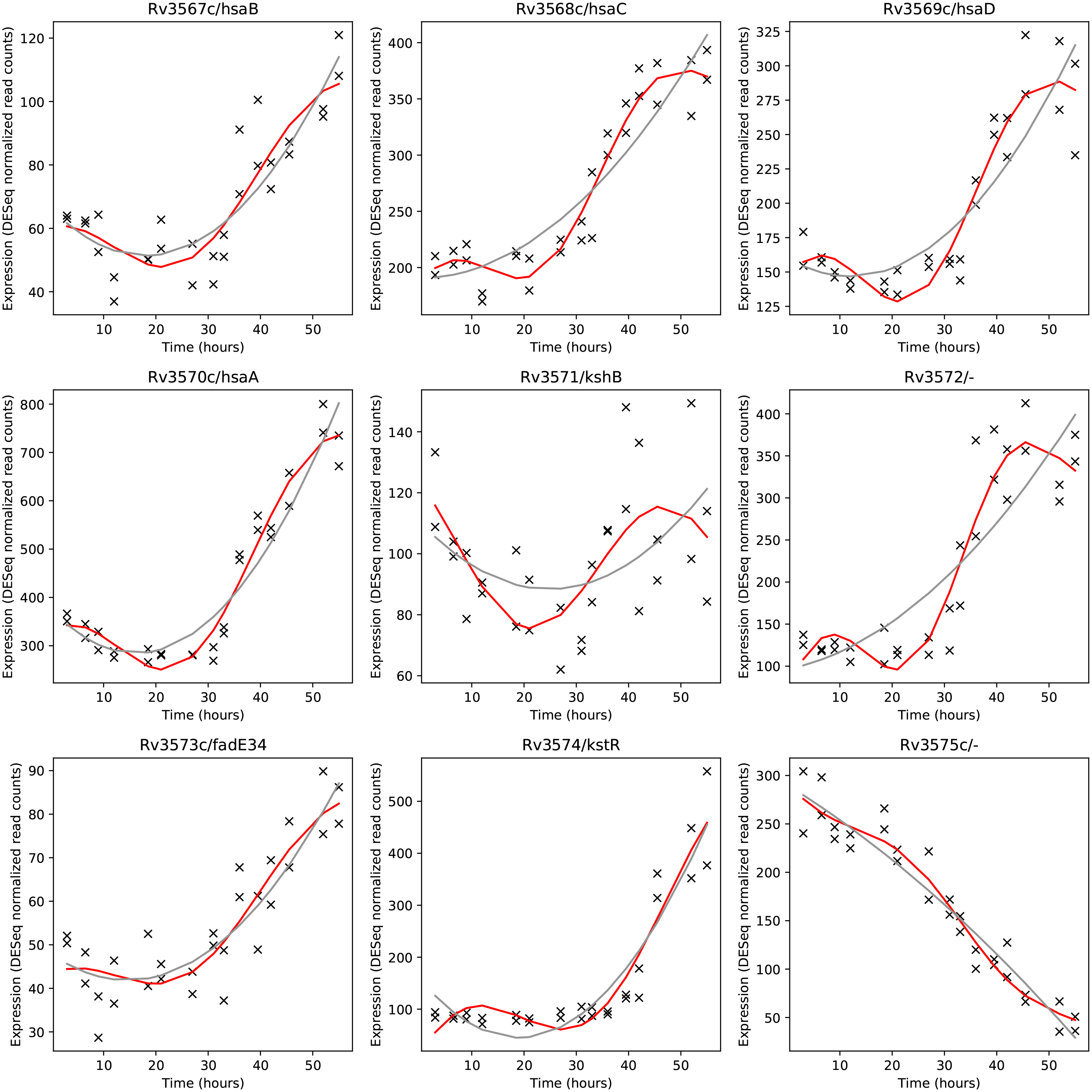
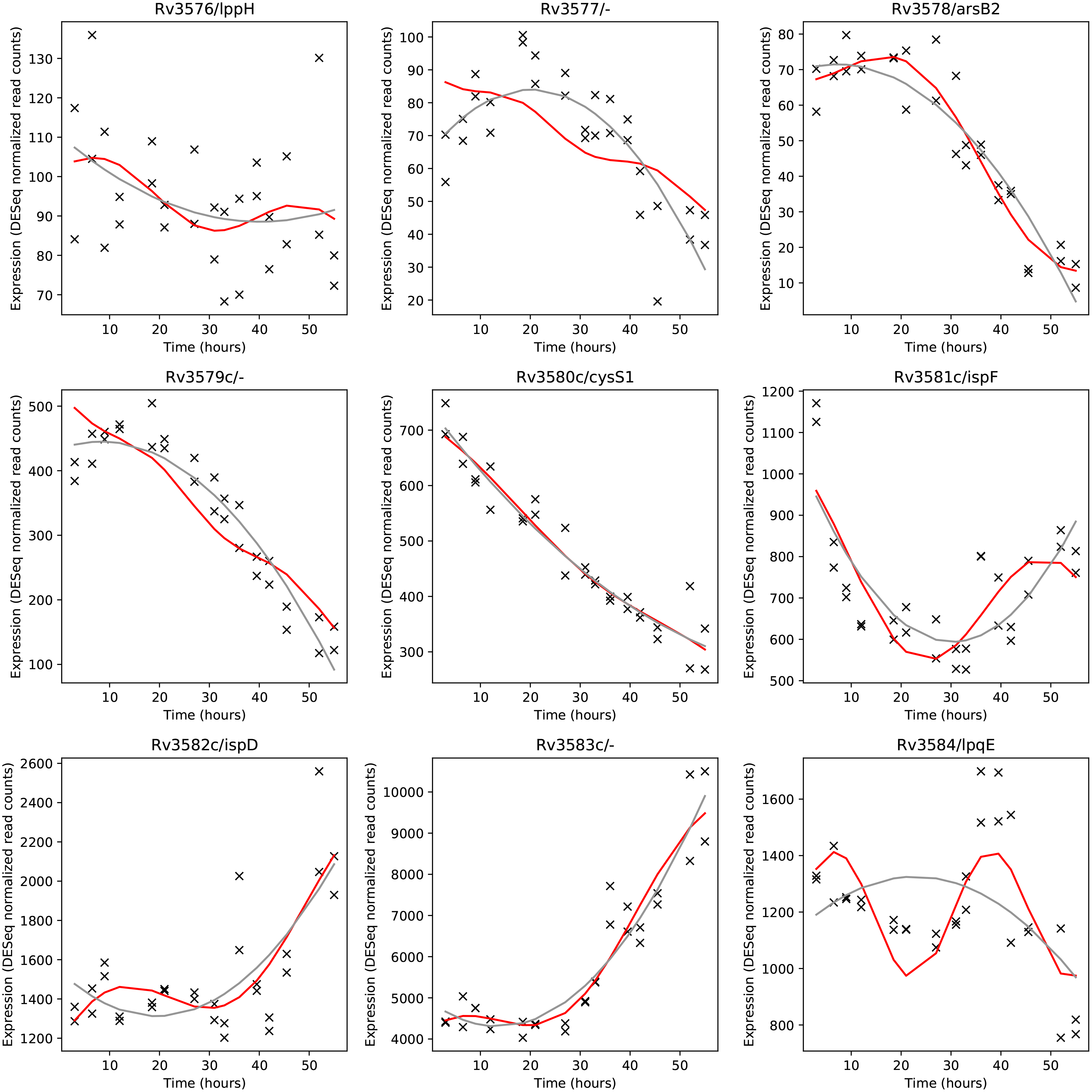
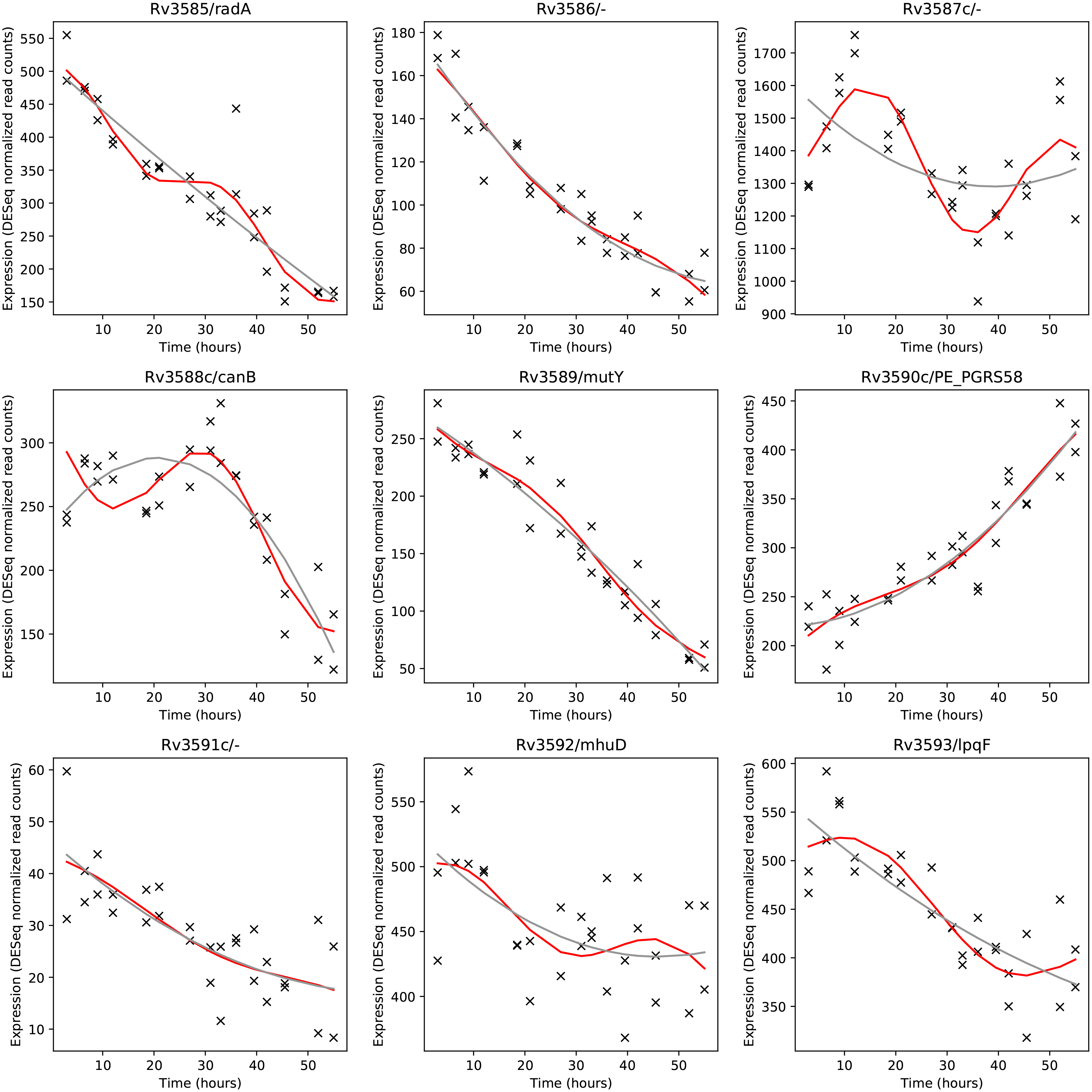
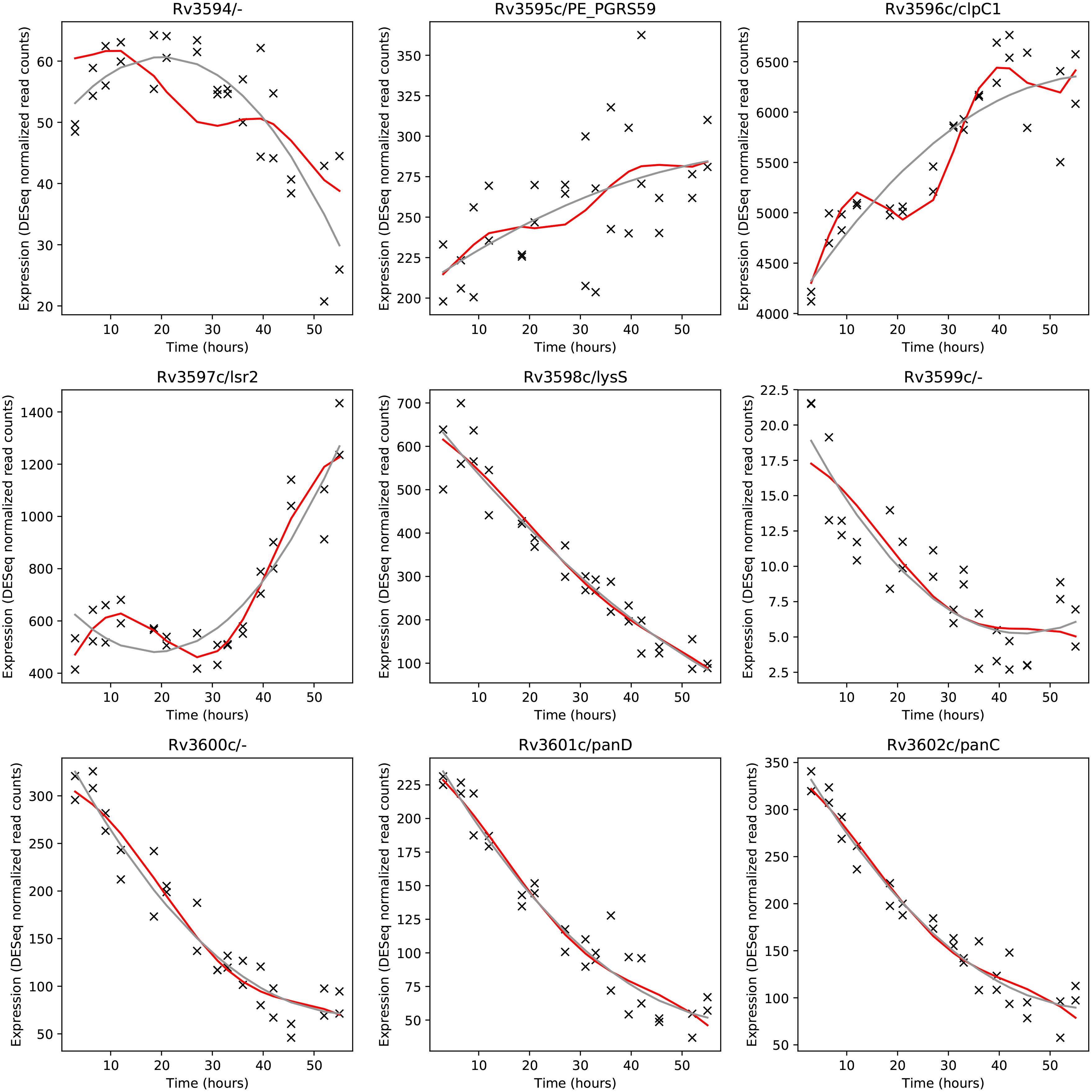
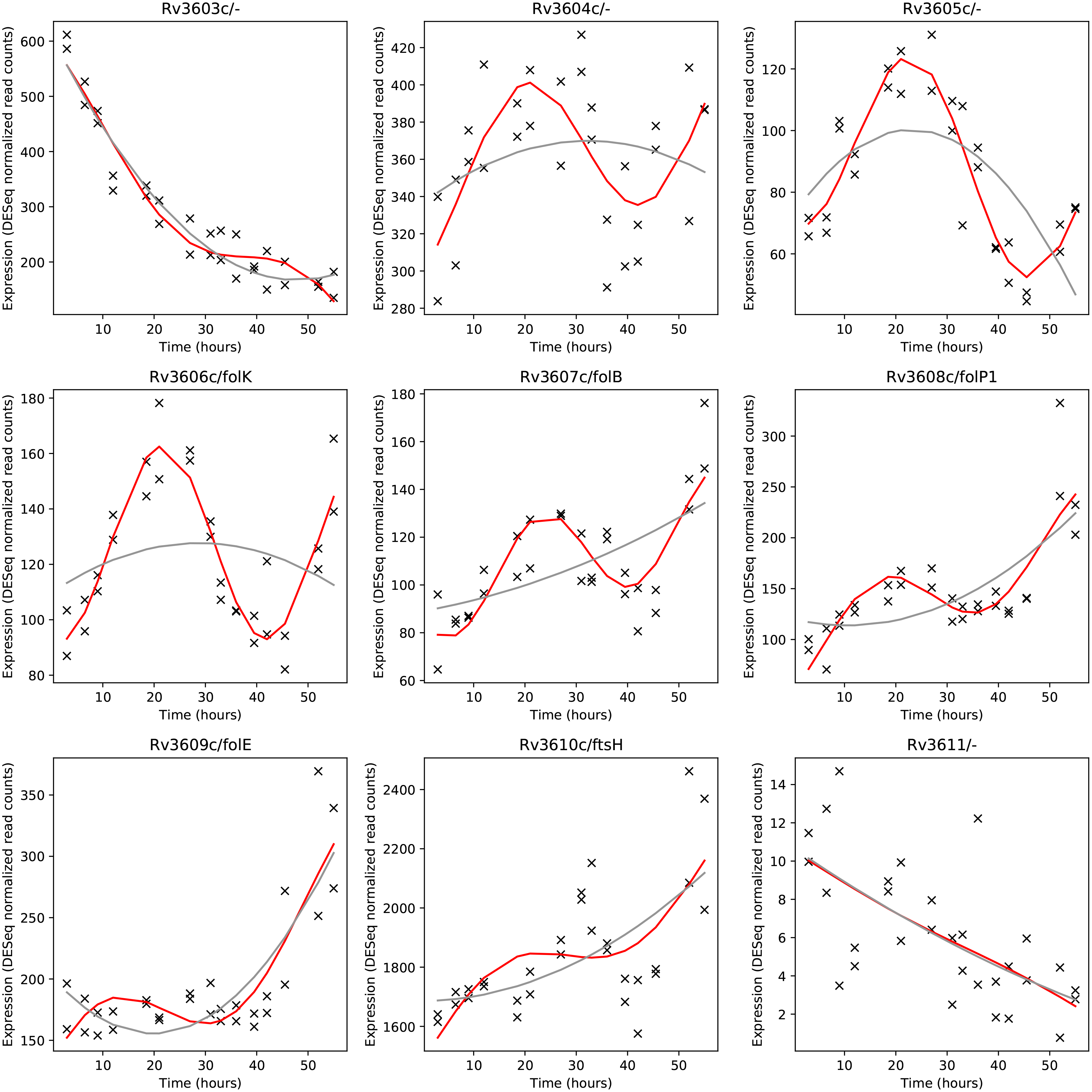
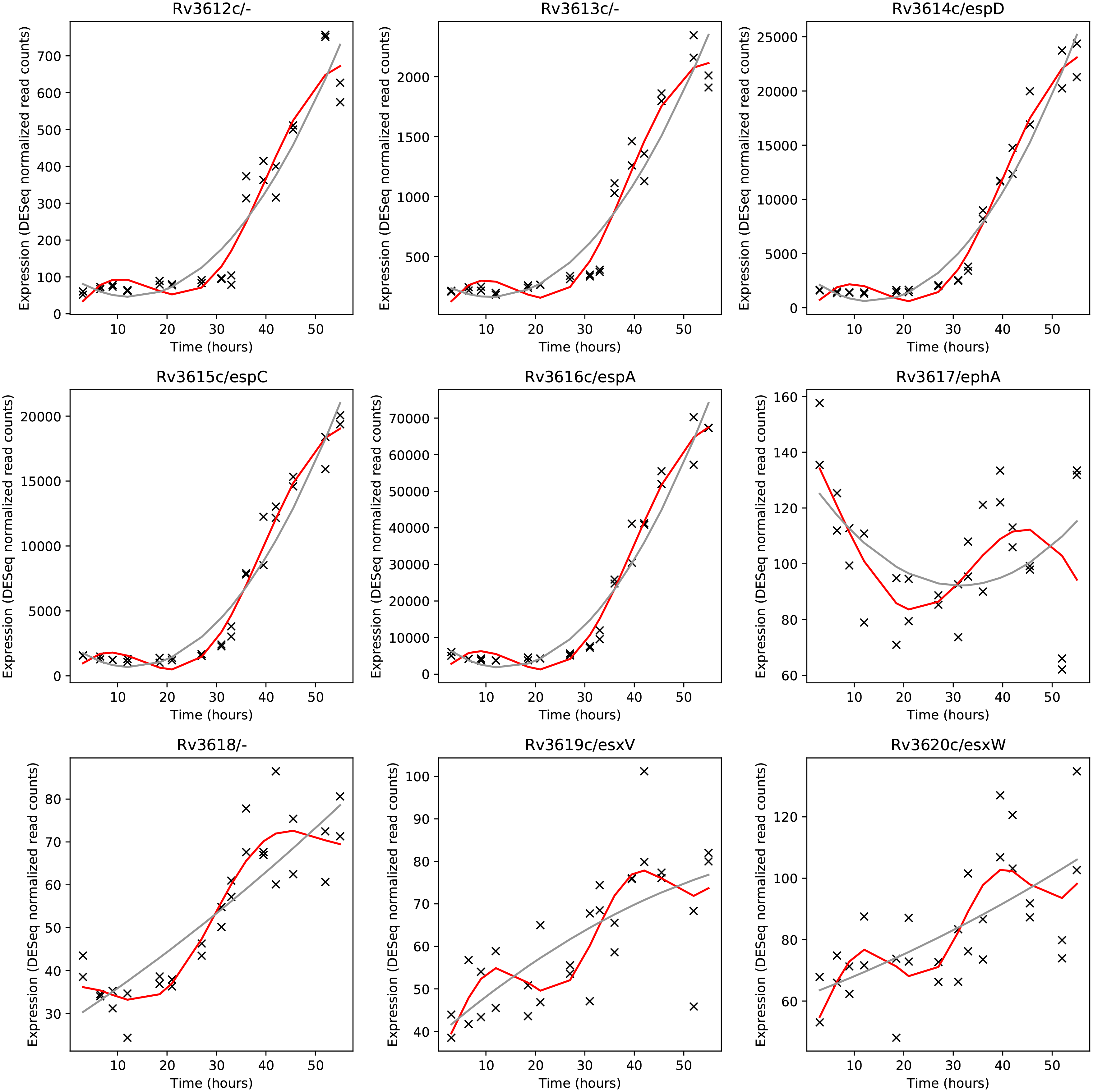
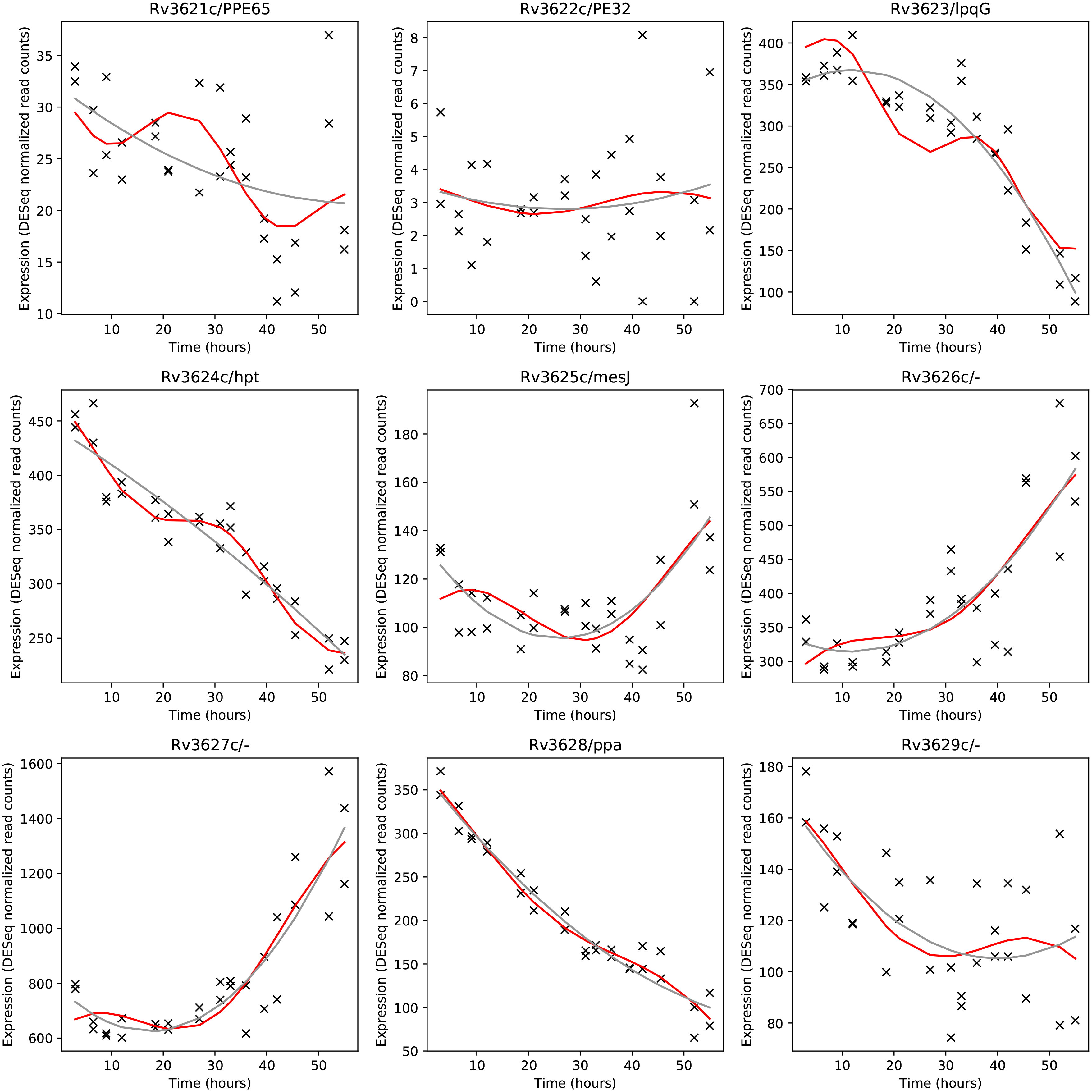
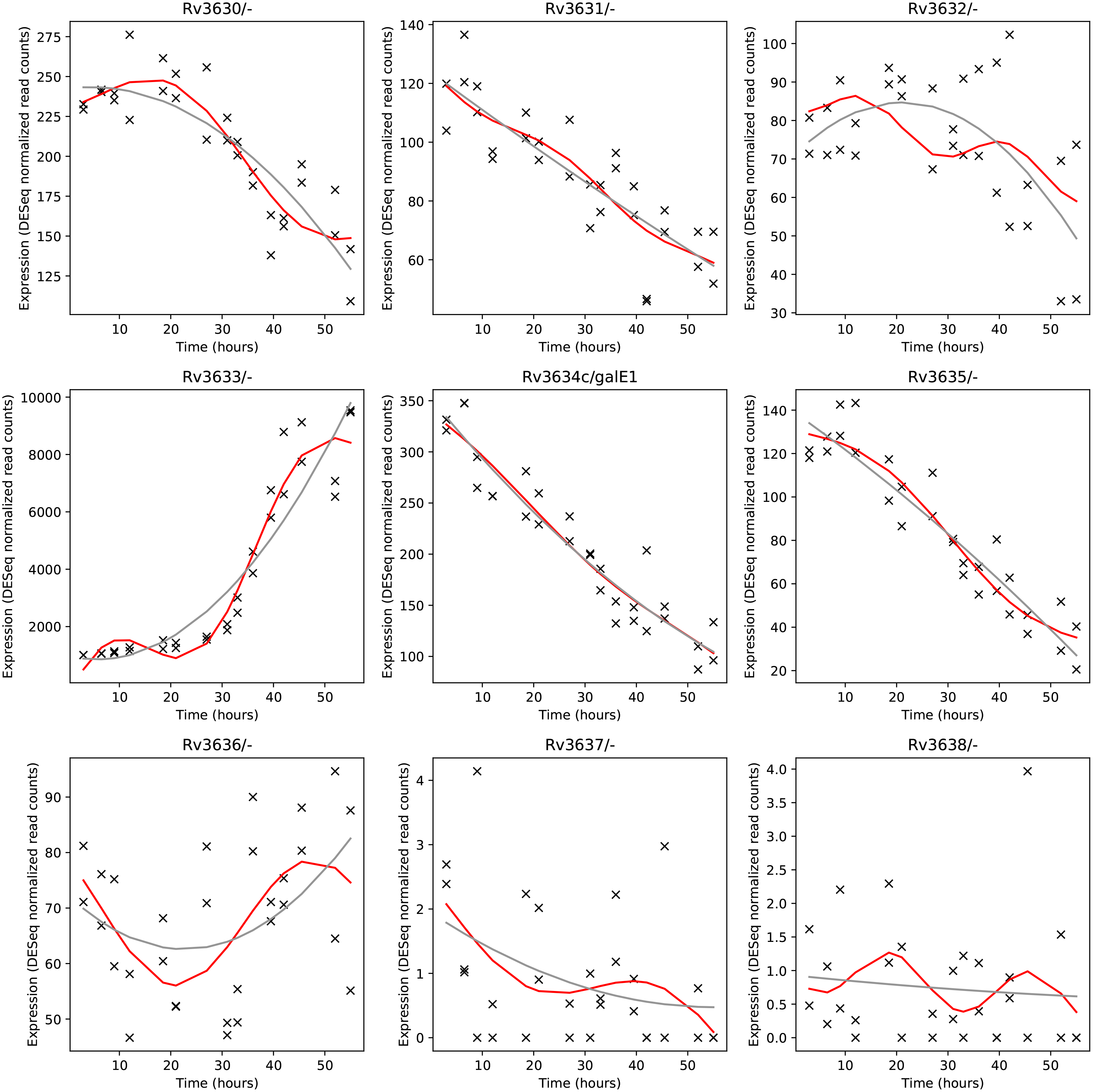
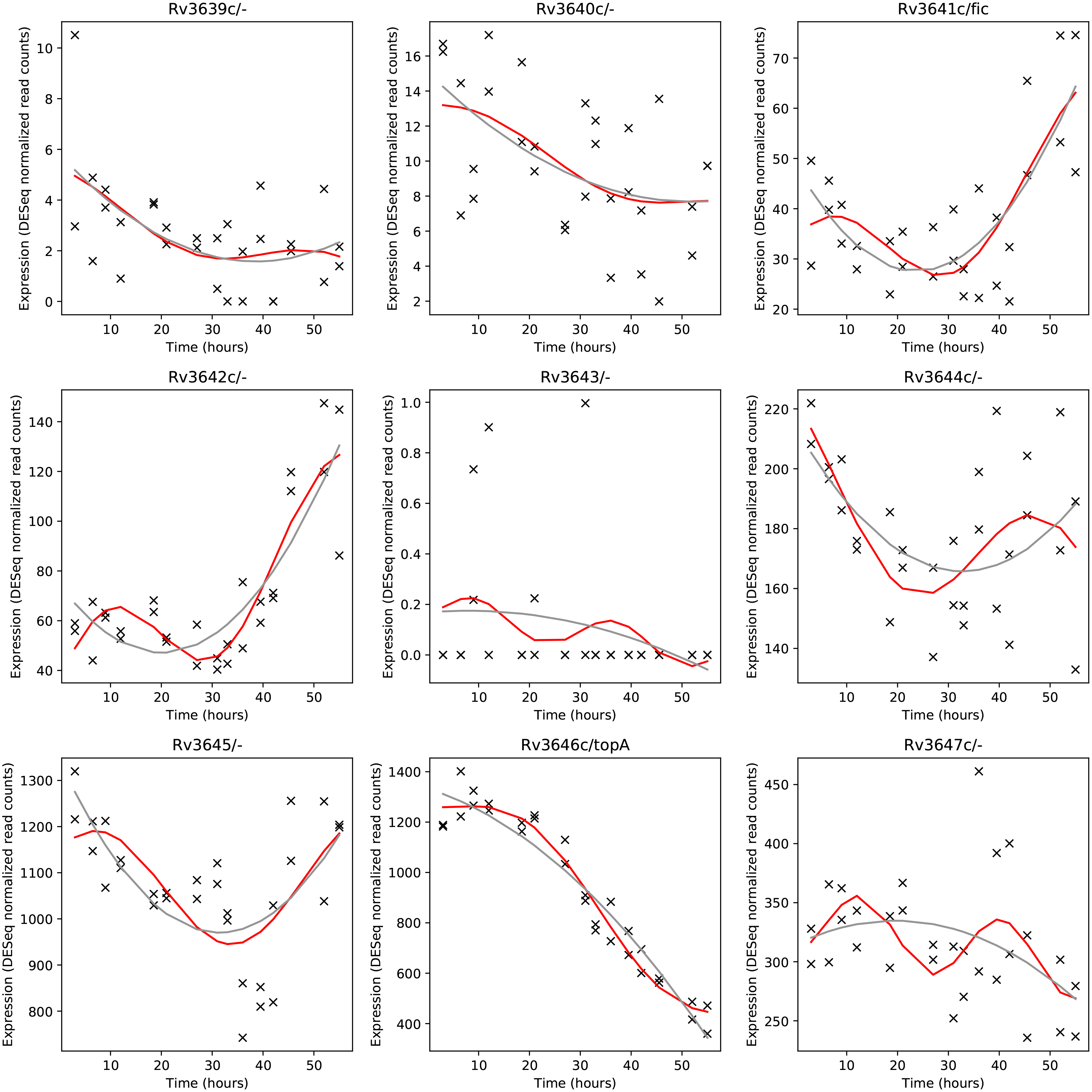
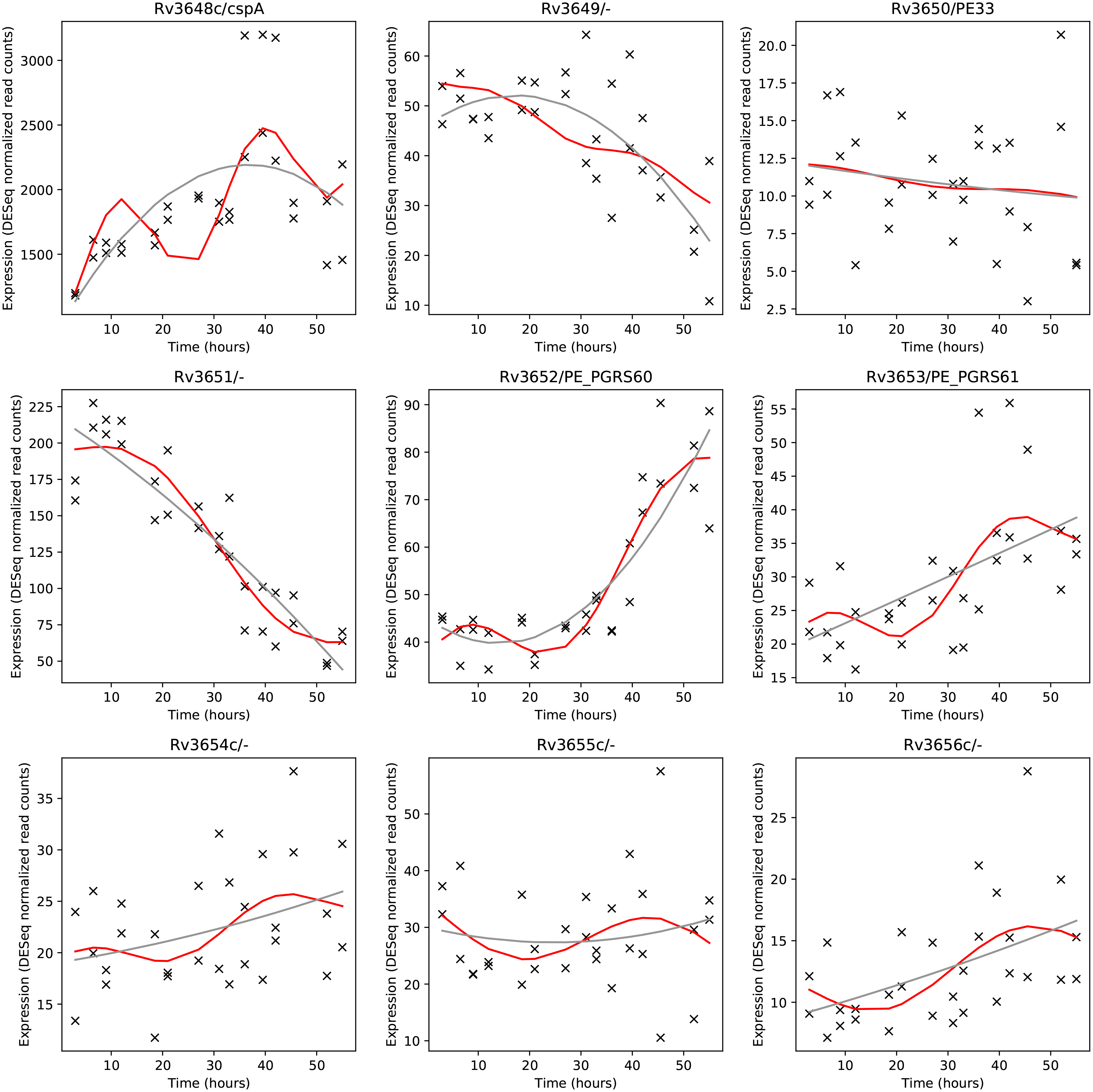
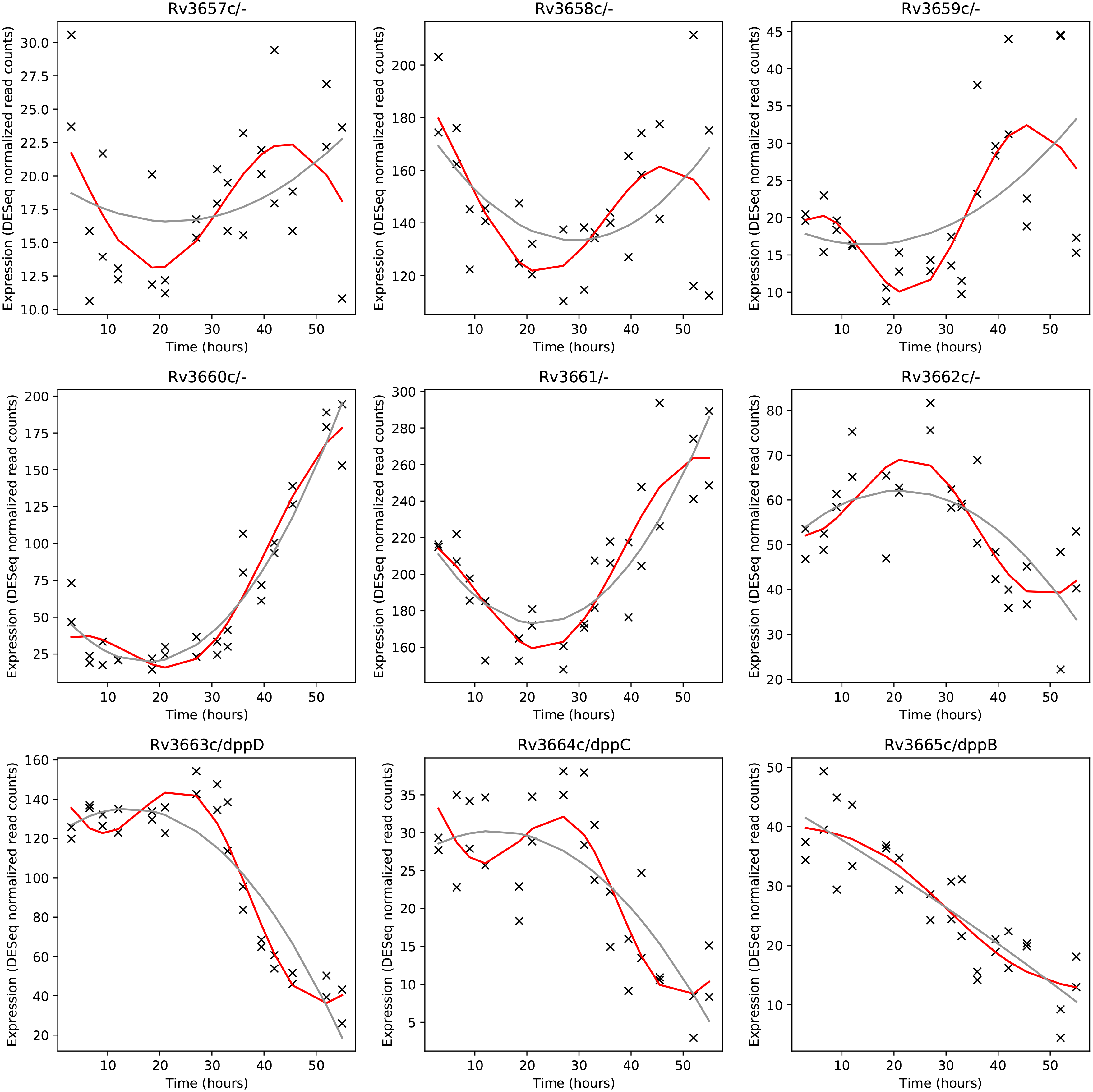
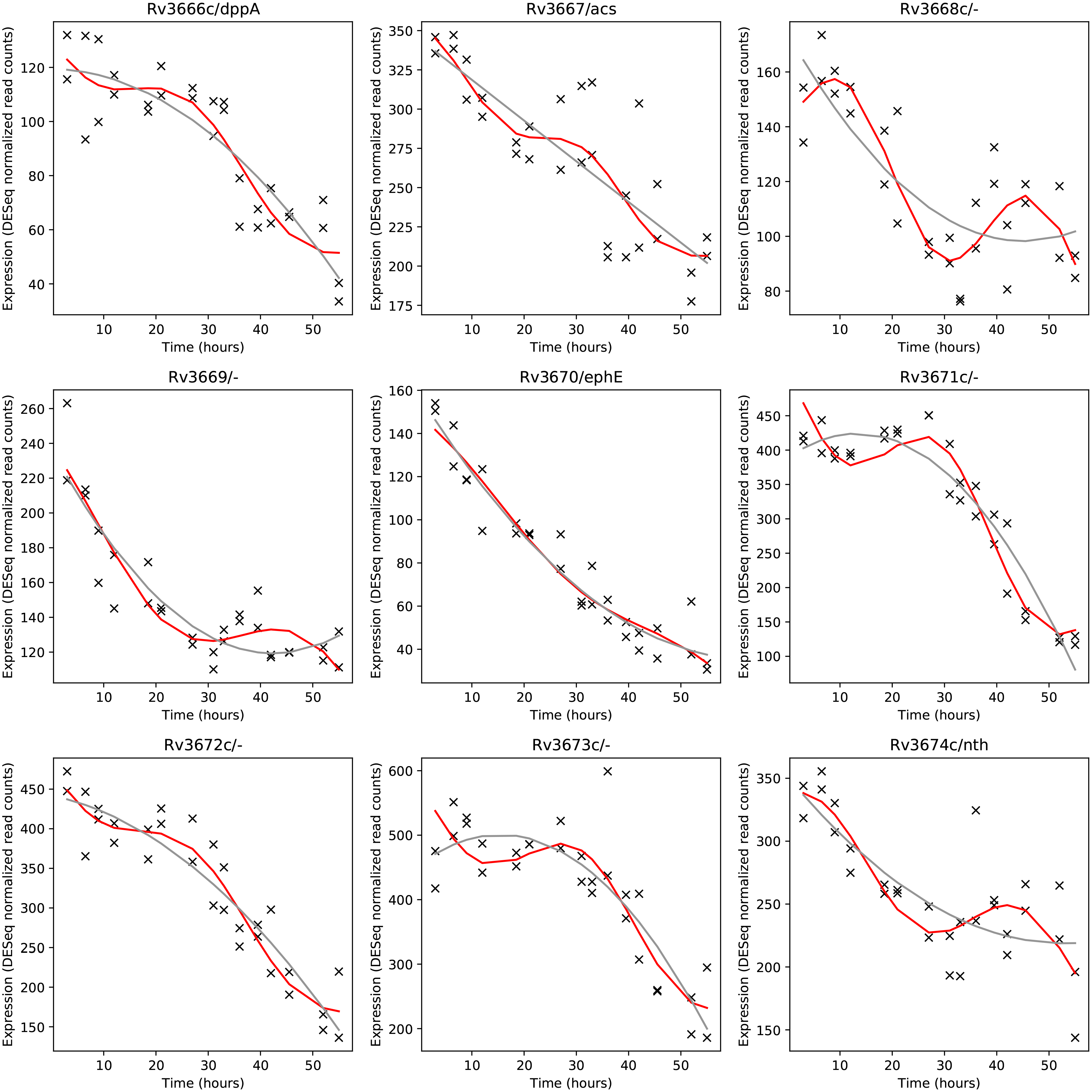
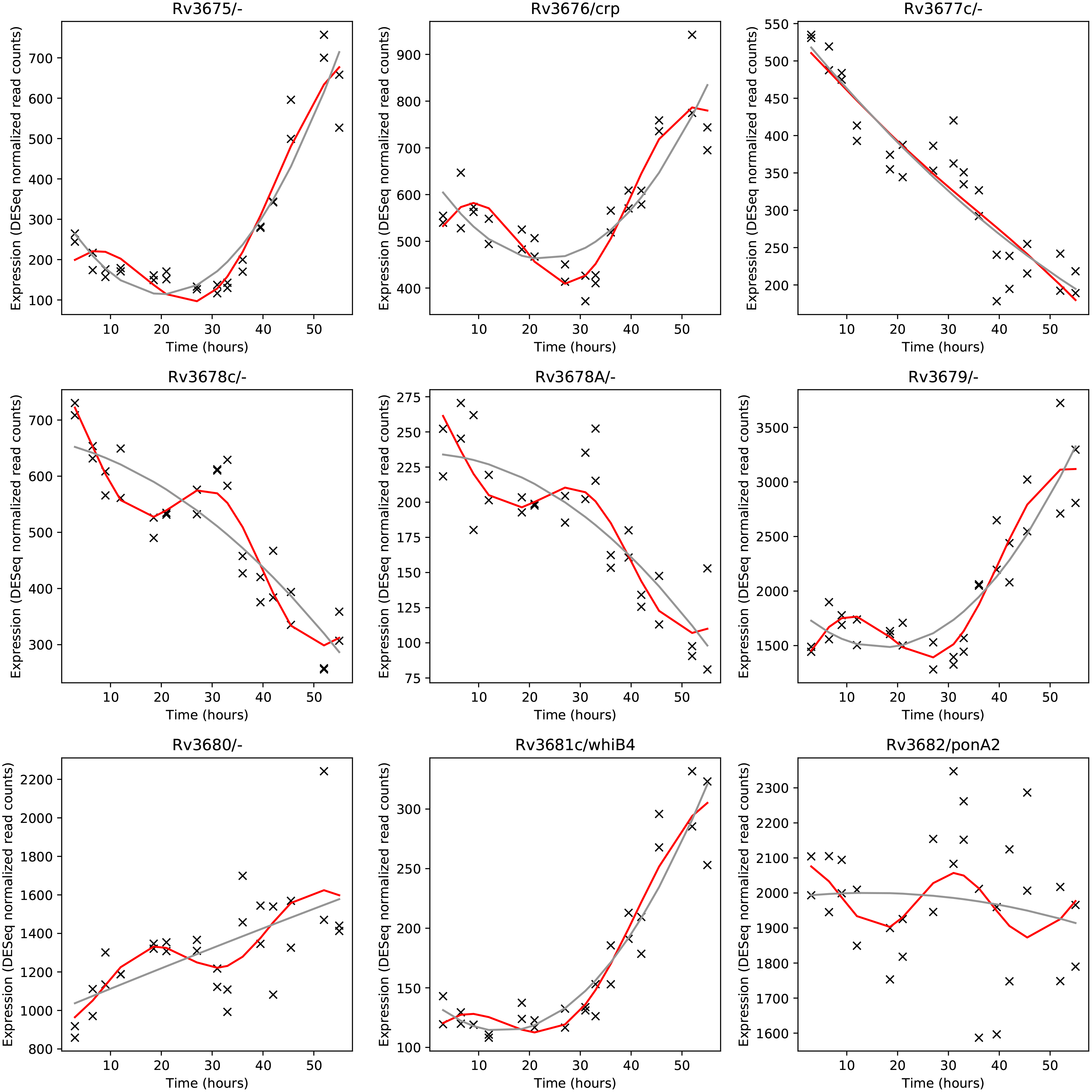
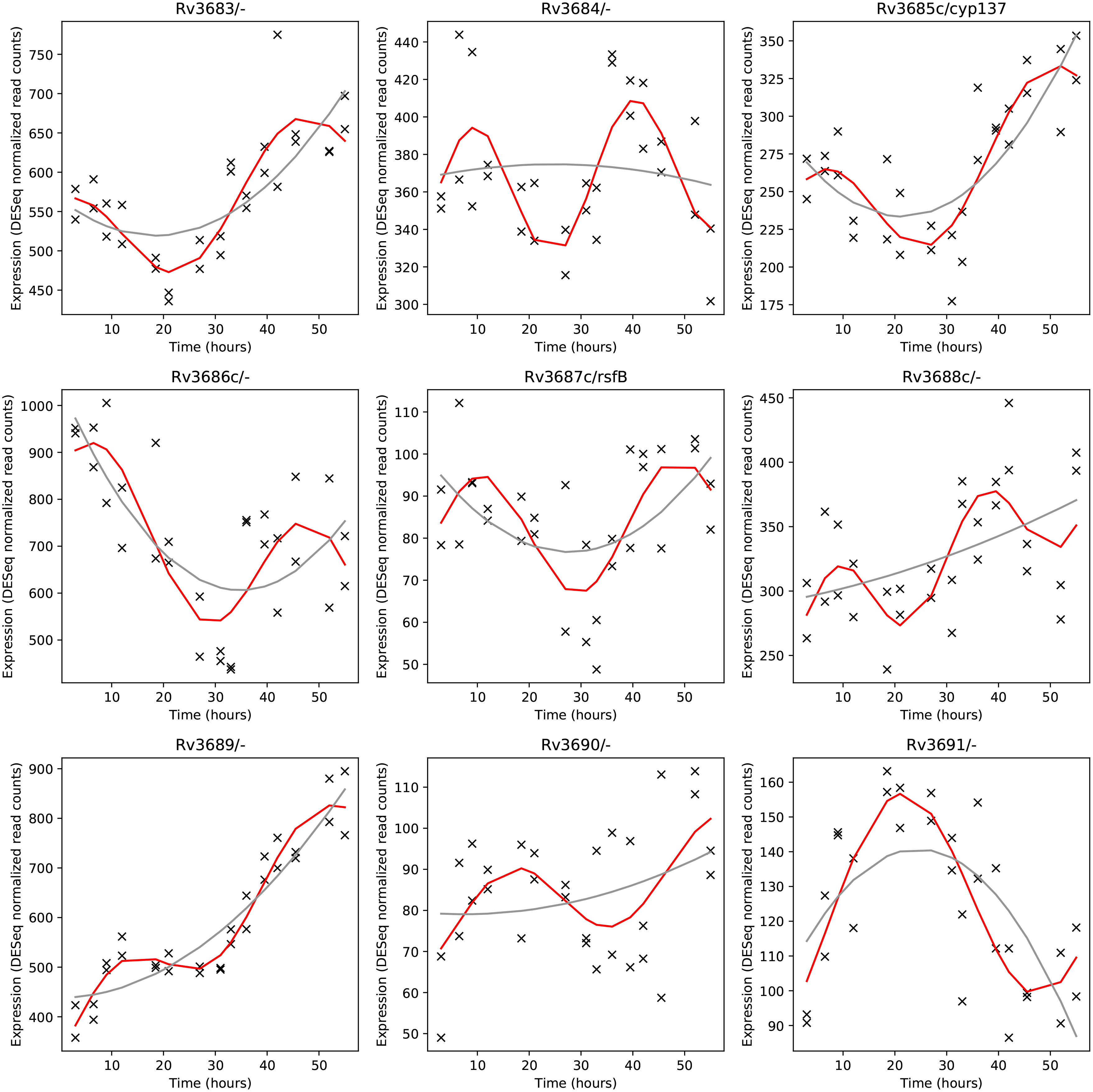
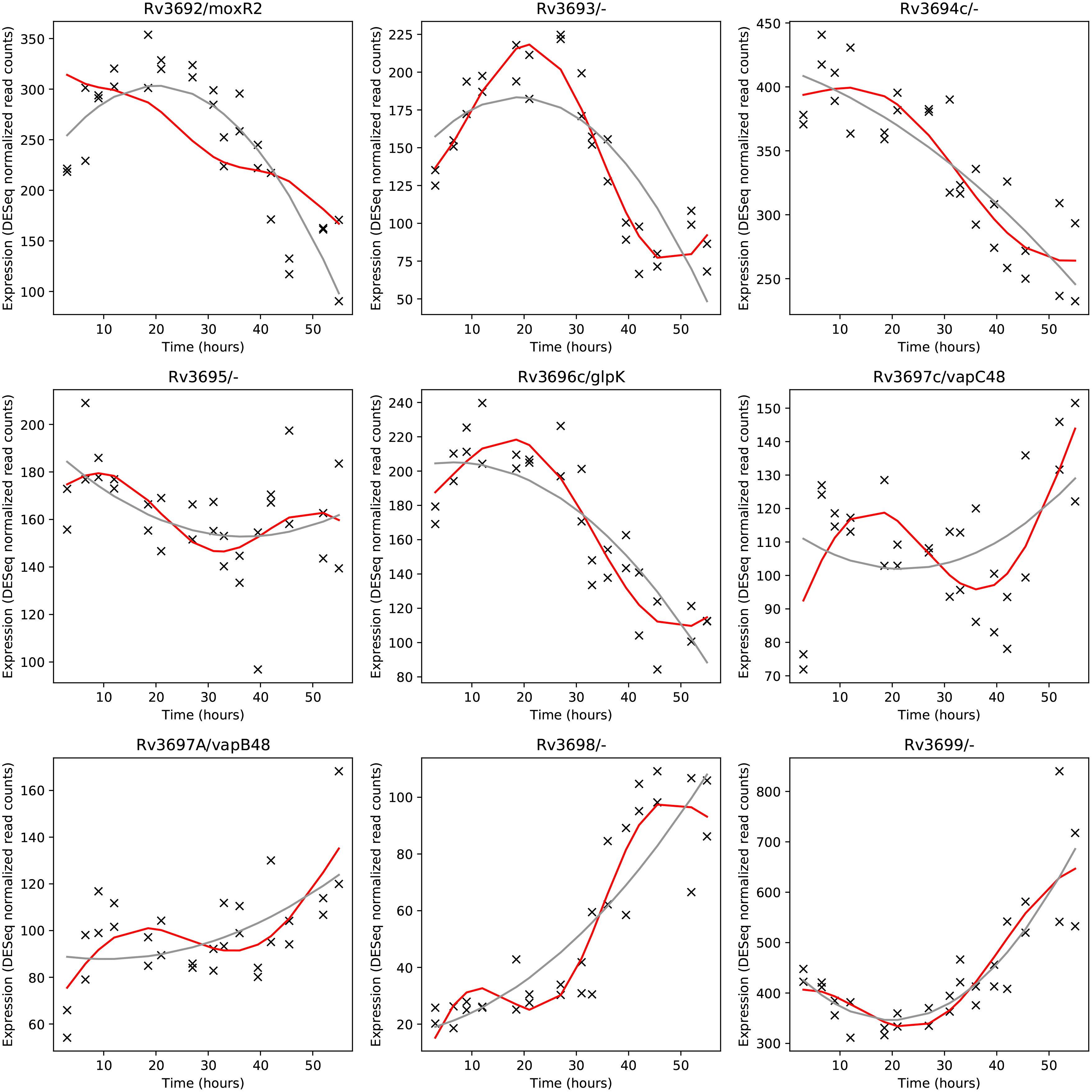
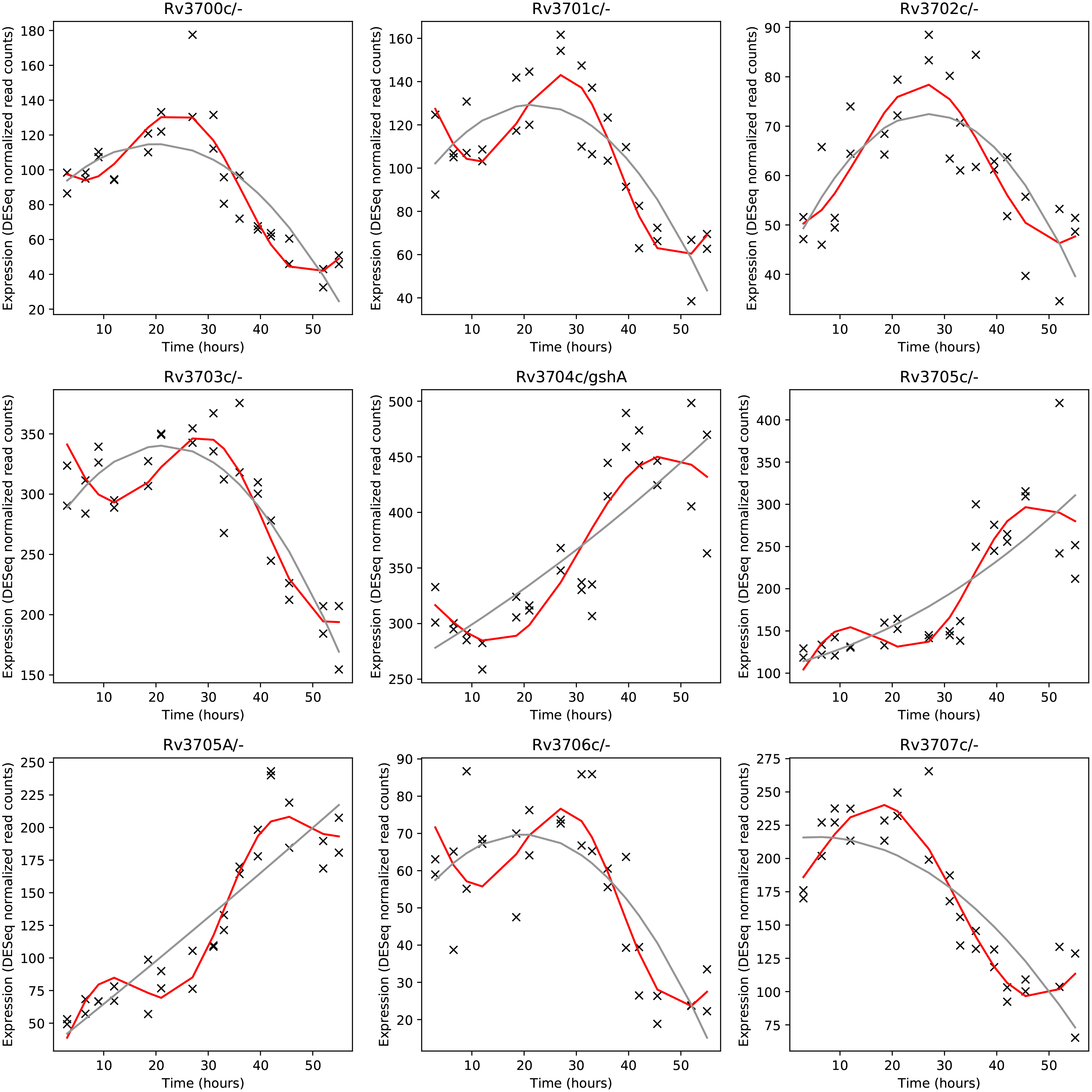
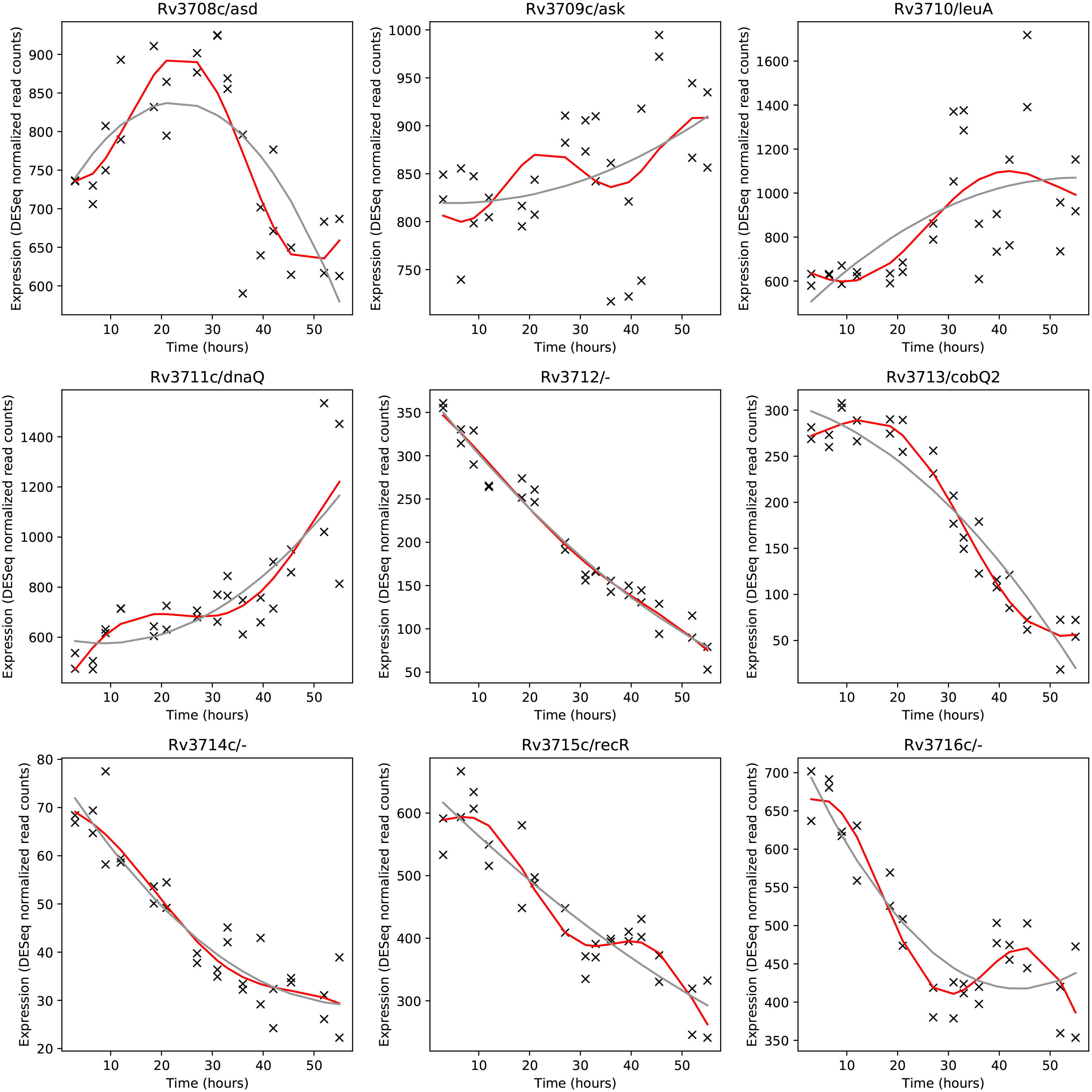
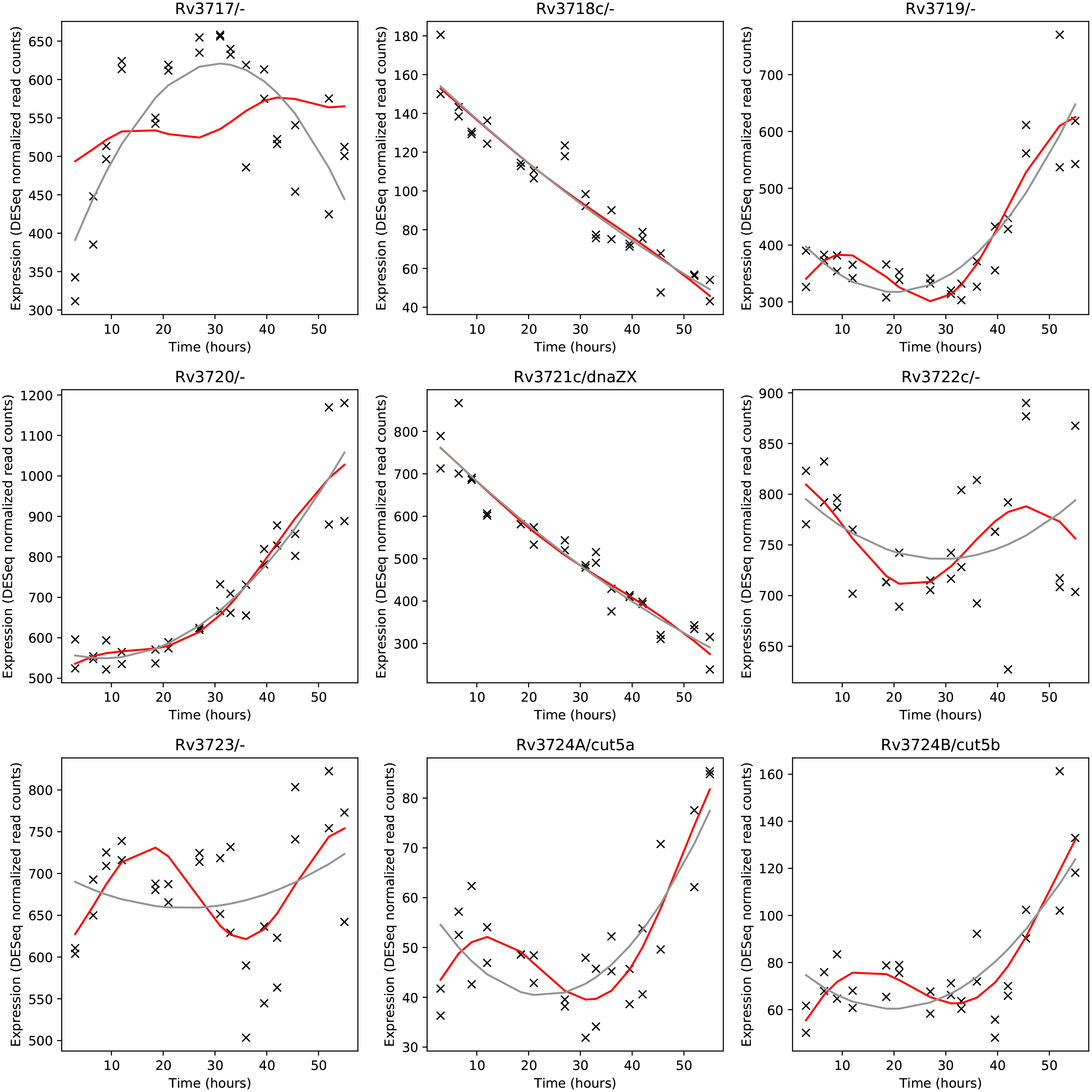
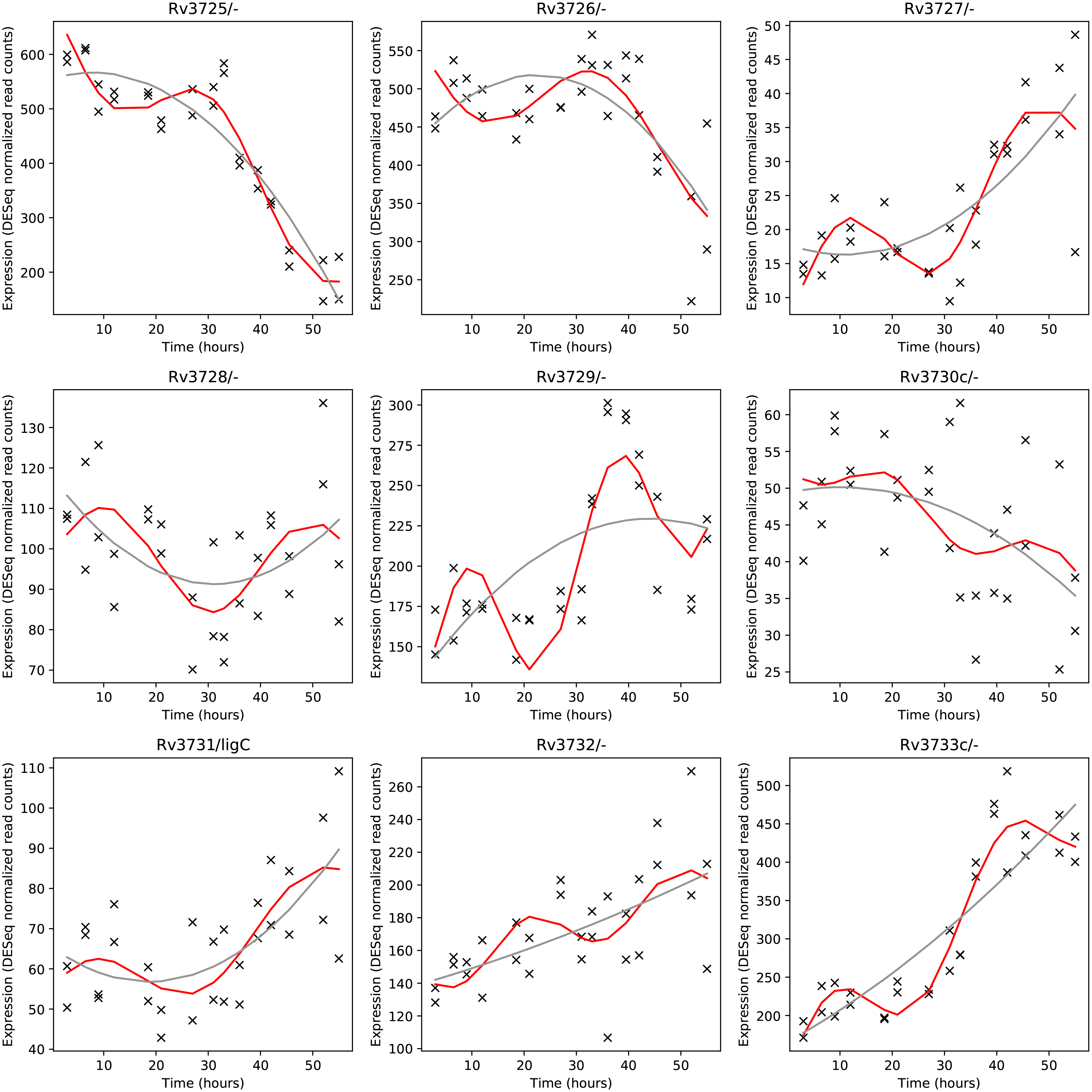
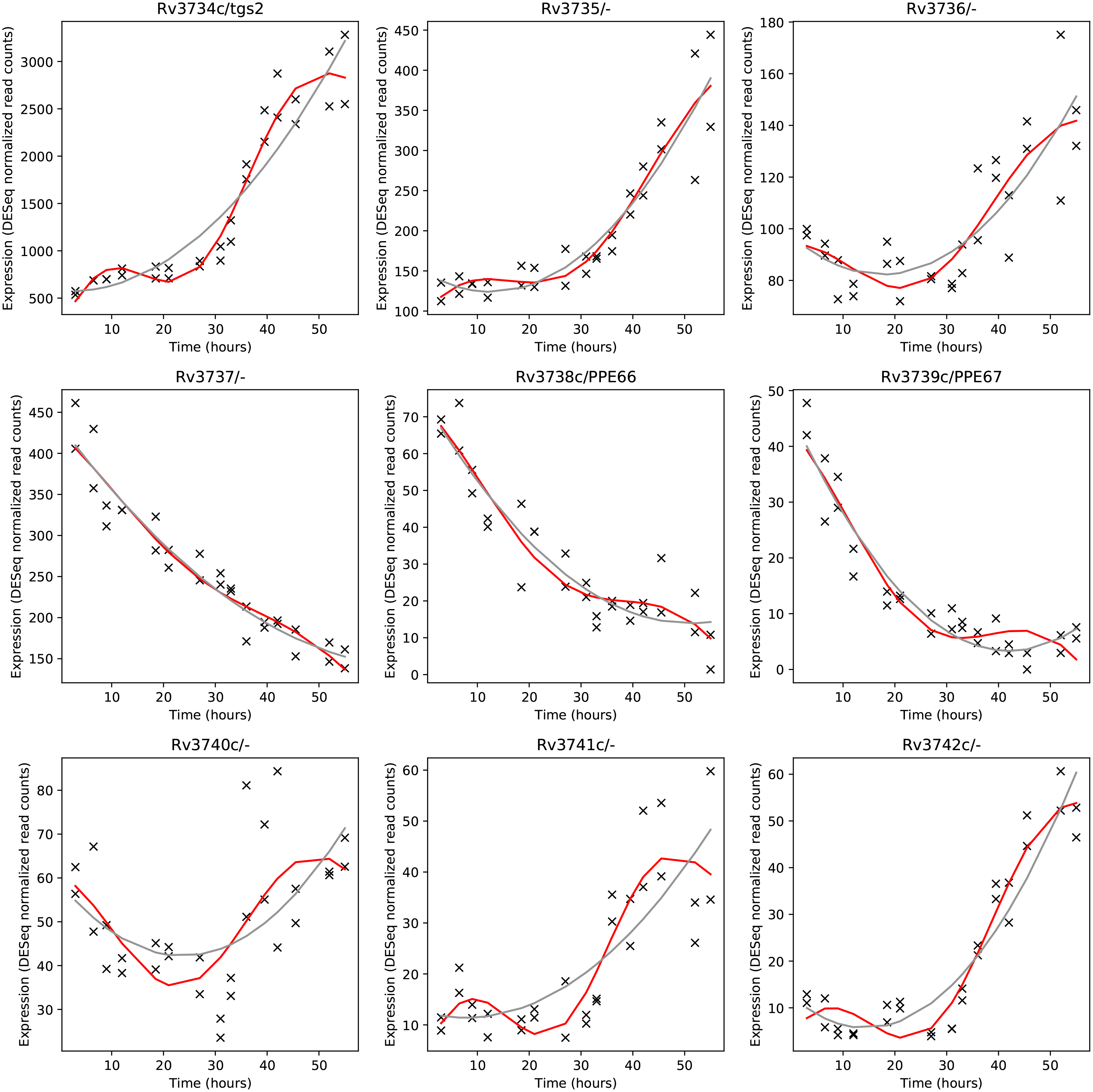
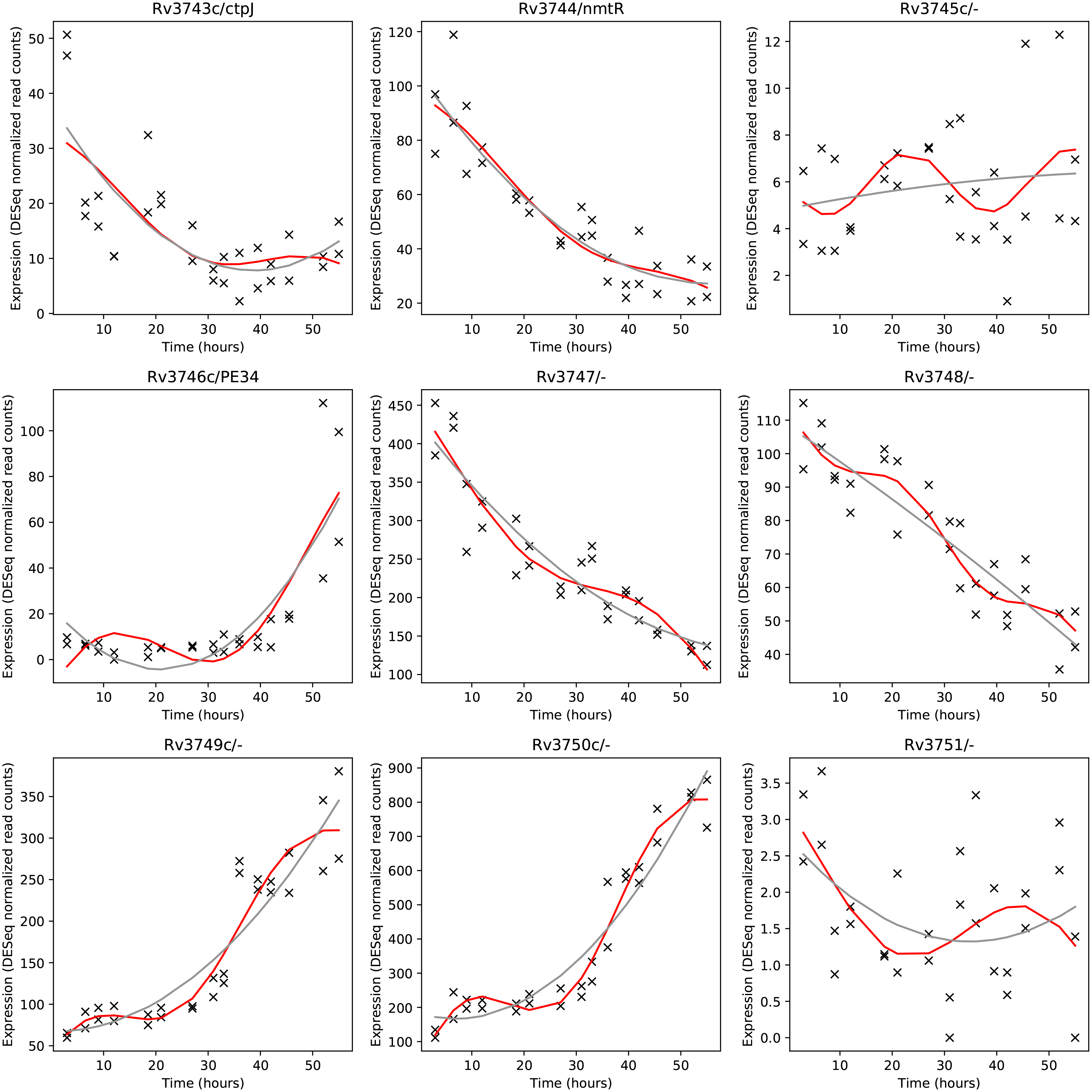
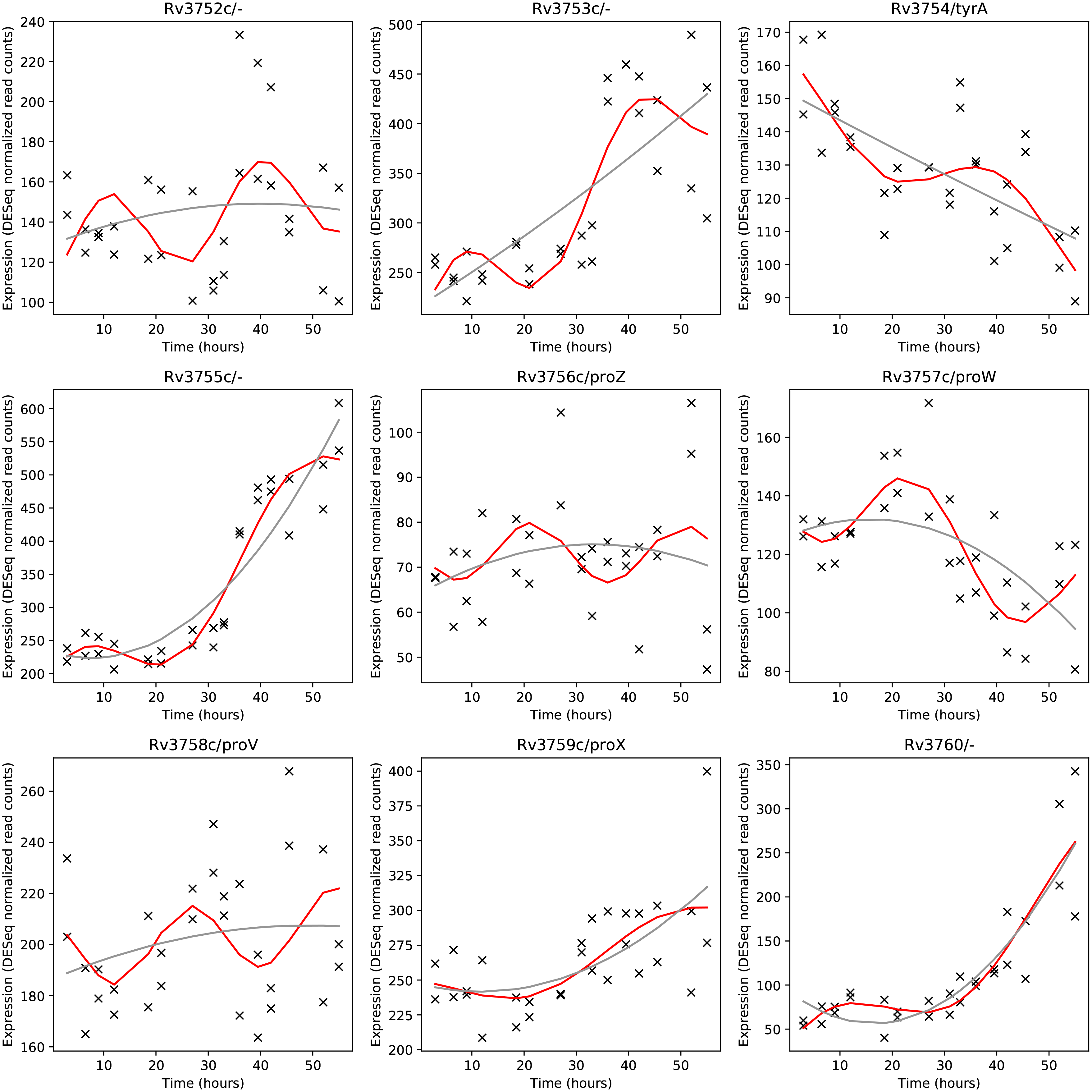
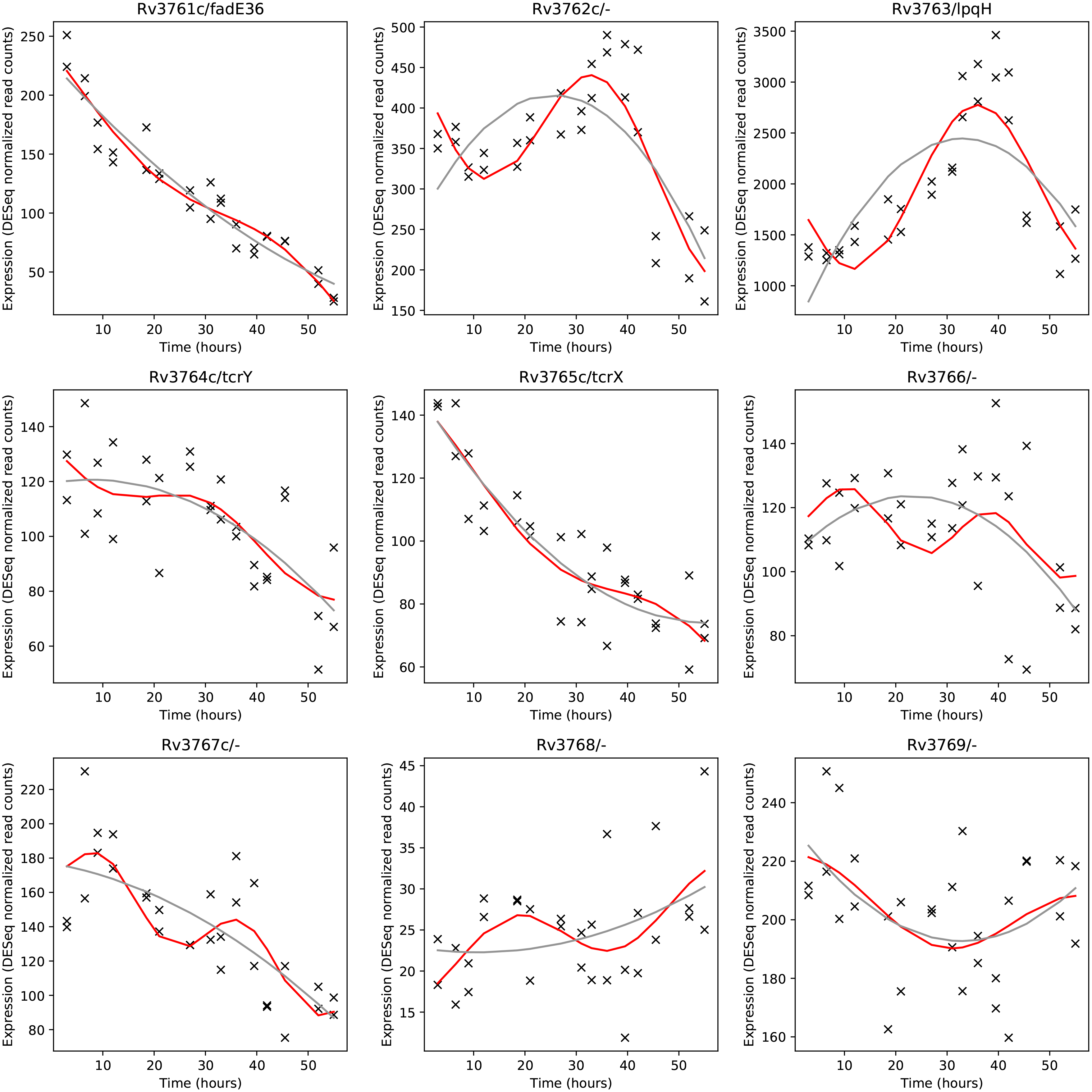
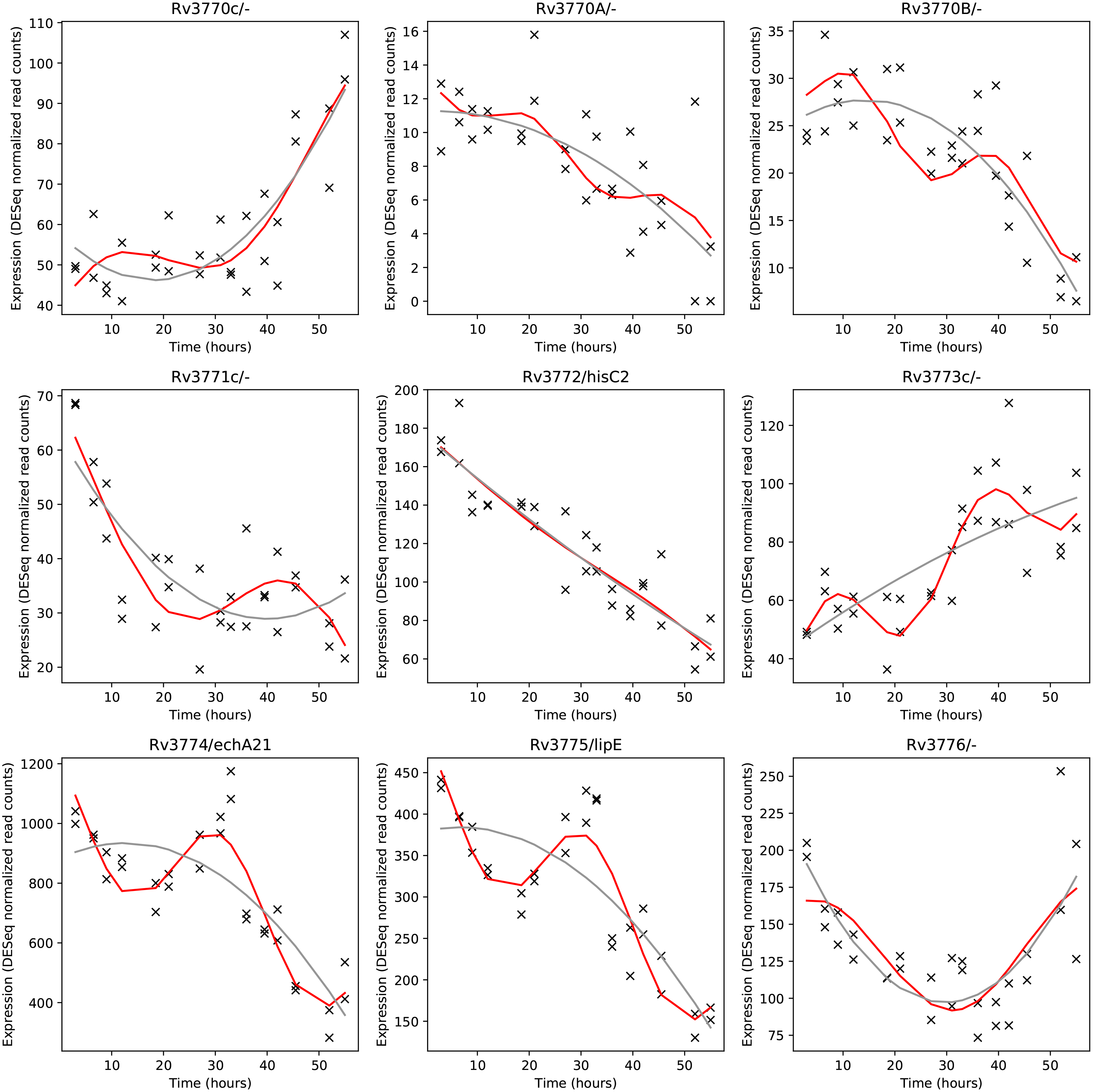
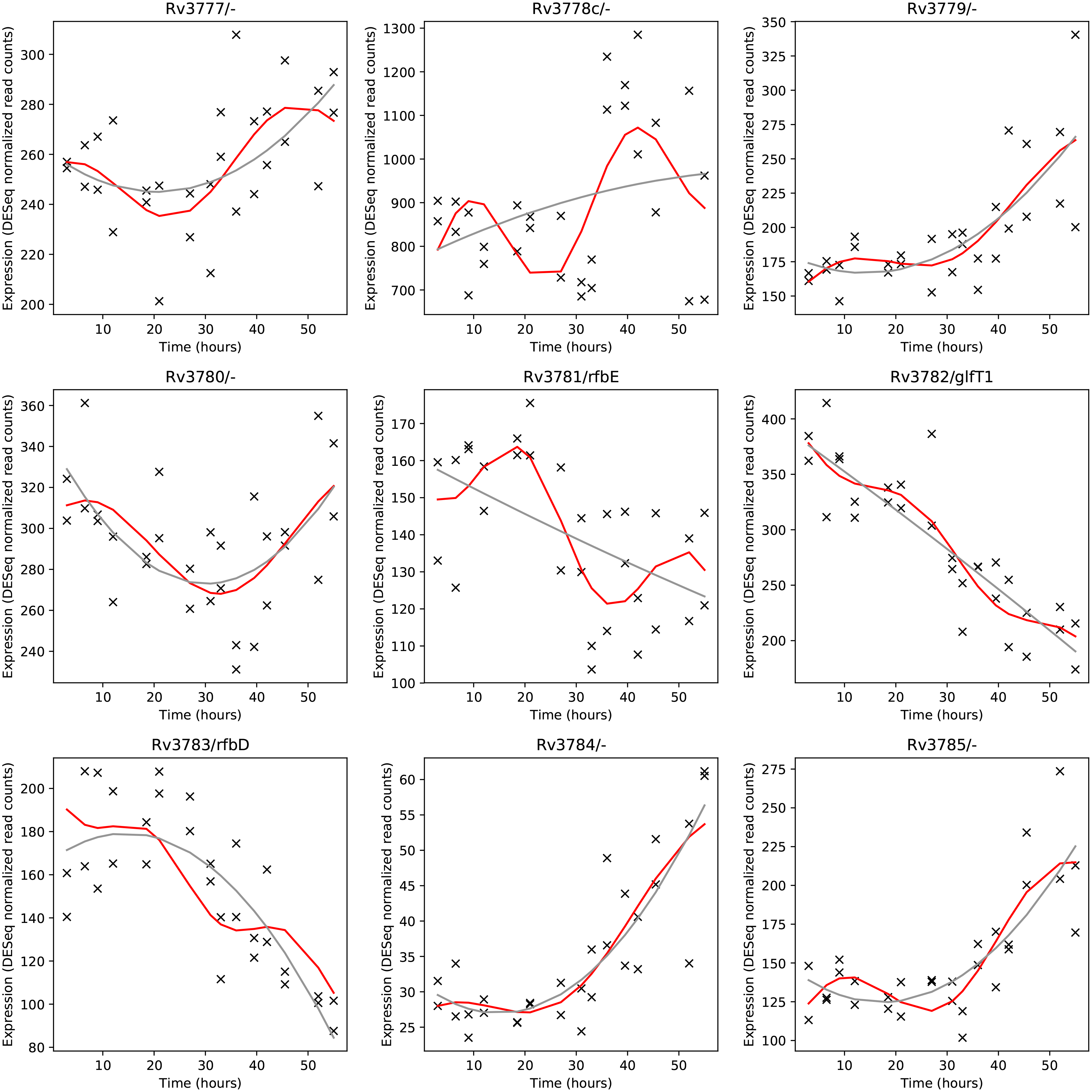
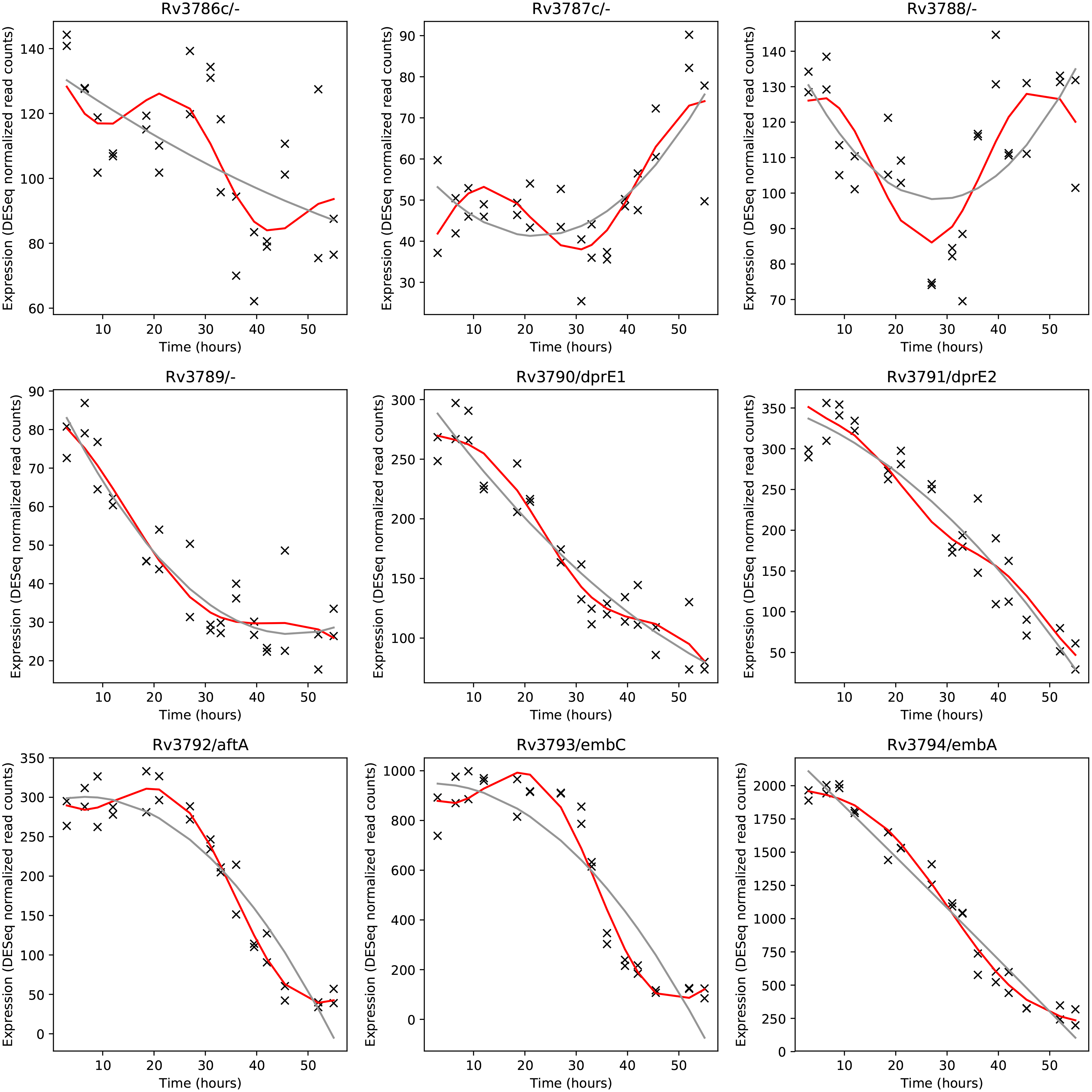
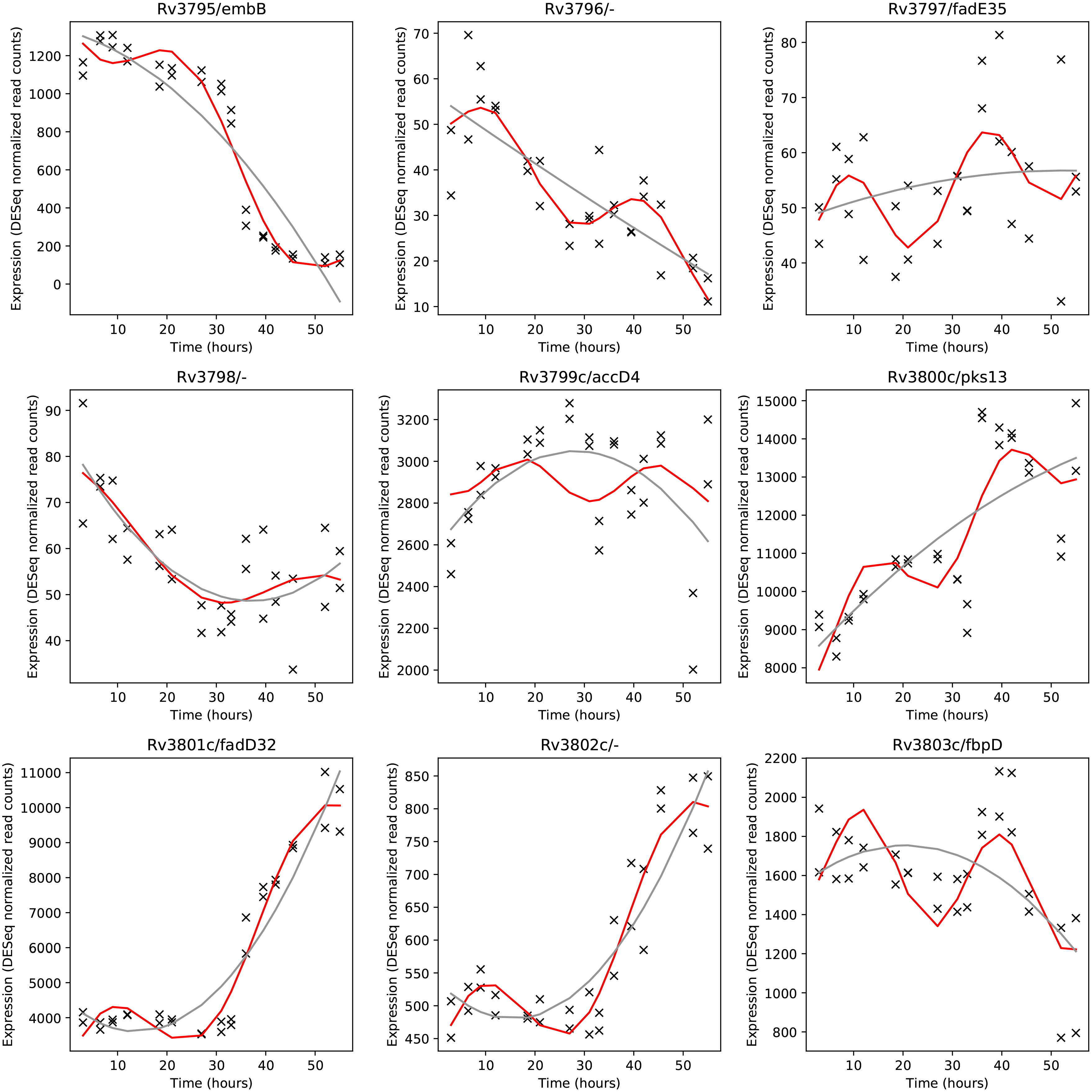
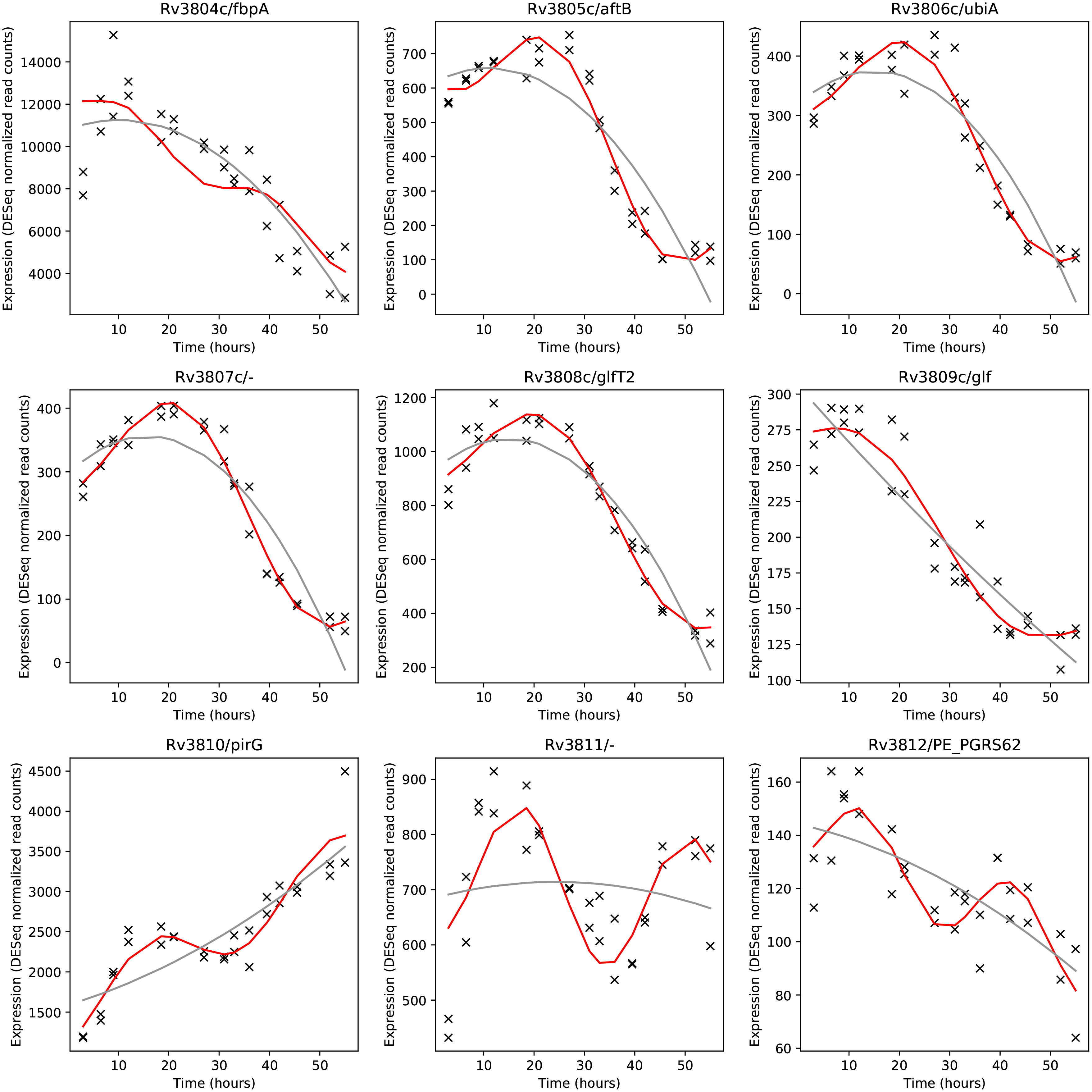
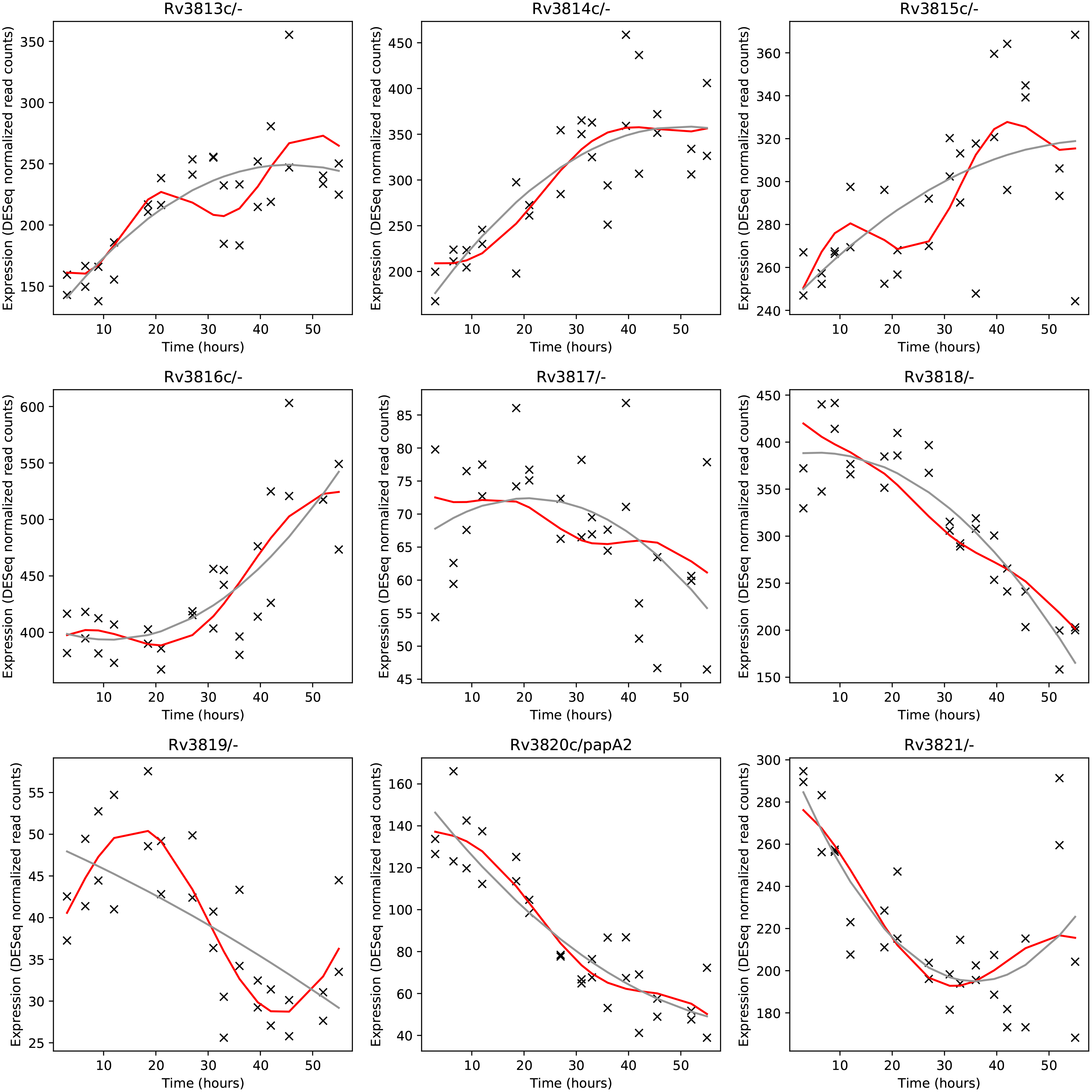
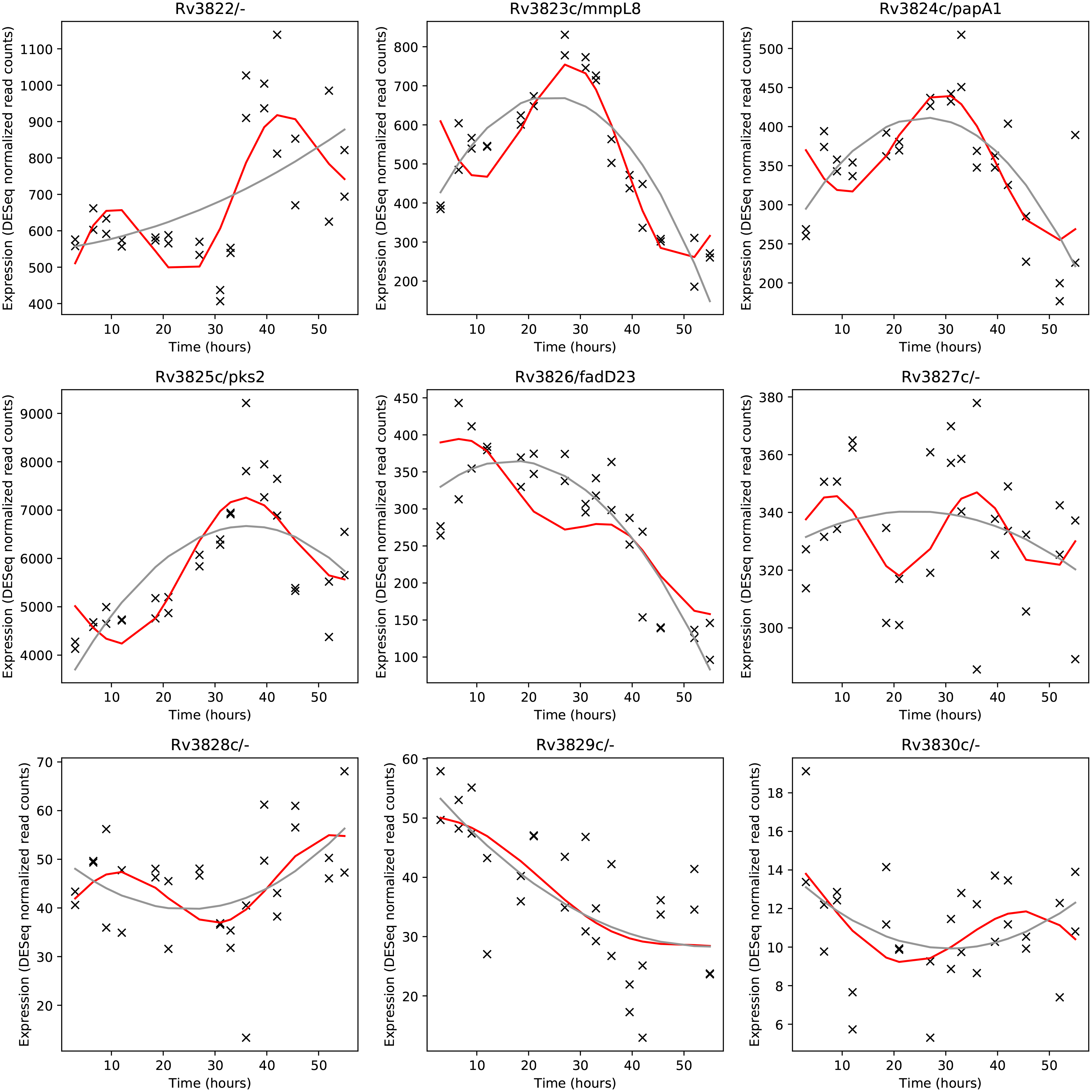
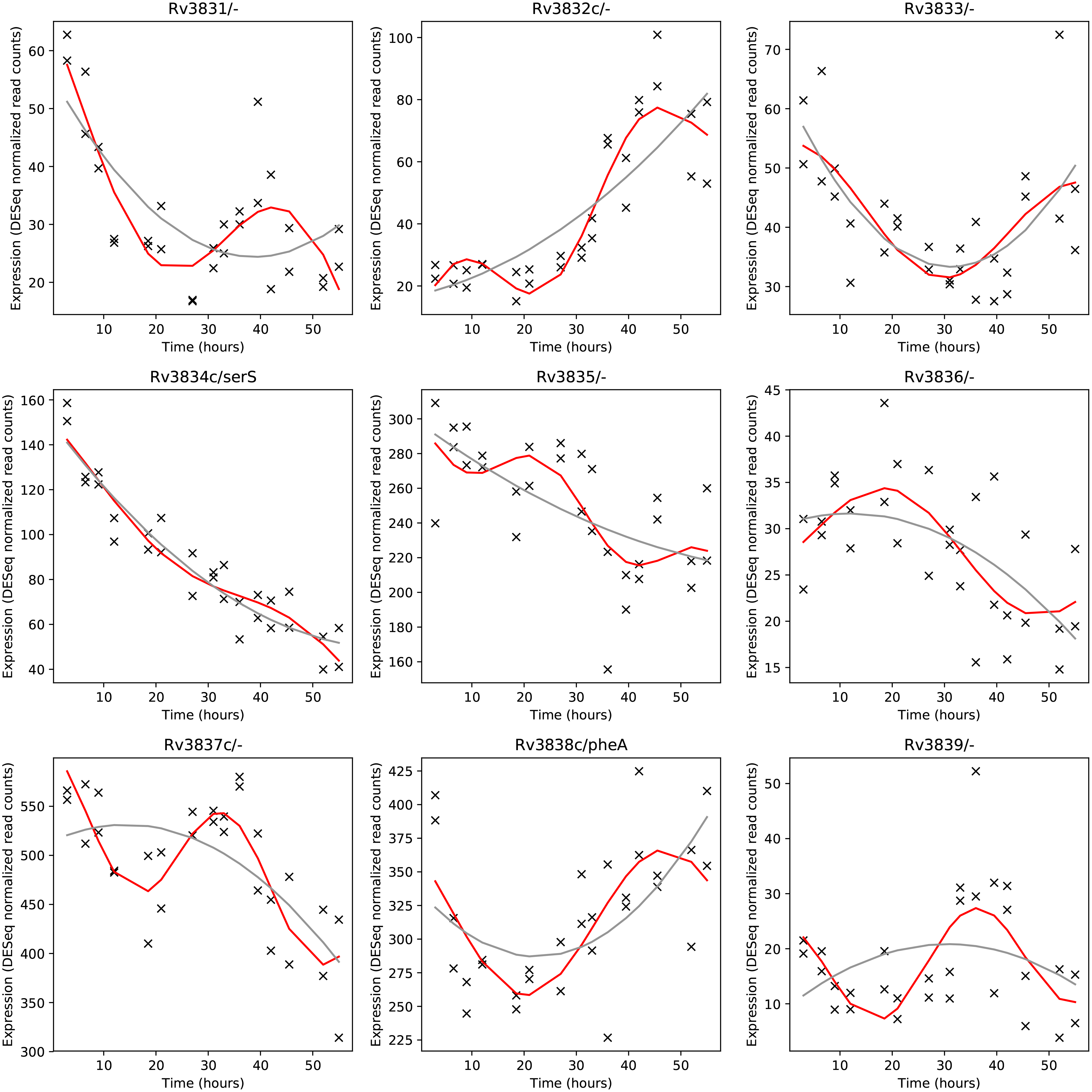
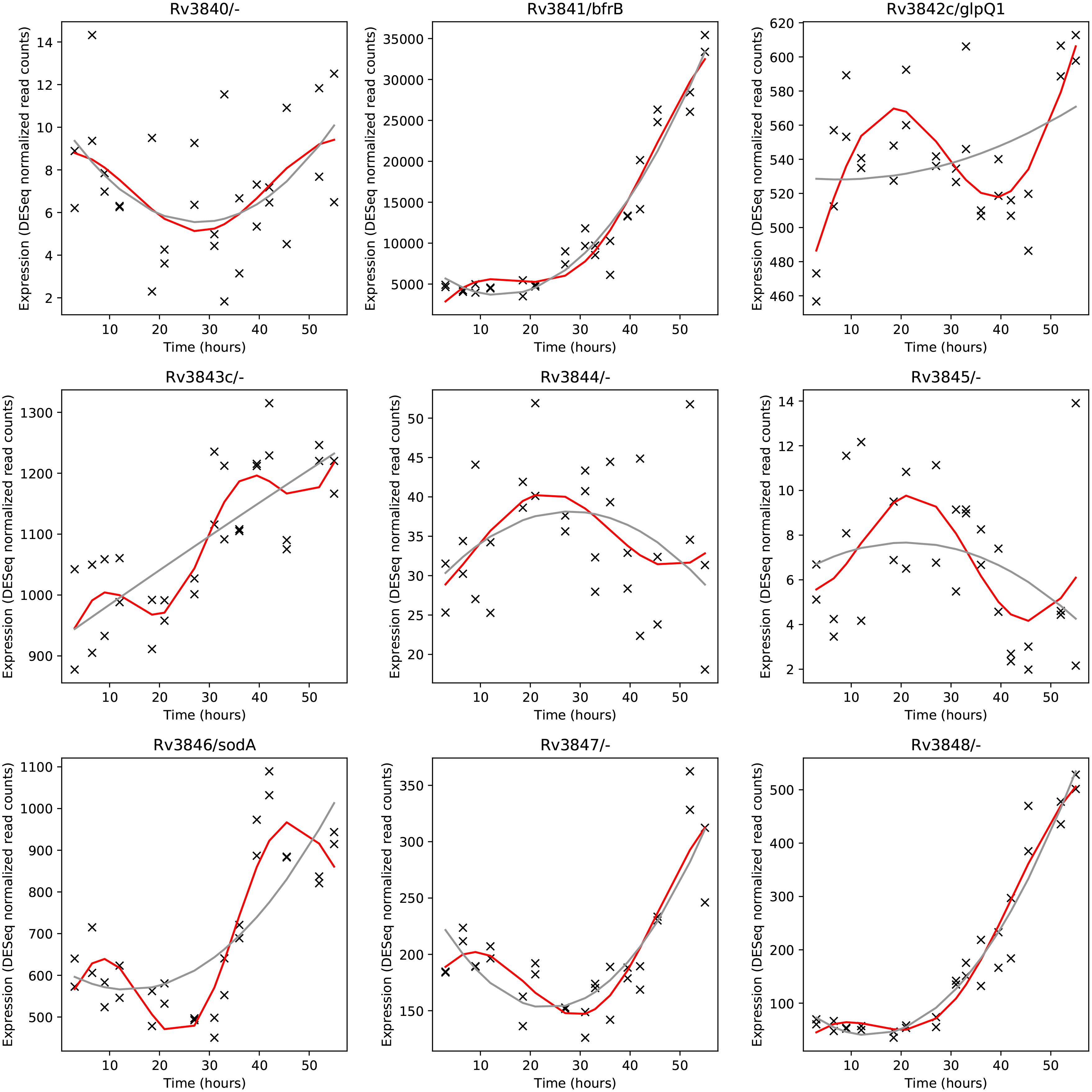
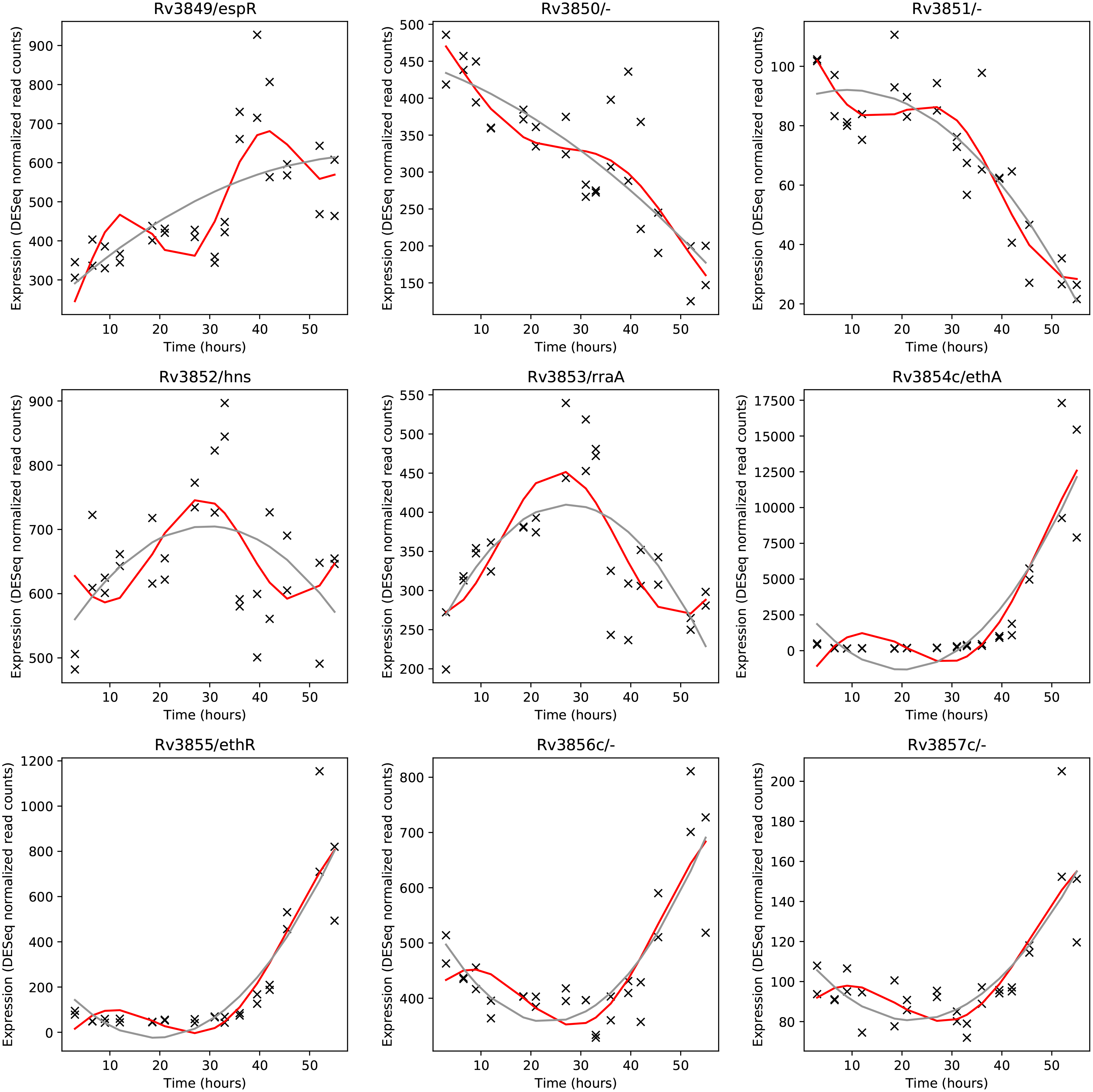
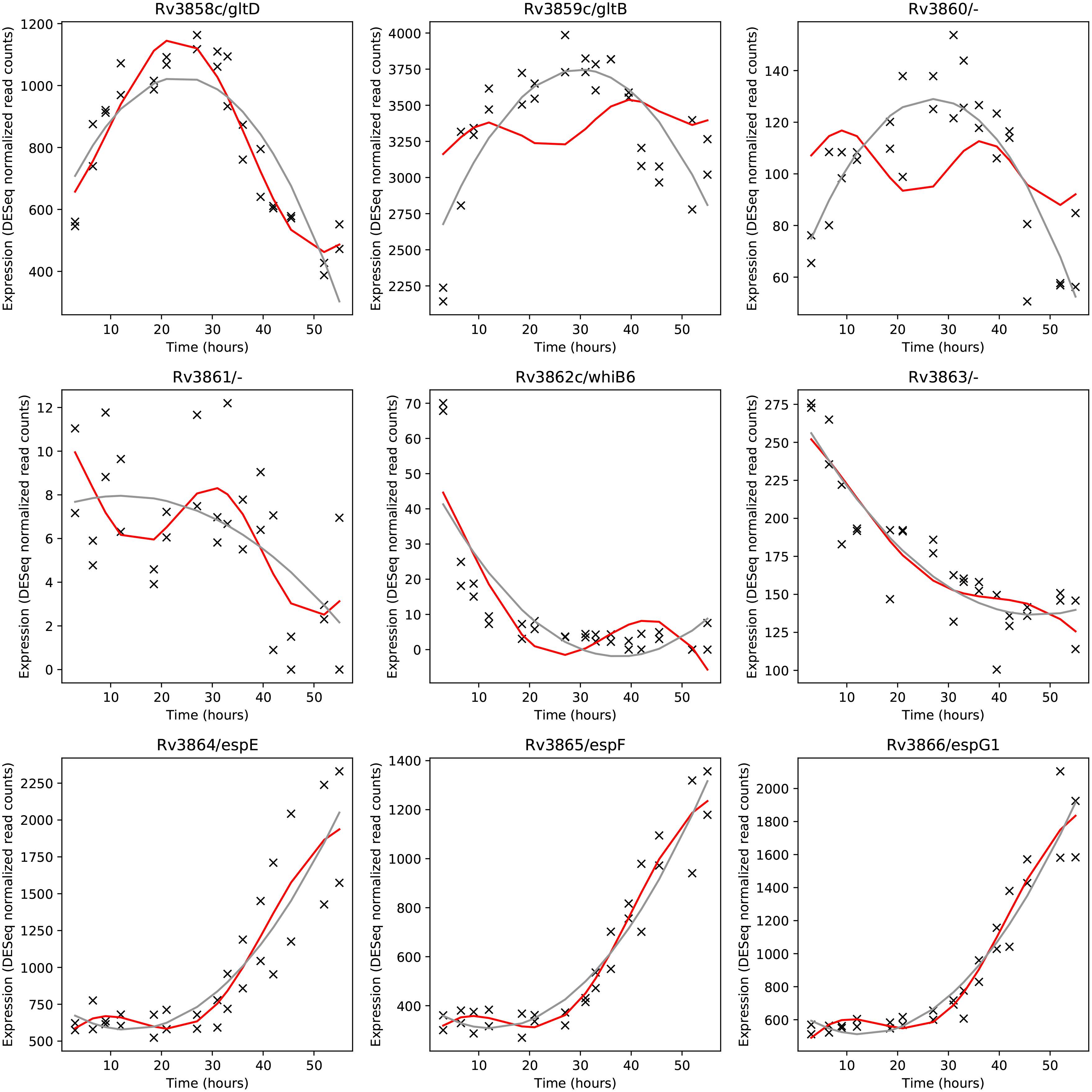
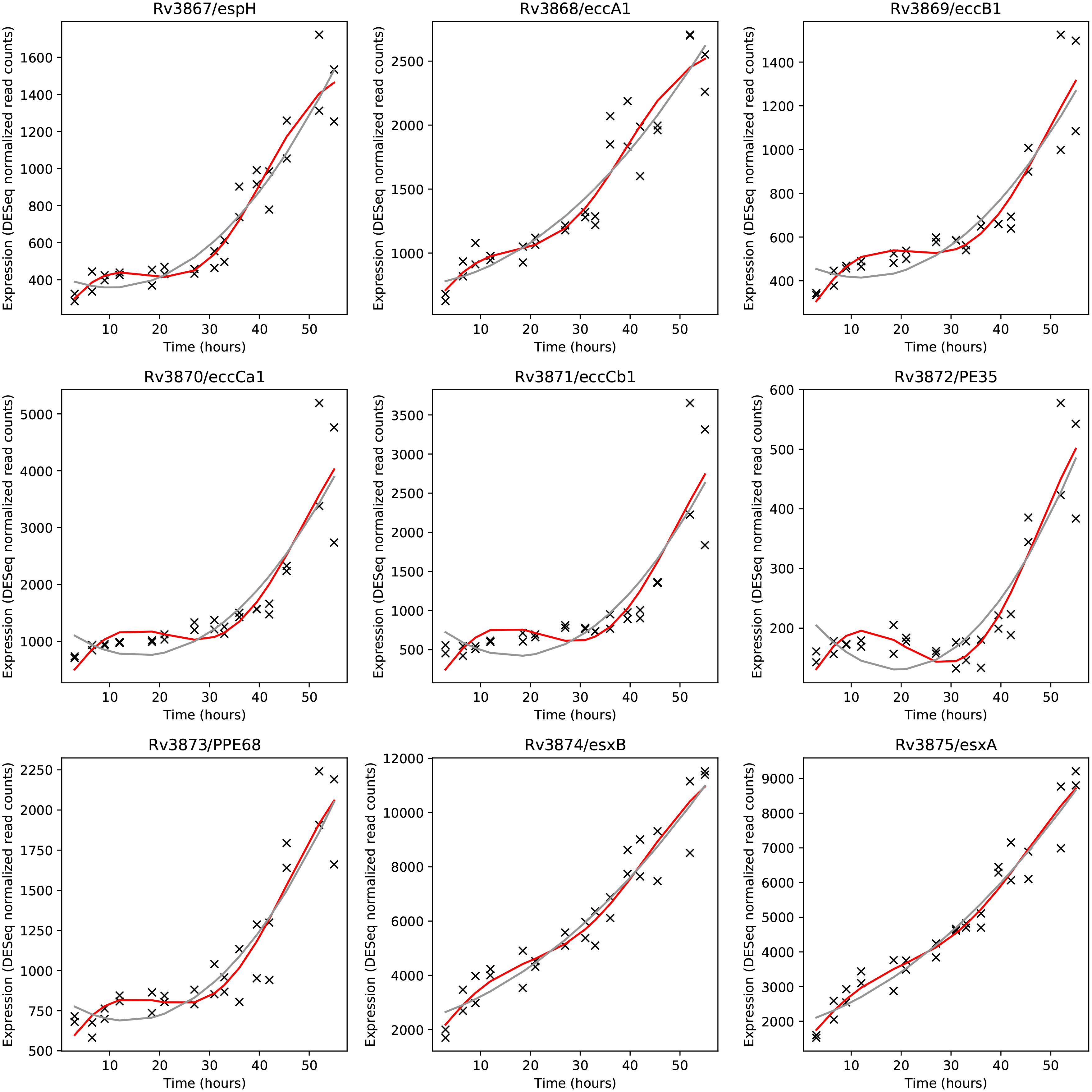
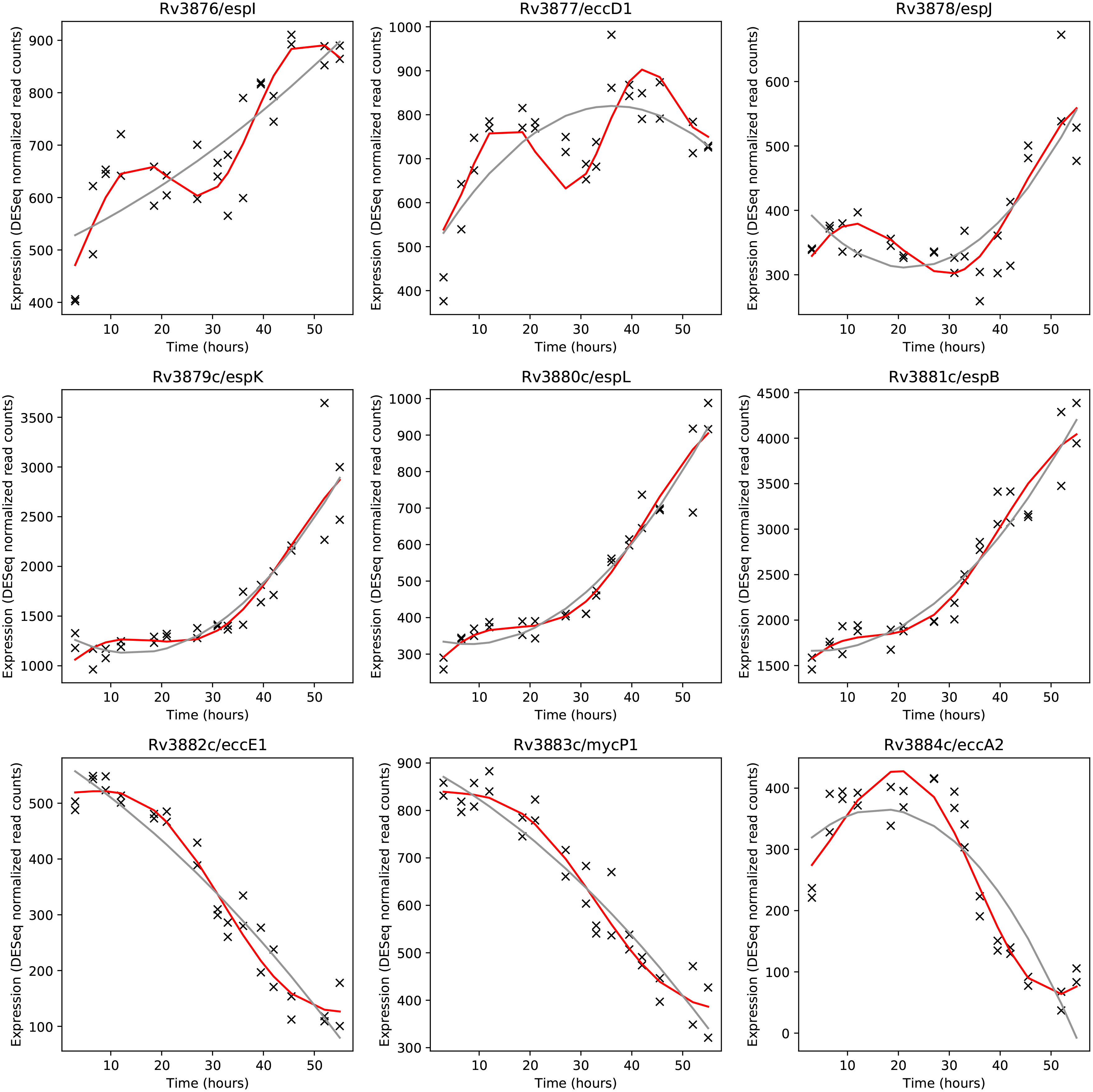
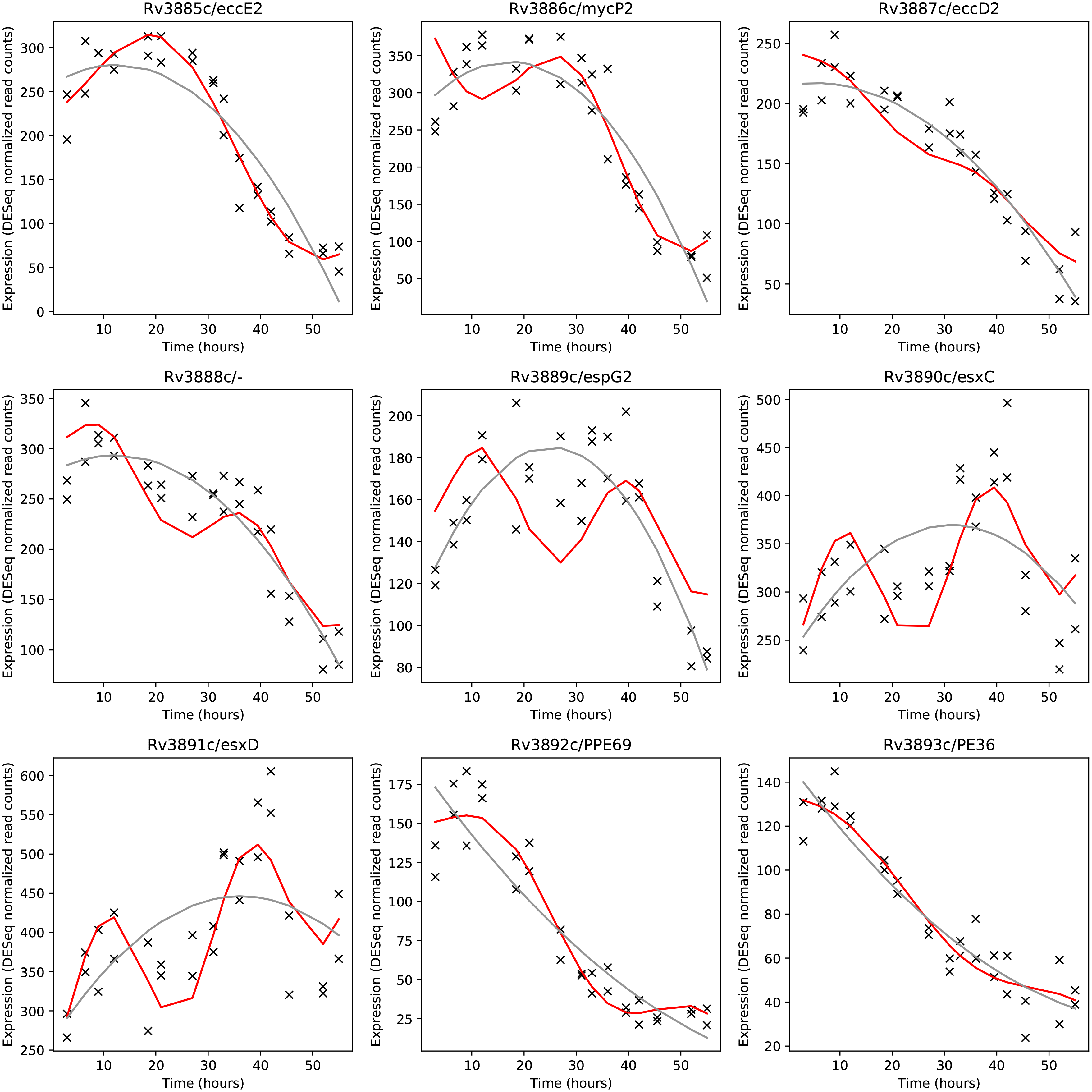
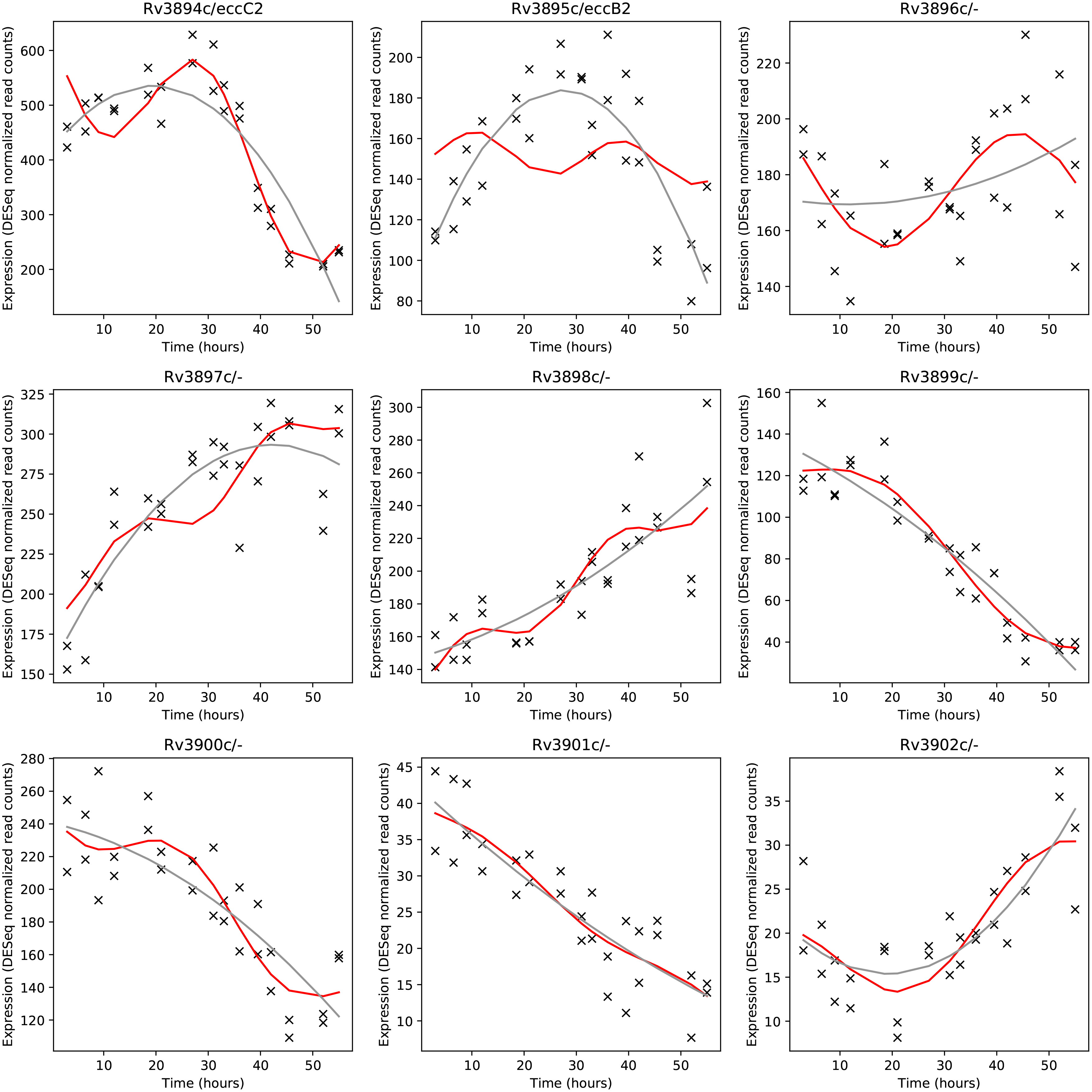
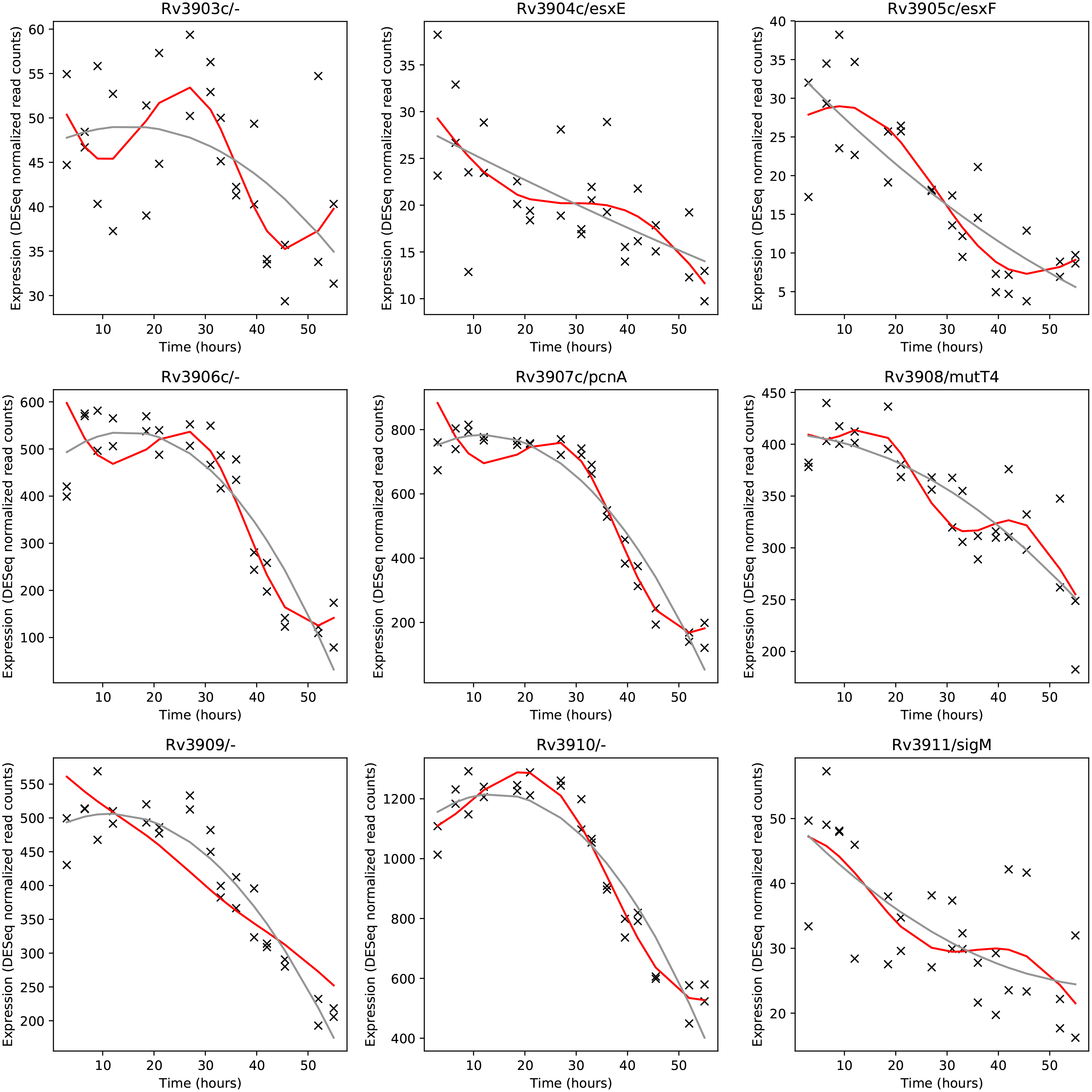
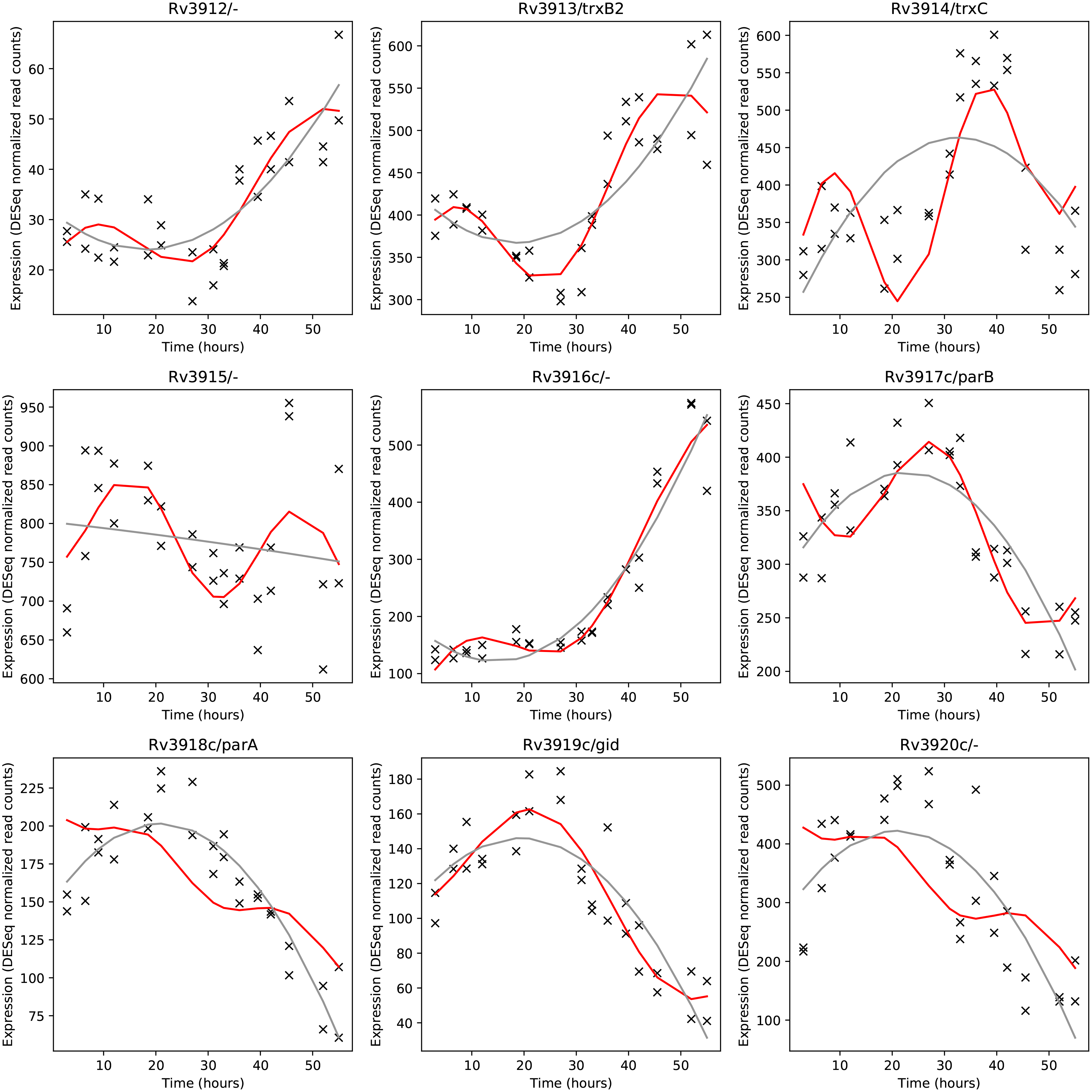
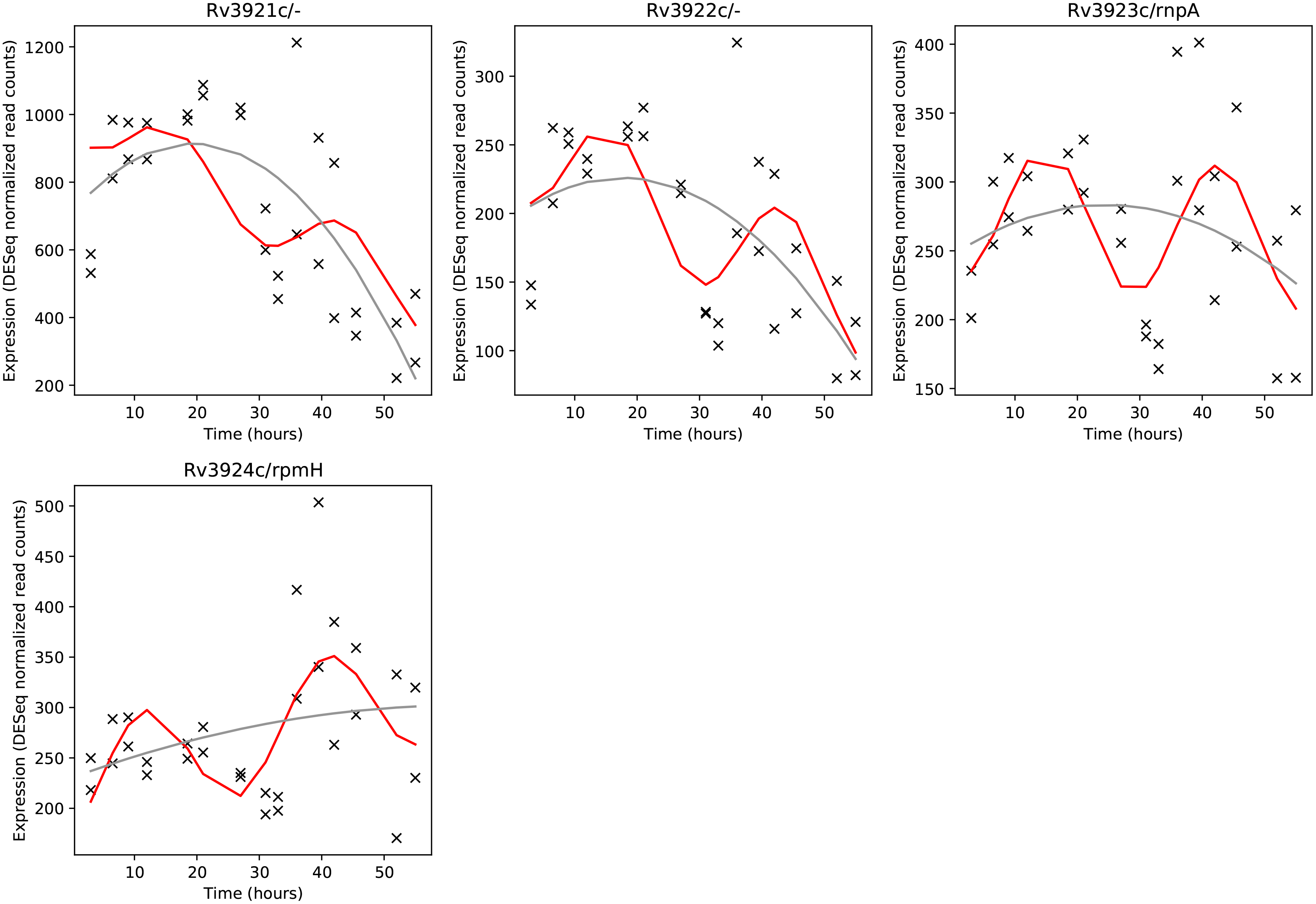
Sinusoidal fits (red) and quadratic fits (gray) to standard normalized DESeq read counts (count-mean/SD) for all genes in *MtbRv*

## Supplemental tables

Table 1: Normalized DESeq2 values for all detected 2948 genes which were deemed to be significantly expressed. Each value represents relative expression, the DESeq2 value normalized to the mean value of that gene across time. Genes determined to have one or two peaks in expression are indicated along with their peak cluster assignment.

Table 2: Standard normalized DESeq2 values for the 485 periodic genes (Count-mean/SD). Periodic gene cluster ID is indicated.

Table3: List of overlapping periodically expressed *Mtb* (this study) and differentially expressed *C crescentus* [7] orthologs.

